# Netrin-1 Acts as a Guardian of Naive Pluripotency in Human Embryonic Stem Cells

**DOI:** 10.64898/2026.07.16.738622

**Authors:** Agathe de Neufville, Etienne Masfaraud, Charbel Alfeghaly, Jan B. Stöckl, Nathalie Doerflinger, Guillaume Marcy, Cloé Rognard, Pierre Osteil, Marie Lantelme, Fabrice Lavial, Claire Chazaud, Thomas Fröhlich, Pierre Savatier, Irène Aksoy

## Abstract

We investigated the role of Netrin-1 (NTN1) in human naïve pluripotency using complementary loss- and gain-of-function approaches. In primate embryos and human embryonic stem cells (hESCs), Netrin-1 expression is associated with the naïve pluripotent state. Disruption of *NTN1* had no detectable effect on hESCs maintained on murine embryonic fibroblasts. However, under sub-optimal culture conditions, NTN1-knockout cells exhibited compromised naïve pluripotency, which was rescued with feeder cells overexpressing Netrin-1. Netrin-1 overexpression in hESCs accelerated acquisition of the naïve state and markedly increased resistance to differentiation. These effects were accompanied by extensive epigenetic remodeling, including H3K27ac and H2K27me3. Proteomic and phospho-proteomic analyses further revealed rapid Netrin-1-dependent alterations in pathways controlling cell adhesion, signaling, and chromatin regulation. Together, these findings extend the role of Netrin-1 beyond its established functions and identify it as a coordinator of extracellular cues, intracellular signaling, and nuclear regulatory mechanisms that support human naïve pluripotency.

## Introduction

Pluripotent stem cells (PSCs) represent the *in vitro* counterparts of the pluripotent cells that constitute the epiblast of the pre- and early post-implantation mammalian embryo. In the mouse, PSCs exist in two distinct states that reflect successive stages of epiblast development: embryonic stem cells (ESCs), which self-renew in a naïve state and correspond to the pre-implantation epiblast, and epiblast stem cells (EpiSCs), which self-renew in a primed state and resemble the post-implantation epiblast at the onset of gastrulation (Endoh and Niwa, 2022; Nichols and Smith, 2009; Ying and Smith, 2017). A comparable dichotomy has been described in human PSCs, which can be maintained in naïve-like or primed-like configurations (Chen and Lai, 2014; Davidson et al., 2015; Rossant and Tam, 2017; Ware, 2017). However, in contrast to the mouse, the stabilization of human naïve pluripotency remains technically demanding and relies on the combined modulation of multiple signaling pathways. Current culture systems for human naïve PSCs are largely based on the activation of LIF and WNT signaling—typically through inhibition of GSK3—and the concomitant suppression of differentiation-associated pathways such as the MEK/ERK cascade (Manor et al., 2015; Weinberger et al., 2016). Additional inhibition of pathways including p38 MAPK, SRC, JNK, tankyrases, and protein kinase C further enhances self-renewal and limits spontaneous differentiation (Bayerl et al., 2021; Guo et al., 2017; Guo et al., 2016). Together, these observations indicate that naïve pluripotency is maintained through a finely tuned signaling equilibrium, in which the balance between self-renewal and differentiation cues must be tightly controlled.

Beyond intracellular signaling, increasing evidence suggests that extracellular cues and cell–environment interactions contribute to the regulation of pluripotent states (Gattazzo et al., 2014; Guilak et al., 2009; Vitillo and Kimber, 2017). In this context, secreted guidance molecules, classically studied in developmental processes, may play underappreciated roles in modulating pluripotent cell identity. Netrins are a family of secreted proteins initially characterized for their roles in axon guidance during nervous system development (Kennedy et al., 1994; Serafini et al., 1994). Among them, Netrin-1, encoded by *NTN1*, is a laminin-related protein that functions as a pleiotropic ligand in a wide range of developmental and pathological contexts (Cirulli and Yebra, 2007; Grandin et al., 2016). Netrin-1 signals through multiple receptors, including Deleted in Colorectal Carcinoma (DCC), Neogenin (NEO1), and members of the UNC5 family, and can elicit distinct cellular responses depending on receptor composition (Bell et al., 2013; Rajagopalan et al., 2004; Xu et al., 2014). This receptor-dependent signaling underlies its ability to exert both attractive and repulsive effects on migrating cells (Ko et al., 2012; Xu *et al*., 2014). In addition to these canonical receptors, Netrin-1 interacts with integrins such as α6β4 and α3β1, thereby influencing cell adhesion and migration (Lee et al., 2016; Stanco et al., 2009b; Yebra et al., 2003a). These properties position Netrin-1 as a regulator of cell behavior at the interface between extracellular cues and intracellular signaling pathways.

Recent work has implicated Netrin-1 in early embryonic development and pluripotency regulation. In the mouse blastocyst, Netrin-1 contributes to the regulation of inner cell mass cell numbers (Huyghe et al., 2020). In mouse ESCs, activation of a Netrin-1–NEO1–UNC5B signaling axis sustains *Nanog* expression and promotes self-renewal in the undifferentiated state (Huyghe *et al*., 2020). Mechanistically, the balance between NEO1 and UNC5B receptors modulates key signaling pathways, including WNT and MAPK, through the regulation of GSK3α/β and ERK1/2 activity, thereby reinforcing pluripotency.

Whether Netrin-1 exerts a similar function in human PSCs, and more broadly how extracellular guidance cues contribute to the stabilization of human naïve pluripotency, remains unclear. In this study, we investigate the role of Netrin-1 in human naïve PSCs using complementary loss- and gain-of-function approaches. By combining single-cell transcriptomics, CUT&RUN profiling, and proteomic and phosphoproteomic analyses, we aim to uncover the molecular mechanisms by which Netrin-1 influences pluripotent cell identity and signaling states in humans.

## Results

### Dynamic Expression of Netrin Ligand and Receptor Families in Primate Embryos

To investigate the function of Netrins in controlling pluripotency in primates, we first established an overview of netrin and receptor gene family expression in human and non-human primate embryos by analyzing available single-cell RNA-sequencing datasets (Blakeley et al., 2015; Nakamura et al., 2016; Stirparo et al., 2018). In the cynomolgus monkey preimplantation embryo, we observed that *NTN1* and *NTNG1* are the only expressed ligand family members (**Fig. 1A**). *NTN1* is enriched in the epiblast (EPI) of early blastocysts (E6) but is low in other tissues, with only minimal expression in the EPI of mid- and late blastocysts (E7) (**Fig. 1B**). Notably, *NANOG*, *OCT4* and *SOX2* are expressed at higher levels in *NTN1^high^* cells than in *NTN1^low^* within the EPI (**Fig. 1C**). In contrast, *NTNG1* is expressed exclusively in the primitive endoderm (PE) (**Fig. 1A**). *NTN2* and *NTN3* were not identified in these datasets. Among receptor genes, only *UNC5B* and *NEO1* are expressed in cynomolgus monkey pre-implantation embryos, with maximum expression observed in the EPI of late blastocysts (**Fig. 1A, 1B**). Upon implantation, *NTN1* is rapidly downregulated in the EPI and remains unexpressed until E16-17, whereas *NEO*, *UNC5B* are expressed at higher levels in the post-implantation EPI than in the preimplantation EPI, and *UNC5D* expression is activated (**Fig. 1D**). Furthermore, *NTN4* and *UNC5D*, which were undetected in preimplantation embryos, are expressed in the post-implantation EPI. Collectively, these results reveal dynamic regulation of netrins and their receptors during pre- and early post-implantation development, and identify *NTN1* as a marker of EPI identity in the cynomolgus macaque preimplantation embryo.

**Figure 1:**
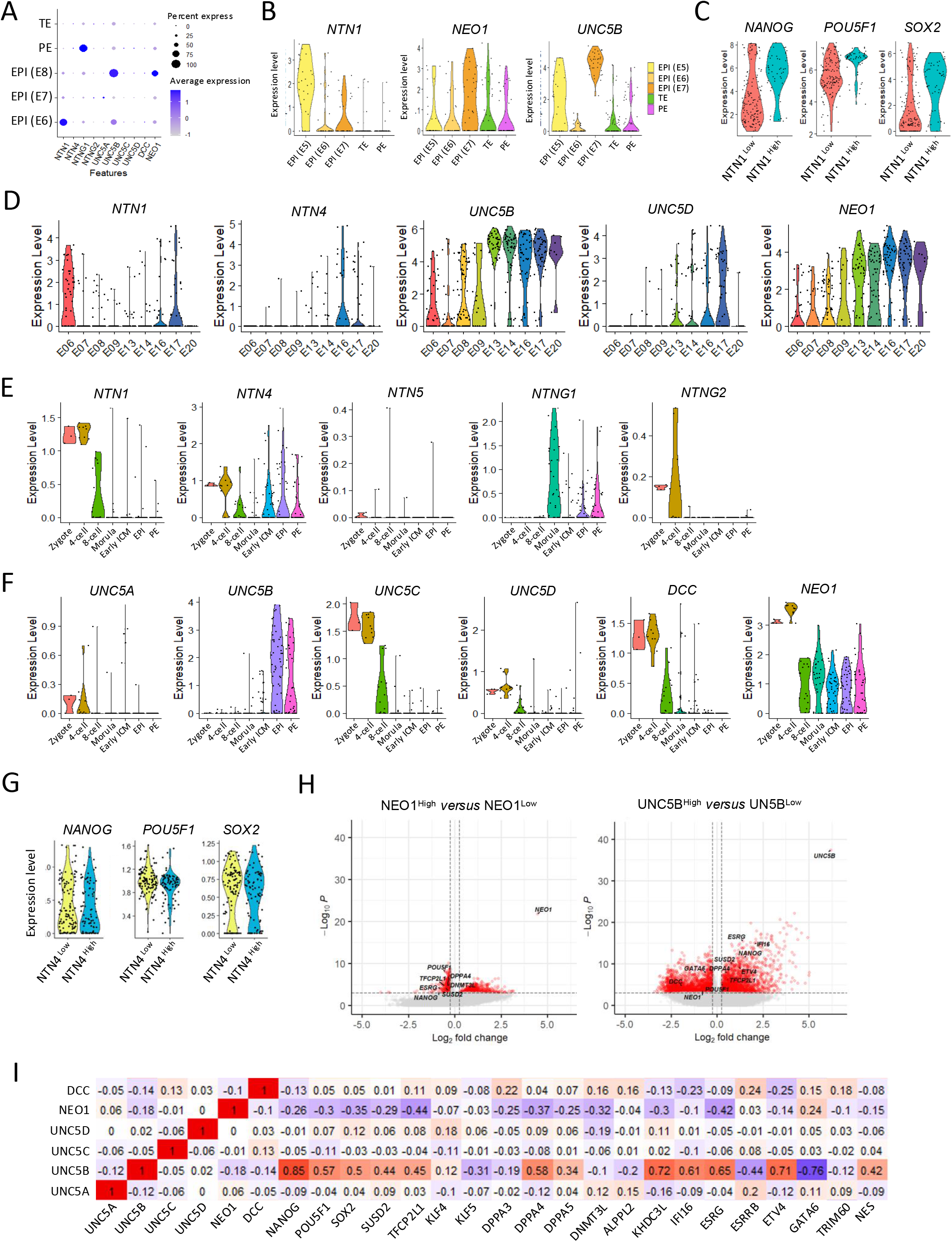
Expression of Netrin ligand and receptor families in primate embryos. (A) Dot representation of Netrins and receptor expression in cynomolgus monkey preimplantation embryos. TE, trophectoderm; PE, primitive endoderm; EPI, epiblast. (B) Violin plots showing *NTN1*, *NEO1* and *UNC5B* expression in cynomolgus monkey preimplantation embryos. (C) Violin plots showing *NANOG*, *POU5F1*, and *SOX2* expression in *NTN1^High^* and *NTN1^Low^* cells. (D) Violin plots showing Netrins and receptor expression in EPI cells in cynomolgus pre-5 (E6-E9) and post-implantation (E13-E20) embryos. (E) Expression of netrin family members in human preimplantation embryos. PE, primitive endoderm; EPI, epiblast. (F) Netrin receptor expression in human preimplantation embryos. PE, primitive endoderm; EPI, epiblast. (G) Violin plots showing *NANOG*, *POU5F1*, and *SOX2* expression in *NTN4^High^* and *NTN4^Low^* cells. (H) Volcan plots showing differentially expressed genes (DEGs) between NEO1^High^ and NEO1^Low^ cells (left panel) and between UNC5B^High^ and UNC5B^Low^ cells (right panel). (I) Heat map of the correlation coefficients among the indicated genes. **(A-D**) Original data are from Nakamura et al. 2017. (**E-I**) Original data are from Stirparo et al., 2018.

Analysis of the human preimplantation embryo revealed expression patterns of netrins and receptors that differ substantially from those observed in the cynomolgus monkey embryo. In particular, most *NTN1^High^* cells were detected during early cleavage stages, rarely at later stages, while *NTN4^High^* cells were observed at all stages and in all lineages including the EPI and trophectoderm (TE) (**Fig. 1E**). *NTN4* expression in the human EPI is not associated with elevated levels of *NANOG*, *OCT4* and *SOX2*, unlike the situation for *NTN1* in cynomolgus monkey embryos (**Fig. 1G**). Consistent with the cynomolgus monkey data, *UNC5B* and *NEO1* are expressed in most EPI and PE human cells, while the other receptors (*i.e. UNC5A*, *UNC5C*, *UNC5D*, and *DCC*) are only expressed at detectable levels during early cleavage stages (**Fig. 1F**).

To further characterize the *UNC5B^High^*, *UNC5B^Low^*, *NEO1^High^*, and *NEO1^Low^* cell populations within the human EPI, we analysed differentially-expressed genes (DEGs). Multiple genes associated with pluripotency–including *NANOG*, *POU5F1*, and SOX2, as well as markers of naïve pluripotency such as *TFCP2L1*, *SUSD2*, *DPPA4*, *DPPA5*, *KHDC3L*, *IFI16*, and *ESRG*, were upregulated in *UNC5B^High^* cells compared to *UNC5B^Low^* cells (**Fig. 1H**). Their expression levels showed a positive correlation with *UNC5B* expression (**Fig. 1I**, **Supplementary 1A-C**). In contrast, although *NEO1^High^* cells are present within the EPI compartment, they expressed *NANOG*, *POU5F1*, *TFCP2L1*, *DPPA4*, and *ESSRG* at lower levels than *NEO1^Low^* cells, with expression levels negatively correlated with NEO1. Altogether, these results suggests that *UNC5B* expression in the human preimplantation embryo is associated with naïve pluripotency and likely characterizes the early EPI.

### *NTN1* and *UNC5B* Expression is Associated with Naïve Pluripotency in Human PSCs

We examined the expression of Netrin ligands and their receptors in the human ESC line Oscar (OS) (Chen et al., 2015) using single-cell RNA sequencing following reprogramming to the naive state in PXGL medium (OS-PXGL) (Guo *et al*., 2017; Guo *et al*., 2016). Naïve OS cells expressed *NTN1* at variable levels (**Fig. 2A,B**). Notably, the 10% cells showing the highest *NTN1* expression (*NTN^High^*) exhibited elevated levels of naïve pluripotency markers–including *TFCP2L1*, *KLF4*, *KLF5*, *DPPA3*, *DPPA5*, *DNMT3L*, *ALPG*, *KHDC3L*, *TRIM60, SUSD2*, *FGF4*, *TCF3*, and *NLRP7*–compared to the 10% cells showing the lowest *NTN1* expression (*NTN1^Low^*) (**Fig. 2E**).

**Figure 2:**
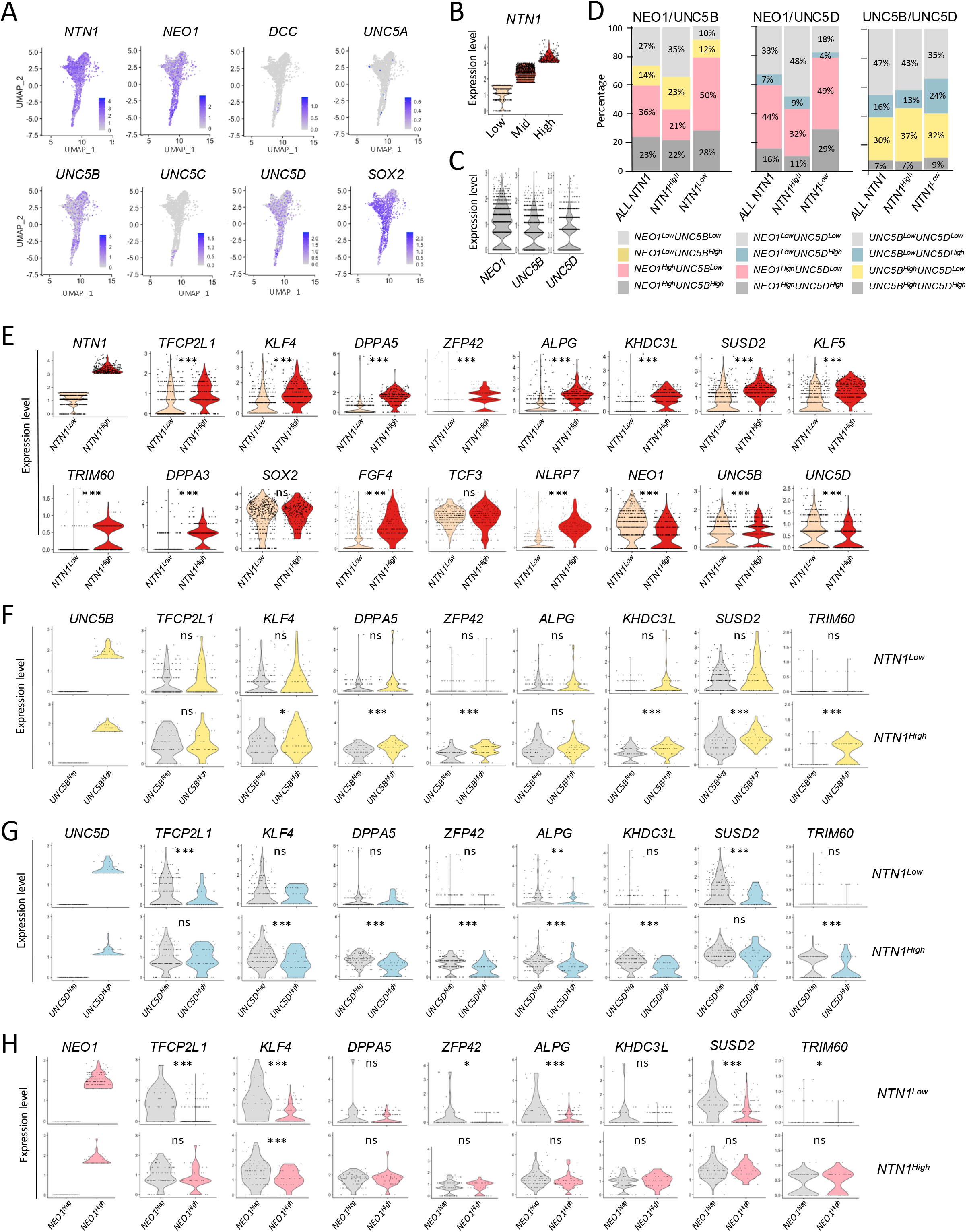
Expression of Netrin ligand and receptor families in human OS-PXGL. (A) UMAP representation of *NTN1* and receptor expression. (B) Violin plots showing *NTN1* expression in *NTN1^High^*, *NTN1^Mid^,* and *NTN1^Low^* cells. (C) Violin plots showing *NEO1*, *UNC5B*, and *UNC5D* expression. (D) Histogram illustrating percentages of cells expressing combinations of NEO1 and UNC5B receptors among all NTN1, *NTN1^High^* and *NTN1^Low^* cells. (E) Violin plots showing naïve pluripotency markers in *NTN1^High^* and *NTN1^Low^* cells. (F) Violin plots showing naïve pluripotency markers in *UNC5B^Neg^-NTN1^Low^*, *UNC5B^High^-NTN1^Low^*, *UNC5B^Neg^-NTN1^High^*, and *UNC5B^High^-NTN1^High^*cells. (G) Violin plots showing naïve pluripotency markers in *UNC5D^Neg^-NTN1^Low^*, *UNC5D^High^-NTN1^Low^*, *UNC5D^Neg^-NTN1^High^*, and *UNC5D^High^-NTN1^High^*cells. (H) Violin plots showing naïve pluripotency markers in *NEO1^Neg^-NTN1^Low^*, *NEO1^High^-NTN1^Low^*, *NEO1^Neg^-NTN1^High^*, and *NEO1^High^-NTN1^High^*cells. (**E-H**) Two-sided Student’s t-test (ns, non-significant; *, p < 0.05; **, p < 0.01; ***, p < 0.001).

OS-PXGL cells displayed heterogeneous expression of the receptor genes *NEO1*, *UNC5B*, and *UNC5D*, while *DCC*, *UNC5A*, and *UNC5C* were not detected (**Fig. 2A,C**). Focusing on *NEO1* and *UNC5B*, we identified four subpopulations: *NEO1^High^/UNC5B^High^*(24%), *NEO1^Low^/UNC5B^High^* (14%), *NEO1^High^/UNC5B^Low^* (36%), *NEO1^Low^/UNC5B^Low^* (27%) (**Fig. 2D**). These proportions were largely maintained when considering *NEO1* and *UNC5D* instead. Notably, the *NEO1^Low^*subpopulation was more abundant among *NTN^High^* cells compared to *NTN1^Low^* cells, consistent with the downregulation of *NEO1* expression observed in *NTN1^High^* relative to *NTN1^Low^* cells (**Fig. 2E**). Most cells expressed either *UNC5B* or *UNC5D*, with fewer than 10% expressing both, regardless of *NTN1* expression status (**Fig. 2D**). These results highlight the heterogenous and dynamics expression of Netrin-1 receptors within the naïve OS cell population.

To evaluate the responsiveness of OS-PXGL cells based on their receptor profiles, we analysed naïve pluripotency marker expression across 12 subpopulations, defined by combinations of *NEO1*, *UNC5B*, and *UNC5D* expression: *NEO1^High^/UNC5B^High^*, *NEO1^Low^/UNC5B^High^*, *NEO1^High^/UNC5B^Low^*, *NEO1^Low^/UNC5B^Low^*, *NEO1^High^/UNC5D^High^*, *NEO1^Low^/UNC5D^High^*, *NEO1^High^/UNC5D^Low^*, *NEO1^Low^/UNC5D^Low^*, *UNC5B^High^/UNC5D^High^*, *UNC5B^Low^/UNC5D^High^*, *UNC5B^High^/UNC5D^Low^*, and *UNC5B^Low^/UNC5D^Low^*. In all subpopulations, *NTN1^High^* cells exhibited increased expression of naïve pluripotency markers compared to *NTN1^Low^* cells (**Supplementary Fig. 2A-C**). This increase was less marked in *NEO1^Low^* subpopulations, where it was restricted to cells expressing low baseline levels of certain naïve markers–such as *KLF4*, *KLF5*, and *SUSD2*–resulting in reduced heterogeneity within these groups. These findings suggest that *NTN1* enhances naïve marker expression across all subpopulations, albeit to varying extents.

To further characterize the influence of each receptor on naïve marker expression in response to Netrin-1, we profiled the 10% of cells expressing the highest level of receptor (referred to as *UNC5B^High^*, *UNC5D^High^*, and *NEO1^High^* cells) versus cells where receptors were not detected (referred to *UNC5B^Neg^, UNC5D^Neg^*, and *NEO^Neg^*) (**Fig. 2F,G**). Comparison of *UNC5B^Neg^*and *UNC5B^High^* cells revealed no significant difference in naïve marker expression within the *NTN1^Low^* population (**Fig. 2F**). In contrast, moderate upregulation of *KLF4*, *DPPA5*, *ALPG*, *KHDC3L*, *SUSD2*, and *TRIM60* was observed in *NTN^High^/UNC5B^High^* compared to *NTN1^High^/UNC5B^Low^*cells. This suggests that the UNC5B receptor enhances naïve marker expression in the presence of Netrin-1. Transcriptomic comparison of *UNC5D^High^*and *UNC5D^Neg^* cells highlighted opposite effects, showing moderate downregulation of all naïve markers examined in both *NTN1^Neg^*and *NTN1^High^* cells. Transcriptomic comparison of *NEO1^High^*and *NEO1^Neg^* cells revealed distinct profiles depending on *NTN1* status. Within the *NTN1^Low^* subpopulation, naïve marker expression was significantly lower in *NEO1^High^* compared to *NEO1^Neg^* cells. In contrast, in the *NTN1^High^* subpopulation, *NEO1^High^* cells showed elevated naïve marker expression, typically reaching levels similar to those of *NEO1^Neg^* cells. These findings suggest that NEO1 acts as an inhibitor of naive pluripotency marker expression, and that Netrin-1 counteracts this inhibitory effect. Collectively, these results indicate that *UNC5B*, *UNC5D*, and *NEO1* exert antagonistic influences on naive OS cells, modulated by *NTN1* expression, with *UNC5B* having an enhancing effect, and *UNC5C* and *NEO1* impeding these effects.

### Self-renewal of *NTN1-KO* OS-PXGL cells

To investigate the influence of Netrin-1 on naïve pluripotency, *NTN1* was disrupted in OS-PXGL cells. Three clones harboring homozygous indels in the *NTN1* coding sequence were isolated: clone *#1* harbors a bi-allelic frameshift deletion of 23 bp; clone *#2* harbors a 1 bp frameshift insertion on one allele and a 16 bp frameshift deletion on the second allele; and clone *#17* harbors two bi-allelic frameshift deletions of 52 and 11 bp, respectively (**Supplementary Fig. 3A,B**). In these three *NTN1^-/-^* clones, Netrin-1 protein abundance was dramatically reduced (**Fig. 3A**). Three control–NTN1^+/+^–clones, designated *#5, #6,* and *#12*, were isolated from the same CRISPR/Cas9-treated cell population. The six clones were undistinguishable under routine culture conditions. They formed dome-shaped colonies (**Fig. 3B, Supplementary Fig. 3C)** and were regularly passaged every three days at a 1:5 split ratio. These findings indicate that *NTN1* disruption had no visible effects on colony morphology and cell growth under these culture conditions.

**Figure 3:**
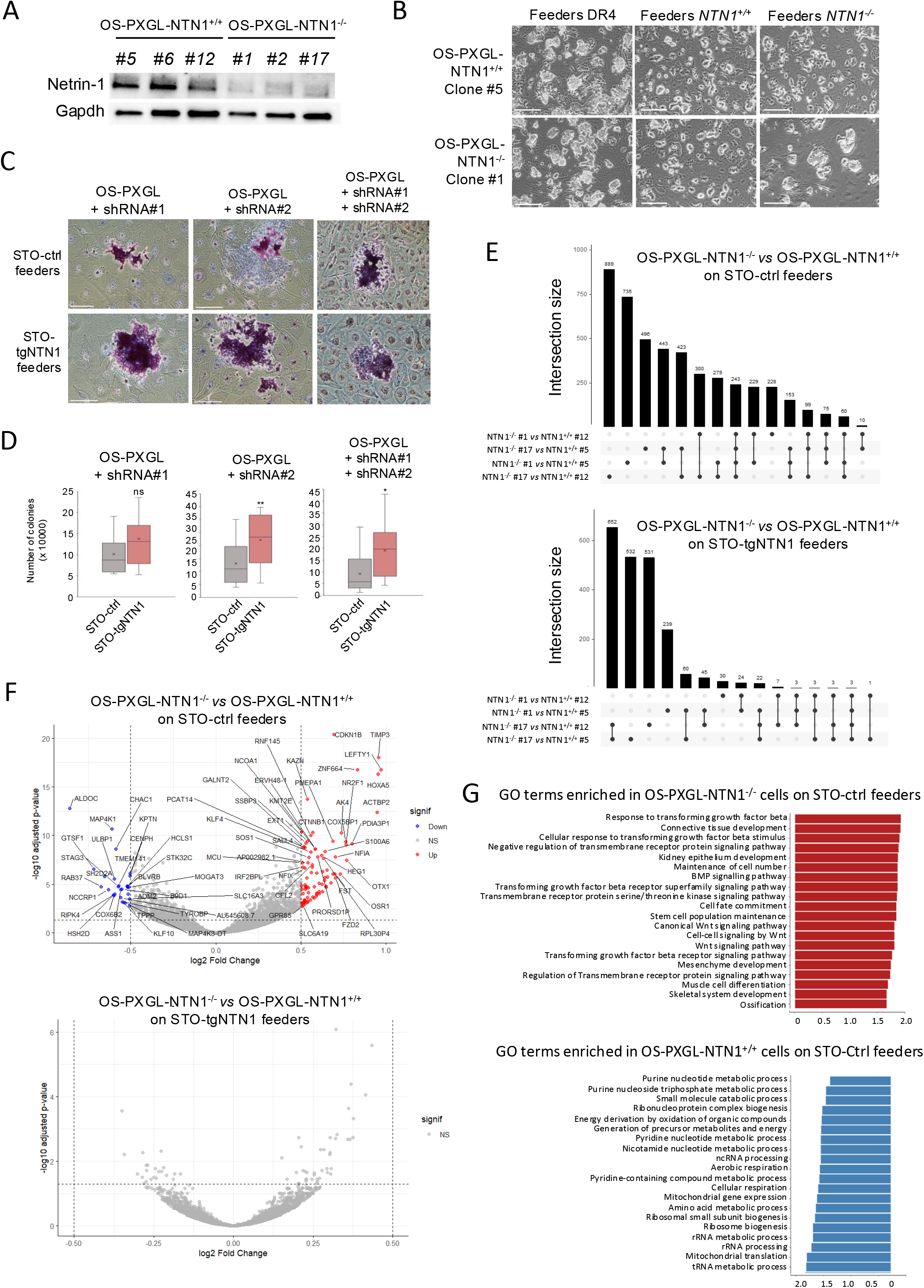
Characterization of *NTN1* knockout OS-PXGL cells. (A) Western blot analysis of Netrin-1 in OS-PXGL-NTN1^+/+^ and OS-PXGL-NTN1^-/-^ clones. (B) Phase contrast images of OS-PXGL-NTN1^+/+^ (clone #5) and OS-PXGL-NTN^-/-^ (clone #1) cultured on DR4 feeders, NTN1^+/+^ and NTN1^-/-^ C57bl/6 feeders. Scale bars, 200 µm. (C) Colony-forming assay with OS-PXGL cells transfected with Cas9 and sgRNAs prior to culturing on STO-Ctrl and STO-tgNTN1 feeders, and stained to reveal alkaline phosphatase (AP) activity. Scale bars, 200 µm. (D) Histograms of total colony numbers obtained with OS-PXGL cells transfected with Cas9 and sgRNAs and cultured on STO-Ctrl and STO-tgNTN1 feeders. (E) UpSet plots showing the overlap of differentially expressed genes across pairwise comparisons of independent NTN1⁻/⁻ and NTN1⁺/⁺ OS-PXGL clones cultured on (top) control feeders (STO-ctrl) or (bottom) NTN1-expressing feeders (STO-tgNTN1). Vertical bars indicate the size of each intersection (number of shared genes), while connected dots below denote the specific comparisons included in each intersection. Comparisons involve independent clone pairs (NTN1⁻/⁻ #1, #17 *versus* NTN1⁺/⁺ #5, #12). The largest intersections represent gene expression signatures consistently shared across multiple comparisons, whereas smaller intersections reflect clone-specific or condition-specific effects. (F) Volcano plots showing differentially expressed genes (DEGs) between OS-PGXL-NTN1^+/+^ (clones #5 and #12) and OS-PGXL-NTN^-/-^ (clone #1 and #17) cultured on STO-ctrl (top panel) and STO-tgNTN1 (bottom panel) feeders. (G) GO term enrichment analysis of the DEGs between OS-PGXL-NTN1^+/+^ (clones #5 and #12) and OS-PGXL-NTN^-/-^ (clone #1 and #17) cultured on STO-ctrl (top panel) and STO-tgNTN1 feeders.

Since feeder cells produce Netrin-1 (Huyghe *et al*., 2020), we asked if this Netrin-1 can rescue the growth and self-renewal capacities of Netrin-1-deficient OS-PXGL. To address this question, control and *NTN1^-/-^* clones were cultured on growth-inactivated mouse embryonic fibroblasts prepared from C57bl/6 *NTN1^+/+^* or *NTN1^-/-^* mouse fetuses (**Supplementary Fig. 3D**). Again, the six clones were undistinguishable, forming dome-shaped colonies and sustaining regular passaging for at least four weeks (**Fig. 3B, Supplementary Fig. 3E**). A transcriptome analysis confirmed that *NTN1^+/+^*and *NTN1^-/-^* clones were indistinguishable in a PCA analysis (**Supplementary Fig. 3F**). These findings indicate that Netrin-1 spontaneously expressed by feeder cells has no visible effect on the self-renewal capacity of OS cells when cultured in PXGL medium using growth-inactivated mouse embryonic fibroblasts as feeder cells.

### *NTN1-KO* OS-PXGL Exhibit Eroded Naïve Characteristics

To investigate the paracrine contribution of Netrin-1 to naïve pluripotency further, we treated OS-PXGL cells with Cas9 and sgRNAs prior to culturing them on STO feeders (STO-ctrl) and STO feeders overexpressing an HA-tagged human Netrin-1 (STO-tgNTN1) (**Supplementary Fig. 3G**). OS-PXGL cells were significantly more clonogenic and formed larger colonies when cultured on STO-tgNTN1 compared with STO-ctrl, suggesting enhanced viability, increased growth, or both (**Fig. 3C,D**). To assess transcriptional changes associated with the loss of Netrin-1, naïve OS-PXGL cells were cultured for 10 passages on STO-ctrl or STO-tgNTN1 feeders (**Supplementary Fig. 3H**), and RNA sequencing was performed on two NTN1^+/+^ clones (#5 and #12) and two NTN1^−/−^ clones (#1 and #17). On STO-ctrl feeders, differential expression analysis identified a set of 243 genes consistently altered across all four pairwise comparisons–i.e. *NTN1^-/-^*#1 and #17 *versus NTN1^+/+^* #5 and #12–indicating a robust *NTN1*-dependent transcriptional signature (**Fig. 3E,F; Supplementary Table S1**). A Gene Ontology (GO) term analysis of genes overexpressed in both *NTN1^-/-^* clones versus both *NTN1^+/+^* clones indicate a strong shift toward developmental and differentiation-associated signaling programs, particularly involving TGF-β/BMP and WNT pathways (**Fig. 3G**). These changes are coupled to processes related to mesenchymal, skeletal, muscle, and connective tissue development, suggesting enhanced lineage commitment and differentiation potential. Concurrently, terms such as “stem cell population maintenance”, “maintenance of cell number”, and “regulation of receptor signaling” point to alterations in the balance between self-renewal and differentiation, consistent with a reconfiguration of signaling networks that govern pluripotency and cell identity. Analysis at gene level refined these conclusions: genes upregulated in *NTN1^-/-^* cells (161) were enriched for regulators of developmental patterning and morphogen signaling, including transcription factors such as *PRRX1*, *HOXA5*, *DLX1*, *FOXG1*, *NFIA*, *NR2F1*, *NR2F2*, *PRRX1*, *ZFHX4, TSHZ1*, *OTX1,* and *OSR1*, as well as signaling modulators such as *FZD2*, *LEFTY1*, *FST*, and *GAS1*. These changes suggest increased responsiveness to WNT, TGF-β/NODAL, and Hedgehog pathways and are consistent with transcriptional priming toward developmental programs. In parallel, genes associated with extracellular matrix organization and cell adhesion, including *COL1A1*, *SDC2*, *HEG1*, and *TIMP3*, were elevated in *NTN1^-/-^*cells, suggesting remodeling of cell–matrix interactions. Differential expression analysis identified an additional set of 288 genes altered across three out of four pairwise comparisons (**Fig. 3E**; **Supplementary Fig. 3I**; **Supplementary Table S1**). This lower-stringency dataset expanded the “extracellular matrix organization and cell adhesion” signature to include multiple collagen genes (*COL1A2*, *COL3A1*, *COL5A1*, *COL8A1*), as well as *FN1*, *CRIM1*, and *AMOT*.

The GO terms enriched in OS-PXGL-NTN1^+/+^ cells are dominated by core metabolic and biosynthetic processes, including nucleotide and amino acid metabolism, mitochondrial respiration, and energy production, indicating a metabolically active state with robust oxidative phosphorylation (**Fig. 3G**). In parallel, strong enrichment for ribosome biogenesis, rRNA/tRNA processing, and mitochondrial gene expression/translation reflects elevated protein synthesis capacity and translational activity. Together, this profile is consistent with cells maintaining a highly active, growth-supportive state, favoring biosynthesis and energy metabolism rather than differentiation-associated signaling. Genes reduced in *NTN1^-/-^* cells (82) were strongly enriched for serine/glycine, amino-acid stress, and redox-defense genes: *PHGDH*, *PSAT1*, *SHMT2*, *SLC7A11*, *SLC1A5*, *SLC7A5*, *CHAC1*, *NUPR1*, *EIF4EBP1*, *TSPO*, *ALDH1L2*, *PYCR1*, and *ASS1*. Additional KO-down genes identified in the lower-stringency dataset included *ATF4*, *PSPH*, *PFKFB3*, *PCK2*, *IGFBP2*, likely reinforcing the downregulation of *PHGDH*, *PSAT1*, *SLC7A11*, *SLC1A5*, *SLC7A5*, *CHAC1*. *ATF4* is a central regulator of cellular stress adaptation, especially amino-acid response and related metabolic rewiring (Oomen et al., 2025). Notably, *KLF17*, one of the hallmark markers of human naive pluripotency was down-regulated in the lower-stringency dataset. Together, these results indicate that loss of Netrin-1 in OS-PXGL cells cultured on STO feeders is associated with coordinated changes in transcriptional programs, extracellular signaling, and metabolic state, linked to early developmental/ differentiation competence.

In sharp contrast, differential expression analysis of OS-PXGL cultured on STO-tgNTN1 revealed a much narrower gap between NTN1^+/+^ and NTN1^-/-^ cells, with only 3 DEGs consistently altered across all four pairwise comparisons, and 35 altered across three out of four pairwise comparisons (**Fig. 3E,F; Supplementary Fig. 3I** and **Table S1**). These results indicate that the erosion of naïve identity observed in NTN1^-/-^ OS-PXGL cells can be almost completely rescued by feeder cells overexpressing Netrin-1, highlighting the paracrine role of overexpressed Netrin-1 in our system.

### Stabilization of Naïve Pluripotency by Netrin-1 Overexpression

To assess the role of Netrin-1 in reprogramming human PSCs to the naïve state, we engineered OS cells to overexpress HA-tagged *NTN1* cDNA (OS-tgNTN1^WT^) (**Fig. 4A**). After 16 passages (P16) in PXGL, OS-tgNTN1^WT^ cells predominantly formed dome-shaped colonies characteristic of naïve pluripotency, whereas control cells retained a flat morphology. These morphological differences gradually diminished and became nearly undetectable by passage 31 (**Supplementary Fig. 4A**), suggesting that Netrin-1 overexpression accelerates PXGL-induced reprogramming.

**Figure 4:**
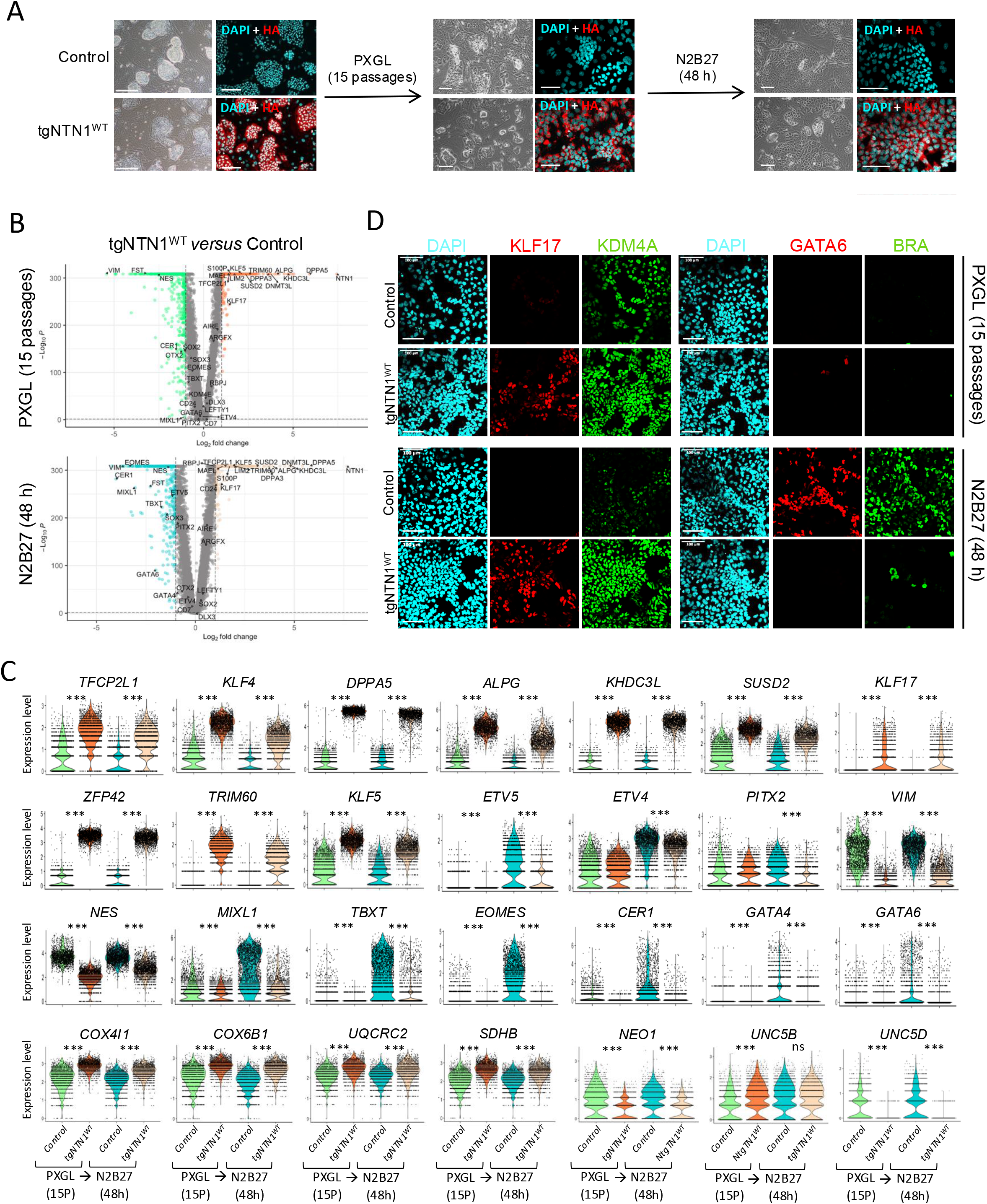
Characterization of OS-tgNTN1^WT^ cells. (A) Phase-contrast and confocal images of control and tgNTN1^WT^ cells cultured in PXGL for 15 passages, followed by N2B27 for 48 h. Immunostaining shows HA:NTN1^WT^ expression. Scale bars, 100 µm. (B) Volcano plot showing DEGs between OS-tgNTN1^WT^ and control cells under PXGL (15 passages) and N2B27 (48 h) conditions. Orange and green dots indicate upregulated and downregulated genes, respectively, in OS-tgNTN1^WT^ cells compared with controls. (C) Violin plots showing naïve, formative, and primed pluripotency markers, lineage markers, and Netrin-1 receptors in control and OS-tgNTN1^WT^ cells. Two-sided Student’s t-test (ns, non-significant; *, p < 0.05; **, p < 0.01; ***, p < 0.001). (D) Confocal images of control and OS-tgNTN1^WT^ cells stained for KLF17, KDM4A, GATA6, and BRACHYURY (BRA). Scale bars, 100 µm.

OS-tgNTN1^WT^ and control cells were collected at P16 and analyzed by single-cell RNA sequencing. Compared with control, OS-tgNTN1^WT^ cells displayed elevated expression of naïve pluripotency markers–including *TFCP2L1*, *KLF4*, *DPPA5*, *ALPG*, *KHDC3L*, *SUSD2*, *ZFP42, TRIM60,* and *KLF5*–and reduced expression of formative/primed markers such as *ETV4, ETV5,* and *PITX2* (**Fig. 4B,C; Supplementary Table S2**). Immunostaining confirmed increased expression of naïve markers KDM4A and KLF17 in OS-tgNTN1^WT^ cells cultured in PXGL for 15 days (**Fig. 4D**). Moreover, a GO term analysis of DEGs identified between OS-tgNTN1^WT^ and control cells showed enrichment in “oxidative phosphorylation”, “aerobic respiration”, and “mitochondrial respiratory chain complex assembly” terms, and an impoverishment in “cell growth”, “gastrulation”, “Wnt signaling”, and “hippo signaling” terms, all of which are associated with naïve pluripotency (**Supplementary Fig. 4B**). Furthermore, colony-forming assays further demonstrated significant increase in both colony number and size with tgNTN1^WT^ cells, indicating that Netrin-1 enhances proliferative capacity as well as clonogenicity (**Supplementary Fig. 4C-E**). Collectively, these findings show that Netrin-1 overexpression facilitates reprogramming of OS cells to a naïve pluripotent state.

In a parallel experiment, OS-tgNTN1^WT^ and control cells at P16 were transferred to LIF-and kinase inhibitor-free medium for 48 hours [N2B27 (48hr)] and subjected to single-cell RNA-seq. In control cells, this transition triggered morphological changes (**Fig. 4A**) accompanied by sharp downregulation of naïve markers (*TFCP2L1*, *KLF4*, *KLF5*, *ALPG*, *SUSD2*) and upregulation of formative/primed markers (*ETV4*, *ETV5*, *PITX2*) (**Fig. 4B,C; Supplementary Table S2**). By contrast, some OS-tgNTN1^WT^ cells retained an undifferentiated morphology, maintained high expression of naïve markers, and showed markedly attenuated induction of formative/primed genes. Notably, activation of early lineage markers–including *MIXL1*, *TBXT*, *EOMES*, *GATA4*, and *GATA6*–seen in control cells was abolished in OS-tgNTN1^WT^ cells. Immunofluorescence for GATA6 and BRACHYURY confirmed substantially reduced expression in OS-tgNTN1^WT^ cells compared with controls under N2B27 conditions (**Fig. 4D**). These results indicate that Netrin-1 overexpression protects OS cells from differentiation induced by LIF and inhibitor withdrawal.

### Epigenome Reconfiguration Induced by Netrin-1 Overexpression

To gain further insight into the role of Netrin-1 during naïve-state reprogramming, we profiled the histone marks H3K4me3, H3K9me3, H3K27me3, and H3K27ac in OS-tgNTN1^WT^ and control cells at passages 16 (P16) and 40 (P40) following transfer to PXGL conditions using CUT&RUN. At P15/P16, OS-tgNTN1^WT^ cells displayed a marked increase in H3K27ac, reflected by a substantially greater number of significant peaks across the genome, indicative of enhanced enhancer and promoter activity and a globally more accessible chromatin landscape (**Fig. 5A; Supplementary Fig. S5**). H3K27me3 levels were also elevated, although to a lesser extent, suggesting reinforced repression of differentiation-associated loci. In contrast, H3K4me3 exhibited both gains and losses at distinct genomic regions, consistent with a reorganization of promoter activity rather than a uniform increase in transcriptional output. No global changes were detected in H3K9me3 distribution, indicating preservation of constitutive heterochromatin. By P40, differences in H3K27ac enrichment between OS-tgNTN1^WT^ and control cells had largely disappeared, while alterations in H3K27me3 and H3K4me3 were markedly attenuated. Together, these findings suggest that tgNTN1^WT^ accelerates the naïve-like epigenetic remodeling induced by the transition of OS cells to PXGL conditions, thereby promoting earlier acquisition of a naïve-like chromatin state.

**Figure 5:**
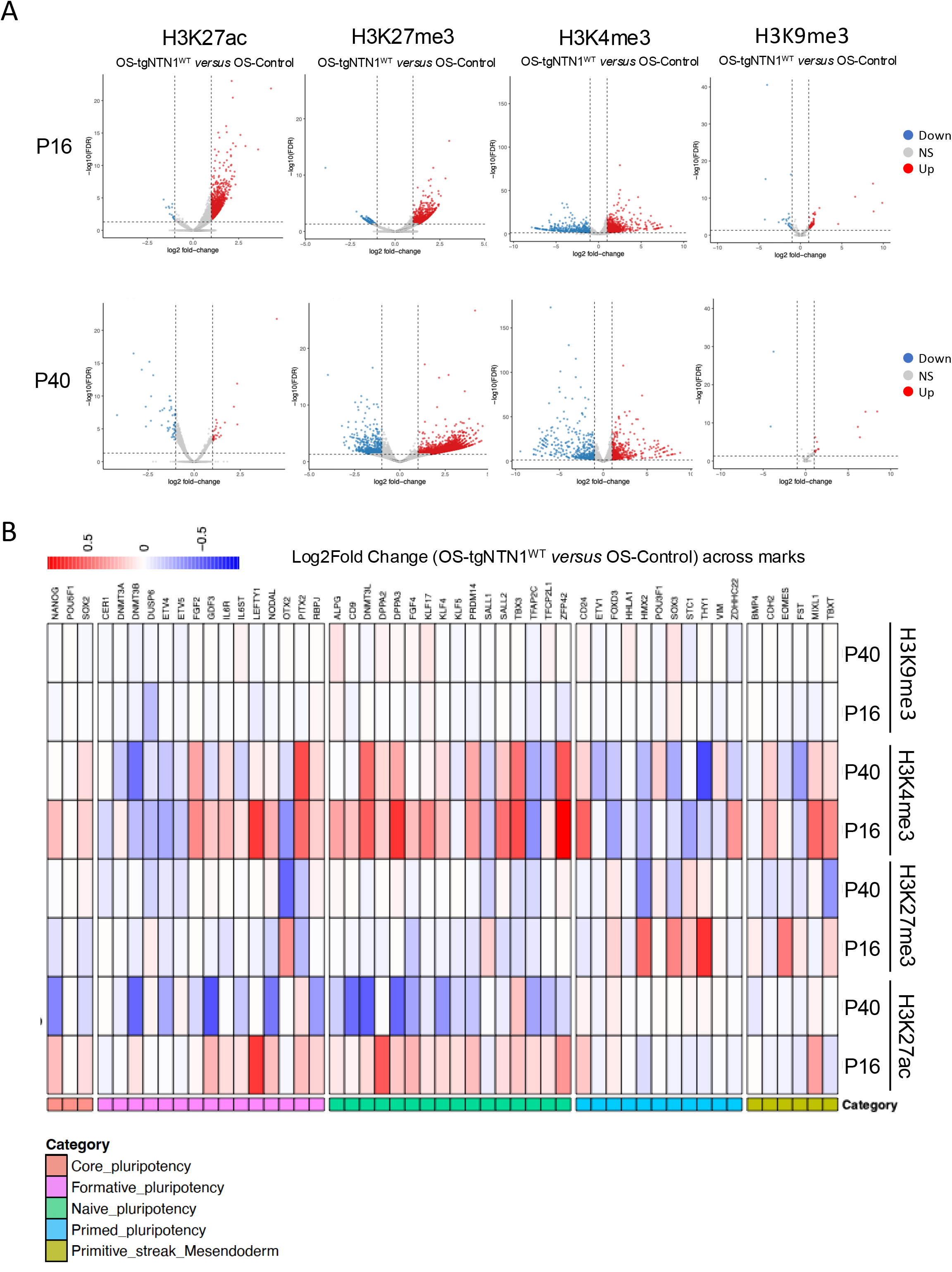
Epigenomic characterization of OS-tgNTN1^WT^ cells by CUT&RUN analysis. (A) Volcano plots showing differentially enriched DNA fragments in OS-tgNTN1^WT^ relative to control cells cultured under PXGL conditions, for the indicated histone marks at passages P16 and P40. (B) Heatmap showing differential enrichment of the indicated histone marks at selected genomic loci in OS-tgNTN1^WT^ relative to control cells cultured under PXGL conditions, at passages P16/P17 and P39/P40.

Examination of the histone marks H3K4me3, H3K9me3, H3K27me3, and H3K27ac at selected pluripotency-associated loci confirmed and extended these observations. At P16, H3K27ac and H3K4me3 were strongly enriched at naïve and formative pluripotency genes in OS-tgNTN1^WT^ cells compared with control cells, consistent with their increased transcriptional activity (**Fig. 5B**). In parallel, the repressive mark H3K27me3 was enriched at primed pluripotency and primitive streak genes. By contrast, at P40, H3K27ac enrichment at naïve pluripotency loci was reduced in OS-tgNTN1^WT^ relative to control cells, suggesting that control cells had caught up with, or even surpassed, OS-tgNTN1^WT^ cells in their reprogramming progression. H3K4me3 displayed a more heterogeneous pattern, with both gains and losses observed in OS-tgNTN1^WT^ relative to control cells across the loci examined, irrespective of their association with naïve or primed pluripotency. H3K9me3 remained essentially unaltered both at P16 and P40. Together, these findings indicate that Netrin-1 overexpression actively modulates histone post-translational modifications at K4 and K27, thereby facilitating the naïve-state reprogramming induced by LIF and small-molecule treatment.

### Stabilization of Naïve Pluripotency by NEO1 and UNC5H Binding-Deficient Netrin-1

To identify the receptors that might mediate the effects of Netrin-1, we analyzed the single-cell transcriptome of OS-tgNTN1^WT^ and control cells according to their expression of the receptors *NEO1* and *UNC5B* under both PXGL (P15) and N2B27 (48h) conditions. No major differences in naïve, primed, or early lineage marker expression were observed between *NEO1^High^* and *NEO^Neg^* OS-tgNTN1^WT^ cells (**Supplementary Fig. 6A**). In contrast, *UNC5B^High^* cells displayed a moderate increase in the expression of some naïve markers together with a modest decrease in some primed and lineage markers compared with *UNC5B^Neg^* cells (**Supplementary Fig. 6B**). Collectively, although UNC5B expression is associated with modest differences in pluripotency marker expression, these findings suggest that the enhancement of naïve pluripotency induced by Netrin-1 overexpression does not critically depend on the expression levels of either *NEO1* or *UNC5B*.

To get some further insight into the receptors that mediate the effects of tgNTN1^wt^, we engineered OS cells overexpressing Netrin-1 mutated on residues critical for its interaction with NEO1 (NEO1-mut), UNC5H (UNC5H-mut), or both (N-mut/U-mut) (Grandin *et al*., 2016; Xu *et al*., 2014). OS-tgNTN1^WT^, OS-tgNTN1^UNC5H-mut^, OS-tgNTN1^NEO1-mut^, and OS-tgNTN1^N-mut/U-mut^ cells expressed Netrin-1 at similar levels in Western blot and immunofluorescence analyses (**Fig. 6A**, **Supplementary Fig. 6C**). After being reprogrammed in PXGL for 15 passages, transfected cells were analyzed in a clonal assay along with the OS-tgNTN1^WT^ and control cells previously described. This revealed a substantial increase in colony scores in all four Netrin-1-expressing lines compared with the control (**Fig. 6B**). All cell lines were then analyzed by single-cell RNA-seq. A UMAP representation of the global dataset shows that all tgNTN1 cells form a distinct cluster, with most tgNTN1^UNC5H-mut^ and tgNTN1^U-mut/N-mut^ cells forming a subcluster and tgNTN1^WT^ and tgNTN1^NEO1-mut^ cells forming another (**Fig. 6C**). Notably, a small subpopulation of control cells (0.02%) clustered with tgNTN1 cells, suggesting that rare OS cells can spontaneously adopt a phenotype characteristic of OS-tgNTN1 cells. An analysis of DEGs revealed the same higher expression of naïve pluripotency markers–including *DPPA5*, *KHDC3L*, *KLF4*, *KLF5*, *ALPG*, *KHDC3L*, *TFCP2L1*, *ALPG*, *ZFP42*, *SUSD2*, *TRIM60*, *KDM4E*, *S100P*, and *LIM2*–and lower expression of formative/primed pluripotency and early lineage markers such as *ETV4*, *LEFTY1, SOX3, PITX2*, *VIM, NES, FST, MIXL1, TBXT,* and *GATA6* in OS-tgNTN1^WT^, OS-tgNTN1^UNC5H-mut^, OS-tgNTN1^NEO1-mut^, and OS-tgNTN1^U-mut/N-mut^ cells (**Fig. 6D**; **Supplementary Fig. 6D,E; Supplementary Table S2**). A similar transcriptomic analysis was performed with all five cell types after withdrawal of LIF and kinase inhibitors for 48 hours (N2B27 48h), revealing strong similarities in the patterns of naïve and primed marker gene expression between OS-tgNTN1^WT^, OS-tgNTN1^UNC5H-mut^, OS-tgNTN1^NEO1-mut^, and OS-tgNTN1^U-mut/N-mut^ compared to control OS cells (**Fig. 6D**; **Supplementary Fig. 6D,E**; **Supplementary Table S2**). Altogether, these results indicate that the ability of Netrin-1 to enhance naïve pluripotency when overexpressed does not critically depend on binding to UNC5B and NEO1 receptors.

**Figure 6:**
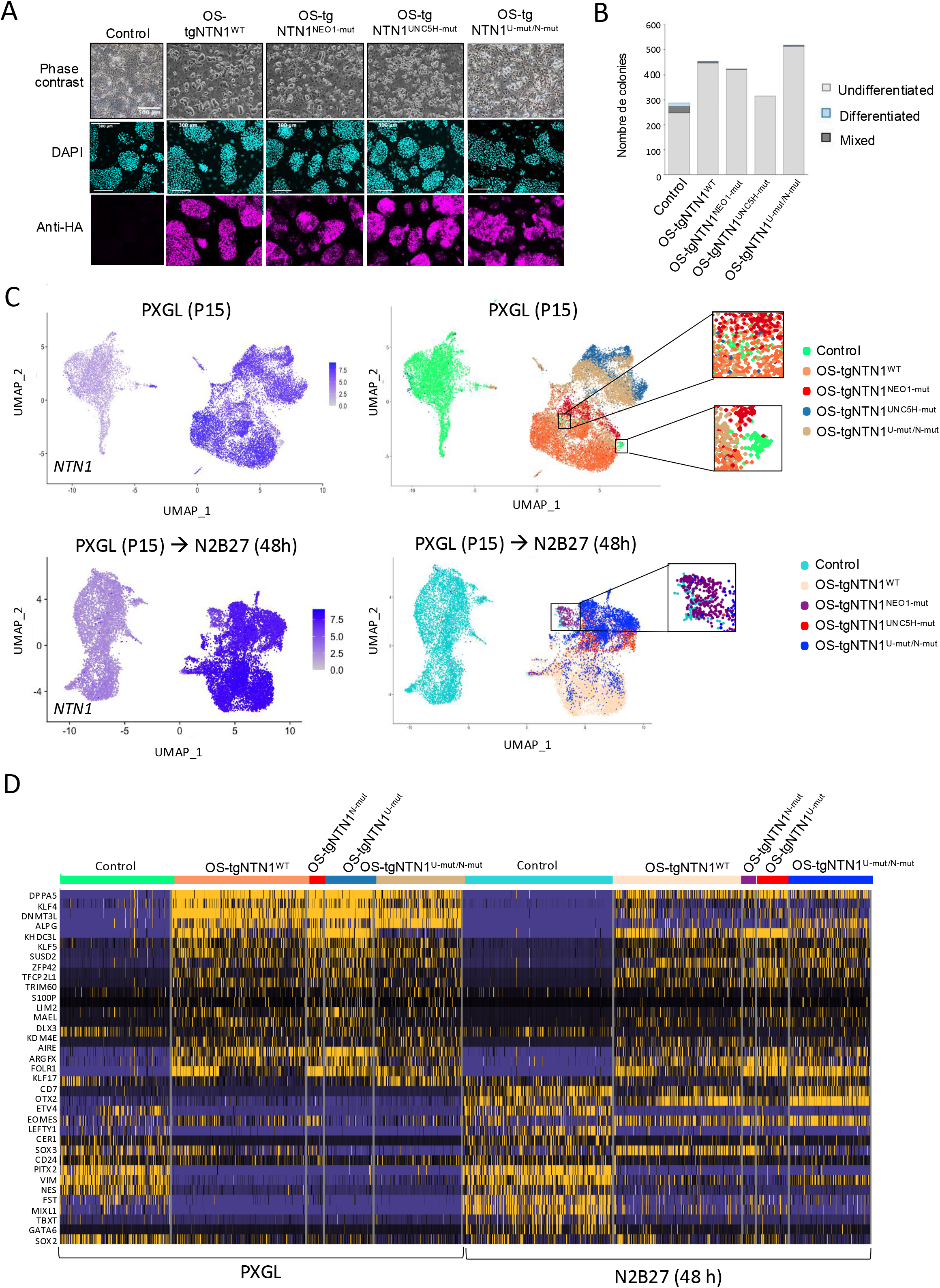
Characterization of OS-tgNTN1^UNC5H-mut^, OS-tgNTN1^NEO1-mut^, and OS-tgNTN1^U-mut/N-mut^ cells. (A) Phase-contrast and confocal images of Control, OS-tgNTN1^WT^, OS-tgNTN1^UNC5H-mut^, OS-tgNTN1^NEO1-mut^, and OS-tgNTN1^U-mut/N-mut^ cells. Immunostaining shows HA:NTN1 expression. Scale bars, 200 µm. (B) Histogram of colony counting with Control, OS-tgNTN1WT, OS-tgNTN1^UNC5H-mut^, OS-tgNTN1^NEO1-mut^, and OS-tgNTN1^U-mut/N-mut^ cells. (C) Left panels: UMAP representation of NTN1 expression in Control, OS-tgNTN1WT, OS-tgNTN1^UNC5H-mut^, OS-tgNTN1^NEO1-mut^, and OS-tgNTN1^U-mut/N-mut^ cells. Right panels: UMAP representation of Control, OS-tgNTN1WT, OS-tgNTN1^UNC5H-mut^, OS-tgNTN1^NEO1-mut^, and OS-tgNTN1^U-mut/N-mut^ cells. Top panels: cells cultured in PXGL conditions. Bottom panels: cells cultured in N2B27 conditions (for 48 h). (D) Heatmap showing naïve- and primed-associated gene expression in Control, OS-tgNTN1WT, OS-tgNTN1^UNC5H-mut^, OS-tgNTN1^NEO1-mut^, and OS-tgNTN1^U-mut/N-mut^ cells cultured in PXGL (at P15) and in N2B27 (48 h).

### Early responses to tgNTN1^wt^ overexpression

To investigate the mechanisms by which Netrin-1 triggers the reprogramming of PSCs toward the naïve state, we performed quantitative proteomic and phosphoproteomic analyses at 4h, 12h and 24 h following Netrin-1 induction. We used OS cells carrying a doxycycline-inducible *tgNTN1^WT^* construct (OS-Dox-tgNTN1^WT^) (**Fig. 7A**). Immunofluorescence analysis confirmed progressive induction, with tgNTN1^WT^ detectable in 19%, 71%, and nearly 99% of cells after 4h, 12h, and 24h, respectively (**Fig. 7B**). Proteomic profiling of control cells (OS-Dox-GFP) revealed no significant changes in protein abundance across the time course (**Fig. 7C**,**D**). In contrast, Netrin-1 induction led to alterations in the proteome, with 92 proteins increased and 65 decreased after 12h and 24h of doxycycline treatment (**Supplementary Table S3**). Proteins associated with integrin signaling and cytoskeletal organization were enriched among the more abundant proteins, including PTPRA (Zeng et al., 2003), RAPH1/Lamellipodin (Lagarrigue et al., 2015), ARHGAP12 (Tambrin et al., 2025), and RALGPS2 (Ceriani et al., 2007). Consistent with these changes, several proteins involved in endosomal trafficking and membrane organization were also increased, including VPS26B (Wang et al., 2024), VPS45 (Frey et al., 2021), STON1/GTF2A1L (Martina et al., 2001), and RTN3 (Wu and Voeltz, 2021). Conversely, components of the Ras–MAPK/AP-1 pathway were reduced, including KRAS, RRAS, and JUN, while the MAPK phosphatase DUSP3 was increased. In parallel, multiple chromatin-associated and transcriptional regulators (BRD9, BRPF1, MED8, MED11, INTS2, ENY2, HMGB1, SET) showed decreased abundance, indicating remodeling of nuclear regulatory components.

**Figure 7:**
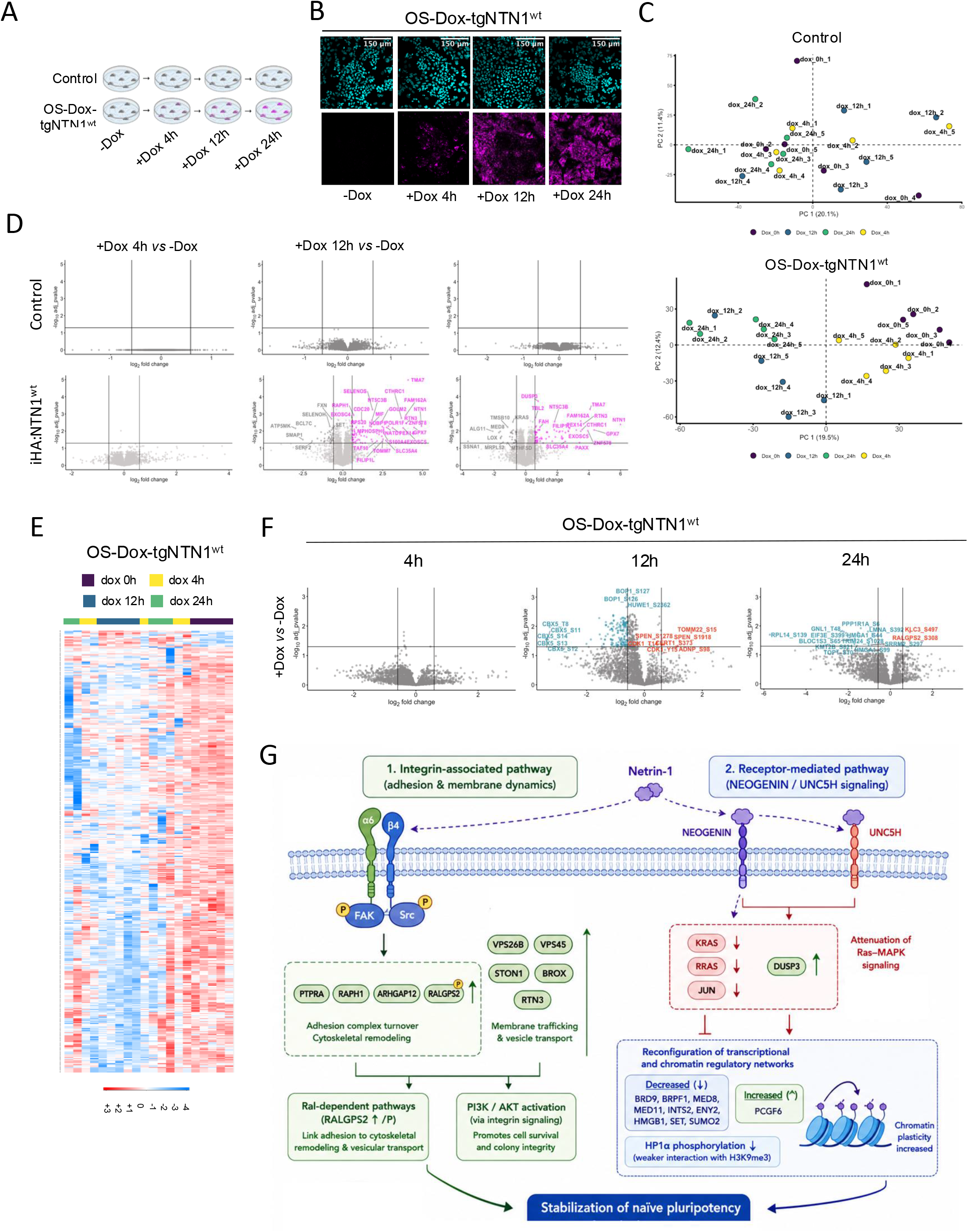
Proteomic and phospho-proteomic analyses of OS-Dox-tgNTN1 cells. (A) Experimental scheme. (B) Confocal images OS-Dox-tgNTN1^WT^ before and after treatment with doxycycline. Immunostaining shows HA:NTN1 expression. Scale bars, 100 µm. (C) Two-dimensional PCA of proteomic profiles from the indicated cell lines (three biological replicates). (D) Volcano plots showing differentially expressed proteins between doxycycline-treated (4h, 12h, and 24h) and untreated OS-Dox-tgNTN1^WT^ cells. Top panels: Control cells; Bottom panels: OS-Dox-HA:NTN1^WT^ cells. (E) Heatmap showing of phospho-peptide abundance in OS-Dox-HA:NTN1^WT^ cells before and after doxycycline treatment. (F) Volcano plots showing differentially expressed phosphopeptides between doxycycline-treated (4h, 12h, and 24h) and untreated OS-Dox-HA:NTN1^WT^ cells.

Phosphoproteomic analysis revealed widespread changes in phosphorylation, with a global trend toward decreased phosphopeptide abundance at 12h and 24h (**Supplementary Table S4; Fig. 7E,F**). Notably, phosphorylation of YAP at Ser61 was reduced, a modification associated with modulating YAP activity (Li et al., 2010; Qin et al., 2016). The most prominent phosphorylation changes were observed in Heterochromatin Protein 1a (HP1a/CBX5), with decreased phosphorylation at five N-terminal residues (Thr8, Ser11, Ser12, Ser13, and Ser14). These reductions were detectable as early as 4h and reached a minimum at 12h. They are commonly associated with reduced affinity to H3K9me3 and subsequent weakening of heterochromatin stability (Hiragami-Hamada et al., 2011; Sales-Gil and Vagnarelli, 2020).

Collectively, these data indicate that Netrin-1 induction is associated with coordinated changes in proteins involved in adhesion, signaling, and chromatin regulation, accompanied by broad alterations in phosphorylation dynamics (**Fig. 7G**).

## Discussion

Our findings identify Netrin-1 signaling as a modulator of naïve pluripotency in human ESCs. *NTN1* expression is associated with increased levels of key naïve markers, an effect that is further enhanced in NTN1^high^ cells co-expressing high levels of UNC5B, supporting a central role for the Netrin-1–UNC5B axis in promoting the naïve state. NEO1 appears to exert a context-dependent effect, as its association with naïve marker expression is primarily observed under conditions of elevated *NTN1*, suggesting that Netrin-1 can reinforce naïve pluripotency through both UNC5B and NEO1. This is consistent with previous observations in mouse ESCs, where Netrin-1 signaling through NEO1 and UNC5 receptors supports self-renewal via inhibition of MEK and GSK3β (Huyghe *et al*., 2020). In contrast, in the human PSCs, UNC5D expression correlates with reduced naïve marker levels in NTN1^high^ cells, raising the possibility that distinct UNC5 receptors exert opposing effects downstream of Netrin-1 to fine-tune the balance between self-renewal and differentiation.

Transcriptomic studies further support a role for Netrin-1 in stabilizing the naïve state. NTN1 knockout is associated with gene expression changes indicative of naïve state erosion, including upregulation of extracellular matrix and adhesion-related genes, and induction of developmental regulators and morphogen signaling components, notably within WNT and TGF-β/NODAL pathways. These changes suggest that loss of Netrin-1 alters both the extracellular signaling environment and the interpretation of differentiation cues. Concomitantly, downregulation of genes involved in amino acid and one-carbon metabolism, as well as mitochondrial function, points to a shift away from the anabolic metabolic program characteristic of naïve pluripotency. Together, these data support a model in which Netrin-1 integrates adhesion-dependent signaling with intracellular pathways, including PI3K–AKT and MAPK, to constrain premature activation of differentiation programs. Despite these effects, the overall contribution of Netrin-1 appears modulatory rather than essential, as *NTN1* inactivation does not impair cell growth under PXGL conditions.

A key finding of this study is the ability of Netrin-1 to reinforce and stabilize naïve pluripotency when overexpressed, as demonstrated by the increased expression of naïve-specific genes. We demonstrated the effects of Netrin-1 in two distinct experimental settings. First, during reprogramming of hESCs toward the naïve state under PXGL culture conditions, Netrin-1 overexpression significantly enhanced the acquisition of naïve pluripotency. Second, following withdrawal of LIF, PKC and Tankyrase inhibitors, Netrin-1 counteracted the destabilizing effects of factor deprivation and prevented differentiation, indicating that Netrin-1 contributes not only to the establishment but also to the maintenance of the naïve state. Our epigenomic analyses provide insight into the mechanisms that may underlie these effects. Rather than promoting a uniform shift toward chromatin activation or repression, Netrin-1 induced coordinated changes in both active and repressive histone modifications, consistent with large-scale remodeling of the pluripotent chromatin landscape. The genome-wide increase in H3K27ac suggests enhanced activity of promoters and enhancers, potentially facilitating activation of transcriptional programs associated with naïve pluripotency. Unexpectedly, this increase was accompanied by a global enrichment in H3K27me3. Although these marks are generally associated with opposing chromatin states, their simultaneous increase may reflect the dual requirement for activation of naïve pluripotency genes and repression of primed-state or lineage-specification programs during reprogramming. This interpretation is consistent with the view that acquisition of naïve pluripotency involves both enhancer activation and extensive Polycomb-mediated regulation of developmental genes (Boyer et al., 2006; Marks et al., 2012; Theunissen et al., 2016). The redistribution of H3K4me3 further supports a model of selective transcriptional rewiring. Rather than exhibiting a global increase, H3K4me3 was gained at some loci and lost at others, suggesting activation of specific gene networks accompanied by repression of alternative transcriptional programs. Such promoter-specific remodeling is consistent with the extensive transcriptional reorganization that accompanies transitions between pluripotent states (Buecker et al., 2014; Pastor et al., 2016).

Our combined proteomic and phosphoproteomic analyses indicate that Netrin-1 overexpression in naïve human PSCs rapidly engages at least two complementary signaling pathways: an integrin-associated pathway linked to adhesion and membrane dynamics, and a receptor-mediated pathway consistent with signaling through NEOGENIN and UNC5H family members. The enrichment of proteins such as PTPRA, RAPH1, ARHGAP12, and RALGPS2, together with increased abundance of endosomal and vesicle-associated factors (VPS26B, VPS45, STON1, BROX, RTN3) supports activation of integrin/Src/FAK-related processes (Caswell and Norman, 2006; Mitra et al., 2005). These changes are consistent with enhanced adhesion complex turnover and membrane trafficking that are known to influence cell survival and colony organization (Caswell and Norman, 2008; Maxfield and McGraw, 2004). The concomitant upregulation and phosphorylation RALGPS2 further suggest engagement of Ral-dependent pathways connecting adhesion signaling to cytoskeletal remodeling and vesicular transport (D’Aloia et al., 2021). In parallel, the reduced expression of *KRAS*, *RRAS*, and *JUN*, together with increased *DUSP3*, points to an attenuation of Ras–MAPK signaling. Given the well-established role of this pathway in destabilizing the naïve state (Nichols and Smith, 2009; Ying et al., 2008), this downregulation is consistent with a role for Netrin-1 in promoting or accelerating reprogramming toward this state. This effect may be mediated, at least in part, by Netrin-1 receptors such as NEOGENIN and UNC5H, which have been implicated in modulation of Ras-related signaling pathways (Forcet et al., 2002; Gibert and Mehlen, 2015). However, our observation that a Netrin-1 mutant with reduced affinity to NEOGENIN and UNC5H receptors retains the capacity to accelerate reprogramming to the naïve state when overexpressed suggests that Ras-related signaling is ancillary, rather than essential to this process. Changes in nuclear proteins further suggest a reconfiguration of transcriptional and chromatin regulatory networks. The decreased abundance of multiple chromatin-associated and transcriptional regulators, including BRD9, BRPF1, MED8, MED11, INTS2, ENY2, HMGB1, SET, and SUMO2, alongside selective increases in factors such as PCGF6, points to a restructuring of gene regulatory programs rather than a uniform shift in transcriptional activity. In addition, the observed decrease in HP1α phosphorylation at residues that regulate its interaction with H3K9me3 (Hiragami-Hamada *et al*., 2011; Sales-Gil and Vagnarelli, 2020) suggests a potential weakening of heterochromatin stability, which may contribute to increased chromatin plasticity. Although these pathways are mechanistically distinct, their effects are likely to converge. Enhanced integrin-associated signaling may support cell survival and structural integrity, while reduced MAPK activity and chromatin reorganization may limit differentiation cues and reinforce the naïve transcriptional state. Together, these changes are expected to stabilize naïve pluripotency by reinforcing colony integrity and limiting lineage priming.

Unexpectedly, Netrin-1 mutants with reduced affinity for NEO1 and UNC5H receptors reprogrammed hPSCs to the naïve state as efficiently as wild-type Netrin-1 when overexpressed. We therefore hypothesized that NEO1 and UNC5B receptors are already occupied by endogenous Netrin-1 produced by hESCs and feeder cells, such as exogenous Netrin-1 acts through a NEO1/UNC5B-independent mechanism to promote naïve pluripotency. Our proteomic analysis instead implicates integrins as candidate mediators of this alternative pathway. Netrin-1 is known to bind the α6β4 integrin, regulating adhesion, migration, and survival signaling (Stanco et al., 2009a; Yebra et al., 2003b), and this interaction is associated with activation of the PI3K/AKT pathway (Son et al., 2013), a key axis in maintaining naïve pluripotency (Duggal et al., 2015; Warrier et al., 2017). However, a direct role for α6β4 integrin in pluripotency control has not been formally established. In primed hESCs, α6β1–laminin interactions are required for single-cell survival and clonal expansion via Fyn–RhoA/ROCK signaling (Nakashima and Tsukahara, 2022; Rodin et al., 2014). These pathways contribute to the regulation of core pluripotency factors, including OCT4, NANOG, and SOX2. Notably, α6β1 integrin also serves as a receptor for Netrin-4 (Larrieu-Lahargue et al., 2011), which we find to be preferentially expressed in the human blastocyst epiblast. This raises the possibility that Netrin-4 engages a signaling circuitry analogous to that of Netrin-1 in the embryonic context.

Overall, our findings extend the role of Netrin-1 beyond its established functions, suggesting that it coordinates extracellular signaling, intracellular signaling cascades, and nuclear regulatory mechanisms in pluripotent cells. Further studies will be required to dissect the relative contributions of individual receptors and downstream effectors in mediating these effects.

## Material and Methods

### Cell lines, media composition, and culture

The male human ESC line Oscar (hPSCreg: INSRMe001-A; Chen *et al*., 2015) was routinely cultured in knockout Dulbecco’s modified Eagle’s medium (KO-DMEM; Gibco, Cat. 10829018) supplemented with 20% knockout serum replacement (KOSR, Gibco, Cat. 10828028), 1 mM GlutaMAX^TM^ (Gibco, Cat. 35050061), 0.1 mM β-mercaptoethanol (Sigma, Cat. M6250), 1% non-essential amino acid (Gibco, Cat. 11140050), 1% Penicillin-Streptomycin (Gibco, Cat. 15140122), and 4–8 ng/mL PeproGMP® Human FGF-basic (FGF2/bFGF) (Gibco, Cat. 100-18B) on growth-inactivated murine embryonic fibroblasts and maintained at 37 °C in 5% CO_2_ and 5% O_2_. Oscar hESCs were kept frozen in liquid nitrogen in KO-DMEM/fetal calf serum/DMSO (45:45:10), and thawed at passage 15-18. Passaging was performed by manual dissociation with Collagenase (Stem Cell technologies, Cat. 07909; split ratio 1:3). Oscar hESCs were converted to the PXGL state using the following medium composition (Guo *et al*., 2017): first in cRM1 media for three days with 1 μM PD0325901 (Miltenyi, Cat. 130-103-923), 10,000 U/mL hLIF, and 1mM Valproic acid, followed by the final media containing DMEM-F12 (Gibco, Cat. A4192001) and Neurobasal (Gibco, Cat. 10888022) (1:1) medium supplemented with N2 (Gibco, Cat. 17502001) and B27 (Gibco, Cat. A3582801) supplements, 1 μM PD0325901 (Miltenyi, Cat. 130-103-923), 10,000 U/mL hLIF, 2 μM Gö6983 (Bio-techne, Cat. 2285), 2 μM XAV939 (Merck, Cat. X3004). Cells were passaged every 4–5 days by single-cell dissociation using Tryple (Gibco, Cat. 12604939). ROCK inhibitor (Y-27632; Miltenyi, Cat. 130-103-922) was added during passaging for 24 hours. The human ESC line OS3, a naïve-like Oscar-derivative (Chen *et al*., 2015), was routinely cultured in KO-DMEM supplemented with 20% KOSR, 1 mM glutamine, 0.1 mM β-mercaptoethanol, 1% non-essential amino acid, 0.5 mg/mL 4’OH-tamoxifen, 10,000 U/mL hLIF, 1 μM PD0325901, and 3 µM CHIR99021 (Miltenyi, Cat 130-103-926). All cells were regularly tested for mycoplasma contamination by PCR (Applied Biosystems™ MycoSEQ™ Mycoplasma Detection Kit, Cat. 4460623.)

### Feeder preparation

All ESCs were cultured on feeder cells prepared either from growth-inactivated mouse embryonic fibroblasts or STO cells. NTN1-KO feeders were prepared from mouse embryos at E12.5 to E13.5. NTN1^fl/fl^ :: ZP3-Cre females were crossed to C57Bl6 males to obtain NTN1^+/-^mice. Then, embryos from mating NTN1^+/-^ male and female were dissected, dissociated into Accutase® (Gibco, Cat. A1110501) for 5min at 37°C, and plated on a 6-well plate into MEF medium comprising DMEM (Gibco, Cat. 10569010), 10% FCS, 0.1 mM β-mercaptoethanol, and 1% non-essential amino acid. Cells were expanded up until passage 3 before use. Embryo heads were kept for genotyping using the following primers: cNetrin1lox_F : 5’–CAGCTCTGAACTCTGGCTG– 3’; Netrin1lox_R : 5’–GGATACAGTAATCTGGGCTC– 3’.

### Generation of plasmids

#### PB-EF1α-NeoR backbone

HA-tagged cDNAs encoding human wild-type Netrin-1 (*HA:NTN1wt*), a mutant with reduced affinity for NEOGENIN/DCC receptors (*HA:NTN1^NEO1-mut^*), and a mutant with reduced affinity for UNC5H receptors (*HA:NTN1^UNC5H-mut^*), were amplified from *Xlone-NTN1-HA*, *Xlone-NTN1-1979-HA-Δneo1*, and *Xlone-NTN1-1950-HA-ΔUNC5B* (Huyghe *et al*., 2020) using the following primers: *5’-*TTCTTAGCTAGCACCGGTGCCACCATGATGCGCGCAGTG*-3’* and *5’-*AACTCAAGCG TAATCTGGAACATCGTATG*-3’*. The resulting 1862 bp fragments were sub-cloned between *Nhe*I and *Hpa*I in *PB-EF1α-NeoR* (Pham et al., 2025) to generate *PB-EF1α-HA:NTN1wt-NeoR, PB-EF1α-HA:NTN1^NEO1-mut^-NeoR,* and *PB-EF1α-HA:NTN1^UNC5H-mut^*, respectively. A mutant with reduced affinity for both NEOGENIN/DCC and UNC5H receptors (*HA:NTN1^U-mut/N-mut^*) was generated after sub-cloning a 844 bp *BsrG*I*/BstX*I fragment containing the *ΔUNC5B* mutation between *Nhe*I and *Hpa*I in *PB-EF1α-HA:NTN1 ^NEO1-mut^-NeoR*.

#### pPB-CAG-rtT3IRES backbones

a 1860 bp fragment encompassing *HA:NTN1wt* was amplified from Xlone-NTN1-HA (Huyghe *et al*., 2020) using the following primers: *5’-TATGAATTAATTAAGCCACCATGATGCGCGCAGTG-3’* and *5’-TCTAGAACCGGTTCAAGCGTAATCTGGAACATCGTATG-3’*, and sub-cloned between *Pac*I and *Age*I in *pPB-CAG-rtT3IRES-PuroR-TRE-pA* (Pham et al., 2025) to generate *pPB-CAG-rtT3IRES-PuroR-TRE-HA:NTN1wt-pA.* A 4282 bp fragment encompassing *Flag*3-NLS-SpCas9-NLS* was amplified from *AAV-Neo-CAG-Cas9* (addgene 86698, Ludovic Vallier’s Lab) using primers *5’-TATTCATTAATTAAGCCACCATGGACTATAAGG-3’* and *5’-CAATTG<u>ACCGGT</u>TTACTTTTT CTTTTTTGCCTGGC-3’*, and sub-cloned between *PacI*I and *Age*I in *PB-CAG-rtTA3-IRES-HygroR-TRE-SV40pA* to generate *PB-CAG-rtTA3-IRES-HygroR-TRE-SpCas9-SV40pA*.

#### LentiGuide-Puro backbone

the lentiGuide-Puro vector (Addgene ref#52963) was digested with *BsmB*I, and annealed oligonucleotides encoding the sgRNA protospacer sequence were ligated into the sgRNA expression cassette upstream of the scaffold. Oligonucleotides were designed based on a 20 bp target sequence located immediately upstream of an NGG PAM sequence in the genomic DNA. Cloning-compatible overhangs were included (CACC at the 5′ end of the sense oligo and AAAC at the 5′ end of the antisense oligo). sgRNAs targeting the human NTN1 gene were designed using CHOPCHOP with CRISPR/Cas9 knockout parameters, focusing on sequences near the translation start site in exon 2.

### Cell transfection, CRISPR knockout, and generation of transgenic lines

Human OS3 cells were transfected with 2.5 μg of either the *PB-EF1a-HA:NTN1^WT^-NeoR*, *PB-EF1a-HA:NTN^UNC5H-mut^-NeoR*, *PB-EF1a-HA:NTN1^NEO1-mut^-NeoR*, *PB-EF1a-HA:NTN1^N-mut/U-mut^-NeoR*, *pPB-CAG-rtT3IRES-PuroR-TRE-HA:NTN1^wt^-pA* or *PB-CAG-rtTA3-IRES-HygroR-TRE-SpCas9-SV40pA* vector alongside 2.5 μg of the PBase plasmid using the NEON transfection system, following the manufacturer’s instructions. Briefly, PSCs were dissociated into single cells and resuspended at a density of 10×10⁶ cells/mL. For each transfection, 100 μL of cell suspension was mixed with the plasmids. Transfection parameters were set to 1,200 V, 20 ms, and 2 pulses. Post-transfection, cells were plated onto multi-resistant growth-inactivated murine embryonic fibroblasts (MEF DR4) in medium supplemented with 10 μM ROCK inhibitor (Y-27632; Miltenyi Biotec) to enhance survival. Selection was performed using 250 ng/mL G418 or 1 ng/mL puromycin.

STO cells (10⁶ per condition) were transfected with 2.5 μg of *PB-EF1a-HA:NTN1^WT^-NeoR* and 2.5 μg of the PBase plasmid using NEON (1,400 V, 10 ms, 3 pulses). Stably transfected STO cells were selected using 250 ng/mL G418.

OS-DoxCas9 cells were transfected with either 2.5 µg of sgRNA#1 (5’-*GTACATCTTGCGGCACTGCGTGG*-3’) and 2.5 µg of sgRNA#2 (5’-*CATGATGCGCGCAGTGTGGGAGG*-3’), or with 5 µg of sgRNA#1, both of which target the *NTN1* locus at positions chr17:9022906 and chr17:9022373, respectively. Cas9 expression was activated 24 hours before transfection and 48 hours after, using 1 mg/mL Dox. After G418 selection, clones were selected and amplified before PCR amplification of *NTN1* using the following primers: 5’-*TCTGCGGCAGGCGGACAG*-3’ and 5’-*CTGCAGTCGCACACCAGG*-3’. OS-PXGL#1 was obtained using sgRNA#1, while OS-PXGL#2 and OS-PXGL#17 were obtained using both sgRNA#1 and sgRNA#2.

### Immunoblotting, immunostaining, and alkaline phosphatase assay

Frozen cell pellets were lysed in NP40 buffer complemented with protease and phosphatase inhibitors. Protein lysates were then cleared by centrifugation (13,300 rpm for 15 min). After SDS–PAGE and electroblotting on polyvinylidene difluoride, the membranes were incubated with specific primary antibodies (**Supplementary Table S6**) overnight at 4°C. Blots were then incubated with horseradish peroxidase-coupled anti-mouse or -rabbit immunoglobulin G (Jackson ImmunoResearch; **Supplementary Table S6**) for 1 hour at RT and developed with Clarity Western ECL Substrate (BIO-RAD).

For immunostaining, cells were fixed in 4% PFA in PBS for 20 minutes at room temperature, followed by four washes with PBS. Permeabilization was carried out in PBS containing 0.5% Triton X-100 for 1 hour. Non-specific binding sites were blocked using PBS containing 0,1% Triton X-100 and 5% of donkey serum for 1 hour at room temperature. Primary antibody (diluted in the blocking solution) incubation was performed overnight at 4°C. After four washes (5 minutes each) with PBS containing 0.1% Triton X-100, cells were incubated with secondary antibodies diluted in the blocking solution at room temperature for 1 hour. Nuclei were stained with DAPI (0.5 μg/mL). Finally, cells were mounted on coverslips using the mounting medium Fluoromount-G (Invitrogen) and analyzed by confocal imaging with a DM 6000 CS SP5 microscope (Leica) using an oil immersion objective (45×/1.25 0.75, PL APO HCX; Leica).

For alkaline phosphatase assay, human PSCs were plated at clonal density (200 cells/cm²) and maintained in PXGL medium for 1 week. After culture, cells were fixed for 30 s in a solution containing 8.2% formaldehyde and 25.5% citrate in acetone. Alkaline phosphatase staining was subsequently performed using the AP Substrate Kit (Sigma-Aldrich, 86R-1KT) following the manufacturer’s protocol.

### RNA extraction and sequencing

Total RNA was isolated from 4–5 ×10^6^ cells using RNeasy mini-kit (Qiagen) with a DNase I (Qiagen #79254) treatment. The libraries were prepared using 200 ng of RNA with the Stranded mRNA ligation it (Illumina). Samples were sequenced on a NovaSeq sequencing machine (Illumina) as paired-end reads of 100 bp. The bcl2fastq conversion software was used for demultiplexing (Illumina), and data trimming was performed using Cutadapt. The sequencing depth for each sample was approximately 30 million reads, which were subsequently mapped to the GRCh38 human genome.

### Cut&Run

CUT&RUN was performed as previously described (Skene and Henikoff, 2017). Briefly, 0.5 million cells per replicate were bound to 20 µl of concanavalin A-coated beads (Bangs Laboratories) in binding buffer (20 mM HEPES, 10 mM KCl, 1 mM CaCl_2_ and 1 mM MnCl_2_). The beads were washed and resuspended in DIG Wash buffer (20 mM HEPES, 150 mM NaCl, 0.5 mM spermidine and 0.05% digitonin). The primary antibodies (1:50) were added to the bead slurry and rotated at room temperature for 1 h. The beads were washed with DIG Wash buffer and protein A–MNase (micrococcal nuclease) fusion protein (1:400, produced by the Institut Curie Recombinant Protein Platform, 0.785 mg.ml^−1^) was added, before rotating at room temperature for 15 min. After two washes, the beads were resuspended in 150 µl of DIG Wash buffer and the MNase was activated with 2 mM CaCl2, before incubating for 30 min at 0 °C. MNase activity was terminated with 150 µl of 2× Stop buffer (200 mM NaCl, 20 mM EDTA, 4 mM EGTA, 50 µg.ml^−1^ RNase A and 40 µg.ml^−1^ glycogen). Cleaved DNA fragments were released by incubating for 20 min at 37 °C, followed by centrifugation for 5 min at 16,000*g* at 4 °C and collection of the supernatant from the beads on a magnetic rack. The DNA was purified by phenol–chloroform and libraries were prepared using the KAPA HyperPrep Kit from Roche following the manufacturer’s protocol. Sequencing was performed on a NovaSeq 6000 instrument (ICGex next-generation sequencing (NGS) platform) to generate 2 × 100 paired-end reads. All the antibodies used in this study are listed in **Supplementary Table 6.**

Sequencing data were processed using the nextflow ChIP-Seq pipeline (https://doi.org/10.5281/zenodo.10357860). Briefly, reads were mapped with bwa (version 0.7.17-r1188; {Li, 2010 #8674) to the human genome (version hg38). Duplicates or low-quality reads were removed. Peaks were called with MACS2 (version 2.2.7.1; Zhang et al., 2008) using the “broad” option. The percentage occupancy was calculated by normalizing the total length of peaks per chromosome by the size of their respective chromosome. For histone modification enrichment, raw counts were generated using multiBigwigSummary from deepTools (version 3.5.4) with ‘--binSize 10000’ and the mean counts per chromosome were calculated. Differential binding analysis was performed using the Bioconductor package DiffBind (version 3.14.0) in R. Differentially enriched regions were identified using the DESeq2 framework implemented in DiffBind.

### Proteomic and phosphoproteomic analyses

Cells were lysed in lysis buffer (6 M guanidine hydrochloride, 5mM TCEP, 10 mM CAA, 100 mM Tris pH 8.5) for 10 min at 95° C and 100 rpm. Samples were then sonicated (Bandelin BR-30, 10 s pulse 20 s breaks, 20 cycles) and then homogenized using QIAshredder devices (Qiagen). Protein content was assessed using a BCA assay. Samples were then diluted to 2 M guanidine hydrochloride with Tris 25 mM, trypsin (1:100 enzyme to protein) was added and the samples were digested overnight at 37° C. Digestion was stopped by acidification with formic acid. Samples were then desalted using Sep-Pak C18 Plus Light (Waters). Phosphopeptides were enriched using a commercial kit (High Select TiO2 Phosphopeptide Enrichment Kit, Thermo Fisher). Samples were dried using vacuum centrifugation and then resuspended in 0.1 formic acid in water and subjected to mass spectrometry-based proteomics. The enriched phospho-peptides were analyzed on a nanoElute 2-LC system coupled online to a timsTOF HT (both Bruker) operated in the data-dependent acquisition (DDA) mode. Peptides were transferred to a PepMap Neo trap column (300 µm× 5 mm, 5 µM particles, C18, Thermo Fisher Scientific) and then separated on a PepSep Ultra analytical column (C18, 25 cm x 75 µm, 1.5 µm particles, Bruker) at 250 nl/min with a 25-min gradient of 2-25% of solvent B followed by a 12-min increase to 37%. Solvent A consisted of 0.1% formic acid in water and solvent B of 0.1% formic acid in acetonitrile. Raw data was then analyzed using MaxQuant 2.6.7.0 (Tyanova et al., 2016) with all human entries in Swiss-Prot as a database (retrieval data: 11.01.2024). Phosphorylations at serine, threonine and tyrosine were set as variable modifications, match between runs and label free quantification were turned on. For the whole proteome analysis, peptides prior to desalting were analyzed using the same instrumentation and separation method, however the data was acquired using data independent acquisition. The raw data was then searched using DIA-NN 2.1.0 (Demichev et al., 2020) and the same sequence database as above. The proteomics and phosphoproteomics data are available in the MassIVE repository.

### Bioinformatics analysis

All RNA-seq data analyses were performed using R software (version 4.4.1). For cell lines bulk RNA-sequencing, data normalization and gene expression levels were carried out using the DESeq2 R package (version 1.44.0) (Love et al., 2014). For each sample, mean expression levels and standard deviations were calculated using the R base package (version 4.4.1) based on three replicates. Heatmaps were generated with the ggplot2 package (version (3.5.1) using log2-transformed normalized counts. Single-cell RNA-sequencing data were analyzed with the Seurat R package (version 4.3.0) (Hao et al., 2021). Briefly, gene expression values were log-transformed and normalized, and PCA was performed using 2000 variable genes as input. Differential expression analysis was conducted using the FindMarkers() function of Seurat (logfc.threshold = 0.8, min.pct = 0.5), filtering for upregulated genes (avg_log2FC > 0) with a p-value ≤ 0.05 for subsequent enrichment analysis. Finally, KEGG pathway enrichment analysis was performed using the clusterProfiler (version 4.12.6) (Yu et al., 2012) and DOSE (version 3.30.5) R packages with the human database org.Hs.eg.db (version 3.18.0).

## Statistical analysis

A two-sided Welch’s t-test for unequal variances was used throughout the study to compare distributions (ns, non-significant; *, p<0.05; **, p<0.01; ***, p<0.001). Cell counts are represented as box plots, with the center line indicating the median; box limits representing the upper and lower quartiles; whiskers extending to 1.5x interquartile range, and individual points denoting outliers (3 biological replicates).

## Data availability

The RNA-Seq datasets will be publicly available as of the date of publication.

## Declaration of generative AI and AI-assisted technologies in the manuscript preparation process

During the preparation of this work, the authors used ChatGPT to generate the schematic shown in Fig. 7G and to improve the language. The authors reviewed and edited the output as needed and take full responsibility for the content of the published article.

## Acknowledgments

This work was supported by the Fondation pour la Recherche Médicale (EQU202303016295 to P.S.), the LabEx REVIVE (ANR-10-LABX-73 to P.S.), the LabEx “DEVweCAN” (ANR-10-LABX-0061 to P.S.), the LabEx “CORTEX” (ANR-11-LABX-0042 to P.S.), the Infrastructure Nationale en Biologie et Sante INGESTEM (ANR-11-INBS-0009 to P.S.), and the Fondation pour la Recherche contre le Cancer (RAC18005CCA to I.A). We also thank the ICGex NGS platform of the Institut Curie supported by the grants ANR-10-EQPX-03 (Equipex), ANR-10-INBS-09-08 (France Génomique Consortium), and ANR-from the Agence Nationale de la Recherche (‘Investissements d’Avenir’ program), the ITMO-Cancer Aviesan (Plan Cancer III) and the SiRIC-Curie program (SiRIC Grant INCa-DGOS-465 and INCa-DGOSInserm_12554) for the high-throughput sequencing. We thank the bioinformatics platform of the Institut Curie for data management, quality control and primary analysis.

## Authors’ contribution

A.d.N. C.R., N.D., J.B.S., C.A. P.O., and M.L. performed plasmid constructs, ES and iPS cell line cultures and quality control, embryo cultures, and cell characterization.

A.d.N., E.M., C.A., G.M., and J.S. performed bioinformatic analyses.

A.d.N., E.M., C.A., T.F., I.A., and P.S. analyzed the data

F.L. provided essential reagents and offered scientific advice

P.S. conceptualized the study and wrote the manuscript

## Declaration of interest

The authors declare no competing interest.

## Notes

### Competing Interest Statement

The authors have declared no competing interest.

