## Supplemental Information for "Netrin-1 Acts as a Guardian of Naive Pluripotency in Human Embryonic Stem Cells"

**Supplementary information**

Supplementary Figure 1
Supplementary Figure 2
Supplementary Figure 3
Supplementary Figure 4
Supplementary Figure 5
Supplementary Figure 6

Supplementary Table 1
Supplementary Table 2
Supplementary Table 3
Supplementary Table 4
Supplementary Table 5

**Supplementary Figure 1**: **Expression of Netrin ligand and receptor families in primate embryos.**
(**A, B**) Correlation plots of the indicated genes. Numbers indicate the correlation coefficient between the two indicated genes.
(**C**) Heat map of the correlation coefficients among the indicated genes.

**Supplementary Figure 2**: **Expression of naïve pluripotency markers according to Oscar-PXGL's receptors status.** Violin plot representation of naïve pluripotency marker differential expression between $NTN1^{Low}$ and $NTN1^{High}$ cells in:
(**A**) $NEO1^{Low}$-$UNC5B^{Low}$, $NEO1^{Low}$-$UNC5B^{High}$, $NEO1^{High}$-$UNC5B^{Low}$, and $NEO1^{High}$-$UNC5B^{High}$ cells;
(**B**) $NEO1^{Low}$-$UNC5D^{Low}$, $NEO1^{Low}$-$UNC5D^{High}$, $NEO1^{High}$-$UNC5D^{Low}$, and $NEO1^{High}$-$UNC5D^{High}$ cells;
(**C**) $UNC5B^{Low}$-$UNC5D^{Low}$, $UNC5B^{Low}$-$UNC5D^{High}$, $UNC5B^{High}$-$UNC5D^{Low}$, and $UNC5B^{High}$-$UNC5D^{High}$ cells.
Two-sided Student's t-test (ns, non-significant; *, $p < 0.05$; **, $p < 0.01$; ***, $p < 0.001$).

**Supplementary Figure 3: *NTN1* knockout in OS-PXGL cells.**
(**A**) DNA sequences of the NTN1 locus between nucleotides 101 and 180 (top line) and between nucleotides 641 and 690 (bottom line) with respect to cap site, showing the ATG codon and the positions of indels in each mutant allele.
(**B**) Description of clones #1, #2, #15, #16, #12, and #17 with respect to wt or mutant alleles.
(**C**) Phase contrast images of OS-PXGL-NTN1$^{+/+}$ (clone #6 and #12) and OS-PXGL-NTN$^{-/-}$ (clone #2 and #17) cultured on DR4 feeders. Scale bars, 100 µm.
(**D**) Genotyping of mouse fetuses by genomic PCR.
(**E**) Phase contrast images of OS-PXGL-NTN1$^{+/+}$ (clones #6 and #12) and OS-PXGL-NTN$^{-/-}$ (clones #2 and #17) cultured on STO-Ctrl and STO-tgNTN1wt feeders. Scale bars, 100 µm.
(**F**) Principal component analysis (PCA) of RNA-seq data obtained from the indicated cell lines.
(**G**) Phase-contrast and confocal images of STO-ctrl and STO-tgNTN1 PXGL. Immunostaining shows HA:Netrin-1 expression.
(**H**) Phase contrast images of OS-PXGL-NTN1$^{+/+}$ (clones #5 and #12) and OS-PXGL-NTN$^{-/-}$ (clones #1 and #17) cultured on STO-ctrl and STO-tgNTN1 feeders. Scale bars, 200 µm.
(**I**) Volcano plot of differentially expressed genes (DEGs) between OS-PGXL-NTN1$^{+/+}$ (clones #5 and #12) and OS-PGXL-NTN$^{-/-}$ (clone #1 and #17) cultured on STO-Ctrl (top panel) and STO-tgNTN1wt (bottom panel) feeders.

**Supplementary Figure 4: Transcriptome characterization of OS-tgNTN1$^{wt}$ cells**.
(**A**) Phase-contrast images of control and OS-tgNTN1$^{WT}$ cells cultured in PXGL for 16 and 31 passages.
(**B**) GO term enrichment analysis of the DEGs between OS-tgNTN1$^{WT}$ and control cells.
(**C**) Colony-forming assay performed with OS-tgNTN1$^{WT}$ and control cells.
(**D**) Histogram of colony counting with OS-tgNTN1$^{WT}$ and control cells.
(**E**) Histogram of colony area with OS-tgNTN1$^{WT}$ and control cells.
(**F**) Violin plot representation of pluripotency (naïve, formative, and primed) and lineage marker expression in $NEO1^{Neg}$ *versus* $NEO1^{High}$ cells. Two-sided Student's t-test (ns, non-significant; *, $p < 0.05$; **, $p < 0.01$; ***, $p < 0.001$).

(**G**) Violin plot of pluripotency (naïve, formative, and primed) and lineage marker expression in *UNC5B*$^{Neg}$ *versus* *UNC5B*$^{High}$ cells. Two-sided Student's t-test (ns, non-significant; *, $p < 0.05$; **, $p < 0.01$; ***, $p < 0.001$).

**Supplementary Fig. 5: Histone mark occupancy in OS-tgNTN1$^{wt}$ and control cells.** Percentage of H3K27ac, H3K27me3, H3K4me3 and H3K9me3 peak occupancy per chromosome for OS-tgNTN1wt and control cells at P16 and P40.

**Supplementary Fig. 6: Transcriptome characterization of OS-tgNTN1$^{UNC5H\text{-}mut}$, OS-tgNTN1$^{NEO1\text{-}mut}$, and OS-tgNTN1$^{U\text{-}mut/N\text{-}mut}$ cells.**
(**A**) Western blot analysis if Netrin-1 expression in OS-tgNTN1$^{wt}$, OS-tgNTN1$^{UNC5H\text{-}mut}$, OS-tgNTN1$^{NEO1\text{-}mut}$, and OS-tgNTN1$^{U\text{-}mut/N\text{-}mut}$ cells.
**(B)** Volcano plot of DEGs between OS-tgNTN1$^{WT}$, OS-tgNTN1$^{UNC5H\text{-}mut}$, OS-tgNTN1$^{NEO1\text{-}mut}$, and OS-tgNTN1$^{U\text{-}mut/N\text{-}mut}$ and control cells under PXGL (15 passages) and N2B27 (48 h) conditions.
(**C**)Violin plot of naïve, formative, and primed pluripotency markers in control, OS-tgNTN1$^{wt}$, OS-tgNTN1$^{UNC5H\text{-}mut}$, OS-tgNTN1$^{NEO1\text{-}mut}$, and OS-tgNTN1$^{U\text{-}mut/N\text{-}mut}$ cells. Two-sided Student's t-test (ns, non-significant; *, $p < 0.05$; **, $p < 0.01$; ***, $p < 0.001$).

**Supplementary Table 1: Transcriptome alterations associated with NTN1 knockout in OS cells**

Sheet 1: DEGs between *NTN1*$^{-/-}$ #1 and #17 *versus NTN1*$^{+/+}$ #5 and #12 (across all four pairwise comparisons) cultured on STO-ctrl

Sheet 2: DEGs between *NTN1*$^{-/-}$ #1 and #17 *versus NTN1*$^{+/+}$ #5 and #12 (across three out of four pairwise comparisons) cultured on STO-ctrl.

Sheet 3: DEGs between *NTN1*$^{-/-}$ #1 and #17 *versus NTN1*$^{+/+}$ #5 and #12 (across all four pairwise comparisons) cultured on STO-tgNTN1wt

Sheet 4: DEGs between *NTN1*$^{-/-}$ #1 and #17 *versus NTN1*$^{+/+}$ #5 and #12 (across three out of four pairwise comparisons) cultured on STO-tgNTN1wt

**Supplementary Table 2: Transcriptome alterations associated with overexpression of tgNTN1 in OS cells**

Sheet 1: DEGs between OS-tgNTN1$^{wt}$ and control cells cultured in PXGL medium

Sheet 2: DEGs between OS-tgNTN1$^{wt}$ and control cells cultured in N2B27 medium

Sheet 3: DEGs between OS-tgNTN1$^{UNC5H\text{-}mut}$ and control cells in PXGL medium

Sheet 4: DEGs between OS-tgNTN1$^{UNC5H\text{-}mut}$ and control cells in N2B27 medium

Sheet 5: DEGs between OS-tgNTN1$^{NEO1\text{-}mut}$ and control cells in PXGL medium

Sheet 6: DEGs between OS-tgNTN1$^{NEO1\text{-}mut}$ and control cells in N2B27 medium

Sheet 7: DEGs between OS-tgNTN1$^{N\text{-}mut/H\text{-}mut}$ and control cells in PXGL medium

Sheet 8: DEGs between OS-tgNTN1$^{N\text{-}mut/H\text{-}mut}$ and control cells in N2B27 medium

**Supplementary Table 3: Differentially expressed proteins (DEPs) between Dox-induced and non-induced OS cells.**

Sheet 1: DEPs between 4h Dox-induced and non-induced OS-Dox-tgNTN1$^{wt}$ cells

Sheet 2: DEPs between 12h Dox-induced and non-induced OS-Dox-tgNTN1$^{wt}$ cells

Sheet 3: DEPs between 24h Dox-induced and non-induced OS-Dox-tgNTN1$^{wt}$ cells

**Supplementary Table 4: Differentially expressed phospho-sites between Dox-induced and non-induced OS cells.**

Sheet 1: Differentially expressed phospho-sites between 4h Dox-induced and non-induced OS-Dox-tgNTN1$^{wt}$ cells

Sheet 2: Differentially expressed phospho-sites between 12h Dox-induced and non-induced OS-Dox-tgNTN1$^{wt}$ cells

Sheet 3: Differentially expressed phospho-sites between 24h Dox-induced and non-induced OS-Dox-tgNTN1$^{wt}$ cells

**Supplementary Table 5: List of primary and secondary antibodies**

| Target | Supplier | Reference | Host species | Dilution for cells | Dilution for preimplan-tation embryos | Dilution for immuno-blotting | Dilution for CUT &RUN |
|---|---|---|---|---|---|---|---|
| HA | Sigma-Aldrich | H6908 | Rabbit | NA | NA | 1:2000 | NA |
| OCT4 | Tebu-Bio | 09-0023 | Rabbit | 1:300 | 1:100 | NA | NA |
| NANOG | R&D Systems | AF2729 | Goat | 1 :300 | NA | NA | NA |
| SOX2 | R&D Systems | AF2018 | Goat | 1:100 | 1:100 | NA | NA |
| DPPA5 | Sigma-Aldrich | AF3125 | Rabbit | 1:100 | NA | NA | NA |
| ALPPL2 | Abcam | Ab96497 | Rabbit | 1:100 | NA | NA | NA |
| KLF17 | Sigma-Aldrich | HPA024629 | Rabbit | 1:100 | NA | NA | NA |
| ACTINB | Sigma-Aldrich | A3854 | Mouse | NA | NA | 1:10000 | NA |
| Netrin-1 | Abcam | ab126729 | Rabbit | NA | NA | 1:1000 | NA |
| GAPDH | Santa-Cruz | sc-47724 | Mouse | NA | NA | 1:1000 | NA |
| H3K4me3 | Active motif | Cat#39060 | Rabbit | NA | NA | NA | 1:50 |
| H3K9me3 | Active motif | Cat#39062 | Rabbit | NA | NA | NA | 1:50 |
| H3K27me3 | Active motif | Cat#39055 | Rabbit | NA | NA | NA | 1:50 |
| H3K27ac | Active motif | Cat#39034 | Rabbit | NA | NA | NA | 1:50 |
| Anti-IgG | Diagenode | Cat#C15410206 | Rabbit | NA | NA | NA | 1:50 |

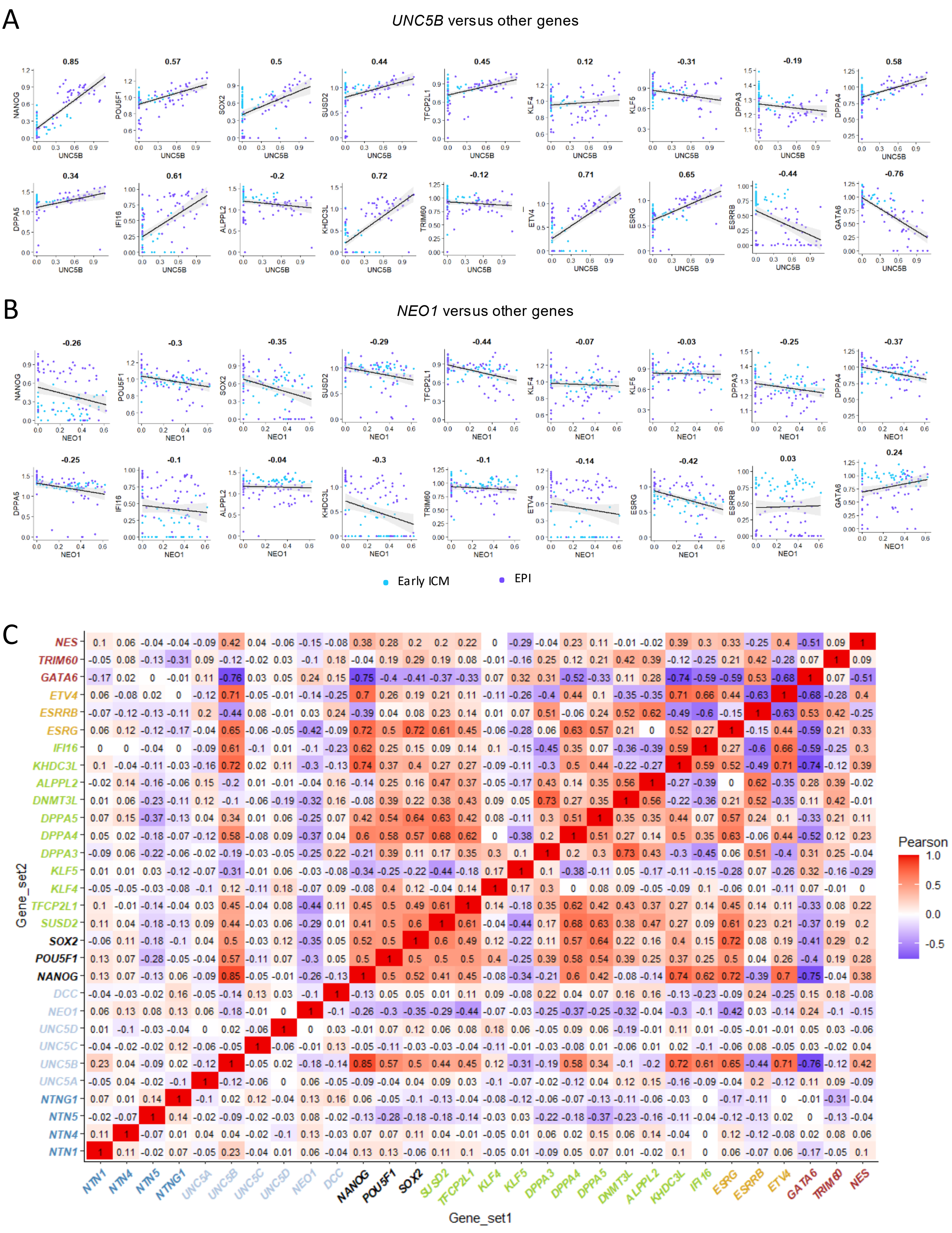

Supplementary Figure 1

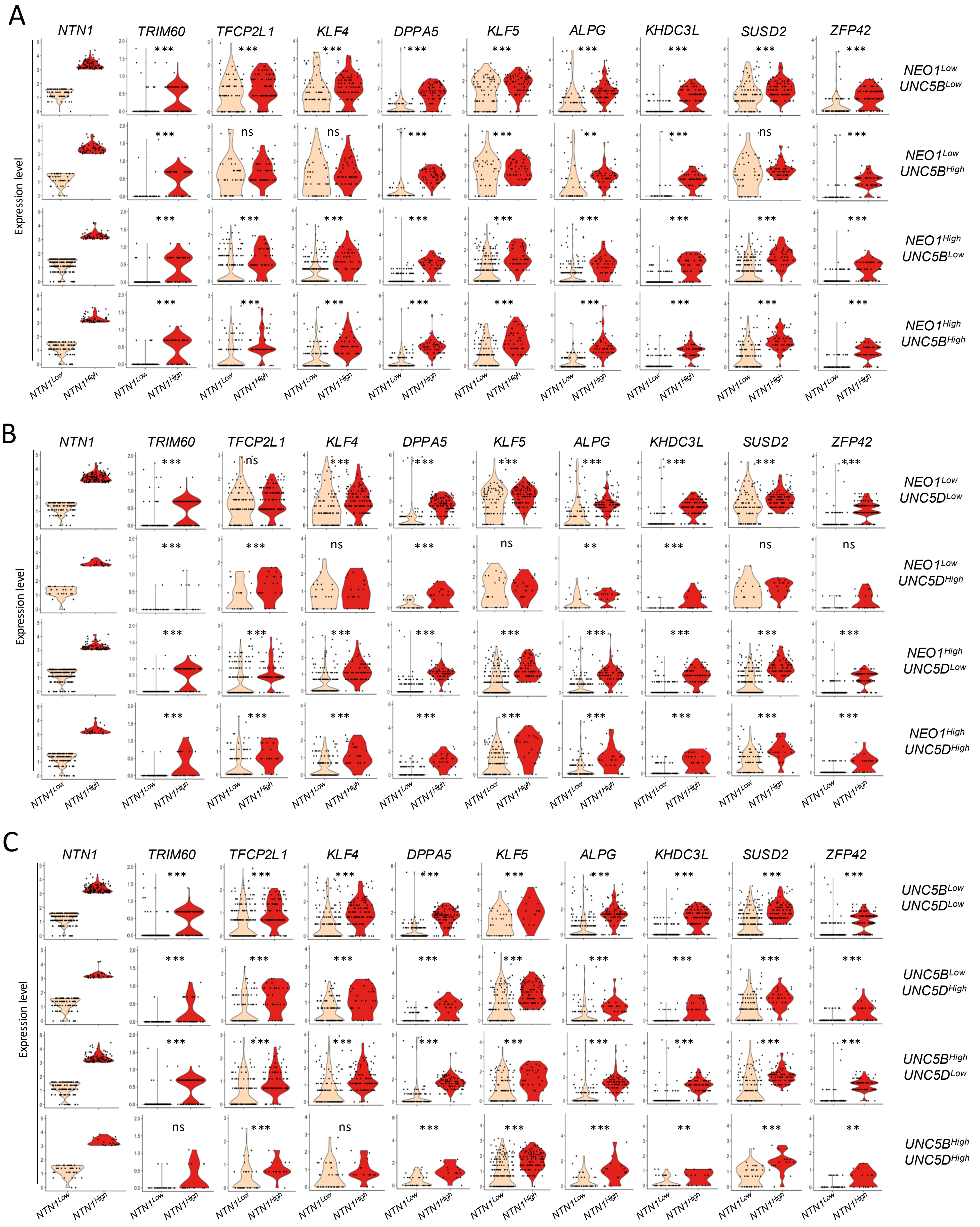

Supplementary Figure 2

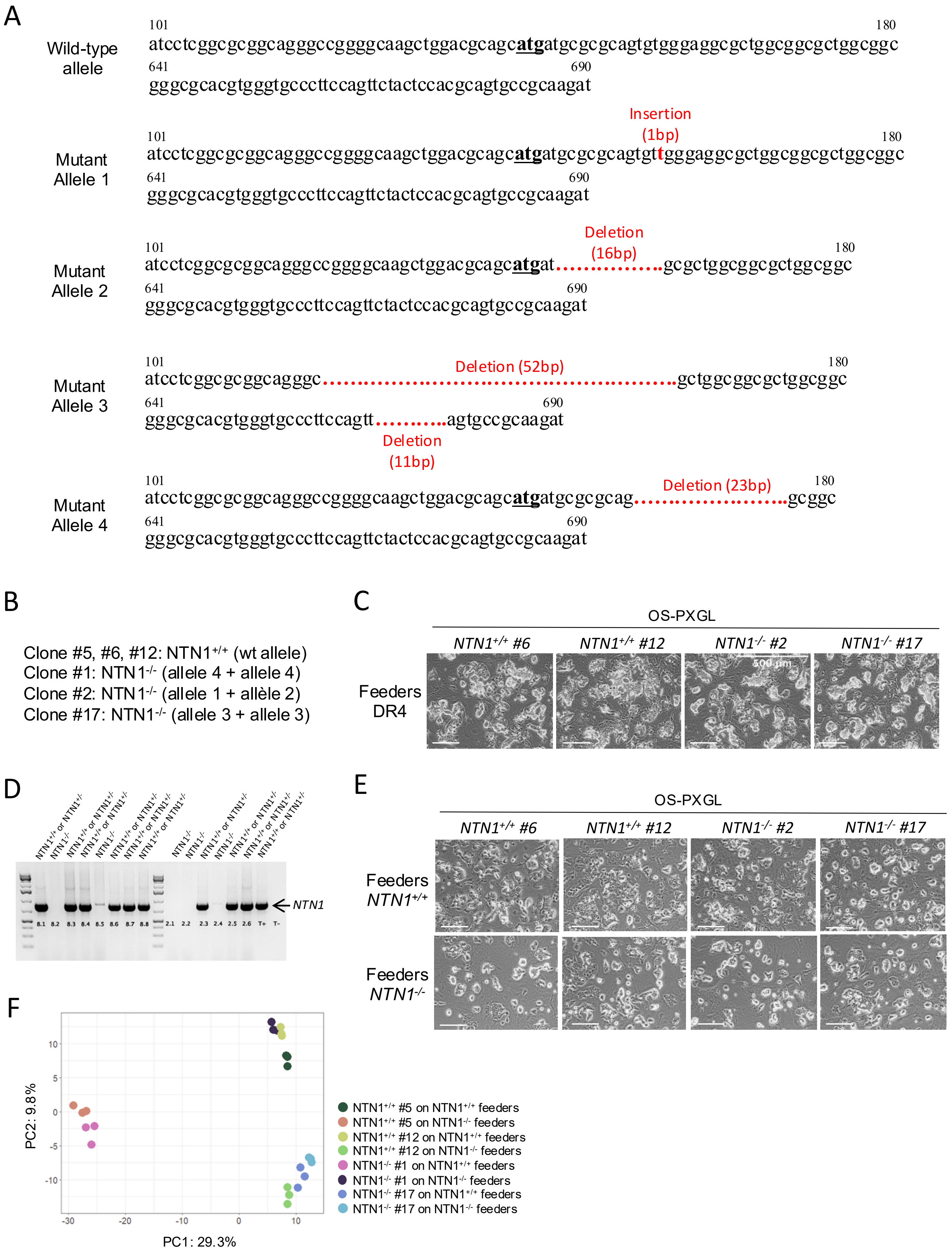

Supplementary Figure 3 (i)

G

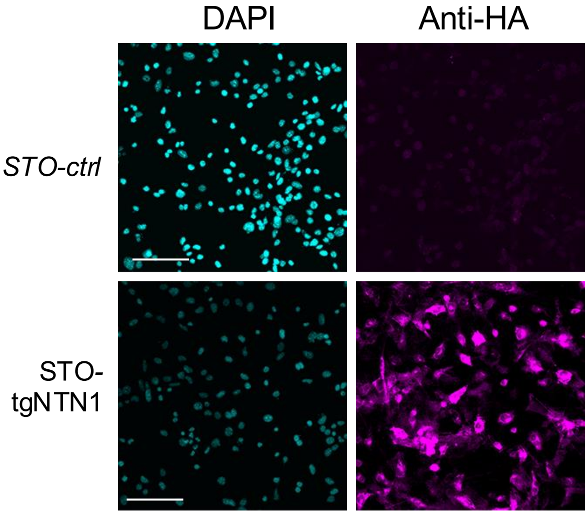

H

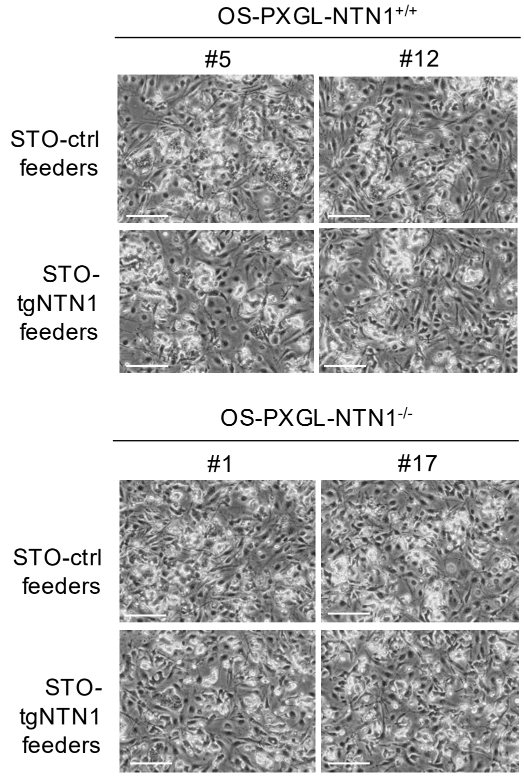

I

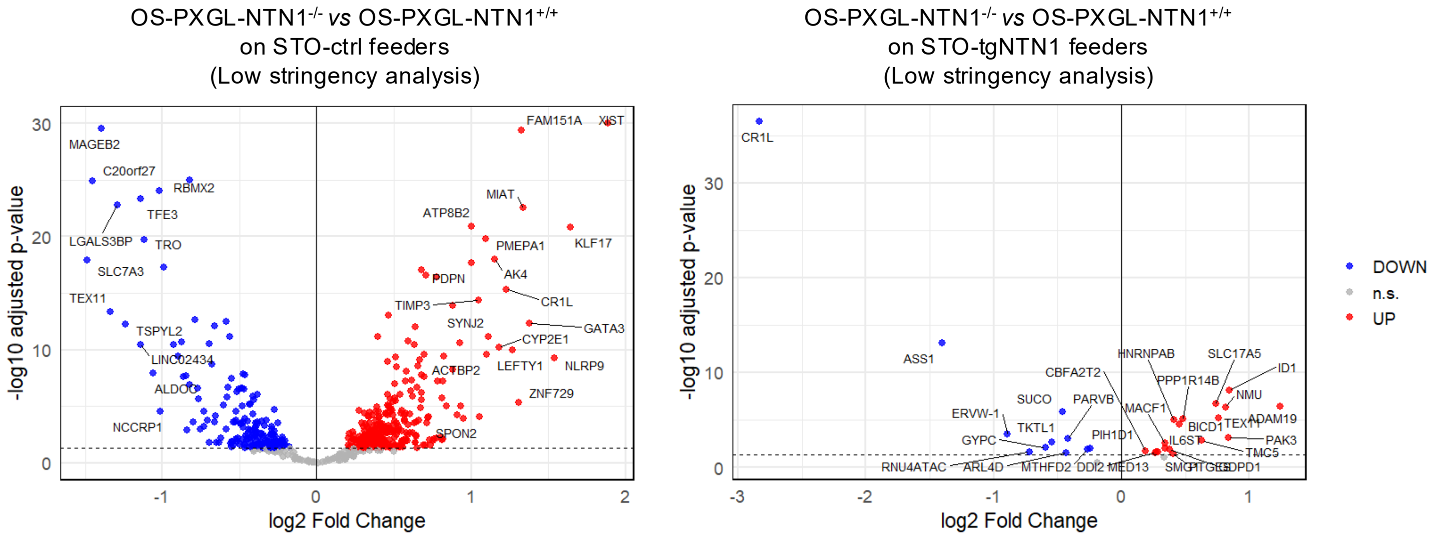

Supplementary Figure 3 (ii)

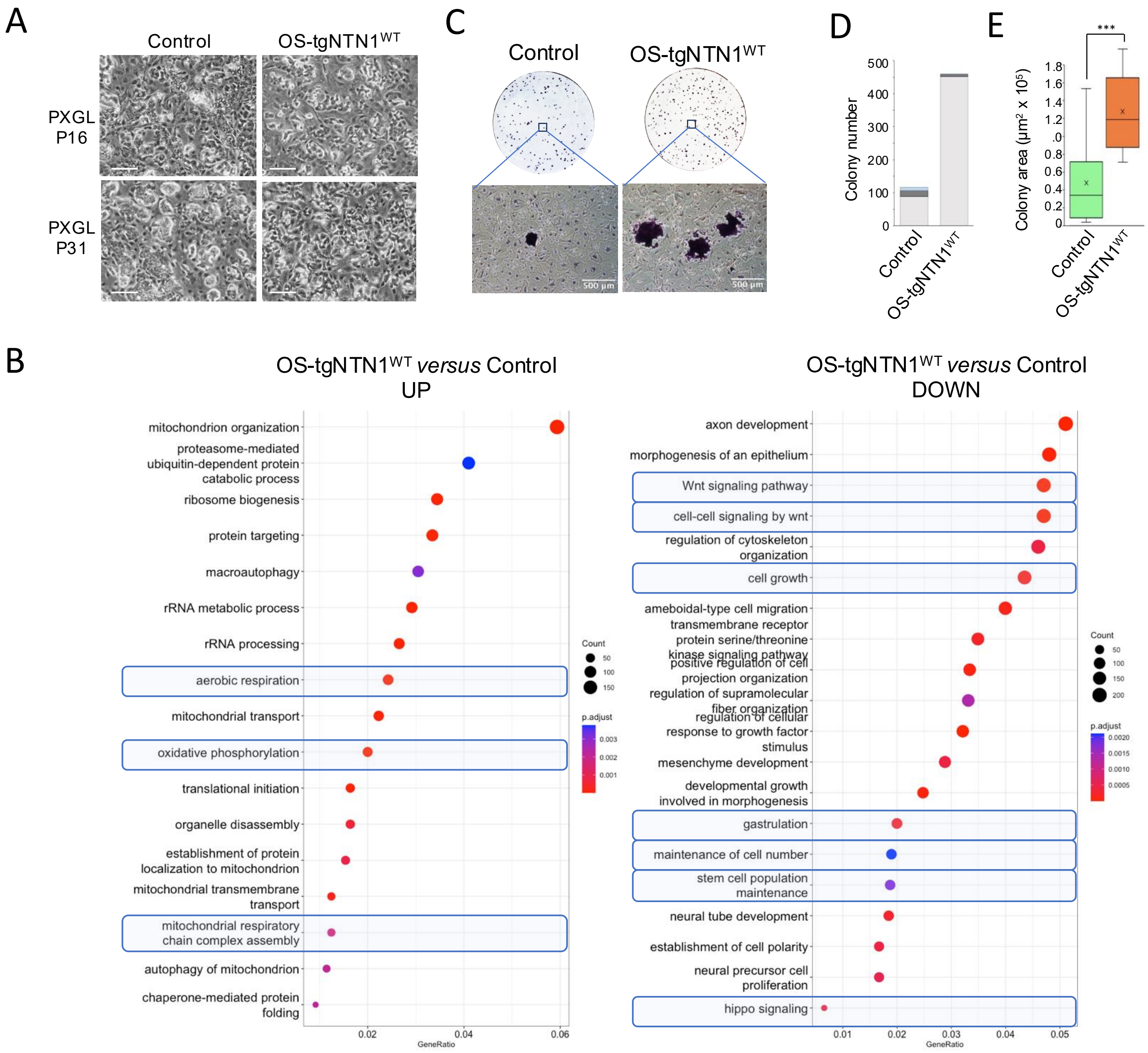

Supplementary Figure 4

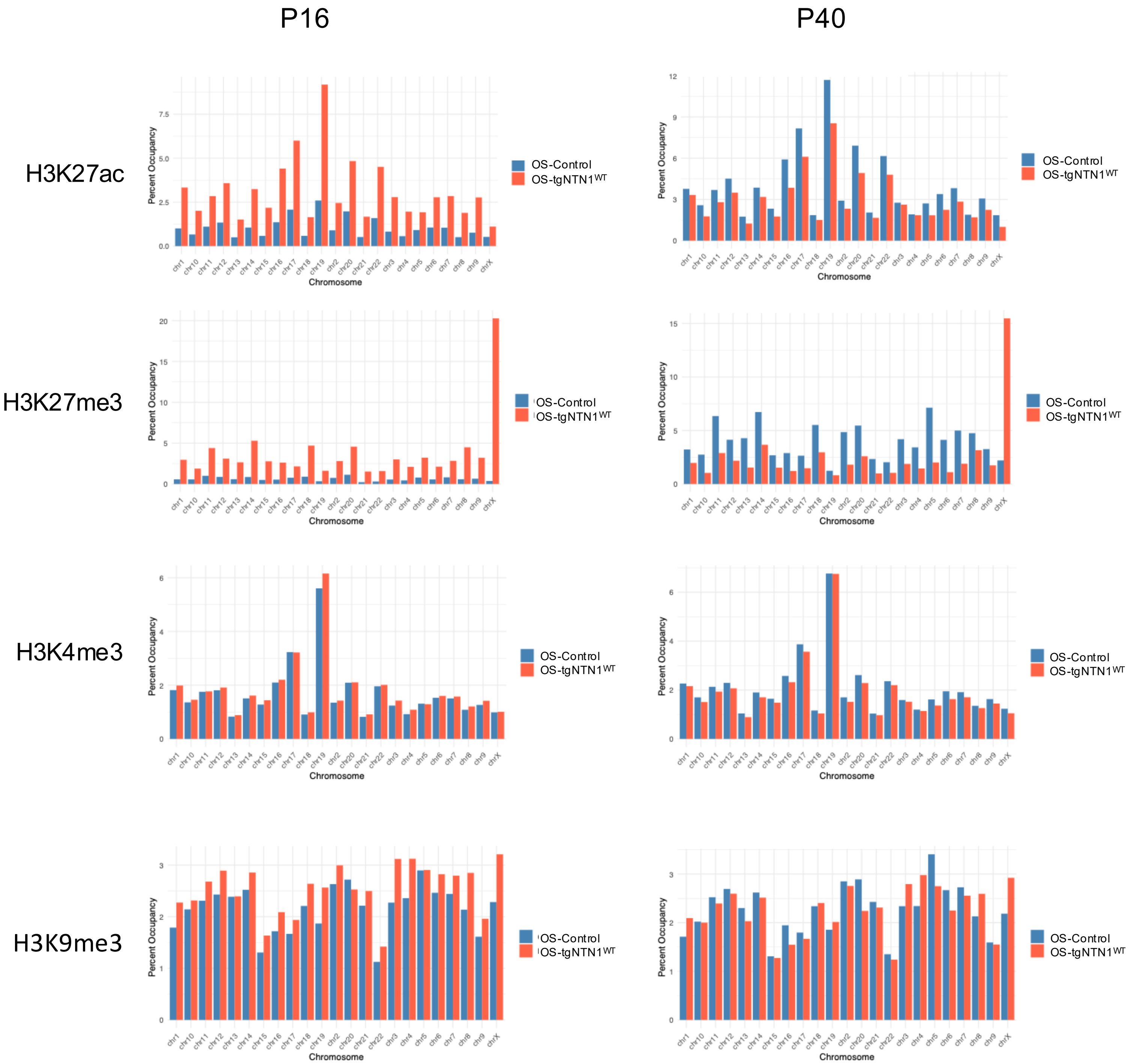

Supplementary Figure 5

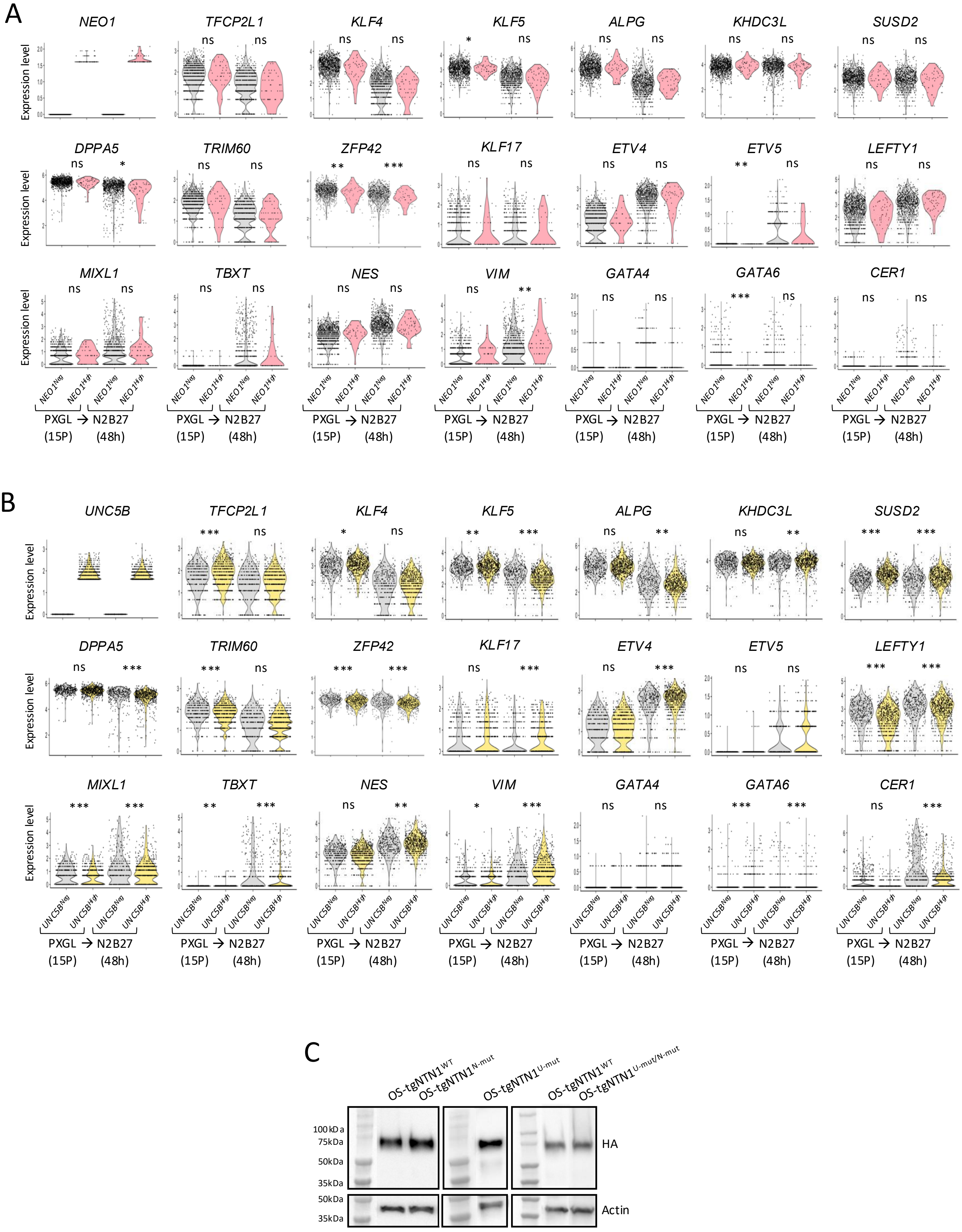

Supplementary Figure 6 (i)

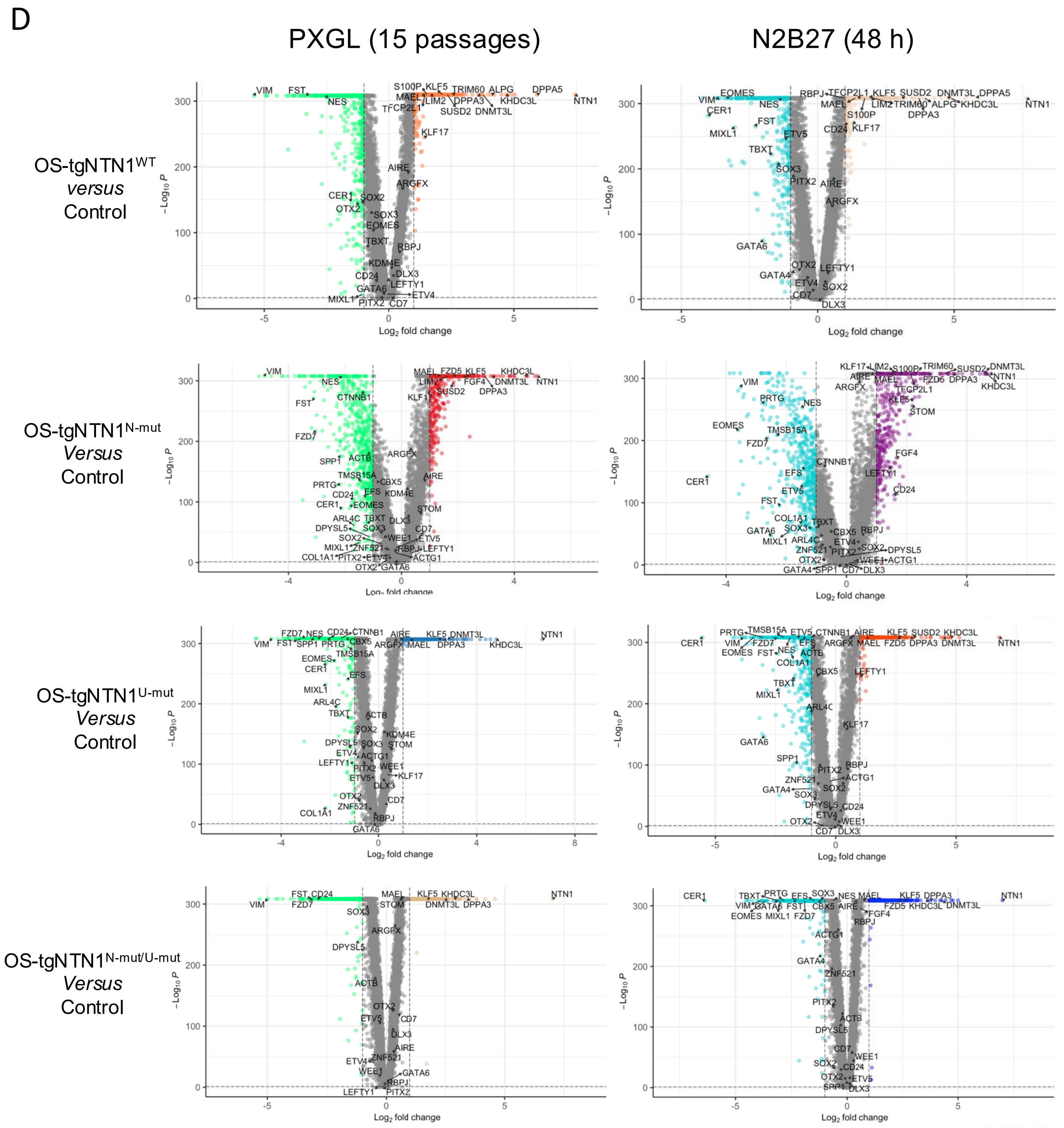

Supplementary Figure 6 (ii)

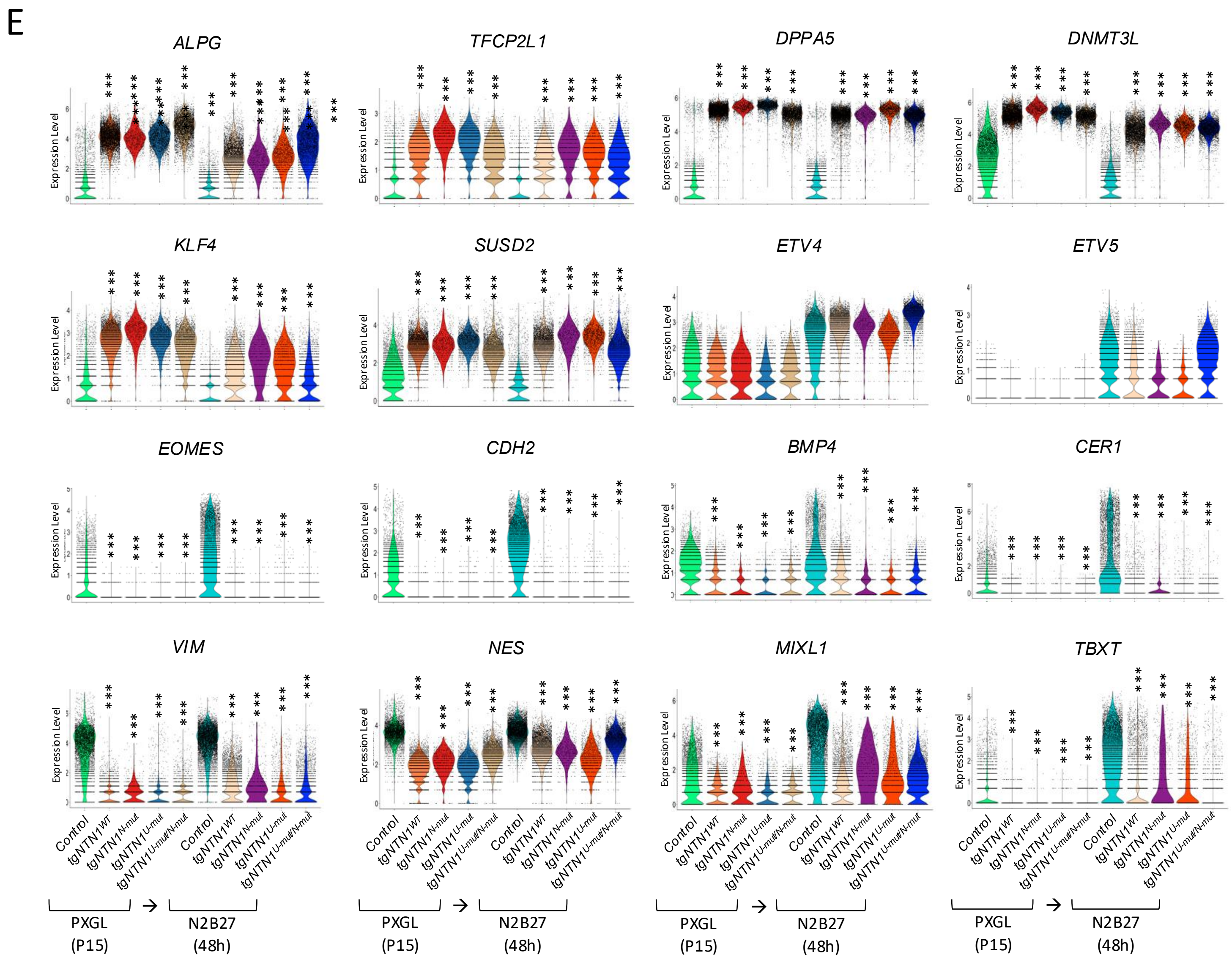

Supplementary Figure 6 (iii)

Supplemental Table S1_Sheet_1

| baseMean | log2FoldChange | lfcSE | stat | pvalue | padj | gene_id | gene_name |
|---|---|---|---|---|---|---|---|
| **169.5084524** | 2.309243341 | 0.146394521 | 15.42158769 | 1.17188E-53 | 5.91719E-50 | ENSG00000120498 | TEX11 |
| **203.8483427** | 1.41642242 | 0.191580866 | 7.375258871 | 1.64026E-13 | 2.76074E-11 | ENSG00000180447 | GAS1 |
| **188.5159699** | 1.375249374 | 0.136974319 | 9.974099777 | 1.97888E-23 | 1.42743E-20 | ENSG00000106004 | HOXA5 |
| **754.791098** | 1.369402831 | 0.15685119 | 8.739466669 | 2.34213E-18 | 8.25085E-16 | ENSG00000213763 | ACTBP2 |
| **109.5292025** | 1.290372626 | 0.171653724 | 7.450041282 | 9.3311E-14 | 1.62468E-11 | ENSG00000091656 | ZFHX4 |
| **42.91981181** | 1.278340768 | 0.203308134 | 6.131657425 | 8.69682E-10 | 6.5542E-08 | ENSG00000180340 | FZD2 |
| **13.76570903** | 1.259062828 | 0.241359789 | 4.933490011 | 8.07732E-07 | 2.81924E-05 | ENSG00000144355 | DLX1 |
| **121.1858494** | 1.257104487 | 0.182277982 | 6.850591234 | 7.35454E-12 | 8.9844E-10 | ENSG00000180867 | PDIA3P1 |
| **159.4337352** | 1.238298343 | 0.12034332 | 10.23194768 | 1.42622E-24 | 1.13707E-21 | ENSG00000101871 | MID1 |
| **228.8991121** | 1.224095787 | 0.147318842 | 8.262184073 | 1.43031E-16 | 4.1666E-14 | ENSG00000175745 | NR2F1 |
| **127.0302405** | 1.207682734 | 0.160081922 | 7.499782599 | 6.39238E-14 | 1.12595E-11 | ENSG00000116132 | PRRX1 |
| **422.3939764** | 1.158512871 | 0.129236209 | 8.947096031 | 3.64957E-19 | 1.38209E-16 | ENSG00000162599 | NFIA |
| **120.1732027** | 1.157158197 | 0.163038818 | 7.039812519 | 1.92499E-12 | 2.65088E-10 | ENSG00000176165 | FOXG1 |
| **654.5365339** | 1.154643006 | 0.123323259 | 9.348356391 | 8.90203E-21 | 4.64993E-18 | ENSG00000100234 | TIMP3 |
| **597.6151913** | 1.14296238 | 0.086416401 | 13.21400353 | 7.28446E-40 | 2.2069E-36 | ENSG00000185222 | TCEAL9 |
| **32.85223777** | 1.080665218 | 0.221114972 | 4.747175813 | 2.06277E-06 | 6.32526E-05 | ENSG00000143867 | OSR1 |
| **180.4944043** | 1.062264496 | 0.157076258 | 6.734910296 | 1.64031E-11 | 1.89675E-09 | ENSG00000248448 | COX5BP1 |
| **761.9846234** | 1.023441372 | 0.122570922 | 8.339259096 | 7.47574E-17 | 2.22344E-14 | ENSG00000185551 | NR2F2 |
| **122.3744278** | 0.968273186 | 0.159559712 | 6.042472042 | 1.51771E-09 | 1.08444E-07 | ENSG00000185052 | SLC24A3 |
| **37.35765168** | 0.945000467 | 0.222181003 | 4.173816189 | 2.99539E-05 | 0.00060905 | ENSG00000109846 | CRYAB |
| **828.5582167** | 0.928259347 | 0.112971042 | 8.208674438 | 2.23644E-16 | 6.15955E-14 | ENSG00000237676 | RPL30P4 |
| **1157.676829** | 0.922129869 | 0.100222274 | 9.198915862 | 3.61579E-20 | 1.61094E-17 | ENSG00000119396 | RAB14 |
| **35.3606338** | 0.918987465 | 0.199284437 | 4.512562276 | 6.40491E-06 | 0.000167857 | ENSG00000115507 | OTX1 |
| **64.35676435** | 0.916291719 | 0.181926431 | 4.99731452 | 5.81342E-07 | 2.13224E-05 | ENSG00000132967 | HMGB1P5 |
| **335.3504135** | 0.909342026 | 0.136450673 | 6.646029072 | 3.01106E-11 | 3.14562E-09 | ENSG00000134072 | CAMK1 |
| **19.36222917** | 0.906271753 | 0.234111072 | 3.747010719 | 0.000178954 | 0.002651669 | ENSG00000175505 | CLCF1 |
| **79.73747105** | 0.90052694 | 0.17385117 | 5.145986697 | 2.66118E-07 | 1.09246E-05 | ENSG00000179981 | TSHZ1 |
| **438.6420113** | 0.900504115 | 0.100406206 | 8.963338461 | 3.14994E-19 | 1.22347E-16 | ENSG00000138814 | PPP3CA |
| **234.4837367** | 0.884572449 | 0.125358034 | 7.043780644 | 1.87093E-12 | 2.60007E-10 | ENSG00000179195 | ZNF664 |
| **67.21319524** | 0.856137124 | 0.179284823 | 4.750418982 | 2.02996E-06 | 6.23728E-05 | ENSG00000185070 | FLRT2 |
| **65.02148975** | 0.846473054 | 0.185364666 | 4.549577546 | 5.37537E-06 | 0.000146979 | ENSG00000134363 | FST |
| **374.3597231** | 0.839326768 | 0.113906501 | 7.361648001 | 1.81653E-13 | 2.99096E-11 | ENSG00000171862 | PTEN |
| **293.8675825** | 0.828927637 | 0.13966999 | 5.920616594 | 3.20737E-09 | 2.15934E-07 | ENSG00000197956 | S100A6 |
| **438.9132666** | 0.82806301 | 0.176258405 | 4.688398568 | 2.75351E-06 | 8.11483E-05 | ENSG00000243709 | LEFTY1 |
| **3376.73191** | 0.822299221 | 0.122364799 | 6.711363849 | 1.92814E-11 | 2.19605E-09 | ENSG00000108821 | COL1A1 |
| **29.30676699** | 0.82031271 | 0.219337111 | 3.675218535 | 0.000237646 | 0.003328787 | ENSG00000214318 | ATP5MC1P6 |
| **109.4183429** | 0.818756156 | 0.143526688 | 5.685876255 | 1.30144E-08 | 7.54111E-07 | ENSG00000169439 | SDC2 |
| **404.1435846** | 0.818707156 | 0.10812068 | 7.569198865 | 3.75533E-14 | 7.29304E-12 | ENSG00000079102 | RUNX1T1 |
| **195.3097428** | 0.815744021 | 0.130833765 | 6.229937194 | 4.66622E-10 | 3.80021E-08 | ENSG00000075420 | FNDC3B |
| **327.3812127** | 0.814220447 | 0.117172916 | 6.943227375 | 3.83242E-12 | 4.91979E-10 | ENSG00000165410 | CFL2 |
| **502.6009224** | 0.809363021 | 0.103126788 | 7.847563848 | 4.24196E-15 | 9.05032E-13 | ENSG00000008294 | SPAG9 |
| **927.9201155** | 0.807568669 | 0.088031605 | 9.171265968 | 4.67496E-20 | 2.02332E-17 | ENSG00000162923 | WDR26 |
| **590.1234664** | 0.807554014 | 0.093475786 | 8.636283373 | 5.80714E-18 | 1.95481E-15 | ENSG00000005483 | KMT2E |

| | | | | | | | |
|---|---|---|---|---|---|---|---|
| **606.2630376** | 0.775670163 | 0.110148802 | 7.035164807 | 1.99026E-12 | 2.71607E-10 | ENSG00000111276 | CDKN1B |
| **1741.687258** | 0.775347103 | 0.088604971 | 8.748706817 | 2.15812E-18 | 7.7836E-16 | ENSG00000168036 | CTNNB1 |
| **2243.659633** | 0.774835826 | 0.104333366 | 7.422139847 | 1.15243E-13 | 1.98375E-11 | ENSG00000008441 | NFIX |
| **63.14891334** | 0.766740461 | 0.180352964 | 4.239216968 | 2.24301E-05 | 0.000475204 | ENSG00000159176 | CSRP1 |
| **178.9342974** | 0.761433478 | 0.131603022 | 5.778852049 | 7.5212E-09 | 4.66931E-07 | ENSG00000136451 | VEZF1 |
| **297.3456488** | 0.755340811 | 0.111276163 | 6.781991156 | 1.18531E-11 | 1.41378E-09 | ENSG00000043143 | JADE2 |
| **134.6086078** | 0.742850438 | 0.158191665 | 4.679306395 | 2.87847E-06 | 8.40136E-05 | ENSG00000231735 | AC092573.1 |
| **380.2328497** | 0.739580332 | 0.097258251 | 7.600283142 | 2.95484E-14 | 5.88945E-12 | ENSG00000145860 | RNF145 |
| **109.7285585** | 0.723600216 | 0.159589676 | 4.521293373 | 6.14629E-06 | 0.000163054 | ENSG00000173706 | HEG1 |
| **96.21988171** | 0.715078667 | 0.162536888 | 4.379792854 | 1.18792E-05 | 0.000282172 | ENSG00000184232 | OAF |
| **483.4953685** | 0.714344809 | 0.118858714 | 6.003364529 | 1.9327E-09 | 1.34915E-07 | ENSG00000213750 | AP002982.1 |
| **168.9930692** | 0.711533772 | 0.134364085 | 5.287119678 | 1.24257E-07 | 5.63548E-06 | ENSG00000130024 | PHF10 |
| **18.29061729** | 0.705295353 | 0.229422558 | 2.951725604 | 0.003160036 | 0.026652687 | ENSG00000017427 | IGF1 |
| **905.6626519** | 0.690587462 | 0.087353939 | 7.905449817 | 2.66968E-15 | 6.03586E-13 | ENSG00000186432 | KPNA4 |
| **967.2027403** | 0.681701049 | 0.097315334 | 7.005026741 | 2.46939E-12 | 3.28125E-10 | ENSG00000070961 | ATP2B1 |
| **20.89785995** | 0.675721075 | 0.224733691 | 2.955713336 | 0.003119469 | 0.026387026 | ENSG00000197757 | HOXC6 |
| **282.3394186** | 0.646096212 | 0.12319471 | 5.242333631 | 1.58558E-07 | 6.96186E-06 | ENSG00000109099 | PMP22 |
| **167.5500201** | 0.635183944 | 0.121201435 | 5.236905853 | 1.63291E-07 | 7.14893E-06 | ENSG00000178409 | BEND3 |
| **507.6852029** | 0.634218366 | 0.10951762 | 5.789672683 | 7.05237E-09 | 4.42603E-07 | ENSG00000133639 | BTG1 |
| **605.7111322** | 0.628559639 | 0.104237772 | 6.027671617 | 1.66339E-09 | 1.17743E-07 | ENSG00000157216 | SSBP3 |
| **771.2461409** | 0.619238631 | 0.091831634 | 6.742123451 | 1.56088E-11 | 1.81879E-09 | ENSG00000073921 | PICALM |
| **561.6523924** | 0.613741196 | 0.106762054 | 5.748458432 | 9.00608E-09 | 5.45696E-07 | ENSG00000138757 | G3BP2 |
| **215.983043** | 0.612479039 | 0.125220851 | 4.889355886 | 1.01166E-06 | 3.42069E-05 | ENSG00000118260 | CREB1 |
| **58.39451404** | 0.610865008 | 0.178780063 | 3.409774379 | 0.000650166 | 0.007660277 | ENSG00000205730 | ITPRIPL2 |
| **7568.390991** | 0.602576404 | 0.093015827 | 6.478155591 | 9.28506E-11 | 8.79063E-09 | ENSG00000140264 | SERF2 |
| **231.3384924** | 0.599846272 | 0.119148841 | 5.032533169 | 4.84041E-07 | 1.82339E-05 | ENSG00000115904 | 1 SOS |
| **363.5412955** | 0.587887491 | 0.094714018 | 6.206258406 | 5.42609E-10 | 4.31987E-08 | ENSG00000126351 | THRA |
| **1789.886568** | 0.574772568 | 0.095265784 | 6.033415326 | 1.6053E-09 | 1.14165E-07 | ENSG00000280623 | PCAT14 |
| **316.7519956** | 0.574057294 | 0.117129781 | 4.899174435 | 9.62402E-07 | 3.27606E-05 | ENSG00000143641 | GALNT2 |
| **577.6184852** | 0.570532242 | 0.086600297 | 6.586799349 | 4.49409E-11 | 4.53843E-09 | ENSG00000119048 | UBE2B |
| **46.94279186** | 0.570390504 | 0.189375205 | 3.00157292 | 0.002685887 | 0.023558666 | ENSG00000168785 | TSPAN5 |
| **574.5480861** | 0.563542046 | 0.120476576 | 4.675825427 | 2.92774E-06 | 8.49604E-05 | ENSG00000162433 | AK4 |
| **264.4431056** | 0.559687736 | 0.108683559 | 5.149334531 | 2.61412E-07 | 1.07605E-05 | ENSG00000213626 | LBH |
| **454.7397706** | 0.545509379 | 0.141879495 | 3.85080654 | 0.000117729 | 0.001882711 | ENSG00000130303 | BST2 |
| **1738.136097** | 0.543094537 | 0.087030022 | 6.239901946 | 4.37845E-10 | 3.62431E-08 | ENSG00000143933 | CALM2 |
| **414.2573106** | 0.540540349 | 0.105892817 | 5.103526628 | 3.33382E-07 | 1.32896E-05 | ENSG00000136826 | KLF4 |
| **578.2953057** | 0.537413892 | 0.094521468 | 5.685499044 | 1.30431E-08 | 7.54111E-07 | ENSG00000084676 | NCOA1 |
| **88.1101016** | 0.533599605 | 0.160751565 | 3.31706388 | 0.000909688 | 0.010056543 | ENSG00000162616 | DNAJB4 |
| **494.8996673** | 0.530740362 | 0.079522088 | 6.675810362 | 2.4587E-11 | 2.69887E-09 | ENSG00000189337 | KAZN |
| **128.9030096** | 0.52908121 | 0.140215941 | 3.772121251 | 0.000161866 | 0.002442171 | ENSG00000028528 | SNX1 |
| **318.5965865** | 0.527183498 | 0.108318327 | 4.866781538 | 1.1343E-06 | 3.75162E-05 | ENSG00000105245 | NUMBL |
| **237.8525326** | 0.521510055 | 0.107012902 | 4.873947711 | 1.0939E-06 | 3.63386E-05 | ENSG00000182197 | EXT1 |
| **1761.31311** | 0.515915255 | 0.098272935 | 5.250190491 | 1.51942E-07 | 6.71025E-06 | ENSG00000148154 | UGCG |
| **226.6290875** | 0.513648554 | 0.106787758 | 4.811093481 | 1.50107E-06 | 4.77693E-05 | ENSG00000109118 | PHF12 |
| **613.4879207** | 0.512359481 | 0.088711924 | 5.773750642 | 7.75261E-09 | 4.77385E-07 | ENSG00000100888 | CHD8 |

| | | | | | | | |
|---|---|---|---|---|---|---|---|
| **824.9561538** | 0.510926633 | 0.090641048 | 5.636729635 | 1.7331E-08 | 9.86954E-07 | ENSG00000101115 | SALL4 |
| **254.3083967** | 0.502813534 | 0.151892969 | 3.308289607 | 0.000938677 | 0.010303479 | ENSG00000181218 | H2AW |
| **511.3430434** | 0.501115727 | 0.105443341 | 4.752222949 | 2.01192E-06 | 6.19443E-05 | ENSG00000152518 | ZFP36L2 |
| **159.1940408** | 0.500460033 | 0.133730128 | 3.740507263 | 0.000183649 | 0.002711422 | ENSG00000132694 | ARHGEF11 |
| **1326.434041** | 0.494687309 | 0.113665424 | 4.351023833 | 1.35503E-05 | 0.000312421 | ENSG00000125304 | TM9SF2 |
| **529.6988377** | 0.488714527 | 0.09549131 | 5.117302973 | 3.09936E-07 | 1.24864E-05 | ENSG00000104517 | UBR5 |
| **168.5721763** | 0.488518509 | 0.119725616 | 4.078709242 | 4.52864E-05 | 0.000847556 | ENSG00000162997 | PRORSD1P |
| **643.8996527** | 0.4774279 | 0.109859586 | 4.345385222 | 1.39031E-05 | 0.000319098 | ENSG00000124225 | PMEPA1 |
| **147.7798918** | 0.473092193 | 0.1542402 | 3.066881015 | 0.002163049 | 0.019858104 | ENSG00000118007 | STAG1 |
| **258.9992553** | 0.470009955 | 0.111359976 | 4.222126216 | 2.42009E-05 | 0.000506346 | ENSG00000156026 | MCU |
| **190.3310759** | 0.467368762 | 0.122976351 | 3.800917272 | 0.000144161 | 0.002217013 | ENSG00000087095 | NLK |
| **454.9550529** | 0.465004601 | 0.099064448 | 4.69367536 | 2.6834E-06 | 7.92361E-05 | ENSG00000127483 | HP1BP3 |
| **1126.823593** | 0.46402244 | 0.08479097 | 5.472378792 | 4.44034E-08 | 2.23463E-06 | ENSG00000185650 | ZFP36L1 |
| **578.7959356** | 0.457322058 | 0.108486795 | 4.214417703 | 2.50423E-05 | 0.000521073 | ENSG00000181555 | SETD2 |
| **417.9550276** | 0.456082592 | 0.078218635 | 5.832566578 | 5.45812E-09 | 3.48859E-07 | ENSG00000139718 | SETD1B |
| **153.3441778** | 0.448263071 | 0.146677715 | 3.056057498 | 0.002242682 | 0.020465153 | ENSG00000078269 | SYNJ2 |
| **492.0588854** | 0.446257469 | 0.101441231 | 4.399503144 | 1.08499E-05 | 0.000265088 | ENSG00000066735 | KIF26A |
| **7527.281272** | 0.444765389 | 0.06801691 | 6.538964125 | 6.19464E-11 | 6.05396E-09 | ENSG00000110321 | EIF4G2 |
| **161.7486306** | 0.442466541 | 0.13936953 | 3.175266385 | 0.00149699 | 0.014948191 | ENSG00000139163 | ETNK1 |
| **301.611062** | 0.438510712 | 0.10781539 | 4.069779443 | 4.70577E-05 | 0.000874637 | ENSG00000108389 | MTMR4 |
| **176.1626949** | 0.435654829 | 0.12557293 | 3.472731794 | 0.00051519 | 0.006344794 | ENSG00000134138 | MEIS2 |
| **228.0128521** | 0.435634163 | 0.118989602 | 3.662813711 | 0.00024946 | 0.00345236 | ENSG00000055208 | TAB2 |
| **710.1393256** | 0.435444935 | 0.072333286 | 6.016599256 | 1.78119E-09 | 1.24914E-07 | ENSG00000072364 | AFF4 |
| **582.5893516** | 0.430329006 | 0.080288147 | 5.360806467 | 8.28512E-08 | 3.94664E-06 | ENSG00000183283 | DAZAP2 |
| **604.0147938** | 0.428029314 | 0.082457852 | 5.192262751 | 2.07754E-07 | 8.86493E-06 | ENSG00000153317 | ASAP1 |
| **1139.672727** | 0.425719646 | 0.07380354 | 5.768811517 | 7.98325E-09 | 4.87622E-07 | ENSG00000153250 | RBMS1 |
| **1446.958351** | 0.421757266 | 0.12772944 | 3.303847429 | 0.000953677 | 0.010423017 | ENSG00000167004 | PDIA3 |
| **3470.467441** | 0.41656249 | 0.064503146 | 6.4581743 | 1.05974E-10 | 9.8484E-09 | ENSG00000141367 | CLTC |
| **106.2071268** | 0.412779109 | 0.147919911 | 2.790505734 | 0.005262577 | 0.038810863 | ENSG00000115539 | PDCL3 |
| **710.6520794** | 0.410962134 | 0.099692226 | 4.122058462 | 3.75502E-05 | 0.000729244 | ENSG00000119669 | IRF2BPL |
| **395.8404553** | 0.410722003 | 0.097417096 | 4.219136416 | 2.4524E-05 | 0.000511693 | ENSG00000129245 | FXR2 |
| **1198.908888** | 0.410252939 | 0.104643506 | 3.919937866 | 8.85718E-05 | 0.001503128 | ENSG00000233056 | ERVH48-1 |
| **353.9004556** | 0.406814687 | 0.098592083 | 4.128167984 | 3.65665E-05 | 0.000716571 | ENSG00000171729 | TMEM51 |
| **162.4421974** | 0.404645979 | 0.119825486 | 3.379044451 | 0.000727382 | 0.008438065 | ENSG00000147050 | KDM6A |
| **3012.713587** | 0.400010046 | 0.07417261 | 5.393038256 | 6.92762E-08 | 3.36345E-06 | ENSG00000099783 | HNRNPM |
| **1365.454105** | 0.398944299 | 0.07405437 | 5.387756662 | 7.13426E-08 | 3.45271E-06 | ENSG00000170027 | YWHAG |
| **307.0715828** | 0.397577954 | 0.120143725 | 3.309691205 | 0.000933989 | 0.010274563 | ENSG00000166794 | PPIB |
| **418.2651324** | 0.396344188 | 0.099320571 | 3.991161203 | 6.57506E-05 | 0.001166264 | ENSG00000134910 | STT3A |
| **1347.834127** | 0.396255542 | 0.107995368 | 3.669392482 | 0.000243128 | 0.003385015 | ENSG00000090621 | PABPC4 |
| **225.7982574** | 0.393886795 | 0.11051821 | 3.562606342 | 0.000367191 | 0.004794284 | ENSG00000072803 | FBXW11 |
| **339.8991891** | 0.391074325 | 0.116617332 | 3.354850993 | 0.000794077 | 0.009057745 | ENSG00000140262 | TCF12 |
| **534.9687462** | 0.385100667 | 0.096829392 | 3.979444627 | 6.90764E-05 | 0.001220969 | ENSG00000160712 | IL6R |
| **225.3072231** | 0.380033425 | 0.113038522 | 3.359852708 | 0.00077984 | 0.008922775 | ENSG00000157212 | PAXIP1 |
| **806.5477324** | 0.378628333 | 0.104510485 | 3.623251107 | 0.000290923 | 0.003927722 | ENSG00000269842 | AC011453.1 |
| **495.8020287** | 0.360955262 | 0.093392965 | 3.867704143 | 0.000109865 | 0.001787575 | ENSG00000156639 | ZFAND3 |

| | | | | | | | |
|---|---|---|---|---|---|---|---|
| **518.7911037** | 0.360543536 | 0.092965764 | 3.876829209 | 0.000105827 | 0.001733039 | ENSG00000089057 | SLC23A2 |
| **1076.780145** | 0.340305518 | 0.067507403 | 5.041190401 | 4.62645E-07 | 1.77197E-05 | ENSG00000115524 | SF3B1 |
| **369.9286621** | 0.335668942 | 0.090983433 | 3.686604306 | 0.000227266 | 0.003220421 | ENSG00000107863 | ARHGAP21 |
| **413.631454** | 0.332477367 | 0.089982534 | 3.695404896 | 0.000219537 | 0.003122574 | ENSG00000004534 | RBM6 |
| **227.3516969** | 0.328316364 | 0.115012842 | 2.853781418 | 0.004320225 | 0.033508838 | ENSG00000106591 | MRPL32 |
| **213.794905** | 0.3235377 | 0.113390878 | 2.8490102 | 0.004385547 | 0.033859466 | ENSG00000197134 | ZNF257 |
| **894.0611801** | 0.32111822 | 0.097270477 | 3.30237177 | 0.000958709 | 0.010455382 | ENSG00000159692 | CTBP1 |
| **483.2183531** | 0.318861978 | 0.101354355 | 3.147342895 | 0.001647616 | 0.016156221 | ENSG00000108312 | UBTF |
| **550.6257212** | 0.318089276 | 0.077252238 | 4.11977656 | 3.7924E-05 | 0.00073462 | ENSG00000160714 | UBE2Q1 |
| **569.0179965** | 0.311078244 | 0.087273794 | 3.559238795 | 0.000371931 | 0.00484438 | ENSG00000131504 | DIAPH1 |
| **2013.83448** | 0.306106429 | 0.087257612 | 3.508388206 | 0.000450831 | 0.005693138 | ENSG00000173402 | DAG1 |
| **362.0661822** | 0.299091803 | 0.093910251 | 3.18605767 | 0.001442258 | 0.014526151 | ENSG00000094804 | CDC6 |
| **614.0092549** | 0.292966359 | 0.096373621 | 3.040317947 | 0.002363285 | 0.021410911 | ENSG00000184432 | COPB2 |
| **250.851072** | 0.292507327 | 0.105476015 | 2.778640986 | 0.005458682 | 0.039830498 | ENSG00000138646 | HERC5 |
| **334.898824** | 0.288709566 | 0.096905979 | 2.983172922 | 0.002852768 | 0.024696617 | ENSG00000185619 | PCGF3 |
| **495.6320147** | 0.284469375 | 0.098289664 | 2.894615898 | 0.003796227 | 0.030571639 | ENSG00000102580 | DNAJC3 |
| **1473.589228** | 0.282039282 | 0.059466373 | 4.742907346 | 2.10673E-06 | 6.44701E-05 | ENSG00000100811 | YY1 |
| **312.4362004** | 0.279889468 | 0.101954255 | 2.74308737 | 0.006086449 | 0.043053036 | ENSG00000068323 | TFE3 |
| **341.679087** | 0.270724889 | 0.09224605 | 2.936474107 | 0.003319665 | 0.027690689 | ENSG00000166411 | IDH3A |
| **7603.809577** | 0.270506116 | 0.081514284 | 3.318515052 | 0.000904974 | 0.010020872 | ENSG00000166598 | HSP90B1 |
| **851.1642306** | 0.269420931 | 0.070722764 | 3.81113367 | 0.000138331 | 0.002149166 | ENSG00000084733 | RAB10 |
| **345.187563** | 0.261629521 | 0.094092942 | 2.785068879 | 0.005351635 | 0.039295479 | ENSG00000055130 | CUL1 |
| **311.2443036** | 0.261550262 | 0.097689541 | 2.677256547 | 0.007422777 | 0.049864178 | ENSG00000116001 | TIA1 |
| **1104.122553** | 0.257388152 | 0.078184959 | 3.293603207 | 0.00098912 | 0.01073583 | ENSG00000086061 | DNAJA1 |
| **9200.270443** | 0.234179605 | 0.077500496 | 3.021597267 | 0.002514448 | 0.022418402 | ENSG00000044574 | HSPA5 |
| **3667.599004** | 0.231921697 | 0.052798587 | 4.393286885 | 1.1165E-05 | 0.000269795 | ENSG00000143761 | ARF1 |
| **2409.912714** | 0.231861194 | 0.060679211 | 3.822009048 | 0.000132369 | 0.002071408 | ENSG00000100345 | MYH9 |
| **1964.7293** | 0.214630647 | 0.073436836 | 2.92203476 | 0.003477527 | 0.028695779 | ENSG00000115306 | SPTBN1 |
| **751.8512159** | -0.205043474 | 0.075011944 | -2.73926087 | 0.006157749 | 0.043405112 | ENSG00000158246 | TENT5B |
| **1090.809779** | -0.258564741 | 0.094474603 | -2.74247577 | 0.006097795 | 0.04309719 | ENSG00000100097 | LGALS1 |
| **697.9851514** | -0.260840365 | 0.083753917 | -3.116255932 | 0.001831632 | 0.017560485 | ENSG00000188112 | C6orf132 |
| **1925.997958** | -0.262408263 | 0.058029446 | -4.520217324 | 6.17762E-06 | 0.000163313 | ENSG00000128739 | SNRPN |
| **2267.428184** | -0.264270346 | 0.06761609 | -3.907759552 | 9.31559E-05 | 0.001569662 | ENSG00000118816 | CCNI |
| **30015.49287** | -0.26553587 | 0.079678529 | -3.331979461 | 0.000862306 | 0.00965426 | ENSG00000137154 | RPS6 |
| **482.9867076** | -0.271578155 | 0.078955885 | -3.436538128 | 0.000589199 | 0.007134447 | ENSG00000181924 | COA4 |
| **457.4634221** | -0.277359014 | 0.097251282 | -2.84881751 | 0.004388204 | 0.03386272 | ENSG00000179085 | DPM3 |
| **1097.812283** | -0.283785858 | 0.074909682 | -3.789759259 | 0.000150793 | 0.002301251 | ENSG00000134684 | YARS1 |
| **536.5679895** | -0.296430253 | 0.080909058 | -3.66490121 | 0.000247434 | 0.003432355 | ENSG00000164904 | ALDH7A1 |
| **491.6595076** | -0.29997407 | 0.080152006 | -3.73964498 | 0.00018428 | 0.002715446 | ENSG00000142687 | KIAA0319L |
| **444.9621602** | -0.307645912 | 0.088784116 | -3.46000087 | 0.000540174 | 0.006630916 | ENSG00000076554 | TPD52 |
| **408.3210831** | -0.316799859 | 0.102913133 | -3.083171957 | 0.002048068 | 0.019044893 | ENSG00000119408 | NEK6 |
| **3607.86119** | -0.329752251 | 0.074642386 | -4.417399386 | 9.98955E-06 | 0.000246052 | ENSG00000105202 | FBL |
| **337.4585994** | -0.331254275 | 0.09876893 | -3.352126948 | 0.000801932 | 0.009126725 | ENSG00000188677 | PARVB |
| **2845.199008** | -0.333914202 | 0.112400089 | -2.9685905 | 0.00299169 | 0.025545727 | ENSG00000244245 | AC133134.1 |
| **2914.327338** | -0.334191002 | 0.080312783 | -4.159170836 | 3.19405E-05 | 0.000642503 | ENSG00000232573 | RPL3P4 |

| | | | | | | | |
|---|---|---|---|---|---|---|---|
| **1114.047943** | -0.338465297 | 0.071440001 | -4.737981617 | 2.15857E-06 | 6.57909E-05 | ENSG00000105373 | NOP53 |
| **893.8044571** | -0.341994852 | 0.077940073 | -4.389442491 | 1.13642E-05 | 0.000273245 | ENSG00000077312 | SNRPA |
| **1745.003834** | -0.348291384 | 0.088052598 | -3.954821245 | 7.65919E-05 | 0.001338194 | ENSG00000172053 | QARS1 |
| **709.7890675** | -0.351627087 | 0.081550097 | -4.315800038 | 1.59026E-05 | 0.000359005 | ENSG00000137876 | RSL24D1 |
| **184.3620483** | -0.353742248 | 0.126733959 | -2.805271699 | 0.005027418 | 0.037478017 | ENSG00000106603 | COA1 |
| **337.3765095** | -0.355369696 | 0.096123977 | -3.695709514 | 0.000219274 | 0.003121763 | ENSG00000168256 | NKIRAS2 |
| **241.9214385** | -0.361321933 | 0.123894026 | -2.912228301 | 0.003588602 | 0.029415665 | ENSG00000168569 | TMEM223 |
| **1369.824064** | -0.363795894 | 0.083615083 | -4.352428634 | 1.34638E-05 | 0.000311373 | ENSG00000185201 | IFITM2 |
| **2554.494926** | -0.383860175 | 0.062701368 | -6.122804767 | 9.19423E-10 | 6.82717E-08 | ENSG00000167460 | TPM4 |
| **768.849616** | -0.385202432 | 0.108925053 | -3.53409136 | 0.00040918 | 0.005257212 | ENSG00000215845 | TSTD1 |
| **1204.416119** | -0.391092608 | 0.068142017 | -5.735166205 | 9.74168E-09 | 5.78694E-07 | ENSG00000140983 | RHOT2 |
| **5290.473835** | -0.391145253 | 0.090608954 | -4.316598713 | 1.58452E-05 | 0.000358243 | ENSG00000197958 | RPL12 |
| **346.2168612** | -0.395820001 | 0.113396595 | -3.480995755 | 0.000499553 | 0.006187437 | ENSG00000219222 | RPL12P47 |
| **384.1605686** | -0.398280865 | 0.109752258 | -3.628755294 | 0.000284791 | 0.003862144 | ENSG00000134809 | TIMM10 |
| **193.8812604** | -0.403329397 | 0.138051601 | -2.915164673 | 0.003555009 | 0.029203513 | ENSG00000106384 | MOGAT3 |
| **243.7311473** | -0.40806148 | 0.10828697 | -3.766362082 | 0.000165644 | 0.002494203 | ENSG00000100814 | CCNB1IP1 |
| **512.0151701** | -0.40964617 | 0.10828338 | -3.773524566 | 0.000160957 | 0.002435746 | ENSG00000223803 | RPS20P14 |
| **1656.027445** | -0.415870921 | 0.089401559 | -4.654908889 | 3.24124E-06 | 9.24638E-05 | ENSG00000103257 | SLC7A5 |
| **274.1420682** | -0.417563727 | 0.107034604 | -3.907906322 | 9.30994E-05 | 0.001569662 | ENSG00000187840 | EIF4EBP1 |
| **402.5004896** | -0.440751172 | 0.114710166 | -3.846165816 | 0.000119981 | 0.0019071 | ENSG00000222041 | CYTOR |
| **545.9095632** | -0.442388729 | 0.088586294 | -4.995410497 | 5.87107E-07 | 2.14301E-05 | ENSG00000126749 | EMG1 |
| **408.106098** | -0.452361601 | 0.09700513 | -4.685352899 | 2.79478E-06 | 8.22045E-05 | ENSG00000198093 | ZNF649 |
| **253.9476697** | -0.453545974 | 0.119981562 | -3.774889082 | 0.000160079 | 0.002424875 | ENSG00000164284 | GRPEL2 |
| **332.7442574** | -0.458163324 | 0.106347246 | -4.302116175 | 1.69175E-05 | 0.000377973 | ENSG00000084092 | NOA1 |
| **115.667217** | -0.458918289 | 0.152273192 | -3.087701888 | 0.002017107 | 0.018861196 | ENSG00000136010 | ALDH1L2 |
| **1060.740281** | -0.472057475 | 0.088215513 | -5.350755628 | 8.75877E-08 | 4.13327E-06 | ENSG00000092621 | PHGDH |
| **423.795961** | -0.475378583 | 0.087214126 | -5.441960836 | 5.26973E-08 | 2.63452E-06 | ENSG00000120699 | EXOSC8 |
| **1302.822233** | -0.481263241 | 0.079360984 | -6.065110048 | 1.31863E-09 | 9.51174E-08 | ENSG00000182199 | SHMT2 |
| **1110.445723** | -0.483238707 | 0.098995348 | -4.881038115 | 1.05529E-06 | 3.54446E-05 | ENSG00000112667 | DNPH1 |
| **432.0407195** | -0.483486623 | 0.109536948 | -4.418335859 | 9.94638E-06 | 0.000245387 | ENSG00000143367 | TUFT1 |
| **241.5927578** | -0.484863483 | 0.116515459 | -4.154489327 | 3.26015E-05 | 0.000653237 | ENSG00000118162 | KPTN |
| **1908.01905** | -0.485965662 | 0.08187681 | -5.932476497 | 2.98399E-09 | 2.02697E-07 | ENSG00000090861 | AARS1 |
| **376.9365174** | -0.49468135 | 0.135663623 | -3.662865088 | 0.00024941 | 0.00345236 | ENSG00000244187 | TMEM141 |
| **81.849748** | -0.502414403 | 0.174619322 | -2.880506722 | 0.003970365 | 0.031541419 | ENSG00000105472 | CLEC11A |
| **382.40924** | -0.505081546 | 0.100251259 | -5.040948591 | 4.6323E-07 | 1.77197E-05 | ENSG00000125520 | SLC2A4RG |
| **437.768864** | -0.509284946 | 0.095090846 | -5.348424645 | 8.87231E-08 | 4.17384E-06 | ENSG00000141582 | CBX4 |
| **404.3428663** | -0.514689158 | 0.116587198 | -4.426723183 | 9.56754E-06 | 0.00023837 | ENSG00000105370 | LIM2 |
| **309.2754196** | -0.5168576 | 0.128424925 | -4.025703298 | 5.68053E-05 | 0.001024389 | ENSG00000154328 | NEIL2 |
| **1620.738487** | -0.5182725 | 0.104334758 | -4.965099999 | 6.86657E-07 | 2.43594E-05 | ENSG00000166986 | MARS1 |
| **1532.786351** | -0.528149206 | 0.120633075 | -4.376641498 | 1.20522E-05 | 0.000284372 | ENSG00000105281 | SLC1A5 |
| **67.11137967** | -0.557289137 | 0.173413703 | -3.204164827 | 0.001354549 | 0.013789455 | ENSG00000078814 | MYH7B |
| **1307.739565** | -0.566934327 | 0.084563987 | -6.699455334 | 2.09198E-11 | 2.36487E-09 | ENSG00000042493 | CAPG |
| **40.61324107** | -0.587504667 | 0.201259269 | -2.911518176 | 0.00359677 | 0.029419844 | ENSG00000169964 | TMEM42 |
| **114.9253737** | -0.620958124 | 0.164922866 | -3.759685629 | 0.000170127 | 0.002546526 | ENSG00000100300 | TSPO |
| **115.9319555** | -0.634200742 | 0.157429277 | -4.114721746 | 3.87647E-05 | 0.000747083 | ENSG00000130707 | ASS1 |

| | | | | | | | |
|---|---|---|---|---|---|---|---|
| **1948.791436** | -0.641344022 | 0.067865275 | -9.448490131 | 3.43753E-21 | 2.00276E-18 | ENSG00000108828 | VAT1 |
| **40.48374892** | -0.645735167 | 0.200142882 | -3.175976658 | 0.00149333 | 0.014921476 | ENSG00000188505 | NCCRP1 |
| **745.1629934** | -0.678277329 | 0.090218612 | -7.519922839 | 5.48086E-14 | 1.00029E-11 | ENSG00000151012 | SLC7A11 |
| **281.5626158** | -0.694147385 | 0.107454965 | -6.44642145 | 1.14522E-10 | 1.05138E-08 | ENSG00000153044 | CENPH |
| **63.72669464** | -0.698897967 | 0.195168751 | -3.706573419 | 0.000210082 | 0.003013568 | ENSG00000183421 | RIPK4 |
| **1220.609946** | -0.709940189 | 0.087167349 | -8.150551636 | 3.62268E-16 | 9.30108E-14 | ENSG00000135069 | PSAT1 |
| **676.5434062** | -0.72685682 | 0.122765909 | -5.916062345 | 3.2974E-09 | 2.21014E-07 | ENSG00000128965 | CHAC1 |
| **58.00508689** | -0.734852684 | 0.188153387 | -3.864567479 | 0.000111286 | 0.001806821 | ENSG00000130270 | ATP8B3 |
| **140.1203151** | -0.742144384 | 0.146710198 | -5.091184958 | 3.55833E-07 | 1.41474E-05 | ENSG00000128165 | ADM2 |
| **369.5016608** | -0.750416513 | 0.123599875 | -6.058014654 | 1.37812E-09 | 9.89371E-08 | ENSG00000176046 | NUPR1 |
| **68.03450448** | -0.764574566 | 0.168213483 | -4.493988713 | 6.99013E-06 | 0.000181003 | ENSG00000108641 | B9D1 |
| **63.58257054** | -0.800780934 | 0.180061488 | -4.430447981 | 9.40375E-06 | 0.000234676 | ENSG00000027869 | SH2D2A |
| **244.0788983** | -0.802115068 | 0.121495695 | -6.617583373 | 3.65118E-11 | 3.76246E-09 | ENSG00000180353 | HCLS1 |
| **1076.948496** | -0.804907057 | 0.069209003 | -11.6250586 | 3.07382E-31 | 3.58171E-28 | ENSG00000183010 | PYCR1 |
| **212.6874674** | -0.805956691 | 0.166087176 | -4.874757928 | 1.08942E-06 | 3.63364E-05 | ENSG00000109107 | ALDOC |
| **64.45304812** | -0.811779505 | 0.187982351 | -4.38602858 | 1.15439E-05 | 0.000276688 | ENSG00000272512 | AL645608.7 |
| **323.6159618** | -0.867063592 | 0.101708581 | -8.504018953 | 1.83138E-17 | 5.90249E-15 | ENSG00000104814 | MAP4K1 |
| **225.1428502** | -1.166267481 | 0.127218082 | -9.127230478 | 7.0274E-20 | 2.87706E-17 | ENSG00000170627 | GTSF1 |
| **23.33318236** | -1.169161478 | 0.219560937 | -5.032551398 | 4.83995E-07 | 1.82339E-05 | ENSG00000172794 | RAB37 |
| **113.9826011** | -1.811300077 | 0.179903327 | -9.608866301 | 7.33521E-22 | 4.44455E-19 | ENSG00000248370 | LINC02434 |

Supplemental Table S1_Sheet_2

| baseMean | log2FoldChange | lfcSE | stat | pvalue | padj | gene_id | gene_name |
|---|---|---|---|---|---|---|---|
| **169.5084524** | 2.309243341 | 0.146394521 | 15.42158769 | 1.17188E-53 | 5.91719E-50 | ENSG00000120498 | TEX11 |
| **47.88628914** | 1.471668706 | 0.190762877 | 7.517557942 | 5.58088E-14 | 1.00642E-11 | ENSG00000184368 | MAP7D2 |
| **203.8483427** | 1.41642242 | 0.191580866 | 7.375258871 | 1.64026E-13 | 2.76074E-11 | ENSG00000180447 | GAS1 |
| **188.5159699** | 1.375249374 | 0.136974319 | 9.974099777 | 1.97888E-23 | 1.42743E-20 | ENSG00000106004 | HOXA5 |
| **754.791098** | 1.369402831 | 0.15685119 | 8.739466669 | 2.34213E-18 | 8.25085E-16 | ENSG00000213763 | ACTBP2 |
| **109.5292025** | 1.290372626 | 0.171653724 | 7.450041282 | 9.3311E-14 | 1.62468E-11 | ENSG00000091656 | ZFHX4 |
| **42.91981181** | 1.278340768 | 0.203308134 | 6.131657425 | 8.69682E-10 | 6.5542E-08 | ENSG00000180340 | FZD2 |
| **13.76570903** | 1.259062828 | 0.241359789 | 4.933490011 | 8.07732E-07 | 2.81924E-05 | ENSG00000144355 | DLX1 |
| **121.1858494** | 1.257104487 | 0.182277982 | 6.850591234 | 7.35454E-12 | 8.9844E-10 | ENSG00000180867 | PDIA3P1 |
| **159.4337352** | 1.238298343 | 0.12034332 | 10.23194768 | 1.42622E-24 | 1.13707E-21 | ENSG00000101871 | MID1 |
| **228.8991121** | 1.224095787 | 0.147318842 | 8.262184073 | 1.43031E-16 | 4.1666E-14 | ENSG00000175745 | NR2F1 |
| **127.0302405** | 1.207682734 | 0.160081922 | 7.499782599 | 6.39238E-14 | 1.12595E-11 | ENSG00000116132 | PRRX1 |
| **71.99789622** | 1.20366444 | 0.178785028 | 6.678622557 | 2.41198E-11 | 2.66692E-09 | ENSG00000215105 | TTC3P1 |
| **60.214064** | 1.187973142 | 0.172403118 | 6.791856089 | 1.107E-11 | 1.34151E-09 | ENSG00000094841 | UPRT |
| **64.36924682** | 1.162589343 | 0.185527027 | 6.16403702 | 7.09134E-10 | 5.42523E-08 | ENSG00000174358 | SLC6A19 |
| **422.3939764** | 1.158512871 | 0.129236209 | 8.947096031 | 3.64957E-19 | 1.38209E-16 | ENSG00000162599 | NFIA |
| **120.1732027** | 1.157158197 | 0.163038818 | 7.039812519 | 1.92499E-12 | 2.65088E-10 | ENSG00000176165 | FOXG1 |
| **654.5365339** | 1.154643006 | 0.123323259 | 9.348356391 | 8.90203E-21 | 4.64993E-18 | ENSG00000100234 | TIMP3 |
| **597.6151913** | 1.14296238 | 0.086416401 | 13.21400353 | 7.28446E-40 | 2.2069E-36 | ENSG00000185222 | TCEAL9 |
| **313.7210927** | 1.139961494 | 0.165596602 | 6.876982011 | 6.11338E-12 | 7.65335E-10 | ENSG00000256045 | MTRNR2L10 |
| **63.95151286** | 1.129521836 | 0.180725547 | 6.200703827 | 5.62112E-10 | 4.43483E-08 | ENSG00000126016 | AMOT |
| **48.92745451** | 1.126245703 | 0.199366636 | 5.572249413 | 2.51471E-08 | 1.36046E-06 | ENSG00000164330 | EBF1 |
| **89.99676501** | 1.092371942 | 0.161145957 | 6.718141264 | 1.84057E-11 | 2.1122E-09 | ENSG00000144810 | COL8A1 |
| **133.0953939** | 1.090316073 | 0.152332154 | 7.137658504 | 9.49341E-13 | 1.39618E-10 | ENSG00000150938 | CRIM1 |
| **32.85223777** | 1.080665218 | 0.221114972 | 4.747175813 | 2.06277E-06 | 6.32526E-05 | ENSG00000143867 | OSR1 |
| **446.7971905** | 1.075625382 | 0.11053688 | 9.725723926 | 2.34233E-22 | 1.54268E-19 | ENSG00000171365 | CLCN5 |
| **180.4944043** | 1.062264496 | 0.157076258 | 6.734910296 | 1.64031E-11 | 1.89675E-09 | ENSG00000248448 | COX5BP1 |
| **20.02581585** | 1.032945139 | 0.238392586 | 4.25178375 | 2.12075E-05 | 0.000456322 | ENSG00000115844 | DLX2 |
| **761.9846234** | 1.023441372 | 0.122570922 | 8.339259096 | 7.47574E-17 | 2.22344E-14 | ENSG00000185551 | NR2F2 |
| **172.1152938** | 1.02151982 | 0.129271469 | 7.890806456 | 3.0024E-15 | 6.59136E-13 | ENSG00000101220 | C20orf27 |
| **3251.141225** | 1.018803324 | 0.116001791 | 8.776433719 | 1.6874E-18 | 6.23433E-16 | ENSG00000168542 | COL3A1 |
| **414.9363788** | 1.009190868 | 0.095632636 | 10.54139777 | 5.56653E-26 | 5.27011E-23 | ENSG00000101928 | MOSPD1 |
| **122.3744278** | 0.968273186 | 0.159559712 | 6.042472042 | 1.51771E-09 | 1.08444E-07 | ENSG00000185052 | SLC24A3 |
| **220.5161242** | 0.963361627 | 0.145898511 | 6.59103502 | 4.36771E-11 | 4.44041E-09 | ENSG00000165194 | PCDH19 |
| **37.35765168** | 0.945000467 | 0.222181003 | 4.173816189 | 2.99539E-05 | 0.00060905 | ENSG00000109846 | CRYAB |
| **371.455881** | 0.929916593 | 0.111280542 | 8.346078021 | 7.05672E-17 | 2.18153E-14 | ENSG00000196628 | TCF4 |
| **828.5582167** | 0.928259347 | 0.112971042 | 8.208674438 | 2.23644E-16 | 6.15955E-14 | ENSG00000237676 | RPL30P4 |
| **1157.676829** | 0.922129869 | 0.100222274 | 9.198915862 | 3.61579E-20 | 1.61094E-17 | ENSG00000119396 | RAB14 |
| **35.3606338** | 0.918987465 | 0.199284437 | 4.512562276 | 6.40491E-06 | 0.000167857 | ENSG00000115507 | OTX1 |
| **64.35676435** | 0.916291719 | 0.181926431 | 4.99731452 | 5.81342E-07 | 2.13224E-05 | ENSG00000132967 | HMGB1P5 |
| **335.3504135** | 0.909342026 | 0.136450673 | 6.646029072 | 3.01106E-11 | 3.14562E-09 | ENSG00000134072 | CAMK1 |
| **896.9927837** | 0.907961876 | 0.072670573 | 12.490541 | 8.408E-36 | 1.41516E-32 | ENSG00000180964 | TCEAL8 |
| **19.36222917** | 0.906271753 | 0.234111072 | 3.747010719 | 0.000178954 | 0.002651669 | ENSG00000175505 | CLCF1 |

| | | | | | | | |
|---|---|---|---|---|---|---|---|
| **93.8978836** | 0.905713864 | 0.157120749 | 5.73809471 | 9.57476E-09 | 5.75068E-07 | ENSG00000165655 | ZNF503 |
| **79.73747105** | 0.90052694 | 0.17385117 | 5.145986697 | 2.66118E-07 | 1.09246E-05 | ENSG00000179981 | TSHZ1 |
| **438.6420113** | 0.900504115 | 0.100406206 | 8.963338461 | 3.14994E-19 | 1.22347E-16 | ENSG00000138814 | PPP3CA |
| **234.4837367** | 0.884572449 | 0.125358034 | 7.043780644 | 1.87093E-12 | 2.60007E-10 | ENSG00000179195 | ZNF664 |
| **1252.686379** | 0.862574693 | 0.089471671 | 9.638042134 | 5.52313E-22 | 3.48602E-19 | ENSG00000163565 | IFI16 |
| **18.78405236** | 0.858018548 | 0.234300973 | 3.537663198 | 0.000403685 | 0.005199842 | ENSG00000081479 | LRP2 |
| **67.21319524** | 0.856137124 | 0.179284823 | 4.750418982 | 2.02996E-06 | 6.23728E-05 | ENSG00000185070 | FLRT2 |
| **154.4437013** | 0.849117942 | 0.155420805 | 5.474769591 | 4.38081E-08 | 2.21202E-06 | ENSG00000165349 | SLC7A3 |
| **65.02148975** | 0.846473054 | 0.185364666 | 4.549577546 | 5.37537E-06 | 0.000146979 | ENSG00000134363 | FST |
| **41.97909165** | 0.84379988 | 0.207424671 | 4.060378401 | 4.89932E-05 | 0.000903958 | ENSG00000020577 | SAMD4A |
| **273.2937409** | 0.8432022 | 0.118561979 | 7.107576058 | 1.18099E-12 | 1.72015E-10 | ENSG00000187079 | TEAD1 |
| **374.3597231** | 0.839326768 | 0.113906501 | 7.361648001 | 1.81653E-13 | 2.99096E-11 | ENSG00000171862 | PTEN |
| **60.94113598** | 0.838034246 | 0.185329309 | 4.522710902 | 6.10526E-06 | 0.00016225 | ENSG00000241506 | PSMC1P1 |
| **33.58937298** | 0.837020986 | 0.215451239 | 3.861089828 | 0.000112882 | 0.001822966 | ENSG00000180530 | NRIP1 |
| **451.7138963** | 0.831815454 | 0.095619367 | 8.696162501 | 3.43297E-18 | 1.18188E-15 | ENSG00000130066 | SAT1 |
| **293.8675825** | 0.828927637 | 0.13966999 | 5.920616594 | 3.20737E-09 | 2.15934E-07 | ENSG00000197956 | S100A6 |
| **438.9132666** | 0.82806301 | 0.176258405 | 4.688398568 | 2.75351E-06 | 8.11483E-05 | ENSG00000243709 | LEFTY1 |
| **3376.73191** | 0.822299221 | 0.122364799 | 6.711363849 | 1.92814E-11 | 2.19605E-09 | ENSG00000108821 | COL1A1 |
| **29.30676699** | 0.82031271 | 0.219337111 | 3.675218535 | 0.000237646 | 0.003328787 | ENSG00000214318 | ATP5MC1P6 |
| **109.4183429** | 0.818756156 | 0.143526688 | 5.685876255 | 1.30144E-08 | 7.54111E-07 | ENSG00000169439 | SDC2 |
| **404.1435846** | 0.818707156 | 0.10812068 | 7.569198865 | 3.75533E-14 | 7.29304E-12 | ENSG00000079102 | RUNX1T1 |
| **195.3097428** | 0.815744021 | 0.130833765 | 6.229937194 | 4.66622E-10 | 3.80021E-08 | ENSG00000075420 | FNDC3B |
| **177.1356918** | 0.815161046 | 0.151931065 | 5.339000686 | 9.34603E-08 | 4.36956E-06 | ENSG00000105270 | CLIP3 |
| **327.3812127** | 0.814220447 | 0.117172916 | 6.943227375 | 3.83242E-12 | 4.91979E-10 | ENSG00000165410 | CFL2 |
| **502.6009224** | 0.809363021 | 0.103126788 | 7.847563848 | 4.24196E-15 | 9.05032E-13 | ENSG00000008294 | SPAG9 |
| **927.9201155** | 0.807568669 | 0.088031605 | 9.171265968 | 4.67496E-20 | 2.02332E-17 | ENSG00000162923 | WDR26 |
| **590.1234664** | 0.807554014 | 0.093475786 | 8.636283373 | 5.80714E-18 | 1.95481E-15 | ENSG00000005483 | KMT2E |
| **128.3044863** | 0.807245611 | 0.144412188 | 5.575703027 | 2.46532E-08 | 1.34333E-06 | ENSG00000139793 | MBNL2 |
| **1112.507554** | 0.799787753 | 0.076064043 | 10.51349476 | 7.48667E-26 | 6.67106E-23 | ENSG00000101972 | STAG2 |
| **108.2433277** | 0.797032995 | 0.146523904 | 5.425534013 | 5.77815E-08 | 2.86037E-06 | ENSG00000068366 | ACSL4 |
| **219.9240672** | 0.786352281 | 0.110536882 | 7.103988224 | 1.21207E-12 | 1.74862E-10 | ENSG00000099399 | MAGEB2 |
| **388.3933171** | 0.78262759 | 0.095394326 | 8.200536088 | 2.39317E-16 | 6.47354E-14 | ENSG00000147044 | CASK |
| **211.7496518** | 0.779212723 | 0.139089227 | 5.601656974 | 2.12312E-08 | 1.17806E-06 | ENSG00000115414 | FN1 |
| **45.54127242** | 0.778668404 | 0.212152106 | 3.714947136 | 0.000203246 | 0.002934773 | ENSG00000102908 | NFAT5 |
| **606.2630376** | 0.775670163 | 0.110148802 | 7.035164807 | 1.99026E-12 | 2.71607E-10 | ENSG00000111276 | CDKN1B |
| **1741.687258** | 0.775347103 | 0.088604971 | 8.748706817 | 2.15812E-18 | 7.7836E-16 | ENSG00000168036 | CTNNB1 |
| **2243.659633** | 0.774835826 | 0.104333366 | 7.422139847 | 1.15243E-13 | 1.98375E-11 | ENSG00000008441 | NFIX |
| **291.6073358** | 0.774567904 | 0.108403047 | 7.141275457 | 9.24688E-13 | 1.37325E-10 | ENSG00000067445 | TRO |
| **266.8832327** | 0.774217167 | 0.132805604 | 5.821176597 | 5.84348E-09 | 3.7192E-07 | ENSG00000140836 | ZFHX3 |
| **82.0392087** | 0.771824702 | 0.163938981 | 4.679448745 | 2.87647E-06 | 8.40136E-05 | ENSG00000076356 | PLXNA2 |
| **63.14891334** | 0.766740461 | 0.180352964 | 4.239216968 | 2.24301E-05 | 0.000475204 | ENSG00000159176 | CSRP1 |
| **178.9342974** | 0.761433478 | 0.131603022 | 5.778852049 | 7.5212E-09 | 4.66931E-07 | ENSG00000136451 | VEZF1 |
| **297.3456488** | 0.755340811 | 0.111276163 | 6.781991156 | 1.18531E-11 | 1.41378E-09 | ENSG00000043143 | JADE2 |
| **135.1954228** | 0.752820691 | 0.153056801 | 4.923768831 | 8.48931E-07 | 2.9293E-05 | ENSG00000182095 | TNRC18 |
| **48.21187645** | 0.748765339 | 0.186890256 | 3.993704956 | 6.50488E-05 | 0.001156525 | ENSG00000172638 | EFEMP2 |

| | | | | | | | |
|---|---|---|---|---|---|---|---|
| **134.6086078** | 0.742850438 | 0.158191665 | 4.679306395 | 2.87847E-06 | 8.40136E-05 | ENSG00000231735 | AC092573.1 |
| **380.2328497** | 0.739580332 | 0.097258251 | 7.600283142 | 2.95484E-14 | 5.88945E-12 | ENSG00000145860 | RNF145 |
| **23.42263668** | 0.736230752 | 0.240104126 | 3.222520107 | 0.001270682 | 0.013129808 | ENSG00000124575 | H1-3 |
| **109.7285585** | 0.723600216 | 0.159589676 | 4.521293373 | 6.14629E-06 | 0.000163054 | ENSG00000173706 | HEG1 |
| **378.1640237** | 0.723423733 | 0.099980034 | 7.229788278 | 4.83748E-13 | 7.47736E-11 | ENSG00000177508 | IRX3 |
| **90.97611915** | 0.717314638 | 0.169862384 | 4.216421228 | 2.4821E-05 | 0.000517178 | ENSG00000151474 | FRMD4A |
| **96.21988171** | 0.715078667 | 0.162536888 | 4.379792854 | 1.18792E-05 | 0.000282172 | ENSG00000184232 | OAF |
| **483.4953685** | 0.714344809 | 0.118858714 | 6.003364529 | 1.9327E-09 | 1.34915E-07 | ENSG00000213750 | AP002982.1 |
| **487.9847741** | 0.713247003 | 0.094179522 | 7.571248839 | 3.69653E-14 | 7.27209E-12 | ENSG00000151746 | BICD1 |
| **168.9930692** | 0.711533772 | 0.134364085 | 5.287119678 | 1.24257E-07 | 5.63548E-06 | ENSG00000130024 | PHF10 |
| **18.29061729** | 0.705295353 | 0.229422558 | 2.951725604 | 0.003160036 | 0.026652687 | ENSG00000017427 | IGF1 |
| **898.0313267** | 0.695048896 | 0.099180626 | 7.006401836 | 2.44525E-12 | 3.27793E-10 | ENSG00000175155 | YPEL2 |
| **284.0682131** | 0.691175331 | 0.133518643 | 5.168976684 | 2.35379E-07 | 9.8768E-06 | ENSG00000007350 | TKTL1 |
| **905.6626519** | 0.690587462 | 0.087353939 | 7.905449817 | 2.66968E-15 | 6.03586E-13 | ENSG00000186432 | KPNA4 |
| **967.2027403** | 0.681701049 | 0.097315334 | 7.005026741 | 2.46939E-12 | 3.28125E-10 | ENSG00000070961 | ATP2B1 |
| **20.89785995** | 0.675721075 | 0.224733691 | 2.955713336 | 0.003119469 | 0.026387026 | ENSG00000197757 | HOXC6 |
| **249.6696243** | 0.672104633 | 0.104241137 | 6.443957199 | 1.16398E-10 | 1.05446E-08 | ENSG00000196935 | SRGAP1 |
| **27.88762644** | 0.668539886 | 0.216748877 | 3.018155776 | 0.002543181 | 0.022634613 | ENSG00000159674 | SPON2 |
| **351.2240927** | 0.668306151 | 0.100130321 | 6.671822768 | 2.52646E-11 | 2.73363E-09 | ENSG00000136770 | DNAJC1 |
| **421.8944906** | 0.666581183 | 0.102978987 | 6.470493383 | 9.76834E-11 | 9.134E-09 | ENSG00000151612 | ZNF827 |
| **858.7484287** | 0.659392541 | 0.071018554 | 9.283553032 | 1.63919E-20 | 8.10693E-18 | ENSG00000102054 | RBBP7 |
| **437.5913376** | 0.658726988 | 0.104333863 | 6.31277493 | 2.74076E-10 | 2.39983E-08 | ENSG00000102081 | FMR1 |
| **135.650134** | 0.656234503 | 0.150278841 | 4.362077605 | 1.28833E-05 | 0.00029978 | ENSG00000164692 | COL1A2 |
| **282.3394186** | 0.646096212 | 0.12319471 | 5.242333631 | 1.58558E-07 | 6.96186E-06 | ENSG00000109099 | PMP22 |
| **173.3716144** | 0.645728232 | 0.122690924 | 5.259716553 | 1.44278E-07 | 6.40914E-06 | ENSG00000038382 | TRIO |
| **167.541578** | 0.643580708 | 0.138613048 | 4.642151156 | 3.448E-06 | 9.81774E-05 | ENSG00000104067 | TJP1 |
| **802.7065022** | 0.642501827 | 0.09092482 | 7.06436167 | 1.61356E-12 | 2.30587E-10 | ENSG00000067113 | PLPP1 |
| **256.464548** | 0.64229236 | 0.119953228 | 5.353142183 | 8.64399E-08 | 4.10467E-06 | ENSG00000148143 | ZNF462 |
| **957.13093** | 0.640640127 | 0.08087982 | 7.920318481 | 2.36903E-15 | 5.43728E-13 | ENSG00000005893 | LAMP2 |
| **167.5500201** | 0.635183944 | 0.121201435 | 5.236905853 | 1.63291E-07 | 7.14893E-06 | ENSG00000178409 | BEND3 |
| **507.6852029** | 0.634218366 | 0.10951762 | 5.789672683 | 7.05237E-09 | 4.42603E-07 | ENSG00000133639 | BTG1 |
| **343.2028049** | 0.633444176 | 0.089734923 | 7.057141239 | 1.69963E-12 | 2.40617E-10 | ENSG00000181904 | C5orf24 |
| **444.0771873** | 0.629126064 | 0.126727689 | 4.966104655 | 6.83111E-07 | 2.42905E-05 | ENSG00000082153 | BZW1 |
| **577.762305** | 0.628626601 | 0.089190996 | 7.047083814 | 1.82706E-12 | 2.56262E-10 | ENSG00000071054 | MAP4K4 |
| **605.7111322** | 0.628559639 | 0.104237772 | 6.027671617 | 1.66339E-09 | 1.17743E-07 | ENSG00000157216 | SSBP3 |
| **771.2461409** | 0.619238631 | 0.091831634 | 6.742123451 | 1.56088E-11 | 1.81879E-09 | ENSG00000073921 | PICALM |
| **64.56450614** | 0.615904708 | 0.179859169 | 3.42560735 | 0.000613426 | 0.007322446 | ENSG00000198633 | ZNF534 |
| **25.60627044** | 0.613995241 | 0.237876562 | 2.769110916 | 0.00562095 | 0.04066196 | ENSG00000070047 | PHRF1 |
| **561.6523924** | 0.613741196 | 0.106762054 | 5.748458432 | 9.00608E-09 | 5.45696E-07 | ENSG00000138757 | G3BP2 |
| **215.983043** | 0.612479039 | 0.125220851 | 4.889355886 | 1.01166E-06 | 3.42069E-05 | ENSG00000118260 | CREB1 |
| **58.39451404** | 0.610865008 | 0.178780063 | 3.409774379 | 0.000650166 | 0.007660277 | ENSG00000205730 | ITPRIPL2 |
| **281.0669317** | 0.606251647 | 0.114117147 | 5.311132279 | 1.08946E-07 | 5.01616E-06 | ENSG00000005810 | MYCBP2 |
| **7568.390991** | 0.602576404 | 0.093015827 | 6.478155591 | 9.28506E-11 | 8.79063E-09 | ENSG00000140264 | SERF2 |
| **106.1807333** | 0.602242309 | 0.154735188 | 3.879470761 | 0.000104684 | 0.001719906 | ENSG00000113369 | ARRDC3 |
| **231.3384924** | 0.599846272 | 0.119148841 | 5.032533169 | 4.84041E-07 | 1.82339E-05 | ENSG00000115904 | 1 SOS |

| | | | | | | | |
|---|---|---|---|---|---|---|---|
| **403.4805124** | 0.597788329 | 0.106478699 | 5.611770717 | 2.00267E-08 | 1.11531E-06 | ENSG00000156966 | B3GNT7 |
| **713.6516796** | 0.595093331 | 0.118549598 | 5.016129299 | 5.27229E-07 | 1.97196E-05 | ENSG00000165272 | AQP3 |
| **130.9631707** | 0.595077289 | 0.141901853 | 4.190285066 | 2.78604E-05 | 0.000571856 | ENSG00000015171 | ZMYND11 |
| **174.324139** | 0.588965086 | 0.120882504 | 4.869821304 | 1.11699E-06 | 3.70245E-05 | ENSG00000130827 | PLXNA3 |
| **363.5412955** | 0.587887491 | 0.094714018 | 6.206258406 | 5.42609E-10 | 4.31987E-08 | ENSG00000126351 | THRA |
| **83.01502876** | 0.580944299 | 0.16702486 | 3.476347671 | 0.000508293 | 0.006285405 | ENSG00000129116 | PALLD |
| **1789.886568** | 0.574772568 | 0.095265784 | 6.033415326 | 1.6053E-09 | 1.14165E-07 | ENSG00000280623 | PCAT14 |
| **316.7519956** | 0.574057294 | 0.117129781 | 4.899174435 | 9.62402E-07 | 3.27606E-05 | ENSG00000143641 | GALNT2 |
| **577.6184852** | 0.570532242 | 0.086600297 | 6.586799349 | 4.49409E-11 | 4.53843E-09 | ENSG00000119048 | UBE2B |
| **46.94279186** | 0.570390504 | 0.189375205 | 3.00157292 | 0.002685887 | 0.023558666 | ENSG00000168785 | TSPAN5 |
| **102.2707424** | 0.570221994 | 0.193088267 | 2.937370455 | 0.003310084 | 0.027625982 | ENSG00000199753 | SNORD104 |
| **574.5480861** | 0.563542046 | 0.120476576 | 4.675825427 | 2.92774E-06 | 8.49604E-05 | ENSG00000162433 | AK4 |
| **264.4431056** | 0.559687736 | 0.108683559 | 5.149334531 | 2.61412E-07 | 1.07605E-05 | ENSG00000213626 | LBH |
| **138.3086853** | 0.557183909 | 0.136826867 | 4.068064568 | 4.74052E-05 | 0.000880018 | ENSG00000119778 | ATAD2B |
| **230.2886539** | 0.554710014 | 0.123208935 | 4.501071717 | 6.76117E-06 | 0.000175976 | ENSG00000144749 | LRIG1 |
| **321.8111499** | 0.552246485 | 0.112076677 | 4.924158977 | 8.47239E-07 | 2.9293E-05 | ENSG00000113658 | SMAD5 |
| **454.7397706** | 0.545509379 | 0.141879495 | 3.85080654 | 0.000117729 | 0.001882711 | ENSG00000130303 | BST2 |
| **1738.136097** | 0.543094537 | 0.087030022 | 6.239901946 | 4.37845E-10 | 3.62431E-08 | ENSG00000143933 | CALM2 |
| **414.2573106** | 0.540540349 | 0.105892817 | 5.103526628 | 3.33382E-07 | 1.32896E-05 | ENSG00000136826 | KLF4 |
| **578.2953057** | 0.537413892 | 0.094521468 | 5.685499044 | 1.30431E-08 | 7.54111E-07 | ENSG00000084676 | NCOA1 |
| **201.7803047** | 0.536900456 | 0.146376879 | 3.666559856 | 0.000245835 | 0.003419573 | ENSG00000100647 | SUSD6 |
| **88.1101016** | 0.533599605 | 0.160751565 | 3.31706388 | 0.000909688 | 0.010056543 | ENSG00000162616 | DNAJB4 |
| **1367.400445** | 0.530825854 | 0.070448984 | 7.534014585 | 4.92036E-14 | 9.20169E-12 | ENSG00000102119 | EMD |
| **494.8996673** | 0.530740362 | 0.079522088 | 6.675810362 | 2.4587E-11 | 2.69887E-09 | ENSG00000189337 | KAZN |
| **128.9030096** | 0.52908121 | 0.140215941 | 3.772121251 | 0.000161866 | 0.002442171 | ENSG00000028528 | SNX1 |
| **318.5965865** | 0.527183498 | 0.108318327 | 4.866781538 | 1.1343E-06 | 3.75162E-05 | ENSG00000105245 | NUMBL |
| **243.4414666** | 0.52269546 | 0.117599646 | 4.444568591 | 8.80684E-06 | 0.000222343 | ENSG00000136848 | DAB2IP |
| **237.8525326** | 0.521510055 | 0.107012902 | 4.873947711 | 1.0939E-06 | 3.63386E-05 | ENSG00000182197 | EXT1 |
| **1761.31311** | 0.515915255 | 0.098272935 | 5.250190491 | 1.51942E-07 | 6.71025E-06 | ENSG00000148154 | UGCG |
| **226.6290875** | 0.513648554 | 0.106787758 | 4.811093481 | 1.50107E-06 | 4.77693E-05 | ENSG00000109118 | PHF12 |
| **613.4879207** | 0.512359481 | 0.088711924 | 5.773750642 | 7.75261E-09 | 4.77385E-07 | ENSG00000100888 | CHD8 |
| **824.9561538** | 0.510926633 | 0.090641048 | 5.636729635 | 1.7331E-08 | 9.86954E-07 | ENSG00000101115 | SALL4 |
| **230.0169323** | 0.510129394 | 0.116041113 | 4.393575931 | 1.11501E-05 | 0.000269795 | ENSG00000108679 | LGALS3BP |
| **65.41572534** | 0.508701216 | 0.179405138 | 2.831881388 | 0.004627501 | 0.035278 | ENSG00000174842 | GLMN |
| **112.9600174** | 0.504385459 | 0.154424851 | 3.262126548 | 0.001105798 | 0.011763078 | ENSG00000184205 | TSPYL2 |
| **254.3083967** | 0.502813534 | 0.151892969 | 3.308289607 | 0.000938677 | 0.010303479 | ENSG00000181218 | H2AW |
| **511.3430434** | 0.501115727 | 0.105443341 | 4.752222949 | 2.01192E-06 | 6.19443E-05 | ENSG00000152518 | ZFP36L2 |
| **159.1940408** | 0.500460033 | 0.133730128 | 3.740507263 | 0.000183649 | 0.002711422 | ENSG00000132694 | ARHGEF11 |
| **1326.434041** | 0.494687309 | 0.113665424 | 4.351023833 | 1.35503E-05 | 0.000312421 | ENSG00000125304 | TM9SF2 |
| **75.37546346** | 0.494162616 | 0.173365697 | 2.847500798 | 0.004406398 | 0.03396851 | ENSG00000164442 | CITED2 |
| **690.2290231** | 0.491162559 | 0.079734321 | 6.159764313 | 7.28533E-10 | 5.54564E-08 | ENSG00000186591 | UBE2H |
| **529.6988377** | 0.488714527 | 0.09549131 | 5.117302973 | 3.09936E-07 | 1.24864E-05 | ENSG00000104517 | UBR5 |
| **168.5721763** | 0.488518509 | 0.119725616 | 4.078709242 | 4.52864E-05 | 0.000847556 | ENSG00000162997 | PRORSD1P |
| **760.3896215** | 0.481332758 | 0.074297204 | 6.475285101 | 9.46331E-11 | 8.90374E-09 | ENSG00000102021 | LUZP4 |
| **3787.729455** | 0.477889209 | 0.09013606 | 5.302312321 | 1.14345E-07 | 5.23292E-06 | ENSG00000065978 | YBX1 |

| | | | | | | | |
|---|---|---|---|---|---|---|---|
| **145.7387973** | 0.477779218 | 0.146312847 | 3.261474053 | 0.001108346 | 0.011773533 | ENSG00000116731 | PRDM2 |
| **643.8996527** | 0.4774279 | 0.109859586 | 4.345385222 | 1.39031E-05 | 0.000319098 | ENSG00000124225 | PMEPA1 |
| **212.0067424** | 0.475529198 | 0.1098332 | 4.333542722 | 1.46729E-05 | 0.000335747 | ENSG00000126705 | AHDC1 |
| **147.7798918** | 0.473092193 | 0.1542402 | 3.066881015 | 0.002163049 | 0.019858104 | ENSG00000118007 | STAG1 |
| **64.79470639** | 0.472496815 | 0.170491351 | 2.770416141 | 0.005598471 | 0.040538071 | ENSG00000184887 | BTBD6 |
| **1064.284201** | 0.471419638 | 0.10407787 | 4.529562337 | 5.9106E-06 | 0.000158187 | ENSG00000176485 | PLAAT3 |
| **258.9992553** | 0.470009955 | 0.111359976 | 4.222126216 | 2.42009E-05 | 0.000506346 | ENSG00000156026 | MCU |
| **2508.146621** | 0.467995576 | 0.085080628 | 5.500370968 | 3.78993E-08 | 1.95272E-06 | ENSG00000176788 | BASP1 |
| **190.3310759** | 0.467368762 | 0.122976351 | 3.800917272 | 0.000144161 | 0.002217013 | ENSG00000087095 | NLK |
| **227.0169337** | 0.465871887 | 0.129239819 | 3.604044229 | 0.000313304 | 0.004192513 | ENSG00000196187 | TMEM63A |
| **1367.707097** | 0.465780854 | 0.085386984 | 5.454906182 | 4.89987E-08 | 2.45772E-06 | ENSG00000228462 | RPS19P7 |
| **454.9550529** | 0.465004601 | 0.099064448 | 4.69367536 | 2.6834E-06 | 7.92361E-05 | ENSG00000127483 | HP1BP3 |
| **773.8089372** | 0.464560721 | 0.091324222 | 5.087305712 | 3.63186E-07 | 1.44019E-05 | ENSG00000091986 | CCDC80 |
| **1126.823593** | 0.46402244 | 0.08479097 | 5.472378792 | 4.44034E-08 | 2.23463E-06 | ENSG00000185650 | ZFP36L1 |
| **442.5718265** | 0.460289549 | 0.084673581 | 5.438353849 | 5.37751E-08 | 2.67956E-06 | ENSG00000116497 | S100PBP |
| **578.7959356** | 0.457322058 | 0.108486795 | 4.214417703 | 2.50423E-05 | 0.000521073 | ENSG00000181555 | SETD2 |
| **417.9550276** | 0.456082592 | 0.078218635 | 5.832566578 | 5.45812E-09 | 3.48859E-07 | ENSG00000139718 | SETD1B |
| **394.3289546** | 0.454304769 | 0.104470763 | 4.349450029 | 1.36479E-05 | 0.000314193 | ENSG00000130635 | COL5A1 |
| **1394.574481** | 0.450869651 | 0.091992059 | 4.901084446 | 9.53091E-07 | 3.25167E-05 | ENSG00000120694 | HSPH1 |
| **893.0243036** | 0.449760784 | 0.098992389 | 4.543384172 | 5.53582E-06 | 0.000151093 | ENSG00000112335 | SNX3 |
| **153.3441778** | 0.448263071 | 0.146677715 | 3.056057498 | 0.002242682 | 0.020465153 | ENSG00000078269 | SYNJ2 |
| **212.7644814** | 0.447687765 | 0.127480779 | 3.508040369 | 0.000451421 | 0.005693138 | ENSG00000120733 | KDM3B |
| **492.0588854** | 0.446257469 | 0.101441231 | 4.399503144 | 1.08499E-05 | 0.000265088 | ENSG00000066735 | KIF26A |
| **1452.842539** | 0.44603029 | 0.068828726 | 6.480280523 | 9.15522E-11 | 8.72222E-09 | ENSG00000180879 | SSR4 |
| **429.7741022** | 0.445074291 | 0.094686103 | 4.701320367 | 2.58485E-06 | 7.69258E-05 | ENSG00000213064 | SFT2D2 |
| **7527.281272** | 0.444765389 | 0.06801691 | 6.538964125 | 6.19464E-11 | 6.05396E-09 | ENSG00000110321 | EIF4G2 |
| **161.7486306** | 0.442466541 | 0.13936953 | 3.175266385 | 0.00149699 | 0.014948191 | ENSG00000139163 | ETNK1 |
| **194.664523** | 0.440821493 | 0.134447407 | 3.277744269 | 0.001046401 | 0.011233799 | ENSG00000180357 | ZNF609 |
| **301.611062** | 0.438510712 | 0.10781539 | 4.069779443 | 4.70577E-05 | 0.000874637 | ENSG00000108389 | MTMR4 |
| **4175.108071** | 0.437612226 | 0.098743605 | 4.431001363 | 9.37965E-06 | 0.00023446 | ENSG00000198668 | CALM1 |
| **2158.848291** | 0.437328058 | 0.073843858 | 5.922361371 | 3.17351E-09 | 2.14609E-07 | ENSG00000142168 | SOD1 |
| **280.7363259** | 0.436887824 | 0.143588111 | 3.039626546 | 0.002368717 | 0.02143448 | ENSG00000116212 | LRRC42 |
| **176.1626949** | 0.435654829 | 0.12557293 | 3.472731794 | 0.00051519 | 0.006344794 | ENSG00000134138 | MEIS2 |
| **228.0128521** | 0.435634163 | 0.118989602 | 3.662813711 | 0.00024946 | 0.00345236 | ENSG00000055208 | TAB2 |
| **710.1393256** | 0.435444935 | 0.072333286 | 6.016599256 | 1.78119E-09 | 1.24914E-07 | ENSG00000072364 | AFF4 |
| **294.5036318** | 0.432447152 | 0.097073808 | 4.45678297 | 8.31988E-06 | 0.000211814 | ENSG00000110422 | HIPK3 |
| **582.5893516** | 0.430329006 | 0.080288147 | 5.360806467 | 8.28512E-08 | 3.94664E-06 | ENSG00000183283 | DAZAP2 |
| **414.2557882** | 0.42992006 | 0.125417379 | 3.42633302 | 0.00061179 | 0.007308669 | ENSG00000070915 | SLC12A3 |
| **604.0147938** | 0.428029314 | 0.082457852 | 5.192262751 | 2.07754E-07 | 8.86493E-06 | ENSG00000153317 | ASAP1 |
| **1139.672727** | 0.425719646 | 0.07380354 | 5.768811517 | 7.98325E-09 | 4.87622E-07 | ENSG00000153250 | RBMS1 |
| **1446.958351** | 0.421757266 | 0.12772944 | 3.303847429 | 0.000953677 | 0.010423017 | ENSG00000167004 | PDIA3 |
| **767.3581434** | 0.421216394 | 0.081700731 | 5.155972088 | 2.52318E-07 | 1.05003E-05 | ENSG00000101558 | VAPA |
| **106.6098773** | 0.420576144 | 0.149296927 | 2.814789876 | 0.004880914 | 0.036733882 | ENSG00000117114 | ADGRL2 |
| **206.2231153** | 0.419813394 | 0.12469274 | 3.367452013 | 0.000758662 | 0.008725837 | ENSG00000100330 | MTMR3 |
| **671.2254247** | 0.419681432 | 0.097322085 | 4.313128977 | 1.6096E-05 | 0.000362292 | ENSG00000177606 | JUN |

| | | | | | | | |
|---|---|---|---|---|---|---|---|
| **348.1685983** | 0.419590609 | 0.129649935 | 3.234730098 | 0.001217578 | 0.012711142 | ENSG00000129680 | MAP7D3 |
| **165.3037855** | 0.418983868 | 0.133961326 | 3.119541229 | 0.001811329 | 0.017432029 | ENSG00000135945 | REV1 |
| **3470.467441** | 0.41656249 | 0.064503146 | 6.4581743 | 1.05974E-10 | 9.8484E-09 | ENSG00000141367 | CLTC |
| **368.6444413** | 0.416441893 | 0.111330254 | 3.73509953 | 0.000187641 | 0.002748922 | ENSG00000137942 | FNBP1L |
| **540.7699834** | 0.415770267 | 0.129016435 | 3.223075896 | 0.001268219 | 0.013113303 | ENSG00000100813 | ACIN1 |
| **304.5588048** | 0.414532748 | 0.10032971 | 4.131092905 | 3.61043E-05 | 0.000710269 | ENSG00000140632 | GLYR1 |
| **93.39779379** | 0.413196717 | 0.153690452 | 2.691305454 | 0.0071173 | 0.048368261 | ENSG00000160271 | RALGDS |
| **106.2071268** | 0.412779109 | 0.147919911 | 2.790505734 | 0.005262577 | 0.038810863 | ENSG00000115539 | PDCL3 |
| **710.6520794** | 0.410962134 | 0.099692226 | 4.122058462 | 3.75502E-05 | 0.000729244 | ENSG00000119669 | IRF2BPL |
| **395.8404553** | 0.410722003 | 0.097417096 | 4.219136416 | 2.4524E-05 | 0.000511693 | ENSG00000129245 | FXR2 |
| **1198.908888** | 0.410252939 | 0.104643506 | 3.919937866 | 8.85718E-05 | 0.001503128 | ENSG00000233056 | ERVH48-1 |
| **353.9004556** | 0.406814687 | 0.098592083 | 4.128167984 | 3.65665E-05 | 0.000716571 | ENSG00000171729 | TMEM51 |
| **162.4421974** | 0.404645979 | 0.119825486 | 3.379044451 | 0.000727382 | 0.008438065 | ENSG00000147050 | KDM6A |
| **241.9895166** | 0.403091968 | 0.133263192 | 3.024001983 | 0.002494548 | 0.022293459 | ENSG00000198466 | ZNF587 |
| **3012.713587** | 0.400010046 | 0.07417261 | 5.393038256 | 6.92762E-08 | 3.36345E-06 | ENSG00000099783 | HNRNPM |
| **266.8136593** | 0.399817643 | 0.101385714 | 3.941697521 | 8.0907E-05 | 0.001400661 | ENSG00000163848 | ZNF148 |
| **1365.454105** | 0.398944299 | 0.07405437 | 5.387756662 | 7.13426E-08 | 3.45271E-06 | ENSG00000170027 | YWHAG |
| **140.4043512** | 0.398268554 | 0.137587983 | 2.897788748 | 0.003758036 | 0.030328574 | ENSG00000109255 | NMU |
| **191.4600891** | 0.398048626 | 0.118273477 | 3.367927538 | 0.000757355 | 0.008717639 | ENSG00000185963 | BICD2 |
| **307.0715828** | 0.397577954 | 0.120143725 | 3.309691205 | 0.000933989 | 0.010274563 | ENSG00000166794 | PPIB |
| **418.2651324** | 0.396344188 | 0.099320571 | 3.991161203 | 6.57506E-05 | 0.001166264 | ENSG00000134910 | STT3A |
| **1347.834127** | 0.396255542 | 0.107995368 | 3.669392482 | 0.000243128 | 0.003385015 | ENSG00000090621 | PABPC4 |
| **225.7982574** | 0.393886795 | 0.11051821 | 3.562606342 | 0.000367191 | 0.004794284 | ENSG00000072803 | FBXW11 |
| **339.8991891** | 0.391074325 | 0.116617332 | 3.354850993 | 0.000794077 | 0.009057745 | ENSG00000140262 | TCF12 |
| **499.4237846** | 0.387152504 | 0.080763734 | 4.795510926 | 1.62261E-06 | 5.09943E-05 | ENSG00000138430 | OLA1 |
| **534.9687462** | 0.385100667 | 0.096829392 | 3.979444627 | 6.90764E-05 | 0.001220969 | ENSG00000160712 | IL6R |
| **434.8653941** | 0.382252721 | 0.089382471 | 4.275354108 | 1.90834E-05 | 0.000420779 | ENSG00000108510 | MED13 |
| **225.3072231** | 0.380033425 | 0.113038522 | 3.359852708 | 0.00077984 | 0.008922775 | ENSG00000157212 | PAXIP1 |
| **806.5477324** | 0.378628333 | 0.104510485 | 3.623251107 | 0.000290923 | 0.003927722 | ENSG00000269842 | AC011453.1 |
| **301.3074118** | 0.378406237 | 0.120840054 | 3.129208935 | 0.001752776 | 0.016976378 | ENSG00000078177 | N4BP2 |
| **3166.958368** | 0.377372441 | 0.124657679 | 3.022470162 | 0.002507208 | 0.022380192 | ENSG00000269564 | AC008753.2 |
| **13062.48318** | 0.368781963 | 0.113915078 | 3.238325467 | 0.001202336 | 0.012569344 | ENSG00000198786 | MT-ND5 |
| **1372.343159** | 0.368745237 | 0.073812053 | 4.992905384 | 5.94777E-07 | 2.16579E-05 | ENSG00000163602 | RYBP |
| **215.5376391** | 0.364910901 | 0.121779632 | 2.995489269 | 0.002740049 | 0.02395053 | ENSG00000145715 | RASA1 |
| **495.8020287** | 0.360955262 | 0.093392965 | 3.867704143 | 0.000109865 | 0.001787575 | ENSG00000156639 | ZFAND3 |
| **518.7911037** | 0.360543536 | 0.092965764 | 3.876829209 | 0.000105827 | 0.001733039 | ENSG00000089057 | SLC23A2 |
| **278.4338799** | 0.359951556 | 0.134168993 | 2.678682395 | 0.007391246 | 0.04973905 | ENSG00000162601 | MYSM1 |
| **659.9137957** | 0.355337914 | 0.076948657 | 4.618607721 | 3.86323E-06 | 0.000108489 | ENSG00000143569 | UBAP2L |
| **365.8151997** | 0.353268743 | 0.112439135 | 3.142276126 | 0.001676399 | 0.016372718 | ENSG00000105722 | ERF |
| **658.6764617** | 0.352252095 | 0.091471154 | 3.848567249 | 0.000118811 | 0.001892476 | ENSG00000204120 | GIGYF2 |
| **1057.883671** | 0.351722345 | 0.086963403 | 4.043818734 | 5.25876E-05 | 0.0009586 | ENSG00000072110 | ACTN1 |
| **898.2531587** | 0.351318729 | 0.082460901 | 4.26049093 | 2.03978E-05 | 0.000441409 | ENSG00000078140 | UBE2K |
| **2697.575603** | 0.349715371 | 0.087579354 | 3.992967944 | 6.52514E-05 | 0.001158767 | ENSG00000240563 | L1TD1 |
| **729.2748659** | 0.348072184 | 0.079479935 | 4.378505608 | 1.19496E-05 | 0.000282391 | ENSG00000177565 | TBL1XR1 |
| **166.9396182** | 0.345185648 | 0.124419792 | 2.77620391 | 0.00549977 | 0.040046175 | ENSG00000153944 | MSI2 |

| | | | | | | | |
|---|---|---|---|---|---|---|---|
| **227.9457112** | 0.342181652 | 0.127223792 | 2.68632888 | 0.007224194 | 0.04892012 | ENSG00000125968 | ID1 |
| **184.3111352** | 0.341500849 | 0.12383904 | 2.760369142 | 0.005773608 | 0.04144269 | ENSG00000116406 | EDEM3 |
| **1076.780145** | 0.340305518 | 0.067507403 | 5.041190401 | 4.62645E-07 | 1.77197E-05 | ENSG00000115524 | SF3B1 |
| **369.9286621** | 0.335668942 | 0.090983433 | 3.686604306 | 0.000227266 | 0.003220421 | ENSG00000107863 | ARHGAP21 |
| **471.4282644** | 0.334452087 | 0.113035366 | 2.955677761 | 0.003119829 | 0.026387026 | ENSG00000102753 | KPNA3 |
| **413.631454** | 0.332477367 | 0.089982534 | 3.695404896 | 0.000219537 | 0.003122574 | ENSG00000004534 | RBM6 |
| **220.6597837** | 0.329428077 | 0.111208217 | 2.962700736 | 0.003049529 | 0.025920519 | ENSG00000171604 | CXXC5 |
| **227.3516969** | 0.328316364 | 0.115012842 | 2.853781418 | 0.004320225 | 0.033508838 | ENSG00000106591 | MRPL32 |
| **4296.942703** | 0.327802239 | 0.084138947 | 3.89592243 | 9.78257E-05 | 0.001630213 | ENSG00000198918 | RPL39 |
| **4173.635825** | 0.325971975 | 0.07832094 | 4.161946454 | 3.15546E-05 | 0.000635624 | ENSG00000144381 | HSPD1 |
| **247.580611** | 0.325447694 | 0.109881428 | 2.96463518 | 0.003030421 | 0.025803717 | ENSG00000123353 | ORMDL2 |
| **1622.450922** | 0.325390448 | 0.075117872 | 4.329370953 | 1.49536E-05 | 0.00034114 | ENSG00000083312 | TNPO1 |
| **4259.669253** | 0.324854756 | 0.054983212 | 5.908269725 | 3.4572E-09 | 2.28688E-07 | ENSG00000124766 | SOX4 |
| **213.794905** | 0.3235377 | 0.113390878 | 2.8490102 | 0.004385547 | 0.033859466 | ENSG00000197134 | ZNF257 |
| **894.0611801** | 0.32111822 | 0.097270477 | 3.30237177 | 0.000958709 | 0.010455382 | ENSG00000159692 | CTBP1 |
| **881.6774678** | 0.319283423 | 0.073024831 | 4.3704638 | 1.23983E-05 | 0.000290726 | ENSG00000154001 | PPP2R5E |
| **483.2183531** | 0.318861978 | 0.101354355 | 3.147342895 | 0.001647616 | 0.016156221 | ENSG00000108312 | UBTF |
| **398.9158346** | 0.318386989 | 0.08956171 | 3.553963487 | 0.000379472 | 0.00492988 | ENSG00000134597 | RBMX2 |
| **550.6257212** | 0.318089276 | 0.077252238 | 4.11977656 | 3.7924E-05 | 0.00073462 | ENSG00000160714 | UBE2Q1 |
| **298.0616017** | 0.317379477 | 0.098740527 | 3.214873483 | 0.00130502 | 0.013447923 | ENSG00000170242 | USP47 |
| **1377.461872** | 0.315961737 | 0.081311106 | 3.885434295 | 0.000102147 | 0.001687377 | ENSG00000102317 | RBM3 |
| **783.4724624** | 0.31558051 | 0.085638194 | 3.680205513 | 0.000233046 | 0.003280842 | ENSG00000092148 | HECTD1 |
| **252.1085991** | 0.315270199 | 0.102243491 | 3.082835438 | 0.002050385 | 0.019053993 | ENSG00000138081 | FBXO11 |
| **569.0179965** | 0.311078244 | 0.087273794 | 3.559238795 | 0.000371931 | 0.00484438 | ENSG00000131504 | DIAPH1 |
| **441.3056478** | 0.30788959 | 0.082360261 | 3.739781574 | 0.00018418 | 0.002715446 | ENSG00000140332 | TLE3 |
| **2013.83448** | 0.306106429 | 0.087257612 | 3.508388206 | 0.000450831 | 0.005693138 | ENSG00000173402 | DAG1 |
| **641.6590529** | 0.300461495 | 0.090943162 | 3.301297866 | 0.000962387 | 0.010487936 | ENSG00000112851 | ERBIN |
| **362.0661822** | 0.299091803 | 0.093910251 | 3.18605767 | 0.001442258 | 0.014526151 | ENSG00000094804 | CDC6 |
| **706.0886465** | 0.297664256 | 0.079069635 | 3.764043287 | 0.000167188 | 0.002508968 | ENSG00000062650 | WAPL |
| **753.6774919** | 0.294295055 | 0.076433604 | 3.852743387 | 0.000116802 | 0.00187031 | ENSG00000135090 | TAOK3 |
| **614.0092549** | 0.292966359 | 0.096373621 | 3.040317947 | 0.002363285 | 0.021410911 | ENSG00000184432 | COPB2 |
| **250.851072** | 0.292507327 | 0.105476015 | 2.778640986 | 0.005458682 | 0.039830498 | ENSG00000138646 | HERC5 |
| **1091.925569** | 0.290807706 | 0.084721903 | 3.431734827 | 0.000599734 | 0.007207481 | ENSG00000180370 | PAK2 |
| **711.363328** | 0.290122177 | 0.101046928 | 2.869972264 | 0.004105078 | 0.032336829 | ENSG00000147224 | PRPS1 |
| **334.898824** | 0.288709566 | 0.096905979 | 2.983172922 | 0.002852768 | 0.024696617 | ENSG00000185619 | PCGF3 |
| **495.6320147** | 0.284469375 | 0.098289664 | 2.894615898 | 0.003796227 | 0.030571639 | ENSG00000102580 | DNAJC3 |
| **212.5893371** | 0.283689804 | 0.102820522 | 2.762647524 | 0.005733465 | 0.041289911 | ENSG00000189079 | ARID2 |
| **535.334567** | 0.282340425 | 0.094613071 | 2.985794433 | 0.002828427 | 0.024510877 | ENSG00000168610 | STAT3 |
| **1473.589228** | 0.282039282 | 0.059466373 | 4.742907346 | 2.10673E-06 | 6.44701E-05 | ENSG00000100811 | YY1 |
| **661.4943272** | 0.280183463 | 0.088569891 | 3.156830418 | 0.001594941 | 0.015739519 | ENSG00000127947 | PTPN12 |
| **312.4362004** | 0.279889468 | 0.101954255 | 2.74308737 | 0.006086449 | 0.043053036 | ENSG00000068323 | TFE3 |
| **1194.136689** | 0.277916092 | 0.090069049 | 3.085502573 | 0.002032085 | 0.018946517 | ENSG00000173457 | PPP1R14B |
| **1378.665003** | 0.277612708 | 0.067339052 | 4.122452951 | 3.74859E-05 | 0.00072893 | ENSG00000110917 | MLEC |
| **995.4553639** | 0.276438445 | 0.096481617 | 2.86403838 | 0.004182773 | 0.032693834 | ENSG00000125944 | HNRNPR |
| **884.6230447** | 0.271518923 | 0.071630469 | 3.789810381 | 0.000150762 | 0.002301251 | ENSG00000153827 | TRIP12 |

| | | | | | | | |
|---|---|---|---|---|---|---|---|
| **341.679087** | 0.270724889 | 0.09224605 | 2.936474107 | 0.003319665 | 0.027690689 | ENSG00000166411 | IDH3A |
| **7603.809577** | 0.270506116 | 0.081514284 | 3.318515052 | 0.000904974 | 0.010020872 | ENSG00000166598 | HSP90B1 |
| **851.1642306** | 0.269420931 | 0.070722764 | 3.81113367 | 0.000138331 | 0.002149166 | ENSG00000084733 | RAB10 |
| **414.4100039** | 0.268946013 | 0.086386053 | 3.116481461 | 0.001830232 | 0.017560485 | ENSG00000067992 | PDK3 |
| **726.5787574** | 0.263517378 | 0.08468393 | 3.112068635 | 0.001857813 | 0.017740424 | ENSG00000109606 | DHX15 |
| **345.187563** | 0.261629521 | 0.094092942 | 2.785068879 | 0.005351635 | 0.039295479 | ENSG00000055130 | CUL1 |
| **311.2443036** | 0.261550262 | 0.097689541 | 2.677256547 | 0.007422777 | 0.049864178 | ENSG00000116001 | TIA1 |
| **569.9608651** | 0.261346823 | 0.089943092 | 2.902979337 | 0.00369631 | 0.029956027 | ENSG00000196792 | STRN3 |
| **1104.122553** | 0.257388152 | 0.078184959 | 3.293603207 | 0.00098912 | 0.01073583 | ENSG00000086061 | DNAJA1 |
| **289.0696719** | 0.257251368 | 0.090472479 | 2.850874009 | 0.004359924 | 0.033730404 | ENSG00000102804 | TSC22D1 |
| **2291.606081** | 0.247880435 | 0.077116474 | 3.214399199 | 0.001307178 | 0.01345651 | ENSG00000197451 | HNRNPAB |
| **2326.520733** | 0.241334379 | 0.055802482 | 4.325101954 | 1.52461E-05 | 0.000347291 | ENSG00000177733 | HNRNPA0 |
| **9200.270443** | 0.234179605 | 0.077500496 | 3.021597267 | 0.002514448 | 0.022418402 | ENSG00000044574 | HSPA5 |
| **3667.599004** | 0.231921697 | 0.052798587 | 4.393286885 | 1.1165E-05 | 0.000269795 | ENSG00000143761 | ARF1 |
| **2409.912714** | 0.231861194 | 0.060679211 | 3.822009048 | 0.000132369 | 0.002071408 | ENSG00000100345 | MYH9 |
| **880.9969179** | 0.230164715 | 0.055644738 | 4.141431841 | 3.45144E-05 | 0.000683431 | ENSG00000078699 | CBFA2T2 |
| **1964.7293** | 0.214630647 | 0.073436836 | 2.92203476 | 0.003477527 | 0.028695779 | ENSG00000115306 | SPTBN1 |
| **802.3011404** | 0.208387909 | 0.074113717 | 2.806183523 | 0.005013213 | 0.037390523 | ENSG00000114439 | BBX |
| **751.8512159** | -0.205043474 | 0.075011944 | -2.73926087 | 0.006157749 | 0.043405112 | ENSG00000158246 | TENT5B |
| **436.5598951** | -0.225375306 | 0.083145663 | -2.723128396 | 0.006466692 | 0.045120889 | ENSG00000196305 | IARS1 |
| **3173.747319** | -0.232413011 | 0.076228009 | -3.047724113 | 0.002305815 | 0.020965479 | ENSG00000134419 | RPS15A |
| **2242.457137** | -0.232452959 | 0.085698536 | -2.709971256 | 0.006728904 | 0.046392351 | ENSG00000082781 | ITGB5 |
| **6873.740818** | -0.233988138 | 0.072352097 | -3.233576136 | 0.001222507 | 0.01274504 | ENSG00000114391 | RPL24 |
| **1185.280215** | -0.234274437 | 0.055098699 | -4.248182741 | 2.15512E-05 | 0.000461111 | ENSG00000141552 | ANAPC11 |
| **1121.075302** | -0.235067689 | 0.077332837 | -3.037104795 | 0.002388624 | 0.021588828 | ENSG00000182180 | MRPS16 |
| **415.1255562** | -0.239069615 | 0.08027038 | -2.971223434 | 0.002966159 | 0.025370625 | ENSG00000155906 | RMND1 |
| **626.4730404** | -0.247471278 | 0.083996452 | -2.942668218 | 0.003253969 | 0.027292984 | ENSG00000079462 | PAFAH1B3 |
| **2453.654256** | -0.251987137 | 0.062309425 | -4.041644809 | 5.30776E-05 | 0.000964052 | ENSG00000266472 | MRPS21 |
| **953.1896269** | -0.258004527 | 0.07668476 | -3.359835579 | 0.000779889 | 0.008922775 | ENSG00000004455 | AK2 |
| **1090.809779** | -0.258564741 | 0.094474603 | -2.74247577 | 0.006097795 | 0.04309719 | ENSG00000100097 | LGALS1 |
| **511.5688493** | -0.259289738 | 0.086858349 | -2.983134997 | 0.002853121 | 0.024696617 | ENSG00000117697 | NSL1 |
| **470.98731** | -0.260205012 | 0.091355837 | -2.855589605 | 0.0042957 | 0.033335688 | ENSG00000225663 | MCRIP1 |
| **697.9851514** | -0.260840365 | 0.083753917 | -3.116255932 | 0.001831632 | 0.017560485 | ENSG00000188112 | C6orf132 |
| **2500.074903** | -0.260899911 | 0.068429145 | -3.811425781 | 0.000138168 | 0.002148832 | ENSG00000100138 | SNU13 |
| **312.0317601** | -0.261780883 | 0.097269272 | -2.679193972 | 0.007379963 | 0.049691056 | ENSG00000165688 | PMPCA |
| **12087.59748** | -0.261815222 | 0.093794773 | -2.790978683 | 0.005254893 | 0.038773075 | ENSG00000122406 | RPL5 |
| **1925.997958** | -0.262408263 | 0.058029446 | -4.520217324 | 6.17762E-06 | 0.000163313 | ENSG00000128739 | SNRPN |
| **2267.428184** | -0.264270346 | 0.06761609 | -3.907759552 | 9.31559E-05 | 0.001569662 | ENSG00000118816 | CCNI |
| **30015.49287** | -0.26553587 | 0.079678529 | -3.331979461 | 0.000862306 | 0.00965426 | ENSG00000137154 | RPS6 |
| **482.9867076** | -0.271578155 | 0.078955885 | -3.436538128 | 0.000589199 | 0.007134447 | ENSG00000181924 | COA4 |
| **1109.607242** | -0.272008885 | 0.07174761 | -3.792315116 | 0.000149249 | 0.002283666 | ENSG00000140740 | UQCRC2 |
| **697.7473978** | -0.272214094 | 0.0861074 | -3.160388106 | 0.001575591 | 0.015568855 | ENSG00000187741 | FANCA |
| **456.7203875** | -0.276722228 | 0.091103261 | -3.034794821 | 0.002406994 | 0.02170306 | ENSG00000064545 | TMEM161A |
| **457.4634221** | -0.277359014 | 0.097251282 | -2.84881751 | 0.004388204 | 0.03386272 | ENSG00000179085 | DPM3 |
| **11775.07339** | -0.278289956 | 0.098768059 | -2.817235269 | 0.004843903 | 0.036505194 | ENSG00000130255 | RPL36 |

| | | | | | | | |
|---|---|---|---|---|---|---|---|
| **6085.055202** | -0.282324244 | 0.099183971 | -2.845735746 | 0.004430895 | 0.034087961 | ENSG00000100316 | RPL3 |
| **1097.812283** | -0.283785858 | 0.074909682 | -3.789759259 | 0.000150793 | 0.002301251 | ENSG00000134684 | YARS1 |
| **1355.853231** | -0.289128526 | 0.078215021 | -3.691470669 | 0.000222961 | 0.003168307 | ENSG00000240463 | RPS19P3 |
| **1939.73129** | -0.289291631 | 0.078544226 | -3.680464239 | 0.00023281 | 0.003280842 | ENSG00000220842 | RPL21P16 |
| **536.5679895** | -0.296430253 | 0.080909058 | -3.66490121 | 0.000247434 | 0.003432355 | ENSG00000164904 | ALDH7A1 |
| **491.6595076** | -0.29997407 | 0.080152006 | -3.73964498 | 0.00018428 | 0.002715446 | ENSG00000142687 | KIAA0319L |
| **217.6443296** | -0.301416751 | 0.112662381 | -2.691043539 | 0.00712289 | 0.048384545 | ENSG00000142207 | URB1 |
| **258.0589161** | -0.306207012 | 0.10730477 | -2.84791824 | 0.004400623 | 0.03394126 | ENSG00000197928 | ZNF677 |
| **444.9621602** | -0.307645912 | 0.088784116 | -3.46000087 | 0.000540174 | 0.006630916 | ENSG00000076554 | TPD52 |
| **258.2788054** | -0.316000772 | 0.090025906 | -3.481653219 | 0.000498329 | 0.006181359 | ENSG00000005059 | MCUB |
| **408.3210831** | -0.316799859 | 0.102913133 | -3.083171957 | 0.002048068 | 0.019044893 | ENSG00000119408 | NEK6 |
| **921.1341355** | -0.318010475 | 0.094534771 | -3.360736824 | 0.000777349 | 0.008907168 | ENSG00000143153 | ATP1B1 |
| **283.5982655** | -0.31832126 | 0.108292899 | -2.953089538 | 0.003146107 | 0.026558766 | ENSG00000183207 | RUVBL2 |
| **1796.868123** | -0.32217167 | 0.069065327 | -4.6607326 | 3.15086E-06 | 9.00551E-05 | ENSG00000159352 | PSMD4 |
| **228.4573925** | -0.328269617 | 0.118122815 | -2.775495875 | 0.00551176 | 0.040101892 | ENSG00000115286 | NDUFS7 |
| **1071.085626** | -0.328916341 | 0.064764531 | -5.074616681 | 3.88278E-07 | 1.52374E-05 | ENSG00000143575 | HAX1 |
| **3607.86119** | -0.329752251 | 0.074642386 | -4.417399386 | 9.98955E-06 | 0.000246052 | ENSG00000105202 | FBL |
| **337.4585994** | -0.331254275 | 0.09876893 | -3.352126948 | 0.000801932 | 0.009126725 | ENSG00000188677 | PARVB |
| **538.55527** | -0.331703821 | 0.120750201 | -2.744819407 | 0.00605442 | 0.042896334 | ENSG00000232445 | EMSLR |
| **2845.199008** | -0.333914202 | 0.112400089 | -2.9685905 | 0.00299169 | 0.025545727 | ENSG00000244245 | AC133134.1 |
| **2914.327338** | -0.334191002 | 0.080312783 | -4.159170836 | 3.19405E-05 | 0.000642503 | ENSG00000232573 | RPL3P4 |
| **432.459649** | -0.338189623 | 0.099141211 | -3.400970664 | 0.00067147 | 0.007890948 | ENSG00000143515 | ATP8B2 |
| **1114.047943** | -0.338465297 | 0.071440001 | -4.737981617 | 2.15857E-06 | 6.57909E-05 | ENSG00000105373 | NOP53 |
| **9284.715647** | -0.339643828 | 0.089840633 | -3.779991844 | 0.000156833 | 0.002379576 | ENSG00000163682 | RPL9 |
| **893.8044571** | -0.341994852 | 0.077940073 | -4.389442491 | 1.13642E-05 | 0.000273245 | ENSG00000077312 | SNRPA |
| **371.5664975** | -0.344294294 | 0.092920152 | -3.688133683 | 0.000225905 | 0.003204128 | ENSG00000143194 | MAEL |
| **1745.003834** | -0.348291384 | 0.088052598 | -3.954821245 | 7.65919E-05 | 0.001338194 | ENSG00000172053 | QARS1 |
| **709.7890675** | -0.351627087 | 0.081550097 | -4.315800038 | 1.59026E-05 | 0.000359005 | ENSG00000137876 | RSL24D1 |
| **184.3620483** | -0.353742248 | 0.126733959 | -2.805271699 | 0.005027418 | 0.037478017 | ENSG00000106603 | COA1 |
| **522.8362377** | -0.354114551 | 0.100944294 | -3.505689094 | 0.000455427 | 0.005729199 | ENSG00000175634 | RPS6KB2 |
| **337.3765095** | -0.355369696 | 0.096123977 | -3.695709514 | 0.000219274 | 0.003121763 | ENSG00000168256 | NKIRAS2 |
| **241.9214385** | -0.361321933 | 0.123894026 | -2.912228301 | 0.003588602 | 0.029415665 | ENSG00000168569 | TMEM223 |
| **172.1577221** | -0.363301175 | 0.131306911 | -2.744254559 | 0.006064849 | 0.042950131 | ENSG00000166452 | AKIP1 |
| **1369.824064** | -0.363795894 | 0.083615083 | -4.352428634 | 1.34638E-05 | 0.000311373 | ENSG00000185201 | IFITM2 |
| **155.136643** | -0.373960733 | 0.123872674 | -2.997829109 | 0.002719101 | 0.023781144 | ENSG00000279088 | AC022400.8 |
| **10199.92764** | -0.376459815 | 0.073570643 | -5.116078368 | 3.11953E-07 | 1.25344E-05 | ENSG00000125144 | MT1G |
| **2554.494926** | -0.383860175 | 0.062701368 | -6.122804767 | 9.19423E-10 | 6.82717E-08 | ENSG00000167460 | TPM4 |
| **169.9737013** | -0.384134765 | 0.131529641 | -2.921653313 | 0.003481789 | 0.028711015 | ENSG00000029639 | TFB1M |
| **203.7170836** | -0.384509731 | 0.133632246 | -2.903560223 | 0.00368946 | 0.029918598 | ENSG00000115255 | REEP6 |
| **768.849616** | -0.385202432 | 0.108925053 | -3.53409136 | 0.00040918 | 0.005257212 | ENSG00000215845 | TSTD1 |
| **148.0503642** | -0.386740173 | 0.133854492 | -2.875506452 | 0.004033799 | 0.031891434 | ENSG00000161888 | SPC24 |
| **483.1152631** | -0.387787931 | 0.11179502 | -3.500611855 | 0.000464191 | 0.005817579 | ENSG00000133710 | SPINK5 |
| **282.1326433** | -0.389032397 | 0.109518926 | -3.555602007 | 0.000377115 | 0.004903461 | ENSG00000141741 | MIEN1 |
| **1204.416119** | -0.391092608 | 0.068142017 | -5.735166205 | 9.74168E-09 | 5.78694E-07 | ENSG00000140983 | RHOT2 |
| **5290.473835** | -0.391145253 | 0.090608954 | -4.316598713 | 1.58452E-05 | 0.000358243 | ENSG00000197958 | RPL12 |

| | | | | | | | |
|---|---|---|---|---|---|---|---|
| **346.2168612** | -0.395820001 | 0.113396595 | -3.480995755 | 0.000499553 | 0.006187437 | ENSG00000219222 | RPL12P47 |
| **384.1605686** | -0.398280865 | 0.109752258 | -3.628755294 | 0.000284791 | 0.003862144 | ENSG00000134809 | TIMM10 |
| **899.7702585** | -0.399691313 | 0.103038588 | -3.869488622 | 0.000109064 | 0.001778363 | ENSG00000229659 | RPL26P6 |
| **193.8812604** | -0.403329397 | 0.138051601 | -2.915164673 | 0.003555009 | 0.029203513 | ENSG00000106384 | MOGAT3 |
| **243.7311473** | -0.40806148 | 0.10828697 | -3.766362082 | 0.000165644 | 0.002494203 | ENSG00000100814 | CCNB1IP1 |
| **392.4663447** | -0.409279565 | 0.111443013 | -3.665602461 | 0.000246757 | 0.003429243 | ENSG00000105612 | DNASE2 |
| **512.0151701** | -0.40964617 | 0.10828338 | -3.773524566 | 0.000160957 | 0.002435746 | ENSG00000223803 | RPS20P14 |
| **107.7218377** | -0.413082463 | 0.148422551 | -2.789922927 | 0.005272059 | 0.038842972 | ENSG00000276529 | AP001505.1 |
| **1656.027445** | -0.415870921 | 0.089401559 | -4.654908889 | 3.24124E-06 | 9.24638E-05 | ENSG00000103257 | SLC7A5 |
| **274.1420682** | -0.417563727 | 0.107034604 | -3.907906322 | 9.30994E-05 | 0.001569662 | ENSG00000187840 | EIF4EBP1 |
| **141.5458978** | -0.419107959 | 0.121460065 | -3.444253006 | 0.00057264 | 0.006972946 | ENSG00000073536 | NLE1 |
| **597.7209282** | -0.425887332 | 0.107022418 | -3.957307135 | 7.57994E-05 | 0.00132664 | ENSG00000162391 | FAM151A |
| **124.9842987** | -0.429403987 | 0.154310859 | -2.776104145 | 0.005501458 | 0.040046175 | ENSG00000103034 | NDRG4 |
| **107.0488138** | -0.430234648 | 0.159674874 | -2.751380294 | 0.00593447 | 0.042263921 | ENSG00000169442 | CD52 |
| **1141.324841** | -0.431760823 | 0.088184688 | -4.891778312 | 9.9929E-07 | 3.38641E-05 | ENSG00000158864 | NDUFS2 |
| **374.9118495** | -0.432099256 | 0.12096797 | -3.61649257 | 0.000298622 | 0.004010217 | ENSG00000169136 | ATF5 |
| **432.954285** | -0.438688286 | 0.114594806 | -3.818238641 | 0.000134408 | 0.00209898 | ENSG00000117411 | B4GALT2 |
| **402.5004896** | -0.440751172 | 0.114710166 | -3.846165816 | 0.000119981 | 0.0019071 | ENSG00000222041 | CYTOR |
| **545.9095632** | -0.442388729 | 0.088586294 | -4.995410497 | 5.87107E-07 | 2.14301E-05 | ENSG00000126749 | EMG1 |
| **287.680571** | -0.450055997 | 0.107995642 | -4.15152354 | 3.30269E-05 | 0.000657689 | ENSG00000162493 | PDPN |
| **408.106098** | -0.452361601 | 0.09700513 | -4.685352899 | 2.79478E-06 | 8.22045E-05 | ENSG00000198093 | ZNF649 |
| **253.9476697** | -0.453545974 | 0.119981562 | -3.774889082 | 0.000160079 | 0.002424875 | ENSG00000164284 | GRPEL2 |
| **332.7442574** | -0.458163324 | 0.106347246 | -4.302116175 | 1.69175E-05 | 0.000377973 | ENSG00000084092 | NOA1 |
| **115.667217** | -0.458918289 | 0.152273192 | -3.087701888 | 0.002017107 | 0.018861196 | ENSG00000136010 | ALDH1L2 |
| **2738.277635** | -0.461048883 | 0.117507912 | -3.920393057 | 8.84047E-05 | 0.001502978 | ENSG00000234741 | GAS5 |
| **253.5968325** | -0.461321323 | 0.121390912 | -3.788889658 | 0.000151322 | 0.002306064 | ENSG00000143373 | ZNF687 |
| **737.6224979** | -0.462417748 | 0.137228514 | -3.353947142 | 0.000796676 | 0.009080546 | ENSG00000197879 | MYO1C |
| **190.1041638** | -0.470854858 | 0.12583938 | -3.727933799 | 0.000193056 | 0.00281586 | ENSG00000283633 | AP000547.3 |
| **1060.740281** | -0.472057475 | 0.088215513 | -5.350755628 | 8.75877E-08 | 4.13327E-06 | ENSG00000092621 | PHGDH |
| **305.4841388** | -0.47476646 | 0.136302255 | -3.435082473 | 0.000592374 | 0.007160693 | ENSG00000197721 | CR1L |
| **423.795961** | -0.475378583 | 0.087214126 | -5.441960836 | 5.26973E-08 | 2.63452E-06 | ENSG00000120699 | EXOSC8 |
| **1302.822233** | -0.481263241 | 0.079360984 | -6.065110048 | 1.31863E-09 | 9.51174E-08 | ENSG00000182199 | SHMT2 |
| **229.4710442** | -0.481276244 | 0.107961643 | -4.440334175 | 8.98193E-06 | 0.000225262 | ENSG00000186665 | C17orf58 |
| **1110.445723** | -0.483238707 | 0.098995348 | -4.881038115 | 1.05529E-06 | 3.54446E-05 | ENSG00000112667 | DNPH1 |
| **432.0407195** | -0.483486623 | 0.109536948 | -4.418335859 | 9.94638E-06 | 0.000245387 | ENSG00000143367 | TUFT1 |
| **617.1192493** | -0.483664659 | 0.098924189 | -4.879226544 | 1.06503E-06 | 3.56925E-05 | ENSG00000196365 | LONP1 |
| **682.9828189** | -0.484031985 | 0.101097586 | -4.787560365 | 1.68821E-06 | 5.28368E-05 | ENSG00000224032 | EPB41L4A-AS |
| **493.3726661** | -0.484863432 | 0.124079738 | -3.904748813 | 9.43233E-05 | 0.001584046 | ENSG00000167397 | VKORC1 |
| **241.5927578** | -0.484863483 | 0.116515459 | -4.154489327 | 3.26015E-05 | 0.000653237 | ENSG00000118162 | KPTN |
| **1908.01905** | -0.485965662 | 0.08187681 | -5.932476497 | 2.98399E-09 | 2.02697E-07 | ENSG00000090861 | AARS1 |
| **240.9584964** | -0.489814473 | 0.14405729 | -3.385519484 | 0.000710437 | 0.00826551 | ENSG00000146733 | PSPH |
| **376.9365174** | -0.49468135 | 0.135663623 | -3.662865088 | 0.00024941 | 0.00345236 | ENSG00000244187 | TMEM141 |
| **123.8237995** | -0.494882635 | 0.148000895 | -3.416726432 | 0.000633789 | 0.007518121 | ENSG00000074181 | NOTCH3 |
| **157.1804434** | -0.496541124 | 0.120551149 | -4.098681058 | 4.15511E-05 | 0.000794718 | ENSG00000139725 | RHOF |
| **76.53577314** | -0.499935597 | 0.167525341 | -2.941962372 | 0.003261396 | 0.027325011 | ENSG00000270916 | RPL12P50 |

| | | | | | | | |
|---|---|---|---|---|---|---|---|
| **2150.790575** | -0.500815066 | 0.064447632 | -7.769430306 | 7.88399E-15 | 1.63598E-12 | ENSG00000135390 | ATP5MC2 |
| **81.849748** | -0.502414403 | 0.174619322 | -2.880506722 | 0.003970365 | 0.031541419 | ENSG00000105472 | CLEC11A |
| **145.073542** | -0.503686277 | 0.142062974 | -3.565414899 | 0.000363281 | 0.004748042 | ENSG00000164118 | CEP44 |
| **382.40924** | -0.505081546 | 0.100251259 | -5.040948591 | 4.6323E-07 | 1.77197E-05 | ENSG00000125520 | SLC2A4RG |
| **995.4738525** | -0.506763939 | 0.100584712 | -5.032114921 | 4.85098E-07 | 1.82339E-05 | ENSG00000008838 | MED24 |
| **86.44918029** | -0.507604664 | 0.157640564 | -3.200169825 | 0.001373466 | 0.013953902 | ENSG00000129219 | PLD2 |
| **437.768864** | -0.509284946 | 0.095090846 | -5.348424645 | 8.87231E-08 | 4.17384E-06 | ENSG00000141582 | CBX4 |
| **404.3428663** | -0.514689158 | 0.116587198 | -4.426723183 | 9.56754E-06 | 0.00023837 | ENSG00000105370 | LIM2 |
| **758.0227092** | -0.515756591 | 0.087181832 | -5.914865618 | 3.32147E-09 | 2.2132E-07 | ENSG00000101444 | AHCY |
| **309.2754196** | -0.5168576 | 0.128424925 | -4.025703298 | 5.68053E-05 | 0.001024389 | ENSG00000154328 | NEIL2 |
| **1620.738487** | -0.5182725 | 0.104334758 | -4.965099999 | 6.86657E-07 | 2.43594E-05 | ENSG00000166986 | MARS1 |
| **1532.786351** | -0.528149206 | 0.120633075 | -4.376641498 | 1.20522E-05 | 0.000284372 | ENSG00000105281 | SLC1A5 |
| **488.1678627** | -0.528401037 | 0.115658391 | -4.566369552 | 4.96244E-06 | 0.000136923 | ENSG00000157992 | KRTCAP3 |
| **318.3780195** | -0.537745183 | 0.101891423 | -5.264454095 | 1.40607E-07 | 6.28291E-06 | ENSG00000178802 | MPI |
| **445.4706263** | -0.538275101 | 0.089204546 | -6.019414365 | 1.75049E-09 | 1.23332E-07 | ENSG00000101310 | SEC23B |
| **229.3577374** | -0.539199622 | 0.108708315 | -4.955829688 | 7.20223E-07 | 2.54311E-05 | ENSG00000146426 | TIAM2 |
| **307.2729042** | -0.539841701 | 0.13895482 | -3.896468011 | 9.76057E-05 | 0.001628338 | ENSG00000111319 | SCNN1A |
| **636.0681622** | -0.540494476 | 0.082555925 | -6.539245344 | 6.183E-11 | 6.05396E-09 | ENSG00000125877 | ITPA |
| **293.011922** | -0.541786145 | 0.12658821 | -4.266541338 | 1.98527E-05 | 0.000432081 | ENSG00000225783 | MIAT |
| **83.26380579** | -0.542363481 | 0.179840373 | -2.99865162 | 0.002711772 | 0.023744463 | ENSG00000283047 | FRG1FP |
| **3617.702946** | -0.546140792 | 0.091212809 | -5.986063928 | 2.1498E-09 | 1.49381E-07 | ENSG00000128272 | ATF4 |
| **73.04794438** | -0.547418724 | 0.154044571 | -3.434894238 | 0.000592785 | 0.007160693 | ENSG00000130649 | CYP2E1 |
| **109.3722411** | -0.556490226 | 0.141601674 | -3.924963206 | 8.6743E-05 | 0.001486406 | ENSG00000157927 | RADIL |
| **67.11137967** | -0.557289137 | 0.173413703 | -3.204164827 | 0.001354549 | 0.013789455 | ENSG00000078814 | MYH7B |
| **5326.438371** | -0.562002745 | 0.069048625 | -8.138448445 | 4.00375E-16 | 1.01081E-13 | ENSG00000117632 | STMN1 |
| **1307.739565** | -0.566934327 | 0.084563987 | -6.699455334 | 2.09198E-11 | 2.36487E-09 | ENSG00000042493 | CAPG |
| **111.869855** | -0.56808025 | 0.153123003 | -3.747387855 | 0.000178686 | 0.002651058 | ENSG00000154134 | ROBO3 |
| **197.8239847** | -0.569444618 | 0.146106925 | -4.081349544 | 4.4775E-05 | 0.00084046 | ENSG00000139174 | PRICKLE1 |
| **253.1101464** | -0.57992524 | 0.123030828 | -4.707469433 | 2.50811E-06 | 7.51011E-05 | ENSG00000052344 | PRSS8 |
| **497.522631** | -0.581036472 | 0.101108688 | -5.737565544 | 9.60471E-09 | 5.75068E-07 | ENSG00000132386 | SERPINF1 |
| **40.61324107** | -0.587504667 | 0.201259269 | -2.911518176 | 0.00359677 | 0.029419844 | ENSG00000169964 | TMEM42 |
| **61.54546014** | -0.589796264 | 0.173522043 | -3.281141834 | 0.001033877 | 0.011132293 | ENSG00000107485 | GATA3 |
| **82.01981826** | -0.592139265 | 0.242519592 | -3.011380786 | 0.002600625 | 0.02301067 | ENSG00000196350 | ZNF729 |
| **221.7201872** | -0.598208788 | 0.108707033 | -5.484493335 | 4.14656E-08 | 2.10779E-06 | ENSG00000101470 | TNNC2 |
| **296.0639816** | -0.599702632 | 0.101897863 | -5.864527866 | 4.50412E-09 | 2.95361E-07 | ENSG00000111269 | CREBL2 |
| **207.388145** | -0.614994109 | 0.132796101 | -4.625253209 | 3.74141E-06 | 0.000105934 | ENSG00000230148 | HOXB-AS1 |
| **114.9253737** | -0.620958124 | 0.164922866 | -3.759685629 | 0.000170127 | 0.002546526 | ENSG00000100300 | TSPO |
| **290.6492534** | -0.621326367 | 0.134456978 | -4.620162572 | 3.83439E-06 | 0.000108163 | ENSG00000183077 | AFMID |
| **32.41515952** | -0.625889836 | 0.21448652 | -2.781642597 | 0.005408456 | 0.039559294 | ENSG00000185792 | NLRP9 |
| **115.9319555** | -0.634200742 | 0.157429277 | -4.114721746 | 3.87647E-05 | 0.000747083 | ENSG00000130707 | ASS1 |
| **1948.791436** | -0.641344022 | 0.067865275 | -9.448490131 | 3.43753E-21 | 2.00276E-18 | ENSG00000108828 | VAT1 |
| **342.226067** | -0.642261667 | 0.143308945 | -4.473930143 | 7.67948E-06 | 0.00019719 | ENSG00000197355 | UAP1L1 |
| **37.56037427** | -0.645267115 | 0.207483759 | -3.085056932 | 0.002035132 | 0.018951162 | ENSG00000171368 | TPPP |
| **40.48374892** | -0.645735167 | 0.200142882 | -3.175976658 | 0.00149333 | 0.014921476 | ENSG00000188505 | NCCRP1 |
| **123.258979** | -0.648382924 | 0.159332536 | -4.054995679 | 5.01352E-05 | 0.000921661 | ENSG00000165752 | STK32C |

| | | | | | | | |
|---|---|---|---|---|---|---|---|
| **3251.025479** | -0.651262648 | 0.066613946 | -9.774467312 | 1.44918E-22 | 9.97824E-20 | ENSG00000168159 | RNF187 |
| **324.6759025** | -0.656846189 | 0.114435675 | -5.745941762 | 9.14108E-09 | 5.51669E-07 | ENSG00000100889 | PCK2 |
| **172.1212198** | -0.66745792 | 0.164916849 | -4.077445626 | 4.55332E-05 | 0.000850477 | ENSG00000170525 | PFKFB3 |
| **745.1629934** | -0.678277329 | 0.090218612 | -7.519922839 | 5.48086E-14 | 1.00029E-11 | ENSG00000151012 | SLC7A11 |
| **281.5626158** | -0.694147385 | 0.107454965 | -6.44642145 | 1.14522E-10 | 1.05138E-08 | ENSG00000153044 | CENPH |
| **301.4946307** | -0.695973093 | 0.111409427 | -6.232081804 | 4.60277E-10 | 3.7688E-08 | ENSG00000167700 | MFSD3 |
| **63.72669464** | -0.698897967 | 0.195168751 | -3.706573419 | 0.000210082 | 0.003013568 | ENSG00000183421 | RIPK4 |
| **75.01587066** | -0.709303995 | 0.179835489 | -3.92106225 | 8.81595E-05 | 0.001500494 | ENSG00000188493 | C19orf54 |
| **1220.609946** | -0.709940189 | 0.087167349 | -8.150551636 | 3.62268E-16 | 9.30108E-14 | ENSG00000135069 | PSAT1 |
| **144.3317521** | -0.712607612 | 0.123099786 | -5.735890599 | 9.70013E-09 | 5.78494E-07 | ENSG00000175573 | C11orf68 |
| **676.5434062** | -0.72685682 | 0.122765909 | -5.916062345 | 3.2974E-09 | 2.21014E-07 | ENSG00000128965 | CHAC1 |
| **349.1127948** | -0.729683392 | 0.109182761 | -6.666715923 | 2.61591E-11 | 2.79055E-09 | ENSG00000249242 | TMEM150C |
| **58.00508689** | -0.734852684 | 0.188153387 | -3.864567479 | 0.000111286 | 0.001806821 | ENSG00000130270 | ATP8B3 |
| **956.0179529** | -0.739434161 | 0.120767875 | -6.129942644 | 8.79108E-10 | 6.59244E-08 | ENSG00000115457 | IGFBP2 |
| **191.3786336** | -0.741102605 | 0.131432054 | -5.623202927 | 1.87449E-08 | 1.05557E-06 | ENSG00000272325 | NUDT3 |
| **140.1203151** | -0.742144384 | 0.146710198 | -5.091184958 | 3.55833E-07 | 1.41474E-05 | ENSG00000128165 | ADM2 |
| **369.5016608** | -0.750416513 | 0.123599875 | -6.058014654 | 1.37812E-09 | 9.89371E-08 | ENSG00000176046 | NUPR1 |
| **35.95250742** | -0.753799568 | 0.208034026 | -3.508037786 | 0.000451425 | 0.005693138 | ENSG00000011600 | TYROBP |
| **68.03450448** | -0.764574566 | 0.168213483 | -4.493988713 | 6.99013E-06 | 0.000181003 | ENSG00000108641 | B9D1 |
| **171.5232204** | -0.767632052 | 0.135595245 | -5.644325835 | 1.6583E-08 | 9.4792E-07 | ENSG00000141934 | PLPP2 |
| **268.0318263** | -0.781498579 | 0.102918464 | -7.565965941 | 3.84993E-14 | 7.38212E-12 | ENSG00000089050 | RBBP9 |
| **63.58257054** | -0.800780934 | 0.180061488 | -4.430447981 | 9.40375E-06 | 0.000234676 | ENSG00000027869 | SH2D2A |
| **244.0788983** | -0.802115068 | 0.121495695 | -6.617583373 | 3.65118E-11 | 3.76246E-09 | ENSG00000180353 | HCLS1 |
| **1076.948496** | -0.804907057 | 0.069209003 | -11.6250586 | 3.07382E-31 | 3.58171E-28 | ENSG00000183010 | PYCR1 |
| **212.6874674** | -0.805956691 | 0.166087176 | -4.874757928 | 1.08942E-06 | 3.63364E-05 | ENSG00000109107 | ALDOC |
| **64.45304812** | -0.811779505 | 0.187982351 | -4.38602858 | 1.15439E-05 | 0.000276688 | ENSG00000272512 | AL645608.7 |
| **323.6159618** | -0.867063592 | 0.101708581 | -8.504018953 | 1.83138E-17 | 5.90249E-15 | ENSG00000104814 | MAP4K1 |
| **239.1324489** | -0.902538056 | 0.116397197 | -7.740655032 | 9.89059E-15 | 2.02463E-12 | ENSG00000106948 | AKNA |
| **86.17369686** | -0.912262796 | 0.163186982 | -5.490925846 | 3.99832E-08 | 2.04617E-06 | ENSG00000066923 | STAG3 |
| **36.98199801** | -0.929593479 | 0.211246922 | -4.458010689 | 8.27238E-06 | 0.000211056 | ENSG00000155090 | KLF10 |
| **130.4523541** | -1.038725819 | 0.152765997 | -6.752357474 | 1.45462E-11 | 1.70811E-09 | ENSG00000111981 | ULBP1 |
| **225.1428502** | -1.166267481 | 0.127218082 | -9.127230478 | 7.0274E-20 | 2.87706E-17 | ENSG00000170627 | GTSF1 |
| **23.33318236** | -1.169161478 | 0.219560937 | -5.032551398 | 4.83995E-07 | 1.82339E-05 | ENSG00000172794 | RAB37 |
| **84.30317465** | -1.435515866 | 0.169843991 | -8.185049272 | 2.72192E-16 | 7.10891E-14 | ENSG00000171872 | KLF17 |
| **54.49369857** | -1.650275261 | 0.181573269 | -8.56892628 | 1.04454E-17 | 3.43973E-15 | ENSG00000112273 | HDGFL1 |
| **113.9826011** | -1.811300077 | 0.179903327 | -9.608866301 | 7.33521E-22 | 4.44455E-19 | ENSG00000248370 | LINC02434 |
| **96.24398152** | -1.919743471 | 0.173188196 | -10.48928264 | 9.67602E-26 | 8.14291E-23 | ENSG00000003436 | TFPI |
| **566.3147239** | -3.576114256 | 0.167060577 | -20.44816751 | 6.23747E-93 | 9.44852E-89 | ENSG00000229807 | XIST |

Supplemental Table S1_Sheet_3

| baseMean | log2FoldChange | lfcSE | stat | pvalue | padj | gene_id | gene_name |
|---|---|---|---|---|---|---|---|
| **87.57482141** | -0.558269046 | 0.160174923 | -3.480318705 | 0.000500818 | 0.021240448 | ENSG00000136732 | GYPC |
| **55.67753162** | -0.630138253 | 0.19402518 | -3.277276797 | 0.001048136 | 0.032585032 | ENSG00000242950 | ERVW-1 |
| **714.0403546** | -0.257781099 | 0.066807444 | -3.857841737 | 0.000114393 | 0.008524776 | ENSG00000104872 | PIH1D1 |

Supplemental Table S1_Sheet_4

| baseMean | log2FoldChange | lfcSE | stat | pvalue | padj | gene_id | gene_name |
|---|---|---|---|---|---|---|---|
| **305.4841388** | 3.55482749 | 0.168563288 | 18.42640244 | 8.07E-76 | 7.33E-72 | ENSG00000197721 | CR1L |
| **67.07689273** | 1.35927396 | 0.222995713 | 6.123896045 | 9.13E-10 | 7.55E-07 | ENSG00000264229 | RNU4ATAC |
| **204.0199145** | 0.675866985 | 0.132392326 | 5.082088566 | 3.73E-07 | 0.000109475 | ENSG00000137145 | DENND4C |
| **417.3022627** | 0.675223841 | 0.137336753 | 4.899167609 | 9.62E-07 | 0.000249986 | ENSG00000148344 | PTGES |
| **169.5084524** | 0.64358689 | 0.131406423 | 4.806683355 | 1.53E-06 | 0.00036712 | ENSG00000120498 | TEX11 |
| **58.98414308** | 0.60029296 | 0.197228322 | 3.108710402 | 0.001879058 | 0.045332571 | ENSG00000077264 | PAK3 |
| **284.0682131** | 0.482015075 | 0.130226699 | 3.674500771 | 0.000238315 | 0.013606576 | ENSG00000007350 | TKTL1 |
| **143.4912995** | 0.477390937 | 0.154674969 | 3.128659884 | 0.001756054 | 0.043618281 | ENSG00000103534 | TMC5 |
| **115.9319555** | 0.468001377 | 0.141391855 | 3.312153411 | 0.000925808 | 0.030605517 | ENSG00000130707 | ASS1 |
| **417.5539558** | 0.449779982 | 0.096168547 | 4.665365286 | 3.08E-06 | 0.000614212 | ENSG00000134352 | IL6ST |
| **1593.660216** | 0.409963844 | 0.088199151 | 4.649994949 | 3.32E-06 | 0.000622729 | ENSG00000127603 | MACF1 |
| **333.7051932** | 0.405188312 | 0.12172049 | 3.327729087 | 0.00087557 | 0.029156791 | ENSG00000119899 | SLC17A5 |
| **319.6352955** | 0.398353126 | 0.112088238 | 3.539481538 | 0.000400914 | 0.018132873 | ENSG00000153982 | GDPD1 |
| **547.040314** | 0.384947289 | 0.084987693 | 4.535082775 | 5.76E-06 | 0.00093314 | ENSG00000197312 | DDI2 |
| **614.4468623** | 0.292650113 | 0.092836008 | 3.154726143 | 0.001606488 | 0.041490297 | ENSG00000157106 | SMG1 |
| **487.9847741** | 0.289362988 | 0.088754119 | 3.250520773 | 0.001151939 | 0.034594869 | ENSG00000151746 | BICD1 |
| **434.8653941** | 0.266474147 | 0.08689009 | 3.070315429 | 0.002138328 | 0.047998865 | ENSG00000108510 | MED13 |
| **880.9969179** | 0.254593545 | 0.054806017 | 4.640540318 | 3.47E-06 | 0.000631823 | ENSG00000078699 | CBFA2T2 |
| **634.6757273** | 0.229206348 | 0.074600751 | 3.071318164 | 0.002131159 | 0.047956359 | ENSG00000094975 | SUCO |
| **714.0403546** | -0.257781099 | 0.066807444 | -3.857841737 | 0.000114393 | 0.008524776 | ENSG00000104872 | PIH1D1 |
| **337.4585994** | -0.293301874 | 0.09347911 | -3.139146782 | 0.001694406 | 0.042788446 | ENSG00000188677 | PARVB |
| **2291.606081** | -0.320114195 | 0.076914673 | -4.161700052 | 3.16E-05 | 0.003300839 | ENSG00000197451 | HNRNPAB |
| **1674.082586** | -0.328769404 | 0.07881609 | -4.171417453 | 3.03E-05 | 0.00323758 | ENSG00000065911 | MTHFD2 |
| **1194.136689** | -0.342928347 | 0.089952822 | -3.811737612 | 0.000137993 | 0.009649979 | ENSG00000173457 | PPP1R14B |
| **112.6592751** | -0.404115658 | 0.128093772 | -3.155350152 | 0.001603056 | 0.041490297 | ENSG00000175906 | ARL4D |
| **227.9457112** | -0.434730426 | 0.125528284 | -3.461863413 | 0.000536449 | 0.021967842 | ENSG00000125968 | ID1 |
| **140.4043512** | -0.445653494 | 0.137765281 | -3.23488451 | 0.001216919 | 0.035687144 | ENSG00000109255 | NMU |
| **87.57482141** | -0.558269046 | 0.160174923 | -3.480318705 | 0.000500818 | 0.021240448 | ENSG00000136732 | GYPC |
| **55.67753162** | -0.630138253 | 0.19402518 | -3.277276797 | 0.001048136 | 0.032585032 | ENSG00000242950 | ERVW-1 |
| **5885.130515** | -0.664594343 | 0.111017965 | -5.988945082 | 2.11E-09 | 1.48E-06 | ENSG00000205358 | MT1H |
| **124.8394543** | -0.669412895 | 0.209320027 | -3.300311739 | 0.000965775 | 0.031319972 | ENSG00000135074 | ADAM19 |

Supplemental Table S2_Sheet_1

| | p_val | avg_log2FC | pct.1 | pct.2 | p_val_adj |
|---|---|---|---|---|---|
| **AGRN** | 0 | -1.51175387214881 | 0.94 | 0.998 | 0 |
| **MXRA8** | 0 | -1.60507164992287 | 0.757 | 0.986 | 0 |
| **ATAD3B** | 0 | 1.85390392556141 | 1 | 0.956 | 0 |
| **FNDC10** | 0 | 1.63783803238037 | 0.977 | 0.588 | 0 |
| **HES2** | 0 | 1.0573762708181 | 0.651 | 0.068 | 0 |
| **CLSTN1** | 0 | -1.12797684819481 | 0.996 | 0.999 | 0 |
| **CTNNBIP1** | 0 | -1.14098451194057 | 0.911 | 0.992 | 0 |
| **DFFA** | 0 | 1.02705237755463 | 1 | 0.996 | 0 |
| **SRM** | 0 | 0.972037021878845 | 1 | 1 | 0 |
| **MTOR** | 0 | 0.947660713922573 | 0.998 | 0.954 | 0 |
| **DRAXIN** | 0 | -2.02975614941495 | 0.635 | 0.965 | 0 |
| **MIIP** | 0 | 0.888438187700828 | 1 | 0.99 | 0 |
| **TNFRSF8** | 0 | 1.50801020942278 | 0.738 | 0.067 | 0 |
| **DHRS3** | 0 | -1.42680862563001 | 0.376 | 0.812 | 0 |
| **EFHD2** | 0 | 1.3833350317845 | 0.999 | 0.96 | 0 |
| **EPHA2** | 0 | -2.08410901695482 | 0.041 | 0.752 | 0 |
| **KLHDC7A** | 0 | -1.10700340603936 | 0.003 | 0.525 | 0 |
| **CDA** | 0 | 1.27436294929311 | 0.988 | 0.794 | 0 |
| **PINK1** | 0 | 1.19484433795775 | 1 | 0.957 | 0 |
| **ECE1** | 0 | -2.18488976925175 | 0.667 | 0.994 | 0 |
| **ID3** | 0 | -3.47976175086542 | 0.039 | 0.806 | 0 |
| **FUCA1** | 0 | 1.89745146348357 | 0.999 | 0.814 | 0 |
| **CNR2** | 0 | 2.01077474235312 | 0.898 | 0.137 | 0 |
| **GRHL3** | 0 | -1.70481914060004 | 0.081 | 0.6 | 0 |
| **MAN1C1** | 0 | 1.3038131351233 | 0.999 | 0.886 | 0 |
| **STMN1** | 0 | 1.59976743142086 | 1 | 0.983 | 0 |
| **LIN28A** | 0 | -1.66086928877914 | 1 | 1 | 0 |
| **SLC9A1** | 0 | 1.05045828611108 | 0.982 | 0.811 | 0 |
| **RPA2** | 0 | 1.27156402833874 | 0.963 | 0.58 | 0 |
| **SMPDL3B** | 0 | -2.27519667861454 | 0.035 | 0.819 | 0 |
| **PTPRU** | 0 | 1.86442840151946 | 1 | 0.937 | 0 |
| **NKAIN1** | 0 | -1.48791797128145 | 0.089 | 0.604 | 0 |

| | | | | | |
|---|---|---|---|---|---|
| **EIF3I** | 0 | 1.37934276504115 | 0.947 | 0.445 | 0 |
| **MARCKSL1** | 0 | -1.15535238096178 | 1 | 1 | 0 |
| **KIAA1522** | 0 | -1.48052743686935 | 0.941 | 0.999 | 0 |
| **TMEM54** | 0 | 1.5828003913938 | 0.896 | 0.25 | 0 |
| **ZNF362** | 0 | -1.18162000987274 | 0.822 | 0.986 | 0 |
| **PHC2** | 0 | -1.82426304424256 | 0.022 | 0.589 | 0 |
| **MYCBP** | 0 | 1.45380888845509 | 0.953 | 0.444 | 0 |
| **AKIRIN1** | 0 | 1.12451133432121 | 1 | 0.997 | 0 |
| **PPT1** | 0 | -1.02254582855151 | 0.963 | 0.996 | 0 |
| **YBX1** | 0 | -0.945009298793123 | 1 | 1 | 0 |
| **CLDN19** | 0 | -1.24312373901205 | 0.046 | 0.608 | 0 |
| **HPDL** | 0 | 2.20602803015841 | 0.946 | 0.154 | 0 |
| **PRDX1** | 0 | 2.60674727079587 | 1 | 0.991 | 0 |
| **PIK3R3** | 0 | -1.66928887722775 | 0.574 | 0.914 | 0 |
| **OSBPL9** | 0 | -0.905882603688861 | 0.977 | 0.998 | 0 |
| **NRDC** | 0 | 0.954653646112424 | 1 | 1 | 0 |
| **ZYG11A** | 0 | 1.53509953458241 | 1 | 0.844 | 0 |
| **DHCR24** | 0 | -1.42522959169158 | 0.998 | 1 | 0 |
| **PLPP3** | 0 | -1.40191045209973 | 0.577 | 0.939 | 0 |
| **PGM1** | 0 | -1.41517290153022 | 0.505 | 0.953 | 0 |
| **ROR1** | 0 | -2.60745690598501 | 0.037 | 0.832 | 0 |
| **AK4** | 0 | 1.79178387992507 | 0.992 | 0.884 | 0 |
| **WLS** | 0 | -4.89061147402189 | 0.388 | 0.988 | 0 |
| **CTH** | 0 | 1.49683721891332 | 0.97 | 0.714 | 0 |
| **ST6GALNAC5** | 0 | -1.40081660068709 | 0.073 | 0.632 | 0 |
| **FUBP1** | 0 | -0.97743857752907 | 0.993 | 1 | 0 |
| **LMO4** | 0 | 1.3281214862262 | 0.999 | 0.939 | 0 |
| **TGFBR3** | 0 | 1.56785771346486 | 0.857 | 0.176 | 0 |
| **BRDT** | 0 | 2.89859269282102 | 0.98 | 0.255 | 0 |
| **FNBP1L** | 0 | -0.964587616312201 | 1 | 1 | 0 |
| **PTBP2** | 0 | -1.37957464613758 | 0.699 | 0.957 | 0 |
| **AGL** | 0 | 2.09310755048595 | 1 | 0.958 | 0 |
| **SLC35A3** | 0 | 1.14667385993189 | 0.819 | 0.215 | 0 |
| **SLC25A24** | 0 | -1.2401797265727 | 0.857 | 0.993 | 0 |

| | | | | | |
|---|---|---|---|---|---|
| **FAM102B** | 0 | 1.15352642735689 | 0.985 | 0.837 | 0 |
| **HENMT1** | 0 | 2.03225662744746 | 0.94 | 0.114 | 0 |
| **TAF13** | 0 | 3.0893076017968 | 1 | 0.839 | 0 |
| **GNAI3** | 0 | -0.856397319916381 | 1 | 1 | 0 |
| **CAPZA1** | 0 | 1.1499299872781 | 0.889 | 0.347 | 0 |
| **OLFML3** | 0 | -4.95415613257267 | 0.453 | 0.996 | 0 |
| **SYT6** | 0 | -1.41000027847987 | 0.157 | 0.736 | 0 |
| **BCAS2** | 0 | 1.09215290108829 | 0.941 | 0.551 | 0 |
| **NOTCH2** | 0 | -1.4716484525677 | 0.427 | 0.882 | 0 |
| **SV2A** | 0 | -1.5655180433211 | 0.13 | 0.776 | 0 |
| **PLEKHO1** | 0 | -2.04158786727371 | 0.306 | 0.902 | 0 |
| **HORMAD1** | 0 | 2.75951003508727 | 0.996 | 0.202 | 0 |
| **MLLT11** | 0 | -1.79161562430349 | 0.451 | 0.907 | 0 |
| **S100A10** | 0 | 1.26268318718015 | 1 | 0.992 | 0 |
| **RAB13** | 0 | -1.1374753746264 | 0.963 | 0.996 | 0 |
| **CKS1B** | 0 | 2.1296319265542 | 1 | 0.932 | 0 |
| **FLAD1** | 0 | 1.32280023097872 | 1 | 0.966 | 0 |
| **THBS3** | 0 | -1.38507666329447 | 0.105 | 0.643 | 0 |
| **FDPS** | 0 | -0.951258975951816 | 0.977 | 0.995 | 0 |
| **MEX3A** | 0 | -2.33822214895428 | 0.21 | 0.898 | 0 |
| **NAXE** | 0 | -1.34747218781 | 0.731 | 0.985 | 0 |
| **NES** | 0 | -2.50145025238475 | 0.976 | 1 | 0 |
| **CRABP2** | 0 | -3.31454417995224 | 0.846 | 0.994 | 0 |
| **DUSP23** | 0 | 1.93359417215265 | 0.933 | 0.098 | 0 |
| **IGSF8** | 0 | -1.51320861291758 | 0.674 | 0.94 | 0 |
| **ATP1A2** | 0 | -1.35749830943994 | 0.002 | 0.504 | 0 |
| **PEA15** | 0 | -1.97071247643955 | 0.739 | 0.994 | 0 |
| **VANGL2** | 0 | -1.32455316410369 | 0.757 | 0.973 | 0 |
| **F11R** | 0 | 1.27908202164966 | 1 | 0.988 | 0 |
| **TSTD1** | 0 | 1.11670377591501 | 0.646 | 0.04 | 0 |
| **RGS5** | 0 | -3.14127860450211 | 0.096 | 0.851 | 0 |
| **MGST3** | 0 | -1.61281536215355 | 0.864 | 0.997 | 0 |
| **ALDH9A1** | 0 | -0.896476515123238 | 0.931 | 0.996 | 0 |
| **MAEL** | 0 | 1.42598584260879 | 0.61 | 0.022 | 0 |

| | | | | | |
|---|---|---|---|---|---|
| **TOR3A** | 0 | 0.945313827264915 | 0.988 | 0.876 | 0 |
| **RO60** | 0 | -1.19097826908902 | 0.885 | 0.997 | 0 |
| **GLRX2** | 0 | -1.12061082223348 | 0.054 | 0.638 | 0 |
| **CDC73** | 0 | -1.0213797416529 | 0.937 | 0.998 | 0 |
| **LAD1** | 0 | 1.37443062797826 | 0.739 | 0.085 | 0 |
| **TIMM17A** | 0 | 0.867513029016221 | 0.999 | 0.978 | 0 |
| **ELF3** | 0 | 1.83085632064605 | 0.996 | 0.697 | 0 |
| **KDM5B** | 0 | 0.913671417808302 | 1 | 0.995 | 0 |
| **SNRPE** | 0 | 1.47226283426435 | 1 | 0.865 | 0 |
| **LRRN2** | 0 | -1.12715779427513 | 0.035 | 0.554 | 0 |
| **CR1L** | 0 | 2.5158795582419 | 0.997 | 0.415 | 0 |
| **IRF6** | 0 | 1.20480259564818 | 0.745 | 0.048 | 0 |
| **SERTAD4** | 0 | -1.221689523406 | 0.019 | 0.589 | 0 |
| **TRAF5** | 0 | -1.83588723367863 | 0.041 | 0.765 | 0 |
| **SMYD2** | 0 | -1.52606528703346 | 0.739 | 0.979 | 0 |
| **PTPN14** | 0 | -1.26467553539325 | 0.868 | 0.998 | 0 |
| **DUSP10** | 0 | -1.31963696674629 | 0.253 | 0.772 | 0 |
| **CAPN2** | 0 | -2.5847539740113 | 0.165 | 0.869 | 0 |
| **CNIH4** | 0 | 1.83710713253631 | 0.999 | 0.776 | 0 |
| **H3F3A** | 0 | -0.823400814466762 | 1 | 1 | 0 |
| **GNG4** | 0 | 1.56820520865039 | 0.787 | 0.144 | 0 |
| **LGALS8** | 0 | 1.03354339435648 | 0.616 | 0.016 | 0 |
| **ZBTB18** | 0 | 1.09142792794681 | 0.961 | 0.7 | 0 |
| **COX20** | 0 | 1.06505045597449 | 0.864 | 0.353 | 0 |
| **ACP1** | 0 | 1.28323463017121 | 0.973 | 0.594 | 0 |
| **ID2** | 0 | -1.97760331112233 | 0.291 | 0.855 | 0 |
| **IAH1** | 0 | 2.02714442741989 | 0.924 | 0.101 | 0 |
| **WDR35** | 0 | -1.19992875820609 | 0.198 | 0.731 | 0 |
| **EFR3B** | 0 | 1.68368371301581 | 0.996 | 0.829 | 0 |
| **CAPN13** | 0 | -1.26125427931528 | 0.024 | 0.667 | 0 |
| **YIPF4** | 0 | 0.975519174060643 | 0.997 | 0.955 | 0 |
| **FEZ2** | 0 | 0.987047243871797 | 0.996 | 0.952 | 0 |
| **CDC42EP3** | 0 | -1.27249327625187 | 0.145 | 0.675 | 0 |
| **ATL2** | 0 | 0.985376061803379 | 0.999 | 0.976 | 0 |

| | | | | | |
|---|---|---|---|---|---|
| **PKDCC** | 0 | -3.86155844304184 | 0.026 | 0.767 | 0 |
| **EML4** | 0 | -1.02565744571542 | 0.951 | 0.997 | 0 |
| **PSME4** | 0 | 0.807261532018829 | 1 | 0.998 | 0 |
| **RTN4** | 0 | -1.04542704375911 | 0.999 | 1 | 0 |
| **CCDC88A** | 0 | -1.42096804233539 | 0.749 | 0.991 | 0 |
| **XPO1** | 0 | 1.31587414576663 | 0.999 | 0.926 | 0 |
| **B3GNT2** | 0 | -1.52908136370032 | 0.897 | 0.989 | 0 |
| **UGP2** | 0 | 0.992202330482205 | 1 | 0.938 | 0 |
| **CEP68** | 0 | -1.19363757149858 | 0.629 | 0.933 | 0 |
| **SPRED2** | 0 | -1.24404666122737 | 0.331 | 0.795 | 0 |
| **ANTXR1** | 0 | -1.3099650955601 | 0.173 | 0.728 | 0 |
| **GFPT1** | 0 | 1.39905520238776 | 1 | 0.998 | 0 |
| **SNRPG** | 0 | 1.47367151206399 | 1 | 0.917 | 0 |
| **FAM136A** | 0 | 1.00993807921032 | 1 | 0.959 | 0 |
| **MTHFD2** | 0 | 1.73488334656704 | 1 | 0.994 | 0 |
| **HK2** | 0 | 1.32889160193556 | 1 | 0.965 | 0 |
| **CAPG** | 0 | 0.914996079234931 | 1 | 0.936 | 0 |
| **REEP1** | 0 | 1.48939907679729 | 0.779 | 0.085 | 0 |
| **CNOT11** | 0 | 0.935194992049987 | 1 | 0.998 | 0 |
| **MAP4K4** | 0 | -1.17615476453296 | 0.997 | 1 | 0 |
| **DBI** | 0 | -0.986216685276157 | 0.977 | 0.999 | 0 |
| **GLI2** | 0 | -1.91247557703128 | 0.07 | 0.858 | 0 |
| **TFCP2L1** | 0 | 1.4395092658543 | 0.976 | 0.631 | 0 |
| **NIFK** | 0 | 1.10643922671469 | 0.999 | 0.941 | 0 |
| **TSN** | 0 | -0.947654430922789 | 0.996 | 0.999 | 0 |
| **BIN1** | 0 | -2.31336033363547 | 0.424 | 0.972 | 0 |
| **PLEKHB2** | 0 | 0.812237621962876 | 1 | 0.999 | 0 |
| **LYPD1** | 0 | -2.22231870646737 | 0.048 | 0.65 | 0 |
| **ZEB2** | 0 | -1.76591990584619 | 0.046 | 0.601 | 0 |
| **KIF5C** | 0 | -2.23908296986479 | 0.673 | 0.998 | 0 |
| **NMI** | 0 | 1.21090011466096 | 0.955 | 0.522 | 0 |
| **FMNL2** | 0 | 0.87392852107067 | 1 | 0.988 | 0 |
| **RPRM** | 0 | -1.82855370057086 | 0.431 | 0.868 | 0 |
| **GALNT3** | 0 | 1.31188112461032 | 0.683 | 0.005 | 0 |

| | | | | | |
|---|---|---|---|---|---|
| **CERS6** | 0 | -1.33128202756932 | 0.245 | 0.802 | 0 |
| **SP5** | 0 | -3.79314333288753 | 0.141 | 0.946 | 0 |
| **GAD1** | 0 | -2.47088743748699 | 0.012 | 0.874 | 0 |
| **UBE2E3** | 0 | -1.6696198834255 | 0.798 | 0.997 | 0 |
| **NCKAP1** | 0 | -0.816187475479632 | 0.997 | 1 | 0 |
| **CFLAR** | 0 | 1.08094266338642 | 0.997 | 0.863 | 0 |
| **FZD7** | 0 | -4.12289396223224 | 0.076 | 0.9 | 0 |
| **ICA1L** | 0 | 1.78300981312982 | 0.902 | 0.139 | 0 |
| **CCNYL1** | 0 | -1.21932782131572 | 0.644 | 0.972 | 0 |
| **FZD5** | 0 | 1.60450733674834 | 1 | 0.952 | 0 |
| **IDH1** | 0 | 1.72534098420835 | 1 | 0.975 | 0 |
| **CPS1** | 0 | -1.04032293104012 | 0.722 | 0.98 | 0 |
| **BARD1** | 0 | 1.25362734646445 | 0.988 | 0.812 | 0 |
| **TMBIM1** | 0 | 2.20839137524793 | 0.989 | 0.448 | 0 |
| **OBSL1** | 0 | -2.39142973905985 | 0.272 | 0.971 | 0 |
| **ACSL3** | 0 | 1.03194146220377 | 1 | 0.999 | 0 |
| **SERPINE2** | 0 | -1.34655728538072 | 0.511 | 0.91 | 0 |
| **FBXO36** | 0 | 1.01608461890789 | 0.997 | 0.873 | 0 |
| **ALPG** | 0 | 4.10050761031499 | 1 | 0.61 | 0 |
| **NGEF** | 0 | 1.78904657390464 | 0.831 | 0.117 | 0 |
| **GBX2** | 0 | -1.41580030230055 | 0.132 | 0.682 | 0 |
| **LRRN1** | 0 | -2.9725978339271 | 0.381 | 0.977 | 0 |
| **ARL8B** | 0 | 0.9860528172749 | 0.996 | 0.937 | 0 |
| **EDEM1** | 0 | 1.16224705786146 | 0.985 | 0.814 | 0 |
| **BRK1** | 0 | 1.00944358335114 | 0.699 | 0.079 | 0 |
| **CHCHD4** | 0 | 1.12619507692259 | 0.973 | 0.706 | 0 |
| **3,00 LSM** | 0 | 0.911191659712868 | 0.65 | 0.065 | 0 |
| **SLC6A6** | 0 | 1.46159156572806 | 0.997 | 0.863 | 0 |
| **RAB5A** | 0 | 1.19256361205067 | 1 | 0.949 | 0 |
| **TRIM71** | 0 | -1.13141097006657 | 1 | 0.999 | 0 |
| **TRANK1** | 0 | 1.61622899388134 | 0.649 | 0.09 | 0 |
| **CTNNB1** | 0 | -1.43511623522178 | 0.985 | 0.999 | 0 |
| **HIGD1A** | 0 | 1.16514724068058 | 0.955 | 0.613 | 0 |
| **ZNF502** | 0 | 1.25037329321239 | 0.727 | 0.008 | 0 |

| | | | | | |
|---|---|---|---|---|---|
| **PTH1R** | 0 | 1.82854116319036 | 0.81 | 0.268 | 0 |
| **PRKAR2A** | 0 | 1.20447325641705 | 1 | 0.972 | 0 |
| **WNT5A** | 0 | -1.75737845809833 | 0.015 | 0.721 | 0 |
| **APPL1** | 0 | 0.89561666235232 | 0.992 | 0.886 | 0 |
| **PDE12** | 0 | 1.05276692126768 | 0.942 | 0.559 | 0 |
| **MAGI1** | 0 | -1.37088005160823 | 0.433 | 0.873 | 0 |
| **DCBLD2** | 0 | -1.17948675096121 | 0.861 | 0.992 | 0 |
| **TBC1D23** | 0 | 1.19539086946415 | 1 | 0.975 | 0 |
| **NIT2** | 0 | 1.51311663789431 | 0.985 | 0.646 | 0 |
| **ALCAM** | 0 | -1.77326613723478 | 0.04 | 0.7 | 0 |
| **DPPA2** | 0 | 2.18111561684534 | 0.993 | 0.457 | 0 |
| **NECTIN3** | 0 | -1.87025807476917 | 0.318 | 0.917 | 0 |
| **PHLDB2** | 0 | -1.3586716632146 | 0.186 | 0.731 | 0 |
| **ATG3** | 0 | 2.35965513820915 | 1 | 0.995 | 0 |
| **NAA50** | 0 | 1.95167587604408 | 0.987 | 0.447 | 0 |
| **ZDHHC23** | 0 | 1.16012349648737 | 0.782 | 0.222 | 0 |
| **B4GALT4** | 0 | 1.55279101872384 | 0.941 | 0.396 | 0 |
| **FSTL1** | 0 | -2.3338614328892 | 0.863 | 0.978 | 0 |
| **NDUFB4** | 0 | 1.36742267385618 | 0.921 | 0.363 | 0 |
| **SNX4** | 0 | 1.10431108404757 | 0.999 | 0.987 | 0 |
| **MGLL** | 0 | 1.30289259590592 | 0.669 | 0.048 | 0 |
| **PLXND1** | 0 | 1.7795126920069 | 0.996 | 0.761 | 0 |
| **TRH** | 0 | 1.5269948850801 | 0.774 | 0.083 | 0 |
| **AMOTL2** | 0 | -2.02806737745008 | 0.867 | 0.996 | 0 |
| **RBP1** | 0 | -2.31288539098307 | 0.35 | 0.939 | 0 |
| **SPSB4** | 0 | -2.49755925199649 | 0.148 | 0.927 | 0 |
| **CHST2** | 0 | 2.92221516281746 | 0.992 | 0.512 | 0 |
| **WWTR1** | 0 | 1.24863118434345 | 0.996 | 0.854 | 0 |
| **PFN2** | 0 | -1.30311686698992 | 0.997 | 1 | 0 |
| **RAP2B** | 0 | 1.16912399745765 | 1 | 0.996 | 0 |
| **LRRC34** | 0 | 0.891292713600871 | 0.549 | 0.005 | 0 |
| **GNB4** | 0 | -1.51494646920441 | 0.674 | 0.972 | 0 |
| **USP13** | 0 | 1.03552371939417 | 0.994 | 0.952 | 0 |
| **LIPH** | 0 | 1.80421115293123 | 0.784 | 0.077 | 0 |

| | | | | | |
|---|---|---|---|---|---|
| **SENP2** | 0 | 0.929396457993029 | 1 | 0.996 | 0 |
| **LPP** | 0 | -2.19814995996134 | 0.314 | 0.933 | 0 |
| **HES1** | 0 | -2.44683726532032 | 0.027 | 0.724 | 0 |
| **RNF168** | 0 | 0.968526720325573 | 1 | 0.971 | 0 |
| **NRROS** | 0 | 0.976811249552226 | 0.613 | 0.056 | 0 |
| **PAK2** | 0 | 1.11982856313689 | 0.995 | 0.87 | 0 |
| **ZNF595** | 0 | 1.50361892436967 | 0.985 | 0.75 | 0 |
| **ZNF732** | 0 | 1.29712097422544 | 0.694 | 0.005 | 0 |
| **NSD2** | 0 | -1.31852947268865 | 0.306 | 0.803 | 0 |
| **MFSD10** | 0 | -1.30089985448063 | 0.643 | 0.963 | 0 |
| **S100P** | 0 | 1.52085154473596 | 0.634 | 0.064 | 0 |
| **PI4K2B** | 0 | 1.578202698669 | 0.996 | 0.864 | 0 |
| **SEL1L3** | 0 | -1.20433101233805 | 0.058 | 0.589 | 0 |
| **RELL1** | 0 | 1.16438152689192 | 0.98 | 0.753 | 0 |
| **UCHL1** | 0 | -2.88167690041712 | 0.487 | 0.992 | 0 |
| **LIMCH1** | 0 | -2.24382476294963 | 0.171 | 0.837 | 0 |
| **TEC** | 0 | 1.20848041565733 | 0.747 | 0.113 | 0 |
| **OCIAD2** | 0 | -0.950245500019684 | 0.049 | 0.567 | 0 |
| **PAICS** | 0 | 1.24431116187709 | 0.997 | 0.806 | 0 |
| **IGFBP7** | 0 | -1.94548428161529 | 0.09 | 0.615 | 0 |
| **GRSF1** | 0 | 0.925107291264022 | 1 | 0.987 | 0 |
| **G3BP2** | 0 | -0.952093198903684 | 1 | 1 | 0 |
| **SEPTIN11** | 0 | -1.23737308135571 | 0.979 | 1 | 0 |
| **TMEM150C** | 0 | 1.64815392931011 | 0.863 | 0.163 | 0 |
| **MAPK10** | 0 | -2.20812839585646 | 0.006 | 0.706 | 0 |
| **SPP1** | 0 | -2.55775926267412 | 0.236 | 0.813 | 0 |
| **GRID2** | 0 | -1.57324742097545 | 0.102 | 0.653 | 0 |
| **EIF4E** | 0 | 0.856923810482945 | 0.997 | 0.96 | 0 |
| **PAPSS1** | 0 | -0.995575243101915 | 0.973 | 0.998 | 0 |
| **CAMK2D** | 0 | -1.67902257434819 | 0.129 | 0.801 | 0 |
| **PRSS12** | 0 | 2.19663236972844 | 0.984 | 0.562 | 0 |
| **MAD2L1** | 0 | 1.51553052042662 | 0.937 | 0.613 | 0 |
| **PCDH18** | 0 | -2.36540583873455 | 0.561 | 0.929 | 0 |
| **NOCT** | 0 | 1.16631867181115 | 0.992 | 0.863 | 0 |

| | | | | | |
|---|---|---|---|---|---|
| **GAB1** | 0 | 1.28972352329379 | 0.932 | 0.571 | 0 |
| **ABCE1** | 0 | 1.13655274128111 | 0.993 | 0.792 | 0 |
| **OTUD4** | 0 | 1.7009599310815 | 0.999 | 0.83 | 0 |
| **SFRP2** | 0 | -2.58039703147326 | 0.042 | 0.722 | 0 |
| **TMEM144** | 0 | 1.29482707557787 | 0.774 | 0.205 | 0 |
| **TRIM61** | 0 | 1.50825721927071 | 0.927 | 0.332 | 0 |
| **TRIM60** | 0 | 2.53506620706078 | 0.987 | 0.189 | 0 |
| **TRIM75P** | 0 | 2.60244097521036 | 0.983 | 0.217 | 0 |
| **FAT1** | 0 | -2.5278737623014 | 0.749 | 0.999 | 0 |
| **ZFP42** | 0 | 4.21432052332842 | 1 | 0.385 | 0 |
| **SLC9A3** | 0 | 1.19440210749863 | 0.661 | 0.052 | 0 |
| **IRX1** | 0 | -2.00618366565354 | 0.282 | 0.797 | 0 |
| **SUB1** | 0 | 2.14666795612171 | 0.99 | 0.378 | 0 |
| **SELENOP** | 0 | -2.55513286109155 | 0.12 | 0.941 | 0 |
| **CCL28** | 0 | 0.937146538480969 | 0.571 | 0.004 | 0 |
| **FST** | 0 | -3.24254409598877 | 0.165 | 0.828 | 0 |
| **PLPP1** | 0 | 1.16489765207572 | 1 | 0.968 | 0 |
| **ZSWIM6** | 0 | -1.33474909626703 | 0.113 | 0.706 | 0 |
| **ERBIN** | 0 | 1.57452279811396 | 1 | 0.957 | 0 |
| **MARVELD2** | 0 | 1.25895601864688 | 0.89 | 0.298 | 0 |
| **MAP1B** | 0 | -2.79993671078584 | 0.88 | 1 | 0 |
| **ARSB** | 0 | -1.20596448876611 | 0.548 | 0.914 | 0 |
| **VCAN** | 0 | -3.16034530462674 | 0.61 | 0.996 | 0 |
| **CAST** | 0 | -1.31241042842076 | 0.265 | 0.808 | 0 |
| **PAM** | 0 | -1.72963090525371 | 0.053 | 0.723 | 0 |
| **EFNA5** | 0 | -2.08210885180693 | 0.264 | 0.899 | 0 |
| **MCC** | 0 | -1.50875386397935 | 0.049 | 0.74 | 0 |
| **ZNF608** | 0 | -2.2100287683872 | 0.211 | 0.946 | 0 |
| **ADAMTS19** | 0 | -2.16861651330444 | 0.004 | 0.724 | 0 |
| **HINT1** | 0 | 1.03369283372021 | 1 | 0.977 | 0 |
| **WNT8A** | 0 | -2.5074444217042 | 0.094 | 0.785 | 0 |
| **CXXC5** | 0 | -1.69568903870194 | 0.828 | 0.994 | 0 |
| **PCDHA4** | 0 | -1.02543339838848 | 0.065 | 0.609 | 0 |
| **PCDHB2** | 0 | -1.99761729894733 | 0.252 | 0.917 | 0 |

| | | | | | |
|---|---|---|---|---|---|
| **PCDHGB7** | 0 | -1.54979245239746 | 0.172 | 0.758 | 0 |
| **DIAPH1** | 0 | -1.2469661272256 | 0.995 | 0.999 | 0 |
| **PCDH1** | 0 | -2.37128034931023 | 0.685 | 0.987 | 0 |
| **GNPDA1** | 0 | -1.27936622125016 | 0.807 | 0.984 | 0 |
| **NDFIP1** | 0 | -1.11452203001211 | 0.758 | 0.968 | 0 |
| **DPYSL3** | 0 | -2.22222730801285 | 0.872 | 0.999 | 0 |
| **UBLCP1** | 0 | -0.878324414035315 | 0.955 | 0.997 | 0 |
| **PANK3** | 0 | 1.44458159055238 | 0.995 | 0.784 | 0 |
| **ATP6V0E1** | 0 | 0.819287735849317 | 1 | 0.988 | 0 |
| **NSD1** | 0 | 0.999697411163921 | 1 | 0.967 | 0 |
| **DBN1** | 0 | -1.34746761520876 | 0.995 | 1 | 0 |
| **PDLIM7** | 0 | -1.49304723988973 | 0.968 | 0.997 | 0 |
| **SERPINB1** | 0 | 1.86721329098673 | 0.918 | 0.106 | 0 |
| **SERPINB6** | 0 | 2.70473681559768 | 1 | 0.704 | 0 |
| **NQO2** | 0 | 0.888887699130358 | 1 | 0.969 | 0 |
| **DSP** | 0 | -1.40080788303802 | 0.995 | 0.997 | 0 |
| **EDN1** | 0 | -2.95957807236048 | 0.015 | 0.814 | 0 |
| **JARID2** | 0 | -1.16607958795242 | 0.971 | 0.991 | 0 |
| **DCDC2** | 0 | 1.21958488165749 | 0.697 | 0.112 | 0 |
| **HIST1H1A** | 0 | 4.74126808948193 | 0.997 | 0.424 | 0 |
| **HIST1H2BB** | 0 | 2.0536191307322 | 0.69 | 0.056 | 0 |
| **ZFP57** | 0 | 2.36551057700449 | 0.977 | 0.238 | 0 |
| **TRIM26** | 0 | 1.26452119292478 | 0.999 | 0.928 | 0 |
| **NRM** | 0 | -1.21369857901404 | 0.43 | 0.848 | 0 |
| **FLOT1** | 0 | -1.29237543660727 | 0.901 | 0.997 | 0 |
| **DDAH2** | 0 | -1.42483864629587 | 0.82 | 0.988 | 0 |
| **HSPA1B** | 0 | 1.48161187015697 | 0.86 | 0.417 | 0 |
| **AGPAT1** | 0 | -0.900069880275536 | 0.988 | 1 | 0 |
| **PBX2** | 0 | -0.873174103034627 | 0.98 | 0.998 | 0 |
| **CUTA** | 0 | -1.38182095441966 | 0.991 | 1 | 0 |
| **ITPR3** | 0 | 1.29462344912756 | 0.942 | 0.565 | 0 |
| **HMGA1** | 0 | -0.979760357151321 | 0.987 | 0.992 | 0 |
| **ILRUN** | 0 | 1.79585369331317 | 1 | 0.996 | 0 |
| **SNRPC** | 0 | 0.995408234153668 | 1 | 1 | 0 |

| | | | | | |
|---|---|---|---|---|---|
| **MAPK13** | 0 | 1.35845105411934 | 0.863 | 0.277 | 0 |
| **MDFI** | 0 | 1.26010084202225 | 0.988 | 0.823 | 0 |
| **TOMM6** | 0 | 1.31309776783389 | 1 | 0.956 | 0 |
| **CCND3** | 0 | 1.48024031477268 | 0.981 | 0.709 | 0 |
| **PTK7** | 0 | -1.31154586848207 | 0.935 | 0.989 | 0 |
| **HSP90AB1** | 0 | 0.887192902325112 | 1 | 0.999 | 0 |
| **AARS2** | 0 | 0.980218876679913 | 0.994 | 0.917 | 0 |
| **TNFRSF21** | 0 | -1.47318108130828 | 0.938 | 0.996 | 0 |
| **DPPA5** | 0 | 5.91605871049798 | 1 | 0.524 | 0 |
| **KHDC3L** | 0 | 4.68747357239352 | 0.999 | 0.416 | 0 |
| **DDX43** | 0 | 1.61542080205469 | 0.819 | 0.029 | 0 |
| **CGAS** | 0 | 1.63297507523425 | 0.704 | 0.028 | 0 |
| **IRAK1BP1** | 0 | 1.51947138386933 | 0.763 | 0.01 | 0 |
| **EPHA7** | 0 | -1.6775638916619 | 0.709 | 0.952 | 0 |
| **CCNC** | 0 | 1.30346829609977 | 1 | 0.981 | 0 |
| **LIN28B** | 0 | 1.25228530856911 | 1 | 0.993 | 0 |
| **PRDM1** | 0 | 1.23632095553306 | 0.92 | 0.464 | 0 |
| **ATG5** | 0 | 1.72019866454896 | 1 | 0.949 | 0 |
| **SEC63** | 0 | 1.33105491738099 | 1 | 0.989 | 0 |
| **SNX3** | 0 | 1.45083101354752 | 0.997 | 0.828 | 0 |
| **CD164** | 0 | 0.947570951991038 | 0.946 | 0.608 | 0 |
| **WASF1** | 0 | -1.6580132430709 | 0.504 | 0.94 | 0 |
| **GTF3C6** | 0 | 1.54692398491719 | 0.989 | 0.664 | 0 |
| **SLC16A10** | 0 | 2.47748259272433 | 0.994 | 0.544 | 0 |
| **MARCKS** | 0 | -3.78593891868896 | 0.29 | 0.997 | 0 |
| **PKIB** | 0 | 1.23916658027775 | 0.64 | 0.047 | 0 |
| **SMPDL3A** | 0 | 1.66206285968305 | 0.955 | 0.403 | 0 |
| **ECHDC1** | 0 | -1.78507583218659 | 0.201 | 0.935 | 0 |
| **SGK1** | 0 | 1.61180745886832 | 0.998 | 0.817 | 0 |
| **HEBP2** | 0 | 1.20419634068449 | 1 | 0.988 | 0 |
| **FUCA2** | 0 | -1.25012239999769 | 0.824 | 0.965 | 0 |
| **PHACTR2** | 0 | -1.57574659602759 | 0.39 | 0.897 | 0 |
| **UTRN** | 0 | -1.85091922958354 | 0.507 | 0.936 | 0 |
| **RMND1** | 0 | 1.34766418472163 | 1 | 0.952 | 0 |

| | | | | | |
|---|---|---|---|---|---|
| **ARMT1** | 0 | 1.20658872452273 | 1 | 0.988 | 0 |
| **TCP1** | 0 | 2.74230704902971 | 1 | 0.924 | 0 |
| **IGF2R** | 0 | -2.0767261033254 | 0.469 | 0.978 | 0 |
| **DACT2** | 0 | 1.66128394717399 | 0.845 | 0.095 | 0 |
| **C7orf50** | 0 | -0.95671033087727 | 0.99 | 1 | 0 |
| **TNRC18** | 0 | 0.953542884550781 | 1 | 0.971 | 0 |
| **BZW2** | 0 | 0.902787473317122 | 1 | 0.977 | 0 |
| **CYCS** | 0 | 2.06755857234939 | 0.995 | 0.613 | 0 |
| **CBX3** | 0 | 2.49357427492939 | 1 | 0.834 | 0 |
| **CPVL** | 0 | -1.1484688261266 | 0.986 | 0.998 | 0 |
| **WIPF3** | 0 | 1.44566873624812 | 0.86 | 0.233 | 0 |
| **DPY19L1** | 0 | 1.48186441322881 | 0.964 | 0.538 | 0 |
| **STARD3NL** | 0 | -1.18835506867597 | 0.772 | 0.978 | 0 |
| **HUS1** | 0 | 0.878391292513386 | 0.998 | 0.98 | 0 |
| **SUN3** | 0 | 1.33403492109535 | 0.72 | 0.031 | 0 |
| **UPP1** | 0 | 1.42425506988788 | 1 | 0.952 | 0 |
| **CHCHD2** | 0 | 1.74526367703816 | 1 | 0.997 | 0 |
| **ZNF727** | 0 | 1.59838211780324 | 0.794 | 0.006 | 0 |
| **ZNF736** | 0 | 1.33653980601432 | 0.775 | 0.008 | 0 |
| **ZNF680** | 0 | 1.01424442082834 | 0.975 | 0.783 | 0 |
| **ZNF273** | 0 | 0.967365682867635 | 0.637 | 0.055 | 0 |
| **AUTS2** | 0 | -1.73666858174359 | 0.599 | 0.94 | 0 |
| **CDK6** | 0 | -2.45470423238555 | 0.12 | 0.943 | 0 |
| **TFPI2** | 0 | 1.34743600424165 | 0.832 | 0.312 | 0 |
| **BAIAP2L1** | 0 | 0.967161974721358 | 0.995 | 0.888 | 0 |
| **ARPC1B** | 0 | 1.62506057411362 | 1 | 0.982 | 0 |
| **STAG3** | 0 | 1.5320218286319 | 0.737 | 0.019 | 0 |
| **TRIP6** | 0 | -0.881204715437669 | 1 | 1 | 0 |
| **COL26A1** | 0 | -2.93501631006655 | 0.198 | 0.97 | 0 |
| **CUX1** | 0 | -1.17521146852526 | 0.692 | 0.962 | 0 |
| **PRKAR2B** | 0 | -1.67807956867324 | 0.45 | 0.917 | 0 |
| **LAMB1** | 0 | -1.65358845900501 | 0.806 | 0.996 | 0 |
| **LRRN3** | 0 | -1.49781769439421 | 0.013 | 0.641 | 0 |
| **CAV1** | 0 | -1.70093167687232 | 0.038 | 0.587 | 0 |

| | | | | | |
|---|---|---|---|---|---|
| **ST7** | 0 | -1.03974980192611 | 0.129 | 0.659 | 0 |
| **PTPRZ1** | 0 | -1.64013120856791 | 0.006 | 0.557 | 0 |
| **AASS** | 0 | 1.21166545764437 | 0.998 | 0.953 | 0 |
| **NDUFA5** | 0 | 1.3791517252329 | 0.975 | 0.586 | 0 |
| **HYAL4** | 0 | 2.15452151420336 | 0.95 | 0.129 | 0 |
| **KLHDC10** | 0 | -1.26893877802404 | 0.147 | 0.774 | 0 |
| **MEST** | 0 | 1.96259284147439 | 0.996 | 0.768 | 0 |
| **PODXL** | 0 | -4.50161273846194 | 0.699 | 0.997 | 0 |
| **AKR1B1** | 0 | -0.905128473210009 | 0.969 | 0.998 | 0 |
| **CALD1** | 0 | -1.71749264425835 | 0.996 | 1 | 0 |
| **TRIM24** | 0 | -1.94570076853608 | 0.972 | 1 | 0 |
| **KLRG2** | 0 | 0.891299028201223 | 1 | 0.926 | 0 |
| **TCAF1** | 0 | -2.03427449070435 | 0.325 | 0.954 | 0 |
| **ZNF398** | 0 | 1.01482741008158 | 0.986 | 0.834 | 0 |
| **REPIN1** | 0 | -1.56538225683155 | 0.479 | 0.955 | 0 |
| **ABCB8** | 0 | 0.964328322015975 | 0.999 | 0.963 | 0 |
| **UBE3C** | 0 | 0.810700021119765 | 1 | 0.983 | 0 |
| **PPP1R3B** | 0 | -1.541858884301 | 0.75 | 0.964 | 0 |
| **TUSC3** | 0 | -0.926758785583519 | 0.981 | 1 | 0 |
| **ZDHHC2** | 0 | -1.10632998574802 | 0.05 | 0.601 | 0 |
| **LOXL2** | 0 | -1.93320278763001 | 0.531 | 0.975 | 0 |
| **DPYSL2** | 0 | -2.94314179659485 | 0.551 | 0.999 | 0 |
| **CLU** | 0 | -2.22234696118008 | 0.908 | 1 | 0 |
| **RBPMS** | 0 | -0.999518019517401 | 0.986 | 0.999 | 0 |
| **GSR** | 0 | -1.52406198878489 | 0.762 | 0.993 | 0 |
| **NRG1** | 0 | 1.64591820923726 | 0.821 | 0.145 | 0 |
| **UNC5D** | 0 | -0.92283637792231 | 0.02 | 0.527 | 0 |
| **EIF4EBP1** | 0 | 1.17668545177774 | 1 | 0.996 | 0 |
| **ASH2L** | 0 | 1.07067946451682 | 1 | 0.99 | 0 |
| **FGFR1** | 0 | -3.11961095128212 | 0.598 | 0.996 | 0 |
| **TGS1** | 0 | 0.858211992499768 | 1 | 0.987 | 0 |
| **NSMAF** | 0 | 1.14564517590075 | 0.999 | 0.937 | 0 |
| **CHD7** | 0 | -2.46644741771141 | 0.443 | 0.958 | 0 |
| **MTFR1** | 0 | 1.30184326926608 | 0.996 | 0.868 | 0 |

| | | | | | |
|---|---|---|---|---|---|
| **ARFGEF1** | 0 | 0.893876296443159 | 0.999 | 0.969 | 0 |
| **PREX2** | 0 | -1.72496738015507 | 0.112 | 0.778 | 0 |
| **PRDM14** | 0 | 1.34691008194851 | 0.997 | 0.889 | 0 |
| **CRISPLD1** | 0 | -1.57296665465451 | 0.562 | 0.933 | 0 |
| **PKIA** | 0 | -0.928442517847572 | 0.026 | 0.544 | 0 |
| **FABP5** | 0 | 3.65313800007057 | 0.997 | 0.552 | 0 |
| **TMEM64** | 0 | -3.55494050448211 | 0.611 | 0.995 | 0 |
| **GRHL2** | 0 | 1.09538480255827 | 0.561 | 0.008 | 0 |
| **CTHRC1** | 0 | 1.83826225569859 | 0.876 | 0.07 | 0 |
| **EIF3E** | 0 | 1.02684159111732 | 0.739 | 0.156 | 0 |
| **DEPTOR** | 0 | 1.3622977397781 | 0.997 | 0.802 | 0 |
| **ZHX2** | 0 | -1.85101219718938 | 0.072 | 0.828 | 0 |
| **LRATD2** | 0 | -1.79567291286355 | 0.819 | 0.981 | 0 |
| **MYC** | 0 | -1.69797613876867 | 0.705 | 0.966 | 0 |
| **TOP1MT** | 0 | 1.1750845705107 | 0.998 | 0.954 | 0 |
| **MAFA** | 0 | 1.34393530565604 | 0.832 | 0.265 | 0 |
| **SMARCA2** | 0 | 1.21124420414227 | 0.998 | 0.879 | 0 |
| **ERMP1** | 0 | -1.17464122162302 | 0.447 | 0.859 | 0 |
| **GLDC** | 0 | 2.04025986391581 | 1 | 0.809 | 0 |
| **KDM4C** | 0 | 1.20890906333241 | 0.986 | 0.853 | 0 |
| **PSIP1** | 0 | -1.01617140766622 | 0.954 | 0.992 | 0 |
| **DNAJA1** | 0 | 1.23578880008129 | 0.997 | 0.89 | 0 |
| **UBAP2** | 0 | 0.965538137856459 | 1 | 0.977 | 0 |
| **STOML2** | 0 | 0.810271441545827 | 1 | 0.999 | 0 |
| **TPM2** | 0 | -1.60341458319806 | 0.979 | 0.999 | 0 |
| **CLTA** | 0 | -0.864845503124753 | 1 | 1 | 0 |
| **IGFBPL1** | 0 | -3.5351120904965 | 0.173 | 0.981 | 0 |
| **ANKRD18A** | 0 | -1.73643244565192 | 0.051 | 0.769 | 0 |
| **PCSK5** | 0 | 1.55778848901288 | 0.928 | 0.421 | 0 |
| **GCNT1** | 0 | 1.80839303460774 | 0.957 | 0.544 | 0 |
| **UBQLN1** | 0 | 0.837340414433463 | 0.999 | 0.959 | 0 |
| **C9orf64** | 0 | 1.31036833727491 | 0.736 | 0.017 | 0 |
| **HNRNPK** | 0 | 0.970989350221994 | 1 | 0.942 | 0 |
| **BARX1** | 0 | 1.67669895233153 | 0.766 | 0.019 | 0 |

| | | | | | |
|---|---|---|---|---|---|
| **PTCH1** | 0 | -1.70694372043866 | 0.38 | 0.882 | 0 |
| **TGFBR1** | 0 | -1.22678380097647 | 0.964 | 0.999 | 0 |
| **SEC61B** | 0 | 1.08321073283029 | 0.999 | 0.921 | 0 |
| **ABCA1** | 0 | 1.17731823669309 | 1 | 0.946 | 0 |
| **KLF4** | 0 | 2.74963157694861 | 1 | 0.685 | 0 |
| **EPB41L4B** | 0 | 1.3508734079983 | 0.965 | 0.643 | 0 |
| **PALM2-AKAP2** | 0 | -3.43798518179583 | 0.269 | 0.997 | 0 |
| **PTGR1** | 0 | 1.10953307732774 | 0.991 | 0.858 | 0 |
| **ALAD** | 0 | 1.06802521092879 | 0.984 | 0.807 | 0 |
| **CNTRL** | 0 | -1.30092817161508 | 0.263 | 0.801 | 0 |
| **GSN** | 0 | -2.54883271661683 | 0.027 | 0.831 | 0 |
| **AIF1L** | 0 | -1.31513352778421 | 0.981 | 0.999 | 0 |
| **VAV2** | 0 | 0.999380414329069 | 0.989 | 0.918 | 0 |
| **WDR5** | 0 | 1.08156187162864 | 0.991 | 0.865 | 0 |
| **AGPAT2** | 0 | 1.07411591288753 | 0.995 | 0.899 | 0 |
| **GDI2** | 0 | -1.11627292455156 | 0.988 | 1 | 0 |
| **ECHDC3** | 0 | 1.47750860898209 | 0.788 | 0.012 | 0 |
| **CAMK1D** | 0 | 1.0796113320493 | 0.672 | 0.109 | 0 |
| **FRMD4A** | 0 | -1.48681531180482 | 0.081 | 0.61 | 0 |
| **OLAH** | 0 | 2.89405646467782 | 0.976 | 0.28 | 0 |
| **VIM** | 0 | -5.41584532078574 | 0.391 | 0.981 | 0 |
| **NEBL** | 0 | -1.44271737499739 | 0.067 | 0.59 | 0 |
| **ITGB1** | 0 | -1.47045522669511 | 0.959 | 1 | 0 |
| **NCOA4** | 0 | 0.861928277127961 | 0.997 | 0.95 | 0 |
| **CCDC6** | 0 | -1.28318374272322 | 0.927 | 0.998 | 0 |
| **JMJD1C** | 0 | -1.33638401798122 | 0.794 | 0.992 | 0 |
| **REEP3** | 0 | -1.13454622609465 | 0.577 | 0.925 | 0 |
| **HNRNPH3** | 0 | -0.871427401298317 | 1 | 1 | 0 |
| **SLC25A16** | 0 | 1.61830663095363 | 1 | 0.916 | 0 |
| **HK1** | 0 | -1.4909058696789 | 0.857 | 0.999 | 0 |
| **TSPAN15** | 0 | -1.04551756381303 | 0.093 | 0.634 | 0 |
| **H2AFY2** | 0 | -2.34516866462653 | 0.522 | 0.97 | 0 |
| **PSAP** | 0 | -1.27182558529713 | 1 | 1 | 0 |
| **VCL** | 0 | -1.19803585556921 | 0.998 | 1 | 0 |

| | | | | | |
|---|---|---|---|---|---|
| **ZCCHC24** | 0 | -1.39648885611167 | 0.171 | 0.718 | 0 |
| **PRXL2A** | 0 | -1.41924194344679 | 0.939 | 0.999 | 0 |
| **LIPA** | 0 | -1.39380036117036 | 0.27 | 0.831 | 0 |
| **FGFBP3** | 0 | -1.92362365206674 | 0.013 | 0.54 | 0 |
| **PDLIM1** | 0 | 1.34828461103225 | 1 | 0.963 | 0 |
| **PIK3AP1** | 0 | 1.07483291108253 | 0.643 | 0.048 | 0 |
| **SCD** | 0 | -1.07898981059086 | 1 | 1 | 0 |
| **NFKB2** | 0 | 1.78138550044463 | 1 | 0.897 | 0 |
| **C10orf95** | 0 | 1.62359996280262 | 0.877 | 0.287 | 0 |
| **NT5C2** | 0 | -1.31077944213995 | 0.816 | 0.987 | 0 |
| **INA** | 0 | 1.07527741412626 | 0.976 | 0.771 | 0 |
| **PCGF6** | 0 | 1.32488319261672 | 0.96 | 0.568 | 0 |
| **ATP5MD** | 0 | 1.77778137537556 | 0.968 | 0.405 | 0 |
| **GSTO2** | 0 | 1.03576145823818 | 0.672 | 0.072 | 0 |
| **XPNPEP1** | 0 | -1.06059635246574 | 0.715 | 0.961 | 0 |
| **ADD3** | 0 | -1.23166352175906 | 0.939 | 0.999 | 0 |
| **SMC3** | 0 | -1.12928717454445 | 0.964 | 0.998 | 0 |
| **TCF7L2** | 0 | -1.59922182415163 | 0.456 | 0.915 | 0 |
| **SHTN1** | 0 | -1.2929635326216 | 0.357 | 0.848 | 0 |
| **FGFR2** | 0 | -1.11714745372527 | 0.968 | 0.995 | 0 |
| **NKX1-2** | 0 | -2.37649612081062 | 0.565 | 0.876 | 0 |
| **UTF1** | 0 | 3.00811693520393 | 0.998 | 0.429 | 0 |
| **VENTX** | 0 | 1.27637153141113 | 0.918 | 0.395 | 0 |
| **PRAP1** | 0 | 2.19888757421334 | 0.892 | 0.188 | 0 |
| **IFITM3** | 0 | -1.71372858477473 | 0.985 | 0.999 | 0 |
| **PKP3** | 0 | 1.31567843887354 | 0.982 | 0.65 | 0 |
| **SIGIRR** | 0 | 0.990766888097871 | 0.984 | 0.751 | 0 |
| **TSPAN4** | 0 | -1.56294412585816 | 0.91 | 0.994 | 0 |
| **CDKN1C** | 0 | -1.23771321223874 | 0.726 | 0.97 | 0 |
| **TRIM21** | 0 | -0.86335323225548 | 0.046 | 0.555 | 0 |
| **TRIM6** | 0 | 1.19378247415271 | 0.804 | 0.203 | 0 |
| **TRIM5** | 0 | -1.45553516179638 | 0.294 | 0.867 | 0 |
| **TRIM22** | 0 | -1.77798199278711 | 0.296 | 0.846 | 0 |
| **TPP1** | 0 | -1.21545247111018 | 0.928 | 0.993 | 0 |

| | | | | | |
|---|---|---|---|---|---|
| **ST5** | 0 | -1.16504488595203 | 0.659 | 0.958 | 0 |
| **ADM** | 0 | -2.46571823397394 | 0.116 | 0.751 | 0 |
| **3,00 DKK** | 0 | -1.81149646026221 | 0.338 | 0.874 | 0 |
| **RRAS2** | 0 | 0.859787374045633 | 0.999 | 0.929 | 0 |
| **PDE3B** | 0 | -1.03004854952245 | 0.037 | 0.549 | 0 |
| **RCN1** | 0 | 1.11918210789822 | 0.999 | 0.96 | 0 |
| **C11orf96** | 0 | -3.09431310098521 | 0.182 | 0.932 | 0 |
| **TSPAN18** | 0 | -1.48731283557358 | 0.94 | 0.997 | 0 |
| **TP53I11** | 0 | -1.7851493249805 | 0.131 | 0.822 | 0 |
| **ATG13** | 0 | 0.885047524336781 | 1 | 0.992 | 0 |
| **SERPING1** | 0 | -2.36270353103395 | 0.363 | 0.979 | 0 |
| **TMEM258** | 0 | 1.23854101289153 | 0.999 | 0.869 | 0 |
| **FADS2** | 0 | -1.98954766334347 | 0.999 | 1 | 0 |
| **FADS1** | 0 | -1.4294163698 | 0.941 | 0.999 | 0 |
| **ASRGL1** | 0 | 0.975348060429643 | 1 | 0.954 | 0 |
| **EEF1G** | 0 | 1.2694972457609 | 1 | 0.999 | 0 |
| **MTA2** | 0 | -1.03735498079265 | 0.946 | 0.998 | 0 |
| **PLAAT5** | 0 | -1.82097431990934 | 0.389 | 0.846 | 0 |
| **RCOR2** | 0 | -1.71560632787831 | 0.314 | 0.86 | 0 |
| **RASGRP2** | 0 | 1.58870000355432 | 0.978 | 0.57 | 0 |
| **TM7SF2** | 0 | -1.32530877486863 | 0.855 | 0.991 | 0 |
| **GSTP1** | 0 | -0.989650297803634 | 1 | 1 | 0 |
| **NDUFV1** | 0 | 1.05558243050076 | 1 | 1 | 0 |
| **GAL** | 0 | 1.48846941229522 | 0.99 | 0.702 | 0 |
| **CCND1** | 0 | -1.8415758407785 | 0.953 | 0.997 | 0 |
| **FGF4** | 0 | 2.14435082741998 | 0.988 | 0.652 | 0 |
| **RAB6A** | 0 | 1.20369187753928 | 1 | 0.989 | 0 |
| **UCP2** | 0 | 1.3103043439231 | 0.996 | 0.912 | 0 |
| **CTSC** | 0 | 0.867312006344267 | 1 | 0.987 | 0 |
| **AMOTL1** | 0 | -2.23872494699861 | 0.263 | 0.91 | 0 |
| **SESN3** | 0 | -1.617216420889 | 0.87 | 0.996 | 0 |
| **TRPC6** | 0 | -2.71306946714128 | 0.208 | 0.87 | 0 |
| **YAP1** | 0 | -0.879798876913023 | 0.998 | 1 | 0 |
| **DCUN1D5** | 0 | 1.21215103318521 | 1 | 0.922 | 0 |

| | | | | | |
|---|---|---|---|---|---|
| **EXPH5** | 0 | 1.55696896993477 | 0.87 | 0.195 | 0 |
| **PPP2R1B** | 0 | -1.62715371178055 | 0.812 | 0.997 | 0 |
| **DIXDC1** | 0 | -1.27089862749965 | 0.227 | 0.75 | 0 |
| **NCAM1** | 0 | -1.24056226542709 | 0.007 | 0.505 | 0 |
| **USP28** | 0 | 1.42118192708443 | 1 | 0.98 | 0 |
| **TAGLN** | 0 | -4.90199975887034 | 0.366 | 0.954 | 0 |
| **HYOU1** | 0 | 0.881008870344767 | 1 | 1 | 0 |
| **UBASH3B** | 0 | 1.34102637556595 | 0.82 | 0.243 | 0 |
| **FAM118B** | 0 | -1.32828717578133 | 0.583 | 0.962 | 0 |
| **SNX19** | 0 | 1.28157019291436 | 0.748 | 0.047 | 0 |
| **TEAD4** | 0 | 1.378858488669 | 1 | 0.965 | 0 |
| **CCND2** | 0 | -2.4312567818713 | 0.124 | 0.703 | 0 |
| **GDF3** | 0 | 1.7498657333233 | 0.997 | 0.804 | 0 |
| **DPPA3** | 0 | 3.66609730571099 | 0.996 | 0.309 | 0 |
| **CLEC4D** | 0 | -1.94822827901666 | 0.185 | 0.692 | 0 |
| **DUSP16** | 0 | 1.67628628147484 | 0.994 | 0.87 | 0 |
| **CREBL2** | 0 | 1.21140968001811 | 0.995 | 0.894 | 0 |
| **PLBD1** | 0 | 1.46136363099075 | 1 | 0.917 | 0 |
| **MGST1** | 0 | -1.35997079394655 | 0.91 | 0.998 | 0 |
| **ETNK1** | 0 | -1.16585401086798 | 0.765 | 0.98 | 0 |
| **PPFIBP1** | 0 | -1.24608541753949 | 0.284 | 0.781 | 0 |
| **YAF2** | 0 | -1.46187205645254 | 0.123 | 0.757 | 0 |
| **TWF1** | 0 | 1.09222380711772 | 0.877 | 0.353 | 0 |
| **SLC38A1** | 0 | -1.23733400072367 | 0.695 | 0.982 | 0 |
| **SLC38A2** | 0 | -1.08492102079389 | 0.945 | 0.998 | 0 |
| **PFKM** | 0 | -1.47625790826473 | 0.681 | 0.98 | 0 |
| **ADCY6** | 0 | -1.10814250951468 | 0.042 | 0.549 | 0 |
| **TUBA1C** | 0 | 1.50896165136016 | 1 | 0.997 | 0 |
| **LIMA1** | 0 | 1.29452471302964 | 0.994 | 0.848 | 0 |
| **FIGNL2** | 0 | -2.03224878361533 | 0.109 | 0.743 | 0 |
| **KRT8** | 0 | -2.02591586819905 | 0.986 | 1 | 0 |
| **CBX5** | 0 | -1.46546938047845 | 0.875 | 0.992 | 0 |
| **ZNF385A** | 0 | -1.28345578661718 | 0.053 | 0.639 | 0 |
| **ITGA5** | 0 | -1.48937640049055 | 0.024 | 0.545 | 0 |

| | | | | | |
|---|---|---|---|---|---|
| **ESYT1** | 0 | -1.31701063213065 | 0.959 | 0.999 | 0 |
| **MYL6** | 0 | -0.994680269160892 | 1 | 1 | 0 |
| **SHMT2** | 0 | 1.17751620262637 | 0.977 | 0.745 | 0 |
| **HMGA2** | 0 | -2.50720942203021 | 0.532 | 0.993 | 0 |
| **LLPH** | 0 | 1.00261710803973 | 0.877 | 0.402 | 0 |
| **MDM2** | 0 | 1.32550173707156 | 1 | 0.975 | 0 |
| **YEATS4** | 0 | 1.04728360104325 | 0.985 | 0.841 | 0 |
| **MYRFL** | 0 | -1.20062003895316 | 0.008 | 0.538 | 0 |
| **ATXN7L3B** | 0 | -0.993180049492861 | 0.968 | 0.999 | 0 |
| **PHLDA1** | 0 | -2.88972910475875 | 0.348 | 0.923 | 0 |
| **CCDC59** | 0 | 1.06024567683844 | 0.986 | 0.789 | 0 |
| **UBE2N** | 0 | 1.21608021905004 | 0.927 | 0.421 | 0 |
| **NDUFA12** | 0 | 1.10856080877736 | 1 | 0.94 | 0 |
| **LTA4H** | 0 | -0.819458588835815 | 0.995 | 0.999 | 0 |
| **TMPO** | 0 | 1.15094339788903 | 1 | 0.99 | 0 |
| **ACTR6** | 0 | 1.51765808411954 | 0.965 | 0.468 | 0 |
| **SPIC** | 0 | 2.54276154600547 | 0.909 | 0.242 | 0 |
| **CHPT1** | 0 | 0.901327115742634 | 0.997 | 0.944 | 0 |
| **SYCP3** | 0 | 1.69525794151872 | 0.681 | 0.1 | 0 |
| **HSP90B1** | 0 | 2.53293137480331 | 0.992 | 0.772 | 0 |
| **CHST11** | 0 | -1.87386503192499 | 0.674 | 0.963 | 0 |
| **ALDH1L2** | 0 | 1.39422445122182 | 0.787 | 0.139 | 0 |
| **C12orf75** | 0 | 1.32820193922135 | 0.991 | 0.919 | 0 |
| **CKAP4** | 0 | -1.03894923252361 | 0.986 | 1 | 0 |
| **PPTC7** | 0 | 1.0736586921559 | 0.982 | 0.809 | 0 |
| **ALDH2** | 0 | 1.5256522542005 | 0.999 | 0.942 | 0 |
| **OAS1** | 0 | 1.11769352116736 | 0.603 | 0.023 | 0 |
| **SLC8B1** | 0 | 1.21726773843151 | 0.942 | 0.57 | 0 |
| **TBX3** | 0 | 2.13058821650049 | 0.905 | 0.139 | 0 |
| **VSIG10** | 0 | 1.30828996661752 | 1 | 0.932 | 0 |
| **TAOK3** | 0 | 1.18823616280397 | 0.836 | 0.337 | 0 |
| **HSPB8** | 0 | -1.44721337475526 | 0.024 | 0.572 | 0 |
| **MSI1** | 0 | -1.74990669853596 | 0.213 | 0.846 | 0 |
| **ORAI1** | 0 | 0.942810575116758 | 0.998 | 0.936 | 0 |

| | | | | | |
|---|---|---|---|---|---|
| **RHOF** | 0 | 1.34591056990965 | 0.951 | 0.449 | 0 |
| **CLIP1** | 0 | 0.979647112119911 | 0.999 | 0.932 | 0 |
| **DENR** | 0 | 1.17265510838686 | 1 | 0.975 | 0 |
| **VPS37B** | 0 | 1.45620643504977 | 0.999 | 0.942 | 0 |
| **PUS1** | 0 | 1.06464561446268 | 0.999 | 0.935 | 0 |
| **FBRSL1** | 0 | 0.889493170315196 | 0.995 | 0.901 | 0 |
| **ZMYM2** | 0 | -0.940752062197075 | 0.999 | 1 | 0 |
| **SACS** | 0 | -1.49248882983291 | 0.821 | 0.991 | 0 |
| **WASF3** | 0 | -1.49756464614632 | 0.669 | 0.963 | 0 |
| **FLT1** | 0 | -1.78757106559786 | 0.058 | 0.599 | 0 |
| **PDS5B** | 0 | -1.02559693221655 | 0.872 | 0.993 | 0 |
| **NBEA** | 0 | -2.05096596043814 | 0.025 | 0.804 | 0 |
| **DCLK1** | 0 | -2.9200154373433 | 0.141 | 0.851 | 0 |
| **UFM1** | 0 | 1.17224681428743 | 1 | 0.948 | 0 |
| **LHFPL6** | 0 | -1.90751531914164 | 0.078 | 0.769 | 0 |
| **FOXO1** | 0 | -2.27310423188179 | 0.621 | 0.982 | 0 |
| **RGCC** | 0 | 1.99880507271338 | 0.887 | 0.236 | 0 |
| **DNAJC15** | 0 | 1.92504460042674 | 0.996 | 0.992 | 0 |
| **LCP1** | 0 | 2.16295029280334 | 0.99 | 0.645 | 0 |
| **ESD** | 0 | -0.939173588004838 | 0.985 | 1 | 0 |
| **ITM2B** | 0 | -1.17232956043755 | 0.985 | 0.998 | 0 |
| **RCBTB1** | 0 | 1.58684607959605 | 0.999 | 0.954 | 0 |
| **WDFY2** | 0 | -1.32821250004369 | 0.703 | 0.962 | 0 |
| **KLF5** | 0 | 2.1460199934226 | 1 | 0.799 | 0 |
| **KLF12** | 0 | -1.34269367884473 | 0.167 | 0.762 | 0 |
| **TBC1D4** | 0 | -1.2896530450974 | 0.234 | 0.739 | 0 |
| **EDNRB** | 0 | -1.39690898590468 | 0.003 | 0.563 | 0 |
| **SPRY2** | 0 | -2.4821808295968 | 0.688 | 0.992 | 0 |
| **GPC6** | 0 | -2.42825076657354 | 0.288 | 0.964 | 0 |
| **DZIP1** | 0 | -1.40182171036675 | 0.715 | 0.976 | 0 |
| **IPO5** | 0 | -0.836605281698692 | 0.998 | 1 | 0 |
| **FARP1** | 0 | -1.3223211121839 | 0.674 | 0.949 | 0 |
| **ZIC5** | 0 | -2.51151328453772 | 0.013 | 0.592 | 0 |
| **ZIC2** | 0 | -2.62519853005677 | 0.018 | 0.689 | 0 |

| | | | | | |
|---|---|---|---|---|---|
| **PCCA** | 0 | -1.75891859126247 | 0.887 | 0.998 | 0 |
| **EFNB2** | 0 | -2.30421792748594 | 0.448 | 0.941 | 0 |
| **ARGLU1** | 0 | -1.10583241890712 | 0.994 | 0.999 | 0 |
| **COL4A1** | 0 | -2.72666897361701 | 0.608 | 0.992 | 0 |
| **COL4A2** | 0 | -2.87454265865691 | 0.624 | 0.996 | 0 |
| **ANKRD10** | 0 | -1.67418665377657 | 0.917 | 0.993 | 0 |
| **LAMP1** | 0 | -1.43625926156539 | 1 | 1 | 0 |
| **ARHGEF40** | 0 | -1.73979918665643 | 0.088 | 0.72 | 0 |
| **ZNF219** | 0 | -2.03107421488232 | 0.548 | 0.96 | 0 |
| **SALL2** | 0 | -2.34478903195392 | 0.741 | 0.998 | 0 |
| **SLC7A7** | 0 | 2.58126879204424 | 0.994 | 0.477 | 0 |
| **AJUBA** | 0 | -1.79821771585452 | 0.651 | 0.959 | 0 |
| **PSME1** | 0 | -1.11266844616875 | 0.792 | 0.965 | 0 |
| **REC8** | 0 | 3.33676959672338 | 0.997 | 0.337 | 0 |
| **EGLN3** | 0 | 1.19561318977892 | 0.989 | 0.785 | 0 |
| **SPTSSA** | 0 | 1.47080026456241 | 1 | 1 | 0 |
| **NFKBIA** | 0 | 1.09255447319863 | 1 | 0.956 | 0 |
| **MIPOL1** | 0 | -1.22247930179054 | 0.271 | 0.803 | 0 |
| **GEMIN2** | 0 | 0.996977576776471 | 0.892 | 0.454 | 0 |
| **ARF6** | 0 | 1.01483009874238 | 1 | 0.992 | 0 |
| **PYGL** | 0 | 1.91814165107553 | 1 | 0.928 | 0 |
| **FRMD6** | 0 | 1.31079286187373 | 0.955 | 0.705 | 0 |
| **GNPNAT1** | 0 | 1.57831276932414 | 0.998 | 0.884 | 0 |
| **GMFB** | 0 | 1.25562974740048 | 0.794 | 0.1 | 0 |
| **GCH1** | 0 | 1.25036557806499 | 0.937 | 0.52 | 0 |
| **WDHD1** | 0 | 1.03299973729041 | 1 | 0.99 | 0 |
| **TBPL2** | 0 | 0.932817951309448 | 0.562 | 0.008 | 0 |
| **TMEM30B** | 0 | 1.50198001213462 | 0.771 | 0.022 | 0 |
| **HSPA2** | 0 | 2.0818042079223 | 0.904 | 0.164 | 0 |
| **PLEKHG3** | 0 | -1.735467890178 | 0.097 | 0.623 | 0 |
| **GPX2** | 0 | 2.73313483208855 | 0.963 | 0.301 | 0 |
| **RAB15** | 0 | 1.25961417410393 | 1 | 0.998 | 0 |
| **ACTN1** | 0 | -1.90879324554106 | 0.985 | 1 | 0 |
| **DCAF5** | 0 | -1.3563932435356 | 0.571 | 0.949 | 0 |

| | | | | | |
|---|---|---|---|---|---|
| **DCAF4** | 0 | 1.50477359763608 | 1 | 0.912 | 0 |
| **ACOT4** | 0 | 1.20061325962958 | 0.683 | 0.014 | 0 |
| **VRTN** | 0 | -3.53271186186673 | 0.244 | 0.961 | 0 |
| **NPC2** | 0 | -1.44700581726566 | 1 | 1 | 0 |
| **SLIRP** | 0 | 1.30223005008475 | 1 | 0.999 | 0 |
| **SNW1** | 0 | 1.20355171477131 | 0.988 | 0.763 | 0 |
| **CALM1** | 0 | 1.14009055590335 | 0.971 | 0.691 | 0 |
| **CCDC85C** | 0 | -1.4464922984729 | 0.985 | 0.997 | 0 |
| **WARS** | 0 | 2.13263669975725 | 1 | 0.998 | 0 |
| **CKB** | 0 | -0.807711603368507 | 1 | 1 | 0 |
| **ATP5MPL** | 0 | 1.28494071816498 | 0.981 | 0.663 | 0 |
| **JAG2** | 0 | 1.24074072325774 | 0.806 | 0.241 | 0 |
| **MTA1** | 0 | -0.85639711925712 | 0.99 | 1 | 0 |
| **CRIP1** | 0 | 4.50276600714852 | 1 | 0.727 | 0 |
| **NIPA2** | 0 | 0.990667756456713 | 0.998 | 0.96 | 0 |
| **SNRPN** | 0 | 1.47129895899166 | 0.997 | 0.837 | 0 |
| **LPCAT4** | 0 | 1.50715208662055 | 0.995 | 0.813 | 0 |
| **SPRED1** | 0 | -1.35607885305444 | 0.096 | 0.755 | 0 |
| **SPINT1** | 0 | 0.938296586537342 | 1 | 0.853 | 0 |
| **TMEM87A** | 0 | 1.00799865634143 | 0.955 | 0.638 | 0 |
| **PDIA3** | 0 | 1.23016456587977 | 1 | 0.989 | 0 |
| **B2M** | 0 | -1.67378638776143 | 0.476 | 0.918 | 0 |
| **SORD** | 0 | -1.15398038934438 | 0.493 | 0.876 | 0 |
| **MYEF2** | 0 | -1.86012346727789 | 0.472 | 0.968 | 0 |
| **EID1** | 0 | -1.74691537281719 | 0.952 | 1 | 0 |
| **ATP8B4** | 0 | 1.78093475670415 | 0.787 | 0.07 | 0 |
| **SCG3** | 0 | -1.39733973326661 | 0.081 | 0.684 | 0 |
| **MYO5A** | 0 | 1.05485840518704 | 0.987 | 0.846 | 0 |
| **RSL24D1** | 0 | 1.26683889623654 | 0.805 | 0.103 | 0 |
| **ADAM10** | 0 | 0.964720702699359 | 0.992 | 0.861 | 0 |
| **GTF2A2** | 0 | 1.46611917513098 | 1 | 0.918 | 0 |
| **TPM1** | 0 | -1.37846602020696 | 1 | 1 | 0 |
| **SNX1** | 0 | -1.27673607480874 | 0.756 | 0.979 | 0 |
| **PPIB** | 0 | 0.917230522719625 | 0.999 | 0.996 | 0 |

| | | | | | |
|---|---|---|---|---|---|
| **PCLAF** | 0 | -2.38305709369658 | 0.014 | 0.81 | 0 |
| **OAZ2** | 0 | -2.54838537216493 | 0.812 | 0.999 | 0 |
| **CLPX** | 0 | 1.16962515017004 | 1 | 0.987 | 0 |
| **IGDCC3** | 0 | -1.91956828849249 | 0.252 | 0.889 | 0 |
| **IGDCC4** | 0 | -1.3833551425426 | 0.021 | 0.664 | 0 |
| **SMAD3** | 0 | -1.52333627454063 | 0.589 | 0.953 | 0 |
| **UACA** | 0 | 1.50268117103632 | 0.996 | 0.895 | 0 |
| **MYO9A** | 0 | 1.03851794559568 | 0.931 | 0.557 | 0 |
| **CD276** | 0 | -1.23652173592278 | 0.968 | 0.999 | 0 |
| **ETFA** | 0 | -1.22439057443447 | 0.73 | 0.971 | 0 |
| **PEAK1** | 0 | -1.69739729598682 | 0.21 | 0.855 | 0 |
| **DNAJA4** | 0 | 1.7188428341743 | 0.853 | 0.042 | 0 |
| **MEX3B** | 0 | -1.28570978548815 | 0.056 | 0.588 | 0 |
| **RAMAC** | 0 | 1.11473787768183 | 0.959 | 0.605 | 0 |
| **SEC11A** | 0 | -0.910686641220795 | 0.999 | 1 | 0 |
| **MFGE8** | 0 | -2.07900129490615 | 0.358 | 0.945 | 0 |
| **FES** | 0 | 1.13398685116905 | 0.877 | 0.325 | 0 |
| **FAM174B** | 0 | -1.54446112775898 | 0.288 | 0.793 | 0 |
| **IGF1R** | 0 | -1.52948899012802 | 0.915 | 0.999 | 0 |
| **CHSY1** | 0 | -1.5146422643206 | 0.915 | 0.998 | 0 |
| **HBM** | 0 | 1.06743056966403 | 0.613 | 0.02 | 0 |
| **HBA2** | 0 | 1.18662431930013 | 0.647 | 0.052 | 0 |
| **NME4** | 0 | -1.14451438829272 | 0.995 | 1 | 0 |
| **CACNA1H** | 0 | 1.6779085858141 | 0.851 | 0.057 | 0 |
| **SYNGR3** | 0 | 1.60978400384297 | 0.994 | 0.716 | 0 |
| **SLC9A3R2** | 0 | 1.94125840120051 | 0.996 | 0.641 | 0 |
| **PGP** | 0 | 0.938791849638 | 1 | 0.997 | 0 |
| **IL32** | 0 | 1.77689966236834 | 0.826 | 0.207 | 0 |
| **NMRAL1** | 0 | 1.69271237113843 | 0.997 | 0.75 | 0 |
| **HMOX2** | 0 | 0.873289539121084 | 0.998 | 0.959 | 0 |
| **C16orf72** | 0 | 0.800706746593605 | 1 | 0.996 | 0 |
| **LITAF** | 0 | -1.46763590975671 | 0.994 | 0.999 | 0 |
| **RSL1D1** | 0 | 0.912388819753831 | 1 | 0.951 | 0 |
| **GSPT1** | 0 | 1.1812789631472 | 0.999 | 0.898 | 0 |

| | | | | | |
|---|---|---|---|---|---|
| **SMG1** | 0 | 1.16774528990753 | 0.959 | 0.593 | 0 |
| **NDUFAB1** | 0 | 1.0848451534527 | 1 | 1 | 0 |
| **QPRT** | 0 | -1.01284293053291 | 0.997 | 1 | 0 |
| **SEPHS2** | 0 | 1.761405939137 | 1 | 0.984 | 0 |
| **GPT2** | 0 | 2.00818750112215 | 0.999 | 0.935 | 0 |
| **DNAJA2** | 0 | 0.9612833951147 | 1 | 1 | 0 |
| **NETO2** | 0 | 1.08841124387824 | 0.977 | 0.8 | 0 |
| **ZNF423** | 0 | -2.2377838796655 | 0.54 | 0.981 | 0 |
| **SALL1** | 0 | -2.54894355583826 | 0.031 | 0.803 | 0 |
| **TOX3** | 0 | -1.30502585803779 | 0.103 | 0.668 | 0 |
| **AC007906.2** | 0 | -1.20374417411236 | 0.028 | 0.582 | 0 |
| **MMP2** | 0 | -3.66838655505842 | 0.206 | 0.959 | 0 |
| **MT1E** | 0 | 2.59976444196351 | 0.982 | 0.312 | 0 |
| **MT1F** | 0 | 1.94448183670411 | 1 | 0.878 | 0 |
| **MT1G** | 0 | 3.96507657665334 | 0.998 | 0.377 | 0 |
| **MT1H** | 0 | 4.90960461845224 | 1 | 0.433 | 0 |
| **MT1X** | 0 | 2.71532776569981 | 1 | 0.962 | 0 |
| **CCDC102A** | 0 | -1.23475712618446 | 0.398 | 0.855 | 0 |
| **USB1** | 0 | -1.09168408848859 | 0.847 | 0.988 | 0 |
| **MMP15** | 0 | -1.44080792611805 | 0.869 | 0.996 | 0 |
| **CMTM3** | 0 | -1.70651712679024 | 0.857 | 0.991 | 0 |
| **CMTM4** | 0 | 1.10345327823421 | 0.999 | 0.956 | 0 |
| **RRAD** | 0 | 2.44116921745978 | 0.877 | 0.212 | 0 |
| **PLEKHG4** | 0 | 0.954178756841327 | 0.564 | 0.016 | 0 |
| **HAS3** | 0 | -1.22178383003345 | 0.029 | 0.581 | 0 |
| **ZFHX3** | 0 | 1.58148362848304 | 0.807 | 0.161 | 0 |
| **VAT1L** | 0 | -3.34885073646387 | 0.012 | 0.708 | 0 |
| **WWOX** | 0 | -1.79852455751592 | 0.606 | 0.958 | 0 |
| **MAF** | 0 | -2.56854345086682 | 0.148 | 0.837 | 0 |
| **COX4I1** | 0 | 0.853352987576678 | 1 | 0.993 | 0 |
| **PIEZO1** | 0 | 1.24767808013326 | 0.73 | 0.033 | 0 |
| **DBNDD1** | 0 | -1.614632229907 | 0.672 | 0.965 | 0 |
| **RFLNB** | 0 | 2.76012114667593 | 0.983 | 0.608 | 0 |
| **SHPK** | 0 | 1.51219257789343 | 0.994 | 0.885 | 0 |

| | | | | | |
|---|---|---|---|---|---|
| **ZFP3** | 0 | 1.24863987032002 | 0.661 | 0.023 | 0 |
| **C1QBP** | 0 | 0.935202874131928 | 1 | 0.993 | 0 |
| **TXNDC17** | 0 | 0.861241548716707 | 1 | 0.993 | 0 |
| **ATP1B2** | 0 | -1.59100745336191 | 0.133 | 0.712 | 0 |
| **TP53** | 0 | -1.36016945075752 | 0.992 | 0.999 | 0 |
| **TRAPPC1** | 0 | -0.887669301830106 | 0.987 | 1 | 0 |
| **NTN1** | 0 | 7.44140547537527 | 1 | 0.985 | 0 |
| **GAS7** | 0 | 1.21723886657032 | 1 | 0.881 | 0 |
| **TMEM97** | 0 | -0.959997265407561 | 0.996 | 0.999 | 0 |
| **RAB34** | 0 | -1.16070998049589 | 0.932 | 0.997 | 0 |
| **FLOT2** | 0 | -1.35794657928391 | 0.931 | 0.998 | 0 |
| **CPD** | 0 | -1.12414901596122 | 0.938 | 0.997 | 0 |
| **UTP6** | 0 | 0.825296311705286 | 0.999 | 0.956 | 0 |
| **TMEM98** | 0 | -1.5813736578376 | 0.359 | 0.876 | 0 |
| **AP2B1** | 0 | 1.38718177460941 | 0.999 | 0.89 | 0 |
| **EIF1** | 0 | 1.85403355815237 | 1 | 0.895 | 0 |
| **FKBP10** | 0 | -2.07171361740332 | 0.617 | 0.961 | 0 |
| **GRN** | 0 | -0.908447461022735 | 1 | 1 | 0 |
| **FAM171A2** | 0 | -1.26488947620006 | 0.707 | 0.952 | 0 |
| **FZD2** | 0 | -2.59009496051017 | 0.055 | 0.782 | 0 |
| **GJC1** | 0 | -1.73213134060427 | 0.581 | 0.891 | 0 |
| **HOXB2** | 0 | 1.70394936676423 | 0.926 | 0.434 | 0 |
| **IGF2BP1** | 0 | -1.59660939330498 | 0.951 | 1 | 0 |
| **LRRC59** | 0 | 0.954709418681214 | 1 | 1 | 0 |
| **TOM1L1** | 0 | -1.13366337019682 | 0.676 | 0.966 | 0 |
| **AKAP1** | 0 | 1.1094599011299 | 1 | 0.977 | 0 |
| **MSI2** | 0 | -1.71890513682899 | 0.391 | 0.946 | 0 |
| **RNF43** | 0 | -1.11244669781749 | 0.098 | 0.616 | 0 |
| **YPEL2** | 0 | 1.93538025763569 | 0.996 | 0.589 | 0 |
| **NOL11** | 0 | 1.52391578535597 | 0.955 | 0.416 | 0 |
| **GALK1** | 0 | 1.22768342492046 | 0.948 | 0.618 | 0 |
| **ACOX1** | 0 | 0.947444148861717 | 0.995 | 0.898 | 0 |
| **SEPTIN9** | 0 | -1.06912070433065 | 0.99 | 1 | 0 |
| **SYNGR2** | 0 | 0.954354650373071 | 1 | 0.992 | 0 |

| | | | | | |
|---|---|---|---|---|---|
| **SOCS3** | 0 | -2.01855585888652 | 0.4 | 0.95 | 0 |
| **TIMP2** | 0 | -1.6055226250442 | 0.784 | 0.977 | 0 |
| **LGALS3BP** | 0 | -2.7497280370769 | 0.191 | 0.965 | 0 |
| **RPTOR** | 0 | 1.05292665082774 | 0.979 | 0.843 | 0 |
| **MCRIP1** | 0 | -1.3836235411132 | 0.925 | 0.998 | 0 |
| **PYCR1** | 0 | 2.21074555310709 | 0.997 | 0.558 | 0 |
| **SLC16A3** | 0 | 2.03372420884357 | 0.937 | 0.481 | 0 |
| **TBCD** | 0 | 1.59762286137076 | 1 | 0.998 | 0 |
| **MYL12A** | 0 | 1.02403254554221 | 0.928 | 0.512 | 0 |
| **PPP4R1** | 0 | -0.893274860941891 | 0.984 | 1 | 0 |
| **PIEZO2** | 0 | -2.19545439055225 | 0.037 | 0.778 | 0 |
| **GNAL** | 0 | 1.32928578270752 | 0.927 | 0.504 | 0 |
| **IMPA2** | 0 | 1.54587364438571 | 0.999 | 0.951 | 0 |
| **CDH2** | 0 | -2.99434863413024 | 0.038 | 0.711 | 0 |
| **RNF125** | 0 | 2.42498949660897 | 0.999 | 0.773 | 0 |
| **GALNT1** | 0 | -1.1629796281883 | 0.902 | 0.996 | 0 |
| **TPGS2** | 0 | -1.1633284393519 | 0.965 | 0.999 | 0 |
| **HAUS1** | 0 | 1.12645960558022 | 0.967 | 0.676 | 0 |
| **C18orf32** | 0 | 1.38479183303062 | 0.949 | 0.43 | 0 |
| **ACAA2** | 0 | 0.883716791905129 | 1 | 0.997 | 0 |
| **MAPK4** | 0 | -1.26404417860574 | 0.028 | 0.573 | 0 |
| **TCF4** | 0 | -1.85946281630454 | 0.296 | 0.897 | 0 |
| **ATP8B1** | 0 | 1.44502767169787 | 0.724 | 0.03 | 0 |
| **SLC66A2** | 0 | 1.07151941981832 | 0.999 | 0.918 | 0 |
| **HSBP1L1** | 0 | 1.17267620188834 | 0.791 | 0.19 | 0 |
| **CFD** | 0 | 1.63961216712056 | 0.926 | 0.274 | 0 |
| **CNN2** | 0 | -1.83913795902001 | 0.995 | 1 | 0 |
| **GAMT** | 0 | 1.08970427577407 | 0.955 | 0.614 | 0 |
| **REEP6** | 0 | 1.94290309140406 | 0.992 | 0.627 | 0 |
| **ZNF57** | 0 | 1.68944988452293 | 0.859 | 0.032 | 0 |
| **MATK** | 0 | 1.4790559051909 | 0.847 | 0.137 | 0 |
| **TNFSF9** | 0 | 2.8366211428071 | 0.992 | 0.246 | 0 |
| **CD70** | 0 | 3.56557955882761 | 0.992 | 0.294 | 0 |
| **TRIP10** | 0 | 0.87694122059999 | 1 | 0.978 | 0 |

| | | | | | |
|---|---|---|---|---|---|
| **PRR36** | 0 | -1.69223703949387 | 0.268 | 0.842 | 0 |
| **CD320** | 0 | 1.38807126663934 | 1 | 0.991 | 0 |
| **RAB11B** | 0 | -0.963264983376181 | 0.997 | 1 | 0 |
| **ZNF560** | 0 | 1.16469420541265 | 0.709 | 0.006 | 0 |
| **PDE4A** | 0 | 1.35713819101779 | 0.907 | 0.395 | 0 |
| **AP1M2** | 0 | 1.8042951722452 | 0.977 | 0.305 | 0 |
| **DOCK6** | 0 | 0.997671036377877 | 0.982 | 0.841 | 0 |
| **ACP5** | 0 | 1.43352123057957 | 0.771 | 0.023 | 0 |
| **ZNF878** | 0 | 1.55522548962262 | 0.868 | 0.252 | 0 |
| **ZNF844** | 0 | 1.73472068618922 | 0.829 | 0.009 | 0 |
| **ZNF625** | 0 | 0.96125091003091 | 0.998 | 0.958 | 0 |
| **TNPO2** | 0 | -0.937676172570057 | 0.95 | 0.997 | 0 |
| **RTBDN** | 0 | 1.17179185402548 | 0.723 | 0.081 | 0 |
| **NOTCH3** | 0 | -1.54866170805126 | 0.899 | 0.996 | 0 |
| **AP1M1** | 0 | -1.02798254796568 | 0.815 | 0.98 | 0 |
| **BST2** | 0 | -4.0499542206977 | 0.106 | 0.934 | 0 |
| **SLC5A5** | 0 | 1.25511098317655 | 0.766 | 0.105 | 0 |
| **ARRDC2** | 0 | 1.59452464293791 | 0.939 | 0.34 | 0 |
| **PIK3R2** | 0 | -1.0240240007964 | 0.997 | 1 | 0 |
| **UPF1** | 0 | 0.862737005600554 | 1 | 0.997 | 0 |
| **DDX49** | 0 | 0.853723311127848 | 0.997 | 0.953 | 0 |
| **PBX4** | 0 | 1.37205743337431 | 0.848 | 0.19 | 0 |
| **ZNF253** | 0 | 1.49823106298367 | 0.807 | 0.015 | 0 |
| **ZNF90** | 0 | 1.58569343768875 | 0.995 | 0.734 | 0 |
| **ZNF486** | 0 | 2.28753251394292 | 0.995 | 0.469 | 0 |
| **ZNF626** | 0 | 1.26639467139908 | 0.744 | 0.023 | 0 |
| **ZNF257** | 0 | 1.22145515554888 | 0.68 | 0.003 | 0 |
| **ZNF676** | 0 | 2.90136924750419 | 0.97 | 0.233 | 0 |
| **ZNF729** | 0 | 2.57103052683707 | 0.802 | 0.049 | 0 |
| **ZNF98** | 0 | 1.48264593264102 | 0.766 | 0.01 | 0 |
| **ZNF492** | 0 | 2.03381285346122 | 0.917 | 0.052 | 0 |
| **ZNF728** | 0 | 1.10598191749981 | 0.634 | 0.009 | 0 |
| **CHST8** | 0 | 1.69931477828122 | 0.94 | 0.404 | 0 |
| **DMKN** | 0 | -1.98495655247226 | 0.405 | 0.976 | 0 |

| | | | | | |
|---|---|---|---|---|---|
| **ZNF146** | 0 | 0.994967424612814 | 1 | 0.999 | 0 |
| **ZNF568** | 0 | 2.00269231107566 | 0.967 | 0.187 | 0 |
| **ZNF607** | 0 | 1.22611858986975 | 0.822 | 0.231 | 0 |
| **PPP1R14A** | 0 | 1.54904909293572 | 0.997 | 0.778 | 0 |
| **YIF1B** | 0 | 0.811348215859645 | 1 | 0.999 | 0 |
| **C19orf33** | 0 | 2.42262800434028 | 0.979 | 0.355 | 0 |
| **KCNK6** | 0 | 1.23126250767411 | 0.793 | 0.162 | 0 |
| **MAP4K1** | 0 | 1.18703132494536 | 0.817 | 0.268 | 0 |
| **TIMM50** | 0 | 0.808020935409728 | 1 | 1 | 0 |
| **DLL3** | 0 | 1.09814571644224 | 0.862 | 0.265 | 0 |
| **PLD3** | 0 | -1.55182859849935 | 0.992 | 1 | 0 |
| **AXL** | 0 | -2.18124885074164 | 0.026 | 0.706 | 0 |
| **ATP1A3** | 0 | 1.4906103826212 | 0.879 | 0.228 | 0 |
| **MEGF8** | 0 | -1.19473280248741 | 0.482 | 0.872 | 0 |
| **CBLC** | 0 | 1.04835522630259 | 0.698 | 0.105 | 0 |
| **BCAM** | 0 | -2.23730264229244 | 0.632 | 0.965 | 0 |
| **KLC3** | 0 | 1.03048321287879 | 0.8 | 0.215 | 0 |
| **ARHGAP35** | 0 | -1.10454641961349 | 0.887 | 0.991 | 0 |
| **TMEM160** | 0 | 1.23841470150707 | 1 | 0.998 | 0 |
| **CA11** | 0 | -1.73509060690113 | 0.051 | 0.687 | 0 |
| **RASIP1** | 0 | 1.18642519995356 | 0.75 | 0.16 | 0 |
| **RCN3** | 0 | 1.3149625292993 | 0.787 | 0.139 | 0 |
| **PTOV1** | 0 | -0.990918826804477 | 0.99 | 0.999 | 0 |
| **FAM71E1** | 0 | 1.31161910392415 | 0.825 | 0.222 | 0 |
| **KLK13** | 0 | 1.9720387291262 | 0.975 | 0.492 | 0 |
| **LIM2** | 0 | 1.67507442162385 | 0.895 | 0.11 | 0 |
| **ZNF649** | 0 | 1.4921535709127 | 0.996 | 0.904 | 0 |
| **ZNF841** | 0 | 1.4352983629565 | 0.859 | 0.239 | 0 |
| **ZNF880** | 0 | 1.1625948162807 | 0.802 | 0.165 | 0 |
| **ZNF578** | 0 | 1.22363740702979 | 0.882 | 0.304 | 0 |
| **ZNF808** | 0 | 1.23320521418415 | 0.994 | 0.839 | 0 |
| **ZNF600** | 0 | 1.80049501385848 | 0.986 | 0.626 | 0 |
| **ZNF677** | 0 | 1.95109609488584 | 0.947 | 0.071 | 0 |
| **MBOAT7** | 0 | -0.991601549749696 | 0.962 | 0.999 | 0 |

| | | | | | |
|---|---|---|---|---|---|
| **NLRP7** | 0 | 5.03901461071521 | 1 | 0.684 | 0 |
| **NLRP2** | 0 | 2.32794390134265 | 1 | 0.995 | 0 |
| **TNNT1** | 0 | 1.26408479731152 | 1 | 0.885 | 0 |
| **TNNI3** | 0 | 1.40476132027443 | 0.879 | 0.252 | 0 |
| **ZNF667** | 0 | 1.0378743188319 | 0.633 | 0.006 | 0 |
| **PEG3** | 0 | 1.87953820387508 | 0.745 | 0.032 | 0 |
| **ZNF552** | 0 | 1.01582413385379 | 0.997 | 0.862 | 0 |
| **ZCCHC3** | 0 | -1.70431768446921 | 0.575 | 0.95 | 0 |
| **SOX12** | 0 | -1.64896553826239 | 0.601 | 0.94 | 0 |
| **TRIB3** | 0 | 2.04213218642215 | 0.993 | 0.665 | 0 |
| **SNRPB** | 0 | -0.941695158408005 | 1 | 1 | 0 |
| **CPXM1** | 0 | -1.35856108021831 | 0.842 | 0.981 | 0 |
| **HSPA12B** | 0 | 1.1362610513449 | 0.725 | 0.137 | 0 |
| **CDC25B** | 0 | -1.30838501910854 | 0.962 | 0.99 | 0 |
| **CDS2** | 0 | -1.62349894270293 | 0.845 | 0.998 | 0 |
| **FERMT1** | 0 | 1.44705299273666 | 0.853 | 0.309 | 0 |
| **PLCB4** | 0 | -1.2827724966062 | 0.084 | 0.69 | 0 |
| **JAG1** | 0 | -1.30590641061561 | 0.935 | 0.995 | 0 |
| **BTBD3** | 0 | -1.93454768436107 | 0.639 | 0.978 | 0 |
| **SNRPB2** | 0 | 1.0549086372892 | 0.995 | 0.835 | 0 |
| **OVOL2** | 0 | 1.45132709539517 | 0.911 | 0.326 | 0 |
| **RBBP9** | 0 | 1.4301310369711 | 0.996 | 0.863 | 0 |
| **RIN2** | 0 | -1.561073038877 | 0.588 | 0.959 | 0 |
| **XRN2** | 0 | -0.984066043727395 | 1 | 1 | 0 |
| **CST3** | 0 | -2.31211134937832 | 0.661 | 0.988 | 0 |
| **SYNDIG1** | 0 | -1.51682947142599 | 0.047 | 0.721 | 0 |
| **APMAP** | 0 | -1.09125849125687 | 0.977 | 0.999 | 0 |
| **PYGB** | 0 | 2.27063875819218 | 1 | 1 | 0 |
| **ASXL1** | 0 | -1.62607711976551 | 0.588 | 0.945 | 0 |
| **MAPRE1** | 0 | -0.817510202672597 | 1 | 1 | 0 |
| **CBFA2T2** | 0 | 1.2417627872048 | 1 | 0.972 | 0 |
| **DYNLRB1** | 0 | -1.18233947468224 | 1 | 1 | 0 |
| **MMP24OS** | 0 | -1.30727547697091 | 0.692 | 0.968 | 0 |
| **MMP24** | 0 | -1.7857961339783 | 0.682 | 0.953 | 0 |

| | | | | | |
|---|---|---|---|---|---|
| **ERGIC3** | 0 | -1.05634691356816 | 1 | 1 | 0 |
| **CPNE1** | 0 | -1.14693916286845 | 0.99 | 1 | 0 |
| **SCAND1** | 0 | -0.961910613272491 | 0.98 | 0.999 | 0 |
| **MYL9** | 0 | -3.96251425809161 | 0.718 | 0.999 | 0 |
| **TGIF2** | 0 | -2.27379046001278 | 0.638 | 0.995 | 0 |
| **SAMHD1** | 0 | 2.17333371806609 | 1 | 0.906 | 0 |
| **SRC** | 0 | -1.38581607348896 | 0.763 | 0.981 | 0 |
| **BLCAP** | 0 | -0.963464393321953 | 0.885 | 0.99 | 0 |
| **PLCG1** | 0 | -0.975407099262477 | 0.976 | 0.999 | 0 |
| **MYBL2** | 0 | 1.19779952441277 | 1 | 0.989 | 0 |
| **HNF4A** | 0 | 1.27643790989553 | 0.725 | 0.026 | 0 |
| **ADA** | 0 | 1.21861319391807 | 0.945 | 0.617 | 0 |
| **TOMM34** | 0 | -1.05616890403753 | 0.815 | 0.985 | 0 |
| **STK4** | 0 | -1.1163631837055 | 0.85 | 0.987 | 0 |
| **DBNDD2** | 0 | -1.45365530338628 | 0.335 | 0.862 | 0 |
| **PIGT** | 0 | -1.67504228766636 | 0.903 | 0.998 | 0 |
| **ZMYND8** | 0 | -1.04993692635117 | 0.996 | 0.999 | 0 |
| **NCOA3** | 0 | 1.07496479898412 | 0.997 | 0.969 | 0 |
| **SULF2** | 0 | -1.89347535853583 | 0.637 | 0.97 | 0 |
| **STAU1** | 0 | -1.0551819489506 | 1 | 1 | 0 |
| **TMEM189** | 0 | -0.801251158260281 | 0.993 | 1 | 0 |
| **PTPN1** | 0 | -0.913448176909887 | 0.978 | 1 | 0 |
| **BMP7** | 0 | -2.76003150743768 | 0.556 | 0.995 | 0 |
| **VAPB** | 0 | -1.04116937040169 | 0.986 | 0.999 | 0 |
| **NELFCD** | 0 | -0.914727714663356 | 0.989 | 0.999 | 0 |
| **ATP5F1E** | 0 | -1.5846217019835 | 0.989 | 1 | 0 |
| **LSM14B** | 0 | -0.846904234836494 | 0.995 | 1 | 0 |
| **LAMA5** | 0 | -1.38332997891406 | 0.983 | 0.999 | 0 |
| **NKAIN4** | 0 | -1.93058622222178 | 0.369 | 0.869 | 0 |
| **KCNQ2** | 0 | 1.97102144051634 | 0.949 | 0.218 | 0 |
| **EEF1A2** | 0 | 1.31774651810303 | 0.781 | 0.092 | 0 |
| **DNAJC5** | 0 | -0.877273710244576 | 0.972 | 1 | 0 |
| **NRIP1** | 0 | -1.68270432702965 | 0.387 | 0.86 | 0 |
| **APP** | 0 | -1.09505811915462 | 1 | 1 | 0 |

| | | | | | |
|---|---|---|---|---|---|
| **HUNK** | 0 | -1.27200820095031 | 0.171 | 0.707 | 0 |
| **PDXK** | 0 | 1.1411411898778 | 1 | 0.988 | 0 |
| **DNMT3L** | 0 | 3.76172543421579 | 1 | 0.884 | 0 |
| **SUMO3** | 0 | 0.832808772382911 | 1 | 0.998 | 0 |
| **COL18A1** | 0 | -3.81879192787274 | 0.731 | 1 | 0 |
| **SPATC1L** | 0 | -1.42002847786006 | 0.084 | 0.776 | 0 |
| **LSS** | 0 | -1.06561679959913 | 0.955 | 0.997 | 0 |
| **HDHD5** | 0 | 0.852104917316545 | 1 | 0.929 | 0 |
| **MED15** | 0 | 1.87683175118565 | 1 | 0.933 | 0 |
| **HIC2** | 0 | 1.4427232435229 | 1 | 0.929 | 0 |
| **MIF** | 0 | 0.840811945166372 | 1 | 1 | 0 |
| **SUSD2** | 0 | 2.34024249418619 | 0.999 | 0.798 | 0 |
| **TPST2** | 0 | 1.41615706909115 | 0.992 | 0.815 | 0 |
| **CCDC117** | 0 | 1.32698760574093 | 0.98 | 0.643 | 0 |
| **XBP1** | 0 | 1.03568472242883 | 1 | 0.997 | 0 |
| **NEFH** | 0 | 2.73036602145748 | 0.997 | 0.622 | 0 |
| **NF2** | 0 | 1.02091933660796 | 1 | 0.996 | 0 |
| **ASCC2** | 0 | 0.947871944984222 | 0.999 | 0.967 | 0 |
| **TCN2** | 0 | -1.90189648541655 | 0.464 | 0.963 | 0 |
| **TOMM22** | 0 | 1.24064193103611 | 1 | 0.99 | 0 |
| **SNU13** | 0 | 0.900282341890511 | 1 | 0.999 | 0 |
| **PHETA2** | 0 | 0.921453438396736 | 0.558 | 0.009 | 0 |
| **NDUFA6** | 0 | 1.21625789931331 | 1 | 0.97 | 0 |
| **RRP7A** | 0 | 0.849917463873368 | 1 | 0.972 | 0 |
| **A4GALT** | 0 | 2.63505794876606 | 1 | 0.756 | 0 |
| **PARVB** | 0 | 1.69888919196559 | 0.895 | 0.294 | 0 |
| **ARHGAP8** | 0 | 1.74023181227751 | 0.997 | 0.745 | 0 |
| **KIAA0930** | 0 | 0.939931198763194 | 0.989 | 0.852 | 0 |
| **FBLN1** | 0 | -1.9057996753456 | 0.875 | 0.998 | 0 |
| **CRELD2** | 0 | 1.48248156142336 | 0.981 | 0.928 | 0 |
| **TRABD** | 0 | 0.913526345755244 | 0.997 | 0.936 | 0 |
| **PLXNB2** | 0 | 0.896625087885345 | 0.999 | 0.945 | 0 |
| **CSF2RA** | 0 | 1.78494202268574 | 0.855 | 0.234 | 0 |
| **DHRSX** | 0 | 1.59820703219652 | 0.953 | 0.834 | 0 |

| | | | | | |
|---|---|---|---|---|---|
| **CD99** | 0 | -2.92011678305511 | 0.163 | 0.966 | 0 |
| **MXRA5** | 0 | 2.56436952665448 | 0.935 | 0.319 | 0 |
| **TBL1X** | 0 | -1.59689751767682 | 0.278 | 0.855 | 0 |
| **WWC3** | 0 | -1.62204928200255 | 0.418 | 0.906 | 0 |
| **TMSB4X** | 0 | -2.89841932987723 | 0.997 | 1 | 0 |
| **AP1S2** | 0 | -1.69703425857715 | 0.404 | 0.881 | 0 |
| **CTPS2** | 0 | -1.3189626251041 | 0.848 | 0.998 | 0 |
| **RBBP7** | 0 | -1.17606217208705 | 0.96 | 0.998 | 0 |
| **SH3KBP1** | 0 | -1.74380095895546 | 0.776 | 0.99 | 0 |
| **SMS** | 0 | -2.53948532600117 | 0.87 | 1 | 0 |
| **PHEX** | 0 | 1.76312732846139 | 0.855 | 0.054 | 0 |
| **ACOT9** | 0 | -1.13751568137073 | 0.71 | 0.952 | 0 |
| **EIF2S3** | 0 | 1.388821494279 | 0.905 | 0.305 | 0 |
| **PCYT1B** | 0 | -1.79055740665699 | 0.043 | 0.814 | 0 |
| **MAGEB2** | 0 | 1.48158416730203 | 0.594 | 0.016 | 0 |
| **USP9X** | 0 | -0.932777736779419 | 1 | 1 | 0 |
| **CASK** | 0 | -1.44688177116272 | 0.817 | 0.999 | 0 |
| **USP11** | 0 | -1.26349578046859 | 0.919 | 0.998 | 0 |
| **TIMP1** | 0 | -2.29975831779793 | 0.863 | 0.999 | 0 |
| **GSPT2** | 0 | -1.72959279397311 | 0.529 | 0.975 | 0 |
| **MAGED1** | 0 | -2.36330517329787 | 0.877 | 1 | 0 |
| **WNK3** | 0 | -1.29493903127067 | 0.413 | 0.871 | 0 |
| **MAGED2** | 0 | -1.84019705703991 | 0.989 | 1 | 0 |
| **TRO** | 0 | -1.43609080589537 | 0.699 | 0.98 | 0 |
| **MAGEH1** | 0 | -1.65717657059163 | 0.353 | 0.925 | 0 |
| **UBQLN2** | 0 | -1.36216440473111 | 0.897 | 1 | 0 |
| **ARHGEF9** | 0 | -1.77927316914309 | 0.211 | 0.893 | 0 |
| **ZC4H2** | 0 | -1.18487495870017 | 0.296 | 0.808 | 0 |
| **MSN** | 0 | -1.19776943715788 | 0.959 | 1 | 0 |
| **HEPH** | 0 | -2.26899008932884 | 0.009 | 0.895 | 0 |
| **STARD8** | 0 | -1.04923493200608 | 0.016 | 0.529 | 0 |
| **EFNB1** | 0 | -3.37493975696743 | 0.191 | 0.904 | 0 |
| **EDA** | 0 | -1.42249685640072 | 0.503 | 0.92 | 0 |
| **DLG3** | 0 | 1.00945280775333 | 0.999 | 0.987 | 0 |

| | | | | | |
|---|---|---|---|---|---|
| **SNX12** | 0 | -1.26830267520308 | 0.931 | 1 | 0 |
| **FOXO4** | 0 | -2.4485210988191 | 0.192 | 0.883 | 0 |
| **NONO** | 0 | 1.83049264009309 | 1 | 0.857 | 0 |
| **PHKA1** | 0 | 1.87455188747684 | 0.999 | 0.871 | 0 |
| **NAP1L2** | 0 | -1.18436784665873 | 0.279 | 0.778 | 0 |
| **SLC16A2** | 0 | -2.22447370747744 | 0.496 | 0.969 | 0 |
| **RLIM** | 0 | 1.5019478323687 | 0.862 | 0.134 | 0 |
| **ABCB7** | 0 | -1.33776989820187 | 0.909 | 0.998 | 0 |
| **COX7B** | 0 | 1.96454081437558 | 0.997 | 0.748 | 0 |
| **BRWD3** | 0 | 0.864257880596501 | 1 | 0.984 | 0 |
| **SH3BGRL** | 0 | -1.87678775373548 | 0.468 | 0.975 | 0 |
| **ZNF711** | 0 | -1.44388076163629 | 0.306 | 0.836 | 0 |
| **KLHL4** | 0 | -1.01031067215067 | 0.003 | 0.492 | 0 |
| **NAP1L3** | 0 | -2.13245568810192 | 0.263 | 0.907 | 0 |
| **TNMD** | 0 | -1.46773318680388 | 0.021 | 0.596 | 0 |
| **TSPAN6** | 0 | -1.64804687945324 | 0.908 | 0.997 | 0 |
| **ARMCX1** | 0 | -1.55820274626997 | 0.715 | 0.991 | 0 |
| **ARMCX2** | 0 | -1.56038635784162 | 0.848 | 0.998 | 0 |
| **BEX4** | 0 | -1.66443240418538 | 0.884 | 1 | 0 |
| **TCEAL8** | 0 | -1.20624094577849 | 0.955 | 1 | 0 |
| **TCEAL9** | 0 | -1.74305271777198 | 0.755 | 0.994 | 0 |
| **BEX3** | 0 | -2.86660418791728 | 0.905 | 1 | 0 |
| **NRK** | 0 | -3.00069569000139 | 0.014 | 0.76 | 0 |
| **COL4A6** | 0 | -1.88938041694806 | 0.006 | 0.726 | 0 |
| **COL4A5** | 0 | -2.68552954398241 | 0.068 | 0.896 | 0 |
| **ACSL4** | 0 | -1.54368476658357 | 0.634 | 0.987 | 0 |
| **PGRMC1** | 0 | -1.71986717705626 | 0.983 | 1 | 0 |
| **SLC25A5** | 0 | 0.99975865372985 | 0.67 | 0.087 | 0 |
| **SEPTIN6** | 0 | -2.14789878086572 | 0.179 | 0.938 | 0 |
| **SMARCA1** | 0 | -1.57770898256071 | 0.639 | 0.971 | 0 |
| **STK26** | 0 | -1.28523158636615 | 0.806 | 0.993 | 0 |
| **MBNL3** | 0 | -2.57691422208674 | 0.343 | 0.958 | 0 |
| **HS6ST2** | 0 | -2.1773528774582 | 0.063 | 0.864 | 0 |
| **GPC4** | 0 | -2.98281109632043 | 0.619 | 0.993 | 0 |

| | | | | | |
|---|---|---|---|---|---|
| **ZIC3** | 0 | -1.97290768812622 | 0.424 | 0.941 | 0 |
| **LDOC1** | 0 | -1.62779956284379 | 0.471 | 0.962 | 0 |
| **IDS** | 0 | -1.51184402106907 | 0.36 | 0.892 | 0 |
| **VMA21** | 0 | -0.882038491586628 | 0.98 | 1 | 0 |
| **ZNF185** | 0 | -1.60579098973455 | 0.347 | 0.878 | 0 |
| **CCNQ** | 0 | 1.18719159502408 | 0.986 | 0.809 | 0 |
| **DUSP9** | 0 | 1.550303036625 | 0.783 | 0.043 | 0 |
| **SLC6A8** | 0 | 1.67417407805109 | 1 | 0.979 | 0 |
| **FLNA** | 0 | -1.02407065400567 | 1 | 1 | 0 |
| **LAGE3** | 0 | 0.802750417377827 | 1 | 0.994 | 0 |
| **MT-ND2** | 0 | -1.85089833972357 | 0.771 | 0.991 | 0 |
| **MT-CO2** | 0 | -1.64096563591344 | 0.963 | 0.999 | 0 |
| **MT-CO3** | 0 | -2.24936680183978 | 0.956 | 1 | 0 |
| **MT-CYB** | 0 | -1.78622525413805 | 0.929 | 0.999 | 0 |
| **SEPHS1** | 4.8197 | -1.1429273142392 | 0.975 | 0.995 | 7.8383327 |
| **PTPN21** | 1.0777 | -1.15985714865522 | 0.198 | 0.715 | 1.7527959 |
| **SLC35A5** | 1.1618 | 0.974325549535834 | 0.995 | 0.914 | 1.8895519 |
| **S100A11** | 1.4079 | -1.22187338032174 | 0.995 | 0.999 | 2.2897061 |
| **CTSV** | 2.3050 | -1.64749362513454 | 0.867 | 0.981 | 3.7487773 |
| **EPDR1** | 4.7170 | -1.0283643199315 | 0.012 | 0.491 | 7.6712743 |
| **ATP8B2** | 5.5031 | 0.918122611607667 | 0.995 | 0.959 | 8.9498290 |
| **SLC16A1** | 9.9642 | -0.979419846084753 | 1 | 1 | 1.6204914 |
| **RALGAPA2** | 1.4660 | -1.1389910652693 | 0.744 | 0.963 | 2.3842424 |
| **NUAK2** | 1.8236 | -1.49149113152051 | 0.24 | 0.731 | 2.9658418 |
| **ARHGDIA** | 2.9011 | -0.838072974371113 | 0.981 | 0.999 | 4.7181884 |
| **PRTG** | 5.0041 | -2.5016962772978 | 0.025 | 0.511 | 8.1382436 |
| **LPIN1** | 6.6657 | 1.04152253404865 | 0.918 | 0.514 | 1.0840499 |
| **CIB1** | 7.3491 | 0.905122764844055 | 0.99 | 0.884 | 1.1951972 |
| **GPNMB** | 1.3690 | 1.10491576703941 | 0.957 | 0.576 | 2.2265500 |
| **KIZ** | 1.6474 | -1.18308647610047 | 0.432 | 0.847 | 2.6792259 |
| **TUBGCP2** | 1.8274 | 0.849942098171306 | 0.993 | 0.916 | 2.9719312 |
| **PRRX2** | 1.9685 | 1.07683213783317 | 0.865 | 0.358 | 3.2014263 |
| **PGM2** | 2.1748 | 1.01106597077122 | 0.986 | 0.868 | 3.5368906 |
| **FOXR1** | 2.8866 | 0.963342158258963 | 0.528 | 0.005 | 4.6946119 |

| | | | | | |
|---|---|---|---|---|---|
| **TMEM184C** | 5.9797 | 0.959053956379074 | 0.923 | 0.562 | 9.72491721 |
| **EIF1AX** | 6.0652 | -0.919642363102024 | 0.97 | 0.996 | 9.86397676 |
| **AMER1** | 1.2937 | -1.14811651218631 | 0.406 | 0.834 | 2.10401514 |
| **MFSD4B** | 1.4006 | 1.04554141228285 | 0.982 | 0.816 | 2.27794665 |
| **HCN2** | 3.2811 | 0.979181140119722 | 0.974 | 0.673 | 5.33609023 |
| **EML1** | 4.8621 | -1.09874639756441 | 0.853 | 0.979 | 7.90737552 |
| **EIF2AK2** | 5.3652 | -1.00633227088624 | 0.637 | 0.939 | 8.72543692 |
| **SYK** | 1.0234 | 1.06168751141017 | 0.752 | 0.19 | 1.66436114 |
| **MFSD13A** | 1.2535 | 1.11768712995189 | 0.821 | 0.318 | 2.03863170 |
| **RIMKLB** | 1.5366 | 1.01155696109058 | 0.999 | 0.984 | 2.49912022 |
| **FBXL16** | 4.7666 | 1.02150736088037 | 0.715 | 0.159 | 7.75195795 |
| **SOGA1** | 1.0604 | -0.99636091925345 | 0.88 | 0.991 | 1.72463950 |
| **CITED2** | 4.4751 | -1.71891148584072 | 0.795 | 0.975 | 7.27799247 |
| **PAK1** | 5.2930 | 0.932996066659186 | 0.986 | 0.855 | 8.60802483 |
| **MPP6** | 5.5526 | -1.0467133787922 | 0.975 | 0.995 | 9.03026092 |
| **S100A13** | 5.9414 | 1.05612037388576 | 0.997 | 0.98 | 9.66261279 |
| **CFAP46** | 1.5980 | 0.884012621013892 | 0.588 | 0.049 | 2.59895800 |
| **CDK14** | 2.0675 | -1.40631421166668 | 0.15 | 0.662 | 3.36248014 |
| **CDHR1** | 1.2577 | 1.37448575035209 | 0.956 | 0.682 | 2.04547835 |
| **COL1A2** | 1.5998 | -2.41099672565079 | 0.568 | 0.95 | 2.60188857 |
| **SHOX** | 1.7694 | 0.844663164092301 | 0.538 | 0.017 | 2.87762493 |
| **CHORDC1** | 2.2650 | 0.99112168412852 | 0.79 | 0.263 | 3.68369681 |
| **OSER1** | 3.0913 | -0.935923303422346 | 0.874 | 0.984 | 5.02754135 |
| **ZNF354C** | 5.3778 | 0.804472924511099 | 0.52 | 0.006 | 8.74601222 |
| **GRAMD1B** | 2.5290 | -1.20163031607201 | 0.019 | 0.492 | 4.11303779 |
| **RB1** | 2.6806 | -1.07287171674915 | 0.323 | 0.781 | 4.35946045 |
| **CAP2** | 3.1321 | -1.18540001374905 | 0.595 | 0.908 | 5.09383693 |
| **ELMOD2** | 3.2809 | -1.04930214187663 | 0.531 | 0.896 | 5.33576840 |
| **SFT2D1** | 3.7860 | 0.999895758159724 | 0.892 | 0.492 | 6.15721923 |
| **CACHD1** | 5.8909 | -1.3371485512081 | 0.66 | 0.958 | 9.58039384 |
| **MID1IP1** | 9.1327 | -0.902526499648437 | 0.99 | 0.999 | 1.48525403 |
| **TNKS1BP1** | 1.2136 | -0.980276575539951 | 0.837 | 0.979 | 1.97382697 |
| **CACNA1B** | 2.2578 | 1.06848480165948 | 0.837 | 0.324 | 3.67200565 |
| **SVBP** | 2.4469 | -0.975091088300763 | 0.881 | 0.985 | 3.97939745 |

| | | | | | |
|---|---|---|---|---|---|
| **DDAH1** | 7.0818 | -1.41879563933353 | 0.339 | 0.809 | 1.1517179 |
| **DYNLT3** | 2.2258 | 0.8171961764634 | 0.519 | 0.007 | 3.6199685 |
| **VAMP7** | 4.1855 | 0.846385840985232 | 0.993 | 0.905 | 6.8070387 |
| **JUND** | 1.0365 | -0.960187231363926 | 1 | 1 | 1.6857163 |
| **NUDT8** | 1.4408 | 0.974004321549308 | 0.653 | 0.103 | 2.3431776 |
| **PJA1** | 3.3604 | -1.06452956064522 | 0.815 | 0.985 | 5.4650960 |
| **TMA7** | 4.0288 | 0.991950651246806 | 0.855 | 0.373 | 6.5521625 |
| **CISD1** | 5.2187 | 0.90292688360665 | 0.988 | 0.886 | 8.4872069 |
| **CDIP1** | 1.6116 | -0.998862739079005 | 0.838 | 0.978 | 2.6211004 |
| **SLC30A5** | 3.2416 | 0.920992362736505 | 0.945 | 0.622 | 5.2718738 |
| **IPMK** | 3.8589 | 0.945164801860196 | 0.992 | 0.845 | 6.2757484 |
| **PDK1** | 5.0465 | -1.49285746958771 | 0.742 | 0.954 | 8.2072794 |
| **EXTL2** | 1.0746 | -0.985083314586729 | 0.76 | 0.955 | 1.7476579 |
| **WBP1** | 1.4474 | -0.974150901711368 | 0.884 | 0.986 | 2.3539824 |
| **SHISA3** | 2.2091 | -1.78643356309887 | 0.029 | 0.501 | 3.5927398 |
| **CETN3** | 2.3991 | 0.938025166497439 | 0.967 | 0.756 | 3.9017364 |
| **TINCR** | 4.8809 | 0.950238740646526 | 0.714 | 0.158 | 7.9379630 |
| **TRERF1** | 5.5977 | -0.982748310418447 | 0.02 | 0.489 | 9.1035475 |
| **ST14** | 1.0196 | 1.06985006872356 | 0.704 | 0.162 | 1.6582909 |
| **PPP2CB** | 1.0889 | -0.946093663263416 | 0.703 | 0.942 | 1.7710120 |
| **LIMD2** | 1.2204 | -1.13315709152367 | 0.488 | 0.865 | 1.9848275 |
| **TP53I3** | 3.3029 | -1.10392097341573 | 0.589 | 0.901 | 5.3715216 |
| **CACFD1** | 3.8649 | -1.05881074336824 | 0.638 | 0.925 | 6.2856109 |
| **NAGK** | 6.9788 | 0.917438906005501 | 0.997 | 0.973 | 1.1349742 |
| **REEP2** | 1.1837 | -1.35362315740264 | 0.218 | 0.709 | 1.9251428 |
| **CXCL12** | 1.3751 | -1.7704179287808 | 0.907 | 0.927 | 2.2363462 |
| **DNAJB1** | 3.7445 | 0.90610261961031 | 0.996 | 0.962 | 6.0897069 |
| **GDF11** | 6.8001 | 0.812280549796664 | 1 | 0.985 | 1.1059127 |
| **NTF3** | 9.4076 | -1.12174127323291 | 0.033 | 0.506 | 1.5299704 |
| **NR5A2** | 9.4865 | 1.16253092712674 | 0.606 | 0.078 | 1.5428033 |
| **MT-ND4** | 1.3162 | -1.37695676420904 | 0.973 | 0.999 | 2.1405665 |
| **VPS26A** | 2.5563 | 0.841267540290954 | 0.997 | 0.942 | 4.1574084 |
| **MAPRE2** | 3.0495 | -1.11862315345941 | 0.614 | 0.908 | 4.9594821 |
| **PDCL** | 3.5988 | 0.806831780266938 | 0.997 | 0.988 | 5.8528897 |

| | | | | | |
|---|---|---|---|---|---|
| **ADK** | 7.0968 | -0.972054658273494 | 0.784 | 0.969 | 1.15416679 |
| **SLC35B2** | 7.1440 | -0.910527374844021 | 0.877 | 0.989 | 1.16184476 |
| **EYA2** | 9.0533 | -1.32620776832437 | 0.119 | 0.613 | 1.47234441 |
| **DLGAP4** | 1.0663 | -1.11708882455161 | 0.938 | 0.994 | 1.73419984 |
| **ACTR2** | 1.4032 | 0.891139314801828 | 0.975 | 0.75 | 2.28213710 |
| **SAT1** | 2.9000 | -1.61966644396865 | 0.927 | 0.994 | 4.71633151 |
| **BCL3** | 9.3188 | 1.10267186743425 | 0.9 | 0.483 | 1.51552715 |
| **PTMS** | 1.0266 | -1.07701573235563 | 1 | 1 | 1.66957390 |
| **TUBB2B** | 1.8654 | -1.02732311890644 | 1 | 1 | 3.03371880 |
| **ZNF229** | 2.2316 | 1.00617202551073 | 0.793 | 0.281 | 3.62926289 |
| **NAPRT** | 3.0207 | 0.997103795287937 | 0.921 | 0.515 | 4.91263650 |
| **PRKAB2** | 1.2212 | 0.933585799313109 | 0.982 | 0.861 | 1.98610571 |
| **RGS10** | 2.6862 | 1.01060820383823 | 0.88 | 0.448 | 4.36859907 |
| **LRAT** | 3.0797 | -1.64367519561607 | 0.067 | 0.545 | 5.00853763 |
| **OST4** | 3.5918 | 0.923936066748019 | 0.746 | 0.221 | 5.84147763 |
| **ERICH5** | 4.1583 | 0.908592254470791 | 0.612 | 0.076 | 6.76272150 |
| **CALM2** | 4.2195 | -1.01433163606796 | 0.985 | 1 | 6.86233395 |
| **SSR1** | 3.7131 | 0.905933221949848 | 0.977 | 0.761 | 6.03864630 |
| **SLC23A2** | 4.4974 | 1.00040144070433 | 0.987 | 0.912 | 7.31412801 |
| **FXYD6** | 5.0819 | 1.0803938224571 | 0.983 | 0.777 | 8.26476530 |
| **PTCD1** | 5.6458 | 0.8313770623628 | 0.992 | 0.904 | 9.18177711 |
| **RAVER2** | 8.3427 | -1.06193088325018 | 0.322 | 0.783 | 1.35678882 |
| **DMBX1** | 2.1357 | 1.11183087402457 | 0.825 | 0.285 | 3.47332779 |
| **ZNF580** | 5.2005 | -1.19914660575678 | 0.42 | 0.837 | 8.45758660 |
| **TP53BP2** | 5.9531 | -0.91056013568892 | 0.872 | 0.984 | 9.68163138 |
| **AGPAT3** | 9.6782 | 0.919855602470422 | 0.689 | 0.136 | 1.57397184 |
| **TMSB15A** | 1.0448 | -1.75831160405056 | 0.087 | 0.571 | 1.69916505 |
| **IGLON5** | 1.1585 | -1.11563690441367 | 0.218 | 0.711 | 1.88406925 |
| **TFE3** | 1.3312 | -0.940531612916932 | 0.889 | 0.989 | 2.16502960 |
| **STC1** | 2.2156 | -1.12792630778946 | 0.027 | 0.492 | 3.60333097 |
| **PLD6** | 6.1903 | 0.896232076211038 | 0.536 | 0.028 | 1.00672925 |
| **LGI1** | 2.7957 | -1.74255508828434 | 0.003 | 0.443 | 4.54678719 |
| **KIRREL1** | 6.9979 | 1.00189688436922 | 0.982 | 0.876 | 1.13807137 |
| **EXD2** | 7.7263 | -1.05043582824407 | 0.52 | 0.874 | 1.25654267 |

| | | | | | |
|---|---|---|---|---|---|
| **TNC** | 9.3512 | -1.49247472572351 | 0.008 | 0.453 | 1.52078921 |
| **CRABP1** | 2.7343 | -3.30746107114266 | 0.408 | 0.815 | 4.44688182 |
| **PDCD4** | 5.1373 | -1.13836826038891 | 0.426 | 0.842 | 8.35481345 |
| **GSTK1** | 5.2609 | -1.11750437404059 | 0.611 | 0.91 | 8.55584687 |
| **STAT3** | 1.0588 | -0.901785188113273 | 0.995 | 0.999 | 1.72197393 |
| **F2RL1** | 1.9675 | 0.895077344697859 | 0.998 | 0.956 | 3.19989290 |
| **RAPGEF2** | 1.9791 | 0.866553879634688 | 0.992 | 0.908 | 3.21861979 |
| **PSMC4** | 5.1434 | 0.811126767335414 | 1 | 0.998 | 8.36475360 |
| **LUZP4** | 6.3340 | 0.992676021289044 | 0.502 | 0.008 | 1.03010324 |
| **MCM5** | 8.8151 | 0.890235205227446 | 0.999 | 0.982 | 1.43360653 |
| **POMGNT1** | 8.8671 | -0.88192800919675 | 0.85 | 0.99 | 1.44206200 |
| **KIAA1324L** | 9.3516 | -1.01459731211423 | 0.561 | 0.895 | 1.52085216 |
| **NEPRO** | 1.1747 | 0.810664195631742 | 0.995 | 0.942 | 1.91053139 |
| **SEMA3A** | 1.4365 | -1.02443286202481 | 0.128 | 0.623 | 2.33626877 |
| **IL6R** | 1.4801 | 1.03564825019176 | 0.84 | 0.368 | 2.40721244 |
| **CTTNBP2** | 1.8691 | -0.842047432110342 | 0.014 | 0.461 | 3.03977448 |
| **SMIM26** | 2.1245 | -0.812979709740361 | 0.96 | 0.997 | 3.45521895 |
| **PROM1** | 3.6860 | 1.01194444568076 | 0.992 | 0.858 | 5.99461424 |
| **HIP1** | 3.9011 | -1.02238618096632 | 0.662 | 0.928 | 6.34449612 |
| **NRXN2** | 5.0372 | 0.846724376305069 | 0.595 | 0.068 | 8.19207210 |
| **PDS5A** | 5.0809 | 0.879141449042997 | 0.978 | 0.785 | 8.26316112 |
| **SLAIN2** | 2.0862 | 0.814800917897081 | 0.992 | 0.918 | 3.39294813 |
| **ARL4C** | 6.9840 | -2.68839588539544 | 0.138 | 0.616 | 1.13581110 |
| **TLE4** | 9.7176 | -1.92407695384203 | 0.305 | 0.758 | 1.58038070 |
| **RPA3** | 6.9725 | 0.933663532186792 | 0.785 | 0.277 | 1.13394459 |
| **BTG2** | 1.1654 | -1.64619091528265 | 0.347 | 0.801 | 1.89534728 |
| **ST13** | 1.2867 | 0.843686456100345 | 0.97 | 0.753 | 2.09259141 |
| **GAR1** | 2.3881 | 0.859112792215611 | 0.985 | 0.892 | 3.88382345 |
| **TCEAL4** | 2.7125 | -0.828262395907069 | 0.941 | 0.998 | 4.41139122 |
| **COL2A1** | 4.3716 | -2.31624960038861 | 0.003 | 0.436 | 7.10965252 |
| **LIMK1** | 5.8305 | -0.849580155931656 | 0.943 | 0.995 | 9.48229857 |
| **NUPR1** | 6.6045 | 1.3734274592408 | 0.689 | 0.175 | 1.07409749 |
| **PRKG1** | 1.0063 | -1.03938897376874 | 0.006 | 0.442 | 1.63663849 |
| **LYPD5** | 1.6096 | 0.819868910743978 | 0.522 | 0.025 | 2.61781589 |

| | | | | | |
|---|---|---|---|---|---|
| **TSC22D1** | 1.8268 | 0.943953658405622 | 0.94 | 0.65 | 2.97107748 |
| **EPPK1** | 3.9189 | 0.872825797061629 | 1 | 0.985 | 6.37340243 |
| **CILP2** | 3.0948 | 1.09964828646511 | 0.842 | 0.392 | 5.03319448 |
| **YWHAB** | 5.3414 | 0.865476146832047 | 0.995 | 0.911 | 8.68677644 |
| **LUZP1** | 1.9584 | -0.927873908016882 | 0.906 | 0.995 | 3.18502757 |
| **TACC2** | 9.2624 | -1.01260384045944 | 0.529 | 0.879 | 1.50634602 |
| **TLN2** | 1.0002 | 0.825517394851711 | 0.993 | 0.947 | 1.62665319 |
| **OBI1** | 1.0204 | -1.01438567546362 | 0.761 | 0.957 | 1.65962118 |
| **NLGN4X** | 1.3387 | -1.81914387245941 | 0.088 | 0.56 | 2.17713759 |
| **WDR34** | 3.7343 | 0.890425371194343 | 0.993 | 0.936 | 6.07314358 |
| **SNN** | 3.8447 | -0.980799103664161 | 0.717 | 0.942 | 6.25268281 |
| **AMZ2** | 5.3176 | 0.909038235850366 | 0.931 | 0.617 | 8.64816842 |
| **HDGF** | 7.6462 | -0.902019234193327 | 0.961 | 0.997 | 1.24350408 |
| **PHLDB1** | 1.5153 | -1.13513843288189 | 0.631 | 0.908 | 2.46448184 |
| **TMLHE** | 2.7903 | -1.0240781254118 | 0.403 | 0.819 | 4.53791042 |
| **BCAR1** | 5.4375 | -1.12886916102471 | 0.877 | 0.977 | 8.84314754 |
| **CIZ1** | 8.4901 | -1.20069238237354 | 0.332 | 0.765 | 1.38074924 |
| **SNX24** | 1.3901 | 0.866469417388094 | 0.982 | 0.847 | 2.26078983 |
| **FHDC1** | 1.8228 | 1.09475315851275 | 0.758 | 0.254 | 2.96450151 |
| **ARHGAP31** | 2.5188 | 0.89650757214516 | 0.593 | 0.079 | 4.09638845 |
| **CRTC3** | 3.0500 | -1.07381240596428 | 0.246 | 0.715 | 4.96029881 |
| **ACADSB** | 6.5831 | -1.0150648427132 | 0.197 | 0.676 | 1.07061244 |
| **TSPAN9** | 8.6487 | -1.03695619029029 | 0.758 | 0.952 | 1.40654891 |
| **ATP12A** | 9.7958 | 1.04315603986458 | 0.927 | 0.504 | 1.59309556 |
| **MYL7** | 9.9319 | -1.71388609268111 | 0.015 | 0.45 | 1.61522884 |
| **IL34** | 1.0765 | 0.950031601603451 | 0.714 | 0.195 | 1.75086106 |
| **CA13** | 1.9322 | 1.22534532216753 | 0.845 | 0.443 | 3.14247362 |
| **DNAJB4** | 2.9287 | 0.979704688293875 | 0.912 | 0.555 | 4.76303909 |
| **TTC3** | 3.9407 | -1.09443734576867 | 0.944 | 0.992 | 6.40886274 |
| **UCK2** | 5.8240 | -0.939782233075062 | 0.897 | 0.992 | 9.47167806 |
| **TLNRD1** | 6.9510 | -1.05060228812737 | 0.769 | 0.946 | 1.13044728 |
| **ZFP36L2** | 1.0325 | -1.20345894877928 | 0.902 | 0.99 | 1.67928786 |
| **CBX4** | 2.0864 | 0.988612429621 | 0.981 | 0.819 | 3.39321139 |
| **AKAP3** | 6.5853 | 0.820587270946296 | 0.531 | 0.037 | 1.07097356 |

| | | | | | |
|---|---|---|---|---|---|
| **AFAP1L1** | 7.7328 | 0.978730365287015 | 0.672 | 0.156 | 1.25759958 |
| **SRXN1** | 8.3610 | -1.14238557891653 | 0.609 | 0.91 | 1.35975223 |
| **TRIM38** | 1.1710 | 1.22962707768458 | 0.873 | 0.495 | 1.90448358 |
| **RET** | 1.8246 | -1.47424196419414 | 0.345 | 0.756 | 2.96739077 |
| **EIF2AK4** | 2.0034 | 0.883498680030299 | 0.956 | 0.719 | 3.25820939 |
| **ABR** | 2.7021 | 0.832332955914977 | 0.997 | 0.925 | 4.39456008 |
| **MT-ATP6** | 5.1427 | -1.38703902335864 | 0.955 | 0.999 | 8.36357590 |
| **EPHB1** | 8.6353 | -0.954742753620834 | 0.046 | 0.502 | 1.40436270 |
| **PXDN** | 8.7880 | 0.910059243013362 | 0.997 | 0.874 | 1.42920847 |
| **USP47** | 8.8036 | -0.972470090752392 | 0.732 | 0.946 | 1.43173550 |
| **MT-ND4L** | 9.6311 | -1.34725587127467 | 0.881 | 0.996 | 1.56630649 |
| **USP46** | 1.1725 | -0.929596881889324 | 0.704 | 0.935 | 1.90692201 |
| **MFAP2** | 1.5258 | -1.14122716731838 | 0.96 | 0.992 | 2.48144214 |
| **NRARP** | 3.6514 | -1.18564060868894 | 0.23 | 0.7 | 5.93829014 |
| **C1GALT1C1** | 3.6686 | -0.97166409753738 | 0.765 | 0.957 | 5.96636961 |
| **FKBP11** | 4.7219 | 1.02485267157421 | 0.91 | 0.531 | 7.67935942 |
| **AXIN2** | 7.1317 | -1.0994127729316 | 0.794 | 0.965 | 1.15984291 |
| **DPF1** | 9.7619 | 1.14572254678335 | 0.866 | 0.49 | 1.58757923 |
| **LRRC61** | 1.4113 | -0.978451640420019 | 0.27 | 0.735 | 2.29522260 |
| **C11orf1** | 2.1985 | -0.917930305087651 | 0.755 | 0.958 | 3.57546981 |
| **QDPR** | 3.3000 | -0.846011288457123 | 0.934 | 0.993 | 5.36689635 |
| **DPH6** | 6.6642 | 0.914289692286356 | 0.961 | 0.722 | 1.08380970 |
| **RPUSD4** | 9.3714 | -0.88798066826339 | 0.838 | 0.978 | 1.52408400 |
| **ZNF718** | 1.3563 | 0.822142401540804 | 0.608 | 0.098 | 2.20577540 |
| **RBMXL1** | 2.2451 | 0.897158481766368 | 0.905 | 0.537 | 3.65130444 |
| **HDAC3** | 2.2729 | -0.816277565069409 | 0.895 | 0.984 | 3.69647630 |
| **SGCB** | 2.2910 | -0.961147802112843 | 0.678 | 0.929 | 3.72594407 |
| **HERPUD2** | 2.3455 | -1.01267556610438 | 0.609 | 0.9 | 3.81460671 |
| **MED12** | 6.4730 | -1.05644008226884 | 0.108 | 0.579 | 1.05270844 |
| **LGALS3** | 1.0487 | 1.1983493066333 | 0.668 | 0.158 | 1.70551534 |
| **FUNDC1** | 1.6499 | 0.894599699775929 | 0.959 | 0.749 | 2.68335079 |
| **ENO2** | 1.4207 | 0.964028316777178 | 0.995 | 0.974 | 2.31059335 |
| **CHMP5** | 1.7007 | 0.811071306629307 | 0.976 | 0.813 | 2.76588553 |
| **CRYGD** | 2.1909 | 0.868496733831118 | 0.49 | 0.016 | 3.56308391 |

| | | | | | |
|---|---|---|---|---|---|
| **FAM20A** | 2.8700 | -1.78544504795641 | 0.526 | 0.846 | 4.6675533 |
| **NOP9** | 4.2708 | 0.863813073837275 | 0.942 | 0.69 | 6.9456882 |
| **SARNP** | 8.4262 | 0.83723165439768 | 0.967 | 0.754 | 1.3703653 |
| **1,00 PGK** | 2.6193 | 1.01773014165142 | 0.736 | 0.248 | 4.2598895 |
| **CASP2** | 3.0014 | -1.05384477986306 | 0.732 | 0.941 | 4.8812742 |
| **ZFAND3** | 4.3376 | -0.812990654191844 | 0.948 | 0.995 | 7.0543994 |
| **IFI6** | 4.4076 | -1.16638783745967 | 0.08 | 0.543 | 7.1681006 |
| **SPTB** | 8.5635 | -1.37016492524199 | 0.241 | 0.687 | 1.3926844 |
| **MINDY2** | 1.5936 | 0.80927658778891 | 0.985 | 0.871 | 2.5918001 |
| **OSBPL10** | 1.9343 | 0.953922157380159 | 0.977 | 0.85 | 3.1458993 |
| **LDLRAP1** | 2.3435 | 0.824502532752386 | 0.996 | 0.914 | 3.8113944 |
| **CCDC97** | 3.7393 | -0.872982087564168 | 0.822 | 0.971 | 6.0813324 |
| **HYLS1** | 5.5405 | 1.15399162435087 | 0.652 | 0.149 | 9.0105507 |
| **COA6** | 9.6918 | 0.853419315976639 | 0.962 | 0.753 | 1.5761846 |
| **SAT2** | 1.4972 | -1.06573874088551 | 0.906 | 0.986 | 2.4349516 |
| **SSC4D** | 3.2763 | -0.912909056729712 | 0.117 | 0.59 | 5.3284091 |
| **FGFR4** | 6.0374 | 0.894743606697771 | 0.981 | 0.819 | 9.8187729 |
| **MEIOB** | 6.3851 | 0.884081814045914 | 0.461 | 0.003 | 1.0384223 |
| **SLC25A22** | 7.8723 | 0.823445832344428 | 0.986 | 0.869 | 1.2802802 |
| **OLFM1** | 7.9653 | -1.30276115968908 | 0.424 | 0.81 | 1.2954003 |
| **UBE2D1** | 4.0370 | -0.901420121776708 | 0.546 | 0.873 | 6.5654664 |
| **CADM4** | 4.6527 | -1.08950680317702 | 0.14 | 0.605 | 7.5667856 |
| **ACKR3** | 4.9318 | 1.03918533881333 | 0.822 | 0.351 | 8.0205950 |
| **RAP1GDS1** | 5.7321 | -0.820870332494126 | 0.84 | 0.983 | 9.3221708 |
| **DTNA** | 6.0904 | -1.41087190616934 | 0.02 | 0.443 | 9.9048328 |
| **MCUB** | 8.9048 | 0.967893848673956 | 0.86 | 0.484 | 1.4481980 |
| **KRR1** | 2.1911 | 0.852658778885769 | 0.945 | 0.708 | 3.5635290 |
| **FOXJ1** | 2.4074 | -1.05078077728163 | 0.028 | 0.457 | 3.9152089 |
| **MARCH8** | 6.6682 | 0.91055264459129 | 0.964 | 0.772 | 1.0844545 |
| **CDH1** | 1.0421 | 0.820378366525181 | 1 | 0.868 | 1.6947920 |
| **MAP7D1** | 1.0521 | -1.63876036977146 | 0.453 | 0.808 | 1.7111032 |
| **GCA** | 2.1053 | -1.02960433060554 | 0.363 | 0.787 | 3.4239285 |
| **LHX5** | 6.5081 | -1.77110789623801 | 0.005 | 0.412 | 1.0584199 |
| **ANKRD50** | 1.0328 | -0.958338553495182 | 0.673 | 0.929 | 1.6797462 |

| | | | | | |
|---|---|---|---|---|---|
| **FBRS** | 1.4115 | 1.0397469736371 | 0.956 | 0.739 | 2.2955914 |
| **PRUNE2** | 2.0842 | 0.971904842482802 | 0.973 | 0.778 | 3.3896080 |
| **RBP7** | 6.1596 | 0.867672643991144 | 0.558 | 0.068 | 1.0017416 |
| **GLI3** | 8.6953 | -2.72547229424335 | 0.03 | 0.455 | 1.4141317 |
| **NELFE** | 1.4614 | 0.875944769260941 | 0.787 | 0.317 | 2.3766819 |
| **SELENOM** | 1.5185 | 0.961674552266447 | 0.879 | 0.502 | 2.4695629 |
| **PON2** | 4.5306 | -0.956065969120556 | 0.738 | 0.943 | 7.3682680 |
| **PTPRG** | 8.5298 | -1.14852674890567 | 0.682 | 0.933 | 1.3872036 |
| **ERGIC1** | 1.1347 | -0.837205851196712 | 0.755 | 0.953 | 1.8453924 |
| **PHTF1** | 1.4258 | -1.04087970899871 | 0.634 | 0.912 | 2.3188945 |
| **CFAP36** | 4.1518 | 0.891135034259329 | 0.777 | 0.301 | 6.7520984 |
| **ILK** | 4.4501 | -0.903268258004278 | 0.851 | 0.971 | 7.2373173 |
| **TMEM45A** | 4.5519 | -1.06909419685587 | 0.213 | 0.683 | 7.4028102 |
| **ARSI** | 5.1594 | 0.938906426119458 | 0.926 | 0.63 | 8.3908277 |
| **LZTS2** | 7.2449 | -0.997000094436554 | 0.777 | 0.951 | 1.1782446 |
| **GMPPA** | 1.0376 | 0.872994939758043 | 0.967 | 0.836 | 1.6875619 |
| **TSKU** | 1.0480 | -0.844196077006028 | 0.925 | 0.992 | 1.7044430 |
| **MTRES1** | 1.1301 | 0.874884926050061 | 0.748 | 0.259 | 1.8379319 |
| **FZD3** | 1.4331 | -1.00724885330617 | 0.898 | 0.987 | 2.3307946 |
| **FNDC3A** | 1.6071 | -0.913047059316775 | 0.852 | 0.983 | 2.6137207 |
| **ANKRD33B** | 1.6537 | -1.26761295770571 | 0.08 | 0.531 | 2.6895746 |
| **ZNF300** | 1.9202 | 0.917877983081807 | 0.779 | 0.287 | 3.1228999 |
| **SRMS** | 1.9692 | 0.856327665789073 | 0.504 | 0.031 | 3.2025874 |
| **ZNF618** | 3.1471 | -0.942879467079264 | 0.765 | 0.945 | 5.1182896 |
| **GREM2** | 1.9712 | 1.0833832024514 | 0.755 | 0.253 | 3.2059010 |
| **GPR160** | 2.4504 | 1.13137862405475 | 0.772 | 0.303 | 3.9851771 |
| **CLN8** | 2.7133 | 0.883998980142425 | 0.944 | 0.698 | 4.4126959 |
| **CHKA** | 3.6372 | -0.947386793210469 | 0.802 | 0.962 | 5.9151904 |
| **SPATA22** | 6.9065 | 0.808390306753946 | 0.455 | 0.005 | 1.1232096 |
| **MCAM** | 8.1636 | -1.45497620584866 | 0.456 | 0.82 | 1.3276564 |
| **PCDHB16** | 1.4538 | -0.8381426694582 | 0.053 | 0.492 | 2.3643602 |
| **RIC8B** | 1.6414 | -1.20283174319757 | 0.431 | 0.801 | 2.6694094 |
| **ALPP** | 3.9765 | 0.926058846614985 | 0.501 | 0.032 | 6.4670204 |
| **RGS3** | 4.3726 | -1.01053355841748 | 0.039 | 0.471 | 7.1113159 |

| | | | | | |
|---|---|---|---|---|---|
| **STRA6** | 4.6116 | -1.09226431140631 | 0.127 | 0.587 | 7.4998786 |
| **MAPK12** | 1.4059 | 0.886535412754452 | 0.883 | 0.438 | 2.2865507 |
| **CHD6** | 1.5183 | -0.908951794715849 | 0.781 | 0.952 | 2.4693465 |
| **TBX22** | 1.7077 | -1.1166093231811 | 0.007 | 0.408 | 2.7772409 |
| **LARGE2** | 2.8498 | 0.870709447938892 | 0.996 | 0.887 | 4.6346322 |
| **PLEKHG1** | 3.8946 | 0.904320424863832 | 0.62 | 0.121 | 6.3338438 |
| **HIPK1** | 1.4061 | -0.805479671028863 | 0.861 | 0.983 | 2.2868609 |
| **STARD10** | 1.6669 | 0.961173308974635 | 0.913 | 0.583 | 2.7110406 |
| **FBP1** | 2.2435 | 1.13082799480866 | 0.586 | 0.102 | 3.6486465 |
| **SLC2A4RG** | 7.7363 | -0.965870882639647 | 0.857 | 0.97 | 1.2581603 |
| **HRK** | 9.8643 | 1.03089736479743 | 0.933 | 0.63 | 1.6042315 |
| **THAP12** | 1.6385 | 0.876147536477046 | 0.839 | 0.421 | 2.6648067 |
| **PLEKHA5** | 4.8919 | -0.999720675190677 | 0.68 | 0.929 | 7.9557823 |
| **OSBPL6** | 5.5467 | -0.896887222192327 | 0.099 | 0.553 | 9.0206339 |
| **DNMBP** | 6.9771 | -1.00355487401266 | 0.716 | 0.934 | 1.1346924 |
| **RGS2** | 8.9740 | 1.14096564107763 | 0.885 | 0.535 | 1.4594496 |
| **NEK6** | 1.2551 | 0.811112801250153 | 0.994 | 0.948 | 2.0413051 |
| **SPRY4** | 3.0890 | -1.00684482584501 | 0.02 | 0.428 | 5.0237843 |
| **SH2D3A** | 2.0749 | 0.902270255163541 | 0.847 | 0.381 | 3.3744815 |
| **TRIM44** | 2.6803 | -0.875242846619145 | 0.792 | 0.963 | 4.3590344 |
| **KLF17** | 3.8215 | 1.41236643320943 | 0.454 | 0.009 | 6.2149513 |
| **FGD1** | 6.6148 | -0.835579445413658 | 0.853 | 0.987 | 1.0757757 |
| **MAP3K13** | 1.0717 | 0.948133667848393 | 0.895 | 0.55 | 1.7430110 |
| **IFI27L2** | 1.2957 | -1.08435617545227 | 0.193 | 0.645 | 2.1072191 |
| **SDC1** | 1.3417 | -1.04233562387538 | 0.079 | 0.521 | 2.1821276 |
| **TGFB3** | 1.5617 | 0.904245321657543 | 0.633 | 0.131 | 2.5398791 |
| **KIDINS220** | 7.1447 | -0.841374580121407 | 0.858 | 0.975 | 1.1619505 |
| **FCGBP** | 1.1376 | 0.982389462839461 | 0.524 | 0.054 | 1.8502019 |
| **RTL8A** | 1.2493 | -0.991130210020641 | 0.321 | 0.742 | 2.0318726 |
| **ARHGDIG** | 1.4545 | 0.829968792403597 | 0.607 | 0.118 | 2.3655642 |
| **STON2** | 1.5218 | -1.35190668155997 | 0.298 | 0.721 | 2.4750340 |
| **PTPN11** | 2.3897 | 0.830596780893697 | 0.902 | 0.55 | 3.8863928 |
| **NSMCE4A** | 8.7242 | -0.869115787869185 | 0.884 | 0.978 | 1.4188314 |
| **ADCK2** | 9.8901 | 0.870698152555329 | 0.91 | 0.567 | 1.6084387 |

| | | | | | |
|---|---|---|---|---|---|
| **C11orf95** | 3.4096 | -1.04773310262168 | 0.617 | 0.9 | 5.5451257 |
| **PTPRA** | 5.8675 | -0.808264654373761 | 0.903 | 0.983 | 9.5423769 |
| **PLA2G2A** | 6.7271 | -0.939588818082565 | 0.004 | 0.391 | 1.0940423 |
| **ARFRP1** | 1.6386 | -0.815529481038765 | 0.877 | 0.988 | 2.6648661 |
| **PLEKHG5** | 2.6420 | 0.858766567166854 | 0.72 | 0.231 | 4.2967179 |
| **SLC12A8** | 5.7613 | -1.07945592557899 | 0.158 | 0.603 | 9.3696290 |
| **HDAC7** | 3.3237 | -0.988258855667122 | 0.682 | 0.916 | 5.4054143 |
| **ANK2** | 7.4621 | -1.01555812239854 | 0.661 | 0.916 | 1.2135753 |
| **LAMA2** | 1.7112 | -1.42679607987077 | 0.265 | 0.662 | 2.7830675 |
| **PIPOX** | 1.8838 | -1.13261070720695 | 0.201 | 0.646 | 3.0636294 |
| **KCNMB1** | 1.9523 | -1.00231749398872 | 0.009 | 0.397 | 3.1751844 |
| **SHLD2** | 2.3757 | 0.866680044102289 | 0.888 | 0.528 | 3.8637109 |
| **PIK3R5** | 3.3111 | 0.892404109023992 | 0.645 | 0.166 | 5.3849563 |
| **VANGL1** | 5.7328 | -0.990859098768259 | 0.65 | 0.906 | 9.3233579 |
| **WBP1L** | 7.8649 | -0.853004766435232 | 0.902 | 0.984 | 1.2790798 |
| **KLHL15** | 8.2660 | -0.960528601088106 | 0.416 | 0.809 | 1.3443065 |
| **FLNC** | 3.4396 | -1.97127309849733 | 0.51 | 0.834 | 5.5939012 |
| **NRN1L** | 1.3902 | -1.30532756784032 | 0.089 | 0.523 | 2.2608851 |
| **PACS1** | 1.7914 | -0.898992532755558 | 0.89 | 0.977 | 2.9133785 |
| **KRT19** | 1.8562 | -1.07606582910832 | 0.805 | 0.966 | 3.0188493 |
| **SH3BGRL3** | 2.7020 | -1.00053869094229 | 0.958 | 0.993 | 4.3943011 |
| **C19orf85** | 3.7574 | 1.22814226010046 | 0.676 | 0.202 | 6.1107691 |
| **NFE2L3** | 6.6702 | 0.959294824870341 | 0.786 | 0.336 | 1.0847874 |
| **ACP2** | 7.4873 | -0.918660603934577 | 0.732 | 0.946 | 1.2176665 |
| **DLG1** | 8.9668 | -0.805306534373047 | 0.92 | 0.986 | 1.4582753 |
| **RND3** | 9.1693 | -1.19538157795537 | 0.078 | 0.505 | 1.4912182 |
| **CASP10** | 1.1230 | 0.901306357853249 | 0.774 | 0.319 | 1.8264005 |
| **CNN1** | 1.1233 | -2.10746685644706 | 0.267 | 0.684 | 1.8268414 |
| **HDHD2** | 1.4348 | -0.801858755405998 | 0.812 | 0.966 | 2.3335084 |
| **OAT** | 1.8071 | 0.952789252476512 | 0.872 | 0.504 | 2.9388909 |
| **GALC** | 2.0144 | -0.976193659467246 | 0.511 | 0.84 | 3.2761103 |
| **TAF9B** | 5.6292 | -0.92968769830669 | 0.475 | 0.82 | 9.1548750 |
| **ELMO3** | 8.1055 | 0.837489004848713 | 0.668 | 0.178 | 1.3182110 |
| **SERPINB9** | 1.4531 | 1.03149081203347 | 1 | 0.929 | 2.3632765 |

| | | | | | |
|---|---|---|---|---|---|
| **RASSF9** | 4.2239 | -1.07651237381389 | 0.041 | 0.449 | 6.8694607 |
| **MUC4** | 8.0382 | 0.934457169285889 | 0.895 | 0.526 | 1.3072639 |
| **ZNF695** | 2.0382 | 0.821576741201494 | 0.962 | 0.755 | 3.3147586 |
| **ZKSCAN1** | 2.1644 | -0.986880737019283 | 0.551 | 0.853 | 3.5199943 |
| **SPR** | 2.4924 | 0.903908757834744 | 0.851 | 0.427 | 4.0534732 |
| **VASH2** | 1.6817 | 0.836342290345784 | 0.991 | 0.893 | 2.7350432 |
| **MSRB2** | 2.1801 | -0.882339382269061 | 0.752 | 0.948 | 3.5455641 |
| **C1orf54** | 3.1552 | -0.80787561376801 | 0.11 | 0.553 | 5.1314190 |
| **LZTS1** | 5.2192 | 0.844866421854244 | 0.878 | 0.496 | 8.4881172 |
| **TDRD1** | 7.4176 | 0.945618041813102 | 0.456 | 0.02 | 1.2063364 |
| **PARVA** | 1.4321 | -1.04500668983821 | 0.709 | 0.963 | 2.3290539 |
| **BRSK1** | 3.1370 | -0.995976939546949 | 0.135 | 0.568 | 5.1018144 |
| **SMG6** | 7.4827 | -0.912343144504129 | 0.539 | 0.844 | 1.2169242 |
| **ZNF260** | 8.3281 | 0.853785912166637 | 0.842 | 0.441 | 1.3544029 |
| **CARMIL1** | 1.7693 | -1.04540196703566 | 0.37 | 0.774 | 2.8775415 |
| **CSRP1** | 2.1859 | -2.05551631679124 | 0.838 | 0.96 | 3.5550914 |
| **VASP** | 2.3191 | -0.834189993310317 | 0.999 | 0.999 | 3.7715661 |
| **SLC35F1** | 2.6373 | -0.990570755799288 | 0.238 | 0.671 | 4.2890780 |
| **NFATC4** | 3.1260 | -1.16031654537686 | 0.208 | 0.641 | 5.0838892 |
| **ELF4** | 3.6482 | -1.34573532729128 | 0.145 | 0.571 | 5.9332197 |
| **GALNT6** | 5.5463 | 0.832195235877619 | 0.931 | 0.644 | 9.0200031 |
| **PPP1R1A** | 7.6649 | -1.25145823377022 | 0.596 | 0.883 | 1.2465453 |
| **FAM217B** | 1.0136 | -0.836272996592946 | 0.703 | 0.936 | 1.6484658 |
| **THY1** | 1.8082 | -1.08924592633415 | 0.03 | 0.422 | 2.9407785 |
| **TMEM126B** | 2.3357 | 0.820226060462014 | 0.763 | 0.318 | 3.7986362 |
| **ADGRL2** | 4.1215 | -0.921024064382754 | 0.894 | 0.992 | 6.7028798 |
| **ZNF609** | 4.2672 | -0.858605310483334 | 0.687 | 0.916 | 6.9398823 |
| **IGFBP5** | 5.0910 | -4.04173546934023 | 0.109 | 0.532 | 8.2796225 |
| **IRS2** | 6.9688 | -1.18493490018935 | 0.273 | 0.693 | 1.1333384 |
| **PHLDA2** | 8.3422 | 1.05797044787379 | 0.904 | 0.607 | 1.3567076 |
| **ISYNA1** | 2.4832 | -0.92553679493028 | 0.984 | 0.997 | 4.0384765 |
| **HHIPL1** | 2.9838 | -1.03289971357184 | 0.236 | 0.66 | 4.8525765 |
| **HNMT** | 3.2468 | -0.878894083151258 | 0.137 | 0.572 | 5.2803281 |
| **KIF26A** | 3.8680 | -1.04350992520749 | 0.877 | 0.966 | 6.2906120 |

| | | | | | |
|---|---|---|---|---|---|
| **KCNG3** | 2.5887 | 0.961775848806386 | 0.77 | 0.318 | 4.21008825 |
| **NAT14** | 5.9530 | -0.850540872835167 | 0.905 | 0.987 | 9.68136988 |
| **JADE1** | 1.1609 | -0.808202496689236 | 0.881 | 0.983 | 1.88806691 |
| **CDYL** | 6.7097 | -0.830788370455757 | 0.825 | 0.972 | 1.09121342 |
| **CDCP1** | 8.7665 | 0.923223631253225 | 0.797 | 0.359 | 1.42570411 |
| **SDF2L1** | 2.0369 | 1.32515504318812 | 0.985 | 0.976 | 3.31276878 |
| **BMP4** | 2.4613 | -1.33021319722188 | 0.523 | 0.854 | 4.00294275 |
| **CACNA2D1** | 8.3660 | 0.83247571084214 | 0.964 | 0.784 | 1.36056260 |
| **HEG1** | 1.3155 | -1.15084068181381 | 0.251 | 0.671 | 2.13943365 |
| **PMEL** | 1.7247 | -1.13274950265769 | 0.036 | 0.427 | 2.80488990 |
| **MAP3K3** | 1.8457 | -0.918234397134426 | 0.531 | 0.852 | 3.00169139 |
| **ADGRE5** | 3.8260 | 0.856808701348621 | 0.855 | 0.473 | 6.22234390 |
| **RHBDL3** | 4.1768 | -0.978749145888819 | 0.114 | 0.538 | 6.79286250 |
| **SLC25A15** | 5.3449 | -0.849755947257912 | 0.707 | 0.931 | 8.69255408 |
| **SH3YL1** | 6.1092 | 0.803492434991647 | 0.64 | 0.173 | 9.93545351 |
| **S100A16** | 7.3784 | 1.27674241487607 | 0.706 | 0.26 | 1.19995769 |
| **GAS1** | 7.9673 | -0.932923079840329 | 0.184 | 0.621 | 1.29573221 |
| **MANEAL** | 8.3301 | 0.879379668059171 | 0.961 | 0.758 | 1.35472645 |
| **PCNX4** | 8.7173 | -1.02975254677703 | 0.488 | 0.815 | 1.41769660 |
| **NUAK1** | 2.8185 | -1.01003131818592 | 0.505 | 0.838 | 4.58373038 |
| **ENC1** | 4.4961 | -2.25097590620896 | 0.1 | 0.516 | 7.31209258 |
| **CADM1** | 1.0494 | -1.20728199217076 | 0.579 | 0.867 | 1.70665297 |
| **NTS** | 1.3644 | -1.48406943861305 | 0.108 | 0.529 | 2.21901020 |
| **BMI1** | 1.5910 | 0.942458493996859 | 0.807 | 0.415 | 2.58750650 |
| **NCEH1** | 1.7681 | 0.892808259430138 | 0.811 | 0.39 | 2.87552630 |
| **GRPR** | 4.5821 | -1.02365110443225 | 0.012 | 0.379 | 7.45189672 |
| **APOE** | 7.4503 | -0.867652620409396 | 1 | 1 | 1.21165068 |
| **STOX1** | 2.0546 | -0.824390076711092 | 0.055 | 0.456 | 3.34145642 |
| **ZNF316** | 6.0248 | -0.868421676758997 | 0.303 | 0.712 | 9.79822803 |
| **EEF1AKMT4** | 9.6159 | 0.800883151026602 | 0.951 | 0.753 | 1.56383722 |
| **DHRS7** | 1.4847 | -0.884879266077847 | 0.754 | 0.952 | 2.41470314 |
| **DPP10** | 6.6014 | -1.14494836864095 | 0.31 | 0.694 | 1.07359792 |
| **RTN4RL1** | 1.1415 | -1.07591494873219 | 0.308 | 0.705 | 1.85643068 |
| **SEMA3B** | 1.9942 | -0.837295638400805 | 0.1 | 0.519 | 3.24325179 |

| | | | | | |
|---|---|---|---|---|---|
| **UHRF2** | 2.2062 | -1.05856147910615 | 0.311 | 0.706 | 3.5880965 |
| **AQP3** | 2.8308 | 0.986625398568336 | 0.611 | 0.156 | 4.6037304 |
| **MPV17L** | 4.0341 | 0.831352099014997 | 0.708 | 0.248 | 6.5608001 |
| **NSG1** | 4.6150 | 0.80258706528922 | 0.536 | 0.086 | 7.5055201 |
| **ZNF354A** | 4.2912 | 0.83173830505455 | 0.833 | 0.449 | 6.9789181 |
| **PALLD** | 4.3867 | -1.75447070119679 | 0.253 | 0.666 | 7.1342524 |
| **TICRR** | 1.2764 | 0.859411517766091 | 0.989 | 0.893 | 2.0759390 |
| **AGBL5** | 1.4132 | -0.915368953805451 | 0.661 | 0.901 | 2.2983370 |
| **CDH13** | 1.5607 | -1.3330781061976 | 0.157 | 0.574 | 2.5383130 |
| **MAN1A1** | 2.7868 | -1.04417634866103 | 0.107 | 0.523 | 4.5322774 |
| **PTPRS** | 3.6920 | -1.09207479430056 | 0.989 | 0.999 | 6.0044397 |
| **FIBCD1** | 6.7343 | -1.25014942732008 | 0.375 | 0.731 | 1.0952060 |
| **FOXI3** | 8.6381 | -1.18744350052354 | 0.964 | 0.972 | 1.4048285 |
| **ZNF682** | 8.6908 | 0.83962260270242 | 0.822 | 0.424 | 1.4133979 |
| **C14orf132** | 9.2453 | -1.22154615670257 | 0.09 | 0.495 | 1.5035689 |
| **HABP4** | 1.7544 | -1.00849613664058 | 0.251 | 0.667 | 2.8533404 |
| **PSRC1** | 3.6743 | 0.946853856930907 | 0.974 | 0.812 | 5.9756591 |
| **CXCL14** | 7.7052 | -2.1438486081555 | 0.277 | 0.678 | 1.2530986 |
| **RAB6B** | 1.7803 | -1.14785214727787 | 0.499 | 0.826 | 2.8953911 |
| **SH2D4A** | 9.2295 | 0.876793524640979 | 0.861 | 0.487 | 1.5009969 |
| **ARHGEF2** | 1.2090 | -0.881177426719902 | 0.709 | 0.928 | 1.9663445 |
| **KIF9** | 1.2167 | 0.832558661742094 | 0.932 | 0.677 | 1.9787385 |
| **STK40** | 1.2973 | -0.910906167839329 | 0.619 | 0.877 | 2.1098182 |
| **SLC7A6** | 2.3731 | 0.859716623127861 | 0.895 | 0.588 | 3.8594077 |
| **TMEM154** | 1.3856 | 0.925447237085282 | 0.733 | 0.276 | 2.2534820 |
| **TSPEAR** | 1.7315 | -0.87172363724136 | 0.003 | 0.349 | 2.8159706 |
| **ARHGAP42** | 1.3965 | 0.82145566677353 | 0.905 | 0.636 | 2.2711836 |
| **SIPA1L2** | 2.0671 | -0.956313836168166 | 0.724 | 0.938 | 3.3618748 |
| **MID1** | 3.9634 | -0.966943876057015 | 0.698 | 0.927 | 6.4457670 |
| **APOBEC3G** | 8.6756 | 0.869768314231224 | 0.576 | 0.126 | 1.4109144 |
| **ANGPTL2** | 1.0295 | -0.926213256094056 | 0.028 | 0.397 | 1.6743097 |
| **TTL** | 1.7144 | -0.80464644271781 | 0.875 | 0.97 | 2.7882656 |
| **SPTLC3** | 7.3696 | -0.817625958598351 | 0.015 | 0.371 | 1.1985339 |
| **SPON1** | 1.0387 | -1.76091543777622 | 0.007 | 0.353 | 1.6892448 |

| | | | | | |
|---|---|---|---|---|---|
| **PTN** | 1.9797 | -1.4044582307123 | 0.027 | 0.392 | 3.21970838 |
| **CCNE1** | 3.2858 | 0.955304455637956 | 0.949 | 0.728 | 5.34370589 |
| **NEU1** | 3.9731 | -0.981961622112101 | 0.962 | 0.991 | 6.46155983 |
| **HIPK2** | 6.0583 | -1.05063669898495 | 0.632 | 0.871 | 9.85271817 |
| **TEAD1** | 6.7786 | -0.903677790262168 | 0.59 | 0.862 | 1.10240640 |
| **FAM104B** | 1.6340 | -0.815960833054268 | 0.642 | 0.903 | 2.65746747 |
| **STOM** | 3.4605 | 0.863720303728574 | 0.99 | 0.903 | 5.62785514 |
| **IDH2** | 1.3040 | -0.912463016513587 | 0.931 | 0.991 | 2.12074395 |
| **KRTCAP3** | 1.9801 | 0.873710768857382 | 0.747 | 0.313 | 3.22024048 |
| **BRINP1** | 3.6407 | -1.15475003447732 | 0.152 | 0.55 | 5.92098691 |
| **C1orf198** | 4.2205 | -1.13871781119702 | 0.674 | 0.899 | 6.86380329 |
| **GALNT2** | 5.6194 | -0.948927266953776 | 0.92 | 0.98 | 9.13895489 |
| **LAMB2** | 7.1678 | -1.17204529915954 | 0.77 | 0.931 | 1.16570080 |
| **ADCY3** | 1.0560 | 0.814021325687749 | 0.741 | 0.289 | 1.71750510 |
| **SIX5** | 1.1221 | -0.925062804454464 | 0.586 | 0.847 | 1.82489619 |
| **CCPG1** | 1.4079 | 0.861097495155374 | 0.942 | 0.776 | 2.28982689 |
| **FAM20C** | 4.1817 | -1.00250236037305 | 0.593 | 0.88 | 6.80076932 |
| **DPYSL5** | 9.9696 | -1.97506057310005 | 0.027 | 0.386 | 1.62136687 |
| **CERS4** | 1.4139 | -1.05052639556336 | 0.954 | 0.992 | 2.29954875 |
| **ARRB1** | 2.1133 | 0.888611720714015 | 0.791 | 0.37 | 3.43702077 |
| **SNX7** | 5.5591 | 0.812142746247738 | 0.837 | 0.47 | 9.04085110 |
| **CKAP2** | 6.3452 | -0.898780138940292 | 0.964 | 0.997 | 1.03192032 |
| **RFX3** | 1.1545 | -0.936610701551299 | 0.302 | 0.697 | 1.87769355 |
| **LRRC42** | 7.1303 | -0.811313214242974 | 0.772 | 0.933 | 1.15960527 |
| **SH3GLB2** | 1.0325 | -0.81443582077759 | 0.766 | 0.948 | 1.67920449 |
| **USP51** | 2.5703 | -0.889889576222798 | 0.363 | 0.741 | 4.18008017 |
| **EXT1** | 3.2076 | -0.916642934896805 | 0.881 | 0.98 | 5.21654271 |
| **MND1** | 4.7281 | 0.969471937311148 | 0.942 | 0.83 | 7.68932496 |
| **SLITRK2** | 6.9448 | 0.801476489896925 | 0.388 | 0.007 | 1.12944710 |
| **MT-ND3** | 2.0912 | 1.34208022299826 | 0.952 | 0.806 | 3.40103730 |
| **SLC39A6** | 5.9479 | -0.974183609717654 | 0.563 | 0.829 | 9.67309879 |
| **CCDC69** | 8.2864 | 0.8204881785993 | 0.817 | 0.436 | 1.34762967 |
| **ADD2** | 1.0325 | -0.916019302069501 | 0.852 | 0.973 | 1.67919621 |
| **GRHL1** | 3.1902 | 0.815767923829265 | 0.691 | 0.25 | 5.18830322 |

| | | | | | |
|---|---|---|---|---|---|
| **SIK2** | 5.2129 | -0.851117780671401 | 0.438 | 0.788 | 8.4778290 |
| **DUSP7** | 9.9549 | 0.846585423749002 | 0.891 | 0.607 | 1.6189661 |
| **IRAK1** | 1.1956 | 0.890936968980561 | 0.845 | 0.511 | 1.9445348 |
| **RARA** | 3.9256 | -0.90943372824182 | 0.22 | 0.62 | 6.3842450 |
| **GYG2** | 4.2859 | -0.842945932325653 | 0.718 | 0.924 | 6.9701708 |
| **MAP2** | 1.5160 | -1.78821076838344 | 0.007 | 0.338 | 2.4655277 |
| **KCNC3** | 2.5311 | 0.86480792261241 | 0.705 | 0.274 | 4.1164593 |
| **ZFP36L1** | 4.0820 | -0.853175171084155 | 0.989 | 0.999 | 6.6387021 |
| **SLC43A2** | 6.7421 | -0.906229641283904 | 0.805 | 0.952 | 1.0964703 |
| **ZNF613** | 9.9368 | 0.824564015023592 | 0.855 | 0.529 | 1.6160318 |
| **LRFN5** | 9.9915 | 0.803376755628827 | 0.464 | 0.056 | 1.6249214 |
| **GINS1** | 1.7494 | -0.864104024155174 | 0.988 | 0.996 | 2.8451893 |
| **SPIN3** | 7.1697 | -1.00761260329615 | 0.417 | 0.771 | 1.1660084 |
| **EGLN1** | 1.3076 | -0.806802811829146 | 0.643 | 0.917 | 2.1266748 |
| **FOS** | 1.6674 | -1.29239852103569 | 0.288 | 0.693 | 2.7117652 |
| **IKBIP** | 3.8742 | -0.869898878648002 | 0.416 | 0.765 | 6.3006532 |
| **RBCK1** | 5.4026 | -0.86335323225548 | 0.658 | 0.886 | 8.7863664 |
| **PCBD1** | 1.3420 | -0.809236260298634 | 0.826 | 0.956 | 2.1825413 |
| **ZSWIM9** | 1.5571 | -0.849152627638655 | 0.268 | 0.664 | 2.5323786 |
| **ULK4** | 5.2972 | 0.834080561422738 | 0.934 | 0.713 | 8.6148480 |
| **CENPA** | 1.0751 | 0.994186969244967 | 0.977 | 0.851 | 1.7485639 |
| **ARHGEF10L** | 2.2658 | -0.872339349357462 | 0.312 | 0.692 | 3.6849567 |
| **PPP1R9B** | 3.7353 | -0.957165704419594 | 0.449 | 0.77 | 6.0748747 |
| **ARHGAP6** | 4.5534 | -0.956343950040758 | 0.206 | 0.601 | 7.4053309 |
| **DERL3** | 6.3572 | 0.846697877388227 | 0.489 | 0.08 | 1.0338745 |
| **AJAP1** | 2.3973 | -0.897805797957712 | 0.153 | 0.546 | 3.8988330 |
| **P3H2** | 2.6221 | -0.913773549938975 | 0.006 | 0.33 | 4.2644659 |
| **PPP1R18** | 5.7275 | -1.12433941298404 | 0.309 | 0.691 | 9.3147672 |
| **APBB2** | 1.0417 | -0.965668486114097 | 0.284 | 0.663 | 1.6942417 |
| **RGMA** | 2.7476 | -1.41586442010437 | 0.492 | 0.779 | 4.4685206 |
| **CAVIN3** | 3.4134 | -0.988087427478616 | 0.358 | 0.739 | 5.5513262 |
| **ZRANB3** | 3.9967 | -0.853455468891797 | 0.354 | 0.719 | 6.4998940 |
| **AIRE** | 1.0016 | 0.835965074992858 | 0.559 | 0.132 | 1.6290271 |
| **MT-ND1** | 1.4034 | -1.09876523975403 | 0.733 | 0.956 | 2.2823735 |

| | | | | | |
|---|---|---|---|---|---|
| **MTCL1** | 1.7764 | -0.893963314290034 | 0.602 | 0.866 | 2.88909791 |
| **CNKSR2** | 1.8727 | -0.858423344477537 | 0.116 | 0.507 | 3.04558245 |
| **HOMER2** | 3.9817 | 0.896748720849741 | 0.941 | 0.774 | 6.47546048 |
| **FBXL13** | 9.8616 | 0.873319534308216 | 0.661 | 0.239 | 1.60379598 |
| **MYH9** | 1.0415 | -0.81513246388198 | 0.993 | 0.999 | 1.69390373 |
| **NET1** | 6.8829 | -0.802858384391521 | 0.74 | 0.913 | 1.11937296 |
| **GPC3** | 1.6934 | -1.32314544953016 | 0.761 | 0.942 | 2.75404533 |
| **NME7** | 3.0265 | -0.801971758876454 | 0.296 | 0.676 | 4.92213581 |
| **CNMD** | 4.5767 | 0.964082058253839 | 0.937 | 0.698 | 7.44320840 |
| **P3H1** | 5.7941 | -0.914118356918615 | 0.424 | 0.744 | 9.42299232 |
| **TENM4** | 1.1180 | -1.33038843158544 | 0.416 | 0.764 | 1.81831744 |
| **CDKN2AIP** | 1.8746 | -0.804253089990213 | 0.768 | 0.932 | 3.04874471 |
| **RNF135** | 2.1060 | 0.801997460055956 | 0.852 | 0.555 | 3.42509157 |
| **PCDH8** | 3.2197 | -0.908750072111608 | 0.071 | 0.437 | 5.23629113 |
| **TSC22D3** | 5.1273 | -1.05510638124156 | 0.398 | 0.757 | 8.33855478 |
| **TMEM37** | 6.0843 | 0.860337854236155 | 0.97 | 0.822 | 9.89497308 |
| **KATNAL1** | 1.0075 | -0.821187028799997 | 0.4 | 0.751 | 1.63855760 |
| **EZR** | 1.1417 | -1.08977825393766 | 1 | 1 | 1.85675351 |
| **CA2** | 3.1142 | 0.945873395615299 | 0.949 | 0.845 | 5.06471610 |
| **MX1** | 2.2445 | 0.921437386669453 | 0.373 | 0.013 | 3.65025459 |
| **ARX** | 4.2244 | 0.940480343067601 | 0.393 | 0.025 | 6.87018795 |
| **MMD** | 1.3784 | -0.80676154267219 | 0.586 | 0.843 | 2.24184498 |
| **FAM13A** | 2.5966 | -0.842678930057913 | 0.434 | 0.776 | 4.22294567 |
| **MYORG** | 7.0031 | -0.811137358931627 | 0.16 | 0.541 | 1.13892069 |
| **SLC5A3** | 1.3272 | -1.10409182230308 | 0.617 | 0.857 | 2.15857110 |
| **NRXN1** | 1.8112 | -0.800034027617045 | 0.116 | 0.502 | 2.94563073 |
| **SHROOM3** | 2.3660 | -0.946831987684908 | 0.742 | 0.93 | 3.84790930 |
| **TMEM47** | 1.0349 | -0.834059785044793 | 0.165 | 0.548 | 1.68306224 |
| **TUNAR** | 1.0397 | -1.03389099681876 | 0.355 | 0.715 | 1.69102080 |
| **AMOT** | 1.1989 | -1.20701168484457 | 0.105 | 0.475 | 1.94984521 |
| **ADGRG2** | 1.4091 | 0.815129755646914 | 0.802 | 0.411 | 2.29163144 |
| **ZBED5** | 1.5181 | -0.814973239308722 | 0.554 | 0.833 | 2.46896223 |
| **BEND4** | 3.3283 | -0.959515960624515 | 0.846 | 0.914 | 5.41291949 |
| **DDR1** | 4.2829 | -1.01354399799691 | 0.89 | 0.977 | 6.96542275 |

| | | | | | |
|---|---|---|---|---|---|
| **AMH** | 5.3069 | -1.11808262725494 | 0.185 | 0.559 | 8.6306722 |
| **EBF2** | 6.7253 | -1.18673471508849 | 0.124 | 0.489 | 1.0937395 |
| **CHST6** | 1.0053 | -0.891018414416137 | 0.477 | 0.767 | 1.6350171 |
| **COTL1** | 1.4370 | -0.973775380519122 | 0.949 | 0.981 | 2.3370958 |
| **PTGIS** | 1.5111 | -1.18466535597341 | 0.095 | 0.454 | 2.4575487 |
| **VIT** | 1.7059 | -1.3403573530604 | 0.019 | 0.34 | 2.7744238 |
| **CDON** | 2.6695 | -1.11160348658327 | 0.256 | 0.631 | 4.3415203 |
| **PLPP2** | 3.2492 | 0.941298354524284 | 0.474 | 0.086 | 5.2842841 |
| **PAX3** | 5.6208 | -1.38004528197581 | 0.002 | 0.3 | 9.1412437 |
| **NCS1** | 5.4230 | -0.868406459981765 | 0.643 | 0.873 | 8.8194760 |
| **PCED1A** | 1.2007 | -1.00506706630903 | 0.382 | 0.721 | 1.9527441 |
| **SRSF8** | 3.0413 | 0.879167270218168 | 0.96 | 0.819 | 4.9461908 |
| **CNKSR3** | 4.6488 | -0.877598279823901 | 0.113 | 0.484 | 7.5603944 |
| **RAD51** | 5.2947 | 0.836072385099953 | 0.968 | 0.878 | 8.6108480 |
| **S100A4** | 6.4992 | -0.90724597856305 | 0.028 | 0.353 | 1.0569773 |
| **LHFPL2** | 8.7406 | -0.852674463729453 | 0.245 | 0.622 | 1.4214908 |
| **GNPAT** | 2.7282 | -0.888681679438046 | 0.595 | 0.847 | 4.4369041 |
| **SLC39A14** | 6.8717 | 0.837760957632767 | 0.851 | 0.586 | 1.1175460 |
| **RIT1** | 9.8454 | -0.804469178866354 | 0.476 | 0.788 | 1.6011586 |
| **INKA2** | 1.5535 | -0.916966533771501 | 0.284 | 0.657 | 2.5264770 |
| **CPE** | 1.4651 | -2.69158261723842 | 0.069 | 0.414 | 2.3827930 |
| **GADL1** | 2.2926 | 1.19337015435881 | 0.421 | 0.05 | 3.7285684 |
| **IGFBP2** | 4.6212 | -1.00369840249803 | 0.998 | 1 | 7.5154772 |
| **RCBTB2** | 8.5280 | -0.840652480454058 | 0.247 | 0.624 | 1.3869167 |
| **CCNA1** | 5.3509 | 1.17730732823271 | 0.491 | 0.105 | 8.7022993 |
| **HSPA5** | 1.8762 | 1.18420456939398 | 0.957 | 0.904 | 3.0513220 |
| **RTN4RL2** | 8.7401 | -0.840698026595169 | 0.15 | 0.517 | 1.4214168 |
| **DLK1** | 1.0984 | -0.857045427050305 | 0.82 | 0.341 | 1.7864667 |
| **SCUBE3** | 2.6173 | -0.936495117738611 | 0.145 | 0.507 | 4.2565176 |
| **HES6** | 2.7195 | -0.900642910490734 | 0.297 | 0.655 | 4.4228015 |
| **PKIG** | 2.7253 | -0.824076703620126 | 0.827 | 0.939 | 4.4322488 |
| **TNNC1** | 3.8021 | -0.856940493710877 | 0.163 | 0.529 | 6.1834869 |
| **LPAR1** | 8.6249 | -0.857216399357604 | 0.095 | 0.45 | 1.4026803 |
| **TYRO3** | 5.9588 | -0.862073500815705 | 0.822 | 0.947 | 9.6909129 |

| | | | | | |
|---|---|---|---|---|---|
| **PCDH7** | 8.7784 | 1.01886319999577 | 0.468 | 0.084 | 1.4276374 |
| **ALDH1B1** | 5.4974 | -0.82920371717622 | 0.82 | 0.923 | 8.9405686 |
| **DYRK1B** | 6.1365 | -1.02056540026013 | 0.454 | 0.73 | 9.9798250 |
| **PMP22** | 8.6666 | -1.26633729668373 | 0.237 | 0.585 | 1.4094625 |
| **IFI16** | 2.8131 | -0.92180920094567 | 0.306 | 0.675 | 4.5749641 |
| **SP8** | 3.8784 | -0.895312973653025 | 0.902 | 0.945 | 6.3075496 |
| **XKR6** | 4.1920 | -0.882936297464153 | 0.7 | 0.896 | 6.8174830 |
| **KIAA1549** | 8.3499 | -0.864587535802227 | 0.337 | 0.675 | 1.3579569 |
| **USP44** | 1.1320 | -0.843673529104064 | 0.197 | 0.559 | 1.8411227 |
| **SYNGAP1** | 6.3822 | -1.18133146124961 | 0.28 | 0.613 | 1.0379469 |
| **FYN** | 8.1063 | -0.966926046376117 | 0.804 | 0.938 | 1.3183377 |
| **ZNF577** | 1.2013 | 0.810203191734665 | 0.753 | 0.392 | 1.9537938 |
| **SYNGR1** | 3.3521 | -0.966874216978639 | 0.355 | 0.676 | 5.4516481 |
| **PPM1H** | 5.6663 | -0.848829334012698 | 0.558 | 0.824 | 9.2151392 |
| **SYNE3** | 8.3310 | -0.82596817141639 | 0.056 | 0.381 | 1.3548778 |
| **NPHP3** | 8.8516 | -0.85824181843788 | 0.4 | 0.718 | 1.4395388 |
| **ROR2** | 1.0317 | -1.1313052701413 | 0.498 | 0.774 | 1.6779286 |
| **FBN3** | 1.8445 | -0.923902998447474 | 0.401 | 0.735 | 2.9998709 |
| **UBE2J1** | 4.0685 | -0.837205591471256 | 0.757 | 0.928 | 6.6166048 |
| **NEO1** | 6.7760 | -0.938475006715383 | 0.546 | 0.797 | 1.1019888 |
| **PCOLCE2** | 7.0279 | -0.887249828966863 | 0.71 | 0.885 | 1.1429476 |
| **CRB2** | 1.3619 | -1.51203806758986 | 0.01 | 0.292 | 2.2149208 |
| **SEMA3C** | 1.4623 | -0.870254677307561 | 0.01 | 0.295 | 2.3781815 |
| **TUBB6** | 2.0942 | -0.84687491282919 | 0.93 | 0.981 | 3.4058072 |
| **TMEFF1** | 4.5903 | -0.985126079555892 | 0.876 | 0.967 | 7.4653092 |
| **IGSF1** | 2.3676 | 0.939725967853435 | 0.862 | 0.668 | 3.8504331 |
| **CCN1** | 3.1543 | -1.44064345486455 | 0.823 | 0.948 | 5.1299140 |
| **SH3D21** | 7.6927 | 0.839095530968515 | 0.745 | 0.383 | 1.2510744 |
| **PAMR1** | 9.8567 | -1.48297750027293 | 0.043 | 0.35 | 1.6029964 |
| **CYP26A1** | 2.5146 | -1.35303943720332 | 0.019 | 0.307 | 4.0894999 |
| **GALNT7** | 6.3441 | -0.923897287598715 | 0.944 | 0.98 | 1.0317480 |
| **LTBP1** | 2.0549 | -0.927375479690438 | 0.648 | 0.852 | 3.3419763 |
| **ARHGAP11A** | 1.0767 | 0.924332407088684 | 0.897 | 0.7 | 1.7511823 |
| **PRICKLE1** | 1.5061 | -1.0390622063539 | 0.552 | 0.81 | 2.4494987 |

| | | | | | |
|---|---|---|---|---|---|
| **SLC1A3** | 5.6868 | -0.898921552157049 | 0.243 | 0.587 | 9.24846559 |
| **BOC** | 7.4254 | -1.85676342289513 | 0.061 | 0.374 | 1.20760297 |
| **COMT** | 2.6636 | -0.892325043505602 | 0.498 | 0.781 | 4.33196750 |
| **ARSE** | 7.2186 | -0.809112479164203 | 0.199 | 0.567 | 1.17396853 |
| **MGAT5** | 8.3041 | -0.893957001893634 | 0.436 | 0.725 | 1.35049652 |
| **PIM1** | 9.7132 | -0.859366096284532 | 0.973 | 0.989 | 1.57965777 |
| **FJX1** | 1.1446 | -1.78758742730523 | 0.128 | 0.457 | 1.86156780 |
| **AGPAT4** | 8.2328 | -0.808747409614199 | 0.68 | 0.881 | 1.33891011 |
| **LIX1** | 1.8297 | -1.28883751896944 | 0.003 | 0.259 | 2.97573383 |
| **STC2** | 2.1036 | 1.18047731492087 | 0.911 | 0.806 | 3.42108833 |
| **3,00 F** | 5.6090 | 1.16842130461165 | 0.8 | 0.533 | 9.12200733 |
| **C1orf21** | 1.2654 | -0.976825254488665 | 0.071 | 0.385 | 2.05806893 |
| **SLC7A2** | 1.2971 | -0.934742167094553 | 0.372 | 0.683 | 2.10957993 |
| **CER1** | 2.3045 | -1.48392698976301 | 0.029 | 0.314 | 3.74791971 |
| **SMAGP** | 3.3445 | -1.07869486478572 | 0.438 | 0.709 | 5.43930820 |
| **TCF7L1** | 4.5925 | -1.32613382845379 | 0.511 | 0.741 | 7.46882002 |
| **SVIL** | 7.9638 | -0.82752327777411 | 0.429 | 0.722 | 1.29516468 |
| **DIAPH3** | 1.7490 | -0.971690841141216 | 0.768 | 0.902 | 2.84441210 |
| **RGMB** | 2.5087 | -0.973145811664217 | 0.57 | 0.802 | 4.07993911 |
| **TIMP4** | 3.8067 | -0.907074609684798 | 0.229 | 0.565 | 6.19093618 |
| **SAMD4A** | 1.8271 | -0.831433465100265 | 0.028 | 0.309 | 2.97144237 |
| **LMO1** | 2.9461 | -1.2137678666421 | 0.226 | 0.557 | 4.79138388 |
| **SOX2** | 3.1465 | -1.09973200608822 | 0.954 | 0.959 | 5.11718489 |
| **JADE2** | 4.2357 | 0.810401986941131 | 0.943 | 0.794 | 6.88860344 |
| **ZNF48** | 4.6174 | -0.809838961156743 | 0.822 | 0.935 | 7.50937663 |
| **OTX2** | 6.7688 | -1.20016332453558 | 0.81 | 0.918 | 1.10082351 |
| **RASL10B** | 9.2248 | -0.828303516070567 | 0.156 | 0.491 | 1.50023911 |
| **PTPN13** | 1.4016 | -0.999528230239445 | 0.305 | 0.624 | 2.27957828 |
| **EPCAM** | 1.6073 | -1.47846530033168 | 0.996 | 0.95 | 2.61398372 |
| **IGFBP4** | 2.4201 | -0.890677195237306 | 0.167 | 0.503 | 3.93589814 |
| **TET2** | 4.4685 | 0.891428875230593 | 0.679 | 0.337 | 7.26726983 |
| **CREG1** | 1.2723 | -0.802410468030173 | 0.408 | 0.698 | 2.06917318 |
| **SYNPO** | 1.3578 | -0.917410306718017 | 0.024 | 0.295 | 2.20823892 |
| **FAM111B** | 2.3068 | -1.13311683869017 | 0.381 | 0.69 | 3.75161551 |

| | | | | | |
|---|---|---|---|---|---|
| **SALL3** | 2.5035 | -0.853254378884214 | 0.886 | 0.93 | 4.0714580 |
| **HPD** | 1.0265 | -1.23845689671777 | 0.339 | 0.625 | 1.6694542 |
| **SUN2** | 1.0867 | -0.822095894777643 | 0.13 | 0.456 | 1.7673309 |
| **ZNF703** | 1.1660 | -1.11738379470741 | 0.602 | 0.809 | 1.8963421 |
| **HMGB2** | 2.1238 | -1.07682717542747 | 0.98 | 0.981 | 3.4540348 |
| **TSHZ3** | 4.4919 | -0.811307813429345 | 0.185 | 0.508 | 7.3052968 |
| **FGF18** | 5.0552 | -0.947289541467411 | 0.25 | 0.576 | 8.2214311 |
| **WFDC2** | 5.2670 | -1.03188454976308 | 0.208 | 0.533 | 8.5658548 |
| **ARVCF** | 6.4809 | -0.80893265477765 | 0.467 | 0.728 | 1.0539960 |
| **GPRC5A** | 1.4646 | 1.08124852908543 | 0.514 | 0.149 | 2.3819409 |
| **AHDC1** | 1.4090 | -1.20459913498153 | 0.578 | 0.77 | 2.2916125 |
| **DEPDC1** | 1.4222 | 0.905000969424502 | 0.543 | 0.199 | 2.3130172 |
| **MYL4** | 1.4822 | -2.00358207057375 | 0.015 | 0.263 | 2.4106483 |
| **TUBA1A** | 2.5328 | -1.09704534131501 | 0.767 | 0.895 | 4.1192182 |
| **NAV1** | 2.5314 | -0.946887191931978 | 0.53 | 0.755 | 4.1169206 |
| **GLIPR2** | 1.3134 | -1.11805414812156 | 0.954 | 0.977 | 2.1360764 |
| **ANKRD1** | 4.7582 | -1.43007563340825 | 0.028 | 0.288 | 7.7383402 |
| **RAI1** | 4.7729 | -0.8053716929217 | 0.132 | 0.441 | 7.7622580 |
| **NCAPD2** | 1.3154 | -0.917428456476463 | 0.874 | 0.956 | 2.1393953 |
| **TRNP1** | 1.3217 | -0.849479844946554 | 0.708 | 0.831 | 2.1494867 |
| **CAVIN1** | 3.2538 | -1.32201667455022 | 0.667 | 0.822 | 5.2917406 |
| **THBS2** | 5.6707 | -0.820570621515231 | 0.117 | 0.427 | 9.2223663 |
| **SHF** | 1.9145 | -0.95723007474561 | 0.215 | 0.521 | 3.1136206 |
| **GREB1L** | 5.0348 | -1.96306537147121 | 0.09 | 0.376 | 8.1882242 |
| **CCDC92** | 7.7242 | -0.968621564715374 | 0.153 | 0.458 | 1.2561895 |
| **STOX2** | 1.5538 | -0.805226040370628 | 0.443 | 0.707 | 2.5270508 |
| **LMCD1** | 5.6554 | -1.0074887730483 | 0.077 | 0.367 | 9.1973808 |
| **PMAIP1** | 1.4423 | -1.0743174445894 | 0.957 | 0.934 | 2.3456576 |
| **CCN2** | 2.3984 | -1.3583363983056 | 0.767 | 0.924 | 3.9006449 |
| **IFFO2** | 2.4743 | -0.914945602058563 | 0.336 | 0.636 | 4.0239586 |
| **FBN1** | 2.2414 | -0.973646934804643 | 0.108 | 0.397 | 3.6452092 |
| **GALNT10** | 2.7639 | -0.81715890494137 | 0.489 | 0.737 | 4.4949743 |
| **FOSL2** | 1.0085 | -0.937353813699257 | 0.076 | 0.355 | 1.6402644 |
| **WNT4** | 1.3417 | -2.04673190773344 | 0.019 | 0.25 | 2.1821679 |

| | | | | | |
|---|---|---|---|---|---|
| **CENPI** | 2.6176 | -0.817977867376606 | 0.712 | 0.858 | 4.2570251 |
| **CHODL** | 4.3589 | -0.836605545991504 | 0.499 | 0.738 | 7.0889493 |
| **VAMP8** | 4.5391 | -1.08932121998522 | 0.995 | 0.988 | 7.3819412 |
| **HIST1H2BH** | 6.2101 | 0.830820517916975 | 0.971 | 0.846 | 1.0099622 |
| **ID1** | 9.9236 | -0.82793832254683 | 0.984 | 0.982 | 1.6138863 |
| **TNFRSF12A** | 9.9058 | -0.97976272437079 | 0.894 | 0.964 | 1.6109954 |
| **SDK2** | 1.6642 | -1.22246531693633 | 0.164 | 0.439 | 2.7065154 |
| **WNT1** | 7.7788 | -2.34006630623231 | 0.002 | 0.2 | 1.2650705 |
| **JUN** | 1.1370 | -1.19740710140929 | 0.517 | 0.747 | 1.8492579 |
| **ZNF521** | 4.4703 | -0.854996679899853 | 0.01 | 0.215 | 7.2700648 |
| **LRP1** | 7.3791 | -0.927278940143029 | 0.852 | 0.928 | 1.2000710 |
| **SMIM10L2A** | 2.6335 | -1.00817975640147 | 0.533 | 0.73 | 4.2828963 |
| **CDH3** | 3.9450 | -1.0215958965285 | 0.915 | 0.906 | 6.4158713 |
| **ACTG2** | 8.1504 | -1.44416373480876 | 0.682 | 0.857 | 1.3255052 |
| **TPBG** | 1.2767 | -1.3800656252626 | 0.377 | 0.641 | 2.0764248 |
| **PREX1** | 7.6535 | -0.806329482400556 | 0.249 | 0.53 | 1.2447010 |
| **PCK2** | 4.4553 | 1.05344496008122 | 0.806 | 0.653 | 7.2457141 |
| **FOXB1** | 1.1157 | -0.900405159349645 | 0.34 | 0.616 | 1.8146009 |
| **EFS** | 2.3558 | -1.19261545047276 | 0.47 | 0.673 | 3.8312670 |
| **HPGD** | 1.2127 | -1.68033551646767 | 0.961 | 0.907 | 1.9722503 |
| **SUSD3** | 2.0469 | -0.982368749199372 | 0.459 | 0.661 | 3.3289245 |
| **CAPRIN2** | 8.7395 | -0.839936209919793 | 0.574 | 0.77 | 1.4213156 |
| **IGFBP3** | 2.4607 | -1.12442431812859 | 0.187 | 0.444 | 4.0019179 |
| **SLC12A3** | 1.0300 | 0.871074701979025 | 0.227 | 0.013 | 1.6752050 |
| **MSX1** | 2.0174 | -0.842957203825956 | 0.01 | 0.189 | 3.2810088 |
| **GREM1** | 3.4408 | -0.976427959460112 | 0.002 | 0.168 | 5.5958798 |
| **GDF15** | 7.8799 | 0.825038858121082 | 0.336 | 0.083 | 1.2815134 |
| **CEP55** | 8.9976 | -0.829030122878358 | 0.523 | 0.706 | 1.4632869 |
| **TFPI** | 4.0575 | -1.42334851238556 | 0.41 | 0.632 | 6.5987586 |
| **CDH6** | 1.0929 | -1.6039488389912 | 0.024 | 0.211 | 1.7773904 |
| **SNAI2** | 4.4328 | -0.95771802104246 | 0.016 | 0.194 | 7.2091799 |
| **KRT7** | 5.1535 | -0.822186291761716 | 0.226 | 0.489 | 8.3812352 |
| **GADD45G** | 1.6018 | -0.85747833548155 | 0.312 | 0.551 | 2.6050703 |
| **LOXL1** | 1.7414 | -1.31333867994044 | 0.612 | 0.771 | 2.8321958 |

| | | | | | |
|---|---|---|---|---|---|
| **FGF8** | 2.7637 | -1.23025044090151 | 0.521 | 0.679 | 4.4946067 |
| **ANOS1** | 4.1792 | -1.30103467732146 | 0.14 | 0.366 | 6.7967627 |
| **H2AFX** | 6.0019 | -0.810166656549046 | 0.975 | 0.981 | 9.7609434 |
| **COLEC12** | 3.8574 | -1.7023282901364 | 0.37 | 0.591 | 6.2733220 |
| **POMC** | 3.2245 | -1.41476837487524 | 0.754 | 0.792 | 5.2440884 |
| **ADAMTSL4** | 4.9145 | -1.11537609464961 | 0.383 | 0.575 | 7.9925727 |
| **TBXT** | 2.5128 | -0.896854589267425 | 0.033 | 0.211 | 4.0866259 |
| **ACTA2** | 1.3890 | -1.92329925022078 | 0.005 | 0.15 | 2.2589686 |
| **MYOF** | 2.0222 | -0.849409051353825 | 0.113 | 0.329 | 3.2887688 |
| **AHNAK** | 3.0546 | -0.810701438146621 | 0.96 | 0.985 | 4.9678530 |
| **SPATS2L** | 5.0812 | -1.05279501101955 | 0.224 | 0.452 | 8.2637036 |
| **SDC3** | 2.5450 | -0.812905096141162 | 0.701 | 0.806 | 4.1390239 |
| **ABCC4** | 6.5444 | -0.879337165319024 | 0.703 | 0.806 | 1.0643168 |
| **ACTA1** | 2.0436 | -1.09893491634172 | 0.122 | 0.312 | 3.3235298 |
| **COL5A1** | 4.0615 | -1.49798993797909 | 0.547 | 0.705 | 6.6053202 |
| **IER3** | 2.2453 | -1.30000069039263 | 0.141 | 0.335 | 3.6515467 |
| **NPPB** | 1.3045 | -1.13146653677421 | 0.029 | 0.168 | 2.1216260 |
| **SDC2** | 3.9428 | -1.29104844612314 | 0.356 | 0.525 | 6.4122775 |
| **NID2** | 4.4088 | -0.822851923009661 | 0.095 | 0.262 | 7.1701365 |
| **NR2F1** | 1.1687 | -0.903743564494313 | 0.017 | 0.137 | 1.9007643 |
| **MKI67** | 2.0861 | -0.908441121784018 | 0.84 | 0.875 | 3.3926599 |
| **CYP1B1** | 1.6535 | -1.02484861859331 | 0.248 | 0.42 | 2.6891855 |
| **OLIG3** | 6.8551 | -0.837385547928942 | 0.123 | 0.284 | 1.1148542 |
| **PCBP4** | 1.7673 | -0.989517377413392 | 0.291 | 0.442 | 2.8742680 |
| **BGN** | 1.3907 | -1.01213417626028 | 0.266 | 0.439 | 2.2617173 |
| **COL1A1** | 2.6159 | -2.38399042470064 | 0.887 | 0.914 | 4.2543056 |
| **PCSK1** | 1.8282 | -0.928744832048321 | 0.642 | 0.714 | 2.9733011 |
| **CNTFR** | 3.0913 | -1.22335072552526 | 0.178 | 0.323 | 5.0274316 |
| **RHOB** | 5.9006 | -0.998447913687629 | 0.987 | 0.985 | 9.5961569 |
| **COL3A1** | 2.1416 | -1.93878401562384 | 0.067 | 0.16 | 3.4828842 |
| **CNTNAP2** | 3.4127 | -1.32033916431142 | 0.766 | 0.778 | 5.5500798 |
| **PRSS23** | 1.0715 | -1.75831345427838 | 0.676 | 0.699 | 1.7425990 |
| **SFRP1** | 1.6053 | -1.91104080721742 | 0.6 | 0.556 | 0.0026108 |
| **MIXL1** | 6.6022 | -1.06276327056555 | 0.619 | 0.602 | 0.0107371 |

| | | | | | |
|---|---|---|---|---|---|
| **FN1** | 3.2588 | -0.81762243406778 | 0.976 | 0.984 | 0.52998895 |
| **SOX21** | 0.4416 | -1.01213781897608 | 0.267 | 0.248 | 1 |
| **ANXA1** | 0.6523 | -1.39393004976743 | 0.223 | 0.213 | 1 |

Supplemental Table S2_Sheet_2

| | p_val | avg_log2FC | pct.1 | pct.2 | p_val_adj |
|---|---|---|---|---|---|
| **MXRA8** | 0 | -1.55741465193061 | 0.489 | 0.959 | 0 |
| **ATAD3B** | 0 | 1.83054802903255 | 0.999 | 0.917 | 0 |
| **FNDC10** | 0 | 1.48497172414238 | 0.924 | 0.485 | 0 |
| **ERRFI1** | 0 | 1.46725624100408 | 0.991 | 0.756 | 0 |
| **CTNNBIP1** | 0 | -1.0964644158027 | 0.902 | 0.997 | 0 |
| **DFFA** | 0 | 1.12133718778904 | 1 | 0.997 | 0 |
| **MTOR** | 0 | 0.838360224269783 | 0.998 | 0.97 | 0 |
| **EPHA2** | 0 | -2.44447141343473 | 0.486 | 0.991 | 0 |
| **MFAP2** | 0 | -1.10091209497631 | 0.977 | 1 | 0 |
| **KLHDC7A** | 0 | -1.530752537948 | 0.029 | 0.634 | 0 |
| **ID3** | 0 | -3.6801906609182 | 0.518 | 0.997 | 0 |
| **FUCA1** | 0 | 1.62870339763521 | 0.986 | 0.722 | 0 |
| **CNR2** | 0 | 1.39487592548761 | 0.649 | 0.015 | 0 |
| **MAN1C1** | 0 | 1.78069950957311 | 0.987 | 0.682 | 0 |
| **STMN1** | 0 | 1.06521983221465 | 1 | 0.99 | 0 |
| **ARID1A** | 0 | -0.847779574327996 | 0.988 | 1 | 0 |
| **PTPRU** | 0 | 1.50497314209146 | 0.997 | 0.909 | 0 |
| **KHDRBS1** | 0 | -0.845646479942502 | 0.977 | 1 | 0 |
| **EIF3I** | 0 | 1.03619061063003 | 0.87 | 0.452 | 0 |
| **MARCKSL1** | 0 | -0.987646233544853 | 1 | 1 | 0 |
| **KIAA1522** | 0 | -1.05533736209088 | 0.973 | 0.991 | 0 |
| **ZNF362** | 0 | -1.16008337212696 | 0.86 | 0.995 | 0 |
| **PHC2** | 0 | -1.74756208988989 | 0.061 | 0.736 | 0 |
| **MAP7D1** | 0 | -1.92676080905122 | 0.426 | 0.982 | 0 |
| **MYCBP** | 0 | 1.11193723259059 | 0.926 | 0.556 | 0 |
| **AKIRIN1** | 0 | 1.37783988734741 | 1 | 0.999 | 0 |
| **PPT1** | 0 | -1.01060233492263 | 0.968 | 1 | 0 |
| **CLDN19** | 0 | -1.53217575285189 | 0.058 | 0.722 | 0 |
| **SVBP** | 0 | -0.891994790466662 | 0.948 | 0.997 | 0 |
| **UROD** | 0 | -0.924113255660311 | 0.997 | 1 | 0 |
| **HPDL** | 0 | 1.98819793460149 | 0.901 | 0.134 | 0 |
| **PRDX1** | 0 | 1.89321499107838 | 1 | 0.999 | 0 |

| | | | | | |
|---|---|---|---|---|---|
| **DMBX1** | 0 | 1.03980631220401 | 0.545 | 0.037 | 0 |
| **ZYG11A** | 0 | 1.92022612905522 | 0.986 | 0.722 | 0 |
| **DHCR24** | 0 | -1.637154646797 | 0.999 | 1 | 0 |
| **MYSM1** | 0 | 1.31800355466746 | 0.994 | 0.905 | 0 |
| **L1TD1** | 0 | 1.28145669371883 | 1 | 0.998 | 0 |
| **PGM1** | 0 | -1.64614203572311 | 0.879 | 0.997 | 0 |
| **CACHD1** | 0 | -1.46918382698085 | 0.703 | 0.966 | 0 |
| **AK4** | 0 | 2.27221355127315 | 0.998 | 0.929 | 0 |
| **WLS** | 0 | -3.7499707451123 | 0.474 | 0.995 | 0 |
| **CTH** | 0 | 1.63844137796108 | 0.982 | 0.698 | 0 |
| **ST6GALNAC5** | 0 | -1.18457033502302 | 0.088 | 0.648 | 0 |
| **LMO4** | 0 | 1.80687675436307 | 0.999 | 0.949 | 0 |
| **BRDT** | 0 | 3.12911232924594 | 0.968 | 0.176 | 0 |
| **TMED5** | 0 | 0.947408121012265 | 0.996 | 0.925 | 0 |
| **PTBP2** | 0 | -1.09414986632156 | 0.836 | 0.987 | 0 |
| **AGL** | 0 | 2.25786491116439 | 0.999 | 0.922 | 0 |
| **HENMT1** | 0 | 1.96406723046396 | 0.879 | 0.033 | 0 |
| **TAF13** | 0 | 2.63274895735362 | 1 | 0.79 | 0 |
| **PHTF1** | 0 | -1.088463473973 | 0.703 | 0.959 | 0 |
| **OLFML3** | 0 | -4.22931215292426 | 0.622 | 0.997 | 0 |
| **BCAS2** | 0 | 1.05929372537248 | 0.92 | 0.544 | 0 |
| **VANGL1** | 0 | -1.24727059971063 | 0.766 | 0.973 | 0 |
| **SV2A** | 0 | -1.4470800904951 | 0.368 | 0.908 | 0 |
| **PLEKHO1** | 0 | -1.7482436919419 | 0.585 | 0.978 | 0 |
| **HORMAD1** | 0 | 2.61161346483657 | 0.976 | 0.094 | 0 |
| **MLLT11** | 0 | -1.34806968303483 | 0.525 | 0.92 | 0 |
| **S100A10** | 0 | 1.38381969621067 | 1 | 0.998 | 0 |
| **RAB13** | 0 | -0.837264974205248 | 0.989 | 1 | 0 |
| **ATP8B2** | 0 | 1.02009753727452 | 0.993 | 0.953 | 0 |
| **IL6R** | 0 | 1.25833222435845 | 0.793 | 0.213 | 0 |
| **CKS1B** | 0 | 1.32235603347768 | 1 | 0.983 | 0 |
| **FDPS** | 0 | -0.929372934583783 | 0.987 | 1 | 0 |
| **MEX3A** | 0 | -1.71895900956505 | 0.617 | 0.991 | 0 |
| **NAXE** | 0 | -0.933340770761462 | 0.913 | 0.996 | 0 |

| | | | | | |
|---|---|---|---|---|---|
| **NES** | 0 | -1.37083506058839 | 0.997 | 1 | 0 |
| **CRABP2** | 0 | -3.09165863369522 | 0.922 | 1 | 0 |
| **HDGF** | 0 | -1.56095771725604 | 0.922 | 0.999 | 0 |
| **DUSP23** | 0 | 1.74344655803075 | 0.853 | 0.031 | 0 |
| **ATP1A2** | 0 | -1.14680490140471 | 0.011 | 0.549 | 0 |
| **PEA15** | 0 | -1.22283555908685 | 0.958 | 1 | 0 |
| **F11R** | 0 | 1.18808894691756 | 1 | 0.995 | 0 |
| **TSTD1** | 0 | 1.04670273432049 | 0.615 | 0.026 | 0 |
| **RGS5** | 0 | -3.31577136158958 | 0.19 | 0.925 | 0 |
| **MAEL** | 0 | 1.38307485988698 | 0.586 | 0.016 | 0 |
| **PRDX6** | 0 | 0.951233509496713 | 1 | 0.997 | 0 |
| **RGS2** | 0 | 1.4502676768451 | 0.904 | 0.491 | 0 |
| **RO60** | 0 | -1.02015233589646 | 0.926 | 0.997 | 0 |
| **GLRX2** | 0 | -1.27888603933826 | 0.123 | 0.748 | 0 |
| **BTG2** | 0 | -1.61455022952375 | 0.283 | 0.802 | 0 |
| **SNRPE** | 0 | 1.15107709076935 | 0.999 | 0.913 | 0 |
| **LRRN2** | 0 | -1.2512193685957 | 0.113 | 0.652 | 0 |
| **NUCKS1** | 0 | -0.861053943006256 | 1 | 1 | 0 |
| **CR1L** | 0 | 2.38827742512919 | 0.97 | 0.284 | 0 |
| **IRF6** | 0 | 1.03737026327007 | 0.636 | 0.033 | 0 |
| **TRAF5** | 0 | -1.20583965426566 | 0.185 | 0.733 | 0 |
| **VASH2** | 0 | 1.13803859707814 | 0.995 | 0.963 | 0 |
| **CAPN2** | 0 | -2.36588955226124 | 0.37 | 0.995 | 0 |
| **CNIH4** | 0 | 1.77770699358418 | 0.995 | 0.692 | 0 |
| **GNPAT** | 0 | -1.05343245980935 | 0.722 | 0.973 | 0 |
| **LGALS8** | 0 | 1.00536740358225 | 0.599 | 0.017 | 0 |
| **COX20** | 0 | 1.27789265635086 | 0.902 | 0.404 | 0 |
| **ACP1** | 0 | 1.02937091090849 | 0.946 | 0.623 | 0 |
| **ID2** | 0 | -1.8323877790004 | 0.329 | 0.923 | 0 |
| **IAH1** | 0 | 1.93734995490427 | 0.863 | 0.029 | 0 |
| **RHOB** | 0 | -1.65543764782995 | 0.871 | 0.997 | 0 |
| **EFR3B** | 0 | 1.45841109267121 | 0.927 | 0.653 | 0 |
| **CAPN13** | 0 | -1.23344154722275 | 0.129 | 0.715 | 0 |
| **CEBPZ** | 0 | 0.821154674201419 | 1 | 1 | 0 |

| | | | | | |
|---|---|---|---|---|---|
| **PKDCC** | 0 | -2.82635179653856 | 0.246 | 0.936 | 0 |
| **EML4** | 0 | -0.831776674990216 | 0.989 | 1 | 0 |
| **RTN4** | 0 | -0.879374090080569 | 1 | 1 | 0 |
| **CCDC88A** | 0 | -1.08603977824756 | 0.875 | 0.998 | 0 |
| **XPO1** | 0 | 1.0879799932057 | 0.999 | 0.965 | 0 |
| **UGP2** | 0 | 1.4732584056286 | 0.998 | 0.932 | 0 |
| **ANTXR1** | 0 | -1.47655223245862 | 0.281 | 0.886 | 0 |
| **GFPT1** | 0 | 1.6012153348392 | 1 | 0.996 | 0 |
| **SNRPG** | 0 | 1.21461657345402 | 0.999 | 0.97 | 0 |
| **NAGK** | 0 | 1.00935149511109 | 0.996 | 0.955 | 0 |
| **MTHFD2** | 0 | 1.29895896752039 | 1 | 1 | 0 |
| **HK2** | 0 | 0.954514100532737 | 0.999 | 0.985 | 0 |
| **VAMP8** | 0 | -1.37942151888784 | 0.996 | 1 | 0 |
| **TBC1D8** | 0 | 1.19495300281747 | 0.819 | 0.321 | 0 |
| **MAP4K4** | 0 | -0.908230055004341 | 0.994 | 1 | 0 |
| **TFCP2L1** | 0 | 1.57863740823404 | 0.937 | 0.385 | 0 |
| **NIFK** | 0 | 0.865958653880087 | 0.995 | 0.956 | 0 |
| **TSN** | 0 | -0.901530358248537 | 0.998 | 1 | 0 |
| **BIN1** | 0 | -1.77953603050145 | 0.612 | 0.989 | 0 |
| **ZEB2** | 0 | -1.50984609065498 | 0.096 | 0.692 | 0 |
| **KIF5C** | 0 | -1.36368022728715 | 0.899 | 1 | 0 |
| **NMI** | 0 | 1.90738900021308 | 0.967 | 0.354 | 0 |
| **RIF1** | 0 | 0.937172312357314 | 0.999 | 0.994 | 0 |
| **FMNL2** | 0 | 0.85581370254789 | 1 | 0.971 | 0 |
| **GPD2** | 0 | -1.21448243579779 | 0.873 | 0.993 | 0 |
| **GALNT3** | 0 | 1.55215445207331 | 0.742 | 0.008 | 0 |
| **SP5** | 0 | -2.92906030733216 | 0.684 | 1 | 0 |
| **GAD1** | 0 | -2.12905330161817 | 0.295 | 0.965 | 0 |
| **PDK1** | 0 | -1.79816842575056 | 0.945 | 0.998 | 0 |
| **UBE2E3** | 0 | -1.30881857388136 | 0.94 | 1 | 0 |
| **SLC39A10** | 0 | 0.943980848041237 | 0.999 | 0.998 | 0 |
| **CFLAR** | 0 | 1.2259556430926 | 0.987 | 0.762 | 0 |
| **FZD7** | 0 | -2.97266681945979 | 0.339 | 0.96 | 0 |
| **ICA1L** | 0 | 1.62572071234083 | 0.839 | 0.081 | 0 |

| | | | | | |
|---|---|---|---|---|---|
| **WDR12** | 0 | 0.832577393470251 | 0.997 | 0.961 | 0 |
| **FZD5** | 0 | 2.73608390297217 | 0.999 | 0.867 | 0 |
| **CRYGD** | 0 | 1.20466741533449 | 0.649 | 0.019 | 0 |
| **IDH1** | 0 | 1.30208476014761 | 1 | 0.993 | 0 |
| **CPS1** | 0 | -1.39413874508389 | 0.553 | 0.972 | 0 |
| **TMBIM1** | 0 | 1.64590817874213 | 0.926 | 0.39 | 0 |
| **PTPRN** | 0 | 1.20090822505091 | 0.624 | 0.055 | 0 |
| **GMPPA** | 0 | 0.972823450696887 | 0.982 | 0.86 | 0 |
| **OBSL1** | 0 | -1.57296876104011 | 0.74 | 0.997 | 0 |
| **SERPINE2** | 0 | -2.88599472555248 | 0.865 | 0.997 | 0 |
| **FBXO36** | 0 | 1.85811117621334 | 0.969 | 0.676 | 0 |
| **ALPG** | 0 | 3.91937735124497 | 0.99 | 0.407 | 0 |
| **NGEF** | 0 | 1.53227182291385 | 0.686 | 0.038 | 0 |
| **ARL4C** | 0 | -1.37741351498219 | 0.317 | 0.805 | 0 |
| **GBX2** | 0 | -1.46984675562103 | 0.536 | 0.894 | 0 |
| **SLC6A6** | 0 | 1.27939213345259 | 0.998 | 0.91 | 0 |
| **RAB5A** | 0 | 1.71815322949686 | 1 | 0.94 | 0 |
| **EOMES** | 0 | -3.35356201918452 | 0.12 | 0.812 | 0 |
| **OSBPL10** | 0 | 1.50705101160167 | 0.987 | 0.764 | 0 |
| **CTDSPL** | 0 | 1.52026198765469 | 0.972 | 0.809 | 0 |
| **ZNF502** | 0 | 1.34983376232619 | 0.737 | 0.008 | 0 |
| **CDCP1** | 0 | 1.22561922644037 | 0.874 | 0.368 | 0 |
| **SHISA5** | 0 | 0.845082464981294 | 1 | 0.999 | 0 |
| **PRKAR2A** | 0 | 0.933852064575439 | 1 | 0.972 | 0 |
| **TNNC1** | 0 | -1.47366580237706 | 0.084 | 0.704 | 0 |
| **WNT5A** | 0 | -1.74968383848994 | 0.23 | 0.901 | 0 |
| **IL17RD** | 0 | -1.383550692694 | 0.876 | 0.986 | 0 |
| **FLNB** | 0 | 0.912386479475407 | 1 | 0.997 | 0 |
| **PTPRG** | 0 | -1.27224626912883 | 0.842 | 0.993 | 0 |
| **DCBLD2** | 0 | -1.05213966234995 | 0.906 | 0.998 | 0 |
| **TBC1D23** | 0 | 1.88954798345239 | 0.997 | 0.956 | 0 |
| **NIT2** | 0 | 1.19808368854804 | 0.928 | 0.529 | 0 |
| **TOMM70** | 0 | 0.848057166417341 | 0.999 | 0.993 | 0 |
| **ALCAM** | 0 | -1.68596269519534 | 0.219 | 0.781 | 0 |

| | | | | | |
|---|---|---|---|---|---|
| **DPPA2** | 0 | 3.2164241740151 | 0.996 | 0.426 | 0 |
| **DPPA4** | 0 | 1.22349293191855 | 1 | 1 | 0 |
| **NECTIN3** | 0 | -1.49229079361671 | 0.375 | 0.907 | 0 |
| **ATG3** | 0 | 2.51826049890094 | 1 | 0.996 | 0 |
| **SLC35A5** | 0 | 1.09072063004173 | 0.98 | 0.837 | 0 |
| **CCDC80** | 0 | 2.02245443578479 | 0.978 | 0.614 | 0 |
| **NAA50** | 0 | 1.47036703707733 | 0.953 | 0.472 | 0 |
| **B4GALT4** | 0 | 1.52280224564331 | 0.877 | 0.256 | 0 |
| **FSTL1** | 0 | -1.63844420265822 | 0.913 | 0.999 | 0 |
| **NDUFB4** | 0 | 1.07407150360105 | 0.87 | 0.413 | 0 |
| **MGLL** | 0 | 1.71711688367282 | 0.757 | 0.042 | 0 |
| **RBP1** | 0 | -2.55492573427984 | 0.643 | 0.994 | 0 |
| **SPSB4** | 0 | -2.20479988309311 | 0.232 | 0.938 | 0 |
| **CHST2** | 0 | 1.67006632583774 | 0.985 | 0.723 | 0 |
| **LRRC34** | 0 | 1.23365434186737 | 0.675 | 0.003 | 0 |
| **GPR160** | 0 | 1.2130680043258 | 0.661 | 0.116 | 0 |
| **NCEH1** | 0 | 1.17863522201319 | 0.718 | 0.152 | 0 |
| **GNB4** | 0 | -2.53288528821983 | 0.842 | 0.998 | 0 |
| **EPHB3** | 0 | -1.70847743125217 | 0.096 | 0.683 | 0 |
| **SENP2** | 0 | 1.00984653011924 | 1 | 0.992 | 0 |
| **ST6GAL1** | 0 | 0.858837065234319 | 0.999 | 0.998 | 0 |
| **LPP** | 0 | -1.41264474879031 | 0.627 | 0.966 | 0 |
| **P3H2** | 0 | -2.3080218430738 | 0.036 | 0.593 | 0 |
| **HES1** | 0 | -2.21759765172702 | 0.257 | 0.955 | 0 |
| **MUC4** | 0 | 1.41606926900469 | 0.846 | 0.273 | 0 |
| **ZNF595** | 0 | 1.58574002325695 | 0.985 | 0.686 | 0 |
| **ZNF732** | 0 | 1.28293704632797 | 0.692 | 0.003 | 0 |
| **NSD2** | 0 | -1.18227498102122 | 0.443 | 0.892 | 0 |
| **MFSD10** | 0 | -1.09760527541511 | 0.687 | 0.961 | 0 |
| **S100P** | 0 | 1.72495937614469 | 0.641 | 0.025 | 0 |
| **PROM1** | 0 | 1.28203103063496 | 0.979 | 0.752 | 0 |
| **RBPJ** | 0 | 0.987064045127691 | 0.995 | 0.946 | 0 |
| **UCHL1** | 0 | -1.81934046744251 | 0.826 | 0.997 | 0 |
| **OCIAD2** | 0 | -1.27934000780822 | 0.131 | 0.729 | 0 |

| | | | | | |
|---|---|---|---|---|---|
| **PAICS** | 0 | 1.01381922226002 | 0.997 | 0.886 | 0 |
| **REST** | 0 | 0.938484322669146 | 1 | 1 | 0 |
| **MAPK10** | 0 | -2.33912237038224 | 0.033 | 0.787 | 0 |
| **GRID2** | 0 | -1.92005542696898 | 0.113 | 0.765 | 0 |
| **EIF4E** | 0 | 0.896467861593548 | 0.995 | 0.952 | 0 |
| **CAMK2D** | 0 | -1.35497871434057 | 0.244 | 0.766 | 0 |
| **PRSS12** | 0 | 1.53271184045121 | 0.86 | 0.373 | 0 |
| **MAD2L1** | 0 | 1.28910356472975 | 0.971 | 0.786 | 0 |
| **NOCT** | 0 | 1.53282213970709 | 0.997 | 0.853 | 0 |
| **ABCE1** | 0 | 1.0873350863785 | 0.994 | 0.874 | 0 |
| **OTUD4** | 0 | 1.60138092595822 | 0.994 | 0.824 | 0 |
| **FHDC1** | 0 | 1.49331461214922 | 0.826 | 0.241 | 0 |
| **SFRP2** | 0 | -2.91186586854695 | 0.666 | 0.97 | 0 |
| **RAPGEF2** | 0 | 0.995935882135169 | 0.996 | 0.93 | 0 |
| **TRIM60** | 0 | 2.04194876122977 | 0.902 | 0.041 | 0 |
| **TRIM75P** | 0 | 1.39773511821499 | 0.645 | 0.023 | 0 |
| **HMGB2** | 0 | -1.36841251117697 | 0.994 | 0.999 | 0 |
| **FAT1** | 0 | -1.44199264590827 | 0.895 | 0.998 | 0 |
| **ZFP42** | 0 | 4.27231900685245 | 1 | 0.307 | 0 |
| **TRIML2** | 0 | 1.48828939834202 | 0.988 | 0.749 | 0 |
| **IRX1** | 0 | -1.59182461113161 | 0.151 | 0.737 | 0 |
| **SUB1** | 0 | 1.57362776588409 | 0.948 | 0.39 | 0 |
| **SELENOP** | 0 | -1.69282026720062 | 0.14 | 0.851 | 0 |
| **CCL28** | 0 | 0.839670012239989 | 0.51 | 0.005 | 0 |
| **PLPP1** | 0 | 1.7840317649376 | 0.997 | 0.858 | 0 |
| **ZSWIM6** | 0 | -1.32836290903337 | 0.161 | 0.745 | 0 |
| **ERBIN** | 0 | 1.19653148732784 | 0.998 | 0.946 | 0 |
| **MARVELD2** | 0 | 1.05611581926962 | 0.771 | 0.237 | 0 |
| **MAP1B** | 0 | -1.25850224123917 | 0.984 | 1 | 0 |
| **ARSB** | 0 | -1.04127478583658 | 0.699 | 0.957 | 0 |
| **RASGRF2** | 0 | 1.58453193815462 | 0.833 | 0.187 | 0 |
| **CAST** | 0 | -1.13777071194044 | 0.248 | 0.774 | 0 |
| **PAM** | 0 | -1.91635768135982 | 0.112 | 0.884 | 0 |
| **MCC** | 0 | -1.70775785363642 | 0.295 | 0.888 | 0 |

| | | | | | |
|---|---|---|---|---|---|
| **ZNF608** | 0 | -2.30693123398479 | 0.278 | 0.972 | 0 |
| **ADAMTS19** | 0 | -1.59286611972914 | 0.008 | 0.579 | 0 |
| **CXCL14** | 0 | -2.65877760228421 | 0.392 | 0.85 | 0 |
| **WNT8A** | 0 | -2.4283727526034 | 0.371 | 0.906 | 0 |
| **ECSCR** | 0 | -1.08047375592407 | 0.015 | 0.563 | 0 |
| **CXXC5** | 0 | -1.65324848253147 | 0.921 | 0.999 | 0 |
| **DND1** | 0 | 1.61748148900234 | 0.713 | 0.09 | 0 |
| **PCDHB2** | 0 | -1.7934956051019 | 0.191 | 0.893 | 0 |
| **PCDHGB7** | 0 | -1.84434593855452 | 0.11 | 0.832 | 0 |
| **NDFIP1** | 0 | -0.886866910182949 | 0.897 | 0.996 | 0 |
| **LARS** | 0 | 0.881166527844904 | 1 | 1 | 0 |
| **PDGFRB** | 0 | -1.18469879839404 | 0.052 | 0.641 | 0 |
| **ZNF300** | 0 | 1.10698187076299 | 0.816 | 0.284 | 0 |
| **RNF145** | 0 | -1.05852139255092 | 0.4 | 0.862 | 0 |
| **PANK3** | 0 | 1.10849827992253 | 0.982 | 0.795 | 0 |
| **ERGIC1** | 0 | -1.03813252616769 | 0.743 | 0.968 | 0 |
| **ATP6V0E1** | 0 | 0.895250236755588 | 0.999 | 0.961 | 0 |
| **STC2** | 0 | 1.5271341326963 | 0.992 | 0.924 | 0 |
| **SFXN1** | 0 | -0.914964831602747 | 0.986 | 1 | 0 |
| **FGFR4** | 0 | 1.61069339629421 | 0.985 | 0.849 | 0 |
| **NSD1** | 0 | 1.03746368573125 | 1 | 0.981 | 0 |
| **DBN1** | 0 | -0.948555759121081 | 0.998 | 1 | 0 |
| **PDLIM7** | 0 | -1.16510942311855 | 0.965 | 0.999 | 0 |
| **ZNF354C** | 0 | 0.858454082899267 | 0.535 | 0.003 | 0 |
| **CANX** | 0 | 1.00242829985792 | 0.999 | 0.998 | 0 |
| **SERPINB1** | 0 | 1.44816400583103 | 0.757 | 0.044 | 0 |
| **SERPINB6** | 0 | 1.80625918163312 | 0.998 | 0.792 | 0 |
| **TUBB2B** | 0 | -1.98149495651433 | 0.998 | 1 | 0 |
| **CDYL** | 0 | -1.11492947097162 | 0.719 | 0.978 | 0 |
| **DSP** | 0 | -1.38555214610088 | 0.994 | 1 | 0 |
| **GCNT2** | 0 | 1.17027534647088 | 0.997 | 0.963 | 0 |
| **MYLIP** | 0 | 1.24842965201455 | 0.865 | 0.439 | 0 |
| **SOX4** | 0 | 1.14346008240848 | 1 | 0.993 | 0 |
| **DCDC2** | 0 | 1.34154467034142 | 0.679 | 0.065 | 0 |

| | | | | | |
|---|---|---|---|---|---|
| **MRS2** | 0 | 1.08650781875404 | 1 | 0.998 | 0 |
| **HIST1H1A** | 0 | 4.84201982835756 | 0.996 | 0.396 | 0 |
| **HIST1H2BB** | 0 | 2.04400257760602 | 0.698 | 0.09 | 0 |
| **ZFP57** | 0 | 2.82995790462191 | 0.994 | 0.274 | 0 |
| **TRIM26** | 0 | 1.8271130058661 | 0.992 | 0.823 | 0 |
| **NRM** | 0 | -1.19718930527403 | 0.56 | 0.934 | 0 |
| **FLOT1** | 0 | -1.19962216179087 | 0.961 | 1 | 0 |
| **DDAH2** | 0 | -1.38332842312951 | 0.889 | 0.997 | 0 |
| **AGPAT1** | 0 | -0.906698547913412 | 0.987 | 1 | 0 |
| **PBX2** | 0 | -1.09597879720104 | 0.971 | 1 | 0 |
| **CUTA** | 0 | -1.39045790033598 | 0.983 | 1 | 0 |
| **ILRUN** | 0 | 1.51917682891597 | 1 | 0.994 | 0 |
| **ZFAND3** | 0 | -0.873172402162561 | 0.979 | 0.999 | 0 |
| **TOMM6** | 0 | 1.21184128438835 | 0.999 | 0.957 | 0 |
| **TRERF1** | 0 | -1.26083995440941 | 0.04 | 0.668 | 0 |
| **PTK7** | 0 | -1.40702863027241 | 0.975 | 1 | 0 |
| **HSP90AB1** | 0 | 0.813471161822097 | 1 | 1 | 0 |
| **DPPA5** | 0 | 5.85296238944621 | 1 | 0.547 | 0 |
| **KHDC3L** | 0 | 4.88286875372561 | 0.997 | 0.395 | 0 |
| **DDX43** | 0 | 1.83548820805093 | 0.819 | 0.017 | 0 |
| **CGAS** | 0 | 2.34096179066606 | 0.891 | 0.045 | 0 |
| **IRAK1BP1** | 0 | 1.28958243573982 | 0.672 | 0.009 | 0 |
| **CCNC** | 0 | 1.24381822850494 | 1 | 0.993 | 0 |
| **LIN28B** | 0 | 1.46585974728819 | 1 | 0.995 | 0 |
| **ATG5** | 0 | 1.09633707140605 | 0.998 | 0.949 | 0 |
| **CRYBG1** | 0 | 1.08686987271799 | 0.595 | 0.04 | 0 |
| **SEC63** | 0 | 1.26758321477592 | 1 | 0.996 | 0 |
| **SNX3** | 0 | 0.985978559094641 | 0.974 | 0.775 | 0 |
| **WASF1** | 0 | -1.89238673012253 | 0.549 | 0.986 | 0 |
| **GTF3C6** | 0 | 1.55712038391044 | 0.988 | 0.659 | 0 |
| **SLC16A10** | 0 | 1.95071982510287 | 0.908 | 0.256 | 0 |
| **MARCKS** | 0 | -3.2047442263723 | 0.426 | 0.999 | 0 |
| **PKIB** | 0 | 1.23041639221724 | 0.601 | 0.049 | 0 |
| **ECHDC1** | 0 | -1.87974868828003 | 0.185 | 0.943 | 0 |

| | | | | | |
|---|---|---|---|---|---|
| **ARHGAP18** | 0 | 1.14709090395413 | 0.733 | 0.202 | 0 |
| **SGK1** | 0 | 2.12180421112873 | 0.993 | 0.649 | 0 |
| **FUCA2** | 0 | -1.26652283669747 | 0.756 | 0.949 | 0 |
| **AKAP12** | 0 | 1.1733333530101 | 1 | 0.997 | 0 |
| **RMND1** | 0 | 1.3665457354201 | 0.997 | 0.909 | 0 |
| **ARMT1** | 0 | 1.58029655699716 | 1 | 0.992 | 0 |
| **ESR1** | 0 | 0.95651059374518 | 1 | 0.971 | 0 |
| **TCP1** | 0 | 2.16093962737577 | 1 | 0.948 | 0 |
| **IGF2R** | 0 | -1.41606589745022 | 0.617 | 0.982 | 0 |
| **DACT2** | 0 | 1.25941238772697 | 0.669 | 0.038 | 0 |
| **C7orf50** | 0 | -1.12329616657926 | 0.985 | 1 | 0 |
| **BZW2** | 0 | 0.917179487140719 | 1 | 0.988 | 0 |
| **GPNMB** | 0 | 1.32268890891629 | 0.813 | 0.252 | 0 |
| **MPP6** | 0 | -1.10151869880962 | 0.964 | 0.999 | 0 |
| **CYCS** | 0 | 1.62015526403548 | 0.986 | 0.641 | 0 |
| **CBX3** | 0 | 1.89431052990062 | 1 | 0.94 | 0 |
| **SKAP2** | 0 | -1.39685151653158 | 0.084 | 0.715 | 0 |
| **HOXA1** | 0 | -1.65506024941266 | 0.425 | 0.876 | 0 |
| **WIPF3** | 0 | 1.39743110256201 | 0.811 | 0.207 | 0 |
| **MYL7** | 0 | -2.81743325594244 | 0.149 | 0.816 | 0 |
| **HUS1** | 0 | 1.28807856561291 | 1 | 0.969 | 0 |
| **SUN3** | 0 | 0.935777900758785 | 0.522 | 0.008 | 0 |
| **UPP1** | 0 | 2.88149722812996 | 1 | 0.891 | 0 |
| **CHCHD2** | 0 | 1.47157694701444 | 1 | 1 | 0 |
| **ZNF727** | 0 | 1.4885725973235 | 0.75 | 0.003 | 0 |
| **ZNF736** | 0 | 1.13406616006312 | 0.671 | 0.004 | 0 |
| **ZNF680** | 0 | 1.10469904467945 | 0.97 | 0.736 | 0 |
| **LIMK1** | 0 | -0.918486366461089 | 0.986 | 1 | 0 |
| **HIP1** | 0 | -1.06439054939664 | 0.75 | 0.975 | 0 |
| **CDK6** | 0 | -1.74041200667114 | 0.281 | 0.933 | 0 |
| **NPTX2** | 0 | 1.01241475873569 | 0.556 | 0.008 | 0 |
| **ARPC1B** | 0 | 1.35869982786417 | 1 | 0.995 | 0 |
| **STAG3** | 0 | 1.35107849356372 | 0.661 | 0.021 | 0 |
| **COL26A1** | 0 | -2.00952595384303 | 0.765 | 0.988 | 0 |

| | | | | | |
|---|---|---|---|---|---|
| **CUX1** | 0 | -1.29255568313418 | 0.805 | 0.988 | 0 |
| **PTPRZ1** | 0 | -2.58728056640484 | 0.336 | 0.863 | 0 |
| **AASS** | 0 | 2.16986305694882 | 0.984 | 0.806 | 0 |
| **NDUFA5** | 0 | 1.01888776058618 | 0.931 | 0.576 | 0 |
| **HYAL4** | 0 | 1.6756178656641 | 0.808 | 0.02 | 0 |
| **FLNC** | 0 | -1.36257392623254 | 0.909 | 0.994 | 0 |
| **KLHDC10** | 0 | -1.22937757678118 | 0.1 | 0.748 | 0 |
| **MEST** | 0 | 1.92076860402006 | 0.995 | 0.813 | 0 |
| **AKR1B1** | 0 | -0.808879960050561 | 0.991 | 1 | 0 |
| **CALD1** | 0 | -1.8974884005598 | 0.997 | 1 | 0 |
| **TCAF1** | 0 | -2.11053530223198 | 0.455 | 0.989 | 0 |
| **ZNF398** | 0 | 1.72274650573899 | 0.978 | 0.761 | 0 |
| **REPIN1** | 0 | -1.30699973186346 | 0.444 | 0.913 | 0 |
| **UBE3C** | 0 | 0.838384110722234 | 0.999 | 0.982 | 0 |
| **PPP1R3B** | 0 | -2.20277526275407 | 0.831 | 0.999 | 0 |
| **ZDHHC2** | 0 | -1.7992264382209 | 0.141 | 0.835 | 0 |
| **LOXL2** | 0 | -1.45647236623325 | 0.777 | 0.993 | 0 |
| **DPYSL2** | 0 | -1.67659939234428 | 0.937 | 1 | 0 |
| **NRG1** | 0 | 1.40668578783275 | 0.669 | 0.053 | 0 |
| **UNC5D** | 0 | -1.14256771827691 | 0.037 | 0.607 | 0 |
| **EIF4EBP1** | 0 | 0.894885899738517 | 1 | 1 | 0 |
| **ASH2L** | 0 | 1.31722995311685 | 0.998 | 0.988 | 0 |
| **FGFR1** | 0 | -1.00733744905573 | 0.996 | 1 | 0 |
| **NSMAF** | 0 | 1.55688366360375 | 0.999 | 0.907 | 0 |
| **CHD7** | 0 | -2.78065185303216 | 0.364 | 0.991 | 0 |
| **MTFR1** | 0 | 1.08500747883304 | 0.99 | 0.872 | 0 |
| **SGK3** | 0 | 1.19718763302483 | 0.944 | 0.659 | 0 |
| **ARFGEF1** | 0 | 0.961008471103214 | 0.999 | 0.958 | 0 |
| **PRDM14** | 0 | 2.21107709332639 | 0.986 | 0.765 | 0 |
| **TERF1** | 0 | 2.00334099230809 | 0.994 | 0.883 | 0 |
| **FABP5** | 0 | 3.75861770648342 | 1 | 0.517 | 0 |
| **TMEM64** | 0 | -2.36286493126453 | 0.706 | 0.988 | 0 |
| **ESRP1** | 0 | 1.13827841940322 | 1 | 0.877 | 0 |
| **CTHRC1** | 0 | 1.79851700044854 | 0.822 | 0.034 | 0 |

| | | | | | |
|---|---|---|---|---|---|
| **ANGPT1** | 0 | 1.31698540665833 | 0.761 | 0.109 | 0 |
| **ENPP2** | 0 | 1.25065761287186 | 0.76 | 0.166 | 0 |
| **DEPTOR** | 0 | 1.88955281477536 | 0.945 | 0.366 | 0 |
| **ZHX2** | 0 | -1.9878550654774 | 0.123 | 0.912 | 0 |
| **EPPK1** | 0 | 1.272671827479 | 0.997 | 0.82 | 0 |
| **SMARCA2** | 0 | 1.88520663455001 | 0.97 | 0.574 | 0 |
| **GLDC** | 0 | 2.25660465763821 | 0.993 | 0.647 | 0 |
| **KDM4C** | 0 | 1.25980332716298 | 0.969 | 0.784 | 0 |
| **MYORG** | 0 | -1.27658247329966 | 0.203 | 0.752 | 0 |
| **CNTFR** | 0 | -2.33060990398839 | 0.224 | 0.729 | 0 |
| **TPM2** | 0 | -1.39078515423248 | 0.99 | 1 | 0 |
| **GLIPR2** | 0 | -2.24931297154106 | 0.578 | 0.942 | 0 |
| **IGFBPL1** | 0 | -2.30661394396714 | 0.293 | 0.927 | 0 |
| **CARNMT1** | 0 | 0.982808318904747 | 0.991 | 0.889 | 0 |
| **PCSK5** | 0 | 2.04605923852928 | 0.92 | 0.189 | 0 |
| **GCNT1** | 0 | 1.30598324913361 | 0.837 | 0.331 | 0 |
| **PRUNE2** | 0 | 1.79997272176433 | 0.972 | 0.585 | 0 |
| **C9orf64** | 0 | 1.16640183595048 | 0.67 | 0.012 | 0 |
| **BARX1** | 0 | 1.71856857158967 | 0.8 | 0.131 | 0 |
| **FBP1** | 0 | 1.39714081438587 | 0.582 | 0.063 | 0 |
| **ANP32B** | 0 | -1.08724155720556 | 0.965 | 1 | 0 |
| **SEC61B** | 0 | 1.17743708649587 | 0.997 | 0.946 | 0 |
| **ABCA1** | 0 | 1.66142593530037 | 0.971 | 0.633 | 0 |
| **KLF4** | 0 | 2.43115849066311 | 0.931 | 0.353 | 0 |
| **EPB41L4B** | 0 | 1.54724607132811 | 0.956 | 0.556 | 0 |
| **TNC** | 0 | -2.99116774250087 | 0.124 | 0.834 | 0 |
| **CNTRL** | 0 | -1.36221273285247 | 0.377 | 0.891 | 0 |
| **GSN** | 0 | -2.36616048781651 | 0.17 | 0.956 | 0 |
| **STOM** | 0 | 2.49386410927294 | 0.979 | 0.778 | 0 |
| **HSPA5** | 0 | 1.75390047439751 | 0.995 | 0.912 | 0 |
| **CIZ1** | 0 | -1.27109038785339 | 0.432 | 0.929 | 0 |
| **PRRX2** | 0 | 1.19146046646932 | 0.809 | 0.27 | 0 |
| **ASS1** | 0 | 1.89748684711679 | 0.999 | 0.948 | 0 |
| **NET1** | 0 | -1.21067620181877 | 0.822 | 0.984 | 0 |

| | | | | | |
|---|---|---|---|---|---|
| **GDI2** | 0 | -1.00649183743358 | 0.991 | 1 | 0 |
| **ECHDC3** | 0 | 1.29585981914817 | 0.701 | 0.012 | 0 |
| **OLAH** | 0 | 2.12882335604142 | 0.765 | 0.032 | 0 |
| **VIM** | 0 | -3.64939203764158 | 0.721 | 0.997 | 0 |
| **NEBL** | 0 | -1.82832899731199 | 0.207 | 0.897 | 0 |
| **ITGB1** | 0 | -1.03393284104684 | 0.998 | 1 | 0 |
| **RET** | 0 | -1.40326556875832 | 0.31 | 0.826 | 0 |
| **NCOA4** | 0 | 0.954169320704348 | 0.997 | 0.931 | 0 |
| **REEP3** | 0 | -1.09510242666046 | 0.648 | 0.963 | 0 |
| **SLC25A16** | 0 | 2.07875809759074 | 0.999 | 0.714 | 0 |
| **TET1** | 0 | 0.958188333793832 | 0.999 | 0.997 | 0 |
| **HK1** | 0 | -1.16611765576615 | 0.978 | 1 | 0 |
| **H2AFY2** | 0 | -2.99552332133092 | 0.478 | 0.997 | 0 |
| **PCBD1** | 0 | -1.12075219582008 | 0.81 | 0.966 | 0 |
| **PSAP** | 0 | -0.907009624944996 | 0.999 | 1 | 0 |
| **VCL** | 0 | -1.13437870679384 | 0.999 | 1 | 0 |
| **ZCCHC24** | 0 | -1.36038450591103 | 0.387 | 0.864 | 0 |
| **PRXL2A** | 0 | -1.38488370286748 | 0.897 | 0.998 | 0 |
| **LIPA** | 0 | -1.37702260992139 | 0.386 | 0.896 | 0 |
| **PDLIM1** | 0 | 1.37838703006221 | 1 | 0.96 | 0 |
| **FRAT2** | 0 | 1.90325428764929 | 0.999 | 0.984 | 0 |
| **SCD** | 0 | -0.843970950046452 | 1 | 1 | 0 |
| **CUEDC2** | 0 | -1.0847767291627 | 0.612 | 0.947 | 0 |
| **NT5C2** | 0 | -1.11965440863244 | 0.883 | 0.988 | 0 |
| **ATP5MD** | 0 | 1.31787220304662 | 0.881 | 0.322 | 0 |
| **GSTO2** | 0 | 0.990476918975598 | 0.626 | 0.055 | 0 |
| **ADD3** | 0 | -0.949851781230419 | 0.978 | 1 | 0 |
| **SMC3** | 0 | -0.887858465608661 | 0.991 | 1 | 0 |
| **UTF1** | 0 | 3.45833233175941 | 0.964 | 0.361 | 0 |
| **VENTX** | 0 | 1.39846214185367 | 0.929 | 0.482 | 0 |
| **SIGIRR** | 0 | 1.14073832144303 | 0.957 | 0.652 | 0 |
| **TSPAN4** | 0 | -1.09513353524539 | 0.925 | 0.996 | 0 |
| **TRIM6** | 0 | 1.29082844083794 | 0.813 | 0.211 | 0 |
| **TRIM5** | 0 | -1.29061443372884 | 0.351 | 0.888 | 0 |

| | | | | | |
|---|---|---|---|---|---|
| **3,00 DKK** | 0 | -1.5021152161836 | 0.531 | 0.955 | 0 |
| **RRAS2** | 0 | 1.09900374222855 | 1 | 0.955 | 0 |
| **PDE3B** | 0 | -1.26129625532888 | 0.123 | 0.728 | 0 |
| **ANO5** | 0 | 0.864460496257132 | 0.494 | 0.009 | 0 |
| **LGR4** | 0 | 1.00802638243503 | 0.998 | 0.984 | 0 |
| **RCN1** | 0 | 1.0337384015489 | 1 | 0.995 | 0 |
| **TP53I11** | 0 | -1.44934196436059 | 0.406 | 0.928 | 0 |
| **SERPING1** | 0 | -1.52109353933226 | 0.586 | 0.974 | 0 |
| **TMEM258** | 0 | 0.995415277711422 | 0.998 | 0.923 | 0 |
| **FADS2** | 0 | -1.72314737887369 | 1 | 1 | 0 |
| **FADS1** | 0 | -1.39875294064736 | 0.986 | 1 | 0 |
| **ASRGL1** | 0 | 1.14784269265438 | 0.995 | 0.931 | 0 |
| **EEF1G** | 0 | 1.15706090581135 | 1 | 1 | 0 |
| **MTA2** | 0 | -0.923933408280171 | 0.993 | 1 | 0 |
| **PLAAT5** | 0 | -1.37738583850267 | 0.333 | 0.828 | 0 |
| **RCOR2** | 0 | -1.19824961058805 | 0.762 | 0.977 | 0 |
| **NDUFV1** | 0 | 1.02170602550652 | 1 | 1 | 0 |
| **CCND1** | 0 | -1.16045831217523 | 0.994 | 0.999 | 0 |
| **FGF4** | 0 | 1.60269509462715 | 0.957 | 0.639 | 0 |
| **RAB6A** | 0 | 1.23159321420327 | 1 | 0.987 | 0 |
| **UCP2** | 0 | 1.34044877062954 | 0.997 | 0.919 | 0 |
| **PAK1** | 0 | 1.28249327516211 | 0.995 | 0.897 | 0 |
| **TENM4** | 0 | -1.48933386999722 | 0.295 | 0.819 | 0 |
| **AMOTL1** | 0 | -2.36775335692552 | 0.672 | 1 | 0 |
| **DCUN1D5** | 0 | 1.27845941964852 | 1 | 0.964 | 0 |
| **USP28** | 0 | 1.48354648050537 | 1 | 0.996 | 0 |
| **TAGLN** | 0 | -2.95585753505997 | 0.779 | 0.997 | 0 |
| **HYOU1** | 0 | 1.19364932344313 | 1 | 1 | 0 |
| **HYLS1** | 0 | 1.35705725904456 | 0.704 | 0.123 | 0 |
| **FAM118B** | 0 | -1.11198842636572 | 0.678 | 0.97 | 0 |
| **SNX19** | 0 | 1.10401648004612 | 0.65 | 0.029 | 0 |
| **TEAD4** | 0 | 0.934314248252267 | 1 | 0.989 | 0 |
| **CCND2** | 0 | -1.76664767852797 | 0.68 | 0.976 | 0 |
| **PTMS** | 0 | -1.40596214084226 | 0.999 | 1 | 0 |

| | | | | | |
|---|---|---|---|---|---|
| **GDF3** | 0 | 2.66638751904868 | 0.987 | 0.716 | 0 |
| **DPPA3** | 0 | 3.70606770129368 | 0.977 | 0.207 | 0 |
| **PHC1** | 0 | 0.928080575636895 | 1 | 1 | 0 |
| **DUSP16** | 0 | 1.54435235759984 | 0.975 | 0.763 | 0 |
| **PLBD1** | 0 | 2.45254328609329 | 1 | 0.896 | 0 |
| **YAF2** | 0 | -1.39413572336218 | 0.084 | 0.768 | 0 |
| **TWF1** | 0 | 1.09612852606159 | 0.839 | 0.327 | 0 |
| **COL2A1** | 0 | -1.96426022100772 | 0.022 | 0.575 | 0 |
| **PFKM** | 0 | -1.20592552527608 | 0.749 | 0.983 | 0 |
| **FKBP11** | 0 | 1.33127253687894 | 0.91 | 0.462 | 0 |
| **TUBA1C** | 0 | 1.45253119228841 | 1 | 1 | 0 |
| **FIGNL2** | 0 | -1.13366139835682 | 0.121 | 0.672 | 0 |
| **RARG** | 0 | -1.62581190844042 | 0.329 | 0.862 | 0 |
| **CBX5** | 0 | -1.1582905821076 | 0.934 | 0.999 | 0 |
| **ITGA5** | 0 | -2.19881744565774 | 0.259 | 0.93 | 0 |
| **HMGA2** | 0 | -1.45253889459129 | 0.891 | 1 | 0 |
| **ATXN7L3B** | 0 | -0.889101141097095 | 0.978 | 1 | 0 |
| **CCDC59** | 0 | 1.07237263260721 | 0.989 | 0.853 | 0 |
| **NTS** | 0 | -3.18821350960122 | 0.618 | 0.995 | 0 |
| **TMPO** | 0 | 0.894667689240593 | 1 | 0.998 | 0 |
| **ACTR6** | 0 | 1.27659654377101 | 0.924 | 0.486 | 0 |
| **SYCP3** | 0 | 1.58494099179574 | 0.587 | 0.021 | 0 |
| **HSP90B1** | 0 | 2.52586675948847 | 0.998 | 0.779 | 0 |
| **ALDH1L2** | 0 | 1.31180146084905 | 0.78 | 0.144 | 0 |
| **MMAB** | 0 | -1.07233344623946 | 0.899 | 0.994 | 0 |
| **VSIG10** | 0 | 1.80690977356282 | 0.998 | 0.935 | 0 |
| **MSI1** | 0 | -1.94634577492281 | 0.215 | 0.944 | 0 |
| **RHOF** | 0 | 1.47757470859252 | 0.952 | 0.456 | 0 |
| **DENR** | 0 | 1.19945783516122 | 1 | 0.976 | 0 |
| **VPS37B** | 0 | 2.03797665580784 | 0.994 | 0.709 | 0 |
| **FBRSL1** | 0 | 0.935433861937122 | 0.986 | 0.861 | 0 |
| **SKA3** | 0 | 1.15518262471822 | 0.987 | 0.939 | 0 |
| **WASF3** | 0 | -1.51745362998324 | 0.8 | 0.99 | 0 |
| **PDS5B** | 0 | -0.981109988062089 | 0.92 | 0.998 | 0 |

| | | | | | |
|---|---|---|---|---|---|
| **NBEA** | 0 | -1.81628083390131 | 0.058 | 0.781 | 0 |
| **DCLK1** | 0 | -2.21420343752829 | 0.842 | 0.961 | 0 |
| **UFM1** | 0 | 1.19754748956985 | 0.999 | 0.944 | 0 |
| **LHFPL6** | 0 | -2.06168859860084 | 0.157 | 0.847 | 0 |
| **SLC25A15** | 0 | -0.950082100657784 | 0.817 | 0.979 | 0 |
| **RGCC** | 0 | 1.56073837188249 | 0.695 | 0.126 | 0 |
| **DNAJC15** | 0 | 1.41306928596045 | 0.999 | 0.997 | 0 |
| **LCP1** | 0 | 2.23772268387688 | 0.996 | 0.685 | 0 |
| **ESD** | 0 | -1.18318184587882 | 0.989 | 0.999 | 0 |
| **RB1** | 0 | -1.16907834851231 | 0.411 | 0.883 | 0 |
| **CNMD** | 0 | 1.59515466359466 | 0.857 | 0.344 | 0 |
| **KLF5** | 0 | 1.92471892362238 | 0.994 | 0.701 | 0 |
| **KLF12** | 0 | -1.47186148720899 | 0.101 | 0.777 | 0 |
| **EDNRB** | 0 | -2.12243990800634 | 0.135 | 0.766 | 0 |
| **OBI1** | 0 | -1.36323022723431 | 0.615 | 0.952 | 0 |
| **GPC6** | 0 | -2.55201308138631 | 0.429 | 0.992 | 0 |
| **DZIP1** | 0 | -1.43846648600295 | 0.726 | 0.992 | 0 |
| **IPO5** | 0 | -1.32513380719424 | 1 | 1 | 0 |
| **FARP1** | 0 | -1.08381190494596 | 0.703 | 0.964 | 0 |
| **ZIC5** | 0 | -2.05226852400944 | 0.078 | 0.706 | 0 |
| **ZIC2** | 0 | -2.55598958790034 | 0.114 | 0.85 | 0 |
| **PCCA** | 0 | -1.36911021743784 | 0.917 | 0.998 | 0 |
| **ARGLU1** | 0 | -0.953727856651486 | 0.997 | 1 | 0 |
| **COL4A1** | 0 | -1.79474013416852 | 0.821 | 0.999 | 0 |
| **COL4A2** | 0 | -2.06020143049158 | 0.829 | 1 | 0 |
| **ANKRD10** | 0 | -1.40437325803789 | 0.899 | 0.997 | 0 |
| **LAMP1** | 0 | -1.048503925575 | 1 | 1 | 0 |
| **ARHGEF40** | 0 | -2.12149334395404 | 0.201 | 0.935 | 0 |
| **ZNF219** | 0 | -1.76520176440036 | 0.643 | 0.96 | 0 |
| **SALL2** | 0 | -2.02022848737471 | 0.881 | 0.998 | 0 |
| **AJUBA** | 0 | -1.31194111762082 | 0.728 | 0.976 | 0 |
| **REC8** | 0 | 2.97779210166369 | 0.989 | 0.209 | 0 |
| **MIPOL1** | 0 | -1.12490648880836 | 0.219 | 0.766 | 0 |
| **ARF6** | 0 | 1.07051682664775 | 1 | 0.985 | 0 |

| | | | | | |
|---|---|---|---|---|---|
| **PYGL** | 0 | 1.68033078583207 | 1 | 0.975 | 0 |
| **TMX1** | 0 | 0.849855896730717 | 1 | 0.975 | 0 |
| **GNPNAT1** | 0 | 1.52723119835445 | 0.997 | 0.937 | 0 |
| **WDHD1** | 0 | 1.16228199241018 | 1 | 0.999 | 0 |
| **TMEM30B** | 0 | 1.64044518352006 | 0.804 | 0.013 | 0 |
| **HSPA2** | 0 | 2.6337625276147 | 0.935 | 0.121 | 0 |
| **GPX2** | 0 | 1.9274691002218 | 0.727 | 0.045 | 0 |
| **RAB15** | 0 | 1.0851777596891 | 1 | 0.993 | 0 |
| **DCAF5** | 0 | -1.13228049080896 | 0.701 | 0.976 | 0 |
| **DCAF4** | 0 | 1.61292873444343 | 0.998 | 0.863 | 0 |
| **ACOT4** | 0 | 0.872446045049806 | 0.535 | 0.01 | 0 |
| **VRTN** | 0 | -2.40130697060963 | 0.738 | 1 | 0 |
| **NPC2** | 0 | -0.909504497273006 | 1 | 1 | 0 |
| **SLIRP** | 0 | 1.13415007241881 | 1 | 1 | 0 |
| **GLRX5** | 0 | -0.890255107459495 | 1 | 1 | 0 |
| **TUNAR** | 0 | -2.32807295487409 | 0.315 | 0.901 | 0 |
| **C14orf132** | 0 | -1.79386625304839 | 0.147 | 0.799 | 0 |
| **DLK1** | 0 | 1.71108716524881 | 0.828 | 0.138 | 0 |
| **TNFAIP2** | 0 | -1.87207538745278 | 0.192 | 0.785 | 0 |
| **MTA1** | 0 | -0.815543041729627 | 0.994 | 1 | 0 |
| **CRIP1** | 0 | 3.46588635765141 | 1 | 0.817 | 0 |
| **SNRPN** | 0 | 1.71319337935039 | 0.993 | 0.844 | 0 |
| **DPH6** | 0 | 1.07239218055105 | 0.967 | 0.7 | 0 |
| **SPINT1** | 0 | 1.3735767904059 | 1 | 0.892 | 0 |
| **PDIA3** | 0 | 1.39941983135706 | 1 | 0.997 | 0 |
| **B2M** | 0 | -1.66121451139803 | 0.512 | 0.967 | 0 |
| **SORD** | 0 | -1.13193663118592 | 0.537 | 0.916 | 0 |
| **SHF** | 0 | -1.4569557124806 | 0.333 | 0.853 | 0 |
| **MYEF2** | 0 | -1.317348417358 | 0.545 | 0.919 | 0 |
| **EID1** | 0 | -1.47123780402687 | 0.983 | 1 | 0 |
| **ATP8B4** | 0 | 2.00412072605241 | 0.806 | 0.041 | 0 |
| **RSL24D1** | 0 | 1.0859744294708 | 0.726 | 0.138 | 0 |
| **PRTG** | 0 | -2.26660185502643 | 0.135 | 0.896 | 0 |
| **GTF2A2** | 0 | 1.03532592681331 | 0.999 | 0.958 | 0 |

| | | | | | |
|---|---|---|---|---|---|
| **SNX1** | 0 | -1.2142051338979 | 0.875 | 0.996 | 0 |
| **PPIB** | 0 | 1.19541465441657 | 1 | 0.999 | 0 |
| **PCLAF** | 0 | -2.76438726362562 | 0.165 | 0.973 | 0 |
| **CLPX** | 0 | 1.05784901730227 | 0.998 | 0.978 | 0 |
| **IGDCC3** | 0 | -1.87922450800202 | 0.57 | 0.979 | 0 |
| **MYO9A** | 0 | 1.06509492164453 | 0.872 | 0.471 | 0 |
| **CD276** | 0 | -1.06677317667173 | 0.989 | 1 | 0 |
| **ETFA** | 0 | -1.51391933966058 | 0.708 | 0.993 | 0 |
| **TSPAN3** | 0 | -1.11874250535094 | 0.994 | 1 | 0 |
| **PEAK1** | 0 | -1.22564041383173 | 0.367 | 0.863 | 0 |
| **DNAJA4** | 0 | 1.74793317081475 | 0.842 | 0.033 | 0 |
| **MFGE8** | 0 | -1.35450322075567 | 0.61 | 0.96 | 0 |
| **ARPIN** | 0 | -1.18394779894933 | 0.316 | 0.816 | 0 |
| **FAM174B** | 0 | -1.87724679188924 | 0.487 | 0.942 | 0 |
| **HBM** | 0 | 1.02312029027893 | 0.588 | 0.017 | 0 |
| **HBA2** | 0 | 1.76156968764193 | 0.808 | 0.149 | 0 |
| **NME4** | 0 | -0.999783399245982 | 1 | 1 | 0 |
| **CACNA1H** | 0 | 1.68076693767793 | 0.84 | 0.052 | 0 |
| **SYNGR3** | 0 | 1.35146462494235 | 0.935 | 0.566 | 0 |
| **IL32** | 0 | 1.42528198923676 | 0.841 | 0.333 | 0 |
| **NMRAL1** | 0 | 1.78156571771501 | 0.997 | 0.72 | 0 |
| **HMOX2** | 0 | 0.923394959367248 | 0.998 | 0.943 | 0 |
| **C16orf72** | 0 | 0.800358262922396 | 1 | 0.998 | 0 |
| **GSPT1** | 0 | 1.04076033871886 | 0.998 | 0.912 | 0 |
| **SMG1** | 0 | 1.06099569039881 | 0.911 | 0.543 | 0 |
| **NDUFAB1** | 0 | 1.10698598420545 | 1 | 1 | 0 |
| **SEPHS2** | 0 | 1.47153410167728 | 1 | 0.989 | 0 |
| **GPT2** | 0 | 1.09295716850179 | 1 | 0.981 | 0 |
| **DNAJA2** | 0 | 0.926872349969875 | 1 | 1 | 0 |
| **NETO2** | 0 | 1.2869641470359 | 0.971 | 0.781 | 0 |
| **ZNF423** | 0 | -1.67079645833305 | 0.754 | 0.995 | 0 |
| **SALL1** | 0 | -2.91289849699434 | 0.192 | 0.955 | 0 |
| **MMP2** | 0 | -3.59128595561396 | 0.449 | 0.998 | 0 |
| **MT1E** | 0 | 2.25221991857662 | 0.935 | 0.148 | 0 |

| | | | | | |
|---|---|---|---|---|---|
| **MT1F** | 0 | 2.07149869754016 | 0.988 | 0.768 | 0 |
| **MT1G** | 0 | 3.58245400856727 | 0.99 | 0.268 | 0 |
| **MT1H** | 0 | 4.51814847823881 | 0.996 | 0.362 | 0 |
| **MT1X** | 0 | 2.78591666003955 | 1 | 0.979 | 0 |
| **CCDC102A** | 0 | -1.26379466175594 | 0.305 | 0.841 | 0 |
| **USB1** | 0 | -1.13621474395543 | 0.911 | 0.996 | 0 |
| **CMTM3** | 0 | -1.01960748855194 | 0.978 | 1 | 0 |
| **CDH1** | 0 | 1.55530853881111 | 0.999 | 0.77 | 0 |
| **HAS3** | 0 | -1.7333398999008 | 0.385 | 0.827 | 0 |
| **BCAR1** | 0 | -1.14668325957579 | 0.941 | 0.997 | 0 |
| **CFDP1** | 0 | -0.983780291436161 | 0.992 | 1 | 0 |
| **VAT1L** | 0 | -2.41763833441657 | 0.022 | 0.644 | 0 |
| **WWOX** | 0 | -1.59416833509571 | 0.582 | 0.963 | 0 |
| **PIEZO1** | 0 | 1.16685872124394 | 0.677 | 0.047 | 0 |
| **RFLNB** | 0 | 2.87704308818586 | 0.988 | 0.676 | 0 |
| **SHPK** | 0 | 1.37244346947037 | 0.984 | 0.834 | 0 |
| **ZFP3** | 0 | 1.20396183941015 | 0.626 | 0.011 | 0 |
| **TRAPPC1** | 0 | -0.911073322956689 | 0.984 | 1 | 0 |
| **NTN1** | 0 | 7.6575896366897 | 1 | 0.986 | 0 |
| **TMEM97** | 0 | -1.02898687258319 | 0.996 | 1 | 0 |
| **FLOT2** | 0 | -1.29658366359638 | 0.909 | 0.999 | 0 |
| **CPD** | 0 | -0.913563538297923 | 0.977 | 0.998 | 0 |
| **UTP6** | 0 | 0.903495926407844 | 1 | 0.958 | 0 |
| **TMEM98** | 0 | -1.57578516433063 | 0.663 | 0.985 | 0 |
| **SLFN12** | 0 | 1.52136913721614 | 0.775 | 0.076 | 0 |
| **AP2B1** | 0 | 1.35292105153051 | 0.995 | 0.873 | 0 |
| **IGFBP4** | 0 | -2.04959096458225 | 0.463 | 0.932 | 0 |
| **KRT19** | 0 | -1.13194489744739 | 0.753 | 0.982 | 0 |
| **EIF1** | 0 | 1.6612618004743 | 1 | 0.922 | 0 |
| **FKBP10** | 0 | -1.70126640917301 | 0.838 | 0.997 | 0 |
| **VAT1** | 0 | 1.04112203513347 | 1 | 1 | 0 |
| **FAM171A2** | 0 | -1.15152863087134 | 0.702 | 0.952 | 0 |
| **FZD2** | 0 | -2.98140442352084 | 0.264 | 0.98 | 0 |
| **GJC1** | 0 | -1.4315720230501 | 0.889 | 0.999 | 0 |

| | | | | | |
|---|---|---|---|---|---|
| **HOXB2** | 0 | 1.72413896853711 | 0.902 | 0.478 | 0 |
| **PRAC1** | 0 | -1.16247259069287 | 0.074 | 0.635 | 0 |
| **IGF2BP1** | 0 | -1.13852243660391 | 0.993 | 1 | 0 |
| **NGFR** | 0 | -1.9535024013596 | 0.288 | 0.85 | 0 |
| **AKAP1** | 0 | 1.00700381954262 | 0.995 | 0.957 | 0 |
| **MSI2** | 0 | -1.4596934897379 | 0.494 | 0.959 | 0 |
| **PITPNC1** | 0 | 1.03616412361459 | 0.997 | 0.961 | 0 |
| **NOL11** | 0 | 1.21759067264587 | 0.905 | 0.455 | 0 |
| **JPT1** | 0 | 0.807755995643242 | 1 | 0.998 | 0 |
| **SEPTIN9** | 0 | -1.04812268416317 | 0.996 | 1 | 0 |
| **LGALS3BP** | 0 | -1.93808025162101 | 0.174 | 0.914 | 0 |
| **MCRIP1** | 0 | -0.834793256731886 | 0.982 | 0.999 | 0 |
| **ARHGDIA** | 0 | -0.852189813711382 | 0.994 | 1 | 0 |
| **PYCR1** | 0 | 2.08576258192263 | 0.998 | 0.596 | 0 |
| **TBCD** | 0 | 1.36935590568895 | 1 | 0.999 | 0 |
| **MYL12A** | 0 | 1.03097498571245 | 0.945 | 0.654 | 0 |
| **PIEZO2** | 0 | -2.04910503253565 | 0.072 | 0.855 | 0 |
| **IMPA2** | 0 | 1.18955044412222 | 1 | 0.975 | 0 |
| **ESCO1** | 0 | 0.933922784069836 | 0.991 | 0.905 | 0 |
| **CDH2** | 0 | -3.44057692253815 | 0.208 | 0.941 | 0 |
| **RNF125** | 0 | 2.83108125837052 | 0.988 | 0.673 | 0 |
| **DTNA** | 0 | -1.98129717456052 | 0.083 | 0.645 | 0 |
| **GALNT1** | 0 | -1.26987777517337 | 0.942 | 1 | 0 |
| **TPGS2** | 0 | -0.959694722287913 | 0.983 | 1 | 0 |
| **HAUS1** | 0 | 1.10994283388848 | 0.977 | 0.782 | 0 |
| **TCF4** | 0 | -1.65676854324151 | 0.563 | 0.974 | 0 |
| **CFD** | 0 | 1.31193321361456 | 0.745 | 0.093 | 0 |
| **CNN2** | 0 | -1.1464830518867 | 1 | 1 | 0 |
| **REEP6** | 0 | 1.94942145919312 | 0.942 | 0.414 | 0 |
| **ZNF57** | 0 | 1.22045378235136 | 0.682 | 0.012 | 0 |
| **MATK** | 0 | 1.68918890416945 | 0.83 | 0.086 | 0 |
| **TNFSF9** | 0 | 2.04416152815497 | 0.823 | 0.07 | 0 |
| **CD70** | 0 | 2.69630402358176 | 0.887 | 0.089 | 0 |
| **RAB11B** | 0 | -0.929268449260797 | 0.999 | 1 | 0 |

| | | | | | |
|---|---|---|---|---|---|
| **ZNF560** | 0 | 1.21450102754379 | 0.71 | 0.005 | 0 |
| **AP1M2** | 0 | 1.28178228131617 | 0.912 | 0.353 | 0 |
| **DOCK6** | 0 | 1.26733662700589 | 0.99 | 0.805 | 0 |
| **ACP5** | 0 | 1.18406825214077 | 0.662 | 0.058 | 0 |
| **ZNF878** | 0 | 1.38200714805969 | 0.749 | 0.118 | 0 |
| **ZNF844** | 0 | 1.25822822181139 | 0.681 | 0.007 | 0 |
| **ZNF625** | 0 | 1.03482336850355 | 0.994 | 0.931 | 0 |
| **PRKACA** | 0 | -1.0805794429005 | 0.973 | 1 | 0 |
| **NOTCH3** | 0 | -1.05584421810098 | 0.958 | 0.995 | 0 |
| **AP1M1** | 0 | -1.04923936877082 | 0.855 | 0.992 | 0 |
| **BST2** | 0 | -2.26526524796652 | 0.262 | 0.832 | 0 |
| **SLC5A5** | 0 | 1.17483966795869 | 0.681 | 0.056 | 0 |
| **ZNF253** | 0 | 1.5409351506498 | 0.807 | 0.013 | 0 |
| **ZNF90** | 0 | 2.35782260073682 | 0.999 | 0.721 | 0 |
| **ZNF486** | 0 | 2.3612470604644 | 0.985 | 0.407 | 0 |
| **ZNF626** | 0 | 1.06471732843343 | 0.655 | 0.021 | 0 |
| **ZNF257** | 0 | 0.93593473063399 | 0.563 | 0.006 | 0 |
| **ZNF676** | 0 | 2.77676712055829 | 0.957 | 0.129 | 0 |
| **ZNF729** | 0 | 1.91868377615412 | 0.691 | 0.015 | 0 |
| **ZNF98** | 0 | 1.40442192987066 | 0.724 | 0.004 | 0 |
| **ZNF492** | 0 | 1.95368659248239 | 0.876 | 0.009 | 0 |
| **ZNF728** | 0 | 1.06851850879668 | 0.586 | 0.005 | 0 |
| **CHST8** | 0 | 1.4770368603641 | 0.78 | 0.14 | 0 |
| **ZNF146** | 0 | 1.0410051105041 | 1 | 1 | 0 |
| **ZNF568** | 0 | 1.8593319742923 | 0.907 | 0.086 | 0 |
| **PPP1R14A** | 0 | 1.03642793439149 | 0.993 | 0.898 | 0 |
| **C19orf33** | 0 | 1.8720097429514 | 0.897 | 0.233 | 0 |
| **KCNK6** | 0 | 1.21705819825768 | 0.759 | 0.148 | 0 |
| **FCGBP** | 0 | 1.96773369029526 | 0.745 | 0.046 | 0 |
| **PSMC4** | 0 | 1.03961712461113 | 1 | 1 | 0 |
| **PLD3** | 0 | -0.847291906375004 | 0.999 | 1 | 0 |
| **AXL** | 0 | -2.57904951723617 | 0.182 | 0.915 | 0 |
| **ATP1A3** | 0 | 1.28602998471355 | 0.718 | 0.083 | 0 |
| **BCAM** | 0 | -1.96058577716874 | 0.656 | 0.946 | 0 |

| | | | | | |
|---|---|---|---|---|---|
| **ARHGAP35** | 0 | -1.08593516563318 | 0.92 | 0.995 | 0 |
| **TMEM160** | 0 | 1.09906187572553 | 0.998 | 0.999 | 0 |
| **CA11** | 0 | -1.57293928890115 | 0.054 | 0.725 | 0 |
| **RASIP1** | 0 | 1.33048290259503 | 0.801 | 0.184 | 0 |
| **RCN3** | 0 | 1.37119577457911 | 0.784 | 0.113 | 0 |
| **PTOV1** | 0 | -0.848370361175224 | 0.995 | 1 | 0 |
| **KLK13** | 0 | 1.69234073112673 | 0.909 | 0.315 | 0 |
| **LIM2** | 0 | 1.9644867611836 | 0.843 | 0.022 | 0 |
| **ZNF649** | 0 | 1.80165308109768 | 0.992 | 0.812 | 0 |
| **ZNF841** | 0 | 1.26354246630186 | 0.728 | 0.162 | 0 |
| **ZNF880** | 0 | 1.34047157422776 | 0.821 | 0.127 | 0 |
| **ZNF578** | 0 | 1.29835237424527 | 0.872 | 0.314 | 0 |
| **ZNF600** | 0 | 1.25001183301264 | 0.985 | 0.779 | 0 |
| **ZNF677** | 0 | 1.99947726741881 | 0.915 | 0.018 | 0 |
| **NLRP7** | 0 | 5.4071238519755 | 1 | 0.58 | 0 |
| **NLRP2** | 0 | 2.97113761132407 | 1 | 0.99 | 0 |
| **TNNI3** | 0 | 1.22106799826635 | 0.804 | 0.187 | 0 |
| **C19orf85** | 0 | 2.24207841669644 | 0.788 | 0.074 | 0 |
| **ZNF580** | 0 | -1.27385542443733 | 0.412 | 0.895 | 0 |
| **ZNF667** | 0 | 1.03523160669352 | 0.62 | 0.006 | 0 |
| **PEG3** | 0 | 2.26648876587937 | 0.781 | 0.026 | 0 |
| **ZNF552** | 0 | 1.20823054493494 | 0.979 | 0.694 | 0 |
| **ZCCHC3** | 0 | -1.87890067944464 | 0.786 | 0.999 | 0 |
| **SOX12** | 0 | -1.52639742918549 | 0.659 | 0.971 | 0 |
| **TRIB3** | 0 | 1.90251684915512 | 0.957 | 0.697 | 0 |
| **SRXN1** | 0 | -1.38899609790656 | 0.703 | 0.961 | 0 |
| **SNRPB** | 0 | -0.900258934163642 | 1 | 1 | 0 |
| **CDS2** | 0 | -1.1976527922395 | 0.946 | 1 | 0 |
| **FERMT1** | 0 | 1.47649298872593 | 0.758 | 0.125 | 0 |
| **BTBD3** | 0 | -1.25484267700318 | 0.753 | 0.984 | 0 |
| **SNRPB2** | 0 | 1.11811765847746 | 0.994 | 0.873 | 0 |
| **OVOL2** | 0 | 1.27514357736648 | 0.843 | 0.34 | 0 |
| **RIN2** | 0 | -1.77129087603985 | 0.283 | 0.913 | 0 |
| **XRN2** | 0 | -0.891877630150109 | 1 | 1 | 0 |

| | | | | | |
|---|---|---|---|---|---|
| **CST3** | 0 | -1.70718508565465 | 0.528 | 0.953 | 0 |
| **PYGB** | 0 | 1.42970360663207 | 0.999 | 0.999 | 0 |
| **GINS1** | 0 | -0.958983821745737 | 0.997 | 1 | 0 |
| **ASXL1** | 0 | -1.82769957927193 | 0.668 | 0.995 | 0 |
| **NOL4L** | 0 | -1.23413854526653 | 0.453 | 0.899 | 0 |
| **MAPRE1** | 0 | -0.832815742650028 | 1 | 1 | 0 |
| **CBFA2T2** | 0 | 1.78129856073323 | 0.994 | 0.918 | 0 |
| **RALY** | 0 | -0.806422386331899 | 0.98 | 0.999 | 0 |
| **DYNLRB1** | 0 | -1.18051743729445 | 1 | 1 | 0 |
| **ERGIC3** | 0 | -0.934263233496911 | 1 | 1 | 0 |
| **CPNE1** | 0 | -1.29501236487609 | 0.99 | 1 | 0 |
| **MYL9** | 0 | -3.18843238513083 | 0.832 | 1 | 0 |
| **TGIF2** | 0 | -1.55173177549741 | 0.933 | 1 | 0 |
| **SAMHD1** | 0 | 2.67438137822407 | 0.982 | 0.611 | 0 |
| **MANBAL** | 0 | -0.828355287491505 | 0.989 | 1 | 0 |
| **SRC** | 0 | -1.44269075981014 | 0.812 | 0.997 | 0 |
| **NNAT** | 0 | 1.5072836573817 | 0.488 | 0.005 | 0 |
| **HNF4A** | 0 | 0.95208409202536 | 0.525 | 0.019 | 0 |
| **PKIG** | 0 | -1.09784534367785 | 0.899 | 0.988 | 0 |
| **TOMM34** | 0 | -1.1313020111148 | 0.823 | 0.991 | 0 |
| **STK4** | 0 | -1.1284842214379 | 0.871 | 0.992 | 0 |
| **DBNDD2** | 0 | -1.76700595482574 | 0.311 | 0.904 | 0 |
| **PIGT** | 0 | -1.58517546690986 | 0.896 | 1 | 0 |
| **WFDC2** | 0 | -2.97286120501022 | 0.48 | 0.931 | 0 |
| **SULF2** | 0 | -1.72333962529934 | 0.825 | 0.988 | 0 |
| **STAU1** | 0 | -0.839701663839919 | 1 | 1 | 0 |
| **PTGIS** | 0 | -1.25886734685272 | 0.159 | 0.716 | 0 |
| **TMEM189** | 0 | -0.834987003830181 | 0.995 | 1 | 0 |
| **PTPN1** | 0 | -0.859020428081533 | 0.978 | 0.999 | 0 |
| **BMP7** | 0 | -1.77979425530216 | 0.786 | 0.985 | 0 |
| **RAE1** | 0 | -0.941741734602646 | 0.983 | 1 | 0 |
| **NELFCD** | 0 | -1.12232109995014 | 0.99 | 1 | 0 |
| **ATP5F1E** | 0 | -1.79044440178243 | 0.995 | 1 | 0 |
| **CDH4** | 0 | -1.12882857563734 | 0.036 | 0.578 | 0 |

| | | | | | |
|---|---|---|---|---|---|
| **PSMA7** | 0 | -0.836297525718861 | 1 | 1 | 0 |
| **KCNQ2** | 0 | 1.73836932622918 | 0.95 | 0.312 | 0 |
| **SLC2A4RG** | 0 | -0.952700741990528 | 0.895 | 0.995 | 0 |
| **DNAJC5** | 0 | -1.06047252531654 | 0.977 | 1 | 0 |
| **NRIP1** | 0 | -1.25768849170347 | 0.523 | 0.93 | 0 |
| **PDXK** | 0 | 1.08352819564244 | 1 | 0.997 | 0 |
| **DNMT3L** | 0 | 4.75925739358119 | 1 | 0.708 | 0 |
| **COL18A1** | 0 | -2.71631130213375 | 0.87 | 1 | 0 |
| **SPATC1L** | 0 | -1.41038105355206 | 0.1 | 0.794 | 0 |
| **ATP6V1E1** | 0 | 0.891162470234336 | 1 | 0.985 | 0 |
| **TBX1** | 0 | 0.957927687125782 | 0.547 | 0.021 | 0 |
| **MED15** | 0 | 1.37643969360756 | 1 | 0.942 | 0 |
| **SDF2L1** | 0 | 1.76811630581038 | 0.997 | 0.98 | 0 |
| **DDT** | 0 | 1.01337396798293 | 1 | 1 | 0 |
| **SUSD2** | 0 | 3.08133969492132 | 0.996 | 0.641 | 0 |
| **ADORA2A** | 0 | 1.40410754311362 | 0.748 | 0.099 | 0 |
| **TPST2** | 0 | 1.48224529641618 | 0.994 | 0.807 | 0 |
| **CCDC117** | 0 | 1.29355100206058 | 0.986 | 0.673 | 0 |
| **XBP1** | 0 | 1.23921930226074 | 1 | 0.998 | 0 |
| **NEFH** | 0 | 2.70897928377261 | 0.954 | 0.371 | 0 |
| **THOC5** | 0 | 0.859580107481957 | 1 | 0.983 | 0 |
| **NF2** | 0 | 1.16760537465049 | 0.999 | 0.995 | 0 |
| **TCN2** | 0 | -1.36610437446726 | 0.426 | 0.934 | 0 |
| **SELENOM** | 0 | 1.14439527203346 | 0.946 | 0.616 | 0 |
| **LIMK2** | 0 | 1.12848572267959 | 0.998 | 0.955 | 0 |
| **TOMM22** | 0 | 1.08777323908089 | 0.999 | 0.996 | 0 |
| **SYNGR1** | 0 | -1.37535260341215 | 0.459 | 0.901 | 0 |
| **SNU13** | 0 | 0.957113285389866 | 1 | 1 | 0 |
| **NDUFA6** | 0 | 1.12142728354136 | 1 | 0.981 | 0 |
| **A4GALT** | 0 | 2.65106922410202 | 0.99 | 0.666 | 0 |
| **ARFGAP3** | 0 | 1.07934595411087 | 0.961 | 0.668 | 0 |
| **TTLL12** | 0 | 1.12098493019032 | 1 | 0.991 | 0 |
| **PARVB** | 0 | 1.33927965646384 | 0.887 | 0.488 | 0 |
| **ARHGAP8** | 0 | 1.46767088744636 | 0.99 | 0.697 | 0 |

| | | | | | |
|---|---|---|---|---|---|
| **FBLN1** | 0 | -1.11012225525644 | 0.975 | 1 | 0 |
| **CRELD2** | 0 | 1.79120130503992 | 0.996 | 0.934 | 0 |
| **CD99** | 0 | -2.16188056786939 | 0.329 | 0.969 | 0 |
| **MXRA5** | 0 | 1.20476849833741 | 0.808 | 0.227 | 0 |
| **NLGN4X** | 0 | -2.25145684917088 | 0.167 | 0.79 | 0 |
| **TBL1X** | 0 | -1.50179998689292 | 0.375 | 0.925 | 0 |
| **WWC3** | 0 | -1.70907198013255 | 0.594 | 0.976 | 0 |
| **TMSB4X** | 0 | -2.94735463706641 | 1 | 1 | 0 |
| **PIR** | 0 | -1.37130667607099 | 0.547 | 0.943 | 0 |
| **AP1S2** | 0 | -1.20133773803859 | 0.613 | 0.95 | 0 |
| **CTPS2** | 0 | -0.834575405946436 | 0.934 | 0.996 | 0 |
| **RBBP7** | 0 | -0.941655024350911 | 0.96 | 0.997 | 0 |
| **SH3KBP1** | 0 | -1.11822696833657 | 0.806 | 0.981 | 0 |
| **SMS** | 0 | -1.20068239110742 | 0.993 | 1 | 0 |
| **PHEX** | 0 | 1.91328458811796 | 0.887 | 0.041 | 0 |
| **EIF2S3** | 0 | 1.05038825182075 | 0.845 | 0.372 | 0 |
| **PCYT1B** | 0 | -1.71789221310133 | 0.316 | 0.926 | 0 |
| **MAGEB2** | 0 | 1.42372949045874 | 0.574 | 0.026 | 0 |
| **DYNLT3** | 0 | 0.945156266356697 | 0.578 | 0.006 | 0 |
| **CASK** | 0 | -1.0457736314719 | 0.891 | 0.999 | 0 |
| **USP11** | 0 | -0.925647521518329 | 0.963 | 1 | 0 |
| **TIMP1** | 0 | -2.20650917250933 | 0.92 | 1 | 0 |
| **RBM3** | 0 | -1.27240786781871 | 0.995 | 1 | 0 |
| **PIM2** | 0 | 1.58483792371017 | 0.99 | 0.973 | 0 |
| **GSPT2** | 0 | -1.21630346113032 | 0.61 | 0.961 | 0 |
| **MAGED1** | 0 | -1.88674463099805 | 0.966 | 1 | 0 |
| **WNK3** | 0 | -1.0579471453389 | 0.557 | 0.922 | 0 |
| **MAGED2** | 0 | -1.71570524465074 | 0.994 | 1 | 0 |
| **MAGEH1** | 0 | -1.3173431928062 | 0.483 | 0.931 | 0 |
| **UBQLN2** | 0 | -1.19133819878927 | 0.935 | 1 | 0 |
| **ARHGEF9** | 0 | -1.40927114578539 | 0.204 | 0.83 | 0 |
| **AMER1** | 0 | -1.17749316388371 | 0.553 | 0.928 | 0 |
| **ZC4H2** | 0 | -1.10200560198034 | 0.406 | 0.86 | 0 |
| **MSN** | 0 | -1.62878718499476 | 0.975 | 1 | 0 |

| | | | | | |
|---|---|---|---|---|---|
| **HEPH** | 0 | -1.91295799349685 | 0.022 | 0.869 | 0 |
| **EFNB1** | 0 | -2.50545224930554 | 0.433 | 0.974 | 0 |
| **SNX12** | 0 | -1.07368405899913 | 0.932 | 0.999 | 0 |
| **FOXO4** | 0 | -1.59403050006532 | 0.65 | 0.925 | 0 |
| **MED12** | 0 | -1.35087366492227 | 0.17 | 0.804 | 0 |
| **NONO** | 0 | 1.12630377281935 | 0.996 | 0.901 | 0 |
| **PHKA1** | 0 | 2.50710408245494 | 1 | 0.773 | 0 |
| **SLC16A2** | 0 | -1.76076677465808 | 0.653 | 0.987 | 0 |
| **RLIM** | 0 | 0.983162629428777 | 0.67 | 0.121 | 0 |
| **ABCB7** | 0 | -1.05333268353587 | 0.951 | 1 | 0 |
| **COX7B** | 0 | 1.5708635406106 | 0.993 | 0.745 | 0 |
| **TAF9B** | 0 | -1.10986967686481 | 0.602 | 0.928 | 0 |
| **TBX22** | 0 | -1.4387687182488 | 0.023 | 0.595 | 0 |
| **SH3BGRL** | 0 | -1.4437826267711 | 0.581 | 0.968 | 0 |
| **ZNF711** | 0 | -1.16821380664671 | 0.478 | 0.905 | 0 |
| **NAP1L3** | 0 | -1.59067665068865 | 0.363 | 0.894 | 0 |
| **ARMCX2** | 0 | -1.02659688630597 | 0.868 | 0.994 | 0 |
| **TMSB15A** | 0 | -2.03412873656455 | 0.23 | 0.823 | 0 |
| **BEX4** | 0 | -1.6154707858834 | 0.925 | 1 | 0 |
| **TCEAL8** | 0 | -1.13244393835588 | 0.986 | 1 | 0 |
| **TCEAL9** | 0 | -1.65269168302456 | 0.873 | 0.999 | 0 |
| **BEX3** | 0 | -2.3559356301192 | 0.979 | 1 | 0 |
| **NRK** | 0 | -2.89354438063815 | 0.305 | 0.795 | 0 |
| **TSC22D3** | 0 | -1.41600298051418 | 0.479 | 0.902 | 0 |
| **COL4A6** | 0 | -1.36308232914451 | 0.027 | 0.646 | 0 |
| **COL4A5** | 0 | -1.77613294283542 | 0.152 | 0.856 | 0 |
| **ACSL4** | 0 | -1.27030470022284 | 0.804 | 0.996 | 0 |
| **PGRMC1** | 0 | -1.82105025686497 | 0.989 | 1 | 0 |
| **SMARCA1** | 0 | -1.65148642862774 | 0.684 | 0.992 | 0 |
| **MBNL3** | 0 | -1.742261091185 | 0.482 | 0.944 | 0 |
| **HS6ST2** | 0 | -1.9755373806209 | 0.185 | 0.921 | 0 |
| **FAM122B** | 0 | -1.29654975776202 | 0.728 | 0.965 | 0 |
| **LDOC1** | 0 | -1.29513433273061 | 0.624 | 0.963 | 0 |
| **IDS** | 0 | -1.20315823979179 | 0.452 | 0.913 | 0 |

| | | | | | |
|---|---|---|---|---|---|
| **TMEM185A** | 0 | -1.24696459040987 | 0.794 | 0.99 | 0 |
| **VMA21** | 0 | -0.97564289814238 | 0.983 | 1 | 0 |
| **ZNF185** | 0 | -1.46141027329655 | 0.425 | 0.891 | 0 |
| **DUSP9** | 0 | 1.38380715564093 | 0.738 | 0.069 | 0 |
| **SLC6A8** | 0 | 1.90949289454159 | 0.998 | 0.95 | 0 |
| **GDI1** | 0 | -0.836663714306941 | 0.993 | 1 | 0 |
| **DDX3Y** | 0 | 1.11335377824015 | 0.983 | 0.822 | 0 |
| **LYPD1** | 4.6138 | -2.06471622433865 | 0.221 | 0.724 | 7.5561784 |
| **DLG3** | 4.7568 | 0.989524956710147 | 0.995 | 0.988 | 7.7902953 |
| **PREX2** | 1.3369 | -1.29383411525482 | 0.364 | 0.818 | 2.1895462 |
| **C18orf32** | 2.5124 | 1.0237310676994 | 0.831 | 0.363 | 4.1145917 |
| **PAPPA** | 2.6190 | -1.23663486360201 | 0.011 | 0.524 | 4.2892813 |
| **UBASH3B** | 6.5529 | 1.1433089496638 | 0.909 | 0.553 | 1.0731723 |
| **CITED2** | 2.5011 | -1.15077860101403 | 0.839 | 0.99 | 4.0961052 |
| **SFT2D1** | 5.4654 | 1.00478684325464 | 0.913 | 0.563 | 8.9508185 |
| **USO1** | 7.1825 | 0.868279363870145 | 1 | 0.999 | 1.1762869 |
| **GOLM1** | 1.2744 | -0.803088716106298 | 0.995 | 1 | 2.0872048 |
| **UGCG** | 1.6150 | 0.962580790808892 | 1 | 0.979 | 2.6449379 |
| **DNAJB6** | 2.9132 | 0.926927167120022 | 0.993 | 0.949 | 4.7710318 |
| **PTPRR** | 3.3557 | 0.850194075561441 | 0.501 | 0.016 | 5.4957149 |
| **UBE2N** | 8.5709 | 0.967360054690928 | 0.91 | 0.542 | 1.4036664 |
| **PDE12** | 8.7330 | 0.994921216512209 | 0.894 | 0.497 | 1.4302155 |
| **EEF1A2** | 9.9486 | 0.911945455468263 | 0.615 | 0.086 | 1.6292900 |
| **CBLC** | 1.1133 | 0.9162460122389 | 0.638 | 0.105 | 1.8232833 |
| **PGM2** | 7.3690 | 1.13673240467404 | 0.981 | 0.871 | 1.2068232 |
| **ATP8B1** | 9.2771 | 0.870440394275855 | 0.506 | 0.018 | 1.5193264 |
| **MAP4K1** | 3.9378 | 1.17009281263696 | 0.771 | 0.268 | 6.4490643 |
| **UACA** | 4.1343 | 1.04012538087395 | 0.974 | 0.832 | 6.7707574 |
| **SPC24** | 6.5857 | -0.986023893111723 | 0.87 | 0.981 | 1.0785455 |
| **BACE2** | 8.0434 | -1.20552399851906 | 0.42 | 0.886 | 1.3172761 |
| **PCDH10** | 1.2872 | -2.44291451236944 | 0.05 | 0.563 | 2.1081675 |
| **LINGO1** | 2.1788 | 1.00337228065401 | 0.991 | 0.854 | 3.5682380 |
| **FGD4** | 3.2176 | 1.11222221252088 | 0.969 | 0.708 | 5.2694669 |
| **FLT1** | 5.2961 | -1.67967334734896 | 0.154 | 0.673 | 8.6734553 |

| | | | | | |
|---|---|---|---|---|---|
| **PRKAB2** | 8.1757 | 1.01801141404176 | 0.975 | 0.819 | 1.3389345 |
| **COPRS** | 9.6590 | -0.883251005747416 | 0.937 | 0.996 | 1.5818631 |
| **C6orf132** | 1.0919 | 1.03780496967768 | 0.993 | 0.82 | 1.7882564 |
| **PTGR1** | 1.5098 | 0.929966173931053 | 0.981 | 0.811 | 2.4727537 |
| **SMG6** | 2.5809 | -0.987026215809145 | 0.663 | 0.94 | 4.2268590 |
| **SACS** | 8.3583 | -1.0226258981292 | 0.885 | 0.987 | 1.3688467 |
| **MAP3K13** | 1.6188 | 1.08057161409518 | 0.903 | 0.549 | 2.6512615 |
| **CIB1** | 2.1704 | 0.920158917628002 | 0.978 | 0.833 | 3.5545095 |
| **RAP1GDS1** | 3.2832 | -0.911804902296945 | 0.804 | 0.984 | 5.3770514 |
| **HIGD1A** | 5.8134 | 0.958783652204498 | 0.928 | 0.595 | 9.5206796 |
| **NFKB2** | 6.4163 | 1.3521592099058 | 0.968 | 0.834 | 1.0508114 |
| **SEMA4B** | 7.1287 | 0.977077473953749 | 0.983 | 0.82 | 1.1674754 |
| **PI4K2B** | 1.6632 | 0.969272312947542 | 0.978 | 0.83 | 2.7238837 |
| **STK26** | 3.0407 | -0.915868017074907 | 0.927 | 0.995 | 4.9798815 |
| **MANEAL** | 6.0057 | 1.03826644903287 | 0.979 | 0.769 | 9.8356038 |
| **HCN2** | 3.0952 | 0.999093116648359 | 0.965 | 0.708 | 5.0690610 |
| **ZSCAN2** | 3.9619 | 1.19699915681916 | 0.972 | 0.892 | 6.4884421 |
| **UBE2D1** | 1.4688 | -0.951659019991014 | 0.622 | 0.93 | 2.4055686 |
| **PCGF6** | 6.6610 | 1.00497184241355 | 0.927 | 0.638 | 1.0908760 |
| **IFITM3** | 1.3884 | -0.855483437286515 | 0.997 | 1 | 2.2737988 |
| **RBMXL1** | 1.7400 | 0.976693610047457 | 0.868 | 0.435 | 2.8496400 |
| **ZNF385A** | 1.3546 | -1.10681364811425 | 0.218 | 0.736 | 2.2185681 |
| **NOTCH2** | 2.4877 | -1.02384128402727 | 0.608 | 0.922 | 4.0741253 |
| **LPIN1** | 1.8614 | 1.06862946661202 | 0.831 | 0.323 | 3.0485090 |
| **AGRN** | 1.9149 | -0.982213303679756 | 0.981 | 0.999 | 3.1360491 |
| **BRSK2** | 2.5134 | 0.908324645124883 | 0.593 | 0.084 | 4.1163547 |
| **ADK** | 5.1741 | -0.927582024513231 | 0.816 | 0.975 | 8.4737439 |
| **MDM2** | 6.1453 | 1.20555720219593 | 1 | 0.987 | 1.0064267 |
| **EIF3E** | 1.3516 | 0.972531221566151 | 0.712 | 0.201 | 2.2136078 |
| **GSTM3** | 2.4443 | -1.42363988436251 | 0.192 | 0.694 | 4.0030348 |
| **ZIC3** | 1.3745 | -1.25375857935371 | 0.695 | 0.942 | 2.2511051 |
| **PAK2** | 3.5763 | 0.901073224830961 | 0.977 | 0.838 | 5.8569835 |
| **PLEC** | 4.4041 | 0.888111571325068 | 0.996 | 0.954 | 7.2127118 |
| **TMEM200A** | 5.4297 | 0.86041428717061 | 0.533 | 0.043 | 8.8923199 |

| | | | | | |
|---|---|---|---|---|---|
| **FAM104B** | 6.2698 | -0.999648268284094 | 0.652 | 0.941 | 1.0268114 |
| **ASCC2** | 7.7449 | 0.800196641581098 | 0.998 | 0.949 | 1.2683908 |
| **CFL2** | 1.5803 | -0.993672442904064 | 0.709 | 0.95 | 2.5880896 |
| **UHRF2** | 2.4128 | -1.12183200472263 | 0.359 | 0.819 | 3.9514969 |
| **B4GALT5** | 1.4879 | -0.81800263049971 | 0.982 | 1 | 2.4368935 |
| **TSPAN6** | 2.2295 | -0.83550799116885 | 0.984 | 0.999 | 3.6513598 |
| **SLC35B2** | 1.0888 | -0.899966008712821 | 0.838 | 0.985 | 1.7831449 |
| **ELF3** | 1.1063 | 1.38891304534483 | 0.823 | 0.366 | 1.8118401 |
| **RBM7** | 1.9572 | 0.93372169995217 | 0.958 | 0.726 | 3.2053715 |
| **XXYLT1** | 2.9913 | -0.903265586990841 | 0.807 | 0.969 | 4.8990060 |
| **SLC25A24** | 3.5344 | -0.863309462650212 | 0.931 | 0.996 | 5.7884156 |
| **ANK2** | 4.5519 | -1.09145681911442 | 0.641 | 0.933 | 7.4546905 |
| **BTBD2** | 5.4707 | -0.840365103805628 | 0.974 | 0.999 | 8.9594260 |
| **NOTCH1** | 7.3601 | -1.02445291285801 | 0.839 | 0.979 | 1.2053685 |
| **NIPAL1** | 7.8530 | 1.07031163706477 | 0.682 | 0.171 | 1.2860914 |
| **ZNF229** | 8.4355 | 1.00153087236349 | 0.756 | 0.255 | 1.3814878 |
| **PPME1** | 2.5768 | -0.979328762582093 | 0.737 | 0.959 | 4.2201004 |
| **ECE1** | 2.7708 | -1.01028674745321 | 0.917 | 0.997 | 4.5378219 |
| **ARHGDIG** | 3.1531 | 0.934193097067811 | 0.629 | 0.116 | 5.1638939 |
| **PTPRA** | 1.0025 | -0.873124272556603 | 0.896 | 0.987 | 1.6418758 |
| **TSKU** | 2.0220 | -1.19212764418665 | 0.938 | 0.996 | 3.3115714 |
| **LEFTY2** | 3.1823 | 1.68626107690049 | 0.99 | 0.945 | 5.2117145 |
| **LLPH** | 1.0313 | 0.958166863341551 | 0.849 | 0.438 | 1.6889765 |
| **PARP8** | 1.7261 | -0.981810125949917 | 0.098 | 0.619 | 2.8268374 |
| **SLC7A7** | 3.6203 | 1.35537832947896 | 0.89 | 0.57 | 5.9290116 |
| **UTRN** | 4.4200 | -1.16511422769841 | 0.606 | 0.933 | 7.2386960 |
| **KRTCAP3** | 6.2457 | 1.01661400126848 | 0.708 | 0.188 | 1.0228656 |
| **TTI1** | 1.7229 | -0.808087635461974 | 0.941 | 0.994 | 2.8216936 |
| **EPDR1** | 2.5409 | -0.921467245911121 | 0.035 | 0.532 | 4.1612865 |
| **AMOTL2** | 2.5560 | -0.839021188183042 | 0.994 | 0.999 | 4.1860764 |
| **KIF26A** | 3.4545 | -0.931655336539785 | 0.906 | 0.996 | 5.6574490 |
| **RAVER2** | 6.5438 | -0.986405368327781 | 0.507 | 0.879 | 1.0716944 |
| **BRSK1** | 7.0827 | -1.0184224985987 | 0.147 | 0.664 | 1.1599486 |
| **IGF1R** | 7.8517 | -0.801885912264954 | 0.987 | 1 | 1.2858879 |

| | | | | | |
|---|---|---|---|---|---|
| **PINK1** | 2.0741 | 0.966015291171789 | 0.994 | 0.94 | 3.3967861 |
| **PIK3AP1** | 2.3450 | 0.856487642589284 | 0.536 | 0.051 | 3.8404505 |
| **GMFB** | 5.3107 | 0.840022172440668 | 0.6 | 0.098 | 8.6974953 |
| **DLGAP4** | 6.8909 | -0.853562695684472 | 0.957 | 0.996 | 1.1285274 |
| **AGBL5** | 8.7293 | -0.913729005416365 | 0.838 | 0.978 | 1.4295997 |
| **SYNGAP1** | 9.5955 | -1.24703415488438 | 0.332 | 0.78 | 1.5714703 |
| **KCTD14** | 1.0434 | -1.01770437407929 | 0.185 | 0.698 | 1.7088990 |
| **ACE** | 1.8814 | 1.03885350177732 | 0.725 | 0.2 | 3.0812926 |
| **USP3** | 1.9094 | -0.963094332454336 | 0.056 | 0.558 | 3.1271430 |
| **CER1** | 2.2427 | -4.02223168595711 | 0.165 | 0.695 | 3.6729066 |
| **PKIA** | 1.5200 | -0.871836275074261 | 0.038 | 0.533 | 2.4893634 |
| **ZNF614** | 5.8637 | 0.936268146656461 | 0.948 | 0.697 | 9.6030433 |
| **SHISA9** | 8.3342 | -1.30027973850767 | 0.242 | 0.746 | 1.3648929 |
| **ZNF808** | 9.7269 | 0.906811216514295 | 0.982 | 0.835 | 1.5929851 |
| **FRMD6** | 1.5703 | 1.10817123430982 | 0.957 | 0.728 | 2.5717824 |
| **PHLDB2** | 1.6377 | -1.26343591296899 | 0.537 | 0.891 | 2.6821819 |
| **RASSF9** | 3.9745 | -1.43676463143681 | 0.149 | 0.659 | 6.5091027 |
| **SALL3** | 1.3293 | -1.17133396212991 | 0.767 | 0.973 | 2.1771123 |
| **ABCB8** | 3.1396 | 0.898397667301274 | 0.985 | 0.891 | 5.1418145 |
| **HSPA1B** | 4.6377 | 1.42331802177327 | 0.832 | 0.459 | 7.5952189 |
| **GPRC5A** | 8.6831 | 1.56759302884365 | 0.527 | 0.049 | 1.4220470 |
| **ZNF682** | 1.3854 | 1.07210544233178 | 0.844 | 0.414 | 2.2688923 |
| **HIPK1** | 1.6691 | -0.826005989318636 | 0.869 | 0.978 | 2.7335665 |
| **SLC9A3R2** | 1.8302 | 1.06469752196365 | 0.936 | 0.65 | 2.9973666 |
| **FES** | 3.8827 | 0.943817728715874 | 0.818 | 0.345 | 6.3587947 |
| **PLEKHG3** | 8.6853 | -1.18694243659061 | 0.493 | 0.889 | 1.4223968 |
| **EEA1** | 9.7786 | 0.953679761017121 | 0.959 | 0.768 | 1.6014547 |
| **STARD8** | 1.0970 | -1.24487728998856 | 0.36 | 0.829 | 1.7966148 |
| **LDLRAP1** | 1.0992 | 0.918540436897975 | 0.992 | 0.884 | 1.8002300 |
| **SERTAD4** | 1.3727 | -1.15049748597563 | 0.113 | 0.622 | 2.2481088 |
| **GNPDA1** | 1.6395 | -0.855733611422121 | 0.903 | 0.987 | 2.6850835 |
| **ANGPTL2** | 2.8114 | -0.968092639123549 | 0.054 | 0.55 | 4.6043288 |
| **BRWD3** | 5.7912 | 1.04168434911885 | 0.992 | 0.978 | 9.4843962 |
| **SH3BGRL3** | 9.2342 | -0.962370762837581 | 0.96 | 0.997 | 1.5122858 |

| | | | | | |
|---|---|---|---|---|---|
| **TNFRSF8** | 9.6727 | 0.919986642544917 | 0.489 | 0.029 | 1.5841133 |
| **TRIM44** | 1.1414 | -0.873758337865403 | 0.821 | 0.98 | 1.8694277 |
| **NCAPD2** | 1.5184 | -0.995666869358448 | 0.962 | 0.998 | 2.4867411 |
| **KCNMB1** | 3.2386 | -1.72758270067438 | 0.044 | 0.532 | 5.3039075 |
| **MAML3** | 3.6443 | -1.43458557768964 | 0.218 | 0.718 | 5.9683665 |
| **SDCBP** | 5.6230 | 0.80361371364198 | 0.998 | 0.974 | 9.2089213 |
| **ANKRD18A** | 1.8005 | -1.16473430393226 | 0.182 | 0.683 | 2.9487930 |
| **ASPH** | 2.0090 | -0.839407188746403 | 0.94 | 0.997 | 3.2901767 |
| **OAT** | 3.7850 | 1.06287394598812 | 0.865 | 0.488 | 6.1987076 |
| **FN3K** | 4.6993 | 1.07423607475366 | 0.915 | 0.6 | 7.6961831 |
| **SLC35F6** | 9.2732 | 0.820231661728449 | 0.998 | 0.97 | 1.5186792 |
| **KLF17** | 1.8521 | 1.28368074150681 | 0.439 | 0.004 | 3.0331886 |
| **PIK3R1** | 3.6472 | -1.33167871787197 | 0.369 | 0.798 | 5.9730278 |
| **JAG2** | 1.0241 | 1.02628703456967 | 0.73 | 0.233 | 1.6772979 |
| **LARGE2** | 1.3488 | 0.958562903446432 | 0.981 | 0.789 | 2.2090695 |
| **SLCO4C1** | 4.3394 | 1.03387222174436 | 0.933 | 0.603 | 7.1067014 |
| **CD24** | 7.4722 | 1.09480503104469 | 0.998 | 0.983 | 1.2237333 |
| **MARVELD3** | 1.1983 | 0.916338371038126 | 0.712 | 0.2 | 1.9626012 |
| **MAPK13** | 3.5065 | 1.01102328557378 | 0.712 | 0.222 | 5.7426631 |
| **ZNF296** | 6.7261 | 1.24890694129842 | 0.865 | 0.518 | 1.1015403 |
| **MCM5** | 8.6137 | 0.805210891018496 | 1 | 0.995 | 1.4106816 |
| **ARRDC2** | 8.9944 | 1.0746024901484 | 0.791 | 0.33 | 1.4730178 |
| **STK40** | 9.0103 | -0.921394204512146 | 0.748 | 0.959 | 1.4756245 |
| **DRAXIN** | 1.0433 | -1.43552981338206 | 0.815 | 0.95 | 1.7086506 |
| **STARD3NL** | 1.2230 | -0.813007030671275 | 0.903 | 0.993 | 2.0029686 |
| **MBOAT1** | 1.4907 | 1.00227956647333 | 0.901 | 0.517 | 2.4414097 |
| **ITPKB** | 2.8008 | -0.927055454589171 | 0.049 | 0.535 | 4.5870206 |
| **WDR35** | 6.3101 | -1.01418786850896 | 0.277 | 0.748 | 1.0334115 |
| **BCL7C** | 1.0933 | -1.05983239530971 | 0.899 | 0.992 | 1.7906206 |
| **HNMT** | 3.7095 | -0.999920086119519 | 0.208 | 0.7 | 6.0751744 |
| **KCNG3** | 6.8631 | 0.947217929852423 | 0.562 | 0.083 | 1.1239850 |
| **ZRANB3** | 7.3392 | -1.00095540998639 | 0.285 | 0.755 | 1.2019484 |
| **TBL2** | 7.5791 | -0.933397909464577 | 0.707 | 0.943 | 1.2412384 |
| **PSRC1** | 1.2594 | 1.06664820112935 | 0.97 | 0.707 | 2.0626822 |

| | | | | | |
|---|---|---|---|---|---|
| **IGDCC4** | 1.5230 | -0.852295789539588 | 0.033 | 0.509 | 2.4943593 |
| **SLC23A2** | 1.7554 | 1.09034572191741 | 0.969 | 0.845 | 2.8749151 |
| **APPL1** | 4.9090 | 0.821101841228844 | 0.988 | 0.91 | 8.0395754 |
| **NIN** | 5.2975 | -0.9992599070436 | 0.687 | 0.941 | 8.6757691 |
| **EDEM1** | 7.1376 | 0.944010334712337 | 0.987 | 0.891 | 1.1689325 |
| **ACTN1** | 1.0270 | -0.815162017787059 | 1 | 1 | 1.6819951 |
| **CALM1** | 1.9096 | 0.884492548329577 | 0.967 | 0.793 | 3.1274333 |
| **LIMD2** | 3.2019 | -0.965348407429471 | 0.889 | 0.982 | 5.2438971 |
| **AGAP3** | 4.5881 | -0.954489467732642 | 0.269 | 0.758 | 7.5140071 |
| **FST** | 6.0396 | -2.28581134791739 | 0.313 | 0.743 | 9.8911502 |
| **CENPV** | 8.6860 | -1.10377318280413 | 0.866 | 0.98 | 1.4225092 |
| **NAPRT** | 2.7914 | 0.944849016486127 | 0.879 | 0.471 | 4.5716236 |
| **BAIAP2L1** | 1.2445 | 0.832070805452831 | 0.99 | 0.927 | 2.0381826 |
| **RPUSD4** | 1.5472 | -0.831299503515221 | 0.847 | 0.988 | 2.5338941 |
| **RAB31** | 1.6307 | -0.973207255022178 | 0.364 | 0.802 | 2.6706641 |
| **FBXL16** | 1.9236 | 0.924773324601551 | 0.63 | 0.142 | 3.1503125 |
| **ZNF778** | 3.2355 | 0.941358577384677 | 0.897 | 0.581 | 5.2988678 |
| **TMEM131L** | 3.4828 | 0.985748305331658 | 0.965 | 0.798 | 5.7039065 |
| **MIXL1** | 3.4975 | -3.05072864630175 | 0.743 | 0.93 | 5.7279749 |
| **NR6A1** | 3.8426 | -1.08237597042587 | 0.924 | 0.983 | 6.2930768 |
| **SLC7A8** | 4.1880 | 1.04493897228866 | 0.985 | 0.841 | 6.8587906 |
| **TSPAN13** | 4.4715 | 0.889307548370075 | 0.985 | 0.857 | 7.3230827 |
| **SCML2** | 4.5774 | -0.91245834059101 | 0.614 | 0.921 | 7.4965221 |
| **POMGNT1** | 5.5832 | -0.813379782977713 | 0.878 | 0.988 | 9.1436175 |
| **ZNF770** | 5.8719 | 0.83631993042825 | 0.61 | 0.125 | 9.6164840 |
| **MPV17L** | 6.6248 | 0.944359663300922 | 0.687 | 0.205 | 1.0849558 |
| **PBX4** | 9.6209 | 0.935863508429331 | 0.638 | 0.144 | 1.5756254 |
| **EPS8L1** | 2.4056 | 1.00024307175894 | 0.779 | 0.32 | 3.9397212 |
| **PM20D2** | 6.1098 | -1.02394661992438 | 0.166 | 0.658 | 1.0006080 |
| **TTYH1** | 1.0768 | 1.16613068046669 | 0.546 | 0.076 | 1.7635283 |
| **ST5** | 1.9777 | -0.896986699763696 | 0.789 | 0.973 | 3.2389825 |
| **FAM102B** | 2.0765 | 1.10667738266729 | 0.905 | 0.644 | 3.4007223 |
| **FAM107B** | 2.3184 | 0.96275998103194 | 0.974 | 0.803 | 3.7969086 |
| **OLFM1** | 2.7585 | -1.26222722934343 | 0.652 | 0.913 | 4.5176803 |

| | | | | | |
|---|---|---|---|---|---|
| **TNS2** | 6.7752 | -1.25424198650606 | 0.073 | 0.55 | 1.1095761 |
| **SRMS** | 4.4226 | 0.879859990891956 | 0.46 | 0.022 | 7.2429085 |
| **SYT6** | 4.8535 | -1.09989193780672 | 0.212 | 0.695 | 7.9486105 |
| **DERL3** | 2.1439 | 1.05128326016893 | 0.649 | 0.171 | 3.5110752 |
| **TBC1D4** | 4.6894 | -1.02609331076808 | 0.457 | 0.856 | 7.6799669 |
| **TRIOBP** | 5.5673 | -0.844026830240712 | 0.878 | 0.985 | 9.1176322 |
| **PWWP3A** | 9.2748 | 0.865097034655567 | 0.994 | 0.952 | 1.5189483 |
| **FGFBP3** | 1.4667 | -1.00749972437168 | 0.017 | 0.472 | 2.4021274 |
| **LDB2** | 3.2230 | -1.48793186736683 | 0.607 | 0.882 | 5.2783272 |
| **TRIM24** | 5.8127 | -0.86904341520837 | 0.994 | 1 | 9.5194681 |
| **C10orf95** | 9.6423 | 0.94660230616875 | 0.495 | 0.045 | 1.5791223 |
| **CHSY1** | 1.0348 | -0.881475024930042 | 0.976 | 1 | 1.6947302 |
| **MME** | 1.1175 | -1.17666479395624 | 0.2 | 0.674 | 1.8301366 |
| **INA** | 2.3806 | 1.0270728726295 | 0.923 | 0.614 | 3.8988251 |
| **AKT3** | 2.9336 | -1.04728255315167 | 0.524 | 0.88 | 4.8044898 |
| **TMEM144** | 3.6870 | 1.01150878671307 | 0.615 | 0.138 | 6.0382536 |
| **ST13** | 1.2826 | 0.8004759732055 | 0.977 | 0.811 | 2.1006710 |
| **ISLR** | 1.2937 | -1.12939041275457 | 0.022 | 0.477 | 2.1187919 |
| **TMEM154** | 1.6476 | 1.18555738533139 | 0.779 | 0.336 | 2.6983018 |
| **PDS5A** | 1.6633 | 0.816489942333517 | 0.978 | 0.843 | 2.7239966 |
| **PXDN** | 2.1904 | 0.855363442987982 | 1 | 0.912 | 3.5873053 |
| **KATNAL1** | 2.4077 | -0.953284198355426 | 0.433 | 0.835 | 3.9432472 |
| **DPP6** | 2.4591 | 0.988668834891587 | 0.605 | 0.129 | 4.0274172 |
| **SARS** | 2.8496 | 0.823550025794784 | 0.98 | 0.86 | 4.6668185 |
| **OAF** | 3.3897 | -1.0035193385978 | 0.046 | 0.514 | 5.5513412 |
| **LIMCH1** | 5.7419 | -1.49316842591107 | 0.401 | 0.803 | 9.4036547 |
| **JAKMIP2** | 6.0566 | -1.01557063716715 | 0.032 | 0.492 | 9.9189646 |
| **RASL10B** | 6.9171 | -1.14161100712126 | 0.247 | 0.717 | 1.1328174 |
| **PHLDB1** | 6.9410 | -0.904157091964733 | 0.717 | 0.949 | 1.1367349 |
| **CHORDC1** | 9.2296 | 0.927221934059776 | 0.782 | 0.339 | 1.5115387 |
| **GYG2** | 1.0049 | -0.913191723068813 | 0.805 | 0.983 | 1.6458047 |
| **ELP5** | 1.0462 | -0.844492057924357 | 0.862 | 0.978 | 1.7134982 |
| **FAM217B** | 1.0685 | -0.869076064240247 | 0.731 | 0.944 | 1.7499132 |
| **ZNF589** | 1.6812 | 0.997191875517112 | 0.982 | 0.904 | 2.7534628 |

| | | | | | |
|---|---|---|---|---|---|
| **CREBL2** | 1.8764 | 0.957145338634313 | 0.977 | 0.877 | 3.0731056 |
| **CNN1** | 2.4104 | -1.86744343667746 | 0.602 | 0.904 | 3.9476322 |
| **P2RY1** | 2.6602 | -1.29295651700179 | 0.111 | 0.586 | 4.3566467 |
| **AFF2** | 4.7707 | 1.17942186172198 | 0.701 | 0.235 | 7.8129937 |
| **C20orf96** | 5.0345 | -0.881970702496457 | 0.086 | 0.566 | 8.2450876 |
| **APOBEC3D** | 1.0111 | 0.849435017432625 | 0.604 | 0.122 | 1.6559596 |
| **SLC25A38** | 1.2775 | 0.809410957412523 | 0.986 | 0.913 | 2.0921864 |
| **HS2ST1** | 1.4092 | -0.811194374369696 | 0.838 | 0.978 | 2.3079424 |
| **TMC6** | 1.4295 | -1.14883248002292 | 0.686 | 0.91 | 2.3412507 |
| **CTSV** | 2.1154 | -1.15440661095081 | 0.813 | 0.971 | 3.4645350 |
| **HSBP1L1** | 3.9341 | 0.80843817206236 | 0.517 | 0.059 | 6.4430178 |
| **ZBTB12** | 5.3835 | -0.956219105637056 | 0.141 | 0.623 | 8.8166111 |
| **CAP2** | 1.8757 | -0.964995619369269 | 0.695 | 0.947 | 3.0718844 |
| **HERPUD2** | 2.5824 | -0.888760566549624 | 0.671 | 0.929 | 4.2292961 |
| **CLDN7** | 2.6834 | 0.971048150210534 | 0.992 | 0.743 | 4.3946690 |
| **DMXL1** | 3.0941 | 0.951689894045447 | 0.922 | 0.672 | 5.0672625 |
| **SYS1** | 3.2302 | -0.87341711161046 | 0.732 | 0.947 | 5.2901336 |
| **GATAD2B** | 6.1932 | -0.812089014203734 | 0.838 | 0.976 | 1.0142693 |
| **HOXA7** | 6.2073 | -0.808087022348342 | 0.072 | 0.547 | 1.0165767 |
| **LAMB2** | 1.2396 | -1.11507546112582 | 0.63 | 0.904 | 2.0301575 |
| **FBXO17** | 3.1157 | -0.963857688854621 | 0.445 | 0.841 | 5.1026045 |
| **SYNDIG1** | 6.2869 | -0.933898554018874 | 0.035 | 0.492 | 1.0296068 |
| **MPP1** | 1.0946 | -0.984611982891059 | 0.493 | 0.866 | 1.7927821 |
| **CTSZ** | 2.3339 | -1.04614660481961 | 0.749 | 0.934 | 3.8223881 |
| **LRIG1** | 2.3340 | 0.84817813417774 | 0.99 | 0.97 | 3.8224255 |
| **SCG3** | 5.1403 | -1.04951841266976 | 0.29 | 0.744 | 8.4183946 |
| **PTAR1** | 6.6081 | 0.856290010557489 | 0.993 | 0.956 | 1.0822244 |
| **AFAP1L1** | 2.5919 | 0.937065008551008 | 0.671 | 0.187 | 4.2447941 |
| **EFNA2** | 2.9125 | 0.803621472484751 | 0.578 | 0.1 | 4.7698855 |
| **DNMT3B** | 3.1409 | 1.13603220454351 | 1 | 1 | 5.1439357 |
| **LSS** | 4.2890 | -0.803596132528283 | 0.983 | 0.998 | 7.0241460 |
| **CDIP1** | 1.8969 | -0.8792318115803 | 0.888 | 0.988 | 3.1066240 |
| **EGLN1** | 2.2131 | -0.919814088944149 | 0.842 | 0.985 | 3.6244846 |
| **CCSAP** | 2.8788 | 0.964338477936006 | 0.826 | 0.412 | 4.7147452 |

| | | | | | |
|---|---|---|---|---|---|
| **ZNF260** | 1.3727 | 0.89065189142276 | 0.814 | 0.397 | 2.2480924 |
| **RTL8A** | 1.7623 | -1.01411529331324 | 0.409 | 0.8 | 2.8862339 |
| **PIFO** | 1.9540 | 0.971461922259055 | 0.892 | 0.49 | 3.2000801 |
| **CTSK** | 3.3124 | -2.01562703623181 | 0.199 | 0.647 | 5.4248754 |
| **TMLHE** | 3.3299 | -0.97506709504745 | 0.359 | 0.781 | 5.4535400 |
| **ZNF618** | 5.9682 | -0.852466933874123 | 0.817 | 0.964 | 9.7742639 |
| **HUNK** | 7.0995 | -1.02426000711229 | 0.503 | 0.862 | 1.1626852 |
| **USP47** | 1.5673 | -0.835663567834555 | 0.823 | 0.971 | 2.5668726 |
| **EPOP** | 1.7749 | -1.15025388726309 | 0.476 | 0.835 | 2.9068119 |
| **DPY19L1** | 2.5373 | 0.971754583039374 | 0.861 | 0.49 | 4.1554882 |
| **ULK4** | 6.2546 | 0.988078621763621 | 0.909 | 0.609 | 1.0243237 |
| **NDUFAF7** | 2.7442 | 0.859474610680651 | 0.969 | 0.817 | 4.4941881 |
| **ZDHHC23** | 2.9310 | 0.826897206925591 | 0.538 | 0.082 | 4.8001263 |
| **TP53BP2** | 5.0511 | -0.81545375049651 | 0.836 | 0.971 | 8.2722494 |
| **TMEM184C** | 7.9350 | 0.891770257917597 | 0.85 | 0.508 | 1.2995152 |
| **ELMOD2** | 2.3688 | -0.88302406251616 | 0.542 | 0.882 | 3.8795165 |
| **LAD1** | 2.8945 | 0.85605561368957 | 0.567 | 0.105 | 4.7403482 |
| **ETV5** | 4.0834 | -1.21718607937996 | 0.359 | 0.771 | 6.6875112 |
| **WDR5** | 4.5683 | 0.834011506948262 | 0.967 | 0.841 | 7.4815238 |
| **CEMIP2** | 1.3970 | 0.894217880525002 | 0.94 | 0.698 | 2.2880233 |
| **SLC16A3** | 3.9248 | 1.07378181579108 | 0.98 | 0.76 | 6.4277602 |
| **CEBPZOS** | 4.1748 | 0.862969176843055 | 0.969 | 0.832 | 6.8372321 |
| **FAM111B** | 5.4333 | -1.32427706658652 | 0.445 | 0.821 | 8.8982580 |
| **CDC42EP1** | 1.4040 | -0.922632316239594 | 0.152 | 0.654 | 2.2993642 |
| **EFCAB10** | 1.8785 | 0.837327146997718 | 0.614 | 0.141 | 3.0764480 |
| **PDIA5** | 4.3465 | 0.835767155042735 | 0.981 | 0.904 | 7.1183066 |
| **DDR1** | 5.0311 | -0.835194415714832 | 0.885 | 0.985 | 8.2395759 |
| **CAV1** | 5.4866 | -1.47636846577243 | 0.411 | 0.835 | 8.9854371 |
| **TMEM54** | 5.7397 | 0.981158948243811 | 0.769 | 0.303 | 9.3999939 |
| **SEPTIN5** | 6.8775 | 0.931590676109199 | 0.683 | 0.213 | 1.1263349 |
| **B3GNT2** | 1.1762 | -1.07786125604818 | 0.946 | 0.989 | 1.9262648 |
| **CNKSR3** | 5.0894 | -1.16647385329889 | 0.096 | 0.559 | 8.3349649 |
| **LUZP4** | 7.2682 | 0.875484336496132 | 0.405 | 0.006 | 1.1903134 |
| **FOXO1** | 2.3228 | -1.33182693034702 | 0.716 | 0.93 | 3.8041507 |

| | | | | | |
|---|---|---|---|---|---|
| **SDC1** | 3.3097 | -1.02349300235989 | 0.304 | 0.754 | 5.4204588 |
| **CPXM1** | 6.3437 | -1.05634380069704 | 0.888 | 0.984 | 1.0389189 |
| **TRO** | 1.1021 | -0.905213758261004 | 0.734 | 0.95 | 1.8049370 |
| **ARX** | 1.8147 | 0.837912001956781 | 0.399 | 0.004 | 2.9720203 |
| **CHMP7** | 2.0826 | -0.827199029432281 | 0.767 | 0.953 | 3.4107634 |
| **DNAJA1** | 5.1661 | 0.81835869789219 | 0.992 | 0.908 | 8.4605654 |
| **DUSP7** | 5.4255 | 0.907648898719499 | 0.957 | 0.77 | 8.8853534 |
| **HIP1R** | 9.9554 | 0.88191625200194 | 0.762 | 0.305 | 1.6304056 |
| **EXPH5** | 1.2591 | 0.840327003507428 | 0.521 | 0.077 | 2.0620587 |
| **OVCH2** | 1.4624 | -0.935297151244299 | 0.012 | 0.439 | 2.3950914 |
| **TSC22D1** | 1.5806 | 0.938320900021721 | 0.829 | 0.441 | 2.5886238 |
| **EFEMP2** | 2.4152 | -0.952949021331464 | 0.24 | 0.695 | 3.9553786 |
| **RPA3** | 3.8399 | 0.890888849108427 | 0.757 | 0.325 | 6.2886603 |
| **LAMA5** | 1.1606 | -0.951393742432685 | 0.989 | 0.996 | 1.9008204 |
| **KLRG2** | 1.8855 | 0.841885573957925 | 0.995 | 0.925 | 3.0879046 |
| **MT-ND3** | 5.2767 | 1.69894209587468 | 0.916 | 0.722 | 8.6416807 |
| **MT-CO3** | 7.0427 | -1.1490619595672 | 0.952 | 1 | 1.1533901 |
| **SKIL** | 7.4892 | 0.871812507790552 | 0.999 | 0.992 | 1.2265168 |
| **TCF15** | 9.8248 | -0.907935506796355 | 0.134 | 0.597 | 1.6090147 |
| **ST6GALNAC6** | 4.5293 | -0.962137617079843 | 0.4 | 0.806 | 7.4176960 |
| **HIPK2** | 5.0758 | -0.99059825053842 | 0.689 | 0.919 | 8.3126580 |
| **SGCB** | 9.6152 | -0.840489322503635 | 0.737 | 0.95 | 1.5746927 |
| **CREG1** | 1.0343 | -0.92850415926349 | 0.56 | 0.886 | 1.6938794 |
| **SNCG** | 1.1678 | -0.97074490555607 | 0.096 | 0.553 | 1.9126670 |
| **DERA** | 3.4126 | -0.817257577354087 | 0.857 | 0.983 | 5.5889158 |
| **LRP8** | 7.1751 | -1.17369888639317 | 0.886 | 0.972 | 1.1750753 |
| **PITPNM2** | 7.9158 | 0.921675207244116 | 0.927 | 0.626 | 1.2963851 |
| **TMEM87A** | 8.6934 | 0.853030744730219 | 0.907 | 0.64 | 1.4237265 |
| **SLC39A14** | 1.2441 | 0.920470097770128 | 0.936 | 0.741 | 2.0375940 |
| **AMZ2** | 1.7772 | 0.8230531867676 | 0.925 | 0.677 | 2.9106470 |
| **TSPAN18** | 3.0362 | -1.18837196440596 | 0.966 | 0.973 | 4.9724036 |
| **PLAGL2** | 9.4295 | -0.838089902220533 | 0.701 | 0.93 | 1.5442764 |
| **PLPP2** | 2.3094 | 0.93106066106652 | 0.467 | 0.048 | 3.7821456 |
| **ST14** | 3.9536 | 0.93981226931541 | 0.655 | 0.193 | 6.4749181 |

| | | | | | |
|---|---|---|---|---|---|
| **PIK3R5** | 5.0071 | 0.832807562786397 | 0.63 | 0.165 | 8.20026575 |
| **AHDC1** | 5.2164 | -1.03924450102343 | 0.559 | 0.875 | 8.54297709 |
| **NKD1** | 6.9805 | -1.47279539507041 | 0.232 | 0.664 | 1.14320967 |
| **KLC3** | 1.2625 | 0.806368076835953 | 0.661 | 0.192 | 2.06771072 |
| **SLC35A3** | 2.4464 | 0.821270259784376 | 0.665 | 0.209 | 4.00654005 |
| **NFATC4** | 7.2330 | -1.11579864650673 | 0.269 | 0.715 | 1.18455432 |
| **IGLON5** | 1.4993 | -0.974537455858881 | 0.379 | 0.794 | 2.45551185 |
| **ADGRA2** | 2.1394 | 0.835033773292744 | 0.542 | 0.096 | 3.50383960 |
| **PREX1** | 2.7562 | -1.44923365478997 | 0.546 | 0.858 | 4.51385268 |
| **CEP68** | 2.8166 | -0.83487986427255 | 0.752 | 0.957 | 4.61276474 |
| **GRAMD1B** | 4.4420 | -0.848207351029666 | 0.056 | 0.493 | 7.27479234 |
| **AXIN2** | 5.9926 | -0.956504424093492 | 0.842 | 0.966 | 9.81420140 |
| **USP13** | 7.5777 | 0.821493050526258 | 0.98 | 0.932 | 1.24100599 |
| **TRIM61** | 1.2147 | 0.884396576734096 | 0.663 | 0.215 | 1.98937197 |
| **KCTD20** | 1.0869 | -0.816134208086934 | 0.864 | 0.984 | 1.78015504 |
| **SH2D3A** | 1.0937 | 0.890914918684108 | 0.837 | 0.42 | 1.79128910 |
| **RASGRP2** | 2.5765 | 1.12503887269173 | 0.776 | 0.37 | 4.21955634 |
| **AC007906.2** | 2.9061 | -0.998611814071406 | 0.132 | 0.584 | 4.75934689 |
| **ZNF703** | 4.7499 | -1.35508941956009 | 0.619 | 0.894 | 7.77904982 |
| **STC1** | 5.5727 | -1.26058880375927 | 0.073 | 0.518 | 9.12650554 |
| **SERPINF1** | 7.6084 | 1.01020510974881 | 0.825 | 0.464 | 1.24603908 |
| **SRSF12** | 2.3884 | -0.923062818124033 | 0.283 | 0.73 | 3.91152789 |
| **HCLS1** | 3.2274 | 0.94049206666411 | 0.532 | 0.093 | 5.28560533 |
| **EFS** | 3.3190 | -1.03290044664014 | 0.608 | 0.891 | 5.43557002 |
| **FOXD3** | 5.7165 | 1.16499469213327 | 0.709 | 0.271 | 9.36193000 |
| **TFPI2** | 7.2021 | 1.06971239853566 | 0.868 | 0.515 | 1.17949247 |
| **THAP12** | 2.1712 | 0.835243110681815 | 0.855 | 0.497 | 3.55588164 |
| **SLF2** | 4.2854 | -0.830956958609901 | 0.827 | 0.967 | 7.01833015 |
| **LOXL1** | 4.5435 | -1.15027500101758 | 0.516 | 0.842 | 7.44096645 |
| **C2CD5** | 5.4108 | 0.889118045725168 | 0.974 | 0.93 | 8.86140371 |
| **FAM71E1** | 7.3322 | 0.830100418809221 | 0.588 | 0.143 | 1.20079881 |
| **FUNDC1** | 1.6255 | 0.819482320798446 | 0.948 | 0.772 | 2.66222218 |
| **CKAP2** | 2.4371 | -0.957129939385609 | 0.967 | 0.998 | 3.99139085 |
| **RARA** | 3.3532 | -0.895773139728474 | 0.295 | 0.736 | 5.49159857 |

| | | | | | |
|---|---|---|---|---|---|
| **SPSB2** | 3.8954 | 0.878292777445521 | 0.919 | 0.649 | 6.3796089 |
| **SHISAL2B** | 6.5486 | -0.989537932592013 | 0.166 | 0.614 | 1.0724726 |
| **MEGF8** | 6.9910 | -0.873103498181642 | 0.561 | 0.878 | 1.1449165 |
| **ITSN1** | 9.3046 | 0.804239080310015 | 0.943 | 0.75 | 1.5238291 |
| **IL13RA1** | 9.5701 | -1.30250452981865 | 0.107 | 0.548 | 1.5673017 |
| **ADCY2** | 1.3045 | -1.16662606611741 | 0.029 | 0.441 | 2.1364776 |
| **BCAS4** | 2.3518 | -0.858008832204795 | 0.758 | 0.945 | 3.8516504 |
| **SH2D4A** | 3.2071 | 0.943785308024714 | 0.868 | 0.501 | 5.2524228 |
| **GALC** | 3.5258 | -0.921206303280394 | 0.543 | 0.869 | 5.7742378 |
| **MMP25** | 6.7180 | -1.12124234569465 | 0.357 | 0.744 | 1.1002135 |
| **TSPAN9** | 7.4263 | -0.81738458496445 | 0.873 | 0.985 | 1.2162145 |
| **GNG4** | 2.3934 | 1.04041930734735 | 0.664 | 0.222 | 3.9196805 |
| **TP53I3** | 3.6769 | -0.858038962275608 | 0.658 | 0.918 | 6.0217709 |
| **CCNQ** | 4.5696 | 0.8042455526488 | 0.965 | 0.809 | 7.4836917 |
| **SUFU** | 5.3971 | -0.852181060064458 | 0.225 | 0.675 | 8.8389661 |
| **SLC25A44** | 6.1770 | 0.801113367462904 | 0.991 | 0.937 | 1.0116077 |
| **TBXT** | 9.0193 | -1.67899458562697 | 0.444 | 0.844 | 1.4771039 |
| **TWSG1** | 1.2841 | -0.822958898797504 | 0.922 | 0.987 | 2.1031004 |
| **NRARP** | 1.4390 | -1.10063308245686 | 0.449 | 0.8 | 2.3567719 |
| **MYO10** | 1.7779 | 0.920920476342145 | 0.972 | 0.938 | 2.9117806 |
| **PDE9A** | 4.5967 | -1.01563868526596 | 0.261 | 0.689 | 7.5280517 |
| **TGFBR3** | 5.4719 | 0.949433650158511 | 0.735 | 0.305 | 8.9613997 |
| **CDKN2AIP** | 1.3364 | -0.86336035053396 | 0.812 | 0.956 | 2.1886592 |
| **ATP5MPL** | 1.4776 | 0.825240350521406 | 0.888 | 0.602 | 2.4198668 |
| **C1GALT1C1** | 1.6254 | -0.871709530605776 | 0.659 | 0.912 | 2.6619995 |
| **PPA2** | 2.3008 | -0.80964886207407 | 0.647 | 0.913 | 3.7681411 |
| **ST3GAL3** | 4.4573 | -0.856829180126987 | 0.132 | 0.578 | 7.2998644 |
| **MT-CYB** | 9.7016 | -1.02317583574449 | 0.896 | 0.998 | 1.5888432 |
| **RFX3** | 1.0056 | -0.916708809292517 | 0.312 | 0.741 | 1.6469910 |
| **TRIM14** | 1.6107 | 0.890188922373625 | 0.87 | 0.562 | 2.6378792 |
| **PMAIP1** | 6.4831 | 0.819188021575068 | 0.997 | 0.988 | 1.0617438 |
| **IER5** | 1.1253 | -0.983200919064053 | 0.536 | 0.875 | 1.8430659 |
| **LMO1** | 1.8871 | -1.07759834513055 | 0.091 | 0.518 | 3.0905665 |
| **FAM20C** | 2.7236 | -0.972717491681471 | 0.632 | 0.9 | 4.4605913 |

| | | | | | |
|---|---|---|---|---|---|
| **MGAT5B** | 3.8763 | -0.900547879644659 | 0.195 | 0.635 | 6.3482287 |
| **PTPN21** | 5.4471 | -0.901432982780334 | 0.289 | 0.716 | 8.9207728 |
| **MMP24OS** | 6.3938 | -0.8684060302343 | 0.642 | 0.912 | 1.0471153 |
| **CHST11** | 1.1990 | -0.977585427657874 | 0.816 | 0.971 | 1.9636893 |
| **3,00 F** | 1.2476 | 1.18736428909155 | 0.809 | 0.446 | 2.0433324 |
| **ZNF577** | 1.3355 | 0.994742756673959 | 0.737 | 0.327 | 2.1872963 |
| **ARHGAP44** | 2.1620 | 0.860606512178964 | 0.854 | 0.518 | 3.5408471 |
| **MT-ND2** | 6.4832 | -1.08544233012992 | 0.802 | 0.989 | 1.0617664 |
| **TIMP2** | 4.0916 | -0.936775457785815 | 0.902 | 0.973 | 6.7009369 |
| **RIMKLB** | 7.7342 | 0.879532471352436 | 0.999 | 0.991 | 1.2666324 |
| **IPMK** | 9.0633 | 0.812616392497186 | 0.95 | 0.787 | 1.4843085 |
| **ELAC1** | 1.2330 | -0.856263458619176 | 0.213 | 0.649 | 2.0193880 |
| **TRIM22** | 3.5486 | -1.27998757916962 | 0.146 | 0.575 | 5.8116466 |
| **MEX3B** | 6.7838 | -0.919779728264333 | 0.109 | 0.546 | 1.1109905 |
| **CALHM2** | 1.5888 | -0.871830877423947 | 0.12 | 0.556 | 2.6019944 |
| **NKAIN1** | 1.6319 | -1.08314893714418 | 0.315 | 0.72 | 2.6727053 |
| **CDR2** | 5.5001 | 0.833584305323908 | 0.926 | 0.705 | 9.0075452 |
| **TAF1D** | 1.8344 | 0.820410410399956 | 0.997 | 0.975 | 3.0042999 |
| **TC2N** | 4.3640 | 0.861746881395692 | 0.928 | 0.679 | 7.1469659 |
| **TMEM216** | 7.5592 | -0.825607938609602 | 0.263 | 0.7 | 1.2379825 |
| **TNFAIP3** | 8.5314 | 0.874750893702655 | 0.543 | 0.116 | 1.3971917 |
| **APLNR** | 2.1688 | -3.25109904459353 | 0.312 | 0.701 | 3.5519719 |
| **FADS3** | 3.0146 | 0.835603112849877 | 0.694 | 0.256 | 4.9370347 |
| **FGF13** | 1.5562 | -0.942785227431393 | 0.946 | 0.991 | 2.5487258 |
| **WNT3A** | 2.8048 | -1.79685338120909 | 0.097 | 0.508 | 4.5934397 |
| **PCED1A** | 2.8196 | -1.00588671167633 | 0.326 | 0.733 | 4.6178215 |
| **CD83** | 3.3341 | 0.862841860454294 | 0.609 | 0.18 | 5.4604003 |
| **1,00 DKK** | 4.5808 | -2.77275422881181 | 0.025 | 0.418 | 7.5019822 |
| **CERS4** | 1.2586 | -0.906417673464331 | 0.987 | 0.999 | 2.0613563 |
| **TMEM92** | 3.1816 | -0.909756381465157 | 0.317 | 0.723 | 5.2105134 |
| **BAHCC1** | 4.2694 | -0.847883767975514 | 0.055 | 0.467 | 6.9921343 |
| **GPX3** | 5.2109 | -0.843590499702409 | 0.674 | 0.915 | 8.5339304 |
| **PCNA** | 5.3368 | -0.804775175989223 | 0.997 | 0.998 | 8.7400977 |
| **2,00 NPR** | 6.0046 | -0.813346552477058 | 0.139 | 0.577 | 9.8338213 |

| | | | | | |
|---|---|---|---|---|---|
| **WDR44** | 7.0527 | 0.867482888849957 | 0.96 | 0.891 | 1.1550233 |
| **NHSL2** | 8.0420 | -0.831903264188739 | 0.122 | 0.556 | 1.3170402 |
| **GAS7** | 1.0391 | 0.832183742178844 | 0.998 | 0.952 | 1.7017949 |
| **CAPZA1** | 2.8688 | 0.809606197396155 | 0.756 | 0.36 | 4.6983489 |
| **CYP26A1** | 1.5758 | -2.69450506476771 | 0.151 | 0.576 | 2.5807965 |
| **SOX3** | 1.7926 | -1.48245151782428 | 0.099 | 0.52 | 2.9357835 |
| **TCEA2** | 2.9650 | -0.84445628634824 | 0.692 | 0.92 | 4.8558226 |
| **SPRY4** | 3.5344 | -1.05938257856568 | 0.885 | 0.976 | 5.7884486 |
| **RAB17** | 4.3142 | -1.26212581265874 | 0.095 | 0.526 | 7.0654073 |
| **FAM171A1** | 5.7651 | -0.827749804919153 | 0.58 | 0.875 | 9.4415885 |
| **SIX5** | 1.4605 | -0.834410515858541 | 0.598 | 0.89 | 2.3918739 |
| **RTN4RL1** | 1.9286 | -0.803286149258512 | 0.048 | 0.452 | 3.1585501 |
| **CCNA1** | 2.6801 | 1.23385922477462 | 0.433 | 0.046 | 4.3892270 |
| **ROR1** | 3.0499 | -1.07576007473466 | 0.443 | 0.802 | 4.9949098 |
| **IGFBP7** | 5.1195 | -1.50930862008483 | 0.252 | 0.67 | 8.3843502 |
| **ATP12A** | 6.3324 | 1.03384761793842 | 0.823 | 0.415 | 1.0370651 |
| **DMD** | 7.8444 | -1.0994271752085 | 0.59 | 0.865 | 1.2846884 |
| **AMOT** | 1.6146 | -0.911781635292402 | 0.136 | 0.576 | 2.6442798 |
| **FOXB1** | 3.2625 | -1.4192871315095 | 0.424 | 0.765 | 5.3430295 |
| **NR0B1** | 3.9550 | -1.65196739627111 | 0.024 | 0.409 | 6.4771995 |
| **MFSD13A** | 1.7085 | 0.885117351533934 | 0.716 | 0.293 | 2.7980766 |
| **TLE4** | 1.9776 | -0.940701160554046 | 0.489 | 0.833 | 3.2387424 |
| **C12orf75** | 2.2975 | 0.827513243076925 | 0.988 | 0.937 | 3.7626628 |
| **ELF4** | 5.2698 | -1.03174885005268 | 0.187 | 0.615 | 8.6303616 |
| **IER3** | 6.0534 | -1.4483962917263 | 0.351 | 0.742 | 9.9138003 |
| **FGF17** | 6.1594 | -3.18004368553772 | 0.23 | 0.621 | 1.0087285 |
| **P3H4** | 2.3778 | -0.858770767651874 | 0.485 | 0.824 | 3.8942180 |
| **NECAB3** | 2.7903 | -0.832384169826605 | 0.628 | 0.898 | 4.5697203 |
| **SOGA1** | 4.2027 | -0.81897313674129 | 0.884 | 0.982 | 6.8828009 |
| **ENOX2** | 6.1927 | -0.835134520821431 | 0.502 | 0.829 | 1.0141873 |
| **KIF1A** | 9.0816 | 1.10398039555893 | 0.543 | 0.129 | 1.4872984 |
| **BGN** | 1.0885 | -1.71461562081535 | 0.344 | 0.718 | 1.7827320 |
| **MAF** | 1.2290 | -1.11051868549087 | 0.137 | 0.563 | 2.0127877 |
| **SH3GL3** | 2.3494 | -0.974745302587548 | 0.286 | 0.691 | 3.8477495 |

| | | | | | |
|---|---|---|---|---|---|
| **NME7** | 2.8760 | -0.842943306435736 | 0.294 | 0.704 | 4.7101191 |
| **MAPRE2** | 2.8889 | -0.844999762641314 | 0.724 | 0.935 | 4.7312036 |
| **H2AFY** | 2.9891 | -0.873634227226796 | 0.567 | 0.866 | 4.8952513 |
| **IGSF8** | 4.2094 | -0.892895269709408 | 0.775 | 0.936 | 6.8937645 |
| **HSD17B11** | 4.9245 | 0.814724004398554 | 0.891 | 0.614 | 8.0648770 |
| **PKP3** | 1.4539 | 0.852020067000645 | 0.946 | 0.647 | 2.3810626 |
| **ARHGAP6** | 2.6998 | -0.807871711313548 | 0.105 | 0.526 | 4.4216250 |
| **BSPRY** | 3.4334 | 0.815613980767406 | 0.687 | 0.238 | 5.6229370 |
| **ZDBF2** | 6.3375 | 0.816862980427418 | 0.85 | 0.523 | 1.0378975 |
| **ACP2** | 1.6396 | -0.80144331943209 | 0.681 | 0.915 | 2.6852672 |
| **GAB1** | 3.8559 | 0.866060067047951 | 0.793 | 0.441 | 6.3148784 |
| **COMT** | 2.4517 | -0.818515457275198 | 0.623 | 0.899 | 4.0151604 |
| **MSX1** | 3.9162 | -1.52781490398692 | 0.01 | 0.378 | 6.4136717 |
| **MCAM** | 8.8267 | -1.01043970086083 | 0.815 | 0.954 | 1.4455532 |
| **CENPA** | 1.4988 | 0.937342274903082 | 0.983 | 0.874 | 2.4547042 |
| **EFNA5** | 2.6552 | -0.902174026200825 | 0.543 | 0.875 | 4.3485005 |
| **DUSP10** | 5.6600 | -0.974703002399215 | 0.311 | 0.721 | 9.2693848 |
| **CAAP1** | 9.4287 | 0.82689228328419 | 0.898 | 0.635 | 1.5441432 |
| **PDZD2** | 1.0166 | 0.834603777740529 | 0.69 | 0.262 | 1.6649771 |
| **LARP1B** | 1.3423 | 0.807533135005697 | 0.902 | 0.687 | 2.1983091 |
| **CD82** | 2.5904 | -1.0160016246624 | 0.344 | 0.712 | 4.2423497 |
| **USP18** | 3.9941 | 0.969765458524789 | 0.776 | 0.414 | 6.5412909 |
| **ZKSCAN1** | 1.0435 | -0.862580125099643 | 0.467 | 0.811 | 1.7090163 |
| **CRABP1** | 3.3982 | -1.62286520201282 | 0.63 | 0.879 | 5.5652859 |
| **REEP2** | 6.8782 | -0.936420253556368 | 0.291 | 0.687 | 1.1264497 |
| **INSYN1** | 2.1376 | -1.06935326002828 | 0.125 | 0.528 | 3.5008649 |
| **CSF2RA** | 1.5573 | 1.2419417997157 | 0.548 | 0.144 | 2.5504653 |
| **ARMCX1** | 1.8081 | -0.82782062359632 | 0.717 | 0.937 | 2.9611981 |
| **CALB1** | 2.7336 | 0.833528626523265 | 0.556 | 0.132 | 4.4769472 |
| **HES4** | 3.6743 | 0.907453811920529 | 0.706 | 0.282 | 6.0174539 |
| **RRAD** | 4.1883 | 1.16945815553259 | 0.386 | 0.031 | 6.8593097 |
| **CDX2** | 3.5548 | -1.60455163946187 | 0.067 | 0.465 | 5.8218374 |
| **CUEDC1** | 1.3795 | -0.865706522309242 | 0.244 | 0.653 | 2.2592963 |
| **UST** | 1.9472 | -0.879002480499201 | 0.815 | 0.957 | 3.1890828 |

| | | | | | |
|---|---|---|---|---|---|
| **RGS3** | 2.6654 | -0.842696544262742 | 0.083 | 0.485 | 4.3651635 |
| **PTPN13** | 9.0444 | -0.920434392006899 | 0.463 | 0.797 | 1.4812019 |
| **SPIC** | 1.3378 | 0.913995454747475 | 0.367 | 0.022 | 2.1909898 |
| **EPHX1** | 2.9504 | -1.18239887593121 | 0.968 | 0.991 | 4.8319111 |
| **COL5A1** | 3.7879 | -1.28852975055008 | 0.533 | 0.845 | 6.2034851 |
| **COTL1** | 5.8456 | -0.842826316090863 | 0.985 | 0.997 | 9.5734130 |
| **MKI67** | 7.7246 | -1.14787700946713 | 0.901 | 0.981 | 1.2650690 |
| **CACNA1B** | 8.4955 | 0.826239745535967 | 0.641 | 0.224 | 1.3913106 |
| **ZNF354A** | 1.5122 | 0.800733206379674 | 0.8 | 0.429 | 2.4765308 |
| **ADM** | 6.9180 | -1.4492473822129 | 0.27 | 0.652 | 1.1329758 |
| **TAOK3** | 8.7953 | 0.819109287671016 | 0.674 | 0.267 | 1.4404164 |
| **PITX2** | 1.2643 | -0.950310712374554 | 0.397 | 0.784 | 2.0706258 |
| **VASN** | 6.1679 | -0.821407003369099 | 0.177 | 0.586 | 1.0101314 |
| **BNC2** | 6.9343 | 0.86677901002871 | 0.788 | 0.39 | 1.1356451 |
| **TMEM50B** | 1.9445 | 0.821350586408756 | 0.933 | 0.731 | 3.1846084 |
| **TM7SF2** | 5.9748 | -0.935705198275965 | 0.898 | 0.952 | 9.7850262 |
| **CADM4** | 7.7137 | -0.833886599370106 | 0.176 | 0.58 | 1.2632881 |
| **TMEM88** | 1.6647 | -0.868863206244208 | 0.118 | 0.516 | 2.7263469 |
| **PCNX4** | 4.3637 | -0.82465111256654 | 0.537 | 0.841 | 7.1464976 |
| **ZSWIM9** | 5.2733 | -0.810106882672101 | 0.319 | 0.71 | 8.6362165 |
| **SLIT2** | 6.8836 | -1.03107202737275 | 0.011 | 0.36 | 1.1273280 |
| **ME1** | 7.0694 | -0.844841236251709 | 0.455 | 0.8 | 1.1577711 |
| **RRAS** | 1.0177 | -0.850182747336502 | 0.214 | 0.619 | 1.6668095 |
| **CACFD1** | 1.5174 | -0.812322895219223 | 0.529 | 0.827 | 2.4851103 |
| **IFI27L2** | 3.0197 | -0.86559082137592 | 0.24 | 0.644 | 4.9454889 |
| **TSPAN7** | 5.3903 | -1.14741803215924 | 0.104 | 0.511 | 8.8278159 |
| **SOCS3** | 6.6330 | -0.854140509934438 | 0.751 | 0.954 | 1.0862873 |
| **ZNF613** | 8.2194 | 0.841880476575321 | 0.795 | 0.458 | 1.3461007 |
| **ZFHX3** | 1.2149 | 0.982124468865655 | 0.577 | 0.165 | 1.9897932 |
| **SLC43A2** | 1.4809 | -0.814696169743427 | 0.803 | 0.941 | 2.4254287 |
| **GLS2** | 1.3691 | 0.809337789261322 | 0.623 | 0.212 | 2.2422254 |
| **FBXO25** | 1.4528 | 0.802382326066167 | 0.818 | 0.497 | 2.3793299 |
| **NEO1** | 4.6461 | -0.819900313995538 | 0.599 | 0.861 | 7.6089336 |
| **EDN1** | 3.6695 | -1.57898436842567 | 0.065 | 0.44 | 6.0096356 |

| | | | | | |
|---|---|---|---|---|---|
| **GJA5** | 1.0359 | -0.854658454026156 | 0.031 | 0.393 | 1.6965886 |
| **YPEL2** | 1.7828 | 1.09212373497721 | 0.648 | 0.249 | 2.9197968 |
| **ASF1B** | 1.8923 | -0.813148849922547 | 0.926 | 0.973 | 3.0991305 |
| **SMPDL3A** | 4.3569 | 0.818136983930891 | 0.571 | 0.174 | 7.1353746 |
| **RASGEF1A** | 4.4247 | 0.808775740366089 | 0.898 | 0.655 | 7.2464058 |
| **TRH** | 6.2627 | 0.824349645271505 | 0.522 | 0.123 | 1.0256542 |
| **DDB2** | 3.0904 | -0.862457145146637 | 0.728 | 0.914 | 5.0612845 |
| **RHOBTB3** | 6.6895 | -1.49449802765588 | 0.087 | 0.476 | 1.0955544 |
| **NR5A2** | 8.6147 | 0.885424680974755 | 0.542 | 0.152 | 1.4108441 |
| **COL13A1** | 1.6285 | -1.27022206548822 | 0.016 | 0.359 | 2.6670080 |
| **AEBP1** | 3.1327 | -0.836308446274774 | 0.637 | 0.884 | 5.1305827 |
| **PTH1R** | 1.2432 | 0.982124274386219 | 0.516 | 0.136 | 2.0360622 |
| **SPIN3** | 1.4172 | -1.03124068075832 | 0.43 | 0.771 | 2.3210848 |
| **CBR3** | 2.2199 | 0.9247253116445 | 0.776 | 0.408 | 3.6356269 |
| **ZFP36L2** | 1.5409 | -0.808388593542589 | 0.961 | 0.994 | 2.5236360 |
| **FGF8** | 4.1331 | -1.43200968803606 | 0.511 | 0.786 | 6.7689216 |
| **NUPR1** | 2.2233 | 0.998787749878729 | 0.603 | 0.208 | 3.6411997 |
| **ARHGEF10L** | 2.4149 | -0.811893652220068 | 0.391 | 0.744 | 3.9550085 |
| **SMPDL3B** | 4.4467 | -0.85414872908509 | 0.188 | 0.581 | 7.2825176 |
| **EVA1B** | 1.0374 | -0.920557121585275 | 0.329 | 0.702 | 1.6989582 |
| **ANKRD1** | 1.3058 | -1.58130686029535 | 0.101 | 0.489 | 2.1386163 |
| **GPC4** | 1.8600 | -0.891033267077944 | 0.971 | 1 | 3.0461987 |
| **NRBP2** | 4.1564 | 0.952253933446444 | 0.621 | 0.235 | 6.8070061 |
| **GRPR** | 3.1204 | -1.01625850406257 | 0.051 | 0.41 | 5.1103392 |
| **TCF7L2** | 1.1531 | -0.829549645436728 | 0.771 | 0.941 | 1.8885471 |
| **GCA** | 1.8387 | -0.851720131642609 | 0.385 | 0.727 | 3.0113861 |
| **AGPAT4** | 2.3405 | -0.866079471700693 | 0.568 | 0.852 | 3.8330647 |
| **CLU** | 2.6169 | -0.818189248428988 | 0.959 | 0.995 | 4.2857611 |
| **LIPG** | 2.6506 | -1.25097263445403 | 0.104 | 0.484 | 4.3409736 |
| **MYC** | 4.1159 | -0.886717454014863 | 0.959 | 0.991 | 6.7406153 |
| **MAGI1** | 3.4362 | -0.937900765652557 | 0.7 | 0.888 | 5.6275840 |
| **NEXN** | 4.6461 | -0.817200693680472 | 0.166 | 0.558 | 7.6089295 |
| **HSPB8** | 6.2462 | -0.899203110422648 | 0.16 | 0.591 | 1.0229535 |
| **ADRA2C** | 3.1836 | -1.02736758495558 | 0.889 | 0.952 | 5.2138932 |

| | | | | | |
|---|---|---|---|---|---|
| **LIX1L** | 4.3839 | -1.07460499126658 | 0.613 | 0.861 | 7.1795908 |
| **DAB1** | 6.8666 | -1.16201342569357 | 0.205 | 0.575 | 1.1245581 |
| **GFPT2** | 2.5451 | -0.833556339327767 | 0.875 | 0.983 | 4.1681165 |
| **RGMB** | 2.5331 | -0.905362670751552 | 0.649 | 0.874 | 4.1486106 |
| **KIFC3** | 4.2645 | -0.800335526934456 | 0.348 | 0.722 | 6.9840082 |
| **CCDC92** | 1.6259 | -0.821761881215457 | 0.061 | 0.416 | 2.6628519 |
| **TUBB2A** | 5.5449 | -1.06153192509468 | 0.989 | 0.999 | 9.0810075 |
| **CYP1B1** | 7.4736 | -1.42926260655452 | 0.214 | 0.585 | 1.2239652 |
| **KCNG1** | 9.2924 | -1.12718405976285 | 0.552 | 0.821 | 1.5218220 |
| **TRIM38** | 1.3980 | 0.917887295750455 | 0.691 | 0.335 | 2.2895896 |
| **ZNF483** | 2.7568 | 0.935539735796451 | 0.833 | 0.601 | 4.5149562 |
| **CREB3L1** | 7.2781 | 0.84781262803177 | 0.829 | 0.547 | 1.1919420 |
| **VWA5A** | 1.7825 | -1.06108761922324 | 0.295 | 0.652 | 2.9192369 |
| **SERPINB9** | 4.9842 | 1.06951865826758 | 0.991 | 0.952 | 8.1626343 |
| **BMP2** | 4.7302 | -2.29888558574549 | 0.147 | 0.508 | 7.7467682 |
| **CAVIN3** | 8.7531 | -0.877050757777889 | 0.332 | 0.695 | 1.4335005 |
| **PRR36** | 1.0927 | -1.02066084432453 | 0.432 | 0.715 | 1.7895451 |
| **TJP3** | 1.1604 | 0.805291830461584 | 0.902 | 0.623 | 1.9004606 |
| **NKX1-2** | 1.4787 | -1.16547736454232 | 0.907 | 0.931 | 2.4217490 |
| **C11orf96** | 7.9730 | -1.30523227050202 | 0.14 | 0.506 | 1.3057488 |
| **CRLF1** | 1.2933 | -2.21078208532114 | 0.509 | 0.735 | 2.1180514 |
| **EGLN3** | 2.8262 | 0.962106184471464 | 0.952 | 0.839 | 4.6286188 |
| **GRHL3** | 2.5663 | -1.13307692447652 | 0.147 | 0.508 | 4.2028674 |
| **COL6A2** | 8.4445 | -1.292664315376 | 0.918 | 0.955 | 1.3829641 |
| **RASL11B** | 4.8176 | -0.988716572365704 | 0.405 | 0.744 | 7.8899210 |
| **DHRSX** | 5.3056 | 0.957282029548812 | 0.93 | 0.856 | 8.6890495 |
| **TRPC6** | 7.6996 | -1.28178511624808 | 0.187 | 0.537 | 1.2609777 |
| **WIPI1** | 1.0953 | 0.846035428369163 | 0.737 | 0.393 | 1.7938150 |
| **BEX1** | 1.2127 | -0.913309997529904 | 0.137 | 0.496 | 1.9861654 |
| **SAMD4A** | 6.8291 | -0.820235636005187 | 0.199 | 0.598 | 1.1184143 |
| **OTUD1** | 6.4327 | 0.811659291876088 | 0.629 | 0.276 | 1.0534868 |
| **TACR1** | 1.9329 | -1.61337362901768 | 0.076 | 0.413 | 3.1655899 |
| **NPAS1** | 1.4291 | 0.940690078057441 | 0.576 | 0.228 | 2.3404596 |
| **AP3B2** | 4.2160 | -1.06623861506706 | 0.205 | 0.547 | 6.9047046 |

| | | | | | |
|---|---|---|---|---|---|
| **TGFB1I1** | 9.4337 | -0.898662348346117 | 0.264 | 0.612 | 1.5449582 |
| **FZD10** | 2.4319 | -1.26811720211147 | 0.701 | 0.876 | 3.9828284 |
| **ACSS3** | 4.2064 | -1.08193588141717 | 0.397 | 0.705 | 6.8888941 |
| **PKP2** | 9.7478 | 0.835375360849895 | 0.724 | 0.379 | 1.5963991 |
| **EOGT** | 2.6423 | -0.877213600151399 | 0.887 | 0.973 | 4.3273874 |
| **ACKR3** | 3.6487 | 0.925441035788918 | 0.637 | 0.276 | 5.9756108 |
| **GMPR** | 5.2572 | -1.4627469611991 | 0.234 | 0.552 | 8.6097856 |
| **SAMD3** | 7.8424 | -0.937620995383554 | 0.008 | 0.289 | 1.2843633 |
| **PLK2** | 7.7540 | 0.970281773113791 | 0.592 | 0.237 | 1.2698816 |
| **CDHR1** | 4.6291 | 0.864502475575576 | 0.695 | 0.356 | 7.5810856 |
| **COL6A1** | 2.4169 | -1.38893851317613 | 0.751 | 0.88 | 3.9582283 |
| **ENC1** | 7.6537 | -1.21733168948095 | 0.619 | 0.88 | 1.2534605 |
| **ZNF850** | 3.2235 | 0.802832357230572 | 0.949 | 0.875 | 5.2792498 |
| **CNTNAP2** | 2.9506 | -2.21424893159731 | 0.848 | 0.869 | 4.8322288 |
| **LAMA2** | 6.3962 | -0.811225817582061 | 0.254 | 0.583 | 1.0475146 |
| **ZNF503** | 3.6159 | -0.846965778105816 | 0.079 | 0.398 | 5.9218587 |
| **EBF2** | 7.3855 | -0.861892769805346 | 0.092 | 0.412 | 1.2095383 |
| **MAPK4** | 1.6853 | -0.803198666526333 | 0.179 | 0.514 | 2.7601693 |
| **DEPDC1** | 4.4535 | 0.831417793481374 | 0.604 | 0.273 | 7.2935184 |
| **EFNB2** | 1.1947 | -0.927766855875384 | 0.745 | 0.903 | 1.9566629 |
| **ROR2** | 2.4962 | -1.08534530391125 | 0.655 | 0.846 | 4.0881668 |
| **FOXC1** | 8.2162 | -1.33817645711415 | 0.106 | 0.419 | 1.3455702 |
| **GSC** | 1.6420 | -1.64901447176792 | 0.031 | 0.305 | 2.6892627 |
| **METTL7A** | 8.9638 | 0.889002493623016 | 0.592 | 0.279 | 1.4680098 |
| **MEIS3** | 2.3393 | -1.4694901653182 | 0.728 | 0.818 | 3.8311417 |
| **THBS1** | 6.7698 | -0.809004959453384 | 0.171 | 0.498 | 1.1086909 |
| **SNAI2** | 1.2021 | -1.30552780717421 | 0.027 | 0.292 | 1.9686859 |
| **PLAU** | 1.5258 | 1.11284841047069 | 0.674 | 0.369 | 2.4988111 |
| **MFAP4** | 1.9361 | -0.942808029052946 | 0.074 | 0.362 | 3.1708334 |
| **FAM20A** | 9.5702 | -1.13271539119536 | 0.19 | 0.493 | 1.5673119 |
| **RAB20** | 4.5429 | -0.915691201629476 | 0.738 | 0.877 | 7.4400551 |
| **HIST1H4J** | 1.9481 | -1.06808366317867 | 0.495 | 0.722 | 3.1904156 |
| **CDX1** | 6.6511 | -1.0947983060714 | 0.563 | 0.787 | 1.0892632 |
| **HAPLN1** | 1.9605 | 1.16641919529964 | 0.509 | 0.177 | 3.2108568 |

| | | | | | |
|---|---|---|---|---|---|
| **EVX1** | 4.6227 | -0.822998574191007 | 0.035 | 0.293 | 7.5705982 |
| **COL5A2** | 6.0539 | -1.42672857410941 | 0.157 | 0.449 | 9.9146045 |
| **ITGB8** | 4.1728 | -0.824761132879177 | 0.1 | 0.397 | 6.8338137 |
| **HMOX1** | 5.3987 | 0.842120635529616 | 0.721 | 0.418 | 8.8414813 |
| **LGALS3** | 3.5268 | 0.983224732211682 | 0.504 | 0.2 | 5.7759709 |
| **CCNG2** | 3.3143 | 0.815023796248974 | 0.813 | 0.583 | 5.4279072 |
| **DDIT4** | 1.0147 | 0.855909732682481 | 0.937 | 0.821 | 1.6618167 |
| **USP44** | 1.3256 | 0.899187186141533 | 0.771 | 0.59 | 2.1709739 |
| **SPATS2L** | 9.8369 | -1.00587504980173 | 0.718 | 0.863 | 1.6109966 |
| **BMP4** | 3.1278 | -1.17203887179097 | 0.681 | 0.86 | 5.1224921 |
| **FOXA2** | 1.2048 | -1.06547128838953 | 0.025 | 0.25 | 1.9732120 |
| **GLI3** | 1.8470 | -0.823667638289901 | 0.26 | 0.552 | 3.0249423 |
| **PODXL** | 2.5319 | -1.38904719054427 | 0.984 | 0.976 | 4.1465050 |
| **LHX1** | 2.1677 | -1.63492515794536 | 0.012 | 0.222 | 3.5500449 |
| **VIT** | 5.4001 | -0.886615786468658 | 0.085 | 0.341 | 8.8438219 |
| **DDAH1** | 1.4025 | -0.975377829771439 | 0.614 | 0.799 | 2.2969585 |
| **ADAMTSL4** | 7.0153 | -1.09097129393681 | 0.436 | 0.646 | 1.1488997 |
| **GDF15** | 4.3041 | 1.11745471858328 | 0.377 | 0.117 | 7.0489668 |
| **COL7A1** | 1.0731 | -0.858557638957545 | 0.178 | 0.452 | 1.7574969 |
| **B3GNT7** | 5.2713 | -0.993815713015945 | 0.712 | 0.808 | 8.6329620 |
| **GATA6** | 3.7533 | -1.99835887385391 | 0.101 | 0.351 | 6.1468672 |
| **COL1A1** | 2.4522 | -0.801226544577496 | 0.899 | 0.957 | 4.0160678 |
| **APCDD1** | 3.0934 | -1.02402970502816 | 0.348 | 0.599 | 5.0661820 |
| **HPD** | 2.8945 | -0.906826881682458 | 0.252 | 0.502 | 4.7404202 |
| **CCN2** | 2.3634 | -0.895078789236588 | 0.903 | 0.942 | 3.8706459 |
| **DHRS3** | 5.5325 | -0.824580994002802 | 0.403 | 0.615 | 9.0606445 |
| **TPM1** | 2.7597 | -0.864681050772818 | 1 | 1 | 4.5197155 |
| **ANOS1** | 2.5231 | -1.52132237337561 | 0.183 | 0.411 | 4.1321615 |
| **TPX2** | 3.1798 | -0.859803069241728 | 0.988 | 0.997 | 5.2076142 |
| **COLEC12** | 2.0774 | -1.41610710475322 | 0.39 | 0.626 | 3.4022193 |
| **CCDC141** | 4.2292 | -0.867326518824748 | 0.141 | 0.375 | 6.9263182 |
| **TBX6** | 9.6276 | -1.83122284738795 | 0.118 | 0.324 | 1.5767197 |
| **MSGN1** | 2.6403 | -1.67860122558001 | 0.047 | 0.235 | 4.3241231 |
| **QPCT** | 4.2032 | -0.821198051186009 | 0.769 | 0.852 | 6.8837270 |

| | | | | | |
|---|---|---|---|---|---|
| **GAL** | 5.9661 | -1.36471165127151 | 0.997 | 0.981 | 9.77077166 |
| **PCDH18** | 1.3000 | -0.940933662333648 | 0.618 | 0.759 | 2.12901885 |
| **MESP1** | 8.1989 | -0.813437001166554 | 0.09 | 0.283 | 1.34274777 |
| **CSRP1** | 1.6127 | -0.828293479135768 | 0.974 | 0.993 | 2.64118896 |
| **PLXNA2** | 2.3101 | -1.5251922473564 | 0.202 | 0.404 | 3.78326802 |
| **FOXJ1** | 1.1394 | -0.801532306090302 | 0.173 | 0.376 | 1.86603062 |
| **MT-ND5** | 9.8769 | 0.98963307223672 | 0.811 | 0.75 | 1.61754503 |
| **SNAI1** | 2.4777 | -0.856143119655906 | 0.184 | 0.351 | 4.05783831 |
| **GATA4** | 1.1907 | -0.862002634860924 | 0.168 | 0.327 | 1.95008929 |
| **SPP1** | 1.9392 | -0.951473552498011 | 0.718 | 0.847 | 3.17597374 |
| **GYPC** | 4.0174 | -0.981719494209878 | 0.826 | 0.829 | 6.57937711 |
| **DLL1** | 1.4092 | -0.893641889478853 | 0.089 | 0.205 | 2.30798939 |
| **CLDN10** | 1.0157 | -1.3071118373104 | 0.337 | 0.462 | 1.66349484 |
| **ACTA1** | 2.0581 | -1.09261298634225 | 0.412 | 0.534 | 3.37061907 |
| **ACTG2** | 5.6466 | -1.32306481854658 | 0.776 | 0.82 | 9.24746776 |
| **NPPB** | 2.4897 | -1.02384880112831 | 0.156 | 0.274 | 4.0775219 |
| **HIST1H1E** | 3.2497 | -0.909456276469148 | 0.918 | 0.931 | 5.3221253 |
| **DLL3** | 2.3169 | -1.20530181261878 | 0.74 | 0.578 | 3.79440221 |
| **ARL4D** | 0.0004 | -0.92452152349672 | 0.556 | 0.56 | 1 |
| **LZTS1** | 0.0089 | -0.934767429002002 | 0.744 | 0.614 | 1 |

Supplemental Table S2_Sheet_3

| | p_val | avg_log2FC | pct.1 | pct.2 | p_val_adj |
|---|---|---|---|---|---|
| **AGRN** | 0 | -1.81221316738546 | 0.899 | 0.998 | 0 |
| **MXRA8** | 0 | -1.84552174824832 | 0.684 | 0.986 | 0 |
| **ATAD3B** | 0 | 1.76274376462673 | 1 | 0.956 | 0 |
| **FNDC10** | 0 | 1.16356319127576 | 0.948 | 0.588 | 0 |
| **SKI** | 0 | -0.953709301985817 | 0.954 | 0.999 | 0 |
| **ENO1** | 0 | -0.833615630062416 | 1 | 1 | 0 |
| **CLSTN1** | 0 | -0.965817366348623 | 0.999 | 0.999 | 0 |
| **CTNNBIP1** | 0 | -1.20014407598067 | 0.906 | 0.992 | 0 |
| **KIF1B** | 0 | -0.836406634525696 | 0.986 | 1 | 0 |
| **DFFA** | 0 | 1.09439334548332 | 1 | 0.996 | 0 |
| **SRM** | 0 | 0.872229854007516 | 1 | 1 | 0 |
| **MTOR** | 0 | 0.908726463093068 | 0.998 | 0.954 | 0 |
| **DRAXIN** | 0 | -2.50580870499806 | 0.385 | 0.965 | 0 |
| **TNFRSF8** | 0 | 2.17834349923834 | 0.939 | 0.067 | 0 |
| **DHRS3** | 0 | -1.48271606908824 | 0.316 | 0.812 | 0 |
| **EFHD2** | 0 | 1.1246517293166 | 0.997 | 0.96 | 0 |
| **EPHA2** | 0 | -2.05288798154993 | 0.047 | 0.752 | 0 |
| **MFAP2** | 0 | -1.91321428535409 | 0.822 | 0.992 | 0 |
| **KLHDC7A** | 0 | -1.09245854647721 | 0.008 | 0.525 | 0 |
| **CDA** | 0 | 1.45291921178337 | 0.998 | 0.794 | 0 |
| **PINK1** | 0 | 1.41123401241858 | 1 | 0.957 | 0 |
| **ECE1** | 0 | -2.41819513776515 | 0.57 | 0.994 | 0 |
| **LUZP1** | 0 | -1.2698583765915 | 0.827 | 0.995 | 0 |
| **ID3** | 0 | -3.41242358742776 | 0.062 | 0.806 | 0 |
| **FUCA1** | 0 | 1.88577122378569 | 0.999 | 0.814 | 0 |
| **CNR2** | 0 | 2.05469069349769 | 0.91 | 0.137 | 0 |
| **GRHL3** | 0 | -1.72289695182294 | 0.053 | 0.6 | 0 |
| **MAN1C1** | 0 | 1.22658274225725 | 0.998 | 0.886 | 0 |
| **STMN1** | 0 | 1.34251409145807 | 1 | 0.983 | 0 |
| **LIN28A** | 0 | -2.11611482062716 | 1 | 1 | 0 |
| **ARID1A** | 0 | -1.05676291343141 | 0.984 | 1 | 0 |
| **SLC9A1** | 0 | 1.00961622847793 | 0.978 | 0.811 | 0 |

| | | | | | |
|---|---|---|---|---|---|
| **RPA2** | 0 | 1.09992550335105 | 0.937 | 0.58 | 0 |
| **SMPDL3B** | 0 | -2.26229881623691 | 0.027 | 0.819 | 0 |
| **PTPRU** | 0 | 1.38943266818307 | 1 | 0.937 | 0 |
| **NKAIN1** | 0 | -1.50299215778928 | 0.073 | 0.604 | 0 |
| **EIF3I** | 0 | 1.32396888281912 | 0.926 | 0.445 | 0 |
| **MARCKSL1** | 0 | -1.69156680103493 | 1 | 1 | 0 |
| **KIAA1522** | 0 | -1.38986413230297 | 0.967 | 0.999 | 0 |
| **TMEM54** | 0 | 1.20905480128685 | 0.807 | 0.25 | 0 |
| **ZNF362** | 0 | -1.32061703014147 | 0.781 | 0.986 | 0 |
| **PHC2** | 0 | -1.77779266014329 | 0.043 | 0.589 | 0 |
| **MYCBP** | 0 | 1.37136566152226 | 0.938 | 0.444 | 0 |
| **AKIRIN1** | 0 | 0.969660124673673 | 1 | 0.997 | 0 |
| **MACF1** | 0 | -0.993037270480599 | 0.945 | 0.994 | 0 |
| **PPT1** | 0 | -1.01751280814272 | 0.961 | 0.996 | 0 |
| **YBX1** | 0 | -0.882549345702277 | 1 | 1 | 0 |
| **CLDN19** | 0 | -1.19898082255798 | 0.068 | 0.608 | 0 |
| **HPDL** | 0 | 1.85449102573748 | 0.887 | 0.154 | 0 |
| **PRDX1** | 0 | 2.633480886186 | 1 | 0.991 | 0 |
| **PIK3R3** | 0 | -1.54202932016499 | 0.6 | 0.914 | 0 |
| **CMPK1** | 0 | 0.848454850401746 | 0.996 | 0.932 | 0 |
| **OSBPL9** | 0 | -1.07213617230601 | 0.969 | 0.998 | 0 |
| **NRDC** | 0 | 0.973899299172662 | 1 | 1 | 0 |
| **ZFYVE9** | 0 | 0.821551072616018 | 1 | 0.982 | 0 |
| **ZYG11A** | 0 | 1.40081195229715 | 0.999 | 0.844 | 0 |
| **DHCR24** | 0 | -1.34894028864487 | 1 | 1 | 0 |
| **PLPP3** | 0 | -1.46209410953333 | 0.542 | 0.939 | 0 |
| **PGM1** | 0 | -1.36865006743219 | 0.529 | 0.953 | 0 |
| **ROR1** | 0 | -2.53888680445273 | 0.061 | 0.832 | 0 |
| **AK4** | 0 | 2.21770747541914 | 1 | 0.884 | 0 |
| **WLS** | 0 | -4.5311429461346 | 0.542 | 0.988 | 0 |
| **CTH** | 0 | 1.71763320673194 | 0.979 | 0.714 | 0 |
| **ST6GALNAC** | 0 | -1.42212005268058 | 0.054 | 0.632 | 0 |
| **FUBP1** | 0 | -0.905276653683837 | 0.998 | 1 | 0 |
| **LMO4** | 0 | 1.15690425013538 | 0.999 | 0.939 | 0 |

| | | | | | |
|---|---|---|---|---|---|
| **ZNF644** | 0 | -0.992469171624085 | 0.991 | 1 | 0 |
| **TGFBR3** | 0 | 1.85321672211065 | 0.911 | 0.176 | 0 |
| **BRDT** | 0 | 3.10290685785331 | 0.997 | 0.255 | 0 |
| **TMED5** | 0 | 0.907413228284289 | 1 | 0.952 | 0 |
| **FNBP1L** | 0 | -1.21915895592114 | 0.999 | 1 | 0 |
| **PTBP2** | 0 | -1.18692726810009 | 0.753 | 0.957 | 0 |
| **AGL** | 0 | 2.26503426274662 | 1 | 0.958 | 0 |
| **SLC35A3** | 0 | 1.13414272497573 | 0.786 | 0.215 | 0 |
| **SLC25A24** | 0 | -1.53490122565021 | 0.759 | 0.993 | 0 |
| **FAM102B** | 0 | 1.40176077153742 | 0.992 | 0.837 | 0 |
| **HENMT1** | 0 | 1.97883401683664 | 0.92 | 0.114 | 0 |
| **TAF13** | 0 | 3.15703338450404 | 1 | 0.839 | 0 |
| **GNAI3** | 0 | -0.897436041711703 | 1 | 1 | 0 |
| **SLC16A1** | 0 | -1.02380989994492 | 1 | 1 | 0 |
| **OLFML3** | 0 | -4.45811874889165 | 0.579 | 0.996 | 0 |
| **SYT6** | 0 | -1.41686078897818 | 0.145 | 0.736 | 0 |
| **NOTCH2** | 0 | -1.42569697334685 | 0.431 | 0.882 | 0 |
| **SV2A** | 0 | -1.62508290124745 | 0.09 | 0.776 | 0 |
| **PLEKHO1** | 0 | -2.16286947346321 | 0.227 | 0.902 | 0 |
| **HORMAD1** | 0 | 2.69252073755561 | 0.99 | 0.202 | 0 |
| **MLLT11** | 0 | -1.86557210224923 | 0.381 | 0.907 | 0 |
| **POGZ** | 0 | -1.00581474420582 | 0.858 | 0.988 | 0 |
| **CGN** | 0 | -1.24450449955946 | 0.543 | 0.917 | 0 |
| **S100A10** | 0 | 1.16590366747798 | 1 | 0.992 | 0 |
| **RAB13** | 0 | -1.34522821971513 | 0.935 | 0.996 | 0 |
| **UBAP2L** | 0 | -0.893552590825394 | 0.997 | 1 | 0 |
| **CKS1B** | 0 | 2.08751629941429 | 1 | 0.932 | 0 |
| **FLAD1** | 0 | 1.08979127473733 | 1 | 0.966 | 0 |
| **THBS3** | 0 | -1.43824088389996 | 0.066 | 0.643 | 0 |
| **FDPS** | 0 | -0.995535897536243 | 0.975 | 0.995 | 0 |
| **MEX3A** | 0 | -2.36351738827375 | 0.173 | 0.898 | 0 |
| **SLC25A44** | 0 | 0.912204518850749 | 1 | 0.975 | 0 |
| **PMF1** | 0 | -0.895898579440954 | 0.949 | 0.996 | 0 |
| **NAXE** | 0 | -1.51018281935152 | 0.642 | 0.985 | 0 |

| | | | | | |
|---|---|---|---|---|---|
| **NES** | 0 | -2.48742950444067 | 0.982 | 1 | 0 |
| **CRABP2** | 0 | -3.6446006297001 | 0.771 | 0.994 | 0 |
| **HDGF** | 0 | -1.05539121864072 | 0.946 | 0.997 | 0 |
| **DUSP23** | 0 | 1.88721430811961 | 0.91 | 0.098 | 0 |
| **IGSF8** | 0 | -1.42990538293583 | 0.658 | 0.94 | 0 |
| **PEA15** | 0 | -1.87932700047058 | 0.786 | 0.994 | 0 |
| **VANGL2** | 0 | -1.65949028674714 | 0.611 | 0.973 | 0 |
| **F11R** | 0 | 1.23960231049464 | 1 | 0.988 | 0 |
| **TSTD1** | 0 | 2.1679390982784 | 0.908 | 0.04 | 0 |
| **RGS5** | 0 | -3.01658592172805 | 0.117 | 0.851 | 0 |
| **MGST3** | 0 | -1.56844035488778 | 0.886 | 0.997 | 0 |
| **MAEL** | 0 | 1.42401231778172 | 0.617 | 0.022 | 0 |
| **ANKRD45** | 0 | 1.48760118253342 | 0.683 | 0.086 | 0 |
| **GLUL** | 0 | -1.1655786343546 | 0.986 | 1 | 0 |
| **RO60** | 0 | -1.27567008462484 | 0.866 | 0.997 | 0 |
| **GLRX2** | 0 | -1.03477175070283 | 0.099 | 0.638 | 0 |
| **CDC73** | 0 | -0.940285175772652 | 0.957 | 0.998 | 0 |
| **NR5A2** | 0 | 1.45183968697005 | 0.743 | 0.078 | 0 |
| **TIMM17A** | 0 | 0.930918591173291 | 0.999 | 0.978 | 0 |
| **ELF3** | 0 | 1.59596295507135 | 0.99 | 0.697 | 0 |
| **BTG2** | 0 | -2.01331760832798 | 0.225 | 0.801 | 0 |
| **SNRPE** | 0 | 1.54872536214367 | 1 | 0.865 | 0 |
| **SOX13** | 0 | -1.14921609866742 | 0.602 | 0.924 | 0 |
| **NUAK2** | 0 | -1.50418363528208 | 0.219 | 0.731 | 0 |
| **NUCKS1** | 0 | -1.07812802920878 | 1 | 1 | 0 |
| **MAPKAPK2** | 0 | -0.958752697937146 | 0.951 | 0.998 | 0 |
| **CR1L** | 0 | 2.6369895314013 | 0.998 | 0.415 | 0 |
| **IRF6** | 0 | 1.1253791687476 | 0.684 | 0.048 | 0 |
| **SERTAD4** | 0 | -1.16410628573043 | 0.035 | 0.589 | 0 |
| **TRAF5** | 0 | -1.67562178852753 | 0.063 | 0.765 | 0 |
| **SMYD2** | 0 | -1.26317392951653 | 0.833 | 0.979 | 0 |
| **PTPN14** | 0 | -1.48748119469833 | 0.801 | 0.998 | 0 |
| **DUSP10** | 0 | -1.29291497986195 | 0.257 | 0.772 | 0 |
| **CAPN2** | 0 | -2.2699942367025 | 0.306 | 0.869 | 0 |

| | | | | | |
|---|---|---|---|---|---|
| **TP53BP2** | 0 | -1.00461978518998 | 0.825 | 0.984 | 0 |
| **CNIH4** | 0 | 1.74056382322811 | 0.999 | 0.776 | 0 |
| **H3F3A** | 0 | -1.01265015828241 | 1 | 1 | 0 |
| **COA6** | 0 | 1.06514503872496 | 0.97 | 0.753 | 0 |
| **LGALS8** | 0 | 1.20992452954507 | 0.642 | 0.016 | 0 |
| **ACP1** | 0 | 1.06334788835381 | 0.949 | 0.594 | 0 |
| **ID2** | 0 | -2.19072609278297 | 0.152 | 0.855 | 0 |
| **KIDINS220** | 0 | -1.03978201410031 | 0.773 | 0.975 | 0 |
| **IAH1** | 0 | 2.10726502892887 | 0.903 | 0.101 | 0 |
| **WDR35** | 0 | -1.17558892680339 | 0.203 | 0.731 | 0 |
| **NCOA1** | 0 | 1.09376979040223 | 0.989 | 0.916 | 0 |
| **EFR3B** | 0 | 1.64579216804518 | 0.997 | 0.829 | 0 |
| **KRTCAP3** | 0 | 2.04624587779378 | 0.955 | 0.313 | 0 |
| **CAPN13** | 0 | -1.23200742696338 | 0.042 | 0.667 | 0 |
| **YIPF4** | 0 | 1.0686344629654 | 0.998 | 0.955 | 0 |
| **PKDCC** | 0 | -3.5766215336561 | 0.103 | 0.767 | 0 |
| **EML4** | 0 | -1.12933163129565 | 0.946 | 0.997 | 0 |
| **ZFP36L2** | 0 | -1.40892285940836 | 0.899 | 0.99 | 0 |
| **PSME4** | 0 | 1.06568570654899 | 1 | 0.998 | 0 |
| **RTN4** | 0 | -1.07931612366404 | 1 | 1 | 0 |
| **CCDC88A** | 0 | -1.41663756333174 | 0.751 | 0.991 | 0 |
| **XPO1** | 0 | 1.11541175284789 | 0.999 | 0.926 | 0 |
| **B3GNT2** | 0 | -1.89374298395464 | 0.803 | 0.989 | 0 |
| **UGP2** | 0 | 1.0151969988633 | 0.999 | 0.938 | 0 |
| **CEP68** | 0 | -1.78277449702385 | 0.307 | 0.933 | 0 |
| **SPRED2** | 0 | -1.38138616469773 | 0.238 | 0.795 | 0 |
| **ANTXR1** | 0 | -1.32090896404433 | 0.151 | 0.728 | 0 |
| **GFPT1** | 0 | 1.32908390000981 | 1 | 0.998 | 0 |
| **SNRPG** | 0 | 1.50501784960341 | 1 | 0.917 | 0 |
| **FAM136A** | 0 | 1.12067458709128 | 1 | 0.959 | 0 |
| **MTHFD2** | 0 | 1.36176891086865 | 1 | 0.994 | 0 |
| **WBP1** | 0 | -1.09733795308345 | 0.846 | 0.986 | 0 |
| **HK2** | 0 | 1.35161037262091 | 1 | 0.965 | 0 |
| **TMSB10** | 0 | -1.10766963162741 | 1 | 1 | 0 |

| | | | | | |
|---|---|---|---|---|---|
| **TCF7L1** | 0 | -1.94659992798029 | 0.177 | 0.741 | 0 |
| **CAPG** | 0 | 0.952009427353266 | 1 | 0.936 | 0 |
| **REEP1** | 0 | 1.71833449181828 | 0.83 | 0.085 | 0 |
| **CNOT11** | 0 | 0.963621080464845 | 1 | 0.998 | 0 |
| **MAP4K4** | 0 | -1.20305289220338 | 0.998 | 1 | 0 |
| **DBI** | 0 | -0.997206883610679 | 0.986 | 0.999 | 0 |
| **GLI2** | 0 | -1.94276415078555 | 0.052 | 0.858 | 0 |
| **TFCP2L1** | 0 | 1.71086773039761 | 0.991 | 0.631 | 0 |
| **NIFK** | 0 | 1.06758637097996 | 0.998 | 0.941 | 0 |
| **TSN** | 0 | -1.13405558407795 | 0.992 | 0.999 | 0 |
| **BIN1** | 0 | -2.61377150006632 | 0.32 | 0.972 | 0 |
| **PLEKHB2** | 0 | 0.871773855479217 | 1 | 0.999 | 0 |
| **LYPD1** | 0 | -2.19457684085367 | 0.04 | 0.65 | 0 |
| **ZEB2** | 0 | -1.71257030929973 | 0.057 | 0.601 | 0 |
| **KIF5C** | 0 | -2.14821388029558 | 0.69 | 0.998 | 0 |
| **NMI** | 0 | 1.51674789867663 | 0.981 | 0.522 | 0 |
| **RPRM** | 0 | -2.02104020748272 | 0.304 | 0.868 | 0 |
| **CERS6** | 0 | -1.40319755643784 | 0.193 | 0.802 | 0 |
| **SP5** | 0 | -3.53399769673338 | 0.208 | 0.946 | 0 |
| **GAD1** | 0 | -2.32726655677921 | 0.052 | 0.874 | 0 |
| **UBE2E3** | 0 | -1.48084820336506 | 0.878 | 0.997 | 0 |
| **SLC39A10** | 0 | 0.912841206820542 | 0.999 | 0.993 | 0 |
| **CFLAR** | 0 | 1.15942767579603 | 0.997 | 0.863 | 0 |
| **FZD7** | 0 | -4.01941580265003 | 0.114 | 0.9 | 0 |
| **ICA1L** | 0 | 2.09002154259585 | 0.927 | 0.139 | 0 |
| **CCNYL1** | 0 | -1.65279851919231 | 0.45 | 0.972 | 0 |
| **FZD5** | 0 | 1.21537060556339 | 1 | 0.952 | 0 |
| **CRYGD** | 0 | 1.46421616739745 | 0.67 | 0.016 | 0 |
| **IDH1** | 0 | 1.42186017894726 | 1 | 0.975 | 0 |
| **BARD1** | 0 | 1.11693307438238 | 0.985 | 0.812 | 0 |
| **TMBIM1** | 0 | 2.38822347560167 | 0.997 | 0.448 | 0 |
| **PTPRN** | 0 | 1.32352817014264 | 0.724 | 0.131 | 0 |
| **GMPPA** | 0 | 1.0008874170086 | 0.983 | 0.836 | 0 |
| **OBSL1** | 0 | -2.36322380898321 | 0.267 | 0.971 | 0 |

| | | | | | |
|---|---|---|---|---|---|
| **SERPINE2** | 0 | -1.41414379529685 | 0.466 | 0.91 | 0 |
| **FBXO36** | 0 | 0.992314229637616 | 0.997 | 0.873 | 0 |
| **ALPG** | 0 | 4.01579572944578 | 1 | 0.61 | 0 |
| **NGEF** | 0 | 1.80508420820049 | 0.835 | 0.117 | 0 |
| **ARL4C** | 0 | -2.75952903885934 | 0.089 | 0.616 | 0 |
| **GBX2** | 0 | -1.46704257409049 | 0.097 | 0.682 | 0 |
| **LRRN1** | 0 | -3.2638684536476 | 0.27 | 0.977 | 0 |
| **EDEM1** | 0 | 1.33898902658457 | 0.989 | 0.814 | 0 |
| **SLC6A6** | 0 | 1.81229151592433 | 0.999 | 0.863 | 0 |
| **RAB5A** | 0 | 1.19407870713121 | 1 | 0.949 | 0 |
| **TOP2B** | 0 | -0.827330746377559 | 0.981 | 0.998 | 0 |
| **TRIM71** | 0 | -1.5126510954223 | 1 | 0.999 | 0 |
| **ACVR2B** | 0 | -0.93355806447884 | 0.993 | 1 | 0 |
| **EIF1B** | 0 | -0.944765991267614 | 0.996 | 1 | 0 |
| **CTNNB1** | 0 | -1.80740015363363 | 0.974 | 0.999 | 0 |
| **HIGD1A** | 0 | 1.06767832298004 | 0.941 | 0.613 | 0 |
| **ZNF502** | 0 | 0.995797350262847 | 0.579 | 0.008 | 0 |
| **KIF9** | 0 | 1.42577245896636 | 0.975 | 0.677 | 0 |
| **SHISA5** | 0 | 0.87043024972084 | 1 | 0.999 | 0 |
| **PRKAR2A** | 0 | 1.12563615102817 | 1 | 0.972 | 0 |
| **MAPKAPK3** | 0 | -1.10443457708382 | 0.886 | 0.99 | 0 |
| **NT5DC2** | 0 | -1.29239577655178 | 0.997 | 1 | 0 |
| **WNT5A** | 0 | -1.71577221412401 | 0.026 | 0.721 | 0 |
| **MAGI1** | 0 | -1.59297338247954 | 0.286 | 0.873 | 0 |
| **PDZRN3** | 0 | 2.03782301549208 | 0.943 | 0.631 | 0 |
| **ZNF717** | 0 | 0.937430658679249 | 0.552 | 0.002 | 0 |
| **DCBLD2** | 0 | -1.45846047999226 | 0.759 | 0.992 | 0 |
| **TBC1D23** | 0 | 1.04057899599293 | 1 | 0.975 | 0 |
| **NIT2** | 0 | 1.4664305748626 | 0.983 | 0.646 | 0 |
| **TOMM70** | 0 | 0.942247232351001 | 1 | 0.993 | 0 |
| **ALCAM** | 0 | -1.74716479499125 | 0.033 | 0.7 | 0 |
| **DPPA2** | 0 | 2.16463609503173 | 0.99 | 0.457 | 0 |
| **NECTIN3** | 0 | -1.86815987859945 | 0.294 | 0.917 | 0 |
| **ATG3** | 0 | 2.10698045460043 | 1 | 0.995 | 0 |

| | | | | | |
|---|---|---|---|---|---|
| **CCDC80** | 0 | 1.22786116811784 | 0.984 | 0.75 | 0 |
| **NAA50** | 0 | 1.56816465490248 | 0.964 | 0.447 | 0 |
| **B4GALT4** | 0 | 1.36372283447991 | 0.893 | 0.396 | 0 |
| **ARHGAP31** | 0 | 1.06477119996016 | 0.646 | 0.079 | 0 |
| **FSTL1** | 0 | -2.41546948135816 | 0.831 | 0.978 | 0 |
| **NDUFB4** | 0 | 1.3052002703599 | 0.895 | 0.363 | 0 |
| **HCLS1** | 0 | 1.584004281172 | 0.73 | 0.081 | 0 |
| **SNX4** | 0 | 0.886651211624844 | 1 | 0.987 | 0 |
| **MGLL** | 0 | 1.61343657280328 | 0.763 | 0.048 | 0 |
| **PLXND1** | 0 | 1.95141306097569 | 1 | 0.761 | 0 |
| **TRH** | 0 | 1.20789253139083 | 0.666 | 0.083 | 0 |
| **RAB6B** | 0 | -1.54390260260754 | 0.3 | 0.826 | 0 |
| **AMOTL2** | 0 | -2.40555428860433 | 0.809 | 0.996 | 0 |
| **PPP2R3A** | 0 | -1.13813087171654 | 0.498 | 0.917 | 0 |
| **RBP1** | 0 | -2.440443334505 | 0.261 | 0.939 | 0 |
| **SPSB4** | 0 | -2.34001700532841 | 0.214 | 0.927 | 0 |
| **CHST2** | 0 | 2.14839393525543 | 0.97 | 0.512 | 0 |
| **PFN2** | 0 | -1.36094199435594 | 0.999 | 1 | 0 |
| **LRRC34** | 0 | 1.14565349657926 | 0.628 | 0.005 | 0 |
| **GNB4** | 0 | -1.58027791407899 | 0.627 | 0.972 | 0 |
| **LIPH** | 0 | 2.0901654180203 | 0.873 | 0.077 | 0 |
| **LPP** | 0 | -2.18719152148829 | 0.305 | 0.933 | 0 |
| **HES1** | 0 | -2.36372852472429 | 0.05 | 0.724 | 0 |
| **NRROS** | 0 | 1.29507317266466 | 0.717 | 0.056 | 0 |
| **PAK2** | 0 | 1.03935019003405 | 0.991 | 0.87 | 0 |
| **ZNF595** | 0 | 2.03728419851068 | 0.999 | 0.75 | 0 |
| **ZNF732** | 0 | 1.06354502874449 | 0.597 | 0.005 | 0 |
| **RNF212** | 0 | 1.22608389583477 | 0.644 | 0.005 | 0 |
| **TNIP2** | 0 | 0.979940499448544 | 0.997 | 0.945 | 0 |
| **NSG1** | 0 | 1.38288438542803 | 0.709 | 0.086 | 0 |
| **S100P** | 0 | 2.32849324385396 | 0.887 | 0.064 | 0 |
| **PROM1** | 0 | 1.1920694091678 | 0.996 | 0.858 | 0 |
| **PI4K2B** | 0 | 1.37326225531879 | 0.992 | 0.864 | 0 |
| **SEL1L3** | 0 | -1.19467159124981 | 0.064 | 0.589 | 0 |

| | | | | | |
|---|---|---|---|---|---|
| **RELL1** | 0 | 1.24330621507821 | 0.977 | 0.753 | 0 |
| **PDS5A** | 0 | 0.933695226909692 | 0.976 | 0.785 | 0 |
| **UCHL1** | 0 | -2.56393331539321 | 0.637 | 0.992 | 0 |
| **LIMCH1** | 0 | -1.76112941892387 | 0.401 | 0.837 | 0 |
| **COX7B2** | 0 | 1.44582127559576 | 0.616 | 0.005 | 0 |
| **TEC** | 0 | 1.25394979508179 | 0.743 | 0.113 | 0 |
| **SGCB** | 0 | -1.08936128481032 | 0.598 | 0.929 | 0 |
| **PAICS** | 0 | 1.39384274778475 | 0.997 | 0.806 | 0 |
| **IGFBP7** | 0 | -1.86850284019326 | 0.077 | 0.615 | 0 |
| **GRSF1** | 0 | 0.970948780494087 | 1 | 0.987 | 0 |
| **G3BP2** | 0 | -1.14165085630888 | 1 | 1 | 0 |
| **TMEM150C** | 0 | 1.99689771395812 | 0.927 | 0.163 | 0 |
| **MAPK10** | 0 | -2.13374594483217 | 0.03 | 0.706 | 0 |
| **SPP1** | 0 | -2.72016035237411 | 0.176 | 0.813 | 0 |
| **GRID2** | 0 | -1.59752149816227 | 0.091 | 0.653 | 0 |
| **EIF4E** | 0 | 0.932532621341142 | 1 | 0.96 | 0 |
| **DDIT4L** | 0 | 1.07869766055954 | 0.585 | 0.036 | 0 |
| **PAPSS1** | 0 | -0.924366651892264 | 0.969 | 0.998 | 0 |
| **MCUB** | 0 | 1.56136997560188 | 0.945 | 0.484 | 0 |
| **GAR1** | 0 | 0.98027770251196 | 0.988 | 0.892 | 0 |
| **CAMK2D** | 0 | -1.57380095749953 | 0.176 | 0.801 | 0 |
| **PRSS12** | 0 | 3.17154216259656 | 1 | 0.562 | 0 |
| **MAD2L1** | 0 | 1.44197505647762 | 0.947 | 0.613 | 0 |
| **FGF2** | 0 | -1.38077469297514 | 0.871 | 0.986 | 0 |
| **JADE1** | 0 | -1.35191391322069 | 0.715 | 0.983 | 0 |
| **PCDH18** | 0 | -2.52367434319582 | 0.484 | 0.929 | 0 |
| **OTUD4** | 0 | 1.19633235649218 | 0.989 | 0.83 | 0 |
| **SFRP2** | 0 | -2.52336893308846 | 0.055 | 0.722 | 0 |
| **TMEM144** | 0 | 1.3119927165305 | 0.748 | 0.205 | 0 |
| **APELA** | 0 | 1.399049133004 | 0.998 | 0.861 | 0 |
| **TRIM61** | 0 | 1.50307388288221 | 0.907 | 0.332 | 0 |
| **TRIM60** | 0 | 2.56286725959346 | 0.989 | 0.189 | 0 |
| **TRIM75P** | 0 | 2.69094732485696 | 0.987 | 0.217 | 0 |
| **FAT1** | 0 | -2.44528205280829 | 0.751 | 0.999 | 0 |

| | | | | | |
|---|---|---|---|---|---|
| **ZFP42** | 0 | 3.92914096531734 | 1 | 0.385 | 0 |
| **CLPTM1L** | 0 | 1.00187966030491 | 1 | 0.995 | 0 |
| **IRX2** | 0 | -1.49528030946885 | 0.858 | 0.989 | 0 |
| **IRX1** | 0 | -2.28407538050473 | 0.138 | 0.797 | 0 |
| **SUB1** | 0 | 1.7332748083932 | 0.96 | 0.378 | 0 |
| **DAB2** | 0 | 1.0198464288746 | 0.581 | 0.022 | 0 |
| **SELENOP** | 0 | -2.3007035981013 | 0.237 | 0.941 | 0 |
| **CCL28** | 0 | 0.902213648927535 | 0.536 | 0.004 | 0 |
| **FST** | 0 | -3.36547402969692 | 0.108 | 0.828 | 0 |
| **PLPP1** | 0 | 1.08533977559861 | 1 | 0.968 | 0 |
| **ZSWIM6** | 0 | -1.30185221774904 | 0.122 | 0.706 | 0 |
| **ERBIN** | 0 | 1.40205396109473 | 1 | 0.957 | 0 |
| **MAP1B** | 0 | -3.22312782910389 | 0.799 | 1 | 0 |
| **ARSB** | 0 | -1.1771986189159 | 0.54 | 0.914 | 0 |
| **VCAN** | 0 | -3.78978661334364 | 0.418 | 0.996 | 0 |
| **CAST** | 0 | -1.70873538656723 | 0.113 | 0.808 | 0 |
| **PAM** | 0 | -1.65665366892432 | 0.088 | 0.723 | 0 |
| **EFNA5** | 0 | -2.34127678277533 | 0.137 | 0.899 | 0 |
| **MCC** | 0 | -1.48266939941135 | 0.058 | 0.74 | 0 |
| **ZNF608** | 0 | -2.1661041430561 | 0.213 | 0.946 | 0 |
| **ADAMTS19** | 0 | -2.13795491500637 | 0.016 | 0.724 | 0 |
| **HINT1** | 0 | 1.23658749307815 | 1 | 0.977 | 0 |
| **WNT8A** | 0 | -2.41953443046584 | 0.086 | 0.785 | 0 |
| **REEP2** | 0 | -1.40580927220375 | 0.179 | 0.709 | 0 |
| **CXXC5** | 0 | -1.94721118015745 | 0.735 | 0.994 | 0 |
| **DND1** | 0 | 1.38800928019284 | 0.713 | 0.123 | 0 |
| **PCDHB2** | 0 | -1.90309741627969 | 0.273 | 0.917 | 0 |
| **TAF7** | 0 | -0.905211953001234 | 0.985 | 1 | 0 |
| **PCDHGB7** | 0 | -1.56298067404328 | 0.15 | 0.758 | 0 |
| **DIAPH1** | 0 | -1.34239027767328 | 0.992 | 0.999 | 0 |
| **HDAC3** | 0 | -0.943226534519253 | 0.855 | 0.984 | 0 |
| **PCDH1** | 0 | -2.44854141673366 | 0.658 | 0.987 | 0 |
| **GNPDA1** | 0 | -1.21845804701199 | 0.8 | 0.984 | 0 |
| **LARS** | 0 | 0.935131541109834 | 1 | 1 | 0 |

| | | | | | |
|---|---|---|---|---|---|
| DPYSL3 | 0 | -3.77942592754712 | 0.656 | 0.999 | 0 |
| AFAP1L1 | 0 | 1.49874726541591 | 0.818 | 0.156 | 0 |
| UBLCP1 | 0 | -0.8784054479057 | 0.948 | 0.997 | 0 |
| PANK3 | 0 | 1.31956164225819 | 0.995 | 0.784 | 0 |
| ERGIC1 | 0 | -1.05742337633733 | 0.628 | 0.953 | 0 |
| NSD1 | 0 | 0.852499186179235 | 0.999 | 0.967 | 0 |
| DBN1 | 0 | -1.73534646738686 | 0.986 | 1 | 0 |
| PDLIM7 | 0 | -1.66686910674686 | 0.963 | 0.997 | 0 |
| CANX | 0 | 1.00148253129069 | 1 | 0.997 | 0 |
| GFPT2 | 0 | -1.34841277993522 | 0.555 | 0.931 | 0 |
| SERPINB1 | 0 | 1.98379562170999 | 0.915 | 0.106 | 0 |
| SERPINB6 | 0 | 2.80371212480287 | 1 | 0.704 | 0 |
| TUBB2B | 0 | -1.82276427185392 | 0.997 | 1 | 0 |
| SSR1 | 0 | 1.02520240417794 | 0.973 | 0.761 | 0 |
| EDN1 | 0 | -2.81980206026226 | 0.046 | 0.814 | 0 |
| NOL7 | 0 | 0.846850587914644 | 0.998 | 0.961 | 0 |
| JARID2 | 0 | -1.32239379466602 | 0.968 | 0.991 | 0 |
| HDGFL1 | 0 | 1.31320209021546 | 0.626 | 0.005 | 0 |
| HIST1H1A | 0 | 4.74575440363698 | 0.998 | 0.424 | 0 |
| HIST1H2BB | 0 | 1.9898845407715 | 0.623 | 0.056 | 0 |
| ZFP57 | 0 | 1.99448482495302 | 0.956 | 0.238 | 0 |
| TRIM26 | 0 | 1.13129645184268 | 0.998 | 0.928 | 0 |
| NRM | 0 | -1.34934468161386 | 0.34 | 0.848 | 0 |
| FLOT1 | 0 | -1.40492616217094 | 0.894 | 0.997 | 0 |
| DDAH2 | 0 | -1.93216851932674 | 0.64 | 0.988 | 0 |
| PBX2 | 0 | -0.879789334349049 | 0.997 | 0.998 | 0 |
| BAK1 | 0 | 1.06916623135365 | 1 | 0.998 | 0 |
| HMGA1 | 0 | -1.18388923640622 | 0.979 | 0.992 | 0 |
| ILRUN | 0 | 2.04724511250444 | 1 | 0.996 | 0 |
| SNRPC | 0 | 1.09296123098036 | 1 | 1 | 0 |
| ANKS1A | 0 | -0.943399880554051 | 0.776 | 0.98 | 0 |
| MAPK13 | 0 | 1.44796130333718 | 0.865 | 0.277 | 0 |
| TOMM6 | 0 | 1.20545899578148 | 1 | 0.956 | 0 |
| CCND3 | 0 | 1.77499360902403 | 0.991 | 0.709 | 0 |

| | | | | | |
|---|---|---|---|---|---|
| **PTK7** | 0 | -1.66805133552516 | 0.856 | 0.989 | 0 |
| **DNPH1** | 0 | 0.995726864239056 | 0.993 | 0.908 | 0 |
| **AARS2** | 0 | 1.10034289230808 | 0.997 | 0.917 | 0 |
| **TNFRSF21** | 0 | -1.47988252904714 | 0.935 | 0.996 | 0 |
| **ADGRB3** | 0 | 1.77697462104436 | 0.774 | 0.003 | 0 |
| **DPPA5** | 0 | 6.15549992877459 | 1 | 0.524 | 0 |
| **KHDC3L** | 0 | 5.32894281879333 | 1 | 0.416 | 0 |
| **OOEP** | 0 | 1.45400774285203 | 0.741 | 0.005 | 0 |
| **DDX43** | 0 | 1.82740034642446 | 0.857 | 0.029 | 0 |
| **CGAS** | 0 | 1.33921260139823 | 0.618 | 0.028 | 0 |
| **IRAK1BP1** | 0 | 1.79007930213616 | 0.822 | 0.01 | 0 |
| **CCNC** | 0 | 1.38406186956004 | 1 | 0.981 | 0 |
| **LIN28B** | 0 | 1.21080916489527 | 1 | 0.993 | 0 |
| **ATG5** | 0 | 2.01229540061612 | 1 | 0.949 | 0 |
| **CD24** | 0 | -2.18231528979942 | 0.805 | 0.982 | 0 |
| **SEC63** | 0 | 1.51013171723121 | 1 | 0.989 | 0 |
| **SNX3** | 0 | 1.54944712659315 | 0.998 | 0.828 | 0 |
| **AFG1L** | 0 | 1.14834936903543 | 0.918 | 0.587 | 0 |
| **WASF1** | 0 | -1.64181168803496 | 0.48 | 0.94 | 0 |
| **GTF3C6** | 0 | 1.91070858571854 | 0.997 | 0.664 | 0 |
| **SLC16A10** | 0 | 2.7517105639229 | 1 | 0.544 | 0 |
| **MARCKS** | 0 | -3.72117594638411 | 0.32 | 0.997 | 0 |
| **GJA1** | 0 | -1.18439963996353 | 1 | 1 | 0 |
| **PKIB** | 0 | 1.5288478003462 | 0.721 | 0.047 | 0 |
| **SMPDL3A** | 0 | 1.65539749480577 | 0.959 | 0.403 | 0 |
| **ECHDC1** | 0 | -2.00784406301249 | 0.111 | 0.935 | 0 |
| **LAMA2** | 0 | -1.61919469706142 | 0.134 | 0.662 | 0 |
| **SGK1** | 0 | 1.26257848086649 | 0.994 | 0.817 | 0 |
| **HEBP2** | 0 | 1.24395071558396 | 1 | 0.988 | 0 |
| **CITED2** | 0 | -2.16509175348603 | 0.678 | 0.975 | 0 |
| **AIG1** | 0 | -1.57876857590997 | 0.268 | 0.886 | 0 |
| **FUCA2** | 0 | -1.24900717261228 | 0.813 | 0.965 | 0 |
| **PHACTR2** | 0 | -1.70357869923457 | 0.302 | 0.897 | 0 |
| **UTRN** | 0 | -1.80811809654178 | 0.532 | 0.936 | 0 |

| | | | | | |
|---|---|---|---|---|---|
| **NUP43** | 0 | 0.870351741389339 | 0.999 | 0.971 | 0 |
| **RMND1** | 0 | 1.38770997724503 | 1 | 0.952 | 0 |
| **ARMT1** | 0 | 1.38697094798004 | 1 | 0.988 | 0 |
| **TCP1** | 0 | 2.50207248459196 | 1 | 0.924 | 0 |
| **IGF2R** | 0 | -1.94270237098661 | 0.523 | 0.978 | 0 |
| **SFT2D1** | 0 | 1.08546481709419 | 0.904 | 0.492 | 0 |
| **DACT2** | 0 | 1.82769010396769 | 0.863 | 0.095 | 0 |
| **FAM20C** | 0 | -1.71356718911914 | 0.269 | 0.88 | 0 |
| **SNX8** | 0 | 1.05079027954503 | 1 | 0.981 | 0 |
| **MPP6** | 0 | -1.10182006380859 | 0.967 | 0.995 | 0 |
| **CYCS** | 0 | 1.96949510106133 | 0.995 | 0.613 | 0 |
| **CBX3** | 0 | 2.36999861601979 | 1 | 0.834 | 0 |
| **CPVL** | 0 | -0.975838828968673 | 0.992 | 0.998 | 0 |
| **WIPF3** | 0 | 1.28519959951893 | 0.813 | 0.233 | 0 |
| **DPY19L1** | 0 | 1.2852401084748 | 0.932 | 0.538 | 0 |
| **HERPUD2** | 0 | -1.40106975073043 | 0.377 | 0.9 | 0 |
| **STARD3NL** | 0 | -1.19942382137951 | 0.749 | 0.978 | 0 |
| **HUS1** | 0 | 1.24569902865938 | 0.999 | 0.98 | 0 |
| **SUN3** | 0 | 1.79670638953565 | 0.831 | 0.031 | 0 |
| **UPP1** | 0 | 1.77103509331846 | 1 | 0.952 | 0 |
| **SEC61G** | 0 | 0.864044133173176 | 1 | 0.999 | 0 |
| **CHCHD2** | 0 | 1.85310865869328 | 1 | 0.997 | 0 |
| **ZNF727** | 0 | 1.57956422258424 | 0.793 | 0.006 | 0 |
| **ZNF736** | 0 | 1.14497086993501 | 0.675 | 0.008 | 0 |
| **AUTS2** | 0 | -2.09989405113305 | 0.456 | 0.94 | 0 |
| **LIMK1** | 0 | -1.00716193293451 | 0.918 | 0.995 | 0 |
| **CACNA2D1** | 0 | 1.19944101039783 | 0.983 | 0.784 | 0 |
| **CDK6** | 0 | -2.43088553686221 | 0.111 | 0.943 | 0 |
| **COL1A2** | 0 | -3.22168698228147 | 0.393 | 0.95 | 0 |
| **SLC25A13** | 0 | -0.919336169169632 | 0.964 | 0.999 | 0 |
| **ARPC1B** | 0 | 1.5769265793858 | 1 | 0.982 | 0 |
| **STAG3** | 0 | 1.64184986087995 | 0.765 | 0.019 | 0 |
| **TRIP6** | 0 | -1.1361664469827 | 0.995 | 1 | 0 |
| **COL26A1** | 0 | -2.99655516400709 | 0.162 | 0.97 | 0 |

| | | | | | |
|---|---|---|---|---|---|
| **CUX1** | 0 | -1.28537326939407 | 0.629 | 0.962 | 0 |
| **PRKAR2B** | 0 | -1.81423876754599 | 0.34 | 0.917 | 0 |
| **LAMB1** | 0 | -1.8428343947173 | 0.779 | 0.996 | 0 |
| **LRRN3** | 0 | -1.48086420014237 | 0.019 | 0.641 | 0 |
| **PTPRZ1** | 0 | -1.5635003482242 | 0.034 | 0.557 | 0 |
| **AASS** | 0 | 1.4684866042837 | 0.999 | 0.953 | 0 |
| **NDUFA5** | 0 | 1.31948159180528 | 0.963 | 0.586 | 0 |
| **HYAL4** | 0 | 2.40077939801189 | 0.957 | 0.129 | 0 |
| **KLHDC10** | 0 | -1.1979071261706 | 0.176 | 0.774 | 0 |
| **MEST** | 0 | 1.8066753043317 | 0.996 | 0.768 | 0 |
| **PODXL** | 0 | -4.86321449682106 | 0.549 | 0.997 | 0 |
| **AKR1B1** | 0 | -1.00052058846278 | 0.964 | 0.998 | 0 |
| **CALD1** | 0 | -1.93031885440602 | 0.996 | 1 | 0 |
| **TRIM24** | 0 | -2.00207259824614 | 0.973 | 1 | 0 |
| **KLRG2** | 0 | 0.961838090907368 | 1 | 0.926 | 0 |
| **CASP2** | 0 | -1.27868714337147 | 0.649 | 0.941 | 0 |
| **TCAF1** | 0 | -2.06385472993419 | 0.286 | 0.954 | 0 |
| **REPIN1** | 0 | -1.51663585143529 | 0.485 | 0.955 | 0 |
| **ABCB8** | 0 | 0.914525125970499 | 1 | 0.963 | 0 |
| **PPP1R3B** | 0 | -2.08292586946679 | 0.639 | 0.964 | 0 |
| **TUSC3** | 0 | -0.875671132001443 | 0.98 | 1 | 0 |
| **ZDHHC2** | 0 | -1.10193317023928 | 0.047 | 0.601 | 0 |
| **XPO7** | 0 | 0.868212056568408 | 1 | 0.998 | 0 |
| **PIWIL2** | 0 | 1.20240451201235 | 0.624 | 0.006 | 0 |
| **LOXL2** | 0 | -1.43149248084758 | 0.685 | 0.975 | 0 |
| **DPYSL2** | 0 | -3.09704517175003 | 0.487 | 0.999 | 0 |
| **CLU** | 0 | -3.21480291073775 | 0.811 | 1 | 0 |
| **RBPMS** | 0 | -0.80067629951755 | 0.996 | 0.999 | 0 |
| **GSR** | 0 | -1.49648453587079 | 0.745 | 0.993 | 0 |
| **NRG1** | 0 | 2.13740187896367 | 0.897 | 0.145 | 0 |
| **EIF4EBP1** | 0 | 1.09241207596032 | 1 | 0.996 | 0 |
| **ASH2L** | 0 | 1.21348386661843 | 1 | 0.99 | 0 |
| **FGFR1** | 0 | -3.07659164570914 | 0.624 | 0.996 | 0 |
| **TGS1** | 0 | 0.802582377414409 | 1 | 0.987 | 0 |

| | | | | | |
|---|---|---|---|---|---|
| **CHD7** | 0 | -2.19346331698033 | 0.561 | 0.958 | 0 |
| **MTFR1** | 0 | 1.6166816463579 | 1 | 0.868 | 0 |
| **PREX2** | 0 | -1.69425070413626 | 0.119 | 0.778 | 0 |
| **CRISPLD1** | 0 | -1.70124646702179 | 0.503 | 0.933 | 0 |
| **FABP5** | 0 | 3.52837131550265 | 0.999 | 0.552 | 0 |
| **CA13** | 0 | 1.81598622619714 | 0.94 | 0.443 | 0 |
| **TMEM64** | 0 | -3.0922462168466 | 0.748 | 0.995 | 0 |
| **CTHRC1** | 0 | 1.7695809554277 | 0.835 | 0.07 | 0 |
| **EIF3H** | 0 | 0.832200285947537 | 1 | 0.997 | 0 |
| **EXT1** | 0 | -1.33034642164606 | 0.757 | 0.98 | 0 |
| **DEPTOR** | 0 | 1.50588589059066 | 1 | 0.802 | 0 |
| **ZHX2** | 0 | -1.8500680660187 | 0.07 | 0.828 | 0 |
| **MYC** | 0 | -2.14710126752009 | 0.573 | 0.966 | 0 |
| **EFR3A** | 0 | 1.22445586585712 | 1 | 0.973 | 0 |
| **TOP1MT** | 0 | 1.12722867436274 | 0.999 | 0.954 | 0 |
| **NRBP2** | 0 | 1.53760684964964 | 0.926 | 0.541 | 0 |
| **SMARCA2** | 0 | 1.60629755088897 | 1 | 0.879 | 0 |
| **GLDC** | 0 | 1.94212572562817 | 0.999 | 0.809 | 0 |
| **KDM4C** | 0 | 1.14973288784216 | 0.992 | 0.853 | 0 |
| **DMAC1** | 0 | 0.988470243894456 | 1 | 0.988 | 0 |
| **PSIP1** | 0 | -1.2163147114332 | 0.922 | 0.992 | 0 |
| **AQP3** | 0 | 1.71848302535933 | 0.827 | 0.156 | 0 |
| **UBAP2** | 0 | 1.21472427850253 | 1 | 0.977 | 0 |
| **STOML2** | 0 | 0.92696234993462 | 1 | 0.999 | 0 |
| **TPM2** | 0 | -1.70469610976585 | 0.973 | 0.999 | 0 |
| **IGFBPL1** | 0 | -3.19176408799257 | 0.229 | 0.981 | 0 |
| **ANKRD18A** | 0 | -1.60310440090826 | 0.073 | 0.769 | 0 |
| **PCSK5** | 0 | 1.82499727243847 | 0.976 | 0.421 | 0 |
| **GCNT1** | 0 | 1.90334093337593 | 0.979 | 0.544 | 0 |
| **UBQLN1** | 0 | 0.898134451608658 | 0.999 | 0.959 | 0 |
| **C9orf64** | 0 | 1.66567990657579 | 0.819 | 0.017 | 0 |
| **HNRNPK** | 0 | 0.917228728032814 | 1 | 0.942 | 0 |
| **GAS1** | 0 | -1.12277345432497 | 0.062 | 0.621 | 0 |
| **BARX1** | 0 | 1.60702263872728 | 0.743 | 0.019 | 0 |

| | | | | | |
|---|---|---|---|---|---|
| **FBP1** | 0 | 2.23362236194505 | 0.815 | 0.102 | 0 |
| **PTCH1** | 0 | -1.49860986075931 | 0.433 | 0.882 | 0 |
| **NANS** | 0 | 0.841406654977462 | 0.999 | 0.988 | 0 |
| **TGFBR1** | 0 | -1.36484437054168 | 0.957 | 0.999 | 0 |
| **SEC61B** | 0 | 1.14218177726782 | 1 | 0.921 | 0 |
| **ABCA1** | 0 | 1.14409210484653 | 1 | 0.946 | 0 |
| **KLF4** | 0 | 2.4553145762466 | 0.999 | 0.685 | 0 |
| **PALM2-AKAI** | 0 | -2.93094577820843 | 0.53 | 0.997 | 0 |
| **GSN** | 0 | -2.35911287647412 | 0.076 | 0.831 | 0 |
| **HSPA5** | 0 | 2.04855573101416 | 0.996 | 0.904 | 0 |
| **AIF1L** | 0 | -1.59013884160467 | 0.976 | 0.999 | 0 |
| **VAV2** | 0 | 1.44792661699563 | 0.998 | 0.918 | 0 |
| **OLFM1** | 0 | -1.50188411695749 | 0.292 | 0.81 | 0 |
| **AGPAT2** | 0 | 1.1093353214295 | 0.999 | 0.899 | 0 |
| **GDI2** | 0 | -0.872760158408431 | 0.997 | 1 | 0 |
| **ECHDC3** | 0 | 1.64920693296312 | 0.818 | 0.012 | 0 |
| **SEPHS1** | 0 | -1.22867055408246 | 0.978 | 0.995 | 0 |
| **FRMD4A** | 0 | -1.4730793891718 | 0.073 | 0.61 | 0 |
| **OLAH** | 0 | 3.44909267340694 | 0.994 | 0.28 | 0 |
| **VIM** | 0 | -4.65264313808121 | 0.504 | 0.981 | 0 |
| **NEBL** | 0 | -1.42964822357607 | 0.066 | 0.59 | 0 |
| **BMI1** | 0 | 1.14963084733074 | 0.865 | 0.415 | 0 |
| **MSRB2** | 0 | -1.08303528967992 | 0.627 | 0.948 | 0 |
| **ITGB1** | 0 | -1.10662186749169 | 0.987 | 1 | 0 |
| **MARCH8** | 0 | 1.24750151947034 | 0.985 | 0.772 | 0 |
| **TIMM23** | 0 | 0.959425038577637 | 1 | 0.997 | 0 |
| **NCOA4** | 0 | 0.96554805439952 | 0.998 | 0.95 | 0 |
| **IPMK** | 0 | 1.18902191634778 | 0.997 | 0.845 | 0 |
| **CISD1** | 0 | 1.22403102124653 | 0.997 | 0.886 | 0 |
| **CCDC6** | 0 | -1.11491037553138 | 0.956 | 0.998 | 0 |
| **NRBF2** | 0 | 1.08848261753715 | 1 | 0.983 | 0 |
| **JMJD1C** | 0 | -1.21049465798829 | 0.831 | 0.992 | 0 |
| **REEP3** | 0 | -1.08487432146457 | 0.578 | 0.925 | 0 |
| **HNRNPH3** | 0 | -0.96722010353841 | 1 | 1 | 0 |

| | | | | | |
|---|---|---|---|---|---|
| **SLC25A16** | 0 | 1.82067613784323 | 1 | 0.916 | 0 |
| **VPS26A** | 0 | 1.18269588052523 | 1 | 0.942 | 0 |
| **H2AFY2** | 0 | -1.80802823647613 | 0.743 | 0.97 | 0 |
| **PSAP** | 0 | -1.04795590687924 | 1 | 1 | 0 |
| **ZCCHC24** | 0 | -1.4018589755736 | 0.165 | 0.718 | 0 |
| **PRXL2A** | 0 | -1.13576597182876 | 0.977 | 0.999 | 0 |
| **CDHR1** | 0 | 1.89300141664972 | 0.971 | 0.682 | 0 |
| **LIPA** | 0 | -1.349270944402 | 0.286 | 0.831 | 0 |
| **FGFBP3** | 0 | -1.89956157227091 | 0.023 | 0.54 | 0 |
| **PDLIM1** | 0 | 1.19625516297863 | 1 | 0.963 | 0 |
| **PIK3AP1** | 0 | 1.42990110212124 | 0.764 | 0.048 | 0 |
| **NFKB2** | 0 | 1.39509810959335 | 1 | 0.897 | 0 |
| **C10orf95** | 0 | 1.56834936746874 | 0.868 | 0.287 | 0 |
| **MFSD13A** | 0 | 1.16774288390076 | 0.825 | 0.318 | 0 |
| **NT5C2** | 0 | -1.26362786893699 | 0.824 | 0.987 | 0 |
| **INA** | 0 | 1.22625516132966 | 0.983 | 0.771 | 0 |
| **PCGF6** | 0 | 1.52371980235598 | 0.97 | 0.568 | 0 |
| **ATP5MD** | 0 | 2.04252087404231 | 0.984 | 0.405 | 0 |
| **SLK** | 0 | 1.02037602494741 | 0.996 | 0.951 | 0 |
| **GSTO2** | 0 | 1.38006700698315 | 0.78 | 0.072 | 0 |
| **SMC3** | 0 | -0.971233426124236 | 0.98 | 0.998 | 0 |
| **TCF7L2** | 0 | -1.75130091069762 | 0.352 | 0.915 | 0 |
| **SHTN1** | 0 | -1.35953566125414 | 0.315 | 0.848 | 0 |
| **PRDX3** | 0 | 1.03567015412994 | 1 | 0.954 | 0 |
| **RGS10** | 0 | 1.12090114187002 | 0.892 | 0.448 | 0 |
| **FGFR2** | 0 | -1.14098093625265 | 0.961 | 0.995 | 0 |
| **NKX1-2** | 0 | -2.66233639047941 | 0.45 | 0.876 | 0 |
| **CFAP46** | 0 | 1.1757336829607 | 0.678 | 0.049 | 0 |
| **UTF1** | 0 | 3.21467719852587 | 1 | 0.429 | 0 |
| **VENTX** | 0 | 1.46347266212192 | 0.955 | 0.395 | 0 |
| **TUBGCP2** | 0 | 1.1516767311089 | 0.999 | 0.916 | 0 |
| **PRAP1** | 0 | 2.35014107096977 | 0.927 | 0.188 | 0 |
| **FUOM** | 0 | 1.18855727523442 | 0.979 | 0.763 | 0 |
| **PKP3** | 0 | 1.90623936803258 | 0.997 | 0.65 | 0 |

| | | | | | |
|---|---|---|---|---|---|
| **SIGIRR** | 0 | 1.1980234020323 | 0.987 | 0.751 | 0 |
| **HRAS** | 0 | -1.06772091696829 | 0.908 | 0.999 | 0 |
| **TALDO1** | 0 | 1.2329392342975 | 1 | 0.998 | 0 |
| **SLC25A22** | 0 | 1.08503979528925 | 0.991 | 0.869 | 0 |
| **TSPAN4** | 0 | -1.1711925408938 | 0.951 | 0.994 | 0 |
| **CD81** | 0 | 1.1514547887015 | 1 | 1 | 0 |
| **PHLDA2** | 0 | 1.51852469337201 | 0.961 | 0.607 | 0 |
| **TRIM6** | 0 | 1.67260417406488 | 0.906 | 0.203 | 0 |
| **ARFIP2** | 0 | 1.04106224037553 | 1 | 0.986 | 0 |
| **TMEM9B** | 0 | 1.11766699022722 | 0.995 | 0.898 | 0 |
| **DENND5A** | 0 | 0.947947555472212 | 0.997 | 0.957 | 0 |
| **TMEM41B** | 0 | 1.04302930302894 | 1 | 0.997 | 0 |
| **ADM** | 0 | -2.31020360122356 | 0.175 | 0.751 | 0 |
| **3,00 DKK** | 0 | -1.61412932416958 | 0.429 | 0.874 | 0 |
| **RRAS2** | 0 | 1.4185220635454 | 1 | 0.929 | 0 |
| **PSMA1** | 0 | 0.918068031677772 | 1 | 0.977 | 0 |
| **GTF2H1** | 0 | 1.48405049105522 | 1 | 0.898 | 0 |
| **PRMT3** | 0 | 1.04077339068707 | 0.997 | 0.907 | 0 |
| **LGR4** | 0 | 0.952636073233877 | 1 | 0.994 | 0 |
| **LIN7C** | 0 | 0.878699310316529 | 1 | 0.993 | 0 |
| **RCN1** | 0 | 1.59940982310148 | 1 | 0.96 | 0 |
| **EIF3M** | 0 | 0.932772844150987 | 1 | 1 | 0 |
| **APIP** | 0 | 1.5194759619057 | 0.995 | 0.798 | 0 |
| **PDHX** | 0 | 1.0286298539492 | 0.998 | 0.944 | 0 |
| **C11orf96** | 0 | -2.83004832311731 | 0.26 | 0.932 | 0 |
| **TP53I11** | 0 | -1.72877481234727 | 0.141 | 0.822 | 0 |
| **LARGE2** | 0 | 1.43265385778614 | 1 | 0.887 | 0 |
| **AMBRA1** | 0 | 1.1016385905303 | 1 | 0.979 | 0 |
| **ATG13** | 0 | 1.28401763875789 | 1 | 0.992 | 0 |
| **ARHGAP1** | 0 | 1.00503974277211 | 1 | 1 | 0 |
| **ARFGAP2** | 0 | 0.930199021751436 | 0.981 | 0.857 | 0 |
| **TIMM10** | 0 | 1.23928153065538 | 1 | 0.995 | 0 |
| **SERPING1** | 0 | -2.04819426215327 | 0.505 | 0.979 | 0 |
| **TMX2** | 0 | 0.990545479218267 | 1 | 0.998 | 0 |

| | | | | | |
|---|---|---|---|---|---|
| **TMEM258** | 0 | 1.7073702062196 | 1 | 0.869 | 0 |
| **FADS2** | 0 | -1.55668496570474 | 0.999 | 1 | 0 |
| **ASRGL1** | 0 | 1.94796481603701 | 1 | 0.954 | 0 |
| **EEF1G** | 0 | 1.59072850985943 | 1 | 0.999 | 0 |
| **POLR2G** | 0 | 0.846390903557255 | 1 | 0.998 | 0 |
| **PLAAT5** | 0 | -1.84733972106495 | 0.389 | 0.846 | 0 |
| **RCOR2** | 0 | -1.7212631770335 | 0.307 | 0.86 | 0 |
| **CATSPERZ** | 0 | 1.00128554523717 | 0.62 | 0.065 | 0 |
| **ESRRA** | 0 | 1.25092620524075 | 0.998 | 0.92 | 0 |
| **PRDX5** | 0 | 0.860654212830875 | 1 | 0.998 | 0 |
| **RASGRP2** | 0 | 1.93548793365641 | 0.981 | 0.57 | 0 |
| **TM7SF2** | 0 | -1.22402435948157 | 0.847 | 0.991 | 0 |
| **YIF1A** | 0 | 0.826589234080314 | 1 | 0.994 | 0 |
| **CTSF** | 0 | 0.943568744801093 | 0.559 | 0.003 | 0 |
| **NDUFV1** | 0 | 1.59512848856963 | 1 | 1 | 0 |
| **NUDT8** | 0 | 1.34568890780748 | 0.748 | 0.103 | 0 |
| **TESMIN** | 0 | 1.18125050125944 | 0.907 | 0.485 | 0 |
| **CCND1** | 0 | -2.08517564904939 | 0.952 | 0.997 | 0 |
| **FGF4** | 0 | 2.31756480944315 | 0.997 | 0.652 | 0 |
| **CTTN** | 0 | 1.02250003177469 | 1 | 0.999 | 0 |
| **ANAPC15** | 0 | 0.91911582186785 | 1 | 0.994 | 0 |
| **ARAP1** | 0 | 1.20258859494124 | 0.999 | 0.972 | 0 |
| **RAB6A** | 0 | 1.76240784320505 | 1 | 0.989 | 0 |
| **UCP2** | 0 | 1.41323681750853 | 0.996 | 0.912 | 0 |
| **THAP12** | 0 | 1.24559502706125 | 0.899 | 0.421 | 0 |
| **NDUFC2** | 0 | 1.12651240590029 | 1 | 1 | 0 |
| **CCDC90B** | 0 | 1.07844103238459 | 0.948 | 0.651 | 0 |
| **TMEM126B** | 0 | 1.1040200930146 | 0.829 | 0.318 | 0 |
| **CTSC** | 0 | 1.46161617573108 | 1 | 0.987 | 0 |
| **CHORDC1** | 0 | 1.25063565648786 | 0.828 | 0.263 | 0 |
| **TAF1D** | 0 | 1.21901233434715 | 1 | 0.98 | 0 |
| **AMOTL1** | 0 | -1.99827255536232 | 0.362 | 0.91 | 0 |
| **SESN3** | 0 | -1.51842274035771 | 0.929 | 0.996 | 0 |
| **ARHGAP42** | 0 | 1.62144284404791 | 0.986 | 0.636 | 0 |

| | | | | | |
|---|---|---|---|---|---|
| **TRPC6** | 0 | -2.3036659155831 | 0.336 | 0.87 | 0 |
| **DCUN1D5** | 0 | 1.75547509085619 | 1 | 0.922 | 0 |
| **AASDHPPT** | 0 | 1.16158540942454 | 1 | 0.986 | 0 |
| **SLC35F2** | 0 | 0.966151681521787 | 0.997 | 0.906 | 0 |
| **EXPH5** | 0 | 2.03701352544359 | 0.933 | 0.195 | 0 |
| **PPP2R1B** | 0 | -1.25560048600105 | 0.919 | 0.997 | 0 |
| **DLAT** | 0 | 1.14050688877242 | 1 | 0.994 | 0 |
| **USP28** | 0 | 1.8903455416821 | 1 | 0.98 | 0 |
| **TAGLN** | 0 | -4.62812149270881 | 0.511 | 0.954 | 0 |
| **FXYD6** | 0 | 1.79561388853543 | 0.996 | 0.777 | 0 |
| **TMPRSS4** | 0 | 1.19405278807675 | 0.551 | 0.009 | 0 |
| **FOXR1** | 0 | 1.44815775858361 | 0.72 | 0.005 | 0 |
| **SLC37A4** | 0 | 1.21252172268355 | 1 | 0.981 | 0 |
| **HYOU1** | 0 | 1.67204179621503 | 1 | 1 | 0 |
| **ARHGEF12** | 0 | 0.910332349094259 | 0.993 | 0.898 | 0 |
| **UBASH3B** | 0 | 1.73228637892044 | 0.876 | 0.243 | 0 |
| **HYLS1** | 0 | 1.61862667365617 | 0.748 | 0.149 | 0 |
| **FAM118B** | 0 | -1.02889212428855 | 0.711 | 0.962 | 0 |
| **SRPRA** | 0 | 0.85160913241054 | 1 | 0.993 | 0 |
| **ZBTB44** | 0 | 0.907882489745311 | 1 | 0.994 | 0 |
| **SNX19** | 0 | 1.53114002699141 | 0.803 | 0.047 | 0 |
| **TEAD4** | 0 | 1.5623253649146 | 1 | 0.965 | 0 |
| **CCND2** | 0 | -2.35872551718224 | 0.143 | 0.703 | 0 |
| **TIGAR** | 0 | 0.992178119093588 | 1 | 0.986 | 0 |
| **GDF3** | 0 | 2.02288458425922 | 0.999 | 0.804 | 0 |
| **DPPA3** | 0 | 3.26340353538519 | 0.998 | 0.309 | 0 |
| **CLEC4D** | 0 | -2.01862987978455 | 0.122 | 0.692 | 0 |
| **DUSP16** | 0 | 1.61737568423851 | 0.998 | 0.87 | 0 |
| **CREBL2** | 0 | 1.27831417808152 | 0.997 | 0.894 | 0 |
| **PLBD1** | 0 | 1.49624273900378 | 1 | 0.917 | 0 |
| **PLEKHA5** | 0 | -1.29875887926095 | 0.523 | 0.929 | 0 |
| **ETNK1** | 0 | -1.33669966246129 | 0.676 | 0.98 | 0 |
| **PPFIBP1** | 0 | -1.37313100860088 | 0.197 | 0.781 | 0 |
| **YAF2** | 0 | -1.47449899584409 | 0.101 | 0.757 | 0 |

| | | | | | |
|---|---|---|---|---|---|
| **TWF1** | 0 | 1.19818420443255 | 0.868 | 0.353 | 0 |
| **SLC38A2** | 0 | -1.25907813955138 | 0.918 | 0.998 | 0 |
| **PFKM** | 0 | -1.53735441007824 | 0.654 | 0.98 | 0 |
| **ADCY6** | 0 | -1.11551604512895 | 0.035 | 0.549 | 0 |
| **FKBP11** | 0 | 2.1878502786883 | 0.989 | 0.531 | 0 |
| **TUBA1C** | 0 | 1.46883007966895 | 1 | 0.997 | 0 |
| **LIMA1** | 0 | 1.62044952140121 | 1 | 0.848 | 0 |
| **FIGNL2** | 0 | -2.03136278690374 | 0.086 | 0.743 | 0 |
| **KRT8** | 0 | -1.88151537678208 | 0.997 | 1 | 0 |
| **CBX5** | 0 | -1.69753338782968 | 0.819 | 0.992 | 0 |
| **PPP1R1A** | 0 | -1.76370038281804 | 0.339 | 0.883 | 0 |
| **ESYT1** | 0 | -1.53492205954663 | 0.939 | 0.999 | 0 |
| **MYL6** | 0 | -1.46745624361191 | 1 | 1 | 0 |
| **HMGA2** | 0 | -2.3500535044242 | 0.624 | 0.993 | 0 |
| **LLPH** | 0 | 1.05149864967104 | 0.86 | 0.402 | 0 |
| **YEATS4** | 0 | 1.08788049318571 | 0.989 | 0.841 | 0 |
| **MYRFL** | 0 | -1.18665531185228 | 0.013 | 0.538 | 0 |
| **ATXN7L3B** | 0 | -1.29557006068761 | 0.936 | 0.999 | 0 |
| **PHLDA1** | 0 | -2.95353247373409 | 0.303 | 0.923 | 0 |
| **UBE2N** | 0 | 1.3008653607194 | 0.922 | 0.421 | 0 |
| **NDUFA12** | 0 | 1.53746717989846 | 1 | 0.94 | 0 |
| **NTN4** | 0 | 1.1146645020657 | 0.654 | 0.066 | 0 |
| **LTA4H** | 0 | -0.904612338331221 | 0.995 | 0.999 | 0 |
| **TMPO** | 0 | 0.932166071412913 | 1 | 0.99 | 0 |
| **ACTR6** | 0 | 1.66501874211719 | 0.969 | 0.468 | 0 |
| **SPIC** | 0 | 3.31992720045396 | 0.988 | 0.242 | 0 |
| **CHPT1** | 0 | 1.05110620965249 | 0.999 | 0.944 | 0 |
| **SYCP3** | 0 | 2.66772934163659 | 0.94 | 0.1 | 0 |
| **HSP90B1** | 0 | 2.74134234101528 | 0.999 | 0.772 | 0 |
| **CHST11** | 0 | -1.7819940998333 | 0.69 | 0.963 | 0 |
| **ALDH1L2** | 0 | 1.22971688305496 | 0.726 | 0.139 | 0 |
| **NUAK1** | 0 | -1.41132428911681 | 0.28 | 0.838 | 0 |
| **CKAP4** | 0 | -1.15793736727365 | 0.98 | 1 | 0 |
| **PPTC7** | 0 | 1.11097668983856 | 0.978 | 0.809 | 0 |

| | | | | | |
|---|---|---|---|---|---|
| **ALDH2** | 0 | 1.14619535233411 | 0.999 | 0.942 | 0 |
| **OAS1** | 0 | 1.38595052752586 | 0.683 | 0.023 | 0 |
| **SLC8B1** | 0 | 1.4989617114092 | 0.966 | 0.57 | 0 |
| **HRK** | 0 | 1.39269376733436 | 0.969 | 0.63 | 0 |
| **VSIG10** | 0 | 1.3074747349736 | 1 | 0.932 | 0 |
| **TAOK3** | 0 | 1.36608536846304 | 0.871 | 0.337 | 0 |
| **HSPB8** | 0 | -1.41688970041276 | 0.033 | 0.572 | 0 |
| **MSI1** | 0 | -1.91439036955483 | 0.092 | 0.846 | 0 |
| **CLIP1** | 0 | 1.03882283510264 | 0.997 | 0.932 | 0 |
| **DENR** | 0 | 0.929144848521445 | 1 | 0.975 | 0 |
| **MICU2** | 0 | 0.883987376292307 | 0.997 | 0.953 | 0 |
| **SACS** | 0 | -1.39766394444933 | 0.848 | 0.991 | 0 |
| **ATP12A** | 0 | 1.79559835514112 | 0.983 | 0.504 | 0 |
| **FLT1** | 0 | -1.74280873860209 | 0.065 | 0.599 | 0 |
| **NBEA** | 0 | -1.85164314782017 | 0.102 | 0.804 | 0 |
| **DCLK1** | 0 | -2.64806949992338 | 0.24 | 0.851 | 0 |
| **UFM1** | 0 | 1.77344232995523 | 1 | 0.948 | 0 |
| **LHFPL6** | 0 | -1.83681835052041 | 0.099 | 0.769 | 0 |
| **FOXO1** | 0 | -2.55588859376361 | 0.534 | 0.982 | 0 |
| **RGCC** | 0 | 2.85519996648765 | 0.973 | 0.236 | 0 |
| **EPSTI1** | 0 | 1.27505770305206 | 0.707 | 0.117 | 0 |
| **DNAJC15** | 0 | 3.23092851123792 | 1 | 0.992 | 0 |
| **TSC22D1** | 0 | 1.21484724278 | 0.967 | 0.65 | 0 |
| **LCP1** | 0 | 3.91538879884503 | 1 | 0.645 | 0 |
| **RCBTB1** | 0 | 1.95054064331709 | 1 | 0.954 | 0 |
| **CNMD** | 0 | 2.50754569383057 | 0.998 | 0.698 | 0 |
| **MZT1** | 0 | 1.15478569188124 | 0.992 | 0.84 | 0 |
| **KLF5** | 0 | 2.69251940129116 | 1 | 0.799 | 0 |
| **KLF12** | 0 | -1.19837652363797 | 0.235 | 0.762 | 0 |
| **EDNRB** | 0 | -1.37576962267286 | 0.015 | 0.563 | 0 |
| **SPRY2** | 0 | -2.26851307841255 | 0.79 | 0.992 | 0 |
| **GPC6** | 0 | -2.43796632441158 | 0.242 | 0.964 | 0 |
| **DNAJC3** | 0 | 1.26013224819796 | 1 | 0.994 | 0 |
| **DOCK9** | 0 | 1.65957083290303 | 0.999 | 0.779 | 0 |

| | | | | | |
|---|---|---|---|---|---|
| **UBAC2** | 0 | 0.836843577736088 | 1 | 0.987 | 0 |
| **ZIC5** | 0 | -2.46665250630413 | 0.035 | 0.592 | 0 |
| **ZIC2** | 0 | -2.57561064318031 | 0.029 | 0.689 | 0 |
| **PCCA** | 0 | -1.37196021455111 | 0.95 | 0.998 | 0 |
| **EFNB2** | 0 | -2.49289216418482 | 0.346 | 0.941 | 0 |
| **COL4A1** | 0 | -2.33999908985085 | 0.75 | 0.992 | 0 |
| **COL4A2** | 0 | -2.37003161534995 | 0.803 | 0.996 | 0 |
| **LAMP1** | 0 | -1.03404931282226 | 1 | 1 | 0 |
| **GRTP1** | 0 | 1.23228034813041 | 0.901 | 0.403 | 0 |
| **ARHGEF40** | 0 | -1.69690888679436 | 0.106 | 0.72 | 0 |
| **ZNF219** | 0 | -2.14180784412165 | 0.468 | 0.96 | 0 |
| **SALL2** | 0 | -2.63361508087944 | 0.643 | 0.998 | 0 |
| **OXA1L** | 0 | 0.958586082578078 | 1 | 1 | 0 |
| **SLC7A7** | 0 | 2.63621543171377 | 0.996 | 0.477 | 0 |
| **AJUBA** | 0 | -1.89538146126267 | 0.596 | 0.959 | 0 |
| **PSME1** | 0 | -1.25268837063227 | 0.705 | 0.965 | 0 |
| **REC8** | 0 | 2.99765322840068 | 0.99 | 0.337 | 0 |
| **NOP9** | 0 | 1.01119245489764 | 0.957 | 0.69 | 0 |
| **EGLN3** | 0 | 1.65293927527579 | 0.999 | 0.785 | 0 |
| **SPTSSA** | 0 | 1.38020338756006 | 1 | 1 | 0 |
| **ARF6** | 0 | 0.97386070535212 | 1 | 0.992 | 0 |
| **PYGL** | 0 | 1.89693618156036 | 1 | 0.928 | 0 |
| **FRMD6** | 0 | 1.49939360765412 | 0.965 | 0.705 | 0 |
| **GNPNAT1** | 0 | 1.94519368098153 | 1 | 0.884 | 0 |
| **BMP4** | 0 | -1.85338817045537 | 0.313 | 0.854 | 0 |
| **GMFB** | 0 | 1.13168851985222 | 0.732 | 0.1 | 0 |
| **GCH1** | 0 | 1.86917756417845 | 0.982 | 0.52 | 0 |
| **WDHD1** | 0 | 0.89496207901793 | 1 | 0.99 | 0 |
| **MAPK1IP1L** | 0 | 0.807047966895135 | 1 | 0.999 | 0 |
| **LGALS3** | 0 | 1.49115463328775 | 0.715 | 0.158 | 0 |
| **TBPL2** | 0 | 1.10489105397853 | 0.604 | 0.008 | 0 |
| **TMEM30B** | 0 | 1.33747299632953 | 0.69 | 0.022 | 0 |
| **HIF1A** | 0 | -0.983050696169613 | 0.942 | 0.999 | 0 |
| **HSPA2** | 0 | 2.61612861500611 | 0.961 | 0.164 | 0 |

| | | | | | |
|---|---|---|---|---|---|
| **PLEKHG3** | 0 | -1.79033425134859 | 0.059 | 0.623 | 0 |
| **SPTB** | 0 | -1.52939988242131 | 0.134 | 0.687 | 0 |
| **GPX2** | 0 | 2.77212571096158 | 0.979 | 0.301 | 0 |
| **RAB15** | 0 | 1.73903954240413 | 1 | 0.998 | 0 |
| **TMEM229B** | 0 | 1.78789244099089 | 0.91 | 0.301 | 0 |
| **ZFP36L1** | 0 | -1.15595869963607 | 0.98 | 0.999 | 0 |
| **ACTN1** | 0 | -1.85836933778074 | 0.994 | 1 | 0 |
| **DCAF5** | 0 | -1.24102962753095 | 0.584 | 0.949 | 0 |
| **EXD2** | 0 | -1.16080937805986 | 0.433 | 0.874 | 0 |
| **DCAF4** | 0 | 1.66379577287647 | 1 | 0.912 | 0 |
| **ACOT4** | 0 | 1.73825788946965 | 0.821 | 0.014 | 0 |
| **VRTN** | 0 | -3.51249245492095 | 0.202 | 0.961 | 0 |
| **NPC2** | 0 | -1.35709848923778 | 1 | 1 | 0 |
| **TGFB3** | 0 | 1.18107124992848 | 0.705 | 0.131 | 0 |
| **SLIRP** | 0 | 1.48001736802511 | 1 | 0.999 | 0 |
| **SNW1** | 0 | 1.59915690954359 | 0.993 | 0.763 | 0 |
| **CCDC85C** | 0 | -1.58209635411387 | 0.981 | 0.997 | 0 |
| **WARS** | 0 | 1.81737286966749 | 1 | 0.998 | 0 |
| **CKB** | 0 | -1.08635388691155 | 1 | 1 | 0 |
| **ATP5MPL** | 0 | 1.2125657069728 | 0.964 | 0.663 | 0 |
| **KIF26A** | 0 | -1.35956415569296 | 0.831 | 0.966 | 0 |
| **MTA1** | 0 | -1.19027980719544 | 0.981 | 1 | 0 |
| **CRIP1** | 0 | 4.2160141622278 | 1 | 0.727 | 0 |
| **NIPA2** | 0 | 1.0351326698352 | 0.998 | 0.96 | 0 |
| **TJP1** | 0 | -0.870351880213787 | 1 | 1 | 0 |
| **LPCAT4** | 0 | 1.37608031441959 | 0.988 | 0.813 | 0 |
| **DPH6** | 0 | 1.06334385408336 | 0.965 | 0.722 | 0 |
| **SPRED1** | 0 | -1.37037434856493 | 0.082 | 0.755 | 0 |
| **EIF2AK4** | 0 | 0.977422760947499 | 0.959 | 0.719 | 0 |
| **SPINT1** | 0 | 0.985843202310133 | 1 | 0.853 | 0 |
| **TMEM87A** | 0 | 0.999745152721704 | 0.948 | 0.638 | 0 |
| **SNAP23** | 0 | 0.884980041366409 | 1 | 0.995 | 0 |
| **PDIA3** | 0 | 1.45295355636263 | 1 | 0.989 | 0 |
| **SORD** | 0 | -1.30059081181927 | 0.395 | 0.876 | 0 |

| | | | | | |
|---|---|---|---|---|---|
| **MYEF2** | 0 | -2.19391416380478 | 0.329 | 0.968 | 0 |
| **EID1** | 0 | -2.05423953534612 | 0.921 | 1 | 0 |
| **SPPL2A** | 0 | 0.884207356341815 | 0.997 | 0.931 | 0 |
| **SCG3** | 0 | -1.24720799514099 | 0.162 | 0.684 | 0 |
| **MYO5A** | 0 | 1.41807199582435 | 0.997 | 0.846 | 0 |
| **RSL24D1** | 0 | 1.14007708234238 | 0.724 | 0.103 | 0 |
| **CCPG1** | 0 | 1.75251157472172 | 0.984 | 0.776 | 0 |
| **GTF2A2** | 0 | 1.51366928095989 | 1 | 0.918 | 0 |
| **ANXA2** | 0 | 0.855937005984005 | 1 | 1 | 0 |
| **TPM1** | 0 | -1.19609855382934 | 1 | 1 | 0 |
| **SNX1** | 0 | -1.21553771072075 | 0.763 | 0.979 | 0 |
| **PPIB** | 0 | 1.35807048684591 | 1 | 0.996 | 0 |
| **PCLAF** | 0 | -2.30988212633507 | 0.038 | 0.81 | 0 |
| **OAZ2** | 0 | -2.28834836697249 | 0.888 | 0.999 | 0 |
| **CLPX** | 0 | 1.11482396718915 | 1 | 0.987 | 0 |
| **IGDCC3** | 0 | -1.98472208520905 | 0.195 | 0.889 | 0 |
| **IGDCC4** | 0 | -1.34772772942642 | 0.029 | 0.664 | 0 |
| **SMAD3** | 0 | -1.66984345332687 | 0.49 | 0.953 | 0 |
| **C15orf61** | 0 | 1.1412424326891 | 0.881 | 0.472 | 0 |
| **UACA** | 0 | 2.42291949006343 | 1 | 0.895 | 0 |
| **MYO9A** | 0 | 1.06731408647464 | 0.918 | 0.557 | 0 |
| **CD276** | 0 | -1.19310921602515 | 0.98 | 0.999 | 0 |
| **ARID3B** | 0 | -0.952379248520453 | 0.999 | 1 | 0 |
| **ETFA** | 0 | -1.22618536624704 | 0.74 | 0.971 | 0 |
| **PEAK1** | 0 | -1.68317702318579 | 0.184 | 0.855 | 0 |
| **DNAJA4** | 0 | 1.63531955598895 | 0.814 | 0.042 | 0 |
| **CRABP1** | 0 | -3.38819268618017 | 0.332 | 0.815 | 0 |
| **TLNRD1** | 0 | -1.56116276905557 | 0.523 | 0.946 | 0 |
| **MEX3B** | 0 | -1.28748324759911 | 0.054 | 0.588 | 0 |
| **RAMAC** | 0 | 1.27941309526622 | 0.96 | 0.605 | 0 |
| **SEC11A** | 0 | -1.10662772530671 | 0.999 | 1 | 0 |
| **MFGE8** | 0 | -2.15253425112322 | 0.3 | 0.945 | 0 |
| **FES** | 0 | 1.06365211768309 | 0.832 | 0.325 | 0 |
| **FAM174B** | 0 | -1.57652671383425 | 0.257 | 0.793 | 0 |

| | | | | | |
|---|---|---|---|---|---|
| **RGMA** | 0 | -1.76685490964636 | 0.297 | 0.779 | 0 |
| **IGF1R** | 0 | -2.0010029243219 | 0.85 | 0.999 | 0 |
| **CHSY1** | 0 | -1.78730568695415 | 0.876 | 0.998 | 0 |
| **NME4** | 0 | -1.38292524385246 | 0.993 | 1 | 0 |
| **CACNA1H** | 0 | 1.41196310861257 | 0.749 | 0.057 | 0 |
| **SYNGR3** | 0 | 1.53238734810505 | 0.992 | 0.716 | 0 |
| **SLC9A3R2** | 0 | 2.04468354223563 | 0.995 | 0.641 | 0 |
| **PGP** | 0 | 0.88682751294347 | 1 | 0.997 | 0 |
| **ATP6V0C** | 0 | -0.806975409620133 | 0.999 | 0.999 | 0 |
| **CLDN6** | 0 | -0.860467723264764 | 1 | 1 | 0 |
| **IL32** | 0 | 1.78704141647792 | 0.859 | 0.207 | 0 |
| **NMRAL1** | 0 | 1.65101370508306 | 0.998 | 0.75 | 0 |
| **HMOX2** | 0 | 1.08747057240711 | 1 | 0.959 | 0 |
| **CDIP1** | 0 | -1.25580485482335 | 0.724 | 0.978 | 0 |
| **LITAF** | 0 | -2.49537430990952 | 0.962 | 0.999 | 0 |
| **SNN** | 0 | -1.26702983024272 | 0.572 | 0.942 | 0 |
| **RSL1D1** | 0 | 0.916145756630804 | 1 | 0.951 | 0 |
| **GSPT1** | 0 | 1.1347979338084 | 0.998 | 0.898 | 0 |
| **SMG1** | 0 | 1.03878388771303 | 0.93 | 0.593 | 0 |
| **NDUFAB1** | 0 | 1.17327596786192 | 1 | 1 | 0 |
| **QPRT** | 0 | -1.09525047454906 | 0.997 | 1 | 0 |
| **SEPHS2** | 0 | 1.76830216805785 | 1 | 0.984 | 0 |
| **GPT2** | 0 | 1.55180440541765 | 0.999 | 0.935 | 0 |
| **DNAJA2** | 0 | 1.02002312178538 | 1 | 1 | 0 |
| **ZNF423** | 0 | -2.38552061874825 | 0.459 | 0.981 | 0 |
| **SALL1** | 0 | -2.38314424813824 | 0.094 | 0.803 | 0 |
| **TOX3** | 0 | -1.39297652361058 | 0.047 | 0.668 | 0 |
| **AC007906.2** | 0 | -1.17687906029768 | 0.032 | 0.582 | 0 |
| **MMP2** | 0 | -3.35920628397684 | 0.282 | 0.959 | 0 |
| **MT1E** | 0 | 2.55840589497528 | 0.989 | 0.312 | 0 |
| **MT1F** | 0 | 1.99164982009936 | 0.998 | 0.878 | 0 |
| **MT1G** | 0 | 4.02069541622576 | 0.999 | 0.377 | 0 |
| **MT1H** | 0 | 4.94869844853777 | 1 | 0.433 | 0 |
| **MT1X** | 0 | 2.62280762291098 | 1 | 0.962 | 0 |

| | | | | | |
|---|---|---|---|---|---|
| **CCDC102A** | 0 | -1.31639355557997 | 0.323 | 0.855 | 0 |
| **USB1** | 0 | -1.01989790111205 | 0.859 | 0.988 | 0 |
| **MMP15** | 0 | -1.36180885022568 | 0.914 | 0.996 | 0 |
| **CMTM3** | 0 | -2.13969895311952 | 0.749 | 0.991 | 0 |
| **CMTM4** | 0 | 1.18390447436793 | 0.999 | 0.956 | 0 |
| **RRAD** | 0 | 2.93545011200175 | 0.964 | 0.212 | 0 |
| **CIAO2B** | 0 | 1.04329093775147 | 1 | 0.981 | 0 |
| **PLEKHG4** | 0 | 1.14063537461697 | 0.621 | 0.016 | 0 |
| **HAS3** | 0 | -1.20657692140314 | 0.035 | 0.581 | 0 |
| **ST3GAL2** | 0 | -1.05752406662862 | 0.891 | 0.991 | 0 |
| **BCAR1** | 0 | -1.42929824725215 | 0.76 | 0.977 | 0 |
| **VAT1L** | 0 | -3.19002795110406 | 0.04 | 0.708 | 0 |
| **WWOX** | 0 | -1.66048316682155 | 0.659 | 0.958 | 0 |
| **MAF** | 0 | -2.53019603363674 | 0.114 | 0.837 | 0 |
| **COX4I1** | 0 | 0.90068893457744 | 1 | 0.993 | 0 |
| **SLC7A5** | 0 | -1.80969440884224 | 0.972 | 0.999 | 0 |
| **DBNDD1** | 0 | -1.92887820588085 | 0.536 | 0.965 | 0 |
| **RFLNB** | 0 | 1.52171952686625 | 0.963 | 0.608 | 0 |
| **NXN** | 0 | -0.974642451630195 | 0.984 | 0.999 | 0 |
| **SPATA22** | 0 | 0.993776121207744 | 0.553 | 0.005 | 0 |
| **ASPA** | 0 | 1.21353666783523 | 0.57 | 0.03 | 0 |
| **SHPK** | 0 | 1.25244766220426 | 0.995 | 0.885 | 0 |
| **ZFP3** | 0 | 1.56447333626445 | 0.759 | 0.023 | 0 |
| **C1QBP** | 0 | 0.98268736604346 | 1 | 0.993 | 0 |
| **SAT2** | 0 | -1.22110446746333 | 0.849 | 0.986 | 0 |
| **ATP1B2** | 0 | -1.54019018337375 | 0.115 | 0.712 | 0 |
| **TP53** | 0 | -1.42505335848881 | 0.988 | 0.999 | 0 |
| **TRAPPC1** | 0 | -0.870233336918353 | 0.991 | 1 | 0 |
| **NTN1** | 0 | 6.91443829335772 | 1 | 0.985 | 0 |
| **GAS7** | 0 | 1.45464271739556 | 1 | 0.881 | 0 |
| **RAB34** | 0 | -0.990115433058631 | 0.955 | 0.997 | 0 |
| **FLOT2** | 0 | -1.37996945234569 | 0.931 | 0.998 | 0 |
| **PIPOX** | 0 | -1.28766771568289 | 0.106 | 0.646 | 0 |
| **UTP6** | 0 | 0.891769239151637 | 1 | 0.956 | 0 |

| | | | | | |
|---|---|---|---|---|---|
| **TMEM98** | 0 | -1.44451564183764 | 0.42 | 0.876 | 0 |
| **AP2B1** | 0 | 1.16320701353941 | 0.997 | 0.89 | 0 |
| **EIF1** | 0 | 1.68670167245708 | 1 | 0.895 | 0 |
| **FKBP10** | 0 | -2.10432353330043 | 0.563 | 0.961 | 0 |
| **ACLY** | 0 | -0.964608839241828 | 0.992 | 1 | 0 |
| **STAT3** | 0 | -1.23491671594351 | 0.978 | 0.999 | 0 |
| **GRN** | 0 | -1.17772070391374 | 1 | 1 | 0 |
| **FAM171A2** | 0 | -1.68723050555942 | 0.511 | 0.952 | 0 |
| **FZD2** | 0 | -2.52741743584489 | 0.06 | 0.782 | 0 |
| **ATP5MC1** | 0 | 0.834551896258555 | 0.999 | 0.982 | 0 |
| **IGF2BP1** | 0 | -2.39834815677645 | 0.906 | 1 | 0 |
| **LRRC59** | 0 | 1.04755086504461 | 1 | 1 | 0 |
| **SPAG9** | 0 | -1.00999576564083 | 0.886 | 0.994 | 0 |
| **TOM1L1** | 0 | -1.12331533743079 | 0.677 | 0.966 | 0 |
| **AKAP1** | 0 | 1.22484031694977 | 1 | 0.977 | 0 |
| **MSI2** | 0 | -1.15108106680097 | 0.657 | 0.946 | 0 |
| **RNF43** | 0 | -1.13435389077324 | 0.082 | 0.616 | 0 |
| **YPEL2** | 0 | 1.9042352218483 | 0.998 | 0.589 | 0 |
| **AXIN2** | 0 | -1.49552373627945 | 0.629 | 0.965 | 0 |
| **NOL11** | 0 | 1.38027843203339 | 0.93 | 0.416 | 0 |
| **JPT1** | 0 | 0.904356751106664 | 1 | 0.999 | 0 |
| **SYNGR2** | 0 | 0.968165614002582 | 1 | 0.992 | 0 |
| **SOCS3** | 0 | -1.66982899406659 | 0.508 | 0.95 | 0 |
| **TIMP2** | 0 | -1.83120142101468 | 0.705 | 0.977 | 0 |
| **LGALS3BP** | 0 | -2.43542426527226 | 0.327 | 0.965 | 0 |
| **RPTOR** | 0 | 1.03697100574404 | 0.982 | 0.843 | 0 |
| **MCRIP1** | 0 | -1.25402207684774 | 0.95 | 0.998 | 0 |
| **PYCR1** | 0 | 2.22546130790474 | 0.999 | 0.558 | 0 |
| **SLC16A3** | 0 | 2.09260429081398 | 0.926 | 0.481 | 0 |
| **TBCD** | 0 | 1.38398427757377 | 1 | 0.998 | 0 |
| **MYL12A** | 0 | 1.04463366951895 | 0.924 | 0.512 | 0 |
| **MTCL1** | 0 | -1.27887855585677 | 0.411 | 0.866 | 0 |
| **PPP4R1** | 0 | -1.04738452217456 | 0.975 | 1 | 0 |
| **PIEZO2** | 0 | -2.14546122807472 | 0.043 | 0.778 | 0 |

| | | | | | |
|---|---|---|---|---|---|
| **GNAL** | 0 | 1.70051865694861 | 0.957 | 0.504 | 0 |
| **IMPA2** | 0 | 1.36431285140115 | 1 | 0.951 | 0 |
| **CDH2** | 0 | -2.82520056486525 | 0.107 | 0.711 | 0 |
| **RNF125** | 0 | 2.4183018200628 | 1 | 0.773 | 0 |
| **MAPRE2** | 0 | -1.3645113920905 | 0.481 | 0.908 | 0 |
| **GALNT1** | 0 | -1.02990748926724 | 0.941 | 0.996 | 0 |
| **TPGS2** | 0 | -1.31522759575888 | 0.951 | 0.999 | 0 |
| **HAUS1** | 0 | 1.37435340043011 | 0.98 | 0.676 | 0 |
| **C18orf32** | 0 | 1.4270688089032 | 0.938 | 0.43 | 0 |
| **ACAA2** | 0 | 0.925662090424195 | 1 | 0.997 | 0 |
| **MAPK4** | 0 | -1.20684373834914 | 0.043 | 0.573 | 0 |
| **TCF4** | 0 | -1.88263270752843 | 0.257 | 0.897 | 0 |
| **ATP8B1** | 0 | 1.13847545250713 | 0.614 | 0.03 | 0 |
| **NEDD4L** | 0 | -1.24093199626638 | 0.635 | 0.979 | 0 |
| **SALL3** | 0 | -2.06471898634915 | 0.561 | 0.93 | 0 |
| **SLC66A2** | 0 | 1.21908224503185 | 0.998 | 0.918 | 0 |
| **HSBP1L1** | 0 | 1.41324375816763 | 0.826 | 0.19 | 0 |
| **CFD** | 0 | 1.39819496912328 | 0.854 | 0.274 | 0 |
| **CNN2** | 0 | -2.18646297728444 | 0.996 | 1 | 0 |
| **REEP6** | 0 | 1.41854056834251 | 0.969 | 0.627 | 0 |
| **TCF3** | 0 | -1.20817805661633 | 0.948 | 0.999 | 0 |
| **ZNF57** | 0 | 1.48510974436695 | 0.766 | 0.032 | 0 |
| **PTPRS** | 0 | -1.7985283455108 | 0.932 | 0.999 | 0 |
| **TINCR** | 0 | 1.09104316360228 | 0.739 | 0.158 | 0 |
| **TNFSF9** | 0 | 2.4352595615143 | 0.978 | 0.246 | 0 |
| **CD70** | 0 | 2.96199953726962 | 0.974 | 0.294 | 0 |
| **INSR** | 0 | 1.06531176219377 | 0.992 | 0.85 | 0 |
| **PRR36** | 0 | -1.88501053002193 | 0.15 | 0.842 | 0 |
| **CD320** | 0 | 1.74085657778545 | 1 | 0.991 | 0 |
| **ZNF560** | 0 | 0.913818023713478 | 0.554 | 0.006 | 0 |
| **PDE4A** | 0 | 1.35337088350432 | 0.888 | 0.395 | 0 |
| **AP1M2** | 0 | 1.8304040102242 | 0.966 | 0.305 | 0 |
| **ACP5** | 0 | 1.39541929953747 | 0.716 | 0.023 | 0 |
| **ZNF878** | 0 | 1.30044437974307 | 0.802 | 0.252 | 0 |

| | | | | | |
|---|---|---|---|---|---|
| **ZNF844** | 0 | 1.32604114082186 | 0.712 | 0.009 | 0 |
| **TNPO2** | 0 | -0.934778201976766 | 0.951 | 0.997 | 0 |
| **PRKACA** | 0 | -0.91834332354479 | 0.988 | 1 | 0 |
| **NOTCH3** | 0 | -2.29010751003965 | 0.735 | 0.996 | 0 |
| **BST2** | 0 | -3.42109104189372 | 0.216 | 0.934 | 0 |
| **SLC5A5** | 0 | 1.23112174468526 | 0.754 | 0.105 | 0 |
| **ARRDC2** | 0 | 1.180369995094 | 0.852 | 0.34 | 0 |
| **PIK3R2** | 0 | -1.15192300112111 | 0.996 | 1 | 0 |
| **JUND** | 0 | -1.42389177332046 | 1 | 1 | 0 |
| **FKBP8** | 0 | -0.944612143803151 | 0.93 | 0.995 | 0 |
| **DDX49** | 0 | 1.01960934748883 | 0.997 | 0.953 | 0 |
| **PBX4** | 0 | 1.32180585591636 | 0.813 | 0.19 | 0 |
| **ZNF253** | 0 | 1.16517009220565 | 0.657 | 0.015 | 0 |
| **ZNF90** | 0 | 1.3689974752209 | 0.984 | 0.734 | 0 |
| **ZNF486** | 0 | 2.05916903423814 | 0.986 | 0.469 | 0 |
| **ZNF626** | 0 | 0.934098152664241 | 0.576 | 0.023 | 0 |
| **ZNF257** | 0 | 1.36663041516476 | 0.746 | 0.003 | 0 |
| **ZNF676** | 0 | 3.12419214153517 | 0.995 | 0.233 | 0 |
| **ZNF98** | 0 | 1.67245269362816 | 0.846 | 0.01 | 0 |
| **ZNF492** | 0 | 2.04437252165292 | 0.912 | 0.052 | 0 |
| **ZNF728** | 0 | 1.18958838052574 | 0.681 | 0.009 | 0 |
| **CHST8** | 0 | 1.49026383536703 | 0.918 | 0.404 | 0 |
| **DMKN** | 0 | -2.48822856898684 | 0.2 | 0.976 | 0 |
| **ZNF146** | 0 | 0.859084526798995 | 1 | 0.999 | 0 |
| **ZNF568** | 0 | 2.22547027976173 | 0.973 | 0.187 | 0 |
| **PPP1R14A** | 0 | 1.31812328133767 | 0.993 | 0.778 | 0 |
| **YIF1B** | 0 | 1.12089805277682 | 1 | 0.999 | 0 |
| **C19orf33** | 0 | 2.59924416718628 | 0.988 | 0.355 | 0 |
| **KCNK6** | 0 | 1.14387433654783 | 0.749 | 0.162 | 0 |
| **MAP4K1** | 0 | 1.26454313433627 | 0.813 | 0.268 | 0 |
| **PLD3** | 0 | -1.73071449467022 | 0.99 | 1 | 0 |
| **AXL** | 0 | -2.17068918254993 | 0.031 | 0.706 | 0 |
| **HNRNPUL1** | 0 | -0.85418734474476 | 1 | 1 | 0 |
| **ATP1A3** | 0 | 1.48792799469667 | 0.86 | 0.228 | 0 |

| | | | | | |
|---|---|---|---|---|---|
| **MEGF8** | 0 | -1.45105209893331 | 0.298 | 0.872 | 0 |
| **PINLYP** | 0 | 1.08748352378242 | 0.646 | 0.026 | 0 |
| **BCL3** | 0 | 1.33181636536954 | 0.906 | 0.483 | 0 |
| **BCAM** | 0 | -2.72819057951698 | 0.372 | 0.965 | 0 |
| **VASP** | 0 | -1.28693637841525 | 0.99 | 0.999 | 0 |
| **ARHGAP35** | 0 | -1.40852241060058 | 0.775 | 0.991 | 0 |
| **TMEM160** | 0 | 1.34977373370858 | 1 | 0.998 | 0 |
| **CA11** | 0 | -1.66471300142069 | 0.076 | 0.687 | 0 |
| **RASIP1** | 0 | 1.65905351663562 | 0.837 | 0.16 | 0 |
| **PIH1D1** | 0 | 0.924465948529286 | 1 | 0.994 | 0 |
| **SCAF1** | 0 | -0.945711794810651 | 0.974 | 0.999 | 0 |
| **PTOV1** | 0 | -1.12684627164951 | 0.983 | 0.999 | 0 |
| **NR1H2** | 0 | 0.955285490380135 | 0.997 | 0.925 | 0 |
| **FAM71E1** | 0 | 1.18111699815543 | 0.784 | 0.222 | 0 |
| **KLK13** | 0 | 1.51615529982992 | 0.936 | 0.492 | 0 |
| **IGLON5** | 0 | -1.30988905851955 | 0.098 | 0.711 | 0 |
| **LIM2** | 0 | 1.86028652386061 | 0.911 | 0.11 | 0 |
| **ZNF649** | 0 | 1.0331359943217 | 0.989 | 0.904 | 0 |
| **ZNF880** | 0 | 1.1039036054772 | 0.758 | 0.165 | 0 |
| **ZNF600** | 0 | 1.225147668631 | 0.954 | 0.626 | 0 |
| **ZNF677** | 0 | 1.78038518448806 | 0.901 | 0.071 | 0 |
| **MBOAT7** | 0 | -0.925515577184434 | 0.968 | 0.999 | 0 |
| **NLRP7** | 0 | 4.95096457602402 | 1 | 0.684 | 0 |
| **NLRP2** | 0 | 2.34320732790405 | 1 | 0.995 | 0 |
| **TNNI3** | 0 | 1.5307659162489 | 0.901 | 0.252 | 0 |
| **PEG3** | 0 | 1.83399858219698 | 0.806 | 0.032 | 0 |
| **ZCCHC3** | 0 | -1.89366476156743 | 0.449 | 0.95 | 0 |
| **SOX12** | 0 | -2.07133630448814 | 0.405 | 0.94 | 0 |
| **TRIB3** | 0 | 2.01548272983895 | 0.997 | 0.665 | 0 |
| **PSMF1** | 0 | -0.907283764476239 | 1 | 1 | 0 |
| **SNRPB** | 0 | -1.00889502845265 | 1 | 1 | 0 |
| **IDH3B** | 0 | -0.874438752027915 | 1 | 1 | 0 |
| **PTPRA** | 0 | -0.984335211083187 | 0.855 | 0.983 | 0 |
| **CDC25B** | 0 | -1.42493610236928 | 0.946 | 0.99 | 0 |

| | | | | | |
|---|---|---|---|---|---|
| **CDS2** | 0 | -1.99138502594913 | 0.741 | 0.998 | 0 |
| **FERMT1** | 0 | 1.69936478116765 | 0.884 | 0.309 | 0 |
| **PLCB4** | 0 | -1.32514899130175 | 0.05 | 0.69 | 0 |
| **JAG1** | 0 | -1.89466882234357 | 0.835 | 0.995 | 0 |
| **BTBD3** | 0 | -1.86474098913214 | 0.651 | 0.978 | 0 |
| **SNRPB2** | 0 | 0.948796268045927 | 0.993 | 0.835 | 0 |
| **OVOL2** | 0 | 1.23647274018026 | 0.848 | 0.326 | 0 |
| **RBBP9** | 0 | 1.61074185114973 | 0.998 | 0.863 | 0 |
| **SEC23B** | 0 | 1.10559902857861 | 1 | 0.998 | 0 |
| **RIN2** | 0 | -1.26479945541683 | 0.713 | 0.959 | 0 |
| **RALGAPA2** | 0 | -1.3111854566575 | 0.643 | 0.963 | 0 |
| **XRN2** | 0 | -0.828543639213773 | 1 | 1 | 0 |
| **CST3** | 0 | -2.30711705303853 | 0.619 | 0.988 | 0 |
| **SYNDIG1** | 0 | -1.48848513232511 | 0.053 | 0.721 | 0 |
| **APMAP** | 0 | -1.42355789931185 | 0.947 | 0.999 | 0 |
| **PYGB** | 0 | 2.26933030118885 | 1 | 1 | 0 |
| **ASXL1** | 0 | -1.59472809319901 | 0.571 | 0.945 | 0 |
| **MAPRE1** | 0 | -0.913009675903898 | 1 | 1 | 0 |
| **CBFA2T2** | 0 | 1.10110454390917 | 1 | 0.972 | 0 |
| **DYNLRB1** | 0 | -1.27500632155098 | 1 | 1 | 0 |
| **MMP24OS** | 0 | -1.26454733428276 | 0.694 | 0.968 | 0 |
| **ERGIC3** | 0 | -1.1417522742608 | 1 | 1 | 0 |
| **CPNE1** | 0 | -1.23247313938963 | 0.988 | 1 | 0 |
| **SCAND1** | 0 | -0.895162315754976 | 0.981 | 0.999 | 0 |
| **DLGAP4** | 0 | -1.42997035201722 | 0.878 | 0.994 | 0 |
| **MYL9** | 0 | -3.78015543905338 | 0.761 | 0.999 | 0 |
| **TGIF2** | 0 | -2.19651397848454 | 0.666 | 0.995 | 0 |
| **SOGA1** | 0 | -1.08935733250599 | 0.852 | 0.991 | 0 |
| **SAMHD1** | 0 | 2.08474649057783 | 1 | 0.906 | 0 |
| **SRC** | 0 | -1.56814215590687 | 0.684 | 0.981 | 0 |
| **BLCAP** | 0 | -1.18410031931236 | 0.827 | 0.99 | 0 |
| **CTNNBL1** | 0 | -0.956027639055905 | 0.933 | 0.993 | 0 |
| **1,00 TOP** | 0 | -0.92600357329354 | 0.998 | 1 | 0 |
| **PLCG1** | 0 | -1.22338680113538 | 0.952 | 0.999 | 0 |

| | | | | | |
|---|---|---|---|---|---|
| **MYBL2** | 0 | 0.801668460687009 | 1 | 0.989 | 0 |
| **HNF4A** | 0 | 1.44284269294104 | 0.748 | 0.026 | 0 |
| **ADA** | 0 | 1.16556765611699 | 0.946 | 0.617 | 0 |
| **YWHAB** | 0 | 0.949414744618068 | 0.998 | 0.911 | 0 |
| **STK4** | 0 | -1.01085120608007 | 0.868 | 0.987 | 0 |
| **DBNDD2** | 0 | -1.54602862631005 | 0.262 | 0.862 | 0 |
| **PIGT** | 0 | -1.58547596007505 | 0.924 | 0.998 | 0 |
| **ZMYND8** | 0 | -0.986419866931015 | 0.997 | 0.999 | 0 |
| **NCOA3** | 0 | 1.12827521413276 | 0.997 | 0.969 | 0 |
| **SULF2** | 0 | -2.250746399724 | 0.454 | 0.97 | 0 |
| **STAU1** | 0 | -1.01746870710194 | 1 | 1 | 0 |
| **B4GALT5** | 0 | -1.10318085543131 | 0.977 | 0.999 | 0 |
| **PTPN1** | 0 | -0.856570541644721 | 0.975 | 1 | 0 |
| **BMP7** | 0 | -2.93165367826008 | 0.465 | 0.995 | 0 |
| **RAE1** | 0 | -0.85537914061171 | 0.976 | 0.999 | 0 |
| **VAPB** | 0 | -1.00638575932584 | 0.986 | 0.999 | 0 |
| **NELFCD** | 0 | -0.960625845323377 | 0.988 | 0.999 | 0 |
| **ATP5F1E** | 0 | -1.47989809421063 | 0.989 | 1 | 0 |
| **FAM217B** | 0 | -1.03920432932997 | 0.603 | 0.936 | 0 |
| **LSM14B** | 0 | -1.11963061564335 | 0.988 | 1 | 0 |
| **LAMA5** | 0 | -1.49239813079576 | 0.976 | 0.999 | 0 |
| **NKAIN4** | 0 | -1.80863400170509 | 0.404 | 0.869 | 0 |
| **KCNQ2** | 0 | 1.86747357677952 | 0.928 | 0.218 | 0 |
| **SLC2A4RG** | 0 | -1.25098242149018 | 0.745 | 0.97 | 0 |
| **DNAJC5** | 0 | -1.06821963281012 | 0.945 | 1 | 0 |
| **NRIP1** | 0 | -1.49234017795964 | 0.467 | 0.86 | 0 |
| **APP** | 0 | -1.45880854040292 | 1 | 1 | 0 |
| **HUNK** | 0 | -1.41766693698066 | 0.081 | 0.707 | 0 |
| **TTC3** | 0 | -1.31724330790068 | 0.906 | 0.992 | 0 |
| **PDXK** | 0 | 1.19366795809287 | 1 | 0.988 | 0 |
| **DNMT3L** | 0 | 3.41272030502195 | 1 | 0.884 | 0 |
| **COL18A1** | 0 | -3.90723831864574 | 0.651 | 1 | 0 |
| **SPATC1L** | 0 | -1.49541107414985 | 0.039 | 0.776 | 0 |
| **MED15** | 0 | 1.79929401083541 | 1 | 0.933 | 0 |

| | | | | | |
|---|---|---|---|---|---|
| **HIC2** | 0 | 1.29332634445201 | 1 | 0.929 | 0 |
| **SDF2L1** | 0 | 1.80489745644954 | 0.999 | 0.976 | 0 |
| **DERL3** | 0 | 2.22268578238083 | 0.881 | 0.08 | 0 |
| **MIF** | 0 | 0.883640029433732 | 1 | 1 | 0 |
| **DDT** | 0 | 0.953638636579327 | 1 | 0.99 | 0 |
| **SUSD2** | 0 | 2.52326297861303 | 1 | 0.798 | 0 |
| **TPST2** | 0 | 1.19526518484411 | 0.992 | 0.815 | 0 |
| **CCDC117** | 0 | 1.09039154478641 | 0.958 | 0.643 | 0 |
| **XBP1** | 0 | 1.34057596687013 | 1 | 0.997 | 0 |
| **NEFH** | 0 | 2.38694532355865 | 0.997 | 0.622 | 0 |
| **NF2** | 0 | 1.14750048774567 | 1 | 0.996 | 0 |
| **ASCC2** | 0 | 0.887124457812011 | 0.997 | 0.967 | 0 |
| **TCN2** | 0 | -2.08098697454921 | 0.357 | 0.963 | 0 |
| **TOMM22** | 0 | 1.34808074624178 | 1 | 0.99 | 0 |
| **SNU13** | 0 | 1.09744052478539 | 1 | 0.999 | 0 |
| **PHETA2** | 0 | 0.981717078455619 | 0.567 | 0.009 | 0 |
| **NDUFA6** | 0 | 1.30130260966214 | 1 | 0.97 | 0 |
| **A4GALT** | 0 | 2.56225109648967 | 1 | 0.756 | 0 |
| **TTLL12** | 0 | 0.849163533231598 | 1 | 0.993 | 0 |
| **PARVB** | 0 | 2.44951614303721 | 0.981 | 0.294 | 0 |
| **ARHGAP8** | 0 | 1.21543023496823 | 0.983 | 0.745 | 0 |
| **FBLN1** | 0 | -2.09451378556234 | 0.838 | 0.998 | 0 |
| **CRELD2** | 0 | 1.93442165422649 | 0.999 | 0.928 | 0 |
| **TRABD** | 0 | 0.935999493112817 | 0.998 | 0.936 | 0 |
| **PLXNB2** | 0 | 0.981143866069679 | 1 | 0.945 | 0 |
| **CSF2RA** | 0 | 2.10982262605009 | 0.937 | 0.234 | 0 |
| **DHRSX** | 0 | 2.34262128107275 | 0.999 | 0.834 | 0 |
| **CD99** | 0 | -2.70187973333366 | 0.2 | 0.966 | 0 |
| **TBL1X** | 0 | -1.62700766423905 | 0.243 | 0.855 | 0 |
| **WWC3** | 0 | -1.93886158984104 | 0.246 | 0.906 | 0 |
| **TMSB4X** | 0 | -2.90386387416288 | 0.998 | 1 | 0 |
| **AP1S2** | 0 | -1.60505057967 | 0.423 | 0.881 | 0 |
| **CTPS2** | 0 | -1.22782359854173 | 0.854 | 0.998 | 0 |
| **RBBP7** | 0 | -0.843845884069756 | 0.986 | 0.998 | 0 |

| | | | | | |
|---|---|---|---|---|---|
| **SH3KBP1** | 0 | -1.52416075104414 | 0.803 | 0.99 | 0 |
| **SMS** | 0 | -2.86384456135074 | 0.857 | 1 | 0 |
| **ACOT9** | 0 | -1.19381604943046 | 0.665 | 0.952 | 0 |
| **SAT1** | 0 | -1.75552594129457 | 0.861 | 0.994 | 0 |
| **EIF2S3** | 0 | 1.44731507699924 | 0.897 | 0.305 | 0 |
| **PCYT1B** | 0 | -1.75876938557339 | 0.054 | 0.814 | 0 |
| **MAGEB2** | 0 | 2.1337002651684 | 0.774 | 0.016 | 0 |
| **CASK** | 0 | -0.918621889546529 | 0.978 | 0.999 | 0 |
| **FUNDC1** | 0 | 1.0672771246342 | 0.972 | 0.749 | 0 |
| **USP11** | 0 | -1.00960677228691 | 0.961 | 0.998 | 0 |
| **TIMP1** | 0 | -1.79966318766523 | 0.949 | 0.999 | 0 |
| **GSPT2** | 0 | -1.32757828624707 | 0.712 | 0.975 | 0 |
| **MAGED1** | 0 | -2.04547448265832 | 0.96 | 1 | 0 |
| **WNK3** | 0 | -1.28118053021505 | 0.39 | 0.871 | 0 |
| **MAGED2** | 0 | -1.69491405029007 | 0.999 | 1 | 0 |
| **TRO** | 0 | -1.56083119998503 | 0.642 | 0.98 | 0 |
| **MAGEH1** | 0 | -1.63054819603948 | 0.344 | 0.925 | 0 |
| **UBQLN2** | 0 | -1.13559322114371 | 0.951 | 1 | 0 |
| **ARHGEF9** | 0 | -1.70640187362143 | 0.227 | 0.893 | 0 |
| **ZC4H2** | 0 | -1.18088771284083 | 0.282 | 0.808 | 0 |
| **MSN** | 0 | -0.871124567823986 | 0.992 | 1 | 0 |
| **HEPH** | 0 | -2.20692368060191 | 0.032 | 0.895 | 0 |
| **EFNB1** | 0 | -3.06012098031614 | 0.272 | 0.904 | 0 |
| **EDA** | 0 | -1.24418096866485 | 0.559 | 0.92 | 0 |
| **DLG3** | 0 | 1.37805332843046 | 1 | 0.987 | 0 |
| **SNX12** | 0 | -1.27115042453977 | 0.94 | 1 | 0 |
| **FOXO4** | 0 | -2.32799305449541 | 0.2 | 0.883 | 0 |
| **NONO** | 0 | 1.84243316799509 | 1 | 0.857 | 0 |
| **PHKA1** | 0 | 1.80307749391811 | 1 | 0.871 | 0 |
| **NAP1L2** | 0 | -1.28942321771597 | 0.201 | 0.778 | 0 |
| **SLC16A2** | 0 | -2.24043934229327 | 0.439 | 0.969 | 0 |
| **RLIM** | 0 | 1.44197085003128 | 0.826 | 0.134 | 0 |
| **COX7B** | 0 | 2.29951864976812 | 0.999 | 0.748 | 0 |
| **BRWD3** | 0 | 1.00935570049299 | 1 | 0.984 | 0 |

| | | | | | |
|---|---|---|---|---|---|
| **SH3BGRL** | 0 | -1.26511620950865 | 0.727 | 0.975 | 0 |
| **ZNF711** | 0 | -1.40014469522171 | 0.307 | 0.836 | 0 |
| **NAP1L3** | 0 | -2.0922307241214 | 0.278 | 0.907 | 0 |
| **TNMD** | 0 | -1.39157492818102 | 0.045 | 0.596 | 0 |
| **TSPAN6** | 0 | -1.23099022903793 | 0.978 | 0.997 | 0 |
| **TIMM8A** | 0 | 0.883392505085893 | 1 | 0.965 | 0 |
| **ARMCX1** | 0 | -1.27514588674694 | 0.84 | 0.991 | 0 |
| **ARMCX2** | 0 | -1.16125268224125 | 0.936 | 0.998 | 0 |
| **BEX4** | 0 | -1.64318461780263 | 0.9 | 1 | 0 |
| **TCEAL8** | 0 | -0.856947691352867 | 0.991 | 1 | 0 |
| **TCEAL9** | 0 | -1.83322211630143 | 0.732 | 0.994 | 0 |
| **BEX3** | 0 | -3.08772150312721 | 0.887 | 1 | 0 |
| **NRK** | 0 | -2.86200522727033 | 0.037 | 0.76 | 0 |
| **RADX** | 0 | 0.930139672319471 | 0.567 | 0.015 | 0 |
| **COL4A6** | 0 | -1.85344823351317 | 0.026 | 0.726 | 0 |
| **COL4A5** | 0 | -2.46880056847121 | 0.149 | 0.896 | 0 |
| **ACSL4** | 0 | -1.30030811762144 | 0.704 | 0.987 | 0 |
| **LUZP4** | 0 | 1.57276118122282 | 0.764 | 0.008 | 0 |
| **PGRMC1** | 0 | -1.49321753188236 | 0.999 | 1 | 0 |
| **SEPTIN6** | 0 | -2.01854692330644 | 0.211 | 0.938 | 0 |
| **MCTS1** | 0 | 0.984205375427988 | 1 | 0.987 | 0 |
| **XIAP** | 0 | 1.06214517787165 | 0.927 | 0.611 | 0 |
| **IGSF1** | 0 | 1.75332064947325 | 0.967 | 0.668 | 0 |
| **STK26** | 0 | -1.19168710785361 | 0.856 | 0.993 | 0 |
| **MBNL3** | 0 | -2.3321906396771 | 0.421 | 0.958 | 0 |
| **HS6ST2** | 0 | -2.05563846690309 | 0.093 | 0.864 | 0 |
| **GPC4** | 0 | -2.83111077824642 | 0.751 | 0.993 | 0 |
| **ZIC3** | 0 | -1.96573575973434 | 0.403 | 0.941 | 0 |
| **LDOC1** | 0 | -1.58210896206785 | 0.497 | 0.962 | 0 |
| **IDS** | 0 | -1.49109637885431 | 0.33 | 0.892 | 0 |
| **ZNF185** | 0 | -1.62915543399754 | 0.322 | 0.878 | 0 |
| **CCNQ** | 0 | 1.39395443328531 | 0.991 | 0.809 | 0 |
| **DUSP9** | 0 | 1.2863524868021 | 0.684 | 0.043 | 0 |
| **SLC6A8** | 0 | 1.58779238917035 | 1 | 0.979 | 0 |

| | | | | | |
|---|---|---|---|---|---|
| **IDH3G** | 0 | 0.819196479728905 | 1 | 0.994 | 0 |
| **IRAK1** | 0 | 1.6833854764761 | 0.977 | 0.511 | 0 |
| **FLNA** | 0 | -1.37487312325204 | 1 | 1 | 0 |
| **VAMP7** | 0 | 0.92583323364957 | 0.992 | 0.905 | 0 |
| **MT-ND1** | 0 | -1.64783786135859 | 0.65 | 0.956 | 0 |
| **MT-ND2** | 0 | -2.3926232422266 | 0.743 | 0.991 | 0 |
| **MT-CO2** | 0 | -2.15065376826165 | 0.962 | 0.999 | 0 |
| **MT-ATP6** | 0 | -1.95626608755391 | 0.949 | 0.999 | 0 |
| **MT-CO3** | 0 | -2.87175944393187 | 0.952 | 1 | 0 |
| **MT-ND4L** | 0 | -2.08971478091958 | 0.823 | 0.996 | 0 |
| **MT-ND4** | 0 | -2.10344594220203 | 0.954 | 0.999 | 0 |
| **MT-CYB** | 0 | -2.15115393609705 | 0.917 | 0.999 | 0 |
| **SNAPC3** | 5.8279 | 0.971636719362043 | 0.987 | 0.853 | 9.47796938 |
| **VPS41** | 5.8825 | -0.945973769495776 | 0.836 | 0.987 | 9.56674879 |
| **HIPK1** | 1.0335 | -0.95223479746059 | 0.795 | 0.983 | 1.68092114 |
| **PTPRG** | 1.1861 | -1.3070453569046 | 0.611 | 0.933 | 1.92906666 |
| **CERS4** | 1.3097 | -1.33814581630277 | 0.912 | 0.992 | 2.13000252 |
| **GAB1** | 1.9084 | 1.0993153197712 | 0.916 | 0.571 | 3.10363291 |
| **FCGBP** | 3.8134 | 1.30985717456948 | 0.602 | 0.054 | 6.20184801 |
| **ATP1A2** | 5.7129 | -1.32978602764437 | 0.016 | 0.504 | 9.29098882 |
| **ADGRG2** | 6.0507 | 1.24119612337349 | 0.869 | 0.411 | 9.84034582 |
| **CDYL** | 1.2949 | -1.04314565853742 | 0.734 | 0.972 | 2.10604202 |
| **NREP** | 1.7756 | -0.945938764319648 | 0.995 | 0.999 | 2.88777523 |
| **CAMK1D** | 2.9544 | 1.1440546438596 | 0.66 | 0.109 | 4.80483302 |
| **IL6R** | 5.2846 | 1.13907634398624 | 0.852 | 0.368 | 8.59448649 |
| **PAK1** | 5.7673 | 0.957309946835648 | 0.986 | 0.855 | 9.37938137 |
| **STARD10** | 9.4574 | 1.06843108137216 | 0.937 | 0.583 | 1.53807074 |
| **GYG2** | 1.0578 | -1.13567572392924 | 0.583 | 0.924 | 1.72044061 |
| **MTRES1** | 1.7582 | 1.06097510668104 | 0.783 | 0.259 | 2.85949471 |
| **MEIOB** | 2.1156 | 1.03673731362248 | 0.526 | 0.003 | 3.44072310 |
| **SLC30A5** | 3.1401 | 0.989318808499418 | 0.936 | 0.622 | 5.10680298 |
| **TNNT1** | 3.3452 | 1.06072709909833 | 1 | 0.885 | 5.44032129 |
| **ZSWIM9** | 4.0009 | -1.05147246867867 | 0.135 | 0.664 | 6.50682134 |
| **LIMD2** | 4.1315 | -1.17062986384237 | 0.439 | 0.865 | 6.71916489 |

| | | | | | |
|---|---|---|---|---|---|
| **PKIA** | 5.1462 | -0.902136859698119 | 0.041 | 0.544 | 8.3693966 |
| **SLC9A3** | 8.6172 | 1.09984276954021 | 0.601 | 0.052 | 1.4014244 |
| **GABRD** | 3.2405 | 0.970308708716275 | 0.545 | 0.016 | 5.2701385 |
| **EXTL2** | 7.9628 | -1.02455082196848 | 0.719 | 0.955 | 1.2949961 |
| **FMNL2** | 1.1985 | 0.822613734124183 | 0.999 | 0.988 | 1.9491796 |
| **PDCL** | 1.9108 | 0.814827659449286 | 0.999 | 0.988 | 3.1076659 |
| **PJA1** | 4.9606 | -1.0840331320605 | 0.828 | 0.985 | 8.0675008 |
| **OBI1** | 5.9389 | -1.10487481989554 | 0.695 | 0.957 | 9.6585160 |
| **LTBP1** | 7.6433 | -1.31834444916003 | 0.422 | 0.852 | 1.2430309 |
| **HK1** | 1.0863 | -0.9492362002205 | 0.965 | 0.999 | 1.7667120 |
| **SKA3** | 1.5796 | 1.24997861033864 | 0.974 | 0.837 | 2.5689865 |
| **VOPP1** | 1.8250 | -0.998023076859802 | 0.686 | 0.946 | 2.9681095 |
| **EPDR1** | 2.7479 | -1.02634406292969 | 0.013 | 0.491 | 4.4690666 |
| **NSMAF** | 6.7235 | 0.848851667960645 | 0.998 | 0.937 | 1.0934530 |
| **CAPZA1** | 1.2606 | 1.01926328616622 | 0.829 | 0.347 | 2.0502352 |
| **ZNF273** | 1.3824 | 0.947642899184203 | 0.595 | 0.055 | 2.2483006 |
| **ABCB7** | 1.9153 | -0.849535363824821 | 0.98 | 0.998 | 3.1148794 |
| **SSC4D** | 2.4944 | -0.971368985253564 | 0.073 | 0.59 | 4.0567571 |
| **EIF3E** | 7.7146 | 1.00279992085651 | 0.703 | 0.156 | 1.2546390 |
| **NRXN2** | 8.4629 | 0.965817383142321 | 0.619 | 0.068 | 1.3763368 |
| **PLPP2** | 5.0168 | 1.23677725593428 | 0.629 | 0.086 | 8.1588617 |
| **NRARP** | 1.0084 | -1.27480242536323 | 0.191 | 0.7 | 1.6400017 |
| **STARD8** | 1.3397 | -1.00649595505894 | 0.034 | 0.529 | 2.1788401 |
| **ADAM10** | 1.5232 | 0.899704093527628 | 0.992 | 0.861 | 2.4773174 |
| **RHOF** | 1.9837 | 1.04740459588986 | 0.892 | 0.449 | 3.2261717 |
| **ZNF667** | 4.1130 | 0.870321136334593 | 0.522 | 0.006 | 6.6890749 |
| **INKA2** | 5.7120 | -1.18115827977589 | 0.137 | 0.657 | 9.2894686 |
| **ACADSB** | 6.8704 | -1.06514191697549 | 0.155 | 0.676 | 1.1173481 |
| **HIP1** | 7.3332 | -1.08059399007421 | 0.609 | 0.928 | 1.1926141 |
| **CEBPZOS** | 9.3035 | 0.934431362556533 | 0.992 | 0.851 | 1.5130294 |
| **MAVS** | 1.2422 | -0.925812200798067 | 0.89 | 0.99 | 2.0202626 |
| **EIF1AX** | 1.9978 | -0.904130673765067 | 0.977 | 0.996 | 3.2490787 |
| **ARSE** | 2.7442 | -1.11226135487738 | 0.06 | 0.567 | 4.4629416 |
| **TSPAN18** | 1.6848 | -1.03954322770547 | 0.98 | 0.997 | 2.7400808 |

| | | | | | |
|---|---|---|---|---|---|
| **GALNT3** | 3.5057 | 0.984369861702204 | 0.518 | 0.005 | 5.7013426 |
| **PDE3B** | 3.9713 | -1.00330229203463 | 0.05 | 0.549 | 6.4585827 |
| **SLC35A5** | 4.2714 | 0.932987100124952 | 0.991 | 0.914 | 6.9467310 |
| **SEPTIN11** | 2.0205 | -0.929645007417588 | 0.996 | 1 | 3.2860793 |
| **PCDHA4** | 4.6734 | -0.975116301869339 | 0.095 | 0.609 | 7.6004749 |
| **FKBP3** | 1.7435 | 0.832372386679196 | 1 | 0.936 | 2.8354776 |
| **BCAS4** | 1.9175 | -1.05841222675571 | 0.641 | 0.947 | 3.1185831 |
| **DZIP1** | 2.0674 | -1.11074644701397 | 0.806 | 0.976 | 3.3622958 |
| **PDE12** | 2.4969 | 0.986072221981543 | 0.917 | 0.559 | 4.0608036 |
| **BHMT** | 3.0063 | 0.92620194677586 | 0.524 | 0.01 | 4.8892775 |
| **ZFAND3** | 1.9108 | -0.899587182686573 | 0.933 | 0.995 | 3.1076023 |
| **EPHB1** | 1.0540 | -0.983776812211826 | 0.023 | 0.502 | 1.7142263 |
| **FZD3** | 2.0004 | -1.11262448772776 | 0.857 | 0.987 | 3.2533268 |
| **ZDHHC23** | 3.1974 | 1.10798073017952 | 0.75 | 0.222 | 5.1999587 |
| **PGM2** | 4.7644 | 0.963930284301465 | 0.988 | 0.868 | 7.7484058 |
| **TMEM97** | 5.0855 | -0.901873851007539 | 0.998 | 0.999 | 8.2706679 |
| **PSMC4** | 5.2256 | 0.902560916044181 | 0.999 | 0.998 | 8.4985142 |
| **LRRN2** | 1.2966 | -1.07817918793066 | 0.057 | 0.554 | 2.1086871 |
| **PLTP** | 1.4076 | -0.992957841088514 | 0.956 | 0.998 | 2.2892059 |
| **OXCT1** | 2.3314 | -1.08445880980773 | 0.486 | 0.891 | 3.7917089 |
| **MFSD4B** | 2.6528 | 1.0110686743561 | 0.98 | 0.816 | 4.3143200 |
| **RB1** | 3.8169 | -1.06001038854233 | 0.305 | 0.781 | 6.2074906 |
| **ITPR3** | 4.6059 | 1.14621480882881 | 0.921 | 0.565 | 7.4906183 |
| **GDPD1** | 6.9409 | 1.09608991168877 | 0.705 | 0.169 | 1.1288091 |
| **AGPAT5** | 8.1022 | -1.06398383423315 | 0.824 | 0.973 | 1.3176648 |
| **RPP40** | 1.0503 | 0.974882426811548 | 0.957 | 0.728 | 1.7081258 |
| **CCDC59** | 1.8214 | 0.925199357965986 | 0.967 | 0.789 | 2.9621879 |
| **TFAM** | 2.5351 | 0.882825727644926 | 0.989 | 0.835 | 4.1229202 |
| **MIPOL1** | 2.6153 | -1.08897977029785 | 0.334 | 0.803 | 4.2533700 |
| **DDHD1** | 4.6861 | 1.01717879553558 | 0.975 | 0.822 | 7.6211539 |
| **UBXN7** | 5.1455 | -0.918414116155753 | 0.789 | 0.971 | 8.3682732 |
| **NCAM1** | 5.3836 | -1.19111251949963 | 0.027 | 0.505 | 8.7554165 |
| **SFXN1** | 6.0782 | -0.862382689949849 | 0.937 | 0.999 | 9.8850326 |
| **RBM7** | 6.7312 | 0.947368853140175 | 0.953 | 0.694 | 1.0947058 |

| | | | | | |
|---|---|---|---|---|---|
| **CCDC69** | 2.6074 | 1.06322300702666 | 0.866 | 0.436 | 4.24055160 |
| **STK17B** | 3.4394 | 1.07269649600115 | 0.861 | 0.417 | 5.59359708 |
| **NUPR1** | 5.1745 | 1.42070612798326 | 0.698 | 0.175 | 8.41536454 |
| **LSS** | 7.6245 | -0.946242471708737 | 0.967 | 0.997 | 1.23997349 |
| **LRATD2** | 7.9326 | -1.49535568577914 | 0.875 | 0.981 | 1.29008034 |
| **HBM** | 1.4264 | 1.03909203968014 | 0.535 | 0.02 | 2.31977560 |
| **KIAA0930** | 3.6151 | 0.899284550982172 | 0.987 | 0.852 | 5.87927080 |
| **LRRC56** | 3.7024 | 1.03847913616095 | 0.574 | 0.05 | 6.02133142 |
| **ERMP1** | 4.4613 | -1.11331441314348 | 0.438 | 0.859 | 7.25556663 |
| **BCAS2** | 4.8909 | 0.957562899025235 | 0.911 | 0.551 | 7.95413629 |
| **MORF4L2** | 7.5106 | 0.845189989255516 | 0.996 | 0.96 | 1.22144920 |
| **PLCB3** | 7.5370 | 0.869896584616254 | 0.998 | 0.919 | 1.22574634 |
| **MSLN** | 8.0163 | 1.02027372426693 | 0.523 | 0.014 | 1.30370404 |
| **SRSF8** | 4.9839 | 1.00192702916105 | 0.976 | 0.819 | 8.10547490 |
| **AMER1** | 5.0112 | -1.11293746321273 | 0.395 | 0.834 | 8.14972652 |
| **APOBEC3G** | 5.7540 | 1.10810951876518 | 0.662 | 0.126 | 9.35783689 |
| **TP53BP1** | 5.9291 | -1.05399827261727 | 0.657 | 0.94 | 9.64260620 |
| **TBRG1** | 9.2552 | 0.855998765187644 | 0.987 | 0.892 | 1.50517894 |
| **CXCL12** | 2.2447 | -1.77918812938256 | 0.925 | 0.927 | 3.65064015 |
| **APOC1** | 3.1447 | 0.987924583739544 | 0.955 | 0.715 | 5.11426993 |
| **PSKH1** | 3.2723 | -0.916275697945051 | 0.811 | 0.985 | 5.32186810 |
| **MARVELD2** | 4.3753 | 1.04455335693837 | 0.801 | 0.298 | 7.11558322 |
| **AJAP1** | 6.5242 | -1.04835406964675 | 0.057 | 0.546 | 1.06104362 |
| **SHLD2** | 1.3453 | 0.99682488551514 | 0.898 | 0.528 | 2.18788647 |
| **NSD2** | 5.7500 | -1.18593242210331 | 0.365 | 0.803 | 9.35123860 |
| **FIBCD1** | 6.3702 | -1.41755401496466 | 0.255 | 0.731 | 1.03599774 |
| **NTF3** | 7.3200 | -1.11583926570628 | 0.032 | 0.506 | 1.19046277 |
| **KLHL4** | 1.0439 | -0.956643500518806 | 0.021 | 0.492 | 1.69781840 |
| **CAAP1** | 1.0884 | 0.990219244792731 | 0.943 | 0.663 | 1.77011910 |
| **IGFBP2** | 3.7228 | -1.33520796938236 | 0.993 | 1 | 6.05442702 |
| **CHD6** | 7.0271 | -1.00423319517558 | 0.708 | 0.952 | 1.14282509 |
| **PPP2CB** | 7.9853 | -0.935811499188212 | 0.698 | 0.942 | 1.29865710 |
| **ZKSCAN1** | 8.3851 | -1.12717989291211 | 0.449 | 0.853 | 1.36367324 |
| **MAGEF1** | 1.4626 | -1.03881409616863 | 0.686 | 0.945 | 2.37866838 |

| | | | | | |
|---|---|---|---|---|---|
| **UBE2Q2** | 2.2067 | 0.806129929000738 | 0.998 | 0.961 | 3.5889023 |
| **SNX7** | 3.1206 | 1.03382138185639 | 0.873 | 0.47 | 5.0751461 |
| **SLC25A33** | 5.1533 | 1.00548076173482 | 0.962 | 0.76 | 8.3808401 |
| **SCAF4** | 1.0021 | -0.864338795082396 | 0.933 | 0.996 | 1.6297744 |
| **FBN3** | 1.7781 | -1.20683188516071 | 0.228 | 0.735 | 2.8917774 |
| **ARHGEF2** | 1.8516 | -1.1233150299548 | 0.63 | 0.928 | 3.0113630 |
| **MANEAL** | 4.4243 | 1.01923536537814 | 0.975 | 0.758 | 7.1953966 |
| **HIPK2** | 6.9009 | -1.26084463302651 | 0.505 | 0.871 | 1.1223063 |
| **RIC8B** | 1.9772 | -1.28323680952468 | 0.362 | 0.801 | 3.2156325 |
| **TAF4B** | 3.0055 | 1.18266572429663 | 0.845 | 0.43 | 4.8878892 |
| **PBX3** | 3.2068 | 0.94649000810645 | 0.997 | 0.952 | 5.2152347 |
| **SVBP** | 1.4631 | -0.953745803214858 | 0.869 | 0.985 | 2.3795326 |
| **PIEZO1** | 1.5301 | 0.897990712885111 | 0.546 | 0.033 | 2.4885315 |
| **C1orf198** | 1.7174 | -1.34764832481908 | 0.553 | 0.899 | 2.7931092 |
| **XPNPEP1** | 2.1958 | -0.964512035758087 | 0.749 | 0.961 | 3.5711150 |
| **TP53I3** | 3.3489 | -1.08788192094581 | 0.566 | 0.901 | 5.4464446 |
| **ZNF316** | 4.8397 | -0.984380788086304 | 0.209 | 0.712 | 7.8708151 |
| **ANKRD33B** | 8.2536 | -1.30620993723543 | 0.052 | 0.531 | 1.3422916 |
| **GALK1** | 1.1531 | 1.05396320507952 | 0.917 | 0.618 | 1.8754164 |
| **SEMA3A** | 3.2226 | -1.02478904848339 | 0.122 | 0.623 | 5.2409369 |
| **NOCT** | 4.0844 | 0.973549951827986 | 0.981 | 0.863 | 6.6424835 |
| **SLC7A6** | 4.4749 | 1.01590676718992 | 0.919 | 0.588 | 7.2776771 |
| **FN3K** | 9.7043 | 0.950443335765418 | 0.995 | 0.898 | 1.5782140 |
| **PEMT** | 2.9450 | 0.821984817029357 | 0.997 | 0.972 | 4.7894706 |
| **PRRX2** | 5.4639 | 1.03633649522618 | 0.843 | 0.358 | 8.8861007 |
| **FBRS** | 7.4497 | 1.07082478029769 | 0.962 | 0.739 | 1.2115459 |
| **CPS1** | 1.7789 | -0.930458596266432 | 0.755 | 0.98 | 2.8931845 |
| **LRAT** | 2.5609 | -1.56199875968854 | 0.064 | 0.545 | 4.1649366 |
| **GJC1** | 3.0890 | -1.63656771508177 | 0.615 | 0.891 | 5.0237370 |
| **ZNF680** | 5.3640 | 0.938485629283045 | 0.97 | 0.783 | 8.7235135 |
| **EML1** | 8.1036 | -1.03682283428248 | 0.869 | 0.979 | 1.3178949 |
| **TMEM45A** | 9.7251 | -1.10546952421914 | 0.17 | 0.683 | 1.5816029 |
| **SLC23A2** | 1.1533 | 0.911078435521425 | 0.993 | 0.912 | 1.8757634 |
| **ACACB** | 1.5200 | 1.12676513348205 | 0.768 | 0.278 | 2.4719877 |

| | | | | | |
|---|---|---|---|---|---|
| **DNAJA1** | 1.9011 | 0.889021666507456 | 0.991 | 0.89 | 3.0918451 |
| **WASF3** | 2.0878 | -1.16760577508917 | 0.779 | 0.963 | 3.3954879 |
| **EZR** | 3.5461 | -1.48547068230044 | 1 | 1 | 5.7670511 |
| **FAM53C** | 3.7361 | -0.803289751302301 | 0.946 | 0.997 | 6.0760228 |
| **AK5** | 4.0630 | 0.925132552649991 | 0.504 | 0.012 | 6.6077479 |
| **PITPNC1** | 4.7315 | 0.831018186671742 | 0.997 | 0.946 | 7.6948733 |
| **PTPN21** | 6.8355 | -1.10305813334896 | 0.227 | 0.715 | 1.1116636 |
| **SHMT2** | 7.6401 | 0.957716037341795 | 0.959 | 0.745 | 1.2425111 |
| **SDC1** | 8.9689 | -1.08214504972271 | 0.047 | 0.521 | 1.4586283 |
| **TULP2** | 1.1243 | 0.852720753038881 | 0.522 | 0.024 | 1.8285098 |
| **PPARG** | 1.1366 | 1.13022432666675 | 0.58 | 0.067 | 1.8485142 |
| **RET** | 2.8354 | -1.48542304351893 | 0.312 | 0.756 | 4.6112717 |
| **MMP24** | 3.5034 | -1.30879816777125 | 0.849 | 0.953 | 5.6977161 |
| **LPIN1** | 4.1916 | 1.00856441481142 | 0.888 | 0.514 | 6.8169227 |
| **ABCB6** | 4.4404 | 1.06690917804716 | 0.862 | 0.465 | 7.2214267 |
| **SSBP3** | 5.6383 | -0.974676659818178 | 0.717 | 0.95 | 9.1696532 |
| **SIAH2** | 8.4750 | -1.00276535577635 | 0.089 | 0.581 | 1.3783045 |
| **ITGA5** | 1.5484 | -1.40721695197224 | 0.067 | 0.545 | 2.5182574 |
| **RYBP** | 1.6326 | -0.913196698282221 | 0.964 | 0.992 | 2.6552227 |
| **PNKD** | 1.7168 | 0.810238465034972 | 0.992 | 0.921 | 2.7921274 |
| **MBOAT1** | 2.1579 | 1.02163033507822 | 0.911 | 0.508 | 3.5095003 |
| **TMEM126A** | 2.3605 | 0.846093116581493 | 0.986 | 0.892 | 3.8389501 |
| **FXN** | 2.9711 | 0.832482000807567 | 0.993 | 0.942 | 4.8319620 |
| **SEMA3B** | 3.0069 | -0.909143677415274 | 0.047 | 0.519 | 4.8901961 |
| **HSPA12B** | 3.4871 | 1.0233229301841 | 0.659 | 0.137 | 5.6712158 |
| **MX1** | 6.5606 | 1.23329663989843 | 0.503 | 0.013 | 1.0669573 |
| **AFAP1** | 8.8596 | 0.928570465863664 | 0.984 | 0.838 | 1.4408437 |
| **GEMIN2** | 8.9625 | 0.956104003803116 | 0.868 | 0.454 | 1.4575804 |
| **CNN1** | 6.4028 | -2.17511475900063 | 0.203 | 0.684 | 1.0412947 |
| **ID1** | 9.2225 | -1.35228600286829 | 0.951 | 0.982 | 1.4998573 |
| **PHF20** | 1.6310 | -0.879389734272048 | 0.791 | 0.972 | 2.6525762 |
| **DPF1** | 3.5152 | 1.22443926106382 | 0.855 | 0.49 | 5.7168164 |
| **PUS1** | 4.8047 | 0.831934464285889 | 0.995 | 0.935 | 7.8140017 |
| **SMG6** | 5.4751 | -1.02286901638957 | 0.453 | 0.844 | 8.9042372 |

| | | | | | |
|---|---|---|---|---|---|
| **PRTG** | 6.1116 | -2.43613212383066 | 0.048 | 0.511 | 9.9393197 |
| **PCMTD2** | 9.4242 | 0.805034069362883 | 1 | 0.989 | 1.5326717 |
| **ABHD12B** | 1.2267 | 0.911024265014711 | 0.595 | 0.069 | 1.9950504 |
| **GNG2** | 1.4745 | -0.941457240364721 | 0.067 | 0.544 | 2.3981128 |
| **BRK1** | 2.5996 | 0.878298903787808 | 0.591 | 0.079 | 4.2278483 |
| **RFX3** | 3.9990 | -1.06413142086989 | 0.199 | 0.697 | 6.5035764 |
| **BNIP3L** | 8.0941 | -0.900929975820842 | 0.886 | 0.989 | 1.3163490 |
| **FRK** | 1.2396 | 0.884141002162515 | 0.573 | 0.063 | 2.0161195 |
| **IDH2** | 1.8244 | -1.09624874784374 | 0.913 | 0.991 | 2.9671173 |
| **CACFD1** | 1.9039 | -1.01956957376111 | 0.63 | 0.925 | 3.0963210 |
| **USP18** | 4.9766 | 1.12056416311508 | 0.773 | 0.279 | 8.0934996 |
| **ENC1** | 8.0691 | -2.34807237364227 | 0.054 | 0.516 | 1.3122819 |
| **WDR34** | 8.8940 | 0.861645922037388 | 0.991 | 0.936 | 1.4464446 |
| **SLC25A5** | 2.0470 | 0.893020517519703 | 0.595 | 0.087 | 3.3291349 |
| **PIK3R5** | 2.8706 | 1.00695073340707 | 0.677 | 0.166 | 4.6684605 |
| **RND3** | 3.0876 | -1.24156813371655 | 0.043 | 0.505 | 5.0214978 |
| **SMARCA1** | 3.3602 | -1.15565605226622 | 0.803 | 0.971 | 5.4647477 |
| **ABHD17B** | 8.3815 | -0.908194744470138 | 0.965 | 0.998 | 1.3630887 |
| **TMEM200C** | 1.3484 | 0.986368208566664 | 0.646 | 0.121 | 2.1930112 |
| **SRXN1** | 2.0939 | -1.1515568187845 | 0.579 | 0.91 | 3.4053248 |
| **CNTRL** | 2.6474 | -1.087762932947 | 0.341 | 0.801 | 4.3055713 |
| **FOXI3** | 3.1218 | -1.40655380312429 | 0.942 | 0.972 | 5.0770960 |
| **C5orf15** | 8.5941 | -0.816579122489587 | 0.932 | 0.993 | 1.3976680 |
| **PLEKHG1** | 1.4633 | 0.955945422030826 | 0.643 | 0.121 | 2.3798273 |
| **NRIP3** | 1.7658 | 1.03459047540853 | 0.718 | 0.214 | 2.8717575 |
| **LZTS2** | 1.7947 | -1.04888481870144 | 0.741 | 0.951 | 2.9187894 |
| **ARGFX** | 2.8109 | 0.818382058124065 | 0.475 | 0.003 | 4.5715154 |
| **PPP1R18** | 3.1306 | -1.28622168218658 | 0.203 | 0.691 | 5.0914142 |
| **NRN1L** | 7.1072 | -1.3517140757201 | 0.058 | 0.523 | 1.1558594 |
| **GEMIN6** | 8.7304 | 0.809433465919079 | 0.995 | 0.952 | 1.4198405 |
| **LDLRAP1** | 1.3760 | 0.856065799189518 | 0.995 | 0.914 | 2.2378980 |
| **IFI27L2** | 1.4772 | -1.11424541974221 | 0.154 | 0.645 | 2.4025038 |
| **RTBDN** | 2.1452 | 0.937514096110279 | 0.593 | 0.081 | 3.4888810 |
| **ZNF229** | 5.6364 | 1.0396245149279 | 0.764 | 0.281 | 9.1665880 |

| | | | | | |
|---|---|---|---|---|---|
| **DNMT3B** | 6.7100 | -1.38862277936241 | 0.998 | 0.993 | 1.0912568 |
| **OSMR** | 7.6107 | 1.15458180622743 | 0.624 | 0.113 | 1.2377420 |
| **FAM20A** | 2.5500 | -1.75999684417922 | 0.498 | 0.846 | 4.1471992 |
| **TNKS1BP1** | 3.2603 | -0.91220100545813 | 0.854 | 0.979 | 5.3023877 |
| **CHCHD4** | 3.2901 | 0.923931915107959 | 0.955 | 0.706 | 5.3507690 |
| **SLC15A1** | 6.4366 | 0.974331769622236 | 0.604 | 0.094 | 1.0467859 |
| **NDUFAF7** | 6.6403 | 0.841152557997317 | 0.99 | 0.914 | 1.0799282 |
| **ADGRE5** | 7.2589 | 1.03120336154109 | 0.866 | 0.473 | 1.1805244 |
| **IRF8** | 7.2661 | 1.08589665996378 | 0.49 | 0.014 | 1.1816866 |
| **EYA2** | 1.2120 | -1.27007627161315 | 0.133 | 0.613 | 1.9711248 |
| **CAV1** | 1.5242 | -1.52503417039909 | 0.11 | 0.587 | 2.4788873 |
| **FBXO6** | 1.9599 | 0.840403383468873 | 0.536 | 0.042 | 3.1874306 |
| **FOXB1** | 2.9141 | -1.39347902510623 | 0.135 | 0.616 | 4.7392626 |
| **RHOV** | 3.9520 | 0.96557076967329 | 0.62 | 0.112 | 6.4271406 |
| **HEG1** | 7.3527 | -1.22975983220013 | 0.19 | 0.671 | 1.1957838 |
| **RAVER2** | 9.1903 | -1.01571689896762 | 0.329 | 0.783 | 1.4946272 |
| **TLN1** | 9.6774 | -0.836314168892536 | 0.957 | 0.997 | 1.5738367 |
| **HDHD2** | 1.1501 | -0.87894953103626 | 0.774 | 0.966 | 1.8704641 |
| **EPHA7** | 1.4878 | -1.49869286844535 | 0.756 | 0.952 | 2.4197597 |
| **CTSV** | 1.8973 | -1.52213882615049 | 0.909 | 0.981 | 3.0857285 |
| **C11orf1** | 5.6529 | -0.928585691778704 | 0.729 | 0.958 | 9.1933846 |
| **RTN4RL1** | 6.3010 | -1.18061611780076 | 0.239 | 0.705 | 1.0247407 |
| **COMT** | 2.1363 | -1.20238490247851 | 0.338 | 0.781 | 3.4743522 |
| **BAIAP2L1** | 2.6088 | 0.823780508868689 | 0.99 | 0.888 | 4.2427161 |
| **TEAD1** | 5.1115 | -1.04960636929753 | 0.493 | 0.862 | 8.3128887 |
| **CARMIL1** | 9.1332 | -1.12795388667403 | 0.327 | 0.774 | 1.4853443 |
| **TMEFF1** | 1.0937 | -1.28681755556428 | 0.789 | 0.967 | 1.7788162 |
| **SERPINF1** | 1.3667 | 0.998651132774373 | 0.972 | 0.802 | 2.2227683 |
| **PHLDB2** | 1.4737 | -1.18998921507927 | 0.246 | 0.731 | 2.3967278 |
| **RBMXL1** | 1.5780 | 0.947997050229105 | 0.89 | 0.537 | 2.5663635 |
| **NELFE** | 1.9945 | 0.92349874299934 | 0.788 | 0.317 | 3.2436813 |
| **MED12** | 3.6033 | -1.04846942847602 | 0.103 | 0.579 | 5.8601533 |
| **ST5** | 1.8878 | -0.972961585651421 | 0.722 | 0.958 | 3.0702515 |
| **GATAD2B** | 2.2685 | -0.867461499619053 | 0.771 | 0.971 | 3.6894230 |

| | | | | | |
|---|---|---|---|---|---|
| **CPXM1** | 3.4589 | -1.17103249207145 | 0.89 | 0.981 | 5.6253433 |
| **BRSK2** | 3.5419 | 0.886383807765637 | 0.578 | 0.078 | 5.7603326 |
| **ARHGEF10L** | 4.5967 | -1.00341403998656 | 0.211 | 0.692 | 7.4757146 |
| **CRY2** | 5.1666 | 0.898882167686616 | 0.973 | 0.865 | 8.4025908 |
| **RAD51** | 8.5243 | 0.978915679638967 | 0.989 | 0.878 | 1.3863160 |
| **ELF4** | 2.9152 | -1.37716875099374 | 0.103 | 0.571 | 4.7410315 |
| **TMSB15A** | 1.4758 | -1.69564403802627 | 0.103 | 0.571 | 2.4001887 |
| **COX20** | 1.5824 | 0.961340430057207 | 0.804 | 0.353 | 2.5735414 |
| **KIAA1324L** | 2.1796 | -0.973842140677541 | 0.564 | 0.895 | 3.5447826 |
| **TSPAN9** | 6.9714 | -1.01706691270495 | 0.751 | 0.952 | 1.1337625 |
| **UCK2** | 1.1311 | -0.917668381537271 | 0.913 | 0.992 | 1.8396522 |
| **COL2A1** | 2.0061 | -2.24863557533129 | 0.012 | 0.436 | 3.2625718 |
| **TACC2** | 5.4380 | -0.980725137776798 | 0.511 | 0.879 | 8.8438685 |
| **SPR** | 9.8992 | 0.996437257693054 | 0.858 | 0.427 | 1.6099138 |
| **HS6ST1** | 1.0385 | -1.23232638147054 | 0.935 | 0.99 | 1.6890236 |
| **GRHL1** | 2.2457 | 1.0169092487662 | 0.735 | 0.25 | 3.6522707 |
| **CARNMT1** | 4.6347 | 0.803133882650345 | 0.988 | 0.885 | 7.5375016 |
| **PAX5** | 5.2289 | 0.895475314751885 | 0.543 | 0.054 | 8.5037642 |
| **TPP1** | 6.3925 | -0.985174514667555 | 0.957 | 0.993 | 1.0396158 |
| **FAM155A** | 8.4375 | 0.937404125559647 | 0.557 | 0.065 | 1.3721953 |
| **MINDY2** | 2.9186 | 0.824264624984502 | 0.98 | 0.871 | 4.7466313 |
| **PTMS** | 8.1400 | -1.00983692164602 | 1 | 1 | 1.3238220 |
| **CALM1** | 9.3407 | 0.88536677274172 | 0.949 | 0.691 | 1.5190913 |
| **LHFPL2** | 1.0349 | -1.00118280240828 | 0.146 | 0.622 | 1.6831001 |
| **SGSM3** | 1.1031 | 0.916922243188102 | 0.931 | 0.695 | 1.7941270 |
| **HDAC7** | 1.4721 | -1.03929482538532 | 0.624 | 0.916 | 2.3942003 |
| **HCN2** | 1.5117 | 0.890868567193899 | 0.96 | 0.673 | 2.4586053 |
| **CCN2** | 2.4852 | -1.82115450453225 | 0.625 | 0.924 | 4.0416887 |
| **DAPK3** | 2.6820 | -1.01940624990186 | 0.985 | 0.996 | 4.3617876 |
| **CTTNBP2** | 3.6030 | -0.817993368722228 | 0.028 | 0.461 | 5.8595611 |
| **PSRC1** | 3.8155 | 1.12514021407806 | 0.978 | 0.812 | 6.2052999 |
| **BTG1** | 6.9364 | -1.04412988545201 | 0.705 | 0.957 | 1.1280798 |
| **NFKBIA** | 8.4881 | 0.843413889955345 | 0.997 | 0.956 | 1.3804310 |
| **UNC5D** | 1.3950 | -0.842416544546893 | 0.068 | 0.527 | 2.2687995 |

| | | | | | |
|---|---|---|---|---|---|
| **COLEC12** | 1.5350 | -2.27998387685458 | 0.13 | 0.591 | 2.49637595 |
| **SELENOM** | 1.6603 | 0.968642933253033 | 0.877 | 0.502 | 2.70026707 |
| **ZNF385A** | 2.1563 | -1.0909196315842 | 0.164 | 0.639 | 3.50680387 |
| **MFSD10** | 2.4927 | -0.925957908123227 | 0.786 | 0.963 | 4.05396286 |
| **WDR5** | 3.8192 | 0.804802387772613 | 0.983 | 0.865 | 6.21132472 |
| **VEPH1** | 4.0844 | -0.808302140289515 | 0.053 | 0.502 | 6.64252186 |
| **STK40** | 8.1277 | -1.01926618560428 | 0.536 | 0.877 | 1.32182154 |
| **UGCG** | 8.8038 | 0.812966189295337 | 1 | 0.988 | 1.43177508 |
| **OSBPL6** | 9.9542 | -0.906521562908847 | 0.088 | 0.553 | 1.61886467 |
| **STRA6** | 1.0741 | -1.10146955092511 | 0.118 | 0.587 | 1.74690562 |
| **PRDM1** | 1.5259 | 0.981055584768736 | 0.88 | 0.464 | 2.48165358 |
| **PIM1** | 1.9210 | -1.15755625894981 | 0.96 | 0.989 | 3.12427590 |
| **RASSF9** | 4.0037 | -1.09272187421818 | 0.022 | 0.449 | 6.51134378 |
| **AKAP3** | 6.0895 | 0.847861626415989 | 0.513 | 0.037 | 9.90337629 |
| **FAM83B** | 7.1585 | 0.827131967403398 | 0.559 | 0.071 | 1.16419211 |
| **SHISA3** | 7.4055 | -1.70266861660081 | 0.057 | 0.501 | 1.20436772 |
| **OST4** | 1.1232 | 0.890617490700497 | 0.71 | 0.221 | 1.82670289 |
| **MXRA5** | 1.4683 | 1.30176003012015 | 0.76 | 0.319 | 2.38793064 |
| **KLC3** | 2.1707 | 0.929975272816027 | 0.717 | 0.215 | 3.53024756 |
| **CAVIN1** | 5.3512 | -1.78409645031901 | 0.487 | 0.822 | 8.70278609 |
| **ST14** | 7.9326 | 1.04223067381691 | 0.656 | 0.162 | 1.29008938 |
| **GLI3** | 2.9131 | -2.71316073297092 | 0.029 | 0.455 | 4.73757462 |
| **TMX4** | 3.0913 | 0.816144858080826 | 0.993 | 0.939 | 5.02739202 |
| **TFPI2** | 3.2822 | 1.18212645440799 | 0.769 | 0.312 | 5.33797792 |
| **SCD** | 4.6802 | -0.810349628406951 | 1 | 1 | 7.61143252 |
| **MAP7D1** | 1.1381 | -1.62569522947868 | 0.458 | 0.808 | 1.85103050 |
| **ENDOG** | 1.8210 | 0.856953039403024 | 0.973 | 0.816 | 2.96151996 |
| **OCIAD2** | 1.8980 | -0.867998716173721 | 0.1 | 0.567 | 3.08685071 |
| **CADM4** | 5.3118 | -1.07299489470696 | 0.142 | 0.605 | 8.63866901 |
| **CD164** | 6.5002 | 0.863867047944605 | 0.916 | 0.608 | 1.05712859 |
| **ZNHIT2** | 8.4229 | 0.911898883733676 | 0.941 | 0.716 | 1.36982841 |
| **NEU1** | 1.1017 | -1.09245566914121 | 0.93 | 0.991 | 1.79178665 |
| **PRKG1** | 1.7697 | -1.00522588318691 | 0.021 | 0.442 | 2.87812303 |
| **MATK** | 3.3567 | 0.992485114883342 | 0.638 | 0.137 | 5.45907655 |

| | | | | | |
|---|---|---|---|---|---|
| **TMA7** | 4.9580 | 0.960943264923389 | 0.809 | 0.373 | 8.0633064 |
| **FOXJ1** | 1.2357 | -1.04202269604381 | 0.031 | 0.457 | 2.0096666 |
| **AP1M1** | 2.2528 | -0.880723302616211 | 0.835 | 0.98 | 3.6638460 |
| **KHDRBS1** | 2.4880 | -0.862150548752174 | 0.962 | 0.994 | 4.0462908 |
| **CACHD1** | 4.6107 | -1.198713940851 | 0.822 | 0.958 | 7.4984677 |
| **TSKS** | 5.4394 | 0.821984784990616 | 0.509 | 0.039 | 8.8461376 |
| **GPR160** | 5.5050 | 1.25449546364546 | 0.759 | 0.303 | 8.9527888 |
| **AIRE** | 7.6136 | 1.08110278975572 | 0.625 | 0.132 | 1.2382102 |
| **RHBDL3** | 1.1483 | -1.01449275002233 | 0.084 | 0.538 | 1.8676233 |
| **USP51** | 2.4249 | -0.990730443073615 | 0.293 | 0.741 | 3.9436830 |
| **COL9A3** | 2.4672 | -0.952855489029376 | 0.997 | 0.994 | 4.0125025 |
| **MPI** | 2.9085 | 0.80855007570044 | 0.984 | 0.903 | 4.7302403 |
| **DDX10** | 4.3774 | 0.920319568463411 | 0.832 | 0.427 | 7.1190506 |
| **S100A16** | 4.7485 | 1.24316002169805 | 0.732 | 0.26 | 7.7226275 |
| **ANKRD10** | 4.7613 | -1.16327431349419 | 0.979 | 0.993 | 7.7433331 |
| **DYRK2** | 5.2314 | -0.949769327650791 | 0.671 | 0.931 | 8.5079430 |
| **MYL7** | 5.3525 | -1.68809263379052 | 0.029 | 0.45 | 8.7048269 |
| **MAFA** | 6.9603 | 1.0588807207223 | 0.749 | 0.265 | 1.1319627 |
| **ZNF841** | 2.1035 | 0.986908848565694 | 0.714 | 0.239 | 3.4209772 |
| **ZNF695** | 2.1756 | 0.891185451654186 | 0.965 | 0.755 | 3.5382182 |
| **STC1** | 7.1526 | -1.05877901279444 | 0.051 | 0.492 | 1.1632301 |
| **STX6** | 1.4770 | -0.826231977589421 | 0.817 | 0.972 | 2.4021995 |
| **TADA3** | 1.9982 | -0.808886101411295 | 0.902 | 0.988 | 3.2497390 |
| **ACSF2** | 2.6718 | 1.07268345854464 | 0.789 | 0.352 | 4.3452251 |
| **C19orf85** | 4.7021 | 1.40739571537517 | 0.68 | 0.202 | 7.6471297 |
| **ZNF358** | 5.2914 | -0.825058572888054 | 0.93 | 0.996 | 8.6055322 |
| **PDRG1** | 6.9677 | -0.822273149912536 | 0.775 | 0.963 | 1.1331711 |
| **OSER1** | 9.0372 | -0.823521500236295 | 0.889 | 0.984 | 1.4697226 |
| **ENAH** | 1.4562 | -0.815410352914207 | 0.998 | 0.999 | 2.3683560 |
| **RCN3** | 1.6663 | 0.928178388745464 | 0.628 | 0.139 | 2.7100570 |
| **EIF2AK2** | 2.2971 | -0.87056079999283 | 0.676 | 0.939 | 3.7358454 |
| **ZBTB18** | 2.9456 | 0.94170662378196 | 0.937 | 0.7 | 4.7905828 |
| **TAF9B** | 3.6429 | -0.95529693752814 | 0.446 | 0.82 | 5.9244489 |
| **LAMB2** | 3.6902 | -1.27937654023704 | 0.707 | 0.931 | 6.0014914 |

| | | | | | |
|---|---|---|---|---|---|
| **TFPI** | 5.7118 | -1.79814277462569 | 0.17 | 0.632 | 9.2891616 |
| **ISYNA1** | 8.3392 | -0.972943056838458 | 0.979 | 0.997 | 1.3562173 |
| **TUNAR** | 8.3442 | -1.20491244199913 | 0.268 | 0.715 | 1.3570279 |
| **MOB2** | 9.3622 | 0.838458375920281 | 0.948 | 0.756 | 1.5225867 |
| **BTK** | 9.8673 | 0.997226465677439 | 0.514 | 0.048 | 1.6047323 |
| **TNC** | 1.2799 | -1.44332268481191 | 0.031 | 0.453 | 2.0815127 |
| **FOS** | 1.4745 | -1.45275334288759 | 0.231 | 0.693 | 2.3979931 |
| **LGI1** | 2.2260 | -1.67147165464307 | 0.026 | 0.443 | 3.6202428 |
| **HAVCR1** | 2.4921 | 0.817580084377171 | 0.436 | 0 | 4.0529059 |
| **ZNF300** | 3.5955 | 0.942017763272422 | 0.765 | 0.287 | 5.8474822 |
| **ZNF618** | 3.9399 | -0.927671231732355 | 0.762 | 0.945 | 6.4074714 |
| **FARP1** | 3.9810 | -1.09133895873172 | 0.747 | 0.949 | 6.4743317 |
| **CHST12** | 4.6782 | 0.883955774260185 | 0.933 | 0.698 | 7.6082842 |
| **FABP3** | 5.6243 | 0.892934258998362 | 0.998 | 0.876 | 9.1468263 |
| **ERBB2** | 1.0286 | -0.937481941175285 | 0.911 | 0.986 | 1.6728208 |
| **EXOSC4** | 1.7152 | 0.802131565397296 | 0.988 | 0.922 | 2.7894497 |
| **GJA10** | 2.1924 | 0.893581122685395 | 0.545 | 0.074 | 3.5655386 |
| **PALLD** | 2.7117 | -1.81664178037356 | 0.214 | 0.666 | 4.4101597 |
| **LARP1B** | 2.7386 | 0.848249592456981 | 0.967 | 0.814 | 4.4538537 |
| **HPD** | 4.3341 | -1.50261904484454 | 0.184 | 0.625 | 7.0485649 |
| **FKRP** | 6.4002 | -0.985731116531646 | 0.416 | 0.805 | 1.0408670 |
| **WDFY2** | 1.0655 | -1.00813103287044 | 0.795 | 0.962 | 1.7329158 |
| **GRAMD1B** | 1.9978 | -1.12978904546798 | 0.059 | 0.492 | 3.2490801 |
| **ANKRD50** | 2.8832 | -0.932124438680178 | 0.661 | 0.929 | 4.6891003 |
| **C14orf132** | 2.9702 | -1.24817327668965 | 0.06 | 0.495 | 4.8305807 |
| **ZNF398** | 9.7577 | 0.868820234476305 | 0.978 | 0.834 | 1.5869037 |
| **HAPLN3** | 1.0118 | -0.875777849498969 | 0.09 | 0.542 | 1.6456007 |
| **IGFBP5** | 1.1801 | -4.01210334642829 | 0.09 | 0.532 | 1.9193362 |
| **DLG1** | 1.6632 | -0.816747341959069 | 0.923 | 0.986 | 2.7049065 |
| **MAP3K3** | 5.0858 | -0.963896547665338 | 0.487 | 0.852 | 8.2711528 |
| **CGREF1** | 1.3137 | 1.24595050182912 | 0.577 | 0.111 | 2.1365394 |
| **H2AFY** | 2.9000 | -0.969731704159283 | 0.1 | 0.543 | 4.7163247 |
| **ERICH5** | 3.4651 | 0.840449888704324 | 0.554 | 0.076 | 5.6352990 |
| **CDC42EP3** | 5.3572 | -1.13612761078748 | 0.22 | 0.675 | 8.7125077 |

| | | | | | |
|---|---|---|---|---|---|
| **ARSI** | 2.8888 | 0.924074189084114 | 0.916 | 0.63 | 4.6981076 |
| **SLC5A3** | 5.7596 | -1.25924288417663 | 0.515 | 0.857 | 9.3668480 |
| **NCS1** | 5.9707 | -1.03670317844356 | 0.546 | 0.873 | 9.7102054 |
| **INPP5D** | 8.9974 | 0.971139148008623 | 0.877 | 0.514 | 1.4632575 |
| **HHIPL1** | 1.6769 | -1.05204623946559 | 0.208 | 0.66 | 2.7271526 |
| **TCEAL3** | 4.8454 | -0.827805516640139 | 0.116 | 0.573 | 7.8801683 |
| **TMEM138** | 4.8521 | 0.850348221790629 | 0.948 | 0.762 | 7.8910249 |
| **C1orf54** | 5.5098 | -0.814622428562141 | 0.101 | 0.553 | 8.9606354 |
| **ALDH1B1** | 6.6795 | -1.02458265602266 | 0.738 | 0.923 | 1.0862998 |
| **AP1S1** | 8.8722 | 0.817392173327993 | 0.997 | 0.975 | 1.4428925 |
| **SH3BP5** | 1.0655 | -1.07500624747785 | 0.233 | 0.684 | 1.7329123 |
| **RTL8A** | 1.0760 | -0.975270981089271 | 0.308 | 0.742 | 1.7500032 |
| **1,00 PGK** | 1.4270 | 1.03881768419809 | 0.705 | 0.248 | 2.3208689 |
| **CTNNA3** | 1.6996 | -0.909214009377107 | 0.043 | 0.466 | 2.7641084 |
| **NAPRT** | 2.1410 | 0.880716194260616 | 0.896 | 0.515 | 3.4820088 |
| **SPRY4** | 3.7866 | -1.00053979539399 | 0.024 | 0.428 | 6.1581725 |
| **ELMOD2** | 4.8346 | -0.910250024706981 | 0.589 | 0.896 | 7.8626688 |
| **MDM2** | 5.5620 | 0.951528984501784 | 0.998 | 0.975 | 9.0456045 |
| **CNKSR2** | 5.7954 | -0.934549214236881 | 0.073 | 0.507 | 9.4251048 |
| **EEF1AKMT4** | 8.0460 | 0.848958047334622 | 0.955 | 0.753 | 1.3085254 |
| **DTNA** | 1.5499 | -1.36601112793575 | 0.032 | 0.443 | 2.5207372 |
| **PTN** | 3.1148 | -1.43150753444305 | 0.006 | 0.392 | 5.0656267 |
| **ZBTB12** | 3.4560 | -0.809818133135516 | 0.054 | 0.479 | 5.6205897 |
| **NIPSNAP1** | 8.1858 | -0.819079295620857 | 0.905 | 0.99 | 1.3312571 |
| **WWC1** | 1.8309 | -0.963944167758278 | 0.198 | 0.654 | 2.9777522 |
| **FYN** | 2.2776 | -1.17438975921937 | 0.724 | 0.938 | 3.7040931 |
| **WBP1L** | 4.3472 | -0.850150538355666 | 0.914 | 0.984 | 7.0699675 |
| **TUBA1A** | 8.5605 | -1.40800498605322 | 0.653 | 0.895 | 1.3921986 |
| **PCDHB16** | 2.3810 | -0.81832639732556 | 0.063 | 0.492 | 3.8723061 |
| **HDAC5** | 3.6423 | -0.826178326998734 | 0.855 | 0.983 | 5.9234780 |
| **ATRN** | 5.7068 | -0.81027738046405 | 0.893 | 0.988 | 9.2810425 |
| **EXOC4** | 8.5484 | -0.861789765838226 | 0.838 | 0.976 | 1.3902374 |
| **PIP4K2A** | 1.2171 | 0.870546963966537 | 0.691 | 0.224 | 1.9794125 |
| **TYRO3** | 1.4568 | -1.04020526471986 | 0.749 | 0.947 | 2.3691964 |

| | | | | | |
|---|---|---|---|---|---|
| **MOGAT3** | 1.8622 | 0.824780837841178 | 0.52 | 0.062 | 3.0285043 |
| **CNP** | 2.7051 | -0.861082569479651 | 0.919 | 0.989 | 4.3994073 |
| **ADGRL2** | 3.1081 | -0.93317964420445 | 0.902 | 0.992 | 5.0547420 |
| **DNAJB4** | 3.2082 | 0.938668224564268 | 0.883 | 0.555 | 5.2175576 |
| **IRF2BPL** | 6.4079 | -1.21070565250161 | 0.868 | 0.958 | 1.0421307 |
| **OSBPL3** | 6.4635 | -1.02945776144078 | 0.334 | 0.753 | 1.0511616 |
| **TCL1B** | 6.8350 | 1.13308138155511 | 0.756 | 0.293 | 1.1115827 |
| **MED19** | 7.4864 | 0.824852530501067 | 0.94 | 0.711 | 1.2175294 |
| **TMEM47** | 8.1684 | -0.917522763129773 | 0.107 | 0.548 | 1.3284395 |
| **LHX5** | 8.3272 | -1.74639904388138 | 0.019 | 0.412 | 1.3542582 |
| **FEN1** | 1.1153 | 0.884325213329085 | 0.997 | 0.94 | 1.8139351 |
| **APOE** | 1.9700 | -0.885149099053202 | 1 | 1 | 3.2038162 |
| **PURB** | 4.8528 | -0.816348501732024 | 0.816 | 0.978 | 7.8922608 |
| **TOMM34** | 1.0414 | -0.818952865370535 | 0.88 | 0.985 | 1.6936440 |
| **RASL11A** | 1.6795 | 0.842312802100081 | 0.615 | 0.138 | 2.7313756 |
| **PDE7A** | 2.1079 | -0.924224305366646 | 0.873 | 0.976 | 3.4282244 |
| **GALNT6** | 2.6795 | 0.856850409643244 | 0.926 | 0.644 | 4.3577386 |
| **PDSS2** | 3.0823 | 0.830746615527149 | 0.978 | 0.857 | 5.0128305 |
| **F2RL1** | 3.3638 | 0.808784331165416 | 0.998 | 0.956 | 5.4707026 |
| **CDC42EP1** | 4.0953 | -0.843264594550783 | 0.035 | 0.441 | 6.6602992 |
| **LRRC61** | 3.3055 | -0.908166375354348 | 0.308 | 0.735 | 5.3757721 |
| **HBA2** | 4.0036 | 0.963207349050172 | 0.5 | 0.052 | 6.5111367 |
| **GINS1** | 4.0470 | -0.960618652977178 | 0.988 | 0.996 | 6.5816714 |
| **AGPAT3** | 4.8441 | 0.877746896084861 | 0.619 | 0.136 | 7.8780979 |
| **SLC39A14** | 1.0132 | 0.945959444668807 | 0.875 | 0.586 | 1.6479248 |
| **SHE** | 1.0879 | 0.842101668687056 | 0.456 | 0.024 | 1.7693952 |
| **SIK2** | 1.3528 | -0.924151005097693 | 0.381 | 0.788 | 2.2002139 |
| **TRIM38** | 3.4874 | 1.02603087152981 | 0.864 | 0.495 | 5.6717106 |
| **MB21D2** | 1.0623 | -0.907232639181396 | 0.261 | 0.699 | 1.7276520 |
| **ADCK2** | 1.1164 | 0.891643590179616 | 0.885 | 0.567 | 1.8157609 |
| **SAP25** | 1.8401 | 0.999592661321244 | 0.609 | 0.145 | 2.9925757 |
| **ZBTB33** | 2.1027 | -0.861723767858971 | 0.791 | 0.97 | 3.4197229 |
| **KLHL15** | 2.4695 | -0.93843637458349 | 0.394 | 0.809 | 4.0162546 |
| **TMLHE** | 3.3484 | -0.926628981047458 | 0.438 | 0.819 | 5.4456495 |

| | | | | | |
|---|---|---|---|---|---|
| **DMBX1** | 3.3899 | 0.859117908531816 | 0.775 | 0.285 | 5.5130358 |
| **NAGK** | 3.4419 | 0.82547738919542 | 0.996 | 0.973 | 5.5976705 |
| **CAP2** | 3.7223 | -1.00288288831469 | 0.643 | 0.908 | 6.0537176 |
| **ZNF552** | 5.0260 | 0.815164135030892 | 0.994 | 0.862 | 8.1738516 |
| **SSBP2** | 1.0023 | -0.971575418340024 | 0.436 | 0.821 | 1.6300617 |
| **CDKN2AIP** | 1.3387 | -0.907263784933329 | 0.711 | 0.932 | 2.1771903 |
| **RBP7** | 3.7026 | 0.84314931057688 | 0.522 | 0.068 | 6.0215652 |
| **MYLPF** | 6.5673 | 0.805905037863594 | 0.561 | 0.095 | 1.0680511 |
| **PLA2G2A** | 6.8737 | -0.891219796107852 | 0.011 | 0.391 | 1.1178729 |
| **KCNMB1** | 2.9348 | -0.988945345607173 | 0.015 | 0.397 | 4.7729439 |
| **SLC22A31** | 9.8743 | 1.04615522033347 | 0.694 | 0.237 | 1.6058594 |
| **FGF18** | 1.5789 | -1.15509672113189 | 0.142 | 0.576 | 2.5678908 |
| **DOCK3** | 2.7316 | -0.881324732359303 | 0.089 | 0.52 | 4.4424996 |
| **NTS** | 3.3935 | -1.49361949548702 | 0.1 | 0.529 | 5.5188694 |
| **SLC12A8** | 8.9507 | -1.04584065944293 | 0.169 | 0.603 | 1.4556560 |
| **RGS3** | 1.0430 | -0.984985168959964 | 0.057 | 0.471 | 1.6962880 |
| **GALC** | 1.1371 | -0.954994746152722 | 0.494 | 0.84 | 1.8493628 |
| **SH3BGRL3** | 2.1973 | -0.970961455949221 | 0.966 | 0.993 | 3.5734904 |
| **NDN** | 4.2258 | -0.878138276936667 | 0.645 | 0.925 | 6.8725384 |
| **AIFM2** | 4.9903 | 0.885188256033919 | 0.681 | 0.216 | 8.1158261 |
| **FAM98A** | 5.0476 | 0.850669204690105 | 0.997 | 0.945 | 8.2090338 |
| **KRT19** | 1.0624 | -1.05121770291102 | 0.811 | 0.966 | 1.7278308 |
| **OTX2** | 1.8915 | -1.50817762996657 | 0.741 | 0.918 | 3.0762011 |
| **TBX22** | 5.4149 | -1.06370128178181 | 0.021 | 0.408 | 8.8063231 |
| **B3GALT5** | 6.2991 | -0.826655613879353 | 0.068 | 0.484 | 1.0244304 |
| **TRIM2** | 9.6427 | -0.908422173045221 | 0.849 | 0.972 | 1.5682073 |
| **RAB42** | 1.0810 | 0.803344243449447 | 0.514 | 0.068 | 1.7580406 |
| **KRT7** | 1.5922 | -1.18131597331839 | 0.074 | 0.489 | 2.5895372 |
| **RASL10B** | 2.0313 | -0.951710909997828 | 0.074 | 0.491 | 3.3035445 |
| **LAD1** | 2.2410 | 0.851351493507067 | 0.538 | 0.085 | 3.6445720 |
| **SIPA1L2** | 2.5880 | -0.970296734630863 | 0.698 | 0.938 | 4.2088885 |
| **SYK** | 4.4430 | 0.90140295440223 | 0.662 | 0.19 | 7.2258016 |
| **STOX1** | 5.3742 | -0.803456043086511 | 0.048 | 0.456 | 8.7402240 |
| **ZNF607** | 5.4043 | 0.859494421588461 | 0.683 | 0.231 | 8.7890870 |

| | | | | | |
|---|---|---|---|---|---|
| **PLAGL2** | 1.3387 | -0.862859770100901 | 0.607 | 0.881 | 2.17714054 |
| **RBCK1** | 2.4114 | -0.934727872697314 | 0.601 | 0.886 | 3.92166513 |
| **LRCH3** | 6.2523 | -0.815315941385917 | 0.818 | 0.965 | 1.01682638 |
| **AHNAK** | 7.1139 | -1.36595706985859 | 0.905 | 0.985 | 1.15693775 |
| **NUCB2** | 1.3941 | 0.963143904803654 | 0.909 | 0.668 | 2.26734294 |
| **DNMT1** | 1.7917 | -0.854706715660908 | 0.97 | 0.992 | 2.91397360 |
| **THY1** | 2.0278 | -1.0799003089282 | 0.035 | 0.422 | 3.29789912 |
| **USO1** | 2.4696 | -0.958761688643837 | 1 | 0.998 | 4.01641280 |
| **FADS3** | 3.2914 | 0.860066708438925 | 0.842 | 0.476 | 5.35285750 |
| **GSTK1** | 3.4314 | -0.957822801820332 | 0.646 | 0.91 | 5.58060558 |
| **USP44** | 5.4901 | -0.941604648016187 | 0.13 | 0.559 | 8.92856881 |
| **ARHGAP6** | 5.5336 | -0.998898642967381 | 0.168 | 0.601 | 8.99943195 |
| **SLC43A2** | 7.7696 | -0.963318727411447 | 0.794 | 0.952 | 1.26357910 |
| **MCM8** | 9.2975 | -0.823865481855659 | 0.944 | 0.984 | 1.51205372 |
| **ZNF729** | 1.2151 | 1.63979300053323 | 0.466 | 0.049 | 1.97615243 |
| **GLIPR2** | 1.2348 | -1.49650876884225 | 0.946 | 0.977 | 2.00829447 |
| **PLPP5** | 2.8924 | 0.961062247897828 | 0.909 | 0.673 | 4.70404370 |
| **GALNT2** | 3.5447 | -0.978994837969894 | 0.897 | 0.98 | 5.76474629 |
| **B2M** | 5.1756 | -1.13941952347941 | 0.689 | 0.918 | 8.41716809 |
| **NAA35** | 1.4087 | 0.803774712122957 | 0.865 | 0.509 | 2.29100000 |
| **DNMBP** | 7.4867 | -0.92374843913226 | 0.727 | 0.934 | 1.21757302 |
| **DPP10** | 2.1302 | -1.11820777467289 | 0.289 | 0.694 | 3.46446980 |
| **PC** | 2.8175 | 1.00279667865757 | 0.848 | 0.523 | 4.58213570 |
| **GPC1** | 1.7173 | -1.02290731295363 | 0.871 | 0.976 | 2.79295078 |
| **CRIP3** | 3.8071 | 1.13860488078255 | 0.611 | 0.173 | 6.19161474 |
| **ZNF43** | 5.5786 | -0.884365625600496 | 0.419 | 0.793 | 9.07254240 |
| **FBXL16** | 1.4539 | 0.861178357063289 | 0.612 | 0.159 | 2.36458384 |
| **ZNF578** | 4.2800 | 0.841764443802272 | 0.745 | 0.304 | 6.96058079 |
| **EPB41L1** | 5.9223 | -0.842216308169559 | 0.877 | 0.982 | 9.63156940 |
| **DPYSL5** | 6.4505 | -1.98278946886808 | 0.02 | 0.386 | 1.04905462 |
| **PTGIS** | 7.4287 | -1.22986708815215 | 0.061 | 0.454 | 1.20813530 |
| **CDH5** | 7.9227 | 0.818392596783414 | 0.459 | 0.038 | 1.28847688 |
| **C11orf80** | 1.0897 | -0.881193231892935 | 0.471 | 0.835 | 1.77230259 |
| **PCNX4** | 1.4532 | -1.00430501503093 | 0.483 | 0.815 | 2.36339604 |

| | | | | | |
|---|---|---|---|---|---|
| **LDHD** | 1.7719 | -1.16674057240391 | 0.278 | 0.698 | 2.8817834 |
| **WFDC2** | 5.2579 | -1.18571523602383 | 0.115 | 0.533 | 8.5509872 |
| **ST7** | 8.7773 | -0.854133066037054 | 0.233 | 0.659 | 1.4274652 |
| **GSDMD** | 1.0326 | 0.875337942936141 | 0.692 | 0.248 | 1.6794684 |
| **SLC39A6** | 1.3570 | -1.00070258619673 | 0.508 | 0.829 | 2.2069852 |
| **ZNF48** | 1.5941 | -0.987191152949165 | 0.737 | 0.935 | 2.5925123 |
| **RIT1** | 3.4703 | -0.893811601054247 | 0.419 | 0.788 | 5.6439012 |
| **CALM2** | 4.0912 | -0.81242514247979 | 0.996 | 1 | 6.6536474 |
| **ZNF703** | 5.7182 | -1.34412714469838 | 0.474 | 0.809 | 9.2996058 |
| **GREB1L** | 5.9646 | -2.11676845535309 | 0.016 | 0.376 | 9.7003677 |
| **IRS1** | 9.2931 | -0.887888722490614 | 0.154 | 0.583 | 1.5113441 |
| **UHRF2** | 1.2177 | -1.03848729284706 | 0.312 | 0.706 | 1.9804578 |
| **CPD** | 2.7749 | -0.86756814238616 | 0.968 | 0.997 | 4.5129807 |
| **BDNF** | 6.1798 | 0.930458543855473 | 0.456 | 0.041 | 1.0050256 |
| **LNPEP** | 7.3296 | -0.877786069660484 | 0.223 | 0.641 | 1.1920175 |
| **IKBIP** | 1.0107 | -0.907229108660262 | 0.391 | 0.765 | 1.6438215 |
| **ALPL** | 2.8769 | -0.84821183052019 | 0.988 | 0.997 | 4.6787964 |
| **USP46** | 6.3834 | -0.800320103245248 | 0.729 | 0.935 | 1.0381437 |
| **NRXN1** | 7.3726 | -0.827817504716717 | 0.089 | 0.502 | 1.1990097 |
| **FANCA** | 1.7674 | 0.83583346226361 | 0.968 | 0.797 | 2.8744216 |
| **PHLDB1** | 1.7845 | -0.970267413058483 | 0.661 | 0.908 | 2.9021740 |
| **SCML2** | 1.2118 | -0.827067209529597 | 0.558 | 0.861 | 1.9709015 |
| **RWDD4** | 1.4461 | 0.804110808814259 | 0.843 | 0.488 | 2.3518221 |
| **ANGPTL2** | 1.4671 | -0.92511499907088 | 0.029 | 0.397 | 2.3860942 |
| **MTM1** | 1.5677 | -0.871848496952486 | 0.775 | 0.956 | 2.5496959 |
| **STOX2** | 1.0557 | -1.01237430115699 | 0.303 | 0.707 | 1.7169197 |
| **DMPK** | 1.2942 | -0.944428742886429 | 0.278 | 0.694 | 2.1048848 |
| **CRTC1** | 1.5003 | -0.859853250408083 | 0.306 | 0.716 | 2.4400049 |
| **SYNGR1** | 1.7519 | -1.0654721081173 | 0.279 | 0.676 | 2.8491441 |
| **GRPR** | 3.5875 | -1.00076717645678 | 0.02 | 0.379 | 5.8344618 |
| **ARHGAP4** | 4.6601 | 0.826850270910191 | 0.511 | 0.08 | 7.5788091 |
| **BRINP1** | 9.7621 | -1.14738890619734 | 0.15 | 0.55 | 1.5876124 |
| **CIZ1** | 1.0003 | -1.03414770654966 | 0.412 | 0.765 | 1.6268965 |
| **CCDC71L** | 2.5444 | -1.03932834913 | 0.611 | 0.891 | 4.1380843 |

| | | | | | |
|---|---|---|---|---|---|
| **LZTS1** | 3.6420 | 0.824287355905463 | 0.849 | 0.496 | 5.9230983 |
| **MMD** | 5.8116 | -0.864279520375563 | 0.527 | 0.843 | 9.4514051 |
| **PLEKHG2** | 2.9936 | -0.92300530776576 | 0.292 | 0.692 | 4.8685078 |
| **CHST6** | 3.5787 | -0.948723034581235 | 0.416 | 0.767 | 5.8201352 |
| **SMAD7** | 3.9030 | -0.957552604761569 | 0.819 | 0.958 | 6.3474841 |
| **STON2** | 2.8798 | -1.23426322172273 | 0.324 | 0.721 | 4.6834904 |
| **DIXDC1** | 5.9062 | -0.983251974382587 | 0.365 | 0.75 | 9.6052564 |
| **MCAM** | 8.0495 | -1.31874840832437 | 0.506 | 0.82 | 1.3090928 |
| **RAB11FIP1** | 1.4331 | -0.823994767263724 | 0.94 | 0.989 | 2.3306876 |
| **FAM172A** | 1.7380 | -0.832483862578406 | 0.571 | 0.867 | 2.8266147 |
| **PMEL** | 2.2001 | -1.08537325907756 | 0.049 | 0.427 | 3.5780987 |
| **PDCD4** | 4.0944 | -0.93935553855609 | 0.504 | 0.842 | 6.6587615 |
| **ADD2** | 3.7208 | -0.905110568235879 | 0.911 | 0.973 | 6.0512224 |
| **CPE** | 4.3231 | -2.72331097197808 | 0.044 | 0.414 | 7.0307009 |
| **FLNC** | 4.6770 | -1.81663467619702 | 0.549 | 0.834 | 7.6062626 |
| **ZSCAN10** | 5.7336 | -0.940020240455111 | 0.981 | 0.96 | 9.3245976 |
| **NIPAL1** | 8.1574 | 0.872193863707173 | 0.865 | 0.505 | 1.3266530 |
| **SYS1** | 8.4458 | -0.810850257377894 | 0.694 | 0.916 | 1.3735541 |
| **ARX** | 9.3230 | 1.05728003573529 | 0.416 | 0.025 | 1.5162064 |
| **SOX2** | 1.5189 | -1.3334495873335 | 0.96 | 0.959 | 2.4703440 |
| **DAGLA** | 5.7144 | 0.897960391080851 | 0.815 | 0.459 | 9.2934900 |
| **VANGL1** | 7.3342 | -0.88989386935289 | 0.67 | 0.906 | 1.1927731 |
| **IGFBP4** | 7.4998 | -0.980736713833775 | 0.103 | 0.503 | 1.2197034 |
| **IRF2BP2** | 8.0005 | -0.919186514801956 | 0.534 | 0.842 | 1.3011339 |
| **DDR1** | 1.0842 | -1.06427297828864 | 0.886 | 0.977 | 1.7633218 |
| **FGFR3** | 1.3301 | -0.909565127121827 | 0.981 | 0.992 | 2.1631477 |
| **S100A4** | 1.5650 | -0.929357994672836 | 0.012 | 0.353 | 2.5452573 |
| **SPTLC3** | 3.2114 | -0.804231503361006 | 0.021 | 0.371 | 5.2227092 |
| **ROR2** | 3.3075 | -1.22962533801633 | 0.428 | 0.774 | 5.3791345 |
| **ZRANB3** | 5.0676 | -0.855088458358411 | 0.332 | 0.719 | 8.2415790 |
| **NNAT** | 6.0234 | 0.82187916753212 | 0.379 | 0.006 | 9.7959345 |
| **ALPP** | 1.3005 | 0.835192703569738 | 0.427 | 0.032 | 2.1150269 |
| **EFS** | 1.4225 | -1.53247237478453 | 0.3 | 0.673 | 2.3134976 |
| **NAV1** | 1.8720 | -1.13922200232346 | 0.416 | 0.755 | 3.0445270 |

| | | | | | |
|---|---|---|---|---|---|
| **SP8** | 3.1662 | -1.00210911656789 | 0.895 | 0.945 | 5.1491971 |
| **SIX5** | 3.7074 | -0.899503425130829 | 0.569 | 0.847 | 6.0294690 |
| **NCEH1** | 5.5147 | 0.883261625547671 | 0.779 | 0.39 | 8.9686832 |
| **HABP4** | 7.4519 | -0.962935661963173 | 0.267 | 0.667 | 1.2119120 |
| **CDK14** | 1.5341 | -1.17034459402536 | 0.266 | 0.662 | 2.4950388 |
| **HOXA1** | 2.2404 | -0.957653100720734 | 0.075 | 0.458 | 3.6436158 |
| **LTBR** | 6.7687 | 0.852552873724949 | 0.589 | 0.161 | 1.1008097 |
| **AGBL5** | 1.0679 | -0.867088440153074 | 0.665 | 0.901 | 1.7367789 |
| **MTSS2** | 1.2037 | -0.958313422847721 | 0.729 | 0.919 | 1.9576551 |
| **SVIL** | 1.2914 | -0.941602658703004 | 0.341 | 0.722 | 2.1003595 |
| **PAK6** | 1.5004 | 0.830878050993409 | 0.591 | 0.161 | 2.4401181 |
| **ASAP1** | 2.0478 | -0.879624610581227 | 0.748 | 0.946 | 3.3303838 |
| **CCDC92** | 7.1579 | -1.08427736164435 | 0.077 | 0.458 | 1.1641012 |
| **MAN1A1** | 1.0448 | -1.00073977067124 | 0.123 | 0.523 | 1.6992224 |
| **BAZ2B** | 1.8972 | -0.82716920704666 | 0.689 | 0.922 | 3.0855176 |
| **P3H1** | 3.8198 | -0.922512476757127 | 0.383 | 0.744 | 6.2122370 |
| **DEPDC1** | 8.1330 | 1.13853654116526 | 0.613 | 0.199 | 1.3226790 |
| **HOMER2** | 2.9589 | 1.04718298550332 | 0.925 | 0.774 | 4.8121504 |
| **EPB41L4B** | 3.0799 | 0.860323517590812 | 0.91 | 0.643 | 5.0088989 |
| **RARA** | 7.6949 | -0.887149710727344 | 0.214 | 0.62 | 1.2514218 |
| **PTGES3L** | 3.7460 | -0.836953198445274 | 0.186 | 0.59 | 6.0921914 |
| **TENT5A** | 4.1979 | 0.897967672231871 | 0.503 | 0.092 | 6.8271424 |
| **GREM2** | 1.6355 | 0.883977285606963 | 0.695 | 0.253 | 2.6599297 |
| **PRSS8** | 1.9758 | -1.24756401091322 | 0.762 | 0.9 | 3.2132780 |
| **CACNA1B** | 2.5352 | 0.844041939141324 | 0.74 | 0.324 | 4.1230325 |
| **AHDC1** | 1.6924 | -1.37886924256565 | 0.476 | 0.77 | 2.7525043 |
| **MACROD1** | 8.8791 | 0.822904705424764 | 0.754 | 0.353 | 1.4440127 |
| **EBF2** | 9.6277 | -1.18485321174397 | 0.108 | 0.489 | 1.5657682 |
| **PCBD1** | 7.5792 | -0.801554781700377 | 0.8 | 0.956 | 1.2326106 |
| **TRMT2A** | 1.1693 | 0.810143447743741 | 0.858 | 0.575 | 1.9016581 |
| **PLPPR3** | 1.1782 | -1.13811365728943 | 0.659 | 0.897 | 1.9161407 |
| **LRP1** | 1.3596 | -1.1956417137216 | 0.778 | 0.928 | 2.2111530 |
| **PKD2** | 8.2205 | -0.913614535641664 | 0.574 | 0.86 | 1.3369064 |
| **MAP2** | 2.4187 | -1.77126168831614 | 0.013 | 0.338 | 3.9336920 |

| | | | | | |
|---|---|---|---|---|---|
| **ANK2** | 2.5958 | -0.874542766182649 | 0.719 | 0.916 | 4.2217091 |
| **TENM3** | 3.1817 | -0.943317238168194 | 0.488 | 0.826 | 5.1745252 |
| **SLC16A9** | 3.4033 | 0.912728574468072 | 0.826 | 0.501 | 5.5347904 |
| **LMCD1** | 1.1740 | -1.08117464798649 | 0.027 | 0.367 | 1.9093960 |
| **ZNF627** | 1.5180 | -0.814604045656366 | 0.444 | 0.785 | 2.4688814 |
| **C3orf14** | 2.3775 | 0.820168310528393 | 0.874 | 0.635 | 3.8666057 |
| **PTPN13** | 2.6237 | -1.09522747306766 | 0.241 | 0.624 | 4.2670597 |
| **CCNE1** | 2.9698 | 0.821447794742593 | 0.943 | 0.728 | 4.8297894 |
| **MGAT5** | 1.7199 | -0.971928034941742 | 0.369 | 0.725 | 2.7972047 |
| **TRIB2** | 2.8500 | -0.806508488926331 | 0.138 | 0.529 | 4.6350676 |
| **SCLY** | 3.4991 | -0.843807393063421 | 0.467 | 0.787 | 5.6907363 |
| **C20orf27** | 4.9686 | -0.824833842341848 | 0.602 | 0.857 | 8.0805081 |
| **TSC22D3** | 8.2001 | -1.05514340225867 | 0.389 | 0.757 | 1.3335837 |
| **TSPEAR** | 9.1638 | -0.819811925214722 | 0.019 | 0.349 | 1.4903240 |
| **NUP210** | 4.4019 | -0.892377288927714 | 0.477 | 0.808 | 7.1589031 |
| **CSRP1** | 1.5089 | -1.9167449249756 | 0.883 | 0.96 | 2.4539688 |
| **CKS2** | 2.9127 | 0.971469501170513 | 0.976 | 0.883 | 4.7369897 |
| **RCBTB2** | 4.7600 | -0.851693796221142 | 0.235 | 0.624 | 7.7412247 |
| **CPT1A** | 1.0760 | 0.812473162755397 | 0.708 | 0.297 | 1.7500537 |
| **SNX25** | 1.2590 | 0.816263439205282 | 0.758 | 0.381 | 2.0475906 |
| **NFATC4** | 1.3480 | -1.01067879472482 | 0.25 | 0.641 | 2.1922817 |
| **THBS2** | 3.8730 | -0.913368038204478 | 0.066 | 0.427 | 6.2986777 |
| **BLVRB** | 4.9833 | -0.800451209840116 | 0.856 | 0.962 | 8.1043912 |
| **C18orf54** | 6.6013 | -0.828366685805228 | 0.635 | 0.883 | 1.0735720 |
| **CRTC3** | 7.9494 | -0.862989413222504 | 0.35 | 0.715 | 1.2928142 |
| **C1orf21** | 1.9312 | -1.02157158489346 | 0.045 | 0.385 | 3.1407611 |
| **CENPA** | 2.1429 | 0.97249858094792 | 0.982 | 0.851 | 3.4851605 |
| **FADS1** | 3.2794 | -0.860243959157345 | 0.981 | 0.999 | 5.3333495 |
| **PRSS35** | 3.9993 | 0.867304086752064 | 0.522 | 0.12 | 6.5042200 |
| **RNF145** | 3.4636 | -0.801154115338505 | 0.523 | 0.817 | 5.6329335 |
| **NOL4L** | 4.8999 | -0.922008818611532 | 0.361 | 0.722 | 7.9688266 |
| **SYNE3** | 7.4788 | -0.837093426721095 | 0.043 | 0.381 | 1.2162842 |
| **COTL1** | 7.8905 | -0.971396942832826 | 0.951 | 0.981 | 1.2832360 |
| **FNDC4** | 1.2392 | 0.815458230158006 | 0.873 | 0.59 | 2.0153472 |

| | | | | | |
|---|---|---|---|---|---|
| **CBX2** | 4.8226 | -0.862924219583842 | 0.822 | 0.953 | 7.8430077 |
| **BIK** | 5.4055 | -1.08697232364169 | 0.409 | 0.731 | 8.7910818 |
| **SLC12A2** | 1.1071 | 0.808827827749075 | 0.782 | 0.435 | 1.8005045 |
| **SPON1** | 1.8490 | -1.71324441788457 | 0.029 | 0.353 | 3.0071428 |
| **TDRD1** | 3.0037 | 0.868984105201033 | 0.371 | 0.02 | 4.8849675 |
| **IFI6** | 6.8259 | -0.999186844702596 | 0.162 | 0.543 | 1.1101075 |
| **NNT** | 1.4076 | 0.844942527023337 | 0.975 | 0.925 | 2.2891908 |
| **TCF7** | 1.1366 | -0.963854103860897 | 0.809 | 0.916 | 1.8485536 |
| **EZH2** | 6.4056 | -0.848785306386294 | 0.933 | 0.978 | 1.0417516 |
| **GNG4** | 1.1030 | 0.824215718008524 | 0.541 | 0.144 | 1.7938947 |
| **BRSK1** | 1.2128 | -0.876019235840579 | 0.195 | 0.568 | 1.9725225 |
| **TCEA2** | 3.6540 | -0.869121681810829 | 0.635 | 0.867 | 5.9425169 |
| **TXNDC16** | 9.8402 | -0.801662283536626 | 0.359 | 0.715 | 1.6003145 |
| **KLHL23** | 1.2520 | -0.878236533308534 | 0.621 | 0.875 | 2.0362485 |
| **TNFRSF12A** | 1.8025 | -1.17738673899565 | 0.86 | 0.964 | 2.9314506 |
| **ARHGAP11A** | 5.9161 | 0.918539107281237 | 0.908 | 0.7 | 9.6214280 |
| **SYNC** | 3.6204 | 0.80788901664499 | 0.544 | 0.148 | 5.8879852 |
| **ARVCF** | 7.0702 | -0.897404215003906 | 0.396 | 0.728 | 1.1498377 |
| **IRS2** | 1.3167 | -1.02917693633542 | 0.337 | 0.693 | 2.1414229 |
| **CD53** | 1.1576 | 0.898346954724111 | 0.682 | 0.284 | 1.8827611 |
| **SYNGAP1** | 1.2780 | -1.18349339408193 | 0.267 | 0.613 | 2.0785410 |
| **VIT** | 2.5232 | -1.3385086756278 | 0.028 | 0.34 | 4.1035231 |
| **PXN** | 3.6735 | -0.807069941982437 | 0.95 | 0.98 | 5.9743360 |
| **ZNF580** | 4.2987 | -0.890555189802114 | 0.559 | 0.837 | 6.9909971 |
| **CTSZ** | 5.6849 | -0.926699505505461 | 0.634 | 0.864 | 9.2454292 |
| **KIZ** | 1.3520 | -0.826489195290509 | 0.587 | 0.847 | 2.1987762 |
| **P3H2** | 2.2073 | -0.880207230840689 | 0.022 | 0.33 | 3.5898045 |
| **CCNA1** | 4.6305 | 1.20908640805391 | 0.481 | 0.105 | 7.5307248 |
| **SGSM2** | 2.7917 | -0.841790827775623 | 0.378 | 0.719 | 4.5401490 |
| **CDCP1** | 5.5542 | 0.801256459894505 | 0.74 | 0.359 | 9.0328306 |
| **NCAPD2** | 1.1529 | -1.03953860841844 | 0.854 | 0.956 | 1.8750078 |
| **RAI1** | 2.1956 | -0.864930430758857 | 0.095 | 0.441 | 3.5707838 |
| **CADM1** | 6.4115 | -1.02417361564367 | 0.619 | 0.867 | 1.0427073 |
| **PAX3** | 2.1558 | -1.36309943104932 | 0.011 | 0.3 | 3.5061329 |

| | | | | | |
|---|---|---|---|---|---|
| **DNAJB5** | 2.2806 | -0.827285405987883 | 0.379 | 0.721 | 3.7089674 |
| **CCN1** | 1.2158 | -1.38806682026338 | 0.798 | 0.948 | 1.9772815 |
| **SEMA3C** | 1.4103 | -0.87060825060831 | 0.009 | 0.295 | 2.2935766 |
| **SQLE** | 2.2593 | -0.843084699920192 | 0.937 | 0.985 | 3.6743589 |
| **SLC7A2** | 3.5155 | -1.15573969270634 | 0.376 | 0.683 | 5.7173072 |
| **PMAIP1** | 3.4954 | -1.28082240936564 | 0.957 | 0.934 | 5.6846409 |
| **CRB2** | 4.7085 | -1.50403837308523 | 0.009 | 0.292 | 7.6575901 |
| **RGMB** | 8.7982 | -1.00456252114776 | 0.518 | 0.802 | 1.4308669 |
| **AMOT** | 1.1209 | -1.18075456157811 | 0.126 | 0.475 | 1.8229621 |
| **FN1** | 9.7739 | -1.88897472543469 | 0.921 | 0.984 | 1.5895377 |
| **HPGD** | 2.1740 | -2.18506733914686 | 0.943 | 0.907 | 3.5356259 |
| **LDLR** | 2.7394 | -0.815340798991836 | 0.955 | 0.988 | 4.4551148 |
| **DYRK1B** | 3.5347 | -0.986086489836883 | 0.443 | 0.73 | 5.7486201 |
| **SYNPO** | 1.8463 | -0.935458818989027 | 0.013 | 0.295 | 3.0027226 |
| **LRRC58** | 8.1311 | -0.819478927339649 | 0.378 | 0.715 | 1.3223763 |
| **TUBB6** | 1.4901 | -0.829732873990981 | 0.939 | 0.981 | 2.4234774 |
| **NEFL** | 1.7549 | 0.817598632064844 | 0.343 | 0.021 | 2.8540942 |
| **FAM151A** | 2.1419 | 1.05265205681104 | 0.316 | 0.008 | 3.4834086 |
| **APBB2** | 3.4121 | -0.866422354919234 | 0.318 | 0.663 | 5.5492264 |
| **MYO10** | 3.6682 | -0.828722330494509 | 0.912 | 0.971 | 5.9656289 |
| **NCKAP5L** | 4.3708 | -0.871239493678093 | 0.504 | 0.787 | 7.1083005 |
| **CAPRIN2** | 7.8942 | -1.0144447678029 | 0.474 | 0.77 | 1.2838444 |
| **XKR6** | 2.0628 | -0.838479566131071 | 0.696 | 0.896 | 3.3548051 |
| **VASN** | 2.5360 | -0.843617914752636 | 0.102 | 0.438 | 4.1243399 |
| **CER1** | 5.4674 | -1.48340577089232 | 0.025 | 0.314 | 8.8917878 |
| **NEO1** | 6.0860 | -0.909970045998036 | 0.53 | 0.797 | 9.8976984 |
| **FILIP1** | 5.1695 | -0.86451477214266 | 0.235 | 0.588 | 8.4071624 |
| **NPHP3** | 6.8628 | -0.815657523067963 | 0.39 | 0.718 | 1.1161069 |
| **TET2** | 1.2649 | 0.912133469579749 | 0.68 | 0.337 | 2.0571991 |
| **SMAGP** | 5.6423 | -1.06789449181442 | 0.416 | 0.709 | 9.1760804 |
| **PCDH7** | 1.5187 | 1.17719983459091 | 0.435 | 0.084 | 2.4699434 |
| **PRICKLE1** | 2.5718 | -1.01424418225267 | 0.574 | 0.81 | 4.1826501 |
| **SCUBE3** | 2.5920 | -0.876819838064482 | 0.166 | 0.507 | 4.2155158 |
| **LIX1** | 5.5102 | -1.28877154079619 | 0.004 | 0.259 | 8.9613265 |

| | | | | | |
|---|---|---|---|---|---|
| **GPC3** | 6.8459 | -1.15669059071454 | 0.808 | 0.942 | 1.1133513 |
| **AMH** | 6.8646 | -1.03571197604051 | 0.227 | 0.559 | 1.1164045 |
| **TSHZ3** | 7.9376 | -0.808981711880269 | 0.167 | 0.508 | 1.2908930 |
| **DUSP3** | 1.9626 | -0.804632170514997 | 0.797 | 0.928 | 3.1918094 |
| **SAMD4A** | 2.2485 | -0.836727342950563 | 0.027 | 0.309 | 3.6567626 |
| **FOSL2** | 9.3228 | -0.97705166702592 | 0.052 | 0.355 | 1.5161680 |
| **CPNE2** | 2.8194 | -0.861301620830471 | 0.273 | 0.612 | 4.5852016 |
| **NOTCH1** | 4.4717 | -0.823584657446043 | 0.703 | 0.892 | 7.2723414 |
| **ATP2B4** | 3.8532 | -0.83002530203789 | 0.966 | 0.991 | 6.2664598 |
| **KLF2** | 7.4672 | 0.826375023218467 | 0.581 | 0.195 | 1.2143935 |
| **EPCAM** | 1.2939 | -1.45942423625044 | 0.997 | 0.95 | 2.1043154 |
| **JUN** | 1.8928 | -1.34096164954627 | 0.444 | 0.747 | 3.0783032 |
| **LRP8** | 3.8043 | -0.85939176706284 | 0.61 | 0.84 | 6.1870611 |
| **FGF8** | 1.7392 | -1.44802132981964 | 0.392 | 0.679 | 2.8285568 |
| **FBN1** | 2.3489 | -0.98271256654415 | 0.086 | 0.397 | 3.8200185 |
| **CTIF** | 2.4198 | -0.938589309836784 | 0.278 | 0.604 | 3.9354112 |
| **SUSD3** | 2.4796 | -1.10028076525073 | 0.359 | 0.661 | 4.0325909 |
| **PPP1R9B** | 2.6368 | -0.803491250166795 | 0.503 | 0.77 | 4.2882719 |
| **CNKSR3** | 1.4239 | -0.805938270375855 | 0.152 | 0.484 | 2.3157045 |
| **FJX1** | 2.2833 | -1.76224717131562 | 0.141 | 0.457 | 3.7133596 |
| **HSPA1B** | 3.9937 | 0.866390555269597 | 0.716 | 0.417 | 6.4950314 |
| **DSEL** | 3.6167 | -0.898175820063822 | 0.434 | 0.725 | 5.8818688 |
| **SLC1A3** | 9.1056 | -0.855176764905395 | 0.262 | 0.587 | 1.4808577 |
| **CYP26A1** | 1.2893 | -1.2811328663832 | 0.034 | 0.307 | 2.0968791 |
| **SUN2** | 1.1002 | -0.808657115886463 | 0.133 | 0.456 | 1.7893142 |
| **TIMP4** | 3.4011 | -0.881229187958498 | 0.248 | 0.565 | 5.5313080 |
| **SYVN1** | 5.6708 | 0.813969789582456 | 0.934 | 0.852 | 9.2224365 |
| **POU3F1** | 1.1067 | -0.805380621588992 | 0.107 | 0.42 | 1.7999200 |
| **GALNT10** | 1.6625 | -0.851953312314777 | 0.458 | 0.737 | 2.7038088 |
| **TRNP1** | 2.0771 | -0.856747294938765 | 0.693 | 0.831 | 3.3780778 |
| **CDON** | 5.7312 | -0.984772227068932 | 0.314 | 0.631 | 9.3207045 |
| **GALNT18** | 1.7746 | 0.826422203191139 | 0.544 | 0.202 | 2.8861249 |
| **RAI14** | 3.0537 | -0.878151282088814 | 0.591 | 0.818 | 4.9662921 |
| **BEND4** | 4.3310 | -0.800601937633636 | 0.868 | 0.914 | 7.0435630 |

| | | | | | |
|---|---|---|---|---|---|
| **MYL4** | 1.0157 | -2.00211571173379 | 0.016 | 0.263 | 1.6518425 |
| **GADL1** | 6.3729 | 1.07424682910161 | 0.353 | 0.05 | 1.0364292 |
| **COL1A1** | 1.6706 | -2.56078182400815 | 0.785 | 0.914 | 2.7169497 |
| **GATA2** | 3.1373 | -0.922084965279479 | 0.036 | 0.301 | 5.1022739 |
| **PMP22** | 4.5831 | -1.12966082307006 | 0.279 | 0.585 | 7.4536395 |
| **TRIM22** | 5.1576 | -0.864129193303867 | 0.613 | 0.846 | 8.3878861 |
| **GALNT17** | 4.6106 | -0.803066533241727 | 0.777 | 0.924 | 7.4983714 |
| **BOC** | 4.6797 | -1.81020623772825 | 0.085 | 0.374 | 7.6107241 |
| **PDK1** | 8.5212 | -0.948285951556064 | 0.869 | 0.954 | 1.3858122 |
| **SHF** | 3.1133 | -0.938267486147678 | 0.216 | 0.521 | 5.0631632 |
| **PCNA** | 1.3284 | -0.82849552128972 | 0.978 | 0.988 | 2.1603848 |
| **GADD45G** | 3.8012 | -0.961044788944182 | 0.247 | 0.551 | 6.1820227 |
| **SMIM10L2A** | 6.9149 | -1.03082776185 | 0.485 | 0.73 | 1.1245815 |
| **CTXN1** | 1.1903 | -1.06324958773834 | 0.896 | 0.949 | 1.9358635 |
| **MT-ND3** | 4.6034 | 0.98754740267553 | 0.935 | 0.806 | 7.4866443 |
| **WNT4** | 5.1392 | -2.05176551706655 | 0.016 | 0.25 | 8.3580330 |
| **CACNG4** | 7.4974 | -0.841057516154449 | 0.024 | 0.268 | 1.2193158 |
| **PRPS2** | 7.6622 | -0.868580765022657 | 0.9 | 0.943 | 1.2461189 |
| **HMGB2** | 7.7559 | -1.0157431201302 | 0.996 | 0.981 | 1.2613477 |
| **ANOS1** | 2.4446 | -1.3921011873109 | 0.087 | 0.366 | 3.9757034 |
| **IGFBP3** | 2.2872 | -1.15146623055894 | 0.146 | 0.444 | 3.7196908 |
| **ANKRD1** | 3.7462 | -1.41890673855326 | 0.036 | 0.288 | 6.0924796 |
| **SDC3** | 4.6506 | -1.00493693219048 | 0.597 | 0.806 | 7.5633177 |
| **TMC5** | 8.6428 | -0.957588987314976 | 0.548 | 0.758 | 1.4055819 |
| **CXCL14** | 2.8320 | -1.71712511453724 | 0.408 | 0.678 | 4.6057148 |
| **MED13L** | 4.6582 | -0.84147069541599 | 0.951 | 0.982 | 7.5757557 |
| **TAGLN2** | 2.5900 | -0.965327507028172 | 0.864 | 0.925 | 4.2121716 |
| **TPBG** | 2.9020 | -1.39820217933618 | 0.359 | 0.641 | 4.7196641 |
| **PMEPA1** | 4.1198 | -0.940421665043895 | 0.953 | 0.952 | 6.7000767 |
| **PCBP4** | 2.8559 | -1.18163021033424 | 0.162 | 0.442 | 4.6447118 |
| **HTR1D** | 4.4776 | -0.836070998531683 | 0.034 | 0.27 | 7.2819571 |
| **HAS2** | 2.7853 | -0.890639757364223 | 0.378 | 0.659 | 4.5298402 |
| **WNT1** | 1.5610 | -2.33699553024038 | 0.003 | 0.2 | 2.5386859 |
| **PPL** | 7.7142 | -0.809079938729608 | 0.403 | 0.663 | 1.2545730 |

| | | | | | |
|---|---|---|---|---|---|
| **CDH6** | 1.3994 | -1.62803326822992 | 0.009 | 0.211 | 2.27588302 |
| **HERPUD1** | 5.6213 | 1.02470564047366 | 0.828 | 0.696 | 9.14199782 |
| **PAMR1** | 2.9828 | -1.39789824407965 | 0.094 | 0.35 | 4.85106759 |
| **SDC4** | 6.2821 | -0.822018696115959 | 0.984 | 0.994 | 1.02165917 |
| **ANXA3** | 1.7810 | -1.44342137164634 | 0.914 | 0.918 | 2.89654498 |
| **SPATS2L** | 5.4068 | -1.12259469464879 | 0.176 | 0.452 | 8.79311822 |
| **GADD45A** | 5.4958 | -0.923381800241389 | 0.423 | 0.684 | 8.93789254 |
| **TLE4** | 1.9984 | -1.40499436280098 | 0.554 | 0.758 | 3.25000937 |
| **FAM111B** | 1.6353 | -0.934637656977167 | 0.431 | 0.69 | 2.65951253 |
| **COL5A1** | 9.6225 | -1.6172653991145 | 0.445 | 0.705 | 1.56491210 |
| **IFFO2** | 6.5196 | -0.824562855240609 | 0.372 | 0.636 | 1.06029338 |
| **MSX1** | 1.5503 | -0.847015927317244 | 0.007 | 0.189 | 2.52132170 |
| **BGN** | 6.0692 | -1.12277057339745 | 0.17 | 0.439 | 9.87039964 |
| **PDZD4** | 3.7452 | -0.840922089919645 | 0.379 | 0.629 | 6.09091991 |
| **GREM1** | 8.3924 | -0.975176853721049 | 0.003 | 0.168 | 1.36486473 |
| **TMEM51** | 1.5244 | -0.852341352068515 | 0.7 | 0.821 | 2.47921172 |
| **TBXT** | 3.7332 | -0.909842858907724 | 0.023 | 0.211 | 6.07142627 |
| **CDKN1A** | 1.3261 | -1.09786772976595 | 0.328 | 0.578 | 2.15676151 |
| **B3GNT7** | 1.9244 | -0.833387874493599 | 0.722 | 0.819 | 3.12978078 |
| **ZNF367** | 8.3363 | -0.820826391326846 | 0.356 | 0.584 | 1.35573383 |
| **QPCT** | 3.3500 | -1.00097297791756 | 0.42 | 0.618 | 5.44825642 |
| **ABCC4** | 7.6491 | -0.945927661038485 | 0.641 | 0.806 | 1.24398530 |
| **NLGN4X** | 8.6425 | -1.34197070790617 | 0.354 | 0.56 | 1.40553467 |
| **EFNA1** | 1.0855 | -0.900689941800832 | 0.464 | 0.695 | 1.76536868 |
| **DOK6** | 5.5072 | -1.00814099310421 | 0.171 | 0.402 | 8.95636057 |
| **SNAI2** | 8.4934 | -0.947281727679936 | 0.021 | 0.194 | 1.38128262 |
| **TENM4** | 1.6189 | -0.955974156221506 | 0.583 | 0.764 | 2.63292591 |
| **PIMREG** | 2.3975 | -0.835018781610651 | 0.492 | 0.661 | 3.89913782 |
| **IER3** | 8.9035 | -1.33309917920326 | 0.116 | 0.335 | 1.44798472 |
| **FZD10** | 1.0260 | -0.914303983657535 | 0.536 | 0.715 | 1.66859617 |
| **SDK2** | 1.3501 | -1.10196330023628 | 0.228 | 0.439 | 2.19580548 |
| **VAMP8** | 1.1520 | -0.921366915999554 | 0.999 | 0.988 | 1.87357591 |
| **IER5L** | 1.2606 | -0.843992259703636 | 0.354 | 0.561 | 2.05025800 |
| **NPPB** | 4.8917 | -1.15626919548185 | 0.017 | 0.168 | 7.95547652 |

| | | | | | |
|---|---|---|---|---|---|
| **CDH3** | 4.5724 | -0.815233011530557 | 0.952 | 0.906 | 7.4362438 |
| **ACTA2** | 3.6055 | -1.91294865311717 | 0.012 | 0.15 | 5.8636436 |
| **ADAMTSL4** | 7.4509 | -1.0162086653482 | 0.392 | 0.575 | 1.2117514 |
| **GAP43** | 5.9719 | -0.977292129558987 | 0.22 | 0.437 | 9.7122153 |
| **SDC2** | 5.1364 | -1.31575612314518 | 0.322 | 0.525 | 8.3533663 |
| **MAL2** | 1.0484 | -0.85298218604037 | 0.489 | 0.644 | 1.7051047 |
| **ACTA1** | 7.3282 | -1.06353285599168 | 0.114 | 0.312 | 1.1917938 |
| **IL6** | 4.8799 | -0.846224157898826 | 0.446 | 0.616 | 7.9362272 |
| **NID2** | 6.0226 | -0.822920023200537 | 0.084 | 0.262 | 9.7946330 |
| **CDH13** | 8.7532 | -0.821457773526037 | 0.379 | 0.574 | 1.4235340 |
| **LOXL1** | 4.3941 | -1.19226317429636 | 0.648 | 0.771 | 7.1462748 |
| **INSM1** | 8.9110 | -1.07337008553207 | 0.409 | 0.552 | 1.4492094 |
| **MN1** | 1.0326 | -0.893134403969302 | 0.497 | 0.643 | 1.6793359 |
| **OLIG3** | 1.3770 | -0.864422811915014 | 0.113 | 0.284 | 2.2394657 |
| **NR2F1** | 4.3194 | -0.901043193208321 | 0.02 | 0.137 | 7.0247854 |
| **COL3A1** | 6.2316 | -1.97730528755974 | 0.037 | 0.16 | 1.0134476 |
| **LEFTY2** | 1.2415 | -1.17671076463053 | 0.882 | 0.932 | 2.0191986 |
| **FOXC2** | 7.0603 | -0.802708488183685 | 0.027 | 0.125 | 1.1482269 |
| **CNTFR** | 1.5383 | -1.16342602474689 | 0.196 | 0.323 | 2.5018971 |
| **LEFTY1** | 2.7385 | -1.27913409274314 | 0.861 | 0.86 | 4.4537745 |
| **POMC** | 2.4833 | -1.1039623759024 | 0.836 | 0.792 | 4.0387394 |
| **MIXL1** | 3.6126 | -1.26305885417123 | 0.521 | 0.602 | 5.8752904 |
| **DLK1** | 2.1273 | -1.59575898742696 | 0.557 | 0.341 | 3.4596528 |
| **ANXA1** | 7.3510 | -1.57515464122207 | 0.135 | 0.213 | 1.1954957 |
| **SFRP1** | 5.7155 | -1.90221924755623 | 0.569 | 0.556 | 9.2952144 |
| **RHOB** | 9.7323 | -0.807439646582517 | 0.997 | 0.985 | 1.5827759 |
| **NRP2** | 1.0877 | -0.880156218678582 | 0.42 | 0.454 | 1.7690483 |
| **CNTNAP2** | 1.4774 | -1.20182820787174 | 0.82 | 0.778 | 0.0024027 |
| **SOX21** | 0.0014 | -0.916758424669969 | 0.328 | 0.248 | 1 |
| **L1CAM** | 0.0016 | -0.814692954589709 | 0.414 | 0.42 | 1 |
| **PRSS23** | 0.6215 | -1.54221104361813 | 0.719 | 0.699 | 1 |

Supplemental Table S2_Sheet_4

| | p_val | avg_log2FC | pct.1 | pct.2 | p_val_adj |
|---|---|---|---|---|---|
| AGRN | 0 | -1.30016502842085 | 0.952 | 0.999 | 0 |
| MXRA8 | 0 | -1.64919338607399 | 0.46 | 0.959 | 0 |
| ATAD3B | 0 | 1.97035037359049 | 0.999 | 0.917 | 0 |
| ERRFI1 | 0 | 1.59082917962405 | 0.992 | 0.756 | 0 |
| CTNNBIP1 | 0 | -1.12679962517736 | 0.884 | 0.997 | 0 |
| DFFA | 0 | 1.31434801089722 | 1 | 0.997 | 0 |
| DRAXIN | 0 | -2.06510889365081 | 0.579 | 0.95 | 0 |
| TNFRSF8 | 0 | 1.61685719466593 | 0.756 | 0.029 | 0 |
| EPHA2 | 0 | -2.51531780159969 | 0.419 | 0.991 | 0 |
| MFAP2 | 0 | -1.81915810278375 | 0.881 | 1 | 0 |
| ECE1 | 0 | -1.58524722097763 | 0.803 | 0.997 | 0 |
| LUZP1 | 0 | -1.10478896749598 | 0.897 | 0.994 | 0 |
| ID3 | 0 | -3.65836806401544 | 0.471 | 0.997 | 0 |
| FUCA1 | 0 | 1.69121075051719 | 0.991 | 0.722 | 0 |
| CNR2 | 0 | 1.93523401032758 | 0.799 | 0.015 | 0 |
| LDLRAP1 | 0 | 1.23053283335904 | 0.996 | 0.884 | 0 |
| MAN1C1 | 0 | 2.34370060033688 | 0.999 | 0.682 | 0 |
| ARID1A | 0 | -1.23792327232478 | 0.957 | 1 | 0 |
| MARCKSL1 | 0 | -1.26515742808683 | 1 | 1 | 0 |
| KIAA1522 | 0 | -1.50430519197235 | 0.93 | 0.991 | 0 |
| ZNF362 | 0 | -1.41098723195494 | 0.785 | 0.995 | 0 |
| PHC2 | 0 | -1.83712720501058 | 0.075 | 0.736 | 0 |
| MAP7D1 | 0 | -2.10138791532823 | 0.388 | 0.982 | 0 |
| MANEAL | 0 | 1.21314024537658 | 0.981 | 0.769 | 0 |
| AKIRIN1 | 0 | 0.989026995458448 | 1 | 0.999 | 0 |
| CLDN19 | 0 | -1.4999291968824 | 0.075 | 0.722 | 0 |
| SVBP | 0 | -1.05802131489563 | 0.909 | 0.997 | 0 |
| HPDL | 0 | 1.93618905091356 | 0.87 | 0.134 | 0 |
| PRDX1 | 0 | 1.72032591360544 | 1 | 0.999 | 0 |
| UQCRH | 0 | 0.928504113596582 | 1 | 0.999 | 0 |
| OSBPL9 | 0 | -0.946015147642365 | 0.983 | 1 | 0 |
| ZYG11A | 0 | 1.68321934304112 | 0.995 | 0.722 | 0 |

| | | | | | |
|---|---|---|---|---|---|
| **DHCR24** | 0 | -1.60577754096181 | 1 | 1 | 0 |
| **TACSTD2** | 0 | 0.889142685385987 | 0.517 | 0.009 | 0 |
| **L1TD1** | 0 | 1.02960326112797 | 1 | 0.998 | 0 |
| **AK4** | 0 | 2.44959461702934 | 1 | 0.929 | 0 |
| **WLS** | 0 | -4.32620869912146 | 0.419 | 0.995 | 0 |
| **CTH** | 0 | 2.36862972129118 | 0.996 | 0.698 | 0 |
| **LMO4** | 0 | 1.52436875108242 | 0.999 | 0.949 | 0 |
| **BRDT** | 0 | 3.21114610299655 | 0.997 | 0.176 | 0 |
| **TMED5** | 0 | 1.21539745860641 | 0.998 | 0.925 | 0 |
| **CNN3** | 0 | -0.879588705304167 | 1 | 1 | 0 |
| **AGL** | 0 | 2.7134222547573 | 1 | 0.922 | 0 |
| **HENMT1** | 0 | 2.05476257948449 | 0.894 | 0.033 | 0 |
| **TAF13** | 0 | 2.76535416700011 | 0.999 | 0.79 | 0 |
| **OLFML3** | 0 | -3.78625497969717 | 0.68 | 0.997 | 0 |
| **ANKRD35** | 0 | 1.35812805699037 | 0.655 | 0.039 | 0 |
| **SV2A** | 0 | -1.58324299846307 | 0.29 | 0.908 | 0 |
| **PLEKHO1** | 0 | -2.10275402220068 | 0.41 | 0.978 | 0 |
| **HORMAD1** | 0 | 2.28079664080427 | 0.958 | 0.094 | 0 |
| **S100A10** | 0 | 1.41255727789547 | 1 | 0.998 | 0 |
| **S100A11** | 0 | 1.37750202066694 | 1 | 1 | 0 |
| **RAB13** | 0 | -1.00545209655733 | 0.973 | 1 | 0 |
| **IL6R** | 0 | 1.43361314914648 | 0.839 | 0.213 | 0 |
| **MEX3A** | 0 | -1.9208394117741 | 0.546 | 0.991 | 0 |
| **SLC25A44** | 0 | 1.20529140808621 | 0.998 | 0.937 | 0 |
| **NAXE** | 0 | -1.28527080683001 | 0.799 | 0.996 | 0 |
| **NES** | 0 | -1.79128617093699 | 0.994 | 1 | 0 |
| **CRABP2** | 0 | -3.14219343121753 | 0.891 | 1 | 0 |
| **HDGF** | 0 | -1.50460290971328 | 0.939 | 0.999 | 0 |
| **DUSP23** | 0 | 1.76908861147174 | 0.87 | 0.031 | 0 |
| **PEA15** | 0 | -1.39616398702879 | 0.932 | 1 | 0 |
| **F11R** | 0 | 1.34663328122016 | 1 | 0.995 | 0 |
| **TSTD1** | 0 | 1.96465502032661 | 0.884 | 0.026 | 0 |
| **RGS5** | 0 | -3.31922622282536 | 0.158 | 0.925 | 0 |
| **MAEL** | 0 | 1.40419439128055 | 0.593 | 0.016 | 0 |

| | | | | | |
|---|---|---|---|---|---|
| **RO60** | 0 | -1.06612955631048 | 0.91 | 0.997 | 0 |
| **BTG2** | 0 | -1.7764524582906 | 0.206 | 0.802 | 0 |
| **NUCKS1** | 0 | -1.21061111027484 | 1 | 1 | 0 |
| **CR1L** | 0 | 2.7104578258575 | 0.994 | 0.284 | 0 |
| **IRF6** | 0 | 1.00921716415247 | 0.642 | 0.033 | 0 |
| **PTPN14** | 0 | -1.22889879425283 | 0.907 | 0.998 | 0 |
| **CAPN2** | 0 | -2.55731948085589 | 0.483 | 0.995 | 0 |
| **CNIH4** | 0 | 1.36871227850044 | 0.984 | 0.692 | 0 |
| **ENAH** | 0 | -1.09755564471705 | 0.995 | 1 | 0 |
| **LGALS8** | 0 | 1.01789977485795 | 0.59 | 0.017 | 0 |
| **ID2** | 0 | -2.08515801465008 | 0.231 | 0.923 | 0 |
| **IAH1** | 0 | 1.98958813476099 | 0.864 | 0.029 | 0 |
| **EFR3B** | 0 | 1.76579375927818 | 0.963 | 0.653 | 0 |
| **SLC35F6** | 0 | 1.00085908521352 | 0.999 | 0.97 | 0 |
| **KRTCAP3** | 0 | 1.93611204762882 | 0.938 | 0.188 | 0 |
| **PKDCC** | 0 | -2.21762532397594 | 0.338 | 0.936 | 0 |
| **EML4** | 0 | -1.00360337967489 | 0.972 | 1 | 0 |
| **RTN4** | 0 | -0.968090353512446 | 1 | 1 | 0 |
| **CCDC88A** | 0 | -1.40025618224584 | 0.781 | 0.998 | 0 |
| **B3GNT2** | 0 | -1.49814758451423 | 0.859 | 0.989 | 0 |
| **UGP2** | 0 | 1.25881062229709 | 0.997 | 0.932 | 0 |
| **CEP68** | 0 | -1.3508648844827 | 0.514 | 0.957 | 0 |
| **ANTXR1** | 0 | -1.5327283137932 | 0.261 | 0.886 | 0 |
| **GFPT1** | 0 | 1.64351158566684 | 1 | 0.996 | 0 |
| **SNRPG** | 0 | 0.914090755283733 | 1 | 0.97 | 0 |
| **MTHFD2** | 0 | 0.970858800861914 | 1 | 1 | 0 |
| **VAMP8** | 0 | -1.29588652452493 | 0.997 | 1 | 0 |
| **COX5B** | 0 | 0.921591087402971 | 1 | 0.999 | 0 |
| **MAP4K4** | 0 | -1.1462858613089 | 0.995 | 1 | 0 |
| **TFCP2L1** | 0 | 1.78760797047745 | 0.951 | 0.385 | 0 |
| **TSN** | 0 | -1.08261699757314 | 0.993 | 1 | 0 |
| **BIN1** | 0 | -2.23688963902179 | 0.427 | 0.989 | 0 |
| **PLEKHB2** | 0 | 0.993856576138783 | 1 | 0.999 | 0 |
| **KIF5C** | 0 | -1.47938619322878 | 0.848 | 1 | 0 |

| | | | | | |
|---|---|---|---|---|---|
| **NMI** | 0 | 2.03642371871055 | 0.978 | 0.354 | 0 |
| **GALNT3** | 0 | 1.256981903031 | 0.642 | 0.008 | 0 |
| **SP5** | 0 | -2.27267116899907 | 0.82 | 1 | 0 |
| **GAD1** | 0 | -2.12404311686334 | 0.33 | 0.965 | 0 |
| **UBE2E3** | 0 | -1.20976655057916 | 0.955 | 1 | 0 |
| **NCKAP1** | 0 | -0.85798145360028 | 1 | 1 | 0 |
| **SLC39A10** | 0 | 0.964360853129321 | 1 | 0.998 | 0 |
| **CFLAR** | 0 | 1.41908915951731 | 0.991 | 0.762 | 0 |
| **FZD7** | 0 | -2.77635927673727 | 0.389 | 0.96 | 0 |
| **ICA1L** | 0 | 1.9788141299484 | 0.895 | 0.081 | 0 |
| **FZD5** | 0 | 2.66710178123979 | 1 | 0.867 | 0 |
| **CRYGD** | 0 | 1.86425231556731 | 0.819 | 0.019 | 0 |
| **IDH1** | 0 | 1.11012504458276 | 1 | 0.993 | 0 |
| **CPS1** | 0 | -1.2623982886449 | 0.594 | 0.972 | 0 |
| **TMBIM1** | 0 | 1.9238551438429 | 0.959 | 0.39 | 0 |
| **OBSL1** | 0 | -1.75345512696825 | 0.691 | 0.997 | 0 |
| **SERPINE2** | 0 | -2.92377570674771 | 0.826 | 0.997 | 0 |
| **FBXO36** | 0 | 1.77992752516333 | 0.99 | 0.676 | 0 |
| **ALPG** | 0 | 3.9277847901576 | 0.987 | 0.407 | 0 |
| **ARL4C** | 0 | -1.62344532156289 | 0.193 | 0.805 | 0 |
| **SLC6A6** | 0 | 1.50125951461736 | 0.998 | 0.91 | 0 |
| **RAB5A** | 0 | 1.25043399344489 | 1 | 0.94 | 0 |
| **EOMES** | 0 | -3.06214519553704 | 0.165 | 0.812 | 0 |
| **CTNNB1** | 0 | -1.13874427790407 | 0.999 | 1 | 0 |
| **ZNF502** | 0 | 1.08382512232263 | 0.624 | 0.008 | 0 |
| **SHISA5** | 0 | 1.13061052667488 | 1 | 0.999 | 0 |
| **NT5DC2** | 0 | -1.40198218022166 | 1 | 1 | 0 |
| **WNT5A** | 0 | -2.2372225149425 | 0.215 | 0.901 | 0 |
| **PTPRG** | 0 | -1.55232061700441 | 0.761 | 0.993 | 0 |
| **ZNF717** | 0 | 0.913549034249426 | 0.587 | 0.003 | 0 |
| **DCBLD2** | 0 | -1.437372471665 | 0.791 | 0.998 | 0 |
| **TBC1D23** | 0 | 1.66818553367888 | 0.999 | 0.956 | 0 |
| **NIT2** | 0 | 1.24405339256089 | 0.941 | 0.529 | 0 |
| **TOMM70** | 0 | 0.910736599854601 | 1 | 0.993 | 0 |

| | | | | | |
|---|---|---|---|---|---|
| **ALCAM** | 0 | -1.8071024178027 | 0.158 | 0.781 | 0 |
| **DPPA2** | 0 | 3.26003969997374 | 0.999 | 0.426 | 0 |
| **DPPA4** | 0 | 1.04980785838879 | 1 | 1 | 0 |
| **NECTIN3** | 0 | -1.55547242673697 | 0.325 | 0.907 | 0 |
| **ATG3** | 0 | 2.40682378163017 | 1 | 0.996 | 0 |
| **CCDC80** | 0 | 2.6243030135239 | 0.993 | 0.614 | 0 |
| **FSTL1** | 0 | -2.36148008564323 | 0.783 | 0.999 | 0 |
| **HCLS1** | 0 | 2.16821506517956 | 0.897 | 0.093 | 0 |
| **MGLL** | 0 | 2.18687308501754 | 0.849 | 0.042 | 0 |
| **H1FX** | 0 | -0.97335422592571 | 0.999 | 1 | 0 |
| **AMOTL2** | 0 | -1.3491297761199 | 0.98 | 0.999 | 0 |
| **RBP1** | 0 | -2.75682937926555 | 0.562 | 0.994 | 0 |
| **SPSB4** | 0 | -2.22204213527463 | 0.208 | 0.938 | 0 |
| **SERP1** | 0 | 0.869959579834696 | 1 | 1 | 0 |
| **LRRC34** | 0 | 1.45818431937819 | 0.77 | 0.003 | 0 |
| **NCEH1** | 0 | 1.27406964303098 | 0.768 | 0.152 | 0 |
| **GNB4** | 0 | -2.66977748632131 | 0.77 | 0.998 | 0 |
| **LIPH** | 0 | 1.44846357132848 | 0.698 | 0.032 | 0 |
| **LPP** | 0 | -1.56405155049745 | 0.587 | 0.966 | 0 |
| **HES1** | 0 | -2.30016153268723 | 0.202 | 0.955 | 0 |
| **ZNF595** | 0 | 1.97679897828399 | 0.998 | 0.686 | 0 |
| **S100P** | 0 | 2.65357757867638 | 0.872 | 0.025 | 0 |
| **PROM1** | 0 | 1.4936573490445 | 0.984 | 0.752 | 0 |
| **UCHL1** | 0 | -1.80887717666907 | 0.823 | 0.997 | 0 |
| **COX7B2** | 0 | 1.2620552774298 | 0.578 | 0.004 | 0 |
| **REST** | 0 | 1.054218889732 | 1 | 1 | 0 |
| **GRSF1** | 0 | 0.827109105093208 | 1 | 0.997 | 0 |
| **MAPK10** | 0 | -2.23202239784987 | 0.061 | 0.787 | 0 |
| **GRID2** | 0 | -1.85208986076123 | 0.143 | 0.765 | 0 |
| **DDIT4L** | 0 | 1.22495429212068 | 0.644 | 0.014 | 0 |
| **PRSS12** | 0 | 2.90023248118004 | 0.993 | 0.373 | 0 |
| **NOCT** | 0 | 1.57170831458836 | 0.996 | 0.853 | 0 |
| **TRIM60** | 0 | 2.08628761076969 | 0.916 | 0.041 | 0 |
| **TRIM75P** | 0 | 1.60487801150164 | 0.709 | 0.023 | 0 |

| | | | | | |
|---|---|---|---|---|---|
| **FAT1** | 0 | -1.790612198384 | 0.786 | 0.998 | 0 |
| **ZFP42** | 0 | 4.37430604706338 | 1 | 0.307 | 0 |
| **TRIML2** | 0 | 1.22310001583075 | 0.996 | 0.749 | 0 |
| **IRX1** | 0 | -1.75977680860111 | 0.07 | 0.737 | 0 |
| **PDZD2** | 0 | 1.43593023769951 | 0.828 | 0.262 | 0 |
| **SELENOP** | 0 | -1.69466187606828 | 0.209 | 0.851 | 0 |
| **CCL28** | 0 | 0.936851083332126 | 0.557 | 0.005 | 0 |
| **FST** | 0 | -2.80174863127138 | 0.148 | 0.743 | 0 |
| **PLPP1** | 0 | 1.83599298983826 | 0.999 | 0.858 | 0 |
| **MAP1B** | 0 | -1.89690084130043 | 0.937 | 1 | 0 |
| **VCAN** | 0 | -1.68301515928258 | 0.883 | 0.999 | 0 |
| **PAM** | 0 | -1.78169146088517 | 0.16 | 0.884 | 0 |
| **EFNA5** | 0 | -1.42488297377155 | 0.323 | 0.875 | 0 |
| **MCC** | 0 | -1.71824333118977 | 0.258 | 0.888 | 0 |
| **TMED7** | 0 | 0.973239872983032 | 1 | 1 | 0 |
| **ZNF608** | 0 | -2.29707317078179 | 0.263 | 0.972 | 0 |
| **HINT1** | 0 | 0.932597308586738 | 1 | 0.982 | 0 |
| **WNT8A** | 0 | -2.33509218681366 | 0.375 | 0.906 | 0 |
| **CXXC5** | 0 | -1.8068324980541 | 0.874 | 0.999 | 0 |
| **DND1** | 0 | 2.4652509039331 | 0.857 | 0.09 | 0 |
| **PCDHB2** | 0 | -1.67465633256995 | 0.265 | 0.893 | 0 |
| **PCDHGB7** | 0 | -1.834298005264 | 0.12 | 0.832 | 0 |
| **DIAPH1** | 0 | -0.857498494267376 | 0.998 | 1 | 0 |
| **LARS** | 0 | 0.992476442173607 | 1 | 1 | 0 |
| **DPYSL3** | 0 | -2.01633280078866 | 0.713 | 0.99 | 0 |
| **SFXN1** | 0 | -1.26660343285988 | 0.953 | 1 | 0 |
| **FGFR4** | 0 | 1.43113045644045 | 0.995 | 0.849 | 0 |
| **DBN1** | 0 | -1.39260229173677 | 0.986 | 1 | 0 |
| **PDLIM7** | 0 | -1.34806272637217 | 0.955 | 0.999 | 0 |
| **CANX** | 0 | 1.20486462972126 | 1 | 0.998 | 0 |
| **GFPT2** | 0 | -1.54838064914628 | 0.592 | 0.983 | 0 |
| **SERPINB1** | 0 | 1.55821786320506 | 0.784 | 0.044 | 0 |
| **SERPINB6** | 0 | 2.05940502434728 | 0.999 | 0.792 | 0 |
| **TUBB2A** | 0 | -1.94553572776849 | 0.957 | 0.999 | 0 |

| | | | | | |
|---|---|---|---|---|---|
| **TUBB2B** | 0 | -2.35447484743046 | 1 | 1 | 0 |
| **SOX4** | 0 | 1.05733065611374 | 1 | 0.993 | 0 |
| **HDGFL1** | 0 | 1.17119938604072 | 0.616 | 0.005 | 0 |
| **MRS2** | 0 | 1.15739319002668 | 1 | 0.998 | 0 |
| **HIST1H1A** | 0 | 5.33931990081991 | 0.997 | 0.396 | 0 |
| **HIST1H2BB** | 0 | 2.39232924285136 | 0.716 | 0.09 | 0 |
| **ZFP57** | 0 | 2.55931841932209 | 0.995 | 0.274 | 0 |
| **TRIM26** | 0 | 1.71618780860516 | 0.993 | 0.823 | 0 |
| **FLOT1** | 0 | -1.3345786823947 | 0.945 | 1 | 0 |
| **DDAH2** | 0 | -1.75565371314439 | 0.777 | 0.997 | 0 |
| **PBX2** | 0 | -0.954921537301106 | 0.997 | 1 | 0 |
| **CUTA** | 0 | -0.824009517659952 | 0.999 | 1 | 0 |
| **BAK1** | 0 | 0.981692505422305 | 1 | 0.997 | 0 |
| **ILRUN** | 0 | 1.8706632714435 | 1 | 0.994 | 0 |
| **SNRPC** | 0 | 0.901563126563896 | 1 | 1 | 0 |
| **ZFAND3** | 0 | -1.03230322882326 | 0.956 | 0.999 | 0 |
| **TOMM6** | 0 | 0.963542588977568 | 0.998 | 0.957 | 0 |
| **PTK7** | 0 | -1.34327078561955 | 0.979 | 1 | 0 |
| **ADGRB3** | 0 | 1.14214341511171 | 0.583 | 0.002 | 0 |
| **DPPA5** | 0 | 6.20853920348064 | 1 | 0.547 | 0 |
| **KHDC3L** | 0 | 5.385909363511 | 1 | 0.395 | 0 |
| **OOEP** | 0 | 1.75666458429397 | 0.785 | 0.006 | 0 |
| **DDX43** | 0 | 1.9734208718529 | 0.861 | 0.017 | 0 |
| **CGAS** | 0 | 2.02143691842039 | 0.852 | 0.045 | 0 |
| **IRAK1BP1** | 0 | 1.56686325376672 | 0.776 | 0.009 | 0 |
| **CCNC** | 0 | 0.905528824549692 | 1 | 0.993 | 0 |
| **LIN28B** | 0 | 1.54550551940735 | 1 | 0.995 | 0 |
| **ATG5** | 0 | 1.06637107455574 | 0.996 | 0.949 | 0 |
| **CRYBG1** | 0 | 1.11325750471439 | 0.609 | 0.04 | 0 |
| **SEC63** | 0 | 1.24901718514017 | 1 | 0.996 | 0 |
| **WASF1** | 0 | -1.88938396417246 | 0.519 | 0.986 | 0 |
| **GTF3C6** | 0 | 1.28282955938681 | 0.966 | 0.659 | 0 |
| **SLC16A10** | 0 | 2.47537412441667 | 0.969 | 0.256 | 0 |
| **MARCKS** | 0 | -3.11900409581688 | 0.408 | 0.999 | 0 |

| | | | | | |
|---|---|---|---|---|---|
| **FAM162B** | 0 | 1.4448122223715 | 0.728 | 0.023 | 0 |
| **PKIB** | 0 | 1.54199832003748 | 0.688 | 0.049 | 0 |
| **ECHDC1** | 0 | -1.98075970551304 | 0.141 | 0.943 | 0 |
| **ARHGAP18** | 0 | 1.32526286937755 | 0.794 | 0.202 | 0 |
| **SGK1** | 0 | 1.82716876124142 | 0.988 | 0.649 | 0 |
| **CITED2** | 0 | -1.78424548442864 | 0.711 | 0.99 | 0 |
| **FUCA2** | 0 | -1.4212642808265 | 0.694 | 0.949 | 0 |
| **AKAP12** | 0 | 1.24272282516521 | 1 | 0.997 | 0 |
| **RMND1** | 0 | 1.35831025058869 | 0.997 | 0.909 | 0 |
| **ARMT1** | 0 | 1.95749401502734 | 1 | 0.992 | 0 |
| **TCP1** | 0 | 1.73846872090161 | 1 | 0.948 | 0 |
| **IGF2R** | 0 | -1.33249409015806 | 0.678 | 0.982 | 0 |
| **FAM20C** | 0 | -1.63942811364441 | 0.297 | 0.9 | 0 |
| **MPP6** | 0 | -1.16251007992383 | 0.965 | 0.999 | 0 |
| **CBX3** | 0 | 1.197093855331 | 0.997 | 0.94 | 0 |
| **SKAP2** | 0 | -1.4288371726921 | 0.069 | 0.715 | 0 |
| **HOXA1** | 0 | -2.10730819247298 | 0.241 | 0.876 | 0 |
| **HERPUD2** | 0 | -1.39218641867909 | 0.423 | 0.929 | 0 |
| **MYL7** | 0 | -2.99284495689914 | 0.143 | 0.816 | 0 |
| **HUS1** | 0 | 1.44755309320292 | 0.999 | 0.969 | 0 |
| **SUN3** | 0 | 1.51367650574496 | 0.707 | 0.008 | 0 |
| **UPP1** | 0 | 3.01045871788453 | 1 | 0.891 | 0 |
| **SEC61G** | 0 | 1.18690527386897 | 1 | 0.999 | 0 |
| **CHCHD2** | 0 | 1.55150800872343 | 1 | 1 | 0 |
| **ZNF727** | 0 | 1.44779020778369 | 0.761 | 0.003 | 0 |
| **ZNF736** | 0 | 0.986162737178773 | 0.617 | 0.004 | 0 |
| **LIMK1** | 0 | -1.08955037076183 | 0.972 | 1 | 0 |
| **HIP1** | 0 | -1.23774535689283 | 0.678 | 0.975 | 0 |
| **CDK6** | 0 | -1.77693849755416 | 0.242 | 0.933 | 0 |
| **ARPC1B** | 0 | 1.39980417568554 | 1 | 0.995 | 0 |
| **STAG3** | 0 | 1.29302055984423 | 0.667 | 0.021 | 0 |
| **COL26A1** | 0 | -2.76878583593082 | 0.486 | 0.988 | 0 |
| **CUX1** | 0 | -1.43951795175676 | 0.741 | 0.988 | 0 |
| **AASS** | 0 | 1.70856619959341 | 0.978 | 0.806 | 0 |

| | | | | | |
|---|---|---|---|---|---|
| **HYAL4** | 0 | 1.95943095469018 | 0.881 | 0.02 | 0 |
| **FLNC** | 0 | -1.85863155124975 | 0.758 | 0.994 | 0 |
| **PODXL** | 0 | -3.32118854022306 | 0.771 | 0.976 | 0 |
| **AKR1B1** | 0 | -1.09422149149277 | 0.984 | 1 | 0 |
| **HIPK2** | 0 | -1.40839123057772 | 0.47 | 0.919 | 0 |
| **TCAF1** | 0 | -2.21381898465572 | 0.368 | 0.989 | 0 |
| **ZNF398** | 0 | 1.39203703478696 | 0.985 | 0.761 | 0 |
| **AGPAT5** | 0 | -1.0968179483591 | 0.944 | 1 | 0 |
| **PPP1R3B** | 0 | -2.75389626736773 | 0.673 | 0.999 | 0 |
| **ZDHHC2** | 0 | -1.71545516873939 | 0.177 | 0.835 | 0 |
| **LOXL2** | 0 | -1.43818891383684 | 0.762 | 0.993 | 0 |
| **DPYSL2** | 0 | -2.28686698580121 | 0.859 | 1 | 0 |
| **CLU** | 0 | -1.94559265335943 | 0.783 | 0.995 | 0 |
| **NRG1** | 0 | 1.37742763270702 | 0.632 | 0.053 | 0 |
| **ASH2L** | 0 | 1.19651793825599 | 0.998 | 0.988 | 0 |
| **FGFR1** | 0 | -1.31156305317386 | 0.991 | 1 | 0 |
| **TGS1** | 0 | 0.892792286151182 | 1 | 0.994 | 0 |
| **NSMAF** | 0 | 1.36541602563358 | 0.998 | 0.907 | 0 |
| **CHD7** | 0 | -2.56924800256214 | 0.482 | 0.991 | 0 |
| **MTFR1** | 0 | 1.27090934809525 | 0.992 | 0.872 | 0 |
| **SGK3** | 0 | 1.27480477818895 | 0.959 | 0.659 | 0 |
| **PRDM14** | 0 | 1.73039585212506 | 0.995 | 0.765 | 0 |
| **TERF1** | 0 | 1.63369729228222 | 0.995 | 0.883 | 0 |
| **FABP5** | 0 | 2.88741373948383 | 0.997 | 0.517 | 0 |
| **TMEM64** | 0 | -2.05413205208714 | 0.75 | 0.988 | 0 |
| **CTHRC1** | 0 | 1.7512382087303 | 0.807 | 0.034 | 0 |
| **ANGPT1** | 0 | 1.39258526820756 | 0.778 | 0.109 | 0 |
| **EIF3H** | 0 | 0.834061097560422 | 1 | 0.998 | 0 |
| **DEPTOR** | 0 | 2.26869040258747 | 0.988 | 0.366 | 0 |
| **ZHX2** | 0 | -1.92773116696975 | 0.154 | 0.912 | 0 |
| **SMARCA2** | 0 | 2.23172274624418 | 0.998 | 0.574 | 0 |
| **GLDC** | 0 | 1.98355339093987 | 0.995 | 0.647 | 0 |
| **CAAP1** | 0 | 1.53059570187619 | 0.975 | 0.635 | 0 |
| **UBAP2** | 0 | 1.05486312967324 | 0.997 | 0.967 | 0 |

| | | | | | |
|---|---|---|---|---|---|
| **TPM2** | 0 | -1.47224428075078 | 0.993 | 1 | 0 |
| **GLIPR2** | 0 | -2.5908096790259 | 0.45 | 0.942 | 0 |
| **IGFBPL1** | 0 | -2.23737222904976 | 0.257 | 0.927 | 0 |
| **CARNMT1** | 0 | 1.13662653875479 | 0.995 | 0.889 | 0 |
| **PCSK5** | 0 | 2.3425979055687 | 0.965 | 0.189 | 0 |
| **PRUNE2** | 0 | 1.71807693696998 | 0.984 | 0.585 | 0 |
| **C9orf64** | 0 | 1.47735581329507 | 0.771 | 0.012 | 0 |
| **BARX1** | 0 | 1.54159142735396 | 0.76 | 0.131 | 0 |
| **FBP1** | 0 | 2.3367504310062 | 0.807 | 0.063 | 0 |
| **ANP32B** | 0 | -1.11274110612552 | 0.972 | 1 | 0 |
| **TGFBR1** | 0 | -1.08320297342903 | 0.978 | 1 | 0 |
| **ABCA1** | 0 | 1.5158954383328 | 0.959 | 0.633 | 0 |
| **KLF4** | 0 | 2.13119714977275 | 0.89 | 0.353 | 0 |
| **ECPAS** | 0 | 0.924258562688872 | 0.999 | 0.996 | 0 |
| **UGCG** | 0 | 1.26324763304742 | 1 | 0.979 | 0 |
| **TNC** | 0 | -2.86211085061977 | 0.13 | 0.834 | 0 |
| **GSN** | 0 | -2.1162626918647 | 0.242 | 0.956 | 0 |
| **STOM** | 0 | 2.19045200005234 | 0.993 | 0.778 | 0 |
| **HSPA5** | 0 | 2.25331196119299 | 1 | 0.912 | 0 |
| **PRRX2** | 0 | 1.3095459324765 | 0.843 | 0.27 | 0 |
| **VAV2** | 0 | 1.22487998699896 | 0.987 | 0.861 | 0 |
| **ECHDC3** | 0 | 1.45819530342088 | 0.755 | 0.012 | 0 |
| **OLAH** | 0 | 2.31110368058125 | 0.804 | 0.032 | 0 |
| **VIM** | 0 | -3.68740653992799 | 0.591 | 0.997 | 0 |
| **NEBL** | 0 | -1.83923934003614 | 0.171 | 0.897 | 0 |
| **REEP3** | 0 | -1.20794899744705 | 0.606 | 0.963 | 0 |
| **SLC25A16** | 0 | 2.3395587624691 | 1 | 0.714 | 0 |
| **TET1** | 0 | 1.04363264604887 | 1 | 0.997 | 0 |
| **DDX21** | 0 | 0.885900671043639 | 1 | 1 | 0 |
| **VPS26A** | 0 | 0.944958170823295 | 0.997 | 0.955 | 0 |
| **HK1** | 0 | -1.05614391320364 | 0.992 | 1 | 0 |
| **H2AFY2** | 0 | -2.5498249182502 | 0.682 | 0.997 | 0 |
| **PDLIM1** | 0 | 1.06196010359899 | 1 | 0.96 | 0 |
| **PIK3AP1** | 0 | 1.35310569835977 | 0.741 | 0.051 | 0 |

| | | | | | |
|---|---|---|---|---|---|
| **FRAT2** | 0 | 2.09912941950761 | 1 | 0.984 | 0 |
| **NFKB2** | 0 | 1.72518471023995 | 0.986 | 0.834 | 0 |
| **C10orf95** | 0 | 1.54543373792474 | 0.706 | 0.045 | 0 |
| **PCGF6** | 0 | 1.22223277167874 | 0.969 | 0.638 | 0 |
| **GSTO2** | 0 | 1.40924632481918 | 0.775 | 0.055 | 0 |
| **OAT** | 0 | 1.22624749753891 | 0.912 | 0.488 | 0 |
| **UTF1** | 0 | 4.15017358438319 | 0.996 | 0.361 | 0 |
| **VENTX** | 0 | 1.87401858209633 | 0.988 | 0.482 | 0 |
| **PKP3** | 0 | 1.46526702280203 | 0.991 | 0.647 | 0 |
| **SIGIRR** | 0 | 1.32577430077982 | 0.972 | 0.652 | 0 |
| **TALDO1** | 0 | 1.25871345411196 | 1 | 0.997 | 0 |
| **CD81** | 0 | 1.05314414353583 | 1 | 1 | 0 |
| **TRIM6** | 0 | 1.81660639844839 | 0.92 | 0.211 | 0 |
| **ARFIP2** | 0 | 1.19717332545023 | 0.999 | 0.982 | 0 |
| **TMEM9B** | 0 | 1.05908372309862 | 0.986 | 0.828 | 0 |
| **TMEM41B** | 0 | 1.55379052108467 | 1 | 0.999 | 0 |
| **EIF4G2** | 0 | 0.881538481914607 | 1 | 1 | 0 |
| **3,00 DKK** | 0 | -1.62293984681499 | 0.474 | 0.955 | 0 |
| **RRAS2** | 0 | 1.1936707254268 | 1 | 0.955 | 0 |
| **LGR4** | 0 | 1.66844245734376 | 1 | 0.984 | 0 |
| **LIN7C** | 0 | 1.0309941106326 | 1 | 0.996 | 0 |
| **RCN1** | 0 | 1.32125112932053 | 1 | 0.995 | 0 |
| **EIF3M** | 0 | 0.916569509497771 | 1 | 1 | 0 |
| **APIP** | 0 | 1.28119558069957 | 0.992 | 0.85 | 0 |
| **TP53I11** | 0 | -1.62519018048122 | 0.306 | 0.928 | 0 |
| **LARGE2** | 0 | 1.40468050483758 | 0.997 | 0.789 | 0 |
| **CREB3L1** | 0 | 1.62229414040835 | 0.954 | 0.547 | 0 |
| **AMBRA1** | 0 | 0.98682116489239 | 0.998 | 0.957 | 0 |
| **ATG13** | 0 | 0.859875741355985 | 1 | 0.995 | 0 |
| **ARHGAP1** | 0 | 0.996284946608831 | 1 | 0.999 | 0 |
| **TIMM10** | 0 | 1.05686830943754 | 1 | 1 | 0 |
| **SERPING1** | 0 | -1.38753892423628 | 0.623 | 0.974 | 0 |
| **TMX2** | 0 | 0.884960693739731 | 1 | 0.999 | 0 |
| **OSBP** | 0 | 0.852881895843647 | 1 | 0.995 | 0 |

| | | | | | |
|---|---|---|---|---|---|
| **TMEM109** | 0 | 0.982828326655893 | 0.999 | 0.99 | 0 |
| **TMEM258** | 0 | 1.15678526518792 | 0.997 | 0.923 | 0 |
| **FADS2** | 0 | -1.32455230118356 | 1 | 1 | 0 |
| **FADS3** | 0 | 1.33048323775517 | 0.843 | 0.256 | 0 |
| **ASRGL1** | 0 | 2.52650442530742 | 1 | 0.931 | 0 |
| **EEF1G** | 0 | 1.53612174792991 | 1 | 1 | 0 |
| **POLR2G** | 0 | 0.853256041261336 | 1 | 0.999 | 0 |
| **WDR74** | 0 | 1.05923594762372 | 1 | 0.997 | 0 |
| **ESRRA** | 0 | 1.26890282057687 | 0.996 | 0.881 | 0 |
| **PRDX5** | 0 | 1.20209398994845 | 1 | 0.997 | 0 |
| **CDK2AP2** | 0 | 1.08226820256543 | 0.985 | 0.886 | 0 |
| **NDUFV1** | 0 | 1.54679862494825 | 1 | 1 | 0 |
| **FGF4** | 0 | 1.65073234401921 | 0.968 | 0.639 | 0 |
| **CTTN** | 0 | 0.848115057837291 | 1 | 1 | 0 |
| **RAB6A** | 0 | 1.74259936669671 | 1 | 0.987 | 0 |
| **UCP2** | 0 | 1.29458768670194 | 0.996 | 0.919 | 0 |
| **SERPINH1** | 0 | 0.953433880131624 | 1 | 1 | 0 |
| **PAK1** | 0 | 1.32132585371603 | 0.995 | 0.897 | 0 |
| **NDUFC2** | 0 | 0.896585865381079 | 1 | 1 | 0 |
| **TAF1D** | 0 | 1.30739614827146 | 1 | 0.975 | 0 |
| **C11orf54** | 0 | 1.06405370816584 | 0.988 | 0.832 | 0 |
| **FUT4** | 0 | 1.38166587107816 | 0.954 | 0.647 | 0 |
| **AMOTL1** | 0 | -2.19179965024057 | 0.739 | 1 | 0 |
| **DCUN1D5** | 0 | 1.55060017704277 | 1 | 0.964 | 0 |
| **AASDHPPT** | 0 | 1.06306874620392 | 1 | 0.984 | 0 |
| **EXPH5** | 0 | 1.23688076169046 | 0.704 | 0.077 | 0 |
| **DLAT** | 0 | 1.04028407183687 | 1 | 0.998 | 0 |
| **USP28** | 0 | 1.88276361258915 | 1 | 0.996 | 0 |
| **TAGLN** | 0 | -2.78938487704497 | 0.791 | 0.997 | 0 |
| **TRAPPC4** | 0 | 0.966728611857595 | 0.999 | 0.996 | 0 |
| **HYOU1** | 0 | 1.85823133843579 | 1 | 1 | 0 |
| **HYLS1** | 0 | 1.81191987024958 | 0.783 | 0.123 | 0 |
| **SRPRA** | 0 | 0.987382937143161 | 1 | 0.994 | 0 |
| **SNX19** | 0 | 1.48763932685691 | 0.796 | 0.029 | 0 |

| | | | | | |
|---|---|---|---|---|---|
| **TEAD4** | 0 | 0.899167886235797 | 1 | 0.989 | 0 |
| **TIGAR** | 0 | 1.09334067662597 | 1 | 0.979 | 0 |
| **PTMS** | 0 | -1.53689740713433 | 0.998 | 1 | 0 |
| **GDF3** | 0 | 2.95150957686277 | 1 | 0.716 | 0 |
| **DPPA3** | 0 | 3.47998750269856 | 0.988 | 0.207 | 0 |
| **GPRC5A** | 0 | 1.76108189604233 | 0.641 | 0.049 | 0 |
| **PLBD1** | 0 | 2.93522791044664 | 1 | 0.896 | 0 |
| **YAF2** | 0 | -1.33892599694645 | 0.112 | 0.768 | 0 |
| **PFKM** | 0 | -1.24100314551741 | 0.716 | 0.983 | 0 |
| **FKBP11** | 0 | 2.37912181731865 | 0.988 | 0.462 | 0 |
| **TUBA1C** | 0 | 1.34895414534449 | 1 | 1 | 0 |
| **LIMA1** | 0 | 1.63356110724905 | 0.991 | 0.814 | 0 |
| **RARG** | 0 | -1.69816932260489 | 0.273 | 0.862 | 0 |
| **CBX5** | 0 | -1.12022474955566 | 0.944 | 0.999 | 0 |
| **ITGA5** | 0 | -2.03370692294297 | 0.361 | 0.93 | 0 |
| **MYL6** | 0 | -1.04381962805615 | 1 | 1 | 0 |
| **HMGA2** | 0 | -1.55724762752078 | 0.893 | 1 | 0 |
| **ATXN7L3B** | 0 | -1.03731726223031 | 0.965 | 1 | 0 |
| **NTS** | 0 | -3.60591787438608 | 0.629 | 0.995 | 0 |
| **TMPO** | 0 | 1.11385418585952 | 1 | 0.998 | 0 |
| **ACTR6** | 0 | 1.52564373189088 | 0.937 | 0.486 | 0 |
| **SPIC** | 0 | 1.83000427056653 | 0.681 | 0.022 | 0 |
| **SYCP3** | 0 | 2.66530894469349 | 0.902 | 0.021 | 0 |
| **HSP90B1** | 0 | 2.29305673810025 | 0.995 | 0.779 | 0 |
| **ALDH1L2** | 0 | 1.34620782794952 | 0.78 | 0.144 | 0 |
| **NUAK1** | 0 | -1.40001375719596 | 0.541 | 0.955 | 0 |
| **VSIG10** | 0 | 1.95321401417343 | 1 | 0.935 | 0 |
| **MSI1** | 0 | -2.0063524117805 | 0.174 | 0.944 | 0 |
| **RHOF** | 0 | 1.4482082814798 | 0.94 | 0.456 | 0 |
| **DENR** | 0 | 1.04215490551831 | 0.999 | 0.976 | 0 |
| **VPS37B** | 0 | 1.47271380557492 | 0.978 | 0.709 | 0 |
| **SKA3** | 0 | 1.60471519819475 | 0.995 | 0.939 | 0 |
| **ATP12A** | 0 | 2.07332463840335 | 0.955 | 0.415 | 0 |
| **WASF3** | 0 | -1.23029595146966 | 0.866 | 0.99 | 0 |

| | | | | | |
|---|---|---|---|---|---|
| **NBEA** | 0 | -1.59966497447114 | 0.163 | 0.781 | 0 |
| **UFM1** | 0 | 1.42023045211242 | 1 | 0.944 | 0 |
| **LHFPL6** | 0 | -2.08835282552391 | 0.15 | 0.847 | 0 |
| **RGCC** | 0 | 2.52697965935687 | 0.865 | 0.126 | 0 |
| **EPSTI1** | 0 | 1.20816613044838 | 0.683 | 0.09 | 0 |
| **DNAJC15** | 0 | 2.8627463613313 | 1 | 0.997 | 0 |
| **LCP1** | 0 | 4.01743302748164 | 0.999 | 0.685 | 0 |
| **SUCLA2** | 0 | 1.18112155475558 | 1 | 0.95 | 0 |
| **NUDT15** | 0 | 0.970489827462027 | 1 | 0.999 | 0 |
| **CNMD** | 0 | 2.27621057080723 | 0.96 | 0.344 | 0 |
| **KLF5** | 0 | 2.73015516192888 | 1 | 0.701 | 0 |
| **KLF12** | 0 | -1.41789837588785 | 0.132 | 0.777 | 0 |
| **OBI1** | 0 | -1.31974949958807 | 0.601 | 0.952 | 0 |
| **GPC6** | 0 | -2.46513279389197 | 0.407 | 0.992 | 0 |
| **DZIP1** | 0 | -1.24109851204991 | 0.791 | 0.992 | 0 |
| **DNAJC3** | 0 | 1.33262548515515 | 1 | 0.998 | 0 |
| **DOCK9** | 0 | 1.19888916663289 | 0.993 | 0.74 | 0 |
| **ZIC5** | 0 | -2.0062428195616 | 0.075 | 0.706 | 0 |
| **ZIC2** | 0 | -2.30653416485553 | 0.151 | 0.85 | 0 |
| **FAM155A** | 0 | 1.10883832192151 | 0.629 | 0.043 | 0 |
| **COL4A1** | 0 | -1.36399003918355 | 0.932 | 0.999 | 0 |
| **COL4A2** | 0 | -1.4891036655561 | 0.954 | 1 | 0 |
| **ARHGEF40** | 0 | -1.94904955927561 | 0.259 | 0.935 | 0 |
| **ZNF219** | 0 | -1.94707240939981 | 0.519 | 0.96 | 0 |
| **SALL2** | 0 | -2.54997545261927 | 0.744 | 0.998 | 0 |
| **SLC7A7** | 0 | 1.62479162351837 | 0.936 | 0.57 | 0 |
| **AJUBA** | 0 | -1.49318810968216 | 0.702 | 0.976 | 0 |
| **SLC7A8** | 0 | 1.52862410300249 | 0.994 | 0.841 | 0 |
| **PSME1** | 0 | -1.18168402637446 | 0.805 | 0.99 | 0 |
| **REC8** | 0 | 2.70351674908139 | 0.986 | 0.209 | 0 |
| **EGLN3** | 0 | 1.71906353812283 | 0.99 | 0.839 | 0 |
| **PYGL** | 0 | 1.9920281719735 | 1 | 0.975 | 0 |
| **GNPNAT1** | 0 | 1.72898054117658 | 0.999 | 0.937 | 0 |
| **WDHD1** | 0 | 1.13951078942247 | 1 | 0.999 | 0 |

| | | | | | |
|---|---|---|---|---|---|
| **TBPL2** | 0 | 0.86196734487383 | 0.515 | 0.006 | 0 |
| **TMEM30B** | 0 | 1.49568229352507 | 0.756 | 0.013 | 0 |
| **HSPA2** | 0 | 2.80184165120302 | 0.972 | 0.121 | 0 |
| **PLEKHG3** | 0 | -1.60430145266216 | 0.355 | 0.889 | 0 |
| **GPX2** | 0 | 2.06740907804751 | 0.818 | 0.045 | 0 |
| **RAB15** | 0 | 1.56255758765471 | 1 | 0.993 | 0 |
| **DCAF5** | 0 | -1.28750961057128 | 0.639 | 0.976 | 0 |
| **DCAF4** | 0 | 1.69278125570588 | 0.997 | 0.863 | 0 |
| **ACOT4** | 0 | 1.22151862913231 | 0.678 | 0.01 | 0 |
| **VRTN** | 0 | -3.30352103260367 | 0.432 | 1 | 0 |
| **SLIRP** | 0 | 1.28326157811839 | 1 | 1 | 0 |
| **TUNAR** | 0 | -2.25146100283357 | 0.308 | 0.901 | 0 |
| **C14orf132** | 0 | -1.77771183574723 | 0.134 | 0.799 | 0 |
| **ANKRD9** | 0 | 1.01283275300722 | 0.995 | 0.945 | 0 |
| **TNFAIP2** | 0 | -1.88548532555449 | 0.17 | 0.785 | 0 |
| **KIF26A** | 0 | -1.21873827868953 | 0.853 | 0.996 | 0 |
| **INF2** | 0 | -0.936628122006288 | 0.995 | 1 | 0 |
| **MTA1** | 0 | -1.23959866049365 | 0.971 | 1 | 0 |
| **CRIP1** | 0 | 3.10049074323231 | 1 | 0.817 | 0 |
| **SPINT1** | 0 | 1.39030778505805 | 1 | 0.892 | 0 |
| **PDIA3** | 0 | 1.26159460242413 | 1 | 0.997 | 0 |
| **B2M** | 0 | -1.36506864519191 | 0.652 | 0.967 | 0 |
| **MYEF2** | 0 | -1.63806331553379 | 0.391 | 0.919 | 0 |
| **EID1** | 0 | -1.78103272761142 | 0.952 | 1 | 0 |
| **CCPG1** | 0 | 1.66508669273079 | 0.995 | 0.898 | 0 |
| **PRTG** | 0 | -2.38618915142729 | 0.151 | 0.896 | 0 |
| **SNX1** | 0 | -1.20121674636822 | 0.871 | 0.996 | 0 |
| **PPIB** | 0 | 1.59280544001329 | 1 | 0.999 | 0 |
| **PCLAF** | 0 | -2.63428531013513 | 0.181 | 0.973 | 0 |
| **OAZ2** | 0 | -0.913022294252676 | 0.996 | 1 | 0 |
| **CLPX** | 0 | 1.26149359721476 | 1 | 0.978 | 0 |
| **IGDCC3** | 0 | -1.91000097318822 | 0.496 | 0.979 | 0 |
| **UACA** | 0 | 1.3320812931133 | 0.981 | 0.832 | 0 |
| **ETFA** | 0 | -1.54920490085604 | 0.729 | 0.993 | 0 |

| | | | | | |
|---|---|---|---|---|---|
| **TSPAN3** | 0 | -0.935885695588664 | 0.995 | 1 | 0 |
| **PEAK1** | 0 | -1.28539678487824 | 0.318 | 0.863 | 0 |
| **DNAJA4** | 0 | 1.5727898145288 | 0.815 | 0.033 | 0 |
| **SEC11A** | 0 | -0.817773017412656 | 0.999 | 1 | 0 |
| **MFGE8** | 0 | -1.61260753454537 | 0.484 | 0.96 | 0 |
| **FAM174B** | 0 | -1.99511901938092 | 0.381 | 0.942 | 0 |
| **IGF1R** | 0 | -1.3377132191628 | 0.95 | 1 | 0 |
| **CHSY1** | 0 | -1.32285756316198 | 0.951 | 1 | 0 |
| **HBM** | 0 | 1.10603645763215 | 0.595 | 0.017 | 0 |
| **HBA2** | 0 | 1.64034583925599 | 0.745 | 0.149 | 0 |
| **NME4** | 0 | -1.3229712265745 | 0.997 | 1 | 0 |
| **CACNA1H** | 0 | 1.3489047337352 | 0.749 | 0.052 | 0 |
| **SYNGR3** | 0 | 1.29748559212949 | 0.952 | 0.566 | 0 |
| **SLC9A3R2** | 0 | 1.31929448289812 | 0.959 | 0.65 | 0 |
| **IL32** | 0 | 1.52199760332645 | 0.872 | 0.333 | 0 |
| **NMRAL1** | 0 | 1.76574665146637 | 0.997 | 0.72 | 0 |
| **HMOX2** | 0 | 1.16018356266537 | 0.999 | 0.943 | 0 |
| **CDIP1** | 0 | -1.18874424243687 | 0.801 | 0.988 | 0 |
| **LITAF** | 0 | -1.48594595586557 | 0.981 | 1 | 0 |
| **SHISA9** | 0 | -1.60900177584965 | 0.129 | 0.746 | 0 |
| **NDUFAB1** | 0 | 1.36521958581535 | 1 | 1 | 0 |
| **SEPHS2** | 0 | 1.65730269563281 | 1 | 0.989 | 0 |
| **DNAJA2** | 0 | 1.09698289932827 | 1 | 1 | 0 |
| **ZNF423** | 0 | -2.02128616988724 | 0.578 | 0.995 | 0 |
| **SALL1** | 0 | -2.69472477624661 | 0.238 | 0.955 | 0 |
| **MMP2** | 0 | -3.6824357448921 | 0.432 | 0.998 | 0 |
| **MT1E** | 0 | 2.35890619699072 | 0.946 | 0.148 | 0 |
| **MT1F** | 0 | 2.06529885234171 | 0.992 | 0.768 | 0 |
| **MT1G** | 0 | 3.99982345705664 | 0.999 | 0.268 | 0 |
| **MT1H** | 0 | 4.80835518351029 | 0.999 | 0.362 | 0 |
| **MT1X** | 0 | 2.9966797602911 | 1 | 0.979 | 0 |
| **USB1** | 0 | -1.19947859878547 | 0.897 | 0.996 | 0 |
| **PLEKHG4** | 0 | 0.958977047168168 | 0.566 | 0.019 | 0 |
| **BCAR1** | 0 | -1.4627159484073 | 0.888 | 0.997 | 0 |

| | | | | | |
|---|---|---|---|---|---|
| **WWOX** | 0 | -1.62355800875479 | 0.553 | 0.963 | 0 |
| **PIEZO1** | 0 | 0.958413154870744 | 0.611 | 0.047 | 0 |
| **RFLNB** | 0 | 1.69525202771715 | 0.969 | 0.676 | 0 |
| **SERPINF1** | 0 | 1.70012527228897 | 0.946 | 0.464 | 0 |
| **SPATA22** | 0 | 0.884906389582466 | 0.523 | 0.007 | 0 |
| **SHPK** | 0 | 1.25844949408562 | 0.984 | 0.834 | 0 |
| **ZFP3** | 0 | 1.46406934899326 | 0.747 | 0.011 | 0 |
| **TRAPPC1** | 0 | -0.979109009604147 | 0.976 | 1 | 0 |
| **NTN1** | 0 | 7.20488101385609 | 1 | 0.986 | 0 |
| **FLOT2** | 0 | -1.37408012332641 | 0.875 | 0.999 | 0 |
| **UTP6** | 0 | 1.00761564973744 | 0.997 | 0.958 | 0 |
| **TMEM98** | 0 | -1.4123728212263 | 0.73 | 0.985 | 0 |
| **SLFN12** | 0 | 1.30816573505624 | 0.732 | 0.076 | 0 |
| **IGFBP4** | 0 | -2.20650322505378 | 0.36 | 0.932 | 0 |
| **KRT19** | 0 | -1.38822321105155 | 0.774 | 0.982 | 0 |
| **EIF1** | 0 | 1.01877162593394 | 0.997 | 0.922 | 0 |
| **FKBP10** | 0 | -1.67061700661296 | 0.824 | 0.997 | 0 |
| **VAT1** | 0 | 0.953535099473055 | 1 | 1 | 0 |
| **FAM171A2** | 0 | -1.58049372046588 | 0.493 | 0.952 | 0 |
| **FZD2** | 0 | -2.96316937541669 | 0.225 | 0.98 | 0 |
| **GJC1** | 0 | -1.94059101438533 | 0.776 | 0.999 | 0 |
| **ATP5MC1** | 0 | 0.951542879531098 | 1 | 0.986 | 0 |
| **IGF2BP1** | 0 | -1.64755772979014 | 0.99 | 1 | 0 |
| **LRRC59** | 0 | 0.849512956119983 | 1 | 1 | 0 |
| **ACE** | 0 | 1.46441785740188 | 0.849 | 0.2 | 0 |
| **PITPNC1** | 0 | 1.18955856546006 | 1 | 0.961 | 0 |
| **JPT1** | 0 | 1.23340881081922 | 1 | 0.998 | 0 |
| **SEPTIN9** | 0 | -0.945813138921866 | 0.999 | 1 | 0 |
| **TIMP2** | 0 | -1.57328841748107 | 0.716 | 0.973 | 0 |
| **LGALS3BP** | 0 | -1.66768550338808 | 0.302 | 0.914 | 0 |
| **PYCR1** | 0 | 2.65740031916078 | 1 | 0.596 | 0 |
| **FN3K** | 0 | 1.30122612921194 | 0.957 | 0.6 | 0 |
| **TBCD** | 0 | 1.30543546845988 | 1 | 0.999 | 0 |
| **PIEZO2** | 0 | -2.00298252698463 | 0.083 | 0.855 | 0 |

| | | | | | |
|---|---|---|---|---|---|
| **IMPA2** | 0 | 1.20739924336554 | 0.999 | 0.975 | 0 |
| **CDH2** | 0 | -3.07738430513923 | 0.272 | 0.941 | 0 |
| **RNF125** | 0 | 2.77751231164709 | 0.997 | 0.673 | 0 |
| **GALNT1** | 0 | -1.37695952818457 | 0.946 | 1 | 0 |
| **TPGS2** | 0 | -1.10402145653192 | 0.977 | 1 | 0 |
| **TCF4** | 0 | -1.74429505655869 | 0.523 | 0.974 | 0 |
| **NARS** | 0 | 0.874935681105781 | 1 | 1 | 0 |
| **ATP8B1** | 0 | 0.958634207152394 | 0.539 | 0.018 | 0 |
| **SALL3** | 0 | -2.07337244512106 | 0.455 | 0.973 | 0 |
| **SLC66A2** | 0 | 1.2911372982212 | 0.986 | 0.85 | 0 |
| **HSBP1L1** | 0 | 0.983298068570681 | 0.622 | 0.059 | 0 |
| **CFD** | 0 | 1.24562609586845 | 0.701 | 0.093 | 0 |
| **CNN2** | 0 | -1.34099733027507 | 1 | 1 | 0 |
| **CIRBP** | 0 | -0.843943803870292 | 1 | 1 | 0 |
| **REEP6** | 0 | 1.55880904892117 | 0.921 | 0.414 | 0 |
| **ZNF57** | 0 | 1.09621691694707 | 0.612 | 0.012 | 0 |
| **MATK** | 0 | 1.12485209583856 | 0.661 | 0.086 | 0 |
| **PTPRS** | 0 | -1.15931368885278 | 0.973 | 0.998 | 0 |
| **TNFSF9** | 0 | 2.07743025198954 | 0.809 | 0.07 | 0 |
| **CD70** | 0 | 2.25292046886813 | 0.804 | 0.089 | 0 |
| **ZNF358** | 0 | -1.09630628550757 | 0.883 | 0.996 | 0 |
| **CTXN1** | 0 | -1.53106688366687 | 0.752 | 0.98 | 0 |
| **CD320** | 0 | 1.3844994857684 | 1 | 0.998 | 0 |
| **RAB11B** | 0 | -0.803121656815668 | 0.998 | 1 | 0 |
| **ZNF560** | 0 | 0.996267698442171 | 0.608 | 0.005 | 0 |
| **CDC37** | 0 | 0.810242280042658 | 1 | 1 | 0 |
| **AP1M2** | 0 | 1.40463467474668 | 0.938 | 0.353 | 0 |
| **DOCK6** | 0 | 1.19212897841585 | 0.984 | 0.805 | 0 |
| **ACP5** | 0 | 1.46807917543068 | 0.784 | 0.058 | 0 |
| **ZNF844** | 0 | 0.867041549089449 | 0.525 | 0.007 | 0 |
| **CALR** | 0 | 0.915368676977831 | 1 | 1 | 0 |
| **PRKACA** | 0 | -1.1978114498879 | 0.961 | 1 | 0 |
| **NOTCH3** | 0 | -1.98063215878771 | 0.737 | 0.995 | 0 |
| **SLC5A5** | 0 | 1.26938841575253 | 0.729 | 0.056 | 0 |

| | | | | | |
|---|---|---|---|---|---|
| **ZNF253** | 0 | 1.20691769250649 | 0.692 | 0.013 | 0 |
| **ZNF90** | 0 | 1.63201164251875 | 0.993 | 0.721 | 0 |
| **ZNF486** | 0 | 1.97572098568593 | 0.964 | 0.407 | 0 |
| **ZNF257** | 0 | 0.879835956886694 | 0.545 | 0.006 | 0 |
| **ZNF676** | 0 | 2.95757165866941 | 0.989 | 0.129 | 0 |
| **ZNF98** | 0 | 1.40427861850891 | 0.768 | 0.004 | 0 |
| **ZNF492** | 0 | 1.36076336682667 | 0.736 | 0.009 | 0 |
| **ZNF728** | 0 | 1.12912051621812 | 0.658 | 0.005 | 0 |
| **ZNF829** | 0 | 0.802520273223759 | 0.534 | 0.007 | 0 |
| **ZNF568** | 0 | 2.13230148309661 | 0.94 | 0.086 | 0 |
| **PPP1R14A** | 0 | 1.23125610470732 | 0.998 | 0.898 | 0 |
| **YIF1B** | 0 | 1.03801228955359 | 1 | 0.997 | 0 |
| **C19orf33** | 0 | 2.56180395171975 | 0.975 | 0.233 | 0 |
| **KCNK6** | 0 | 1.26519329004868 | 0.781 | 0.148 | 0 |
| **MAP4K1** | 0 | 1.50698083287494 | 0.851 | 0.268 | 0 |
| **FCGBP** | 0 | 2.74357886047027 | 0.87 | 0.046 | 0 |
| **PSMC4** | 0 | 1.41240600870107 | 1 | 1 | 0 |
| **PLD3** | 0 | -1.17580378501871 | 0.995 | 1 | 0 |
| **AXL** | 0 | -2.57998950885774 | 0.131 | 0.915 | 0 |
| **ATP1A3** | 0 | 1.36053023544996 | 0.764 | 0.083 | 0 |
| **PINLYP** | 0 | 1.07315947476075 | 0.641 | 0.026 | 0 |
| **BCAM** | 0 | -2.50133094946892 | 0.369 | 0.946 | 0 |
| **ARHGAP35** | 0 | -1.44131012673851 | 0.835 | 0.995 | 0 |
| **TMEM160** | 0 | 1.39448853972837 | 1 | 0.999 | 0 |
| **CA11** | 0 | -1.50801378088969 | 0.069 | 0.725 | 0 |
| **RASIP1** | 0 | 1.53238369878975 | 0.834 | 0.184 | 0 |
| **NUCB1** | 0 | 0.916937321731097 | 1 | 0.999 | 0 |
| **RCN3** | 0 | 1.15948488974442 | 0.715 | 0.113 | 0 |
| **PTOV1** | 0 | -1.01976990780774 | 0.995 | 1 | 0 |
| **KLK13** | 0 | 1.40377480757214 | 0.851 | 0.315 | 0 |
| **LIM2** | 0 | 2.23652892784934 | 0.914 | 0.022 | 0 |
| **ZNF649** | 0 | 1.39722528996797 | 0.988 | 0.812 | 0 |
| **ZNF880** | 0 | 1.32873409296229 | 0.825 | 0.127 | 0 |
| **ZNF677** | 0 | 1.75906862713369 | 0.882 | 0.018 | 0 |

| | | | | | |
|---|---|---|---|---|---|
| **NLRP7** | 0 | 5.58487647006752 | 1 | 0.58 | 0 |
| **NLRP2** | 0 | 3.08055817310231 | 1 | 0.99 | 0 |
| **C19orf85** | 0 | 2.62135273908946 | 0.847 | 0.074 | 0 |
| **ZNF667** | 0 | 0.824190793326974 | 0.516 | 0.006 | 0 |
| **PEG3** | 0 | 2.29838291550793 | 0.874 | 0.026 | 0 |
| **ZNF552** | 0 | 1.25476611488626 | 0.986 | 0.694 | 0 |
| **ZCCHC3** | 0 | -1.99170452901545 | 0.709 | 0.999 | 0 |
| **SOX12** | 0 | -1.74279082833256 | 0.568 | 0.971 | 0 |
| **TRIB3** | 0 | 1.91800798305101 | 0.974 | 0.697 | 0 |
| **SRXN1** | 0 | -1.4307019180213 | 0.662 | 0.961 | 0 |
| **IDH3B** | 0 | -0.878864159572393 | 0.999 | 1 | 0 |
| **PTPRA** | 0 | -1.07785201860353 | 0.846 | 0.987 | 0 |
| **CDS2** | 0 | -1.55665655091296 | 0.859 | 1 | 0 |
| **FERMT1** | 0 | 1.4979588379976 | 0.778 | 0.125 | 0 |
| **BTBD3** | 0 | -1.34013165030605 | 0.703 | 0.984 | 0 |
| **XRN2** | 0 | -0.806745469236816 | 1 | 1 | 0 |
| **CST3** | 0 | -1.7750120578761 | 0.428 | 0.953 | 0 |
| **PYGB** | 0 | 1.75081375456897 | 1 | 0.999 | 0 |
| **GINS1** | 0 | -1.07207889087665 | 0.998 | 1 | 0 |
| **ASXL1** | 0 | -1.96266275051095 | 0.606 | 0.995 | 0 |
| **NOL4L** | 0 | -1.42532032719075 | 0.36 | 0.899 | 0 |
| **MAPRE1** | 0 | -1.02137489502989 | 0.998 | 1 | 0 |
| **CBFA2T2** | 0 | 1.7237885684696 | 0.996 | 0.918 | 0 |
| **RALY** | 0 | -1.01999988353465 | 0.967 | 0.999 | 0 |
| **DYNLRB1** | 0 | -1.3481369759119 | 0.999 | 1 | 0 |
| **ERGIC3** | 0 | -0.990492606733484 | 1 | 1 | 0 |
| **CPNE1** | 0 | -1.63669863959594 | 0.967 | 1 | 0 |
| **DLGAP4** | 0 | -1.34897748317296 | 0.865 | 0.996 | 0 |
| **MYL9** | 0 | -3.27039170076289 | 0.831 | 1 | 0 |
| **TGIF2** | 0 | -1.41188211462647 | 0.946 | 1 | 0 |
| **SAMHD1** | 0 | 2.71088713151521 | 0.993 | 0.611 | 0 |
| **SRC** | 0 | -1.59929756941908 | 0.762 | 0.997 | 0 |
| **NNAT** | 0 | 1.27538678461733 | 0.546 | 0.005 | 0 |
| **1,00 TOP** | 0 | -1.08845411692353 | 0.999 | 1 | 0 |

| | | | | | |
|---|---|---|---|---|---|
| **PLCG1** | 0 | -1.1013037199347 | 0.971 | 1 | 0 |
| **HNF4A** | 0 | 1.43899026809791 | 0.738 | 0.019 | 0 |
| **STK4** | 0 | -1.12336581502673 | 0.855 | 0.992 | 0 |
| **DBNDD2** | 0 | -1.86904995940425 | 0.259 | 0.904 | 0 |
| **PIGT** | 0 | -1.56854952755609 | 0.917 | 1 | 0 |
| **WFDC2** | 0 | -3.29647772723491 | 0.277 | 0.931 | 0 |
| **PLTP** | 0 | -1.19795953961433 | 0.864 | 0.995 | 0 |
| **SULF2** | 0 | -2.09247693360789 | 0.711 | 0.988 | 0 |
| **CSE1L** | 0 | -0.857925757174642 | 0.999 | 1 | 0 |
| **STAU1** | 0 | -0.990528512284717 | 1 | 1 | 0 |
| **B4GALT5** | 0 | -1.19363821906182 | 0.948 | 1 | 0 |
| **PTPN1** | 0 | -0.964486610604844 | 0.957 | 0.999 | 0 |
| **BCAS4** | 0 | -1.24360244250469 | 0.602 | 0.945 | 0 |
| **BMP7** | 0 | -2.26073533365313 | 0.597 | 0.985 | 0 |
| **RAE1** | 0 | -1.04423048847607 | 0.969 | 1 | 0 |
| **NELFCD** | 0 | -1.27611154405001 | 0.974 | 1 | 0 |
| **ATP5F1E** | 0 | -1.57762480627995 | 0.997 | 1 | 0 |
| **PSMA7** | 0 | -0.974604297454849 | 1 | 1 | 0 |
| **KCNQ2** | 0 | 1.50016116701828 | 0.937 | 0.312 | 0 |
| **SRMS** | 0 | 1.3106846979808 | 0.617 | 0.022 | 0 |
| **SLC2A4RG** | 0 | -1.09846343042312 | 0.857 | 0.995 | 0 |
| **DNAJC5** | 0 | -1.33050670365483 | 0.935 | 1 | 0 |
| **TTC3** | 0 | -1.09519777808648 | 0.87 | 0.995 | 0 |
| **BACE2** | 0 | -1.59926683798546 | 0.276 | 0.886 | 0 |
| **PDXK** | 0 | 1.14213384796442 | 1 | 0.997 | 0 |
| **DNMT3L** | 0 | 4.62353960190359 | 1 | 0.708 | 0 |
| **COL18A1** | 0 | -3.31037801814448 | 0.693 | 1 | 0 |
| **SPATC1L** | 0 | -1.46990573331086 | 0.063 | 0.794 | 0 |
| **ATP6V1E1** | 0 | 0.830638798120678 | 1 | 0.985 | 0 |
| **USP18** | 0 | 1.43442817798121 | 0.914 | 0.414 | 0 |
| **MED15** | 0 | 1.18593530601427 | 0.996 | 0.942 | 0 |
| **SDF2L1** | 0 | 2.15426551331231 | 1 | 0.98 | 0 |
| **DERL3** | 0 | 2.2025083769699 | 0.898 | 0.171 | 0 |
| **DDT** | 0 | 1.12560820621257 | 1 | 1 | 0 |

| | | | | | |
|---|---|---|---|---|---|
| **SUSD2** | 0 | 3.73952124447865 | 1 | 0.641 | 0 |
| **ADORA2A** | 0 | 1.35532884610096 | 0.739 | 0.099 | 0 |
| **TPST2** | 0 | 1.79977345595717 | 0.999 | 0.807 | 0 |
| **CHEK2** | 0 | 1.27429463127609 | 0.997 | 0.941 | 0 |
| **XBP1** | 0 | 1.70191447573617 | 1 | 0.998 | 0 |
| **NEFH** | 0 | 2.37433665304076 | 0.942 | 0.371 | 0 |
| **THOC5** | 0 | 1.06779856453211 | 1 | 0.983 | 0 |
| **NF2** | 0 | 1.51917636263455 | 1 | 0.995 | 0 |
| **TCN2** | 0 | -1.66525911154445 | 0.277 | 0.934 | 0 |
| **EIF3L** | 0 | 0.891159831294672 | 1 | 1 | 0 |
| **TOMM22** | 0 | 1.27368351687857 | 1 | 0.996 | 0 |
| **APOBEC3G** | 0 | 1.33226641161262 | 0.785 | 0.167 | 0 |
| **SYNGR1** | 0 | -1.54335747285094 | 0.351 | 0.901 | 0 |
| **SNU13** | 0 | 1.18738718851423 | 1 | 1 | 0 |
| **NDUFA6** | 0 | 1.10406785207279 | 1 | 0.981 | 0 |
| **A4GALT** | 0 | 2.86794210745345 | 0.999 | 0.666 | 0 |
| **TTLL12** | 0 | 1.16744351956348 | 1 | 0.991 | 0 |
| **PARVB** | 0 | 1.92258536845558 | 0.972 | 0.488 | 0 |
| **FBLN1** | 0 | -1.3855208569029 | 0.957 | 1 | 0 |
| **CRELD2** | 0 | 2.04746256042086 | 0.999 | 0.934 | 0 |
| **DHRSX** | 0 | 1.933041983629 | 0.995 | 0.856 | 0 |
| **CD99** | 0 | -2.35689446360293 | 0.293 | 0.969 | 0 |
| **GYG2** | 0 | -1.31580315921167 | 0.652 | 0.983 | 0 |
| **TBL1X** | 0 | -1.57621483542116 | 0.374 | 0.925 | 0 |
| **WWC3** | 0 | -1.96524437768863 | 0.448 | 0.976 | 0 |
| **TMSB4X** | 0 | -2.99638331054685 | 0.999 | 1 | 0 |
| **AP1S2** | 0 | -1.29658968399541 | 0.578 | 0.95 | 0 |
| **SMS** | 0 | -1.75367728169981 | 0.976 | 1 | 0 |
| **PCYT1B** | 0 | -1.59708613418447 | 0.351 | 0.926 | 0 |
| **ARX** | 0 | 1.18408624489078 | 0.509 | 0.004 | 0 |
| **MAGEB2** | 0 | 2.29974585980685 | 0.806 | 0.026 | 0 |
| **DYNLT3** | 0 | 0.923362383924002 | 0.561 | 0.006 | 0 |
| **TIMP1** | 0 | -1.63084328315434 | 0.984 | 1 | 0 |
| **PIM2** | 0 | 1.7695787812277 | 0.997 | 0.973 | 0 |

| | | | | | |
|---|---|---|---|---|---|
| **MAGED1** | 0 | -1.79099100235818 | 0.981 | 1 | 0 |
| **MAGED2** | 0 | -1.44273372569647 | 0.999 | 1 | 0 |
| **MAGEH1** | 0 | -1.34759785821255 | 0.456 | 0.931 | 0 |
| **UBQLN2** | 0 | -0.978710868137409 | 0.96 | 1 | 0 |
| **ARHGEF9** | 0 | -1.30790334053129 | 0.253 | 0.83 | 0 |
| **MSN** | 0 | -1.17055326955804 | 0.999 | 1 | 0 |
| **HEPH** | 0 | -1.8191411589242 | 0.065 | 0.869 | 0 |
| **EFNB1** | 0 | -2.24550603828013 | 0.46 | 0.974 | 0 |
| **MED12** | 0 | -1.37056067122002 | 0.166 | 0.804 | 0 |
| **PHKA1** | 0 | 2.7079065959909 | 1 | 0.773 | 0 |
| **SLC16A2** | 0 | -1.62655432699617 | 0.641 | 0.987 | 0 |
| **COX7B** | 0 | 1.30345770285355 | 0.972 | 0.745 | 0 |
| **BRWD3** | 0 | 1.34244183964739 | 0.997 | 0.978 | 0 |
| **NAP1L3** | 0 | -1.58068676830324 | 0.363 | 0.894 | 0 |
| **BEX5** | 0 | 0.997987138344989 | 0.574 | 0.037 | 0 |
| **TMSB15A** | 0 | -2.03892501638611 | 0.212 | 0.823 | 0 |
| **BEX4** | 0 | -1.32166698416163 | 0.967 | 1 | 0 |
| **TCEAL9** | 0 | -1.64383820996321 | 0.864 | 0.999 | 0 |
| **BEX3** | 0 | -2.49739255152651 | 0.96 | 1 | 0 |
| **COL4A5** | 0 | -1.6039592380029 | 0.24 | 0.856 | 0 |
| **LUZP4** | 0 | 1.605950422847 | 0.722 | 0.006 | 0 |
| **PGRMC1** | 0 | -1.45530632926944 | 0.998 | 1 | 0 |
| **SMARCA1** | 0 | -1.61494898042836 | 0.674 | 0.992 | 0 |
| **HS6ST2** | 0 | -1.85628191548639 | 0.222 | 0.921 | 0 |
| **LDOC1** | 0 | -1.18489354216613 | 0.688 | 0.963 | 0 |
| **ZNF185** | 0 | -1.45783268140325 | 0.39 | 0.891 | 0 |
| **DUSP9** | 0 | 1.20826726172209 | 0.676 | 0.069 | 0 |
| **SLC6A8** | 0 | 1.80067966521201 | 0.999 | 0.95 | 0 |
| **IRAK1** | 0 | 1.38377866207148 | 0.947 | 0.586 | 0 |
| **FLNA** | 0 | -0.878354622398243 | 1 | 1 | 0 |
| **GDI1** | 0 | -1.00624302815115 | 0.98 | 1 | 0 |
| **DDX3Y** | 0 | 1.14809657783943 | 0.989 | 0.822 | 0 |
| **MT-ND3** | 0 | 2.12316849794781 | 0.982 | 0.722 | 0 |
| **C7orf50** | 4.6022 | -0.886392666784599 | 0.993 | 1 | 7.5371276 |

| | | | | | |
|---|---|---|---|---|---|
| **PTBP2** | 1.4661 | -1.20952566991075 | 0.796 | 0.987 | 2.4011945 |
| **ARHGAP42** | 1.8708 | 1.21138828621222 | 0.912 | 0.461 | 3.0639514 |
| **TESMIN** | 2.3800 | 1.22501789810527 | 0.919 | 0.492 | 3.8978566 |
| **CDH1** | 2.7381 | 1.21638793678602 | 1 | 0.77 | 4.4841992 |
| **CPD** | 5.4168 | -1.04735618844524 | 0.963 | 0.998 | 8.8712036 |
| **FAM122B** | 2.3603 | -1.32035720664039 | 0.701 | 0.965 | 3.8655604 |
| **NIPA2** | 7.6022 | 0.865706976880008 | 0.997 | 0.965 | 1.2450254 |
| **UTRN** | 8.4070 | -1.38667927606649 | 0.517 | 0.933 | 1.3768160 |
| **PTPRU** | 8.7405 | 1.13130435617254 | 0.989 | 0.909 | 1.4314413 |
| **SH3KBP1** | 9.6499 | -1.23145611734255 | 0.758 | 0.981 | 1.5803772 |
| **NRROS** | 4.6615 | 0.982088186810399 | 0.575 | 0.044 | 7.6341526 |
| **CAST** | 4.7314 | -1.35866513002351 | 0.187 | 0.774 | 7.7486150 |
| **ZCCHC24** | 7.9766 | -1.39620322531821 | 0.344 | 0.864 | 1.3063280 |
| **PCCA** | 1.6771 | -1.08056406159695 | 0.958 | 0.998 | 2.7466422 |
| **EIF4EBP1** | 2.7672 | 0.997032368459051 | 1 | 1 | 4.5319827 |
| **AKT1** | 4.4961 | -0.916613299155253 | 0.997 | 1 | 7.3632764 |
| **NOTCH1** | 7.3399 | -1.24871577833534 | 0.736 | 0.979 | 1.2020554 |
| **DNAJB6** | 9.0343 | 0.995300138823443 | 0.999 | 0.949 | 1.4795554 |
| **AFAP1L1** | 1.1273 | 1.17811193022214 | 0.754 | 0.187 | 1.8463074 |
| **B4GALT4** | 1.3445 | 1.16051888876048 | 0.803 | 0.256 | 2.2019808 |
| **SRSF8** | 1.6501 | 1.18898522672338 | 0.98 | 0.734 | 2.7025110 |
| **GPT2** | 1.7896 | 1.07409326626626 | 1 | 0.981 | 2.9309348 |
| **SEC61B** | 1.9701 | 0.961921019624238 | 0.997 | 0.946 | 3.2264874 |
| **HDGFL3** | 3.2312 | -0.855565259337546 | 0.978 | 1 | 5.2918542 |
| **PCBD1** | 3.6725 | -1.27099178189639 | 0.728 | 0.966 | 6.0144585 |
| **TES** | 3.9397 | 1.05836436083767 | 0.999 | 0.947 | 6.4521427 |
| **ZNF732** | 4.0067 | 0.816251659023602 | 0.492 | 0.003 | 6.5618183 |
| **GPD2** | 1.0009 | -1.15397783798453 | 0.881 | 0.993 | 1.6392624 |
| **OLFM1** | 1.5203 | -1.54228079460329 | 0.49 | 0.913 | 2.4899343 |
| **CSNK1G2** | 5.8507 | -0.853803210256474 | 0.984 | 1 | 9.5818022 |
| **MAPRE2** | 6.8408 | -1.24937434790639 | 0.528 | 0.935 | 1.1203202 |
| **NOC2L** | 2.6955 | 0.834364591499217 | 1 | 0.998 | 4.4144298 |
| **HAUS1** | 3.7573 | 1.06041908801996 | 0.976 | 0.782 | 6.1533777 |
| **CCDC102A** | 5.0086 | -1.25527892561022 | 0.31 | 0.841 | 8.2026990 |

| | | | | | |
|---|---|---|---|---|---|
| **ZBTB44** | 6.3883 | 0.803625251550666 | 1 | 0.994 | 1.0462192 |
| **SLC25A24** | 7.3629 | -1.02059859163805 | 0.906 | 0.996 | 1.2058258 |
| **APMAP** | 9.9930 | -0.948687435125399 | 0.948 | 1 | 1.6365684 |
| **MBNL3** | 1.1247 | -1.47611921828058 | 0.609 | 0.944 | 1.8419531 |
| **AP2B1** | 1.1585 | 0.977130055446689 | 0.991 | 0.873 | 1.8974316 |
| **BRSK2** | 1.8428 | 1.03155933436349 | 0.631 | 0.084 | 3.0180863 |
| **TMEM126A** | 2.1040 | 0.973266131458565 | 0.99 | 0.869 | 3.4458653 |
| **TNPO2** | 2.1110 | -1.00051287587013 | 0.923 | 0.999 | 3.4572609 |
| **LHX8** | 2.2987 | 0.906442667036075 | 0.5 | 0.008 | 3.7646195 |
| **PDK1** | 2.8604 | -1.52274792888763 | 0.948 | 0.998 | 4.6845002 |
| **PTGIS** | 5.7134 | -1.34334743876452 | 0.105 | 0.716 | 9.3569088 |
| **RIN2** | 2.6278 | -1.36231885148516 | 0.474 | 0.913 | 4.3035731 |
| **KLHDC10** | 3.5410 | -1.15297209005274 | 0.141 | 0.748 | 5.7992220 |
| **GBX2** | 1.0617 | -1.57965165176785 | 0.419 | 0.894 | 1.7388556 |
| **MYSM1** | 8.3166 | 1.1410651003556 | 0.996 | 0.905 | 1.3620205 |
| **SPAG9** | 2.3901 | -0.989023496152627 | 0.868 | 0.995 | 3.9144241 |
| **SYVN1** | 2.3974 | 1.32699088213469 | 0.96 | 0.827 | 3.9262509 |
| **SLC23A2** | 6.4072 | 1.13323184017797 | 0.98 | 0.845 | 1.0493114 |
| **EFS** | 7.6679 | -1.38245778505407 | 0.439 | 0.891 | 1.2557866 |
| **DCLK1** | 1.2644 | -1.74998121293016 | 0.903 | 0.961 | 2.0707865 |
| **VANGL1** | 2.2143 | -1.38807987461303 | 0.739 | 0.973 | 3.6264517 |
| **SMG6** | 2.9585 | -1.11986296450373 | 0.599 | 0.94 | 4.8451583 |
| **CTNNBL1** | 3.1980 | -0.854028476697683 | 0.979 | 1 | 5.2374897 |
| **ERGIC1** | 1.0184 | -1.09067301432643 | 0.698 | 0.968 | 1.6679464 |
| **OTUD4** | 1.0316 | 1.02377962099099 | 0.978 | 0.824 | 1.6895674 |
| **REPIN1** | 1.0659 | -1.18157413078789 | 0.5 | 0.913 | 1.7457089 |
| **TAF9B** | 2.7348 | -1.16591346133186 | 0.547 | 0.928 | 4.4787961 |
| **NRM** | 2.7977 | -1.14261397453318 | 0.59 | 0.934 | 4.5819128 |
| **ACSL4** | 3.6626 | -1.1379166429989 | 0.82 | 0.996 | 5.9982909 |
| **CHST8** | 4.1476 | 1.20460596065336 | 0.69 | 0.14 | 6.7926704 |
| **EFR3A** | 7.9104 | 0.959322038905843 | 0.999 | 0.991 | 1.2954978 |
| **GTF2H1** | 9.7707 | 0.970460821051511 | 0.988 | 0.894 | 1.6001603 |
| **HMGB2** | 1.0779 | -1.1195745537295 | 0.998 | 0.999 | 1.7654349 |
| **TBL2** | 1.0881 | -1.15037141740973 | 0.593 | 0.943 | 1.7820475 |

| | | | | | |
|---|---|---|---|---|---|
| **CMTM3** | 1.6944 | -1.06582267977371 | 0.969 | 1 | 2.7750767 |
| **ACACB** | 1.8892 | 1.22184119903645 | 0.783 | 0.253 | 3.0939736 |
| **TNNI3** | 2.3397 | 1.12245636348235 | 0.764 | 0.187 | 3.8317498 |
| **RHOB** | 2.3466 | -1.65514560671517 | 0.917 | 0.997 | 3.8431745 |
| **CACHD1** | 2.9147 | -1.54865416348459 | 0.724 | 0.966 | 4.7735102 |
| **ZEB2** | 3.2476 | -1.54465968493649 | 0.086 | 0.692 | 5.3187243 |
| **KHDRBS1** | 3.5623 | -0.921610975306862 | 0.979 | 1 | 5.8340641 |
| **PRXL2A** | 4.8555 | -0.914277514788244 | 0.962 | 0.998 | 7.9518626 |
| **CNN1** | 7.0213 | -2.17396024901792 | 0.456 | 0.904 | 1.1498848 |
| **SORD** | 1.0152 | -1.17303498061492 | 0.537 | 0.916 | 1.6626358 |
| **PDIA4** | 1.3381 | 0.937912415045747 | 0.999 | 0.997 | 2.1914802 |
| **RTF2** | 2.5529 | -0.808538472898282 | 0.982 | 0.999 | 4.1809805 |
| **APOBEC3D** | 3.3789 | 1.03370333095497 | 0.676 | 0.122 | 5.5337114 |
| **PMM2** | 9.2104 | 0.919075770870047 | 0.997 | 0.964 | 1.5083911 |
| **PDGFRB** | 1.7507 | -1.21298451599942 | 0.037 | 0.641 | 2.8671782 |
| **UBASH3B** | 2.2718 | 1.26203708753608 | 0.927 | 0.553 | 3.7205305 |
| **DPH6** | 4.5022 | 1.10962165641894 | 0.967 | 0.7 | 7.3733973 |
| **GNPAT** | 5.3829 | -1.09212419937771 | 0.688 | 0.973 | 8.8156805 |
| **PDE7A** | 6.8834 | -1.03727676623553 | 0.88 | 0.994 | 1.1272967 |
| **COL6A2** | 7.6443 | -1.89021683903712 | 0.702 | 0.955 | 1.2519105 |
| **FGD4** | 1.8441 | 1.12161415915649 | 0.981 | 0.708 | 3.0200954 |
| **HADHA** | 2.6832 | -0.820726271527982 | 0.989 | 1 | 4.3943985 |
| **RB1** | 2.9108 | -1.13124559372278 | 0.43 | 0.883 | 4.7670752 |
| **ZNRF1** | 3.3240 | -0.837110607500773 | 0.98 | 1 | 5.4438055 |
| **TNNC1** | 5.4256 | -1.41909047163621 | 0.117 | 0.704 | 8.8855609 |
| **HAS3** | 7.4334 | -1.86980762835375 | 0.314 | 0.827 | 1.2173711 |
| **CDYL** | 9.7463 | -1.07203680566748 | 0.752 | 0.978 | 1.5961618 |
| **VANGL2** | 1.0752 | -1.02532037087004 | 0.884 | 0.995 | 1.7609629 |
| **CNTRL** | 2.5800 | -1.25116240605794 | 0.434 | 0.891 | 4.2253402 |
| **ZIC3** | 3.5990 | -1.41300422672559 | 0.608 | 0.942 | 5.8941360 |
| **ARSB** | 6.2737 | -1.07197187625272 | 0.661 | 0.957 | 1.0274553 |
| **SLC35A5** | 7.4382 | 1.01675131881832 | 0.982 | 0.837 | 1.2181626 |
| **NGFR** | 8.0672 | -1.74206474737161 | 0.367 | 0.85 | 1.3211751 |
| **TINCR** | 2.1851 | 0.909468164419306 | 0.592 | 0.062 | 3.5786423 |

| | | | | | |
|---|---|---|---|---|---|
| **ZSWIM6** | 7.1616 | -1.35069107416909 | 0.162 | 0.745 | 1.17285818 |
| **NUDT8** | 8.4455 | 1.0220540477633 | 0.659 | 0.113 | 1.38312208 |
| **IL17RD** | 1.4404 | -1.4451391921404 | 0.845 | 0.986 | 2.35901653 |
| **SART1** | 1.7443 | 0.818211989025226 | 0.999 | 0.994 | 2.85665257 |
| **DMBX1** | 2.8143 | 1.01629655232603 | 0.544 | 0.037 | 4.60907964 |
| **PDHX** | 9.1000 | 0.894246154404708 | 0.995 | 0.956 | 1.49032161 |
| **PVR** | 9.8468 | 1.03827484111384 | 0.997 | 0.898 | 1.61261442 |
| **NT5C2** | 1.5846 | -1.14462124267903 | 0.867 | 0.988 | 2.59523356 |
| **TOMM34** | 2.4390 | -1.0183687740495 | 0.848 | 0.991 | 3.99447139 |
| **ZMYM3** | 3.7786 | -0.868219240765994 | 0.986 | 0.999 | 6.18828276 |
| **NRBP2** | 5.5829 | 1.34536405814764 | 0.76 | 0.235 | 9.14326930 |
| **DNMT1** | 1.1668 | -0.848360899738033 | 0.992 | 0.999 | 1.91092929 |
| **CMTM4** | 5.8694 | 0.980476585080389 | 0.989 | 0.932 | 9.61233219 |
| **SFRP2** | 1.1277 | -2.03940212314352 | 0.865 | 0.97 | 1.84690365 |
| **ZNHIT2** | 1.5636 | 1.13750369672511 | 0.952 | 0.7 | 2.56071880 |
| **INA** | 2.7370 | 1.24926722920403 | 0.94 | 0.614 | 4.48248412 |
| **STK38L** | 5.1804 | 1.03620588658943 | 0.993 | 0.911 | 8.48405450 |
| **VOPP1** | 1.4204 | -0.997662441834879 | 0.803 | 0.979 | 2.32630176 |
| **DDX49** | 1.5300 | 0.92709396825874 | 0.996 | 0.956 | 2.50576036 |
| **KLHDC7A** | 1.8763 | -1.49249807524618 | 0.043 | 0.634 | 3.07290757 |
| **AKT3** | 1.8850 | -1.25344827655002 | 0.401 | 0.88 | 3.08710783 |
| **CTSZ** | 3.5704 | -1.30340172465841 | 0.673 | 0.934 | 5.84731058 |
| **APP** | 5.6430 | -0.835457970393954 | 1 | 1 | 9.24169481 |
| **PTPRN** | 7.8947 | 1.05913810872091 | 0.568 | 0.055 | 1.29291697 |
| **CTSV** | 1.7041 | -1.54024274099599 | 0.772 | 0.971 | 2.79093918 |
| **BTBD2** | 2.2550 | -0.971552545272445 | 0.956 | 0.999 | 3.69317190 |
| **CAT** | 2.7218 | 1.08606181230737 | 0.943 | 0.64 | 4.45753785 |
| **SHE** | 3.8369 | 0.971098162697234 | 0.542 | 0.04 | 6.28376373 |
| **EXOSC4** | 5.7091 | 1.00792831002913 | 0.99 | 0.901 | 9.34983891 |
| **RASL10B** | 7.2335 | -1.33331971703465 | 0.138 | 0.717 | 1.18464265 |
| **FOXO4** | 1.6317 | -1.66901378529008 | 0.573 | 0.925 | 2.67230884 |
| **ELF3** | 6.3042 | 1.55696650881213 | 0.84 | 0.366 | 1.03245334 |
| **NSD2** | 6.6396 | -1.15759360414338 | 0.461 | 0.892 | 1.08736748 |
| **MCM8** | 1.1662 | -0.899116947976794 | 0.98 | 0.999 | 1.90989173 |

| | | | | | |
|---|---|---|---|---|---|
| **PSIP1** | 2.2692 | -0.875681146649165 | 0.98 | 0.998 | 3.71640512 |
| **TAOK3** | 2.2692 | 1.18366214654335 | 0.788 | 0.267 | 3.71642413 |
| **ZNF711** | 3.1691 | -1.21731143746758 | 0.462 | 0.905 | 5.19008395 |
| **HUNK** | 3.4978 | -1.25792964500251 | 0.356 | 0.862 | 5.72836433 |
| **PKD2** | 3.7971 | -1.18313375204968 | 0.553 | 0.94 | 6.21851657 |
| **AKAP1** | 7.8017 | 1.02996518253078 | 0.997 | 0.957 | 1.27769070 |
| **VAT1L** | 1.6932 | -2.25820778672627 | 0.052 | 0.644 | 2.77310697 |
| **PRKAR2A** | 1.7071 | 0.835322305602926 | 0.999 | 0.972 | 2.79577222 |
| **SEMA4B** | 2.1542 | 1.05501062719375 | 0.987 | 0.82 | 3.52794510 |
| **PDE3B** | 2.5566 | -1.20972151561612 | 0.156 | 0.728 | 4.18707469 |
| **STK17B** | 3.0836 | 1.16296294132102 | 0.885 | 0.444 | 5.05009237 |
| **SLC7A5** | 3.1557 | -1.23393793460317 | 0.993 | 1 | 5.16820610 |
| **CALD1** | 3.4459 | -2.02555085372091 | 0.997 | 1 | 5.64336003 |
| **MT-ND5** | 3.6764 | 1.52893073462023 | 0.958 | 0.75 | 6.02094224 |
| **CBR3** | 4.0855 | 1.32465230523417 | 0.887 | 0.408 | 6.69089985 |
| **KCNG1** | 4.3795 | -1.61579441290629 | 0.306 | 0.821 | 7.17245787 |
| **SKI** | 4.9644 | -0.850440136827192 | 0.971 | 1 | 8.13028383 |
| **RAB31** | 6.3325 | -1.28070752924441 | 0.278 | 0.802 | 1.03708247 |
| **ARHGDIG** | 1.0848 | 0.967461820902388 | 0.657 | 0.116 | 1.77672497 |
| **CATSPERZ** | 1.7833 | 0.888590271363979 | 0.585 | 0.068 | 2.92065613 |
| **TMEM150C** | 1.9006 | 1.06851670647554 | 0.613 | 0.083 | 3.11261320 |
| **CSF2RA** | 3.0064 | 1.25160319506515 | 0.677 | 0.144 | 4.92368160 |
| **NR5A2** | 3.7033 | 1.16968246898253 | 0.688 | 0.152 | 6.06494473 |
| **CALB1** | 3.9355 | 1.27551656720012 | 0.689 | 0.132 | 6.44526888 |
| **NCOA4** | 8.9875 | 0.923779418018763 | 0.996 | 0.931 | 1.47188735 |
| **CLDN7** | 1.1368 | 1.19460780024309 | 0.995 | 0.743 | 1.86178041 |
| **ZC4H2** | 1.5582 | -1.1157601035232 | 0.407 | 0.86 | 2.55194712 |
| **EPB41L4B** | 1.9881 | 1.15675559080471 | 0.928 | 0.556 | 3.25602397 |
| **PFKP** | 2.0418 | 0.854397691372246 | 1 | 0.997 | 3.34392329 |
| **GSPT1** | 3.6140 | 0.896724530745529 | 0.991 | 0.912 | 5.91880290 |
| **PLLP** | 3.6860 | 1.04158458054731 | 0.653 | 0.118 | 6.03662937 |
| **PCNX1** | 4.0693 | -0.975185272563934 | 0.861 | 0.99 | 6.66433878 |
| **GCNT1** | 4.4299 | 1.16952987566513 | 0.82 | 0.331 | 7.25486904 |
| **ARHGEF12** | 7.3925 | 0.93341434007471 | 0.99 | 0.907 | 1.21068427 |

| | | | | | |
|---|---|---|---|---|---|
| **TSPAN13** | 1.1059 | 1.02153421610371 | 0.988 | 0.857 | 1.8112898 |
| **TMEM97** | 1.0041 | -0.834929919306278 | 0.998 | 1 | 1.6445014 |
| **FZD8** | 2.0532 | 0.977049782898029 | 0.828 | 0.214 | 3.3625637 |
| **NET1** | 2.1050 | -1.32334111853024 | 0.851 | 0.984 | 3.4474711 |
| **NRK** | 2.1833 | -2.74043781286379 | 0.276 | 0.795 | 3.5756894 |
| **AGMAT** | 3.0201 | 0.809062612156116 | 0.535 | 0.039 | 4.9460851 |
| **RRAD** | 3.1658 | 1.72893157530788 | 0.511 | 0.031 | 5.1847943 |
| **SOCS3** | 4.3477 | -1.28551197841514 | 0.59 | 0.954 | 7.1202990 |
| **EPHB3** | 4.5110 | -1.71526154808312 | 0.104 | 0.683 | 7.3876892 |
| **EDNRB** | 8.0790 | -1.890743431731 | 0.229 | 0.766 | 1.3231064 |
| **ST6GALNAC** | 1.1879 | -1.21593912478968 | 0.068 | 0.648 | 1.9454461 |
| **PIWIL2** | 2.4904 | 0.877477307734985 | 0.469 | 0.009 | 4.0785502 |
| **ABCB7** | 4.6506 | -0.809700033344071 | 0.977 | 1 | 7.6162975 |
| **ANKRD10** | 1.5277 | -1.11305544173229 | 0.955 | 0.997 | 2.5019564 |
| **GLRX2** | 4.0581 | -1.1632082208585 | 0.185 | 0.748 | 6.6460549 |
| **SSR3** | 5.8844 | 0.872248704641875 | 0.999 | 0.99 | 9.6369405 |
| **WIPF3** | 7.7423 | 1.08480704345713 | 0.745 | 0.207 | 1.2679623 |
| **LARP1B** | 1.1159 | 1.04783585998315 | 0.95 | 0.687 | 1.8276437 |
| **OSTC** | 1.9098 | 0.834510309236167 | 0.999 | 0.977 | 3.1277610 |
| **C2CD5** | 7.3607 | 1.07384016018203 | 0.984 | 0.93 | 1.2054698 |
| **UHRF2** | 7.3659 | -1.22364851043055 | 0.306 | 0.819 | 1.2063210 |
| **OVOL2** | 8.3937 | 1.23241891991446 | 0.845 | 0.34 | 1.3746511 |
| **WNK3** | 1.0075 | -1.0991996601425 | 0.532 | 0.922 | 1.6500075 |
| **CCDC9** | 1.0285 | 0.83851818499335 | 0.999 | 0.982 | 1.6843883 |
| **SHF** | 1.2191 | -1.31463593503002 | 0.389 | 0.853 | 1.9965429 |
| **CABYR** | 2.8586 | 0.892703735869825 | 0.578 | 0.068 | 4.6815551 |
| **PPFIBP1** | 3.8801 | -1.21345778673139 | 0.481 | 0.918 | 6.3545029 |
| **PPARG** | 7.4980 | 0.937906747121586 | 0.565 | 0.062 | 1.2279632 |
| **TEC** | 6.5391 | 0.908636767040313 | 0.607 | 0.087 | 1.0709106 |
| **SLC25A13** | 7.1002 | -0.836657609575463 | 0.976 | 0.999 | 1.1628159 |
| **TRAF5** | 7.3632 | -1.24612541663098 | 0.16 | 0.733 | 1.2058865 |
| **EEA1** | 1.7911 | 1.00400012674504 | 0.976 | 0.768 | 2.9333470 |
| **MLLT11** | 1.9138 | -1.23524240346152 | 0.563 | 0.92 | 3.1342737 |
| **CCDC117** | 2.2860 | 0.987024737420483 | 0.955 | 0.673 | 3.7437925 |

| | | | | | |
|---|---|---|---|---|---|
| **CAPN13** | 3.3223 | -1.2093038962312 | 0.147 | 0.715 | 5.44105038 |
| **SMPDL3A** | 6.7095 | 1.07293986140986 | 0.698 | 0.174 | 1.09882220 |
| **SENP2** | 1.4238 | 0.867922445872987 | 0.999 | 0.992 | 2.33191351 |
| **EXT1** | 3.3548 | -1.13363758220627 | 0.784 | 0.98 | 5.49429148 |
| **GMPPA** | 7.5842 | 0.980680714883308 | 0.984 | 0.86 | 1.24207171 |
| **FLT1** | 8.1451 | -1.72285699838064 | 0.103 | 0.673 | 1.33393395 |
| **UQCC1** | 8.9188 | -0.898109800377877 | 0.919 | 0.996 | 1.46064115 |
| **SFT2D1** | 9.0039 | 1.01434273697728 | 0.915 | 0.563 | 1.47456964 |
| **IGLON5** | 9.2316 | -1.19577085708916 | 0.257 | 0.794 | 1.51186353 |
| **STK40** | 1.0715 | -1.05757626263474 | 0.675 | 0.959 | 1.75483078 |
| **DNTTIP1** | 6.2722 | -0.822611131472848 | 0.966 | 1 | 1.02719865 |
| **ABCB8** | 7.2182 | 0.938163055726005 | 0.986 | 0.891 | 1.18212531 |
| **AHDC1** | 1.1129 | -1.27033740946481 | 0.45 | 0.875 | 1.82265852 |
| **PINK1** | 2.2199 | 1.00650985816285 | 0.995 | 0.94 | 3.63555592 |
| **SNX12** | 2.3745 | -0.825408370856964 | 0.956 | 0.999 | 3.88887343 |
| **DACT2** | 2.5984 | 0.935027964160198 | 0.523 | 0.038 | 4.25553771 |
| **FAM102B** | 2.8586 | 1.2758470205446 | 0.925 | 0.644 | 4.68168792 |
| **ZNF518A** | 4.1852 | 0.852582296471511 | 0.998 | 0.945 | 6.85411773 |
| **DTNA** | 5.4150 | -1.95630597869913 | 0.075 | 0.645 | 8.86818158 |
| **COL4A6** | 8.4249 | -1.26984469323105 | 0.081 | 0.646 | 1.37976039 |
| **FBXL15** | 1.2791 | 1.03155887787849 | 0.954 | 0.699 | 2.09478423 |
| **HIPK1** | 1.3403 | -0.922318657326403 | 0.82 | 0.978 | 2.19506360 |
| **STC2** | 1.5085 | 1.39845485505052 | 0.988 | 0.924 | 2.47059491 |
| **NAGK** | 1.5802 | 1.03972324222105 | 0.997 | 0.955 | 2.58795631 |
| **EPHX1** | 1.8899 | -1.60481312962013 | 0.933 | 0.991 | 3.09517549 |
| **ENPP2** | 2.8942 | 1.08962140672851 | 0.694 | 0.166 | 4.73995615 |
| **CCNA1** | 3.1441 | 1.45772840385815 | 0.528 | 0.046 | 5.14918442 |
| **OSBPL10** | 5.9447 | 1.11181355165691 | 0.973 | 0.764 | 9.73572015 |
| **AGBL5** | 8.7067 | -1.00442970907412 | 0.79 | 0.978 | 1.42590163 |
| **CGREF1** | 1.1758 | 1.40544786094386 | 0.625 | 0.121 | 1.92573755 |
| **NLGN4X** | 2.2040 | -1.98350239599132 | 0.335 | 0.79 | 3.60955479 |
| **NOTCH2** | 5.3951 | -1.08308126928029 | 0.566 | 0.922 | 8.83560770 |
| **CFL2** | 1.0484 | -1.05716294122656 | 0.695 | 0.95 | 1.71699860 |
| **SERINC5** | 1.3014 | 0.839136786388541 | 0.999 | 0.987 | 2.13137970 |

| | | | | | |
|---|---|---|---|---|---|
| **CAMK2D** | 1.6314 | -1.36411166360506 | 0.244 | 0.766 | 2.67180930 |
| **PREX2** | 1.7914 | -1.32020492695345 | 0.326 | 0.818 | 2.93383979 |
| **NPC2** | 6.2413 | -0.864412719087812 | 1 | 1 | 1.02214186 |
| **H2AFY** | 8.4574 | -1.15025737082228 | 0.408 | 0.866 | 1.38508288 |
| **TRIM24** | 1.2280 | -0.997854920115656 | 0.986 | 1 | 2.01117042 |
| **SLC22A31** | 1.2286 | 1.17325298735323 | 0.69 | 0.177 | 2.01220698 |
| **LIPA** | 1.5750 | -1.17106656640518 | 0.501 | 0.896 | 2.57950417 |
| **MPPED2** | 2.4370 | 0.840968091310115 | 0.446 | 0.007 | 3.99123673 |
| **ZSCAN2** | 5.0172 | 1.14711205985875 | 0.982 | 0.892 | 8.21672181 |
| **LPIN1** | 6.4068 | 1.10227143124327 | 0.839 | 0.323 | 1.04925294 |
| **SCML2** | 8.2314 | -1.03142874957244 | 0.547 | 0.921 | 1.34807135 |
| **WDR44** | 9.6571 | 1.00443714639895 | 0.988 | 0.891 | 1.58155511 |
| **CCNQ** | 1.0808 | 0.99713205783027 | 0.974 | 0.809 | 1.77012564 |
| **ZNF229** | 1.1131 | 1.08495950777686 | 0.759 | 0.255 | 1.82301158 |
| **ESR1** | 2.3911 | 0.951369389547085 | 0.999 | 0.971 | 3.91599590 |
| **DMKN** | 5.1240 | -0.978906548467167 | 0.822 | 0.996 | 8.39158511 |
| **WDR46** | 8.9693 | 0.854090611459295 | 0.999 | 0.997 | 1.46891247 |
| **RET** | 2.5655 | -1.40641481476632 | 0.32 | 0.826 | 4.20164137 |
| **CXCL14** | 3.0159 | -2.65366189836965 | 0.409 | 0.85 | 4.93925610 |
| **PRR36** | 3.1727 | -1.46248636232233 | 0.192 | 0.715 | 5.19607079 |
| **TMEM185A** | 4.0413 | -1.07584727209301 | 0.837 | 0.99 | 6.61845063 |
| **FIGNL2** | 1.2643 | -1.21789852369085 | 0.114 | 0.672 | 2.07064879 |
| **SACS** | 1.2811 | -1.04067150850622 | 0.882 | 0.987 | 2.09813650 |
| **P3H2** | 1.8232 | -2.28201456972357 | 0.035 | 0.593 | 2.98596291 |
| **IRX2** | 2.0790 | -1.4884177975891 | 0.623 | 0.947 | 3.40489793 |
| **CKAP4** | 2.0848 | -0.806139252382442 | 0.995 | 1 | 3.41442828 |
| **TRIM61** | 1.0674 | 0.980751478244613 | 0.731 | 0.215 | 1.74814801 |
| **NTN4** | 1.1795 | 0.84811366874821 | 0.599 | 0.077 | 1.93175241 |
| **INPP5A** | 2.3928 | -0.845397240361947 | 0.942 | 0.994 | 3.91871005 |
| **FES** | 3.3692 | 0.97189700511804 | 0.84 | 0.345 | 5.51780245 |
| **GPNMB** | 4.7134 | 1.18893686583743 | 0.779 | 0.252 | 7.71927705 |
| **KLRG2** | 4.8480 | 0.96683926404731 | 0.996 | 0.925 | 7.93961684 |
| **EIF4E** | 1.6645 | 0.814295612839092 | 0.995 | 0.952 | 2.72609733 |
| **MANF** | 4.9357 | 1.03883134043797 | 1 | 1 | 8.08332925 |

| | | | | | |
|---|---|---|---|---|---|
| **SOGA1** | 2.2835 | -1.06855457731796 | 0.794 | 0.982 | 3.7397056 |
| **EIF1AD** | 2.3277 | 0.86389681142796 | 0.995 | 0.936 | 3.8121230 |
| **NGEF** | 3.2953 | 0.946799903230696 | 0.504 | 0.038 | 5.3968643 |
| **ANKRD18A** | 3.3816 | -1.25656375183452 | 0.132 | 0.683 | 5.5381687 |
| **MAPK13** | 4.9786 | 1.0654052710991 | 0.735 | 0.222 | 8.1535292 |
| **CYCS** | 1.3605 | 1.01376242722386 | 0.927 | 0.641 | 2.2282525 |
| **KRT8** | 3.2871 | -1.00961041188328 | 0.999 | 1 | 5.3833123 |
| **ZNF730** | 5.5125 | -1.05215424738865 | 0.796 | 0.979 | 9.0278529 |
| **PPT1** | 1.4055 | -0.824870574429246 | 0.983 | 1 | 2.3018504 |
| **CMPK1** | 2.3658 | 0.814936905021903 | 0.996 | 0.937 | 3.8745197 |
| **EDEM1** | 2.8253 | 1.02240156986799 | 0.983 | 0.891 | 4.6271098 |
| **NOP16** | 3.3356 | 0.836561949434738 | 1 | 0.999 | 5.4627794 |
| **SSBP3** | 5.0757 | -0.919238967335888 | 0.821 | 0.986 | 8.3125685 |
| **SH3BGRL** | 5.9816 | -0.980114166693518 | 0.777 | 0.968 | 9.7960874 |
| **PICALM** | 6.7564 | 0.918744651277825 | 1 | 0.998 | 1.1064987 |
| **RASSF9** | 8.8065 | -1.54039820069475 | 0.108 | 0.659 | 1.4422464 |
| **LEFTY2** | 2.4616 | 1.74533646969731 | 0.992 | 0.945 | 4.0313626 |
| **TFPI2** | 2.5636 | 1.2994512888306 | 0.893 | 0.515 | 4.1985189 |
| **RCOR2** | 2.6768 | -1.0506899009015 | 0.798 | 0.977 | 4.3838926 |
| **CKS1B** | 2.7136 | 0.88860952465849 | 1 | 0.983 | 4.4440758 |
| **BMP4** | 4.6403 | -1.85886188512776 | 0.43 | 0.86 | 7.5994936 |
| **RAVER2** | 6.1733 | -1.03969117128787 | 0.463 | 0.879 | 1.0110092 |
| **MFSD13A** | 6.9305 | 1.07174696489309 | 0.787 | 0.293 | 1.1350195 |
| **CDHR1** | 9.9005 | 1.36710006159419 | 0.813 | 0.356 | 1.6214058 |
| **ENC1** | 1.1409 | -1.90172977314479 | 0.431 | 0.88 | 1.8684600 |
| **ARRDC2** | 1.6550 | 1.09063643240503 | 0.815 | 0.33 | 2.7105367 |
| **ATP5PF** | 1.6953 | 0.804779871147359 | 0.995 | 0.982 | 2.7765299 |
| **LRP1** | 2.4087 | -1.05700587396588 | 0.833 | 0.988 | 3.9447953 |
| **PRPSAP1** | 2.4806 | 0.922408760253059 | 0.995 | 0.969 | 4.0625193 |
| **LRP8** | 2.8526 | -1.36197269780038 | 0.834 | 0.972 | 4.6718473 |
| **PPP2R3A** | 3.0908 | -1.0178490384575 | 0.491 | 0.896 | 5.0618320 |
| **PTPRR** | 3.8594 | 0.804663192275266 | 0.453 | 0.016 | 6.3206019 |
| **TRO** | 7.7557 | -1.05101050517279 | 0.664 | 0.95 | 1.2701609 |
| **MBOAT1** | 8.7279 | 1.05920385344361 | 0.917 | 0.517 | 1.4293785 |

| | | | | | |
|---|---|---|---|---|---|
| **IDS** | 1.0526 | -1.02856475648917 | 0.528 | 0.913 | 1.7239848 |
| **FRMD4A** | 1.2013 | -1.23720069846835 | 0.153 | 0.696 | 1.9675321 |
| **EVL** | 1.2496 | -0.836722796746799 | 0.955 | 1 | 2.0465341 |
| **CAP2** | 4.3727 | -1.08683213747703 | 0.673 | 0.947 | 7.1613040 |
| **TTI1** | 2.0825 | -0.858165369477534 | 0.924 | 0.994 | 3.4105629 |
| **ARHGAP8** | 6.2130 | 1.00955173938535 | 0.959 | 0.697 | 1.0175052 |
| **AHCYL2** | 1.3189 | 1.04939666416764 | 0.974 | 0.848 | 2.1600823 |
| **ADORA2B** | 1.7420 | 1.01223267177858 | 0.75 | 0.245 | 2.8529439 |
| **PC** | 2.4122 | 1.15841657054258 | 0.864 | 0.46 | 3.9506145 |
| **ESCO1** | 2.6941 | 0.870592294959216 | 0.993 | 0.905 | 4.4122880 |
| **MSL2** | 4.5378 | -0.943164868106547 | 0.732 | 0.974 | 7.4316597 |
| **TPD52** | 7.9829 | 0.831240419052316 | 0.999 | 0.981 | 1.3073639 |
| **RNF145** | 1.1389 | -1.03067097219482 | 0.42 | 0.862 | 1.8653114 |
| **ZNF300** | 1.1829 | 0.992226171768281 | 0.794 | 0.284 | 1.9372379 |
| **KIDINS220** | 1.6594 | -0.930723702276974 | 0.737 | 0.965 | 2.7176479 |
| **ZSWIM9** | 1.6965 | -1.05767725200419 | 0.172 | 0.71 | 2.7783584 |
| **RBBP9** | 1.8744 | 1.04131882554514 | 0.976 | 0.88 | 3.0697098 |
| **NONO** | 3.5803 | 0.932287283434082 | 0.987 | 0.901 | 5.8635133 |
| **COL2A1** | 4.3206 | -1.88818593599696 | 0.035 | 0.575 | 7.0759875 |
| **MAK16** | 4.8490 | 0.813730084046542 | 0.999 | 0.995 | 7.9412267 |
| **GATAD2B** | 5.1810 | -0.914616009587575 | 0.796 | 0.976 | 8.4849244 |
| **PGAM1** | 6.6913 | 0.813375786530141 | 0.994 | 0.941 | 1.0958465 |
| **PDE12** | 8.2763 | 0.977614143985245 | 0.889 | 0.497 | 1.3554258 |
| **GCH1** | 1.5709 | 1.06389031167414 | 0.827 | 0.369 | 2.5727743 |
| **NIN** | 1.5893 | -1.09426205875784 | 0.623 | 0.941 | 2.6028528 |
| **ADAMTS19** | 2.1398 | -1.52260665039994 | 0.041 | 0.579 | 3.5044859 |
| **MYO5A** | 2.5506 | 0.997508833694106 | 0.947 | 0.717 | 4.1771492 |
| **SNW1** | 3.6952 | 0.971728201797367 | 0.961 | 0.772 | 6.0517227 |
| **COMT** | 4.6839 | -1.08172516271111 | 0.521 | 0.899 | 7.6708831 |
| **USP13** | 4.9090 | 0.941404506298085 | 0.985 | 0.932 | 8.0395024 |
| **SDC1** | 7.4121 | -1.18807590497582 | 0.228 | 0.754 | 1.2138796 |
| **FAM217B** | 1.0055 | -0.96750982368226 | 0.673 | 0.944 | 1.6467907 |
| **RASGRP2** | 1.5949 | 1.32171008744515 | 0.828 | 0.37 | 2.6119720 |
| **MAGI1** | 1.6831 | -1.30139299608912 | 0.542 | 0.888 | 2.7565106 |

| | | | | | |
|---|---|---|---|---|---|
| **TBC1D8** | 3.1030 | 1.02609747912378 | 0.8 | 0.321 | 5.0818191 |
| **NCAPD2** | 3.3597 | -1.0464630309023 | 0.956 | 0.998 | 5.5022924 |
| **MORF4L2** | 1.2858 | 0.841717132121076 | 0.996 | 0.949 | 2.1058465 |
| **CTPS2** | 1.8701 | -0.8097081215585 | 0.929 | 0.996 | 3.0627700 |
| **ATP5MD** | 1.9182 | 0.991720812295479 | 0.785 | 0.322 | 3.1414640 |
| **PGM2** | 1.9802 | 1.06783947625122 | 0.98 | 0.871 | 3.2431062 |
| **LAMA5** | 2.3034 | -1.09246900691524 | 0.981 | 0.996 | 3.7724164 |
| **CRABP1** | 3.3477 | -1.82287294840834 | 0.472 | 0.879 | 5.4825857 |
| **LAMB2** | 1.2820 | -1.22834540410843 | 0.568 | 0.904 | 2.0995688 |
| **IPMK** | 1.4307 | 0.939238389227134 | 0.97 | 0.787 | 2.3430902 |
| **PRMT3** | 1.9439 | 0.833104134450313 | 0.993 | 0.937 | 3.1835674 |
| **ETNK1** | 2.0176 | -0.95707788032855 | 0.77 | 0.973 | 3.3043814 |
| **PDE9A** | 2.3404 | -1.18907531856985 | 0.157 | 0.689 | 3.8329981 |
| **UROD** | 3.3089 | -0.803721336167421 | 0.999 | 1 | 5.4190148 |
| **COL1A1** | 3.8281 | -1.41936079521708 | 0.722 | 0.957 | 6.2694313 |
| **PLAAT5** | 4.1752 | -1.29924641671556 | 0.376 | 0.828 | 6.8378851 |
| **LOXL1** | 1.1431 | -1.33745446955217 | 0.417 | 0.842 | 1.8721865 |
| **CNTFR** | 1.4419 | -2.21327194646784 | 0.236 | 0.729 | 2.3614673 |
| **TTYH1** | 1.6732 | 1.15888666003248 | 0.552 | 0.076 | 2.7402842 |
| **MIPOL1** | 6.4443 | -1.05635156603295 | 0.263 | 0.766 | 1.0553947 |
| **TNFRSF21** | 6.4869 | -1.31218993210593 | 0.963 | 0.996 | 1.0623716 |
| **PALM2-AKAI** | 2.4511 | -1.33187899953229 | 0.927 | 0.992 | 4.0143104 |
| **AMER1** | 2.5061 | -1.03379412539493 | 0.612 | 0.928 | 4.1042583 |
| **TIMM8A** | 2.9597 | 0.889385732263415 | 0.997 | 0.975 | 4.8471210 |
| **ABHD12B** | 5.6262 | 0.896849565333256 | 0.645 | 0.133 | 9.2141624 |
| **ST5** | 7.3489 | -0.97895138559996 | 0.766 | 0.973 | 1.2035385 |
| **ASS1** | 7.8743 | 1.17138514626802 | 0.993 | 0.948 | 1.2895843 |
| **EEF1AKMT4** | 1.1839 | 0.940494486989919 | 0.973 | 0.772 | 1.9389883 |
| **SESTD1** | 2.3186 | 0.879156247071066 | 0.991 | 0.913 | 3.7972670 |
| **DLK1** | 3.1904 | 0.982664721244825 | 0.635 | 0.138 | 5.2250519 |
| **COPRS** | 3.8536 | -0.848028170376191 | 0.94 | 0.996 | 6.3110994 |
| **ESRP1** | 5.9143 | 0.896302407641127 | 0.999 | 0.877 | 9.6859081 |
| **TGFBR3** | 9.9390 | 1.07301145734383 | 0.783 | 0.305 | 1.6277256 |
| **ARHGEF2** | 2.2863 | -0.973476255533809 | 0.709 | 0.968 | 3.7444328 |

| | | | | | |
|---|---|---|---|---|---|
| **XXYLT1** | 2.4882 | -0.903723017627069 | 0.799 | 0.969 | 4.0749316 |
| **ROR1** | 3.2442 | -1.31197024143661 | 0.334 | 0.802 | 5.3131512 |
| **RCBTB1** | 1.0502 | 1.03374260008679 | 0.991 | 0.971 | 1.7199345 |
| **DUSP16** | 1.0600 | 1.20605801270833 | 0.961 | 0.763 | 1.7360883 |
| **TBX22** | 2.4423 | -1.34362338231505 | 0.061 | 0.595 | 3.9998814 |
| **VEZF1** | 3.0422 | -0.918667380237541 | 0.857 | 0.985 | 4.9823626 |
| **SGCB** | 5.6754 | -0.963707361415447 | 0.679 | 0.95 | 9.2947078 |
| **TSPAN4** | 9.5968 | -0.878692069909325 | 0.944 | 0.996 | 1.5716775 |
| **SLC8B1** | 2.1066 | 1.07813914354116 | 0.803 | 0.36 | 3.4500158 |
| **BST2** | 2.3422 | -1.63432135483655 | 0.357 | 0.832 | 3.8359774 |
| **TTC7B** | 3.7416 | -0.952282033264685 | 0.752 | 0.964 | 6.1277494 |
| **STARD3NL** | 4.3113 | -0.851197910114329 | 0.883 | 0.993 | 7.0607557 |
| **ECSCR** | 4.3483 | -1.04132846575283 | 0.038 | 0.563 | 7.1212810 |
| **CDC42EP1** | 5.8252 | -1.11441309964258 | 0.113 | 0.654 | 9.5400894 |
| **COTL1** | 8.9720 | -1.11126096633299 | 0.979 | 0.997 | 1.4693460 |
| **TMEM131L** | 1.3589 | 0.971371090527694 | 0.968 | 0.798 | 2.2256197 |
| **CAMK1D** | 1.8574 | 0.905002207364477 | 0.589 | 0.107 | 3.0419618 |
| **NRIP3** | 2.1792 | 0.826127099785081 | 0.532 | 0.07 | 3.5689283 |
| **HK2** | 2.7613 | 0.890320564652084 | 0.999 | 0.985 | 4.5223360 |
| **KIF9** | 2.8698 | 1.09231605824749 | 0.906 | 0.588 | 4.6999434 |
| **FOXD3** | 3.2250 | 1.37548479805057 | 0.749 | 0.271 | 5.2817104 |
| **NEK9** | 8.1989 | -0.847732617950244 | 0.877 | 0.985 | 1.3427339 |
| **TSC22D3** | 8.6253 | -1.31612179293639 | 0.532 | 0.902 | 1.4125798 |
| **HRK** | 8.6301 | 1.02110734481568 | 0.641 | 0.156 | 1.4133668 |
| **BLCAP** | 1.1230 | -0.881777942712884 | 0.881 | 0.994 | 1.8392054 |
| **SPSB2** | 3.2680 | 1.04335516173369 | 0.938 | 0.649 | 5.3521162 |
| **DUSP6** | 8.3849 | 1.10089356919336 | 0.532 | 0.069 | 1.3732087 |
| **ACTN1** | 1.0201 | -0.850455495001098 | 1 | 1 | 1.6706451 |
| **PRKD3** | 2.2242 | -0.979648955878869 | 0.927 | 0.995 | 3.6427242 |
| **BOD1L1** | 2.8186 | 0.823513268075102 | 0.995 | 0.992 | 4.6161035 |
| **NLRP12** | 3.0482 | 1.15949449951047 | 0.867 | 0.475 | 4.9920647 |
| **CDH4** | 4.9026 | -1.08813923436141 | 0.053 | 0.578 | 8.0290361 |
| **CEBPZOS** | 6.3321 | 0.934835814647617 | 0.978 | 0.832 | 1.0370106 |
| **PIP5K1C** | 1.8104 | -0.931876270097856 | 0.796 | 0.97 | 2.9650350 |

| | | | | | |
|---|---|---|---|---|---|
| **WDR35** | 1.9492 | -1.05300777716952 | 0.258 | 0.748 | 3.1922437 |
| **GPC1** | 3.4166 | -0.901904994795222 | 0.89 | 0.989 | 5.5954969 |
| **CRISPLD2** | 8.1502 | 1.03884869400295 | 0.716 | 0.225 | 1.3347621 |
| **MEGF8** | 9.0779 | -1.01529643122844 | 0.489 | 0.878 | 1.4867029 |
| **SYNGAP1** | 1.0178 | -1.24289697696823 | 0.318 | 0.78 | 1.6669323 |
| **AFF2** | 1.2767 | 1.16672641781708 | 0.72 | 0.235 | 2.0908625 |
| **FBN3** | 2.7794 | -1.09794948043591 | 0.449 | 0.875 | 4.5518902 |
| **SLC2A3** | 3.1373 | -1.18544126252488 | 0.783 | 0.966 | 5.1380254 |
| **BGN** | 3.9313 | -1.90583230438123 | 0.219 | 0.718 | 6.4383980 |
| **PLAGL2** | 4.0804 | -0.942990981665994 | 0.649 | 0.93 | 6.6825057 |
| **ZRANB3** | 4.8968 | -1.03656734354965 | 0.26 | 0.755 | 8.0195698 |
| **SEPHS1** | 8.5618 | -0.989372803767955 | 0.997 | 1 | 1.4021822 |
| **MSI2** | 1.5561 | -0.964097665975998 | 0.727 | 0.959 | 2.5485224 |
| **NCS1** | 3.3109 | -0.95564204389139 | 0.725 | 0.959 | 5.4223163 |
| **GPC4** | 3.7135 | -1.15041036715421 | 0.984 | 1 | 6.0816565 |
| **HOXA7** | 8.3131 | -0.864738316010479 | 0.033 | 0.547 | 1.3614510 |
| **ATP1A2** | 8.4108 | -1.10629793577524 | 0.035 | 0.549 | 1.3774495 |
| **LRRC42** | 1.8648 | -0.869129313485575 | 0.849 | 0.982 | 3.0540539 |
| **CCND2** | 2.5830 | -1.35397780329735 | 0.771 | 0.976 | 4.2301826 |
| **ATL3** | 6.7387 | 0.893611046901333 | 0.961 | 0.782 | 1.1035972 |
| **ARID5B** | 1.0403 | 0.826035279034287 | 0.397 | 0.005 | 1.7037760 |
| **RIC8B** | 1.0923 | -1.03325051882113 | 0.348 | 0.807 | 1.7889211 |
| **NRARP** | 1.0987 | -1.2702892935576 | 0.361 | 0.8 | 1.7994351 |
| **AP1M1** | 1.1129 | -0.857030413879175 | 0.915 | 0.992 | 1.8226769 |
| **PIK3R5** | 5.7774 | 0.864305949014734 | 0.662 | 0.165 | 9.4616508 |
| **TMEM229B** | 6.1283 | 1.14980287794108 | 0.699 | 0.202 | 1.0036317 |
| **TP53BP2** | 1.0577 | -0.884184898739783 | 0.812 | 0.971 | 1.7322281 |
| **TBKBP1** | 1.1220 | -0.985352997769901 | 0.563 | 0.909 | 1.8376204 |
| **TCF7L2** | 1.6770 | -1.12398780576935 | 0.681 | 0.941 | 2.7465150 |
| **JUND** | 3.5360 | -0.848938333862483 | 0.999 | 1 | 5.7909936 |
| **SNX3** | 8.1703 | 0.855066155706591 | 0.973 | 0.775 | 1.3380636 |
| **PLOD1** | 4.8553 | 0.803317666212971 | 0.999 | 0.988 | 7.9516410 |
| **LDB2** | 5.2871 | -1.5126973310077 | 0.544 | 0.882 | 8.6587587 |
| **PGM1** | 6.8769 | -1.41587074315041 | 0.869 | 0.997 | 1.1262356 |

| | | | | | |
|---|---|---|---|---|---|
| **PLEKHG2** | 3.7358 | -0.990308046198213 | 0.449 | 0.85 | 6.11814839 |
| **FOXK1** | 3.8401 | -0.920112348983163 | 0.755 | 0.967 | 6.28902898 |
| **ZNF878** | 5.1301 | 0.948341311651455 | 0.587 | 0.118 | 8.40160820 |
| **ANGPTL2** | 6.3705 | -0.98838026247146 | 0.042 | 0.55 | 1.04330688 |
| **FAM118B** | 7.4122 | -0.862666081842931 | 0.781 | 0.97 | 1.21390082 |
| **RELL1** | 1.6453 | 0.918138129852032 | 0.964 | 0.775 | 2.69457625 |
| **PBX4** | 3.0867 | 0.90146524215175 | 0.627 | 0.144 | 5.05516950 |
| **LRRN2** | 3.8040 | -1.15477754928205 | 0.126 | 0.652 | 6.22981678 |
| **MBTD1** | 3.8492 | -0.848245924482089 | 0.875 | 0.989 | 6.30391279 |
| **PIR** | 5.4627 | -1.00077642440052 | 0.695 | 0.943 | 8.94637713 |
| **ST6GALNAC** | 5.6321 | -1.05026885061493 | 0.354 | 0.806 | 9.22372941 |
| **IGFBP7** | 7.0182 | -1.7807959284364 | 0.161 | 0.67 | 1.14938531 |
| **EIF3I** | 4.2699 | 0.919813279788069 | 0.843 | 0.452 | 6.99295307 |
| **UBXN7** | 5.3261 | -0.809427374427236 | 0.866 | 0.989 | 8.72263395 |
| **STK26** | 5.4398 | -0.839797820301839 | 0.946 | 0.995 | 8.90888681 |
| **MYORG** | 6.5441 | -1.13862702884896 | 0.284 | 0.752 | 1.07173105 |
| **EZH2** | 8.1231 | -0.843816815555742 | 0.961 | 0.994 | 1.33032281 |
| **KDM4C** | 9.3605 | 1.01308683574603 | 0.962 | 0.784 | 1.53298445 |
| **NECAB3** | 5.7991 | -0.994809507714136 | 0.565 | 0.898 | 9.49725097 |
| **DLG1** | 6.6238 | -0.809702971686357 | 0.917 | 0.99 | 1.08478472 |
| **CCND1** | 1.3197 | -0.805233675597411 | 0.997 | 0.999 | 2.16129570 |
| **BNIP3L** | 3.9427 | -0.803378985826053 | 0.982 | 1 | 6.45712343 |
| **FRMD6** | 4.5752 | 1.11721726757211 | 0.946 | 0.728 | 7.49290273 |
| **GAS7** | 5.8517 | 0.856340736281152 | 1 | 0.952 | 9.58347093 |
| **SMYD2** | 5.8842 | -0.912862157568246 | 0.911 | 0.99 | 9.63655480 |
| **PAK2** | 7.9857 | 0.825603701119074 | 0.976 | 0.838 | 1.30781829 |
| **RGS2** | 9.7452 | 1.17855767098721 | 0.865 | 0.491 | 1.59598630 |
| **ARFGEF1** | 1.7360 | 0.827414842339933 | 0.996 | 0.958 | 2.84317260 |
| **SLCO4C1** | 1.7530 | 0.94129590759916 | 0.954 | 0.603 | 2.87097120 |
| **AEBP1** | 4.7061 | -1.06019756193732 | 0.521 | 0.884 | 7.70719364 |
| **LIMD2** | 8.2057 | -0.975937143193843 | 0.86 | 0.982 | 1.34386270 |
| **AGPAT2** | 2.5424 | 0.932372682960223 | 0.989 | 0.957 | 4.16373208 |
| **UBE2Q2** | 5.6382 | 0.811917324698239 | 0.993 | 0.922 | 9.23382859 |
| **AC007906.2** | 6.3091 | -1.09073910962095 | 0.079 | 0.584 | 1.03324924 |

| | | | | | |
|---|---|---|---|---|---|
| **AP1S1** | 1.7187 | 0.848502684353703 | 0.998 | 0.966 | 2.8148058 |
| **NKAIN1** | 1.9100 | -1.22093543752031 | 0.243 | 0.72 | 3.1280647 |
| **SNRPE** | 2.6065 | 0.824720429296762 | 0.988 | 0.913 | 4.2687160 |
| **ZNF625** | 9.9035 | 0.801109322046537 | 0.993 | 0.931 | 1.6219121 |
| **GSTM3** | 1.3424 | -1.35989438361512 | 0.21 | 0.694 | 2.1985645 |
| **HSPB8** | 1.4570 | -1.41923951737017 | 0.075 | 0.591 | 2.3862020 |
| **ZNF618** | 1.7418 | -0.891575205765967 | 0.787 | 0.964 | 2.8526675 |
| **FNDC10** | 1.7500 | 1.0475109325031 | 0.867 | 0.485 | 2.8660190 |
| **MMP25** | 1.3719 | -1.22329414004546 | 0.278 | 0.744 | 2.2468565 |
| **SLC16A9** | 2.1664 | 1.06923375861553 | 0.84 | 0.455 | 3.5479748 |
| **SPTY2D1** | 2.8820 | 0.825611050540195 | 0.982 | 0.869 | 4.7198996 |
| **TBC1D20** | 4.4389 | -0.875763113559159 | 0.824 | 0.975 | 7.2696228 |
| **CDC25B** | 8.0498 | -0.90034271137211 | 0.957 | 0.996 | 1.3183315 |
| **NUPR1** | 1.0416 | 1.30222325394358 | 0.671 | 0.208 | 1.7059291 |
| **DDIT4** | 2.4514 | 1.42861174843943 | 0.974 | 0.821 | 4.0147522 |
| **SYT6** | 2.8330 | -1.11357760414257 | 0.202 | 0.695 | 4.6396400 |
| **NECTIN1** | 4.1498 | -0.986884679993929 | 0.935 | 0.987 | 6.7961941 |
| **UBE2G2** | 6.7348 | -0.834910578236836 | 0.819 | 0.976 | 1.1029710 |
| **CHST11** | 2.7720 | -1.08985617292118 | 0.777 | 0.971 | 4.5397677 |
| **RASGRP4** | 1.6098 | 0.806441483816945 | 0.602 | 0.134 | 2.6364695 |
| **KCTD20** | 2.9122 | -0.878824929515831 | 0.85 | 0.984 | 4.7693623 |
| **CIZ1** | 3.2623 | -0.907790520060188 | 0.626 | 0.929 | 5.3427463 |
| **CREBL2** | 4.7235 | 0.867509700199915 | 0.971 | 0.877 | 7.7357281 |
| **AXIN2** | 1.5405 | -1.03526979008283 | 0.748 | 0.966 | 2.5229544 |
| **FAM171A1** | 1.9059 | -0.944763237684269 | 0.542 | 0.875 | 3.1214451 |
| **SLC35F2** | 2.3737 | 0.88238333507552 | 0.971 | 0.813 | 3.8875712 |
| **TNS2** | 8.7892 | -1.27047161525158 | 0.061 | 0.55 | 1.4394129 |
| **CHD6** | 8.9384 | -0.884183401301545 | 0.771 | 0.959 | 1.4638423 |
| **ARFGAP3** | 2.3082 | 0.914384734447652 | 0.934 | 0.668 | 3.7802469 |
| **NRIP1** | 2.5245 | -0.98500141273072 | 0.642 | 0.93 | 4.1344174 |
| **LINGO1** | 3.0003 | 0.881923049802214 | 0.99 | 0.854 | 4.9136726 |
| **ZNF385A** | 4.0291 | -1.04377900267496 | 0.258 | 0.736 | 6.5986140 |
| **MED19** | 5.1648 | 0.844661362328032 | 0.932 | 0.709 | 8.4585484 |
| **MARVELD3** | 6.4446 | 0.845927840135124 | 0.688 | 0.2 | 1.0554419 |

| | | | | | |
|---|---|---|---|---|---|
| **HEG1** | 1.1107 | -1.15595430731921 | 0.226 | 0.704 | 1.8190459 |
| **TMC6** | 1.4977 | -1.14888507756432 | 0.669 | 0.91 | 2.4528851 |
| **PHTF1** | 4.3644 | -0.905191515967333 | 0.764 | 0.959 | 7.1476172 |
| **FBXL16** | 1.0074 | 0.851609798347744 | 0.603 | 0.142 | 1.6499381 |
| **PRAP1** | 1.0762 | 1.00326091372138 | 0.492 | 0.067 | 1.7626377 |
| **XIAP** | 1.0916 | 0.892516876417656 | 0.915 | 0.635 | 1.7877640 |
| **RBM7** | 1.1600 | 0.882410257620935 | 0.944 | 0.726 | 1.8997809 |
| **EPDR1** | 1.4022 | -0.904164503971026 | 0.046 | 0.532 | 2.2964662 |
| **COLEC12** | 1.6905 | -2.17341078266785 | 0.145 | 0.626 | 2.7685323 |
| **C2CD4C** | 1.8469 | 0.921897523906539 | 0.742 | 0.275 | 3.0247367 |
| **FOXO1** | 1.9447 | -1.3677779707447 | 0.675 | 0.93 | 3.1848988 |
| **PCDH1** | 2.1859 | -1.33686326304957 | 0.915 | 0.949 | 3.5799026 |
| **RAI14** | 2.4646 | -0.965324317832224 | 0.705 | 0.944 | 4.0363208 |
| **ZFP36L2** | 3.5099 | -1.01205857512086 | 0.946 | 0.994 | 5.7482052 |
| **NETO2** | 1.2088 | 0.99844764740241 | 0.959 | 0.781 | 1.9798153 |
| **P2RY1** | 1.5310 | -1.30446632417381 | 0.098 | 0.586 | 2.5074668 |
| **DDR1** | 1.6094 | -0.856714689626711 | 0.88 | 0.985 | 2.6357279 |
| **PTPRZ1** | 1.7306 | -1.75131982703987 | 0.554 | 0.863 | 2.8342923 |
| **NKD1** | 2.2656 | -1.53663989824304 | 0.199 | 0.664 | 3.7104287 |
| **KIF3B** | 3.1294 | -0.844398170220841 | 0.742 | 0.956 | 5.1250461 |
| **ZBTB12** | 3.3731 | -0.966125967702308 | 0.129 | 0.623 | 5.5241572 |
| **USP3** | 4.4060 | -0.938476942569418 | 0.067 | 0.558 | 7.2157646 |
| **BCL7C** | 4.5922 | -1.02784659813042 | 0.91 | 0.992 | 7.5208025 |
| **NAGA** | 9.2272 | 0.815300293731623 | 0.986 | 0.901 | 1.5111442 |
| **PTPN13** | 1.0570 | -1.08091309979308 | 0.379 | 0.797 | 1.7311085 |
| **ERBIN** | 1.0944 | 0.847142531636727 | 0.994 | 0.946 | 1.7924432 |
| **HDHD2** | 4.0300 | -0.825114050850347 | 0.788 | 0.969 | 6.6000183 |
| **TCEA2** | 4.7106 | -0.948774457908213 | 0.65 | 0.92 | 7.7145957 |
| **YPEL2** | 7.9472 | 1.19800056197264 | 0.703 | 0.249 | 1.3015229 |
| **DUSP7** | 8.4813 | 0.91430078503219 | 0.957 | 0.77 | 1.3889928 |
| **KCNMB1** | 9.4805 | -1.69946278027642 | 0.05 | 0.532 | 1.5526294 |
| **NCKAP5L** | 1.0671 | -0.909559577910536 | 0.565 | 0.908 | 1.7477285 |
| **ZNF675** | 6.5712 | 0.818415549135991 | 0.977 | 0.827 | 1.0761688 |
| **ARHGEF10L** | 1.4368 | -0.989694075487204 | 0.289 | 0.744 | 2.3530835 |

| | | | | | |
|---|---|---|---|---|---|
| **MUC4** | 1.4943 | 0.953812951510373 | 0.743 | 0.273 | 2.4472504 |
| **AGAP3** | 1.5895 | -0.953473611850333 | 0.308 | 0.758 | 2.6031633 |
| **IGSF1** | 1.6254 | 1.27487697818264 | 0.868 | 0.569 | 2.6620399 |
| **SERTAD4** | 1.9136 | -1.13771060302882 | 0.135 | 0.622 | 3.1339303 |
| **PCNA** | 2.3454 | -0.903357069959629 | 0.995 | 0.998 | 3.8410930 |
| **LYPD1** | 3.2789 | -1.83170890546463 | 0.273 | 0.724 | 5.3699238 |
| **TMEM154** | 3.5297 | 1.03263920461483 | 0.776 | 0.336 | 5.7806987 |
| **SYS1** | 6.2085 | -0.860573285432043 | 0.737 | 0.947 | 1.0167808 |
| **GALC** | 2.5173 | -0.973307212819819 | 0.523 | 0.869 | 4.1227225 |
| **MAGEF1** | 8.7159 | -0.839827326526268 | 0.83 | 0.979 | 1.4274109 |
| **PAPPA** | 1.8517 | -1.1348407771884 | 0.043 | 0.524 | 3.0325687 |
| **C1orf198** | 1.8599 | -0.835481002036303 | 0.781 | 0.963 | 3.0460909 |
| **FEN1** | 4.7348 | 0.816227850802346 | 0.999 | 0.985 | 7.7542129 |
| **PCDH10** | 6.4535 | -2.31275266373881 | 0.093 | 0.563 | 1.0568971 |
| **UAP1L1** | 2.2718 | 1.00387082159755 | 0.838 | 0.449 | 3.7206803 |
| **SMPDL3B** | 5.7146 | -1.01284619993463 | 0.097 | 0.581 | 9.3588172 |
| **GSPT2** | 6.9551 | -0.894365961491324 | 0.726 | 0.961 | 1.1390375 |
| **ST14** | 9.9513 | 0.924188441066246 | 0.659 | 0.193 | 1.6297344 |
| **CASP2** | 1.0321 | -0.862807997101447 | 0.796 | 0.97 | 1.6904012 |
| **TWSG1** | 1.3890 | -0.887079827725364 | 0.883 | 0.987 | 2.2748389 |
| **HP1BP3** | 2.7834 | -0.853913663864771 | 0.782 | 0.958 | 4.5585199 |
| **RFX3** | 3.1181 | -0.983053424900928 | 0.275 | 0.741 | 5.1065809 |
| **ELF4** | 3.5520 | -1.15307856001894 | 0.133 | 0.615 | 5.8172025 |
| **PKIG** | 3.7566 | -0.869053936013296 | 0.938 | 0.988 | 6.1522722 |
| **CHMP7** | 4.9523 | -0.83350544042708 | 0.758 | 0.953 | 8.1103922 |
| **SPC24** | 4.9669 | -0.847876836325768 | 0.892 | 0.981 | 8.1344039 |
| **PM20D2** | 6.3209 | -1.00861058986027 | 0.185 | 0.658 | 1.0351860 |
| **RTL8A** | 1.8937 | -0.998739610581465 | 0.41 | 0.8 | 3.1013256 |
| **FOXB1** | 2.3583 | -1.54517542526082 | 0.364 | 0.765 | 3.8622480 |
| **HEXIM1** | 2.8506 | -0.886027518459803 | 0.781 | 0.958 | 4.6684526 |
| **ZNF740** | 3.3387 | -0.966356843633638 | 0.399 | 0.795 | 5.4678198 |
| **TUB** | 3.3439 | 0.945581155961579 | 0.783 | 0.358 | 5.4764239 |
| **TWF1** | 4.5936 | 0.842010712369488 | 0.758 | 0.327 | 7.5229504 |
| **1,00 MDC** | 6.5151 | -0.828813753842435 | 0.862 | 0.983 | 1.0669813 |

| | | | | | |
|---|---|---|---|---|---|
| **SAP25** | 6.7973 | 0.849367872493851 | 0.521 | 0.094 | 1.11320259 |
| **COL9A3** | 1.2709 | -0.874167839140111 | 0.997 | 0.998 | 2.08142824 |
| **DLG3** | 1.4731 | 0.884733185822164 | 0.998 | 0.988 | 2.41262438 |
| **AGPAT4** | 1.5017 | -1.06248276772672 | 0.479 | 0.852 | 2.45938237 |
| **GRTP1** | 3.0708 | 0.862096935573057 | 0.76 | 0.297 | 5.02919412 |
| **TRIOBP** | 3.1690 | -0.801745393478368 | 0.88 | 0.985 | 5.19001602 |
| **COL1A2** | 1.8358 | -1.15826785012139 | 0.762 | 0.989 | 3.00654026 |
| **SEMA6A** | 3.5488 | -1.03955050109874 | 0.982 | 0.998 | 5.81189452 |
| **BSPRY** | 5.7952 | 0.845821011008352 | 0.718 | 0.238 | 9.49083569 |
| **ARHGAP4** | 7.6131 | 0.892038412279171 | 0.619 | 0.169 | 1.24679739 |
| **NEFL** | 9.5825 | 0.875025707928414 | 0.368 | 0.011 | 1.56933367 |
| **SELENOM** | 9.7299 | 0.91386376931116 | 0.905 | 0.616 | 1.59348174 |
| **CTSK** | 7.3533 | -2.00344879366083 | 0.202 | 0.647 | 1.20426165 |
| **ZNF589** | 1.2504 | 0.822264713934576 | 0.991 | 0.904 | 2.04779382 |
| **SAMD4A** | 1.4988 | -1.17727454938073 | 0.108 | 0.598 | 2.45472243 |
| **FAM172A** | 2.3394 | -0.893778110009901 | 0.525 | 0.867 | 3.83126557 |
| **MARVELD2** | 3.1678 | 0.839621263376745 | 0.704 | 0.237 | 5.18796885 |
| **CHORDC1** | 9.2693 | 0.869068094986096 | 0.76 | 0.339 | 1.51803408 |
| **SYNDIG1** | 9.7368 | -0.936911480792354 | 0.033 | 0.492 | 1.59460607 |
| **E2F3** | 1.2781 | -0.863461948137803 | 0.822 | 0.971 | 2.09330164 |
| **PSKH1** | 1.3080 | -0.834167652289111 | 0.751 | 0.96 | 2.14224003 |
| **C9orf135** | 1.6370 | 1.04729194441797 | 0.904 | 0.498 | 2.68097073 |
| **CDC42EP3** | 3.1702 | -0.972992210892163 | 0.525 | 0.876 | 5.19196327 |
| **EXTL2** | 4.1517 | -0.808091502089132 | 0.791 | 0.954 | 6.79939704 |
| **SLC39A14** | 4.5225 | 0.948541671726243 | 0.935 | 0.741 | 7.40664567 |
| **ATP5MPL** | 5.2974 | 0.825274948797295 | 0.899 | 0.602 | 8.67566135 |
| **AJAP1** | 2.1983 | -1.05918449654339 | 0.089 | 0.558 | 3.60017500 |
| **RASL11B** | 2.8668 | -1.25007251268817 | 0.313 | 0.744 | 4.69506588 |
| **ST3GAL3** | 3.9062 | -0.899241299133641 | 0.103 | 0.578 | 6.39730691 |
| **RRP12** | 4.8313 | 0.887015803754386 | 0.974 | 0.898 | 7.91237789 |
| **PHLDA2** | 5.6820 | 1.08666647105792 | 0.924 | 0.68 | 9.30552319 |
| **KATNAL1** | 7.2166 | -0.901725252184186 | 0.454 | 0.835 | 1.18187254 |
| **SH3GL3** | 1.4861 | -1.0689869652976 | 0.249 | 0.691 | 2.43379009 |
| **ZNF609** | 1.9199 | -0.822717401097221 | 0.692 | 0.928 | 3.14432691 |

| | | | | | |
|---|---|---|---|---|---|
| **ZNF729** | 5.5178 | 1.05183483299538 | 0.367 | 0.015 | 9.03653872 |
| **TLNRD1** | 8.8568 | -0.830505926306249 | 0.883 | 0.982 | 1.45048315 |
| **THAP12** | 1.4723 | 0.872108854709058 | 0.849 | 0.497 | 2.41130612 |
| **COX20** | 2.2952 | 0.821122760252515 | 0.805 | 0.404 | 3.75899752 |
| **ARPIN** | 3.0343 | -0.94538551137062 | 0.434 | 0.816 | 4.96932449 |
| **SLC16A3** | 4.4421 | 1.28973624181065 | 0.954 | 0.76 | 7.27488789 |
| **UNC5D** | 8.5449 | -0.966220439531009 | 0.132 | 0.607 | 1.39939982 |
| **LYPD6** | 9.4711 | -0.854308467347373 | 0.089 | 0.564 | 1.55109404 |
| **BARD1** | 1.5485 | 0.866839294322434 | 0.961 | 0.814 | 2.53599676 |
| **NIPAL1** | 1.9643 | 0.891091570735951 | 0.612 | 0.171 | 3.21698902 |
| **IFI27L2** | 2.0784 | -0.974497451853841 | 0.173 | 0.644 | 3.40391363 |
| **DBNDD1** | 6.6202 | -1.18559870797551 | 0.644 | 0.886 | 1.08420181 |
| **MEST** | 7.6259 | 0.923503636480149 | 0.95 | 0.813 | 1.24889599 |
| **CRLF1** | 1.3755 | -2.50330190140254 | 0.369 | 0.735 | 2.25272943 |
| **ZNF680** | 2.2819 | 0.809330784148353 | 0.95 | 0.736 | 3.73721372 |
| **SEMA6B** | 1.1210 | -1.03455259287795 | 0.568 | 0.881 | 1.83601878 |
| **IGSF8** | 1.3268 | -0.960347494185692 | 0.731 | 0.936 | 2.17293604 |
| **JAKMIP2** | 3.0560 | -0.998737809066498 | 0.041 | 0.492 | 5.00495917 |
| **PKIA** | 9.2557 | -0.814940582765596 | 0.071 | 0.533 | 1.51581816 |
| **EPOP** | 1.9587 | -1.09917747766638 | 0.505 | 0.835 | 3.20792167 |
| **ZNF43** | 2.5083 | -0.874977621711905 | 0.504 | 0.84 | 4.10796950 |
| **CCNYL1** | 9.3867 | -0.82077733644491 | 0.788 | 0.975 | 1.53726718 |
| **CDCP1** | 2.1245 | 0.884116636656258 | 0.795 | 0.368 | 3.47934715 |
| **CRISPLD1** | 3.2363 | -1.07240106158104 | 0.684 | 0.937 | 5.30015221 |
| **IGDCC4** | 4.1567 | -0.811380321181975 | 0.054 | 0.509 | 6.80752759 |
| **IDH2** | 5.3907 | -0.814108411035971 | 0.936 | 0.998 | 8.82849238 |
| **PLPP2** | 5.4839 | 0.814389302392835 | 0.428 | 0.048 | 8.98112264 |
| **MYO9A** | 6.2487 | 0.860509337755262 | 0.836 | 0.471 | 1.02335290 |
| **CUEDC2** | 1.3090 | -0.811431777225742 | 0.729 | 0.947 | 2.14389393 |
| **HSPG2** | 1.4549 | -0.929857516463744 | 0.136 | 0.602 | 2.38279014 |
| **MFSD4B** | 2.9199 | 0.905125295643286 | 0.899 | 0.625 | 4.78203124 |
| **IER3** | 2.9281 | -1.62156673482043 | 0.326 | 0.742 | 4.79538282 |
| **VASH2** | 3.0384 | 0.902735518232165 | 0.997 | 0.963 | 4.97615059 |
| **OCIAD2** | 6.0693 | -0.968321466518216 | 0.271 | 0.729 | 9.93975714 |

| | | | | | |
|---|---|---|---|---|---|
| **GBA** | 7.5182 | -0.817917543261699 | 0.741 | 0.949 | 1.23126038 |
| **C20orf96** | 8.4919 | -0.850999343770911 | 0.105 | 0.566 | 1.39072080 |
| **AIFM2** | 9.4317 | 0.826445909017279 | 0.73 | 0.292 | 1.54463770 |
| **CYB5R2** | 1.3884 | 0.83055380696298 | 0.824 | 0.392 | 2.27385822 |
| **TMEM50B** | 1.8171 | 0.909116991627676 | 0.93 | 0.731 | 2.97596262 |
| **SLF2** | 2.4724 | -0.821692749495921 | 0.828 | 0.967 | 4.04905608 |
| **REEP2** | 2.4829 | -1.00420277116607 | 0.24 | 0.687 | 4.06631182 |
| **NUCB2** | 5.6482 | 0.939893875185735 | 0.902 | 0.69 | 9.25013260 |
| **ITPR3** | 5.7409 | 0.869430337925442 | 0.94 | 0.685 | 9.40191623 |
| **ZNF580** | 5.9781 | -0.93669465932305 | 0.583 | 0.895 | 9.79034705 |
| **SH3BGRL3** | 6.6380 | -0.849821740088534 | 0.963 | 0.997 | 1.08710649 |
| **SHISAL2B** | 8.4348 | -1.00256488174285 | 0.157 | 0.614 | 1.38136930 |
| **HCN2** | 1.4484 | 0.822016260567318 | 0.949 | 0.708 | 2.37204481 |
| **KIF21A** | 2.4495 | 0.88519922908271 | 0.961 | 0.811 | 4.01158642 |
| **DSC2** | 9.9402 | -0.8623185793856 | 0.798 | 0.967 | 1.62790730 |
| **CER1** | 1.1501 | -3.74021995506265 | 0.24 | 0.695 | 1.88367664 |
| **ZNF614** | 3.3465 | 0.830492441515289 | 0.935 | 0.697 | 5.48068989 |
| **ELAC1** | 3.7350 | -0.879894312229452 | 0.198 | 0.649 | 6.11693491 |
| **AARS2** | 4.4418 | 0.876620194746148 | 0.969 | 0.88 | 7.27441288 |
| **CABLES2** | 5.4967 | -0.901111739649549 | 0.238 | 0.687 | 9.00205510 |
| **CCN2** | 1.2350 | -1.42119934748156 | 0.769 | 0.942 | 2.02270364 |
| **CPNE2** | 1.3305 | -0.937350883080876 | 0.313 | 0.748 | 2.17907949 |
| **LZTS2** | 1.4385 | -0.801383527825426 | 0.855 | 0.973 | 2.35598139 |
| **DOCK11** | 1.6187 | 0.923463842780968 | 0.874 | 0.552 | 2.65103490 |
| **RPP40** | 2.0373 | 0.812634613807482 | 0.942 | 0.748 | 3.33660083 |
| **ALDH1B1** | 2.6820 | -0.844786139846393 | 0.775 | 0.95 | 4.39244995 |
| **MFSD10** | 5.9785 | -0.801752484184371 | 0.798 | 0.961 | 9.79111532 |
| **VASN** | 9.8257 | -0.899618663836012 | 0.128 | 0.586 | 1.60916248 |
| **CERS4** | 1.1046 | -0.944110601805777 | 0.967 | 0.999 | 1.80913628 |
| **LLPH** | 1.6312 | 0.807228067095011 | 0.809 | 0.438 | 2.67149002 |
| **NDUFAF7** | 1.7617 | 0.815427754311966 | 0.967 | 0.817 | 2.88517412 |
| **ACVR1** | 2.4247 | 0.828292357269489 | 0.95 | 0.791 | 3.97108427 |
| **BAHCC1** | 2.8854 | -0.880427531045635 | 0.031 | 0.467 | 4.72544433 |
| **MYC** | 3.0998 | -1.03961573973234 | 0.917 | 0.991 | 5.07654430 |

| | | | | | |
|---|---|---|---|---|---|
| **PTPN21** | 4.6520 | -0.917718573801287 | 0.282 | 0.716 | 7.6186809 |
| **TMEM138** | 5.9419 | 0.858165428501227 | 0.931 | 0.724 | 9.7311168 |
| **CHST2** | 9.8227 | 1.12828436904166 | 0.956 | 0.723 | 1.6086648 |
| **CENPA** | 1.3596 | 1.05387654306053 | 0.982 | 0.874 | 2.2267465 |
| **PHLDB1** | 3.5246 | -0.818440261698055 | 0.739 | 0.949 | 5.7723642 |
| **EPS8L1** | 3.6136 | 0.855217288033147 | 0.744 | 0.32 | 5.9180853 |
| **SOCS1** | 3.9427 | -0.92177079254958 | 0.957 | 0.995 | 6.4570480 |
| **NEO1** | 1.2388 | -0.901496304503456 | 0.547 | 0.861 | 2.0289249 |
| **FAM107B** | 4.8822 | 0.891854533786287 | 0.954 | 0.803 | 7.9956804 |
| **ANK2** | 3.2778 | -0.910203753372854 | 0.705 | 0.933 | 5.3681571 |
| **MFAP3L** | 6.5658 | -1.08030567750572 | 0.612 | 0.891 | 1.0752940 |
| **INSR** | 9.4694 | 0.804425961145858 | 0.983 | 0.88 | 1.5508068 |
| **BRSK1** | 1.1874 | -0.885877201750593 | 0.212 | 0.664 | 1.9447313 |
| **COL6A1** | 2.1322 | -1.63463079208235 | 0.637 | 0.88 | 3.4919620 |
| **LAMA2** | 2.8550 | -1.01233106725273 | 0.139 | 0.583 | 4.6757174 |
| **FARP1** | 4.2311 | -0.825265393262107 | 0.786 | 0.964 | 6.9293974 |
| **TENM4** | 7.4014 | -1.16187943294178 | 0.458 | 0.819 | 1.2121387 |
| **AUTS2** | 7.5262 | -1.00067613958566 | 0.635 | 0.924 | 1.2325676 |
| **MEX3B** | 2.6833 | -0.933785916876305 | 0.099 | 0.546 | 4.3945896 |
| **ZNF483** | 5.9905 | 0.958351114873627 | 0.875 | 0.601 | 9.8106828 |
| **MME** | 6.8985 | -1.07551395406254 | 0.247 | 0.674 | 1.1297794 |
| **TM7SF2** | 1.4006 | -0.994024562021253 | 0.83 | 0.952 | 2.2937801 |
| **RGMA** | 1.9008 | -1.17344597129587 | 0.492 | 0.815 | 3.1130031 |
| **MPZL1** | 2.0223 | -0.818359939605575 | 0.733 | 0.938 | 3.3119295 |
| **CYP1B1** | 2.1804 | -1.7357219939937 | 0.145 | 0.585 | 3.5708868 |
| **ZNF703** | 2.1975 | -1.38577945593117 | 0.587 | 0.894 | 3.5988522 |
| **2,00 NPR** | 8.0481 | -0.830456142500941 | 0.127 | 0.577 | 1.3180450 |
| **POC1A** | 6.3797 | -0.800182299020693 | 0.739 | 0.938 | 1.0448140 |
| **KCNE3** | 1.7093 | 0.852894926407138 | 0.619 | 0.189 | 2.7994073 |
| **TMLHE** | 1.9859 | -0.887941775974383 | 0.414 | 0.781 | 3.2524153 |
| **AIG1** | 2.0159 | -0.986420588251615 | 0.325 | 0.754 | 3.3014554 |
| **NAA50** | 2.5865 | 0.806798468006033 | 0.817 | 0.472 | 4.2359469 |
| **CASP10** | 3.9572 | 0.87802514834065 | 0.758 | 0.35 | 6.4808007 |
| **BRCA2** | 4.5960 | -0.800446298154732 | 0.847 | 0.965 | 7.5268893 |

| | | | | | |
|---|---|---|---|---|---|
| **DDHD2** | 7.8743 | -0.880379398989833 | 0.45 | 0.806 | 1.28957675 |
| **CACNA2D1** | 8.0445 | 0.827344424034079 | 0.933 | 0.73 | 1.31745280 |
| **FBXO17** | 9.7415 | -0.85614868750718 | 0.498 | 0.841 | 1.59537351 |
| **LTBP1** | 1.6777 | -0.955200440575768 | 0.486 | 0.818 | 2.74759012 |
| **MPC1** | 4.1368 | 0.86640583228925 | 0.828 | 0.47 | 6.77484388 |
| **CALHM2** | 1.6302 | -0.867949168536088 | 0.113 | 0.556 | 2.66989060 |
| **UST** | 1.6472 | -0.91961514159155 | 0.813 | 0.957 | 2.69773424 |
| **SRR** | 2.8659 | -0.801896787120166 | 0.113 | 0.556 | 4.69355960 |
| **DPY19L1** | 8.1014 | 0.859632144741433 | 0.84 | 0.49 | 1.32678029 |
| **OVCH2** | 8.6865 | -0.917333177496275 | 0.026 | 0.439 | 1.42259570 |
| **E2F1** | 1.8750 | -1.16327624652211 | 0.908 | 0.972 | 3.07078424 |
| **HSD17B11** | 8.2737 | 0.814792825392898 | 0.897 | 0.614 | 1.35499838 |
| **ADRA2C** | 1.2195 | -1.12952483822186 | 0.846 | 0.952 | 1.99721477 |
| **ADCY6** | 1.2358 | -0.841964487223323 | 0.103 | 0.538 | 2.02401865 |
| **PKN3** | 1.5119 | -0.811553378176468 | 0.629 | 0.899 | 2.47606192 |
| **C15orf61** | 2.7355 | 0.83260963552351 | 0.776 | 0.385 | 4.48001643 |
| **TP53BP1** | 4.3289 | -0.808330170545323 | 0.73 | 0.927 | 7.08947188 |
| **FMNL3** | 4.8629 | -0.865259427299227 | 0.191 | 0.627 | 7.96413134 |
| **MEIS3** | 5.8989 | -1.80859603850103 | 0.572 | 0.818 | 9.66066827 |
| **3,00 F** | 1.4042 | 1.17558223140253 | 0.794 | 0.446 | 2.29973103 |
| **SUB1** | 1.4665 | 0.804028328659531 | 0.759 | 0.39 | 2.40169455 |
| **PCED1A** | 3.6602 | -1.00280492079939 | 0.339 | 0.733 | 5.99436758 |
| **ADCY2** | 2.0762 | -1.1388749423479 | 0.03 | 0.441 | 3.40020163 |
| **COBL** | 3.5881 | -1.22394198786455 | 0.566 | 0.831 | 5.87627510 |
| **PPP1R1A** | 9.6685 | -1.16502646552078 | 0.443 | 0.795 | 1.58342122 |
| **MSRB2** | 9.9382 | -0.811440991849569 | 0.653 | 0.911 | 1.62758414 |
| **UNK** | 4.2234 | -0.825614212729078 | 0.593 | 0.883 | 6.91667547 |
| **TXNDC16** | 1.2317 | -0.836817367622344 | 0.281 | 0.693 | 2.01716341 |
| **CCDC85C** | 1.4936 | -0.879285067435782 | 0.975 | 0.987 | 2.44615021 |
| **PTCH1** | 1.9828 | -0.924315901634122 | 0.548 | 0.888 | 3.24738580 |
| **OAF** | 9.1682 | -0.925581565429497 | 0.088 | 0.514 | 1.50148447 |
| **RAI1** | 1.5165 | -0.829589114993019 | 0.076 | 0.497 | 2.48370883 |
| **OXCT1** | 1.9070 | -0.804935240537965 | 0.6 | 0.895 | 3.12309492 |
| **NUP210** | 2.2287 | -0.985816713632641 | 0.479 | 0.815 | 3.65005977 |

| | | | | | |
|---|---|---|---|---|---|
| **LRFN4** | 4.8903 | -0.859382282960365 | 0.255 | 0.673 | 8.0089845 |
| **GULP1** | 1.0836 | -0.896267669100157 | 0.493 | 0.812 | 1.7747258 |
| **SLC39A6** | 2.4615 | -0.871603635670554 | 0.644 | 0.892 | 4.0313436 |
| **NFATC4** | 4.6709 | -1.03538288346362 | 0.324 | 0.715 | 7.6496782 |
| **FKRP** | 5.0277 | -0.806664313720363 | 0.464 | 0.815 | 8.2338870 |
| **RARA** | 1.2009 | -0.851191554596523 | 0.335 | 0.736 | 1.9667878 |
| **ARG2** | 1.2408 | 0.855655250059189 | 0.932 | 0.744 | 2.0320676 |
| **CD83** | 1.6487 | 0.885129057688277 | 0.574 | 0.18 | 2.7000950 |
| **SPP1** | 3.5839 | -1.95936264354175 | 0.511 | 0.847 | 5.8695106 |
| **BDNF** | 5.4907 | 0.924187583443551 | 0.5 | 0.12 | 8.9922819 |
| **HSPA4L** | 6.0802 | 0.803373160988736 | 0.642 | 0.236 | 9.9576473 |
| **PIK3R1** | 6.7479 | -1.12451858044028 | 0.475 | 0.798 | 1.1051153 |
| **GRHL3** | 7.8058 | -1.28015532655268 | 0.096 | 0.508 | 1.2783665 |
| **TSKU** | 8.1808 | -0.96939869412041 | 0.967 | 0.996 | 1.3397781 |
| **ANKRD45** | 1.1771 | 0.92812169339245 | 0.449 | 0.087 | 1.9277541 |
| **SIX5** | 2.0743 | -0.800445464792975 | 0.613 | 0.89 | 3.3971591 |
| **CREG1** | 4.0723 | -0.826084694241525 | 0.621 | 0.886 | 6.6693175 |
| **ISLR** | 5.4713 | -1.04721592820739 | 0.069 | 0.477 | 8.9604871 |
| **WNT3A** | 6.6683 | -1.75160272838866 | 0.101 | 0.508 | 1.0920730 |
| **NDRG2** | 1.1405 | -0.824232361891711 | 0.191 | 0.613 | 1.8678362 |
| **CYP26A1** | 1.8381 | -2.70174978319309 | 0.165 | 0.576 | 3.0104116 |
| **SHTN1** | 3.8843 | -0.860902307194796 | 0.257 | 0.67 | 6.3614313 |
| **SCG3** | 6.3192 | -0.912914609012732 | 0.378 | 0.744 | 1.0349039 |
| **SYNPO** | 7.2747 | -0.812140095306029 | 0.029 | 0.426 | 1.1913914 |
| **MAF** | 3.7219 | -1.1036394855337 | 0.145 | 0.563 | 6.0954390 |
| **EDN1** | 3.7446 | -1.65121539740177 | 0.045 | 0.44 | 6.1326747 |
| **ADAMTSL4** | 6.9181 | -1.43331639605997 | 0.272 | 0.646 | 1.1329914 |
| **IRF2BPL** | 4.0276 | -1.00593151983239 | 0.959 | 0.982 | 6.5961560 |
| **PCSK1** | 4.8406 | 1.23984134142188 | 0.913 | 0.691 | 7.9275093 |
| **RAB17** | 5.6994 | -1.30593559157794 | 0.11 | 0.526 | 9.3339987 |
| **IER5L** | 1.0749 | -1.02022758906752 | 0.31 | 0.688 | 1.7603833 |
| **RGMB** | 1.4120 | -0.960996887902998 | 0.619 | 0.874 | 2.3125784 |
| **KTI12** | 2.4524 | 0.802977813001218 | 0.874 | 0.587 | 4.0163233 |
| **EMILIN3** | 3.9859 | -0.810405927098397 | 0.092 | 0.506 | 6.5278426 |

| | | | | | |
|---|---|---|---|---|---|
| **URGCP** | 4.2162 | 0.855808108536138 | 0.835 | 0.548 | 6.9049753 |
| **DMPK** | 7.4130 | -0.834375894867473 | 0.255 | 0.663 | 1.2140360 |
| **CNKSR3** | 7.6249 | -1.06949616021701 | 0.141 | 0.559 | 1.2487314 |
| **TEAD1** | 8.8693 | -0.80677013341486 | 0.555 | 0.856 | 1.4525395 |
| **HERPUD1** | 1.5109 | 1.15421359593718 | 0.91 | 0.792 | 2.4744674 |
| **EFEMP2** | 1.7418 | -0.848554131229727 | 0.305 | 0.695 | 2.8525730 |
| **CPXM1** | 2.2374 | -0.914902034563927 | 0.919 | 0.984 | 3.6642993 |
| **FGFBP3** | 2.2578 | -0.906540418318164 | 0.068 | 0.472 | 3.6977278 |
| **BAZ1A** | 2.9514 | -0.882146136505631 | 0.4 | 0.756 | 4.8335083 |
| **TCF7** | 3.3967 | -0.822540667254794 | 0.892 | 0.979 | 5.5628365 |
| **CADM4** | 7.2864 | -0.8404109584177 | 0.165 | 0.58 | 1.1933094 |
| **TMEM88** | 2.1544 | -0.884350365060825 | 0.107 | 0.516 | 3.5283877 |
| **FAXC** | 3.9579 | 0.816588070346569 | 0.804 | 0.434 | 6.4818690 |
| **CERS6** | 1.1872 | -0.823605563963315 | 0.43 | 0.783 | 1.9443496 |
| **KREMEN2** | 1.7048 | -0.908706788310074 | 0.549 | 0.847 | 2.7920364 |
| **L1CAM** | 2.9111 | -1.09160821372504 | 0.405 | 0.772 | 4.7676251 |
| **NUAK2** | 9.7150 | -0.953326094541943 | 0.496 | 0.815 | 1.5910329 |
| **TCF15** | 1.1274 | -0.821331422633824 | 0.187 | 0.597 | 1.8463716 |
| **METTL7A** | 1.1444 | 0.986111884813722 | 0.657 | 0.279 | 1.8743211 |
| **ZKSCAN1** | 1.5813 | -0.827197768768818 | 0.497 | 0.811 | 2.5898512 |
| **CA13** | 8.3016 | 0.813624633391652 | 0.559 | 0.176 | 1.3595659 |
| **TMEM92** | 7.8798 | -0.843443269967032 | 0.343 | 0.723 | 1.2904825 |
| **INSYN1** | 1.1644 | -1.05655470476526 | 0.131 | 0.528 | 1.9070472 |
| **ZBTB3** | 2.1377 | 0.896451059901946 | 0.828 | 0.538 | 3.5009820 |
| **MGAT3** | 2.6132 | -0.953476918333716 | 0.327 | 0.698 | 4.2796832 |
| **PLAAT3** | 5.7232 | 0.841100663470932 | 0.988 | 0.886 | 9.3729922 |
| **SRGAP1** | 6.2288 | 0.80478667987883 | 0.871 | 0.596 | 1.0201006 |
| **TSPAN7** | 8.1006 | -1.18774588070618 | 0.102 | 0.511 | 1.3266397 |
| **ANKRD1** | 2.7362 | -1.71483194830523 | 0.088 | 0.489 | 4.4811589 |
| **PRAC1** | 5.0446 | -0.842066489696548 | 0.214 | 0.635 | 8.2615973 |
| **FAM111B** | 8.3419 | -1.09261032802357 | 0.49 | 0.821 | 1.3661631 |
| **GRPR** | 2.8939 | -1.05704489271568 | 0.033 | 0.41 | 4.7393697 |
| **ZNF296** | 3.8340 | 1.01964956815423 | 0.809 | 0.518 | 6.2789914 |
| **TCF7L1** | 9.0529 | -1.23110050230432 | 0.28 | 0.642 | 1.4825937 |

| | | | | | |
|---|---|---|---|---|---|
| **ARSE** | 4.5878 | -0.830202551314963 | 0.147 | 0.563 | 7.5135495 |
| **LIMCH1** | 1.2053 | -1.28006056762462 | 0.557 | 0.803 | 1.9739618 |
| **ZNF165** | 2.2569 | 0.98535974503096 | 0.649 | 0.264 | 3.6962137 |
| **TGFB3** | 3.6849 | 0.810875069330869 | 0.548 | 0.163 | 6.0348092 |
| **DUSP10** | 4.4020 | -0.919816122374207 | 0.343 | 0.721 | 7.2092811 |
| **DHRS3** | 1.6674 | -1.11407792388369 | 0.236 | 0.615 | 2.7308163 |
| **GJA5** | 2.0270 | -0.899733263706551 | 0.026 | 0.393 | 3.3196739 |
| **TNFRSF10D** | 2.2428 | -0.916477503379602 | 0.132 | 0.531 | 3.6730669 |
| **WWC1** | 1.2657 | -0.916844435311811 | 0.35 | 0.705 | 2.0729069 |
| **CUEDC1** | 2.7991 | -0.835696061912302 | 0.271 | 0.653 | 4.5841131 |
| **NEXN** | 2.8180 | -0.867579925252695 | 0.162 | 0.558 | 4.6151306 |
| **TRIM38** | 3.3529 | 0.913930625217375 | 0.697 | 0.335 | 5.4911164 |
| **FAM43A** | 3.3697 | -0.972346280899274 | 0.079 | 0.466 | 5.5186150 |
| **TBXT** | 6.0651 | -1.80988474245146 | 0.509 | 0.844 | 9.9328409 |
| **PHLDB2** | 3.1199 | -0.94239145470743 | 0.653 | 0.891 | 5.1094924 |
| **BEX1** | 1.2356 | -0.949032628141155 | 0.11 | 0.496 | 2.0236285 |
| **IRS1** | 2.1795 | -0.815843229150448 | 0.237 | 0.624 | 3.5694222 |
| **DDB2** | 5.8363 | -0.840843687252854 | 0.72 | 0.914 | 9.5581955 |
| **APLNR** | 7.3987 | -3.16915276845784 | 0.379 | 0.701 | 1.2116882 |
| **TACR1** | 1.4158 | -1.74185055441932 | 0.046 | 0.413 | 2.3186673 |
| **MGAT5B** | 8.6305 | -0.807194313131783 | 0.255 | 0.635 | 1.4134187 |
| **THBS1** | 1.5819 | -0.967988629895378 | 0.114 | 0.498 | 2.5908090 |
| **FANCB** | 5.4219 | -0.807146961277344 | 0.502 | 0.803 | 8.8795518 |
| **PCBP4** | 6.1348 | -0.974386329710581 | 0.276 | 0.644 | 1.0047006 |
| **OTUD1** | 9.3666 | 0.876386285628585 | 0.637 | 0.276 | 1.5339735 |
| **GLI3** | 6.5425 | -1.08085872910328 | 0.167 | 0.552 | 1.0714724 |
| **EDA** | 1.0418 | -0.808759855044605 | 0.572 | 0.843 | 1.7061811 |
| **GMPR** | 4.3886 | -1.55876125430948 | 0.196 | 0.552 | 7.1873287 |
| **GALNT10** | 1.2791 | -0.800668221894018 | 0.501 | 0.803 | 2.0947974 |
| **STARD8** | 1.7431 | -0.930064456859943 | 0.497 | 0.829 | 2.8548017 |
| **CDX2** | 1.8207 | -1.54587135094319 | 0.088 | 0.465 | 2.9819004 |
| **STC1** | 2.5326 | -1.11526404075018 | 0.127 | 0.518 | 4.1476427 |
| **CAPRIN2** | 4.1385 | -0.874906726362473 | 0.411 | 0.753 | 6.7777170 |
| **PREX1** | 4.4056 | -1.28868834869575 | 0.653 | 0.858 | 7.2151016 |

| | | | | | |
|---|---|---|---|---|---|
| **SMAD3** | 7.9896 | -0.807077632745134 | 0.7 | 0.91 | 1.30846227 |
| **CBX4** | 1.4642 | -0.866615132738923 | 0.401 | 0.759 | 2.39802703 |
| **ROBO1** | 1.1797 | -0.865857729737301 | 0.749 | 0.943 | 1.93210254 |
| **BMP2** | 1.3581 | -2.32352093112802 | 0.138 | 0.508 | 2.22418159 |
| **1,00 DKK** | 3.7242 | -2.59557878774812 | 0.058 | 0.418 | 6.09916127 |
| **RGS3** | 5.0346 | -0.827995593390194 | 0.107 | 0.485 | 8.24528257 |
| **SHB** | 6.5853 | -0.814529527836187 | 0.816 | 0.941 | 1.07847752 |
| **LIPG** | 1.5104 | -1.32711321014851 | 0.113 | 0.484 | 2.47358472 |
| **1,00 PGK** | 1.5104 | 0.86919168409926 | 0.753 | 0.456 | 2.47360411 |
| **FGF17** | 1.9624 | -2.83086945429814 | 0.285 | 0.621 | 3.21385225 |
| **EOGT** | 6.8110 | -0.902048145616812 | 0.844 | 0.973 | 1.11544778 |
| **CENPI** | 7.0873 | -0.814889970482928 | 0.716 | 0.913 | 1.16069385 |
| **RND3** | 9.5392 | -1.00552877773083 | 0.118 | 0.495 | 1.56224500 |
| **CCNG2** | 1.0206 | 0.921693771245764 | 0.844 | 0.583 | 1.67147454 |
| **NKX1-2** | 2.4040 | -1.1500036363069 | 0.893 | 0.931 | 3.93716517 |
| **ETV5** | 2.6940 | -0.963694091288366 | 0.473 | 0.771 | 4.41202360 |
| **MSX1** | 9.6041 | -1.43454967777582 | 0.031 | 0.378 | 1.57286874 |
| **SIAH2** | 1.3454 | -0.864279827489198 | 0.357 | 0.716 | 2.20351785 |
| **PITX2** | 1.7541 | -0.903062441775407 | 0.464 | 0.784 | 2.87272644 |
| **FZD10** | 2.5572 | -1.3074678420692 | 0.68 | 0.876 | 4.18807502 |
| **RHOBTB3** | 2.8167 | -1.48738900489546 | 0.107 | 0.476 | 4.61295539 |
| **SLIT2** | 1.1750 | -1.00770154896188 | 0.024 | 0.36 | 1.92443760 |
| **NR0B1** | 2.4187 | -1.57093802506355 | 0.062 | 0.409 | 3.96115145 |
| **VLDLR** | 8.0770 | -0.892143170901686 | 0.546 | 0.838 | 1.32277367 |
| **DAB1** | 7.1306 | -1.1206089005209 | 0.218 | 0.575 | 1.16779391 |
| **ITGB8** | 1.3584 | -1.05503793729385 | 0.052 | 0.397 | 2.22474685 |
| **ANKRD33B** | 2.5749 | -1.02265963344266 | 0.161 | 0.525 | 4.21705238 |
| **LRRN1** | 2.6053 | -1.42352020464057 | 0.49 | 0.774 | 4.26685345 |
| **CCDC92** | 7.5918 | -0.800634426102973 | 0.065 | 0.416 | 1.24332496 |
| **MIXL1** | 1.3065 | -2.26250982569227 | 0.845 | 0.93 | 2.13973068 |
| **SNCG** | 1.3196 | -0.805246216151611 | 0.189 | 0.553 | 2.16125453 |
| **AMOT** | 2.2362 | -0.919356767118922 | 0.218 | 0.576 | 3.66237124 |
| **TNFRSF12A** | 2.9036 | -0.943862679430959 | 0.95 | 0.965 | 4.75535659 |
| **SEL1L3** | 1.2461 | -0.88428651357099 | 0.155 | 0.514 | 2.04082195 |

| | | | | | |
|---|---|---|---|---|---|
| **GABBR1** | 1.6300 | -1.04257459830778 | 0.597 | 0.845 | 2.6694882 |
| **MKI67** | 2.7932 | -0.979254231759312 | 0.925 | 0.981 | 4.5745082 |
| **PMP22** | 1.0099 | -1.01237721580548 | 0.187 | 0.544 | 1.6539740 |
| **SPRY4** | 2.9414 | -0.844590864351139 | 0.883 | 0.976 | 4.8172559 |
| **EVA1B** | 2.9525 | -0.828751808216638 | 0.36 | 0.702 | 4.8354653 |
| **COL5A1** | 4.1102 | -1.30272583983003 | 0.627 | 0.845 | 6.7313091 |
| **CNTNAP2** | 1.9362 | -2.22731918758876 | 0.815 | 0.869 | 3.1710697 |
| **TGFB1I1** | 1.1920 | -0.914573061026192 | 0.272 | 0.612 | 1.9522927 |
| **HPD** | 1.1326 | -1.07938191500388 | 0.16 | 0.502 | 1.8549281 |
| **TSHZ3** | 3.4066 | -0.81390461651107 | 0.121 | 0.469 | 5.5790672 |
| **AMH** | 6.9031 | -0.817251466099307 | 0.194 | 0.547 | 1.1305354 |
| **MAML3** | 9.5947 | -1.06517146249158 | 0.415 | 0.718 | 1.5713387 |
| **CTDSPL** | 1.8252 | 0.824280508402206 | 0.904 | 0.809 | 2.9891519 |
| **B3GNT7** | 2.3122 | -1.16558645719206 | 0.618 | 0.808 | 3.7868237 |
| **FGF8** | 2.4697 | -1.19775616049228 | 0.529 | 0.786 | 4.0446410 |
| **PRKAR2B** | 1.9363 | -0.916573864299331 | 0.535 | 0.821 | 3.1711397 |
| **ABCG2** | 2.1036 | -1.32441979473422 | 0.343 | 0.645 | 3.4451331 |
| **COL13A1** | 4.3783 | -1.20509770873614 | 0.05 | 0.359 | 7.1704057 |
| **FOXC1** | 2.7725 | -1.38975620791734 | 0.096 | 0.419 | 4.5406080 |
| **MFAP4** | 4.3040 | -0.997567689184438 | 0.054 | 0.362 | 7.0487033 |
| **COL5A2** | 5.9074 | -1.50840336223662 | 0.128 | 0.449 | 9.6746247 |
| **LRATD2** | 8.2835 | -0.964701994820038 | 0.935 | 0.965 | 1.3565929 |
| **HOXB2** | 8.2604 | 0.824889883296541 | 0.753 | 0.478 | 1.3528098 |
| **EFNA1** | 6.8195 | -0.925435777564371 | 0.279 | 0.621 | 1.1168382 |
| **COL7A1** | 6.4772 | -0.96625373711557 | 0.122 | 0.452 | 1.0607806 |
| **CAV1** | 4.1777 | -0.970866091917407 | 0.536 | 0.835 | 6.8419030 |
| **SAMD3** | 1.6957 | -0.921912260448974 | 0.013 | 0.289 | 2.7771898 |
| **CCDC141** | 8.5675 | -1.13395460898073 | 0.07 | 0.375 | 1.4031074 |
| **ROR2** | 4.1716 | -1.04066623911252 | 0.65 | 0.846 | 6.8318933 |
| **AP3B2** | 6.8125 | -1.00366897938449 | 0.254 | 0.547 | 1.1156869 |
| **GAL** | 1.2273 | -1.70081027052008 | 0.993 | 0.981 | 2.0099857 |
| **FAM20A** | 2.6617 | -1.12029412747652 | 0.19 | 0.493 | 4.3591678 |
| **ZNF367** | 1.8596 | -0.82192362421528 | 0.471 | 0.739 | 3.0455237 |
| **PLXNA2** | 3.7876 | -1.81453604132675 | 0.109 | 0.404 | 6.2030397 |

| | | | | | |
|---|---|---|---|---|---|
| **PALLD** | 1.7694 | -1.10475041811165 | 0.234 | 0.535 | 2.8977798 |
| **ADM** | 1.5812 | -1.1621432153215 | 0.388 | 0.652 | 2.5896934 |
| **EVX1** | 6.0383 | -0.858540488689148 | 0.029 | 0.293 | 9.8890296 |
| **SNAI2** | 2.8086 | -1.39509364452082 | 0.031 | 0.292 | 4.5996544 |
| **FXYD6** | 4.9796 | 0.964540758098284 | 0.845 | 0.734 | 8.1552397 |
| **FAM189A2** | 1.4110 | -0.85073912924338 | 0.153 | 0.441 | 2.3108280 |
| **SPATS2L** | 3.0759 | -1.05413202543771 | 0.728 | 0.863 | 5.0374992 |
| **SEMA3F** | 1.9295 | -0.902198295005206 | 0.468 | 0.714 | 3.1600668 |
| **TAGLN2** | 5.0431 | -0.824082801101594 | 0.906 | 0.955 | 8.2591903 |
| **VIT** | 1.9618 | -0.922930813736232 | 0.079 | 0.341 | 3.2129085 |
| **TRIM22** | 4.0041 | -0.89725152911618 | 0.292 | 0.575 | 6.5576213 |
| **HS6ST1** | 1.0638 | -0.901811022661806 | 0.93 | 0.971 | 1.7422043 |
| **GATA6** | 8.4877 | -2.22144148896016 | 0.091 | 0.351 | 1.3900380 |
| **PGF** | 1.0129 | -0.827169892258282 | 0.402 | 0.659 | 1.6589423 |
| **PCDH18** | 1.0846 | -1.04320966682074 | 0.536 | 0.759 | 1.7763705 |
| **GSC** | 1.9106 | -1.54137633777646 | 0.058 | 0.305 | 3.1291150 |
| **GTSF1** | 3.0762 | 0.839839043885114 | 0.283 | 0.069 | 5.0380407 |
| **MSGN1** | 3.9206 | -1.97870693681462 | 0.016 | 0.235 | 6.4209258 |
| **HAND1** | 7.1700 | -1.70697720215935 | 0.037 | 0.267 | 1.1742456 |
| **CACNG4** | 1.2274 | -0.812863384086116 | 0.081 | 0.324 | 2.0102011 |
| **IL13RA1** | 1.9669 | -0.948744953191319 | 0.296 | 0.548 | 3.2212263 |
| **ACSS3** | 4.1988 | -0.880668177999805 | 0.485 | 0.705 | 6.8765376 |
| **TPX2** | 4.8835 | -0.857056892981124 | 0.991 | 0.997 | 7.9977961 |
| **FOXA2** | 5.6751 | -1.0298992385363 | 0.035 | 0.25 | 9.2942591 |
| **LHX1** | 7.2194 | -1.60145396869667 | 0.019 | 0.222 | 1.1823288 |
| **PLAU** | 8.2998 | 1.05660381773733 | 0.615 | 0.369 | 1.3592714 |
| **DLC1** | 5.8497 | -0.935530500196469 | 0.466 | 0.675 | 9.5801130 |
| **ANOS1** | 6.9863 | -1.45582575625742 | 0.174 | 0.411 | 1.1441524 |
| **TRPC6** | 3.3062 | -0.919339421101927 | 0.316 | 0.537 | 5.4145872 |
| **FN1** | 5.3277 | -0.828984762263872 | 0.882 | 0.97 | 8.7253012 |
| **HIST1H4J** | 1.5685 | -0.836584778052357 | 0.557 | 0.722 | 2.5688473 |
| **EPCAM** | 9.7322 | -0.85022311051339 | 0.999 | 0.974 | 1.5938508 |
| **SOX3** | 3.6563 | -1.01207150084639 | 0.303 | 0.52 | 5.9880033 |
| **FOXJ1** | 3.9613 | -0.804565580645681 | 0.144 | 0.376 | 6.4875483 |

| | | | | | |
|---|---|---|---|---|---|
| **TBC1D2** | 4.3329 | -0.812277345857625 | 0.236 | 0.456 | 7.09606498 |
| **GDF15** | 2.1933 | 1.07489193854414 | 0.321 | 0.117 | 3.59207080 |
| **OTX2** | 2.0792 | -0.877061552666723 | 0.755 | 0.879 | 3.40518788 |
| **PCSK9** | 2.0606 | 0.842476917655386 | 0.66 | 0.45 | 3.37465794 |
| **C11orf96** | 4.3997 | -0.914267601946539 | 0.283 | 0.506 | 7.20541129 |
| **CHAC1** | 1.5352 | 0.827996150031646 | 0.675 | 0.501 | 2.51430085 |
| **APCDD1** | 1.1223 | -0.897717593460351 | 0.403 | 0.599 | 1.83801550 |
| **TPM1** | 1.0148 | -0.822098330493894 | 1 | 1 | 1.66208665 |
| **TBX6** | 2.9483 | -1.77933442668652 | 0.145 | 0.324 | 4.82849369 |
| **EMILIN2** | 6.7602 | -0.84866546083245 | 0.18 | 0.37 | 1.10712119 |
| **DLL1** | 8.1463 | -0.972582669230301 | 0.049 | 0.205 | 1.33412423 |
| **NPPB** | 3.0552 | -1.29354160640398 | 0.103 | 0.274 | 5.00353312 |
| **ACTA1** | 1.2932 | -1.12660516359509 | 0.347 | 0.534 | 2.11788427 |
| **CSRP1** | 1.7361 | -0.919669491256891 | 0.976 | 0.993 | 2.84328170 |
| **NRP2** | 2.2160 | -0.829745365695372 | 0.392 | 0.577 | 3.62930439 |
| **SNAI1** | 1.5816 | -0.817185472723643 | 0.21 | 0.351 | 2.59021506 |
| **UBE2C** | 2.1415 | -0.839785345560676 | 0.978 | 0.993 | 3.50720355 |
| **CDX1** | 8.1038 | -0.92603402911371 | 0.758 | 0.787 | 1.32716703 |
| **FAM89A** | 1.1344 | -0.843149619735771 | 0.075 | 0.196 | 1.85791231 |
| **CAVIN1** | 5.3700 | -0.842376259153026 | 0.796 | 0.819 | 8.79448867 |
| **HAS2** | 1.0777 | -0.809753082265737 | 0.826 | 0.682 | 1.76496795 |
| **CLDN10** | 0.0002 | -1.20834084097469 | 0.458 | 0.462 | 1 |
| **APELA** | 0.0006 | -0.869437900853469 | 0.76 | 0.675 | 1 |
| **DLL3** | 0.0462 | -1.51608381384297 | 0.705 | 0.578 | 1 |
| **ARL4D** | 0.9817 | -0.870466333941037 | 0.613 | 0.56 | 1 |

Supplemental Table S2_Sheet_5

| | p_val | avg_log2FC | pct.1 | pct.2 | p_val_adj |
|---|---|---|---|---|---|
| **ATAD3B** | 0 | 1.94209689957208 | 1 | 0.956 | 0 |
| **TNFRSF8** | 0 | 2.0699612284696 | 0.846 | 0.067 | 0 |
| **FUCA1** | 0 | 1.97412294983203 | 0.999 | 0.814 | 0 |
| **CNR2** | 0 | 2.26398370728559 | 0.906 | 0.137 | 0 |
| **PTPRU** | 0 | 1.79943846480379 | 0.997 | 0.937 | 0 |
| **MARCKSL1** | 0 | -1.3600702208223 | 0.999 | 1 | 0 |
| **HPDL** | 0 | 1.98130731401245 | 0.901 | 0.154 | 0 |
| **PRDX1** | 0 | 2.06459186475557 | 1 | 0.991 | 0 |
| **WLS** | 0 | -4.55855168859975 | 0.556 | 0.988 | 0 |
| **BRDT** | 0 | 2.79637315180068 | 0.988 | 0.255 | 0 |
| **AGL** | 0 | 2.12025553907063 | 1 | 0.958 | 0 |
| **HENMT1** | 0 | 1.79570428527443 | 0.863 | 0.114 | 0 |
| **TAF13** | 0 | 2.61888319726583 | 1 | 0.839 | 0 |
| **OLFML3** | 0 | -4.49102646197968 | 0.671 | 0.996 | 0 |
| **HORMAD1** | 0 | 2.15077487263194 | 0.961 | 0.202 | 0 |
| **NES** | 0 | -2.24195363713204 | 0.987 | 1 | 0 |
| **DUSP23** | 0 | 2.02330233073903 | 0.91 | 0.098 | 0 |
| **IAH1** | 0 | 1.88070957070937 | 0.871 | 0.101 | 0 |
| **MTHFD2** | 0 | 1.84896907804943 | 0.999 | 0.994 | 0 |
| **REEP1** | 0 | 1.80831008165204 | 0.837 | 0.085 | 0 |
| **KIF5C** | 0 | -2.13342313052607 | 0.713 | 0.998 | 0 |
| **TMBIM1** | 0 | 2.60331463100508 | 0.994 | 0.448 | 0 |
| **ALPG** | 0 | 4.05814530578755 | 1 | 0.61 | 0 |
| **LRRN1** | 0 | -3.21073814210217 | 0.332 | 0.977 | 0 |
| **DPPA2** | 0 | 2.08590924684593 | 0.983 | 0.457 | 0 |
| **ATG3** | 0 | 2.42327400204213 | 1 | 0.995 | 0 |
| **TRH** | 0 | 1.82169525695024 | 0.796 | 0.083 | 0 |
| **AMOTL2** | 0 | -2.36917622621754 | 0.841 | 0.996 | 0 |
| **CHST2** | 0 | 2.98928507009138 | 0.991 | 0.512 | 0 |
| **UCHL1** | 0 | -2.66002040189857 | 0.634 | 0.992 | 0 |
| **PRSS12** | 0 | 2.32501627667493 | 0.994 | 0.562 | 0 |
| **TRIM60** | 0 | 2.30208473085032 | 0.973 | 0.189 | 0 |

| | | | | | |
|---|---|---|---|---|---|
| **TRIM75P** | 0 | 2.57156896175988 | 0.972 | 0.217 | 0 |
| **FAT1** | 0 | -2.39134411379152 | 0.798 | 0.999 | 0 |
| **ZFP42** | 0 | 4.41024470641164 | 1 | 0.385 | 0 |
| **MAP1B** | 0 | -2.3395331707672 | 0.951 | 1 | 0 |
| **DPYSL3** | 0 | -2.84876419033173 | 0.853 | 0.999 | 0 |
| **SERPINB1** | 0 | 1.59209333772783 | 0.805 | 0.106 | 0 |
| **SERPINB6** | 0 | 2.88763891030495 | 1 | 0.704 | 0 |
| **HIST1H1A** | 0 | 4.83780807170506 | 0.99 | 0.424 | 0 |
| **ZFP57** | 0 | 2.22451255208549 | 0.952 | 0.238 | 0 |
| **CCND3** | 0 | 1.96286905085682 | 0.995 | 0.709 | 0 |
| **DPPA5** | 0 | 6.02023446182621 | 0.997 | 0.524 | 0 |
| **KHDC3L** | 0 | 5.00681929361695 | 0.995 | 0.416 | 0 |
| **DDX43** | 0 | 1.60307419896538 | 0.77 | 0.029 | 0 |
| **IRAK1BP1** | 0 | 1.28130256532943 | 0.645 | 0.01 | 0 |
| **SLC16A10** | 0 | 2.55313325975706 | 0.997 | 0.544 | 0 |
| **MARCKS** | 0 | -3.81812263414412 | 0.302 | 0.997 | 0 |
| **DACT2** | 0 | 1.73525018080149 | 0.829 | 0.095 | 0 |
| **SUN3** | 0 | 1.68892343235708 | 0.801 | 0.031 | 0 |
| **CHCHD2** | 0 | 1.89231202338482 | 1 | 0.997 | 0 |
| **ZNF727** | 0 | 1.34272668047888 | 0.69 | 0.006 | 0 |
| **CDK6** | 0 | -2.37040614075871 | 0.155 | 0.943 | 0 |
| **ARPC1B** | 0 | 1.80884884254213 | 1 | 0.982 | 0 |
| **COL26A1** | 0 | -2.67420312397493 | 0.32 | 0.97 | 0 |
| **HYAL4** | 0 | 1.96300969705658 | 0.89 | 0.129 | 0 |
| **PODXL** | 0 | -4.46447642622727 | 0.775 | 0.997 | 0 |
| **DPYSL2** | 0 | -2.76786366743826 | 0.646 | 0.999 | 0 |
| **FGFR1** | 0 | -2.90952402747263 | 0.698 | 0.996 | 0 |
| **TMEM64** | 0 | -3.56882503613444 | 0.66 | 0.995 | 0 |
| **IGFBPL1** | 0 | -3.36649999525317 | 0.236 | 0.981 | 0 |
| **BARX1** | 0 | 1.44314578717806 | 0.7 | 0.019 | 0 |
| **KLF4** | 0 | 2.88336456899342 | 0.999 | 0.685 | 0 |
| **PALM2-AKAI** | 0 | -3.30746840122815 | 0.37 | 0.997 | 0 |
| **OLAH** | 0 | 2.6682981418518 | 0.95 | 0.28 | 0 |
| **SLC25A16** | 0 | 1.97156716536796 | 1 | 0.916 | 0 |

| | | | | | |
|---|---|---|---|---|---|
| **NFKB2** | 0 | 2.12366478106611 | 1 | 0.897 | 0 |
| **C10orf95** | 0 | 2.32666073869256 | 0.951 | 0.287 | 0 |
| **UTF1** | 0 | 3.2263931499731 | 0.994 | 0.429 | 0 |
| **PRAP1** | 0 | 2.94719625512343 | 0.955 | 0.188 | 0 |
| **SERPING1** | 0 | -2.23944385409 | 0.415 | 0.979 | 0 |
| **DPPA3** | 0 | 3.5539638962896 | 0.994 | 0.309 | 0 |
| **HMGA2** | 0 | -2.7386153313853 | 0.514 | 0.993 | 0 |
| **SPIC** | 0 | 3.17646479000446 | 0.969 | 0.242 | 0 |
| **RGCC** | 0 | 2.41069331688738 | 0.93 | 0.236 | 0 |
| **LCP1** | 0 | 3.27626386944823 | 0.992 | 0.645 | 0 |
| **KLF5** | 0 | 2.44900149332659 | 1 | 0.799 | 0 |
| **SALL2** | 0 | -2.34695415604879 | 0.771 | 0.998 | 0 |
| **SLC7A7** | 0 | 2.36430426629295 | 0.988 | 0.477 | 0 |
| **REC8** | 0 | 3.12334394997851 | 0.991 | 0.337 | 0 |
| **SPTSSA** | 0 | 1.55285856639139 | 1 | 1 | 0 |
| **PYGL** | 0 | 1.93318595914949 | 1 | 0.928 | 0 |
| **TMEM30B** | 0 | 1.57148258668224 | 0.766 | 0.022 | 0 |
| **HSPA2** | 0 | 2.09338709090665 | 0.877 | 0.164 | 0 |
| **GPX2** | 0 | 3.15622902892252 | 0.97 | 0.301 | 0 |
| **RAB15** | 0 | 1.62126174966926 | 1 | 0.998 | 0 |
| **VRTN** | 0 | -3.3853084900969 | 0.311 | 0.961 | 0 |
| **SLIRP** | 0 | 1.39240198733974 | 1 | 0.999 | 0 |
| **WARS** | 0 | 2.33535352128846 | 1 | 0.998 | 0 |
| **CRIP1** | 0 | 4.60077853795265 | 1 | 0.727 | 0 |
| **EID1** | 0 | -1.84999203878827 | 0.928 | 1 | 0 |
| **OAZ2** | 0 | -2.46342958721627 | 0.858 | 0.999 | 0 |
| **DNAJA4** | 0 | 1.72447464998301 | 0.827 | 0.042 | 0 |
| **SLC9A3R2** | 0 | 2.263593853271 | 0.995 | 0.641 | 0 |
| **IL32** | 0 | 2.37073537294259 | 0.932 | 0.207 | 0 |
| **NDUFAB1** | 0 | 1.44059252822272 | 1 | 1 | 0 |
| **SEPHS2** | 0 | 1.86200301487799 | 1 | 0.984 | 0 |
| **DNAJA2** | 0 | 1.14651049177173 | 1 | 1 | 0 |
| **ZNF423** | 0 | -2.42267257147348 | 0.434 | 0.981 | 0 |
| **MT1E** | 0 | 3.42195344444708 | 0.997 | 0.312 | 0 |

| | | | | | |
|---|---|---|---|---|---|
| **MT1G** | 0 | 4.52849165871968 | 0.999 | 0.377 | 0 |
| **MT1H** | 0 | 5.31524579423377 | 1 | 0.433 | 0 |
| **MT1X** | 0 | 3.15037487041705 | 1 | 0.962 | 0 |
| **RRAD** | 0 | 3.18636876986039 | 0.937 | 0.212 | 0 |
| **NTN1** | 0 | 5.07254753577592 | 1 | 0.985 | 0 |
| **PYCR1** | 0 | 2.1487677407872 | 0.996 | 0.558 | 0 |
| **TBCD** | 0 | 1.43116203099033 | 1 | 0.998 | 0 |
| **RNF125** | 0 | 2.33880428019978 | 1 | 0.773 | 0 |
| **ATP8B1** | 0 | 1.4158523510604 | 0.683 | 0.03 | 0 |
| **CNN2** | 0 | -2.01113681159472 | 0.99 | 1 | 0 |
| **ZNF57** | 0 | 1.70544932944796 | 0.809 | 0.032 | 0 |
| **TNFSF9** | 0 | 2.70935336295835 | 0.99 | 0.246 | 0 |
| **CD70** | 0 | 2.93665184108004 | 0.972 | 0.294 | 0 |
| **CD320** | 0 | 2.14618035169155 | 1 | 0.991 | 0 |
| **AP1M2** | 0 | 1.88890112466207 | 0.956 | 0.305 | 0 |
| **ACP5** | 0 | 1.51017460059048 | 0.735 | 0.023 | 0 |
| **ZNF676** | 0 | 2.74012721366647 | 0.897 | 0.233 | 0 |
| **ZNF729** | 0 | 2.72289312765813 | 0.742 | 0.049 | 0 |
| **ZNF728** | 0 | 1.28202265271193 | 0.661 | 0.009 | 0 |
| **DMKN** | 0 | -2.24975738291759 | 0.317 | 0.976 | 0 |
| **ZNF568** | 0 | 2.01398126717799 | 0.944 | 0.187 | 0 |
| **C19orf33** | 0 | 2.92884190831877 | 0.992 | 0.355 | 0 |
| **ATP1A3** | 0 | 2.0707632879083 | 0.941 | 0.228 | 0 |
| **LIM2** | 0 | 1.68209650259809 | 0.871 | 0.11 | 0 |
| **ZNF677** | 0 | 1.65510094694235 | 0.857 | 0.071 | 0 |
| **NLRP7** | 0 | 5.14145326200311 | 1 | 0.684 | 0 |
| **NLRP2** | 0 | 2.62168829975307 | 1 | 0.995 | 0 |
| **PEG3** | 0 | 2.04186256167737 | 0.742 | 0.032 | 0 |
| **PYGB** | 0 | 2.31870316450531 | 1 | 1 | 0 |
| **CPNE1** | 0 | -1.5913127687294 | 0.966 | 1 | 0 |
| **MYL9** | 0 | -3.45527732953175 | 0.858 | 0.999 | 0 |
| **TGIF2** | 0 | -2.44504084392295 | 0.558 | 0.995 | 0 |
| **SAMHD1** | 0 | 2.19723539727213 | 0.997 | 0.906 | 0 |
| **BMP7** | 0 | -2.79597227049441 | 0.545 | 0.995 | 0 |

| | | | | | |
|---|---|---|---|---|---|
| **DNMT3L** | 0 | 3.79777704123063 | 1 | 0.884 | 0 |
| **COL18A1** | 0 | -3.6446967486184 | 0.813 | 1 | 0 |
| **MED15** | 0 | 1.73691703615131 | 0.996 | 0.933 | 0 |
| **SUSD2** | 0 | 2.41700180302137 | 1 | 0.798 | 0 |
| **NEFH** | 0 | 3.04199016328476 | 0.999 | 0.622 | 0 |
| **SNU13** | 0 | 1.16900081827112 | 1 | 0.999 | 0 |
| **NDUFA6** | 0 | 1.41985143417287 | 1 | 0.97 | 0 |
| **A4GALT** | 0 | 2.62284211884884 | 1 | 0.756 | 0 |
| **CD99** | 0 | -2.75394038573957 | 0.23 | 0.966 | 0 |
| **TMSB4X** | 0 | -2.62982649728221 | 1 | 1 | 0 |
| **SMS** | 0 | -2.57652002240481 | 0.891 | 1 | 0 |
| **MAGED1** | 0 | -2.73658992190417 | 0.855 | 1 | 0 |
| **MAGED2** | 0 | -2.22213282708638 | 0.943 | 1 | 0 |
| **ARMCX2** | 0 | -2.05887023993914 | 0.724 | 0.998 | 0 |
| **BEX3** | 0 | -3.25162329434631 | 0.826 | 1 | 0 |
| **PGRMC1** | 0 | -1.78989783327895 | 0.973 | 1 | 0 |
| **GPC4** | 0 | -3.54417969617081 | 0.588 | 0.993 | 0 |
| **S100P** | 9.3367 | 2.12905185903475 | 0.733 | 0.064 | 1.5184392 |
| **OBSL1** | 1.3457 | -2.30969084676281 | 0.304 | 0.971 | 2.1885966 |
| **SEC63** | 1.3792 | 1.39296541128771 | 1 | 0.989 | 2.2430019 |
| **PIK3AP1** | 2.1376 | 1.33518879127954 | 0.718 | 0.048 | 3.4764986 |
| **LGALS3BP** | 1.1003 | -2.43481945789284 | 0.34 | 0.965 | 1.7895302 |
| **NGEF** | 1.7078 | 1.73850247038117 | 0.802 | 0.117 | 2.7775402 |
| **CACNA1H** | 2.2404 | 1.34757095391535 | 0.733 | 0.057 | 3.6436553 |
| **FGF4** | 2.3275 | 2.4381298275587 | 0.996 | 0.652 | 3.7853524 |
| **CRABP2** | 3.1559 | -3.17343682227192 | 0.925 | 0.994 | 5.1324806 |
| **COL4A2** | 3.2690 | -2.65654002187757 | 0.691 | 0.996 | 5.3164972 |
| **TFCP2L1** | 4.3912 | 1.97242112243259 | 0.999 | 0.631 | 7.1415095 |
| **TIMM50** | 4.5069 | 1.0872381771087 | 1 | 1 | 7.3296856 |
| **ILRUN** | 8.1502 | 1.55098461010781 | 1 | 0.996 | 1.3254786 |
| **VCAN** | 1.0106 | -2.92042995275789 | 0.734 | 0.996 | 1.6436770 |
| **SP5** | 1.0834 | -3.49928438263307 | 0.234 | 0.946 | 1.7619601 |
| **PFN2** | 1.0938 | -1.41386704216396 | 0.996 | 1 | 1.7788953 |
| **BEX4** | 3.2314 | -1.82750643894698 | 0.791 | 1 | 5.2553865 |

| | | | | | |
|---|---|---|---|---|---|
| **GPC6** | 4.1957 | -2.37067486118223 | 0.306 | 0.964 | 6.8234865 |
| **UPP1** | 1.1649 | 1.95551855351966 | 1 | 0.952 | 1.8945859 |
| **EFHD2** | 1.2656 | 1.77419021810582 | 1 | 0.96 | 2.0583854 |
| **TCN2** | 1.3331 | -2.2435219283219 | 0.276 | 0.963 | 2.1680936 |
| **TRIM24** | 8.9457 | -2.02533428222664 | 0.963 | 1 | 1.4548468 |
| **PIEZO1** | 1.6518 | 1.1660485772951 | 0.682 | 0.033 | 2.6864169 |
| **USP28** | 2.4610 | 1.35227044251548 | 1 | 0.98 | 4.0024228 |
| **LIN28A** | 2.7196 | -1.58737897689254 | 1 | 1 | 4.4228957 |
| **BAK1** | 1.6505 | 1.27206126491323 | 1 | 0.998 | 2.6842564 |
| **CDS2** | 1.3504 | -1.68182837838679 | 0.836 | 0.998 | 2.1962795 |
| **DNAJC15** | 1.5748 | 2.36111284988449 | 0.997 | 0.992 | 2.5611082 |
| **GPT2** | 3.6612 | 1.87144634187327 | 0.999 | 0.935 | 5.9542530 |
| **SMPDL3A** | 3.9301 | 1.82334920166073 | 0.953 | 0.403 | 6.3915682 |
| **BIN1** | 4.6838 | -2.48728485184396 | 0.377 | 0.972 | 7.6172972 |
| **FADS2** | 5.3724 | -1.93652149519628 | 0.996 | 1 | 8.7371822 |
| **HEPH** | 6.8102 | -2.21895333736968 | 0.034 | 0.895 | 1.1075491 |
| **TCEAL9** | 7.3079 | -2.04868581994882 | 0.597 | 0.994 | 1.1884937 |
| **GLDC** | 4.9714 | 1.7187125500298 | 0.997 | 0.809 | 8.0851228 |
| **SELENOP** | 2.1756 | -2.3419234138359 | 0.19 | 0.941 | 3.5382585 |
| **FCGBP** | 2.3375 | 1.73552330464909 | 0.707 | 0.054 | 3.8015410 |
| **MAP4K4** | 4.3596 | -1.38069621607371 | 0.988 | 1 | 7.0901722 |
| **MBNL3** | 5.4491 | -2.57927735537122 | 0.292 | 0.958 | 8.8619237 |
| **CHD7** | 1.1166 | -2.51969841468444 | 0.376 | 0.958 | 1.8160292 |
| **COL4A1** | 8.1250 | -2.55314949032666 | 0.687 | 0.992 | 1.3213829 |
| **SLC6A8** | 1.0429 | 1.72225651950419 | 1 | 0.979 | 1.6961066 |
| **TRO** | 2.2341 | -1.95628168553007 | 0.437 | 0.98 | 3.6333527 |
| **ZFP3** | 2.6488 | 1.25868855055944 | 0.64 | 0.023 | 4.3077691 |
| **ELF3** | 2.8433 | 1.9461668726356 | 0.994 | 0.697 | 4.6241010 |
| **LITAF** | 4.8114 | -2.12893341865115 | 0.979 | 0.999 | 7.8249274 |
| **IGF2BP1** | 5.2034 | -1.6199373315403 | 0.935 | 1 | 8.4624367 |
| **ZNF98** | 1.1644 | 1.08091854029774 | 0.603 | 0.01 | 1.8936863 |
| **PLXND1** | 1.8068 | 2.20539302124771 | 0.997 | 0.761 | 2.9385479 |
| **BST2** | 1.8353 | -3.79956846887194 | 0.231 | 0.934 | 2.9848203 |
| **FOXR1** | 2.1641 | 1.12894695886223 | 0.588 | 0.005 | 3.5195878 |

| | | | | | |
|---|---|---|---|---|---|
| **VIM** | 4.5361 | -5.10392002824342 | 0.673 | 0.981 | 7.3771989 |
| **ARMCX1** | 6.8620 | -1.97442874104484 | 0.457 | 0.991 | 1.1159697 |
| **SLC16A2** | 1.7898 | -2.39381922051729 | 0.358 | 0.969 | 2.9109118 |
| **HCLS1** | 3.6410 | 1.67292998933409 | 0.738 | 0.081 | 5.9215006 |
| **YPEL2** | 3.7543 | 2.01430892614797 | 0.995 | 0.589 | 6.1057428 |
| **NDUFV1** | 4.9581 | 1.10188011245525 | 1 | 1 | 8.0633675 |
| **IGF2R** | 1.7710 | -1.97041565212944 | 0.521 | 0.978 | 2.8803183 |
| **LGALS3** | 1.9538 | 1.92939344121529 | 0.829 | 0.158 | 3.1774909 |
| **C9orf64** | 3.1965 | 1.09686153925259 | 0.619 | 0.017 | 5.1985209 |
| **SPSB4** | 4.7522 | -2.42330923724239 | 0.205 | 0.927 | 7.7286643 |
| **NUPR1** | 6.3593 | 2.49516718294641 | 0.84 | 0.175 | 1.0342155 |
| **EFR3B** | 4.5649 | 1.65348798783631 | 1 | 0.829 | 7.4239595 |
| **ZIC3** | 6.7307 | -2.25493810332841 | 0.266 | 0.941 | 1.0946292 |
| **PLBD1** | 6.7325 | 1.72322031722725 | 1 | 0.917 | 1.0949217 |
| **COL4A5** | 7.6500 | -2.61742698979844 | 0.093 | 0.896 | 1.2441267 |
| **FOXO1** | 4.4942 | -2.42973070453734 | 0.606 | 0.982 | 7.3090074 |
| **NRDC** | 7.2993 | 1.01952106581728 | 1 | 1 | 1.1870868 |
| **GFPT1** | 9.4982 | 1.44069195406539 | 0.999 | 0.998 | 1.5446965 |
| **CTNNB1** | 1.0594 | -1.53378463212869 | 0.986 | 0.999 | 1.7229127 |
| **TMEM54** | 1.6546 | 1.61587230474217 | 0.898 | 0.25 | 2.6909994 |
| **VENTX** | 1.8537 | 1.78780338563065 | 0.961 | 0.395 | 3.0147965 |
| **CR1L** | 7.1962 | 1.9695554081795 | 0.97 | 0.415 | 1.1703189 |
| **SH3BGRL** | 9.0852 | -1.88704630915378 | 0.391 | 0.975 | 1.4775320 |
| **FBLN1** | 1.0419 | -1.88373849604763 | 0.876 | 0.998 | 1.6945757 |
| **ECE1** | 2.0572 | -2.02377857423712 | 0.727 | 0.994 | 3.3457106 |
| **CFD** | 2.3922 | 1.66861031727726 | 0.906 | 0.274 | 3.8904808 |
| **JPT1** | 2.7875 | 1.25278960164995 | 1 | 0.999 | 4.5333888 |
| **UBE2E3** | 6.4900 | -1.5772010208779 | 0.863 | 0.997 | 1.0554844 |
| **IGF1R** | 8.5387 | -1.76120957643852 | 0.904 | 0.999 | 1.3886615 |
| **SLC66A2** | 1.2293 | 1.5433248139099 | 0.999 | 0.918 | 1.9992213 |
| **ARMT1** | 1.2721 | 1.40847412791153 | 1 | 0.988 | 2.0689265 |
| **GDF3** | 2.1014 | 2.14874132664918 | 0.995 | 0.804 | 3.4176542 |
| **SLC5A5** | 4.3149 | 1.33632825284194 | 0.774 | 0.105 | 7.0173708 |
| **SNX19** | 1.4340 | 1.14301040117608 | 0.667 | 0.047 | 2.3321173 |

| | | | | | |
|---|---|---|---|---|---|
| **MYEF2** | 1.7327 | -2.07065880163318 | 0.395 | 0.968 | 2.8179555 |
| **LIPH** | 4.0550 | 1.68158883712983 | 0.714 | 0.077 | 6.5946810 |
| **FZD7** | 4.3567 | -3.97640378605998 | 0.14 | 0.9 | 7.0853794 |
| **SNRPC** | 8.7032 | 0.986432587308583 | 1 | 1 | 1.4154119 |
| **EFNA5** | 1.0604 | -2.39931368932782 | 0.128 | 0.899 | 1.7245895 |
| **PINK1** | 1.8035 | 1.40373716248048 | 1 | 0.957 | 2.9331772 |
| **TEAD4** | 2.3390 | 1.3536369546147 | 1 | 0.965 | 3.8040647 |
| **DUSP9** | 3.5045 | 1.26832964199019 | 0.661 | 0.043 | 5.6993881 |
| **PLEKHB2** | 3.8768 | 1.11540697739261 | 1 | 0.999 | 6.3049854 |
| **IRF6** | 7.2276 | 1.14498995509958 | 0.678 | 0.048 | 1.1754367 |
| **HIST1H2BB** | 8.4204 | 2.17394125588983 | 0.678 | 0.056 | 1.3694098 |
| **TMEM160** | 9.1854 | 1.32006657634439 | 1 | 0.998 | 1.4938219 |
| **FLAD1** | 1.1830 | 1.4096554465123 | 1 | 0.966 | 1.9239549 |
| **LOXL2** | 1.2779 | -2.28229173706351 | 0.448 | 0.975 | 2.0783497 |
| **FABP5** | 1.3600 | 2.35479408619048 | 0.975 | 0.552 | 2.2119005 |
| **MGLL** | 4.6249 | 1.30600968648503 | 0.667 | 0.048 | 7.5215556 |
| **SEPTIN6** | 1.6623 | -1.97202316943668 | 0.257 | 0.938 | 2.7035119 |
| **MFGE8** | 1.9847 | -2.08268940994292 | 0.337 | 0.945 | 3.2277202 |
| **ERBIN** | 2.5309 | 1.42773217728223 | 1 | 0.957 | 4.1160591 |
| **ATG5** | 4.2707 | 1.46109336347676 | 1 | 0.949 | 6.9454844 |
| **CSF2RA** | 6.8867 | 1.89458735505573 | 0.862 | 0.234 | 1.1199882 |
| **FZD5** | 1.4298 | 1.56572757066896 | 0.997 | 0.952 | 2.3253171 |
| **SYNGR3** | 1.4336 | 1.59475254385836 | 0.987 | 0.716 | 2.3315735 |
| **AQP3** | 4.1561 | 1.70279207828049 | 0.815 | 0.156 | 6.7591512 |
| **RBP1** | 5.2325 | -2.32611098474856 | 0.329 | 0.939 | 8.5096895 |
| **UBQLN2** | 1.3118 | -1.46853441528715 | 0.839 | 1 | 2.1334376 |
| **SH3KBP1** | 2.4041 | -1.89938551822258 | 0.709 | 0.99 | 3.9098354 |
| **GAD1** | 2.9403 | -2.33967960822823 | 0.071 | 0.874 | 4.7818219 |
| **APP** | 3.4915 | -1.37731411867464 | 0.999 | 1 | 5.6783784 |
| **ZNF253** | 4.1025 | 0.980276286101071 | 0.589 | 0.015 | 6.6720057 |
| **TGFBR3** | 5.8412 | 1.47590012286465 | 0.831 | 0.176 | 9.4996245 |
| **MMP2** | 6.7612 | -3.07914373540619 | 0.428 | 0.959 | 1.0995885 |
| **TIMP1** | 2.7047 | -1.75265951884204 | 0.943 | 0.999 | 4.3987474 |
| **TCP1** | 2.7815 | 1.75026319741846 | 1 | 0.924 | 4.5236545 |

| | | | | | |
|---|---|---|---|---|---|
| **CTPS2** | 2.8819 | -1.4954548907187 | 0.766 | 0.998 | 4.6869259 |
| **AFAP1L1** | 3.1380 | 1.52596045210553 | 0.807 | 0.156 | 5.1034868 |
| **MEX3A** | 1.1033 | -2.37119331369259 | 0.174 | 0.898 | 1.7943690 |
| **FKBP10** | 1.4257 | -2.31089113734029 | 0.464 | 0.961 | 2.3187317 |
| **SLC25A39** | 2.3340 | 0.931466612578929 | 1 | 1 | 3.7958555 |
| **ARHGEF9** | 3.2611 | -1.88294777143954 | 0.128 | 0.893 | 5.3036681 |
| **RMND1** | 5.3464 | 1.33933998852944 | 1 | 0.952 | 8.6949998 |
| **SYNGR2** | 5.9729 | 1.25208793587797 | 0.999 | 0.992 | 9.7137837 |
| **PPP2R1B** | 6.8709 | -1.42183434140329 | 0.876 | 0.997 | 1.1174190 |
| **CCND1** | 8.4842 | -1.7425452838264 | 0.961 | 0.997 | 1.3798002 |
| **YAP1** | 3.5953 | -1.12513716646009 | 0.992 | 1 | 5.8470456 |
| **C11orf96** | 3.7039 | -2.82945446222405 | 0.306 | 0.932 | 6.0238130 |
| **ECHDC1** | 9.5508 | -1.77407990146625 | 0.211 | 0.935 | 1.5532468 |
| **DCAF4** | 1.2396 | 1.38881457469446 | 1 | 0.912 | 2.0159651 |
| **BTBD3** | 1.3530 | -2.02085577353841 | 0.593 | 0.978 | 2.2004243 |
| **ASH2L** | 3.8065 | 1.2527730644527 | 1 | 0.99 | 6.1906284 |
| **GSPT2** | 4.4596 | -1.78727607324232 | 0.45 | 0.975 | 7.2527186 |
| **CYC1** | 7.1231 | 0.92990173307543 | 1 | 0.999 | 1.1584455 |
| **HSPA9** | 1.2327 | 1.11200975526537 | 1 | 0.996 | 2.0048247 |
| **TCAF1** | 2.4836 | -1.84163710122987 | 0.404 | 0.954 | 4.0391329 |
| **MAEL** | 1.2570 | 1.47541874888885 | 0.597 | 0.022 | 2.0443242 |
| **NARS** | 1.2950 | 0.882772198023331 | 1 | 1 | 2.1060990 |
| **AK4** | 1.4952 | 2.25796727721745 | 0.995 | 0.884 | 2.4316933 |
| **TBX3** | 1.8114 | 1.77828604867784 | 0.787 | 0.139 | 2.9458987 |
| **NDUFA12** | 5.0512 | 1.31362562175088 | 1 | 0.94 | 8.2147743 |
| **NRBF2** | 5.5400 | 1.36621434240942 | 1 | 0.983 | 9.0097269 |
| **GLI2** | 1.3114 | -1.92745332368042 | 0.065 | 0.858 | 2.1328183 |
| **VSIG10** | 1.3827 | 1.27113686993179 | 1 | 0.932 | 2.2487093 |
| **WWC3** | 1.6456 | -2.04908961818516 | 0.194 | 0.906 | 2.6762790 |
| **CCNQ** | 2.1851 | 1.39212488050144 | 0.991 | 0.809 | 3.5536776 |
| **ALDH1L2** | 2.8982 | 1.50733349426513 | 0.787 | 0.139 | 4.7133825 |
| **HNF4A** | 2.9647 | 1.07559056741609 | 0.605 | 0.026 | 4.8215347 |
| **NRG1** | 3.1234 | 1.64792589957977 | 0.786 | 0.145 | 5.0795912 |
| **MFSD13A** | 3.5899 | 1.59991455115368 | 0.912 | 0.318 | 5.8382549 |

| | | | | | |
|---|---|---|---|---|---|
| **DRAXIN** | 6.3533 | -2.23502365655067 | 0.54 | 0.965 | 1.0332421 |
| **MAGEH1** | 2.1317 | -1.8071772174271 | 0.225 | 0.925 | 3.4668348 |
| **RCBTB1** | 3.9368 | 1.75940055110273 | 0.995 | 0.954 | 6.4024463 |
| **MTA1** | 5.2747 | -1.21252214796512 | 0.968 | 1 | 8.5783391 |
| **EEF1A2** | 6.5862 | 1.37130798094176 | 0.734 | 0.092 | 1.0711216 |
| **ECHDC3** | 7.1798 | 1.05595640597402 | 0.563 | 0.012 | 1.1676571 |
| **UBAP2** | 2.3522 | 1.22647833294079 | 0.997 | 0.977 | 3.8254326 |
| **NAP1L3** | 4.1067 | -2.20064794782745 | 0.212 | 0.907 | 6.6788484 |
| **NDUFB10** | 1.4276 | 0.878599683006644 | 1 | 1 | 2.3218667 |
| **ALDH2** | 3.3790 | 1.42780755968989 | 0.999 | 0.942 | 5.4954180 |
| **NF2** | 6.5332 | 1.0600910579517 | 1 | 0.996 | 1.0624948 |
| **OAS1** | 1.2910 | 1.14391836331336 | 0.59 | 0.023 | 2.0996470 |
| **SDHB** | 3.4859 | 1.14773652902686 | 1 | 0.996 | 5.6692600 |
| **APMAP** | 9.4377 | -1.38518526658827 | 0.947 | 0.999 | 1.5348609 |
| **TSPAN6** | 1.6591 | -1.6612239663762 | 0.904 | 0.997 | 2.6983008 |
| **ASRGL1** | 2.3437 | 1.32512298189589 | 1 | 0.954 | 3.8117012 |
| **ZNF257** | 2.8547 | 0.925939883364391 | 0.53 | 0.003 | 4.6426442 |
| **SLC25A24** | 3.7289 | -1.5190759176785 | 0.773 | 0.993 | 6.0643777 |
| **SLC8B1** | 4.3876 | 1.62777638465581 | 0.965 | 0.57 | 7.1356274 |
| **EFNB1** | 6.0674 | -3.20126135244266 | 0.231 | 0.904 | 9.8674506 |
| **PKIB** | 7.4317 | 1.28124343508201 | 0.643 | 0.047 | 1.2086206 |
| **PGP** | 1.2443 | 1.0371043245923 | 1 | 0.997 | 2.0237296 |
| **PCDH1** | 1.6697 | -2.10930531887712 | 0.77 | 0.987 | 2.7154947 |
| **TOP1MT** | 6.7948 | 1.36404268182716 | 1 | 0.954 | 1.1050421 |
| **PIGT** | 7.2505 | -1.37439738943274 | 0.948 | 0.998 | 1.1791515 |
| **NMRAL1** | 9.9951 | 1.46135077191495 | 0.994 | 0.75 | 1.6255150 |
| **TSPAN18** | 1.7564 | -1.72315153251645 | 0.901 | 0.997 | 2.8565152 |
| **CLU** | 1.9884 | -2.00915118916681 | 0.946 | 1 | 3.2338419 |
| **CHSY1** | 1.5211 | -1.56291556170288 | 0.925 | 0.998 | 2.4738624 |
| **PCDHB2** | 3.1225 | -1.86113937726496 | 0.315 | 0.917 | 5.0782305 |
| **NECTIN3** | 3.5623 | -1.91580403952618 | 0.276 | 0.917 | 5.7934323 |
| **AMOTL1** | 5.1328 | -2.17200664595064 | 0.28 | 0.91 | 8.3475298 |
| **MYO1G** | 8.1998 | 1.00990448940899 | 0.547 | 0.011 | 1.3335464 |
| **DFFA** | 8.6652 | 1.12538631428313 | 1 | 0.996 | 1.4092264 |

| | | | | | |
|---|---|---|---|---|---|
| **COL1A2** | 1.0369 | -3.6979647495776 | 0.417 | 0.95 | 1.6864684 |
| **TCF4** | 1.0383 | -1.9438271843592 | 0.238 | 0.897 | 1.6887034 |
| **PLD3** | 2.0391 | -1.52614774307127 | 0.991 | 1 | 3.3162623 |
| **BCAM** | 3.3412 | -2.3182173211031 | 0.568 | 0.965 | 5.4339470 |
| **CLPX** | 3.6068 | 1.21995074754991 | 1 | 0.987 | 5.8658358 |
| **ACOT4** | 7.9478 | 0.946711206719501 | 0.556 | 0.014 | 1.2925604 |
| **DUSP16** | 1.3362 | 1.6437094455337 | 0.997 | 0.87 | 2.1732188 |
| **KRT8** | 1.5546 | -2.11411461738451 | 0.994 | 1 | 2.5283312 |
| **TMEM150C** | 1.6238 | 1.36577581496186 | 0.791 | 0.163 | 2.6407942 |
| **STK26** | 1.8028 | -1.66425611707189 | 0.572 | 0.993 | 2.9319713 |
| **HS6ST2** | 1.8459 | -2.06603631400841 | 0.094 | 0.864 | 3.0021178 |
| **LPP** | 2.9244 | -2.0727363754433 | 0.38 | 0.933 | 4.7560951 |
| **SHPK** | 3.0255 | 1.52258954182822 | 0.997 | 0.885 | 4.9204793 |
| **PEA15** | 1.2754 | -1.87229880067294 | 0.788 | 0.994 | 2.0742800 |
| **ROR1** | 2.4245 | -2.55578246730456 | 0.057 | 0.832 | 3.9429929 |
| **SLC9A3** | 8.0461 | 1.32628646646974 | 0.641 | 0.052 | 1.3085511 |
| **ZNF502** | 8.9661 | 0.937154592880699 | 0.535 | 0.008 | 1.4581615 |
| **PPP1R3B** | 1.4182 | -2.27808047302089 | 0.562 | 0.964 | 2.3064618 |
| **H2AFY2** | 2.0302 | -2.1388695562146 | 0.624 | 0.97 | 3.3018068 |
| **SRM** | 2.6701 | 1.04789990120215 | 1 | 1 | 4.3424782 |
| **DLAT** | 4.3853 | 1.04239233024585 | 1 | 0.994 | 7.1318227 |
| **KCNQ2** | 4.8543 | 1.45099430610746 | 0.841 | 0.218 | 7.8946332 |
| **STAU1** | 6.6736 | -0.873798490222535 | 1 | 1 | 1.0853424 |
| **PDE4A** | 9.9657 | 1.62987295880919 | 0.93 | 0.395 | 1.6207337 |
| **SMPDL3B** | 1.3979 | -2.25980168046515 | 0.043 | 0.819 | 2.2734381 |
| **ACTN1** | 2.2188 | -1.5859692161372 | 0.996 | 1 | 3.6085575 |
| **IGDCC3** | 2.5063 | -1.95140393081154 | 0.22 | 0.889 | 4.0761240 |
| **MATK** | 3.7429 | 1.38103826216086 | 0.774 | 0.137 | 6.0870782 |
| **DYNLRB1** | 7.5658 | -1.02521782896843 | 1 | 1 | 1.2304388 |
| **NSMAF** | 9.8645 | 1.28963339861145 | 1 | 0.937 | 1.6042760 |
| **PGM1** | 1.2262 | -1.81020516925325 | 0.372 | 0.953 | 1.9942432 |
| **SLC38A1** | 1.5880 | -1.86273951433846 | 0.509 | 0.982 | 2.5825658 |
| **LDOC1** | 1.7798 | -1.62959731063584 | 0.453 | 0.962 | 2.8944960 |
| **HIC2** | 6.3617 | 1.29940797302946 | 0.996 | 0.929 | 1.0346159 |

| | | | | | |
|---|---|---|---|---|---|
| **MT1F** | 1.0517 | 1.67942979368762 | 0.996 | 0.878 | 1.7104512 |
| **GSN** | 2.1589 | -2.46745687627334 | 0.074 | 0.831 | 3.5111142 |
| **ACSL3** | 2.3525 | 1.07801544522859 | 1 | 0.999 | 3.8258904 |
| **FNDC10** | 3.2598 | 1.59944441618392 | 0.977 | 0.588 | 5.3014744 |
| **AFG3L2** | 7.1509 | 0.887487072912298 | 1 | 0.999 | 1.1629537 |
| **TRPC6** | 1.8011 | -2.7160720171967 | 0.181 | 0.87 | 2.9291882 |
| **TUBA1C** | 2.9641 | 1.39372909241213 | 1 | 0.997 | 4.8206473 |
| **F11R** | 4.9535 | 1.09218442857391 | 1 | 0.988 | 8.0560366 |
| **ZHX2** | 6.8214 | -1.85604766032003 | 0.066 | 0.828 | 1.1093782 |
| **HBA2** | 9.9399 | 1.26771171258244 | 0.632 | 0.052 | 1.6165330 |
| **ERGIC3** | 1.9237 | -0.940045525648246 | 1 | 1 | 3.1286541 |
| **EXPH5** | 3.9369 | 1.43402757889891 | 0.809 | 0.195 | 6.4026172 |
| **PTPN14** | 5.3481 | -1.38039768546377 | 0.841 | 0.998 | 8.6977714 |
| **NDUFB7** | 9.4089 | 0.817294607244323 | 1 | 1 | 1.5301816 |
| **ZNF219** | 1.3241 | -1.94944804429448 | 0.547 | 0.96 | 2.1534176 |
| **TIMM10** | 1.4864 | 1.01346704103703 | 1 | 0.995 | 2.4174348 |
| **TPST2** | 2.6248 | 1.38945680472826 | 0.994 | 0.815 | 4.2688250 |
| **MCRIP1** | 5.0634 | -1.3532798659845 | 0.939 | 0.998 | 8.2346706 |
| **EDN1** | 6.5425 | -2.88502100824443 | 0.047 | 0.814 | 1.0640116 |
| **SMARCA1** | 1.6899 | -1.8525718920202 | 0.444 | 0.971 | 2.7483602 |
| **NME4** | 2.1969 | -1.31293779239326 | 0.999 | 1 | 3.5728503 |
| **TCEAL8** | 2.5321 | -1.35320387430026 | 0.875 | 1 | 4.1179819 |
| **CLSTN1** | 4.8001 | -1.18705176018482 | 0.997 | 0.999 | 7.8064914 |
| **FNBP1L** | 5.3520 | -1.11912186471053 | 0.996 | 1 | 8.7039849 |
| **USP11** | 5.7306 | -1.34601497302059 | 0.897 | 0.998 | 9.3197540 |
| **PDXK** | 7.6389 | 1.13749921898694 | 1 | 0.988 | 1.2423301 |
| **NOP16** | 2.2248 | 1.12640358203828 | 1 | 0.993 | 3.6182485 |
| **TRIB3** | 2.9261 | 1.95016345940087 | 0.982 | 0.665 | 4.7588524 |
| **ITGB1** | 5.6833 | -1.33559636753712 | 0.955 | 1 | 9.2427786 |
| **MCL1** | 7.2072 | 0.884828855322881 | 1 | 1 | 1.1721162 |
| **UQCRQ** | 1.6460 | 1.14144604254918 | 1 | 0.992 | 2.6769925 |
| **REEP6** | 1.6951 | 1.6577592301714 | 0.97 | 0.627 | 2.7567629 |
| **ZNF736** | 7.7235 | 0.85065587598014 | 0.519 | 0.008 | 1.2560824 |
| **AGRN** | 1.0445 | -1.55458558861036 | 0.941 | 0.998 | 1.6986733 |

| | | | | | |
|---|---|---|---|---|---|
| **NPC2** | 1.5047 | -1.2712107701158 | 1 | 1 | 2.44716165 |
| **LAMB1** | 1.7299 | -1.70595336733798 | 0.836 | 0.996 | 2.81343028 |
| **PCYT1B** | 2.9580 | -1.76250944739949 | 0.062 | 0.814 | 4.81066582 |
| **CEP68** | 3.1933 | -1.5958905556233 | 0.407 | 0.933 | 5.19332838 |
| **BHMT** | 4.2719 | 1 | 0.521 | 0.01 | 6.94749723 |
| **CFAP46** | 5.6773 | 0.983369325455879 | 0.62 | 0.049 | 9.23306687 |
| **TRIM61** | 8.0415 | 1.4153552945179 | 0.893 | 0.332 | 1.30779762 |
| **CTHRC1** | 8.2813 | 1.25029720492202 | 0.656 | 0.07 | 1.34679556 |
| **ARRDC2** | 8.2887 | 1.4608310210122 | 0.895 | 0.34 | 1.34800443 |
| **RTBDN** | 1.0620 | 1.1718865177756 | 0.678 | 0.081 | 1.72719678 |
| **NOTCH3** | 1.2434 | -1.80273221326978 | 0.879 | 0.996 | 2.02216280 |
| **JAG1** | 1.2742 | -1.77716993726381 | 0.885 | 0.995 | 2.07229345 |
| **RBBP9** | 1.3294 | 1.40725983849013 | 0.992 | 0.863 | 2.16209213 |
| **MARVELD2** | 4.3351 | 1.45168384849883 | 0.88 | 0.298 | 7.05032159 |
| **CST3** | 4.8152 | -2.22511253696139 | 0.678 | 0.988 | 7.83110171 |
| **NDUFC2** | 4.9763 | 0.872401980391285 | 1 | 1 | 8.09309525 |
| **RGS5** | 6.2833 | -2.98907336509096 | 0.161 | 0.851 | 1.02186793 |
| **AKAP1** | 4.6849 | 1.14626304765905 | 0.999 | 0.977 | 7.61913413 |
| **DNAJA3** | 5.9735 | 1.04960067270518 | 1 | 0.99 | 9.71472319 |
| **CITED2** | 8.8353 | -2.3327199452297 | 0.62 | 0.975 | 1.43688761 |
| **PCLAF** | 9.2876 | -2.28891490227982 | 0.07 | 0.81 | 1.51044440 |
| **AP1S2** | 1.5206 | -1.94824358255527 | 0.245 | 0.881 | 2.47307575 |
| **CRYGD** | 2.3316 | 1.08085650905643 | 0.534 | 0.016 | 3.79188828 |
| **TNFRSF21** | 2.5190 | -1.69875107199648 | 0.906 | 0.996 | 4.09673850 |
| **IFITM3** | 3.9225 | -1.85013453137922 | 0.99 | 0.999 | 6.37918697 |
| **GAS7** | 4.1572 | 1.41113760685228 | 0.999 | 0.881 | 6.76094391 |
| **PKP3** | 5.0544 | 1.53281551127933 | 0.99 | 0.65 | 8.22010360 |
| **PLEKHO1** | 6.0935 | -1.99907965489514 | 0.313 | 0.902 | 9.90994750 |
| **PFKM** | 6.1559 | -1.44670512329777 | 0.686 | 0.98 | 1.00114035 |
| **YIPF4** | 7.7749 | 1.27174383992012 | 0.997 | 0.955 | 1.26444153 |
| **GAL** | 1.2422 | 1.69602915206178 | 0.992 | 0.702 | 2.02033350 |
| **CPXM1** | 1.5799 | -1.82594588143211 | 0.707 | 0.981 | 2.56953961 |
| **PPP1R14A** | 1.8874 | 1.34789264062211 | 0.99 | 0.778 | 3.06962313 |
| **EIF4A3** | 3.6032 | 0.890341184743264 | 1 | 1 | 5.85996337 |

| | | | | | |
|---|---|---|---|---|---|
| **EFNB2** | 4.3261 | -2.27423814946948 | 0.413 | 0.941 | 7.0355777 |
| **ZCCHC3** | 1.8185 | -1.77674446464386 | 0.494 | 0.95 | 2.9575771 |
| **MAGEB2** | 1.9902 | 1.38659424049253 | 0.528 | 0.016 | 3.2368177 |
| **LGALS8** | 2.2619 | 0.959317364182095 | 0.531 | 0.016 | 3.6785298 |
| **FOXO4** | 2.4039 | -2.30580716192448 | 0.218 | 0.883 | 3.9095977 |
| **SLK** | 4.5375 | 1.32649136902182 | 0.997 | 0.951 | 7.3794527 |
| **NBEA** | 4.6154 | -1.98413752018885 | 0.062 | 0.804 | 7.5060964 |
| **RHOF** | 4.7947 | 1.41066470156847 | 0.941 | 0.449 | 7.7976559 |
| **PHLDA1** | 7.1180 | -2.54300902521231 | 0.468 | 0.923 | 1.1576101 |
| **MYBL2** | 7.5852 | 1.01354059823 | 1 | 0.989 | 1.2335887 |
| **ATP5F1D** | 7.5879 | 0.849414930993307 | 1 | 1 | 1.2340245 |
| **RAB13** | 8.2187 | -1.28372807181706 | 0.956 | 0.996 | 1.3366154 |
| **MIF** | 8.7536 | 0.86134360829582 | 1 | 1 | 1.4236054 |
| **TBC1D23** | 4.2907 | 1.15558025491554 | 1 | 0.975 | 6.9781200 |
| **CKAP4** | 7.4297 | -1.43168606646892 | 0.96 | 1 | 1.2082935 |
| **PRKAR2A** | 7.8440 | 1.03973398557827 | 0.999 | 0.972 | 1.2756826 |
| **MFAP2** | 1.0893 | -1.78674010382192 | 0.857 | 0.992 | 1.7716680 |
| **CCDC88A** | 1.3579 | -1.44836510076832 | 0.748 | 0.991 | 2.2084860 |
| **KIAA1522** | 2.1145 | -1.33131693887774 | 0.95 | 0.999 | 3.4389546 |
| **ABCB8** | 2.4589 | 1.1413387610522 | 0.999 | 0.963 | 3.9989193 |
| **TRIM71** | 3.6512 | -1.1408117539801 | 0.997 | 0.999 | 5.9380686 |
| **RASGRP2** | 5.7771 | 1.69862695636529 | 0.981 | 0.57 | 9.3953272 |
| **MSI2** | 9.3357 | -1.56840340884775 | 0.455 | 0.946 | 1.5182665 |
| **ZYG11A** | 1.0139 | 1.4499098983579 | 1 | 0.844 | 1.6490049 |
| **FAM71E1** | 1.0707 | 1.31952910029444 | 0.824 | 0.222 | 1.7414394 |
| **MSN** | 2.0248 | -1.32978139985536 | 0.864 | 1 | 3.2930341 |
| **LIN28B** | 3.2035 | 1.0921223061341 | 1 | 0.993 | 5.2098696 |
| **XRN2** | 6.3968 | -0.809461720767935 | 1 | 1 | 1.0403176 |
| **GDI2** | 1.1940 | -1.04437199990285 | 0.994 | 1 | 1.9418239 |
| **ESRRA** | 1.8526 | 1.18808768612273 | 0.997 | 0.92 | 3.0129058 |
| **EGLN3** | 3.1749 | 1.49173545868595 | 0.995 | 0.785 | 5.1634802 |
| **PCSK5** | 1.3391 | 1.61702439973593 | 0.916 | 0.421 | 2.1778451 |
| **ID3** | 1.4705 | -3.39239044289979 | 0.087 | 0.806 | 2.3916018 |
| **IGF2BP3** | 1.5261 | -1.06806125766051 | 0.995 | 1 | 2.4820443 |

| | | | | | |
|---|---|---|---|---|---|
| **SEPTIN9** | 2.7725 | -1.06236487202264 | 0.995 | 1 | 4.5089428 |
| **ATP5F1E** | 6.7377 | -1.12533577555515 | 0.995 | 1 | 1.0957601 |
| **PEAK1** | 9.5901 | -1.72383323032529 | 0.167 | 0.855 | 1.5596535 |
| **SESN3** | 2.0682 | -1.77450973396731 | 0.87 | 0.996 | 3.3636724 |
| **ARHGAP8** | 2.3085 | 1.395854891476 | 0.988 | 0.745 | 3.7543344 |
| **TNNI3** | 3.5944 | 1.33736479874965 | 0.849 | 0.252 | 5.8456161 |
| **CAPN2** | 3.8257 | -2.46499472725116 | 0.236 | 0.869 | 6.2218020 |
| **RRP7A** | 4.9313 | 1.09686683347832 | 0.999 | 0.972 | 8.0197954 |
| **PRXL2A** | 1.0076 | -1.15436274493206 | 0.977 | 0.999 | 1.6387704 |
| **TP53** | 1.4885 | -1.25239692742271 | 0.982 | 0.999 | 2.4208856 |
| **EDA** | 1.8195 | -1.66676468865714 | 0.329 | 0.92 | 2.9590733 |
| **CNOT11** | 3.0227 | 0.872613578606447 | 1 | 0.998 | 4.9159676 |
| **DCLK1** | 4.5255 | -2.70968898075413 | 0.236 | 0.851 | 7.3599081 |
| **TEC** | 9.6241 | 1.20738249569413 | 0.707 | 0.113 | 1.5651795 |
| **HUS1** | 2.8303 | 1.23586737019313 | 1 | 0.98 | 4.6029308 |
| **MGST1** | 1.2213 | -1.42699733003443 | 0.943 | 0.998 | 1.9862548 |
| **PTGR1** | 1.2293 | 1.24038929712806 | 0.992 | 0.858 | 1.9993143 |
| **FADS1** | 1.6901 | -1.61959156831553 | 0.924 | 0.999 | 2.7486520 |
| **IDS** | 3.2575 | -1.62119568456612 | 0.255 | 0.892 | 5.2976887 |
| **SRC** | 7.2632 | -1.41979053249737 | 0.712 | 0.981 | 1.1812285 |
| **WWOX** | 7.7532 | -1.86198809083267 | 0.579 | 0.958 | 1.2609034 |
| **UACA** | 8.5663 | 1.59408878651699 | 0.99 | 0.895 | 1.3931434 |
| **SNX24** | 1.3593 | 1.23941336898245 | 0.988 | 0.847 | 2.2107718 |
| **TBL1X** | 1.7510 | -1.68716062278589 | 0.202 | 0.855 | 2.8477592 |
| **ABCB7** | 2.1603 | -1.2380507202199 | 0.886 | 0.998 | 3.5134334 |
| **SPRY2** | 3.3902 | -1.76948114980996 | 0.859 | 0.992 | 5.5135821 |
| **ESYT1** | 6.4481 | -1.25707892584297 | 0.969 | 0.999 | 1.0486618 |
| **KCNK6** | 7.2886 | 1.27270557931119 | 0.761 | 0.162 | 1.1853526 |
| **RFLNB** | 9.5000 | 1.89043676199309 | 0.968 | 0.608 | 1.5449999 |
| **REPIN1** | 1.6130 | -1.42567287029081 | 0.513 | 0.955 | 2.6233322 |
| **ICA1L** | 3.6944 | 1.21205047711499 | 0.722 | 0.139 | 6.0082460 |
| **CHST11** | 5.8301 | -1.84297489704078 | 0.671 | 0.963 | 9.4815134 |
| **ERICH5** | 7.8043 | 1.01778248366952 | 0.649 | 0.076 | 1.2692284 |
| **ZNF649** | 1.0933 | 1.27853839370465 | 0.999 | 0.904 | 1.7781212 |

| | | | | | |
|---|---|---|---|---|---|
| **CUTA** | 1.7423 | -1.05137638411977 | 0.999 | 1 | 2.8336082 |
| **MSLN** | 1.8198 | 0.98532765317728 | 0.504 | 0.014 | 2.9597021 |
| **MSI1** | 3.8845 | -1.75261955060216 | 0.189 | 0.846 | 6.3173997 |
| **WNT8A** | 9.6070 | -2.53491405029088 | 0.068 | 0.785 | 1.5623995 |
| **NRXN2** | 1.1106 | 1.0148154780026 | 0.632 | 0.068 | 1.8062827 |
| **PCCA** | 1.1667 | -1.39352616441138 | 0.937 | 0.998 | 1.8975516 |
| **LAMA5** | 1.7976 | -1.48357717575956 | 0.968 | 0.999 | 2.9234816 |
| **ALPP** | 2.3517 | 1.24257805697101 | 0.549 | 0.032 | 3.8245930 |
| **RAB34** | 2.7680 | -1.19185646909081 | 0.932 | 0.997 | 4.5017101 |
| **AUTS2** | 3.0754 | -1.99381633901455 | 0.465 | 0.94 | 5.0015473 |
| **ZNF185** | 3.4020 | -1.75877451195637 | 0.27 | 0.878 | 5.5326919 |
| **MAFA** | 4.3771 | 1.5295523938994 | 0.844 | 0.265 | 7.1185091 |
| **MX1** | 4.8321 | 1.51078335133571 | 0.496 | 0.013 | 7.8584828 |
| **TPGS2** | 5.6235 | -1.14053541812967 | 0.966 | 0.999 | 9.1455239 |
| **AGPAT2** | 7.5953 | 1.37485649996936 | 0.997 | 0.899 | 1.2352239 |
| **TTLL12** | 8.1071 | 0.967753529353089 | 1 | 0.993 | 1.3184615 |
| **IMPA2** | 9.8502 | 1.21748699828729 | 0.997 | 0.951 | 1.6019387 |
| **WASF1** | 1.6360 | -1.6062519989576 | 0.505 | 0.94 | 2.6607676 |
| **ZNF486** | 3.6886 | 1.65577660554688 | 0.902 | 0.469 | 5.9988436 |
| **PSME4** | 4.7322 | 0.942390956037336 | 1 | 0.998 | 7.6960694 |
| **RO60** | 5.8861 | -1.1496565897924 | 0.893 | 0.997 | 9.5726790 |
| **CASK** | 6.4878 | -1.25798214511411 | 0.928 | 0.999 | 1.0551241 |
| **SLC6A6** | 6.7757 | 1.40677575031999 | 0.996 | 0.863 | 1.1019373 |
| **ACOX1** | 8.7418 | 1.20476510935359 | 0.995 | 0.898 | 1.4216912 |
| **FLOT2** | 1.5064 | -1.23222504665268 | 0.943 | 0.998 | 2.4499819 |
| **ABHD12B** | 1.5673 | 1.06772808252832 | 0.638 | 0.069 | 2.5490113 |
| **PRR36** | 1.7275 | -1.80818572832352 | 0.195 | 0.842 | 2.8095066 |
| **SPATC1L** | 2.0889 | -1.47401006756198 | 0.045 | 0.776 | 3.3972820 |
| **ZNF711** | 4.8646 | -1.6663249451535 | 0.151 | 0.836 | 7.9114607 |
| **MIIP** | 6.9019 | 1.02363661817515 | 0.999 | 0.99 | 1.1224630 |
| **XBP1** | 9.1451 | 1.13268903406565 | 0.999 | 0.997 | 1.4872685 |
| **GALNT3** | 9.5894 | 0.87457901325672 | 0.468 | 0.005 | 1.5595333 |
| **RASIP1** | 1.0447 | 1.27226735811362 | 0.747 | 0.16 | 1.6990479 |
| **DBNDD1** | 1.3740 | -1.58207415568637 | 0.663 | 0.965 | 2.2345393 |

| | | | | | |
|---|---|---|---|---|---|
| **PIEZO2** | 1.5436 | -2.12497666017205 | 0.054 | 0.778 | 2.5104411 |
| **LMO4** | 1.9722 | 1.18480908247438 | 0.996 | 0.939 | 3.2074416 |
| **RCOR2** | 2.3782 | -1.83569439464271 | 0.243 | 0.86 | 3.8678118 |
| **S100A16** | 2.6803 | 1.90608851095771 | 0.823 | 0.26 | 4.3589765 |
| **DCBLD2** | 2.7163 | -1.30049302760605 | 0.829 | 0.992 | 4.4176300 |
| **ZNF362** | 3.8363 | -1.36910045522269 | 0.749 | 0.986 | 6.2390127 |
| **NRROS** | 4.8579 | 1.03408422549651 | 0.597 | 0.056 | 7.9004206 |
| **SALL1** | 6.1784 | -2.39315352540482 | 0.111 | 0.803 | 1.0048076 |
| **PHKA1** | 1.0242 | 1.69845307091519 | 0.987 | 0.871 | 1.6656728 |
| **FZD2** | 1.3313 | -2.5239135049865 | 0.078 | 0.782 | 2.1651653 |
| **FBP1** | 1.3666 | 1.49131402870358 | 0.665 | 0.102 | 2.2226036 |
| **SEC11A** | 1.9198 | -0.939734399970694 | 0.999 | 1 | 3.1222366 |
| **FXYD4** | 2.2862 | 1.08095692238952 | 0.597 | 0.056 | 3.7182051 |
| **TOMM40** | 2.7652 | 0.894655224177339 | 1 | 0.996 | 4.4971527 |
| **ZNF644** | 5.2492 | -0.934345893950848 | 0.992 | 1 | 8.5368575 |
| **GNB4** | 5.5845 | -1.50603742620252 | 0.638 | 0.972 | 9.0822282 |
| **LARS** | 8.0405 | 0.840879086053195 | 1 | 1 | 1.3076406 |
| **DBN1** | 1.3633 | -1.03356392247552 | 0.994 | 1 | 2.2171637 |
| **NT5DC2** | 1.6031 | -1.06593064078297 | 0.995 | 1 | 2.6071460 |
| **PREX2** | 2.0255 | -1.78783311774486 | 0.07 | 0.778 | 3.2941369 |
| **FLOT1** | 2.8187 | -1.1030139169505 | 0.941 | 0.997 | 4.5841403 |
| **HOXB2** | 3.0605 | 1.56243958800447 | 0.913 | 0.434 | 4.9773687 |
| **TP53I11** | 3.5529 | -1.72673646184932 | 0.159 | 0.822 | 5.7780964 |
| **HSBP1L1** | 4.0559 | 1.3087947905316 | 0.776 | 0.19 | 6.5961887 |
| **ZNF595** | 5.0069 | 1.550345964937 | 0.965 | 0.75 | 8.1428646 |
| **PLCG1** | 7.2174 | -1.12337661248586 | 0.964 | 0.999 | 1.1737706 |
| **DPF1** | 7.3678 | 1.6548470880335 | 0.915 | 0.49 | 1.1982297 |
| **BUD31** | 1.8475 | 0.929255580463161 | 1 | 0.997 | 3.0046101 |
| **FLNA** | 2.4594 | -1.06075937381377 | 1 | 1 | 3.9997535 |
| **HSP90AA1** | 4.0128 | 0.91400795572408 | 1 | 1 | 6.5260318 |
| **ANKRD18A** | 5.1140 | -1.72489212602935 | 0.056 | 0.769 | 8.3169449 |
| **TRIM6** | 6.4692 | 1.273756177746 | 0.788 | 0.203 | 1.0520996 |
| **FST** | 8.4365 | -3.27820982413769 | 0.177 | 0.828 | 1.3720350 |
| **MLLT11** | 9.6739 | -1.8585212026901 | 0.394 | 0.907 | 1.5732759 |

| | | | | | |
|---|---|---|---|---|---|
| UTRN | 1.1842 | -1.88693024778781 | 0.481 | 0.936 | 1.9259939 |
| MAF | 1.2675 | -2.44568291489675 | 0.19 | 0.837 | 2.0614590 |
| CACHD1 | 1.4785 | -1.7622819266816 | 0.541 | 0.958 | 2.4045632 |
| EML1 | 2.0804 | -1.44616430213342 | 0.773 | 0.979 | 3.3834914 |
| TAGLN | 2.8142 | -4.44120238530527 | 0.65 | 0.954 | 4.5767435 |
| ZNF880 | 3.8638 | 1.0930150828563 | 0.752 | 0.165 | 6.2838534 |
| SOX12 | 4.3632 | -1.81326232478665 | 0.521 | 0.94 | 7.0960097 |
| TOMM22 | 4.9774 | 0.955848110579947 | 1 | 0.99 | 8.0948484 |
| SYCP3 | 5.8533 | 1.46720471504753 | 0.646 | 0.1 | 9.5193338 |
| COX5B | 6.4047 | 0.838364799575714 | 1 | 0.998 | 1.0416054 |
| NRK | 7.1319 | -2.92093346675407 | 0.053 | 0.76 | 1.1598624 |
| NAXE | 1.0085 | -1.27646715028924 | 0.73 | 0.985 | 1.6402131 |
| MMP15 | 1.4634 | -1.5241885058275 | 0.848 | 0.996 | 2.3800118 |
| RTN4 | 2.0008 | -0.973113027103921 | 1 | 1 | 3.2539614 |
| ATG13 | 2.4312 | 0.917524558930909 | 1 | 0.992 | 3.9538651 |
| FES | 3.5319 | 1.20700744170486 | 0.872 | 0.325 | 5.7439934 |
| C1QBP | 6.1665 | 0.946246206961376 | 1 | 0.993 | 1.0028670 |
| SNX12 | 6.7074 | -1.10045154488069 | 0.95 | 1 | 1.0908292 |
| RPRM | 8.8658 | -2.00126087091117 | 0.32 | 0.868 | 1.4418521 |
| VPS37B | 1.0451 | 1.1377555790569 | 1 | 0.942 | 1.6997188 |
| PLAAT5 | 4.3312 | -2.1163197206335 | 0.283 | 0.846 | 7.0438645 |
| CHPT1 | 4.7407 | 1.09953324283581 | 0.997 | 0.944 | 7.7098881 |
| CREBL2 | 9.5401 | 1.2463272303801 | 0.991 | 0.894 | 1.5515088 |
| TSPAN4 | 1.0766 | -1.38951697859781 | 0.925 | 0.994 | 1.7509724 |
| NR5A2 | 2.0134 | 1.3268576234991 | 0.618 | 0.078 | 3.2744706 |
| TGS1 | 2.9277 | 0.99103799533727 | 1 | 0.987 | 4.7613195 |
| IDH2 | 3.2634 | -1.53425714685381 | 0.829 | 0.991 | 5.3073095 |
| AJUBA | 3.6412 | -1.74826056673393 | 0.664 | 0.959 | 5.9217955 |
| GALNT1 | 2.1689 | -1.24202244903828 | 0.888 | 0.996 | 3.5274061 |
| GARS | 2.2559 | 1.00558704183407 | 1 | 0.998 | 3.6687731 |
| EIF4EBP1 | 5.4936 | 1.17844831619022 | 1 | 0.996 | 8.9343840 |
| XPO7 | 7.1411 | 0.868791539999466 | 1 | 0.998 | 1.1613719 |
| QPRT | 8.7694 | -0.97593714674297 | 0.992 | 1 | 1.4261769 |
| ZNF608 | 1.8547 | -1.65834319055578 | 0.443 | 0.946 | 3.0163375 |

| | | | | | |
|---|---|---|---|---|---|
| **ZMYND8** | 3.0953 | -1.10717653804292 | 0.994 | 0.999 | 5.0339991 |
| **MYL6** | 6.4552 | -0.904906368430741 | 1 | 1 | 1.0498250 |
| **BARD1** | 6.8557 | 1.25826834205296 | 0.987 | 0.812 | 1.1149456 |
| **PTCH1** | 1.8442 | -1.75768472610367 | 0.339 | 0.882 | 2.9992914 |
| **KLK13** | 3.5074 | 1.58662422380671 | 0.919 | 0.492 | 5.7041043 |
| **DHCR24** | 4.1856 | -1.26297659607461 | 1 | 1 | 6.8070636 |
| **MGST3** | 4.5095 | -1.43633280884981 | 0.888 | 0.997 | 7.3339399 |
| **PBX4** | 8.2897 | 1.21364786874753 | 0.766 | 0.19 | 1.3481663 |
| **EIF1B** | 8.4308 | -0.992293056949417 | 0.988 | 1 | 1.3711128 |
| **TRAF5** | 8.7095 | -1.77623200766055 | 0.078 | 0.765 | 1.4164371 |
| **LSM14B** | 8.7460 | -0.932626267579917 | 0.994 | 1 | 1.4223728 |
| **HBM** | 1.9483 | 0.866934096171846 | 0.496 | 0.02 | 3.1685307 |
| **PKDCC** | 2.3105 | -3.73912625356819 | 0.078 | 0.767 | 3.7576780 |
| **JMJD1C** | 4.1898 | -1.25168385795861 | 0.824 | 0.992 | 6.8139051 |
| **LIMA1** | 6.7372 | 1.34255187842135 | 0.992 | 0.848 | 1.0956796 |
| **RIN2** | 2.2111 | -1.51203097826731 | 0.545 | 0.959 | 3.5959596 |
| **PQBP1** | 2.5442 | 0.950085881463199 | 1 | 0.998 | 4.1376427 |
| **CGAS** | 2.7574 | 1.07482907609716 | 0.513 | 0.028 | 4.4844808 |
| **PDK1** | 3.3929 | -1.86543035131746 | 0.627 | 0.954 | 5.5180273 |
| **TSN** | 4.0342 | -0.92548108176845 | 0.996 | 0.999 | 6.5608652 |
| **COL4A6** | 5.3265 | -1.86335929142166 | 0.019 | 0.726 | 8.6626227 |
| **CAST** | 8.3746 | -1.65914638204621 | 0.137 | 0.808 | 1.3619693 |
| **SLC9A1** | 2.2533 | 1.2054246739685 | 0.983 | 0.811 | 3.6646522 |
| **CTNNBIP1** | 2.2932 | -1.3477910800856 | 0.848 | 0.992 | 3.7294480 |
| **NKAIN4** | 3.3436 | -2.05330603192422 | 0.301 | 0.869 | 5.4377214 |
| **FAM171A2** | 4.0897 | -1.54443939103942 | 0.57 | 0.952 | 6.6511043 |
| **MPV17L2** | 5.1490 | 1.08122475121236 | 0.996 | 0.899 | 8.3738649 |
| **KLRG2** | 1.0696 | 1.03703386724194 | 1 | 0.926 | 1.7395369 |
| **ARHGAP31** | 1.1978 | 1.07362322763568 | 0.619 | 0.079 | 1.9480775 |
| **OOEP** | 1.7647 | 0.812541753804519 | 0.443 | 0.005 | 2.8700574 |
| **MXRA5** | 1.8606 | 1.87413535387974 | 0.832 | 0.319 | 3.0260186 |
| **SOD1** | 2.1870 | 0.8989561000274 | 1 | 1 | 3.5567385 |
| **ENO1** | 2.2725 | -0.806321660167916 | 1 | 1 | 3.6958588 |
| **BCL3** | 3.1288 | 1.46286602436946 | 0.929 | 0.483 | 5.0883768 |

| | | | | | |
|---|---|---|---|---|---|
| **CFLAR** | 3.4120 | 1.18742893507383 | 0.995 | 0.863 | 5.5489683 |
| **CHST8** | 5.4405 | 1.44959273788981 | 0.884 | 0.404 | 8.8479500 |
| **NMI** | 9.1421 | 1.29804978757311 | 0.948 | 0.522 | 1.4867874 |
| **RAP2B** | 2.1572 | 1.05752709109907 | 1 | 0.996 | 3.5083880 |
| **NOTCH2** | 3.1505 | -1.6370241562923 | 0.34 | 0.882 | 5.1237886 |
| **CXXC5** | 3.9115 | -1.25686063821359 | 0.907 | 0.994 | 6.3613085 |
| **ATXN7L3B** | 4.0571 | -1.09795185365391 | 0.956 | 0.999 | 6.5982210 |
| **HYOU1** | 7.0424 | 0.932732854848198 | 1 | 1 | 1.1453215 |
| **PARVB** | 9.6123 | 1.38251807460183 | 0.817 | 0.294 | 1.5632570 |
| **ANXA2** | 1.3581 | 0.871397366949528 | 1 | 1 | 2.2088180 |
| **TSTD1** | 6.4141 | 0.914304793756608 | 0.536 | 0.04 | 1.0431267 |
| **SOCS3** | 7.3535 | -1.66071833836237 | 0.53 | 0.95 | 1.1959006 |
| **B3GNT2** | 8.2957 | -1.52388376503335 | 0.888 | 0.989 | 1.3491345 |
| **MDFI** | 1.3275 | 1.39081417446917 | 0.991 | 0.823 | 2.1590002 |
| **NOCT** | 2.7224 | 1.19909486232797 | 0.995 | 0.863 | 4.4275697 |
| **DCAF5** | 3.0033 | -1.34708986701806 | 0.552 | 0.949 | 4.8843372 |
| **CBX5** | 4.6984 | -1.26170874281428 | 0.913 | 0.992 | 7.6411698 |
| **SV2A** | 7.9259 | -1.59120027212487 | 0.107 | 0.776 | 1.2890032 |
| **PTBP2** | 1.5878 | -1.45264846816671 | 0.614 | 0.957 | 2.5823489 |
| **PDE12** | 2.0934 | 1.13806609503672 | 0.952 | 0.559 | 3.4045516 |
| **ADAMTS19** | 2.8033 | -2.1090294076219 | 0.034 | 0.724 | 4.5590812 |
| **CHCHD3** | 3.4628 | 0.956044589795266 | 1 | 0.993 | 5.6316226 |
| **CUX1** | 3.8265 | -1.294024451477 | 0.618 | 0.962 | 6.2231099 |
| **UBE2G2** | 1.0312 | -1.23935760081255 | 0.708 | 0.985 | 1.6770765 |
| **MCC** | 1.2827 | -1.48819260760383 | 0.056 | 0.74 | 2.0860614 |
| **GCH1** | 1.5737 | 1.32793578359238 | 0.924 | 0.52 | 2.5594415 |
| **GHITM** | 3.7839 | 0.869744093607996 | 1 | 0.998 | 6.1538304 |
| **LIMCH1** | 4.4747 | -2.02892067126874 | 0.264 | 0.837 | 7.2773388 |
| **NXN** | 1.0656 | -1.02254511833849 | 0.982 | 0.999 | 1.7330385 |
| **STOML2** | 2.1243 | 0.807185390837159 | 1 | 0.999 | 3.4548907 |
| **UPF1** | 2.6833 | 0.851120227923667 | 1 | 0.997 | 4.3639504 |
| **PTK7** | 4.1577 | -1.32177202207669 | 0.933 | 0.989 | 6.7617122 |
| **CAMK2D** | 4.2874 | -1.61195169711728 | 0.168 | 0.801 | 6.9727132 |
| **ZNF600** | 1.4410 | 1.27921956588727 | 0.972 | 0.626 | 2.3435110 |

| | | | | | |
|---|---|---|---|---|---|
| **LRRC59** | 2.0923 | 0.839510782575781 | 1 | 1 | 3.4028586 |
| **DLL3** | 2.2873 | 1.02627914733844 | 0.827 | 0.265 | 3.7199712 |
| **PYCARD** | 3.0559 | 1.1360171342819 | 0.694 | 0.134 | 4.9699229 |
| **AIF1L** | 3.7318 | -1.25329416162006 | 0.983 | 0.999 | 6.0691712 |
| **ARFGEF1** | 4.9293 | 0.982923395485894 | 0.997 | 0.969 | 8.0165443 |
| **SLC35F6** | 5.6929 | 1.020211455692 | 1 | 0.99 | 9.2585154 |
| **CTH** | 6.8206 | 1.51828487016192 | 0.969 | 0.714 | 1.1092420 |
| **RALGAPA2** | 1.0065 | -1.38135451984315 | 0.621 | 0.963 | 1.6370271 |
| **EPHA2** | 1.1625 | -1.9765022895919 | 0.089 | 0.752 | 1.8906206 |
| **HMOX2** | 1.1922 | 0.994061070808334 | 0.999 | 0.959 | 1.9389391 |
| **PIK3R2** | 1.5320 | -1.04567325101034 | 0.997 | 1 | 2.4916078 |
| **G3BP2** | 2.4061 | -0.88502625643172 | 1 | 1 | 3.9131462 |
| **HRK** | 2.8007 | 1.40234723473007 | 0.968 | 0.63 | 4.5548715 |
| **SACS** | 2.8384 | -1.28332937206576 | 0.875 | 0.991 | 4.6160963 |
| **WIPF3** | 5.6856 | 1.28036489666309 | 0.773 | 0.233 | 9.2465697 |
| **ARHGAP35** | 9.8397 | -1.13286926922168 | 0.866 | 0.991 | 1.6002352 |
| **ATP5MC3** | 1.4964 | 0.85840729934585 | 1 | 0.998 | 2.4337399 |
| **DLGAP4** | 1.6333 | -1.38041956027511 | 0.895 | 0.994 | 2.6562464 |
| **CALD1** | 2.4802 | -1.65119464410642 | 0.995 | 1 | 4.0335718 |
| **DDAH2** | 2.6336 | -1.46119617017965 | 0.798 | 0.988 | 4.2830914 |
| **PPTC7** | 4.2290 | 1.12917059398923 | 0.975 | 0.809 | 6.8777648 |
| **TUSC3** | 5.0423 | -0.999992459000327 | 0.959 | 1 | 8.2003387 |
| **NIT2** | 5.5159 | 1.13318186439399 | 0.959 | 0.646 | 8.9705531 |
| **ID2** | 7.0744 | -1.92551130861963 | 0.31 | 0.855 | 1.1505133 |
| **P4HA1** | 7.7326 | -1.20482014749342 | 0.965 | 0.999 | 1.2575640 |
| **MFSD4B** | 2.2727 | 1.23491559245742 | 0.987 | 0.816 | 3.6961294 |
| **SYNDIG1** | 2.5018 | -1.5167277649687 | 0.048 | 0.721 | 4.0687121 |
| **LHFPL6** | 2.9570 | -1.82053751982897 | 0.124 | 0.769 | 4.8090421 |
| **AARS2** | 4.9982 | 1.21000176529349 | 1 | 0.917 | 8.1287338 |
| **GRSF1** | 7.3472 | 0.829987629904889 | 1 | 0.987 | 1.1948779 |
| **CCDC6** | 8.3800 | -1.04553787753667 | 0.96 | 0.998 | 1.3628556 |
| **OVOL2** | 1.3349 | 1.32808541657186 | 0.855 | 0.326 | 2.1711026 |
| **TNNT1** | 1.5661 | 1.27826045189705 | 0.999 | 0.885 | 2.5470118 |
| **PHACTR2** | 1.7598 | -1.47159701294636 | 0.42 | 0.897 | 2.8620961 |

| | | | | | |
|---|---|---|---|---|---|
| **NAP1L2** | 4.0233 | -1.40502847452831 | 0.129 | 0.778 | 6.5430928 |
| **TINCR** | 4.3233 | 1.08253880406747 | 0.725 | 0.158 | 7.0310563 |
| **WNT5A** | 4.5424 | -1.69208456214789 | 0.052 | 0.721 | 7.3874669 |
| **ASXL1** | 8.6441 | -1.51878018970261 | 0.601 | 0.945 | 1.4057991 |
| **TPM1** | 1.4455 | -1.26809309588777 | 1 | 1 | 2.3508534 |
| **PAX5** | 1.5865 | 0.990093921968886 | 0.55 | 0.054 | 2.5802333 |
| **MAGI1** | 3.3819 | -1.46861088249665 | 0.362 | 0.873 | 5.5000960 |
| **GNPNAT1** | 4.3164 | 1.24416161363599 | 0.991 | 0.884 | 7.0198734 |
| **HES1** | 4.9530 | -2.38367168914078 | 0.057 | 0.724 | 8.0550691 |
| **SAT1** | 5.9167 | -1.88388902612463 | 0.841 | 0.994 | 9.6223923 |
| **VAV2** | 7.8005 | 1.08030378632246 | 0.996 | 0.918 | 1.2686085 |
| **TIMM17A** | 7.8172 | 0.949653208185262 | 1 | 0.978 | 1.2713137 |
| **SRMS** | 1.0281 | 0.890025627479112 | 0.499 | 0.031 | 1.6720523 |
| **AIG1** | 1.1280 | -1.50398321139379 | 0.354 | 0.886 | 1.8345979 |
| **MYC** | 1.7091 | -1.73339402655133 | 0.665 | 0.966 | 2.7796144 |
| **FAM155A** | 1.9770 | 0.96604646068068 | 0.571 | 0.065 | 3.2152608 |
| **NCOA3** | 1.9983 | 1.15357613817156 | 0.996 | 0.969 | 3.2498356 |
| **TM7SF2** | 2.3045 | -1.37499248121408 | 0.858 | 0.991 | 3.7478741 |
| **IDH1** | 4.8734 | 1.03425425806532 | 1 | 0.975 | 7.9257719 |
| **PLPP3** | 1.7865 | -1.42953007627211 | 0.548 | 0.939 | 2.9054125 |
| **PRKAR2B** | 1.9920 | -1.54490312796024 | 0.496 | 0.917 | 3.2396085 |
| **AXL** | 2.2705 | -2.17124575953532 | 0.039 | 0.706 | 3.6925764 |
| **ACSL4** | 5.7192 | -1.32023677260376 | 0.683 | 0.987 | 9.3012237 |
| **TMEM98** | 5.7437 | -1.45703832573452 | 0.402 | 0.876 | 9.3410976 |
| **STK4** | 9.5455 | -1.09985364039134 | 0.845 | 0.987 | 1.5523897 |
| **CCNYL1** | 1.5229 | -1.19967991948999 | 0.634 | 0.972 | 2.4767763 |
| **SFRP2** | 2.9434 | -2.52990501535846 | 0.072 | 0.722 | 4.7869601 |
| **WNK3** | 2.9518 | -1.35910186839596 | 0.35 | 0.871 | 4.8005867 |
| **IGSF8** | 5.0715 | -1.53760473033866 | 0.627 | 0.94 | 8.2478046 |
| **ETNK1** | 6.0574 | -1.19702071895723 | 0.729 | 0.98 | 9.8511529 |
| **ST5** | 6.9053 | -1.272236599094 | 0.598 | 0.958 | 1.1230233 |
| **NEFL** | 7.3765 | 1.09436664871818 | 0.466 | 0.021 | 1.1996474 |
| **CBFA2T2** | 1.4201 | 1.19714040487839 | 1 | 0.972 | 2.3096282 |
| **SGK1** | 2.1058 | 1.50472010857958 | 0.995 | 0.817 | 3.4247108 |

| | | | | | |
|---|---|---|---|---|---|
| **S100A13** | 2.8511 | 1.21042613682144 | 1 | 0.98 | 4.6368895 |
| **NQO2** | 3.4233 | 0.99738269964125 | 1 | 0.969 | 5.5674536 |
| **TCF7L2** | 4.5779 | -1.50994487856833 | 0.447 | 0.915 | 7.4451099 |
| **GLUL** | 5.3024 | -1.04881979924891 | 0.992 | 1 | 8.6234285 |
| **SPRED1** | 1.1840 | -1.31827396587126 | 0.119 | 0.755 | 1.9256410 |
| **MAPK10** | 1.3552 | -2.12331269904301 | 0.04 | 0.706 | 2.2040416 |
| **VAT1L** | 2.0747 | -3.24931813114187 | 0.052 | 0.708 | 3.3741246 |
| **ARID3B** | 5.4114 | -0.809276149472238 | 0.999 | 1 | 8.8007042 |
| **WDHD1** | 6.1378 | 0.872244786481105 | 1 | 0.99 | 9.9819406 |
| **RTL8C** | 6.3694 | -1.32475491342154 | 0.425 | 0.9 | 1.0358666 |
| **DOCK9** | 6.8608 | 1.17562258564847 | 0.988 | 0.779 | 1.1157748 |
| **MAN1C1** | 7.6650 | 1.29923228856541 | 0.991 | 0.886 | 1.2465693 |
| **CRISPLD1** | 1.4039 | -1.56604086007985 | 0.523 | 0.933 | 2.2832435 |
| **SYT6** | 1.4490 | -1.53366291263594 | 0.092 | 0.736 | 2.3565660 |
| **SERPINE2** | 2.0732 | -1.42968010074823 | 0.46 | 0.91 | 3.3717210 |
| **SULF2** | 4.2783 | -1.68088955969542 | 0.726 | 0.97 | 6.9578679 |
| **NCOR2** | 6.3952 | -0.864825574263321 | 0.997 | 1 | 1.0400633 |
| **SMAD3** | 6.4965 | -1.37454243095807 | 0.615 | 0.953 | 1.0565411 |
| **GRPEL1** | 8.1971 | 0.941557775450625 | 0.999 | 0.972 | 1.3330992 |
| **TTC3** | 6.9667 | -1.3558132108228 | 0.893 | 0.992 | 1.1330099 |
| **VANGL2** | 1.1049 | -1.36400802931028 | 0.758 | 0.973 | 1.7969021 |
| **NSG1** | 2.1009 | 1.06455416087778 | 0.594 | 0.086 | 3.4167995 |
| **ORAI1** | 2.7531 | 1.01145431591434 | 0.997 | 0.936 | 4.4773795 |
| **SORD** | 4.1228 | -1.31550182572793 | 0.385 | 0.876 | 6.7049657 |
| **ZC4H2** | 4.2678 | -1.29010311821031 | 0.218 | 0.808 | 6.9407745 |
| **SLC25A44** | 4.4600 | 1.04479885916778 | 0.999 | 0.975 | 7.2534444 |
| **VAPB** | 8.3314 | -0.8410134450251 | 0.99 | 0.999 | 1.3549435 |
| **PSMC4** | 1.2343 | 0.959580609663776 | 1 | 0.998 | 2.0074730 |
| **STARD3NL** | 2.2226 | -1.174339849315 | 0.757 | 0.978 | 3.6146513 |
| **PGM2** | 2.8536 | 1.12081147213498 | 0.987 | 0.868 | 4.6409603 |
| **DLG3** | 3.5509 | 1.03594291104266 | 0.999 | 0.987 | 5.7749800 |
| **TRIM26** | 3.6264 | 1.06549057973943 | 0.997 | 0.928 | 5.8976518 |
| **ACAA2** | 5.1835 | 0.881450412296843 | 1 | 0.997 | 8.4299578 |
| **AKAP3** | 8.6641 | 0.822037679492269 | 0.496 | 0.037 | 1.4090433 |

| | | | | | |
|---|---|---|---|---|---|
| **RELL1** | 8.7124 | 1.20785922920227 | 0.966 | 0.753 | 1.4169049 |
| **ZIC2** | 1.0091 | -2.59293641971992 | 0.036 | 0.689 | 1.6412139 |
| **NCKAP1** | 1.5036 | -0.833672366011945 | 0.997 | 1 | 2.4453438 |
| **SNX8** | 1.5699 | 0.911620342858817 | 0.999 | 0.981 | 2.5532133 |
| **ARAP1** | 3.1708 | 1.00665393307085 | 0.994 | 0.972 | 5.1567168 |
| **CLIP1** | 3.8109 | 1.01302301896163 | 0.995 | 0.932 | 6.1978141 |
| **UBAC2** | 4.4124 | 0.865091160251286 | 0.999 | 0.987 | 7.1759023 |
| **TRIP6** | 4.8666 | -0.951013148840826 | 0.995 | 1 | 7.9146061 |
| **CERS6** | 5.4597 | -1.36931327695638 | 0.203 | 0.802 | 8.8792010 |
| **NR1H2** | 6.3139 | 0.982428691236334 | 0.995 | 0.925 | 1.0268370 |
| **CAPG** | 1.1154 | 1.01530260204037 | 0.999 | 0.936 | 1.8139879 |
| **PCDH18** | 1.1400 | -2.24934981282815 | 0.57 | 0.929 | 1.8540520 |
| **GALK1** | 1.6132 | 1.2060418989824 | 0.933 | 0.618 | 2.6236129 |
| **RBPMS** | 1.6702 | -0.864485152749676 | 0.992 | 0.999 | 2.7162705 |
| **LIPA** | 2.8961 | -1.36486027906449 | 0.274 | 0.831 | 4.7099801 |
| **TALDO1** | 4.6955 | 0.847939372848585 | 1 | 0.998 | 7.6363861 |
| **HES2** | 2.7113 | 0.965533488564151 | 0.556 | 0.068 | 4.4094634 |
| **GPC3** | 2.7705 | -1.91064791204556 | 0.51 | 0.942 | 4.5056873 |
| **SYK** | 6.7674 | 1.09273000508454 | 0.738 | 0.19 | 1.1005954 |
| **HSPH1** | 9.2961 | 0.994384138383283 | 1 | 0.999 | 1.5118383 |
| **UBASH3B** | 9.6234 | 1.153540626454 | 0.765 | 0.243 | 1.5650542 |
| **RBBP7** | 2.1688 | -1.00593629063167 | 0.937 | 0.998 | 3.5272386 |
| **HDHD5** | 2.4012 | 0.956623610486701 | 0.995 | 0.929 | 3.9051138 |
| **PAPSS1** | 3.4695 | -0.951097388067816 | 0.97 | 0.998 | 5.6424917 |
| **KLF17** | 3.6068 | 1.1804115086308 | 0.411 | 0.009 | 5.8657693 |
| **F2RL1** | 4.2582 | 1.14649518638099 | 0.999 | 0.956 | 6.9251616 |
| **DLG1** | 4.6714 | -1.09633637945428 | 0.851 | 0.986 | 7.5971582 |
| **FIGNL2** | 1.0734 | -1.96133685623309 | 0.133 | 0.743 | 1.7458008 |
| **HYLS1** | 1.1829 | 1.22723435344638 | 0.674 | 0.149 | 1.9237527 |
| **CAPN13** | 1.2064 | -1.26196481845682 | 0.022 | 0.667 | 1.9620246 |
| **SPINT1** | 2.6817 | 1.0510007003032 | 0.999 | 0.853 | 4.3613646 |
| **ENO2** | 3.3555 | 1.16163231017543 | 0.999 | 0.974 | 5.4572095 |
| **SLC23A2** | 3.4946 | 1.15205762290617 | 0.988 | 0.912 | 5.6833063 |
| **FBXO36** | 5.5340 | 1.06510408243012 | 0.997 | 0.873 | 9.0000473 |

| | | | | | |
|---|---|---|---|---|---|
| **PLXNB2** | 6.4326 | 1.02988141932987 | 0.997 | 0.945 | 1.0461391 |
| **JADE2** | 7.4355 | 1.3958118841172 | 0.975 | 0.794 | 1.2092414 |
| **CCR8** | 1.0075 | 1.12969574618277 | 0.585 | 0.08 | 1.6386585 |
| **CMTM4** | 1.0476 | 1.02583779104725 | 0.991 | 0.956 | 1.7037391 |
| **LPCAT4** | 1.3580 | 1.15249903536842 | 0.978 | 0.813 | 2.2085170 |
| **NT5C2** | 1.5295 | -1.18800847376071 | 0.848 | 0.987 | 2.4875363 |
| **GOT1** | 1.5702 | 1.11816618838794 | 0.992 | 0.894 | 2.5537695 |
| **DDT** | 2.3725 | 0.81114762308935 | 1 | 0.99 | 3.8585121 |
| **C7orf50** | 2.4738 | -0.93535213201737 | 0.999 | 1 | 4.0232947 |
| **PRRX2** | 2.8306 | 1.20168271723023 | 0.868 | 0.358 | 4.6034502 |
| **PJA1** | 4.9883 | -1.24832984351165 | 0.693 | 0.985 | 8.1125484 |
| **NRM** | 5.7906 | -1.33259836019343 | 0.342 | 0.848 | 9.4172992 |
| **TOM1L1** | 9.3298 | -1.11482375695056 | 0.678 | 0.966 | 1.5173168 |
| **INA** | 9.6535 | 1.20913376686233 | 0.978 | 0.771 | 1.5699543 |
| **TPP1** | 9.7508 | -1.23099736251976 | 0.912 | 0.993 | 1.5857744 |
| **EVL** | 1.1245 | -1.16937880071142 | 0.928 | 0.996 | 1.8288340 |
| **PRDM14** | 1.1245 | 1.29764076581908 | 0.995 | 0.889 | 1.8288412 |
| **NFKBIA** | 1.4820 | 1.08286776704435 | 0.995 | 0.956 | 2.4103276 |
| **MARCH8** | 1.6272 | 1.1546597415211 | 0.969 | 0.772 | 2.6463474 |
| **OSBPL9** | 1.9100 | -0.88997634816716 | 0.979 | 0.998 | 3.1062587 |
| **NRIP1** | 4.0661 | -1.63640592080713 | 0.391 | 0.86 | 6.6127837 |
| **LDLRAP1** | 4.4556 | 1.0143001577964 | 0.996 | 0.914 | 7.2461856 |
| **SMYD2** | 9.0096 | -1.26832632451589 | 0.826 | 0.979 | 1.4652441 |
| **TRABD** | 1.1983 | 0.914304914489727 | 0.994 | 0.936 | 1.9488345 |
| **SERINC5** | 1.9492 | 0.981335337212219 | 0.999 | 0.976 | 3.1700994 |
| **ST14** | 2.1117 | 1.13450084849055 | 0.685 | 0.162 | 3.4343790 |
| **ALCAM** | 1.2081 | -1.70857568972801 | 0.075 | 0.7 | 1.9648426 |
| **DHRS3** | 1.2082 | -1.55777674503193 | 0.275 | 0.812 | 1.9649719 |
| **NIPA2** | 1.2933 | 0.897298112669812 | 0.997 | 0.96 | 2.1034409 |
| **PAM** | 1.5007 | -1.65070945077571 | 0.106 | 0.723 | 2.4407219 |
| **BTK** | 1.5578 | 0.961874701990991 | 0.503 | 0.048 | 2.5335145 |
| **HSPA12B** | 1.6569 | 1.05436144330135 | 0.654 | 0.137 | 2.6947162 |
| **VDAC1** | 2.2822 | 0.941901554429104 | 0.999 | 0.976 | 3.7115471 |
| **TIMP2** | 3.1368 | -1.64425670122425 | 0.807 | 0.977 | 5.1015098 |

| | | | | | |
|---|---|---|---|---|---|
| **CDH2** | 3.4727 | -2.89054567207079 | 0.097 | 0.711 | 5.6477264 |
| **KLF12** | 5.2620 | -1.34253723631656 | 0.163 | 0.762 | 8.5576067 |
| **BMP4** | 5.6223 | -1.76287579223757 | 0.37 | 0.854 | 9.1435693 |
| **SEPHS1** | 5.7789 | -1.24989024109727 | 0.968 | 0.995 | 9.3982371 |
| **KLC3** | 1.1375 | 1.06639233404866 | 0.749 | 0.215 | 1.8500397 |
| **UBE2R2** | 3.0233 | -0.904327132789022 | 0.99 | 1 | 4.9169312 |
| **BLCAP** | 3.9120 | -1.03161568944038 | 0.864 | 0.99 | 6.3621512 |
| **IGDCC4** | 4.7561 | -1.36713823031132 | 0.031 | 0.664 | 7.7349965 |
| **PTOV1** | 6.0714 | -0.881913541859439 | 0.984 | 0.999 | 9.8740157 |
| **CA11** | 8.7843 | -1.71201377766489 | 0.063 | 0.687 | 1.4286012 |
| **FGFR4** | 1.1445 | 1.17282742848208 | 0.986 | 0.819 | 1.8613222 |
| **ZNF593** | 1.4448 | 0.855608965355863 | 0.999 | 0.976 | 2.3497153 |
| **NPW** | 1.5235 | 0.862258578086188 | 0.494 | 0.044 | 2.4776998 |
| **CBX3** | 1.8926 | 1.17670813687275 | 0.974 | 0.834 | 3.0780604 |
| **MTA2** | 2.7053 | -0.995652980131875 | 0.955 | 0.998 | 4.3997755 |
| **FUCA2** | 2.7574 | -1.32047832858866 | 0.789 | 0.965 | 4.4843783 |
| **DBNDD2** | 3.5447 | -1.30290995709621 | 0.391 | 0.862 | 5.7648411 |
| **CDA** | 1.2761 | 1.22281225402295 | 0.987 | 0.794 | 2.0754627 |
| **MBOAT7** | 2.0354 | -0.938732372140742 | 0.955 | 0.999 | 3.3103212 |
| **TGFBR1** | 3.5083 | -1.10532673763083 | 0.977 | 0.999 | 5.7056078 |
| **LSS** | 4.2370 | -1.0663095765759 | 0.952 | 0.997 | 6.8906748 |
| **CCNE1** | 7.5538 | 1.29477381527666 | 0.979 | 0.728 | 1.2284902 |
| **GTF2A2** | 8.6037 | 1.03625669139127 | 0.996 | 0.918 | 1.3992302 |
| **BNIP3L** | 1.6269 | -1.0285019144456 | 0.893 | 0.989 | 2.6459593 |
| **CIAO2B** | 2.4205 | 0.887568982812396 | 0.999 | 0.981 | 3.9365555 |
| **DZIP1** | 2.6113 | -1.22928736110879 | 0.747 | 0.976 | 4.2468312 |
| **YAF2** | 6.3635 | -1.36385568357792 | 0.159 | 0.757 | 1.0349051 |
| **WDFY2** | 8.5629 | -1.26702291468497 | 0.69 | 0.962 | 1.3926006 |
| **CKS1B** | 1.0708 | 1.20592416183379 | 0.995 | 0.932 | 1.7414859 |
| **RAB42** | 1.2257 | 0.865649647973902 | 0.537 | 0.068 | 1.9934939 |
| **GSTO2** | 3.4499 | 0.838490639339125 | 0.547 | 0.072 | 5.6106940 |
| **FAM118B** | 9.1610 | -1.05150906986233 | 0.696 | 0.962 | 1.4898573 |
| **SDF2L1** | 9.7466 | 1.1447245297938 | 1 | 0.976 | 1.5850973 |
| **SEC61G** | 1.2007 | 0.875893308502072 | 1 | 0.999 | 1.9528299 |

| | | | | | |
|---|---|---|---|---|---|
| **MFSD10** | 1.7860 | -1.1419376344901 | 0.683 | 0.963 | 2.9046187 |
| **3,00 DKK** | 2.1229 | -1.65313400336274 | 0.377 | 0.874 | 3.4525963 |
| **PITPNC1** | 2.8608 | 0.936584236345121 | 0.997 | 0.946 | 4.6526780 |
| **SNX4** | 3.9213 | 0.915577450769599 | 0.995 | 0.987 | 6.3772861 |
| **ACOT9** | 3.0269 | -1.20678206517375 | 0.647 | 0.952 | 4.9227800 |
| **MT2A** | 4.1136 | 1.06996437873734 | 0.999 | 0.977 | 6.6900020 |
| **CCND2** | 4.5814 | -2.47240525480199 | 0.102 | 0.703 | 7.4508260 |
| **PDLIM1** | 9.6075 | 0.995798068367986 | 1 | 0.963 | 1.5624721 |
| **WDR5** | 1.1291 | 0.9861051468383 | 0.99 | 0.865 | 1.8363541 |
| **TUNAR** | 2.1105 | -1.5095887898341 | 0.119 | 0.715 | 3.4323960 |
| **AKAP12** | 6.1697 | 0.891726282835779 | 1 | 0.999 | 1.0033841 |
| **CDIP1** | 7.7988 | -1.10327257423491 | 0.795 | 0.978 | 1.2683264 |
| **LRRN3** | 7.8107 | -1.47706015122442 | 0.022 | 0.641 | 1.2702654 |
| **SALL3** | 1.5241 | -1.59920875057502 | 0.689 | 0.93 | 2.4787524 |
| **DDIT4L** | 1.5262 | 0.824352419223363 | 0.463 | 0.036 | 2.4821937 |
| **PRDX3** | 2.2216 | 0.90858853797169 | 0.999 | 0.954 | 3.6131074 |
| **EXTL2** | 3.6623 | -1.09217289345896 | 0.685 | 0.955 | 5.9560779 |
| **KIAA1324L** | 4.4805 | -1.1611260340223 | 0.455 | 0.895 | 7.2867075 |
| **ARHGEF40** | 8.3818 | -1.64616551063578 | 0.129 | 0.72 | 1.3631464 |
| **PLCB4** | 1.0355 | -1.27688227441101 | 0.083 | 0.69 | 1.6841634 |
| **SLC25A33** | 1.3335 | 1.13728344975959 | 0.965 | 0.76 | 2.1686873 |
| **ADM** | 1.4920 | -2.24668312111873 | 0.195 | 0.751 | 2.4265891 |
| **GJA1** | 1.6471 | -1.05841949585013 | 1 | 1 | 2.6787373 |
| **ACO2** | 2.3325 | 0.832034176495924 | 1 | 0.986 | 3.7933540 |
| **PBX2** | 2.4828 | -0.891717833379175 | 0.99 | 0.998 | 4.0378036 |
| **CD24** | 3.1072 | -1.85648059997796 | 0.884 | 0.982 | 5.0533450 |
| **PCDHGB7** | 4.3691 | -1.47850464656093 | 0.194 | 0.758 | 7.1056241 |
| **IRX1** | 6.4763 | -2.03084020285162 | 0.242 | 0.797 | 1.0532561 |
| **CXCL12** | 1.0185 | -1.99735094708614 | 0.866 | 0.927 | 1.6564369 |
| **NAV2** | 1.6678 | -1.45955586178142 | 0.85 | 0.971 | 2.7123846 |
| **ZSWIM6** | 2.7037 | -1.32761145281232 | 0.112 | 0.706 | 4.3971885 |
| **C19orf85** | 3.3294 | 1.49086100365207 | 0.713 | 0.202 | 5.4146596 |
| **YARS** | 3.5239 | 0.837998186943308 | 0.999 | 0.976 | 5.7310313 |
| **MSRB2** | 9.6781 | -1.13436841665732 | 0.603 | 0.948 | 1.5739494 |

| | | | | | |
|---|---|---|---|---|---|
| **COX7B** | 1.7584 | 1.16459191607274 | 0.953 | 0.748 | 2.8597425 |
| **AP1S1** | 7.1165 | 1.09989547073167 | 0.999 | 0.975 | 1.1573605 |
| **LRATD2** | 7.8544 | -1.58862210420412 | 0.862 | 0.981 | 1.2773688 |
| **HCN2** | 8.0485 | 1.06857114593024 | 0.966 | 0.673 | 1.3089342 |
| **DUSP10** | 1.8097 | -1.4126298971666 | 0.208 | 0.772 | 2.9431795 |
| **NSD2** | 3.4123 | -1.32132948533769 | 0.289 | 0.803 | 5.5494522 |
| **SNN** | 3.6591 | -1.12458669460784 | 0.63 | 0.942 | 5.9508471 |
| **PREB** | 3.6621 | 0.838703312112291 | 0.999 | 0.972 | 5.9557864 |
| **LAD1** | 5.4864 | 0.998724906763347 | 0.55 | 0.085 | 8.9226484 |
| **ARHGDIG** | 6.1751 | 0.929721303989404 | 0.607 | 0.118 | 1.0042604 |
| **ANTXR1** | 1.0157 | -1.33703073985316 | 0.158 | 0.728 | 1.6518555 |
| **DENR** | 1.1055 | 0.830687731431058 | 1 | 0.975 | 1.7979930 |
| **HUNK** | 1.1573 | -1.34864913770557 | 0.123 | 0.707 | 1.8821770 |
| **TMEM45A** | 2.0798 | -1.32053676895272 | 0.088 | 0.683 | 3.3825066 |
| **KIF1B** | 2.1046 | -0.813516829992263 | 0.979 | 1 | 3.4227819 |
| **HEBP2** | 3.0283 | 0.929883820377971 | 1 | 0.988 | 4.9249869 |
| **NUAK1** | 3.1115 | -1.31244341339378 | 0.34 | 0.838 | 5.0602343 |
| **TOR3A** | 3.5472 | 0.949497641627846 | 0.99 | 0.876 | 5.7688501 |
| **PRDM1** | 1.0787 | 1.17585551247379 | 0.898 | 0.464 | 1.7544504 |
| **CDHR1** | 1.4281 | 1.43490514633281 | 0.959 | 0.682 | 2.3226605 |
| **NINL** | 1.8282 | -1.16239474031458 | 0.672 | 0.956 | 2.9733499 |
| **CDC73** | 2.1103 | -0.849490030198136 | 0.959 | 0.998 | 3.4320159 |
| **CNTRL** | 2.3831 | -1.26760093249801 | 0.26 | 0.801 | 3.8756997 |
| **DHRSX** | 2.8505 | 1.50743951926747 | 0.942 | 0.834 | 4.6359095 |
| **TMEM97** | 9.9179 | -0.990133775965184 | 0.99 | 0.999 | 1.6129525 |
| **MCRIP2** | 1.3152 | 1.03084687086653 | 0.946 | 0.662 | 2.1390498 |
| **PTCD1** | 2.3256 | 0.952729647818203 | 0.988 | 0.904 | 3.7822275 |
| **IGLON5** | 2.3629 | -1.26534735124068 | 0.129 | 0.711 | 3.8428741 |
| **EXT1** | 3.5341 | -1.2117995784281 | 0.784 | 0.98 | 5.7475969 |
| **RGS10** | 4.5390 | 1.08742073266015 | 0.875 | 0.448 | 7.3818331 |
| **ACVR2B** | 6.4174 | -0.850073887440714 | 0.994 | 1 | 1.0436669 |
| **TUBB2B** | 1.6596 | -1.08647968971857 | 0.999 | 1 | 2.6991009 |
| **DDX49** | 2.0715 | 0.860659203876063 | 0.997 | 0.953 | 3.3689565 |
| **CNN3** | 3.8285 | -0.8256364293562 | 1 | 1 | 6.2264114 |

| | | | | | |
|---|---|---|---|---|---|
| **TOX3** | 4.0439 | -1.35836256956776 | 0.076 | 0.668 | 6.5766192 |
| **SLC16A1** | 6.1699 | -1.0073495632206 | 1 | 1 | 1.0034110 |
| **NSD1** | 1.8369 | 0.886281919403653 | 0.999 | 0.967 | 2.9873803 |
| **PLEKHG1** | 2.1279 | 0.887769510995728 | 0.614 | 0.121 | 3.4606482 |
| **SNX3** | 2.8189 | 0.978631568677461 | 0.988 | 0.828 | 4.5844502 |
| **ATP13A2** | 2.9350 | 0.907777140968686 | 0.994 | 0.933 | 4.7732549 |
| **PCOLCE2** | 3.3136 | -1.35472439705556 | 0.494 | 0.885 | 5.3890330 |
| **MAPK13** | 4.4140 | 1.11225461515008 | 0.762 | 0.277 | 7.1785664 |
| **NEDD4L** | 4.8535 | -1.02135985259114 | 0.695 | 0.979 | 7.8933342 |
| **TRAPPC1** | 6.5855 | -0.817877156831547 | 0.992 | 1 | 1.0710148 |
| **TNPO2** | 9.8809 | -0.914928149009853 | 0.944 | 0.997 | 1.6069322 |
| **TBC1D16** | 1.0006 | -0.942319042408289 | 0.937 | 0.996 | 1.6273077 |
| **KRTCAP3** | 1.4761 | 1.15902232682254 | 0.8 | 0.313 | 2.4007199 |
| **MEGF8** | 2.0698 | -1.16316546672591 | 0.452 | 0.872 | 3.3661712 |
| **TMPO** | 2.1643 | 0.820107303452425 | 1 | 0.99 | 3.5198690 |
| **B4GALT5** | 2.3764 | -0.864129877854048 | 0.979 | 0.999 | 3.8648891 |
| **USP9X** | 2.8843 | -0.878332549116744 | 1 | 1 | 4.6908671 |
| **S100A10** | 4.0767 | 0.862035890038908 | 0.999 | 0.992 | 6.6300018 |
| **EIF2AK2** | 5.0217 | -1.03703680333252 | 0.593 | 0.939 | 8.1669086 |
| **EPB41L4B** | 5.9970 | 1.18993738284056 | 0.938 | 0.643 | 9.7529488 |
| **CLEC4D** | 7.2888 | -2.01458148038062 | 0.136 | 0.692 | 1.1853937 |
| **ZNF398** | 1.1303 | 1.01072750671925 | 0.974 | 0.834 | 1.8382177 |
| **TMLHE** | 1.4098 | -1.15189107861205 | 0.275 | 0.819 | 2.2928366 |
| **CBLC** | 1.9614 | 0.852247417119684 | 0.584 | 0.105 | 3.1898756 |
| **ZNF878** | 2.5019 | 1.11945634666632 | 0.743 | 0.252 | 4.0689187 |
| **MMP24OS** | 4.4685 | -1.11259022553145 | 0.72 | 0.968 | 7.2672642 |
| **HK1** | 5.7236 | -0.972323183213513 | 0.947 | 0.999 | 9.3083753 |
| **SCD** | 6.3444 | -0.931557537654691 | 1 | 1 | 1.0318043 |
| **MYO5A** | 7.1045 | 1.04374588772401 | 0.981 | 0.846 | 1.1554167 |
| **AXIN2** | 8.2702 | -1.23027055882662 | 0.726 | 0.965 | 1.3449867 |
| **MPP6** | 1.8715 | -1.06625462919838 | 0.965 | 0.995 | 3.0436481 |
| **NKX1-2** | 2.1279 | -2.16830924619057 | 0.654 | 0.876 | 3.4606928 |
| **MTFR1** | 2.2927 | 0.897625763426457 | 0.977 | 0.868 | 3.7287294 |
| **UBQLN1** | 4.0508 | 0.806587735099409 | 0.996 | 0.959 | 6.5878682 |

| | | | | | |
|---|---|---|---|---|---|
| **SNX1** | 8.4523 | -1.08383274380587 | 0.805 | 0.979 | 1.3746064 |
| **WASF3** | 1.3969 | -1.24485253704218 | 0.743 | 0.963 | 2.2718585 |
| **ST6GALNAC** | 1.4684 | -1.43193911371103 | 0.05 | 0.632 | 2.3881740 |
| **PPFIBP1** | 2.1643 | -1.28321228777142 | 0.243 | 0.781 | 3.5198694 |
| **OXCT1** | 2.1769 | -1.13402040177336 | 0.459 | 0.891 | 3.5403575 |
| **VASP** | 2.3376 | -1.01439068907416 | 0.999 | 0.999 | 3.8017088 |
| **TNIP2** | 3.4931 | 0.928905803714727 | 0.994 | 0.945 | 5.6809901 |
| **PROM1** | 4.4734 | 1.14968697627083 | 0.992 | 0.858 | 7.2751112 |
| **WDR35** | 8.5689 | -1.21193967749059 | 0.171 | 0.731 | 1.3935662 |
| **CAP2** | 8.7210 | -1.2242221588319 | 0.541 | 0.908 | 1.4182986 |
| **ADA** | 9.9532 | 1.10838346532525 | 0.941 | 0.617 | 1.6186902 |
| **PPT1** | 1.0424 | -0.946698734846137 | 0.977 | 0.996 | 1.6953828 |
| **ETFA** | 1.7308 | -1.11214394339928 | 0.762 | 0.971 | 2.8149163 |
| **SENP2** | 3.4021 | 0.875424879521646 | 1 | 0.996 | 5.5329139 |
| **TNKS1BP1** | 8.2139 | -0.999125506843423 | 0.828 | 0.979 | 1.3358324 |
| **EXD2** | 1.6652 | -1.12377903186242 | 0.447 | 0.874 | 2.7081483 |
| **KLF3** | 1.6920 | 1.1460558666947 | 0.968 | 0.705 | 2.7517015 |
| **FAM102B** | 1.9111 | 1.12624606154498 | 0.977 | 0.837 | 3.1081519 |
| **BAZ2B** | 3.6683 | -1.10860709719296 | 0.578 | 0.922 | 5.9657682 |
| **SKI** | 8.0621 | -0.832784675044716 | 0.984 | 0.999 | 1.3111431 |
| **FGD1** | 1.0161 | -0.987629225325922 | 0.78 | 0.987 | 1.6525118 |
| **FGFR2** | 1.0630 | -1.04546840800911 | 0.974 | 0.995 | 1.7287632 |
| **MMP24** | 1.9092 | -1.38913805225673 | 0.829 | 0.953 | 3.1050341 |
| **ZC3HAV1** | 2.4201 | 0.8482261800126 | 1 | 0.957 | 3.9358236 |
| **CD276** | 2.5182 | -1.12761249364933 | 0.975 | 0.999 | 4.0954884 |
| **AMER1** | 2.7152 | -1.1694602905216 | 0.362 | 0.834 | 4.4157606 |
| **SLC16A3** | 4.9373 | 1.15073847636438 | 0.891 | 0.481 | 8.0296699 |
| **DEPTOR** | 9.6928 | 1.06286592450715 | 0.991 | 0.802 | 1.5763467 |
| **DMBX1** | 9.7169 | 0.939800600223231 | 0.811 | 0.285 | 1.5802739 |
| **MTOR** | 1.0874 | 0.817131412605147 | 0.999 | 0.954 | 1.7685129 |
| **KLHDC10** | 1.1180 | -1.09240143938806 | 0.231 | 0.774 | 1.8183441 |
| **SYT2** | 1.3784 | 0.863263663208261 | 0.516 | 0.078 | 2.2417831 |
| **ATP8B2** | 2.1249 | 0.86184319970193 | 0.995 | 0.959 | 3.4557962 |
| **TRIM5** | 2.2326 | -1.1861375910297 | 0.425 | 0.867 | 3.6309652 |

| | | | | | |
|---|---|---|---|---|---|
| **IL6R** | 2.5429 | 1.06494329513138 | 0.827 | 0.368 | 4.1356765 |
| **FAM20C** | 5.9522 | -1.28806893873593 | 0.457 | 0.88 | 9.6800811 |
| **FKBP8** | 6.2651 | -0.834737632612591 | 0.951 | 0.995 | 1.0189038 |
| **FEZ2** | 7.2333 | 0.969405502117031 | 0.994 | 0.952 | 1.1763584 |
| **FBRSL1** | 7.2459 | 0.931542117589077 | 0.988 | 0.901 | 1.1784108 |
| **ZCCHC24** | 8.5677 | -1.37938726155497 | 0.187 | 0.718 | 1.3933701 |
| **TSKU** | 1.1389 | -0.950947555395141 | 0.897 | 0.992 | 1.8522509 |
| **AGPAT3** | 1.6733 | 0.873707025660845 | 0.625 | 0.136 | 2.7213917 |
| **SHMT2** | 1.6817 | 1.02367967358434 | 0.944 | 0.745 | 2.7350297 |
| **ZNF618** | 2.2957 | -1.06174447046774 | 0.689 | 0.945 | 3.7335652 |
| **TBC1D4** | 2.3990 | -1.31682648996145 | 0.213 | 0.739 | 3.9016414 |
| **AIFM2** | 3.3702 | 0.979406411179234 | 0.704 | 0.216 | 5.4810996 |
| **CCDC102A** | 3.6466 | -1.1431249420986 | 0.407 | 0.855 | 5.9305321 |
| **INF2** | 3.7095 | -0.944508485411745 | 0.984 | 0.999 | 6.0328044 |
| **KLHL15** | 4.0584 | -1.11378442293923 | 0.284 | 0.809 | 6.6002042 |
| **LYPD1** | 4.1496 | -2.13323846556059 | 0.087 | 0.65 | 6.7486197 |
| **USP51** | 7.7945 | -1.15100267727135 | 0.196 | 0.741 | 1.2676221 |
| **ALAD** | 9.2648 | 0.967871921577371 | 0.964 | 0.807 | 1.5067434 |
| **EML4** | 1.4924 | -0.931869581346118 | 0.955 | 0.997 | 2.4271886 |
| **RPP40** | 1.7506 | 0.942201692016052 | 0.946 | 0.728 | 2.8471411 |
| **TCEAL4** | 1.7642 | -0.867732970918515 | 0.924 | 0.998 | 2.8691511 |
| **NUDT8** | 4.0209 | 0.868976974578507 | 0.559 | 0.103 | 6.5392068 |
| **SLC25A22** | 4.9730 | 0.928910399167423 | 0.986 | 0.869 | 8.0877410 |
| **MAP4K1** | 5.1208 | 1.0588403290269 | 0.752 | 0.268 | 8.3281180 |
| **SOGA1** | 6.7433 | -0.987941661725066 | 0.866 | 0.991 | 1.0966785 |
| **ABCA1** | 1.2186 | 0.889794187138752 | 0.997 | 0.946 | 1.9818834 |
| **TNRC18** | 1.5996 | 0.820824952333547 | 0.997 | 0.971 | 2.6015398 |
| **GSR** | 1.6343 | -0.895465208036142 | 0.908 | 0.993 | 2.6579670 |
| **UCK2** | 2.5630 | -1.0067652880873 | 0.888 | 0.992 | 4.1682714 |
| **GDPD1** | 2.9719 | 1.05158422731015 | 0.646 | 0.169 | 4.8332930 |
| **IPMK** | 3.2587 | 0.920086909074304 | 0.986 | 0.845 | 5.2997608 |
| **IDH3A** | 3.5274 | 0.945946104137918 | 0.987 | 0.822 | 5.7367278 |
| **UPRT** | 3.9626 | -1.10465175416255 | 0.304 | 0.816 | 6.4443960 |
| **CYBA** | 6.3134 | 0.881325216097748 | 0.999 | 0.982 | 1.0267512 |

| | | | | | |
|---|---|---|---|---|---|
| **MACF1** | 6.6222 | -0.90636339112946 | 0.955 | 0.994 | 1.0769834 |
| **MIPOL1** | 6.7384 | -1.126283396249 | 0.329 | 0.803 | 1.0958741 |
| **APOC1** | 7.8581 | 0.993054553939233 | 0.961 | 0.715 | 1.2779704 |
| **SLC27A4** | 1.8883 | 0.823412513455802 | 0.996 | 0.962 | 3.0710168 |
| **THBS3** | 2.2251 | -1.42056803982801 | 0.081 | 0.643 | 3.6187849 |
| **TGFB3** | 3.0343 | 0.920300284912122 | 0.606 | 0.131 | 4.9347559 |
| **EPHA7** | 3.5277 | -1.57406610580959 | 0.722 | 0.952 | 5.7371550 |
| **LIMK1** | 3.8009 | -0.894887192105748 | 0.924 | 0.995 | 6.1815111 |
| **PNKD** | 5.0524 | 0.857784846930742 | 0.992 | 0.921 | 8.2168223 |
| **CAPN1** | 7.3177 | -0.853958464793667 | 0.999 | 0.999 | 1.1900885 |
| **DIXDC1** | 9.5859 | -1.23885676081245 | 0.229 | 0.75 | 1.5589624 |
| **TP53I3** | 1.2715 | -1.11808606992269 | 0.534 | 0.901 | 2.0679169 |
| **TIGAR** | 1.4778 | 0.879026169255332 | 1 | 0.986 | 2.4034349 |
| **CRABP1** | 1.6647 | -3.38253883268188 | 0.376 | 0.815 | 2.7073326 |
| **DDAH1** | 1.9323 | -1.44280174590377 | 0.314 | 0.809 | 3.1426285 |
| **POLR3K** | 3.2579 | 0.872570744045543 | 0.997 | 0.964 | 5.2983553 |
| **CASP2** | 5.9938 | -1.13237225555946 | 0.699 | 0.941 | 9.7478617 |
| **TMEM37** | 6.2776 | 1.13769378425449 | 0.987 | 0.822 | 1.0209377 |
| **ATF4** | 1.4032 | 0.81512358112129 | 1 | 0.992 | 2.2821256 |
| **SEPTIN11** | 1.5608 | -0.926702012550077 | 0.986 | 1 | 2.5384555 |
| **PHLDA2** | 1.6977 | 1.18352306682366 | 0.922 | 0.607 | 2.7610541 |
| **OLFM1** | 1.9384 | -1.4238226780875 | 0.353 | 0.81 | 3.1525456 |
| **CILP2** | 2.3212 | 1.17148425048995 | 0.809 | 0.392 | 3.7750997 |
| **RASL11A** | 2.6553 | 0.876715082471699 | 0.603 | 0.138 | 4.3184463 |
| **SLC12A3** | 2.9914 | 1.07826561633814 | 0.353 | 0.013 | 4.8650314 |
| **MTCL1** | 3.3603 | -1.17301980786518 | 0.451 | 0.866 | 5.4649374 |
| **HIP1** | 4.8613 | -1.04902956300676 | 0.612 | 0.928 | 7.9060857 |
| **SCG3** | 2.6904 | -1.27250408297149 | 0.154 | 0.684 | 4.3754393 |
| **FAM174B** | 3.0938 | -1.4437022622349 | 0.323 | 0.793 | 5.0315408 |
| **MXRA8** | 7.8521 | -1.28578430762167 | 0.836 | 0.986 | 1.2769966 |
| **USP13** | 8.9579 | 1.00563109275084 | 0.99 | 0.952 | 1.4568308 |
| **FYN** | 1.5235 | -1.31282390708867 | 0.664 | 0.938 | 2.4777984 |
| **SLC2A4RG** | 4.9284 | -1.06651467636079 | 0.797 | 0.97 | 8.0151120 |
| **SPTB** | 1.2804 | -1.49887543306274 | 0.16 | 0.687 | 2.0823905 |

| | | | | | |
|---|---|---|---|---|---|
| **CTSB** | 1.7165 | 0.837569535104612 | 1 | 0.976 | 2.7916227 |
| **LRFN5** | 1.7239 | 0.852037313375512 | 0.457 | 0.056 | 2.8036527 |
| **PHC2** | 3.0019 | -1.79306149497007 | 0.037 | 0.589 | 4.8820876 |
| **DNAJB4** | 3.0983 | 1.08216811238083 | 0.899 | 0.555 | 5.0388280 |
| **CCN1** | 3.1828 | -1.87120174970022 | 0.724 | 0.948 | 5.1762796 |
| **PDCL** | 1.3223 | 0.812581393614281 | 0.999 | 0.988 | 2.1505239 |
| **SLC35F1** | 1.4852 | -1.15979013999709 | 0.125 | 0.671 | 2.4154816 |
| **MAGEF1** | 1.8478 | -1.02496406694991 | 0.694 | 0.945 | 3.0050929 |
| **FN1** | 1.8899 | -2.34223093940847 | 0.851 | 0.984 | 3.0736581 |
| **PLEKHA5** | 2.4520 | -1.09677706809668 | 0.592 | 0.929 | 3.9878270 |
| **TAOK3** | 2.6208 | 1.02933651891177 | 0.77 | 0.337 | 4.2622893 |
| **SLC7A6** | 2.8631 | 1.02741915985256 | 0.907 | 0.588 | 4.6563589 |
| **CPVL** | 3.7264 | -0.896174500013719 | 0.992 | 0.998 | 6.0602750 |
| **SAT2** | 3.7956 | -1.11745922802521 | 0.899 | 0.986 | 6.1729139 |
| **ZNF544** | 4.2480 | -0.838843121321352 | 0.953 | 0.997 | 6.9085846 |
| **SNRPG** | 4.8833 | 0.883079705963863 | 0.995 | 0.917 | 7.9418290 |
| **LRRC61** | 5.0809 | -1.05224519574586 | 0.207 | 0.735 | 8.2631422 |
| **TMEFF1** | 5.1591 | -1.33398957045916 | 0.775 | 0.967 | 8.3903695 |
| **EDNRB** | 8.2499 | -1.37891672467312 | 0.014 | 0.563 | 1.3416850 |
| **AC007906.2** | 9.2238 | -1.18679158057815 | 0.031 | 0.582 | 1.5000728 |
| **ZDHHC23** | 1.9579 | 0.980728890980593 | 0.691 | 0.222 | 3.1841810 |
| **ZIC5** | 2.5906 | -2.46034703430309 | 0.047 | 0.592 | 4.2132267 |
| **SLC37A4** | 2.8681 | 0.865553914592471 | 0.995 | 0.981 | 4.6644568 |
| **SMC3** | 3.8067 | -0.827634801426566 | 0.991 | 0.998 | 6.1909920 |
| **NUAK2** | 3.8276 | -1.48255702683748 | 0.238 | 0.731 | 6.2249646 |
| **TNNC2** | 3.9617 | 0.863297426757877 | 0.645 | 0.174 | 6.4429925 |
| **FAM13A** | 5.5726 | -1.10796286270539 | 0.274 | 0.776 | 9.0627318 |
| **SLC30A2** | 1.0655 | 1.08572797191745 | 0.61 | 0.156 | 1.7328884 |
| **REEP2** | 1.5948 | -1.3777691142096 | 0.187 | 0.709 | 2.5937658 |
| **FSTL1** | 2.4067 | -2.18839621241487 | 0.882 | 0.978 | 3.9141681 |
| **ABI2** | 4.7425 | -0.83488286067007 | 0.978 | 0.998 | 7.7128900 |
| **BTBD2** | 5.4683 | -0.935679834828142 | 0.946 | 0.995 | 8.8931563 |
| **HERPUD2** | 1.0831 | -1.05112163694101 | 0.552 | 0.9 | 1.7614721 |
| **CRELD2** | 4.3708 | 1.00286953018932 | 0.99 | 0.928 | 7.1083851 |

| | | | | | |
|---|---|---|---|---|---|
| **SERTAD4** | 5.2062 | -1.17168545930675 | 0.048 | 0.589 | 8.4668706 |
| **NAPRT** | 5.9972 | 0.985847500374599 | 0.901 | 0.515 | 9.7533900 |
| **ARGLU1** | 6.0836 | -1.00283202531949 | 0.996 | 0.999 | 9.8938391 |
| **UFM1** | 6.4897 | 0.918992347711862 | 0.996 | 0.948 | 1.0554275 |
| **HAS3** | 1.3591 | -1.20125578227541 | 0.04 | 0.581 | 2.2103566 |
| **TRIOBP** | 1.4222 | -0.918561926468837 | 0.762 | 0.972 | 2.3130613 |
| **SLC39A8** | 3.0285 | 1.19199167011432 | 0.601 | 0.143 | 4.9253810 |
| **CCNA1** | 4.0540 | 2.50795648597253 | 0.53 | 0.105 | 6.5930241 |
| **PCDHA4** | 4.3072 | -1.01939548122067 | 0.07 | 0.609 | 7.0048319 |
| **TNMD** | 8.0960 | -1.37904891217427 | 0.053 | 0.596 | 1.3166556 |
| **NEBL** | 8.7336 | -1.45846244697234 | 0.052 | 0.59 | 1.4203564 |
| **SEMA3A** | 9.5235 | -1.08996969686091 | 0.085 | 0.623 | 1.5488207 |
| **NKAIN1** | 1.0528 | -1.50742040279257 | 0.068 | 0.604 | 1.7121902 |
| **KIRREL1** | 1.2320 | 0.906863971969907 | 0.99 | 0.876 | 2.0037536 |
| **EEF1AKMT4** | 1.5564 | 0.892729289181883 | 0.969 | 0.753 | 2.5311942 |
| **WBP1** | 2.0421 | -0.934312757274447 | 0.87 | 0.986 | 3.3210858 |
| **CACFD1** | 2.4648 | -1.01515618610303 | 0.627 | 0.925 | 4.0085494 |
| **GRK6** | 2.4745 | 0.891764315289572 | 0.986 | 0.954 | 4.0243101 |
| **RNF43** | 2.6218 | -1.13458214724723 | 0.08 | 0.616 | 4.2638541 |
| **CCDC85C** | 7.2037 | -1.2091911477438 | 0.99 | 0.997 | 1.1715489 |
| **CPS1** | 8.2346 | -0.887744336106753 | 0.722 | 0.98 | 1.3391984 |
| **FGF2** | 9.5644 | -1.14826939091231 | 0.87 | 0.986 | 1.5554596 |
| **JARID2** | 9.5758 | -1.02536303905799 | 0.984 | 0.991 | 1.5573239 |
| **SPP1** | 1.3514 | -2.20825297150662 | 0.375 | 0.813 | 2.1978615 |
| **ZNF385A** | 1.7450 | -1.18652290485914 | 0.103 | 0.639 | 2.8379256 |
| **CA13** | 2.4580 | 1.22078244575629 | 0.818 | 0.443 | 3.9975056 |
| **TPM2** | 3.1559 | -1.15812177070542 | 0.994 | 0.999 | 5.1324656 |
| **FRMD4A** | 3.7856 | -1.47887085362915 | 0.076 | 0.61 | 6.1565512 |
| **ANK2** | 4.3615 | -1.10935547281052 | 0.599 | 0.916 | 7.0932334 |
| **AASS** | 4.7689 | 0.973991093377401 | 0.991 | 0.953 | 7.7558062 |
| **PTPRA** | 5.0809 | -0.867147482521456 | 0.877 | 0.983 | 8.2631909 |
| **RAP2C** | 5.5535 | -0.959715507177476 | 0.788 | 0.982 | 9.0317882 |
| **FBXL16** | 9.3287 | 0.916406813210131 | 0.61 | 0.159 | 1.5171320 |
| **HINT1** | 1.2391 | 0.821608732301468 | 0.999 | 0.977 | 2.0152633 |

| | | | | | |
|---|---|---|---|---|---|
| **RGMA** | 1.5820 | -1.66664388444242 | 0.345 | 0.779 | 2.5728376 |
| **PCGF6** | 2.1928 | 0.942884566496162 | 0.897 | 0.568 | 3.5662319 |
| **FAM217B** | 2.2105 | -0.924313835333946 | 0.656 | 0.936 | 3.5949746 |
| **LAMA2** | 2.7048 | -1.57058045872368 | 0.174 | 0.662 | 4.3989704 |
| **FERMT1** | 4.2060 | 1.00272979275557 | 0.752 | 0.309 | 6.8403309 |
| **AP1M1** | 8.3810 | -0.910558330318253 | 0.833 | 0.98 | 1.3630138 |
| **ENDOG** | 8.8869 | 0.873989114570979 | 0.977 | 0.816 | 1.4452906 |
| **GJC1** | 9.0415 | -1.58478892671793 | 0.614 | 0.891 | 1.4704228 |
| **OTUD4** | 1.9713 | 0.887547519571032 | 0.972 | 0.83 | 3.2059674 |
| **MAPK12** | 2.5669 | 0.950370217184294 | 0.86 | 0.438 | 4.1746360 |
| **GRHL3** | 3.8599 | -1.72649311276535 | 0.076 | 0.6 | 6.2773602 |
| **SVBP** | 4.3980 | -0.909486867999369 | 0.868 | 0.985 | 7.1525141 |
| **GNG4** | 5.0989 | 0.961619562000459 | 0.581 | 0.144 | 8.2924355 |
| **CLASP1** | 5.5867 | -0.821590358643067 | 0.89 | 0.986 | 9.0857984 |
| **PTPRS** | 6.4624 | -1.20991442407466 | 0.981 | 0.999 | 1.0509889 |
| **GFPT2** | 6.8536 | -1.21552683625972 | 0.702 | 0.931 | 1.1146136 |
| **ZKSCAN1** | 1.6883 | -1.08048357549341 | 0.455 | 0.853 | 2.7457241 |
| **PI4K2B** | 1.8153 | 0.942003913080778 | 0.975 | 0.864 | 2.9522676 |
| **CAMK1D** | 2.3483 | 0.824535744168111 | 0.534 | 0.109 | 3.8191412 |
| **DND1** | 2.3642 | 0.972986873523967 | 0.554 | 0.123 | 3.8450606 |
| **GYG2** | 2.9985 | -1.02217824436983 | 0.627 | 0.924 | 4.8765505 |
| **C6orf132** | 3.4374 | 0.844165859537336 | 0.996 | 0.935 | 5.5903474 |
| **PLPP1** | 4.4098 | 1.00179555909858 | 1 | 0.968 | 7.1716607 |
| **POGZ** | 4.6861 | -0.862437168033645 | 0.886 | 0.988 | 7.6210743 |
| **CLDN19** | 5.1443 | -1.17733304533228 | 0.081 | 0.608 | 8.3662275 |
| **FGFBP3** | 1.0310 | -1.91811673890337 | 0.016 | 0.54 | 1.6768425 |
| **PTPRZ1** | 1.1040 | -1.59008339846619 | 0.032 | 0.557 | 1.7955175 |
| **GALC** | 1.2541 | -1.05662543732775 | 0.463 | 0.84 | 2.0397052 |
| **TMSB15A** | 2.3087 | -1.82413413029079 | 0.049 | 0.571 | 3.7546711 |
| **RTN3** | 3.2571 | -0.920953252642047 | 0.996 | 0.997 | 5.2971346 |
| **CERS4** | 3.4891 | -1.19272760962291 | 0.952 | 0.992 | 5.6743832 |
| **MID1** | 5.1037 | -1.07574801499716 | 0.561 | 0.927 | 8.3002124 |
| **PLPPR2** | 5.6581 | 0.920677547338594 | 0.788 | 0.334 | 9.2017810 |
| **GRID2** | 6.4008 | -1.47643804394299 | 0.143 | 0.653 | 1.0409643 |

| | | | | | |
|---|---|---|---|---|---|
| **ZEB2** | 6.6818 | -1.67698599447077 | 0.075 | 0.601 | 1.0866688 |
| **GSTA4** | 7.4518 | -0.989564302252814 | 0.678 | 0.941 | 1.2118970 |
| **PCED1A** | 7.8102 | -1.23179145763671 | 0.221 | 0.721 | 1.2701870 |
| **LIMD2** | 7.9182 | -1.0623336635382 | 0.483 | 0.865 | 1.2877464 |
| **LUZP1** | 8.3480 | -0.899566780876374 | 0.911 | 0.995 | 1.3576437 |
| **HSPB8** | 9.4629 | -1.35761146944313 | 0.044 | 0.572 | 1.5389524 |
| **GALNT17** | 1.4224 | -1.18018669711466 | 0.602 | 0.924 | 2.3132511 |
| **MAPRE2** | 2.1185 | -1.05001211206545 | 0.61 | 0.908 | 3.4453802 |
| **FOS** | 2.6128 | -1.44477069763313 | 0.164 | 0.693 | 4.2492272 |
| **EDEM1** | 4.6524 | 0.965355434355668 | 0.95 | 0.814 | 7.5662696 |
| **OTUD1** | 5.7133 | 1.1493223965051 | 0.722 | 0.291 | 9.2915589 |
| **PLEKHG3** | 1.0315 | -1.701400045279 | 0.115 | 0.623 | 1.6775945 |
| **ANKRD50** | 3.0536 | -0.960582753046537 | 0.656 | 0.929 | 4.9662311 |
| **CHD6** | 3.3294 | -0.939736487450677 | 0.726 | 0.952 | 5.4147129 |
| **OCIAD2** | 3.3376 | -0.950143793562402 | 0.044 | 0.567 | 5.4279654 |
| **GAMT** | 5.0883 | 0.918860060319455 | 0.899 | 0.614 | 8.2752444 |
| **KIDINS220** | 6.4636 | -0.876156726390851 | 0.835 | 0.975 | 1.0511858 |
| **ERMP1** | 8.0474 | -1.04198516754008 | 0.492 | 0.859 | 1.3087552 |
| **CCN2** | 8.5122 | -1.81835494552053 | 0.619 | 0.924 | 1.3843518 |
| **STMN1** | 9.6274 | 0.925900774064322 | 0.996 | 0.983 | 1.5657067 |
| **FOCAD** | 1.2655 | -0.827914978079219 | 0.857 | 0.978 | 2.0581740 |
| **PPIF** | 2.1490 | 0.911220186106227 | 0.997 | 0.934 | 3.4950374 |
| **CCDC97** | 3.7519 | -0.859670628696303 | 0.813 | 0.971 | 6.1018569 |
| **SLC38A2** | 5.9937 | -0.878688001933007 | 0.956 | 0.998 | 9.7477120 |
| **TEAD1** | 8.6700 | -1.0271767071638 | 0.488 | 0.862 | 1.4100139 |
| **HHIPL1** | 9.0725 | -1.14217935542746 | 0.156 | 0.66 | 1.4754639 |
| **CMTM3** | 1.5107 | -1.29821276155627 | 0.933 | 0.991 | 2.4568724 |
| **KLHDC7A** | 1.5817 | -1.08717682155892 | 0.012 | 0.525 | 2.5724303 |
| **ZNF432** | 2.3215 | 0.926489547672841 | 0.818 | 0.419 | 3.7756028 |
| **GBX2** | 3.1205 | -1.28102913160206 | 0.181 | 0.682 | 5.0749280 |
| **TUBA1A** | 3.3671 | -1.46918001562321 | 0.628 | 0.895 | 5.4760397 |
| **FAM20A** | 3.5465 | -1.7471794070282 | 0.509 | 0.846 | 5.7677866 |
| **GLRX2** | 3.8021 | -0.997206379451646 | 0.124 | 0.638 | 6.1833882 |
| **TADA3** | 4.1031 | -0.822968981645485 | 0.875 | 0.988 | 6.6728729 |

| | | | | | |
|---|---|---|---|---|---|
| **PPP2R3A** | 4.3371 | -0.924240012247908 | 0.578 | 0.917 | 7.0534433 |
| **MYRFL** | 4.6087 | -1.16913874874375 | 0.025 | 0.538 | 7.4952477 |
| **HRAS** | 5.0910 | -0.833706391990682 | 0.91 | 0.999 | 8.2795163 |
| **CPT1A** | 5.6483 | 0.921620501526067 | 0.747 | 0.297 | 9.1859494 |
| **TNKS** | 5.7701 | -0.836382711833861 | 0.91 | 0.989 | 9.3840641 |
| **C1orf198** | 6.3671 | -1.27339567419387 | 0.581 | 0.899 | 1.0354892 |
| **ZBTB33** | 7.2918 | -0.925849966460937 | 0.731 | 0.97 | 1.1858680 |
| **ELMOD2** | 8.7670 | -0.949129546325684 | 0.565 | 0.896 | 1.4257891 |
| **PSRC1** | 9.5967 | 1.12360245287892 | 0.973 | 0.812 | 1.5607151 |
| **STARD8** | 9.8928 | -1.04658233562475 | 0.018 | 0.529 | 1.6088782 |
| **DPH6** | 1.4062 | 0.890099326415903 | 0.952 | 0.722 | 2.2869571 |
| **JAG2** | 1.7159 | 0.977210904605832 | 0.69 | 0.241 | 2.7906783 |
| **ITPR3** | 1.8912 | 1.01553811134844 | 0.893 | 0.565 | 3.0758192 |
| **KLF2** | 2.0983 | 1.16817856398417 | 0.66 | 0.195 | 3.4125560 |
| **SPR** | 2.2851 | 0.978040194933753 | 0.84 | 0.427 | 3.7162756 |
| **ANKRD10** | 2.3646 | -1.15463369688595 | 0.965 | 0.993 | 3.8456222 |
| **TTYH1** | 2.5047 | 1.00713497187557 | 0.682 | 0.2 | 4.0735297 |
| **WDR34** | 3.1938 | 0.833490233563677 | 0.987 | 0.936 | 5.1941035 |
| **MCUB** | 5.6405 | 0.961187777660844 | 0.836 | 0.484 | 9.1732674 |
| **MED14** | 5.7579 | -0.883786201324787 | 0.912 | 0.992 | 9.3642093 |
| **GCA** | 6.7735 | -1.04380509124668 | 0.348 | 0.787 | 1.1015796 |
| **USB1** | 7.2777 | -0.829224022377417 | 0.876 | 0.988 | 1.1835880 |
| **PKIA** | 7.3447 | -0.916957311788799 | 0.035 | 0.544 | 1.1944706 |
| **CDYL** | 7.6850 | -0.900285098541421 | 0.766 | 0.972 | 1.2498212 |
| **SPIN4** | 1.3540 | -1.05784802997628 | 0.815 | 0.975 | 2.2021610 |
| **PSIP1** | 1.3857 | -0.822330586395991 | 0.974 | 0.992 | 2.2536811 |
| **SSBP3** | 1.5485 | -0.916752693392523 | 0.708 | 0.95 | 2.5184748 |
| **FOXI3** | 2.5489 | -1.29685488512003 | 0.951 | 0.972 | 4.1453611 |
| **TAF9B** | 3.0461 | -0.98016900060985 | 0.43 | 0.82 | 4.9540053 |
| **BTG2** | 3.5290 | -1.59334337726078 | 0.381 | 0.801 | 5.7392339 |
| **DOCK6** | 4.3484 | 0.8079404001518 | 0.975 | 0.841 | 7.0719535 |
| **REEP3** | 4.7766 | -0.908201540760866 | 0.643 | 0.925 | 7.7681845 |
| **PIK3R3** | 6.4764 | -1.35493981296473 | 0.676 | 0.914 | 1.0532730 |
| **KMT2E** | 6.8200 | -0.901230346906807 | 0.669 | 0.938 | 1.1091472 |

| | | | | | |
|---|---|---|---|---|---|
| **TRIM21** | 1.6431 | -0.862276571750444 | 0.045 | 0.555 | 2.6722486 |
| **UNC5D** | 1.7423 | -0.916430022148698 | 0.022 | 0.527 | 2.8336201 |
| **PLAGL2** | 1.9432 | -0.932496149840407 | 0.572 | 0.881 | 3.1603129 |
| **EGLN1** | 2.9248 | -0.968291140272662 | 0.578 | 0.917 | 4.7567399 |
| **CENPA** | 3.0767 | 1.1107543466816 | 0.991 | 0.851 | 5.0036477 |
| **GAS1** | 3.7758 | -1.03704197276831 | 0.112 | 0.621 | 6.1406691 |
| **ZDHHC2** | 4.0643 | -1.03387948190024 | 0.096 | 0.601 | 6.6098261 |
| **ARL4C** | 4.9939 | -2.70625246081491 | 0.125 | 0.616 | 8.1217244 |
| **FBN3** | 5.0902 | -1.17731279824065 | 0.266 | 0.735 | 8.2782350 |
| **MAPK4** | 5.3070 | -1.17959393233879 | 0.059 | 0.573 | 8.6308411 |
| **CCDC69** | 7.2570 | 0.980382925435934 | 0.822 | 0.436 | 1.1802080 |
| **SPRED2** | 8.4855 | -1.0968699910093 | 0.375 | 0.795 | 1.3800045 |
| **PTPN21** | 1.0620 | -1.0811147516127 | 0.243 | 0.715 | 1.7272521 |
| **MSL2** | 1.5593 | -0.864556089456392 | 0.699 | 0.956 | 2.5359785 |
| **SSC4D** | 3.0512 | -0.963190268579796 | 0.081 | 0.59 | 4.9622902 |
| **PPP1R1A** | 6.0818 | -1.2987062907031 | 0.549 | 0.883 | 9.8908573 |
| **STAT3** | 7.5559 | -0.820061209929668 | 0.994 | 0.999 | 1.2288230 |
| **PON2** | 1.2691 | -0.93734540938665 | 0.708 | 0.943 | 2.0640148 |
| **TP53BP2** | 1.6961 | -0.824401273058709 | 0.877 | 0.984 | 2.7583678 |
| **EPPK1** | 2.6550 | 0.805069771616953 | 0.996 | 0.985 | 4.3179208 |
| **USO1** | 6.1623 | -1.01360593797733 | 0.997 | 0.998 | 1.0021821 |
| **KIF26A** | 7.3418 | -1.08332827959176 | 0.866 | 0.966 | 1.1939991 |
| **CBX4** | 1.3248 | 0.956593528051138 | 0.969 | 0.819 | 2.1545423 |
| **ARSB** | 2.1447 | -0.931871012746124 | 0.623 | 0.914 | 3.4880596 |
| **KIZ** | 2.9324 | -1.03641169152197 | 0.486 | 0.847 | 4.7690201 |
| **LTBP1** | 2.9543 | -1.16623051788457 | 0.499 | 0.852 | 4.8046874 |
| **CAV1** | 7.2623 | -1.59655564825483 | 0.092 | 0.587 | 1.1810718 |
| **QSER1** | 8.7945 | -0.828845035798403 | 0.899 | 0.985 | 1.4302553 |
| **ADCY3** | 1.0422 | 0.871973079097562 | 0.739 | 0.289 | 1.6950302 |
| **ARSI** | 2.1200 | 0.956437809180528 | 0.898 | 0.63 | 3.4477864 |
| **PSME1** | 3.5579 | -0.90925311676438 | 0.832 | 0.965 | 5.7863729 |
| **SELENOM** | 4.6240 | 0.908862916663342 | 0.855 | 0.502 | 7.5200509 |
| **STRA6** | 4.8386 | -1.15083562824609 | 0.088 | 0.587 | 7.8690254 |
| **ZBTB18** | 6.3540 | 0.930351202223041 | 0.919 | 0.7 | 1.0333582 |

| | | | | | |
|---|---|---|---|---|---|
| **CCPG1** | 6.8942 | 0.982952708503686 | 0.95 | 0.776 | 1.12121482 |
| **HIPK2** | 8.1230 | -1.14677594650855 | 0.576 | 0.871 | 1.32105446 |
| **DPY19L1** | 1.1474 | 0.930855502008848 | 0.86 | 0.538 | 1.86613861 |
| **ATP1A2** | 1.7134 | -1.33830630174349 | 0.013 | 0.504 | 2.78650504 |
| **ADD3** | 2.1776 | -0.813014226335101 | 0.988 | 0.999 | 3.54156622 |
| **AGPAT5** | 4.0712 | -0.91805985438914 | 0.842 | 0.973 | 6.62100449 |
| **SAR1B** | 9.5714 | 0.824889728798858 | 0.969 | 0.855 | 1.55661219 |
| **SIGIRR** | 9.9575 | 0.843201786896423 | 0.969 | 0.751 | 1.61939412 |
| **RABGAP1L** | 1.0821 | -1.00959332157406 | 0.689 | 0.937 | 1.75985769 |
| **AIRE** | 1.6327 | 0.926404158365552 | 0.545 | 0.132 | 2.65534670 |
| **KIAA0930** | 1.6988 | 0.821198779222689 | 0.966 | 0.852 | 2.76288042 |
| **RET** | 3.1612 | -1.40405554305296 | 0.358 | 0.756 | 5.14110785 |
| **C6orf120** | 5.0055 | -0.835761002801199 | 0.797 | 0.977 | 8.14056809 |
| **PTPRG** | 5.2097 | -1.09084565802862 | 0.68 | 0.933 | 8.47261502 |
| **CDC42EP3** | 5.3862 | -1.16626830306957 | 0.185 | 0.675 | 8.75973691 |
| **ITGA5** | 6.0124 | -1.43414719547037 | 0.053 | 0.545 | 9.77812494 |
| **ADCY6** | 6.9464 | -1.08016316521771 | 0.058 | 0.549 | 1.12970013 |
| **DPP10** | 1.2606 | -1.22673389665163 | 0.252 | 0.694 | 2.05018670 |
| **JUND** | 1.4183 | -0.833455659645899 | 1 | 1 | 2.30658370 |
| **NREP** | 1.9892 | -0.806871093857589 | 0.978 | 0.999 | 3.23513159 |
| **RAB6B** | 2.0130 | -1.21719126190282 | 0.452 | 0.826 | 3.27376979 |
| **CD9** | 2.2117 | 0.802849328788733 | 1 | 0.973 | 3.59693383 |
| **KRT19** | 2.2885 | -1.04039233462251 | 0.793 | 0.966 | 3.72192140 |
| **UHRF2** | 2.5813 | -1.13744895475678 | 0.265 | 0.706 | 4.19800644 |
| **RTL8A** | 2.8185 | -0.991623644103286 | 0.301 | 0.742 | 4.58377649 |
| **PARVA** | 4.7336 | -1.04241955788159 | 0.722 | 0.963 | 7.69835159 |
| **ATP1B2** | 6.4144 | -1.32965920581658 | 0.26 | 0.712 | 1.04318137 |
| **PHTF1** | 2.9811 | -0.99242477773475 | 0.645 | 0.912 | 4.84831395 |
| **LRRN2** | 3.4557 | -1.07375108197343 | 0.067 | 0.554 | 5.62012310 |
| **SLC12A8** | 3.5104 | -1.13329769401623 | 0.123 | 0.603 | 5.70910850 |
| **MEX3B** | 3.7591 | -1.20351637665668 | 0.107 | 0.588 | 6.11346973 |
| **RHBDL3** | 3.9631 | -1.083687053906 | 0.056 | 0.538 | 6.44531439 |
| **GSTK1** | 3.9846 | -0.990416073872163 | 0.634 | 0.91 | 6.48029631 |
| **SEL1L3** | 4.0732 | -1.13718158086685 | 0.105 | 0.589 | 6.62438770 |

| | | | | | |
|---|---|---|---|---|---|
| **AGPAT4** | 5.9461 | -1.02800740351254 | 0.548 | 0.881 | 9.67015025 |
| **PHLDB2** | 7.7592 | -1.14409038497036 | 0.262 | 0.731 | 1.26188489 |
| **SH3BGRL3** | 8.0524 | -0.982191547812225 | 0.966 | 0.993 | 1.30956866 |
| **DNMBP** | 8.3876 | -0.964037346719064 | 0.713 | 0.934 | 1.36407794 |
| **MCM8** | 1.1503 | -0.847519461721229 | 0.944 | 0.984 | 1.87082183 |
| **ZNF580** | 1.1681 | -1.06266996405171 | 0.443 | 0.837 | 1.89978210 |
| **B2M** | 1.7996 | -1.18383737203242 | 0.655 | 0.918 | 2.92677583 |
| **PRTG** | 2.3904 | -2.48696009884916 | 0.03 | 0.511 | 3.88756808 |
| **ILK** | 4.9201 | -0.852324583395516 | 0.846 | 0.971 | 8.00158884 |
| **NCAM1** | 6.4616 | -1.21153100282948 | 0.027 | 0.505 | 1.05085039 |
| **ZSWIM9** | 6.7653 | -0.959903354779943 | 0.186 | 0.664 | 1.10025248 |
| **STC1** | 7.5275 | -1.15732496466268 | 0.014 | 0.492 | 1.22420397 |
| **ACTR6** | 7.9065 | 0.850526207881832 | 0.826 | 0.468 | 1.28583965 |
| **GRHL1** | 9.7829 | 0.886855880241858 | 0.672 | 0.25 | 1.59100815 |
| **SERPINB9** | 1.0633 | 1.10958729962703 | 0.997 | 0.929 | 1.72933614 |
| **USP46** | 1.4447 | -0.832690957838682 | 0.743 | 0.935 | 2.34960268 |
| **CALM2** | 1.8734 | -0.847826486019534 | 0.988 | 1 | 3.04684507 |
| **MED12** | 1.9739 | -1.0711212705551 | 0.097 | 0.579 | 3.21015687 |
| **CRTC3** | 2.7329 | -1.01895504410507 | 0.261 | 0.715 | 4.44460857 |
| **KLHL4** | 5.9085 | -0.988652093721341 | 0.018 | 0.492 | 9.60901256 |
| **C12orf75** | 7.1349 | 0.920995867959389 | 0.975 | 0.919 | 1.16035331 |
| **FABP3** | 9.9089 | 0.872135127166463 | 0.996 | 0.876 | 1.61149001 |
| **PHLDB1** | 2.1096 | -1.01636654013915 | 0.633 | 0.908 | 3.43087389 |
| **CHIC1** | 2.2218 | -0.919530033954874 | 0.216 | 0.685 | 3.61347325 |
| **BCAR1** | 2.3530 | -1.01875119231701 | 0.89 | 0.977 | 3.82671318 |
| **SLC7A2** | 4.0281 | -1.26637573703572 | 0.248 | 0.683 | 6.55100759 |
| **PDE3B** | 4.6797 | -0.98240663210973 | 0.072 | 0.549 | 7.61062227 |
| **ADK** | 4.7447 | -0.818234816334816 | 0.81 | 0.969 | 7.71636660 |
| **CORO1A** | 6.2899 | 0.826497280490323 | 0.558 | 0.152 | 1.02292752 |
| **RB1** | 9.0369 | -0.923637039055424 | 0.403 | 0.781 | 1.46968570 |
| **SH2D3A** | 1.0987 | 0.811147201574284 | 0.819 | 0.381 | 1.78688021 |
| **TRIM22** | 1.8848 | -1.22663480418624 | 0.51 | 0.846 | 3.06539145 |
| **RPTOR** | 3.1174 | 0.843852043500227 | 0.963 | 0.843 | 5.06998383 |
| **CHST6** | 4.3097 | -1.02487466863903 | 0.376 | 0.767 | 7.00897229 |

| | | | | | |
|---|---|---|---|---|---|
| **SKA3** | 4.6517 | 0.953204409809933 | 0.972 | 0.837 | 7.5651701 |
| **ADGRE5** | 7.2545 | 0.939143971304158 | 0.818 | 0.473 | 1.1798055 |
| **OGG1** | 7.3283 | -0.851085986319644 | 0.633 | 0.907 | 1.1918069 |
| **PXMP4** | 1.0287 | 0.868689879720971 | 0.845 | 0.505 | 1.6730861 |
| **FAM151A** | 1.1383 | 1.62908553082219 | 0.271 | 0.008 | 1.8512635 |
| **PLPP2** | 1.1988 | 0.915158294375384 | 0.448 | 0.086 | 1.9497323 |
| **TCEAL3** | 1.4714 | -0.868516352601553 | 0.093 | 0.573 | 2.3930979 |
| **HEG1** | 1.8345 | -1.20460593784148 | 0.234 | 0.671 | 2.9835369 |
| **SC5D** | 1.9732 | -0.912483819437671 | 0.87 | 0.983 | 3.2090873 |
| **POLI** | 2.0926 | -0.879676753099094 | 0.682 | 0.923 | 3.4032517 |
| **RAVER2** | 2.1242 | -0.936465694417264 | 0.367 | 0.783 | 3.4547338 |
| **PFKFB3** | 2.5492 | -0.938435156113403 | 0.528 | 0.869 | 4.1458803 |
| **TMEM229B** | 5.2351 | 0.980314474498429 | 0.73 | 0.301 | 8.5139156 |
| **EYA2** | 7.7651 | -1.26024237755323 | 0.154 | 0.613 | 1.2628483 |
| **LPCAT1** | 1.3869 | -0.920554619994022 | 0.997 | 0.997 | 2.2556489 |
| **EXOC4** | 2.7352 | -0.810978063125334 | 0.851 | 0.976 | 4.4483400 |
| **IL34** | 2.8072 | 0.801172141672355 | 0.61 | 0.195 | 4.5653517 |
| **ARSE** | 4.4573 | -1.05995215941718 | 0.093 | 0.567 | 7.2490078 |
| **PALLD** | 4.7315 | -1.78386656913523 | 0.218 | 0.666 | 7.6949770 |
| **CHST12** | 5.2935 | 0.813010513608917 | 0.921 | 0.698 | 8.6088192 |
| **ZNF609** | 5.5898 | -0.846848806688125 | 0.659 | 0.916 | 9.0908194 |
| **LZTS1** | 5.8148 | 0.871540385822472 | 0.845 | 0.496 | 9.4567348 |
| **CARMIL1** | 6.9577 | -1.0393964868678 | 0.364 | 0.774 | 1.1315353 |
| **PTPRN** | 7.2340 | 0.805922381085655 | 0.532 | 0.131 | 1.1764783 |
| **IGFBP7** | 9.1226 | -1.78690630365875 | 0.164 | 0.615 | 1.4836101 |
| **ADGRL2** | 9.9583 | -0.883921477905318 | 0.91 | 0.992 | 1.6195300 |
| **LPIN1** | 1.0166 | 0.895366458809407 | 0.859 | 0.514 | 1.6534315 |
| **EPDR1** | 2.2606 | -1.0058587878747 | 0.028 | 0.491 | 3.6765481 |
| **ROR2** | 2.8193 | -1.28299343339858 | 0.372 | 0.774 | 4.5851818 |
| **SHTN1** | 3.5535 | -0.968672755151461 | 0.488 | 0.848 | 5.7790760 |
| **CPD** | 4.5580 | -0.877107070083524 | 0.968 | 0.997 | 7.4126917 |
| **LRAT** | 5.0875 | -1.57728914502145 | 0.083 | 0.545 | 8.2738339 |
| **CHD9** | 5.1044 | -0.874327015474176 | 0.694 | 0.933 | 8.3013723 |
| **SDR39U1** | 6.0369 | -0.804646185471581 | 0.851 | 0.971 | 9.8179626 |

| | | | | | |
|---|---|---|---|---|---|
| **EPHB1** | 9.5818 | -0.969725944011898 | 0.039 | 0.502 | 1.5583000 |
| **FOXB1** | 1.6400 | -1.34954032609522 | 0.152 | 0.616 | 2.6672141 |
| **XPNPEP1** | 7.0914 | -0.808486827013073 | 0.773 | 0.961 | 1.1532761 |
| **TRERF1** | 1.2283 | -0.961113538199615 | 0.028 | 0.489 | 1.9977265 |
| **LZTS2** | 1.2381 | -0.901047606923259 | 0.761 | 0.951 | 2.0135841 |
| **VEZF1** | 1.5991 | -0.887018848647817 | 0.643 | 0.901 | 2.6006577 |
| **TMEM47** | 1.7601 | -0.95534902997847 | 0.081 | 0.548 | 2.8624705 |
| **C1orf54** | 1.9212 | -0.843997030695654 | 0.085 | 0.553 | 3.1245813 |
| **ST7** | 2.0973 | -0.910135591445423 | 0.205 | 0.659 | 3.4108776 |
| **CAPRIN2** | 2.7644 | -1.19205134556599 | 0.372 | 0.77 | 4.4957935 |
| **SHISA3** | 2.7700 | -1.762722915045 | 0.044 | 0.501 | 4.5049483 |
| **GLIPR2** | 3.3410 | -1.42582928708222 | 0.944 | 0.977 | 5.4334687 |
| **NRARP** | 7.5877 | -0.990980805511224 | 0.255 | 0.7 | 1.2339951 |
| **NTF3** | 8.1040 | -1.09100425770904 | 0.045 | 0.506 | 1.3179584 |
| **FLT1** | 8.1481 | -1.59451484416153 | 0.152 | 0.599 | 1.3251287 |
| **PIPOX** | 9.3556 | -1.12459829834286 | 0.2 | 0.646 | 1.5215131 |
| **RASD1** | 1.0896 | 0.932271387706239 | 0.537 | 0.152 | 1.7721200 |
| **B4GALT4** | 1.0905 | 0.90062740606199 | 0.761 | 0.396 | 1.7735701 |
| **PIM1** | 1.1411 | -0.954769323439653 | 0.964 | 0.989 | 1.8558752 |
| **CDC25B** | 1.1500 | -0.926817339779253 | 0.982 | 0.99 | 1.8703284 |
| **GRAMD1B** | 1.3044 | -1.17412137026186 | 0.035 | 0.492 | 2.1214314 |
| **C1GALT1C1** | 1.3634 | -0.843039282469895 | 0.782 | 0.957 | 2.2173976 |
| **TRIM38** | 1.5467 | 1.00486845960843 | 0.839 | 0.495 | 2.5154954 |
| **GALNT7** | 2.5821 | -1.07034241174236 | 0.92 | 0.98 | 4.1993079 |
| **TLNRD1** | 3.8215 | -0.905152857243689 | 0.77 | 0.946 | 6.2149523 |
| **SGCB** | 4.2212 | -0.832246296386695 | 0.693 | 0.929 | 6.8650093 |
| **GREM2** | 8.0112 | 0.950804935832192 | 0.673 | 0.253 | 1.3028697 |
| **SLC2A1** | 8.8431 | -0.904011895496525 | 0.986 | 0.997 | 1.4381636 |
| **TSPAN15** | 9.4181 | -0.900935872470897 | 0.171 | 0.634 | 1.5316772 |
| **GDF15** | 1.0138 | 1.49953468541056 | 0.43 | 0.083 | 1.6487974 |
| **C14orf132** | 1.5760 | -1.29759151359707 | 0.041 | 0.495 | 2.5630900 |
| **EFS** | 2.0512 | -1.58591854245397 | 0.248 | 0.673 | 3.3358993 |
| **IFI27L2** | 2.5288 | -1.05391321097181 | 0.196 | 0.645 | 4.1127048 |
| **RND3** | 2.8790 | -1.24383412063626 | 0.05 | 0.505 | 4.6822463 |

| | | | | | |
|---|---|---|---|---|---|
| **FBRS** | 6.2937 | 1.06914372574867 | 0.921 | 0.739 | 1.0235565 |
| **DCX** | 8.8588 | -0.808382848742266 | 0.025 | 0.473 | 1.4407162 |
| **CADM1** | 1.2479 | -1.13173061334784 | 0.556 | 0.867 | 2.0295398 |
| **RIC8B** | 1.4711 | -1.09811634170898 | 0.453 | 0.801 | 2.3925567 |
| **TRAPPC6A** | 2.6868 | -0.821767335756827 | 0.833 | 0.977 | 4.3695940 |
| **SOX13** | 4.3943 | -0.865175837694867 | 0.713 | 0.924 | 7.1465255 |
| **SLC12A2** | 4.4012 | 0.870959591757571 | 0.784 | 0.435 | 7.1577489 |
| **CTSV** | 5.1534 | -1.35175355416228 | 0.93 | 0.981 | 8.3810741 |
| **HDAC7** | 5.9944 | -0.914911443891599 | 0.674 | 0.916 | 9.7488541 |
| **CADM4** | 7.1423 | -1.04761999746853 | 0.158 | 0.605 | 1.1615605 |
| **STARD10** | 1.4405 | 0.861034431688827 | 0.876 | 0.583 | 2.3427660 |
| **SCO2** | 1.8572 | 0.829569082336358 | 0.917 | 0.696 | 3.0204624 |
| **GRTP1** | 2.3584 | 0.835553797880996 | 0.784 | 0.403 | 3.8354777 |
| **NCS1** | 3.2900 | -0.943818415695775 | 0.572 | 0.873 | 5.3505852 |
| **ZNF316** | 4.9007 | -0.859724783053298 | 0.291 | 0.712 | 7.9700547 |
| **NET1** | 6.6060 | -0.848167347223928 | 0.677 | 0.913 | 1.0743469 |
| **VANGL1** | 6.9505 | -0.904588576181293 | 0.641 | 0.906 | 1.1303653 |
| **ZFHX3** | 7.7109 | 0.823688555898026 | 0.554 | 0.161 | 1.2540273 |
| **PTH1R** | 9.8725 | 1.02021322088951 | 0.669 | 0.268 | 1.6055688 |
| **FNDC3A** | 1.4282 | -0.801619740350862 | 0.872 | 0.983 | 2.3227655 |
| **INKA2** | 2.1309 | -1.00483092842974 | 0.209 | 0.657 | 3.4655639 |
| **CAVIN3** | 3.2279 | -1.04747318368139 | 0.322 | 0.739 | 5.2495614 |
| **TSPAN9** | 3.5890 | -0.870234774763063 | 0.778 | 0.952 | 5.8368612 |
| **FRMD6** | 5.1633 | 0.992798755005788 | 0.925 | 0.705 | 8.3972326 |
| **KIF9** | 6.8745 | 0.888665329561273 | 0.912 | 0.677 | 1.1180127 |
| **RNF145** | 8.4233 | -0.88218212515374 | 0.49 | 0.817 | 1.3698898 |
| **UCP2** | 2.6771 | 0.828337732007619 | 0.977 | 0.912 | 4.3538282 |
| **OSMR** | 3.2804 | 0.849164371753912 | 0.479 | 0.113 | 5.3350012 |
| **ATP12A** | 5.5845 | 0.937115964160526 | 0.868 | 0.504 | 9.0822260 |
| **TRIM25** | 1.8206 | -1.00027034480373 | 0.579 | 0.85 | 2.9609114 |
| **ELF4** | 2.3336 | -1.33667854625268 | 0.133 | 0.571 | 3.7952560 |
| **CTTNBP2** | 6.5850 | -0.823729300728029 | 0.026 | 0.461 | 1.0709342 |
| **SPIN3** | 7.4365 | -1.01297511106057 | 0.39 | 0.771 | 1.2094110 |
| **SGSM2** | 8.7134 | -0.954090000595568 | 0.307 | 0.719 | 1.4170688 |

| | | | | | |
|---|---|---|---|---|---|
| **OBI1** | 9.3103 | -0.823056602280047 | 0.817 | 0.957 | 1.5141481 |
| **COL2A1** | 1.4657 | -2.30621754340673 | 0.006 | 0.436 | 2.3837034 |
| **C11orf95** | 2.2036 | -0.92343021448893 | 0.642 | 0.9 | 3.5837359 |
| **KCNH2** | 2.2390 | 0.834227590234552 | 0.858 | 0.532 | 3.6413055 |
| **ARMCX4** | 2.4507 | -0.815762379107863 | 0.15 | 0.595 | 3.9855873 |
| **FZD3** | 2.7654 | -0.875343918450166 | 0.912 | 0.987 | 4.4973707 |
| **PCDHA12** | 2.8176 | -0.807806116757416 | 0.076 | 0.517 | 4.5824090 |
| **PDE7A** | 3.1297 | -0.835926147256028 | 0.893 | 0.976 | 5.0898399 |
| **CACNA1B** | 3.6819 | 0.814529878569244 | 0.721 | 0.324 | 5.9880280 |
| **ACADS** | 3.7767 | 0.813894846640479 | 0.731 | 0.348 | 6.1421171 |
| **ARHGEF2** | 4.3316 | -0.864935526604576 | 0.671 | 0.928 | 7.0445772 |
| **GULP1** | 5.7674 | -0.960967799466803 | 0.24 | 0.659 | 9.3795963 |
| **STON2** | 8.1192 | -1.24824176217018 | 0.328 | 0.721 | 1.3204272 |
| **DIAPH2** | 8.8604 | -0.905165096333112 | 0.167 | 0.613 | 1.4409806 |
| **SNAPC3** | 1.1811 | 0.814538672306434 | 0.963 | 0.853 | 1.9209754 |
| **TNC** | 1.2891 | -1.47893425305174 | 0.023 | 0.453 | 2.0965082 |
| **NPHP3** | 1.5964 | -0.967011588046089 | 0.323 | 0.718 | 2.5963142 |
| **SDSL** | 1.7551 | -0.824531016945505 | 0.125 | 0.57 | 2.8543542 |
| **STC2** | 2.0214 | 1.3695905179876 | 0.917 | 0.806 | 3.2875415 |
| **CCDC80** | 2.7624 | 0.854890949332408 | 0.946 | 0.75 | 4.4925226 |
| **FOXJ1** | 2.8760 | -1.05061250361418 | 0.026 | 0.457 | 4.6773310 |
| **CDH13** | 4.9754 | -1.3490804827063 | 0.14 | 0.574 | 8.0915856 |
| **NFATC4** | 6.4384 | -1.12422610642011 | 0.22 | 0.641 | 1.0470900 |
| **SLC16A9** | 6.9963 | 0.911069708833101 | 0.81 | 0.501 | 1.1378082 |
| **MAN1A1** | 7.0764 | -1.08447404293551 | 0.088 | 0.523 | 1.1508350 |
| **SDC1** | 1.1545 | -1.01581329718493 | 0.08 | 0.521 | 1.8777183 |
| **MB21D2** | 1.1977 | -0.864742404215124 | 0.295 | 0.699 | 1.9479348 |
| **GINS1** | 1.2701 | -0.85123536767873 | 0.992 | 0.996 | 2.0656912 |
| **EZR** | 1.9068 | -1.11145624944399 | 1 | 1 | 3.1011488 |
| **PFKP** | 1.9527 | -0.85292504654169 | 0.935 | 0.994 | 3.1758198 |
| **NRXN1** | 2.1420 | -0.893467963515776 | 0.065 | 0.502 | 3.4836608 |
| **DYRK2** | 2.1535 | -0.811086797243003 | 0.733 | 0.931 | 3.5022453 |
| **ENC1** | 2.8494 | -2.28525667585674 | 0.083 | 0.516 | 4.6341210 |
| **TRIM2** | 3.5723 | -0.826958656590708 | 0.824 | 0.972 | 5.8097780 |

| | | | | | |
|---|---|---|---|---|---|
| **WWTR1** | 3.9655 | 0.815072345084584 | 0.955 | 0.854 | 6.44911119 |
| **DMD** | 5.9141 | -1.04303359852565 | 0.468 | 0.843 | 9.61812378 |
| **BEND4** | 6.2864 | -1.00096681495995 | 0.8 | 0.914 | 1.02236053 |
| **GLI3** | 6.2885 | -2.72501631808905 | 0.03 | 0.455 | 1.02270984 |
| **ARHGEF10L** | 7.4483 | -0.898348775403239 | 0.284 | 0.692 | 1.21131724 |
| **MTM1** | 1.1402 | -0.833739357617389 | 0.788 | 0.956 | 1.85431195 |
| **AGBL5** | 1.9783 | -0.852990712883078 | 0.638 | 0.901 | 3.21736482 |
| **TFPI2** | 2.1243 | 0.977284157163752 | 0.689 | 0.312 | 3.45484923 |
| **KLHL23** | 2.3733 | -0.943753155549194 | 0.597 | 0.875 | 3.85971250 |
| **NLGN4X** | 2.9763 | -1.72553610109451 | 0.137 | 0.56 | 4.84037854 |
| **TMC5** | 3.0277 | -1.22531466586412 | 0.402 | 0.758 | 4.92405901 |
| **RFX7** | 3.5080 | -0.861568926128478 | 0.329 | 0.726 | 5.70510084 |
| **GNAL** | 4.7640 | 0.930037441832433 | 0.806 | 0.504 | 7.74782014 |
| **FARP1** | 6.6662 | -0.945826576746845 | 0.753 | 0.949 | 1.08412483 |
| **PDCD4** | 1.1967 | -0.910794734660899 | 0.537 | 0.842 | 1.94624467 |
| **LPIN2** | 1.3364 | 0.890882945078241 | 0.881 | 0.606 | 2.17352769 |
| **FIBCD1** | 1.4748 | -1.17633151428652 | 0.376 | 0.731 | 2.39850100 |
| **SLC1A5** | 1.5664 | 0.823307914039821 | 0.997 | 0.985 | 2.54748737 |
| **DUSP7** | 1.9246 | 0.909060671971209 | 0.846 | 0.607 | 3.13005684 |
| **CNN1** | 1.9581 | -2.01855648349144 | 0.297 | 0.684 | 3.18457460 |
| **NRN1L** | 2.1334 | -1.2951647828309 | 0.093 | 0.523 | 3.46962234 |
| **PPP1R18** | 3.4153 | -1.14894102738806 | 0.289 | 0.691 | 5.55439400 |
| **ITM2B** | 3.5722 | -0.851530725659716 | 0.994 | 0.998 | 5.80950140 |
| **JADE3** | 4.3751 | -0.803886903462401 | 0.487 | 0.831 | 7.11531130 |
| **C11orf80** | 1.0376 | -0.832417655520133 | 0.47 | 0.835 | 1.68745374 |
| **MAP7D1** | 1.0535 | -1.43282537141027 | 0.54 | 0.808 | 1.71332317 |
| **ZBTB12** | 1.5766 | -0.808822088240888 | 0.053 | 0.479 | 2.56402613 |
| **TACC2** | 1.9520 | -0.803162751906173 | 0.61 | 0.879 | 3.17460113 |
| **LGI1** | 2.5121 | -1.70149613568184 | 0.025 | 0.443 | 4.08547368 |
| **AJAP1** | 5.8490 | -0.947391616205133 | 0.119 | 0.546 | 9.51232254 |
| **PCNX4** | 6.2839 | -0.942935565995768 | 0.523 | 0.815 | 1.02195395 |
| **CNKSR2** | 7.5143 | -0.919734320094461 | 0.079 | 0.507 | 1.22206345 |
| **DOCK3** | 1.1378 | -0.887661124128466 | 0.09 | 0.52 | 1.85052899 |
| **RGS3** | 1.5975 | -0.989494846246219 | 0.047 | 0.471 | 2.59808709 |

| | | | | | |
|---|---|---|---|---|---|
| **PRKG1** | 1.9453 | -1.00070461297659 | 0.025 | 0.442 | 3.1637319 |
| **NME7** | 2.1951 | -0.813087644537115 | 0.253 | 0.676 | 3.5699215 |
| **UBE2J1** | 2.3086 | -0.911105793681455 | 0.764 | 0.928 | 3.7544846 |
| **SMAD6** | 2.4299 | -0.858070370916132 | 0.194 | 0.616 | 3.9519007 |
| **RAB3B** | 3.5412 | -0.902961258138223 | 0.977 | 0.983 | 5.7591327 |
| **ANKRD33B** | 4.6369 | -1.21614930838033 | 0.105 | 0.531 | 7.5409984 |
| **RAD51** | 7.1671 | 0.805875764118599 | 0.979 | 0.878 | 1.1655855 |
| **ELF1** | 7.9146 | 0.871974227381729 | 0.894 | 0.685 | 1.2871589 |
| **AMH** | 9.8220 | -1.16344945644281 | 0.136 | 0.559 | 1.5973550 |
| **SIK2** | 1.5373 | -0.822444907105792 | 0.419 | 0.788 | 2.5001753 |
| **TSPYL2** | 2.0726 | -0.897801621746765 | 0.41 | 0.771 | 3.3706749 |
| **ARMCX3** | 2.8300 | -0.958476850445706 | 0.483 | 0.827 | 4.6024504 |
| **LHX5** | 2.9734 | -1.76901662762985 | 0.006 | 0.412 | 4.8357363 |
| **SLC5A3** | 4.1042 | -1.09852102614825 | 0.585 | 0.857 | 6.6747026 |
| **OSBPL6** | 5.1299 | -0.848207380702364 | 0.128 | 0.553 | 8.3427979 |
| **SEMA3B** | 5.7917 | -0.837590344323932 | 0.094 | 0.519 | 9.4191681 |
| **CDK14** | 6.9919 | -1.17355670893818 | 0.265 | 0.662 | 1.1371046 |
| **TCF7L1** | 7.6094 | -1.45103693108017 | 0.432 | 0.741 | 1.2375172 |
| **ZFP36L1** | 7.9892 | -0.817624348756323 | 0.992 | 0.999 | 1.2992878 |
| **SRSF8** | 1.8674 | 1.09742518733728 | 0.956 | 0.819 | 3.0370049 |
| **CHML** | 1.8726 | -0.823827776764851 | 0.637 | 0.882 | 3.0455702 |
| **SRXN1** | 2.0214 | -0.871578018801273 | 0.671 | 0.91 | 3.2874323 |
| **STK40** | 2.1496 | -0.839578696813831 | 0.624 | 0.877 | 3.4960206 |
| **RASSF9** | 3.4836 | -1.07681042803693 | 0.037 | 0.449 | 5.6654383 |
| **PCDHB16** | 6.0991 | -0.809368632887821 | 0.075 | 0.492 | 9.9191259 |
| **ACADSB** | 6.3425 | -0.857238234531812 | 0.273 | 0.676 | 1.0314922 |
| **SLC39A6** | 6.4177 | -0.917576755640576 | 0.549 | 0.829 | 1.0437140 |
| **RIT1** | 6.8765 | -0.830327807355635 | 0.465 | 0.788 | 1.1183332 |
| **ST3GAL3** | 8.0982 | -0.831088843898411 | 0.094 | 0.516 | 1.3170230 |
| **SIAH2** | 1.1772 | -0.88128381501018 | 0.151 | 0.581 | 1.9145424 |
| **ZNF512** | 2.2029 | -0.865511266926769 | 0.525 | 0.815 | 3.5826431 |
| **CNKSR3** | 5.6092 | -0.942964834484733 | 0.07 | 0.484 | 9.1222665 |
| **CCNG1** | 6.7121 | -0.821722889432696 | 0.987 | 0.993 | 1.0916000 |
| **IGFBP5** | 8.5310 | -3.98887737762587 | 0.124 | 0.532 | 1.3874071 |

| | | | | | |
|---|---|---|---|---|---|
| **SMIM10L2A** | 9.1746 | -1.240990081964 | 0.371 | 0.73 | 1.4920722 |
| **H2AFY** | 1.3025 | -0.912670157282301 | 0.128 | 0.543 | 2.1183214 |
| **PRRG1** | 1.6942 | -0.831842861022454 | 0.205 | 0.62 | 2.7553030 |
| **IKBIP** | 8.0492 | -0.83401743171281 | 0.434 | 0.765 | 1.3090485 |
| **ARHGAP6** | 9.2897 | -0.95558789719479 | 0.185 | 0.601 | 1.5107988 |
| **RASL10B** | 3.1879 | -0.945284448331609 | 0.083 | 0.491 | 5.1846319 |
| **BRSK1** | 3.3662 | -0.935447883585645 | 0.16 | 0.568 | 5.4745097 |
| **CIZ1** | 3.8548 | -0.957842191041631 | 0.46 | 0.765 | 6.2692069 |
| **HPGD** | 1.7734 | -2.22691142954424 | 0.926 | 0.907 | 2.8841869 |
| **IFI6** | 2.5202 | -1.03558787310868 | 0.133 | 0.543 | 4.0987250 |
| **DTNA** | 2.9078 | -1.36322014011095 | 0.041 | 0.443 | 4.7290703 |
| **SSBP2** | 4.0161 | -0.830392032467359 | 0.501 | 0.821 | 6.5315063 |
| **CLCN5** | 6.6350 | -0.887171532984122 | 0.556 | 0.87 | 1.0790642 |
| **MYO10** | 9.5575 | -0.882106681000855 | 0.855 | 0.971 | 1.5543447 |
| **RBCK1** | 1.2604 | -0.817974423455498 | 0.627 | 0.886 | 2.0498270 |
| **SYNGAP1** | 1.6711 | -1.22799862475525 | 0.231 | 0.613 | 2.7177126 |
| **LHFPL2** | 2.3046 | -0.856658815977525 | 0.213 | 0.622 | 3.7479849 |
| **SP8** | 6.0567 | -0.909547974985821 | 0.891 | 0.945 | 9.8500395 |
| **UNC119B** | 1.0122 | -0.855080529824671 | 0.641 | 0.874 | 1.6462960 |
| **PMEL** | 1.2165 | -1.13461390503478 | 0.035 | 0.427 | 1.9784500 |
| **SPRY4** | 2.3509 | -0.982948804871326 | 0.036 | 0.428 | 3.8232961 |
| **CCDC14** | 2.9511 | -0.805473683457631 | 0.859 | 0.967 | 4.7995239 |
| **GNPAT** | 1.2700 | -0.864251019169121 | 0.596 | 0.847 | 2.0654945 |
| **P3H1** | 1.3344 | -0.870459772933996 | 0.425 | 0.744 | 2.1701420 |
| **RCBTB2** | 1.3568 | -0.859919047563972 | 0.227 | 0.624 | 2.2065666 |
| **RTN4RL1** | 1.4344 | -0.951175427424404 | 0.354 | 0.705 | 2.3328414 |
| **GALNT2** | 1.6591 | -0.86221983891941 | 0.907 | 0.98 | 2.6983182 |
| **HNMT** | 5.7324 | -0.818350513284504 | 0.178 | 0.572 | 9.3227040 |
| **PLEKHG2** | 8.5807 | -0.851753351577375 | 0.331 | 0.692 | 1.3954916 |
| **USP44** | 1.6026 | -0.882466794418076 | 0.154 | 0.559 | 2.6063331 |
| **NTS** | 5.1310 | -1.44597987799888 | 0.134 | 0.529 | 8.3446313 |
| **PANX1** | 2.1190 | -0.805045351520359 | 0.53 | 0.818 | 3.4461989 |
| **TBX22** | 2.2391 | -1.08212847335323 | 0.027 | 0.408 | 3.6416023 |
| **PTGES3L** | 2.5415 | -0.817275179976551 | 0.195 | 0.59 | 4.1333402 |

| | | | | | |
|---|---|---|---|---|---|
| **OTX2** | 2.6849 | -1.25678577035268 | 0.796 | 0.918 | 4.3665816 |
| **ZFP36L2** | 2.8976 | -0.87755177337579 | 0.956 | 0.99 | 4.7124086 |
| **PTN** | 2.9346 | -1.4171652658018 | 0.016 | 0.392 | 4.7725491 |
| **LAMB2** | 8.5150 | -1.03195242297285 | 0.779 | 0.931 | 1.3848092 |
| **AHDC1** | 2.5799 | -1.29021923516002 | 0.494 | 0.77 | 4.1957294 |
| **NUP210** | 2.7750 | -0.841202016781024 | 0.509 | 0.808 | 4.5131218 |
| **TANC1** | 4.1015 | -0.842992370656666 | 0.304 | 0.68 | 6.6703364 |
| **PLA2G2A** | 4.1778 | -0.900935872470897 | 0.017 | 0.391 | 6.7943859 |
| **GRPR** | 7.2193 | -1.02615132723156 | 0.01 | 0.379 | 1.1740869 |
| **SH3GL3** | 9.7211 | -0.829944040899459 | 0.181 | 0.571 | 1.5809461 |
| **AHNAK** | 3.1214 | -1.1660523613967 | 0.939 | 0.985 | 5.0763419 |
| **MDM2** | 4.1734 | 0.853560597243945 | 0.994 | 0.975 | 6.7872867 |
| **FNDC4** | 8.3302 | 0.911732849483175 | 0.846 | 0.59 | 1.3547408 |
| **MCAM** | 2.5853 | -1.18319793656063 | 0.558 | 0.82 | 4.2045729 |
| **TAF4B** | 3.6349 | 0.938162870360248 | 0.729 | 0.43 | 5.9114943 |
| **BEX1** | 5.9957 | -0.820918203029659 | 0.027 | 0.396 | 9.7508489 |
| **TYRO3** | 7.1688 | -0.838218981795036 | 0.793 | 0.947 | 1.1658677 |
| **RFX3** | 1.0983 | -0.82173013189586 | 0.323 | 0.697 | 1.7862253 |
| **THY1** | 2.3993 | -1.05685731753873 | 0.05 | 0.422 | 3.9020393 |
| **SLC2A3** | 4.2040 | -0.983981738996801 | 0.457 | 0.766 | 6.8371095 |
| **CDON** | 5.6688 | -1.07896988173479 | 0.266 | 0.631 | 9.2192222 |
| **PRICKLE1** | 6.5088 | -1.04773720587979 | 0.537 | 0.81 | 1.0585403 |
| **COTL1** | 6.9218 | -0.909235414374885 | 0.955 | 0.981 | 1.1257083 |
| **HABP4** | 9.6654 | -0.889347936800741 | 0.306 | 0.667 | 1.5718941 |
| **XKR6** | 3.1951 | -0.826256483053702 | 0.661 | 0.896 | 5.1963434 |
| **CCDC92** | 4.2083 | -1.07523022571707 | 0.081 | 0.458 | 6.8441014 |
| **ANGPTL2** | 4.3077 | -0.920624706642348 | 0.034 | 0.397 | 7.0057009 |
| **LRP1** | 4.8103 | -1.09582437526767 | 0.817 | 0.928 | 7.8231123 |
| **HSPA1A** | 9.5401 | 0.847271450426424 | 0.549 | 0.214 | 1.5515206 |
| **SMAGP** | 1.6666 | -1.0831921812153 | 0.394 | 0.709 | 2.7104596 |
| **DPYSL5** | 2.4545 | -1.96937005971037 | 0.027 | 0.386 | 3.9919100 |
| **AMOT** | 4.0252 | -1.18901942361302 | 0.102 | 0.475 | 6.5463381 |
| **PTPN13** | 4.6263 | -1.0424275623117 | 0.266 | 0.624 | 7.5238868 |
| **PTGIS** | 4.6472 | -1.18904155589541 | 0.087 | 0.454 | 7.5577774 |

| | | | | | |
|---|---|---|---|---|---|
| **ARPIN** | 5.4055 | -0.802179347311682 | 0.433 | 0.748 | 8.79101089 |
| **SLC7A3** | 8.1190 | -0.925558972739696 | 0.243 | 0.622 | 1.32040225 |
| **DDR1** | 9.0444 | -0.933596378381602 | 0.904 | 0.977 | 1.47090633 |
| **SPTLC3** | 1.0172 | -0.813844870899283 | 0.017 | 0.371 | 1.65428644 |
| **WDR27** | 3.1629 | -0.889413604717111 | 0.486 | 0.789 | 5.14390115 |
| **KCNMB1** | 5.0510 | -0.95644363401326 | 0.037 | 0.397 | 8.21449521 |
| **DMPK** | 5.6811 | -0.821335540654176 | 0.341 | 0.694 | 9.23929527 |
| **RNF24** | 2.1610 | -0.861121311835573 | 0.908 | 0.948 | 3.51453438 |
| **RARA** | 3.8943 | -0.820738207903978 | 0.261 | 0.62 | 6.33330482 |
| **LPAR1** | 2.0491 | -0.873103119557657 | 0.083 | 0.45 | 3.33246762 |
| **COMT** | 2.8158 | -0.87265464488143 | 0.481 | 0.781 | 4.57946659 |
| **STOX2** | 4.0570 | -0.869574388830862 | 0.386 | 0.707 | 6.59793241 |
| **THBS2** | 5.8527 | -0.945202013102909 | 0.066 | 0.427 | 9.51831317 |
| **LRP8** | 6.3021 | -0.870043744231533 | 0.596 | 0.84 | 1.02492486 |
| **GPC1** | 9.5894 | -0.826712605651172 | 0.899 | 0.976 | 1.55953013 |
| **SOX2** | 1.6093 | -1.08766979537493 | 0.96 | 0.959 | 2.61734508 |
| **SYNGR1** | 2.2847 | -0.917810561857483 | 0.354 | 0.676 | 3.71564919 |
| **SVIL** | 3.0067 | -0.818775023897344 | 0.403 | 0.722 | 4.88986785 |
| **RIC3** | 4.7978 | -0.935392659752038 | 0.266 | 0.62 | 7.80271646 |
| **TLE4** | 1.5301 | -1.55432787413421 | 0.484 | 0.758 | 2.48853559 |
| **PRPS2** | 3.1296 | -0.907333699472032 | 0.866 | 0.943 | 5.08973923 |
| **ARVCF** | 6.5602 | -0.835514405727224 | 0.432 | 0.728 | 1.06689525 |
| **TSPEAR** | 5.7612 | -0.851897052760926 | 0.016 | 0.349 | 9.36955727 |
| **PDLIM7** | 8.7677 | -0.970070841744693 | 0.995 | 0.997 | 1.42590382 |
| **ID1** | 1.0262 | -0.897489294395073 | 0.982 | 0.982 | 1.66906220 |
| **C1orf21** | 1.3033 | -1.00776258106207 | 0.041 | 0.385 | 2.11956806 |
| **S100A4** | 1.3353 | -0.920388744678346 | 0.019 | 0.353 | 2.17171031 |
| **MYL7** | 7.9637 | -1.54291559721843 | 0.101 | 0.45 | 1.29513807 |
| **PMAIP1** | 1.3288 | -1.11571638280334 | 0.947 | 0.934 | 2.16113392 |
| **MAP2** | 2.5762 | -1.7767993377414 | 0.016 | 0.338 | 4.18973767 |
| **RGS2** | 7.3413 | 0.842632209510501 | 0.782 | 0.535 | 1.19391955 |
| **LMCD1** | 7.7229 | -1.0765916516948 | 0.035 | 0.367 | 1.25597530 |
| **DYRK1B** | 9.4424 | -0.910741328065243 | 0.469 | 0.73 | 1.53562534 |
| **IRS2** | 1.0490 | -0.9501582298242 | 0.394 | 0.693 | 1.70606762 |

| | | | | | |
|---|---|---|---|---|---|
| **IGFBP4** | 1.7548 | -0.88405460310463 | 0.154 | 0.503 | 2.8539494 |
| **CPE** | 1.9326 | -2.67433437814954 | 0.078 | 0.414 | 3.1430740 |
| **PCDH8** | 4.0240 | -0.866714624634375 | 0.094 | 0.437 | 6.5442685 |
| **SLC1A3** | 5.2989 | -0.873903711419517 | 0.256 | 0.587 | 8.6176236 |
| **GADD45G** | 8.0368 | -1.01673817359296 | 0.212 | 0.551 | 1.3070276 |
| **SYNE3** | 2.9990 | -0.832695803349225 | 0.052 | 0.381 | 4.8773411 |
| **IGFBP2** | 3.2283 | -0.978251363774937 | 0.997 | 1 | 5.2502260 |
| **CD99L2** | 6.3954 | -0.914402707911233 | 0.31 | 0.618 | 1.0400855 |
| **SPON1** | 9.3747 | -1.72403331961805 | 0.032 | 0.353 | 1.5246164 |
| **FLNC** | 2.1386 | -1.57557409742554 | 0.642 | 0.834 | 3.4780236 |
| **P3H2** | 1.5731 | -0.895635549974609 | 0.018 | 0.33 | 2.5584513 |
| **TSC22D3** | 1.7765 | -0.861004232275334 | 0.446 | 0.757 | 2.8891280 |
| **TIMP4** | 1.8781 | -0.88943890111957 | 0.238 | 0.565 | 3.0544011 |
| **EPCAM** | 3.2862 | -1.32475584859208 | 0.994 | 0.95 | 5.3443791 |
| **GREB1L** | 7.8120 | -2.03571906779295 | 0.057 | 0.376 | 1.2704791 |
| **FGF18** | 8.5268 | -0.935718416074764 | 0.253 | 0.576 | 1.3867176 |
| **DUSP4** | 1.5148 | -0.840547764583233 | 0.106 | 0.44 | 2.4636350 |
| **COLEC12** | 2.2029 | -1.92361281640788 | 0.276 | 0.591 | 3.5827021 |
| **NCAPD2** | 2.9936 | -0.898352170284884 | 0.872 | 0.956 | 4.8686286 |
| **CNIH2** | 8.8700 | -0.842215825525257 | 0.606 | 0.823 | 1.4425297 |
| **NEO1** | 2.2990 | -0.812402961101274 | 0.576 | 0.797 | 3.7388954 |
| **FJX1** | 3.1613 | -1.76416059223222 | 0.136 | 0.457 | 5.1412766 |
| **PMP22** | 9.6820 | -1.15648724372118 | 0.28 | 0.585 | 1.5745972 |
| **FAM111B** | 2.5256 | -1.09317289071862 | 0.422 | 0.69 | 4.1075374 |
| **SEMA3C** | 2.9886 | -0.884976807689255 | 0.004 | 0.295 | 4.8604416 |
| **SUSD3** | 1.1661 | -1.05659190519452 | 0.395 | 0.661 | 1.8965678 |
| **VIT** | 4.2182 | -1.30840514671438 | 0.037 | 0.34 | 6.8601909 |
| **PAX3** | 5.4144 | -1.36821397923062 | 0.01 | 0.3 | 8.8054586 |
| **SDC2** | 1.7729 | -1.55079006855797 | 0.209 | 0.525 | 2.8834035 |
| **HPD** | 1.7972 | -1.13685507221617 | 0.35 | 0.625 | 2.9228013 |
| **PABPC1L** | 2.6299 | -0.968791989102858 | 0.488 | 0.769 | 4.2771372 |
| **CRB2** | 5.9112 | -1.51704390281173 | 0.008 | 0.292 | 9.6135391 |
| **BOC** | 1.2394 | -1.8427602581888 | 0.068 | 0.374 | 2.0156774 |
| **PAMR1** | 3.0607 | -1.46501162811477 | 0.052 | 0.35 | 4.9777202 |

| | | | | | |
|---|---|---|---|---|---|
| **NAV1** | 4.5257 | -0.887152813861227 | 0.536 | 0.755 | 7.3602500 |
| **EBF2** | 5.7509 | -1.05937286331716 | 0.186 | 0.489 | 9.3527794 |
| **FOSL2** | 9.9908 | -0.977029477350361 | 0.054 | 0.355 | 1.6248137 |
| **SYNPO** | 1.1544 | -0.936287822938242 | 0.013 | 0.295 | 1.8774034 |
| **TENM4** | 2.1222 | -1.01562467220645 | 0.478 | 0.764 | 3.4514756 |
| **TPBG** | 3.3387 | -1.4114599037317 | 0.358 | 0.641 | 5.4297783 |
| **DNMT3B** | 3.7277 | -0.835168186420211 | 0.996 | 0.993 | 6.0624791 |
| **RAI1** | 3.8756 | -0.804934570261407 | 0.132 | 0.441 | 6.3029366 |
| **OGT** | 7.4049 | -0.812350175380765 | 0.961 | 0.987 | 1.2042682 |
| **PNPLA3** | 1.3267 | -0.847244859378057 | 0.336 | 0.617 | 2.1577065 |
| **BRINP1** | 3.1250 | -0.929831541548046 | 0.27 | 0.55 | 5.0822537 |
| **LIX1** | 2.4724 | -1.29237700941852 | 0.001 | 0.259 | 4.0209512 |
| **SDK2** | 4.9416 | -1.22770133272151 | 0.151 | 0.439 | 8.0366287 |
| **ANKRD1** | 3.5340 | -1.45111664138492 | 0.022 | 0.288 | 5.7474763 |
| **CER1** | 5.1425 | -1.44036774783668 | 0.043 | 0.314 | 8.3633894 |
| **POMC** | 5.2096 | -1.50052765043121 | 0.689 | 0.792 | 8.4725193 |
| **VAMP8** | 5.5193 | -1.02593283708776 | 0.996 | 0.988 | 8.9761242 |
| **FZD10** | 1.8578 | -1.00320793039924 | 0.472 | 0.715 | 3.0213427 |
| **SAMD4A** | 1.9465 | -0.811938407942905 | 0.041 | 0.309 | 3.1657475 |
| **CTIF** | 2.9161 | -0.844238254734916 | 0.342 | 0.604 | 4.7424712 |
| **COL5A1** | 6.3976 | -1.61111587100905 | 0.443 | 0.705 | 1.0404478 |
| **RGMB** | 6.5907 | -0.809061244008007 | 0.624 | 0.802 | 1.0718547 |
| **ANOS1** | 1.6173 | -1.3921912082613 | 0.097 | 0.366 | 2.6302715 |
| **CAVIN1** | 1.9344 | -1.15670475379041 | 0.696 | 0.822 | 3.1460084 |
| **EFNA1** | 2.7362 | -0.980368497958363 | 0.452 | 0.695 | 4.4500305 |
| **CYP26A1** | 4.4374 | -1.30420732560127 | 0.047 | 0.307 | 7.2165706 |
| **WFDC2** | 1.3798 | -0.878335394928229 | 0.248 | 0.533 | 2.2439952 |
| **MYL4** | 1.3703 | -1.99614395926659 | 0.021 | 0.263 | 2.2286770 |
| **WNT4** | 1.3740 | -2.0567927635486 | 0.013 | 0.25 | 2.2346886 |
| **HIST1H2BA** | 1.6690 | 0.983462719842535 | 0.19 | 0.03 | 2.7144224 |
| **DLK1** | 1.7567 | -1.25336236659524 | 0.69 | 0.341 | 2.8569931 |
| **CDH3** | 2.0368 | -0.924303127895325 | 0.941 | 0.906 | 3.3125889 |
| **ZNF367** | 2.7290 | -0.867585871975169 | 0.34 | 0.584 | 4.4383207 |
| **ZNF519** | 6.2037 | -0.800044319574339 | 0.517 | 0.735 | 1.0089095 |

| | | | | | |
|---|---|---|---|---|---|
| **FBN1** | 6.4500 | -0.93251322678442 | 0.132 | 0.397 | 1.0489704 |
| **SDC3** | 4.0017 | -0.882812665628132 | 0.641 | 0.806 | 6.5079751 |
| **TFPI** | 9.9671 | -1.39326509208671 | 0.395 | 0.632 | 1.6209500 |
| **ZNF703** | 2.0376 | -0.878690444779468 | 0.655 | 0.809 | 3.3138208 |
| **CXCL14** | 3.3185 | -1.6064099447692 | 0.475 | 0.678 | 5.3969846 |
| **CSRP1** | 1.5262 | -1.49269141903476 | 0.95 | 0.96 | 2.4821844 |
| **HMGCS1** | 4.8677 | -0.887490375106351 | 0.941 | 0.968 | 7.9164625 |
| **MAL2** | 8.4643 | -0.94597051860778 | 0.461 | 0.644 | 1.3765562 |
| **SPATS2L** | 1.6292 | -1.0806918832729 | 0.203 | 0.452 | 2.6495958 |
| **LMO1** | 4.7797 | -0.997379680632262 | 0.326 | 0.557 | 7.7732271 |
| **ACTG2** | 1.6903 | -1.35558452035149 | 0.778 | 0.857 | 2.7490704 |
| **WNT1** | 2.8459 | -2.33748901238676 | 0.004 | 0.2 | 4.6283563 |
| **CDH6** | 2.6282 | -1.62040125152134 | 0.013 | 0.211 | 4.2743424 |
| **ZNF521** | 8.2441 | -0.844091157767403 | 0.017 | 0.215 | 1.3407456 |
| **MYOF** | 3.0008 | -0.882661068746966 | 0.101 | 0.329 | 4.8802398 |
| **GAP43** | 1.2861 | -0.998909515730972 | 0.195 | 0.437 | 2.0916483 |
| **TC2N** | 1.7718 | 1.09484184726621 | 0.884 | 0.731 | 2.8815915 |
| **TBXT** | 1.3123 | -0.897037016886531 | 0.019 | 0.211 | 2.1343516 |
| **GREM1** | 8.1648 | -0.979431697877862 | 0 | 0.168 | 1.3278490 |
| **SNAI2** | 8.4008 | -0.962398070895601 | 0.014 | 0.194 | 1.3662309 |
| **MSX1** | 3.2099 | -0.838202823256053 | 0.013 | 0.189 | 5.2203982 |
| **CYP1B1** | 5.2289 | -1.04361123122062 | 0.212 | 0.42 | 8.5037919 |
| **NID2** | 1.9666 | -0.846422280174853 | 0.074 | 0.262 | 3.1983644 |
| **PCSK1** | 2.9565 | -0.931967594159459 | 0.574 | 0.714 | 4.8082861 |
| **IER3** | 4.4034 | -1.29661700635817 | 0.132 | 0.335 | 7.1612782 |
| **PCBP4** | 2.7042 | -1.0065625973097 | 0.256 | 0.442 | 4.3979805 |
| **CNTNAP2** | 9.8018 | -1.5994898033048 | 0.69 | 0.778 | 1.5940807 |
| **ACTA2** | 2.3989 | -1.91810701782986 | 0.009 | 0.15 | 3.9014749 |
| **ADAMTSL4** | 2.7679 | -0.948939932555293 | 0.437 | 0.575 | 4.5015656 |
| **OLIG3** | 8.0538 | -0.850602411352947 | 0.109 | 0.284 | 1.3097966 |
| **MN1** | 1.3655 | -0.839100506066554 | 0.518 | 0.643 | 2.2207434 |
| **LOXL1** | 9.7437 | -1.08642933915567 | 0.667 | 0.771 | 1.5846211 |
| **NPPB** | 4.8217 | -1.12436450232844 | 0.039 | 0.168 | 7.8416769 |
| **PCK2** | 1.6307 | 0.94443007893407 | 0.695 | 0.653 | 2.6520154 |

| | | | | | |
|---|---|---|---|---|---|
| **BGN** | 3.4311 | -0.956828714175772 | 0.275 | 0.439 | 5.58010165 |
| **NR2F1** | 7.2362 | -0.892568166189734 | 0.026 | 0.137 | 1.17682819 |
| **FGF8** | 3.3994 | -0.890748195063803 | 0.615 | 0.679 | 5.52848197 |
| **ACTA1** | 1.1494 | -0.83873319335242 | 0.171 | 0.312 | 1.86934575 |
| **COL3A1** | 2.1822 | -1.94836969712525 | 0.053 | 0.16 | 3.54899449 |
| **ANXA3** | 2.9347 | -0.826028188585222 | 0.965 | 0.918 | 4.77285482 |
| **TKTL1** | 4.4497 | 0.853664874652464 | 0.132 | 0.055 | 7.23659007 |
| **NRP2** | 3.3189 | -0.897356755042344 | 0.388 | 0.454 | 0.00053975 |
| **PRSS23** | 4.5678 | -1.72721462282178 | 0.629 | 0.699 | 0.00074286 |
| **ANXA1** | 4.5889 | -1.60522435059181 | 0.146 | 0.213 | 0.07463080 |
| **CNTFR** | 8.2363 | -0.984373725215703 | 0.279 | 0.323 | 0.13394842 |
| **COL1A1** | 0.0002 | -2.1009037850767 | 0.915 | 0.914 | 1 |
| **SLC34A2** | 0.1061 | 0.806487315495383 | 0.103 | 0.086 | 1 |
| **CDH11** | 0.3909 | -0.819020942209879 | 0.189 | 0.167 | 1 |
| **SFRP1** | 0.9554 | -1.61942197146453 | 0.668 | 0.556 | 1 |

Supplemental Table S2_Sheet_6

| | p_val | avg_log2FC | pct.1 | pct.2 | p_val_adj |
|---|---|---|---|---|---|
| **CNR2** | 0 | 2.11143533058203 | 0.788 | 0.015 | 0 |
| **HPDL** | 0 | 2.14968067099385 | 0.902 | 0.134 | 0 |
| **BRDT** | 0 | 3.07074430830557 | 0.97 | 0.176 | 0 |
| **HENMT1** | 0 | 1.69420514393125 | 0.766 | 0.033 | 0 |
| **TAF13** | 0 | 3.19623308284294 | 1 | 0.79 | 0 |
| **HORMAD1** | 0 | 2.95709560953195 | 0.971 | 0.094 | 0 |
| **DUSP23** | 0 | 1.55465460773352 | 0.799 | 0.031 | 0 |
| **MAEL** | 0 | 1.64928020036774 | 0.636 | 0.016 | 0 |
| **IAH1** | 0 | 1.71957782504395 | 0.817 | 0.029 | 0 |
| **CRYGD** | 0 | 1.3094898498046 | 0.667 | 0.019 | 0 |
| **ALPG** | 0 | 3.80332071455792 | 0.98 | 0.407 | 0 |
| **ZNF502** | 0 | 1.22116372440523 | 0.668 | 0.008 | 0 |
| **DPPA2** | 0 | 3.70871607090026 | 0.993 | 0.426 | 0 |
| **HCLS1** | 0 | 1.79278802118846 | 0.797 | 0.093 | 0 |
| **MGLL** | 0 | 1.90353796513465 | 0.738 | 0.042 | 0 |
| **ZNF732** | 0 | 1.55291864343074 | 0.739 | 0.003 | 0 |
| **S100P** | 0 | 2.14294999961276 | 0.655 | 0.025 | 0 |
| **TRIM60** | 0 | 2.50444151256925 | 0.931 | 0.041 | 0 |
| **TRIM75P** | 0 | 1.86102682877455 | 0.748 | 0.023 | 0 |
| **ZFP42** | 0 | 4.6040807540382 | 0.995 | 0.307 | 0 |
| **SUB1** | 0 | 2.35546948555139 | 0.997 | 0.39 | 0 |
| **HIST1H1A** | 0 | 5.22651655663377 | 0.968 | 0.396 | 0 |
| **ZFP57** | 0 | 2.93529755481497 | 0.986 | 0.274 | 0 |
| **DPPA5** | 0 | 5.70881559168377 | 0.993 | 0.547 | 0 |
| **KHDC3L** | 0 | 5.27579324634107 | 0.993 | 0.395 | 0 |
| **DDX43** | 0 | 2.23523698278863 | 0.86 | 0.017 | 0 |
| **CGAS** | 0 | 1.98674790961643 | 0.78 | 0.045 | 0 |
| **IRAK1BP1** | 0 | 1.17054673281099 | 0.635 | 0.009 | 0 |
| **FAM162B** | 0 | 1.17759016659592 | 0.643 | 0.023 | 0 |
| **TCP1** | 0 | 2.40592681817107 | 1 | 0.948 | 0 |
| **CYCS** | 0 | 2.64615375591454 | 1 | 0.641 | 0 |
| **SUN3** | 0 | 1.1960727434424 | 0.611 | 0.008 | 0 |

| | | | | | |
|---|---|---|---|---|---|
| **ZNF727** | 0 | 1.58226676661531 | 0.687 | 0.003 | 0 |
| **ZNF736** | 0 | 1.34086019995543 | 0.77 | 0.004 | 0 |
| **HYAL4** | 0 | 1.64351112771285 | 0.763 | 0.02 | 0 |
| **FABP5** | 0 | 3.83794584684268 | 0.993 | 0.517 | 0 |
| **PCSK5** | 0 | 2.07579745472835 | 0.919 | 0.189 | 0 |
| **C9orf64** | 0 | 1.14362897969485 | 0.673 | 0.012 | 0 |
| **PIK3AP1** | 0 | 1.45451450207046 | 0.745 | 0.051 | 0 |
| **C10orf95** | 0 | 1.76753562088457 | 0.723 | 0.045 | 0 |
| **ATP5MD** | 0 | 2.37295977177789 | 0.992 | 0.322 | 0 |
| **UTF1** | 0 | 4.0668631156608 | 0.97 | 0.361 | 0 |
| **SNX19** | 0 | 1.13236135693605 | 0.651 | 0.029 | 0 |
| **DPPA3** | 0 | 3.9196190212852 | 0.963 | 0.207 | 0 |
| **REC8** | 0 | 2.73943990485373 | 0.976 | 0.209 | 0 |
| **GMFB** | 0 | 2.08493101052408 | 0.946 | 0.098 | 0 |
| **TMEM30B** | 0 | 1.52099681491515 | 0.76 | 0.013 | 0 |
| **HSPA2** | 0 | 2.36690235489509 | 0.897 | 0.121 | 0 |
| **GPX2** | 0 | 2.53163763795587 | 0.802 | 0.045 | 0 |
| **RSL24D1** | 0 | 1.98834660284935 | 0.958 | 0.138 | 0 |
| **DNAJA4** | 0 | 1.64367393004227 | 0.799 | 0.033 | 0 |
| **CACNA1H** | 0 | 1.29425224968755 | 0.736 | 0.052 | 0 |
| **MT1E** | 0 | 2.69897350499349 | 0.966 | 0.148 | 0 |
| **MT1G** | 0 | 3.95321017424557 | 0.983 | 0.268 | 0 |
| **MT1H** | 0 | 4.77904159031344 | 0.992 | 0.362 | 0 |
| **ZFP3** | 0 | 1.3278369771394 | 0.694 | 0.011 | 0 |
| **NTN1** | 0 | 5.21911478446024 | 1 | 0.986 | 0 |
| **ZNF57** | 0 | 1.15243673975102 | 0.667 | 0.012 | 0 |
| **TNFSF9** | 0 | 2.20878739915065 | 0.816 | 0.07 | 0 |
| **CD70** | 0 | 1.91288146483779 | 0.778 | 0.089 | 0 |
| **ZNF560** | 0 | 1.24255689243177 | 0.731 | 0.005 | 0 |
| **ZNF844** | 0 | 1.72903789816705 | 0.77 | 0.007 | 0 |
| **ZNF253** | 0 | 1.6468534629732 | 0.822 | 0.013 | 0 |
| **ZNF626** | 0 | 1.27446497163438 | 0.756 | 0.021 | 0 |
| **ZNF257** | 0 | 1.05986398397085 | 0.635 | 0.006 | 0 |
| **ZNF676** | 0 | 3.29887281212251 | 0.885 | 0.129 | 0 |

| | | | | | |
|---|---|---|---|---|---|
| **ZNF729** | 0 | 2.69030082368721 | 0.694 | 0.015 | 0 |
| **ZNF98** | 0 | 1.58848516646681 | 0.79 | 0.004 | 0 |
| **ZNF492** | 0 | 2.57992858747296 | 0.934 | 0.009 | 0 |
| **ZNF723** | 0 | 1.05853597790454 | 0.572 | 0.001 | 0 |
| **ZNF728** | 0 | 1.21780973909183 | 0.663 | 0.005 | 0 |
| **ZNF568** | 0 | 1.75921532713105 | 0.858 | 0.086 | 0 |
| **C19orf33** | 0 | 2.25366573867824 | 0.958 | 0.233 | 0 |
| **FCGBP** | 0 | 2.93869920264253 | 0.844 | 0.046 | 0 |
| **ATP1A3** | 0 | 1.61568123787471 | 0.812 | 0.083 | 0 |
| **LIM2** | 0 | 1.74316610106692 | 0.8 | 0.022 | 0 |
| **ZNF677** | 0 | 1.85926804078503 | 0.89 | 0.018 | 0 |
| **NLRP7** | 0 | 5.4364405310301 | 1 | 0.58 | 0 |
| **C19orf85** | 0 | 2.43010098286221 | 0.78 | 0.074 | 0 |
| **PEG3** | 0 | 2.51791941587241 | 0.745 | 0.026 | 0 |
| **HNF4A** | 0 | 1.23469336725137 | 0.623 | 0.019 | 0 |
| **SRMS** | 0 | 1.51711216651709 | 0.685 | 0.022 | 0 |
| **EIF2S3** | 0 | 2.40428038227047 | 0.988 | 0.372 | 0 |
| **MAGEB2** | 0 | 1.87690009235726 | 0.675 | 0.026 | 0 |
| **SLC25A5** | 0 | 1.71647678086291 | 0.882 | 0.101 | 0 |
| **CNIH4** | 7.5341 | 2.33340527664631 | 1 | 0.692 | 1.2338614 |
| **PRDX1** | 8.5836 | 2.29421344664095 | 1 | 0.999 | 1.4057494 |
| **CBX3** | 1.0774 | 2.28602811627402 | 1 | 0.94 | 1.7645768 |
| **NLRP2** | 1.1812 | 3.19282037819268 | 1 | 0.99 | 1.9345480 |
| **CRIP1** | 1.3402 | 3.6368813137536 | 1 | 0.817 | 2.1949695 |
| **GALNT3** | 6.2564 | 1.10913710667112 | 0.569 | 0.008 | 1.0246241 |
| **BRK1** | 1.0026 | 1.21144010549666 | 0.756 | 0.077 | 1.6421150 |
| **BEX3** | 1.7004 | -2.7139042379921 | 0.902 | 1 | 2.7848117 |
| **EIF1** | 6.0917 | 2.18923428074414 | 1 | 0.922 | 9.9764206 |
| **SUSD2** | 7.2913 | 3.74212644657935 | 0.992 | 0.641 | 1.1941089 |
| **GTF3C6** | 6.9479 | 2.0291749566671 | 0.997 | 0.659 | 1.1378587 |
| **MARCKS** | 4.4265 | -3.20333732246698 | 0.372 | 0.999 | 7.2493011 |
| **ATG3** | 3.9281 | 2.51475891611648 | 1 | 0.996 | 6.4331350 |
| **H2AFY2** | 9.8807 | -2.92644962158408 | 0.496 | 0.997 | 1.6181762 |
| **DNMT3L** | 1.3475 | 4.7391472631637 | 0.995 | 0.708 | 2.2069057 |

| | | | | | |
|---|---|---|---|---|---|
| **MAGED1** | 3.4236 | -2.39180907547864 | 0.875 | 1 | 5.6068822 |
| **TMSB4X** | 5.0711 | -2.73611574102909 | 1 | 1 | 8.3050642 |
| **UPP1** | 6.5790 | 3.27374068529374 | 1 | 0.891 | 1.0774580 |
| **PLBD1** | 2.2363 | 2.76527775621261 | 1 | 0.896 | 3.6624907 |
| **NONO** | 1.7122 | 1.87514800124022 | 1 | 0.901 | 2.8040895 |
| **HSP90B1** | 2.3377 | 2.29678416715761 | 1 | 0.779 | 3.8285615 |
| **NAA50** | 6.8951 | 1.89289494516979 | 0.983 | 0.472 | 1.1292170 |
| **ATAD3B** | 8.8281 | 2.16813560164274 | 1 | 0.917 | 1.4457806 |
| **SLC5A5** | 4.0845 | 1.19854770687929 | 0.699 | 0.056 | 6.6891971 |
| **WLS** | 8.6562 | -4.19458544180491 | 0.452 | 0.995 | 1.4176297 |
| **MAGED2** | 3.9525 | -2.07988035246135 | 0.934 | 1 | 6.4730715 |
| **MATK** | 5.1081 | 1.49278406789761 | 0.758 | 0.086 | 8.3656266 |
| **CRABP2** | 6.0784 | -2.60177070177643 | 0.973 | 1 | 9.9546751 |
| **HIST1H2BB** | 1.7488 | 2.61493559273613 | 0.75 | 0.09 | 2.8641520 |
| **GLDC** | 1.8411 | 2.61685886596044 | 0.993 | 0.647 | 3.0153275 |
| **NOL11** | 2.3548 | 1.79565316920544 | 0.986 | 0.455 | 3.8565821 |
| **CHD7** | 2.5452 | -2.80491130563431 | 0.335 | 0.991 | 4.1683134 |
| **EEF1G** | 6.9046 | 1.18878836729455 | 1 | 1 | 1.1307677 |
| **IRF6** | 6.1124 | 1.02994496524237 | 0.633 | 0.033 | 1.0010347 |
| **TOMM22** | 8.9072 | 1.59206898122647 | 1 | 0.996 | 1.4587373 |
| **OLFML3** | 1.4710 | -3.80059926677664 | 0.699 | 0.997 | 2.4092168 |
| **FZD2** | 4.4353 | -3.01992058656889 | 0.225 | 0.98 | 7.2637307 |
| **PHKA1** | 9.3921 | 2.64000345859 | 0.997 | 0.773 | 1.5381515 |
| **MT1X** | 3.4456 | 3.14979908116186 | 0.998 | 0.979 | 5.6429335 |
| **ZNF667** | 5.2009 | 0.851173939453137 | 0.53 | 0.006 | 8.5175449 |
| **VRTN** | 5.4865 | -2.56843872371835 | 0.736 | 1 | 8.9853916 |
| **COL18A1** | 6.7426 | -3.09165459603528 | 0.804 | 1 | 1.1042399 |
| **IL6R** | 9.2517 | 1.69911051457097 | 0.898 | 0.213 | 1.5151606 |
| **ATP5F1E** | 1.3645 | -1.55248088495881 | 0.998 | 1 | 2.2347132 |
| **CKS1B** | 4.2061 | 1.76099390530689 | 1 | 0.983 | 6.8884402 |
| **SNRPE** | 7.6782 | 1.68422349157765 | 1 | 0.913 | 1.2574627 |
| **PPP1R3B** | 9.5071 | -2.81912331944776 | 0.668 | 0.999 | 1.5569788 |
| **FZD5** | 1.6287 | 2.7067750253863 | 0.998 | 0.867 | 2.6674292 |
| **COX7B** | 3.5306 | 1.98316956046509 | 0.997 | 0.745 | 5.7821877 |

| | | | | | |
|---|---|---|---|---|---|
| **NMI** | 8.5845 | 1.98331490182666 | 0.963 | 0.354 | 1.40588790 |
| **SLIRP** | 3.0587 | 1.52321432157358 | 1 | 1 | 5.00930239 |
| **VSIG10** | 7.8391 | 2.07925071251927 | 0.997 | 0.935 | 1.28381913 |
| **AMOTL1** | 9.4801 | -2.3575926552182 | 0.638 | 1 | 1.55256846 |
| **CPNE1** | 9.7970 | -1.71649486896343 | 0.964 | 1 | 1.60445822 |
| **PCLAF** | 1.2860 | -2.63716429006197 | 0.223 | 0.973 | 2.10611723 |
| **TUBA1C** | 7.4650 | 1.75427912064535 | 1 | 1 | 1.22254710 |
| **CHCHD2** | 1.6228 | 1.3457943236211 | 1 | 1 | 2.65769605 |
| **3,00 LSM** | 2.6291 | 1.24829807175866 | 0.738 | 0.088 | 4.30570965 |
| **SERPINB1** | 8.8833 | 1.209096987615 | 0.648 | 0.044 | 1.45481882 |
| **SYCP3** | 3.8181 | 1.37137847183126 | 0.574 | 0.021 | 6.25303411 |
| **SLC25A16** | 5.8519 | 2.4408984728745 | 0.99 | 0.714 | 9.58372480 |
| **SALL2** | 2.7750 | -2.29647144404828 | 0.839 | 0.998 | 4.54467500 |
| **PANK3** | 1.1963 | 1.69883354298667 | 1 | 0.795 | 1.95920760 |
| **ZNF829** | 3.2437 | 0.811951063217244 | 0.521 | 0.007 | 5.31222151 |
| **DYNLRB1** | 5.5249 | -1.28246191789682 | 1 | 1 | 9.04817623 |
| **ZNF354C** | 3.4068 | 0.815953049264538 | 0.503 | 0.003 | 5.57942581 |
| **ACP5** | 1.2146 | 1.22567355502095 | 0.675 | 0.058 | 1.98922510 |
| **DND1** | 3.9277 | 1.9549909272746 | 0.726 | 0.09 | 6.43253993 |
| **RLIM** | 7.5517 | 1.52998702604509 | 0.775 | 0.121 | 1.23675590 |
| **ZNF90** | 1.0312 | 2.38104720741081 | 0.99 | 0.721 | 1.68892250 |
| **ARF6** | 1.2814 | 1.44767775845979 | 1 | 0.985 | 2.09866477 |
| **NDUFA5** | 1.7995 | 1.71349966141219 | 0.988 | 0.576 | 2.94710713 |
| **LDHB** | 3.1586 | 1.37488637354717 | 1 | 1 | 5.17287535 |
| **MMP2** | 3.4271 | -2.92281939453565 | 0.672 | 0.998 | 5.61267620 |
| **NDUFAB1** | 3.8325 | 1.39532995325408 | 1 | 1 | 6.27657747 |
| **SNRPC** | 7.4719 | 1.15279212788712 | 1 | 1 | 1.22368868 |
| **DENR** | 1.3590 | 1.48294910214234 | 1 | 0.976 | 2.22576279 |
| **ID3** | 2.4196 | -2.88798669724923 | 0.731 | 0.997 | 3.96259702 |
| **SLFN12** | 3.5788 | 1.31728966152098 | 0.709 | 0.076 | 5.86100945 |
| **HSP90AB1** | 4.2986 | 1.17657181936029 | 1 | 1 | 7.03995638 |
| **AKIRIN1** | 3.1263 | 1.44256128686624 | 1 | 0.999 | 5.12002160 |
| **L1TD1** | 3.5197 | 1.66444373781654 | 1 | 0.998 | 5.76425478 |
| **FBP1** | 6.8409 | 1.73152097730869 | 0.667 | 0.063 | 1.12034898 |

| | | | | | |
|---|---|---|---|---|---|
| **CR1L** | 1.4260 | 2.10314754069698 | 0.927 | 0.284 | 2.3354295 |
| **LRRC34** | 3.1309 | 0.823887963070461 | 0.494 | 0.003 | 5.1275343 |
| **ARMT1** | 6.6202 | 1.96348303277302 | 1 | 0.992 | 1.0841940 |
| **GPC6** | 1.3056 | -2.35422981730565 | 0.465 | 0.992 | 2.1382161 |
| **SNRPN** | 1.4854 | 1.87814314112361 | 0.993 | 0.844 | 2.4326636 |
| **RNF125** | 1.5776 | 2.50367413517794 | 0.98 | 0.673 | 2.5836867 |
| **NT5DC2** | 1.8975 | -1.29805232652874 | 1 | 1 | 3.1076991 |
| **FADS2** | 4.0112 | -1.74785683382184 | 0.998 | 1 | 6.5691791 |
| **EIF3I** | 5.1765 | 1.62721368638185 | 0.966 | 0.452 | 8.4776260 |
| **NME4** | 5.9909 | -1.57315977150112 | 0.995 | 1 | 9.8114292 |
| **EID1** | 2.8790 | -1.48909497340613 | 0.97 | 1 | 4.7150607 |
| **SNRPG** | 3.8918 | 1.5544743187813 | 1 | 0.97 | 6.3736238 |
| **MAD2L1** | 2.1783 | 1.91743339615759 | 0.988 | 0.786 | 3.5674485 |
| **A4GALT** | 2.5171 | 2.61653049814661 | 0.983 | 0.666 | 4.1222790 |
| **NDUFB4** | 2.8644 | 1.63796218873367 | 0.956 | 0.413 | 4.6910919 |
| **NOCT** | 4.8767 | 1.82974742893884 | 0.998 | 0.853 | 7.9867295 |
| **COX7A1** | 6.3886 | 0.866604601126312 | 0.496 | 0.005 | 1.0462632 |
| **PKIB** | 6.7134 | 1.43830457594348 | 0.638 | 0.049 | 1.0994629 |
| **SNU13** | 1.4836 | 1.04848015399598 | 1 | 1 | 2.4298072 |
| **HMGA2** | 2.2698 | -1.90194498512009 | 0.809 | 1 | 3.7173662 |
| **AGL** | 2.3304 | 2.37425292806346 | 0.997 | 0.922 | 3.8165180 |
| **MT1F** | 3.0162 | 2.42679376168788 | 0.988 | 0.768 | 4.9397130 |
| **NEFH** | 4.2587 | 2.81054823053633 | 0.931 | 0.371 | 6.9745662 |
| **NCOA4** | 1.2084 | 1.5250041269831 | 1 | 0.931 | 1.9791572 |
| **RGCC** | 1.7027 | 2.26244511197285 | 0.775 | 0.126 | 2.7886408 |
| **SMARCA1** | 1.7982 | -2.08908250449975 | 0.394 | 0.992 | 2.9450021 |
| **ZNF423** | 1.8999 | -1.89868324134522 | 0.636 | 0.995 | 3.1116057 |
| **CCNA1** | 4.2777 | 2.39130969216269 | 0.623 | 0.046 | 7.0056527 |
| **MARCKSL1** | 3.4340 | -1.09535668985344 | 1 | 1 | 5.6238708 |
| **CAPN2** | 3.6617 | -2.39204875880705 | 0.416 | 0.995 | 5.9968118 |
| **SNRPB2** | 4.1763 | 1.55612898562926 | 1 | 0.873 | 6.8395516 |
| **CUTA** | 4.7978 | -1.46898322780395 | 0.98 | 1 | 7.8574904 |
| **UBE2N** | 8.4425 | 1.57445678398422 | 0.98 | 0.542 | 1.3826295 |
| **TIMP1** | 1.6479 | -1.7205633188652 | 0.963 | 1 | 2.6988274 |

| | | | | | |
|---|---|---|---|---|---|
| **TCAF1** | 1.6828 | -1.93606742983805 | 0.536 | 0.989 | 2.7560404 |
| **NSMAF** | 2.2094 | 1.69341943495752 | 1 | 0.907 | 3.6184196 |
| **IDH1** | 4.2165 | 1.66271113321472 | 1 | 0.993 | 6.9054605 |
| **OAT** | 1.8581 | 1.93451397236956 | 0.968 | 0.488 | 3.0430743 |
| **TUBB2B** | 2.3945 | -2.34691629606188 | 0.995 | 1 | 3.9214831 |
| **GDF3** | 3.7717 | 3.18226913615285 | 0.986 | 0.716 | 6.1769561 |
| **ACE** | 6.8788 | 1.75432062670288 | 0.866 | 0.2 | 1.1265464 |
| **GNB4** | 9.7533 | -2.63631221652652 | 0.734 | 0.998 | 1.5973067 |
| **PIGT** | 2.2455 | -1.63927181527882 | 0.87 | 1 | 3.6776164 |
| **SPATA22** | 3.5045 | 0.853473201126577 | 0.492 | 0.007 | 5.7394128 |
| **SAMHD1** | 4.2771 | 2.55165153373573 | 0.978 | 0.611 | 7.0046701 |
| **TOMM6** | 1.4360 | 1.5056512060688 | 1 | 0.957 | 2.3517940 |
| **PGRMC1** | 2.1860 | -1.67770345477428 | 0.975 | 1 | 3.5801375 |
| **LARS** | 2.2739 | 1.20247350370758 | 1 | 1 | 3.7241227 |
| **ACTR6** | 2.5516 | 1.68283872754234 | 0.964 | 0.486 | 4.1787679 |
| **MTA1** | 1.2222 | -1.3593083467089 | 0.976 | 1 | 2.0016979 |
| **WASF1** | 1.6586 | -1.93742646753147 | 0.496 | 0.986 | 2.7164262 |
| **SLC16A10** | 1.7816 | 1.77715789562126 | 0.885 | 0.256 | 2.9177674 |
| **LMO4** | 2.8237 | 1.93241152571103 | 0.998 | 0.949 | 4.6245120 |
| **HYLS1** | 3.0895 | 1.52061324433203 | 0.768 | 0.123 | 5.0597826 |
| **DNAJA2** | 3.5794 | 1.06290238350108 | 1 | 1 | 5.8621451 |
| **HUS1** | 3.8418 | 1.7664839411581 | 1 | 0.969 | 6.2918088 |
| **ILRUN** | 3.8503 | 1.51731936304803 | 1 | 0.994 | 6.3056666 |
| **TLE2** | 1.3338 | 1.28065865610754 | 0.662 | 0.065 | 2.1844247 |
| **ZNF273** | 1.6358 | 0.959617747440964 | 0.618 | 0.046 | 2.6790316 |
| **LCP1** | 3.1546 | 3.22331088368427 | 0.988 | 0.685 | 5.1663830 |
| **MAN1C1** | 5.0651 | 2.32897873747651 | 0.985 | 0.682 | 8.2951706 |
| **SNRPB** | 9.4300 | -0.99522575612977 | 1 | 1 | 1.5443523 |
| **AK4** | 1.7846 | 2.72327646549155 | 0.998 | 0.929 | 2.9227956 |
| **ZNF595** | 3.9804 | 1.96800753829112 | 0.988 | 0.686 | 6.5187123 |
| **XRN2** | 8.0583 | -0.96160715690505 | 1 | 1 | 1.3197214 |
| **HSPA5** | 8.3283 | 2.02672160949413 | 1 | 0.912 | 1.3639390 |
| **CCDC80** | 1.5881 | 2.14052684856602 | 0.985 | 0.614 | 2.6009099 |
| **TFCP2L1** | 3.5041 | 1.98743222051981 | 0.944 | 0.385 | 5.7388051 |

| | | | | | |
|---|---|---|---|---|---|
| **CCNC** | 2.1179 | 1.31078210348859 | 1 | 0.993 | 3.4685436 |
| **FKBP10** | 1.4653 | -1.89311082921683 | 0.738 | 0.997 | 2.3998043 |
| **GSN** | 3.6586 | -2.17443779521502 | 0.255 | 0.956 | 5.9918221 |
| **DCAF4** | 4.2939 | 1.69064411242253 | 0.997 | 0.863 | 7.0322695 |
| **MSI1** | 1.1833 | -1.9209422944581 | 0.228 | 0.944 | 1.9379814 |
| **ASXL1** | 1.9133 | -1.81498676486243 | 0.65 | 0.995 | 3.1334617 |
| **EIF4E** | 2.6968 | 1.40672696929266 | 0.998 | 0.952 | 4.4166846 |
| **ZNF880** | 5.6367 | 1.20388075844175 | 0.773 | 0.127 | 9.2313295 |
| **TMPO** | 1.8808 | 1.23754574826737 | 1 | 0.998 | 3.0802768 |
| **RBP1** | 1.0950 | -2.60367725957299 | 0.614 | 0.994 | 1.7933750 |
| **ZCCHC3** | 1.4013 | -1.73399975808307 | 0.804 | 0.999 | 2.2950154 |
| **BCAS2** | 1.9974 | 1.53896442700003 | 0.976 | 0.544 | 3.2712753 |
| **FRAT2** | 2.7970 | 2.18625362818754 | 0.997 | 0.984 | 4.5807965 |
| **CRYBG1** | 3.3503 | 1.07958690391231 | 0.587 | 0.04 | 5.4869067 |
| **VIM** | 3.9786 | -3.34008765804492 | 0.721 | 0.997 | 6.5157892 |
| **ZYG11A** | 4.0949 | 2.02211878631795 | 0.986 | 0.722 | 6.7062230 |
| **ECHDC3** | 2.7596 | 0.872069723870071 | 0.494 | 0.012 | 4.5194191 |
| **NDUFV1** | 5.5155 | 1.11154038161681 | 1 | 1 | 9.0328576 |
| **PDK1** | 5.7957 | -2.3694957858508 | 0.853 | 0.998 | 9.4916741 |
| **TGIF2** | 5.2820 | -1.62556352265959 | 0.931 | 1 | 8.6504442 |
| **TCEAL9** | 1.0983 | -1.99521411214682 | 0.658 | 0.999 | 1.7987311 |
| **BIN1** | 2.1500 | -1.99863915507729 | 0.523 | 0.989 | 3.5212002 |
| **NSD1** | 4.3110 | 1.43593950904961 | 1 | 0.981 | 7.0602383 |
| **KLF4** | 8.3365 | 2.44117047524259 | 0.914 | 0.353 | 1.3652760 |
| **MYCBP** | 4.3573 | 1.51622183978953 | 0.973 | 0.556 | 7.1360928 |
| **NDUFA6** | 4.5879 | 1.28978060407023 | 1 | 0.981 | 7.5136730 |
| **LIN28B** | 1.0992 | 1.41437524587519 | 1 | 0.995 | 1.8002014 |
| **USP28** | 2.6408 | 1.44679360332875 | 1 | 0.996 | 4.3248854 |
| **TNFRSF8** | 3.1110 | 1.17584167584897 | 0.547 | 0.029 | 5.0950311 |
| **BEX4** | 3.8558 | -1.70971193069984 | 0.844 | 1 | 6.3147705 |
| **F11R** | 5.8569 | 1.36104297300542 | 1 | 0.995 | 9.5918724 |
| **TRIM6** | 5.8690 | 1.48666654139171 | 0.854 | 0.211 | 9.6117383 |
| **NNAT** | 8.3030 | 1.01189184729355 | 0.46 | 0.005 | 1.3597835 |
| **SFT2D1** | 1.7960 | 1.5133739289255 | 0.973 | 0.563 | 2.9414221 |

| | | | | | |
|---|---|---|---|---|---|
| **PLEKHO1** | 2.4122 | -1.94232396179751 | 0.465 | 0.978 | 3.9505081 |
| **SP5** | 3.4008 | -2.42673708166491 | 0.78 | 1 | 5.5696132 |
| **FZD7** | 1.3915 | -2.89402189485717 | 0.362 | 0.96 | 2.2789941 |
| **CDS2** | 2.2240 | -1.4239296323085 | 0.89 | 1 | 3.6422718 |
| **SPSB4** | 2.7411 | -2.13642837157555 | 0.252 | 0.938 | 4.4891995 |
| **DUSP9** | 2.3946 | 1.15683389864037 | 0.65 | 0.069 | 3.9217166 |
| **MYL9** | 2.8166 | -2.89919802394943 | 0.917 | 1 | 4.6127965 |
| **HIGD1A** | 4.0530 | 1.4776138503892 | 0.983 | 0.595 | 6.6376887 |
| **SMS** | 6.2703 | -1.67933182074948 | 0.99 | 1 | 1.0268986 |
| **NOP16** | 8.3460 | 1.32237593729153 | 1 | 0.999 | 1.3668316 |
| **PYGL** | 8.9258 | 1.63461123210013 | 0.998 | 0.975 | 1.4617882 |
| **HNRNPK** | 1.0915 | 1.28728332125204 | 1 | 0.979 | 1.7875649 |
| **FSTL1** | 1.1442 | -2.04370974169327 | 0.849 | 0.999 | 1.8739707 |
| **SALL1** | 1.2507 | -2.52861964722097 | 0.25 | 0.955 | 2.0483806 |
| **OLAH** | 1.2782 | 1.52048530096199 | 0.548 | 0.032 | 2.0933312 |
| **OOEP** | 2.6059 | 0.800874655328183 | 0.459 | 0.006 | 4.2678255 |
| **MEX3A** | 3.1396 | -1.85128156960364 | 0.526 | 0.991 | 5.1417941 |
| **YBX1** | 1.0609 | -0.86482794198078 | 1 | 1 | 1.7375726 |
| **ZHX2** | 1.4537 | -1.9463027562947 | 0.144 | 0.912 | 2.3808258 |
| **NGEF** | 3.5393 | 1.15520013494481 | 0.565 | 0.038 | 5.7964271 |
| **OBSL1** | 6.1164 | -1.90007930130154 | 0.667 | 0.997 | 1.0016933 |
| **DDAH2** | 1.8568 | -1.66264595589534 | 0.81 | 0.997 | 3.0409209 |
| **XPO1** | 3.7101 | 1.23758924349681 | 1 | 0.965 | 6.0761711 |
| **SEPHS2** | 4.9774 | 1.61094383785147 | 0.998 | 0.989 | 8.1515624 |
| **DCUN1D5** | 5.0414 | 1.32099620775719 | 1 | 0.964 | 8.2563969 |
| **AXL** | 6.5042 | -2.54010114966922 | 0.183 | 0.915 | 1.0652084 |
| **TBC1D23** | 7.3077 | 1.70003389882155 | 1 | 0.956 | 1.1967851 |
| **CCDC59** | 1.2901 | 1.44845358918581 | 0.997 | 0.853 | 2.1129045 |
| **TSC22D1** | 4.0655 | 1.72123967895117 | 0.941 | 0.441 | 6.6582070 |
| **RAB6A** | 1.1450 | 1.26249188919947 | 1 | 0.987 | 1.8752739 |
| **PIEZO1** | 1.9963 | 1.01256407947093 | 0.592 | 0.047 | 3.2694037 |
| **ZNF770** | 2.9210 | 1.14221641212185 | 0.741 | 0.125 | 4.7837847 |
| **RMND1** | 7.8081 | 1.56898151374607 | 1 | 0.909 | 1.2787327 |
| **CDH2** | 9.3322 | -3.1042497312417 | 0.264 | 0.941 | 1.5283410 |

| | | | | | |
|---|---|---|---|---|---|
| **SERPINB6** | 1.2861 | 1.83332631005471 | 0.998 | 0.792 | 2.1063512 |
| **GSPT1** | 5.6590 | 1.2948296477705 | 1 | 0.912 | 9.2677988 |
| **TMX1** | 1.0003 | 1.1380572328914 | 1 | 0.975 | 1.6382415 |
| **HSBP1L1** | 4.5315 | 1.07937894717291 | 0.616 | 0.059 | 7.4213432 |
| **ERGIC3** | 4.6811 | -0.98994926408177 | 1 | 1 | 7.6663391 |
| **TCN2** | 4.1169 | -1.78134870191928 | 0.208 | 0.934 | 6.7423941 |
| **ASRGL1** | 4.5126 | 1.58543707641671 | 0.995 | 0.931 | 7.3903447 |
| **GJC1** | 1.2651 | -1.69173119421807 | 0.854 | 0.999 | 2.0719303 |
| **HDGF** | 1.4610 | -1.44202157851718 | 0.926 | 0.999 | 2.3927626 |
| **WWC3** | 3.1532 | -1.92019199798237 | 0.443 | 0.976 | 5.1641203 |
| **UFM1** | 4.4738 | 1.43156494446995 | 1 | 0.944 | 7.3268270 |
| **NF2** | 1.3768 | 1.21247472557762 | 1 | 0.995 | 2.2549342 |
| **SEC63** | 2.3549 | 1.26307507544303 | 1 | 0.996 | 3.8566301 |
| **ATP6V0E1** | 9.1231 | 1.31146811785219 | 0.998 | 0.961 | 1.4940990 |
| **MSN** | 1.0476 | -1.58848442303193 | 0.829 | 1 | 1.7156595 |
| **ZNF718** | 1.7404 | 0.985402176837859 | 0.69 | 0.096 | 2.8503908 |
| **HEPH** | 1.7556 | -1.85041591471582 | 0.047 | 0.869 | 2.8752629 |
| **AFAP1L1** | 6.9902 | 1.40135242841963 | 0.812 | 0.187 | 1.1447855 |
| **AP2B1** | 8.2882 | 1.32942141217501 | 1 | 0.873 | 1.3573708 |
| **ACP1** | 8.3249 | 1.37767358787872 | 0.978 | 0.623 | 1.3633836 |
| **NIFK** | 1.5803 | 1.27410472524746 | 1 | 0.956 | 2.5881383 |
| **FAM136A** | 4.1563 | 1.29599798110879 | 1 | 0.964 | 6.8068567 |
| **ARHGEF40** | 5.4083 | -1.89590364113142 | 0.279 | 0.935 | 8.8572576 |
| **STAU1** | 1.2674 | -0.83048257476593 | 1 | 1 | 2.0756952 |
| **COL26A1** | 1.4543 | -2.12264083651168 | 0.723 | 0.988 | 2.3817387 |
| **SERPING1** | 1.9555 | -1.68226158232913 | 0.503 | 0.974 | 3.2026093 |
| **RGS5** | 1.2038 | -2.87583146823061 | 0.227 | 0.925 | 1.9714936 |
| **TWF1** | 2.1226 | 1.35453605522151 | 0.895 | 0.327 | 3.4762622 |
| **ERRFI1** | 2.7202 | 1.90094027494512 | 0.988 | 0.756 | 4.4549208 |
| **NTS** | 2.7998 | -2.83969465995824 | 0.646 | 0.995 | 4.5853455 |
| **CTHRC1** | 3.4902 | 0.99503145701839 | 0.54 | 0.034 | 5.7159762 |
| **SMARCA2** | 3.9133 | 1.8795009021489 | 0.949 | 0.574 | 6.4089505 |
| **FBXO36** | 8.6929 | 1.81486647178159 | 0.963 | 0.676 | 1.4236477 |
| **CD99** | 9.2713 | -1.97553075840236 | 0.389 | 0.969 | 1.5183759 |

| | | | | | |
|---|---|---|---|---|---|
| **BARX1** | 1.0416 | 1.34172157217147 | 0.746 | 0.131 | 1.7059871 |
| **LGALS3BP** | 2.1995 | -1.86358357058459 | 0.206 | 0.914 | 3.6021752 |
| **HS6ST2** | 2.2593 | -1.88255582929413 | 0.19 | 0.921 | 3.7002001 |
| **HSPA9** | 3.4970 | 1.0565640652934 | 1 | 1 | 5.7271971 |
| **RAB5A** | 5.3274 | 1.34543742917333 | 1 | 0.94 | 8.7247513 |
| **TSTD1** | 6.4225 | 0.863883152630089 | 0.511 | 0.026 | 1.0518213 |
| **RASIP1** | 1.1957 | 1.37606366845993 | 0.8 | 0.184 | 1.9582102 |
| **PAM** | 1.3272 | -1.89403851382922 | 0.117 | 0.884 | 2.1736656 |
| **DHCR24** | 1.8757 | -1.51453186153682 | 1 | 1 | 3.0719175 |
| **EPHA2** | 2.2667 | -2.09035266238867 | 0.569 | 0.991 | 3.7122615 |
| **VPS37B** | 3.6521 | 1.66962101748312 | 0.981 | 0.709 | 5.9811265 |
| **WAC** | 3.7083 | 1.08581927759493 | 0.695 | 0.108 | 6.0732108 |
| **P4HA1** | 4.2976 | -1.39122773147505 | 0.976 | 1 | 7.0382171 |
| **DFFA** | 1.5169 | 1.18167527042519 | 1 | 0.997 | 2.4842539 |
| **TUNAR** | 2.0541 | -2.5015996197423 | 0.198 | 0.901 | 3.3641560 |
| **ICA1L** | 3.3149 | 1.14007172618387 | 0.65 | 0.081 | 5.4288549 |
| **ZNF146** | 3.4104 | 0.960837130532949 | 1 | 1 | 5.5852144 |
| **UCHL1** | 4.1842 | -1.93546303152577 | 0.81 | 0.997 | 6.8524872 |
| **GTF2A2** | 8.4399 | 1.27216042037089 | 1 | 0.958 | 1.3822093 |
| **MTHFD2** | 1.2888 | 1.45135775849937 | 1 | 1 | 2.1106919 |
| **CAPZA1** | 3.1683 | 1.35841544983451 | 0.912 | 0.36 | 5.1887331 |
| **KLK13** | 3.1982 | 1.62695065969077 | 0.883 | 0.315 | 5.2378379 |
| **ANGPT1** | 4.4033 | 1.10940463727282 | 0.706 | 0.109 | 7.2113641 |
| **TBCD** | 5.1628 | 1.24154343481778 | 1 | 0.999 | 8.4551660 |
| **ZNF649** | 5.9709 | 1.65567387549588 | 0.986 | 0.812 | 9.7786920 |
| **C1QBP** | 2.5181 | 0.986622668694865 | 1 | 0.999 | 4.1239676 |
| **SPINT1** | 2.6366 | 1.42139501194856 | 1 | 0.892 | 4.3180548 |
| **PYCR1** | 6.9259 | 1.66067164970864 | 0.992 | 0.596 | 1.1342604 |
| **KIF5C** | 1.7936 | -1.45086311562277 | 0.89 | 1 | 2.9374877 |
| **ADORA2A** | 2.2159 | 1.11711629549904 | 0.677 | 0.099 | 3.6291137 |
| **VENTX** | 2.6544 | 1.74914612297565 | 0.948 | 0.482 | 4.3472440 |
| **NIPA2** | 2.9608 | 1.12484014653506 | 1 | 0.965 | 4.8489564 |
| **CTNNBIP1** | 3.8991 | -1.44492362982244 | 0.805 | 0.997 | 6.3856295 |
| **EIF3E** | 9.3309 | 1.25022688477464 | 0.802 | 0.201 | 1.5281232 |

| | | | | | |
|---|---|---|---|---|---|
| **TPST2** | 9.6815 | 1.57849862926133 | 0.992 | 0.807 | 1.5855474 |
| **KLF5** | 9.6955 | 2.20223721770995 | 0.986 | 0.701 | 1.5878428 |
| **AGMAT** | 2.3659 | 0.813535821996187 | 0.545 | 0.039 | 3.8746382 |
| **TMEM258** | 4.9327 | 1.2297516922401 | 0.998 | 0.923 | 8.0783590 |
| **TERF1** | 5.2834 | 2.04483240071391 | 0.995 | 0.883 | 8.6527620 |
| **DPPA4** | 8.3015 | 1.18224341041473 | 1 | 1 | 1.3595425 |
| **ATG5** | 1.3812 | 1.28736645960412 | 0.998 | 0.949 | 2.2621446 |
| **DPYSL2** | 2.3075 | -1.29197994098136 | 0.973 | 1 | 3.7790514 |
| **MAP7D1** | 2.7427 | -1.69217170991615 | 0.519 | 0.982 | 4.4917584 |
| **FLOT2** | 2.7832 | -1.35488540777958 | 0.885 | 0.999 | 4.5581987 |
| **HINT1** | 3.0231 | 1.2064610909867 | 0.998 | 0.982 | 4.9510314 |
| **GNPNAT1** | 9.1147 | 1.47008947070178 | 0.998 | 0.937 | 1.4927162 |
| **PBX2** | 1.4681 | -1.18817648479521 | 0.985 | 1 | 2.4044134 |
| **RBM7** | 1.6350 | 1.39871473788594 | 0.985 | 0.726 | 2.6776738 |
| **CHORDC1** | 2.7732 | 1.37871577965368 | 0.895 | 0.339 | 4.5416973 |
| **RPA2** | 4.1201 | 1.33279123743019 | 0.986 | 0.654 | 6.7475316 |
| **HBA2** | 4.8951 | 1.49323383446028 | 0.765 | 0.149 | 8.0167773 |
| **BMP7** | 7.2171 | -2.0729974449651 | 0.677 | 0.985 | 1.1819508 |
| **DZIP1** | 1.7236 | -1.47282914012034 | 0.702 | 0.992 | 2.8228817 |
| **RHOF** | 2.0412 | 1.56936442747569 | 0.946 | 0.456 | 3.3429592 |
| **AIRE** | 6.3580 | 0.939812573605632 | 0.474 | 0.02 | 1.0412589 |
| **ZNF486** | 8.0221 | 1.81327810938085 | 0.902 | 0.407 | 1.3137798 |
| **EFNB1** | 8.7100 | -2.1503062035429 | 0.509 | 0.974 | 1.4264457 |
| **H2AFZ** | 2.0637 | 1.08861727018047 | 1 | 0.995 | 3.3798372 |
| **IRX4** | 2.4206 | 1.00742848316786 | 0.445 | 0.012 | 3.9643704 |
| **LOXL2** | 3.5103 | -1.78014622824105 | 0.65 | 0.993 | 5.7488382 |
| **TMBIM1** | 5.7388 | 1.89148290799828 | 0.917 | 0.39 | 9.3985083 |
| **TFAM** | 8.7001 | 1.18417779925831 | 0.997 | 0.913 | 1.4248222 |
| **DDIT4L** | 1.3148 | 0.843209440013169 | 0.45 | 0.014 | 2.1533492 |
| **PIEZO2** | 1.3312 | -1.98097568735981 | 0.081 | 0.855 | 2.1801278 |
| **ZNF765** | 1.7039 | 0.955163135553749 | 0.665 | 0.094 | 2.7905835 |
| **CFLAR** | 2.1313 | 1.51497231300119 | 0.985 | 0.762 | 3.4905199 |
| **CTH** | 2.4960 | 1.84008730364844 | 0.971 | 0.698 | 4.0877896 |
| **NEFL** | 4.0365 | 0.86487275261656 | 0.438 | 0.011 | 6.6107095 |

| | | | | | |
|---|---|---|---|---|---|
| **SRC** | 4.9839 | -1.40968820226209 | 0.814 | 0.997 | 8.1622023 |
| **HES1** | 5.7734 | -1.92861119562686 | 0.343 | 0.955 | 9.4552006 |
| **NEBL** | 6.2955 | -1.79847948157646 | 0.183 | 0.897 | 1.0310250 |
| **TIMM23** | 7.6991 | 0.938847739728611 | 1 | 1 | 1.2608865 |
| **PSMC4** | 1.0677 | 1.21322867844514 | 1 | 1 | 1.7486434 |
| **SGK1** | 2.9849 | 2.09739521697146 | 0.986 | 0.649 | 4.8884732 |
| **ECHDC1** | 3.3948 | -1.61822387401494 | 0.259 | 0.943 | 5.5596812 |
| **NCEH1** | 3.4974 | 1.26829576023048 | 0.741 | 0.152 | 5.7277091 |
| **NUCKS1** | 3.5423 | -0.88616226297282 | 1 | 1 | 5.8012715 |
| **STRADB** | 5.9012 | 0.874346756002516 | 0.631 | 0.078 | 9.6644409 |
| **SKA3** | 6.0495 | 1.7127202815144 | 0.995 | 0.939 | 9.9073283 |
| **TMEM97** | 1.2516 | -1.2724553989378 | 0.985 | 1 | 2.0497679 |
| **DNAJC5** | 1.2723 | -1.12341565497098 | 0.966 | 1 | 2.0837681 |
| **CCDC117** | 4.2481 | 1.36666911289617 | 0.973 | 0.673 | 6.9572217 |
| **DNAJC15** | 8.4319 | 2.01487005758923 | 0.997 | 0.997 | 1.3808995 |
| **UBE2E3** | 1.1636 | -1.22966282614661 | 0.944 | 1 | 1.9057894 |
| **PRDM14** | 2.6432 | 2.10137546287069 | 0.985 | 0.765 | 4.3289173 |
| **COL4A2** | 3.2883 | -1.63748842183897 | 0.893 | 1 | 5.3853566 |
| **DACT2** | 5.0649 | 0.993586392623396 | 0.53 | 0.038 | 8.2949058 |
| **NRG1** | 7.6834 | 1.01849006895529 | 0.565 | 0.053 | 1.2583162 |
| **GAR1** | 8.2407 | 1.20827988735563 | 0.992 | 0.911 | 1.3495918 |
| **ATP6V1E1** | 1.5935 | 1.06542838875379 | 1 | 0.985 | 2.6096976 |
| **IGFBPL1** | 1.6505 | -2.21644615332897 | 0.289 | 0.927 | 2.7031360 |
| **VPS26A** | 1.7160 | 1.17818128555195 | 1 | 0.955 | 2.8103288 |
| **XBP1** | 1.8752 | 1.25333606905601 | 1 | 0.998 | 3.0710876 |
| **PTMS** | 2.2943 | -1.21599370949028 | 1 | 1 | 3.7574439 |
| **NUP43** | 3.3738 | 1.03515324775915 | 1 | 0.988 | 5.5253462 |
| **DEPTOR** | 3.9647 | 1.64340573177868 | 0.897 | 0.366 | 6.4930511 |
| **IGDCC3** | 4.2455 | -1.73973587036087 | 0.567 | 0.979 | 6.9529830 |
| **ID2** | 9.2105 | -1.94677550492682 | 0.303 | 0.923 | 1.5084188 |
| **B4GALT4** | 1.8553 | 1.2903487316304 | 0.848 | 0.256 | 3.0385771 |
| **PAICS** | 2.2291 | 1.23786457484362 | 0.998 | 0.886 | 3.6506116 |
| **DNAJA1** | 3.7007 | 1.28175037844119 | 0.997 | 0.908 | 6.0607211 |
| **MPP6** | 7.0620 | -1.31162628864089 | 0.934 | 0.999 | 1.1565473 |

| | | | | | |
|---|---|---|---|---|---|
| **FUNDC1** | 7.6563 | 1.35259657917948 | 0.98 | 0.772 | 1.2538834 |
| **TXNDC17** | 8.3396 | 0.976158724211232 | 1 | 1 | 1.3657876 |
| **C7orf50** | 1.6027 | -1.15756547093926 | 0.983 | 1 | 2.6248259 |
| **CALM1** | 1.6631 | 1.36702818313288 | 0.986 | 0.793 | 2.7237172 |
| **FLOT1** | 1.8517 | -1.12108221183509 | 0.978 | 1 | 3.0326303 |
| **TRIM26** | 1.9744 | 1.64509914310332 | 0.978 | 0.823 | 3.2335119 |
| **TCF4** | 2.7301 | -1.67252614399977 | 0.53 | 0.974 | 4.4711210 |
| **DBNDD2** | 3.7052 | -1.78823865395734 | 0.301 | 0.904 | 6.0681496 |
| **WDHD1** | 4.3643 | 1.1055451520181 | 1 | 0.999 | 7.1475386 |
| **OTUD4** | 4.3938 | 1.27086159977162 | 0.998 | 0.824 | 7.1958443 |
| **GAD1** | 6.2034 | -1.78968867710566 | 0.425 | 0.965 | 1.0159340 |
| **PEA15** | 2.2746 | -1.49305516093479 | 0.936 | 1 | 3.7252760 |
| **FAT1** | 3.7722 | -1.52051405624938 | 0.848 | 0.998 | 6.1778472 |
| **TET1** | 4.4385 | 1.14339496573305 | 1 | 0.997 | 7.2690114 |
| **ZNF878** | 5.3688 | 1.23966637304392 | 0.682 | 0.118 | 8.7925548 |
| **FAM174B** | 7.1423 | -1.90017716835584 | 0.426 | 0.942 | 1.1696992 |
| **C18orf32** | 9.9757 | 1.29991348007777 | 0.897 | 0.363 | 1.6337211 |
| **PTGR1** | 1.4457 | 1.38363333675295 | 0.992 | 0.811 | 2.3676758 |
| **UTP6** | 2.2138 | 1.12907871133264 | 1 | 0.958 | 3.6255761 |
| **PTPRD** | 3.1968 | 0.979183555989659 | 0.509 | 0.036 | 5.2355260 |
| **GALNT1** | 5.7313 | -1.21821192971706 | 0.956 | 1 | 9.3862259 |
| **BTBD2** | 7.6649 | -1.32196431913189 | 0.936 | 0.999 | 1.2552885 |
| **PRTG** | 9.2489 | -2.15958417237823 | 0.184 | 0.896 | 1.5146938 |
| **COX4I1** | 9.2667 | 0.935865063605462 | 1 | 0.997 | 1.5176178 |
| **NOTCH3** | 1.0117 | -1.48720298467825 | 0.88 | 0.995 | 1.6568896 |
| **TPGS2** | 1.4358 | -1.14439569354838 | 0.983 | 1 | 2.3514907 |
| **PRKACA** | 7.8029 | -1.16413088589141 | 0.971 | 1 | 1.2778940 |
| **PUS1** | 1.3332 | 1.15201278108011 | 0.998 | 0.955 | 2.1834435 |
| **SPIC** | 1.3792 | 1.18752751930803 | 0.46 | 0.022 | 2.2587293 |
| **CST3** | 3.1496 | -1.80908670945879 | 0.445 | 0.953 | 5.1582557 |
| **ZNF841** | 3.9887 | 1.25418644474635 | 0.745 | 0.162 | 6.5324077 |
| **ARPC1B** | 6.9234 | 1.20167574443311 | 1 | 0.995 | 1.1338585 |
| **TBX1** | 1.7628 | 0.851145857655722 | 0.459 | 0.021 | 2.8870370 |
| **GRSF1** | 4.7359 | 0.962482272776631 | 1 | 0.997 | 7.7560933 |

| | | | | | |
|---|---|---|---|---|---|
| **TRIML2** | 1.5598 | 1.70079161859499 | 0.981 | 0.749 | 2.5545201[illegible] |
| **WFDC2** | 1.8130 | -2.98810815270645 | 0.416 | 0.931 | 2.9692810[illegible] |
| **TMEM98** | 5.2199 | -1.41668586551351 | 0.709 | 0.985 | 8.5486383[illegible] |
| **PIP4K2A** | 9.9037 | 0.983676798807238 | 0.684 | 0.12 | 1.6219410[illegible] |
| **UGP2** | 1.5597 | 1.36979267675308 | 1 | 0.932 | 2.5543281[illegible] |
| **PKDCC** | 2.3496 | -2.36945605984265 | 0.352 | 0.936 | 3.8480601[illegible] |
| **PCDHB2** | 2.7700 | -1.67110028426165 | 0.266 | 0.893 | 4.5365501[illegible] |
| **STMN1** | 2.8142 | 1.21253734879109 | 0.998 | 0.99 | 4.6088485[illegible] |
| **SH3BGRL** | 3.2293 | -1.49495145031559 | 0.438 | 0.968 | 5.2886278[illegible] |
| **PCYT1B** | 3.8224 | -1.63811837643791 | 0.369 | 0.926 | 6.2599561[illegible] |
| **NELFCD** | 5.8803 | -0.96188761768824 | 0.998 | 1 | 9.6302228[illegible] |
| **NFKB2** | 8.0517 | 2.07227632945047 | 0.973 | 0.834 | 1.3186388[illegible] |
| **FUCA1** | 1.6060 | 1.69011621793572 | 0.976 | 0.722 | 2.6302617[illegible] |
| **ZNF625** | 2.0208 | 1.19374052811354 | 0.998 | 0.931 | 3.3096198[illegible] |
| **ARHGAP35** | 3.0127 | -1.15780779814919 | 0.902 | 0.995 | 4.9339177[illegible] |
| **CBFA2T2** | 6.9267 | 1.72684412257428 | 0.99 | 0.918 | 1.1344001[illegible] |
| **S100A10** | 8.1451 | 1.3358836277768 | 1 | 0.998 | 1.3339305[illegible] |
| **PRXL2A** | 8.5807 | -1.2025263313254 | 0.915 | 0.998 | 1.4052707[illegible] |
| **SLC6A8** | 1.5207 | 1.67329453352435 | 0.993 | 0.95 | 2.4905298[illegible] |
| **TIMM17A** | 2.0138 | 1.02278614854147 | 1 | 0.987 | 3.2980757[illegible] |
| **KLF17** | 3.7350 | 1.23286443966008 | 0.381 | 0.004 | 6.1168686[illegible] |
| **NRROS** | 8.1865 | 0.918325283536263 | 0.518 | 0.044 | 1.3407183[illegible] |
| **SOX12** | 2.3862 | -1.60586828601565 | 0.599 | 0.971 | 3.9079939[illegible] |
| **TMEM64** | 2.4945 | -2.22967365310169 | 0.673 | 0.988 | 4.0853688[illegible] |
| **WASF3** | 2.6970 | -1.45368616236606 | 0.795 | 0.99 | 4.4169504[illegible] |
| **RABGGTB** | 3.4801 | 1.252457315379 | 1 | 0.997 | 5.6995128[illegible] |
| **NES** | 6.9127 | -1.35155457457137 | 0.998 | 1 | 1.1321065[illegible] |
| **GPRC5A** | 1.1721 | 1.77898277983365 | 0.526 | 0.049 | 1.9196478[illegible] |
| **NMRAL1** | 6.6761 | 1.34698439465162 | 0.986 | 0.72 | 1.0933466[illegible] |
| **LLPH** | 1.3202 | 1.24798938790012 | 0.912 | 0.438 | 2.1621135[illegible] |
| **IL32** | 1.3249 | 1.7811190885309 | 0.887 | 0.333 | 2.1698922[illegible] |
| **AKAP12** | 1.7726 | 1.24881709789116 | 0.998 | 0.997 | 2.9030495[illegible] |
| **MARVELD2** | 2.7773 | 1.24154750171213 | 0.822 | 0.237 | 4.5484399[illegible] |
| **CARNMT1** | 3.1782 | 1.14651608068758 | 0.998 | 0.889 | 5.2050084[illegible] |

| | | | | | |
|---|---|---|---|---|---|
| **FADS1** | 4.0226 | -1.39746421344511 | 0.978 | 1 | 6.5879087 |
| **BCAM** | 5.1713 | -2.04853861745425 | 0.592 | 0.946 | 8.4691871 |
| **MFAP2** | 5.2584 | -1.37506801983421 | 0.937 | 1 | 8.6117497 |
| **MAPRE1** | 6.7658 | -0.82886209760705 | 1 | 1 | 1.1080385 |
| **ZNF219** | 7.1402 | -1.76119544198769 | 0.606 | 0.96 | 1.1693568 |
| **IDS** | 8.2797 | -1.44892682467986 | 0.305 | 0.913 | 1.3559827 |
| **TTLL12** | 8.3747 | 1.03513054802733 | 1 | 0.991 | 1.3715394 |
| **KRR1** | 9.8707 | 1.17441749223894 | 0.986 | 0.811 | 1.6165366 |
| **MAP4K4** | 2.3476 | -0.98166232164854 | 0.998 | 1 | 3.8446999 |
| **DDX21** | 5.0464 | 0.874136559226202 | 1 | 1 | 8.2645512 |
| **SEC61B** | 6.1898 | 1.10467576947648 | 0.998 | 0.946 | 1.0137170 |
| **LIMK1** | 1.0631 | -1.08468887585262 | 0.983 | 1 | 1.7410718 |
| **ZNF362** | 4.6131 | -1.32424404068369 | 0.785 | 0.995 | 7.5549744 |
| **WWOX** | 5.1155 | -1.58019175299403 | 0.557 | 0.963 | 8.3777488 |
| **CISD1** | 7.3461 | 1.24026759159774 | 0.995 | 0.892 | 1.2030799 |
| **STOML2** | 9.4166 | 0.842564910861251 | 1 | 1 | 1.5421660 |
| **DCBLD2** | 1.4095 | -1.19587158989033 | 0.865 | 0.998 | 2.3083482 |
| **CLPX** | 1.4554 | 1.1265534053001 | 0.998 | 0.978 | 2.3836506 |
| **PITPNC1** | 2.8445 | 1.15366920403764 | 0.992 | 0.961 | 4.6585549 |
| **UBQLN2** | 2.9569 | -1.18784949370605 | 0.866 | 1 | 4.8426205 |
| **DMBX1** | 4.6213 | 1.0791522126147 | 0.489 | 0.037 | 7.5684452 |
| **BAK1** | 5.0410 | 1.02823656934872 | 0.998 | 0.997 | 8.2557258 |
| **PLPP1** | 5.1833 | 1.86235144738848 | 0.997 | 0.858 | 8.4887002 |
| **COX20** | 5.4028 | 1.29721247766956 | 0.892 | 0.404 | 8.8482296 |
| **SLC16A2** | 9.2079 | -1.65858052195061 | 0.616 | 0.987 | 1.5079791 |
| **ARHGEF9** | 1.0561 | -1.52549733727841 | 0.124 | 0.83 | 1.7296673 |
| **ZNF398** | 1.0858 | 1.5726697435279 | 0.97 | 0.761 | 1.7782716 |
| **CCND2** | 1.1170 | -2.06118336741686 | 0.619 | 0.976 | 1.8294444 |
| **MAGEH1** | 1.4428 | -1.47929739530922 | 0.306 | 0.931 | 2.3629281 |
| **PTK7** | 1.4999 | -1.23329856012047 | 0.993 | 1 | 2.4564667 |
| **EIF3J** | 2.2354 | 0.979359707758388 | 1 | 0.987 | 3.6610440 |
| **SNX3** | 2.7695 | 1.20328136086574 | 0.985 | 0.775 | 4.5356235 |
| **SELENOP** | 3.0342 | -1.76442808849802 | 0.152 | 0.851 | 4.9691195 |
| **SMG1** | 3.2794 | 1.28528413879863 | 0.944 | 0.543 | 5.3707554 |

| | | | | | |
|---|---|---|---|---|---|
| **PGM1** | 3.4418 | -2.07510395713267 | 0.753 | 0.997 | 5.6367359 |
| **PRSS12** | 4.4260 | 1.70902878908841 | 0.87 | 0.373 | 7.2484943 |
| **DTD1** | 6.4798 | -1.05563605050977 | 0.983 | 1 | 1.0612038 |
| **TMEM87A** | 6.5748 | 1.23687762216311 | 0.958 | 0.64 | 1.0767610 |
| **RCN1** | 8.0101 | 1.16224375066855 | 1 | 0.995 | 1.3118185 |
| **ELAVL2** | 1.3914 | 0.824300298223887 | 0.418 | 0.016 | 2.2788576 |
| **FGFR4** | 1.9682 | 1.60136187666369 | 0.98 | 0.849 | 3.2233746 |
| **CDK6** | 2.0974 | -1.50759227944739 | 0.347 | 0.933 | 3.4349168 |
| **FADS3** | 2.3353 | 1.23546210616685 | 0.824 | 0.256 | 3.8246449 |
| **TPM2** | 2.6896 | -1.28028541760925 | 0.993 | 1 | 4.4047781 |
| **ZNF682** | 2.8520 | 1.48605420342539 | 0.893 | 0.414 | 4.6707345 |
| **VAMP8** | 3.0223 | -1.29241379197066 | 0.993 | 1 | 4.9496357 |
| **3,00 DKK** | 3.0326 | -1.61001051752015 | 0.469 | 0.955 | 4.9665475 |
| **UQCRFS1** | 3.2171 | 1.22270614409012 | 0.946 | 0.518 | 5.2686673 |
| **RPA3** | 4.9018 | 1.24485612160216 | 0.861 | 0.325 | 8.0277176 |
| **STOM** | 6.9629 | 1.93780485539955 | 0.983 | 0.778 | 1.1403299 |
| **TMSB15A** | 2.1693 | -2.21099329365503 | 0.13 | 0.823 | 3.5526630 |
| **GSTO2** | 2.4395 | 0.845528734563799 | 0.538 | 0.055 | 3.9952753 |
| **RIF1** | 2.6019 | 1.01118847322608 | 1 | 0.994 | 4.2611954 |
| **LHFPL6** | 3.8371 | -2.02760002006541 | 0.174 | 0.847 | 6.2840220 |
| **MFGE8** | 4.9118 | -1.47225414253359 | 0.575 | 0.96 | 8.0440680 |
| **TAGLN** | 5.4760 | -2.77356915169252 | 0.888 | 0.997 | 8.9681869 |
| **IGF2R** | 8.5505 | -1.32233956402229 | 0.657 | 0.982 | 1.4003267 |
| **SLC35A3** | 1.2919 | 1.12885246321593 | 0.773 | 0.209 | 2.1158390 |
| **PTOV1** | 1.3098 | -0.91344060597586 | 0.998 | 1 | 2.1450976 |
| **PDIA3** | 2.6791 | 0.958944854442316 | 1 | 0.997 | 4.3877176 |
| **FAAP20** | 3.8657 | -1.0558529531625 | 0.951 | 0.999 | 6.3309105 |
| **RSL1D1** | 4.2645 | 0.93925154455404 | 1 | 0.976 | 6.9840968 |
| **ABCB7** | 5.6561 | -1.16158774125914 | 0.866 | 1 | 9.2631362 |
| **NIT2** | 1.0231 | 1.26342465160169 | 0.942 | 0.529 | 1.6756883 |
| **SPATC1L** | 1.2310 | -1.47704460855566 | 0.058 | 0.794 | 2.0160645 |
| **COL4A5** | 1.9746 | -1.67506180276956 | 0.176 | 0.856 | 3.2338090 |
| **TSPAN4** | 3.5069 | -1.261163845029 | 0.876 | 0.996 | 5.7433411 |
| **ATP5MC3** | 4.0196 | 0.91519520878577 | 1 | 1 | 6.5830280 |

| | | | | | |
|---|---|---|---|---|---|
| **SERPINE2** | 4.1547 | -2.67276045442653 | 0.86 | 0.997 | 6.8043144 |
| **SERPINB5** | 4.6681 | 0.820058184316483 | 0.377 | 0.007 | 7.6449632 |
| **CPS1** | 9.5194 | -1.32665937033153 | 0.572 | 0.972 | 1.5589966 |
| **GLIPR2** | 1.0539 | -2.46858068825747 | 0.443 | 0.942 | 1.7260301 |
| **COL4A1** | 1.0970 | -1.36154246224626 | 0.853 | 0.999 | 1.7965590 |
| **PIM2** | 1.3157 | 1.72473764654669 | 0.985 | 0.973 | 2.1547766 |
| **DLK1** | 1.3259 | 1.12670882118731 | 0.69 | 0.138 | 2.1714292 |
| **CANX** | 2.4823 | 1.0876028357709 | 1 | 0.998 | 4.0653571 |
| **SV2A** | 6.6443 | -1.47788643161033 | 0.335 | 0.908 | 1.0881474 |
| **NOC2L** | 8.2659 | 0.865718437884229 | 1 | 0.998 | 1.3537155 |
| **C14orf132** | 9.4490 | -1.89678797814234 | 0.09 | 0.799 | 1.5474759 |
| **SHPK** | 2.4884 | 1.50423262138815 | 0.98 | 0.834 | 4.0753131 |
| **FDPS** | 2.7753 | -1.07677427712288 | 0.99 | 1 | 4.5451428 |
| **PRDX6** | 3.5963 | 0.89093641954444 | 1 | 0.997 | 5.8897844 |
| **TRAPPC1** | 3.7481 | -1.0370088812411 | 0.973 | 1 | 6.1383090 |
| **SYNGR1** | 6.3611 | -1.5816447195943 | 0.343 | 0.901 | 1.0417686 |
| **ARMCX2** | 9.7383 | -1.41136115097209 | 0.679 | 0.994 | 1.5948508 |
| **AGPAT5** | 1.1065 | -1.15107212187712 | 0.941 | 1 | 1.8121276 |
| **TSPAN3** | 1.1634 | -0.98747520162508 | 0.997 | 1 | 1.9054579 |
| **DNAJA3** | 1.4739 | 0.947438382377499 | 1 | 0.996 | 2.4139006 |
| **COA6** | 1.7099 | 1.20079222281913 | 0.986 | 0.751 | 2.8004170 |
| **ASH2L** | 1.8194 | 1.26613492347928 | 0.998 | 0.988 | 2.9797773 |
| **ZNF578** | 1.9258 | 1.27984848463725 | 0.865 | 0.314 | 3.1539918 |
| **RAB11B** | 1.9809 | -0.87094508920419 | 1 | 1 | 3.2441541 |
| **ZDHHC2** | 2.1385 | -1.7131279512946 | 0.181 | 0.835 | 3.5023551 |
| **GLRX5** | 4.0499 | -0.86412778589359 | 1 | 1 | 6.6326435 |
| **MYEF2** | 4.9422 | -1.6917296333709 | 0.386 | 0.919 | 8.0939512 |
| **RIN2** | 5.4462 | -1.58431559723575 | 0.347 | 0.913 | 8.9192842 |
| **CXXC5** | 2.1214 | -1.19612977552034 | 0.959 | 0.999 | 3.4743145 |
| **BTBD3** | 2.2497 | -1.35104033020984 | 0.692 | 0.984 | 3.6843535 |
| **MAPK10** | 2.2719 | -2.24628292149744 | 0.074 | 0.787 | 3.7208109 |
| **EIF3H** | 2.8902 | 0.844119522905948 | 1 | 0.998 | 4.7333121 |
| **SALL3** | 6.3412 | -1.69985179803765 | 0.562 | 0.973 | 1.0385135 |
| **NECTIN3** | 8.2300 | -1.52036622380879 | 0.318 | 0.907 | 1.3478351 |

| | | | | | |
|---|---|---|---|---|---|
| **ATP5MPL** | 9.6725 | 1.26045379262179 | 0.937 | 0.602 | 1.5840794 |
| **DOCK6** | 1.2226 | 1.24271733683327 | 0.99 | 0.805 | 2.0023770 |
| **SH3KBP1** | 1.8852 | -1.40141925268061 | 0.69 | 0.981 | 3.0874179 |
| **EOMES** | 2.5692 | -3.23386526760997 | 0.159 | 0.812 | 4.2077077 |
| **EGLN1** | 7.1236 | -1.3672168071279 | 0.753 | 0.985 | 1.1666368 |
| **NOLC1** | 1.0367 | 1.01273250468838 | 1 | 0.982 | 1.6978387 |
| **MUC4** | 1.1163 | 1.33754913709576 | 0.834 | 0.273 | 1.8282570 |
| **PDLIM1** | 2.3305 | 1.29319302841554 | 1 | 0.96 | 3.8167501 |
| **ENPP2** | 2.3680 | 1.12303646875769 | 0.726 | 0.166 | 3.8781695 |
| **ITGB1** | 2.6364 | -1.13723377688643 | 0.988 | 1 | 4.3177205 |
| **COX6B1** | 4.1841 | 0.946407494614647 | 1 | 0.994 | 6.8523301 |
| **INF2** | 5.5263 | -1.06224412903272 | 0.993 | 1 | 9.0505252 |
| **PCDHGB7** | 1.0021 | -1.7146959026577 | 0.193 | 0.832 | 1.6412268 |
| **MYSM1** | 1.1654 | 1.32155166093411 | 0.997 | 0.905 | 1.9087017 |
| **TEAD4** | 1.2438 | 0.96949749562114 | 1 | 0.989 | 2.0369799 |
| **MED15** | 2.3405 | 1.20614040170832 | 0.998 | 0.942 | 3.8331213 |
| **TNC** | 2.5273 | -2.75358890258578 | 0.169 | 0.834 | 4.1389860 |
| **TGS1** | 2.6289 | 1.03163931447883 | 0.998 | 0.994 | 4.3053514 |
| **AMOTL2** | 5.3710 | -1.17391191932679 | 0.992 | 0.999 | 8.7962399 |
| **FAM171A2** | 6.1475 | -1.41649281214654 | 0.533 | 0.952 | 1.0067798 |
| **NAP1L3** | 8.5267 | -1.60943662290249 | 0.311 | 0.894 | 1.3964216 |
| **CNN2** | 1.5282 | -1.05250064680434 | 1 | 1 | 2.5028582 |
| **NETO2** | 1.6281 | 1.45660717648803 | 0.978 | 0.781 | 2.6663712 |
| **CTNNB1** | 1.9544 | -0.90178644987561 | 1 | 1 | 3.2007456 |
| **CFDP1** | 2.4379 | -0.90483578488261 | 0.992 | 1 | 3.9925612 |
| **CKAP4** | 2.4666 | -1.07549255530912 | 0.99 | 1 | 4.0396715 |
| **ALDH1L2** | 4.1000 | 1.16401445157601 | 0.689 | 0.144 | 6.7146168 |
| **NRM** | 5.5416 | -1.3166776110825 | 0.525 | 0.934 | 9.0755416 |
| **ZNF185** | 6.4447 | -1.57102657965895 | 0.371 | 0.891 | 1.0554514 |
| **RAB13** | 8.2332 | -0.99491255233479 | 0.985 | 1 | 1.3483587 |
| **SRSF10** | 8.2332 | 1.08093510157083 | 0.79 | 0.231 | 1.3483653 |
| **B2M** | 1.9267 | -1.41458793612601 | 0.609 | 0.967 | 3.1555084 |
| **AP1M2** | 3.6491 | 1.19122151165703 | 0.878 | 0.353 | 5.9761639 |
| **RCN3** | 4.3809 | 0.958672725887309 | 0.638 | 0.113 | 7.1746411 |

| | | | | | |
|---|---|---|---|---|---|
| **HAUS1** | 6.2798 | 1.15017516586517 | 0.988 | 0.782 | 1.02844428 |
| **RBMXL1** | 6.7798 | 1.19802121701154 | 0.905 | 0.435 | 1.11033989 |
| **ENAH** | 1.1837 | -0.90663103699302 | 0.995 | 1 | 1.93854831 |
| **MRS2** | 1.2290 | 1.14799867991442 | 1 | 0.998 | 2.01273368 |
| **REST** | 2.0870 | 0.973359966110153 | 1 | 1 | 3.41803178 |
| **OSBPL9** | 2.5855 | -0.879158926473050 | 0.992 | 1 | 4.23442046 |
| **AP1S2** | 4.2891 | -1.40019645323973 | 0.525 | 0.95 | 7.02439680 |
| **RTN4** | 5.2602 | -0.890115544713495 | 0.998 | 1 | 8.61469130 |
| **IGFBP4** | 7.1595 | -1.79907961814549 | 0.398 | 0.932 | 1.17251560 |
| **SLCO4C1** | 7.9988 | 1.4386019148928 | 0.959 | 0.603 | 1.30996663 |
| **ARID1A** | 9.0550 | -0.945815496986820 | 0.988 | 1 | 1.48295138 |
| **USP11** | 1.1692 | -1.07194578218368 | 0.927 | 1 | 1.91490920 |
| **YAP1** | 1.6214 | -0.81433534708984 | 1 | 1 | 2.65550629 |
| **REEP6** | 1.8006 | 1.41884363144166 | 0.885 | 0.414 | 2.94897168 |
| **ZNF607** | 1.9331 | 1.01409932398134 | 0.685 | 0.153 | 3.16596963 |
| **FUCA2** | 2.2661 | -1.55936970091212 | 0.599 | 0.949 | 3.71127091 |
| **IFITM3** | 3.4249 | -1.18109874694092 | 0.998 | 1 | 5.60907500 |
| **NAXE** | 3.5506 | -1.10384169675637 | 0.866 | 0.996 | 5.81482125 |
| **STK26** | 3.9109 | -1.33922204113074 | 0.663 | 0.995 | 6.40501179 |
| **TBL1X** | 1.0193 | -1.37742932331296 | 0.386 | 0.925 | 1.66932213 |
| **ACTB** | 1.0227 | 0.924635105796844 | 1 | 1 | 1.67502350 |
| **LDOC1** | 2.8496 | -1.27577629737868 | 0.631 | 0.963 | 4.66684527 |
| **UBAP2** | 3.4352 | 1.01054968668194 | 0.998 | 0.967 | 5.62589028 |
| **JPT1** | 4.2111 | 1.02442159083288 | 1 | 0.998 | 6.89658794 |
| **ZIC2** | 4.6718 | -2.22858948839408 | 0.271 | 0.85 | 7.65100905 |
| **PCGF6** | 5.0755 | 1.22464532792103 | 0.953 | 0.638 | 8.31223672 |
| **CUX1** | 1.0105 | -1.3247573034779 | 0.76 | 0.988 | 1.65496820 |
| **ITGA5** | 1.0820 | -1.73271663420642 | 0.386 | 0.93 | 1.77202410 |
| **BNIP3L** | 1.2047 | -1.11974644554895 | 0.966 | 1 | 1.97301922 |
| **ZNF300** | 1.4141 | 1.17306834089795 | 0.838 | 0.284 | 2.31590687 |
| **RBM22** | 1.4431 | 1.10202060220863 | 0.985 | 0.851 | 2.36350445 |
| **GCNT1** | 2.0693 | 1.4017276074553 | 0.824 | 0.331 | 3.38889785 |
| **CITED2** | 4.3968 | -1.38025206704299 | 0.689 | 0.99 | 7.20064927 |
| **STK4** | 2.3217 | -1.12498994009866 | 0.861 | 0.992 | 3.80237059 |

| | | | | | |
|---|---|---|---|---|---|
| **DDT** | 2.4929 | 0.862182872048893 | 1 | 1 | 4.0827830 |
| **WDR5** | 4.8978 | 1.10179568627972 | 0.98 | 0.841 | 8.0211607 |
| **SH3BGRL3** | 7.2002 | -1.2787620157575 | 0.914 | 0.997 | 1.1791827 |
| **ERBIN** | 9.9338 | 1.27322474023448 | 0.995 | 0.946 | 1.6268747 |
| **HMOX2** | 1.0731 | 1.03275807853454 | 0.993 | 0.943 | 1.7574224 |
| **ANTXR1** | 1.2464 | -1.39150723616346 | 0.305 | 0.886 | 2.0412728 |
| **PDE12** | 1.8702 | 1.19311834784191 | 0.919 | 0.497 | 3.0628920 |
| **THAP12** | 3.0040 | 1.16076077878344 | 0.939 | 0.497 | 4.9196876 |
| **PRUNE2** | 3.0947 | 1.48673805315793 | 0.944 | 0.585 | 5.0682020 |
| **CDH1** | 7.1302 | 1.39138346573041 | 0.995 | 0.77 | 1.1677166 |
| **MCRIP1** | 8.1766 | -0.98256447042010 | 0.964 | 0.999 | 1.3390821 |
| **MAP1B** | 1.2788 | -1.25967399860491 | 0.992 | 1 | 2.0943451 |
| **TEC** | 2.0380 | 0.898231349562991 | 0.579 | 0.087 | 3.3377089 |
| **NBEA** | 2.0966 | -1.70791771534661 | 0.11 | 0.781 | 3.4336027 |
| **C16orf72** | 4.1123 | 0.887725474420803 | 1 | 0.998 | 6.7348507 |
| **ETFA** | 5.4329 | -1.17794292885556 | 0.802 | 0.993 | 8.8975432 |
| **CTPS2** | 5.6592 | -1.06264509320066 | 0.856 | 0.996 | 9.2681559 |
| **CNN3** | 8.2985 | -0.84667084704156 | 1 | 1 | 1.3590509 |
| **PARN** | 1.0770 | 0.940772675997621 | 1 | 0.982 | 1.7638722 |
| **RO60** | 1.3965 | -1.00207662891675 | 0.936 | 0.997 | 2.2870881 |
| **AKAP1** | 2.0685 | 1.23722355878114 | 0.997 | 0.957 | 3.3877038 |
| **KRT19** | 2.7530 | -1.3079370438369 | 0.712 | 0.982 | 4.5086998 |
| **ARHGDIG** | 7.2447 | 0.912611107408527 | 0.633 | 0.116 | 1.1864661 |
| **TP53I11** | 9.1329 | -1.35750757602178 | 0.387 | 0.928 | 1.4957079 |
| **PFKM** | 1.1168 | -1.15868678466606 | 0.746 | 0.983 | 1.8290959 |
| **ABI2** | 2.2026 | -0.94900053634987 | 0.983 | 1 | 3.6072082 |
| **ZNF608** | 4.9003 | -1.50413702281305 | 0.57 | 0.972 | 8.0252932 |
| **PLD3** | 5.4863 | -0.95140986070491 | 0.993 | 1 | 8.9849150 |
| **MORF4L2** | 5.5491 | 1.06002959358718 | 0.998 | 0.949 | 9.0877984 |
| **MCM8** | 1.3807 | -1.03371851846963 | 0.954 | 0.999 | 2.2612170 |
| **NANOGP8** | 2.5969 | 0.870755992373852 | 0.619 | 0.114 | 4.2529497 |
| **REPIN1** | 1.6775 | -1.2954580932798 | 0.423 | 0.913 | 2.7472772 |
| **ARHGAP8** | 1.7594 | 1.22270848884341 | 0.978 | 0.697 | 2.8814706 |
| **IGF1R** | 2.1203 | -1.08182981391653 | 0.963 | 1 | 3.4725587 |

| | | | | | |
|---|---|---|---|---|---|
| **UBE3C** | 4.4433 | 0.847952402728934 | 1 | 0.982 | 7.2768473 |
| **WNT8A** | 4.8332 | -2.27262569052669 | 0.357 | 0.906 | 7.9153447 |
| **SVBP** | 8.2764 | -1.02496629203061 | 0.904 | 0.997 | 1.3554262 |
| **SEC61G** | 8.6673 | 0.9383445465199 | 1 | 0.999 | 1.4194477 |
| **SORD** | 8.8688 | -1.30133810151644 | 0.445 | 0.916 | 1.4524540 |
| **SULF2** | 2.5583 | -1.61609170307206 | 0.836 | 0.988 | 4.1898473 |
| **USB1** | 2.6431 | -1.10851941181473 | 0.902 | 0.996 | 4.3287506 |
| **WDR12** | 2.9201 | 0.957164404998876 | 0.997 | 0.961 | 4.7823420 |
| **PHAX** | 3.8188 | 0.916530134475947 | 1 | 0.991 | 6.2540870 |
| **SLC25A44** | 4.0456 | 1.17674333656094 | 0.997 | 0.937 | 6.6254867 |
| **CEBPZ** | 4.1333 | 0.852963784788348 | 1 | 1 | 6.7691921 |
| **SLC2A4RG** | 5.0152 | -1.11431292639076 | 0.846 | 0.995 | 8.2134840 |
| **ZNF711** | 5.5169 | -1.35282000113684 | 0.337 | 0.905 | 9.0350723 |
| **TIMM8A** | 5.7588 | 1.10370783685087 | 0.997 | 0.975 | 9.4312368 |
| **TBC1D16** | 7.2574 | -1.00936907181765 | 0.936 | 1 | 1.1885536 |
| **RCOR2** | 8.7313 | -1.30321550344932 | 0.679 | 0.977 | 1.4299403 |
| **MZT1** | 1.0932 | 1.11758882688718 | 0.986 | 0.88 | 1.7904099 |
| **TRIM61** | 1.2785 | 1.20848852973965 | 0.736 | 0.215 | 2.0939267 |
| **MT-ND3** | 1.3543 | 2.53416598778896 | 0.953 | 0.722 | 2.2179617 |
| **WIPF3** | 1.4731 | 1.11646466636027 | 0.738 | 0.207 | 2.4125189 |
| **UQCRH** | 1.5513 | 0.854224060985377 | 1 | 0.999 | 2.5405890 |
| **GINS1** | 1.6575 | -0.92127983037925 | 0.995 | 1 | 2.7144973 |
| **ZNF260** | 1.7587 | 1.15810586153997 | 0.876 | 0.397 | 2.8803450 |
| **CCDC88A** | 2.1612 | -1.12427543320426 | 0.863 | 0.998 | 3.5394138 |
| **OAZ2** | 2.4029 | -0.96746696818429 | 0.986 | 1 | 3.9353396 |
| **AKR1B1** | 3.6402 | -0.88581140806100 | 0.995 | 1 | 5.9616624 |
| **CD164** | 4.8166 | 1.12748553353586 | 0.963 | 0.652 | 7.8882869 |
| **EFR3B** | 5.2025 | 1.67931381484806 | 0.929 | 0.653 | 8.5202870 |
| **CCND1** | 6.7908 | -1.09652752455593 | 0.997 | 0.999 | 1.1121435 |
| **HDGFL3** | 1.1691 | -0.92419375375800 | 0.981 | 1 | 1.9147928 |
| **AGRN** | 2.7898 | -1.23453799889534 | 0.964 | 0.999 | 4.5689925 |
| **H1FX** | 4.8851 | -0.96373743646835 | 0.997 | 1 | 8.0004533 |
| **HIP1** | 5.2671 | -1.15150298046636 | 0.699 | 0.975 | 8.6260160 |
| **TBL2** | 6.2971 | -1.23988068302774 | 0.552 | 0.943 | 1.0312841 |

| | | | | | |
|---|---|---|---|---|---|
| **LIPH** | 6.3622 | 0.947235653962666 | 0.426 | 0.032 | 1.0419408 |
| **ZIC3** | 6.9992 | -1.54664203381906 | 0.545 | 0.942 | 1.1462732 |
| **MTOR** | 1.1590 | 0.915713709187111 | 0.997 | 0.97 | 1.8981267 |
| **PPP1R14A** | 1.4829 | 1.10337521342403 | 0.998 | 0.898 | 2.4286835 |
| **TTI1** | 1.6889 | -1.03233734757615 | 0.897 | 0.994 | 2.7659386 |
| **CPXM1** | 2.1986 | -1.44413010544002 | 0.794 | 0.984 | 3.6006887 |
| **NOTCH2** | 2.3076 | -1.25841097996239 | 0.497 | 0.922 | 3.7791583 |
| **TAF1D** | 3.4495 | 1.12645025637816 | 1 | 0.975 | 5.6492480 |
| **CEP68** | 3.5295 | -1.20922756861945 | 0.589 | 0.957 | 5.7804207 |
| **KLF12** | 4.6515 | -1.42258454168546 | 0.117 | 0.777 | 7.6178637 |
| **AJUBA** | 5.7363 | -1.4825989849233 | 0.665 | 0.976 | 9.3944121 |
| **AGPAT1** | 6.0620 | -0.94706967784066 | 0.983 | 1 | 9.9278309 |
| **SDCBP** | 2.4494 | 0.964592374002049 | 0.997 | 0.974 | 4.0114214 |
| **MFSD13A** | 2.5046 | 1.21503049819075 | 0.805 | 0.293 | 4.1017883 |
| **CACHD1** | 3.5108 | -1.65559282880392 | 0.621 | 0.966 | 5.7496727 |
| **ESD** | 3.7662 | -0.83707225790381 | 0.992 | 0.999 | 6.1680414 |
| **FAM20C** | 4.8336 | -1.4626923495897 | 0.409 | 0.9 | 7.9160844 |
| **CTXN1** | 4.9140 | -1.3828723819584 | 0.804 | 0.98 | 8.0476837 |
| **SLC35F6** | 8.7840 | 1.0609817092818 | 0.997 | 0.97 | 1.4385647 |
| **GOLM1** | 4.4235 | -0.96570772945217 | 0.985 | 1 | 7.2443931 |
| **RARG** | 5.4221 | -1.56496279864262 | 0.35 | 0.862 | 8.8798742 |
| **MFSD10** | 6.2498 | -1.20364681203441 | 0.624 | 0.961 | 1.0235398 |
| **PRRX2** | 9.3341 | 1.25255269114388 | 0.792 | 0.27 | 1.5286469 |
| **PLEKHB2** | 1.1270 | 0.867765299329828 | 1 | 0.999 | 1.8457682 |
| **PTPN11** | 1.1964 | 1.02252805830593 | 0.951 | 0.576 | 1.9594578 |
| **SLC30A5** | 1.8606 | 1.08558110584415 | 0.956 | 0.626 | 3.0471509 |
| **TMEM126B** | 3.2308 | 1.12328430421394 | 0.848 | 0.358 | 5.2912420 |
| **KCNK6** | 3.3112 | 0.985051931167398 | 0.665 | 0.148 | 5.4228320 |
| **CHST8** | 4.0373 | 1.08129235028302 | 0.64 | 0.14 | 6.6119995 |
| **CHSY1** | 4.2748 | -1.13585739317608 | 0.973 | 1 | 7.0009511 |
| **CCNB1IP1** | 5.3248 | 0.993811010452326 | 1 | 0.979 | 8.7205154 |
| **EVL** | 6.2551 | -1.00469706692623 | 0.937 | 1 | 1.0244012 |
| **ZNF808** | 7.3833 | 1.11982758617211 | 0.99 | 0.835 | 1.2091776 |
| **OST4** | 2.1243 | 1.0118344886048 | 0.777 | 0.259 | 3.4790578 |

| | | | | | |
|---|---|---|---|---|---|
| **DDX3Y** | 2.5760 | 1.18323958084691 | 0.968 | 0.822 | 4.2187831 |
| **EFS** | 4.0690 | -1.45277572205304 | 0.398 | 0.891 | 6.6638016 |
| **GHITM** | 4.3037 | 0.804630475912741 | 1 | 0.999 | 7.0482363 |
| **ZNF358** | 4.7797 | -1.05072943245522 | 0.871 | 0.996 | 7.8277159 |
| **DLAT** | 6.4404 | 0.806767713220613 | 1 | 0.998 | 1.0547571 |
| **GSPT2** | 6.5423 | -1.23958183728252 | 0.511 | 0.961 | 1.0714380 |
| **SLC35A5** | 8.4232 | 1.17033363591515 | 0.971 | 0.837 | 1.3794781 |
| **CALB1** | 9.2608 | 1.1321998521829 | 0.65 | 0.132 | 1.5166420 |
| **KIAA1522** | 4.4133 | -1.1674954141889 | 0.956 | 0.991 | 7.2277854 |
| **GPD2** | 5.2523 | -1.17029325931209 | 0.878 | 0.993 | 8.6016970 |
| **SCD** | 5.4527 | -0.85829664139419 | 1 | 1 | 8.9299081 |
| **ASPH** | 5.8194 | -1.0571788570801 | 0.91 | 0.997 | 9.5304947 |
| **EPB41L4B** | 6.5513 | 1.23933633946339 | 0.931 | 0.556 | 1.0729179 |
| **PCCA** | 1.0173 | -1.10429409540143 | 0.942 | 0.998 | 1.6661588 |
| **DUSP6** | 1.0223 | 1.1240690561408 | 0.513 | 0.069 | 1.6742768 |
| **ZNF552** | 1.2112 | 1.22531830759002 | 0.978 | 0.694 | 1.9836052 |
| **TTC3** | 1.8082 | -1.05989671855401 | 0.863 | 0.995 | 2.9613856 |
| **MYL12A** | 7.9758 | 1.11616876101044 | 0.956 | 0.654 | 1.3062039 |
| **CHCHD3** | 8.7214 | 0.887141505651419 | 1 | 0.997 | 1.4283166 |
| **B4GALT5** | 9.1686 | -0.99481184647296 | 0.964 | 1 | 1.5015571 |
| **MBNL3** | 4.0453 | -1.501089786333 | 0.482 | 0.944 | 6.6251129 |
| **ZFAND3** | 4.5002 | -0.89582756867101 | 0.958 | 0.999 | 7.3700208 |
| **MYO9A** | 8.5270 | 1.19670525892711 | 0.882 | 0.471 | 1.3964759 |
| **TIMP2** | 1.2703 | -1.35987634614699 | 0.765 | 0.973 | 2.0803846 |
| **ATP8B2** | 1.4470 | 0.999759794776322 | 0.992 | 0.953 | 2.3697634 |
| **AGBL5** | 1.6929 | -1.13495693686497 | 0.733 | 0.978 | 2.7725737 |
| **SLC16A9** | 2.8266 | 1.31846590789289 | 0.875 | 0.455 | 4.6291319 |
| **YAF2** | 3.3052 | -1.30042405232601 | 0.122 | 0.768 | 5.4130750 |
| **SNX2** | 4.4361 | 0.845723587516275 | 1 | 0.999 | 7.2650278 |
| **MFSD4B** | 4.9981 | 1.25975905077103 | 0.927 | 0.625 | 8.1855237 |
| **ATXN7L3B** | 5.3477 | -0.89597011111693 | 0.978 | 1 | 8.7579713 |
| **MAP4K1** | 5.5836 | 1.14112296359623 | 0.787 | 0.268 | 9.1442810 |
| **MYL7** | 6.2850 | -2.66108956502564 | 0.217 | 0.816 | 1.0293082 |
| **KCNQ2** | 7.3209 | 1.08657975092204 | 0.844 | 0.312 | 1.1989489 |

| | | | | | |
|---|---|---|---|---|---|
| **MANBAL** | 9.3149 | -0.82976950367643 | 0.983 | 1 | 1.5255094 |
| **IGF2BP3** | 9.5317 | -0.81767221854700 | 0.99 | 1 | 1.5610197 |
| **ZC4H2** | 1.4561 | -1.21666046350469 | 0.318 | 0.86 | 2.3848162 |
| **DCAF5** | 2.1791 | -1.14780488040449 | 0.67 | 0.976 | 3.5687487 |
| **ACVR2B** | 2.3391 | -0.82108907455683 | 0.995 | 1 | 3.8307834 |
| **CFD** | 2.7139 | 0.872945211368882 | 0.558 | 0.093 | 4.4446427 |
| **PPIB** | 2.8485 | 0.817499123962309 | 1 | 0.999 | 4.6650765 |
| **CPD** | 3.1329 | -1.03440803784551 | 0.958 | 0.998 | 5.1308388 |
| **TTYH1** | 3.2055 | 1.10547224351389 | 0.519 | 0.076 | 5.2496838 |
| **PYGB** | 3.7789 | 1.38308274656354 | 1 | 0.999 | 6.1888549 |
| **BCAS4** | 4.4722 | -1.21231833664312 | 0.607 | 0.945 | 7.3241607 |
| **CA11** | 5.8503 | -1.53012228072522 | 0.083 | 0.725 | 9.5811358 |
| **HK1** | 7.9971 | -1.04764408219998 | 0.986 | 1 | 1.3096939 |
| **TRAPPC6A** | 1.0497 | -1.22212702810037 | 0.619 | 0.953 | 1.7191988 |
| **PEAK1** | 1.3914 | -1.28685188704084 | 0.32 | 0.863 | 2.2787562 |
| **CHCHD4** | 1.6089 | 1.11428982402771 | 0.975 | 0.766 | 2.6350476 |
| **CRISPLD2** | 1.8418 | 1.08229399721543 | 0.746 | 0.225 | 3.0164106 |
| **RRAD** | 2.0547 | 1.46374384025096 | 0.401 | 0.031 | 3.3649897 |
| **CNTRL** | 2.8426 | -1.30357125219307 | 0.411 | 0.891 | 4.6553936 |
| **WNT5A** | 4.3597 | -1.53527548910486 | 0.25 | 0.901 | 7.1400343 |
| **TAF9B** | 4.3976 | -1.18185576488211 | 0.533 | 0.928 | 7.2020786 |
| **MYL6** | 4.5355 | -0.80746376444454 | 1 | 1 | 7.4278264 |
| **MSI2** | 5.6081 | -1.18992186868486 | 0.587 | 0.959 | 9.1844432 |
| **NLGN4X** | 7.1510 | -2.10973972727197 | 0.191 | 0.79 | 1.1711243 |
| **CD320** | 1.0070 | 1.13912281802339 | 1 | 0.998 | 1.6493175 |
| **CASK** | 1.2447 | -0.95436368058315 | 0.939 | 0.999 | 2.0385395 |
| **UBE2G2** | 1.3409 | -1.07801092849712 | 0.729 | 0.976 | 2.1961538 |
| **FLNC** | 1.5482 | -1.43397641146268 | 0.878 | 0.994 | 2.5354925 |
| **NOB1** | 1.6430 | 0.912168739157919 | 0.997 | 0.959 | 2.6909046 |
| **SLC25A24** | 2.2023 | -1.03401240880877 | 0.883 | 0.996 | 3.6067405 |
| **MXRA8** | 2.2433 | -1.33006390582223 | 0.58 | 0.959 | 3.6739229 |
| **EEF1A2** | 3.6326 | 0.818605935967584 | 0.543 | 0.086 | 5.9491691 |
| **RRAS2** | 4.2549 | 0.948326628030545 | 1 | 0.955 | 6.9683337 |
| **CHCHD7** | 4.3508 | 0.968411708895974 | 0.997 | 0.931 | 7.1253627 |

| | | | | | |
|---|---|---|---|---|---|
| **TOMM34** | 7.4299 | -1.05215204000527 | 0.849 | 0.991 | 1.21680347 |
| **TSPAN18** | 9.5962 | -1.67884359990585 | 0.868 | 0.973 | 1.57157365 |
| **SERINC5** | 2.3478 | 0.968736245884377 | 1 | 0.987 | 3.84510481 |
| **TMEM189** | 4.1581 | -0.80469447056708 | 0.998 | 1 | 6.80981573 |
| **TRIM5** | 4.7302 | -1.23143072686386 | 0.384 | 0.888 | 7.74675470 |
| **TMEM154** | 4.7806 | 1.34299043177924 | 0.821 | 0.336 | 7.82928831 |
| **TGFBR3** | 5.0235 | 1.2336175827166 | 0.792 | 0.305 | 8.22710879 |
| **RTL8A** | 6.5874 | -1.30088646693748 | 0.215 | 0.8 | 1.07882917 |
| **WNK3** | 7.3798 | -1.18458891405659 | 0.47 | 0.922 | 1.20859147 |
| **CLDN19** | 7.7647 | -1.47767540500759 | 0.088 | 0.722 | 1.27163773 |
| **ARMCX1** | 8.4348 | -1.26710015152763 | 0.43 | 0.937 | 1.38137560 |
| **ALCAM** | 1.0953 | -1.72545440398228 | 0.19 | 0.781 | 1.79387531 |
| **ISCA1** | 1.2156 | 1.01633213179569 | 0.795 | 0.293 | 1.99082598 |
| **APOBEC3D** | 1.4306 | 0.900938242717492 | 0.599 | 0.122 | 2.34289930 |
| **CETN3** | 1.9315 | 1.13692982786638 | 0.975 | 0.807 | 3.16323610 |
| **B3GNT2** | 2.7950 | -1.37405478776669 | 0.897 | 0.989 | 4.57746813 |
| **ESR1** | 2.9691 | 1.11380409363079 | 0.995 | 0.971 | 4.86249722 |
| **FERMT1** | 3.5154 | 0.936068805303615 | 0.609 | 0.125 | 5.75719001 |
| **CSNK1G2** | 3.6832 | -0.84308981722352 | 0.985 | 1 | 6.03209720 |
| **S100A11** | 4.6068 | 0.848917455996231 | 1 | 1 | 7.54459547 |
| **PREX2** | 9.2529 | -1.46089210693402 | 0.269 | 0.818 | 1.51535169 |
| **CASC4** | 1.1568 | 0.878186349089576 | 0.667 | 0.167 | 1.89458103 |
| **LDLRAP1** | 2.0317 | 1.12412712906834 | 0.985 | 0.884 | 3.32732494 |
| **PLCG1** | 3.9299 | -0.81765803916996 | 0.986 | 1 | 6.43604909 |
| **IDH2** | 4.1510 | -1.14735635869104 | 0.873 | 0.998 | 6.79810581 |
| **FAM122B** | 6.0554 | -1.29335885239169 | 0.685 | 0.965 | 9.91696255 |
| **EFNA5** | 8.3087 | -1.35610224566525 | 0.372 | 0.875 | 1.36072517 |
| **SNX1** | 9.6352 | -0.99705509312084 | 0.912 | 0.996 | 1.57796697 |
| **TMEM185A** | 1.4431 | -1.17405098261879 | 0.765 | 0.99 | 2.36346848 |
| **HMGB2** | 1.5885 | -1.0989583924651 | 0.997 | 0.999 | 2.60162713 |
| **MCC** | 1.6737 | -1.4621147776464 | 0.389 | 0.888 | 2.74109644 |
| **BMPR1A** | 1.7466 | 1.00396624985317 | 0.997 | 0.943 | 2.86054850 |
| **PSPC1** | 3.4676 | 0.854834103945848 | 1 | 0.997 | 5.67904200 |
| **FBLN1** | 9.5318 | -0.98971668247011 | 0.986 | 1 | 1.56103853 |

| | | | | | |
|---|---|---|---|---|---|
| **TCEAL8** | 1.0643 | -0.98674530922835 | 0.948 | 1 | 1.74301585 |
| **PKIG** | 1.3434 | -1.13073330523762 | 0.86 | 0.988 | 2.2001089 |
| **PDS5A** | 1.4080 | 0.9129616611558 | 0.997 | 0.843 | 2.3060314 |
| **TRO** | 1.4498 | -1.20282979742129 | 0.492 | 0.95 | 2.3743948 |
| **CDYL** | 1.6980 | -1.07913013051237 | 0.724 | 0.978 | 2.7808646 |
| **PLAAT5** | 2.6489 | -1.46489694033009 | 0.269 | 0.828 | 4.3381068 |
| **GNPAT** | 3.4424 | -1.08989162533146 | 0.679 | 0.973 | 5.6377146 |
| **TIA1** | 6.5844 | 0.804335266831491 | 1 | 0.982 | 1.0783286 |
| **UBE2J1** | 1.2683 | -1.0738475255787 | 0.751 | 0.991 | 2.0772204 |
| **GPC4** | 2.9682 | -1.42257113957075 | 0.821 | 1 | 4.8610235 |
| **SPC24** | 3.0046 | -1.08974379855766 | 0.834 | 0.981 | 4.9207640 |
| **CCNQ** | 3.0762 | 1.03953504835408 | 0.975 | 0.809 | 5.0379373 |
| **NET1** | 4.8734 | -1.33779331099801 | 0.77 | 0.984 | 7.9813234 |
| **PTPRU** | 5.0077 | 1.23269068799337 | 0.988 | 0.909 | 8.2012452 |
| **TNFAIP2** | 5.4146 | -1.79655558346894 | 0.215 | 0.785 | 8.8676150 |
| **DPY19L1** | 5.5332 | 1.22997538537362 | 0.89 | 0.49 | 9.0618525 |
| **OVOL2** | 6.4604 | 1.37941623943305 | 0.816 | 0.34 | 1.0580208 |
| **PSME3** | 6.5182 | 0.949849322716295 | 0.964 | 0.718 | 1.0675017 |
| **ACSL4** | 2.3400 | -1.14035002441547 | 0.785 | 0.996 | 3.8323730 |
| **TWSG1** | 2.5592 | -1.12651799745223 | 0.844 | 0.987 | 4.1913507 |
| **BACE2** | 3.2726 | -1.36486475714396 | 0.372 | 0.886 | 5.3596516 |
| **KIF26A** | 4.4202 | -1.07219244452985 | 0.865 | 0.996 | 7.2390851 |
| **NELFE** | 8.2835 | 1.00400635609484 | 0.817 | 0.335 | 1.3565984 |
| **LIPA** | 8.3620 | -1.25947826207976 | 0.453 | 0.896 | 1.3694602 |
| **LEFTY2** | 1.4901 | 1.91195819874906 | 0.981 | 0.945 | 2.4404758 |
| **TMED5** | 1.5441 | 0.923254196616881 | 0.997 | 0.925 | 2.5289255 |
| **RAB15** | 1.8111 | 1.09528947361971 | 1 | 0.993 | 2.9661348 |
| **PTPN14** | 2.1237 | -0.99854614468305 | 0.931 | 0.998 | 3.4779860 |
| **SRXN1** | 4.1734 | -1.3221462453614 | 0.692 | 0.961 | 6.8348856 |
| **UBASH3B** | 5.0117 | 1.23775232958517 | 0.907 | 0.553 | 8.2076708 |
| **JADE2** | 8.1406 | 1.19875186155502 | 0.986 | 0.895 | 1.3331968 |
| **PAK1** | 8.5673 | 1.09956487076948 | 0.983 | 0.897 | 1.4030720 |
| **IPMK** | 8.8194 | 1.10937988803336 | 0.954 | 0.787 | 1.4443601 |
| **ANP32B** | 9.0877 | -0.94862502917558 | 0.981 | 1 | 1.4883042 |

| | | | | | |
|---|---|---|---|---|---|
| **CAMK2D** | 1.0846 | -1.45600328272144 | 0.178 | 0.766 | 1.7763788 |
| **BCAR1** | 1.7723 | -1.06785232871715 | 0.963 | 0.997 | 2.9026073 |
| **CDIP1** | 2.2014 | -1.08307418019459 | 0.839 | 0.988 | 3.6053009 |
| **NPC2** | 4.1666 | -0.91489967298709 | 0.998 | 1 | 6.8236645 |
| **SNW1** | 4.8913 | 1.02915196132498 | 0.968 | 0.772 | 8.0105623 |
| **KRTCAP3** | 5.0391 | 0.925914908248705 | 0.694 | 0.188 | 8.2526217 |
| **SOGA1** | 5.6402 | -1.14704935191658 | 0.761 | 0.982 | 9.2370881 |
| **OBI1** | 5.8876 | -1.1213663623422 | 0.685 | 0.952 | 9.6422476 |
| **LINGO1** | 7.0158 | 1.07193319760271 | 0.993 | 0.854 | 1.1489843 |
| **ECPAS** | 9.8415 | 0.827507805008254 | 1 | 0.996 | 1.6117543 |
| **ST13** | 1.3913 | 0.931059176770827 | 0.981 | 0.811 | 2.2786310 |
| **PHC2** | 1.6514 | -1.60049286109835 | 0.098 | 0.736 | 2.7046199 |
| **CTNNBL1** | 2.1569 | -0.82588985284101 | 0.975 | 1 | 3.5324349 |
| **APOBEC3G** | 2.6537 | 1.07808381918039 | 0.658 | 0.167 | 4.3461128 |
| **TTYH3** | 3.1222 | -0.88491118767496 | 0.995 | 1 | 5.1133765 |
| **NOL7** | 3.5159 | 0.841097903755422 | 0.995 | 0.985 | 5.7580945 |
| **GRID2** | 4.4077 | -1.72750689326545 | 0.176 | 0.765 | 7.2185142 |
| **IGFBP2** | 7.0106 | -0.99194521118267 | 0.988 | 1 | 1.1481266 |
| **PHTF1** | 9.8448 | -1.11123742640308 | 0.667 | 0.959 | 1.6122985 |
| **RBMS1** | 1.3610 | 1.16922035934097 | 0.964 | 0.763 | 2.2289577 |
| **HAS3** | 2.2344 | -1.88215464016005 | 0.355 | 0.827 | 3.6593268 |
| **SHISA5** | 3.5312 | 0.820362741809401 | 1 | 0.999 | 5.7832081 |
| **ZNF580** | 9.0773 | -1.21796240107513 | 0.421 | 0.895 | 1.4865977 |
| **NOP9** | 1.3558 | 1.02645042556145 | 0.948 | 0.682 | 2.2205353 |
| **EML4** | 1.5848 | -0.80958528397854 | 0.99 | 1 | 2.5955058 |
| **SCAND1** | 2.3407 | -0.88763611581480 | 0.976 | 1 | 3.8334066 |
| **KLHDC10** | 2.6358 | -1.14289439915254 | 0.161 | 0.748 | 4.3167926 |
| **SHISA9** | 3.6764 | -1.53698375212149 | 0.147 | 0.746 | 6.0209549 |
| **HIPK2** | 8.5512 | -1.23980301795116 | 0.565 | 0.919 | 1.4004413 |
| **CAP2** | 2.5914 | -1.16850990354584 | 0.563 | 0.947 | 4.2439586 |
| **CCDC102A** | 3.7703 | -1.19274612670151 | 0.333 | 0.841 | 6.1747471 |
| **MTFR1** | 4.0469 | 0.985350473030099 | 0.981 | 0.872 | 6.6276797 |
| **RASGRF2** | 4.0920 | 1.21778839063024 | 0.682 | 0.187 | 6.7016092 |
| **RRP12** | 4.1677 | 0.950483636654222 | 0.992 | 0.898 | 6.8256038 |

| | | | | | |
|---|---|---|---|---|---|
| **PTPRA** | 4.7150 | -0.95386819660715 | 0.875 | 0.987 | 7.7219120 |
| **DMXL1** | 8.8687 | 1.13028927112008 | 0.936 | 0.672 | 1.4524333 |
| **C1orf43** | 1.0345 | -0.8639722503565 | 0.944 | 0.998 | 1.6943081 |
| **WASL** | 1.5283 | 0.938071288696811 | 0.998 | 0.99 | 2.5030112 |
| **SOX4** | 1.6081 | 0.973209578392687 | 1 | 0.993 | 2.6336453 |
| **CAST** | 1.6266 | -1.20934557395397 | 0.218 | 0.774 | 2.6638974 |
| **IMPA2** | 1.8056 | 0.994190664890682 | 0.998 | 0.975 | 2.9571198 |
| **DRAXIN** | 2.2168 | -1.63494135587909 | 0.731 | 0.95 | 3.6305098 |
| **SLC23A2** | 2.3202 | 1.22211304604713 | 0.964 | 0.845 | 3.7998283 |
| **ABCB8** | 2.3693 | 0.960690780304631 | 0.986 | 0.891 | 3.8802320 |
| **STC2** | 6.9111 | 1.67798525282273 | 0.975 | 0.924 | 1.1318442 |
| **NGFR** | 7.5206 | -1.6981991942892 | 0.403 | 0.85 | 1.2316582 |
| **CTSV** | 9.8455 | -1.44833783583537 | 0.773 | 0.971 | 1.6124138 |
| **EIF1AX** | 1.1481 | -0.84934872994654 | 0.981 | 0.999 | 1.8802466 |
| **CAPN13** | 2.6949 | -1.25566558583392 | 0.112 | 0.715 | 4.4135779 |
| **VAT1L** | 2.7307 | -2.40437308320498 | 0.027 | 0.644 | 4.4721935 |
| **CIZ1** | 4.2136 | -1.07307576128687 | 0.533 | 0.929 | 6.9006214 |
| **DLGAP4** | 4.3207 | -0.93411303456014 | 0.932 | 0.996 | 7.0760229 |
| **XIAP** | 7.4929 | 1.09285357066041 | 0.917 | 0.635 | 1.2271239 |
| **ARPIN** | 8.1028 | -1.20035998534945 | 0.279 | 0.816 | 1.3270060 |
| **PIR** | 9.0260 | -1.13189411037429 | 0.66 | 0.943 | 1.4781961 |
| **ABCE1** | 9.1763 | 0.910398526103818 | 0.995 | 0.874 | 1.5028089 |
| **PTGIS** | 9.6008 | -1.33081195686047 | 0.125 | 0.716 | 1.5723251 |
| **ELF3** | 9.6832 | 1.51204647817796 | 0.799 | 0.366 | 1.5858329 |
| **OLFM1** | 1.1492 | -1.44121107871865 | 0.552 | 0.913 | 1.8820702 |
| **PPME1** | 1.1894 | -1.07011454766182 | 0.711 | 0.959 | 1.9479077 |
| **MIS18A** | 1.2168 | 0.848902465555619 | 0.995 | 0.944 | 1.9928634 |
| **NR5A2** | 2.6164 | 1.11120901212607 | 0.614 | 0.152 | 4.2850245 |
| **FGD4** | 2.8647 | 1.1119550145177 | 0.959 | 0.708 | 4.6915518 |
| **RFLNB** | 2.9088 | 1.4533399891132 | 0.941 | 0.676 | 4.7637857 |
| **APMAP** | 3.5043 | -0.87729187422826 | 0.954 | 1 | 5.7391077 |
| **NDUFA8** | 4.3487 | 0.877631222400749 | 1 | 0.998 | 7.1219406 |
| **NSD2** | 4.7009 | -1.12937993334585 | 0.469 | 0.892 | 7.6988028 |
| **IRX1** | 4.9668 | -1.58480091606771 | 0.156 | 0.737 | 8.1341376 |

| | | | | | |
|---|---|---|---|---|---|
| **ANK2** | 2.1714 | -1.17812278650241 | 0.594 | 0.933 | 3.5561983 |
| **SLC7A7** | 3.5221 | 1.50463986308114 | 0.89 | 0.57 | 5.7682168 |
| **KCNG1** | 3.5872 | -1.61415668951531 | 0.316 | 0.821 | 5.8748756 |
| **LYSMD3** | 4.5763 | 1.06804847849083 | 0.817 | 0.388 | 7.4947338 |
| **MLLT11** | 6.0869 | -1.22927412887207 | 0.563 | 0.92 | 9.9685215 |
| **INA** | 6.7023 | 1.22668307454534 | 0.934 | 0.614 | 1.0976463 |
| **DUSP7** | 8.8437 | 1.04813096182799 | 0.976 | 0.77 | 1.4483346 |
| **ABHD12B** | 1.0695 | 0.886396532699 | 0.602 | 0.133 | 1.7515941 |
| **ZNF229** | 1.2660 | 0.997798146779599 | 0.736 | 0.255 | 2.0734219 |
| **PBX4** | 1.6416 | 0.900815620121293 | 0.607 | 0.144 | 2.6884495 |
| **THOC5** | 1.7738 | 0.851044648705328 | 1 | 0.983 | 2.9049709 |
| **RBM25** | 3.0602 | 0.809318953785617 | 1 | 0.999 | 5.0117568 |
| **CHST2** | 3.3962 | 1.59846296697987 | 0.97 | 0.723 | 5.5620064 |
| **ZCCHC24** | 4.6509 | -1.27736173871073 | 0.408 | 0.864 | 7.6169124 |
| **OSBPL10** | 5.9898 | 1.14609225984411 | 0.961 | 0.764 | 9.8096415 |
| **AKT3** | 8.3212 | -1.21198329610765 | 0.433 | 0.88 | 1.3627683 |
| **DUSP16** | 1.4769 | 1.47050919585267 | 0.948 | 0.763 | 2.4188465 |
| **RBBP7** | 1.6607 | -0.90293013103586 | 0.929 | 0.997 | 2.7197610 |
| **FGD1** | 2.4804 | -0.94690203344292 | 0.854 | 0.994 | 4.0622015 |
| **GTF2H1** | 2.6465 | 0.908608505601722 | 0.993 | 0.894 | 4.3342361 |
| **TNNC1** | 3.7246 | -1.4000347332993 | 0.124 | 0.704 | 6.0998250 |
| **CHST11** | 4.4001 | -1.23966248561435 | 0.697 | 0.971 | 7.2060618 |
| **COTL1** | 4.7167 | -1.16002050699353 | 0.964 | 0.997 | 7.7246798 |
| **ESRP1** | 7.4820 | 0.988363977712936 | 0.997 | 0.877 | 1.2253346 |
| **TM7SF2** | 1.1550 | -1.33885754126479 | 0.782 | 0.952 | 1.8916293 |
| **PPIL4** | 1.4350 | 0.951856750763378 | 0.953 | 0.734 | 2.3501756 |
| **SENP2** | 1.4396 | 0.944759515217425 | 0.997 | 0.992 | 2.3576588 |
| **LPIN1** | 2.3668 | 1.10131248894218 | 0.809 | 0.323 | 3.8761768 |
| **AP1M1** | 2.4833 | -0.93432559406099 | 0.902 | 0.992 | 4.0669500 |
| **TRERF1** | 4.5023 | -1.20770529221723 | 0.068 | 0.668 | 7.3735453 |
| **CIB1** | 6.6073 | 0.943825256587839 | 0.968 | 0.833 | 1.0820836 |
| **ARL8B** | 7.9964 | 0.88679559023082 | 0.995 | 0.92 | 1.3095832 |
| **ST6GAL1** | 8.3997 | 0.857087209150321 | 1 | 0.998 | 1.3756325 |
| **JAG2** | 1.2357 | 1.06600159301264 | 0.707 | 0.233 | 2.0238328 |

| | | | | | |
|---|---|---|---|---|---|
| **SLC38A1** | 1.6027 | -1.03162856967217 | 0.853 | 0.995 | 2.6248496 |
| **LDB2** | 2.0842 | -1.66921531770021 | 0.501 | 0.882 | 3.4132968 |
| **CTDSPL** | 2.2865 | 1.28891554893801 | 0.946 | 0.809 | 3.7446111 |
| **GFPT2** | 2.6505 | -1.24920466327187 | 0.734 | 0.983 | 4.3408846 |
| **GPC1** | 3.2376 | -0.97661520095390 | 0.848 | 0.989 | 5.3023117 |
| **AGPAT2** | 3.4251 | 1.09382793489568 | 0.99 | 0.957 | 5.6094239 |
| **AMER1** | 1.0454 | -1.08889223579257 | 0.543 | 0.928 | 1.7121090 |
| **ZSWIM6** | 1.1848 | -1.2934600862687 | 0.181 | 0.745 | 1.9403902 |
| **RAVER2** | 1.2078 | -1.05732879161067 | 0.457 | 0.879 | 1.9780179 |
| **PTPRZ1** | 1.4225 | -2.08467048462987 | 0.503 | 0.863 | 2.3297778 |
| **COL4A6** | 1.5140 | -1.31828091815993 | 0.052 | 0.646 | 2.4795129 |
| **PRKAB2** | 1.5788 | 1.07804992352712 | 0.966 | 0.819 | 2.5857150 |
| **IGLON5** | 1.6052 | -1.19661918016871 | 0.245 | 0.794 | 2.6289133 |
| **MBOAT7** | 1.6364 | -0.81386017108618 | 0.968 | 0.999 | 2.6800340 |
| **C2CD5** | 1.7213 | 1.04296486052088 | 0.983 | 0.93 | 2.8191102 |
| **ABCA1** | 2.2684 | 1.32000925224119 | 0.934 | 0.633 | 3.7149890 |
| **NRIP1** | 2.5264 | -1.12992745784542 | 0.528 | 0.93 | 4.1375809 |
| **LSS** | 2.9786 | -0.92632186972149 | 0.973 | 0.998 | 4.8781672 |
| **NRK** | 3.3078 | -2.63251717389167 | 0.288 | 0.795 | 5.4172009 |
| **UCP2** | 5.3384 | 1.0330960400162 | 0.992 | 0.919 | 8.7427565 |
| **CBX5** | 6.1482 | -0.87124571503351 | 0.971 | 0.999 | 1.0069003 |
| **GDI1** | 6.6259 | -0.82196180747788 | 0.993 | 1 | 1.0851272 |
| **B4GALT2** | 7.3265 | -0.88803056409277 | 0.898 | 0.991 | 1.1998622 |
| **EPPK1** | 9.6536 | 1.2356219313642 | 0.992 | 0.82 | 1.5809773 |
| **DEPDC1** | 1.0820 | 1.31470308442447 | 0.731 | 0.273 | 1.7721219 |
| **ARL4C** | 1.1707 | -1.38724182580704 | 0.299 | 0.805 | 1.9173660 |
| **NUDT15** | 1.4615 | 0.807227312132747 | 1 | 0.999 | 2.3935396 |
| **PKP4** | 1.5495 | -0.99496923226346 | 0.623 | 0.928 | 2.5376895 |
| **TMEM165** | 1.7573 | 0.864461997565696 | 0.973 | 0.782 | 2.8779783 |
| **VASH2** | 1.7595 | 1.12717595259341 | 0.993 | 0.963 | 2.8816912 |
| **CRABP1** | 1.8844 | -2.02389493970184 | 0.459 | 0.879 | 3.0860869 |
| **ARFGAP2** | 3.1663 | 0.878577354152657 | 0.981 | 0.874 | 5.1854828 |
| **FOXO4** | 3.8991 | -1.57075382758708 | 0.614 | 0.925 | 6.3857140 |
| **LPP** | 4.6797 | -1.19751255529878 | 0.716 | 0.966 | 7.6640276 |

| | | | | | |
|---|---|---|---|---|---|
| **VANGL1** | 5.3824 | -1.21656964057185 | 0.768 | 0.973 | 8.81490958 |
| **MARVELD3** | 9.7609 | 0.879526676039699 | 0.689 | 0.2 | 1.59854274 |
| **RET** | 1.0269 | -1.37614409263744 | 0.315 | 0.826 | 1.68189288 |
| **TCEA2** | 1.0529 | -1.09301345611984 | 0.555 | 0.92 | 1.72437560 |
| **CXCL14** | 1.0553 | -2.71162744459152 | 0.431 | 0.85 | 1.72831353 |
| **LIMD2** | 1.6988 | -1.06807748474379 | 0.829 | 0.982 | 2.78225944 |
| **DSP** | 2.2875 | -0.90157411707901 | 0.998 | 1 | 3.74636435 |
| **MED12** | 2.4627 | -1.15309373804575 | 0.284 | 0.804 | 4.03327602 |
| **TMEM45A** | 3.4587 | -1.26182386041116 | 0.352 | 0.844 | 5.66442390 |
| **WDR55** | 4.0820 | 0.955308429510664 | 0.995 | 0.921 | 6.68514114 |
| **PSME1** | 6.9825 | -0.94141759282159 | 0.856 | 0.99 | 1.14353239 |
| **TNFRSF21** | 2.4083 | -1.27233562913205 | 0.959 | 0.996 | 3.94417755 |
| **ECE1** | 3.5180 | -1.02706789927815 | 0.937 | 0.997 | 5.76143285 |
| **PDZD2** | 4.2723 | 0.984917908275616 | 0.741 | 0.262 | 6.99678052 |
| **LRP8** | 6.0262 | -1.36458484315581 | 0.824 | 0.972 | 9.86921178 |
| **SYNGR3** | 8.5892 | 1.12247596127053 | 0.888 | 0.566 | 1.40665690 |
| **TBC1D8** | 9.1606 | 1.00588558074115 | 0.783 | 0.321 | 1.50024405 |
| **BZW2** | 1.0166 | 0.800157436712379 | 1 | 0.988 | 1.66504355 |
| **NT5C2** | 3.2436 | -1.0218574519895 | 0.897 | 0.988 | 5.31217381 |
| **UBE2F** | 4.0816 | 0.902422869259977 | 0.942 | 0.715 | 6.68448610 |
| **FOXB1** | 5.4863 | -1.79248527727733 | 0.262 | 0.765 | 8.98491890 |
| **KLHDC7A** | 6.0836 | -1.46424319798243 | 0.049 | 0.634 | 9.96325691 |
| **ASCC2** | 8.7806 | 0.808149204783281 | 0.993 | 0.949 | 1.43801461 |
| **SLC6A6** | 1.6235 | 1.08369405528529 | 0.995 | 0.91 | 2.65888231 |
| **TENM4** | 1.6690 | -1.42380508090222 | 0.31 | 0.819 | 2.73343918 |
| **NUPR1** | 1.9502 | 1.71746130889286 | 0.651 | 0.208 | 3.19390220 |
| **DCLK1** | 2.0791 | -1.46500961365933 | 0.931 | 0.961 | 3.40505323 |
| **LITAF** | 2.4499 | -1.16539985404388 | 0.993 | 1 | 4.01226780 |
| **ASCC3** | 2.6531 | 0.881530046728353 | 0.995 | 0.919 | 4.34507602 |
| **ADAM10** | 2.7320 | 0.867863290979138 | 0.993 | 0.869 | 4.47423202 |
| **FST** | 3.2304 | -2.61990560782783 | 0.23 | 0.743 | 5.29044430 |
| **SPDL1** | 3.8934 | 1.00281570577041 | 0.819 | 0.387 | 6.37632747 |
| **PDE4A** | 5.6929 | 1.05699409927894 | 0.741 | 0.275 | 9.32326341 |
| **TAOK3** | 5.7689 | 0.988574185358677 | 0.733 | 0.267 | 9.44774112 |

| | | | | | |
|---|---|---|---|---|---|
| **GCNT2** | 5.7753 | 0.96763758878886 | 0.983 | 0.963 | 9.4583181 |
| **XXYLT1** | 6.5656 | -0.93117753480962 | 0.761 | 0.969 | 1.0752537 |
| **SEMA4B** | 7.8243 | 0.936936713044571 | 0.98 | 0.82 | 1.2813966 |
| **LRRC42** | 1.6697 | -0.92367014615951 | 0.79 | 0.982 | 2.7345315 |
| **RB1** | 1.7879 | -1.00809903272932 | 0.487 | 0.883 | 2.9281954 |
| **NR6A1** | 1.9769 | -1.17193122199294 | 0.893 | 0.983 | 3.2375957 |
| **PDE3B** | 2.1417 | -1.15903987635359 | 0.188 | 0.728 | 3.5074885 |
| **ESRRA** | 2.4022 | 0.916013081643245 | 0.986 | 0.881 | 3.9341043 |
| **PAK2** | 4.7676 | 0.883763154745014 | 0.975 | 0.838 | 7.8079518 |
| **AMZ2** | 7.1905 | 0.949844983731332 | 0.934 | 0.677 | 1.1775972 |
| **PDE9A** | 9.2928 | -1.24844859075441 | 0.134 | 0.689 | 1.5218891 |
| **GALC** | 1.3157 | -1.06124183570789 | 0.457 | 0.869 | 2.1547792 |
| **DPH6** | 1.4226 | 0.993455703954121 | 0.951 | 0.7 | 2.3298887 |
| **SURF2** | 1.8417 | 0.863849229569288 | 0.993 | 0.916 | 3.0161659 |
| **VEGFB** | 2.9482 | -0.91871147864045 | 0.868 | 0.996 | 4.8282714 |
| **NXN** | 3.4198 | -0.80700432073683 | 0.978 | 1 | 5.6007476 |
| **TSKU** | 3.7691 | -1.19679088388917 | 0.946 | 0.996 | 6.1726826 |
| **SIGIRR** | 5.1267 | 0.984859217292837 | 0.934 | 0.652 | 8.3960959 |
| **UGCG** | 6.2690 | 0.990995027590636 | 0.997 | 0.979 | 1.0266769 |
| **RASSF3** | 9.3449 | 0.863115815150053 | 1 | 0.993 | 1.5304266 |
| **NIPAL1** | 9.8507 | 0.953050275236782 | 0.619 | 0.171 | 1.6132572 |
| **NUP50** | 1.2316 | 0.812363552616152 | 0.992 | 0.931 | 2.0170310 |
| **BOD1L1** | 1.6676 | 0.879268416283235 | 0.998 | 0.992 | 2.7311348 |
| **DCAKD** | 2.8177 | -0.88350583232505 | 0.856 | 0.985 | 4.6147075 |
| **DNAJB6** | 3.5922 | 0.951538225769053 | 0.986 | 0.949 | 5.8830532 |
| **OCIAD2** | 3.7205 | -1.1773962474709 | 0.174 | 0.729 | 6.0931048 |
| **APPL1** | 4.3836 | 0.844932558946068 | 0.986 | 0.91 | 7.1791443 |
| **ZNF483** | 5.1073 | 1.28023704285565 | 0.88 | 0.601 | 8.3643043 |
| **SLC9A3R2** | 6.2550 | 1.12169333080838 | 0.932 | 0.65 | 1.0243867 |
| **TADA3** | 6.7188 | -0.84094578053706 | 0.931 | 0.995 | 1.1003391 |
| **RASGRP2** | 8.0534 | 1.25168242279593 | 0.787 | 0.37 | 1.3189131 |
| **PCBD1** | 8.8489 | -1.06111704663671 | 0.802 | 0.966 | 1.4491999 |
| **FAM171A1** | 1.2432 | -1.0292122369767 | 0.453 | 0.875 | 2.0360938 |
| **SHF** | 1.5706 | -1.21920526364174 | 0.435 | 0.853 | 2.5723271 |

| | | | | | |
|---|---|---|---|---|---|
| **AK6** | 1.6166 | 0.877117427608373 | 0.978 | 0.852 | 2.6476032[illegible] |
| **HIPK1** | 1.7941 | -0.84710259381642[illegible] | 0.846 | 0.978 | 2.9382744[illegible] |
| **RASL10B** | 2.3097 | -1.2597464954122 | 0.183 | 0.717 | 3.7826155[illegible] |
| **AASS** | 3.0127 | 1.43936218186815 | 0.946 | 0.806 | 4.9340611[illegible] |
| **ARRDC2** | 4.0733 | 1.04914026471963 | 0.775 | 0.33 | 6.6709342[illegible] |
| **CHMP5** | 4.1521 | 0.835381584418074 | 0.98 | 0.831 | 6.7999836[illegible] |
| **BCL7C** | 4.9673 | -1.08861238839956 | 0.878 | 0.992 | 8.1350880[illegible] |
| **PLLP** | 5.9931 | 0.817714757907535 | 0.538 | 0.118 | 9.8150029[illegible] |
| **ZNF618** | 1.5455 | -0.94511527729400 | 0.758 | 0.964 | 2.5310985[illegible] |
| **SAR1B** | 2.0608 | 0.886585857293412 | 0.98 | 0.859 | 3.3749838[illegible] |
| **CLU** | 2.4792 | -1.14999904744782 | 0.912 | 0.995 | 4.0602335[illegible] |
| **TNNI3** | 5.0690 | 0.823594731828267 | 0.66 | 0.187 | 8.3016162[illegible] |
| **HCN2** | 5.2320 | 0.943384572780552 | 0.951 | 0.708 | 8.5685109[illegible] |
| **SKAP2** | 5.8597 | -1.21358362660045 | 0.173 | 0.715 | 9.5965140[illegible] |
| **FAM118B** | 6.4660 | -0.89322265299959 | 0.753 | 0.97 | 1.0589512[illegible] |
| **PDGFRB** | 7.0215 | -1.14224476160395 | 0.069 | 0.641 | 1.1499248[illegible] |
| **COL6A2** | 9.2596 | -1.70976708725848 | 0.8 | 0.955 | 1.5164592[illegible] |
| **SMPDL3A** | 9.2948 | 0.966722378633959 | 0.616 | 0.174 | 1.5222133[illegible] |
| **FAM104B** | 1.4765 | -0.98692891328049 | 0.638 | 0.941 | 2.4180972[illegible] |
| **EEF1AKMT[illegible]** | 1.8401 | 0.921549890935405 | 0.942 | 0.772 | 3.0136458[illegible] |
| **ACTR2** | 2.7812 | 0.866091046565541 | 0.978 | 0.766 | 4.5549148[illegible] |
| **IGSF8** | 3.0025 | -1.10327408221479 | 0.682 | 0.936 | 4.9173255[illegible] |
| **ST6GALNA[illegible]** | 3.3865 | -1.17452329585456 | 0.09 | 0.648 | 5.5462297[illegible] |
| **FES** | 4.1842 | 0.888224416262555 | 0.802 | 0.345 | 6.8525060[illegible] |
| **DOCK9** | 4.3613 | 0.894993761314764 | 0.97 | 0.74 | 7.1425743[illegible] |
| **ZNF675** | 4.8428 | 0.915296390990773 | 0.981 | 0.827 | 7.9312144[illegible] |
| **GSTM3** | 6.6527 | -1.46626988635003 | 0.161 | 0.694 | 1.0895222[illegible] |
| **STARD3NL** | 1.0588 | -0.82727831257743 | 0.912 | 0.993 | 1.7340980[illegible] |
| **NKAIN1** | 1.0876 | -1.30301677970491 | 0.201 | 0.72 | 1.7813109[illegible] |
| **EIF2AK2** | 1.7452 | -0.90127890705796 | 0.694 | 0.961 | 2.8581558[illegible] |
| **LOXL1** | 2.5932 | -1.33726419317975 | 0.447 | 0.842 | 4.2470053[illegible] |
| **ARFGEF1** | 3.2449 | 0.875238412928985 | 0.993 | 0.958 | 5.3142166[illegible] |
| **LIMK2** | 6.5850 | 0.980606145596748 | 0.997 | 0.955 | 1.0784361[illegible] |
| **GFPT1** | 6.9899 | 0.876876309507605 | 1 | 0.996 | 1.1447453[illegible] |

| | | | | | |
|---|---|---|---|---|---|
| **SLC66A2** | 7.8861 | 0.988406153517716 | 0.959 | 0.85 | 1.29151923 |
| **ZNF587B** | 9.1187 | 0.895821485024539 | 0.958 | 0.797 | 1.49337580 |
| **ZNF680** | 1.1212 | 0.981202823125884 | 0.948 | 0.736 | 1.83634444 |
| **WDR44** | 1.1747 | 0.985505928812443 | 0.971 | 0.891 | 1.92382847 |
| **SPARC** | 1.6098 | 0.805984878761011 | 1 | 1 | 2.63648353 |
| **ZEB2** | 2.9243 | -1.40818402478424 | 0.146 | 0.692 | 4.78923247 |
| **VCL** | 8.5658 | -0.91649242233354 | 1 | 1 | 1.40282542 |
| **PKP3** | 8.9457 | 1.01115908325214 | 0.958 | 0.647 | 1.46505158 |
| **REEP3** | 9.3880 | -0.92599797538867 | 0.711 | 0.963 | 1.53748418 |
| **UAP1L1** | 1.0602 | 1.10777931254465 | 0.832 | 0.449 | 1.73638140 |
| **MAT2B** | 1.3148 | 0.814102331582617 | 0.997 | 0.977 | 2.15329989 |
| **CHEK2** | 1.5444 | 0.994419369143103 | 0.99 | 0.941 | 2.52938077 |
| **STK10** | 1.6928 | 0.96804808810287 | 0.954 | 0.783 | 2.77243805 |
| **F2RL1** | 3.4146 | 1.0136080062263 | 0.993 | 0.956 | 5.59213359 |
| **NOL4L** | 3.5716 | -1.05884779809872 | 0.513 | 0.899 | 5.84930418 |
| **TMEM54** | 4.0000 | 0.897704564088892 | 0.773 | 0.303 | 6.55084908 |
| **ARHGAP18** | 4.2087 | 0.874914634396342 | 0.657 | 0.202 | 6.89273060 |
| **PLEC** | 4.8148 | 0.846235504430379 | 0.997 | 0.954 | 7.88529640 |
| **ATRN** | 6.2124 | -0.84353978307233 | 0.897 | 0.988 | 1.01742064 |
| **FBRSL1** | 6.2817 | 0.869308730303027 | 0.976 | 0.861 | 1.02875865 |
| **CORO1A** | 7.7436 | 0.98288243369611 | 0.658 | 0.218 | 1.26818462 |
| **ESCO1** | 8.8554 | 0.830994372851991 | 0.993 | 0.905 | 1.45026269 |
| **ADAMTS19** | 1.9549 | -1.5666878809986 | 0.027 | 0.579 | 3.20167462 |
| **ST14** | 1.9899 | 0.916303313464414 | 0.643 | 0.193 | 3.25896568 |
| **KDM4C** | 2.1192 | 1.13471233635547 | 0.946 | 0.784 | 3.47069629 |
| **FAM133B** | 2.9809 | 0.802971500570403 | 0.743 | 0.285 | 4.88185308 |
| **PTBP2** | 4.8656 | -0.96619184592759 | 0.873 | 0.987 | 7.96851179 |
| **TRIOBP** | 5.6909 | -0.88857764151082 | 0.838 | 0.985 | 9.31999863 |
| **DTNA** | 8.9709 | -1.93335603514957 | 0.103 | 0.645 | 1.46917179 |
| **RGS2** | 9.7490 | 1.26723579002808 | 0.854 | 0.491 | 1.59660373 |
| **TSPAN6** | 1.3894 | -0.80593061359233 | 0.975 | 0.999 | 2.27552100 |
| **PITPNM2** | 2.0390 | 1.07847957610387 | 0.92 | 0.626 | 3.33943368 |
| **UHRF2** | 2.4209 | -1.09751664310694 | 0.362 | 0.819 | 3.96477930 |
| **EDNRB** | 3.6750 | -1.68023102319382 | 0.291 | 0.766 | 6.01869775 |

| | | | | | |
|---|---|---|---|---|---|
| **NOTCH1** | 4.8356 | -0.99904889453874 | 0.814 | 0.979 | 7.9193022 |
| **GYG2** | 4.9590 | -0.96994810552904 | 0.765 | 0.983 | 8.1213651 |
| **CLDN7** | 5.5528 | 1.11002773155889 | 0.978 | 0.743 | 9.0938316 |
| **MPV17L2** | 6.4563 | 0.868496692743746 | 0.976 | 0.865 | 1.0573485 |
| **DUSP5** | 1.0479 | 1.18159522068796 | 0.949 | 0.769 | 1.7162994 |
| **PCNX1** | 1.7768 | -0.81581001526274 | 0.926 | 0.99 | 2.9098739 |
| **PRR36** | 1.8923 | -1.38285453070706 | 0.232 | 0.715 | 3.0991575 |
| **ROR1** | 4.1538 | -1.22732968563085 | 0.323 | 0.802 | 6.8027791 |
| **TRAF5** | 9.5084 | -1.14122050690777 | 0.232 | 0.733 | 1.5572041 |
| **UNC5D** | 1.0632 | -1.0928316325466 | 0.063 | 0.607 | 1.7413090 |
| **MDM2** | 1.0894 | 1.30648265019973 | 0.998 | 0.987 | 1.7841357 |
| **GATAD2B** | 1.1036 | -0.84116486944435 | 0.819 | 0.976 | 1.8074077 |
| **PRAP1** | 1.1554 | 0.815724353151326 | 0.421 | 0.067 | 1.8922127 |
| **COL6A1** | 1.1898 | -1.76573107889382 | 0.536 | 0.88 | 1.9486068 |
| **SACS** | 1.9753 | -0.93398115212527 | 0.892 | 0.987 | 3.2350872 |
| **KAT6B** | 2.0553 | 0.895210811272057 | 0.978 | 0.859 | 3.3660653 |
| **PINX1** | 2.4228 | 0.832134425606297 | 0.981 | 0.873 | 3.9679034 |
| **NIN** | 2.4668 | -0.99446223367933 | 0.635 | 0.941 | 4.0399657 |
| **WBP1** | 2.8537 | -0.87860463881330 | 0.902 | 0.985 | 4.6736233 |
| **ERCC1** | 3.7700 | -0.82679878755509 | 0.89 | 0.987 | 6.1742069 |
| **EXOSC4** | 4.1391 | 0.82807673623103 | 0.99 | 0.901 | 6.7787612 |
| **SCGB3A2** | 4.3927 | 1.17963932891057 | 0.99 | 0.964 | 7.1940318 |
| **VOPP1** | 5.3660 | -0.81906356638646 | 0.853 | 0.979 | 8.7879102 |
| **PSRC1** | 6.4943 | 1.16529804601785 | 0.953 | 0.707 | 1.0635720 |
| **SEPHS1** | 9.8750 | -0.96560871714402 | 0.995 | 1 | 1.6172445 |
| **AHDC1** | 1.2270 | -1.1323620112892 | 0.514 | 0.875 | 2.0095976 |
| **MIPOL1** | 1.3354 | -1.01509894209042 | 0.272 | 0.766 | 2.1871434 |
| **TBX22** | 1.7108 | -1.35834048681728 | 0.046 | 0.595 | 2.8018200 |
| **MMP25** | 1.8655 | -1.26864113512514 | 0.269 | 0.744 | 3.0552508 |
| **SYNGAP1** | 1.8854 | -1.22082079200749 | 0.34 | 0.78 | 3.0878114 |
| **PINK1** | 1.9340 | 0.896201211136701 | 0.988 | 0.94 | 3.1674393 |
| **PCED1A** | 3.1113 | -1.1899842115111 | 0.24 | 0.733 | 5.0954341 |
| **HERPUD2** | 3.7736 | -0.91723080649900 | 0.636 | 0.929 | 6.1800705 |
| **HUNK** | 4.8775 | -1.0640034043751 | 0.452 | 0.862 | 7.9880357 |

| | | | | | |
|---|---|---|---|---|---|
| **UTRN** | 5.9029 | -1.10852719394262 | 0.677 | 0.933 | 9.6673246 |
| **PDE7A** | 9.4415 | -0.81515389308331 | 0.929 | 0.994 | 1.5462397 |
| **ADGRL2** | 1.2205 | -0.86631253258911 | 0.964 | 0.997 | 1.9988231 |
| **NIPSNAP2** | 1.3192 | -0.80597972088750 | 0.924 | 0.99 | 2.1604721 |
| **PARVB** | 1.3417 | 1.19423965168538 | 0.819 | 0.488 | 2.1973134 |
| **HRK** | 1.3435 | 0.989170927685485 | 0.565 | 0.156 | 2.2002838 |
| **ZNF296** | 1.4848 | 1.34122084780192 | 0.854 | 0.518 | 2.4318083 |
| **LRFN5** | 1.6967 | 0.938964237585682 | 0.609 | 0.177 | 2.7788265 |
| **MYC** | 2.0656 | -1.15160738571006 | 0.926 | 0.991 | 3.3829814 |
| **ADK** | 2.4048 | -0.85073603479390 | 0.843 | 0.975 | 3.9383426 |
| **CNN1** | 2.4289 | -1.83460053918224 | 0.562 | 0.904 | 3.9779444 |
| **COP1** | 2.8766 | 0.806934106940239 | 0.939 | 0.668 | 4.7110350 |
| **C1orf198** | 3.2570 | -0.85948651763250 | 0.726 | 0.963 | 5.3341290 |
| **SYT6** | 4.2041 | -1.15914507680209 | 0.176 | 0.695 | 6.8851348 |
| **CTSZ** | 4.7025 | -1.0667968596417 | 0.741 | 0.934 | 7.7013961 |
| **SEPTIN7** | 4.8273 | 0.846558374415586 | 0.88 | 0.544 | 7.9056727 |
| **ZIC5** | 5.3794 | -1.67556178745693 | 0.208 | 0.706 | 8.8098839 |
| **FCHO2** | 5.5543 | 0.866873906125367 | 0.768 | 0.343 | 9.0963599 |
| **TBC1D4** | 5.5805 | -1.02918966754958 | 0.45 | 0.856 | 9.1392206 |
| **ARSB** | 8.4844 | -0.87200159092793 | 0.753 | 0.957 | 1.3894964 |
| **FAM217B** | 9.0150 | -0.88427306147863 | 0.712 | 0.944 | 1.4763930 |
| **BRWD3** | 9.7074 | 0.984247843028913 | 0.992 | 0.978 | 1.5897971 |
| **ASS1** | 1.0617 | 1.36867330279924 | 0.981 | 0.948 | 1.7388788 |
| **CUEDC2** | 1.9376 | -0.89435513184375 | 0.687 | 0.947 | 3.1733535 |
| **SFRP2** | 2.0880 | -1.66755005044655 | 0.897 | 0.97 | 3.4196782 |
| **BECN1** | 2.2371 | 0.807284341708858 | 0.973 | 0.855 | 3.6637457 |
| **EEA1** | 2.3752 | 0.867502389608549 | 0.959 | 0.768 | 3.8898932 |
| **ZNF589** | 2.4165 | 0.921623250148305 | 0.981 | 0.904 | 3.9575465 |
| **MAGI1** | 5.9412 | -1.20129675867657 | 0.584 | 0.888 | 9.7299080 |
| **PDLIM2** | 6.1728 | -0.89524329557071 | 0.898 | 0.991 | 1.0109219 |
| **PRDM1** | 6.2213 | 0.882183727097079 | 0.655 | 0.199 | 1.0188652 |
| **CDCP1** | 6.3465 | 1.00697827407046 | 0.794 | 0.368 | 1.0393809 |
| **LYPD1** | 8.3957 | -1.95816518313617 | 0.254 | 0.724 | 1.3749743 |
| **FAM210B** | 1.1461 | 0.913010119843598 | 0.863 | 0.49 | 1.8770140 |

| | | | | | |
|---|---|---|---|---|---|
| **BTG2** | 1.3653 | -1.38721480914307 | 0.31 | 0.802 | 2.2360742 |
| **GRTP1** | 1.5137 | 0.905201167586738 | 0.746 | 0.297 | 2.4790414 |
| **YPEL2** | 1.9581 | 1.42221959034651 | 0.665 | 0.249 | 3.2068039 |
| **ZNF614** | 2.2723 | 0.938851539399238 | 0.939 | 0.697 | 3.7214017 |
| **PGM2** | 3.0658 | 0.985201395568199 | 0.971 | 0.871 | 5.0210175 |
| **SLC25A38** | 3.1854 | 0.824522380163873 | 0.981 | 0.913 | 5.2167690 |
| **GRB7** | 3.3194 | 0.978942740369019 | 0.841 | 0.442 | 5.4362523 |
| **DDB2** | 3.5988 | -1.06478535119051 | 0.616 | 0.914 | 5.8938886 |
| **TTC7B** | 3.8401 | -0.88148474923242 | 0.719 | 0.964 | 6.2890163 |
| **RASSF9** | 4.3208 | -1.48418193539696 | 0.146 | 0.659 | 7.0762208 |
| **RNF14** | 5.3935 | 0.914062643415634 | 0.932 | 0.721 | 8.8330284 |
| **FOXD3** | 5.8527 | 1.29844822645977 | 0.692 | 0.271 | 9.5850445 |
| **BRSK1** | 6.9147 | -1.02290843728421 | 0.146 | 0.664 | 1.1324341 |
| **RELL1** | 7.1512 | 0.915977444707706 | 0.954 | 0.775 | 1.1711676 |
| **CNEP1R1** | 7.2767 | 0.889369584330223 | 0.907 | 0.622 | 1.1917201 |
| **COL2A1** | 1.3695 | -1.88904175985009 | 0.046 | 0.575 | 2.2429636 |
| **CDH4** | 1.5908 | -1.10000549179221 | 0.046 | 0.578 | 2.6054112 |
| **MTRES1** | 1.9595 | 0.842997423754108 | 0.702 | 0.27 | 3.2091226 |
| **ATP1A2** | 6.2485 | -1.12846497585469 | 0.02 | 0.549 | 1.0233313 |
| **MMAB** | 6.8759 | -0.84157315893440 | 0.917 | 0.994 | 1.1260779 |
| **NDUFAF7** | 7.3234 | 0.883846825509755 | 0.966 | 0.817 | 1.1993582 |
| **NCAPD2** | 7.5853 | -0.93513193531121 | 0.963 | 0.998 | 1.2422499 |
| **CBR3** | 1.1983 | 1.12063261557628 | 0.817 | 0.408 | 1.9625533 |
| **SDC1** | 1.2653 | -1.11493847562493 | 0.257 | 0.754 | 2.0722675 |
| **HDHD2** | 1.4510 | -0.82373110858330 | 0.812 | 0.969 | 2.3764180 |
| **PLTP** | 1.6532 | -0.83338000934208 | 0.92 | 0.995 | 2.7075884 |
| **HSPA1B** | 2.8044 | 1.43065690309343 | 0.807 | 0.459 | 4.5928072 |
| **ZNF318** | 2.9234 | 0.868886244511978 | 0.934 | 0.754 | 4.7877546 |
| **PCNA** | 5.3545 | -0.90234111559571 | 0.997 | 0.998 | 8.7691879 |
| **MME** | 1.0413 | -1.21249145248907 | 0.179 | 0.674 | 1.7054061 |
| **FZD10** | 1.6681 | -1.65633115055768 | 0.547 | 0.876 | 2.7318849 |
| **MBOAT1** | 2.2062 | 0.952016997281315 | 0.885 | 0.517 | 3.6131748 |
| **PPTC7** | 2.3112 | 0.871448215167054 | 0.951 | 0.762 | 3.7851922 |
| **KCTD20** | 2.4675 | -0.86100369590387 | 0.846 | 0.984 | 4.0411747 |

| | | | | | |
|---|---|---|---|---|---|
| **VEZF1** | 4.0737 | -0.82578936114176 | 0.878 | 0.985 | 6.6715680 |
| **PDLIM7** | 6.4767 | -0.87958269392168 | 0.99 | 0.999 | 1.0606927 |
| **IFT52** | 6.9417 | -0.81027047117733 | 0.81 | 0.969 | 1.1368482 |
| **TRIM44** | 7.5500 | -0.82033425323654 | 0.817 | 0.98 | 1.2364650 |
| **TMLHE** | 8.3153 | -0.98421907029130 | 0.323 | 0.781 | 1.3618102 |
| **DMD** | 9.1389 | -1.21668291872603 | 0.496 | 0.865 | 1.4966877 |
| **EXTL2** | 9.8624 | -0.83849433875577 | 0.766 | 0.954 | 1.6151694 |
| **BMP4** | 1.0775 | -1.43111973899949 | 0.477 | 0.86 | 1.7646317 |
| **PPP2R3A** | 1.2599 | -0.89544866199604 | 0.557 | 0.896 | 2.0634802 |
| **APIP** | 1.4436 | 0.850066834327283 | 0.978 | 0.85 | 2.3643374 |
| **ECSCR** | 1.6616 | -1.03404189248324 | 0.042 | 0.563 | 2.7212873 |
| **TMEM184C** | 1.7846 | 0.856235755997493 | 0.844 | 0.508 | 2.9227284 |
| **BST2** | 3.0650 | -1.75296390554113 | 0.47 | 0.832 | 5.0197113 |
| **NUAK1** | 3.4780 | -1.01566594286063 | 0.64 | 0.955 | 5.6959991 |
| **FGF4** | 3.8329 | 1.19512021754506 | 0.919 | 0.639 | 6.2772662 |
| **ADORA2B** | 6.0214 | 0.855967419280838 | 0.663 | 0.245 | 9.8613031 |
| **REEP2** | 6.7549 | -1.09081561013485 | 0.188 | 0.687 | 1.1062649 |
| **SH3GL3** | 9.1374 | -1.11788135930833 | 0.201 | 0.691 | 1.4964340 |
| **COMT** | 1.5225 | -0.94011325897161 | 0.57 | 0.899 | 2.4935307 |
| **MYORG** | 1.5252 | -1.10469552863566 | 0.298 | 0.752 | 2.4978360 |
| **MSRB2** | 1.7801 | -0.92538730078599 | 0.602 | 0.911 | 2.9153103 |
| **MEGF8** | 2.1056 | -0.93772218363336 | 0.538 | 0.878 | 3.4484549 |
| **SCML2** | 2.2458 | -0.85909119188783 | 0.64 | 0.921 | 3.6780977 |
| **SDF2L1** | 2.3614 | 0.936863687623496 | 0.998 | 0.98 | 3.8672678 |
| **STK17B** | 2.5004 | 0.992260502341975 | 0.812 | 0.444 | 4.0949933 |
| **KLHL23** | 3.6307 | -0.92765591713370 | 0.658 | 0.916 | 5.9461476 |
| **ZSCAN2** | 4.0266 | 1.01763693513314 | 0.971 | 0.892 | 6.5944817 |
| **ATP12A** | 4.1449 | 1.28208608091929 | 0.817 | 0.415 | 6.7881488 |
| **RMND5B** | 4.1998 | 0.972489727314748 | 0.948 | 0.804 | 6.8780493 |
| **NAP1L2** | 5.7066 | -0.96748552215685 | 0.142 | 0.654 | 9.3457227 |
| **CRELD2** | 5.7555 | 0.915324142383072 | 0.992 | 0.934 | 9.4258906 |
| **FIGNL2** | 7.8605 | -1.08616888274142 | 0.164 | 0.672 | 1.2873230 |
| **ZNF518A** | 8.7505 | 0.801926801767992 | 0.992 | 0.945 | 1.4330844 |
| **PLAGL2** | 1.0076 | -0.84602949588976 | 0.699 | 0.93 | 1.6502582 |

| | | | | | |
|---|---|---|---|---|---|
| **LRRN2** | 1.1151 | -1.16614544855577 | 0.154 | 0.652 | 1.8262591 |
| **RIDA** | 1.1811 | 0.84069729830708 | 0.844 | 0.465 | 1.9344470 |
| **FARP1** | 1.6874 | -0.86647442789608 | 0.77 | 0.964 | 2.7635240 |
| **ENPP1** | 2.8948 | -1.0481273668138 | 0.687 | 0.948 | 4.7408223 |
| **ZNF354A** | 5.3739 | 0.895008842544511 | 0.81 | 0.429 | 8.8009488 |
| **HOXA1** | 1.4720 | -1.33223624658443 | 0.545 | 0.876 | 2.4107286 |
| **SYS1** | 1.6014 | -0.83777271102494 | 0.738 | 0.947 | 2.6227084 |
| **WDR35** | 2.1322 | -0.98086387220448 | 0.293 | 0.748 | 3.4919537 |
| **ZNF480** | 2.6071 | 0.81991372577452 | 0.917 | 0.67 | 4.2697153 |
| **TFPI2** | 4.0648 | 1.13927544264765 | 0.865 | 0.515 | 6.6570630 |
| **STK40** | 5.9436 | -0.85488866190691 | 0.785 | 0.959 | 9.7338520 |
| **DSC2** | 1.0601 | -0.85140782604301 | 0.775 | 0.967 | 1.7362032 |
| **KIAA1324L** | 2.0058 | -0.88433828392896 | 0.606 | 0.916 | 3.2849075 |
| **ERGIC1** | 2.5058 | -0.81356700585576 | 0.788 | 0.968 | 4.1037566 |
| **ACKR3** | 2.6420 | 1.27718129472516 | 0.69 | 0.276 | 4.3269359 |
| **CCNJ** | 3.4790 | 0.813344124528697 | 0.751 | 0.335 | 5.6975802 |
| **AGAP3** | 3.6582 | -0.93938793753505 | 0.299 | 0.758 | 5.9911306 |
| **GLRX2** | 3.8708 | -0.97175925435167 | 0.284 | 0.748 | 6.3392245 |
| **SOCS1** | 5.7670 | -0.94485564983059 | 0.953 | 0.995 | 9.4446908 |
| **LRIG1** | 6.2900 | 0.844596654745974 | 0.986 | 0.97 | 1.0301274 |
| **FKBP11** | 8.8975 | 0.953448314980206 | 0.819 | 0.462 | 1.4571582 |
| **LPAR6** | 1.0192 | 0.852632174831127 | 0.638 | 0.219 | 1.6692109 |
| **P3H2** | 1.8357 | -2.04004368525614 | 0.074 | 0.593 | 3.0063521 |
| **GBX2** | 2.2522 | -1.19404009845086 | 0.602 | 0.894 | 3.6884987 |
| **SMYD2** | 2.5713 | -0.83004475108875 | 0.944 | 0.99 | 4.2110595 |
| **NSUN5** | 3.0566 | 0.805961535735388 | 0.909 | 0.658 | 5.0058559 |
| **TSC22D3** | 4.0209 | -1.0793961386335 | 0.503 | 0.902 | 6.5850373 |
| **SCG3** | 4.0946 | -1.04293240347157 | 0.303 | 0.744 | 6.7057821 |
| **FNDC10** | 4.5458 | 0.996619388102561 | 0.834 | 0.485 | 7.4447520 |
| **FLT1** | 4.7784 | -1.47420256730157 | 0.184 | 0.673 | 7.8256634 |
| **KLRG2** | 5.5728 | 0.851580243078223 | 0.98 | 0.925 | 9.1266342 |
| **LGR4** | 7.6927 | 0.840486579660304 | 0.997 | 0.984 | 1.2598383 |
| **EPDR1** | 1.2953 | -0.92656281217636 | 0.03 | 0.532 | 2.1213721 |
| **PTPRG** | 1.9928 | -1.01935185810221 | 0.88 | 0.993 | 3.2637438 |

| | | | | | |
|---|---|---|---|---|---|
| **ZBTB12** | 2.0385 | -0.96915192294900 | 0.122 | 0.623 | 3.3384911 |
| **PCIF1** | 2.1841 | -0.80472789916889 | 0.853 | 0.976 | 3.5770349 |
| **ADGRL3** | 2.2250 | 0.838425930981899 | 0.404 | 0.076 | 3.6440037 |
| **PM20D2** | 2.4752 | -1.01582425461772 | 0.171 | 0.658 | 4.0537824 |
| **CRISPLD1** | 2.4833 | -1.01947690490906 | 0.653 | 0.937 | 4.0669647 |
| **1,00 PGK** | 2.5768 | 0.975869958159411 | 0.804 | 0.456 | 4.2200854 |
| **CACNA1B** | 2.6208 | 0.806034214600567 | 0.64 | 0.224 | 4.2921726 |
| **PLBD2** | 4.0710 | -0.80638341642905 | 0.844 | 0.976 | 6.6671371 |
| **TRIB3** | 4.8352 | 1.73668533706378 | 0.892 | 0.697 | 7.9186349 |
| **CALD1** | 1.2986 | -1.6822954621125 | 0.998 | 1 | 2.1268352 |
| **RBCK1** | 1.3561 | -0.86122664855605 | 0.648 | 0.927 | 2.2209410 |
| **ACOX1** | 1.4672 | 0.801604227255942 | 0.975 | 0.858 | 2.4029677 |
| **GRPEL2** | 1.5244 | 0.825951903045726 | 0.932 | 0.72 | 2.4966031 |
| **GEMIN2** | 4.3708 | 0.808126071130937 | 0.844 | 0.494 | 7.1581504 |
| **ANKRD18A** | 4.9653 | -1.11563723306691 | 0.215 | 0.683 | 8.1316932 |
| **SMG6** | 5.8342 | -0.83098409722882 | 0.729 | 0.94 | 9.5547509 |
| **LRP4** | 6.3209 | 0.876768882900882 | 0.992 | 0.945 | 1.0351745 |
| **NCS1** | 9.4287 | -0.82627403828446 | 0.797 | 0.959 | 1.5441545 |
| **PODXL** | 1.5863 | -2.20112699324791 | 0.942 | 0.976 | 2.5979407 |
| **GULP1** | 2.6342 | -0.98464184664561 | 0.44 | 0.812 | 4.3141051 |
| **TIGAR** | 3.2421 | 0.849145075851161 | 1 | 0.979 | 5.3096523 |
| **STC1** | 4.0873 | -1.37347738983664 | 0.024 | 0.518 | 6.6938572 |
| **BGN** | 4.2037 | -1.80845416822257 | 0.296 | 0.718 | 6.8844918 |
| **FOXO1** | 4.3527 | -1.15415776035597 | 0.701 | 0.93 | 7.1285470 |
| **CENPV** | 7.6151 | -0.99130530746468 | 0.905 | 0.98 | 1.2471410 |
| **C18orf21** | 9.2067 | 0.818970062857389 | 0.875 | 0.578 | 1.5077842 |
| **TMEM220** | 1.0411 | 0.89120395220573 | 0.839 | 0.497 | 1.7050415 |
| **EDA** | 1.6348 | -0.97067128894159 | 0.431 | 0.843 | 2.6774646 |
| **RAMAC** | 1.6840 | 0.84937087732402 | 0.851 | 0.53 | 2.7580203 |
| **WDR54** | 1.6912 | -0.88185841090152 | 0.839 | 0.973 | 2.7696882 |
| **SNN** | 1.9512 | -0.88087826589501 | 0.567 | 0.894 | 3.1955316 |
| **DCTN5** | 2.0153 | 0.805232285532306 | 0.951 | 0.839 | 3.3005414 |
| **ELK3** | 4.8099 | 0.848718684230271 | 0.714 | 0.3 | 7.8772409 |
| **CSF2RA** | 5.3509 | 1.03908637352917 | 0.521 | 0.144 | 8.7632703 |

| | | | | | |
|---|---|---|---|---|---|
| **PIP5K1C** | 6.0647 | -0.80451425756877 | 0.819 | 0.97 | 9.9321855 |
| **LAMA5** | 7.4902 | -0.91400024934567 | 0.986 | 0.996 | 1.2266714 |
| **DDX10** | 7.7641 | 0.856649990136163 | 0.812 | 0.458 | 1.2715366 |
| **SGK3** | 1.7496 | 0.91807122632737 | 0.91 | 0.659 | 2.8654291 |
| **MGAT5B** | 1.7896 | -0.96547378090682 | 0.164 | 0.635 | 2.9308457 |
| **AARS2** | 1.8840 | 0.870814550961996 | 0.971 | 0.88 | 3.0854999 |
| **CNMD** | 2.1312 | 1.00892776614052 | 0.738 | 0.344 | 3.4903009 |
| **KRT10** | 2.7129 | -0.81157851727467 | 0.653 | 0.913 | 4.4430411 |
| **SLC39A14** | 4.5575 | 0.886178047054567 | 0.926 | 0.741 | 7.4638647 |
| **EPHX1** | 4.9683 | -1.23534310300081 | 0.939 | 0.991 | 8.1367177 |
| **CDC42EP1** | 5.2229 | -1.00128282628305 | 0.151 | 0.654 | 8.5536199 |
| **CAAP1** | 1.2644 | 0.828674692685452 | 0.907 | 0.635 | 2.0708665 |
| **SERTAD4** | 1.3571 | -1.13506408517309 | 0.151 | 0.622 | 2.2226775 |
| **CTSB** | 1.3957 | 0.80309095070215 | 1 | 0.982 | 2.2858216 |
| **PCDH10** | 1.8041 | -2.32776749293264 | 0.09 | 0.563 | 2.9546008 |
| **CENPA** | 6.0797 | 0.956693896825006 | 0.986 | 0.874 | 9.9567563 |
| **COL1A2** | 6.1882 | -1.21794268239949 | 0.827 | 0.989 | 1.0134575 |
| **RBPJ** | 8.8506 | 0.836301601412519 | 0.99 | 0.946 | 1.4494635 |
| **FBXO17** | 1.9682 | -0.89376359696452 | 0.477 | 0.841 | 3.2234772 |
| **CBX2** | 2.3921 | -0.83974879215989 | 0.816 | 0.967 | 3.9175769 |
| **RIMKLB** | 3.1708 | 0.892988437593404 | 0.998 | 0.991 | 5.1929142 |
| **SMPDL3B** | 3.2964 | -1.01254020835567 | 0.102 | 0.581 | 5.3985699 |
| **FRMD4A** | 3.6357 | -0.96405309141728 | 0.215 | 0.696 | 5.9542909 |
| **LAMB2** | 3.8206 | -1.02880037893143 | 0.675 | 0.904 | 6.2571192 |
| **CNTFR** | 6.6160 | -1.99413996125133 | 0.332 | 0.729 | 1.0835041 |
| **CABLES2** | 8.0410 | -0.91029689517167 | 0.225 | 0.687 | 1.3168795 |
| **CNIH2** | 8.2236 | -0.87295567945927 | 0.636 | 0.913 | 1.3467943 |
| **EFEMP2** | 1.5970 | -0.94086184184726 | 0.245 | 0.695 | 2.6154273 |
| **CCDC169** | 2.1048 | 1.10906580517491 | 0.739 | 0.399 | 3.4470635 |
| **STARD8** | 3.5198 | -1.05299791845022 | 0.409 | 0.829 | 5.7643863 |
| **P3H4** | 4.6040 | -0.88857814044989 | 0.44 | 0.824 | 7.5399758 |
| **ANGPTL2** | 5.9641 | -0.94308727124306 | 0.073 | 0.55 | 9.7674731 |
| **PTPN13** | 7.3088 | -0.98446156758017 | 0.404 | 0.797 | 1.1969651 |
| **PAPPA** | 8.7773 | -1.1554072320966 | 0.046 | 0.524 | 1.4374584 |

| | | | | | |
|---|---|---|---|---|---|
| **DBNDD1** | 9.9367 | -1.13705664099318 | 0.626 | 0.886 | 1.62733498 |
| **GALNT17** | 1.1175 | -1.16534439313524 | 0.404 | 0.784 | 1.83021501 |
| **TMC6** | 1.8462 | -1.03464581346073 | 0.692 | 0.91 | 3.02358825 |
| **ZNF778** | 1.9571 | 0.810073623438459 | 0.875 | 0.581 | 3.20527785 |
| **IFI30** | 2.8593 | 0.871561783827696 | 0.939 | 0.735 | 4.68268635 |
| **TCF15** | 3.0582 | -0.92281926803283 | 0.13 | 0.597 | 5.00842529 |
| **ZNF577** | 3.1860 | 0.894644948627096 | 0.714 | 0.327 | 5.21783039 |
| **PNMA2** | 3.3076 | 1.05252329342536 | 0.653 | 0.27 | 5.41686107 |
| **USP3** | 3.7537 | -0.904070413026599 | 0.074 | 0.558 | 6.14754774 |
| **ZSWIM9** | 3.9760 | -0.889530529015727 | 0.262 | 0.71 | 6.51163011 |
| **ST3GAL3** | 8.5099 | -0.909619343330866 | 0.108 | 0.578 | 1.39367507 |
| **ARFGAP3** | 9.4393 | 0.837987065151482 | 0.932 | 0.668 | 1.54588742 |
| **ACP2** | 1.9173 | -0.835884472351854 | 0.653 | 0.915 | 3.14008064 |
| **GPT2** | 2.2069 | 0.834590558727925 | 1 | 0.981 | 3.61430322 |
| **LGALSL** | 2.6525 | -0.857359326530069 | 0.602 | 0.888 | 4.34402827 |
| **CFL2** | 3.4995 | -0.808146945694715 | 0.768 | 0.95 | 5.73117270 |
| **AC007906.2** | 5.2773 | -1.02354512042827 | 0.115 | 0.584 | 8.64274684 |
| **CREBL2** | 8.6555 | 0.874967318504821 | 0.963 | 0.877 | 1.41752625 |
| **OSTF1** | 9.2728 | 0.80031651428852 | 0.973 | 0.89 | 1.51861110 |
| **NRARP** | 1.5343 | -1.08567163600787 | 0.443 | 0.8 | 2.51277812 |
| **PKIA** | 1.8151 | -0.83289582421726 | 0.064 | 0.533 | 2.97273595 |
| **PJA1** | 1.8639 | -0.804986871676055 | 0.739 | 0.982 | 3.05263473 |
| **FRMD6** | 3.0410 | 1.04942569027241 | 0.931 | 0.728 | 4.98025854 |
| **CAV1** | 1.0818 | -1.4225804690545 | 0.445 | 0.835 | 1.77175098 |
| **ZNF600** | 1.1744 | 0.990069574934715 | 0.961 | 0.779 | 1.92346829 |
| **CTSK** | 2.0099 | -1.965873069172 | 0.235 | 0.647 | 3.29172530 |
| **S100A6** | 2.1192 | 0.927139570733201 | 0.455 | 0.11 | 3.47075525 |
| **CERS4** | 3.9953 | -0.915926352515313 | 0.983 | 0.999 | 6.54313344 |
| **ZRANB3** | 6.1889 | -0.87403008329860 | 0.338 | 0.755 | 1.01357248 |
| **SRSF8** | 7.7435 | 1.04599598567882 | 0.922 | 0.734 | 1.26815548 |
| **RHOB** | 1.0278 | -1.02541867020051 | 0.934 | 0.997 | 1.68333848 |
| **SHISAL2B** | 1.0720 | -1.00141071127137 | 0.157 | 0.614 | 1.75562447 |
| **BLOC1S2** | 1.5092 | 0.815965493812055 | 0.893 | 0.631 | 2.47169432 |
| **SIX5** | 1.6885 | -0.833879937705052 | 0.596 | 0.89 | 2.76540850 |

| | | | | | |
|---|---|---|---|---|---|
| **AGPAT4** | 2.9764 | -0.98175594989822 | 0.526 | 0.852 | 4.87449012 |
| **NAV2** | 5.2870 | -1.09851299486787 | 0.822 | 0.957 | 8.65852971 |
| **PTPN21** | 5.6416 | -0.90392946205481 | 0.294 | 0.716 | 9.23934542 |
| **CCNE1** | 7.0139 | 0.90614275657036 | 0.944 | 0.746 | 1.14867277 |
| **ZNF697** | 1.2431 | 0.861650342558223 | 0.946 | 0.831 | 2.03586819 |
| **GPNMB** | 1.4205 | 0.905772201269101 | 0.653 | 0.252 | 2.32651364 |
| **SPSB2** | 1.5460 | 0.877582997004997 | 0.897 | 0.649 | 2.53196168 |
| **GPC3** | 3.3685 | -1.10620506577677 | 0.44 | 0.825 | 5.51673692 |
| **ADM** | 6.9329 | -1.50230007427787 | 0.223 | 0.652 | 1.13541159 |
| **MAPRE2** | 7.8817 | -0.84973556850170 | 0.709 | 0.935 | 1.29078868 |
| **B3GALT6** | 8.6571 | -0.83228045751998 | 0.547 | 0.856 | 1.41778512 |
| **DDHD2** | 8.9215 | -0.86423266353341 | 0.467 | 0.806 | 1.46108392 |
| **PROM1** | 1.7763 | 1.1329236722065 | 0.934 | 0.752 | 2.90917368 |
| **TXNDC16** | 2.2242 | -0.85838088451142 | 0.269 | 0.693 | 3.64264009 |
| **LHPP** | 2.4386 | -0.85808233886329 | 0.501 | 0.839 | 3.99370062 |
| **SOCS3** | 2.8815 | -0.88040078484417 | 0.734 | 0.954 | 4.71909279 |
| **HNMT** | 2.9685 | -0.88213678399050 | 0.276 | 0.7 | 4.86155070 |
| **JAKMIP2** | 3.3988 | -1.00293060968783 | 0.039 | 0.492 | 5.56632680 |
| **ST6GALNAC** | 3.9258 | -0.86539278186680 | 0.433 | 0.806 | 6.42936729 |
| **SLC35F1** | 3.9419 | -0.88820989737060 | 0.102 | 0.559 | 6.45577790 |
| **ARHGAP11A** | 4.3946 | 0.986344790802028 | 0.975 | 0.814 | 7.19716413 |
| **AEBP1** | 5.8922 | -0.88992579643274 | 0.616 | 0.884 | 9.64969393 |
| **FAM111B** | 6.0315 | -1.18896872963565 | 0.481 | 0.821 | 9.87794810 |
| **ZKSCAN1** | 6.5522 | -0.88110454234107 | 0.472 | 0.811 | 1.07305909 |
| **RASL11B** | 6.8237 | -1.15239210898624 | 0.333 | 0.744 | 1.11752209 |
| **SYNDIG1** | 7.1409 | -0.92624555129543 | 0.041 | 0.492 | 1.16947210 |
| **PRICKLE3** | 7.4597 | -0.86856602059845 | 0.264 | 0.696 | 1.22167732 |
| **IGDCC4** | 7.8591 | -0.81195516629199 | 0.052 | 0.509 | 1.28708744 |
| **LARP1B** | 8.3588 | 0.820330190024986 | 0.904 | 0.687 | 1.36892212 |
| **RAI14** | 2.1996 | -0.83577190492829 | 0.751 | 0.944 | 3.60242042 |
| **SLC27A5** | 2.8313 | -0.86880109261272 | 0.377 | 0.759 | 4.63684395 |
| **NEO1** | 2.8359 | -0.83986471785137 | 0.553 | 0.861 | 4.64436979 |
| **DPYSL3** | 4.9169 | -1.14683333133838 | 0.832 | 0.99 | 8.05245851 |
| **BEX1** | 8.1949 | -1.06399822628903 | 0.047 | 0.496 | 1.34209419 |

| | | | | | |
|---|---|---|---|---|---|
| **P2RY1** | 1.8361 | -1.20850587442748 | 0.144 | 0.586 | 3.0070578 |
| **KATNAL1** | 2.5762 | -0.81014275949301 | 0.516 | 0.835 | 4.2192058 |
| **JMY** | 3.0220 | 0.84802451180928 | 0.856 | 0.586 | 4.9492343 |
| **APPL2** | 3.6620 | -0.86235137936562 | 0.469 | 0.825 | 5.9973184 |
| **IFI27L2** | 3.8627 | -0.93117561233447 | 0.213 | 0.644 | 6.3259963 |
| **AFF2** | 5.5093 | 0.986349530576885 | 0.602 | 0.235 | 9.0225955 |
| **PRTFDC1** | 5.5907 | -0.90839040960014 | 0.146 | 0.593 | 9.1559018 |
| **CNKSR3** | 6.1566 | -1.13575349164372 | 0.11 | 0.559 | 1.0082753 |
| **BAZ1A** | 6.7004 | -0.90010414951133 | 0.367 | 0.756 | 1.0973358 |
| **FHDC1** | 8.7165 | 0.817439511356902 | 0.618 | 0.241 | 1.4275124 |
| **PLK2** | 1.2494 | 1.36409115512895 | 0.609 | 0.237 | 2.0461926 |
| **C8orf58** | 1.3081 | -0.82341149677663 | 0.479 | 0.825 | 2.1423768 |
| **ELF4** | 1.3915 | -0.97289355616068 | 0.171 | 0.615 | 2.2789653 |
| **KCNMB1** | 2.0926 | -1.64068013508231 | 0.095 | 0.532 | 3.4271078 |
| **TMED8** | 3.0038 | -0.80811435865651 | 0.738 | 0.934 | 4.9194851 |
| **NECAB3** | 3.0046 | -0.81135146670476 | 0.655 | 0.898 | 4.9206413 |
| **NFE2L3** | 3.7593 | 0.964373387611063 | 0.88 | 0.597 | 6.1566060 |
| **ARHGEF10L** | 4.4361 | -0.85033872280269 | 0.34 | 0.744 | 7.2650700 |
| **MMP24OS** | 5.5727 | -0.80594529318256 | 0.646 | 0.912 | 9.1264597 |
| **GDF15** | 6.1218 | 1.72515249380394 | 0.442 | 0.117 | 1.0025750 |
| **IGFBP7** | 2.1376 | -1.48030949069232 | 0.249 | 0.67 | 3.5008293 |
| **EPHB3** | 4.0175 | -1.26368701731107 | 0.255 | 0.683 | 6.5794891 |
| **PCNX4** | 4.8848 | -0.84111160397624 | 0.497 | 0.841 | 7.9999874 |
| **C20orf96** | 5.4173 | -0.82122169151479 | 0.129 | 0.566 | 8.8719655 |
| **FGFBP3** | 6.1100 | -0.97739603083484 | 0.036 | 0.472 | 1.0006447 |
| **LRRC61** | 6.5680 | -0.83753121096554 | 0.271 | 0.677 | 1.0756463 |
| **CD24** | 8.9407 | 1.44147603734154 | 0.978 | 0.983 | 1.4642253 |
| **OAF** | 2.7996 | -0.82639865025093 | 0.071 | 0.514 | 4.5849990 |
| **UNK** | 3.7076 | -0.80645713387824 | 0.601 | 0.883 | 6.0720664 |
| **ME1** | 3.8572 | -0.84680430584977 | 0.447 | 0.8 | 6.3169740 |
| **MEST** | 4.1982 | 1.4665863097725 | 0.888 | 0.813 | 6.8754359 |
| **TMEM92** | 4.2266 | -0.86809124621968 | 0.32 | 0.723 | 6.9219508 |
| **KCTD14** | 4.6398 | -0.84337028196435 | 0.291 | 0.698 | 7.5986037 |
| **NKD1** | 8.4360 | -1.36250243411975 | 0.272 | 0.664 | 1.3815729 |

| | | | | | |
|---|---|---|---|---|---|
| **FAM107B** | 1.0122 | 0.868862165005422 | 0.953 | 0.803 | 1.6578055 |
| **NFATC4** | 1.0876 | -1.04128613854963 | 0.325 | 0.715 | 1.7813231 |
| **DDAH1** | 3.4907 | -1.29232334387483 | 0.472 | 0.799 | 5.7168532 |
| **BAHCC1** | 3.7695 | -0.87678202922648 | 0.036 | 0.467 | 6.1733210 |
| **ITPKB** | 6.9869 | -0.81905200144887 | 0.093 | 0.535 | 1.1442542 |
| **CPT1A** | 1.2558 | 0.805638011594136 | 0.814 | 0.474 | 2.0567816 |
| **CNTNAP2** | 1.8221 | -2.44445974987884 | 0.745 | 0.869 | 2.9841884 |
| **MPC1** | 1.9925 | 0.812677638648787 | 0.8 | 0.47 | 3.2632025 |
| **MGAT3** | 2.1760 | -0.98307498484202 | 0.311 | 0.698 | 3.5636674 |
| **HOXB2** | 2.3127 | 1.17311876234169 | 0.761 | 0.478 | 3.7875778 |
| **CADM4** | 2.3648 | -0.87510184182661 | 0.157 | 0.58 | 3.8729711 |
| **ZNF385A** | 3.4271 | -0.85261033339467 | 0.328 | 0.736 | 5.6125814 |
| **TSPAN7** | 7.4179 | -1.27593669099922 | 0.078 | 0.511 | 1.2148413 |
| **SLC8B1** | 7.8679 | 0.801703905214021 | 0.711 | 0.36 | 1.2885396 |
| **2,00 NPR** | 9.6458 | -0.81209376916944 | 0.149 | 0.577 | 1.5796963 |
| **NME7** | 1.9067 | -0.80971707300283 | 0.311 | 0.704 | 3.1226661 |
| **NDRG2** | 3.5642 | -0.80434677306261 | 0.191 | 0.613 | 5.8372301 |
| **EML1** | 7.5166 | -1.03821384126274 | 0.657 | 0.871 | 1.2310022 |
| **CYP26A1** | 9.6399 | -2.43731840580207 | 0.159 | 0.576 | 1.5787364 |
| **UST** | 1.0450 | -0.84265672188285 | 0.807 | 0.957 | 1.7114582 |
| **DDIT4** | 1.1576 | 1.47856322276907 | 0.936 | 0.821 | 1.8958628 |
| **FAM102B** | 2.3130 | 0.921141027282039 | 0.866 | 0.644 | 3.7880522 |
| **ARHGAP6** | 2.8775 | -0.82577491697667 | 0.102 | 0.526 | 4.7125772 |
| **PRAC1** | 3.3195 | -0.90930800521444 | 0.23 | 0.635 | 5.4363724 |
| **BMI1** | 4.8023 | 0.927912948731003 | 0.728 | 0.405 | 7.8647428 |
| **HSPG2** | 2.0709 | -0.84634023151181 | 0.186 | 0.602 | 3.3916189 |
| **LIMCH1** | 2.2148 | -1.25044054388993 | 0.53 | 0.803 | 3.6272083 |
| **TNS2** | 2.5597 | -1.10853425079066 | 0.13 | 0.55 | 4.1921530 |
| **GCA** | 5.2641 | -0.86635543167632 | 0.354 | 0.727 | 8.6210628 |
| **MT-ND2** | 5.5168 | -0.88574229997491 | 0.783 | 0.989 | 9.0348649 |
| **IQCH** | 6.8616 | -0.80352850093067 | 0.098 | 0.519 | 1.1237307 |
| **RASGEF1A** | 7.0777 | 0.824396392238529 | 0.878 | 0.655 | 1.1591290 |
| **USP18** | 2.1613 | 0.841340124848564 | 0.736 | 0.414 | 3.5397242 |
| **ADAMTSL4** | 2.8512 | -1.38577446432801 | 0.274 | 0.646 | 4.6695499 |

| | | | | | |
|---|---|---|---|---|---|
| **CCSAP** | 6.0799 | 0.846509543426639 | 0.748 | 0.412 | 9.9571367 |
| **AIG1** | 6.2530 | -0.93504209180975 | 0.389 | 0.754 | 1.0240618 |
| **ZNF703** | 9.0835 | -1.02727785645878 | 0.655 | 0.894 | 1.4876100 |
| **LMO1** | 1.2645 | -1.04579078722328 | 0.108 | 0.518 | 2.0709876 |
| **STK17A** | 1.4928 | 0.813158825743133 | 0.861 | 0.62 | 2.4449158 |
| **S100A13** | 2.7363 | 0.872358800391175 | 1 | 0.994 | 4.4813448 |
| **HIST1H2BA** | 5.6043 | 1.09737938588262 | 0.222 | 0.021 | 9.1782559 |
| **RFX3** | 6.1048 | -0.80852881427323 | 0.364 | 0.741 | 9.9978762 |
| **SOX3** | 1.6067 | -1.41332266730538 | 0.11 | 0.52 | 2.6314029 |
| **SLC7A8** | 2.6475 | 0.814426647410262 | 0.959 | 0.841 | 4.3359548 |
| **DENND2A** | 4.6501 | -0.80916511303947 | 0.252 | 0.643 | 7.6154831 |
| **TMEM131L** | 6.5759 | 0.805786852184949 | 0.939 | 0.798 | 1.0769356 |
| **TJP3** | 6.8773 | 0.914558453613344 | 0.865 | 0.623 | 1.1262987 |
| **SAMD4A** | 9.9830 | -0.95697096403692 | 0.173 | 0.598 | 1.6349198 |
| **TLE4** | 2.2561 | -0.82692611173054 | 0.53 | 0.833 | 3.6948706 |
| **VSNL1** | 1.2367 | 0.966155739576438 | 0.878 | 0.762 | 2.0253514 |
| **MEX3B** | 2.4704 | -0.86926847097308 | 0.14 | 0.546 | 4.0458846 |
| **PDE4D** | 7.1746 | -0.99155358482303 | 0.403 | 0.766 | 1.1749867 |
| **EOGT** | 2.2151 | -0.91687024137361 | 0.88 | 0.973 | 3.6278076 |
| **OVCH2** | 3.0095 | -0.88084643427331 | 0.042 | 0.439 | 4.9287079 |
| **ZNF75D** | 3.5227 | 0.830841860029724 | 0.802 | 0.547 | 5.7692095 |
| **NUAK2** | 3.5742 | -0.84060592239353 | 0.481 | 0.815 | 5.8535122 |
| **ADRA2C** | 4.9764 | -0.99109068666541 | 0.851 | 0.952 | 8.1499996 |
| **RAB17** | 5.2467 | -1.10974682049006 | 0.113 | 0.526 | 8.5925902 |
| **ZBTB3** | 5.2971 | 0.830847065395136 | 0.817 | 0.538 | 8.6751988 |
| **NR0B1** | 6.2482 | -1.65288281842284 | 0.022 | 0.409 | 1.0232728 |
| **ELF1** | 6.3041 | 0.832817295952405 | 0.897 | 0.705 | 1.0324334 |
| **PCSK6** | 8.2762 | -0.80759827112622 | 0.369 | 0.728 | 1.3554079 |
| **VASN** | 8.9738 | -0.80398244390301 | 0.193 | 0.586 | 1.4696477 |
| **PHLDB2** | 9.0068 | -0.88561711213615 | 0.663 | 0.891 | 1.4750550 |
| **COL1A1** | 9.6130 | -1.04067020692259 | 0.841 | 0.957 | 1.5743335 |
| **HEG1** | 1.0065 | -0.84914172872072 | 0.311 | 0.704 | 1.6484515 |
| **LAMA2** | 2.5897 | -0.90573470945619 | 0.193 | 0.583 | 4.2412934 |
| **3,00 F** | 3.2588 | 1.19173014765324 | 0.719 | 0.446 | 5.3370195 |

| | | | | | |
|---|---|---|---|---|---|
| **ZNF165** | 4.0199 | 0.903783849034789 | 0.601 | 0.264 | 6.5834007 |
| **LIMA1** | 2.7633 | 0.89212605873452 | 0.927 | 0.814 | 4.5255678 |
| **RHOBTB3** | 1.1265 | -1.42792900049058 | 0.078 | 0.476 | 1.8449382 |
| **GRPR** | 1.2727 | -1.07480699971534 | 0.03 | 0.41 | 2.0844189 |
| **CDHR1** | 2.5834 | 0.83155766855518 | 0.694 | 0.356 | 4.2309369 |
| **CAVIN3** | 2.7084 | -0.87231112972942 | 0.328 | 0.695 | 4.4355707 |
| **LTBP1** | 4.8134 | -0.80137295699219 | 0.55 | 0.818 | 7.8830630 |
| **HSPB8** | 5.1203 | -0.91548587047463 | 0.164 | 0.591 | 8.3856566 |
| **VLDLR** | 5.6727 | -0.89496826362931 | 0.526 | 0.838 | 9.2902072 |
| **IER3** | 6.3850 | -1.27650050864927 | 0.399 | 0.742 | 1.0456757 |
| **SPIN3** | 8.1737 | -0.94738743346816 | 0.431 | 0.771 | 1.3386226 |
| **HES6** | 4.3660 | -0.83361957381817 | 0.384 | 0.73 | 7.1502310 |
| **TRPC6** | 5.6739 | -1.29146272330133 | 0.168 | 0.537 | 9.2922491 |
| **INSYN1** | 6.7682 | -1.02493176510209 | 0.157 | 0.528 | 1.1084384 |
| **PIK3R1** | 1.0565 | -0.96002257408245 | 0.503 | 0.798 | 1.7303031 |
| **RAB20** | 1.5757 | -1.01421442650391 | 0.66 | 0.877 | 2.5806220 |
| **EDN1** | 1.6537 | -1.51006506027922 | 0.059 | 0.44 | 2.7084201 |
| **DHRS3** | 1.7840 | -1.06338485502516 | 0.264 | 0.615 | 2.9217234 |
| **EPOP** | 2.0508 | -0.84067743870132 | 0.565 | 0.835 | 3.3586152 |
| **SLC45A4** | 6.7753 | 0.816580568274246 | 0.797 | 0.524 | 1.1096011 |
| **RGS3** | 7.3758 | -0.83836365378051 | 0.1 | 0.485 | 1.2079402 |
| **1,00 DKK** | 1.8148 | -2.73225892942838 | 0.049 | 0.418 | 2.9722068 |
| **IRX2** | 1.8867 | -0.95367383863135 | 0.802 | 0.947 | 3.0899013 |
| **DUSP10** | 3.1655 | -0.81490301141310 | 0.36 | 0.721 | 5.1843007 |
| **FAM13A** | 3.1857 | -0.96240249087114 | 0.589 | 0.854 | 5.2172726 |
| **EGLN3** | 4.2958 | 1.14939786131311 | 0.934 | 0.839 | 7.0352903 |
| **ADCY2** | 4.8203 | -1.07616069899169 | 0.069 | 0.441 | 7.8942735 |
| **GRHL3** | 6.8560 | -1.12118084941236 | 0.125 | 0.508 | 1.1228211 |
| **NKX1-2** | 1.4876 | -1.07788575311537 | 0.892 | 0.931 | 2.4363010 |
| **TNFRSF12A** | 2.2776 | -0.92618450936406 | 0.956 | 0.965 | 3.7301838 |
| **MEIS3** | 2.9480 | -1.55606445547046 | 0.679 | 0.818 | 4.8280392 |
| **ISLR** | 6.2920 | -0.98582678719879 | 0.11 | 0.477 | 1.0304441 |
| **PRKAR2B** | 7.3521 | -0.97489157651120 | 0.535 | 0.821 | 1.2040690 |
| **MAF** | 1.5295 | -0.98005906155876 | 0.191 | 0.563 | 2.5049989 |

| | | | | | |
|---|---|---|---|---|---|
| **CDX2** | 5.1510 | -1.47081996929729 | 0.086 | 0.465 | 8.4358371 |
| **ARSI** | 5.4420 | 0.891453461468588 | 0.824 | 0.556 | 8.9123777 |
| **EVA1B** | 7.3408 | -0.83386989968207 | 0.376 | 0.702 | 1.2022039 |
| **CD82** | 9.5109 | -0.85429896585670 | 0.391 | 0.712 | 1.5576042 |
| **CER1** | 1.8778 | -2.7930781334801 | 0.311 | 0.695 | 3.0754229 |
| **LEFTY1** | 1.9309 | 0.8202454492109 | 0.99 | 0.965 | 3.1622692 |
| **COBL** | 5.4555 | -1.00409338672736 | 0.621 | 0.831 | 8.9345500 |
| **ROR2** | 8.3879 | -1.096132890297 | 0.587 | 0.846 | 1.3736969 |
| **AJAP1** | 3.0981 | -0.83517047577248 | 0.196 | 0.558 | 5.0738083 |
| **WNT3A** | 7.6307 | -1.66282097216701 | 0.169 | 0.508 | 1.2496929 |
| **CCDC92** | 1.9842 | -0.8134798709261 | 0.059 | 0.416 | 3.2496218 |
| **SLIT2** | 4.2159 | -1.02024148255095 | 0.019 | 0.36 | 6.9044699 |
| **ANKRD1** | 4.2335 | -1.5127129794644 | 0.118 | 0.489 | 6.9333414 |
| **GJA5** | 8.3920 | -0.85173253246654 | 0.042 | 0.393 | 1.3743630 |
| **SIAH2** | 1.5546 | -0.80689406238463 | 0.387 | 0.716 | 2.5460071 |
| **CCPG1** | 2.2574 | 0.87735824514975 | 0.983 | 0.898 | 3.6970539 |
| **CCN2** | 6.5990 | -1.03940062644631 | 0.832 | 0.942 | 1.0807210 |
| **RGMA** | 3.0298 | -0.91689383014820 | 0.585 | 0.815 | 4.9620410 |
| **TGFB1I1** | 3.6431 | -0.85205256581824 | 0.264 | 0.612 | 5.9663472 |
| **AP3B2** | 3.9031 | -1.06504761344547 | 0.22 | 0.547 | 6.3922512 |
| **VWA5A** | 3.0084 | -0.95651887775633 | 0.332 | 0.652 | 4.9270186 |
| **COL5A1** | 5.0164 | -1.1376143117169 | 0.609 | 0.845 | 8.2154038 |
| **TRIM22** | 5.6582 | -1.05449886218206 | 0.24 | 0.575 | 9.2665240 |
| **SYNE3** | 7.6156 | -0.80301999239206 | 0.056 | 0.395 | 1.2472086 |
| **COL13A1** | 7.9174 | -1.24446384222735 | 0.029 | 0.359 | 1.2966449 |
| **EFNA1** | 8.5713 | -0.86673173954511 | 0.266 | 0.621 | 1.4037310 |
| **OTX2** | 1.7413 | -1.12998904046039 | 0.65 | 0.879 | 2.8518060 |
| **MSX1** | 2.4541 | -1.43365783934269 | 0.042 | 0.378 | 4.0191659 |
| **MT-ND5** | 2.5849 | 1.70408936347175 | 0.851 | 0.75 | 4.2333405 |
| **PREX1** | 4.2995 | -1.15047505372827 | 0.69 | 0.858 | 7.0413885 |
| **MAML3** | 2.3094 | -0.99182687223722 | 0.423 | 0.718 | 3.7821211 |
| **S100A16** | 6.3867 | 1.15126453044139 | 0.706 | 0.432 | 1.0459550 |
| **CYP1B1** | 1.1734 | -1.19847027648359 | 0.255 | 0.585 | 1.9217399 |
| **CCNG2** | 1.2754 | 1.10557347745664 | 0.788 | 0.583 | 2.0888716 |

| | | | | | |
|---|---|---|---|---|---|
| **IL13RA1** | 3.0358 | -0.98441143540695 | 0.205 | 0.548 | 4.9717778 |
| **TACR1** | 1.2665 | -1.62441243957633 | 0.086 | 0.413 | 2.0742250 |
| **PCSK1** | 2.3153 | 1.15925936401221 | 0.856 | 0.691 | 3.7918161 |
| **LIPG** | 6.1136 | -0.99970724309573 | 0.144 | 0.484 | 1.0012392 |
| **FGF17** | 8.6933 | -2.90109898119193 | 0.371 | 0.621 | 1.4237074 |
| **E2F1** | 2.1774 | -0.83394320129877 | 0.953 | 0.972 | 3.5659471 |
| **FAM20A** | 2.2227 | -1.12962735440117 | 0.173 | 0.493 | 3.6402466 |
| **COLEC12** | 4.6025 | -1.48807866186053 | 0.313 | 0.626 | 7.5375160 |
| **TUBB2A** | 8.6587 | -0.84296025194270 | 0.997 | 0.999 | 1.4180427 |
| **FAM43A** | 8.7499 | -0.83291323596274 | 0.137 | 0.466 | 1.4329728 |
| **PIMREG** | 8.8356 | -0.83636271932656 | 0.567 | 0.79 | 1.4470164 |
| **MFAP4** | 2.7049 | -1.00187539737579 | 0.059 | 0.362 | 4.4298706 |
| **MKI67** | 3.2853 | -0.85048151696623 | 0.954 | 0.981 | 5.3803573 |
| **LRRN1** | 3.8517 | -1.19357995918417 | 0.538 | 0.774 | 6.3080534 |
| **DNMT3B** | 2.0644 | 0.823450081629536 | 1 | 1 | 3.3809121 |
| **ENC1** | 3.5519 | -0.89242169973298 | 0.628 | 0.88 | 5.8169779 |
| **BMP2** | 6.9238 | -2.04720693091834 | 0.21 | 0.508 | 1.1339190 |
| **PPP1R1A** | 1.1269 | -0.85577233681457 | 0.587 | 0.795 | 1.8455683 |
| **APCDD1** | 1.7618 | -1.0721170294295 | 0.315 | 0.599 | 2.8854255 |
| **HS6ST1** | 9.1259 | -0.86338117002976 | 0.909 | 0.971 | 1.4945500 |
| **LRATD2** | 5.9476 | -0.82211591995708 | 0.949 | 0.965 | 9.7404640 |
| **GLI3** | 9.9836 | -0.84331644068448 | 0.257 | 0.552 | 1.6350300 |
| **CHAC1** | 1.0229 | 1.0627134447654 | 0.714 | 0.501 | 1.6752249 |
| **SAMD3** | 1.3218 | -0.91144275665301 | 0.022 | 0.289 | 2.1647355 |
| **TCF7L1** | 2.6027 | -0.92213397487446 | 0.416 | 0.642 | 4.2625551 |
| **B3GNT7** | 3.0844 | -0.99030937244951 | 0.68 | 0.808 | 5.0514483 |
| **APLNR** | 3.2109 | -2.51397846552302 | 0.494 | 0.701 | 5.2585183 |
| **TBXT** | 1.0740 | -1.11279863973025 | 0.601 | 0.844 | 1.7588951 |
| **GAL** | 2.7547 | -1.62607746571806 | 0.992 | 0.981 | 4.5114410 |
| **CRLF1** | 6.1704 | -1.73188013452404 | 0.609 | 0.735 | 1.0105317 |
| **SNAI2** | 6.9105 | -1.32588379549086 | 0.032 | 0.292 | 1.1317341 |
| **COL5A2** | 4.2051 | -1.36159549520655 | 0.176 | 0.449 | 6.8867153 |
| **ANKRD33B** | 9.9730 | -0.84114853109591 | 0.257 | 0.525 | 1.6332800 |
| **TBX6** | 3.9631 | -1.88549714393854 | 0.073 | 0.324 | 6.4904421 |

| | | | | | |
|---|---|---|---|---|---|
| **ACSS3** | 9.9584 | -0.88004082778072 | 0.494 | 0.705 | 1.6308923 |
| **FOXJ1** | 2.1632 | -0.87820048578177 | 0.115 | 0.376 | 3.5427186 |
| **VIT** | 4.1032 | -0.87454247831857 | 0.095 | 0.341 | 6.7199184 |
| **SERPINB9** | 8.0683 | 0.808412523721394 | 0.971 | 0.952 | 1.3213591 |
| **DAB1** | 3.9755 | -0.85853503851626 | 0.364 | 0.575 | 6.5106880 |
| **TPX2** | 1.9611 | -0.81748491316623 | 0.99 | 0.997 | 3.2117129 |
| **CCN1** | 2.4612 | -1.01914289271486 | 0.853 | 0.91 | 4.0307495 |
| **MIXL1** | 2.5640 | -1.81779415363263 | 0.898 | 0.93 | 4.1991927 |
| **HPD** | 3.8805 | -0.86034598217427 | 0.266 | 0.502 | 6.3551844 |
| **IQSEC2** | 2.1444 | 1.01795047928887 | 0.653 | 0.494 | 3.5120443 |
| **C11orf96** | 4.4277 | -0.87953705718392 | 0.249 | 0.506 | 7.2513214 |
| **GSC** | 1.0611 | -1.46477194044204 | 0.081 | 0.305 | 1.7378248 |
| **GMPR** | 1.5660 | -1.11986982673269 | 0.377 | 0.552 | 2.5646941 |
| **ANOS1** | 1.5815 | -1.44985868654699 | 0.208 | 0.411 | 2.5900807 |
| **FOXC1** | 4.1983 | -0.97060337313558 | 0.183 | 0.419 | 6.8756546 |
| **PCDH18** | 3.5324 | -0.85898252959158 | 0.592 | 0.759 | 5.7851591 |
| **HAPLN1** | 4.0885 | 1.06353796429424 | 0.384 | 0.177 | 6.6957927 |
| **FOXA2** | 1.3382 | -1.00534715341309 | 0.054 | 0.25 | 2.1916804 |
| **GATA6** | 1.9739 | -1.72381511085016 | 0.144 | 0.351 | 3.2326599 |
| **LHX1** | 8.1821 | -1.51664726461569 | 0.044 | 0.222 | 1.3399869 |
| **PLXNA2** | 4.0985 | -1.56630196678764 | 0.217 | 0.404 | 6.7121927 |
| **MSGN1** | 4.4561 | -1.59849719528264 | 0.064 | 0.235 | 7.2978916 |
| **DLL1** | 1.5563 | -0.93175616486432 | 0.076 | 0.205 | 2.5487670 |
| **NPPB** | 1.5094 | -0.96452362352195 | 0.135 | 0.274 | 2.4720669 |
| **PLAU** | 7.7507 | 0.943612154881905 | 0.474 | 0.369 | 1.2693428 |
| **ACTA1** | 2.2584 | -0.85755339924000 | 0.426 | 0.534 | 0.0036986 |
| **DLL3** | 4.5447 | -1.19588222243866 | 0.755 | 0.578 | 0.0074429 |
| **ACTG2** | 6.6429 | -1.09581605888716 | 0.792 | 0.82 | 0.1087910 |
| **SLC34A2** | 0.1269 | 1.08246455964382 | 0.113 | 0.138 | 1 |
| **CLDN10** | 0.9003 | -0.98730803985687 | 0.513 | 0.462 | 1 |

Supplemental Table S2_Sheet_7

| | p_val | avg_log2FC | pct.1 | pct.2 | p_val_adj |
|---|---|---|---|---|---|
| **NOC2L** | 0 | 0.818225244874983 | 1 | 0.985 | 0 |
| **AGRN** | 0 | -1.4487889623252 | 0.884 | 0.988 | 0 |
| **VWA1** | 0 | 1.07441644191336 | 1 | 0.958 | 0 |
| **ATAD3C** | 0 | 1.25849788538812 | 0.779 | 0.3 | 0 |
| **ATAD3B** | 0 | 1.54064824326904 | 1 | 0.956 | 0 |
| **TMEM52** | 0 | 0.891319685579904 | 0.678 | 0.322 | 0 |
| **AJAP1** | 0 | -0.82421036330608 | 0.184 | 0.585 | 0 |
| **TNFRSF25** | 0 | 0.958883515795767 | 0.616 | 0.19 | 0 |
| **PLEKHG5** | 0 | 0.855084077263857 | 0.646 | 0.266 | 0 |
| **SLC25A33** | 0 | 0.930382817797909 | 0.974 | 0.811 | 0 |
| **CLSTN1** | 0 | -1.48563225278875 | 0.984 | 0.998 | 0 |
| **CTNNBIP1** | 0 | -0.930024035113595 | 0.831 | 0.949 | 0 |
| **SRM** | 0 | 1.26944108650617 | 1 | 0.999 | 0 |
| **MTOR** | 0 | 0.812931013754329 | 0.997 | 0.948 | 0 |
| **DRAXIN** | 0 | -2.36958653647701 | 0.464 | 0.937 | 0 |
| **MIIP** | 0 | 1.40030121078363 | 1 | 0.961 | 0 |
| **TNFRSF8** | 0 | 1.29337732993602 | 0.654 | 0.046 | 0 |
| **DHRS3** | 0 | -1.14745630139169 | 0.415 | 0.814 | 0 |
| **PDPN** | 0 | -1.59705428539621 | 0.893 | 0.982 | 0 |
| **TMEM51** | 0 | -0.98035953690266 | 0.402 | 0.749 | 0 |
| **EFHD2** | 0 | 1.46696722404752 | 0.999 | 0.927 | 0 |
| **EPHA2** | 0 | -1.60911990553704 | 0.074 | 0.688 | 0 |
| **MFAP2** | 0 | -1.35256805387909 | 0.917 | 0.99 | 0 |
| **SDHB** | 0 | 1.12960379518933 | 1 | 0.989 | 0 |
| **ARHGEF10L** | 0 | -1.08411906770232 | 0.074 | 0.659 | 0 |
| **IFFO2** | 0 | -0.830865864584134 | 0.203 | 0.593 | 0 |
| **PLA2G2A** | 0 | -1.19328050891692 | 0.001 | 0.412 | 0 |
| **ECE1** | 0 | -2.28563897574345 | 0.738 | 0.981 | 0 |
| **LUZP1** | 0 | -0.952205109260533 | 0.782 | 0.954 | 0 |
| **ID3** | 0 | -2.37788995527466 | 0.028 | 0.553 | 0 |
| **PITHD1** | 0 | 0.845380768199206 | 1 | 0.997 | 0 |
| **FUCA1** | 0 | 1.84970192388783 | 1 | 0.749 | 0 |

| | | | | | |
|---|---|---|---|---|---|
| **CNR2** | 0 | 1.4620636739958 | 0.743 | 0.086 | 0 |
| **GRHL3** | 0 | -2.21861992357269 | 0.048 | 0.715 | 0 |
| **STMN1** | 0 | 0.968823767529398 | 1 | 0.988 | 0 |
| **ZNF593** | 0 | 0.807460988906461 | 0.998 | 0.949 | 0 |
| **CEP85** | 0 | 0.80117869868963 | 0.981 | 0.863 | 0 |
| **SH3BGRL3** | 0 | -0.874667424140315 | 0.897 | 0.962 | 0 |
| **LIN28A** | 0 | -0.883841258365386 | 1 | 1 | 0 |
| **SLC9A1** | 0 | 1.30533110717482 | 0.991 | 0.768 | 0 |
| **IFI6** | 0 | -0.871249358939893 | 0.025 | 0.435 | 0 |
| **SMPDL3B** | 0 | -2.2804291769404 | 0.013 | 0.834 | 0 |
| **SESN2** | 0 | 1.47995506874153 | 0.995 | 0.839 | 0 |
| **RCC1** | 0 | 0.870538910699827 | 0.999 | 0.945 | 0 |
| **PTPRU** | 0 | 1.94026428019027 | 1 | 0.932 | 0 |
| **NKAIN1** | 0 | -0.96879035358157 | 0.036 | 0.437 | 0 |
| **MARCKSL1** | 0 | -2.00174937442681 | 1 | 1 | 0 |
| **KIAA1522** | 0 | -1.89506733082078 | 0.872 | 0.992 | 0 |
| **TMEM54** | 0 | 0.806249329210726 | 0.58 | 0.165 | 0 |
| **RNF19B** | 0 | 1.86112098837568 | 1 | 0.904 | 0 |
| **ZNF362** | 0 | -1.68332576341382 | 0.441 | 0.928 | 0 |
| **PHC2** | 0 | -1.33026423859141 | 0.025 | 0.501 | 0 |
| **SH3D21** | 0 | 0.925751890517376 | 0.719 | 0.346 | 0 |
| **C1orf109** | 0 | 0.870619945286803 | 0.995 | 0.921 | 0 |
| **YRDC** | 0 | 0.854968013287966 | 0.998 | 0.931 | 0 |
| **MYCBP** | 0 | 1.00233643331949 | 0.962 | 0.651 | 0 |
| **YBX1** | 0 | -0.917282528990117 | 1 | 1 | 0 |
| **KLF17** | 0 | 1.33259428700637 | 0.468 | 0.033 | 0 |
| **PRDX1** | 0 | 1.55414943935437 | 1 | 0.994 | 0 |
| **UQCRH** | 0 | 0.956335448475532 | 1 | 0.997 | 0 |
| **FAF1** | 0 | 0.910064550686081 | 1 | 0.985 | 0 |
| **NRDC** | 0 | 0.981110441950946 | 1 | 1 | 0 |
| **RAB3B** | 0 | -1.72987052240101 | 0.911 | 0.988 | 0 |
| **ZFYVE9** | 0 | 0.890722846531233 | 0.999 | 0.974 | 0 |
| **ZYG11A** | 0 | 1.06254702340578 | 0.999 | 0.884 | 0 |
| **LRP8** | 0 | -1.42351413143492 | 0.346 | 0.844 | 0 |

| | | | | | |
|---|---|---|---|---|---|
| **FAM151A** | 0 | 1.37362865134566 | 0.349 | 0.012 | 0 |
| **DHCR24** | 0 | -1.61053741057085 | 0.997 | 1 | 0 |
| **PLPP3** | 0 | -1.74475063116797 | 0.387 | 0.938 | 0 |
| **HOOK1** | 0 | -0.94432591913286 | 0.185 | 0.698 | 0 |
| **PATJ** | 0 | -1.30600316607806 | 0.749 | 0.96 | 0 |
| **PGM1** | 0 | -2.05007272373078 | 0.264 | 0.93 | 0 |
| **ROR1** | 0 | -2.38249361613508 | 0.166 | 0.843 | 0 |
| **CACHD1** | 0 | -1.50058430592691 | 0.494 | 0.901 | 0 |
| **RAVER2** | 0 | -0.982270377911844 | 0.144 | 0.673 | 0 |
| **AK4** | 0 | 0.846284568832663 | 0.995 | 0.868 | 0 |
| **WLS** | 0 | -4.64955033544866 | 0.424 | 0.952 | 0 |
| **CTH** | 0 | 1.86998644545411 | 0.987 | 0.61 | 0 |
| **ST6GALNAC5** | 0 | -1.72426190106587 | 0.034 | 0.72 | 0 |
| **DNAJB4** | 0 | 1.26365311522517 | 0.946 | 0.588 | 0 |
| **ADGRL2** | 0 | -1.48597975675434 | 0.806 | 0.981 | 0 |
| **DDAH1** | 0 | -1.35839794631249 | 0.122 | 0.684 | 0 |
| **CCN1** | 0 | -1.60555582243271 | 0.516 | 0.752 | 0 |
| **LMO4** | 0 | 1.248755688263 | 1 | 0.939 | 0 |
| **ZNF644** | 0 | -0.849738825920298 | 0.995 | 1 | 0 |
| **TGFBR3** | 0 | 1.5944948675598 | 0.852 | 0.169 | 0 |
| **BRDT** | 0 | 2.8284498001143 | 0.997 | 0.2 | 0 |
| **FNBP1L** | 0 | -1.18104690001916 | 0.992 | 1 | 0 |
| **ARHGAP29** | 0 | -0.941774795438514 | 0.156 | 0.627 | 0 |
| **PTBP2** | 0 | -1.17852808378511 | 0.744 | 0.96 | 0 |
| **AGL** | 0 | 2.2902243755046 | 1 | 0.956 | 0 |
| **EXTL2** | 0 | -0.981505730269616 | 0.564 | 0.9 | 0 |
| **PRMT6** | 0 | -1.0972171651272 | 0.326 | 0.837 | 0 |
| **NTNG1** | 0 | -0.942319753627628 | 0.016 | 0.497 | 0 |
| **SLC25A24** | 0 | -1.42188072101888 | 0.813 | 0.985 | 0 |
| **HENMT1** | 0 | 1.45457910116218 | 0.779 | 0.088 | 0 |
| **WDR47** | 0 | 1.06066009245934 | 0.81 | 0.457 | 0 |
| **TAF13** | 0 | 2.72367154274564 | 1 | 0.911 | 0 |
| **CELSR2** | 0 | -0.877726670269706 | 0.192 | 0.683 | 0 |
| **SORT1** | 0 | -0.863409909138829 | 0.87 | 0.98 | 0 |

| | | | | | |
|---|---|---|---|---|---|
| **GNAI3** | 0 | -0.875012621448426 | 0.995 | 1 | 0 |
| **WDR77** | 0 | -0.87429504847561 | 0.99 | 0.998 | 0 |
| **PHTF1** | 0 | -0.986706506758611 | 0.601 | 0.904 | 0 |
| **OLFML3** | 0 | -5.3151728803883 | 0.592 | 0.986 | 0 |
| **SYT6** | 0 | -1.51660357265312 | 0.058 | 0.764 | 0 |
| **VANGL1** | 0 | -1.09011839083322 | 0.63 | 0.91 | 0 |
| **IGSF3** | 0 | -0.811163716579782 | 0.522 | 0.838 | 0 |
| **PHGDH** | 0 | 1.13649043840775 | 1 | 0.977 | 0 |
| **NOTCH2** | 0 | -1.46587405839978 | 0.205 | 0.833 | 0 |
| **SV2A** | 0 | -1.26372551178774 | 0.148 | 0.717 | 0 |
| **PLEKHO1** | 0 | -1.30833190805611 | 0.38 | 0.814 | 0 |
| **ENSA** | 0 | -0.81520553805929 | 0.963 | 0.996 | 0 |
| **HORMAD1** | 0 | 3.16511929228549 | 0.999 | 0.206 | 0 |
| **MLLT11** | 0 | -1.30240902691214 | 0.276 | 0.743 | 0 |
| **RFX5** | 0 | -0.834284389143491 | 0.51 | 0.833 | 0 |
| **POGZ** | 0 | -1.05890070272158 | 0.792 | 0.977 | 0 |
| **S100A10** | 0 | 0.831494973883348 | 1 | 0.987 | 0 |
| **S100A16** | 0 | 2.18528255219738 | 0.845 | 0.225 | 0 |
| **S100A13** | 0 | 3.03618976515613 | 1 | 0.96 | 0 |
| **RAB13** | 0 | -1.29480890664279 | 0.832 | 0.98 | 0 |
| **CKS1B** | 0 | 1.45474253279224 | 1 | 0.963 | 0 |
| **FLAD1** | 0 | 1.05085681401919 | 1 | 0.962 | 0 |
| **THBS3** | 0 | -1.12017138518021 | 0.032 | 0.559 | 0 |
| **FDPS** | 0 | -0.926794120912158 | 0.891 | 0.963 | 0 |
| **YY1AP1** | 0 | 1.05319567840162 | 0.997 | 0.93 | 0 |
| **MEX3A** | 0 | -2.15210491143786 | 0.075 | 0.825 | 0 |
| **LMNA** | 0 | 0.874898047254872 | 0.999 | 0.974 | 0 |
| **SLC25A44** | 0 | 1.34533975823178 | 1 | 0.978 | 0 |
| **PMF1** | 0 | -0.83942184843392 | 0.967 | 0.996 | 0 |
| **NAXE** | 0 | -0.921549859168071 | 0.781 | 0.966 | 0 |
| **NES** | 0 | -1.64658035595251 | 0.997 | 1 | 0 |
| **CRABP2** | 0 | -1.66233681320176 | 0.994 | 0.988 | 0 |
| **DUSP23** | 0 | 1.6167240322949 | 0.825 | 0.077 | 0 |
| **PIGM** | 0 | -0.853772609353849 | 0.263 | 0.707 | 0 |

| | | | | | |
|---|---|---|---|---|---|
| **IGSF8** | 0 | -1.49886518903652 | 0.733 | 0.955 | 0 |
| **ATP1A2** | 0 | -0.923774353577801 | 0.002 | 0.397 | 0 |
| **PEA15** | 0 | -1.6443150835765 | 0.905 | 0.989 | 0 |
| **VANGL2** | 0 | -1.36146164262019 | 0.593 | 0.933 | 0 |
| **F11R** | 0 | 1.34307664132491 | 1 | 0.993 | 0 |
| **NECTIN4** | 0 | 1.01049815976387 | 0.51 | 0.073 | 0 |
| **UHMK1** | 0 | 1.43437445481104 | 0.995 | 0.84 | 0 |
| **RGS5** | 0 | -2.8243134308163 | 0.075 | 0.778 | 0 |
| **MGST3** | 0 | -2.4097494442606 | 0.553 | 0.992 | 0 |
| **ALDH9A1** | 0 | -0.849784790421666 | 0.863 | 0.983 | 0 |
| **UCK2** | 0 | -1.1118995945849 | 0.873 | 0.985 | 0 |
| **NME7** | 0 | -0.801497551340321 | 0.183 | 0.648 | 0 |
| **ANKRD45** | 0 | 1.3101180135597 | 0.539 | 0.068 | 0 |
| **RABGAP1L** | 0 | -0.832664343026732 | 0.791 | 0.938 | 0 |
| **SOAT1** | 0 | 0.857204812153147 | 0.989 | 0.886 | 0 |
| **RO60** | 0 | -1.03021717718873 | 0.86 | 0.986 | 0 |
| **GLRX2** | 0 | -1.00168450943006 | 0.08 | 0.63 | 0 |
| **CDC73** | 0 | -0.948342504432812 | 0.914 | 0.993 | 0 |
| **ELF3** | 0 | 1.13443573511443 | 0.964 | 0.607 | 0 |
| **PIK3C2B** | 0 | -0.853074286554503 | 0.231 | 0.67 | 0 |
| **MDM4** | 0 | 1.04150869509623 | 1 | 0.985 | 0 |
| **LRRN2** | 0 | -1.37772979404735 | 0.005 | 0.606 | 0 |
| **NUAK2** | 0 | -1.06712625967911 | 0.199 | 0.633 | 0 |
| **NUCKS1** | 0 | -1.01798051620253 | 1 | 1 | 0 |
| **SERTAD4** | 0 | -1.46141591321052 | 0.008 | 0.659 | 0 |
| **TRAF5** | 0 | -2.01046841167963 | 0.023 | 0.807 | 0 |
| **SMYD2** | 0 | -1.21710152140635 | 0.85 | 0.971 | 0 |
| **PTPN14** | 0 | -1.19114646441609 | 0.828 | 0.978 | 0 |
| **DUSP10** | 0 | -1.40766386116571 | 0.253 | 0.817 | 0 |
| **CAPN2** | 0 | -2.23710948652156 | 0.15 | 0.858 | 0 |
| **CNIH4** | 0 | 0.873361343334781 | 1 | 0.89 | 0 |
| **LEFTY2** | 0 | -3.11812402247443 | 0.872 | 0.952 | 0 |
| **MIXL1** | 0 | -2.21763633263925 | 0.548 | 0.743 | 0 |
| **ITPKB** | 0 | -0.804930334378837 | 0.034 | 0.402 | 0 |

| | | | | | |
|---|---|---|---|---|---|
| **C1orf198** | 0 | -1.26202640228758 | 0.279 | 0.779 | 0 |
| **EGLN1** | 0 | -1.19773648898112 | 0.364 | 0.875 | 0 |
| **SIPA1L2** | 0 | -1.29366511003128 | 0.707 | 0.941 | 0 |
| **PCNX2** | 0 | -0.800540575597552 | 0.31 | 0.718 | 0 |
| **GNG4** | 0 | 1.21548241038106 | 0.645 | 0.123 | 0 |
| **NID1** | 0 | 0.829440703310857 | 0.63 | 0.214 | 0 |
| **CHML** | 0 | -1.21240724739051 | 0.306 | 0.842 | 0 |
| **ZBTB18** | 0 | 1.187860789489 | 0.962 | 0.66 | 0 |
| **DESI2** | 0 | 0.987669378040325 | 0.998 | 0.959 | 0 |
| **ZNF695** | 0 | 0.960779125406191 | 0.976 | 0.775 | 0 |
| **ZNF672** | 0 | -0.848365880825245 | 0.199 | 0.65 | 0 |
| **PXDN** | 0 | 1.0353469479896 | 0.994 | 0.73 | 0 |
| **ID2** | 0 | -1.89912638147694 | 0.117 | 0.811 | 0 |
| **IAH1** | 0 | 1.66210238723275 | 0.836 | 0.087 | 0 |
| **SLC66A3** | 0 | 0.941298211785243 | 0.942 | 0.68 | 0 |
| **LPIN1** | 0 | 1.84938070437475 | 0.988 | 0.48 | 0 |
| **TRIB2** | 0 | -1.01957374612735 | 0.136 | 0.594 | 0 |
| **WDR35** | 0 | -0.938097158459169 | 0.121 | 0.634 | 0 |
| **SDC1** | 0 | -1.04782599283904 | 0.035 | 0.484 | 0 |
| **RHOB** | 0 | -1.79598325462046 | 0.947 | 0.98 | 0 |
| **TP53I3** | 0 | -0.955173146328023 | 0.373 | 0.783 | 0 |
| **EFR3B** | 0 | 1.40527198608564 | 0.994 | 0.771 | 0 |
| **POMC** | 0 | -3.10873281658823 | 0.433 | 0.868 | 0 |
| **CENPA** | 0 | 1.1470241115451 | 0.989 | 0.788 | 0 |
| **KRTCAP3** | 0 | 2.33566806686603 | 0.981 | 0.291 | 0 |
| **FNDC4** | 0 | 1.74752552121677 | 0.943 | 0.504 | 0 |
| **ZNF512** | 0 | -1.05040670598343 | 0.341 | 0.83 | 0 |
| **CAPN13** | 0 | -1.2842528397595 | 0.024 | 0.69 | 0 |
| **LTBP1** | 0 | -0.995497937800123 | 0.437 | 0.824 | 0 |
| **FEZ2** | 0 | 1.4728701206437 | 0.999 | 0.91 | 0 |
| **VIT** | 0 | -1.20546061553404 | 0.002 | 0.352 | 0 |
| **EIF2AK2** | 0 | -1.25290143716429 | 0.332 | 0.881 | 0 |
| **PRKD3** | 0 | -0.868480906609451 | 0.654 | 0.923 | 0 |
| **QPCT** | 0 | -2.18923811578571 | 0.219 | 0.778 | 0 |

| | | | | | |
|---|---|---|---|---|---|
| **CDC42EP3** | 0 | -1.00547591191808 | 0.151 | 0.652 | 0 |
| **HNRNPLL** | 0 | -0.855536949989245 | 0.916 | 0.993 | 0 |
| **PKDCC** | 0 | -2.67938811618771 | 0.263 | 0.726 | 0 |
| **EML4** | 0 | -1.04599139679307 | 0.883 | 0.988 | 0 |
| **SIX2** | 0 | 1.78905316583674 | 0.817 | 0.164 | 0 |
| **CALM2** | 0 | -1.03885267907444 | 0.992 | 0.997 | 0 |
| **KCNK12** | 0 | -1.08685106063058 | 0.108 | 0.496 | 0 |
| **FOXN2** | 0 | 0.830350510331169 | 0.926 | 0.656 | 0 |
| **NRXN1** | 0 | -0.91985370113147 | 0.035 | 0.501 | 0 |
| **RTN4** | 0 | -1.0589834576597 | 0.989 | 1 | 0 |
| **CCDC88A** | 0 | -1.10020004638436 | 0.772 | 0.967 | 0 |
| **FAM161A** | 0 | -0.878229773312664 | 0.2 | 0.676 | 0 |
| **B3GNT2** | 0 | -1.65160470142628 | 0.747 | 0.978 | 0 |
| **CEP68** | 0 | -1.4048555009371 | 0.263 | 0.861 | 0 |
| **SPRED2** | 0 | -1.33020433419808 | 0.325 | 0.852 | 0 |
| **ANTXR1** | 0 | -1.21328370705494 | 0.049 | 0.646 | 0 |
| **GFPT1** | 0 | 1.14415917969983 | 0.999 | 0.966 | 0 |
| **ADD2** | 0 | -0.953987520623093 | 0.851 | 0.945 | 0 |
| **ACTG2** | 0 | -2.06836376496509 | 0.548 | 0.791 | 0 |
| **MTHFD2** | 0 | 2.0421476082293 | 1 | 0.99 | 0 |
| **WBP1** | 0 | -0.907818980646723 | 0.728 | 0.94 | 0 |
| **HK2** | 0 | 1.52549666546507 | 1 | 0.875 | 0 |
| **TMSB10** | 0 | -0.841221902723983 | 1 | 1 | 0 |
| **GGCX** | 0 | -0.824561426681524 | 0.351 | 0.765 | 0 |
| **VAMP8** | 0 | -1.73892894107381 | 0.959 | 0.99 | 0 |
| **REEP1** | 0 | 0.902328520647725 | 0.487 | 0.095 | 0 |
| **FOXI3** | 0 | -1.63817009303021 | 0.806 | 0.971 | 0 |
| **MAP4K4** | 0 | -1.57299689612219 | 0.985 | 0.997 | 0 |
| **DPP10** | 0 | -1.63963729958186 | 0.201 | 0.794 | 0 |
| **STEAP3** | 0 | -1.05731476596393 | 0.541 | 0.829 | 0 |
| **DBI** | 0 | -0.880216680784258 | 0.959 | 0.992 | 0 |
| **TMEM177** | 0 | -1.06837213737046 | 0.329 | 0.827 | 0 |
| **TMEM185B** | 0 | -0.910961705418443 | 0.658 | 0.931 | 0 |
| **GLI2** | 0 | -2.07270733621505 | 0.012 | 0.844 | 0 |

| | | | | | |
|---|---|---|---|---|---|
| **TFCP2L1** | 0 | 1.23585563767256 | 0.934 | 0.444 | 0 |
| **BIN1** | 0 | -1.11285274050253 | 0.802 | 0.945 | 0 |
| **HS6ST1** | 0 | -1.0367214160421 | 0.95 | 0.989 | 0 |
| **IMP4** | 0 | 1.00748068973663 | 1 | 0.996 | 0 |
| **PLEKHB2** | 0 | 1.43744041390605 | 1 | 0.999 | 0 |
| **LYPD1** | 0 | -1.78069960730366 | 0.017 | 0.577 | 0 |
| **ZEB2** | 0 | -1.49973283353999 | 0.039 | 0.611 | 0 |
| **KIF5C** | 0 | -1.50100496109998 | 0.871 | 0.991 | 0 |
| **RND3** | 0 | -1.46114098289485 | 0.018 | 0.541 | 0 |
| **RPRM** | 0 | -2.166656554897 | 0.068 | 0.858 | 0 |
| **TANC1** | 0 | -0.918116507931698 | 0.387 | 0.769 | 0 |
| **BAZ2B** | 0 | -1.31736815444999 | 0.379 | 0.886 | 0 |
| **GCA** | 0 | -1.31538976928534 | 0.109 | 0.777 | 0 |
| **STK39** | 0 | -0.868649270650269 | 0.538 | 0.868 | 0 |
| **CERS6** | 0 | -1.22681072871606 | 0.13 | 0.736 | 0 |
| **FASTKD1** | 0 | 0.916311876643686 | 0.998 | 0.921 | 0 |
| **KLHL23** | 0 | -1.43497213520004 | 0.336 | 0.858 | 0 |
| **SP5** | 0 | -3.64411970578237 | 0.147 | 0.916 | 0 |
| **GAD1** | 0 | -2.47740874913949 | 0.022 | 0.877 | 0 |
| **ITGA6** | 0 | -1.26667298876779 | 0.821 | 0.976 | 0 |
| **PDK1** | 0 | -2.37736661608892 | 0.572 | 0.956 | 0 |
| **CDCA7** | 0 | -1.33741026936447 | 0.425 | 0.809 | 0 |
| **OSBPL6** | 0 | -0.987319353891681 | 0.074 | 0.586 | 0 |
| **UBE2E3** | 0 | -1.9281139204989 | 0.446 | 0.958 | 0 |
| **TFPI** | 0 | -1.32167668253916 | 0.36 | 0.718 | 0 |
| **MYO1B** | 0 | 1.08977965852478 | 0.995 | 0.912 | 0 |
| **NABP1** | 0 | 1.63082651046789 | 0.883 | 0.283 | 0 |
| **FZD7** | 0 | -3.10130133807198 | 0.077 | 0.74 | 0 |
| **ICA1L** | 0 | 1.46487208276473 | 0.772 | 0.093 | 0 |
| **PARD3B** | 0 | -0.928462290165728 | 0.105 | 0.498 | 0 |
| **METTL21A** | 0 | -1.0098467594338 | 0.196 | 0.747 | 0 |
| **CCNYL1** | 0 | -1.49612907153276 | 0.322 | 0.92 | 0 |
| **FZD5** | 0 | 1.15294854339823 | 0.999 | 0.924 | 0 |
| **IDH1** | 0 | 1.09614456516696 | 1 | 0.986 | 0 |

| | | | | | |
|---|---|---|---|---|---|
| **CPS1** | 0 | -1.10197768450605 | 0.639 | 0.956 | 0 |
| **BARD1** | 0 | 1.01469953837831 | 0.982 | 0.759 | 0 |
| **XRCC5** | 0 | -0.819908324678158 | 0.978 | 0.997 | 0 |
| **IGFBP2** | 0 | -0.971739610797188 | 0.998 | 1 | 0 |
| **IGFBP5** | 0 | -3.01819670152554 | 0.04 | 0.356 | 0 |
| **TMBIM1** | 0 | 2.21245648181579 | 0.995 | 0.386 | 0 |
| **ABCB6** | 0 | 1.17773138766307 | 0.869 | 0.458 | 0 |
| **PTPRN** | 0 | 0.833445209138692 | 0.523 | 0.118 | 0 |
| **OBSL1** | 0 | -2.36718880707134 | 0.101 | 0.929 | 0 |
| **SERPINE2** | 0 | -1.24486077460575 | 0.525 | 0.894 | 0 |
| **SLC19A3** | 0 | 0.841791246056175 | 0.632 | 0.282 | 0 |
| **FBXO36** | 0 | 1.03946416952506 | 0.996 | 0.848 | 0 |
| **B3GNT7** | 0 | -1.62213689408687 | 0.593 | 0.861 | 0 |
| **ALPP** | 0 | 1.10017666963097 | 0.471 | 0.04 | 0 |
| **ALPG** | 0 | 4.58637390594273 | 1 | 0.516 | 0 |
| **NGEF** | 0 | 1.34108498243585 | 0.704 | 0.11 | 0 |
| **INPP5D** | 0 | 1.21327245391399 | 0.927 | 0.518 | 0 |
| **DGKD** | 0 | -0.955016849867277 | 0.147 | 0.641 | 0 |
| **ARL4C** | 0 | -1.82125510929819 | 0.025 | 0.421 | 0 |
| **GBX2** | 0 | -1.05953341536865 | 0.142 | 0.622 | 0 |
| **SCLY** | 0 | -0.932773957954689 | 0.317 | 0.771 | 0 |
| **GPC1** | 0 | -0.805601045899363 | 0.79 | 0.931 | 0 |
| **LRRN1** | 0 | -3.6248015547671 | 0.344 | 0.955 | 0 |
| **ARL8B** | 0 | 0.829621638411948 | 0.998 | 0.944 | 0 |
| **EDEM1** | 0 | 1.66229040334682 | 0.992 | 0.761 | 0 |
| **LMCD1** | 0 | -0.855863577828511 | 0.062 | 0.432 | 0 |
| **TIMP4** | 0 | -1.23511789280123 | 0.141 | 0.653 | 0 |
| **NUP210** | 0 | -1.51003122543116 | 0.216 | 0.839 | 0 |
| **SLC6A6** | 0 | 0.862226648746258 | 0.998 | 0.91 | 0 |
| **SH3BP5** | 0 | -0.827037308130135 | 0.3 | 0.675 | 0 |
| **EOMES** | 0 | -1.84257157911209 | 0.016 | 0.433 | 0 |
| **ZNF620** | 0 | -0.955880250254098 | 0.288 | 0.77 | 0 |
| **CTNNB1** | 0 | -1.90203290172454 | 0.932 | 0.996 | 0 |
| **ZNF660** | 0 | -0.906230620474883 | 0.056 | 0.602 | 0 |

| | | | | | |
|---|---|---|---|---|---|
| **ZNF502** | 0 | 0.811851430730876 | 0.457 | 0.027 | 0 |
| **PTH1R** | 0 | 1.01025637647719 | 0.845 | 0.437 | 0 |
| **KLHL18** | 0 | 0.944013664102123 | 0.977 | 0.822 | 0 |
| **CCDC71** | 0 | -0.973530741831236 | 0.218 | 0.746 | 0 |
| **MAPKAPK3** | 0 | -0.883259674672824 | 0.835 | 0.947 | 0 |
| **TEX264** | 0 | 0.822369720335383 | 0.952 | 0.756 | 0 |
| **DUSP7** | 0 | 1.20032718663731 | 0.967 | 0.661 | 0 |
| **NT5DC2** | 0 | -1.07214033468086 | 0.998 | 1 | 0 |
| **WNT5A** | 0 | -1.61778804330626 | 0.015 | 0.697 | 0 |
| **IL17RD** | 0 | -1.02770259523002 | 0.305 | 0.738 | 0 |
| **PDE12** | 0 | 1.00375004111377 | 0.92 | 0.533 | 0 |
| **PTPRG** | 0 | -0.926977257245663 | 0.66 | 0.898 | 0 |
| **MAGI1** | 0 | -0.96034776948575 | 0.426 | 0.792 | 0 |
| **GPR27** | 0 | -0.81427993088286 | 0.445 | 0.773 | 0 |
| **VGLL3** | 0 | -0.985436723351815 | 0.011 | 0.48 | 0 |
| **DCBLD2** | 0 | -1.31214611091045 | 0.774 | 0.983 | 0 |
| **TBC1D23** | 0 | 0.894588176712906 | 1 | 0.978 | 0 |
| **TOMM70** | 0 | 0.851229730389114 | 1 | 0.993 | 0 |
| **TMEM45A** | 0 | -1.00235164757512 | 0.111 | 0.637 | 0 |
| **ALCAM** | 0 | -1.67974160338411 | 0.014 | 0.714 | 0 |
| **DPPA2** | 0 | 1.45374225691543 | 0.942 | 0.363 | 0 |
| **NECTIN3** | 0 | -0.9399874232731 | 0.6 | 0.882 | 0 |
| **PHLDB2** | 0 | -1.57790784438179 | 0.134 | 0.781 | 0 |
| **ATG3** | 0 | 1.84712125978803 | 1 | 0.996 | 0 |
| **HCLS1** | 0 | 0.983419108511524 | 0.49 | 0.088 | 0 |
| **KPNA1** | 0 | 0.933636809728068 | 0.999 | 0.985 | 0 |
| **CCDC14** | 0 | -1.05097593541663 | 0.766 | 0.957 | 0 |
| **HEG1** | 0 | -1.38983753906109 | 0.2 | 0.741 | 0 |
| **SLC12A8** | 0 | -1.66038898844302 | 0.068 | 0.709 | 0 |
| **OSBPL11** | 0 | 0.994380304593286 | 0.997 | 0.921 | 0 |
| **MGLL** | 0 | 1.3393024131444 | 0.632 | 0.08 | 0 |
| **PLXND1** | 0 | 2.22610516085022 | 1 | 0.748 | 0 |
| **TRH** | 0 | 1.55863988976678 | 0.757 | 0.101 | 0 |
| **NPHP3** | 0 | -0.844968490915563 | 0.248 | 0.693 | 0 |

| | | | | | |
|---|---|---|---|---|---|
| **RAB6B** | 0 | -1.53143642020277 | 0.231 | 0.809 | 0 |
| **AMOTL2** | 0 | -2.74061176713618 | 0.462 | 0.936 | 0 |
| **PPP2R3A** | 0 | -0.980864812960323 | 0.47 | 0.884 | 0 |
| **RBP1** | 0 | -2.15976643075166 | 0.511 | 0.929 | 0 |
| **SPSB4** | 0 | -2.16413970596422 | 0.123 | 0.869 | 0 |
| **PCOLCE2** | 0 | -1.39622866957624 | 0.772 | 0.925 | 0 |
| **U2SURP** | 0 | 0.809118498549947 | 1 | 1 | 0 |
| **CHST2** | 0 | 2.60259869306896 | 0.998 | 0.482 | 0 |
| **PLSCR4** | 0 | 1.83011366075857 | 0.87 | 0.334 | 0 |
| **WWTR1** | 0 | 1.05923616308807 | 0.987 | 0.871 | 0 |
| **SLC33A1** | 0 | 0.861568287881804 | 0.953 | 0.788 | 0 |
| **ZMAT3** | 0 | 1.42590617212748 | 0.982 | 0.718 | 0 |
| **ZNF639** | 0 | 0.939987193021915 | 0.998 | 0.947 | 0 |
| **GNB4** | 0 | -1.29430942946815 | 0.596 | 0.936 | 0 |
| **SOX2** | 0 | -1.47651950098093 | 0.88 | 0.942 | 0 |
| **YEATS2** | 0 | -0.857983989141918 | 0.785 | 0.97 | 0 |
| **EEF1AKMT4** | 0 | 0.892243163066424 | 0.967 | 0.78 | 0 |
| **MAGEF1** | 0 | -1.46989410332032 | 0.317 | 0.899 | 0 |
| **LIPH** | 0 | 2.06316852406623 | 0.812 | 0.09 | 0 |
| **SENP2** | 0 | 1.31938092740446 | 1 | 0.996 | 0 |
| **IGF2BP2** | 0 | -0.816537559981079 | 0.963 | 0.998 | 0 |
| **LPP** | 0 | -1.91066752890895 | 0.319 | 0.904 | 0 |
| **P3H2** | 0 | -0.917162575980711 | 0.008 | 0.39 | 0 |
| **MB21D2** | 0 | -0.946072047157577 | 0.102 | 0.646 | 0 |
| **HES1** | 0 | -2.064212187546 | 0.018 | 0.659 | 0 |
| **XXYLT1** | 0 | -1.00416299724564 | 0.527 | 0.896 | 0 |
| **DLG1** | 0 | -0.865806657495062 | 0.923 | 0.988 | 0 |
| **ZNF595** | 0 | 1.39702782407903 | 0.99 | 0.799 | 0 |
| **ZNF732** | 0 | 1.09535394566172 | 0.551 | 0.033 | 0 |
| **RNF212** | 0 | 1.24792004284031 | 0.575 | 0.025 | 0 |
| **NSD2** | 0 | -1.59436165737319 | 0.07 | 0.798 | 0 |
| **NAT8L** | 0 | -1.0667465214058 | 0.144 | 0.59 | 0 |
| **MXD4** | 0 | -0.887447173809247 | 0.951 | 0.975 | 0 |
| **TNIP2** | 0 | 0.844325039661824 | 0.994 | 0.932 | 0 |

| | | | | | |
|---|---|---|---|---|---|
| **MFSD10** | 0 | -0.999088813241996 | 0.605 | 0.92 | 0 |
| **ADRA2C** | 0 | -1.23480826332509 | 0.842 | 0.911 | 0 |
| **NSG1** | 0 | 0.870154915547259 | 0.501 | 0.124 | 0 |
| **S100P** | 0 | 1.58293992080674 | 0.651 | 0.085 | 0 |
| **BLOC1S4** | 0 | -0.852083150502875 | 0.537 | 0.874 | 0 |
| **GRPEL1** | 0 | 1.00790682183893 | 0.999 | 0.967 | 0 |
| **BOD1L1** | 0 | -1.1101353791456 | 0.88 | 0.973 | 0 |
| **LDB2** | 0 | -1.17710707650005 | 0.152 | 0.686 | 0 |
| **QDPR** | 0 | -1.21771972532571 | 0.89 | 0.994 | 0 |
| **ZCCHC4** | 0 | 0.820037019705249 | 0.829 | 0.547 | 0 |
| **RELL1** | 0 | 1.5604032953363 | 0.996 | 0.781 | 0 |
| **PGM2** | 0 | 1.48495491520093 | 0.997 | 0.845 | 0 |
| **KLF3** | 0 | 1.66055640698131 | 0.983 | 0.563 | 0 |
| **UCHL1** | 0 | -2.8580747291114 | 0.28 | 0.918 | 0 |
| **LIMCH1** | 0 | -3.13481531300808 | 0.12 | 0.878 | 0 |
| **SLC30A9** | 0 | -0.845306947858703 | 0.956 | 0.997 | 0 |
| **BEND4** | 0 | -1.29032943447314 | 0.629 | 0.907 | 0 |
| **SHISA3** | 0 | -1.93668065011295 | 0.045 | 0.553 | 0 |
| **NIPAL1** | 0 | 0.968701650730255 | 0.891 | 0.575 | 0 |
| **TEC** | 0 | 1.58090448615145 | 0.805 | 0.153 | 0 |
| **SLAIN2** | 0 | 1.32861301230624 | 0.998 | 0.914 | 0 |
| **OCIAD2** | 0 | -0.885988233469416 | 0.017 | 0.528 | 0 |
| **NMU** | 0 | -0.878115893635596 | 0.643 | 0.87 | 0 |
| **IGFBP7** | 0 | -2.15050723322768 | 0.03 | 0.693 | 0 |
| **G3BP2** | 0 | -0.970630168885736 | 1 | 1 | 0 |
| **USO1** | 0 | -1.4123478044025 | 0.977 | 0.996 | 0 |
| **PAQR3** | 0 | 0.908587814288766 | 0.993 | 0.896 | 0 |
| **MAPK10** | 0 | -2.18587601223446 | 0.006 | 0.747 | 0 |
| **HSD17B11** | 0 | 0.866062680379879 | 0.903 | 0.642 | 0 |
| **SPP1** | 0 | -2.49561108057945 | 0.233 | 0.808 | 0 |
| **PKD2** | 0 | -0.977816535687533 | 0.523 | 0.871 | 0 |
| **GRID2** | 0 | -1.35816581470034 | 0.065 | 0.641 | 0 |
| **RAP1GDS1** | 0 | -0.815764470790931 | 0.812 | 0.968 | 0 |
| **AIMP1** | 0 | 0.903541702528048 | 1 | 0.989 | 0 |

| | | | | | |
|---|---|---|---|---|---|
| **ELOVL6** | 0 | -1.13253197121249 | 0.94 | 0.987 | 0 |
| **LARP7** | 0 | 1.32186114257748 | 0.996 | 0.848 | 0 |
| **ANK2** | 0 | -1.13218496416801 | 0.482 | 0.875 | 0 |
| **CAMK2D** | 0 | -1.40625857307918 | 0.12 | 0.749 | 0 |
| **PRSS12** | 0 | 1.7527368949457 | 0.974 | 0.483 | 0 |
| **FGF2** | 0 | -1.40129445416905 | 0.872 | 0.979 | 0 |
| **LARP1B** | 0 | 1.36477820985734 | 0.996 | 0.805 | 0 |
| **PCDH18** | 0 | -2.26774562686609 | 0.479 | 0.894 | 0 |
| **SLC7A11** | 0 | 1.10825064518702 | 0.996 | 0.942 | 0 |
| **NOCT** | 0 | 1.40610065254048 | 0.999 | 0.884 | 0 |
| **ELMOD2** | 0 | -0.874658807988904 | 0.456 | 0.852 | 0 |
| **TTC29** | 0 | -0.884975887930884 | 0.004 | 0.344 | 0 |
| **SFRP2** | 0 | -2.18109702336719 | 0.023 | 0.688 | 0 |
| **LRAT** | 0 | -1.68000870212187 | 0.02 | 0.551 | 0 |
| **PDGFC** | 0 | -0.893918460265624 | 0.151 | 0.583 | 0 |
| **APELA** | 0 | -1.35009890769599 | 0.872 | 0.917 | 0 |
| **TRIM61** | 0 | 0.980865411465885 | 0.794 | 0.352 | 0 |
| **TRIM60** | 0 | 2.02273757985761 | 0.963 | 0.146 | 0 |
| **TRIM75P** | 0 | 1.48374755931021 | 0.862 | 0.15 | 0 |
| **PALLD** | 0 | -1.22485331508999 | 0.171 | 0.546 | 0 |
| **GALNT7** | 0 | -1.30596822214245 | 0.753 | 0.938 | 0 |
| **HPGD** | 0 | -3.5217562311774 | 0.858 | 0.95 | 0 |
| **FAT1** | 0 | -2.8215187249869 | 0.672 | 0.987 | 0 |
| **ZFP42** | 0 | 3.12587297097876 | 1 | 0.288 | 0 |
| **SLC12A7** | 0 | 1.29776002309723 | 0.883 | 0.469 | 0 |
| **IRX2** | 0 | -1.25670346130881 | 0.847 | 0.984 | 0 |
| **IRX1** | 0 | -2.14636881735035 | 0.124 | 0.792 | 0 |
| **TRIO** | 0 | -0.829912513735178 | 0.606 | 0.859 | 0 |
| **MYO10** | 0 | -1.75885606716687 | 0.737 | 0.976 | 0 |
| **PRLR** | 0 | -0.887577463749978 | 0.024 | 0.45 | 0 |
| **SLC1A3** | 0 | -1.07702503218543 | 0.393 | 0.721 | 0 |
| **DAB2** | 0 | 0.831191493964641 | 0.401 | 0.026 | 0 |
| **OXCT1** | 0 | -0.906141525755003 | 0.448 | 0.825 | 0 |
| **SELENOP** | 0 | -2.90270631660457 | 0.121 | 0.924 | 0 |

| | | | | | |
|---|---|---|---|---|---|
| **HMGCS1** | 0 | -1.52943919734039 | 0.991 | 0.981 | 0 |
| **FST** | 0 | -3.21933434399424 | 0.09 | 0.83 | 0 |
| **ZSWIM6** | 0 | -1.09226886406494 | 0.077 | 0.644 | 0 |
| **SHISAL2B** | 0 | -1.09842386750504 | 0.019 | 0.553 | 0 |
| **ERBIN** | 0 | 1.44560243375056 | 1 | 0.942 | 0 |
| **PIK3R1** | 0 | -0.869328604748232 | 0.093 | 0.519 | 0 |
| **AK6** | 0 | 0.903979054848311 | 0.996 | 0.894 | 0 |
| **MARVELD2** | 0 | 1.31912252568785 | 0.8 | 0.215 | 0 |
| **MAP1B** | 0 | -2.32635461894729 | 0.951 | 0.998 | 0 |
| **ENC1** | 0 | -1.65699904715207 | 0.04 | 0.419 | 0 |
| **F2RL1** | 0 | 1.17177581547345 | 1 | 0.974 | 0 |
| **AGGF1** | 0 | -1.07715886667192 | 0.48 | 0.914 | 0 |
| **LHFPL2** | 0 | -0.855278984804804 | 0.168 | 0.59 | 0 |
| **ARSB** | 0 | -1.18302151388724 | 0.239 | 0.782 | 0 |
| **DHFR** | 0 | -0.805241079879315 | 0.807 | 0.929 | 0 |
| **VCAN** | 0 | -2.2866978667145 | 0.865 | 0.99 | 0 |
| **COX7C** | 0 | -1.1413473791968 | 1 | 1 | 0 |
| **ADGRV1** | 0 | -0.879483463108734 | 0.174 | 0.659 | 0 |
| **FAM172A** | 0 | -0.911069695403984 | 0.4 | 0.819 | 0 |
| **PCSK1** | 0 | -2.30423069473779 | 0.231 | 0.77 | 0 |
| **CAST** | 0 | -1.58836193458213 | 0.049 | 0.767 | 0 |
| **LNPEP** | 0 | -0.828821106831013 | 0.133 | 0.609 | 0 |
| **PAM** | 0 | -1.4918767514732 | 0.084 | 0.735 | 0 |
| **EFNA5** | 0 | -2.12767278705088 | 0.077 | 0.831 | 0 |
| **NREP** | 0 | -1.43530265310711 | 0.957 | 0.996 | 0 |
| **MCC** | 0 | -1.58034356290349 | 0.069 | 0.749 | 0 |
| **SNX24** | 0 | 0.981137755622405 | 0.978 | 0.775 | 0 |
| **ZNF608** | 0 | -1.19831473219637 | 0.522 | 0.93 | 0 |
| **ADAMTS19** | 0 | -1.73379741911971 | 0.001 | 0.597 | 0 |
| **TCF7** | 0 | -1.6274006302532 | 0.82 | 0.954 | 0 |
| **JADE2** | 0 | 1.11064718001306 | 0.971 | 0.789 | 0 |
| **H2AFY** | 0 | -0.948174019979697 | 0.089 | 0.571 | 0 |
| **WNT8A** | 0 | -2.58909202883588 | 0.024 | 0.82 | 0 |
| **REEP2** | 0 | -1.00195615854978 | 0.117 | 0.585 | 0 |

| | | | | | |
|---|---|---|---|---|---|
| **EGR1** | 0 | -1.06532659061322 | 0.051 | 0.412 | 0 |
| **CXXC5** | 0 | -2.07714689955422 | 0.463 | 0.959 | 0 |
| **PCDHA4** | 0 | -0.863176040846879 | 0.02 | 0.538 | 0 |
| **PCDHA12** | 0 | -0.931021994103892 | 0.019 | 0.561 | 0 |
| **PCDHB2** | 0 | -1.71970480167545 | 0.163 | 0.857 | 0 |
| **TAF7** | 0 | -0.959164926959007 | 0.995 | 1 | 0 |
| **PCDHGB7** | 0 | -1.05160531402015 | 0.199 | 0.693 | 0 |
| **DIAPH1** | 0 | -0.837738627893817 | 0.998 | 1 | 0 |
| **PCDH1** | 0 | -1.65150104255147 | 0.903 | 0.98 | 0 |
| **DPYSL3** | 0 | -3.89424832521445 | 0.737 | 0.991 | 0 |
| **CDX1** | 0 | -1.55107186623641 | 0.188 | 0.545 | 0 |
| **SPARC** | 0 | 0.858295730728187 | 1 | 1 | 0 |
| **ADAM19** | 0 | -1.21650092998878 | 0.699 | 0.902 | 0 |
| **WWC1** | 0 | -0.845726681500564 | 0.21 | 0.644 | 0 |
| **FGF18** | 0 | -1.50382063594141 | 0.112 | 0.666 | 0 |
| **STC2** | 0 | 2.11659366450153 | 0.975 | 0.76 | 0 |
| **SFXN1** | 0 | -1.22500500140713 | 0.861 | 0.995 | 0 |
| **FGFR4** | 0 | 0.848398036173806 | 0.949 | 0.73 | 0 |
| **NSD1** | 0 | 1.24953842181531 | 1 | 0.975 | 0 |
| **DBN1** | 0 | -1.24757780083371 | 0.991 | 1 | 0 |
| **NHP2** | 0 | 0.906737881732981 | 1 | 1 | 0 |
| **ZNF354C** | 0 | 0.978376802369369 | 0.552 | 0.021 | 0 |
| **GFPT2** | 0 | -1.25549496263918 | 0.681 | 0.91 | 0 |
| **SERPINB1** | 0 | 1.39366136495064 | 0.769 | 0.098 | 0 |
| **SERPINB9** | 0 | -2.48903428213539 | 0.969 | 0.962 | 0 |
| **SERPINB6** | 0 | 2.02603392894681 | 1 | 0.739 | 0 |
| **NQO2** | 0 | 0.828420448494651 | 0.999 | 0.978 | 0 |
| **TUBB2A** | 0 | -1.39019719094195 | 0.956 | 0.981 | 0 |
| **TUBB2B** | 0 | -2.00964354656053 | 0.995 | 0.999 | 0 |
| **GCNT2** | 0 | -0.865863861689153 | 0.844 | 0.957 | 0 |
| **EDN1** | 0 | -2.26293845800339 | 0.007 | 0.673 | 0 |
| **JARID2** | 0 | -1.65278986064772 | 0.956 | 0.994 | 0 |
| **ATXN1** | 0 | 0.983868873084234 | 0.73 | 0.383 | 0 |
| **CAP2** | 0 | -1.05891943276016 | 0.355 | 0.77 | 0 |

| | | | | | |
|---|---|---|---|---|---|
| **E2F3** | 0 | -0.955599406678753 | 0.707 | 0.921 | 0 |
| **CARMIL1** | 0 | -1.18906622163515 | 0.097 | 0.688 | 0 |
| **TRIM38** | 0 | 0.947280408402624 | 0.873 | 0.553 | 0 |
| **HIST1H1A** | 0 | 3.81514250675902 | 0.997 | 0.343 | 0 |
| **HIST1H2BB** | 0 | 2.27501290723942 | 0.75 | 0.075 | 0 |
| **HIST1H2BH** | 0 | 0.963649711859517 | 0.992 | 0.87 | 0 |
| **ZFP57** | 0 | 1.41988826560656 | 0.846 | 0.183 | 0 |
| **TUBB** | 0 | -0.948797258193656 | 0.999 | 1 | 0 |
| **FLOT1** | 0 | -1.3692583692257 | 0.835 | 0.99 | 0 |
| **DDAH2** | 0 | -1.51893504053312 | 0.619 | 0.953 | 0 |
| **PBX2** | 0 | -1.0000226400676 | 0.956 | 0.99 | 0 |
| **SYNGAP1** | 0 | -0.896536189757415 | 0.207 | 0.588 | 0 |
| **BAK1** | 0 | 0.904666364090054 | 1 | 0.99 | 0 |
| **ILRUN** | 0 | 1.6002577345249 | 1 | 0.996 | 0 |
| **SNRPC** | 0 | 0.888809124019188 | 1 | 1 | 0 |
| **ANKS1A** | 0 | -0.858505565476058 | 0.732 | 0.961 | 0 |
| **PIM1** | 0 | -0.944148955117413 | 0.927 | 0.955 | 0 |
| **MDFI** | 0 | 1.33545478902075 | 0.976 | 0.61 | 0 |
| **CCND3** | 0 | 1.23017610371315 | 0.97 | 0.621 | 0 |
| **TRERF1** | 0 | -0.918298806274601 | 0.015 | 0.481 | 0 |
| **PTK7** | 0 | -1.26674410931727 | 0.88 | 0.978 | 0 |
| **POLH** | 0 | 1.31632079368141 | 0.943 | 0.557 | 0 |
| **VEGFA** | 0 | 1.21650876142061 | 0.864 | 0.443 | 0 |
| **SLC35B2** | 0 | -1.26056167207023 | 0.72 | 0.979 | 0 |
| **AARS2** | 0 | 1.24255911965845 | 0.999 | 0.924 | 0 |
| **MCM3** | 0 | -0.85417058346161 | 0.997 | 0.997 | 0 |
| **DPPA5** | 0 | 3.82467221975039 | 1 | 0.464 | 0 |
| **KHDC3L** | 0 | 2.87404380001326 | 0.993 | 0.312 | 0 |
| **DDX43** | 0 | 1.89637483936622 | 0.879 | 0.077 | 0 |
| **IRAK1BP1** | 0 | 0.875847905633655 | 0.432 | 0.027 | 0 |
| **TPBG** | 0 | -1.0799394732949 | 0.214 | 0.635 | 0 |
| **PNRC1** | 0 | -0.84522185674087 | 0.995 | 0.998 | 0 |
| **UBE2J1** | 0 | -1.10139989222903 | 0.568 | 0.879 | 0 |
| **EPHA7** | 0 | -0.891607380039271 | 0.84 | 0.952 | 0 |

| | | | | | |
|---|---|---|---|---|---|
| **MANEA** | 0 | -0.885441657564969 | 0.383 | 0.729 | 0 |
| **LIN28B** | 0 | 0.869026846811665 | 1 | 0.994 | 0 |
| **PRDM1** | 0 | 1.32621990263149 | 0.929 | 0.399 | 0 |
| **ATG5** | 0 | 1.41734036304226 | 1 | 0.956 | 0 |
| **CD24** | 0 | -2.88931061829932 | 0.675 | 0.923 | 0 |
| **SEC63** | 0 | 1.65045339308289 | 1 | 0.991 | 0 |
| **OSTM1** | 0 | -0.804045230029842 | 0.587 | 0.852 | 0 |
| **SNX3** | 0 | 0.822313150417253 | 0.994 | 0.895 | 0 |
| **SESN1** | 0 | 1.07207083979864 | 0.997 | 0.897 | 0 |
| **FIG4** | 0 | -0.840293334222041 | 0.167 | 0.648 | 0 |
| **WASF1** | 0 | -1.40353891030695 | 0.404 | 0.889 | 0 |
| **SLC16A10** | 0 | 1.65057997839508 | 0.982 | 0.499 | 0 |
| **FYN** | 0 | -0.890777445821479 | 0.876 | 0.947 | 0 |
| **MARCKS** | 0 | -3.26391970590231 | 0.376 | 0.971 | 0 |
| **TSPYL4** | 0 | -0.823488373204488 | 0.203 | 0.674 | 0 |
| **DSE** | 0 | 0.829681042401432 | 0.932 | 0.686 | 0 |
| **SLC35F1** | 0 | -1.29307487372724 | 0.036 | 0.678 | 0 |
| **CEP85L** | 0 | 0.932242159982222 | 0.72 | 0.399 | 0 |
| **MAN1A1** | 0 | -0.883171942484205 | 0.07 | 0.487 | 0 |
| **GJA1** | 0 | -1.45615755522888 | 1 | 1 | 0 |
| **PKIB** | 0 | 1.15763367792491 | 0.628 | 0.072 | 0 |
| **FABP7** | 0 | 0.976518941971893 | 0.832 | 0.573 | 0 |
| **SMPDL3A** | 0 | 1.2081206693442 | 0.805 | 0.25 | 0 |
| **ECHDC1** | 0 | -1.83182863530162 | 0.202 | 0.95 | 0 |
| **PTPRK** | 0 | -0.807294248642537 | 0.63 | 0.886 | 0 |
| **LAMA2** | 0 | -1.57105400203057 | 0.239 | 0.772 | 0 |
| **ENPP1** | 0 | -0.85593241760638 | 0.404 | 0.776 | 0 |
| **CCN2** | 0 | -1.81234427462285 | 0.311 | 0.757 | 0 |
| **ABRACL** | 0 | -1.34287591002403 | 0.955 | 0.999 | 0 |
| **CITED2** | 0 | -2.41139648481668 | 0.366 | 0.929 | 0 |
| **AIG1** | 0 | -1.53970747901885 | 0.104 | 0.781 | 0 |
| **PHACTR2** | 0 | -1.66188985686801 | 0.246 | 0.89 | 0 |
| **UTRN** | 0 | -1.53732081369713 | 0.47 | 0.896 | 0 |
| **UST** | 0 | -1.01232224269512 | 0.506 | 0.848 | 0 |

| | | | | | |
|---|---|---|---|---|---|
| **MTHFD1L** | 0 | 0.862290955386211 | 1 | 0.992 | 0 |
| **RMND1** | 0 | 1.51797573496616 | 1 | 0.96 | 0 |
| **EZR** | 0 | -1.19470004745089 | 1 | 1 | 0 |
| **TCP1** | 0 | 1.49896684719826 | 1 | 0.959 | 0 |
| **IGF2R** | 0 | -1.94860656421105 | 0.537 | 0.964 | 0 |
| **MAP3K4** | 0 | -0.942491561999495 | 0.93 | 0.988 | 0 |
| **TBXT** | 0 | -1.24324823278219 | 0.007 | 0.313 | 0 |
| **AFDN** | 0 | -0.962724657459748 | 0.983 | 1 | 0 |
| **DACT2** | 0 | 1.18581773244546 | 0.613 | 0.08 | 0 |
| **THBS2** | 0 | -1.39545209164761 | 0.022 | 0.53 | 0 |
| **C6orf120** | 0 | -1.39996075843974 | 0.486 | 0.948 | 0 |
| **PHF10** | 0 | -0.922934150446193 | 0.936 | 0.987 | 0 |
| **FAM120B** | 0 | -0.911537907756957 | 0.258 | 0.756 | 0 |
| **FAM20C** | 0 | -1.89801901775101 | 0.388 | 0.918 | 0 |
| **PRKAR1B** | 0 | -1.04813769058384 | 0.301 | 0.775 | 0 |
| **C7orf50** | 0 | -1.05181444241092 | 0.903 | 0.977 | 0 |
| **ZFAND2A** | 0 | 0.845908459522009 | 0.884 | 0.653 | 0 |
| **SNX8** | 0 | 0.962498319322992 | 0.999 | 0.972 | 0 |
| **AIMP2** | 0 | -1.18930256441228 | 0.938 | 0.996 | 0 |
| **EIF2AK1** | 0 | -0.857269616415544 | 0.983 | 1 | 0 |
| **TSPAN13** | 0 | 0.879912643808582 | 0.993 | 0.903 | 0 |
| **SP8** | 0 | -1.76513701304301 | 0.446 | 0.879 | 0 |
| **CDCA7L** | 0 | -0.928971176673598 | 0.763 | 0.937 | 0 |
| **IGF2BP3** | 0 | -0.892452634213638 | 0.999 | 1 | 0 |
| **CCDC126** | 0 | 1.21437907793568 | 0.799 | 0.341 | 0 |
| **MPP6** | 0 | -1.25767764510332 | 0.943 | 0.992 | 0 |
| **CYCS** | 0 | 0.83218936050024 | 1 | 0.828 | 0 |
| **CBX3** | 0 | 1.13162916262557 | 1 | 0.914 | 0 |
| **SKAP2** | 0 | -1.20735412483359 | 0.076 | 0.637 | 0 |
| **HOXA1** | 0 | -1.18513075202023 | 0.17 | 0.6 | 0 |
| **CPVL** | 0 | -1.24176059892291 | 0.951 | 0.997 | 0 |
| **DPY19L1** | 0 | 1.36317081168512 | 0.988 | 0.71 | 0 |
| **HERPUD2** | 0 | -1.02073146851467 | 0.395 | 0.839 | 0 |
| **STARD3NL** | 0 | -0.987201435101649 | 0.676 | 0.938 | 0 |

| | | | | | |
|---|---|---|---|---|---|
| **MPLKIP** | 0 | -0.924909369136718 | 0.761 | 0.965 | 0 |
| **DBNL** | 0 | -1.33304658092329 | 0.74 | 0.986 | 0 |
| **PURB** | 0 | -0.815122422406058 | 0.773 | 0.972 | 0 |
| **TBRG4** | 0 | 0.884944705134214 | 1 | 0.991 | 0 |
| **IGFBP3** | 0 | -2.25713118944111 | 0.055 | 0.564 | 0 |
| **SUN3** | 0 | 1.8541076486261 | 0.828 | 0.062 | 0 |
| **UPP1** | 0 | 1.07981423166197 | 1 | 0.927 | 0 |
| **PSPH** | 0 | 0.980280024921402 | 0.992 | 0.897 | 0 |
| **CHCHD2** | 0 | 1.48759141401969 | 1 | 0.998 | 0 |
| **ZNF727** | 0 | 0.854724232894267 | 0.454 | 0.022 | 0 |
| **ZNF736** | 0 | 1.19241043582666 | 0.662 | 0.043 | 0 |
| **ZNF680** | 0 | 1.14944049176631 | 0.983 | 0.804 | 0 |
| **AUTS2** | 0 | -1.35785731856741 | 0.633 | 0.903 | 0 |
| **GALNT17** | 0 | -1.71305352126688 | 0.306 | 0.879 | 0 |
| **LIMK1** | 0 | -0.877660811366695 | 0.855 | 0.971 | 0 |
| **GTF2I** | 0 | -0.901513968456877 | 0.947 | 0.997 | 0 |
| **HIP1** | 0 | -1.00399903924822 | 0.489 | 0.861 | 0 |
| **SSC4D** | 0 | -1.15499373324712 | 0.054 | 0.654 | 0 |
| **TMEM60** | 0 | -0.838459237111088 | 0.217 | 0.699 | 0 |
| **SEMA3A** | 0 | -1.26765805588201 | 0.099 | 0.681 | 0 |
| **CDK14** | 0 | -1.0625459105094 | 0.105 | 0.575 | 0 |
| **CDK6** | 0 | -2.32681614945269 | 0.141 | 0.919 | 0 |
| **COL1A2** | 0 | -3.50920106608442 | 0.377 | 0.906 | 0 |
| **PPP1R9A** | 0 | -0.875540056036137 | 0.059 | 0.554 | 0 |
| **PON2** | 0 | -1.00135751331676 | 0.672 | 0.936 | 0 |
| **SLC25A13** | 0 | -1.04236609456502 | 0.956 | 0.999 | 0 |
| **NPTX2** | 0 | 1.53901568124477 | 0.658 | 0.026 | 0 |
| **ARPC1B** | 0 | 0.903654377211947 | 1 | 0.927 | 0 |
| **BUD31** | 0 | 0.990473495173297 | 1 | 0.984 | 0 |
| **PTCD1** | 0 | 0.819094457881594 | 0.982 | 0.843 | 0 |
| **ZKSCAN1** | 0 | -0.8071429011001 | 0.38 | 0.765 | 0 |
| **STAG3** | 0 | 0.873546859244428 | 0.438 | 0.038 | 0 |
| **SAP25** | 0 | 0.826713692680459 | 0.524 | 0.141 | 0 |
| **TRIP6** | 0 | -1.09378674542454 | 0.989 | 0.998 | 0 |

| | | | | | |
|---|---|---|---|---|---|
| **MUC3A** | 0 | -1.34749666537801 | 0.033 | 0.511 | 0 |
| **AP1S1** | 0 | 0.983751467709499 | 0.993 | 0.866 | 0 |
| **COL26A1** | 0 | -2.23580378365132 | 0.263 | 0.898 | 0 |
| **CUX1** | 0 | -1.36115075046493 | 0.39 | 0.906 | 0 |
| **EFCAB10** | 0 | 0.879367620970677 | 0.53 | 0.083 | 0 |
| **CCDC71L** | 0 | -1.09169477216506 | 0.192 | 0.65 | 0 |
| **PRKAR2B** | 0 | -1.91695564251437 | 0.205 | 0.872 | 0 |
| **LAMB1** | 0 | -1.05827593598498 | 0.92 | 0.986 | 0 |
| **DNAJB9** | 0 | 1.21279245290481 | 0.949 | 0.793 | 0 |
| **LRRN3** | 0 | -1.52654395568269 | 0.003 | 0.651 | 0 |
| **DOCK4** | 0 | 1.05580461954207 | 0.7 | 0.288 | 0 |
| **TES** | 0 | 0.916573700743555 | 0.959 | 0.747 | 0 |
| **CAV2** | 0 | -0.855624228233344 | 0.048 | 0.438 | 0 |
| **CAV1** | 0 | -2.03835277606767 | 0.022 | 0.68 | 0 |
| **ST7** | 0 | -1.27827182456235 | 0.121 | 0.737 | 0 |
| **PTPRZ1** | 0 | -1.89399244308889 | 0.004 | 0.627 | 0 |
| **AASS** | 0 | 0.907371006070114 | 0.994 | 0.87 | 0 |
| **HYAL4** | 0 | 1.90900842478328 | 0.901 | 0.113 | 0 |
| **KLHDC10** | 0 | -0.983694876120739 | 0.21 | 0.749 | 0 |
| **MEST** | 0 | 0.931684601113031 | 0.999 | 0.889 | 0 |
| **PODXL** | 0 | -4.96003222645995 | 0.68 | 0.993 | 0 |
| **AKR1B1** | 0 | -0.850937425751773 | 0.915 | 0.983 | 0 |
| **CALD1** | 0 | -1.94771088304984 | 0.97 | 0.999 | 0 |
| **TRIM24** | 0 | -2.03677009721325 | 0.962 | 0.999 | 0 |
| **ZC3HAV1** | 0 | 1.42498650570492 | 1 | 0.959 | 0 |
| **HIPK2** | 0 | -1.30737859536604 | 0.342 | 0.845 | 0 |
| **SLC37A3** | 0 | 1.08009339835802 | 0.999 | 0.949 | 0 |
| **GSTK1** | 0 | -0.975577340765391 | 0.492 | 0.851 | 0 |
| **CASP2** | 0 | -1.05378216585878 | 0.66 | 0.924 | 0 |
| **TCAF1** | 0 | -1.707776480838 | 0.36 | 0.921 | 0 |
| **LRRC61** | 0 | -0.861817929138411 | 0.228 | 0.692 | 0 |
| **REPIN1** | 0 | -1.25177106886663 | 0.265 | 0.808 | 0 |
| **AGPAT5** | 0 | -1.10671538531294 | 0.747 | 0.957 | 0 |
| **PPP1R3B** | 0 | -2.07177499369045 | 0.516 | 0.967 | 0 |

| | | | | | |
|---|---|---|---|---|---|
| **XKR6** | 0 | -1.01572096025739 | 0.388 | 0.84 | 0 |
| **FAM167A** | 0 | -1.10097523415565 | 0.004 | 0.524 | 0 |
| **CTSB** | 0 | 1.083042904165 | 1 | 0.974 | 0 |
| **ZDHHC2** | 0 | -1.00904770122534 | 0.018 | 0.579 | 0 |
| **XPO7** | 0 | 0.914228186802856 | 1 | 0.982 | 0 |
| **SLC39A14** | 0 | 1.05877874570222 | 0.979 | 0.74 | 0 |
| **LOXL2** | 0 | -2.24350381053367 | 0.468 | 0.958 | 0 |
| **ENTPD4** | 0 | -0.878724811944621 | 0.371 | 0.774 | 0 |
| **STC1** | 0 | -1.04047318315724 | 0.002 | 0.479 | 0 |
| **NEFL** | 0 | 0.893085541521507 | 0.425 | 0.044 | 0 |
| **DOCK5** | 0 | -0.89285318355657 | 0.081 | 0.576 | 0 |
| **EBF2** | 0 | -1.47052010416501 | 0.013 | 0.593 | 0 |
| **BNIP3L** | 0 | -0.977246167206176 | 0.859 | 0.981 | 0 |
| **DPYSL2** | 0 | -2.8074304689319 | 0.5 | 0.982 | 0 |
| **CLU** | 0 | -2.739250118968 | 0.939 | 0.999 | 0 |
| **PBK** | 0 | 0.916810931542282 | 0.798 | 0.503 | 0 |
| **FZD3** | 0 | -1.09255736268391 | 0.824 | 0.976 | 0 |
| **GSR** | 0 | -1.53366317484477 | 0.609 | 0.97 | 0 |
| **ZNF703** | 0 | -1.28922290403673 | 0.389 | 0.791 | 0 |
| **EIF4EBP1** | 0 | 1.99525629427267 | 1 | 0.972 | 0 |
| **ASH2L** | 0 | 1.34272594696889 | 1 | 0.948 | 0 |
| **FGFR1** | 0 | -3.39225481089247 | 0.642 | 0.982 | 0 |
| **SLC20A2** | 0 | -0.804097009618327 | 0.178 | 0.642 | 0 |
| **TMEM68** | 0 | 0.832245606365826 | 0.98 | 0.879 | 0 |
| **TGS1** | 0 | 1.13548475843829 | 1 | 0.989 | 0 |
| **NSMAF** | 0 | 0.966197452326795 | 0.999 | 0.94 | 0 |
| **CHD7** | 0 | -2.1476671284891 | 0.458 | 0.936 | 0 |
| **MTFR1** | 0 | 1.41887685664205 | 0.998 | 0.875 | 0 |
| **PREX2** | 0 | -1.71499635746829 | 0.065 | 0.777 | 0 |
| **PRDM14** | 0 | 0.94977660605629 | 0.991 | 0.86 | 0 |
| **RDH10** | 0 | 0.826162991101264 | 0.68 | 0.347 | 0 |
| **STAU2** | 0 | -0.92124892389678 | 0.987 | 1 | 0 |
| **CRISPLD1** | 0 | -1.8020711844323 | 0.319 | 0.876 | 0 |
| **PKIA** | 0 | -1.0124443177716 | 0.005 | 0.553 | 0 |

| | | | | | |
|---|---|---|---|---|---|
| **ZC2HC1A** | 0 | -0.807948105213181 | 0.181 | 0.652 | 0 |
| **TPD52** | 0 | -0.895557509514143 | 0.913 | 0.977 | 0 |
| **FABP5** | 0 | 2.45185648780873 | 0.998 | 0.588 | 0 |
| **E2F5** | 0 | -1.28994454783337 | 0.577 | 0.938 | 0 |
| **TMEM64** | 0 | -2.13196900032345 | 0.859 | 0.983 | 0 |
| **UBR5** | 0 | -0.89595967957409 | 0.954 | 0.995 | 0 |
| **CTHRC1** | 0 | 0.931091955042919 | 0.542 | 0.078 | 0 |
| **ENPP2** | 0 | -0.86396607889529 | 0.167 | 0.543 | 0 |
| **ZHX2** | 0 | -1.70185565806067 | 0.011 | 0.78 | 0 |
| **MTSS1** | 0 | -0.886251530139343 | 0.271 | 0.691 | 0 |
| **SQLE** | 0 | -0.864942438868355 | 0.935 | 0.982 | 0 |
| **TRIB1** | 0 | -0.963935502574828 | 0.246 | 0.721 | 0 |
| **LRATD2** | 0 | -1.6048731087897 | 0.746 | 0.949 | 0 |
| **MYC** | 0 | -2.23758353856357 | 0.246 | 0.859 | 0 |
| **TOP1MT** | 0 | 1.04819763362665 | 0.998 | 0.934 | 0 |
| **NAPRT** | 0 | 0.924836299777288 | 0.888 | 0.541 | 0 |
| **NRBP2** | 0 | 1.19166370478102 | 0.848 | 0.489 | 0 |
| **EPPK1** | 0 | 1.06222273938859 | 1 | 0.975 | 0 |
| **KANK1** | 0 | -1.21476003560694 | 0.041 | 0.687 | 0 |
| **SMARCA2** | 0 | 0.931790436733428 | 0.999 | 0.924 | 0 |
| **RCL1** | 0 | 0.816819995872387 | 0.99 | 0.917 | 0 |
| **ERMP1** | 0 | -1.29293266463029 | 0.262 | 0.835 | 0 |
| **UHRF2** | 0 | -1.24440353672695 | 0.205 | 0.761 | 0 |
| **GLDC** | 0 | 0.86398534226578 | 0.982 | 0.791 | 0 |
| **CER1** | 0 | -2.13216709907796 | 0.048 | 0.418 | 0 |
| **PSIP1** | 0 | -0.991175473143372 | 0.93 | 0.988 | 0 |
| **FOCAD** | 0 | -0.842757563700521 | 0.647 | 0.904 | 0 |
| **MTAP** | 0 | -0.84997998264293 | 0.256 | 0.693 | 0 |
| **AQP3** | 0 | 2.26508703168581 | 0.901 | 0.161 | 0 |
| **UBE2R2** | 0 | -0.818528992372351 | 0.994 | 0.999 | 0 |
| **UBAP2** | 0 | 1.23810917468478 | 1 | 0.973 | 0 |
| **MYORG** | 0 | -1.16709705853861 | 0.038 | 0.632 | 0 |
| **DNAJB5** | 0 | -0.825338883386402 | 0.218 | 0.631 | 0 |
| **TLN1** | 0 | -0.958093101983146 | 0.794 | 0.951 | 0 |

| | | | | | |
|---|---|---|---|---|---|
| **GLIPR2** | 0 | -2.66875465153663 | 0.701 | 0.963 | 0 |
| **FBXO10** | 0 | -1.0801309794585 | 0.498 | 0.847 | 0 |
| **IGFBPL1** | 0 | -3.62011326110101 | 0.202 | 0.951 | 0 |
| **ANKRD18A** | 0 | -1.65939910039548 | 0.072 | 0.779 | 0 |
| **C9orf135** | 0 | -1.92647683626771 | 0.71 | 0.892 | 0 |
| **KLF9** | 0 | 0.830740277919037 | 0.68 | 0.34 | 0 |
| **CEMIP2** | 0 | -0.920764797929151 | 0.853 | 0.947 | 0 |
| **PCSK5** | 0 | 1.17511208549786 | 0.834 | 0.259 | 0 |
| **VPS13A** | 0 | -0.84391098384082 | 0.25 | 0.688 | 0 |
| **PSAT1** | 0 | 0.970238272351718 | 1 | 0.998 | 0 |
| **GKAP1** | 0 | 0.91216169490007 | 0.686 | 0.327 | 0 |
| **C9orf64** | 0 | 1.28240927370752 | 0.676 | 0.037 | 0 |
| **CKS2** | 0 | 0.960809371163187 | 0.993 | 0.928 | 0 |
| **AUH** | 0 | -0.808688439683407 | 0.691 | 0.924 | 0 |
| **ROR2** | 0 | -1.28398458696023 | 0.277 | 0.747 | 0 |
| **BARX1** | 0 | 0.973470274501395 | 0.551 | 0.053 | 0 |
| **PTCH1** | 0 | -1.60112266648211 | 0.431 | 0.87 | 0 |
| **ZNF367** | 0 | -1.04855703184413 | 0.176 | 0.576 | 0 |
| **CDC14B** | 0 | -0.93164915103604 | 0.039 | 0.539 | 0 |
| **CTSV** | 0 | -1.32142767705124 | 0.897 | 0.978 | 0 |
| **TGFBR1** | 0 | -1.25054171740044 | 0.931 | 0.999 | 0 |
| **ABCA1** | 0 | 1.58415199407481 | 1 | 0.923 | 0 |
| **KLF4** | 0 | 1.97055317723083 | 0.986 | 0.556 | 0 |
| **PALM2-AKAP** | 0 | -3.48808933111981 | 0.494 | 0.976 | 0 |
| **LPAR1** | 0 | -0.825696675641259 | 0.086 | 0.469 | 0 |
| **PTGR1** | 0 | 1.07786304866008 | 0.986 | 0.798 | 0 |
| **BSPRY** | 0 | -0.853109032322564 | 0.137 | 0.5 | 0 |
| **ALAD** | 0 | 0.967625114380546 | 0.975 | 0.754 | 0 |
| **ZNF618** | 0 | -0.999026093373595 | 0.346 | 0.745 | 0 |
| **TNC** | 0 | -1.25041696538187 | 0.011 | 0.443 | 0 |
| **BRINP1** | 0 | -1.25978568476799 | 0.11 | 0.601 | 0 |
| **CNTRL** | 0 | -1.08708996980152 | 0.126 | 0.671 | 0 |
| **GSN** | 0 | -2.52240942236265 | 0.041 | 0.851 | 0 |
| **PDCL** | 0 | 0.990403895587748 | 0.998 | 0.983 | 0 |

| | | | | | |
|---|---|---|---|---|---|
| **CDK9** | 0 | -0.886006831134308 | 0.92 | 0.994 | 0 |
| **ODF2** | 0 | 0.844724364057875 | 0.821 | 0.494 | 0 |
| **SPTAN1** | 0 | -0.904504701373578 | 0.987 | 0.998 | 0 |
| **SH3GLB2** | 0 | -0.918825712121878 | 0.51 | 0.867 | 0 |
| **NCS1** | 0 | -0.89033919804263 | 0.514 | 0.815 | 0 |
| **FIBCD1** | 0 | -1.70746539113375 | 0.312 | 0.803 | 0 |
| **AIF1L** | 0 | -1.30687255800994 | 0.964 | 0.996 | 0 |
| **MED22** | 0 | -0.973237944996909 | 0.752 | 0.967 | 0 |
| **CACFD1** | 0 | -1.10984629454101 | 0.39 | 0.856 | 0 |
| **VAV2** | 0 | 0.969030912767614 | 0.985 | 0.842 | 0 |
| **WDR5** | 0 | 0.930471574452457 | 0.989 | 0.86 | 0 |
| **OLFM1** | 0 | -1.65729739391415 | 0.147 | 0.742 | 0 |
| **PPP1R26** | 0 | -0.837551756736294 | 0.227 | 0.704 | 0 |
| **AGPAT2** | 0 | 0.884493188260984 | 0.992 | 0.873 | 0 |
| **NRARP** | 0 | -1.11745193604967 | 0.118 | 0.632 | 0 |
| **CACNA1B** | 0 | 0.856597653319563 | 0.661 | 0.262 | 0 |
| **PFKP** | 0 | -1.14743439345762 | 0.952 | 0.992 | 0 |
| **GDI2** | 0 | -1.23280764429717 | 0.954 | 0.997 | 0 |
| **CAMK1D** | 0 | 0.804010528425977 | 0.502 | 0.089 | 0 |
| **SEPHS1** | 0 | -1.41945717122678 | 0.985 | 0.997 | 0 |
| **FRMD4A** | 0 | -1.48005240491461 | 0.061 | 0.616 | 0 |
| **OLAH** | 0 | 1.27193490079883 | 0.782 | 0.183 | 0 |
| **FAM171A1** | 0 | -0.804507194695909 | 0.364 | 0.759 | 0 |
| **VIM** | 0 | -5.01279125628037 | 0.497 | 0.961 | 0 |
| **NEBL** | 0 | -1.28815928803307 | 0.037 | 0.599 | 0 |
| **MSRB2** | 0 | -1.28414476658843 | 0.503 | 0.923 | 0 |
| **KIAA1217** | 0 | 0.840450438602358 | 0.995 | 0.898 | 0 |
| **SVIL** | 0 | -1.23866178526447 | 0.339 | 0.814 | 0 |
| **ITGB1** | 0 | -1.29181713887752 | 0.926 | 0.994 | 0 |
| **RET** | 0 | -1.93304553012614 | 0.306 | 0.826 | 0 |
| **FXYD4** | 0 | 0.841475198346161 | 0.484 | 0.071 | 0 |
| **CXCL12** | 0 | -1.62227048543196 | 0.93 | 0.955 | 0 |
| **DEPP1** | 0 | -0.843417221696965 | 0.175 | 0.574 | 0 |
| **MARCH8** | 0 | 1.70947314565482 | 0.992 | 0.67 | 0 |

| | | | | | |
|---|---|---|---|---|---|
| **PRKG1** | 0 | -0.920128641768261 | 0.004 | 0.426 | 0 |
| **CSTF2T** | 0 | -0.941143747706563 | 0.849 | 0.988 | 0 |
| **CCDC6** | 0 | -1.02823915550201 | 0.919 | 0.991 | 0 |
| **NRBF2** | 0 | 1.23753526688679 | 0.999 | 0.96 | 0 |
| **JMJD1C** | 0 | -1.51879020294682 | 0.835 | 0.992 | 0 |
| **REEP3** | 0 | -1.08039762732377 | 0.684 | 0.945 | 0 |
| **CTNNA3** | 0 | -0.806001943513537 | 0.107 | 0.464 | 0 |
| **SLC25A16** | 0 | 1.69654070623251 | 1 | 0.94 | 0 |
| **STOX1** | 0 | -1.42729987927862 | 0.011 | 0.635 | 0 |
| **DDX21** | 0 | 0.822405706712643 | 1 | 0.998 | 0 |
| **HK1** | 0 | -1.27377112818028 | 0.897 | 0.99 | 0 |
| **TSPAN15** | 0 | -0.898870619450346 | 0.08 | 0.603 | 0 |
| **H2AFY2** | 0 | -1.11306663546156 | 0.758 | 0.909 | 0 |
| **PCBD1** | 0 | -1.03147102598063 | 0.639 | 0.939 | 0 |
| **PLAU** | 0 | 1.18502271633249 | 0.651 | 0.197 | 0 |
| **VCL** | 0 | -1.44556588105061 | 0.963 | 0.995 | 0 |
| **DLG5** | 0 | -0.816395494805672 | 0.85 | 0.97 | 0 |
| **ZCCHC24** | 0 | -1.52931402966355 | 0.065 | 0.731 | 0 |
| **PRXL2A** | 0 | -1.31363026830301 | 0.911 | 0.992 | 0 |
| **CDHR1** | 0 | 1.25130419477132 | 0.973 | 0.638 | 0 |
| **BMPR1A** | 0 | -1.50384721030519 | 0.513 | 0.923 | 0 |
| **GLUD1** | 0 | 0.994060383588157 | 1 | 0.995 | 0 |
| **SHLD2** | 0 | 0.995359803218561 | 0.972 | 0.745 | 0 |
| **LIPA** | 0 | -1.4255976676041 | 0.182 | 0.811 | 0 |
| **FGFBP3** | 0 | -1.37831199107809 | 0.004 | 0.483 | 0 |
| **EXOC6** | 0 | -1.2316587431222 | 0.121 | 0.745 | 0 |
| **LGI1** | 0 | -1.88855276049473 | 0.004 | 0.561 | 0 |
| **PDLIM1** | 0 | 1.09775944500451 | 1 | 0.929 | 0 |
| **PIK3AP1** | 0 | 1.10686364790155 | 0.588 | 0.068 | 0 |
| **MARVELD1** | 0 | -1.04048768973445 | 0.436 | 0.863 | 0 |
| **BLOC1S2** | 0 | 1.17715181173316 | 0.995 | 0.836 | 0 |
| **LZTS2** | 0 | -1.10027018825351 | 0.515 | 0.882 | 0 |
| **FGF8** | 0 | -2.03648674665654 | 0.35 | 0.791 | 0 |
| **NFKB2** | 0 | 1.680331046516 | 1 | 0.914 | 0 |

| | | | | | |
|---|---|---|---|---|---|
| **MFSD13A** | 0 | 1.01862458568537 | 0.679 | 0.235 | 0 |
| **WBP1L** | 0 | -0.946542224216148 | 0.731 | 0.939 | 0 |
| **NT5C2** | 0 | -0.9759007669099 | 0.908 | 0.985 | 0 |
| **INA** | 0 | 1.01151167106489 | 0.938 | 0.621 | 0 |
| **PCGF6** | 0 | 1.38207747170741 | 0.98 | 0.685 | 0 |
| **SH3PXD2A** | 0 | 0.827282157484865 | 0.919 | 0.67 | 0 |
| **1,00 STN** | 0 | 0.834139438039545 | 0.783 | 0.469 | 0 |
| **SLK** | 0 | 0.958246413021665 | 0.997 | 0.938 | 0 |
| **COL17A1** | 0 | 1.07035826801158 | 0.502 | 0.095 | 0 |
| **GSTO2** | 0 | 1.02858003858786 | 0.609 | 0.099 | 0 |
| **XPNPEP1** | 0 | -1.29630435018411 | 0.425 | 0.91 | 0 |
| **ADD3** | 0 | -1.54145772612657 | 0.767 | 0.983 | 0 |
| **MXI1** | 0 | -0.941090210515959 | 0.129 | 0.602 | 0 |
| **TCF7L2** | 0 | -1.68430537956117 | 0.266 | 0.88 | 0 |
| **SHTN1** | 0 | -1.32991673985281 | 0.249 | 0.833 | 0 |
| **RGS10** | 0 | 1.44123421752036 | 0.886 | 0.346 | 0 |
| **NSMCE4A** | 0 | -0.901818587797607 | 0.732 | 0.928 | 0 |
| **ACADSB** | 0 | -0.943336959903692 | 0.14 | 0.653 | 0 |
| **CHST15** | 0 | -0.893309569463661 | 0.741 | 0.912 | 0 |
| **NKX1-2** | 0 | -3.13361512374157 | 0.506 | 0.912 | 0 |
| **EBF3** | 0 | -1.33789910267534 | 0.179 | 0.616 | 0 |
| **UTF1** | 0 | 1.62783134141621 | 0.973 | 0.286 | 0 |
| **VENTX** | 0 | 0.978249745951521 | 0.717 | 0.214 | 0 |
| **PRAP1** | 0 | 1.21855070471501 | 0.694 | 0.137 | 0 |
| **FUOM** | 0 | 0.932940598073325 | 0.926 | 0.631 | 0 |
| **IFITM2** | 0 | -1.66711602310689 | 0.866 | 0.988 | 0 |
| **IFITM3** | 0 | -1.98540780438332 | 0.993 | 0.999 | 0 |
| **EPS8L2** | 0 | 1.13089755771407 | 0.955 | 0.646 | 0 |
| **TSPAN4** | 0 | -1.5737201285495 | 0.608 | 0.909 | 0 |
| **CDKN1C** | 0 | -1.19612455137231 | 0.511 | 0.845 | 0 |
| **CARS** | 0 | 0.896119516222277 | 0.865 | 0.56 | 0 |
| **TRIM21** | 0 | -0.837663909149697 | 0.015 | 0.539 | 0 |
| **TRIM6** | 0 | 0.884241175704286 | 0.651 | 0.209 | 0 |
| **TRIM5** | 0 | -0.938229964918729 | 0.364 | 0.807 | 0 |

| | | | | | |
|---|---|---|---|---|---|
| **CAVIN3** | 0 | -1.10563457358041 | 0.106 | 0.626 | 0 |
| **TPP1** | 0 | -1.16947187045247 | 0.949 | 0.993 | 0 |
| **RIC3** | 0 | -1.11184418246215 | 0.135 | 0.667 | 0 |
| **LMO1** | 0 | -1.14998063946064 | 0.341 | 0.686 | 0 |
| **ST5** | 0 | -1.55120014552898 | 0.571 | 0.95 | 0 |
| **ADM** | 0 | -1.64563477791401 | 0.233 | 0.692 | 0 |
| **3,00 DKK** | 0 | -1.74760501618781 | 0.093 | 0.774 | 0 |
| **PARVA** | 0 | -1.62046832371625 | 0.345 | 0.907 | 0 |
| **TEAD1** | 0 | -0.83967125129015 | 0.696 | 0.902 | 0 |
| **SPON1** | 0 | -1.07251899193388 | 0.001 | 0.293 | 0 |
| **PDE3B** | 0 | -0.869338909536234 | 0.084 | 0.541 | 0 |
| **GTF2H1** | 0 | 0.87949957805019 | 0.999 | 0.933 | 0 |
| **NAV2** | 0 | -2.55962494051902 | 0.68 | 0.966 | 0 |
| **FANCF** | 0 | -1.17792503071214 | 0.189 | 0.784 | 0 |
| **RCN1** | 0 | 1.10175269259315 | 1 | 0.973 | 0 |
| **CD44** | 0 | -0.871724768186145 | 0.11 | 0.47 | 0 |
| **FJX1** | 0 | -1.18140163187826 | 0.023 | 0.346 | 0 |
| **TRIM44** | 0 | -0.919373576644713 | 0.708 | 0.947 | 0 |
| **C11orf96** | 0 | -2.65186067022403 | 0.116 | 0.858 | 0 |
| **EXT2** | 0 | -0.954376689243684 | 0.722 | 0.956 | 0 |
| **TSPAN18** | 0 | -1.99413313946908 | 0.86 | 0.979 | 0 |
| **TP53I11** | 0 | -1.61765085738757 | 0.077 | 0.766 | 0 |
| **MDK** | 0 | -0.896045344681378 | 1 | 1 | 0 |
| **ACP2** | 0 | -1.15215836407991 | 0.389 | 0.866 | 0 |
| **TNKS1BP1** | 0 | -0.80013489851692 | 0.753 | 0.938 | 0 |
| **SERPING1** | 0 | -1.51090909171044 | 0.728 | 0.966 | 0 |
| **TMX2** | 0 | 1.08980917254836 | 1 | 0.999 | 0 |
| **SELENOH** | 0 | -1.0287573734783 | 0.999 | 1 | 0 |
| **FAM111B** | 0 | -1.87587230642617 | 0.127 | 0.685 | 0 |
| **TMEM132A** | 0 | -1.00040258777417 | 0.364 | 0.792 | 0 |
| **DDB1** | 0 | -0.863035025990811 | 1 | 1 | 0 |
| **FADS2** | 0 | -1.83843145047709 | 0.996 | 1 | 0 |
| **FADS1** | 0 | -1.3516856801693 | 0.959 | 0.996 | 0 |
| **RAB3IL1** | 0 | 1.01751580078571 | 0.721 | 0.315 | 0 |

| | | | | | |
|---|---|---|---|---|---|
| **ASRGL1** | 0 | 1.21243723764914 | 0.999 | 0.959 | 0 |
| **AHNAK** | 0 | -1.10793400298409 | 0.945 | 0.985 | 0 |
| **MTA2** | 0 | -0.877521917342337 | 0.977 | 0.997 | 0 |
| **PLAAT5** | 0 | -1.35867087625251 | 0.533 | 0.881 | 0 |
| **RTN3** | 0 | -0.997511204933476 | 0.997 | 0.997 | 0 |
| **RCOR2** | 0 | -1.19540238441338 | 0.299 | 0.749 | 0 |
| **MACROD1** | 0 | 0.93896155969912 | 0.741 | 0.364 | 0 |
| **VEGFB** | 0 | -1.12939229468224 | 0.636 | 0.93 | 0 |
| **PPP1R14B** | 0 | -0.842968531728002 | 0.945 | 0.993 | 0 |
| **TRMT112** | 0 | -1.55349907462072 | 0.825 | 0.996 | 0 |
| **NRXN2** | 0 | 1.55282175846653 | 0.801 | 0.099 | 0 |
| **RASGRP2** | 0 | 1.2264503554898 | 0.886 | 0.411 | 0 |
| **TM7SF2** | 0 | -1.99381032565504 | 0.593 | 0.966 | 0 |
| **CAPN1** | 0 | -1.62325128762477 | 0.924 | 0.998 | 0 |
| **CFL1** | 0 | -0.926676511974417 | 1 | 1 | 0 |
| **C11orf68** | 0 | -0.898460681627378 | 0.333 | 0.776 | 0 |
| **PACS1** | 0 | -1.42254088700761 | 0.618 | 0.955 | 0 |
| **CNIH2** | 0 | -0.958417123944723 | 0.319 | 0.678 | 0 |
| **C11orf80** | 0 | -1.03310189842175 | 0.216 | 0.742 | 0 |
| **SSH3** | 0 | -0.80945201344454 | 0.112 | 0.597 | 0 |
| **CORO1B** | 0 | -1.21882535602433 | 0.889 | 0.995 | 0 |
| **GAL** | 0 | 1.47547217139304 | 0.997 | 0.673 | 0 |
| **CPT1A** | 0 | 1.20328578819548 | 0.891 | 0.506 | 0 |
| **FGF4** | 0 | 1.03162389434117 | 0.989 | 0.637 | 0 |
| **RAB6A** | 0 | 1.15036480254124 | 1 | 0.988 | 0 |
| **TENM4** | 0 | -1.35032951490731 | 0.135 | 0.644 | 0 |
| **CTSC** | 0 | 0.836511347833079 | 1 | 0.994 | 0 |
| **AMOTL1** | 0 | -2.30393057391948 | 0.247 | 0.891 | 0 |
| **SRSF8** | 0 | 0.98302758665288 | 0.929 | 0.758 | 0 |
| **TRPC6** | 0 | -2.7595987778388 | 0.338 | 0.905 | 0 |
| **YAP1** | 0 | -1.8051807038534 | 0.937 | 0.998 | 0 |
| **TMEM123** | 0 | -1.0711590937065 | 0.248 | 0.692 | 0 |
| **EXPH5** | 0 | 1.31310614933519 | 0.765 | 0.182 | 0 |
| **PPP2R1B** | 0 | -1.40358353660392 | 0.813 | 0.988 | 0 |

| | | | | | |
|---|---|---|---|---|---|
| **DIXDC1** | 0 | -0.899377985392234 | 0.206 | 0.62 | 0 |
| **DLAT** | 0 | 0.894450239904885 | 1 | 0.991 | 0 |
| **NCAM1** | 0 | -0.981759869609695 | 0.01 | 0.47 | 0 |
| **CADM1** | 0 | -1.39429009072654 | 0.261 | 0.752 | 0 |
| **TAGLN** | 0 | -3.5366024125683 | 0.595 | 0.8 | 0 |
| **FXYD6** | 0 | 1.11603270344898 | 0.971 | 0.65 | 0 |
| **FOXR1** | 0 | 1.85362476070937 | 0.82 | 0.039 | 0 |
| **HYOU1** | 0 | 1.23181765968851 | 1 | 1 | 0 |
| **CBL** | 0 | -0.809832008524035 | 0.652 | 0.923 | 0 |
| **SC5D** | 0 | -1.1025657578973 | 0.945 | 0.989 | 0 |
| **SORL1** | 0 | -0.822257241188232 | 0.122 | 0.581 | 0 |
| **UBASH3B** | 0 | 1.49244279749795 | 0.808 | 0.205 | 0 |
| **GRAMD1B** | 0 | -1.56362360797289 | 0.025 | 0.608 | 0 |
| **CHEK1** | 0 | -0.815540897274078 | 0.92 | 0.971 | 0 |
| **HYLS1** | 0 | 1.1058360577155 | 0.632 | 0.149 | 0 |
| **CDON** | 0 | -0.902339588836913 | 0.1 | 0.515 | 0 |
| **FAM118B** | 0 | -0.976760888338106 | 0.617 | 0.938 | 0 |
| **TEAD4** | 0 | 1.35672504928085 | 1 | 0.97 | 0 |
| **TSPAN9** | 0 | -0.902351980740986 | 0.693 | 0.911 | 0 |
| **CCND2** | 0 | -1.56960910507007 | 0.168 | 0.621 | 0 |
| **AKAP3** | 0 | 2.01236309456324 | 0.856 | 0.041 | 0 |
| **NTF3** | 0 | -1.35661563073682 | 0.037 | 0.599 | 0 |
| **PLEKHG6** | 0 | -1.18464546445084 | 0.266 | 0.761 | 0 |
| **SCNN1A** | 0 | -1.05996298007851 | 0.61 | 0.844 | 0 |
| **PTMS** | 0 | -0.871770772423442 | 1 | 1 | 0 |
| **ENO2** | 0 | 1.27361093134769 | 0.999 | 0.954 | 0 |
| **LPCAT3** | 0 | 0.809971048849444 | 0.98 | 0.9 | 0 |
| **PEX5** | 0 | 0.947911396524118 | 0.969 | 0.766 | 0 |
| **DPPA3** | 0 | 3.42314617657974 | 0.997 | 0.261 | 0 |
| **CLEC4D** | 0 | -2.29430959348471 | 0.07 | 0.781 | 0 |
| **DUSP16** | 0 | 1.42788472434784 | 0.996 | 0.813 | 0 |
| **CREBL2** | 0 | 1.28353582711377 | 0.991 | 0.816 | 0 |
| **PLBD1** | 0 | 0.943412845644403 | 1 | 0.903 | 0 |
| **PLEKHA5** | 0 | -0.995918303370756 | 0.676 | 0.937 | 0 |

| | | | | | |
|---|---|---|---|---|---|
| **AEBP2** | 0 | 0.984642051601686 | 1 | 0.99 | 0 |
| **CMAS** | 0 | -0.871375076568066 | 0.652 | 0.915 | 0 |
| **PPFIBP1** | 0 | -0.970597782427342 | 0.325 | 0.762 | 0 |
| **CAPRIN2** | 0 | -1.01626375155157 | 0.437 | 0.772 | 0 |
| **RESF1** | 0 | 1.17055017393575 | 1 | 0.996 | 0 |
| **SLC2A13** | 0 | -0.83368382952448 | 0.487 | 0.78 | 0 |
| **YAF2** | 0 | -1.21587520689098 | 0.14 | 0.744 | 0 |
| **PRICKLE1** | 0 | -1.46101147843025 | 0.228 | 0.738 | 0 |
| **ARID2** | 0 | -0.898761580750988 | 0.792 | 0.967 | 0 |
| **HDAC7** | 0 | -1.07094252990389 | 0.578 | 0.886 | 0 |
| **COL2A1** | 0 | -1.67031589343038 | 0.003 | 0.306 | 0 |
| **PFKM** | 0 | -0.921188061265875 | 0.755 | 0.958 | 0 |
| **FKBP11** | 0 | 2.4069942351158 | 0.992 | 0.487 | 0 |
| **TUBA1A** | 0 | -1.15732727316894 | 0.673 | 0.902 | 0 |
| **TUBA1C** | 0 | 0.881737345520998 | 1 | 0.998 | 0 |
| **LIMA1** | 0 | 1.40907172919926 | 0.997 | 0.834 | 0 |
| **SMAGP** | 0 | -1.21265641233109 | 0.237 | 0.726 | 0 |
| **BIN2** | 0 | 0.868018123012151 | 0.482 | 0.027 | 0 |
| **FIGNL2** | 0 | -1.71291241632938 | 0.075 | 0.718 | 0 |
| **KRT8** | 0 | -2.85359709668369 | 0.94 | 0.999 | 0 |
| **CBX5** | 0 | -1.41487651069962 | 0.765 | 0.97 | 0 |
| **ZNF385A** | 0 | -1.08952007595919 | 0.07 | 0.599 | 0 |
| **ITGA5** | 0 | -1.41615710724715 | 0.053 | 0.606 | 0 |
| **PDE1B** | 0 | -1.13493632185408 | 0.01 | 0.499 | 0 |
| **PPP1R1A** | 0 | -2.55910128982823 | 0.18 | 0.9 | 0 |
| **PMEL** | 0 | -1.19008734966888 | 0.043 | 0.436 | 0 |
| **ESYT1** | 0 | -0.956032634177122 | 0.984 | 0.998 | 0 |
| **MYL6** | 0 | -1.26856023283701 | 1 | 1 | 0 |
| **SHMT2** | 0 | 1.07858863158328 | 0.979 | 0.736 | 0 |
| **MARS** | 0 | 0.875062011371014 | 0.97 | 0.786 | 0 |
| **DDIT3** | 0 | 1.62586592658713 | 0.846 | 0.549 | 0 |
| **KIF5A** | 0 | 0.905375950216824 | 0.618 | 0.235 | 0 |
| **XPOT** | 0 | 0.820903191917368 | 0.999 | 0.982 | 0 |
| **HMGA2** | 0 | -2.53065094927548 | 0.304 | 0.915 | 0 |

| | | | | | |
|---|---|---|---|---|---|
| **DYRK2** | 0 | -0.817784768776262 | 0.663 | 0.911 | 0 |
| **MDM2** | 0 | 1.62840716011617 | 1 | 0.982 | 0 |
| **MYRFL** | 0 | -1.35767788824491 | 0.046 | 0.609 | 0 |
| **ATXN7L3B** | 0 | -1.75582088888232 | 0.784 | 0.994 | 0 |
| **PHLDA1** | 0 | -2.48746457118228 | 0.344 | 0.891 | 0 |
| **PAWR** | 0 | -1.10987814286354 | 0.943 | 0.998 | 0 |
| **ACSS3** | 0 | -0.872770360049185 | 0.081 | 0.586 | 0 |
| **RASSF9** | 0 | -1.37361046676533 | 0.006 | 0.545 | 0 |
| **NTS** | 0 | -1.60161482791826 | 0.035 | 0.539 | 0 |
| **BTG1** | 0 | -0.873816136841527 | 0.685 | 0.922 | 0 |
| **NDUFA12** | 0 | 1.45299623852838 | 1 | 0.957 | 0 |
| **ACTR6** | 0 | 0.957535643964753 | 0.957 | 0.695 | 0 |
| **SPIC** | 0 | 1.73423045217232 | 0.852 | 0.157 | 0 |
| **CHPT1** | 0 | 0.896327544043672 | 0.996 | 0.896 | 0 |
| **SYCP3** | 0 | 2.15797382551072 | 0.857 | 0.111 | 0 |
| **DRAM1** | 0 | 1.47450392465948 | 0.944 | 0.475 | 0 |
| **TXNRD1** | 0 | 0.890500651448584 | 1 | 1 | 0 |
| **CHST11** | 0 | -1.73638274902942 | 0.473 | 0.895 | 0 |
| **ALDH1L2** | 0 | 1.01858928722719 | 0.633 | 0.129 | 0 |
| **C12orf75** | 0 | 1.27621946561302 | 0.986 | 0.796 | 0 |
| **NUAK1** | 0 | -1.01527791430611 | 0.434 | 0.831 | 0 |
| **CKAP4** | 0 | -1.18178017712698 | 0.937 | 0.993 | 0 |
| **RIC8B** | 0 | -0.862354410370955 | 0.462 | 0.804 | 0 |
| **MMAB** | 0 | -0.84280313251643 | 0.816 | 0.958 | 0 |
| **PPTC7** | 0 | 1.13198997674308 | 0.98 | 0.78 | 0 |
| **ALDH2** | 0 | 1.51807109553621 | 0.999 | 0.879 | 0 |
| **HSPB8** | 0 | -1.13398272386074 | 0.086 | 0.577 | 0 |
| **PXN** | 0 | -0.898687133631302 | 0.902 | 0.977 | 0 |
| **MSI1** | 0 | -1.71515416660445 | 0.064 | 0.812 | 0 |
| **UNC119B** | 0 | -0.98752761455389 | 0.366 | 0.788 | 0 |
| **ACADS** | 0 | 0.989688218782728 | 0.778 | 0.382 | 0 |
| **RHOF** | 0 | 0.996778028000402 | 0.757 | 0.279 | 0 |
| **CLIP1** | 0 | 0.821824440269603 | 0.993 | 0.901 | 0 |
| **CCDC92** | 0 | -1.11931792680242 | 0.063 | 0.531 | 0 |

| | | | | | |
|---|---|---|---|---|---|
| **ZNF664** | 0 | -0.845111897994403 | 0.727 | 0.923 | 0 |
| **RFLNA** | 0 | 1.0026547236189 | 0.509 | 0.101 | 0 |
| **NCOR2** | 0 | -0.834994441305999 | 0.999 | 1 | 0 |
| **FZD10** | 0 | -1.46537788515118 | 0.229 | 0.681 | 0 |
| **ULK1** | 0 | 1.01724564035485 | 1 | 0.989 | 0 |
| **PUS1** | 0 | 0.903743302423492 | 1 | 0.967 | 0 |
| **FBRSL1** | 0 | 1.15340483742256 | 0.993 | 0.841 | 0 |
| **CRYL1** | 0 | -0.9576650610907 | 0.013 | 0.503 | 0 |
| **SKA3** | 0 | 1.14053445593765 | 0.975 | 0.827 | 0 |
| **PARP4** | 0 | -1.19543289933655 | 0.62 | 0.935 | 0 |
| **FLT1** | 0 | -2.3562291998898 | 0.033 | 0.723 | 0 |
| **NBEA** | 0 | -1.98380020746874 | 0.029 | 0.805 | 0 |
| **DCLK1** | 0 | -2.7560233567359 | 0.125 | 0.854 | 0 |
| **UFM1** | 0 | 1.45387600858402 | 1 | 0.971 | 0 |
| **LHFPL6** | 0 | -1.8700155054792 | 0.035 | 0.735 | 0 |
| **FOXO1** | 0 | -2.68517453907974 | 0.456 | 0.966 | 0 |
| **RGCC** | 0 | 1.54786149713561 | 0.865 | 0.2 | 0 |
| **DNAJC15** | 0 | 2.0862010117349 | 0.991 | 0.99 | 0 |
| **LCP1** | 0 | 4.17903253305927 | 0.999 | 0.502 | 0 |
| **ITM2B** | 0 | -1.05686814634645 | 0.978 | 0.997 | 0 |
| **RB1** | 0 | -0.969008923469009 | 0.281 | 0.768 | 0 |
| **RCBTB2** | 0 | -1.13433832280963 | 0.158 | 0.717 | 0 |
| **RCBTB1** | 0 | 1.96006599440471 | 0.998 | 0.948 | 0 |
| **KLF5** | 0 | 1.81127086209491 | 1 | 0.73 | 0 |
| **KLF12** | 0 | -1.217804023687 | 0.11 | 0.723 | 0 |
| **TBC1D4** | 0 | -1.459041104508 | 0.088 | 0.727 | 0 |
| **SLAIN1** | 0 | -1.01220231546844 | 0.193 | 0.707 | 0 |
| **EDNRB** | 0 | -1.10177061965067 | 0.005 | 0.517 | 0 |
| **OBI1** | 0 | -1.18381201557527 | 0.681 | 0.96 | 0 |
| **SPRY2** | 0 | -3.01149382809656 | 0.413 | 0.962 | 0 |
| **GPC6** | 0 | -2.69565301954085 | 0.167 | 0.939 | 0 |
| **IPO5** | 0 | -1.95607398575246 | 0.849 | 1 | 0 |
| **FARP1** | 0 | -1.1008552283296 | 0.623 | 0.917 | 0 |
| **DOCK9** | 0 | 0.839256497735331 | 0.951 | 0.711 | 0 |

| | | | | | |
|---|---|---|---|---|---|
| **ZIC5** | 0 | -1.72244112785318 | 0.009 | 0.464 | 0 |
| **ZIC2** | 0 | -2.08501500820715 | 0.01 | 0.638 | 0 |
| **PCCA** | 0 | -1.11683515858542 | 0.97 | 0.998 | 0 |
| **EFNB2** | 0 | -2.0918687007098 | 0.305 | 0.88 | 0 |
| **ARGLU1** | 0 | -0.964678527883676 | 0.996 | 1 | 0 |
| **IRS2** | 0 | -0.933461054850295 | 0.176 | 0.621 | 0 |
| **COL4A1** | 0 | -2.64564204840653 | 0.445 | 0.969 | 0 |
| **COL4A2** | 0 | -2.53972130548413 | 0.555 | 0.971 | 0 |
| **LAMP1** | 0 | -0.94718355363977 | 1 | 1 | 0 |
| **TEP1** | 0 | -0.871666770324432 | 0.263 | 0.708 | 0 |
| **ARHGEF40** | 0 | -1.58967747735403 | 0.079 | 0.7 | 0 |
| **ZNF219** | 0 | -1.56690471662135 | 0.374 | 0.84 | 0 |
| **SALL2** | 0 | -2.57493034410704 | 0.598 | 0.984 | 0 |
| **SLC7A7** | 0 | 3.31851588766336 | 1 | 0.467 | 0 |
| **MMP14** | 0 | 0.913448520473065 | 0.999 | 0.958 | 0 |
| **AJUBA** | 0 | -0.975148631594802 | 0.745 | 0.893 | 0 |
| **EFS** | 0 | -1.32204477610646 | 0.238 | 0.643 | 0 |
| **PCK2** | 0 | 1.15984272473366 | 0.874 | 0.504 | 0 |
| **PSME1** | 0 | -0.942974685682101 | 0.826 | 0.97 | 0 |
| **REC8** | 0 | 2.29674443563559 | 0.996 | 0.307 | 0 |
| **NYNRIN** | 0 | -0.825396228570572 | 0.362 | 0.764 | 0 |
| **G2E3** | 0 | 0.843635208338041 | 0.972 | 0.848 | 0 |
| **EGLN3** | 0 | 2.3487333071439 | 0.995 | 0.598 | 0 |
| **SPTSSA** | 0 | 0.99331850918794 | 1 | 1 | 0 |
| **NFKBIA** | 0 | 1.48059771235747 | 1 | 0.967 | 0 |
| **BRMS1L** | 0 | 1.12668647660965 | 0.996 | 0.909 | 0 |
| **MIPOL1** | 0 | -1.06633551100446 | 0.256 | 0.795 | 0 |
| **GEMIN2** | 0 | 0.817198496323933 | 0.89 | 0.605 | 0 |
| **SAV1** | 0 | 1.42302355259992 | 1 | 0.969 | 0 |
| **PYGL** | 0 | 1.57250609202391 | 1 | 0.926 | 0 |
| **ABHD12B** | 0 | 0.905557615662797 | 0.555 | 0.097 | 0 |
| **FRMD6** | 0 | 0.938832537008379 | 0.931 | 0.678 | 0 |
| **TXNDC16** | 0 | -0.910957881683516 | 0.31 | 0.768 | 0 |
| **GNPNAT1** | 0 | 1.16361877596953 | 1 | 0.935 | 0 |

| | | | | | |
|---|---|---|---|---|---|
| **DDHD1** | 0 | 1.00176866467755 | 0.968 | 0.805 | 0 |
| **BMP4** | 0 | -1.47922229860059 | 0.432 | 0.876 | 0 |
| **SOCS4** | 0 | 0.954489591824219 | 1 | 0.989 | 0 |
| **LGALS3** | 0 | 0.923909098397416 | 0.541 | 0.154 | 0 |
| **DHRS7** | 0 | -0.857450706954804 | 0.74 | 0.949 | 0 |
| **HIF1A** | 0 | -1.23096248608186 | 0.717 | 0.965 | 0 |
| **PPP2R5E** | 0 | 0.80284873274417 | 1 | 0.994 | 0 |
| **HSPA2** | 0 | 2.14984416163718 | 0.935 | 0.13 | 0 |
| **PLEKHG3** | 0 | -1.4799763015531 | 0.058 | 0.57 | 0 |
| **SPTB** | 0 | -1.89025933068957 | 0.02 | 0.74 | 0 |
| **GPX2** | 0 | 2.12403351427976 | 0.941 | 0.257 | 0 |
| **ZFP36L1** | 0 | -0.98979252989321 | 0.985 | 0.997 | 0 |
| **ACTN1** | 0 | -1.55031187876768 | 0.993 | 1 | 0 |
| **DCAF5** | 0 | -1.51404493754872 | 0.24 | 0.873 | 0 |
| **EXD2** | 0 | -1.32647737679622 | 0.168 | 0.817 | 0 |
| **SRSF5** | 0 | -0.805766173056676 | 0.902 | 0.968 | 0 |
| **VRTN** | 0 | -3.26815266466381 | 0.198 | 0.948 | 0 |
| **NPC2** | 0 | -1.33856642009607 | 1 | 1 | 0 |
| **FOS** | 0 | -2.22686249980536 | 0.19 | 0.762 | 0 |
| **FLVCR2** | 0 | 1.48404710887262 | 0.854 | 0.449 | 0 |
| **IRF2BPL** | 0 | -1.03613927727576 | 0.823 | 0.924 | 0 |
| **CIPC** | 0 | -0.800875321007349 | 0.299 | 0.745 | 0 |
| **SLIRP** | 0 | 0.851508329584098 | 1 | 0.999 | 0 |
| **GALC** | 0 | -0.849208186436952 | 0.304 | 0.723 | 0 |
| **PTPN21** | 0 | -0.800306632061914 | 0.15 | 0.582 | 0 |
| **PPP4R3A** | 0 | 0.819944730898432 | 1 | 0.999 | 0 |
| **MOAP1** | 0 | -0.828741853113802 | 0.192 | 0.644 | 0 |
| **PRIMA1** | 0 | -1.28526169762706 | 0.333 | 0.74 | 0 |
| **DICER1** | 0 | -0.889413142963112 | 0.948 | 0.995 | 0 |
| **SYNE3** | 0 | -2.21082431902895 | 0.053 | 0.634 | 0 |
| **TCL1B** | 0 | 1.10488130139626 | 0.811 | 0.358 | 0 |
| **TUNAR** | 0 | -1.53682770740326 | 0.158 | 0.729 | 0 |
| **C14orf132** | 0 | -1.07865429437066 | 0.01 | 0.446 | 0 |
| **CCDC85C** | 0 | -1.66109282179488 | 0.923 | 0.983 | 0 |

| | | | | | |
|---|---|---|---|---|---|
| **HHIPL1** | 0 | -1.25520509259579 | 0.128 | 0.693 | 0 |
| **EML1** | 0 | -1.87975966289057 | 0.507 | 0.948 | 0 |
| **EVL** | 0 | -0.839006637938866 | 0.98 | 0.998 | 0 |
| **WARS** | 0 | 1.83425439897684 | 1 | 0.978 | 0 |
| **CKB** | 0 | -1.20000661578311 | 1 | 0.999 | 0 |
| **PPP1R13B** | 0 | -0.939960820419844 | 0.199 | 0.689 | 0 |
| **KIF26A** | 0 | -1.32165238963558 | 0.772 | 0.956 | 0 |
| **JAG2** | 0 | 1.03348074613065 | 0.74 | 0.286 | 0 |
| **PACS2** | 0 | -0.953245795662369 | 0.434 | 0.856 | 0 |
| **MTA1** | 0 | -1.0232432609327 | 0.973 | 0.998 | 0 |
| **CRIP1** | 0 | 3.02298757780291 | 1 | 0.557 | 0 |
| **GABRB3** | 0 | -1.35488097057099 | 0.88 | 0.971 | 0 |
| **TJP1** | 0 | -1.28472497185664 | 0.997 | 1 | 0 |
| **LPCAT4** | 0 | 1.53901900189029 | 0.996 | 0.838 | 0 |
| **SPRED1** | 0 | -1.25286917941262 | 0.075 | 0.74 | 0 |
| **FAM98B** | 0 | -0.939867820188502 | 0.97 | 0.998 | 0 |
| **EIF2AK4** | 0 | 0.96054919038054 | 0.971 | 0.764 | 0 |
| **CHAC1** | 0 | 1.33461737941403 | 0.767 | 0.412 | 0 |
| **LCMT2** | 0 | -0.935580462933132 | 0.238 | 0.742 | 0 |
| **SERF2** | 0 | -1.21831242406356 | 0.989 | 1 | 0 |
| **SORD** | 0 | -0.826730109430125 | 0.341 | 0.73 | 0 |
| **MYEF2** | 0 | -2.34025583477364 | 0.143 | 0.929 | 0 |
| **EID1** | 0 | -1.93262528052621 | 0.899 | 0.997 | 0 |
| **SCG3** | 0 | -1.28031123840673 | 0.035 | 0.666 | 0 |
| **MYO5C** | 0 | -1.31961385217903 | 0.169 | 0.791 | 0 |
| **MYO5A** | 0 | 1.26558319995195 | 0.992 | 0.782 | 0 |
| **ARPP19** | 0 | -0.939050405616031 | 0.98 | 0.997 | 0 |
| **RSL24D1** | 0 | 0.945094813785141 | 0.982 | 0.657 | 0 |
| **CCPG1** | 0 | 1.35992519200945 | 0.976 | 0.737 | 0 |
| **PRTG** | 0 | -2.13037220557054 | 0.039 | 0.508 | 0 |
| **MINDY2** | 0 | 0.802687227057161 | 0.994 | 0.887 | 0 |
| **GTF2A2** | 0 | 1.10117315986408 | 1 | 0.954 | 0 |
| **FOXB1** | 0 | -1.28071274250617 | 0.025 | 0.552 | 0 |
| **ANXA2** | 0 | 0.877240165412861 | 1 | 1 | 0 |

| | | | | | |
|---|---|---|---|---|---|
| **VPS13C** | 0 | -0.914571790781732 | 0.343 | 0.779 | 0 |
| **TPM1** | 0 | -1.2919284703538 | 1 | 1 | 0 |
| **SNX1** | 0 | -1.42451344781356 | 0.541 | 0.938 | 0 |
| **PCLAF** | 0 | -2.36732750616661 | 0.011 | 0.781 | 0 |
| **ZNF609** | 0 | -1.18864986592358 | 0.329 | 0.866 | 0 |
| **OAZ2** | 0 | -2.31614164989514 | 0.917 | 0.994 | 0 |
| **IGDCC3** | 0 | -2.37858696131623 | 0.098 | 0.899 | 0 |
| **IGDCC4** | 0 | -1.27589592710598 | 0.01 | 0.657 | 0 |
| **SMAD3** | 0 | -1.37365523407066 | 0.683 | 0.959 | 0 |
| **IQCH** | 0 | -0.895944691751703 | 0.112 | 0.523 | 0 |
| **UACA** | 0 | 1.96401439767713 | 0.999 | 0.83 | 0 |
| **MYO9A** | 0 | 0.811266629666406 | 0.96 | 0.745 | 0 |
| **CD276** | 0 | -0.987025535563928 | 0.927 | 0.978 | 0 |
| **LOXL1** | 0 | -1.31698751249539 | 0.376 | 0.735 | 0 |
| **STRA6** | 0 | -0.888131896110612 | 0.088 | 0.547 | 0 |
| **ARID3B** | 0 | -1.07359151051308 | 0.987 | 1 | 0 |
| **CSK** | 0 | -1.11517509914615 | 0.818 | 0.965 | 0 |
| **PTPN9** | 0 | -1.1537581137009 | 0.582 | 0.939 | 0 |
| **ETFA** | 0 | -1.34486180183864 | 0.805 | 0.977 | 0 |
| **PEAK1** | 0 | -2.09674136782532 | 0.178 | 0.868 | 0 |
| **DNAJA4** | 0 | 0.816845809685267 | 0.502 | 0.056 | 0 |
| **CRABP1** | 0 | -2.72947646459172 | 0.303 | 0.704 | 0 |
| **TLNRD1** | 0 | -1.1326489738538 | 0.618 | 0.928 | 0 |
| **MEX3B** | 0 | -0.824289002532477 | 0.142 | 0.525 | 0 |
| **EFL1** | 0 | -0.925880445574209 | 0.225 | 0.739 | 0 |
| **HDGFL3** | 0 | -0.907438354166596 | 0.913 | 0.991 | 0 |
| **SEC11A** | 0 | -0.931583070563962 | 0.999 | 1 | 0 |
| **MFGE8** | 0 | -1.69474096390787 | 0.341 | 0.908 | 0 |
| **IDH2** | 0 | -1.15897735609956 | 0.899 | 0.968 | 0 |
| **IQGAP1** | 0 | -0.966724208882513 | 0.589 | 0.897 | 0 |
| **CRTC3** | 0 | -1.2235534108778 | 0.128 | 0.739 | 0 |
| **FAM174B** | 0 | -1.83810198411636 | 0.222 | 0.838 | 0 |
| **RGMA** | 0 | -1.79316415855933 | 0.376 | 0.817 | 0 |
| **IGF1R** | 0 | -2.45324322575869 | 0.783 | 0.992 | 0 |

| | | | | | |
|---|---|---|---|---|---|
| **CHSY1** | 0 | -1.70217163314602 | 0.873 | 0.992 | 0 |
| **NME4** | 0 | -1.03642963459538 | 0.994 | 0.999 | 0 |
| **MEIOB** | 0 | 1.3661868227877 | 0.643 | 0.025 | 0 |
| **NDUFB10** | 0 | 0.910138594590516 | 1 | 1 | 0 |
| **SLC9A3R2** | 0 | 1.83964197616297 | 0.992 | 0.49 | 0 |
| **IL32** | 0 | 1.77363809776804 | 0.894 | 0.203 | 0 |
| **NMRAL1** | 0 | 0.894181385191632 | 0.904 | 0.481 | 0 |
| **PPL** | 0 | -0.809542158700977 | 0.132 | 0.489 | 0 |
| **PMM2** | 0 | 0.872210782745911 | 1 | 0.983 | 0 |
| **C16orf72** | 0 | 0.869753927551622 | 1 | 0.997 | 0 |
| **LITAF** | 0 | -2.26652306969118 | 0.988 | 0.999 | 0 |
| **SNN** | 0 | -1.09528068062435 | 0.221 | 0.688 | 0 |
| **TMC5** | 0 | -1.81635012868664 | 0.228 | 0.821 | 0 |
| **GPRC5B** | 0 | -1.22096462563088 | 0.992 | 0.99 | 0 |
| **CDR2** | 0 | 0.916344493359884 | 0.957 | 0.742 | 0 |
| **NDUFAB1** | 0 | 1.39753520879441 | 1 | 1 | 0 |
| **NUPR1** | 0 | 3.05362685812859 | 0.985 | 0.256 | 0 |
| **QPRT** | 0 | -1.22100177769574 | 0.992 | 1 | 0 |
| **CORO1A** | 0 | 0.928496840210836 | 0.544 | 0.121 | 0 |
| **PRR14** | 0 | 0.876363856205954 | 0.819 | 0.475 | 0 |
| **FBRS** | 0 | 1.36676951619039 | 0.969 | 0.7 | 0 |
| **PRSS8** | 0 | -1.42018688787297 | 0.758 | 0.935 | 0 |
| **SHCBP1** | 0 | 1.01431022081019 | 0.969 | 0.737 | 0 |
| **GPT2** | 0 | 1.57135685546875 | 0.996 | 0.86 | 0 |
| **DNAJA2** | 0 | 0.892903956419648 | 1 | 1 | 0 |
| **ZNF423** | 0 | -2.4825376040548 | 0.432 | 0.956 | 0 |
| **BRD7** | 0 | -1.00001025622317 | 0.956 | 0.998 | 0 |
| **SALL1** | 0 | -2.0802702315308 | 0.028 | 0.757 | 0 |
| **TOX3** | 0 | -1.40374685347395 | 0.001 | 0.659 | 0 |
| **AC007906.2** | 0 | -1.08836575705996 | 0.01 | 0.555 | 0 |
| **CHD9** | 0 | -0.979892885201206 | 0.537 | 0.895 | 0 |
| **MMP2** | 0 | -1.1141473418909 | 0.923 | 0.958 | 0 |
| **MT2A** | 0 | -1.13303033588337 | 0.988 | 0.983 | 0 |
| **MT1E** | 0 | 1.41621729159877 | 0.913 | 0.19 | 0 |

| | | | | | |
|---|---|---|---|---|---|
| **MT1F** | 0 | -0.933047696252161 | 0.81 | 0.92 | 0 |
| **MT1G** | 0 | 2.58026850436979 | 0.994 | 0.276 | 0 |
| **MT1H** | 0 | 2.49932544315859 | 0.992 | 0.318 | 0 |
| **MT1X** | 0 | 1.29164194341443 | 0.999 | 0.962 | 0 |
| **HERPUD1** | 0 | 2.18251897578486 | 0.972 | 0.654 | 0 |
| **CPNE2** | 0 | -0.823306768460924 | 0.206 | 0.568 | 0 |
| **CCDC102A** | 0 | -1.12807341330949 | 0.28 | 0.806 | 0 |
| **KIFC3** | 0 | -1.01348412164677 | 0.201 | 0.687 | 0 |
| **USB1** | 0 | -1.14355335196041 | 0.633 | 0.945 | 0 |
| **MMP15** | 0 | -2.83601085220453 | 0.565 | 0.989 | 0 |
| **RRAD** | 0 | 2.40137854664178 | 0.942 | 0.176 | 0 |
| **C16orf70** | 0 | 0.82936294986385 | 0.99 | 0.896 | 0 |
| **PLEKHG4** | 0 | 0.809487015795259 | 0.487 | 0.05 | 0 |
| **NRN1L** | 0 | -0.925344836561793 | 0.022 | 0.397 | 0 |
| **PSKH1** | 0 | -1.02372963008425 | 0.633 | 0.942 | 0 |
| **PLA2G15** | 0 | 0.802585585254104 | 0.825 | 0.594 | 0 |
| **SLC7A6** | 0 | 1.75988490706971 | 0.985 | 0.65 | 0 |
| **HAS3** | 0 | -0.914401746781046 | 0.019 | 0.456 | 0 |
| **AARS** | 0 | 0.934713136949205 | 0.998 | 0.925 | 0 |
| **IL34** | 0 | 0.865612820546067 | 0.634 | 0.236 | 0 |
| **LDHD** | 0 | -0.949665131660339 | 0.16 | 0.573 | 0 |
| **CHST6** | 0 | -0.936486926688948 | 0.198 | 0.669 | 0 |
| **VAT1L** | 0 | -2.88326531407174 | 0.015 | 0.703 | 0 |
| **WWOX** | 0 | -1.59884670735262 | 0.675 | 0.946 | 0 |
| **MAF** | 0 | -2.46806248542177 | 0.037 | 0.787 | 0 |
| **GINS2** | 0 | -0.906808602289924 | 0.822 | 0.906 | 0 |
| **FANCA** | 0 | 1.01183307702863 | 0.976 | 0.766 | 0 |
| **DBNDD1** | 0 | -1.85466742759132 | 0.401 | 0.934 | 0 |
| **RFLNB** | 0 | 2.08535456095976 | 0.988 | 0.517 | 0 |
| **NXN** | 0 | -1.30796513693567 | 0.96 | 0.998 | 0 |
| **PITPNA** | 0 | -0.878231361358748 | 0.995 | 0.999 | 0 |
| **SLC43A2** | 0 | -1.03833770498415 | 0.613 | 0.893 | 0 |
| **RTN4RL1** | 0 | -1.07981115850888 | 0.284 | 0.75 | 0 |
| **SMG6** | 0 | -0.914244204815151 | 0.445 | 0.851 | 0 |

| | | | | | |
|---|---|---|---|---|---|
| **SGSM2** | 0 | -0.978031934301788 | 0.154 | 0.677 | 0 |
| **METTL16** | 0 | 0.825610551624461 | 0.997 | 0.948 | 0 |
| **WSCD1** | 0 | -0.805396495856414 | 0.123 | 0.57 | 0 |
| **SAT2** | 0 | -1.36407701772176 | 0.854 | 0.979 | 0 |
| **ATP1B2** | 0 | -1.99801681571931 | 0.169 | 0.815 | 0 |
| **TP53** | 0 | -1.63143686174378 | 0.939 | 0.997 | 0 |
| **TRAPPC1** | 0 | -0.85883442065777 | 0.938 | 0.989 | 0 |
| **NTN1** | 0 | 7.00534141657374 | 1 | 0.997 | 0 |
| **PLD6** | 0 | 1.05819325358249 | 0.505 | 0.046 | 0 |
| **RASD1** | 0 | 1.26586692680116 | 0.614 | 0.149 | 0 |
| **PEMT** | 0 | 0.998431814136433 | 0.996 | 0.955 | 0 |
| **NATD1** | 0 | -0.857113997123132 | 0.324 | 0.693 | 0 |
| **RAB34** | 0 | -0.891963181778754 | 0.908 | 0.98 | 0 |
| **FLOT2** | 0 | -1.10588858027165 | 0.909 | 0.991 | 0 |
| **PIPOX** | 0 | -1.01199064511924 | 0.171 | 0.654 | 0 |
| **RAB11FIP4** | 0 | -0.886590016421243 | 0.934 | 0.956 | 0 |
| **RHBDL3** | 0 | -1.2843268531553 | 0.016 | 0.606 | 0 |
| **TMEM98** | 0 | -1.18189426280648 | 0.258 | 0.76 | 0 |
| **RASL10B** | 0 | -0.929053566215752 | 0.051 | 0.479 | 0 |
| **MLLT6** | 0 | -1.02648285920688 | 0.393 | 0.784 | 0 |
| **ERBB2** | 0 | -0.970327847075668 | 0.774 | 0.94 | 0 |
| **MSL1** | 0 | -0.839646846461113 | 0.596 | 0.895 | 0 |
| **KRT19** | 0 | -1.16787207611754 | 0.527 | 0.825 | 0 |
| **EIF1** | 0 | 1.04397668747434 | 1 | 0.94 | 0 |
| **FKBP10** | 0 | -1.54350509872617 | 0.682 | 0.944 | 0 |
| **ACLY** | 0 | -0.978510840689133 | 0.968 | 0.994 | 0 |
| **STAT5B** | 0 | -0.941909490729453 | 0.405 | 0.836 | 0 |
| **STAT3** | 0 | -1.12181550548736 | 0.995 | 0.999 | 0 |
| **CAVIN1** | 0 | -1.40196601867299 | 0.435 | 0.761 | 0 |
| **DUSP3** | 0 | -0.984780850882326 | 0.519 | 0.856 | 0 |
| **G6PC3** | 0 | -1.17271356075236 | 0.928 | 0.997 | 0 |
| **SLC25A39** | 0 | 0.923175824885285 | 1 | 1 | 0 |
| **GRN** | 0 | -1.00091618264584 | 0.998 | 0.999 | 0 |
| **FAM171A2** | 0 | -1.28511333834117 | 0.458 | 0.863 | 0 |

| | | | | | |
|---|---|---|---|---|---|
| **FZD2** | 0 | -1.92891106600359 | 0.011 | 0.643 | 0 |
| **GJC1** | 0 | -1.7547092945924 | 0.441 | 0.851 | 0 |
| **HOXB2** | 0 | 1.3656325051003 | 0.954 | 0.513 | 0 |
| **PRAC1** | 0 | -0.952064936532229 | 0.018 | 0.527 | 0 |
| **IGF2BP1** | 0 | -0.897870241827252 | 0.996 | 1 | 0 |
| **PPP1R9B** | 0 | -0.891039674027514 | 0.355 | 0.735 | 0 |
| **COL1A1** | 0 | -1.532723963531 | 0.471 | 0.779 | 0 |
| **TOM1L1** | 0 | -1.73334576082722 | 0.368 | 0.954 | 0 |
| **TRIM25** | 0 | -1.39526165943027 | 0.375 | 0.851 | 0 |
| **SRSF1** | 0 | -1.11734612455429 | 0.942 | 0.997 | 0 |
| **RNF43** | 0 | -1.56869323860649 | 0.085 | 0.744 | 0 |
| **GDPD1** | 0 | 1.42258529674994 | 0.772 | 0.176 | 0 |
| **YPEL2** | 0 | 2.57518098334681 | 0.999 | 0.526 | 0 |
| **CLTC** | 0 | -0.885672844155668 | 0.956 | 0.987 | 0 |
| **MAP3K3** | 0 | -0.89547913498985 | 0.522 | 0.868 | 0 |
| **LIMD2** | 0 | -1.23588615054567 | 0.19 | 0.761 | 0 |
| **AXIN2** | 0 | -1.3291443291785 | 0.6 | 0.934 | 0 |
| **FAM20A** | 0 | -2.13870866693647 | 0.357 | 0.867 | 0 |
| **JPT1** | 0 | 1.04553048052415 | 1 | 0.992 | 0 |
| **GALK1** | 0 | 0.986880000116431 | 0.923 | 0.613 | 0 |
| **ACOX1** | 0 | 1.0702035040424 | 0.993 | 0.848 | 0 |
| **SEC14L1** | 0 | -0.944928519682645 | 0.91 | 0.988 | 0 |
| **SOCS3** | 0 | -1.68049413663281 | 0.349 | 0.854 | 0 |
| **TIMP2** | 0 | -1.61083285057631 | 0.646 | 0.94 | 0 |
| **LGALS3BP** | 0 | -2.67307657235039 | 0.183 | 0.951 | 0 |
| **CBX4** | 0 | 1.72013542260192 | 0.992 | 0.691 | 0 |
| **TBC1D16** | 0 | -1.4192723134031 | 0.742 | 0.98 | 0 |
| **RPTOR** | 0 | 1.0276487352547 | 0.99 | 0.874 | 0 |
| **CHMP6** | 0 | -0.869868084666697 | 0.528 | 0.878 | 0 |
| **MCRIP1** | 0 | -1.38224057803666 | 0.807 | 0.984 | 0 |
| **PYCR1** | 0 | 2.1738108829282 | 1 | 0.58 | 0 |
| **NOTUM** | 0 | -1.02403542723332 | 0.075 | 0.435 | 0 |
| **TBCD** | 0 | 1.31484578021421 | 1 | 0.999 | 0 |
| **LAMA1** | 0 | 2.00262573677926 | 0.916 | 0.202 | 0 |

| | | | | | |
|---|---|---|---|---|---|
| **TWSG1** | 0 | -0.815047308557557 | 0.711 | 0.891 | 0 |
| **PPP4R1** | 0 | -1.04496137590147 | 0.933 | 0.997 | 0 |
| **PIEZO2** | 0 | -2.85495463264771 | 0.038 | 0.847 | 0 |
| **CHMP1B** | 0 | -0.838393303060586 | 0.909 | 0.99 | 0 |
| **IMPA2** | 0 | 0.942817135962749 | 0.998 | 0.921 | 0 |
| **CABLES1** | 0 | -0.971793745364271 | 0.206 | 0.7 | 0 |
| **CDH2** | 0 | -2.42290202784629 | 0.02 | 0.704 | 0 |
| **RNF125** | 0 | 2.37736182848089 | 1 | 0.761 | 0 |
| **DTNA** | 0 | -0.928792598623267 | 0.018 | 0.319 | 0 |
| **MAPRE2** | 0 | -0.811939002256955 | 0.67 | 0.891 | 0 |
| **GALNT1** | 0 | -1.33102829604914 | 0.842 | 0.986 | 0 |
| **TPGS2** | 0 | -1.65142045902963 | 0.801 | 0.994 | 0 |
| **HDHD2** | 0 | -0.945875745000704 | 0.676 | 0.95 | 0 |
| **CTIF** | 0 | -0.863122152109891 | 0.212 | 0.588 | 0 |
| **MAPK4** | 0 | -1.22106346970866 | 0.01 | 0.529 | 0 |
| **MBD2** | 0 | -0.83010230269818 | 0.978 | 0.999 | 0 |
| **TCF4** | 0 | -1.48479852581261 | 0.402 | 0.883 | 0 |
| **NARS** | 0 | 0.872973839881509 | 1 | 1 | 0 |
| **NEDD4L** | 0 | -1.73916272447562 | 0.352 | 0.946 | 0 |
| **PMAIP1** | 0 | -1.54270058096677 | 0.887 | 0.952 | 0 |
| **RNF152** | 0 | -0.820377823785412 | 0.014 | 0.46 | 0 |
| **PHLPP1** | 0 | -0.825610855652914 | 0.408 | 0.797 | 0 |
| **KDSR** | 0 | -0.855915739653781 | 0.799 | 0.966 | 0 |
| **DOK6** | 0 | -1.64091789530531 | 0.17 | 0.605 | 0 |
| **SALL3** | 0 | -2.37672986833881 | 0.613 | 0.959 | 0 |
| **ATP9B** | 0 | 0.889326957399247 | 0.993 | 0.925 | 0 |
| **SLC66A2** | 0 | 1.14883869901739 | 0.996 | 0.831 | 0 |
| **HSBP1L1** | 0 | 1.20142099205396 | 0.771 | 0.313 | 0 |
| **PLPP2** | 0 | 1.74170686854218 | 0.772 | 0.151 | 0 |
| **SHC2** | 0 | -0.954435562275344 | 0.29 | 0.681 | 0 |
| **HCN2** | 0 | 0.86359596875296 | 0.954 | 0.646 | 0 |
| **RNF126** | 0 | 0.877556943465362 | 1 | 0.984 | 0 |
| **ARID3A** | 0 | 1.05664747617863 | 0.999 | 0.973 | 0 |
| **CNN2** | 0 | -2.08947674907391 | 0.988 | 1 | 0 |

| | | | | | |
|---|---|---|---|---|---|
| **CIRBP** | 0 | -1.05374391802733 | 0.993 | 0.999 | 0 |
| **GAMT** | 0 | 0.908157751790031 | 0.885 | 0.535 | 0 |
| **REEP6** | 0 | 2.24790038615244 | 0.998 | 0.593 | 0 |
| **ZNF57** | 0 | 0.961812339363188 | 0.58 | 0.051 | 0 |
| **TJP3** | 0 | -0.904141367910276 | 0.555 | 0.826 | 0 |
| **NMRK2** | 0 | -1.08033879890593 | 0.679 | 0.873 | 0 |
| **CLPP** | 0 | 0.958027296973018 | 1 | 0.997 | 0 |
| **TNFSF9** | 0 | 1.78452307224518 | 0.952 | 0.184 | 0 |
| **CD70** | 0 | 3.04792137912232 | 0.993 | 0.241 | 0 |
| **PRR36** | 0 | -1.73901396157922 | 0.038 | 0.778 | 0 |
| **FBN3** | 0 | -1.18538386877096 | 0.072 | 0.615 | 0 |
| **CERS4** | 0 | -1.31817209439967 | 0.843 | 0.973 | 0 |
| **DNMT1** | 0 | -1.02589679364397 | 0.948 | 0.988 | 0 |
| **PDE4A** | 0 | 1.68023382482496 | 0.984 | 0.502 | 0 |
| **AP1M2** | 0 | 1.41309751999412 | 0.87 | 0.208 | 0 |
| **KANK2** | 0 | -0.877312107636458 | 0.303 | 0.724 | 0 |
| **PLPPR2** | 0 | 0.904522119866429 | 0.698 | 0.288 | 0 |
| **ZNF627** | 0 | -0.933332589770582 | 0.196 | 0.702 | 0 |
| **ACP5** | 0 | 0.838258788388165 | 0.472 | 0.041 | 0 |
| **ZNF844** | 0 | 0.975726165439717 | 0.552 | 0.038 | 0 |
| **TRIR** | 0 | -1.08839346764964 | 0.98 | 0.999 | 0 |
| **RTBDN** | 0 | 0.870154331950702 | 0.558 | 0.11 | 0 |
| **PRKACA** | 0 | -0.855590587654617 | 0.974 | 0.998 | 0 |
| **NOTCH3** | 0 | -2.69223419410598 | 0.453 | 0.981 | 0 |
| **AP1M1** | 0 | -0.823523842353184 | 0.807 | 0.959 | 0 |
| **NR2F6** | 0 | -0.870736229077674 | 0.998 | 1 | 0 |
| **BST2** | 0 | -4.0254087189974 | 0.341 | 0.942 | 0 |
| **SLC5A5** | 0 | 0.911366832961715 | 0.599 | 0.112 | 0 |
| **ARRDC2** | 0 | 1.39312987613931 | 0.889 | 0.343 | 0 |
| **PIK3R2** | 0 | -1.39704515617859 | 0.983 | 0.999 | 0 |
| **JUND** | 0 | -1.29111277843428 | 0.998 | 1 | 0 |
| **GDF15** | 0 | 2.00689475557571 | 0.688 | 0.077 | 0 |
| **ISYNA1** | 0 | -0.881503788272062 | 0.967 | 0.983 | 0 |
| **FKBP8** | 0 | -0.929669426121617 | 0.857 | 0.983 | 0 |

| | | | | | |
|---|---|---|---|---|---|
| **DDX49** | 0 | 0.803493677652933 | 1 | 0.968 | 0 |
| **PBX4** | 0 | 1.6197218971203 | 0.888 | 0.218 | 0 |
| **ZNF253** | 0 | 0.990856587943874 | 0.563 | 0.038 | 0 |
| **ZNF486** | 0 | 1.4450858403391 | 0.905 | 0.428 | 0 |
| **ZNF626** | 0 | 1.11207044866396 | 0.617 | 0.055 | 0 |
| **ZNF257** | 0 | 1.28717078499943 | 0.65 | 0.026 | 0 |
| **ZNF676** | 0 | 2.43332442437433 | 0.864 | 0.151 | 0 |
| **ZNF98** | 0 | 1.08477622920599 | 0.57 | 0.028 | 0 |
| **ZNF492** | 0 | 2.19661110469759 | 0.945 | 0.105 | 0 |
| **ZNF728** | 0 | 1.15275360094316 | 0.611 | 0.021 | 0 |
| **TSHZ3** | 0 | -0.989237540003561 | 0.126 | 0.569 | 0 |
| **DMKN** | 0 | -2.26211837499999 | 0.343 | 0.967 | 0 |
| **ZNF146** | 0 | 0.875773265998435 | 1 | 0.999 | 0 |
| **ZNF829** | 0 | 0.894221048183095 | 0.536 | 0.027 | 0 |
| **ZNF568** | 0 | 1.99797146608682 | 0.954 | 0.169 | 0 |
| **PPP1R14A** | 0 | 1.46836155227889 | 0.992 | 0.676 | 0 |
| **YIF1B** | 0 | 1.27367185031259 | 1 | 0.997 | 0 |
| **PLEKHG2** | 0 | -0.905816066566038 | 0.172 | 0.642 | 0 |
| **TIMM50** | 0 | 1.05622414246001 | 1 | 0.999 | 0 |
| **PLD3** | 0 | -1.14554126172768 | 1 | 1 | 0 |
| **AXL** | 0 | -1.42500908014593 | 0.015 | 0.501 | 0 |
| **ATP1A3** | 0 | 1.96797324676363 | 0.934 | 0.188 | 0 |
| **MEGF8** | 0 | -1.09277024894595 | 0.302 | 0.814 | 0 |
| **ZNF229** | 0 | 0.884003651119265 | 0.713 | 0.317 | 0 |
| **BCL3** | 0 | 0.984928972525648 | 0.786 | 0.398 | 0 |
| **BCAM** | 0 | -2.20595128543947 | 0.462 | 0.95 | 0 |
| **APOC1** | 0 | 1.20906054408843 | 0.983 | 0.744 | 0 |
| **TRAPPC6A** | 0 | -0.92832014172058 | 0.786 | 0.959 | 0 |
| **ERCC1** | 0 | -0.887984559008594 | 0.666 | 0.913 | 0 |
| **VASP** | 0 | -1.15104254950122 | 0.993 | 0.999 | 0 |
| **PRKD2** | 0 | -0.992480951129918 | 0.646 | 0.931 | 0 |
| **SLC1A5** | 0 | 1.41880932549681 | 1 | 0.984 | 0 |
| **ARHGAP35** | 0 | -1.35455250135315 | 0.61 | 0.961 | 0 |
| **C5AR1** | 0 | 1.04474665903932 | 0.58 | 0.047 | 0 |

| | | | | | |
|---|---|---|---|---|---|
| **CA11** | 0 | -1.66176870380524 | 0.034 | 0.705 | 0 |
| **RCN3** | 0 | 1.27718380116711 | 0.756 | 0.195 | 0 |
| **SCAF1** | 0 | -0.95511828913608 | 0.893 | 0.988 | 0 |
| **KCNC3** | 0 | 0.840970298356699 | 0.698 | 0.367 | 0 |
| **FAM71E1** | 0 | 0.955879135506173 | 0.724 | 0.26 | 0 |
| **LRRC4B** | 0 | 0.991865027458355 | 0.534 | 0.032 | 0 |
| **KLK13** | 0 | 1.3043022092941 | 0.917 | 0.438 | 0 |
| **IGLON5** | 0 | -1.05564571230664 | 0.117 | 0.646 | 0 |
| **LIM2** | 0 | 1.03803638497234 | 0.581 | 0.056 | 0 |
| **ZNF577** | 0 | 1.07565647864189 | 0.849 | 0.52 | 0 |
| **ZNF649** | 0 | 0.955465862446689 | 0.989 | 0.884 | 0 |
| **ZNF677** | 0 | 1.34635152341415 | 0.747 | 0.06 | 0 |
| **MBOAT7** | 0 | -1.05349437414025 | 0.909 | 0.995 | 0 |
| **NLRP7** | 0 | 3.5159591621627 | 1 | 0.63 | 0 |
| **NLRP2** | 0 | 1.1129930137573 | 1 | 0.996 | 0 |
| **TNNI3** | 0 | 1.10127550940777 | 0.832 | 0.336 | 0 |
| **ZNF579** | 0 | -0.814509820361249 | 0.154 | 0.6 | 0 |
| **EPN1** | 0 | -0.998973641489798 | 0.976 | 0.999 | 0 |
| **ZNF444** | 0 | -0.826564666622373 | 0.739 | 0.944 | 0 |
| **ZNF667** | 0 | 1.25394470643208 | 0.677 | 0.032 | 0 |
| **PEG3** | 0 | 1.23382418767878 | 0.58 | 0.033 | 0 |
| **ZCCHC3** | 0 | -1.49691693989103 | 0.49 | 0.916 | 0 |
| **SOX12** | 0 | -1.3027156549313 | 0.489 | 0.865 | 0 |
| **TRIB3** | 0 | 2.51127355324089 | 0.994 | 0.571 | 0 |
| **RBCK1** | 0 | -0.983647296559084 | 0.289 | 0.766 | 0 |
| **SLC52A3** | 0 | -2.19157593763116 | 0.413 | 0.831 | 0 |
| **PSMF1** | 0 | -0.987875207323681 | 0.997 | 1 | 0 |
| **CPXM1** | 0 | -0.967862035731283 | 0.897 | 0.974 | 0 |
| **PCED1A** | 0 | -1.18530329611609 | 0.319 | 0.797 | 0 |
| **PTPRA** | 0 | -0.80804781096883 | 0.834 | 0.969 | 0 |
| **SLC4A11** | 0 | 0.828281539603341 | 0.703 | 0.307 | 0 |
| **CDC25B** | 0 | -1.19715031574267 | 0.973 | 0.992 | 0 |
| **MAVS** | 0 | -1.01201321809727 | 0.838 | 0.984 | 0 |
| **CDS2** | 0 | -1.57743592084556 | 0.752 | 0.987 | 0 |

| | | | | | |
|---|---|---|---|---|---|
| **GPCPD1** | 0 | 0.825061127066632 | 0.98 | 0.856 | 0 |
| **PLCB4** | 0 | -1.40530271301953 | 0.04 | 0.714 | 0 |
| **JAG1** | 0 | -1.76881042445812 | 0.926 | 0.996 | 0 |
| **BTBD3** | 0 | -1.91825541177547 | 0.528 | 0.951 | 0 |
| **BFSP1** | 0 | -0.828132678001904 | 0.079 | 0.525 | 0 |
| **PET117** | 0 | -0.818045457430647 | 0.802 | 0.972 | 0 |
| **ZNF133** | 0 | -0.891219099944664 | 0.416 | 0.825 | 0 |
| **RBBP9** | 0 | 1.49865613158661 | 0.991 | 0.702 | 0 |
| **SMIM26** | 0 | -0.920560792340561 | 0.745 | 0.927 | 0 |
| **RIN2** | 0 | -1.69348112116018 | 0.549 | 0.95 | 0 |
| **RALGAPA2** | 0 | -1.69842084736127 | 0.455 | 0.951 | 0 |
| **XRN2** | 0 | -0.974053052611687 | 1 | 1 | 0 |
| **CST3** | 0 | -2.83369247738633 | 0.586 | 0.978 | 0 |
| **SYNDIG1** | 0 | -1.48156906302219 | 0.007 | 0.706 | 0 |
| **APMAP** | 0 | -1.08809124589011 | 0.962 | 0.999 | 0 |
| **PYGB** | 0 | 1.72331370472564 | 1 | 0.999 | 0 |
| **ID1** | 0 | -1.06725544166836 | 0.971 | 0.982 | 0 |
| **ASXL1** | 0 | -1.59606926793326 | 0.42 | 0.918 | 0 |
| **NOL4L** | 0 | -1.09562447743398 | 0.204 | 0.713 | 0 |
| **COMMD7** | 0 | -0.835617987630651 | 0.672 | 0.921 | 0 |
| **DNMT3B** | 0 | -1.34874102688862 | 0.999 | 0.996 | 0 |
| **PXMP4** | 0 | 0.885056244872955 | 0.783 | 0.449 | 0 |
| **MAP1LC3A** | 0 | 0.820443746989509 | 0.587 | 0.17 | 0 |
| **PROCR** | 0 | 2.6105685029526 | 0.928 | 0.535 | 0 |
| **MMP24** | 0 | -1.13094340263198 | 0.859 | 0.95 | 0 |
| **RBM12** | 0 | -0.878708671037922 | 0.904 | 0.991 | 0 |
| **SCAND1** | 0 | -0.985876531454885 | 0.798 | 0.929 | 0 |
| **EPB41L1** | 0 | -1.27463101993304 | 0.793 | 0.976 | 0 |
| **DLGAP4** | 0 | -1.01032990516191 | 0.928 | 0.984 | 0 |
| **MYL9** | 0 | -3.53889165036775 | 0.805 | 0.991 | 0 |
| **TGIF2** | 0 | -1.91750409585575 | 0.558 | 0.957 | 0 |
| **RAB5IF** | 0 | -1.12560674211339 | 0.949 | 0.999 | 0 |
| **SAMHD1** | 0 | 2.31988766592144 | 1 | 0.844 | 0 |
| **SRC** | 0 | -1.1309029480219 | 0.632 | 0.922 | 0 |

| | | | | | |
|---|---|---|---|---|---|
| **NNAT** | 0 | 2.94200956636639 | 0.814 | 0.015 | 0 |
| **EMILIN3** | 0 | -1.12872135739495 | 0.036 | 0.466 | 0 |
| **MYBL2** | 0 | 1.37525427380601 | 1 | 0.989 | 0 |
| **HNF4A** | 0 | 1.57885136810151 | 0.799 | 0.084 | 0 |
| **PKIG** | 0 | -0.823973377045938 | 0.526 | 0.813 | 0 |
| **ADA** | 0 | 2.36357568837328 | 0.998 | 0.598 | 0 |
| **STK4** | 0 | -0.851748449653504 | 0.875 | 0.98 | 0 |
| **DBNDD2** | 0 | -1.55580386915139 | 0.225 | 0.875 | 0 |
| **PIGT** | 0 | -0.987862088841943 | 0.986 | 0.999 | 0 |
| **SNX21** | 0 | 1.213722327306 | 0.947 | 0.638 | 0 |
| **EYA2** | 0 | -1.65630719494347 | 0.018 | 0.661 | 0 |
| **ZMYND8** | 0 | -0.875046936981864 | 0.997 | 0.999 | 0 |
| **NCOA3** | 0 | 1.53088321876425 | 1 | 0.949 | 0 |
| **SULF2** | 0 | -1.62933280428131 | 0.685 | 0.965 | 0 |
| **RNF114** | 0 | 0.948861691995752 | 1 | 0.999 | 0 |
| **UBE2V1** | 0 | -0.98280962204682 | 0.999 | 1 | 0 |
| **PTPN1** | 0 | -1.03227130300836 | 0.918 | 0.997 | 0 |
| **BCAS4** | 0 | -0.824899685193241 | 0.559 | 0.855 | 0 |
| **ATP9A** | 0 | -0.903255235135855 | 0.621 | 0.886 | 0 |
| **SALL4** | 0 | -0.936813995338421 | 0.997 | 0.999 | 0 |
| **TFAP2C** | 0 | -1.09421481242287 | 0.902 | 0.982 | 0 |
| **BMP7** | 0 | -3.14185790012853 | 0.461 | 0.981 | 0 |
| **STX16** | 0 | -0.821236813999146 | 0.966 | 0.997 | 0 |
| **NELFCD** | 0 | -0.834681526440491 | 0.978 | 0.999 | 0 |
| **CTSZ** | 0 | -1.02281320370027 | 0.522 | 0.82 | 0 |
| **ATP5F1E** | 0 | -1.31134679333548 | 0.974 | 0.999 | 0 |
| **FAM217B** | 0 | -0.928714527673193 | 0.641 | 0.939 | 0 |
| **LSM14B** | 0 | -1.16824200010264 | 0.945 | 0.997 | 0 |
| **LAMA5** | 0 | -1.35303201569213 | 0.987 | 0.999 | 0 |
| **COL9A3** | 0 | -1.03550609753081 | 0.988 | 0.99 | 0 |
| **NKAIN4** | 0 | -1.85910248723825 | 0.173 | 0.782 | 0 |
| **KCNQ2** | 0 | 1.62284610006707 | 0.868 | 0.165 | 0 |
| **DNAJC5** | 0 | -0.868236651519159 | 0.934 | 0.993 | 0 |
| **NRIP1** | 0 | -1.54096263546685 | 0.327 | 0.857 | 0 |

| | | | | | |
|---|---|---|---|---|---|
| **BTG3** | 0 | 0.898347984177347 | 0.981 | 0.81 | 0 |
| **CHODL** | 0 | -0.818868617780386 | 0.382 | 0.73 | 0 |
| **APP** | 0 | -1.22725425335317 | 1 | 1 | 0 |
| **HUNK** | 0 | -1.60197530245726 | 0.025 | 0.711 | 0 |
| **CFAP298** | 0 | -1.12705420630077 | 0.982 | 0.998 | 0 |
| **SLC5A3** | 0 | -1.0894755750859 | 0.655 | 0.91 | 0 |
| **PSMG1** | 0 | 0.848732759817609 | 0.999 | 0.96 | 0 |
| **MX1** | 0 | 3.08639949373304 | 0.901 | 0.058 | 0 |
| **PDXK** | 0 | 1.1009656039014 | 1 | 0.985 | 0 |
| **DNMT3L** | 0 | 2.58280384893438 | 1 | 0.919 | 0 |
| **TSPEAR** | 0 | -0.857180589678349 | 0.001 | 0.39 | 0 |
| **UBE2G2** | 0 | -0.867467795562295 | 0.772 | 0.957 | 0 |
| **COL18A1** | 0 | -4.343578125262 | 0.576 | 0.991 | 0 |
| **SPATC1L** | 0 | -0.968963847846834 | 0.073 | 0.619 | 0 |
| **LSS** | 0 | -1.30890713546862 | 0.943 | 0.997 | 0 |
| **HDHD5** | 0 | 1.13698818365408 | 0.999 | 0.829 | 0 |
| **CECR2** | 0 | -1.02840995695598 | 0.836 | 0.979 | 0 |
| **ARVCF** | 0 | -0.986961361153315 | 0.104 | 0.611 | 0 |
| **MED15** | 0 | 1.26372228598252 | 0.996 | 0.924 | 0 |
| **HIC2** | 0 | 1.2813167555567 | 0.999 | 0.928 | 0 |
| **SDF2L1** | 0 | 1.03888663261947 | 0.991 | 0.919 | 0 |
| **ZNF280A** | 0 | 1.03904300502564 | 0.529 | 0.031 | 0 |
| **DERL3** | 0 | 0.901825283616525 | 0.478 | 0.087 | 0 |
| **SUSD2** | 0 | 1.3736276724658 | 0.998 | 0.832 | 0 |
| **TPST2** | 0 | 0.916294717631904 | 0.965 | 0.703 | 0 |
| **NEFH** | 0 | 2.56012366980606 | 0.999 | 0.653 | 0 |
| **THOC5** | 0 | 1.17740724898113 | 0.999 | 0.954 | 0 |
| **TCN2** | 0 | -2.02538547773431 | 0.554 | 0.959 | 0 |
| **SELENOM** | 0 | 1.09124103176176 | 0.874 | 0.469 | 0 |
| **LIMK2** | 0 | 1.2104936873542 | 0.998 | 0.884 | 0 |
| **PATZ1** | 0 | -0.981080398525925 | 0.851 | 0.969 | 0 |
| **HMOX1** | 0 | 0.910731510137559 | 0.973 | 0.77 | 0 |
| **TRIOBP** | 0 | -1.18993024319872 | 0.589 | 0.947 | 0 |
| **ATF4** | 0 | 1.21516525491386 | 1 | 0.989 | 0 |

| | | | | | |
|---|---|---|---|---|---|
| **PMM1** | 0 | 0.857893036742877 | 0.918 | 0.652 | 0 |
| **SNU13** | 0 | 1.22672499513762 | 1 | 1 | 0 |
| **NDUFA6** | 0 | 1.12769580332899 | 1 | 0.954 | 0 |
| **A4GALT** | 0 | 2.09595574035334 | 1 | 0.713 | 0 |
| **TSPO** | 0 | 0.924567257843305 | 0.658 | 0.265 | 0 |
| **SULT4A1** | 0 | 0.857098857609615 | 0.748 | 0.414 | 0 |
| **PNPLA3** | 0 | -1.00635101113301 | 0.365 | 0.723 | 0 |
| **PARVB** | 0 | 1.54372807335355 | 0.893 | 0.258 | 0 |
| **RTL6** | 0 | -0.948586376270899 | 0.092 | 0.639 | 0 |
| **ARHGAP8** | 0 | 1.22639043924324 | 0.971 | 0.616 | 0 |
| **FBLN1** | 0 | -0.96532915540566 | 0.985 | 0.998 | 0 |
| **CRELD2** | 0 | 0.934235802770911 | 0.968 | 0.819 | 0 |
| **PLXNB2** | 0 | 1.12946612626902 | 0.998 | 0.904 | 0 |
| **CSF2RA** | 0 | 1.51244867309249 | 0.808 | 0.246 | 0 |
| **ASMTL** | 0 | 1.68085957622048 | 0.966 | 0.688 | 0 |
| **DHRSX** | 0 | 2.17386589327303 | 0.998 | 0.876 | 0 |
| **CD99** | 0 | -3.08260675501124 | 0.132 | 0.933 | 0 |
| **GYG2** | 0 | -1.16377735147746 | 0.518 | 0.908 | 0 |
| **ARSE** | 0 | -1.59614489174295 | 0.035 | 0.687 | 0 |
| **MXRA5** | 0 | 1.34114809237922 | 0.772 | 0.221 | 0 |
| **TBL1X** | 0 | -0.90048434667413 | 0.69 | 0.897 | 0 |
| **SHROOM2** | 0 | -0.997284086486178 | 0.638 | 0.926 | 0 |
| **WWC3** | 0 | -2.00909515497182 | 0.189 | 0.866 | 0 |
| **ARHGAP6** | 0 | -1.28008476315184 | 0.119 | 0.686 | 0 |
| **PRPS2** | 0 | -1.10572129612732 | 0.748 | 0.895 | 0 |
| **TMSB4X** | 0 | -2.76072362642229 | 0.995 | 1 | 0 |
| **GPM6B** | 0 | -1.47852132194182 | 0.383 | 0.866 | 0 |
| **AP1S2** | 0 | -2.03634444362943 | 0.196 | 0.892 | 0 |
| **GRPR** | 0 | -1.08168757107377 | 0.003 | 0.39 | 0 |
| **CTPS2** | 0 | -1.55932515007548 | 0.683 | 0.985 | 0 |
| **RBBP7** | 0 | -0.822731478812941 | 0.945 | 0.98 | 0 |
| **SH3KBP1** | 0 | -2.04355373373921 | 0.719 | 0.984 | 0 |
| **SMS** | 0 | -3.16439670910334 | 0.753 | 0.996 | 0 |
| **ACOT9** | 0 | -1.1591499018853 | 0.62 | 0.933 | 0 |

| | | | | | |
|---|---|---|---|---|---|
| **SAT1** | 0 | -2.05888242600507 | 0.799 | 0.99 | 0 |
| **PCYT1B** | 0 | -1.55742470121938 | 0.027 | 0.763 | 0 |
| **MAGEB2** | 0 | 1.09537764257759 | 0.46 | 0.032 | 0 |
| **DMD** | 0 | -0.978929231992328 | 0.587 | 0.851 | 0 |
| **USP9X** | 0 | -0.942443758499717 | 1 | 1 | 0 |
| **JADE3** | 0 | -1.2093520270844 | 0.378 | 0.871 | 0 |
| **USP11** | 0 | -1.54659754108519 | 0.793 | 0.987 | 0 |
| **TIMP1** | 0 | -1.3279059606801 | 0.961 | 0.996 | 0 |
| **PORCN** | 0 | 1.13259512110768 | 0.82 | 0.408 | 0 |
| **PQBP1** | 0 | 0.837845632276379 | 1 | 0.998 | 0 |
| **GRIPAP1** | 0 | -0.85757048254971 | 0.791 | 0.966 | 0 |
| **GSPT2** | 0 | -1.6944803754032 | 0.425 | 0.956 | 0 |
| **MAGED1** | 0 | -2.44784175926557 | 0.929 | 0.998 | 0 |
| **WNK3** | 0 | -1.11218194968759 | 0.371 | 0.848 | 0 |
| **MAGED2** | 0 | -2.08656353112131 | 0.994 | 1 | 0 |
| **TRO** | 0 | -1.52269029994362 | 0.665 | 0.971 | 0 |
| **APEX2** | 0 | -0.905411009901387 | 0.886 | 0.982 | 0 |
| **MAGEH1** | 0 | -1.60382999159831 | 0.22 | 0.891 | 0 |
| **USP51** | 0 | -1.3534533260596 | 0.082 | 0.756 | 0 |
| **UBQLN2** | 0 | -1.19249626027255 | 0.93 | 0.997 | 0 |
| **SPIN3** | 0 | -0.827084728061917 | 0.355 | 0.749 | 0 |
| **SPIN4** | 0 | -1.03154883652907 | 0.767 | 0.96 | 0 |
| **ARHGEF9** | 0 | -1.67143948854699 | 0.151 | 0.852 | 0 |
| **AMER1** | 0 | -1.3381693097956 | 0.169 | 0.798 | 0 |
| **ZC4H2** | 0 | -1.00799834306735 | 0.244 | 0.757 | 0 |
| **HEPH** | 0 | -2.1952887256236 | 0.011 | 0.886 | 0 |
| **STARD8** | 0 | -0.810256464089476 | 0.004 | 0.435 | 0 |
| **EFNB1** | 0 | -3.07710791880779 | 0.237 | 0.856 | 0 |
| **PJA1** | 0 | -1.21444160135791 | 0.772 | 0.973 | 0 |
| **EDA** | 0 | -1.20925656964318 | 0.557 | 0.923 | 0 |
| **FOXO4** | 0 | -2.80555201368568 | 0.203 | 0.871 | 0 |
| **MED12** | 0 | -1.43413923198501 | 0.174 | 0.707 | 0 |
| **OGT** | 0 | -1.02835631433458 | 0.98 | 0.99 | 0 |
| **HDAC8** | 0 | 0.946975437409424 | 0.988 | 0.896 | 0 |

| | | | | | |
|---|---|---|---|---|---|
| **PHKA1** | 0 | 2.12144144148135 | 1 | 0.888 | 0 |
| **NAP1L2** | 0 | -1.08217425673941 | 0.069 | 0.639 | 0 |
| **SLC16A2** | 0 | -2.72688889666905 | 0.66 | 0.967 | 0 |
| **ABCB7** | 0 | -0.849573611482664 | 0.98 | 0.999 | 0 |
| **UPRT** | 0 | -1.02524171376963 | 0.398 | 0.84 | 0 |
| **COX7B** | 0 | 1.04199773971717 | 1 | 0.875 | 0 |
| **TAF9B** | 0 | -0.873270893727921 | 0.334 | 0.785 | 0 |
| **TBX22** | 0 | -1.55266879390189 | 0.002 | 0.563 | 0 |
| **SH3BGRL** | 0 | -1.33052094603712 | 0.698 | 0.968 | 0 |
| **ZNF711** | 0 | -1.13739320952656 | 0.413 | 0.859 | 0 |
| **KLHL4** | 0 | -2.01765022152473 | 0.002 | 0.673 | 0 |
| **NAP1L3** | 0 | -1.77101239535516 | 0.176 | 0.835 | 0 |
| **DIAPH2** | 0 | -1.14372936738896 | 0.106 | 0.676 | 0 |
| **TNMD** | 0 | -1.56169303737995 | 0.008 | 0.628 | 0 |
| **TSPAN6** | 0 | -1.50374118373533 | 0.937 | 0.995 | 0 |
| **TRMT2B** | 0 | -0.872291484680345 | 0.529 | 0.881 | 0 |
| **CENPI** | 0 | -1.00574614957885 | 0.715 | 0.877 | 0 |
| **HNRNPH2** | 0 | -1.03416889208164 | 0.988 | 0.998 | 0 |
| **ARMCX1** | 0 | -1.7887571111031 | 0.418 | 0.938 | 0 |
| **ARMCX3** | 0 | -0.864192869231866 | 0.275 | 0.669 | 0 |
| **ARMCX2** | 0 | -2.11678453788616 | 0.708 | 0.99 | 0 |
| **TMSB15A** | 0 | -1.29484759416694 | 0.1 | 0.537 | 0 |
| **BEX4** | 0 | -1.75336103268491 | 0.779 | 0.993 | 0 |
| **TCEAL8** | 0 | -1.38596449259332 | 0.916 | 0.997 | 0 |
| **TCEAL9** | 0 | -1.73497635313423 | 0.659 | 0.977 | 0 |
| **BEX3** | 0 | -2.62897557240325 | 0.951 | 0.996 | 0 |
| **TCEAL4** | 0 | -0.840368042362026 | 0.954 | 0.996 | 0 |
| **NRK** | 0 | -3.27536171745145 | 0.038 | 0.824 | 0 |
| **COL4A6** | 0 | -1.8602012523561 | 0.002 | 0.764 | 0 |
| **COL4A5** | 0 | -2.60560055251112 | 0.049 | 0.888 | 0 |
| **LUZP4** | 0 | 1.05987456503937 | 0.529 | 0.028 | 0 |
| **PGRMC1** | 0 | -1.14038608489603 | 0.997 | 1 | 0 |
| **SEPTIN6** | 0 | -1.98423146826636 | 0.102 | 0.869 | 0 |
| **ZBTB33** | 0 | -1.29123009058746 | 0.625 | 0.963 | 0 |

| | | | | | |
|---|---|---|---|---|---|
| **C1GALT1C1** | 0 | -1.24453253143568 | 0.652 | 0.953 | 0 |
| **SMARCA1** | 0 | -1.21013748136883 | 0.695 | 0.936 | 0 |
| **ELF4** | 0 | -1.17323420782054 | 0.071 | 0.523 | 0 |
| **IGSF1** | 0 | 1.26705588651734 | 0.929 | 0.621 | 0 |
| **STK26** | 0 | -1.15888282999123 | 0.746 | 0.972 | 0 |
| **MBNL3** | 0 | -2.72940309641503 | 0.358 | 0.949 | 0 |
| **HS6ST2** | 0 | -1.99598214880366 | 0.045 | 0.822 | 0 |
| **GPC4** | 0 | -4.55438192242508 | 0.469 | 0.964 | 0 |
| **GPC3** | 0 | -1.51840227937073 | 0.831 | 0.95 | 0 |
| **RTL8C** | 0 | -0.995460883715697 | 0.846 | 0.953 | 0 |
| **SMIM10L2A** | 0 | -1.69806499494642 | 0.154 | 0.771 | 0 |
| **ZIC3** | 0 | -2.157895494368 | 0.098 | 0.878 | 0 |
| **LDOC1** | 0 | -0.884981153858495 | 0.538 | 0.871 | 0 |
| **FMR1** | 0 | -0.832718279999132 | 0.927 | 0.992 | 0 |
| **AFF2** | 0 | 1.08688398966331 | 0.937 | 0.643 | 0 |
| **IDS** | 0 | -1.4423790749218 | 0.187 | 0.843 | 0 |
| **MAMLD1** | 0 | -1.65902691441341 | 0.004 | 0.396 | 0 |
| **MTM1** | 0 | -1.56379227740894 | 0.559 | 0.952 | 0 |
| **CD99L2** | 0 | -1.1538185702552 | 0.136 | 0.636 | 0 |
| **VMA21** | 0 | -1.12233178265685 | 0.966 | 0.999 | 0 |
| **ZNF185** | 0 | -1.67531340416406 | 0.151 | 0.828 | 0 |
| **CCNQ** | 0 | 1.02138715797349 | 0.974 | 0.743 | 0 |
| **DUSP9** | 0 | 0.935970294497171 | 0.55 | 0.075 | 0 |
| **SLC6A8** | 0 | 1.05203198628203 | 0.999 | 0.904 | 0 |
| **PDZD4** | 0 | -1.02454456550096 | 0.185 | 0.565 | 0 |
| **IRAK1** | 0 | 1.11273803742248 | 0.884 | 0.424 | 0 |
| **FLNA** | 0 | -1.02810192655385 | 1 | 1 | 0 |
| **TMLHE** | 0 | -1.14086849546141 | 0.202 | 0.785 | 0 |
| **DDX3Y** | 0 | 0.869198481976292 | 0.963 | 0.907 | 0 |
| **SDC3** | 6.4421 | -0.936383356232183 | 0.292 | 0.647 | 1.06972700 |
| **BNC2** | 9.6712 | -0.886560339265068 | 0.51 | 0.752 | 1.6059148[illegible] |
| **H2AFX** | 3.1720 | -1.12227923396605 | 0.941 | 0.956 | 5.2672011[illegible] |
| **MVD** | 5.5218 | -0.935049406065 | 0.954 | 0.976 | 9.1691010[illegible] |
| **BTG2** | 9.8537 | -0.960672565399117 | 0.555 | 0.817 | 1.6362141[illegible] |

| | | | | | |
|---|---|---|---|---|---|
| **NAV1** | 4.9061 | -0.871945270436885 | 0.285 | 0.63 | 8.1466029 |
| **TMEFF2** | 5.8576 | -1.12614407508006 | 0.266 | 0.6 | 9.7265847 |
| **LAMB2** | 7.1664 | -0.86674212118924 | 0.83 | 0.927 | 1.1899833 |
| **MAP2** | 1.0213 | -1.17010964031306 | 0.001 | 0.27 | 1.6959735 |
| **STOX2** | 2.7210 | -0.808955683276276 | 0.607 | 0.835 | 4.5182659 |
| **ZFP69B** | 1.3226 | 0.803014208969814 | 0.683 | 0.405 | 2.1961936 |
| **TMEM158** | 1.9503 | -0.90383769313294 | 0.27 | 0.619 | 3.2385169 |
| **NKD1** | 3.3736 | -0.892627828964831 | 0.202 | 0.537 | 5.6019023 |
| **DHCR7** | 3.8916 | -0.916643058515761 | 0.987 | 0.983 | 6.4621493 |
| **STON2** | 6.2825 | -0.8060777760104 | 0.292 | 0.64 | 1.0432197 |
| **COL27A1** | 9.0825 | 0.836094891610443 | 0.686 | 0.404 | 1.5081537 |
| **SLC2A3** | 2.5814 | -1.00265203947925 | 0.798 | 0.882 | 4.2864633 |
| **DDIT4** | 7.4751 | 0.92372991998546 | 0.977 | 0.911 | 1.2412439 |
| **COL5A1** | 6.9474 | -1.24391914912451 | 0.463 | 0.744 | 1.1536176 |
| **PAX3** | 1.5715 | -0.844347860310929 | 0 | 0.243 | 2.6094813 |
| **GLI3** | 5.9899 | -1.65719221993709 | 0.026 | 0.285 | 9.9462357 |
| **AMOT** | 1.9326 | -0.858726688788178 | 0.161 | 0.472 | 3.2092060 |
| **LHX5** | 8.3085 | -0.965476668188111 | 0.004 | 0.238 | 1.3796319 |
| **MYL7** | 1.5414 | -1.08777900310788 | 0.024 | 0.276 | 2.5595278 |
| **DPYSL5** | 3.2021 | -1.15530006295229 | 0.01 | 0.245 | 5.3172153 |
| **TMEFF1** | 1.5165 | -0.860615233262406 | 0.787 | 0.882 | 2.5181672 |
| **ZBTB10** | 7.6711 | -0.830825890125363 | 0.987 | 0.986 | 1.2737870 |
| **CYP1B1** | 2.1179 | -0.987162020002041 | 0.154 | 0.45 | 3.5168650 |
| **3,00 F** | 8.6073 | 1.30029802624179 | 0.783 | 0.618 | 1.4292468 |
| **CBR3** | 2.1432 | -0.904997367926303 | 0.198 | 0.489 | 3.5589028 |
| **TCF7L1** | 3.0003 | -1.07178768870255 | 0.35 | 0.609 | 4.9821222 |
| **CDX2** | 2.4629 | -0.83609077318605 | 0.028 | 0.259 | 4.0897302 |
| **AHDC1** | 1.4290 | -0.914161613267206 | 0.401 | 0.674 | 2.3730163 |
| **HMGB2** | 2.5225 | -1.19117829951435 | 0.939 | 0.925 | 4.1887498 |
| **CYP26A1** | 8.3211 | -1.1912491395481 | 0.021 | 0.239 | 1.3817304 |
| **PAMR1** | 6.6356 | -0.842653930310208 | 0.011 | 0.219 | 1.1018446 |
| **COLEC12** | 4.0819 | -1.32006704648379 | 0.251 | 0.526 | 6.7780400 |
| **ANOS1** | 6.9683 | -0.910066773919737 | 0.054 | 0.283 | 1.1570964 |
| **SOCS1** | 4.0804 | -0.877451962439154 | 0.744 | 0.842 | 6.7755825 |

| | | | | | |
|---|---|---|---|---|---|
| **CPE** | 7.6424 | -1.6216017512125 | 0.042 | 0.266 | 1.26903697 |
| **PLPP4** | 1.0955 | -0.835013228769079 | 0.351 | 0.587 | 1.81913943 |
| **FSTL1** | 4.2573 | -1.27760294397264 | 0.932 | 0.949 | 7.06933891 |
| **MCAM** | 4.7051 | -0.854637392349302 | 0.425 | 0.671 | 7.81296528 |
| **CNN1** | 1.0106 | -1.27182546139644 | 0.231 | 0.494 | 1.67826644 |
| **OLIG3** | 2.7074 | -0.800350931087732 | 0.051 | 0.262 | 4.49567105 |
| **TLE4** | 3.4238 | -1.02404829259921 | 0.305 | 0.561 | 5.68528290 |
| **JUN** | 2.4199 | -1.02222868415464 | 0.523 | 0.732 | 4.01826300 |
| **EFHD1** | 7.9879 | -0.954386172347464 | 0.827 | 0.866 | 1.32640580 |
| **FN1** | 1.1942 | -1.48436352107119 | 0.93 | 0.948 | 1.98306172 |
| **SOX9** | 4.5432 | -0.930174003540802 | 0.041 | 0.236 | 7.54402601 |
| **CSRP1** | 2.2180 | -1.35323519561418 | 0.85 | 0.872 | 3.68311882 |
| **DLK1** | 4.2205 | -0.946381562459717 | 0.46 | 0.197 | 7.00819934 |
| **SDC2** | 1.3248 | -0.931352577358073 | 0.151 | 0.367 | 2.19997312 |
| **RNASE1** | 3.0173 | -0.855769617171141 | 0.081 | 0.273 | 5.01030976 |
| **UHRF1** | 7.8694 | -1.00107408259965 | 0.842 | 0.827 | 1.30672581 |
| **NLGN4X** | 3.8452 | -0.92283070605613 | 0.124 | 0.305 | 6.38506980 |
| **KCNG1** | 7.7170 | -0.839878625518753 | 0.466 | 0.638 | 1.28141610 |
| **MT-CO3** | 8.1595 | -1.94411017024646 | 0.91 | 0.883 | 1.35489685 |
| **MYL4** | 5.4558 | -1.02097575640359 | 0.019 | 0.146 | 9.05943998 |
| **WNT4** | 1.0549 | -0.986813847361595 | 0.004 | 0.113 | 1.75177728 |
| **SOX21** | 1.1500 | -0.856691049850051 | 0.03 | 0.162 | 1.90967517 |
| **PMP22** | 6.8259 | -0.864800384806423 | 0.489 | 0.651 | 1.13344933 |
| **EPCAM** | 7.2196 | -0.963570445442686 | 0.981 | 0.924 | 1.19883007 |
| **CNTNAP2** | 9.8939 | -1.27855636382563 | 0.658 | 0.72 | 1.64289843 |
| **DACT1** | 1.0647 | -0.815410631053992 | 0.173 | 0.337 | 1.76808761 |
| **MT-CYB** | 5.1114 | -1.59617118411157 | 0.814 | 0.772 | 8.48754378 |
| **FLNC** | 3.9455 | -1.03867263104474 | 0.548 | 0.635 | 6.55164509 |
| **BOC** | 5.1143 | -0.927582317458912 | 0.103 | 0.233 | 8.49238881 |
| **NRP2** | 2.5947 | -0.803751366001813 | 0.225 | 0.377 | 4.30860274 |
| **CCNA1** | 4.3056 | 1.71226906526492 | 0.196 | 0.107 | 7.14959170 |
| **ANXA1** | 3.9098 | -0.909663730506964 | 0.058 | 0.134 | 6.49228548 |
| **MT-CO2** | 4.9007 | -0.991704540199478 | 0.947 | 0.896 | 8.13770254 |
| **CXCL14** | 6.8926 | -0.951953270078824 | 0.726 | 0.729 | 1.14452682 |

| | | | | | |
|---|---|---|---|---|---|
| **GREB1L** | 1.6950 | -1.06192744107294 | 0.143 | 0.203 | 2.81458546 |
| **MT-ND4L** | 3.8932 | -0.848121133729392 | 0.825 | 0.744 | 6.46476305 |
| **PRSS23** | 1.2851 | -0.808783493167757 | 0.568 | 0.517 | 0.21340263 |

Supplemental Table S2_Sheet_8

| | p_val | avg_log2FC | pct.1 | pct.2 | p_val_adj |
|---|---|---|---|---|---|
| **NOC2L** | 0 | 0.87108301878428 | 1 | 0.985 | 0 |
| **ISG15** | 0 | 1.05220504227084 | 0.955 | 0.753 | 0 |
| **AGRN** | 0 | -1.1360934598528 | 0.935 | 0.986 | 0 |
| **B3GALT6** | 0 | -0.985545814427316 | 0.335 | 0.767 | 0 |
| **ATAD3B** | 0 | 1.5105144519114 | 0.999 | 0.934 | 0 |
| **FAAP20** | 0 | -0.903669977550987 | 0.944 | 0.995 | 0 |
| **AJAP1** | 0 | -0.838211117697025 | 0.216 | 0.576 | 0 |
| **TNFRSF25** | 0 | 0.874472963249046 | 0.597 | 0.108 | 0 |
| **PLEKHG5** | 0 | 0.958241047980407 | 0.684 | 0.21 | 0 |
| **ERRFI1** | 0 | 2.15307709072313 | 0.996 | 0.751 | 0 |
| **CLSTN1** | 0 | -0.949098957011477 | 0.979 | 0.997 | 0 |
| **CTNNBIP1** | 0 | -1.19198870695999 | 0.77 | 0.978 | 0 |
| **RBP7** | 0 | 0.944850621532159 | 0.584 | 0.093 | 0 |
| **DFFA** | 0 | 1.02751103723503 | 1 | 0.991 | 0 |
| **DRAXIN** | 0 | -1.93677704901803 | 0.447 | 0.849 | 0 |
| **MIIP** | 0 | 1.21304224962221 | 0.998 | 0.948 | 0 |
| **TNFRSF8** | 0 | 1.1716455348474 | 0.594 | 0.021 | 0 |
| **EPHA2** | 0 | -1.60082482740936 | 0.722 | 0.987 | 0 |
| **FBXO42** | 0 | -0.847635394897723 | 0.591 | 0.881 | 0 |
| **MFAP2** | 0 | -1.88931969939647 | 0.913 | 0.999 | 0 |
| **SDHB** | 0 | 0.819869971109303 | 1 | 0.996 | 0 |
| **ARHGEF10L** | 0 | -1.15043033535392 | 0.16 | 0.708 | 0 |
| **KLHDC7A** | 0 | -1.21943265433183 | 0.004 | 0.484 | 0 |
| **HP1BP3** | 0 | -0.958353976885709 | 0.657 | 0.921 | 0 |
| **EIF4G3** | 0 | -0.987997998809067 | 0.725 | 0.941 | 0 |
| **ECE1** | 0 | -1.20764464310834 | 0.846 | 0.988 | 0 |
| **ALPL** | 0 | 0.930816579936975 | 1 | 0.992 | 0 |
| **LUZP1** | 0 | -0.985539270707308 | 0.819 | 0.976 | 0 |
| **ID3** | 0 | -3.58930777147205 | 0.461 | 0.983 | 0 |
| **FUCA1** | 0 | 2.18031906517893 | 0.997 | 0.74 | 0 |
| **CNR2** | 0 | 1.45505884699823 | 0.721 | 0.031 | 0 |
| **LDLRAP1** | 0 | 1.02709805732858 | 0.976 | 0.782 | 0 |

| | | | | | |
|---|---|---|---|---|---|
| **MAN1C1** | 0 | 1.7315636622623 | 0.978 | 0.545 | 0 |
| **SH3BGRL3** | 0 | -1.35609838356048 | 0.86 | 0.99 | 0 |
| **ARID1A** | 0 | -0.853981793914581 | 0.949 | 0.995 | 0 |
| **TENT5B** | 0 | 1.15892144900271 | 0.994 | 0.797 | 0 |
| **AHDC1** | 0 | -1.2934400862647 | 0.444 | 0.856 | 0 |
| **SESN2** | 0 | 1.69572279844537 | 0.986 | 0.817 | 0 |
| **RCC1** | 0 | 0.833131821426946 | 0.998 | 0.969 | 0 |
| **PTPRU** | 0 | 1.74382787265369 | 0.999 | 0.898 | 0 |
| **SDC3** | 0 | -0.92024046869055 | 0.366 | 0.734 | 0 |
| **KHDRBS1** | 0 | -0.817159776849502 | 0.975 | 0.997 | 0 |
| **TXLNA** | 0 | -0.862287451030993 | 0.919 | 0.994 | 0 |
| **EIF3I** | 0 | 0.880384330889113 | 0.981 | 0.756 | 0 |
| **MARCKSL1** | 0 | -1.59947835133884 | 1 | 1 | 0 |
| **KIAA1522** | 0 | -1.25496534968676 | 0.919 | 0.986 | 0 |
| **RNF19B** | 0 | 1.92410720038782 | 1 | 0.909 | 0 |
| **ZNF362** | 0 | -1.93166933562125 | 0.422 | 0.963 | 0 |
| **PHC2** | 0 | -2.04620493409389 | 0.042 | 0.744 | 0 |
| **MAP7D1** | 0 | -1.53612486597861 | 0.539 | 0.94 | 0 |
| **YRDC** | 0 | 1.0729753830371 | 0.999 | 0.956 | 0 |
| **YBX1** | 0 | -0.849911650039092 | 1 | 1 | 0 |
| **CLDN19** | 0 | -1.02102667770104 | 0.153 | 0.586 | 0 |
| **SVBP** | 0 | -0.839828911980438 | 0.945 | 0.996 | 0 |
| **KDM4A** | 0 | 1.05094283622002 | 0.999 | 0.963 | 0 |
| **KLF17** | 0 | 1.14398046602345 | 0.367 | 0.014 | 0 |
| **KLF18** | 0 | 0.849506391085685 | 0.284 | 0.004 | 0 |
| **PLK3** | 0 | 1.46681694726927 | 0.815 | 0.277 | 0 |
| **HPDL** | 0 | 0.887948563311069 | 0.567 | 0.108 | 0 |
| **PRDX1** | 0 | 1.33902293787997 | 1 | 0.999 | 0 |
| **UQCRH** | 0 | 1.19768704889036 | 1 | 0.999 | 0 |
| **NSUN4** | 0 | 0.819446338928804 | 0.951 | 0.746 | 0 |
| **ZYG11B** | 0 | -0.899797106154041 | 0.739 | 0.953 | 0 |
| **ZYG11A** | 0 | 1.93081685676782 | 0.996 | 0.686 | 0 |
| **LRP8** | 0 | -2.06222669938747 | 0.62 | 0.962 | 0 |
| **LRRC42** | 0 | -0.816574429369361 | 0.796 | 0.955 | 0 |

| | | | | | |
|---|---|---|---|---|---|
| **DHCR24** | 0 | -1.68198083629563 | 0.998 | 1 | 0 |
| **PLPP3** | 0 | -0.911256726759198 | 0.534 | 0.866 | 0 |
| **DAB1** | 0 | -1.55319377012229 | 0.178 | 0.652 | 0 |
| **MYSM1** | 0 | 1.28167635203251 | 1 | 0.939 | 0 |
| **PATJ** | 0 | -1.0449542735711 | 0.622 | 0.883 | 0 |
| **L1TD1** | 0 | 1.20169989972378 | 1 | 0.991 | 0 |
| **PGM1** | 0 | -2.81409240197944 | 0.585 | 0.996 | 0 |
| **ROR1** | 0 | -1.26770960792152 | 0.354 | 0.722 | 0 |
| **CACHD1** | 0 | -2.18452398535728 | 0.404 | 0.953 | 0 |
| **RAVER2** | 0 | -1.1479365592209 | 0.242 | 0.768 | 0 |
| **AK4** | 0 | 1.54251230325759 | 0.998 | 0.958 | 0 |
| **GADD45A** | 0 | 1.08096546447162 | 0.95 | 0.752 | 0 |
| **WLS** | 0 | -4.44277076089117 | 0.246 | 0.974 | 0 |
| **CTH** | 0 | 2.09268168596212 | 0.989 | 0.598 | 0 |
| **LHX8** | 0 | 1.17500511403504 | 0.504 | 0.018 | 0 |
| **RABGGTB** | 0 | 1.03348007622316 | 1 | 0.998 | 0 |
| **ST6GALNAC5** | 0 | -1.15969656317926 | 0.059 | 0.57 | 0 |
| **DNAJB4** | 0 | 0.800891963912552 | 0.711 | 0.394 | 0 |
| **ADGRL2** | 0 | -1.11330167680494 | 0.91 | 0.994 | 0 |
| **PRKACB** | 0 | -1.25957827579364 | 0.309 | 0.824 | 0 |
| **DDAH1** | 0 | -1.20698122818861 | 0.242 | 0.642 | 0 |
| **LMO4** | 0 | 1.61098698700595 | 0.998 | 0.956 | 0 |
| **RBMXL1** | 0 | 1.12036824490363 | 0.961 | 0.651 | 0 |
| **CDC7** | 0 | -0.822359574749711 | 0.753 | 0.932 | 0 |
| **BRDT** | 0 | 3.09679237498486 | 0.998 | 0.227 | 0 |
| **TMED5** | 0 | 0.864939409483843 | 0.997 | 0.942 | 0 |
| **3,00 F** | 0 | 2.00357559142876 | 0.909 | 0.525 | 0 |
| **PTBP2** | 0 | -1.03842229529378 | 0.783 | 0.968 | 0 |
| **AGL** | 0 | 2.28735181398219 | 0.999 | 0.919 | 0 |
| **PRMT6** | 0 | -0.815911131585406 | 0.461 | 0.791 | 0 |
| **SLC25A24** | 0 | -1.21127473730602 | 0.837 | 0.993 | 0 |
| **FAM102B** | 0 | 0.893704100494276 | 0.798 | 0.521 | 0 |
| **HENMT1** | 0 | 1.53055529829073 | 0.772 | 0.055 | 0 |
| **TAF13** | 0 | 2.78004420534151 | 1 | 0.911 | 0 |

| | | | | | |
|---|---|---|---|---|---|
| **SARS** | 0 | 0.892296487563015 | 0.964 | 0.804 | 0 |
| **PSRC1** | 0 | 0.813563822572439 | 0.881 | 0.621 | 0 |
| **GSTM3** | 0 | -1.83900509781261 | 0.136 | 0.743 | 0 |
| **PIFO** | 0 | 1.60684912887557 | 0.95 | 0.467 | 0 |
| **PHTF1** | 0 | -0.926563110057027 | 0.666 | 0.911 | 0 |
| **OLFML3** | 0 | -3.829951157163 | 0.664 | 0.992 | 0 |
| **SYT6** | 0 | -1.05218863466939 | 0.103 | 0.587 | 0 |
| **VANGL1** | 0 | -1.64344904832062 | 0.666 | 0.968 | 0 |
| **ZNF697** | 0 | 0.972025887463582 | 0.927 | 0.733 | 0 |
| **NOTCH2** | 0 | -1.19430948853937 | 0.447 | 0.886 | 0 |
| **PRKAB2** | 0 | 0.984246206311375 | 0.969 | 0.798 | 0 |
| **GJA5** | 0 | -0.860103344903136 | 0.008 | 0.382 | 0 |
| **SV2A** | 0 | -1.08813666943803 | 0.377 | 0.808 | 0 |
| **PLEKHO1** | 0 | -1.35768426219088 | 0.593 | 0.943 | 0 |
| **CA14** | 0 | -0.861819054321364 | 0.085 | 0.564 | 0 |
| **HORMAD1** | 0 | 3.15968522083715 | 0.997 | 0.184 | 0 |
| **CTSK** | 0 | -2.18562927268187 | 0.233 | 0.723 | 0 |
| **MLLT11** | 0 | -1.22050175834457 | 0.323 | 0.785 | 0 |
| **S100A6** | 0 | 1.68527689686333 | 0.654 | 0.08 | 0 |
| **S100A16** | 0 | 2.24463537229106 | 0.881 | 0.392 | 0 |
| **S100A13** | 0 | 2.19910748111191 | 1 | 0.996 | 0 |
| **CRTC2** | 0 | 0.821199389452213 | 0.868 | 0.601 | 0 |
| **RAB13** | 0 | -1.03878825760787 | 0.907 | 0.988 | 0 |
| **C1orf43** | 0 | -0.833094660065589 | 0.928 | 0.99 | 0 |
| **IL6R** | 0 | 1.22587651468301 | 0.81 | 0.237 | 0 |
| **SHE** | 0 | 0.87067552824477 | 0.426 | 0.038 | 0 |
| **CKS1B** | 0 | 0.989086760664389 | 1 | 0.992 | 0 |
| **FDPS** | 0 | -0.905524396486888 | 0.934 | 0.97 | 0 |
| **MEX3A** | 0 | -2.31991039239794 | 0.323 | 0.975 | 0 |
| **SLC25A44** | 0 | 1.51713745677739 | 1 | 0.946 | 0 |
| **C1orf61** | 0 | -1.43271893389823 | 0.098 | 0.477 | 0 |
| **MEF2D** | 0 | 0.826533530132039 | 0.825 | 0.489 | 0 |
| **CRABP2** | 0 | -1.59615365677924 | 0.999 | 1 | 0 |
| **HDGF** | 0 | -1.36743025818435 | 0.867 | 0.991 | 0 |

| | | | | | |
|---|---|---|---|---|---|
| **IFI16** | 0 | 1.93055665842965 | 0.952 | 0.563 | 0 |
| **DUSP23** | 0 | 1.22178189096258 | 0.709 | 0.046 | 0 |
| **IGSF8** | 0 | -1.0735943695885 | 0.69 | 0.884 | 0 |
| **ATP1A2** | 0 | -0.858038485452234 | 0.005 | 0.422 | 0 |
| **PEA15** | 0 | -1.73359819603548 | 0.966 | 0.999 | 0 |
| **VANGL2** | 0 | -0.893105578576069 | 0.891 | 0.986 | 0 |
| **F11R** | 0 | 1.81419743284087 | 1 | 0.988 | 0 |
| **TSTD1** | 0 | 0.964414080071825 | 0.58 | 0.034 | 0 |
| **NECTIN4** | 0 | 1.10770912870951 | 0.541 | 0.052 | 0 |
| **UHMK1** | 0 | 0.919806920310161 | 0.976 | 0.844 | 0 |
| **RGS5** | 0 | -2.9043615815827 | 0.076 | 0.791 | 0 |
| **UCK2** | 0 | -0.89427188531871 | 0.926 | 0.997 | 0 |
| **CREG1** | 0 | -1.19871999749329 | 0.461 | 0.876 | 0 |
| **MPZL1** | 0 | -1.18122080180117 | 0.63 | 0.93 | 0 |
| **NME7** | 0 | -0.941056612843691 | 0.21 | 0.688 | 0 |
| **PIGC** | 0 | -0.846958043296797 | 0.359 | 0.757 | 0 |
| **RGS2** | 0 | 1.78768754452524 | 0.92 | 0.447 | 0 |
| **RO60** | 0 | -1.06124170240188 | 0.872 | 0.99 | 0 |
| **GLRX2** | 0 | -1.11886948676945 | 0.168 | 0.724 | 0 |
| **ELF3** | 0 | 1.51498790513313 | 0.785 | 0.279 | 0 |
| **LRRN2** | 0 | -1.33755912729832 | 0.027 | 0.606 | 0 |
| **NFASC** | 0 | -1.09630763889191 | 0.036 | 0.548 | 0 |
| **NUCKS1** | 0 | -1.19730040775439 | 0.996 | 1 | 0 |
| **CD55** | 0 | 0.83774725141847 | 0.75 | 0.379 | 0 |
| **PLXNA2** | 0 | -2.70902120956863 | 0.151 | 0.54 | 0 |
| **IRF6** | 0 | 0.855184693588385 | 0.568 | 0.051 | 0 |
| **TRAF5** | 0 | -1.31460408305384 | 0.129 | 0.731 | 0 |
| **SMYD2** | 0 | -0.978404622231432 | 0.896 | 0.982 | 0 |
| **PTPN14** | 0 | -1.14970516885361 | 0.912 | 0.99 | 0 |
| **CAPN2** | 0 | -2.60337747478096 | 0.27 | 0.974 | 0 |
| **CNIH4** | 0 | 1.16224900528058 | 0.999 | 0.878 | 0 |
| **CNIH3** | 0 | -1.03449330714117 | 0.021 | 0.443 | 0 |
| **ENAH** | 0 | -0.877480141259681 | 0.997 | 1 | 0 |
| **EPHX1** | 0 | -1.59480240857147 | 0.937 | 0.987 | 0 |

| | | | | | |
|---|---|---|---|---|---|
| **LEFTY1** | 0 | 1.39627922995041 | 0.995 | 0.903 | 0 |
| **MIXL1** | 0 | -2.98790882270229 | 0.861 | 0.941 | 0 |
| **WNT3A** | 0 | -1.67799153581977 | 0.041 | 0.521 | 0 |
| **CCSAP** | 0 | 0.96292865485833 | 0.787 | 0.375 | 0 |
| **C1orf198** | 0 | -1.37857479628949 | 0.467 | 0.926 | 0 |
| **GNPAT** | 0 | -1.19707437667776 | 0.643 | 0.95 | 0 |
| **EGLN1** | 0 | -1.49953408032613 | 0.396 | 0.913 | 0 |
| **SIPA1L2** | 0 | -1.07397831013666 | 0.748 | 0.951 | 0 |
| **AKT3** | 0 | -0.979678946848237 | 0.406 | 0.771 | 0 |
| **ZNF695** | 0 | 0.894664184906314 | 0.961 | 0.797 | 0 |
| **ZNF672** | 0 | -0.875675276515479 | 0.231 | 0.666 | 0 |
| **PXDN** | 0 | 0.891446796670329 | 0.997 | 0.829 | 0 |
| **ID2** | 0 | -1.72420487655922 | 0.166 | 0.793 | 0 |
| **IAH1** | 0 | 1.5631019669475 | 0.789 | 0.053 | 0 |
| **ODC1** | 0 | -1.58765432456833 | 0.841 | 0.995 | 0 |
| **SLC66A3** | 0 | 0.824551178174495 | 0.866 | 0.619 | 0 |
| **LPIN1** | 0 | 1.71613981660245 | 0.969 | 0.468 | 0 |
| **WDR35** | 0 | -0.894556275951556 | 0.189 | 0.648 | 0 |
| **SDC1** | 0 | -1.32700005568927 | 0.242 | 0.724 | 0 |
| **RHOB** | 0 | -1.67117835851224 | 0.734 | 0.987 | 0 |
| **TP53I3** | 0 | -0.811684619954867 | 0.528 | 0.817 | 0 |
| **EFR3B** | 0 | 1.77337265111464 | 0.96 | 0.522 | 0 |
| **POMC** | 0 | -1.22595124300198 | 0.456 | 0.772 | 0 |
| **SLC35F6** | 0 | 0.914551114873824 | 0.998 | 0.974 | 0 |
| **CENPA** | 0 | 1.4513421359854 | 0.991 | 0.838 | 0 |
| **AGBL5** | 0 | -0.949634966800923 | 0.842 | 0.979 | 0 |
| **KRTCAP3** | 0 | 2.04297920961466 | 0.955 | 0.221 | 0 |
| **FNDC4** | 0 | 1.21076302982931 | 0.85 | 0.46 | 0 |
| **ZNF512** | 0 | -0.848010471576138 | 0.386 | 0.751 | 0 |
| **CAPN13** | 0 | -1.14359048585068 | 0.079 | 0.642 | 0 |
| **VIT** | 0 | -0.976423243037875 | 0.012 | 0.329 | 0 |
| **EIF2AK2** | 0 | -1.04903211110146 | 0.537 | 0.899 | 0 |
| **CEBPZ** | 0 | 1.05033531874911 | 1 | 1 | 0 |
| **PRKD3** | 0 | -1.30225250640539 | 0.813 | 0.985 | 0 |

| | | | | | |
|---|---|---|---|---|---|
| **QPCT** | 0 | -1.77921064411135 | 0.553 | 0.91 | 0 |
| **CYP1B1** | 0 | -1.95487331476226 | 0.065 | 0.592 | 0 |
| **HNRNPLL** | 0 | -0.886920638611415 | 0.906 | 0.993 | 0 |
| **PKDCC** | 0 | -2.50880015163692 | 0.409 | 0.914 | 0 |
| **EML4** | 0 | -1.10046162366442 | 0.95 | 0.998 | 0 |
| **PREPL** | 0 | -0.826704457899534 | 0.523 | 0.842 | 0 |
| **CCDC88A** | 0 | -1.00029431895694 | 0.823 | 0.976 | 0 |
| **FANCL** | 0 | -0.900210262071609 | 0.232 | 0.667 | 0 |
| **B3GNT2** | 0 | -1.01927106684721 | 0.848 | 0.956 | 0 |
| **UGP2** | 0 | 0.95398320177618 | 0.993 | 0.925 | 0 |
| **CEP68** | 0 | -1.58166042058384 | 0.289 | 0.904 | 0 |
| **SPRED2** | 0 | 0.812320356203885 | 0.987 | 0.951 | 0 |
| **ANTXR1** | 0 | -1.78225754059949 | 0.117 | 0.858 | 0 |
| **GFPT1** | 0 | 0.815735845573116 | 0.998 | 0.987 | 0 |
| **MTHFD2** | 0 | 1.71613147408039 | 1 | 0.999 | 0 |
| **WDR54** | 0 | -0.914124886651437 | 0.774 | 0.948 | 0 |
| **WBP1** | 0 | -0.890039389605066 | 0.787 | 0.951 | 0 |
| **HK2** | 0 | 0.827257681411648 | 0.99 | 0.93 | 0 |
| **TACR1** | 0 | -2.07506471772439 | 0.01 | 0.448 | 0 |
| **VAMP8** | 0 | -1.88258899013392 | 0.939 | 0.999 | 0 |
| **MAP4K4** | 0 | -1.09635049496188 | 0.986 | 1 | 0 |
| **ACOXL** | 0 | 1.01590611259692 | 0.924 | 0.693 | 0 |
| **TMEM177** | 0 | -0.831173230907395 | 0.502 | 0.812 | 0 |
| **TMEM185B** | 0 | -1.03594846553568 | 0.687 | 0.953 | 0 |
| **TFCP2L1** | 0 | 1.61305475847959 | 0.858 | 0.237 | 0 |
| **GYPC** | 0 | -1.83747482890213 | 0.604 | 0.857 | 0 |
| **BIN1** | 0 | -1.40875660027723 | 0.631 | 0.965 | 0 |
| **PLEKHB2** | 0 | 1.65609620666585 | 1 | 0.999 | 0 |
| **LYPD1** | 0 | -1.70743318106665 | 0.227 | 0.646 | 0 |
| **MCM6** | 0 | -0.928214891515516 | 0.981 | 0.995 | 0 |
| **CXCR4** | 0 | 1.19751242814384 | 0.585 | 0.103 | 0 |
| **HNMT** | 0 | -0.885291393040011 | 0.23 | 0.644 | 0 |
| **ZEB2** | 0 | -1.60958108419465 | 0.104 | 0.725 | 0 |
| **KIF5C** | 0 | -1.41924246517842 | 0.891 | 0.997 | 0 |

| | | | | | |
|---|---|---|---|---|---|
| **RND3** | 0 | -1.2261037463901 | 0.079 | 0.527 | 0 |
| **NMI** | 0 | 1.08917862116579 | 0.811 | 0.274 | 0 |
| **RIF1** | 0 | 0.872638457428685 | 0.999 | 0.992 | 0 |
| **PKP4** | 0 | -0.906583122687217 | 0.352 | 0.73 | 0 |
| **BAZ2B** | 0 | -0.95446997531267 | 0.27 | 0.697 | 0 |
| **GCA** | 0 | -0.840424668933306 | 0.204 | 0.598 | 0 |
| **CERS6** | 0 | -0.912269077783677 | 0.352 | 0.751 | 0 |
| **KLHL23** | 0 | -1.68407234677687 | 0.343 | 0.918 | 0 |
| **SP5** | 0 | -2.56844008749529 | 0.517 | 0.983 | 0 |
| **GAD1** | 0 | -2.24148372895253 | 0.091 | 0.892 | 0 |
| **TLK1** | 0 | -0.816358734985337 | 0.976 | 0.998 | 0 |
| **PDK1** | 0 | -2.62837143268159 | 0.721 | 0.995 | 0 |
| **CDCA7** | 0 | -1.35203278362608 | 0.59 | 0.936 | 0 |
| **WIPF1** | 0 | -0.902055212971564 | 0.052 | 0.515 | 0 |
| **ATP5MC3** | 0 | 0.807594426125991 | 1 | 1 | 0 |
| **CCDC141** | 0 | -1.17011134968069 | 0.01 | 0.358 | 0 |
| **SESTD1** | 0 | 1.21469169303085 | 0.993 | 0.86 | 0 |
| **UBE2E3** | 0 | -1.94440451032706 | 0.576 | 0.99 | 0 |
| **GULP1** | 0 | -1.19422484090495 | 0.298 | 0.78 | 0 |
| **COL5A2** | 0 | -1.90976175948347 | 0.092 | 0.528 | 0 |
| **NABP1** | 0 | 1.49283207940028 | 0.887 | 0.303 | 0 |
| **SPATS2L** | 0 | -1.72399395324666 | 0.524 | 0.848 | 0 |
| **CFLAR** | 0 | 1.01280901346676 | 0.974 | 0.711 | 0 |
| **FZD7** | 0 | -1.90341131284648 | 0.553 | 0.872 | 0 |
| **ICA1L** | 0 | 1.18532214340388 | 0.716 | 0.118 | 0 |
| **NRP2** | 0 | -2.50051998641278 | 0.243 | 0.688 | 0 |
| **INO80D** | 0 | 0.864805760669697 | 0.985 | 0.877 | 0 |
| **FZD5** | 0 | 2.4370311114264 | 0.997 | 0.784 | 0 |
| **CRYGD** | 0 | 1.29602597412555 | 0.676 | 0.03 | 0 |
| **IDH1** | 0 | 1.23984387729621 | 1 | 0.996 | 0 |
| **CPS1** | 0 | -1.28578417560319 | 0.534 | 0.946 | 0 |
| **BARD1** | 0 | 0.836920009104786 | 0.924 | 0.702 | 0 |
| **IGFBP2** | 0 | -0.93073929784005 | 0.99 | 0.999 | 0 |
| **TMBIM1** | 0 | 2.03011245747003 | 0.965 | 0.373 | 0 |

| | | | | | |
|---|---|---|---|---|---|
| **PTPRN** | 0 | 1.18770929170847 | 0.628 | 0.085 | 0 |
| **OBSL1** | 0 | -2.51492071353203 | 0.324 | 0.987 | 0 |
| **SERPINE2** | 0 | -4.2499926948585 | 0.766 | 0.998 | 0 |
| **SLC19A3** | 0 | 1.15971982541871 | 0.78 | 0.273 | 0 |
| **FBXO36** | 0 | 2.20914072236776 | 0.992 | 0.525 | 0 |
| **PDE6D** | 0 | -0.862967793069111 | 0.299 | 0.717 | 0 |
| **ALPP** | 0 | 0.986488724286946 | 0.313 | 0.016 | 0 |
| **ALPG** | 0 | 4.99909265505516 | 0.995 | 0.475 | 0 |
| **NGEF** | 0 | 0.932909324177607 | 0.517 | 0.067 | 0 |
| **INPP5D** | 0 | 1.20307798087546 | 0.854 | 0.372 | 0 |
| **ARL4C** | 0 | -1.2207165227276 | 0.085 | 0.577 | 0 |
| **RAB17** | 0 | -1.34088009232108 | 0.055 | 0.495 | 0 |
| **LRRFIP1** | 0 | -0.843182845890944 | 0.63 | 0.882 | 0 |
| **EDEM1** | 0 | 0.981319165490033 | 0.974 | 0.876 | 0 |
| **RPUSD3** | 0 | 0.889777955257171 | 0.99 | 0.903 | 0 |
| **TIMP4** | 0 | -1.1779585719324 | 0.502 | 0.807 | 0 |
| **RAF1** | 0 | 0.829711524650723 | 1 | 0.998 | 0 |
| **NUP210** | 0 | -1.25360262611032 | 0.246 | 0.716 | 0 |
| **SLC6A6** | 0 | 1.30599666054409 | 0.998 | 0.888 | 0 |
| **RAB5A** | 0 | 0.94873102054108 | 1 | 0.973 | 0 |
| **EOMES** | 0 | -3.65499443454215 | 0.093 | 0.814 | 0 |
| **OSBPL10** | 0 | 0.861290848164392 | 0.889 | 0.63 | 0 |
| **CTDSPL** | 0 | 0.913157563205824 | 0.877 | 0.652 | 0 |
| **SLC25A38** | 0 | 1.10017940773489 | 0.996 | 0.949 | 0 |
| **CTNNB1** | 0 | -0.845308950142186 | 0.999 | 1 | 0 |
| **ZNF660** | 0 | -0.830141759176544 | 0.103 | 0.57 | 0 |
| **ZNF502** | 0 | 1.01766749625736 | 0.606 | 0.029 | 0 |
| **KIF9** | 0 | 0.907149866135064 | 0.851 | 0.529 | 0 |
| **KLHL18** | 0 | 0.990657096040197 | 0.932 | 0.688 | 0 |
| **SCAP** | 0 | -0.821647473167705 | 0.866 | 0.977 | 0 |
| **SHISA5** | 0 | 0.861638991791786 | 1 | 0.999 | 0 |
| **LAMB2** | 0 | -0.826606294199666 | 0.741 | 0.899 | 0 |
| **CCDC71** | 0 | -0.860035896118873 | 0.355 | 0.765 | 0 |
| **DUSP7** | 0 | 0.883615444652455 | 0.965 | 0.821 | 0 |

| | | | | | |
|---|---|---|---|---|---|
| **POC1A** | 0 | -0.854259988987084 | 0.592 | 0.851 | 0 |
| **TNNC1** | 0 | -0.893994439888289 | 0.055 | 0.454 | 0 |
| **NT5DC2** | 0 | -1.4745100420217 | 0.992 | 1 | 0 |
| **WNT5A** | 0 | -2.74651598004738 | 0.091 | 0.866 | 0 |
| **IL17RD** | 0 | -2.2199493203021 | 0.686 | 0.991 | 0 |
| **PDE12** | 0 | 0.985228460941867 | 0.901 | 0.498 | 0 |
| **PTPRG** | 0 | -2.03696398407033 | 0.652 | 0.988 | 0 |
| **EOGT** | 0 | -1.02489479815351 | 0.869 | 0.964 | 0 |
| **ROBO1** | 0 | -1.19637042469771 | 0.751 | 0.927 | 0 |
| **GBE1** | 0 | -0.865622404623485 | 0.65 | 0.887 | 0 |
| **DCBLD2** | 0 | -1.3981030061039 | 0.771 | 0.986 | 0 |
| **TBC1D23** | 0 | 1.67634504400141 | 0.998 | 0.927 | 0 |
| **NIT2** | 0 | 0.951167227493664 | 0.91 | 0.615 | 0 |
| **TMEM45A** | 0 | -1.32461119475264 | 0.166 | 0.744 | 0 |
| **ALCAM** | 0 | -1.79147978512493 | 0.066 | 0.714 | 0 |
| **DPPA2** | 0 | 2.54188567882195 | 0.997 | 0.432 | 0 |
| **DPPA4** | 0 | 0.898531960541974 | 0.999 | 0.999 | 0 |
| **NECTIN3** | 0 | -1.13276024805376 | 0.481 | 0.857 | 0 |
| **PHLDB2** | 0 | -1.6618851815373 | 0.531 | 0.918 | 0 |
| **ATG3** | 0 | 2.67949705941716 | 1 | 0.989 | 0 |
| **CCDC80** | 0 | 1.56083891175409 | 0.958 | 0.562 | 0 |
| **B4GALT4** | 0 | 1.08033557693312 | 0.871 | 0.475 | 0 |
| **FSTL1** | 0 | -1.56572768884469 | 0.951 | 0.997 | 0 |
| **HCLS1** | 0 | 1.50386972713726 | 0.743 | 0.131 | 0 |
| **CCDC14** | 0 | -0.964725492966132 | 0.779 | 0.961 | 0 |
| **HEG1** | 0 | -1.44967853000403 | 0.209 | 0.759 | 0 |
| **MGLL** | 0 | 1.86738168469474 | 0.775 | 0.097 | 0 |
| **TRH** | 0 | 0.983717791102356 | 0.6 | 0.096 | 0 |
| **AMOTL2** | 0 | -0.971983109099004 | 0.938 | 0.996 | 0 |
| **RBP1** | 0 | -2.54081862298247 | 0.628 | 0.99 | 0 |
| **SPSB4** | 0 | -2.05989341747937 | 0.144 | 0.864 | 0 |
| **ZBTB38** | 0 | -0.88066242358557 | 0.406 | 0.751 | 0 |
| **CHST2** | 0 | 1.88622368140915 | 0.987 | 0.737 | 0 |
| **DIPK2A** | 0 | -1.10339830778001 | 0.766 | 0.924 | 0 |

| | | | | | |
|---|---|---|---|---|---|
| **PLSCR4** | 0 | 0.976847752055783 | 0.627 | 0.275 | 0 |
| **WWTR1** | 0 | 0.888508770077552 | 0.981 | 0.915 | 0 |
| **P2RY1** | 0 | -1.32167050207397 | 0.069 | 0.585 | 0 |
| **MME** | 0 | -1.24375899883669 | 0.136 | 0.613 | 0 |
| **LRRC34** | 0 | 0.842960401051292 | 0.491 | 0.012 | 0 |
| **NCEH1** | 0 | 0.898921495295152 | 0.669 | 0.233 | 0 |
| **TBL1XR1** | 0 | 1.17994267276204 | 1 | 1 | 0 |
| **ZMAT3** | 0 | 1.05293133226623 | 0.87 | 0.594 | 0 |
| **GNB4** | 0 | -2.67220888701866 | 0.693 | 0.991 | 0 |
| **EEF1AKMT4** | 0 | 1.03460504494776 | 0.967 | 0.748 | 0 |
| **EPHB3** | 0 | -1.72959460710984 | 0.249 | 0.732 | 0 |
| **MAGEF1** | 0 | -1.44775746311097 | 0.4 | 0.922 | 0 |
| **LIPH** | 0 | 1.41591904533086 | 0.622 | 0.063 | 0 |
| **SENP2** | 0 | 1.68405681724505 | 1 | 0.992 | 0 |
| **ST6GAL1** | 0 | 0.808151564822497 | 0.998 | 0.989 | 0 |
| **LPP** | 0 | -1.28724596238897 | 0.687 | 0.955 | 0 |
| **P3H2** | 0 | -3.79239448834483 | 0.03 | 0.715 | 0 |
| **HES1** | 0 | -1.53846781610866 | 0.321 | 0.847 | 0 |
| **XXYLT1** | 0 | -1.16431253439027 | 0.489 | 0.889 | 0 |
| **MUC4** | 0 | 1.2909355388968 | 0.873 | 0.332 | 0 |
| **NRROS** | 0 | 0.863735458243799 | 0.533 | 0.05 | 0 |
| **NCBP2** | 0 | 1.10101814541984 | 1 | 1 | 0 |
| **ZNF595** | 0 | 1.5073439853624 | 0.989 | 0.772 | 0 |
| **ZNF732** | 0 | 1.374532134892 | 0.691 | 0.032 | 0 |
| **NSD2** | 0 | -1.62394210592802 | 0.101 | 0.83 | 0 |
| **MFSD10** | 0 | -1.02451120664744 | 0.592 | 0.905 | 0 |
| **ADRA2C** | 0 | -1.8525221477453 | 0.568 | 0.928 | 0 |
| **MSX1** | 0 | -1.82946751793226 | 0.023 | 0.417 | 0 |
| **CRMP1** | 0 | 1.1431876286333 | 0.995 | 0.882 | 0 |
| **S100P** | 0 | 2.10616878761807 | 0.687 | 0.076 | 0 |
| **GRPEL1** | 0 | 1.00113406748796 | 1 | 0.989 | 0 |
| **ZNF518B** | 0 | -0.900390318846718 | 0.242 | 0.694 | 0 |
| **PROM1** | 0 | 1.14761025931705 | 0.933 | 0.647 | 0 |
| **LDB2** | 0 | -2.03614289450861 | 0.197 | 0.818 | 0 |

| | | | | | |
|---|---|---|---|---|---|
| **SLIT2** | 0 | -1.57468957865861 | 0.006 | 0.474 | 0 |
| **RELL1** | 0 | 1.11271952249417 | 0.99 | 0.844 | 0 |
| **PGM2** | 0 | 1.67335409554096 | 0.99 | 0.837 | 0 |
| **KLF3** | 0 | 1.29465676443927 | 0.893 | 0.594 | 0 |
| **KLHL5** | 0 | -1.05143873160057 | 0.417 | 0.82 | 0 |
| **UCHL1** | 0 | -2.19187551709555 | 0.574 | 0.945 | 0 |
| **LIMCH1** | 0 | -2.1399751164695 | 0.162 | 0.728 | 0 |
| **SLC30A9** | 0 | -0.970616489517573 | 0.933 | 0.998 | 0 |
| **NIPAL1** | 0 | 1.3757029523461 | 0.751 | 0.175 | 0 |
| **TEC** | 0 | 1.50429113814443 | 0.759 | 0.086 | 0 |
| **SLAIN2** | 0 | 1.09028043716512 | 0.995 | 0.926 | 0 |
| **OCIAD2** | 0 | -1.27494312076163 | 0.107 | 0.681 | 0 |
| **USP46** | 0 | -0.801035677480244 | 0.74 | 0.929 | 0 |
| **RASL11B** | 0 | -0.916509769195541 | 0.29 | 0.669 | 0 |
| **REST** | 0 | 1.63597566669863 | 1 | 1 | 0 |
| **IGFBP7** | 0 | -1.35411424181486 | 0.057 | 0.522 | 0 |
| **MAPK10** | 0 | -2.77079449810789 | 0.028 | 0.86 | 0 |
| **PTPN13** | 0 | -0.954856577892664 | 0.407 | 0.747 | 0 |
| **HSD17B11** | 0 | 0.922349624601537 | 0.928 | 0.644 | 0 |
| **PKD2** | 0 | -1.40836482598068 | 0.49 | 0.931 | 0 |
| **ABCG2** | 0 | -1.3322268409034 | 0.406 | 0.676 | 0 |
| **PYURF** | 0 | -1.04307715954479 | 0.85 | 0.992 | 0 |
| **FAM13A** | 0 | -1.01271749261694 | 0.609 | 0.84 | 0 |
| **GRID2** | 0 | -1.50350836722022 | 0.084 | 0.656 | 0 |
| **RAP1GDS1** | 0 | -0.857131741432633 | 0.73 | 0.949 | 0 |
| **LEF1** | 0 | -1.25566665866968 | 0.045 | 0.478 | 0 |
| **ANK2** | 0 | -1.41959306711391 | 0.32 | 0.837 | 0 |
| **CAMK2D** | 0 | -1.05334959015793 | 0.209 | 0.661 | 0 |
| **PRSS12** | 0 | 1.22204729408406 | 0.785 | 0.284 | 0 |
| **LARP1B** | 0 | 1.19607225210352 | 0.962 | 0.7 | 0 |
| **PCDH10** | 0 | -3.54652820607313 | 0.106 | 0.766 | 0 |
| **SLC7A11** | 0 | 1.72158974114886 | 0.995 | 0.828 | 0 |
| **NOCT** | 0 | 1.62477714184271 | 0.999 | 0.907 | 0 |
| **MAML3** | 0 | -1.61824939741637 | 0.214 | 0.752 | 0 |

| | | | | | |
|---|---|---|---|---|---|
| **ELMOD2** | 0 | -0.81664698979598 | 0.516 | 0.831 | 0 |
| **SH3D19** | 0 | -0.844668594682951 | 0.448 | 0.792 | 0 |
| **FHDC1** | 0 | 1.19445401751332 | 0.672 | 0.255 | 0 |
| **TMEM131L** | 0 | 0.951404240624966 | 0.907 | 0.676 | 0 |
| **TRIM61** | 0 | 1.2161532529928 | 0.799 | 0.254 | 0 |
| **TRIM60** | 0 | 2.38651546353605 | 0.962 | 0.086 | 0 |
| **TRIM75P** | 0 | 1.16773775788215 | 0.61 | 0.03 | 0 |
| **CPE** | 0 | -1.26835192077499 | 0.061 | 0.431 | 0 |
| **AADAT** | 0 | -0.805406284545059 | 0.49 | 0.776 | 0 |
| **GALNT7** | 0 | -0.923646870454562 | 0.794 | 0.94 | 0 |
| **HMGB2** | 0 | -1.45394664243179 | 0.94 | 0.989 | 0 |
| **FAT1** | 0 | -2.03857159757933 | 0.718 | 0.994 | 0 |
| **ZFP42** | 0 | 3.97058851199714 | 0.999 | 0.364 | 0 |
| **TRIML2** | 0 | 1.657872417641 | 0.995 | 0.645 | 0 |
| **TRIP13** | 0 | -0.899628128845621 | 0.874 | 0.978 | 0 |
| **IRX4** | 0 | 2.00140038094294 | 0.741 | 0.036 | 0 |
| **IRX1** | 0 | -1.53011677253836 | 0.064 | 0.684 | 0 |
| **ADCY2** | 0 | -1.5104549918723 | 0.006 | 0.515 | 0 |
| **MARCH6** | 0 | -0.990087589874757 | 0.681 | 0.936 | 0 |
| **TRIO** | 0 | -0.997722666581953 | 0.64 | 0.901 | 0 |
| **PDZD2** | 0 | 0.816901494207587 | 0.668 | 0.26 | 0 |
| **RAI14** | 0 | -0.919533184639392 | 0.623 | 0.888 | 0 |
| **SELENOP** | 0 | -2.05745998293606 | 0.085 | 0.822 | 0 |
| **PARP8** | 0 | -0.868123200646642 | 0.122 | 0.576 | 0 |
| **FST** | 0 | -2.34200731765845 | 0.145 | 0.622 | 0 |
| **PLPP1** | 0 | 1.28329306667726 | 0.998 | 0.905 | 0 |
| **PLK2** | 0 | 1.30263646622662 | 0.702 | 0.277 | 0 |
| **PDE4D** | 0 | -1.20198061345174 | 0.408 | 0.78 | 0 |
| **ZSWIM6** | 0 | -1.43217343718706 | 0.126 | 0.745 | 0 |
| **KIF2A** | 0 | -0.994513252081874 | 0.925 | 0.997 | 0 |
| **SHISAL2B** | 0 | -1.50899126379222 | 0.06 | 0.672 | 0 |
| **ERBIN** | 0 | 1.20491362840021 | 0.998 | 0.955 | 0 |
| **PIK3R1** | 0 | -1.16394896235032 | 0.484 | 0.819 | 0 |
| **AK6** | 0 | 0.951630319223994 | 0.995 | 0.914 | 0 |

| | | | | | |
|---|---|---|---|---|---|
| **MARVELD2** | 0 | 1.06960591017056 | 0.734 | 0.184 | 0 |
| **MAP1B** | 0 | -0.980040219150712 | 0.972 | 0.999 | 0 |
| **ENC1** | 0 | -2.50011869183676 | 0.458 | 0.9 | 0 |
| **F2RL1** | 0 | 1.03206807082163 | 0.997 | 0.934 | 0 |
| **AGGF1** | 0 | -1.23001138315665 | 0.32 | 0.87 | 0 |
| **ARSB** | 0 | -1.13177786051404 | 0.312 | 0.775 | 0 |
| **DHFR** | 0 | -1.16994355976371 | 0.793 | 0.975 | 0 |
| **RASGRF2** | 0 | 1.66870772565042 | 0.855 | 0.215 | 0 |
| **FAM172A** | 0 | -1.00509869413835 | 0.404 | 0.82 | 0 |
| **RHOBTB3** | 0 | -2.57809114322017 | 0.055 | 0.495 | 0 |
| **CAST** | 0 | -1.38440983372311 | 0.063 | 0.702 | 0 |
| **SLCO4C1** | 0 | 1.91413390069781 | 0.99 | 0.572 | 0 |
| **PAM** | 0 | -2.07551645792712 | 0.099 | 0.884 | 0 |
| **EFNA5** | 0 | -1.53694104702826 | 0.154 | 0.746 | 0 |
| **MCC** | 0 | -1.67446349218125 | 0.345 | 0.886 | 0 |
| **DTWD2** | 0 | -1.27689219845151 | 0.458 | 0.735 | 0 |
| **ZNF608** | 0 | -1.60290305567838 | 0.589 | 0.966 | 0 |
| **FBN2** | 0 | 0.808408231677648 | 0.444 | 0.047 | 0 |
| **ADAMTS19** | 0 | -0.89209790917643 | 0.01 | 0.344 | 0 |
| **TCF7** | 0 | -1.44303711447527 | 0.876 | 0.99 | 0 |
| **JADE2** | 0 | 1.52104309984366 | 0.993 | 0.86 | 0 |
| **H2AFY** | 0 | -0.91084681664714 | 0.475 | 0.811 | 0 |
| **CXCL14** | 0 | -2.37258735417556 | 0.496 | 0.754 | 0 |
| **WNT8A** | 0 | -2.9806557379601 | 0.059 | 0.813 | 0 |
| **ECSCR** | 0 | -0.804975901797627 | 0.012 | 0.453 | 0 |
| **CXXC5** | 0 | -1.705312666344 | 0.799 | 0.996 | 0 |
| **DND1** | 0 | 1.10058397313804 | 0.576 | 0.114 | 0 |
| **PCDHB2** | 0 | -1.6086410304571 | 0.148 | 0.824 | 0 |
| **PCDHGB7** | 0 | -1.62184951006561 | 0.153 | 0.805 | 0 |
| **LARS** | 0 | 0.838318023909398 | 1 | 1 | 0 |
| **DPYSL3** | 0 | -1.93988105701539 | 0.629 | 0.947 | 0 |
| **JAKMIP2** | 0 | -0.83545752624917 | 0.02 | 0.444 | 0 |
| **AFAP1L1** | 0 | 0.83577088246936 | 0.548 | 0.178 | 0 |
| **PDGFRB** | 0 | -1.21961875464706 | 0.038 | 0.629 | 0 |

| | | | | | |
|---|---|---|---|---|---|
| **CDX1** | 0 | -1.57150221071127 | 0.697 | 0.851 | 0 |
| **ZNF300** | 0 | 0.855989882509197 | 0.748 | 0.288 | 0 |
| **GALNT10** | 0 | -0.946681912484287 | 0.52 | 0.82 | 0 |
| **ADAM19** | 0 | -1.42124777075735 | 0.791 | 0.973 | 0 |
| **CCNG1** | 0 | -1.00506576363749 | 0.999 | 0.999 | 0 |
| **WWC1** | 0 | -0.867250675873088 | 0.223 | 0.576 | 0 |
| **KCNMB1** | 0 | -1.11780521653139 | 0.028 | 0.375 | 0 |
| **STK10** | 0 | 0.838742710893637 | 0.911 | 0.717 | 0 |
| **ERGIC1** | 0 | -1.02258862166139 | 0.597 | 0.901 | 0 |
| **STC2** | 0 | 2.39800748094753 | 0.984 | 0.809 | 0 |
| **SFXN1** | 0 | -1.56605035940466 | 0.906 | 0.999 | 0 |
| **FGFR4** | 0 | 1.48920993358995 | 0.975 | 0.689 | 0 |
| **NSD1** | 0 | 1.07018723091641 | 1 | 0.988 | 0 |
| **DBN1** | 0 | -1.18761162383838 | 0.995 | 1 | 0 |
| **ZNF354C** | 0 | 0.889186986602147 | 0.551 | 0.013 | 0 |
| **GFPT2** | 0 | -1.70921869624991 | 0.617 | 0.967 | 0 |
| **FOXC1** | 0 | -2.00780633940854 | 0.05 | 0.529 | 0 |
| **SERPINB1** | 0 | 1.18686437048248 | 0.689 | 0.069 | 0 |
| **SERPINB9** | 0 | -1.46114507549869 | 0.861 | 0.966 | 0 |
| **SERPINB6** | 0 | 1.36761533152234 | 0.994 | 0.854 | 0 |
| **TUBB2A** | 0 | -2.48374581964846 | 0.943 | 0.999 | 0 |
| **TUBB2B** | 0 | -2.71978918728001 | 0.989 | 1 | 0 |
| **SLC22A23** | 0 | -0.839321520112742 | 0.041 | 0.391 | 0 |
| **DSP** | 0 | -1.01461059313962 | 0.984 | 0.999 | 0 |
| **JARID2** | 0 | 0.835904232395205 | 0.999 | 0.992 | 0 |
| **MYLIP** | 0 | 0.915061150718275 | 0.797 | 0.402 | 0 |
| **GMPR** | 0 | -1.38472464387883 | 0.072 | 0.528 | 0 |
| **CAP2** | 0 | -1.21935372039724 | 0.364 | 0.816 | 0 |
| **DEK** | 0 | -0.940598129241934 | 0.986 | 0.997 | 0 |
| **E2F3** | 0 | -0.930826181374633 | 0.832 | 0.972 | 0 |
| **MRS2** | 0 | 1.52212155591025 | 1 | 0.998 | 0 |
| **TRIM38** | 0 | 1.21646321168297 | 0.74 | 0.322 | 0 |
| **HIST1H1A** | 0 | 4.20311768456667 | 0.988 | 0.431 | 0 |
| **HIST1H2BB** | 0 | 2.38145926419254 | 0.765 | 0.083 | 0 |

| | | | | | |
|---|---|---|---|---|---|
| **GABBR1** | 0 | -1.35400516510775 | 0.601 | 0.916 | 0 |
| **ZFP57** | 0 | 2.5654185682425 | 0.995 | 0.303 | 0 |
| **TRIM26** | 0 | 1.48156182512425 | 0.98 | 0.82 | 0 |
| **NRM** | 0 | -1.06434883191058 | 0.409 | 0.829 | 0 |
| **FLOT1** | 0 | -1.37568976507291 | 0.888 | 0.996 | 0 |
| **IER3** | 0 | -1.68273090603313 | 0.384 | 0.766 | 0 |
| **DDAH2** | 0 | -1.69628556752665 | 0.716 | 0.984 | 0 |
| **AGPAT1** | 0 | -0.900712599130856 | 0.973 | 0.999 | 0 |
| **PBX2** | 0 | -0.97666477195998 | 0.985 | 0.999 | 0 |
| **GPSM3** | 0 | -0.855820884299828 | 0.393 | 0.74 | 0 |
| **CUTA** | 0 | -0.834352206782683 | 0.988 | 0.999 | 0 |
| **SYNGAP1** | 0 | -1.38779315113457 | 0.338 | 0.833 | 0 |
| **BAK1** | 0 | 0.896365701771066 | 0.999 | 0.988 | 0 |
| **ITPR3** | 0 | 1.09121151774105 | 0.906 | 0.599 | 0 |
| **ILRUN** | 0 | 1.68698780681377 | 1 | 0.996 | 0 |
| **SNRPC** | 0 | 1.03762686982236 | 1 | 1 | 0 |
| **ANKS1A** | 0 | -0.837435281856378 | 0.714 | 0.932 | 0 |
| **FKBP5** | 0 | -0.971514871212047 | 0.558 | 0.853 | 0 |
| **ZFAND3** | 0 | -0.898331160867531 | 0.991 | 1 | 0 |
| **TRERF1** | 0 | -1.07361330396094 | 0.046 | 0.555 | 0 |
| **PTK7** | 0 | -1.59337672953558 | 0.927 | 0.999 | 0 |
| **ZNF318** | 0 | 0.815628129716197 | 0.939 | 0.769 | 0 |
| **VEGFA** | 0 | 0.925718829940904 | 0.889 | 0.601 | 0 |
| **SLC35B2** | 0 | -1.23469995177218 | 0.687 | 0.973 | 0 |
| **AARS2** | 0 | 1.23486503120495 | 0.994 | 0.898 | 0 |
| **TNFRSF21** | 0 | -1.51170565112368 | 0.924 | 0.991 | 0 |
| **DPPA5** | 0 | 4.69735689761546 | 0.999 | 0.604 | 0 |
| **KHDC3L** | 0 | 3.64523539377173 | 0.99 | 0.424 | 0 |
| **DDX43** | 0 | 2.73470840991766 | 0.966 | 0.091 | 0 |
| **CGAS** | 0 | 1.26158442915 | 0.663 | 0.065 | 0 |
| **IRAK1BP1** | 0 | 0.947829421600304 | 0.48 | 0.015 | 0 |
| **ME1** | 0 | -0.994698313514859 | 0.304 | 0.725 | 0 |
| **AKIRIN2** | 0 | 1.21717018834923 | 1 | 0.997 | 0 |
| **PM20D2** | 0 | -0.84818317198611 | 0.246 | 0.628 | 0 |

| | | | | | |
|---|---|---|---|---|---|
| **UBE2J1** | 0 | -1.27426103853076 | 0.615 | 0.961 | 0 |
| **FAXC** | 0 | 1.15010059019887 | 0.785 | 0.349 | 0 |
| **ASCC3** | 0 | 1.1042905024099 | 0.998 | 0.947 | 0 |
| **LIN28B** | 0 | 1.45558747258107 | 1 | 0.994 | 0 |
| **SEC63** | 0 | 1.68826695271854 | 1 | 0.997 | 0 |
| **WASF1** | 0 | -2.03408734384348 | 0.335 | 0.949 | 0 |
| **SLC16A10** | 0 | 1.45416548969742 | 0.836 | 0.206 | 0 |
| **FYN** | 0 | -0.825507235004952 | 0.829 | 0.946 | 0 |
| **MARCKS** | 0 | -2.68452926464086 | 0.311 | 0.969 | 0 |
| **TSPYL4** | 0 | -0.856490673696896 | 0.219 | 0.649 | 0 |
| **FAM162B** | 0 | 1.48756380891188 | 0.759 | 0.055 | 0 |
| **MAN1A1** | 0 | -1.42304288778787 | 0.128 | 0.54 | 0 |
| **PKIB** | 0 | 1.26160889013518 | 0.687 | 0.08 | 0 |
| **SMPDL3A** | 0 | 0.860342422083186 | 0.566 | 0.096 | 0 |
| **ECHDC1** | 0 | -1.67358771435338 | 0.195 | 0.911 | 0 |
| **LAMA2** | 0 | -1.0390996540555 | 0.14 | 0.6 | 0 |
| **SAMD3** | 0 | -1.46834326570983 | 0.004 | 0.443 | 0 |
| **ENPP1** | 0 | -1.21873713825044 | 0.553 | 0.903 | 0 |
| **SGK1** | 0 | 2.05331756657421 | 0.985 | 0.663 | 0 |
| **PERP** | 0 | -0.932887461746416 | 0.987 | 0.996 | 0 |
| **CITED2** | 0 | -1.90774303170637 | 0.501 | 0.951 | 0 |
| **AIG1** | 0 | -1.02579158688637 | 0.127 | 0.591 | 0 |
| **FUCA2** | 0 | -0.926563444009294 | 0.749 | 0.906 | 0 |
| **PHACTR2** | 0 | -1.11346315310953 | 0.582 | 0.906 | 0 |
| **UTRN** | 0 | -0.983154081897388 | 0.588 | 0.889 | 0 |
| **UST** | 0 | -1.40188553069406 | 0.475 | 0.897 | 0 |
| **PCMT1** | 0 | 1.05025687640204 | 1 | 1 | 0 |
| **AKAP12** | 0 | 1.66371916837761 | 1 | 0.979 | 0 |
| **RMND1** | 0 | 1.87568542155717 | 0.999 | 0.931 | 0 |
| **ARMT1** | 0 | 1.7339172854351 | 1 | 0.969 | 0 |
| **ESR1** | 0 | 1.76675767848531 | 0.999 | 0.91 | 0 |
| **CNKSR3** | 0 | -1.63702469974223 | 0.101 | 0.618 | 0 |
| **TCP1** | 0 | 1.51821363192316 | 1 | 0.977 | 0 |
| **IGF2R** | 0 | -1.67057395981599 | 0.505 | 0.971 | 0 |

| | | | | | |
|---|---|---|---|---|---|
| **MAP3K4** | 0 | -0.891872468797935 | 0.956 | 0.995 | 0 |
| **AGPAT4** | 0 | -1.25293161231481 | 0.482 | 0.881 | 0 |
| **TBXT** | 0 | -3.0540663089413 | 0.215 | 0.872 | 0 |
| **SFT2D1** | 0 | 0.854604207044181 | 0.972 | 0.774 | 0 |
| **RNASET2** | 0 | 0.945184491444615 | 0.923 | 0.626 | 0 |
| **AFDN** | 0 | -1.17689619144022 | 0.963 | 1 | 0 |
| **DACT2** | 0 | 0.846098312933225 | 0.455 | 0.051 | 0 |
| **C6orf120** | 0 | -1.46914527616144 | 0.533 | 0.966 | 0 |
| **PHF10** | 0 | -1.44836460338835 | 0.942 | 0.999 | 0 |
| **FAM120B** | 0 | -0.896901592055283 | 0.296 | 0.717 | 0 |
| **FAM20C** | 0 | -1.85447724219711 | 0.219 | 0.842 | 0 |
| **PDGFA** | 0 | 1.21000945991219 | 0.997 | 0.883 | 0 |
| **PRKAR1B** | 0 | -0.84997153852505 | 0.481 | 0.775 | 0 |
| **C7orf50** | 0 | -1.35089613488574 | 0.836 | 0.986 | 0 |
| **ELFN1** | 0 | -1.02867714687105 | 0.193 | 0.577 | 0 |
| **TTYH3** | 0 | -0.886892308854562 | 0.99 | 0.999 | 0 |
| **AIMP2** | 0 | -1.00950360884967 | 0.961 | 0.999 | 0 |
| **EIF2AK1** | 0 | -0.950246179647469 | 0.97 | 1 | 0 |
| **ETV1** | 0 | 0.857440263127363 | 0.907 | 0.735 | 0 |
| **TSPAN13** | 0 | 1.17796300969234 | 0.983 | 0.761 | 0 |
| **ITGB8** | 0 | -1.29437123772851 | 0.026 | 0.464 | 0 |
| **GPNMB** | 0 | 0.876450758463655 | 0.582 | 0.157 | 0 |
| **CCDC126** | 0 | 1.24025356436197 | 0.863 | 0.435 | 0 |
| **MPP6** | 0 | -1.05959423986689 | 0.928 | 0.993 | 0 |
| **CYCS** | 0 | 0.933422141913072 | 1 | 0.869 | 0 |
| **NFE2L3** | 0 | 0.989860556772847 | 0.904 | 0.589 | 0 |
| **CBX3** | 0 | 0.819170499594951 | 1 | 0.973 | 0 |
| **SKAP2** | 0 | -1.62826097192166 | 0.075 | 0.705 | 0 |
| **HOXA1** | 0 | -1.82871996632042 | 0.236 | 0.745 | 0 |
| **HOXA7** | 0 | -0.85832356302856 | 0.049 | 0.543 | 0 |
| **EVX1** | 0 | -1.12744976311776 | 0.022 | 0.37 | 0 |
| **DPY19L1** | 0 | 1.007228314884 | 0.963 | 0.732 | 0 |
| **HERPUD2** | 0 | -1.08618074484182 | 0.5 | 0.895 | 0 |
| **STARD3NL** | 0 | -0.993146159244269 | 0.775 | 0.971 | 0 |

| | | | | | |
|---|---|---|---|---|---|
| **MPLKIP** | 0 | -0.855311664396437 | 0.733 | 0.954 | 0 |
| **DBNL** | 0 | -1.17792404621165 | 0.808 | 0.99 | 0 |
| **AEBP1** | 0 | -1.16865314446277 | 0.583 | 0.91 | 0 |
| **MYL7** | 0 | -2.48724174388676 | 0.133 | 0.713 | 0 |
| **ZMIZ2** | 0 | -0.93848683423888 | 0.764 | 0.947 | 0 |
| **PURB** | 0 | -0.802110874483231 | 0.817 | 0.977 | 0 |
| **TBRG4** | 0 | 0.898786560822108 | 1 | 0.996 | 0 |
| **HUS1** | 0 | 1.50135017634277 | 1 | 0.977 | 0 |
| **SUN3** | 0 | 1.69898788744793 | 0.804 | 0.029 | 0 |
| **UPP1** | 0 | 2.61319159878128 | 1 | 0.811 | 0 |
| **PSPH** | 0 | 1.12830918878279 | 0.993 | 0.943 | 0 |
| **CHCHD2** | 0 | 1.22091678848316 | 1 | 1 | 0 |
| **ZNF727** | 0 | 0.935838969065578 | 0.523 | 0.011 | 0 |
| **ZNF736** | 0 | 1.26311754205218 | 0.736 | 0.031 | 0 |
| **GALNT17** | 0 | -1.51115082394517 | 0.105 | 0.649 | 0 |
| **TBL2** | 0 | -1.41189150329009 | 0.646 | 0.955 | 0 |
| **LIMK1** | 0 | -1.07189080509769 | 0.925 | 0.995 | 0 |
| **CLIP2** | 0 | -0.821565185815384 | 0.31 | 0.64 | 0 |
| **GTF2I** | 0 | -0.861965696417513 | 0.921 | 0.996 | 0 |
| **HIP1** | 0 | -1.43199340476619 | 0.541 | 0.942 | 0 |
| **CCL26** | 0 | -0.871188208729896 | 0.043 | 0.458 | 0 |
| **RHBDD2** | 0 | -0.981322818165044 | 0.703 | 0.939 | 0 |
| **GNAI1** | 0 | -1.16815259236142 | 0.381 | 0.725 | 0 |
| **CDK6** | 0 | -1.14605249951289 | 0.471 | 0.887 | 0 |
| **COL1A2** | 0 | -1.69587213341043 | 0.783 | 0.991 | 0 |
| **NPTX2** | 0 | 1.69313255737234 | 0.757 | 0.033 | 0 |
| **ARPC1B** | 0 | 0.9709116218133 | 0.998 | 0.917 | 0 |
| **MUC3A** | 0 | -1.10462225976535 | 0.107 | 0.485 | 0 |
| **SERPINE1** | 0 | 0.912337297010569 | 0.803 | 0.425 | 0 |
| **AP1S1** | 0 | 1.35786994487884 | 0.994 | 0.842 | 0 |
| **COL26A1** | 0 | -1.80134251967993 | 0.613 | 0.923 | 0 |
| **CUX1** | 0 | -1.92378706653702 | 0.501 | 0.976 | 0 |
| **ORAI2** | 0 | -0.938409330234305 | 0.265 | 0.69 | 0 |
| **EFCAB10** | 0 | 1.0409441206061 | 0.653 | 0.102 | 0 |

| | | | | | |
|---|---|---|---|---|---|
| **PRKAR2B** | 0 | -0.873372995395686 | 0.303 | 0.638 | 0 |
| **DNAJB9** | 0 | 1.05621716814693 | 0.93 | 0.82 | 0 |
| **DOCK4** | 0 | 1.30676042297501 | 0.82 | 0.266 | 0 |
| **TES** | 0 | 1.03420519737487 | 0.986 | 0.871 | 0 |
| **CAV2** | 0 | -0.881804087615389 | 0.16 | 0.512 | 0 |
| **CAV1** | 0 | -2.22690276381893 | 0.149 | 0.785 | 0 |
| **TSPAN12** | 0 | -1.1355101070973 | 0.14 | 0.642 | 0 |
| **PTPRZ1** | 0 | -2.72459931347559 | 0.192 | 0.761 | 0 |
| **AASS** | 0 | 1.20410343489734 | 0.957 | 0.786 | 0 |
| **WASL** | 0 | 1.25273211054967 | 0.999 | 0.988 | 0 |
| **HYAL4** | 0 | 1.57068025587918 | 0.794 | 0.046 | 0 |
| **FLNC** | 0 | -1.21576363433448 | 0.931 | 0.993 | 0 |
| **KCP** | 0 | -1.11087744757533 | 0.077 | 0.554 | 0 |
| **SMO** | 0 | -0.96423280556227 | 0.594 | 0.869 | 0 |
| **KLHDC10** | 0 | -1.12795053522722 | 0.166 | 0.732 | 0 |
| **PODXL** | 0 | -2.58683425518767 | 0.821 | 0.968 | 0 |
| **AKR1B1** | 0 | -0.899938928782344 | 0.954 | 0.996 | 0 |
| **CALD1** | 0 | -1.88827214280858 | 0.985 | 0.996 | 0 |
| **ZC3HAV1** | 0 | 0.895214549857456 | 1 | 0.991 | 0 |
| **HIPK2** | 0 | -1.66747539735418 | 0.387 | 0.912 | 0 |
| **SLC37A3** | 0 | 0.955020419122402 | 0.993 | 0.924 | 0 |
| **DENND11** | 0 | -0.871984479023895 | 0.371 | 0.741 | 0 |
| **TCAF1** | 0 | -1.70981845048782 | 0.52 | 0.962 | 0 |
| **CNTNAP2** | 0 | -3.93012856626235 | 0.421 | 0.912 | 0 |
| **EZH2** | 0 | -1.04443797834682 | 0.894 | 0.986 | 0 |
| **ZNF398** | 0 | 1.33957990328834 | 0.994 | 0.833 | 0 |
| **REPIN1** | 0 | -1.15689918320355 | 0.176 | 0.71 | 0 |
| **ABCB8** | 0 | 1.1519593722576 | 0.984 | 0.799 | 0 |
| **AGAP3** | 0 | -0.824998178618156 | 0.254 | 0.669 | 0 |
| **TDRP** | 0 | 0.8549319175798 | 0.971 | 0.858 | 0 |
| **AGPAT5** | 0 | -1.60919501231022 | 0.774 | 0.995 | 0 |
| **PPP1R3B** | 0 | -2.7266162145589 | 0.538 | 0.99 | 0 |
| **XKR6** | 0 | -0.848099309227573 | 0.381 | 0.751 | 0 |
| **DLC1** | 0 | -1.46519520571346 | 0.411 | 0.686 | 0 |

| | | | | | |
|---|---|---|---|---|---|
| **ZDHHC2** | 0 | -1.61646649493653 | 0.064 | 0.732 | 0 |
| **SH2D4A** | 0 | 1.19943493746818 | 0.828 | 0.359 | 0 |
| **FGF17** | 0 | -3.80630220255183 | 0.227 | 0.719 | 0 |
| **SLC39A14** | 0 | 0.894755124689695 | 0.976 | 0.839 | 0 |
| **CHMP7** | 0 | -0.908771294368119 | 0.586 | 0.876 | 0 |
| **LOXL2** | 0 | -2.16364702477265 | 0.684 | 0.995 | 0 |
| **ENTPD4** | 0 | -0.838432819996169 | 0.443 | 0.781 | 0 |
| **STC1** | 0 | -1.45689590924028 | 0.006 | 0.463 | 0 |
| **BNIP3L** | 0 | -1.14124237736357 | 0.949 | 0.999 | 0 |
| **DPYSL2** | 0 | -1.78826033909973 | 0.897 | 0.999 | 0 |
| **CLU** | 0 | -1.33120301145409 | 0.888 | 0.984 | 0 |
| **RNF122** | 0 | -1.06287449571332 | 0.296 | 0.681 | 0 |
| **ZNF703** | 0 | -1.35281169737424 | 0.543 | 0.815 | 0 |
| **EIF4EBP1** | 0 | 1.12899603089844 | 1 | 0.992 | 0 |
| **ASH2L** | 0 | 1.35183922760317 | 0.992 | 0.92 | 0 |
| **DDHD2** | 0 | -0.800859625367689 | 0.408 | 0.763 | 0 |
| **FGFR1** | 0 | -1.21232540587265 | 0.992 | 1 | 0 |
| **HGSNAT** | 0 | -1.00545456915356 | 0.129 | 0.637 | 0 |
| **SNAI2** | 0 | -1.66774357742473 | 0.014 | 0.374 | 0 |
| **TGS1** | 0 | 1.29768157609915 | 1 | 0.99 | 0 |
| **SDCBP** | 0 | 0.820882501721474 | 0.999 | 0.979 | 0 |
| **NSMAF** | 0 | 1.64532290319601 | 0.999 | 0.913 | 0 |
| **CHD7** | 0 | -2.76277904840332 | 0.317 | 0.966 | 0 |
| **ASPH** | 0 | -0.903835970136685 | 0.827 | 0.965 | 0 |
| **MTFR1** | 0 | 0.905812960286485 | 0.985 | 0.868 | 0 |
| **ARFGEF1** | 0 | 0.980793966736257 | 0.987 | 0.86 | 0 |
| **PREX2** | 0 | -0.964191567774426 | 0.254 | 0.611 | 0 |
| **PRDM14** | 0 | 1.42873467746888 | 0.943 | 0.698 | 0 |
| **STAU2** | 0 | -0.987225931884332 | 0.972 | 1 | 0 |
| **CRISPLD1** | 0 | -1.14301387503532 | 0.504 | 0.812 | 0 |
| **PKIA** | 0 | -0.906236321598474 | 0.007 | 0.483 | 0 |
| **FABP5** | 0 | 1.99095870071258 | 0.997 | 0.675 | 0 |
| **E2F5** | 0 | -1.71956984881887 | 0.521 | 0.969 | 0 |
| **CALB1** | 0 | 1.1846653857299 | 0.724 | 0.2 | 0 |

| | | | | | |
|---|---|---|---|---|---|
| **ESRP1** | 0 | 1.56021297925258 | 1 | 0.783 | 0 |
| **AZIN1** | 0 | -0.826854385984711 | 0.978 | 0.994 | 0 |
| **CTHRC1** | 0 | 1.01731688203702 | 0.576 | 0.051 | 0 |
| **ANGPT1** | 0 | 0.990680658913885 | 0.702 | 0.155 | 0 |
| **DEPTOR** | 0 | 1.77928177100402 | 0.941 | 0.302 | 0 |
| **ZHX2** | 0 | -1.52423956001085 | 0.089 | 0.768 | 0 |
| **ATAD2** | 0 | -0.877357387176476 | 0.3 | 0.681 | 0 |
| **WDYHV1** | 0 | -0.904529546934402 | 0.226 | 0.668 | 0 |
| **KHDRBS3** | 0 | -1.22554893288537 | 0.688 | 0.89 | 0 |
| **NRBP2** | 0 | 1.15964202120677 | 0.726 | 0.22 | 0 |
| **EPPK1** | 0 | 1.41894966633853 | 0.981 | 0.72 | 0 |
| **SMARCA2** | 0 | 1.9498697889207 | 0.986 | 0.591 | 0 |
| **ERMP1** | 0 | -1.17111762541396 | 0.351 | 0.822 | 0 |
| **UHRF2** | 0 | -1.42404685153288 | 0.204 | 0.833 | 0 |
| **GLDC** | 0 | 1.77997004569349 | 0.989 | 0.562 | 0 |
| **CER1** | 0 | -6.36746175620922 | 0.137 | 0.819 | 0 |
| **FOCAD** | 0 | -0.833686666975444 | 0.624 | 0.864 | 0 |
| **CAAP1** | 0 | 0.866515681979265 | 0.875 | 0.571 | 0 |
| **AQP3** | 0 | 0.8303181655816 | 0.499 | 0.155 | 0 |
| **UBAP2** | 0 | 1.12117230905127 | 0.999 | 0.971 | 0 |
| **MYORG** | 0 | -1.41132140388522 | 0.166 | 0.752 | 0 |
| **FAM219A** | 0 | -1.18898823991661 | 0.305 | 0.797 | 0 |
| **CNTFR** | 0 | -2.41306643800603 | 0.422 | 0.816 | 0 |
| **GBA2** | 0 | -0.844384669883193 | 0.746 | 0.948 | 0 |
| **GLIPR2** | 0 | -2.36363189997531 | 0.275 | 0.894 | 0 |
| **ALDH1B1** | 0 | -0.84428080189913 | 0.682 | 0.908 | 0 |
| **IGFBPL1** | 0 | -2.11575610509161 | 0.099 | 0.829 | 0 |
| **ANKRD18A** | 0 | -1.02272556670928 | 0.128 | 0.567 | 0 |
| **FAM189A2** | 0 | -1.07287398961575 | 0.081 | 0.47 | 0 |
| **PTAR1** | 0 | 0.920709212493225 | 0.998 | 0.965 | 0 |
| **PCSK5** | 0 | 1.32874789910594 | 0.801 | 0.161 | 0 |
| **PRUNE2** | 0 | 1.38053571795241 | 0.932 | 0.483 | 0 |
| **PSAT1** | 0 | 1.19315578761131 | 1 | 0.999 | 0 |
| **C9orf64** | 0 | 1.25159120953103 | 0.698 | 0.027 | 0 |

| | | | | | |
|---|---|---|---|---|---|
| **CKS2** | 0 | 0.998249245164553 | 0.992 | 0.944 | 0 |
| **SECISBP2** | 0 | -1.02853028824571 | 0.789 | 0.978 | 0 |
| **ROR2** | 0 | -1.78239977915252 | 0.476 | 0.85 | 0 |
| **BARX1** | 0 | 1.16173121822658 | 0.752 | 0.197 | 0 |
| **PTCH1** | 0 | -1.20849204210068 | 0.494 | 0.876 | 0 |
| **ZNF367** | 0 | -0.962473006977711 | 0.24 | 0.634 | 0 |
| **CTSV** | 0 | -1.06603947316617 | 0.835 | 0.961 | 0 |
| **XPA** | 0 | -0.86124948635923 | 0.314 | 0.7 | 0 |
| **ANP32B** | 0 | -1.47573948775881 | 0.891 | 0.999 | 0 |
| **TBC1D2** | 0 | -1.16341494744972 | 0.069 | 0.453 | 0 |
| **TGFBR1** | 0 | -1.14123297436301 | 0.966 | 1 | 0 |
| **ABCA1** | 0 | 1.62295216436883 | 0.959 | 0.549 | 0 |
| **KLF4** | 0 | 1.39748047847621 | 0.668 | 0.209 | 0 |
| **CTNNAL1** | 0 | -0.853719076357364 | 0.965 | 0.998 | 0 |
| **LPAR1** | 0 | 0.892788264494291 | 0.659 | 0.292 | 0 |
| **ECPAS** | 0 | 0.800349341517448 | 0.999 | 0.994 | 0 |
| **ZNF483** | 0 | 1.23881147393139 | 0.888 | 0.502 | 0 |
| **PTGR1** | 0 | 0.914813689396346 | 0.983 | 0.811 | 0 |
| **UGCG** | 0 | 1.20314892250611 | 0.999 | 0.985 | 0 |
| **ZNF618** | 0 | -1.03669379227706 | 0.389 | 0.771 | 0 |
| **AKNA** | 0 | 0.888811674576578 | 0.943 | 0.738 | 0 |
| **TNC** | 0 | -3.60514055630863 | 0.084 | 0.865 | 0 |
| **PAPPA** | 0 | -1.24183771319705 | 0.023 | 0.55 | 0 |
| **CNTRL** | 0 | -1.27625523537391 | 0.192 | 0.763 | 0 |
| **GSN** | 0 | -2.61988582281435 | 0.125 | 0.939 | 0 |
| **STOM** | 0 | 2.53392827706886 | 0.997 | 0.695 | 0 |
| **PDCL** | 0 | 0.930715155955873 | 0.998 | 0.984 | 0 |
| **NR6A1** | 0 | -1.0904235262717 | 0.888 | 0.97 | 0 |
| **HSPA5** | 0 | 0.977621458522065 | 1 | 0.964 | 0 |
| **ENG** | 0 | -0.806205918592433 | 0.097 | 0.465 | 0 |
| **ST6GALNAC6** | 0 | -0.863089309816253 | 0.402 | 0.782 | 0 |
| **CIZ1** | 0 | -1.11500607582237 | 0.355 | 0.826 | 0 |
| **SPTAN1** | 0 | -0.8132897734091 | 0.988 | 0.999 | 0 |
| **LRRC8A** | 0 | -0.817258459694777 | 0.989 | 0.999 | 0 |

| | | | | | |
|---|---|---|---|---|---|
| **NCS1** | 0 | -0.855430025897329 | 0.708 | 0.914 | 0 |
| **ABL1** | 0 | -0.944484993122231 | 0.96 | 0.999 | 0 |
| **UCK1** | 0 | -0.859766264318623 | 0.805 | 0.952 | 0 |
| **CACFD1** | 0 | -0.906823344208478 | 0.297 | 0.705 | 0 |
| **VAV2** | 0 | 1.05917675085886 | 0.953 | 0.769 | 0 |
| **WDR5** | 0 | 0.815555982083363 | 0.979 | 0.845 | 0 |
| **COL5A1** | 0 | -2.71859890127873 | 0.503 | 0.912 | 0 |
| **OLFM1** | 0 | -2.1327431180187 | 0.296 | 0.859 | 0 |
| **TMEM250** | 0 | -0.863058565121455 | 0.824 | 0.984 | 0 |
| **AGPAT2** | 0 | 1.1742612715754 | 0.984 | 0.917 | 0 |
| **NRARP** | 0 | -1.06707563127011 | 0.268 | 0.729 | 0 |
| **ZMYND11** | 0 | -0.823920076081121 | 0.689 | 0.921 | 0 |
| **NET1** | 0 | -1.12641787053506 | 0.84 | 0.963 | 0 |
| **GDI2** | 0 | -0.854482246644062 | 0.973 | 0.998 | 0 |
| **SEPHS1** | 0 | -1.07390047640832 | 0.987 | 1 | 0 |
| **FRMD4A** | 0 | -1.29826500008277 | 0.166 | 0.7 | 0 |
| **OLAH** | 0 | 1.48531264612692 | 0.482 | 0.045 | 0 |
| **FAM171A1** | 0 | -1.21149277009146 | 0.227 | 0.777 | 0 |
| **VIM** | 0 | -3.86047545841298 | 0.608 | 0.991 | 0 |
| **NEBL** | 0 | -1.91665855322496 | 0.083 | 0.849 | 0 |
| **DNAJC1** | 0 | -0.980561467961551 | 0.443 | 0.812 | 0 |
| **MSRB2** | 0 | -1.10793363341755 | 0.535 | 0.903 | 0 |
| **ZEB1** | 0 | -0.969979353318451 | 0.018 | 0.465 | 0 |
| **RET** | 0 | -1.82902215803082 | 0.233 | 0.802 | 0 |
| **RASGEF1A** | 0 | 1.41465622804717 | 0.953 | 0.581 | 0 |
| **SGMS1** | 0 | 0.808684218058923 | 0.745 | 0.373 | 0 |
| **1,00 DKK** | 0 | -3.31122533210933 | 0.031 | 0.44 | 0 |
| **SLC16A9** | 0 | 0.843456754200807 | 0.691 | 0.344 | 0 |
| **CCDC6** | 0 | -0.807411387869606 | 0.942 | 0.993 | 0 |
| **NRBF2** | 0 | 1.0364712646321 | 0.999 | 0.964 | 0 |
| **REEP3** | 0 | -1.21585729758879 | 0.693 | 0.966 | 0 |
| **SLC25A16** | 0 | 2.45689990925315 | 0.999 | 0.714 | 0 |
| **TET1** | 0 | 1.4600806400282 | 0.999 | 0.995 | 0 |
| **DDX21** | 0 | 0.959344732328289 | 1 | 1 | 0 |

| | | | | | |
|---|---|---|---|---|---|
| **HK1** | 0 | -1.60645454880266 | 0.895 | 0.997 | 0 |
| **COL13A1** | 0 | -1.46878727669682 | 0.013 | 0.475 | 0 |
| **H2AFY2** | 0 | -2.91582142331724 | 0.453 | 0.981 | 0 |
| **PCBD1** | 0 | -1.1749887212354 | 0.685 | 0.929 | 0 |
| **DDIT4** | 0 | 2.36561825631756 | 0.956 | 0.694 | 0 |
| **P4HA1** | 0 | -0.942677637159877 | 0.995 | 1 | 0 |
| **PLAU** | 0 | 1.94392045246064 | 0.821 | 0.503 | 0 |
| **VCL** | 0 | -1.67112283567308 | 0.971 | 0.999 | 0 |
| **AP3M1** | 0 | -0.81739717135332 | 0.737 | 0.936 | 0 |
| **ZCCHC24** | 0 | -1.98865709350088 | 0.21 | 0.89 | 0 |
| **PRXL2A** | 0 | -1.17123628726193 | 0.82 | 0.976 | 0 |
| **GLUD1** | 0 | 0.914704912196818 | 1 | 0.999 | 0 |
| **LIPA** | 0 | -1.45903521739 | 0.34 | 0.86 | 0 |
| **EXOC6** | 0 | -1.14006777650464 | 0.139 | 0.698 | 0 |
| **CYP26A1** | 0 | -4.03429781178673 | 0.11 | 0.666 | 0 |
| **LGI1** | 0 | -0.970500116711287 | 0.012 | 0.351 | 0 |
| **PDLIM1** | 0 | 0.983358215739369 | 0.999 | 0.957 | 0 |
| **SORBS1** | 0 | -1.07855791228246 | 0.62 | 0.838 | 0 |
| **ZNF518A** | 0 | 0.967576812037217 | 0.995 | 0.931 | 0 |
| **PIK3AP1** | 0 | 1.78384442673497 | 0.827 | 0.1 | 0 |
| **FRAT2** | 0 | 1.45914808062718 | 0.996 | 0.955 | 0 |
| **MARVELD1** | 0 | -1.00954547347671 | 0.429 | 0.837 | 0 |
| **BLOC1S2** | 0 | 1.03431298554955 | 0.989 | 0.841 | 0 |
| **LZTS2** | 0 | -1.0308921521218 | 0.628 | 0.925 | 0 |
| **NFKB2** | 0 | 2.89400431037545 | 0.997 | 0.75 | 0 |
| **MFSD13A** | 0 | 0.998329775978416 | 0.703 | 0.212 | 0 |
| **INA** | 0 | 1.69117100292338 | 0.96 | 0.525 | 0 |
| **PCGF6** | 0 | 0.896597129417116 | 0.962 | 0.758 | 0 |
| **GSTO2** | 0 | 1.11974255668946 | 0.676 | 0.086 | 0 |
| **XPNPEP1** | 0 | -0.921428630649682 | 0.62 | 0.907 | 0 |
| **ADD3** | 0 | -1.01031305479357 | 0.878 | 0.98 | 0 |
| **DUSP5** | 0 | 1.17643707927849 | 0.99 | 0.886 | 0 |
| **SMC3** | 0 | -0.882372888633712 | 0.986 | 1 | 0 |
| **RGS10** | 0 | 1.05490979985564 | 0.829 | 0.366 | 0 |

| | | | | | |
|---|---|---|---|---|---|
| **INPP5F** | 0 | 0.815690766761766 | 0.999 | 0.996 | 0 |
| **FGFR2** | 0 | 0.854671656210215 | 0.986 | 0.905 | 0 |
| **OAT** | 0 | 1.20006455185235 | 0.976 | 0.761 | 0 |
| **NKX1-2** | 0 | -1.24576443458888 | 0.761 | 0.885 | 0 |
| **UROS** | 0 | -1.06740839712545 | 0.503 | 0.889 | 0 |
| **DHX32** | 0 | -0.917478784744609 | 0.262 | 0.695 | 0 |
| **UTF1** | 0 | 3.16576751963795 | 0.995 | 0.371 | 0 |
| **VENTX** | 0 | 1.18307224270843 | 0.902 | 0.372 | 0 |
| **IFITM2** | 0 | -1.02984988752055 | 0.953 | 0.987 | 0 |
| **IFITM3** | 0 | -1.13775807995859 | 0.996 | 1 | 0 |
| **PKP3** | 0 | 1.04692464908521 | 0.907 | 0.403 | 0 |
| **TSPAN4** | 0 | -1.25306192775278 | 0.502 | 0.811 | 0 |
| **PHLDA2** | 0 | 0.928885011262498 | 0.955 | 0.796 | 0 |
| **TRIM6** | 0 | 1.37972207935454 | 0.817 | 0.195 | 0 |
| **TRIM5** | 0 | -0.95424765295673 | 0.407 | 0.817 | 0 |
| **DCHS1** | 0 | -1.08919235462072 | 0.364 | 0.766 | 0 |
| **OVCH2** | 0 | -1.13051987859278 | 0.013 | 0.544 | 0 |
| **ST5** | 0 | -1.18702221174745 | 0.8 | 0.975 | 0 |
| **NRIP3** | 0 | 0.847693680757259 | 0.54 | 0.062 | 0 |
| **USP47** | 0 | -0.96579401472991 | 0.663 | 0.925 | 0 |
| **3,00 DKK** | 0 | -1.96444549690747 | 0.192 | 0.883 | 0 |
| **PARVA** | 0 | -1.18591482704003 | 0.611 | 0.948 | 0 |
| **PDE3B** | 0 | -1.13980086683003 | 0.198 | 0.715 | 0 |
| **NAV2** | 0 | -1.55250523683267 | 0.564 | 0.916 | 0 |
| **FANCF** | 0 | -0.967151327515879 | 0.291 | 0.755 | 0 |
| **RCN1** | 0 | 0.812641630527183 | 1 | 0.997 | 0 |
| **TCP11L1** | 0 | -0.872492048900147 | 0.509 | 0.785 | 0 |
| **TRIM44** | 0 | -0.917265083477273 | 0.807 | 0.978 | 0 |
| **TSPAN18** | 0 | -1.66273474531444 | 0.764 | 0.928 | 0 |
| **TP53I11** | 0 | -1.81520002312108 | 0.252 | 0.902 | 0 |
| **CREB3L1** | 0 | 1.03911589953005 | 0.803 | 0.387 | 0 |
| **LRP4** | 0 | 1.07821469779419 | 0.997 | 0.936 | 0 |
| **ACP2** | 0 | -1.03843634776914 | 0.392 | 0.808 | 0 |
| **APLNR** | 0 | -4.43359233490643 | 0.133 | 0.746 | 0 |

| | | | | | |
|---|---|---|---|---|---|
| **SERPING1** | 0 | -1.09238935911499 | 0.848 | 0.979 | 0 |
| **TMX2** | 0 | 0.889918192619637 | 1 | 0.998 | 0 |
| **SELENOH** | 0 | -0.912374410518872 | 1 | 1 | 0 |
| **FAM111B** | 0 | -1.61145832301611 | 0.137 | 0.674 | 0 |
| **TMEM132A** | 0 | -0.887938001548785 | 0.275 | 0.71 | 0 |
| **TMEM216** | 0 | -0.98071458593611 | 0.319 | 0.771 | 0 |
| **FADS2** | 0 | -1.76312793425809 | 0.996 | 1 | 0 |
| **FADS1** | 0 | -1.92326443375407 | 0.945 | 0.999 | 0 |
| **FADS3** | 0 | 1.10615676874992 | 0.826 | 0.334 | 0 |
| **ASRGL1** | 0 | 2.18023459780539 | 1 | 0.802 | 0 |
| **EEF1G** | 0 | 0.917992680116319 | 1 | 1 | 0 |
| **SLC3A2** | 0 | 0.930423172598785 | 1 | 1 | 0 |
| **LGALS12** | 0 | -0.829187725028591 | 0.067 | 0.41 | 0 |
| **RTN3** | 0 | -0.997543386828863 | 0.978 | 0.997 | 0 |
| **RCOR2** | 0 | -1.10550526065356 | 0.611 | 0.921 | 0 |
| **VEGFB** | 0 | -1.12201341396987 | 0.636 | 0.938 | 0 |
| **NRXN2** | 0 | 1.09739912203397 | 0.699 | 0.166 | 0 |
| **RASGRP2** | 0 | 1.30859405388405 | 0.838 | 0.399 | 0 |
| **TM7SF2** | 0 | -1.53749152602281 | 0.554 | 0.862 | 0 |
| **EFEMP2** | 0 | -1.09199865459785 | 0.145 | 0.708 | 0 |
| **FOSL1** | 0 | 1.18697164489675 | 0.572 | 0.199 | 0 |
| **C11orf68** | 0 | -0.989533856552335 | 0.338 | 0.789 | 0 |
| **PACS1** | 0 | -1.26845796273884 | 0.552 | 0.948 | 0 |
| **GRK2** | 0 | -0.831081412339661 | 0.889 | 0.987 | 0 |
| **NDUFV1** | 0 | 1.08065639645605 | 1 | 1 | 0 |
| **CPT1A** | 0 | 1.44319475558359 | 0.95 | 0.499 | 0 |
| **DHCR7** | 0 | -0.935352957922302 | 0.987 | 0.99 | 0 |
| **INPPL1** | 0 | -0.896519590223035 | 0.889 | 0.991 | 0 |
| **ARAP1** | 0 | -1.0306785753203 | 0.709 | 0.945 | 0 |
| **RAB6A** | 0 | 1.11949655717791 | 1 | 0.992 | 0 |
| **PPME1** | 0 | -1.03720628170183 | 0.563 | 0.886 | 0 |
| **KCNE3** | 0 | 0.901831286327256 | 0.623 | 0.207 | 0 |
| **TSKU** | 0 | -0.953557153183977 | 0.983 | 0.997 | 0 |
| **TENM4** | 0 | -1.74556521740062 | 0.077 | 0.734 | 0 |

| | | | | | |
|---|---|---|---|---|---|
| **PICALM** | 0 | 0.860951110420276 | 1 | 0.998 | 0 |
| **FUT4** | 0 | 0.833298356685021 | 0.836 | 0.561 | 0 |
| **AMOTL1** | 0 | -2.80137369395035 | 0.581 | 0.991 | 0 |
| **SESN3** | 0 | 1.00018838121014 | 0.975 | 0.885 | 0 |
| **YAP1** | 0 | -1.08254835569567 | 0.979 | 1 | 0 |
| **TMEM123** | 0 | -0.834972466309506 | 0.877 | 0.963 | 0 |
| **SLC35F2** | 0 | 0.951649244189116 | 0.979 | 0.834 | 0 |
| **RDX** | 0 | -1.1795124037405 | 0.497 | 0.899 | 0 |
| **USP28** | 0 | 1.20650692209584 | 1 | 0.981 | 0 |
| **CADM1** | 0 | -1.15132992639222 | 0.407 | 0.749 | 0 |
| **TAGLN** | 0 | -2.26844079599024 | 0.89 | 0.992 | 0 |
| **KMT2A** | 0 | -0.879678986200052 | 0.673 | 0.923 | 0 |
| **BCL9L** | 0 | 0.918891935212457 | 0.998 | 0.995 | 0 |
| **FOXR1** | 0 | 1.19921613812349 | 0.638 | 0.008 | 0 |
| **HYOU1** | 0 | 1.12369405437928 | 1 | 1 | 0 |
| **H2AFX** | 0 | -1.06999508788676 | 0.968 | 0.995 | 0 |
| **THY1** | 0 | 1.27075393539854 | 0.98 | 0.934 | 0 |
| **OAF** | 0 | -1.16726583038605 | 0.022 | 0.497 | 0 |
| **UBASH3B** | 0 | 0.899475228095381 | 0.814 | 0.48 | 0 |
| **GRAMD1B** | 0 | -1.01844554170567 | 0.068 | 0.555 | 0 |
| **VWA5A** | 0 | -1.08726460383553 | 0.281 | 0.67 | 0 |
| **HYLS1** | 0 | 1.53903493176071 | 0.762 | 0.123 | 0 |
| **FAM118B** | 0 | -1.11556192180436 | 0.573 | 0.935 | 0 |
| **ETS1** | 0 | -1.45307950787585 | 0.464 | 0.774 | 0 |
| **TEAD4** | 0 | 1.16551748055378 | 1 | 0.987 | 0 |
| **TSPAN9** | 0 | -0.903503578898671 | 0.829 | 0.954 | 0 |
| **AKAP3** | 0 | 1.47113744493371 | 0.74 | 0.033 | 0 |
| **CHD4** | 0 | -0.826678516704526 | 0.992 | 1 | 0 |
| **PTMS** | 0 | -0.971900888191259 | 1 | 1 | 0 |
| **ENO2** | 0 | 1.36052050326942 | 0.996 | 0.886 | 0 |
| **GDF3** | 0 | 1.54284782870622 | 0.969 | 0.718 | 0 |
| **DPPA3** | 0 | 3.75703174108994 | 0.995 | 0.267 | 0 |
| **RIMKLB** | 0 | 0.877693376480059 | 0.999 | 0.991 | 0 |
| **SMIM10L1** | 0 | -0.875089979767398 | 0.696 | 0.941 | 0 |

| | | | | | |
|---|---|---|---|---|---|
| **DUSP16** | 0 | 1.75719786732345 | 0.988 | 0.77 | 0 |
| **CREBL2** | 0 | 1.24530822346858 | 0.976 | 0.82 | 0 |
| **PLBD1** | 0 | 2.59398441791991 | 0.999 | 0.865 | 0 |
| **CMAS** | 0 | -1.03953654752501 | 0.757 | 0.959 | 0 |
| **PPFIBP1** | 0 | -1.05955303833429 | 0.521 | 0.888 | 0 |
| **CAPRIN2** | 0 | -1.42816224983714 | 0.476 | 0.85 | 0 |
| **RESF1** | 0 | 0.99299300623573 | 1 | 1 | 0 |
| **BICD1** | 0 | 0.851800818511732 | 0.995 | 0.91 | 0 |
| **FGD4** | 0 | 1.58789244298707 | 0.986 | 0.628 | 0 |
| **YAF2** | 0 | -1.29915147702457 | 0.108 | 0.741 | 0 |
| **PRICKLE1** | 0 | -1.04747319696336 | 0.246 | 0.661 | 0 |
| **TWF1** | 0 | 0.802284449823253 | 0.901 | 0.57 | 0 |
| **AMIGO2** | 0 | -1.12402574211103 | 0.098 | 0.428 | 0 |
| **COL2A1** | 0 | -2.04148019686333 | 0.034 | 0.55 | 0 |
| **CACNB3** | 0 | 0.822961545544208 | 0.794 | 0.469 | 0 |
| **FKBP11** | 0 | 2.33668727327663 | 0.993 | 0.471 | 0 |
| **TUBA1C** | 0 | 1.38896405033837 | 1 | 0.999 | 0 |
| **FMNL3** | 0 | -0.914991820388146 | 0.101 | 0.569 | 0 |
| **LIMA1** | 0 | 1.50936883263197 | 0.979 | 0.766 | 0 |
| **BIN2** | 0 | 0.8084134205496 | 0.492 | 0.025 | 0 |
| **GALNT6** | 0 | 0.837887214698074 | 0.854 | 0.513 | 0 |
| **SLC4A8** | 0 | 1.1348904046194 | 0.889 | 0.429 | 0 |
| **FIGNL2** | 0 | -1.07527109792777 | 0.076 | 0.578 | 0 |
| **NR4A1** | 0 | 1.42477208596254 | 0.826 | 0.345 | 0 |
| **KRT7** | 0 | 1.18169429149016 | 0.641 | 0.14 | 0 |
| **KRT8** | 0 | -1.4262760023705 | 0.98 | 1 | 0 |
| **TNS2** | 0 | -1.38626952560469 | 0.035 | 0.586 | 0 |
| **RARG** | 0 | -1.21001709021396 | 0.29 | 0.702 | 0 |
| **CBX5** | 0 | -1.29107879218296 | 0.838 | 0.988 | 0 |
| **ZNF385A** | 0 | -0.901908740584648 | 0.336 | 0.745 | 0 |
| **ITGA5** | 0 | -2.87047422046471 | 0.49 | 0.953 | 0 |
| **PPP1R1A** | 0 | -1.30857286383421 | 0.299 | 0.663 | 0 |
| **CDK2** | 0 | -0.872694761788414 | 0.734 | 0.926 | 0 |
| **RAB5B** | 0 | -0.863941267162093 | 0.821 | 0.977 | 0 |

| | | | | | |
|---|---|---|---|---|---|
| **GLS2** | 0 | 0.970380784978114 | 0.636 | 0.16 | 0 |
| **DDIT3** | 0 | 1.73285889221253 | 0.705 | 0.356 | 0 |
| **RASSF3** | 0 | 1.09095568303268 | 1 | 0.996 | 0 |
| **HMGA2** | 0 | -2.17469149005972 | 0.58 | 0.987 | 0 |
| **MDM2** | 0 | 1.6105689586175 | 1 | 0.99 | 0 |
| **ATXN7L3B** | 0 | -1.53545343528912 | 0.773 | 0.996 | 0 |
| **ACSS3** | 0 | -1.86135343148608 | 0.163 | 0.738 | 0 |
| **RASSF9** | 0 | -1.68227696579953 | 0.022 | 0.648 | 0 |
| **NTS** | 0 | -4.54767927312944 | 0.502 | 0.992 | 0 |
| **TMPO** | 0 | 0.889552192845934 | 1 | 0.999 | 0 |
| **ACTR6** | 0 | 1.46249185563181 | 0.98 | 0.662 | 0 |
| **SPIC** | 0 | 0.811716226622959 | 0.393 | 0.016 | 0 |
| **SYCP3** | 0 | 2.3594080661321 | 0.848 | 0.059 | 0 |
| **DRAM1** | 0 | 1.63962873374081 | 0.944 | 0.414 | 0 |
| **HSP90B1** | 0 | 0.976890428711553 | 1 | 0.914 | 0 |
| **TXNRD1** | 0 | 1.20451977611994 | 1 | 1 | 0 |
| **EID3** | 0 | 0.802513155010834 | 0.552 | 0.188 | 0 |
| **CHST11** | 0 | -1.70623930541707 | 0.468 | 0.931 | 0 |
| **APPL2** | 0 | -0.983530880425433 | 0.326 | 0.759 | 0 |
| **NUAK1** | 0 | -1.0986054961068 | 0.689 | 0.943 | 0 |
| **CKAP4** | 0 | -1.25640775048545 | 0.959 | 0.999 | 0 |
| **RIC8B** | 0 | -0.836479545878836 | 0.385 | 0.753 | 0 |
| **ISCU** | 0 | 0.812529505850228 | 1 | 0.992 | 0 |
| **MMAB** | 0 | -1.11964473184823 | 0.864 | 0.979 | 0 |
| **PPTC7** | 0 | 1.18040709465256 | 0.975 | 0.777 | 0 |
| **VSIG10** | 0 | 1.78165103548443 | 0.999 | 0.87 | 0 |
| **HSPB8** | 0 | -1.15361983991783 | 0.196 | 0.631 | 0 |
| **MSI1** | 0 | -1.90695531939707 | 0.075 | 0.868 | 0 |
| **UNC119B** | 0 | -1.2825062535482 | 0.364 | 0.877 | 0 |
| **RHOF** | 0 | 1.70834846066826 | 0.923 | 0.283 | 0 |
| **DENR** | 0 | 0.887615613636925 | 1 | 0.985 | 0 |
| **VPS37B** | 0 | 1.64300928660986 | 0.976 | 0.591 | 0 |
| **CCDC92** | 0 | -0.848487276327401 | 0.032 | 0.417 | 0 |
| **ZNF664** | 0 | -0.991074011362896 | 0.469 | 0.836 | 0 |

| | | | | | |
|---|---|---|---|---|---|
| FZD10 | 0 | -1.88436846215184 | 0.163 | 0.696 | 0 |
| ULK1 | 0 | 1.10107799684511 | 0.997 | 0.965 | 0 |
| PUS1 | 0 | 0.813301198577699 | 1 | 0.979 | 0 |
| SKA3 | 0 | 1.6481671379012 | 0.994 | 0.963 | 0 |
| ATP12A | 0 | 0.914902495768695 | 0.837 | 0.431 | 0 |
| WASF3 | 0 | -0.997876437417719 | 0.861 | 0.982 | 0 |
| CDX2 | 0 | -1.54597215870214 | 0.039 | 0.452 | 0 |
| FLT1 | 0 | -1.73665758473295 | 0.035 | 0.618 | 0 |
| KATNAL1 | 0 | -0.928240187821601 | 0.363 | 0.776 | 0 |
| PDS5B | 0 | -0.873331641434069 | 0.899 | 0.992 | 0 |
| NBEA | 0 | -1.70709459474246 | 0.043 | 0.74 | 0 |
| DCLK1 | 0 | -1.75528524576741 | 0.85 | 0.87 | 0 |
| CCDC169 | 0 | 0.941513741268337 | 0.747 | 0.335 | 0 |
| CCNA1 | 0 | 3.20674602388197 | 0.442 | 0.049 | 0 |
| UFM1 | 0 | 1.25978921630575 | 1 | 0.974 | 0 |
| LHFPL6 | 0 | -2.2976032420819 | 0.154 | 0.845 | 0 |
| FOXO1 | 0 | -1.79823345474538 | 0.357 | 0.864 | 0 |
| ELF1 | 0 | 1.09839529365447 | 0.909 | 0.658 | 0 |
| RGCC | 0 | 1.65583004333453 | 0.738 | 0.143 | 0 |
| DNAJC15 | 0 | 1.59182245412064 | 0.98 | 0.996 | 0 |
| LCP1 | 0 | 4.76527023816736 | 1 | 0.644 | 0 |
| SUCLA2 | 0 | 0.831228313251028 | 0.987 | 0.892 | 0 |
| NUDT15 | 0 | 1.0617760356379 | 1 | 0.999 | 0 |
| RB1 | 0 | -1.116465568375 | 0.367 | 0.842 | 0 |
| RCBTB2 | 0 | -1.03650715009388 | 0.297 | 0.733 | 0 |
| RCBTB1 | 0 | 0.903860562389628 | 0.982 | 0.955 | 0 |
| KLF5 | 0 | 2.60460381831161 | 0.998 | 0.621 | 0 |
| KLF12 | 0 | -1.53634475611671 | 0.117 | 0.783 | 0 |
| TBC1D4 | 0 | -1.14551009818932 | 0.172 | 0.683 | 0 |
| EDNRB | 0 | -2.09428968495552 | 0.178 | 0.744 | 0 |
| OBI1 | 0 | -1.41321830471525 | 0.479 | 0.939 | 0 |
| GPC6 | 0 | -2.78503445674868 | 0.301 | 0.976 | 0 |
| DZIP1 | 0 | -1.29618731838707 | 0.705 | 0.969 | 0 |
| RAP2A | 0 | -1.15707864557468 | 0.845 | 0.968 | 0 |

| | | | | | |
|---|---|---|---|---|---|
| **FARP1** | 0 | -1.35933820541578 | 0.491 | 0.91 | 0 |
| **ZIC5** | 0 | -1.77404969914879 | 0.017 | 0.612 | 0 |
| **ZIC2** | 0 | -2.59531121047614 | 0.063 | 0.842 | 0 |
| **PCCA** | 0 | -0.903086583846254 | 0.978 | 0.999 | 0 |
| **COL4A1** | 0 | -2.29981298116746 | 0.678 | 0.994 | 0 |
| **COL4A2** | 0 | -2.28914296700505 | 0.747 | 0.995 | 0 |
| **RAB20** | 0 | -1.20666012500253 | 0.425 | 0.731 | 0 |
| **RASA3** | 0 | -0.936166308020325 | 0.446 | 0.727 | 0 |
| **NDRG2** | 0 | -1.10143050555979 | 0.272 | 0.702 | 0 |
| **ARHGEF40** | 0 | -2.11721478906046 | 0.201 | 0.914 | 0 |
| **ZNF219** | 0 | -1.31752096790851 | 0.494 | 0.792 | 0 |
| **SALL2** | 0 | -2.90848475894118 | 0.712 | 0.996 | 0 |
| **SLC7A7** | 0 | 2.09616316067376 | 0.974 | 0.595 | 0 |
| **AJUBA** | 0 | -0.867298654907829 | 0.786 | 0.938 | 0 |
| **SLC7A8** | 0 | 0.843921929603893 | 0.946 | 0.717 | 0 |
| **HOMEZ** | 0 | -0.875115423385913 | 0.016 | 0.49 | 0 |
| **EFS** | 0 | -1.4696080643285 | 0.323 | 0.837 | 0 |
| **PSME1** | 0 | -0.838214456963802 | 0.9 | 0.989 | 0 |
| **REC8** | 0 | 2.60566898537257 | 0.993 | 0.286 | 0 |
| **NFATC4** | 0 | -0.86625049398722 | 0.271 | 0.643 | 0 |
| **EGLN3** | 0 | 2.17096763367131 | 0.975 | 0.709 | 0 |
| **CFL2** | 0 | -0.889410197366274 | 0.678 | 0.922 | 0 |
| **BAZ1A** | 0 | -1.13584617897686 | 0.301 | 0.772 | 0 |
| **NFKBIA** | 0 | 1.0444226835134 | 0.997 | 0.974 | 0 |
| **BRMS1L** | 0 | 1.0699981654388 | 0.987 | 0.881 | 0 |
| **MIPOL1** | 0 | -0.947288537580205 | 0.205 | 0.682 | 0 |
| **PYGL** | 0 | 2.10044973770585 | 1 | 0.973 | 0 |
| **ABHD12B** | 0 | 1.41027051735118 | 0.771 | 0.132 | 0 |
| **FRMD6** | 0 | 1.29564732573539 | 0.965 | 0.752 | 0 |
| **TXNDC16** | 0 | -0.968785683900035 | 0.216 | 0.7 | 0 |
| **BMP4** | 0 | -2.04226176470238 | 0.586 | 0.864 | 0 |
| **SAMD4A** | 0 | -1.60516775131457 | 0.149 | 0.68 | 0 |
| **WDHD1** | 0 | 1.10188473549369 | 1 | 0.992 | 0 |
| **SOCS4** | 0 | 1.58021433937618 | 1 | 0.985 | 0 |

| | | | | | |
|---|---|---|---|---|---|
| **HIF1A** | 0 | -1.20495171349748 | 0.756 | 0.968 | 0 |
| **HSPA2** | 0 | 2.48018752205284 | 0.953 | 0.131 | 0 |
| **GPX2** | 0 | 1.7965609093215 | 0.744 | 0.087 | 0 |
| **RAB15** | 0 | 1.13346230275829 | 1 | 0.989 | 0 |
| **TMEM229B** | 0 | 1.48875685906126 | 0.762 | 0.15 | 0 |
| **DCAF5** | 0 | -1.45823594914126 | 0.485 | 0.949 | 0 |
| **EXD2** | 0 | -0.832048746910078 | 0.266 | 0.668 | 0 |
| **SRSF5** | 0 | -0.848283966188441 | 0.907 | 0.989 | 0 |
| **TTC9** | 0 | -0.949461910962235 | 0.143 | 0.503 | 0 |
| **VRTN** | 0 | -2.99305911433818 | 0.519 | 0.989 | 0 |
| **NPC2** | 0 | -0.812301884796164 | 1 | 1 | 0 |
| **FLVCR2** | 0 | 1.01234841714393 | 0.744 | 0.349 | 0 |
| **TMED8** | 0 | -0.828005056048765 | 0.476 | 0.782 | 0 |
| **SLIRP** | 0 | 1.31013024734128 | 1 | 1 | 0 |
| **GALC** | 0 | -0.94960974940531 | 0.29 | 0.708 | 0 |
| **TMEM251** | 0 | -0.929915478607727 | 0.674 | 0.939 | 0 |
| **UBR7** | 0 | -0.950034170317977 | 0.87 | 0.976 | 0 |
| **IFI27L2** | 0 | -1.09675129016953 | 0.365 | 0.772 | 0 |
| **DICER1** | 0 | -1.05622343117586 | 0.907 | 0.997 | 0 |
| **SYNE3** | 0 | -1.21009140314108 | 0.048 | 0.493 | 0 |
| **TCL1B** | 0 | 0.987782665647124 | 0.887 | 0.531 | 0 |
| **TUNAR** | 0 | -2.56095872551395 | 0.283 | 0.857 | 0 |
| **C14orf132** | 0 | -1.46830129553646 | 0.024 | 0.637 | 0 |
| **EML1** | 0 | -1.21859499074628 | 0.436 | 0.817 | 0 |
| **EVL** | 0 | -1.37412193487586 | 0.942 | 0.999 | 0 |
| **SLC25A29** | 0 | 0.960013393238669 | 0.979 | 0.81 | 0 |
| **RCOR1** | 0 | 1.32325986243191 | 0.999 | 0.997 | 0 |
| **TNFAIP2** | 0 | -1.80531583360603 | 0.207 | 0.763 | 0 |
| **ATP5MPL** | 0 | 0.846248295072887 | 0.96 | 0.757 | 0 |
| **KIF26A** | 0 | -1.56440654204151 | 0.549 | 0.955 | 0 |
| **INF2** | 0 | -1.34092042744551 | 0.986 | 1 | 0 |
| **AKT1** | 0 | -1.02851521967206 | 0.975 | 0.998 | 0 |
| **JAG2** | 0 | 0.948862702231396 | 0.741 | 0.299 | 0 |
| **PACS2** | 0 | -1.09131066590297 | 0.321 | 0.812 | 0 |

| | | | | | |
|---|---|---|---|---|---|
| **MTA1** | 0 | -1.41957112338624 | 0.939 | 0.999 | 0 |
| **CRIP1** | 0 | 2.7518079452559 | 0.999 | 0.867 | 0 |
| **GABRB3** | 0 | -1.28456391680397 | 0.983 | 0.988 | 0 |
| **HERC2** | 0 | -0.820303159377251 | 0.673 | 0.912 | 0 |
| **AVEN** | 0 | -0.810180874643448 | 0.754 | 0.926 | 0 |
| **NANOGP8** | 0 | 1.13203510939255 | 0.819 | 0.266 | 0 |
| **CHST14** | 0 | -0.855406931826497 | 0.796 | 0.967 | 0 |
| **SPINT1** | 0 | 0.919967372261646 | 0.997 | 0.728 | 0 |
| **CHAC1** | 0 | 1.54070576533998 | 0.717 | 0.265 | 0 |
| **TMEM87A** | 0 | 0.807633903110046 | 0.984 | 0.828 | 0 |
| **LCMT2** | 0 | -0.804111823760692 | 0.337 | 0.731 | 0 |
| **TP53BP1** | 0 | -0.843004218214559 | 0.699 | 0.906 | 0 |
| **B2M** | 0 | -1.23067354639364 | 0.761 | 0.973 | 0 |
| **SHF** | 0 | -1.1979963722284 | 0.47 | 0.859 | 0 |
| **MYEF2** | 0 | -1.68006802708017 | 0.159 | 0.784 | 0 |
| **DUT** | 0 | -0.90810598720529 | 0.842 | 0.971 | 0 |
| **EID1** | 0 | -1.81810458292876 | 0.897 | 0.999 | 0 |
| **SCG3** | 0 | -0.859953191473859 | 0.2 | 0.575 | 0 |
| **ARPP19** | 0 | -0.996518909340387 | 0.98 | 0.998 | 0 |
| **RSL24D1** | 0 | 1.05595773316781 | 0.99 | 0.71 | 0 |
| **PRTG** | 0 | -2.80687991289313 | 0.203 | 0.905 | 0 |
| **MINDY2** | 0 | 0.834777233217763 | 0.984 | 0.868 | 0 |
| **MYO1E** | 0 | 0.853314500928946 | 0.992 | 0.926 | 0 |
| **FOXB1** | 0 | -1.45720868464818 | 0.032 | 0.467 | 0 |
| **SNX1** | 0 | -1.59454958927203 | 0.678 | 0.986 | 0 |
| **PCLAF** | 0 | -2.69103803704731 | 0.066 | 0.911 | 0 |
| **ZNF609** | 0 | -1.05083696140392 | 0.492 | 0.882 | 0 |
| **OAZ2** | 0 | -0.848539384244157 | 0.995 | 1 | 0 |
| **CLPX** | 0 | 0.926055453645547 | 0.997 | 0.955 | 0 |
| **IGDCC3** | 0 | -2.52161911254822 | 0.313 | 0.979 | 0 |
| **IGDCC4** | 0 | -1.22973672984881 | 0.018 | 0.621 | 0 |
| **DIS3L** | 0 | -0.884490965562745 | 0.586 | 0.869 | 0 |
| **IQCH** | 0 | -0.852612871931819 | 0.092 | 0.505 | 0 |
| **UACA** | 0 | 0.838127916530893 | 0.92 | 0.709 | 0 |

| | | | | | |
|---|---|---|---|---|---|
| **MYO9A** | 0 | 1.06304087943222 | 0.962 | 0.673 | 0 |
| **NEO1** | 0 | -1.12029530300776 | 0.476 | 0.818 | 0 |
| **LOXL1** | 0 | -1.71866318147936 | 0.304 | 0.858 | 0 |
| **ISLR** | 0 | -1.35699664575388 | 0.046 | 0.583 | 0 |
| **STRA6** | 0 | -0.910210803545989 | 0.218 | 0.631 | 0 |
| **CSK** | 0 | -1.0260269608882 | 0.961 | 0.994 | 0 |
| **PTPN9** | 0 | -1.68290049454885 | 0.459 | 0.963 | 0 |
| **ETFA** | 0 | -1.489418479647 | 0.785 | 0.993 | 0 |
| **TSPAN3** | 0 | -1.27332462924751 | 0.987 | 1 | 0 |
| **PEAK1** | 0 | -1.50696824106568 | 0.215 | 0.839 | 0 |
| **LINGO1** | 0 | 0.905165762775034 | 0.978 | 0.868 | 0 |
| **DNAJA4** | 0 | 1.02720232961139 | 0.628 | 0.052 | 0 |
| **CRABP1** | 0 | -1.55014194280658 | 0.619 | 0.845 | 0 |
| **TLNRD1** | 0 | -0.825456477430156 | 0.863 | 0.979 | 0 |
| **AP3B2** | 0 | -1.19097101885804 | 0.232 | 0.609 | 0 |
| **HDGFL3** | 0 | -1.1201430535941 | 0.942 | 0.999 | 0 |
| **SEC11A** | 0 | -0.807337763798838 | 0.999 | 1 | 0 |
| **AKAP13** | 0 | -0.933173647955272 | 0.694 | 0.893 | 0 |
| **AEN** | 0 | 1.18043623749307 | 1 | 0.998 | 0 |
| **MFGE8** | 0 | -1.57294004015791 | 0.496 | 0.951 | 0 |
| **ARPIN** | 0 | -1.01968736445248 | 0.181 | 0.674 | 0 |
| **IDH2** | 0 | -0.960599998970626 | 0.876 | 0.976 | 0 |
| **ST8SIA2** | 0 | -0.975111617071011 | 0.031 | 0.486 | 0 |
| **FAM174B** | 0 | -2.57362992241874 | 0.243 | 0.904 | 0 |
| **IGF1R** | 0 | -1.61661535667549 | 0.843 | 0.994 | 0 |
| **CHSY1** | 0 | -1.00525708666657 | 0.943 | 0.996 | 0 |
| **FAM234A** | 0 | 1.04400579573179 | 0.999 | 0.996 | 0 |
| **NME4** | 0 | -1.28541279308713 | 0.993 | 1 | 0 |
| **PIGQ** | 0 | -0.980866439411473 | 0.347 | 0.787 | 0 |
| **MEIOB** | 0 | 0.971209168237707 | 0.541 | 0.013 | 0 |
| **SLC9A3R2** | 0 | 1.24079698980602 | 0.903 | 0.503 | 0 |
| **MMP25** | 0 | -0.923399992728833 | 0.247 | 0.634 | 0 |
| **IL32** | 0 | 2.03209695395306 | 0.943 | 0.357 | 0 |
| **NMRAL1** | 0 | 0.909082547169819 | 0.912 | 0.511 | 0 |

| | | | | | |
|---|---|---|---|---|---|
| **CDIP1** | 0 | -0.900922196010603 | 0.833 | 0.971 | 0 |
| **C16orf72** | 0 | 0.843512071902589 | 1 | 0.999 | 0 |
| **SNN** | 0 | -0.800594391843772 | 0.276 | 0.622 | 0 |
| **SHISA9** | 0 | -1.50646222762228 | 0.05 | 0.639 | 0 |
| **TMC7** | 0 | 1.25386914406662 | 0.787 | 0.224 | 0 |
| **TMEM159** | 0 | 1.019681998418 | 0.892 | 0.545 | 0 |
| **CDR2** | 0 | 0.964678646006788 | 0.942 | 0.697 | 0 |
| **NDUFAB1** | 0 | 1.57951169668368 | 1 | 1 | 0 |
| **NUPR1** | 0 | 2.6658049344789 | 0.907 | 0.207 | 0 |
| **CORO1A** | 0 | 1.38713675291528 | 0.723 | 0.138 | 0 |
| **BCL7C** | 0 | -1.5586160509971 | 0.706 | 0.971 | 0 |
| **SHCBP1** | 0 | 1.01758295276918 | 0.965 | 0.754 | 0 |
| **GPT2** | 0 | 1.01464493459375 | 0.987 | 0.949 | 0 |
| **ZNF423** | 0 | -2.17600400521151 | 0.552 | 0.981 | 0 |
| **BRD7** | 0 | -0.847081948296271 | 0.978 | 0.999 | 0 |
| **NKD1** | 0 | -2.07221641931983 | 0.332 | 0.811 | 0 |
| **SALL1** | 0 | -2.81488834174179 | 0.159 | 0.92 | 0 |
| **AC007906.2** | 0 | -0.923802339636075 | 0.037 | 0.487 | 0 |
| **MMP2** | 0 | -2.78068717892749 | 0.834 | 0.995 | 0 |
| **MT1E** | 0 | 1.19577203205043 | 0.715 | 0.098 | 0 |
| **MT1G** | 0 | 2.74211525941069 | 0.983 | 0.258 | 0 |
| **MT1H** | 0 | 2.5149919857547 | 0.957 | 0.318 | 0 |
| **MT1X** | 0 | 1.88559220318235 | 1 | 0.947 | 0 |
| **HERPUD1** | 0 | 1.87471067332491 | 0.946 | 0.717 | 0 |
| **CCDC102A** | 0 | -0.976166635303475 | 0.303 | 0.744 | 0 |
| **KIFC3** | 0 | -1.30470445478905 | 0.274 | 0.776 | 0 |
| **USB1** | 0 | -1.22802330509057 | 0.762 | 0.972 | 0 |
| **MMP15** | 0 | -0.843485146124022 | 0.972 | 0.998 | 0 |
| **CNOT1** | 0 | -1.08406276105723 | 0.563 | 0.916 | 0 |
| **CMTM3** | 0 | -0.848367674542802 | 0.99 | 1 | 0 |
| **RRAD** | 0 | 1.87054941541851 | 0.442 | 0.034 | 0 |
| **PLEKHG4** | 0 | 0.96165909581102 | 0.585 | 0.035 | 0 |
| **PSKH1** | 0 | -1.0680054386943 | 0.515 | 0.903 | 0 |
| **SLC7A6** | 0 | 1.31765665683558 | 0.968 | 0.699 | 0 |

| | | | | | |
|---|---|---|---|---|---|
| **HAS3** | 0 | -2.15568524339463 | 0.082 | 0.74 | 0 |
| **MLKL** | 0 | 1.16227110334767 | 0.58 | 0.026 | 0 |
| **VAT1L** | 0 | -1.54559842895247 | 0.008 | 0.439 | 0 |
| **WWOX** | 0 | -1.5636913799555 | 0.497 | 0.938 | 0 |
| **MAF** | 0 | -1.24462143123806 | 0.079 | 0.536 | 0 |
| **HSDL1** | 0 | -0.89567806630621 | 0.536 | 0.863 | 0 |
| **CRISPLD2** | 0 | 1.06140791218667 | 0.668 | 0.169 | 0 |
| **GINS2** | 0 | -1.14622471778966 | 0.826 | 0.977 | 0 |
| **PIEZO1** | 0 | 0.813602285705074 | 0.515 | 0.059 | 0 |
| **RFLNB** | 0 | 1.83403636863967 | 0.973 | 0.677 | 0 |
| **NXN** | 0 | -1.39879852341877 | 0.956 | 0.999 | 0 |
| **SERPINF1** | 0 | 0.839556477802657 | 0.746 | 0.471 | 0 |
| **SMG6** | 0 | -1.24482200288237 | 0.545 | 0.918 | 0 |
| **METTL16** | 0 | 0.85100666389194 | 0.993 | 0.923 | 0 |
| **SHPK** | 0 | 1.37827690531464 | 0.985 | 0.819 | 0 |
| **ZFP3** | 0 | 1.00599724472359 | 0.542 | 0.028 | 0 |
| **AIPL1** | 0 | 0.802809316650904 | 0.517 | 0.049 | 0 |
| **ELP5** | 0 | -0.876058787644776 | 0.713 | 0.926 | 0 |
| **CLDN7** | 0 | 1.58823249778843 | 0.986 | 0.594 | 0 |
| **SOX15** | 0 | 1.72571609512479 | 0.978 | 0.548 | 0 |
| **TRAPPC1** | 0 | -1.05106158292671 | 0.905 | 0.992 | 0 |
| **NTN1** | 0 | 6.98582770439101 | 1 | 0.992 | 0 |
| **ADORA2B** | 0 | 1.37973759021655 | 0.803 | 0.171 | 0 |
| **RASD1** | 0 | 1.48479496854527 | 0.633 | 0.067 | 0 |
| **RAI1** | 0 | -0.806388573921319 | 0.068 | 0.474 | 0 |
| **SREBF1** | 0 | 1.34026665276245 | 0.998 | 0.941 | 0 |
| **MFAP4** | 0 | -1.74149645349294 | 0.034 | 0.518 | 0 |
| **TMEM97** | 0 | -0.898078329985927 | 0.996 | 1 | 0 |
| **FLOT2** | 0 | -1.03510790870362 | 0.849 | 0.976 | 0 |
| **CPD** | 0 | -0.933737868302079 | 0.963 | 0.998 | 0 |
| **COPRS** | 0 | -0.809167627343904 | 0.89 | 0.98 | 0 |
| **MYO1D** | 0 | -1.0425692130995 | 0.7 | 0.948 | 0 |
| **TMEM98** | 0 | -1.36852007050074 | 0.544 | 0.926 | 0 |
| **SLFN12** | 0 | 1.403962123427 | 0.751 | 0.079 | 0 |

| | | | | | |
|---|---|---|---|---|---|
| **RASL10B** | 0 | -1.73811994371746 | 0.157 | 0.746 | 0 |
| **LHX1** | 0 | -3.06132641027456 | 0.023 | 0.358 | 0 |
| **AATF** | 0 | -0.824607062365219 | 0.942 | 0.992 | 0 |
| **GRB7** | 0 | 1.80904682305709 | 0.965 | 0.409 | 0 |
| **RARA** | 0 | -0.955554145548781 | 0.281 | 0.733 | 0 |
| **IGFBP4** | 0 | -2.12469433596526 | 0.384 | 0.926 | 0 |
| **KRT19** | 0 | -1.40909995000437 | 0.533 | 0.901 | 0 |
| **EIF1** | 0 | 0.887172775998057 | 1 | 0.964 | 0 |
| **P3H4** | 0 | -0.983077459080134 | 0.309 | 0.754 | 0 |
| **FKBP10** | 0 | -1.56095865091672 | 0.881 | 0.996 | 0 |
| **PSMC3IP** | 0 | -0.886780009284595 | 0.535 | 0.833 | 0 |
| **VAT1** | 0 | 0.816660993290626 | 1 | 1 | 0 |
| **ETV4** | 0 | 1.14763390290251 | 0.999 | 0.968 | 0 |
| **SLC25A39** | 0 | 0.877103376937609 | 1 | 1 | 0 |
| **FAM171A2** | 0 | -1.48974318840696 | 0.391 | 0.89 | 0 |
| **FZD2** | 0 | -2.71325170190811 | 0.066 | 0.884 | 0 |
| **GJC1** | 0 | -2.30487388195722 | 0.637 | 0.996 | 0 |
| **DCAKD** | 0 | -0.858074501831991 | 0.75 | 0.944 | 0 |
| **HOXB2** | 0 | 1.38570262062919 | 0.883 | 0.549 | 0 |
| **PRAC1** | 0 | -1.20141608950217 | 0.044 | 0.607 | 0 |
| **NGFR** | 0 | -1.25744551385089 | 0.542 | 0.81 | 0 |
| **ITGA3** | 0 | -0.836445467102047 | 0.085 | 0.459 | 0 |
| **PDK2** | 0 | -0.974661983631884 | 0.21 | 0.685 | 0 |
| **COL1A1** | 0 | -0.801200865687092 | 0.613 | 0.911 | 0 |
| **TOM1L1** | 0 | -1.36807929980431 | 0.302 | 0.852 | 0 |
| **CUEDC1** | 0 | -1.27458085672738 | 0.168 | 0.719 | 0 |
| **VEZF1** | 0 | -0.980622183614498 | 0.802 | 0.978 | 0 |
| **GDPD1** | 0 | 1.15568773662969 | 0.585 | 0.092 | 0 |
| **YPEL2** | 0 | 2.33857262303235 | 0.863 | 0.229 | 0 |
| **MRC2** | 0 | -1.36831176495135 | 0.679 | 0.828 | 0 |
| **ACE** | 0 | 1.20510417930308 | 0.799 | 0.291 | 0 |
| **LIMD2** | 0 | -1.70951511430634 | 0.582 | 0.971 | 0 |
| **FAM20A** | 0 | -1.0936469820698 | 0.061 | 0.4 | 0 |
| **GPRC5C** | 0 | 0.961937309380672 | 0.938 | 0.722 | 0 |

| | | | | | |
|---|---|---|---|---|---|
| **JPT1** | 0 | 1.31335309922004 | 1 | 0.989 | 0 |
| **ACOX1** | 0 | 1.00279919993018 | 0.978 | 0.811 | 0 |
| **QRICH2** | 0 | 1.20610847159812 | 0.799 | 0.357 | 0 |
| **PRPSAP1** | 0 | 1.59155997177043 | 0.998 | 0.933 | 0 |
| **SPHK1** | 0 | 1.67704754528351 | 0.829 | 0.301 | 0 |
| **TMC6** | 0 | -1.23411940754728 | 0.474 | 0.854 | 0 |
| **SOCS3** | 0 | -0.811287507542583 | 0.609 | 0.883 | 0 |
| **TIMP2** | 0 | -1.40200036502256 | 0.69 | 0.928 | 0 |
| **LGALS3BP** | 0 | -1.75943207735955 | 0.164 | 0.859 | 0 |
| **TBC1D16** | 0 | -1.35725252193845 | 0.703 | 0.97 | 0 |
| **CHMP6** | 0 | -0.813631511477924 | 0.626 | 0.893 | 0 |
| **SLC38A10** | 0 | -0.883850811987361 | 0.564 | 0.882 | 0 |
| **BAHCC1** | 0 | -0.94558963001697 | 0.033 | 0.494 | 0 |
| **OXLD1** | 0 | -0.869450910904969 | 0.304 | 0.72 | 0 |
| **MCRIP1** | 0 | -0.831159517203842 | 0.943 | 0.995 | 0 |
| **PYCR1** | 0 | 1.80036356918419 | 0.998 | 0.721 | 0 |
| **TBCD** | 0 | 1.53983995602582 | 1 | 0.999 | 0 |
| **COLEC12** | 0 | -2.8949616762999 | 0.239 | 0.695 | 0 |
| **LAMA1** | 0 | 1.81293716188559 | 0.928 | 0.26 | 0 |
| **TWSG1** | 0 | -1.02564628585694 | 0.859 | 0.984 | 0 |
| **RAB31** | 0 | -1.32263810333068 | 0.571 | 0.891 | 0 |
| **PIEZO2** | 0 | -2.3920253419173 | 0.031 | 0.832 | 0 |
| **IMPA2** | 0 | 1.01522576366496 | 0.997 | 0.963 | 0 |
| **ESCO1** | 0 | 0.942280700735462 | 0.996 | 0.923 | 0 |
| **GATA6** | 0 | -3.1236613806202 | 0.061 | 0.427 | 0 |
| **CABYR** | 0 | 0.943283883632841 | 0.557 | 0.058 | 0 |
| **CDH2** | 0 | -3.74007551257495 | 0.134 | 0.92 | 0 |
| **DSC2** | 0 | -0.810158129371946 | 0.619 | 0.872 | 0 |
| **RNF125** | 0 | 2.76638350555813 | 0.997 | 0.646 | 0 |
| **DTNA** | 0 | -1.88539466207258 | 0.042 | 0.584 | 0 |
| **GALNT1** | 0 | -1.72714469694913 | 0.849 | 0.996 | 0 |
| **SLC39A6** | 0 | -1.00103825661693 | 0.688 | 0.921 | 0 |
| **TPGS2** | 0 | -1.61321854020334 | 0.779 | 0.998 | 0 |
| **HDHD2** | 0 | -1.02923416627516 | 0.682 | 0.95 | 0 |

| | | | | | |
|---|---|---|---|---|---|
| **CTIF** | 0 | -0.932573028019251 | 0.254 | 0.633 | 0 |
| **SMAD7** | 0 | 0.913987690144008 | 0.977 | 0.9 | 0 |
| **LIPG** | 0 | -2.2812225177507 | 0.118 | 0.643 | 0 |
| **MAPK4** | 0 | -1.12450824140006 | 0.06 | 0.465 | 0 |
| **TCF4** | 0 | -1.59807000229045 | 0.698 | 0.966 | 0 |
| **NARS** | 0 | 0.935535800033163 | 1 | 1 | 0 |
| **NEDD4L** | 0 | -1.06543968137777 | 0.612 | 0.911 | 0 |
| **ALPK2** | 0 | -1.0203515792297 | 0.003 | 0.32 | 0 |
| **MALT1** | 0 | -1.08976863545626 | 0.848 | 0.965 | 0 |
| **PMAIP1** | 0 | 0.975552774825702 | 0.998 | 0.967 | 0 |
| **SALL3** | 0 | -2.41942571328162 | 0.254 | 0.935 | 0 |
| **SLC66A2** | 0 | 1.07628407364834 | 0.969 | 0.729 | 0 |
| **HSBP1L1** | 0 | 1.27866607450915 | 0.672 | 0.075 | 0 |
| **PLPP2** | 0 | 1.45482373270701 | 0.705 | 0.083 | 0 |
| **HCN2** | 0 | 1.28455239287668 | 0.972 | 0.63 | 0 |
| **CNN2** | 0 | -1.20417170325041 | 0.998 | 1 | 0 |
| **CIRBP** | 0 | -1.26848410184644 | 0.99 | 1 | 0 |
| **REEP6** | 0 | 2.17291076992841 | 0.982 | 0.413 | 0 |
| **BTBD2** | 0 | -1.11125034906619 | 0.918 | 0.997 | 0 |
| **PIP5K1C** | 0 | -1.28580031598064 | 0.836 | 0.983 | 0 |
| **UHRF1** | 0 | -1.2567679980407 | 0.916 | 0.974 | 0 |
| **TNFSF9** | 0 | 2.16886272972385 | 0.893 | 0.107 | 0 |
| **CD70** | 0 | 2.52561539247306 | 0.897 | 0.128 | 0 |
| **SH2D3A** | 0 | 0.929133067877106 | 0.858 | 0.422 | 0 |
| **ZNF358** | 0 | -1.19798765911797 | 0.805 | 0.987 | 0 |
| **PRR36** | 0 | -1.12428999718215 | 0.094 | 0.522 | 0 |
| **CTXN1** | 0 | -1.07318803587348 | 0.614 | 0.826 | 0 |
| **FBN3** | 0 | -1.26941676745658 | 0.437 | 0.881 | 0 |
| **CERS4** | 0 | -1.60407832710618 | 0.904 | 0.996 | 0 |
| **ZNF560** | 0 | 1.10091570663182 | 0.662 | 0.02 | 0 |
| **DNMT1** | 0 | -1.1455973611216 | 0.964 | 0.997 | 0 |
| **PDE4A** | 0 | 1.28497512006898 | 0.939 | 0.547 | 0 |
| **AP1M2** | 0 | 1.11722205345644 | 0.822 | 0.239 | 0 |
| **SPC24** | 0 | -0.946808746559459 | 0.614 | 0.813 | 0 |

| | | | | | |
|---|---|---|---|---|---|
| **DOCK6** | 0 | 1.10200906828338 | 0.982 | 0.808 | 0 |
| **ELAVL3** | 0 | 0.99068505887725 | 0.476 | 0.048 | 0 |
| **ZNF627** | 0 | -0.891197067270213 | 0.233 | 0.679 | 0 |
| **ACP5** | 0 | 0.938941839467966 | 0.585 | 0.07 | 0 |
| **ZNF844** | 0 | 1.17500230207227 | 0.64 | 0.024 | 0 |
| **TNPO2** | 0 | -0.894878887539697 | 0.956 | 0.998 | 0 |
| **PRDX2** | 0 | -0.825753227344644 | 0.999 | 1 | 0 |
| **PRKACA** | 0 | -1.4072998862081 | 0.907 | 0.998 | 0 |
| **ASF1B** | 0 | -1.13485903241921 | 0.773 | 0.938 | 0 |
| **ADGRE5** | 0 | 0.867955935153646 | 0.725 | 0.332 | 0 |
| **NOTCH3** | 0 | -2.32676908640742 | 0.548 | 0.987 | 0 |
| **AP1M1** | 0 | -1.17698589118413 | 0.835 | 0.984 | 0 |
| **MYO9B** | 0 | -0.816116286627289 | 0.75 | 0.943 | 0 |
| **BST2** | 0 | -2.00645960344598 | 0.574 | 0.908 | 0 |
| **SLC5A5** | 0 | 1.09367876943291 | 0.67 | 0.062 | 0 |
| **KCNN1** | 0 | 0.815021563257951 | 0.786 | 0.44 | 0 |
| **ARRDC2** | 0 | 1.46518454001629 | 0.859 | 0.285 | 0 |
| **IFI30** | 0 | 1.08383404265053 | 0.968 | 0.713 | 0 |
| **GDF15** | 0 | 2.60342602423985 | 0.732 | 0.12 | 0 |
| **CRLF1** | 0 | -2.6389311736105 | 0.469 | 0.847 | 0 |
| **RFXANK** | 0 | -0.985868757021127 | 0.86 | 0.987 | 0 |
| **PBX4** | 0 | 1.24559768751776 | 0.773 | 0.176 | 0 |
| **ZNF253** | 0 | 1.11611245520775 | 0.646 | 0.036 | 0 |
| **ZNF682** | 0 | 0.839505973915556 | 0.822 | 0.509 | 0 |
| **ZNF90** | 0 | 0.921752716955111 | 0.981 | 0.843 | 0 |
| **ZNF486** | 0 | 1.52925068713072 | 0.854 | 0.397 | 0 |
| **ZNF626** | 0 | 1.17234357761047 | 0.683 | 0.036 | 0 |
| **ZNF257** | 0 | 1.00007099548147 | 0.574 | 0.013 | 0 |
| **ZNF676** | 0 | 2.4610093951782 | 0.821 | 0.116 | 0 |
| **ZNF98** | 0 | 1.04841692129217 | 0.599 | 0.017 | 0 |
| **ZNF492** | 0 | 2.36345387206179 | 0.958 | 0.086 | 0 |
| **ZNF728** | 0 | 1.0650548513399 | 0.62 | 0.011 | 0 |
| **ZNF730** | 0 | -0.894518336525468 | 0.812 | 0.969 | 0 |
| **ZNF146** | 0 | 0.833927679119385 | 1 | 1 | 0 |

| | | | | | |
|---|---|---|---|---|---|
| **ZNF829** | 0 | 0.981202387199861 | 0.61 | 0.016 | 0 |
| **ZNF568** | 0 | 1.80050112555983 | 0.915 | 0.124 | 0 |
| **ZNF793** | 0 | 0.872227388405337 | 0.818 | 0.448 | 0 |
| **ZNF607** | 0 | 0.813814096245879 | 0.765 | 0.366 | 0 |
| **PPP1R14A** | 0 | 1.67341459030962 | 0.996 | 0.784 | 0 |
| **YIF1B** | 0 | 0.999343890938624 | 1 | 0.996 | 0 |
| **C19orf33** | 0 | 0.933284119690593 | 0.65 | 0.153 | 0 |
| **PLEKHG2** | 0 | -1.17954610754266 | 0.504 | 0.899 | 0 |
| **TIMM50** | 0 | 0.930382468325671 | 1 | 1 | 0 |
| **EID2** | 0 | -0.930346385407449 | 0.419 | 0.816 | 0 |
| **FCGBP** | 0 | 1.62733288412623 | 0.727 | 0.055 | 0 |
| **PSMC4** | 0 | 0.891083772327723 | 1 | 0.998 | 0 |
| **PLD3** | 0 | -0.866333395646357 | 1 | 1 | 0 |
| **AXL** | 0 | -2.09278148433057 | 0.142 | 0.767 | 0 |
| **ATP1A3** | 0 | 1.71724567220855 | 0.838 | 0.075 | 0 |
| **MEGF8** | 0 | -1.00867414612606 | 0.337 | 0.782 | 0 |
| **ETHE1** | 0 | 1.04741659629126 | 0.952 | 0.688 | 0 |
| **ZNF229** | 0 | 0.936256758539974 | 0.787 | 0.309 | 0 |
| **PVR** | 0 | 1.18342433866493 | 0.993 | 0.899 | 0 |
| **BCL3** | 0 | 0.985599005780534 | 0.771 | 0.445 | 0 |
| **BCAM** | 0 | -1.98083341603027 | 0.46 | 0.874 | 0 |
| **APOE** | 0 | 0.997985703417671 | 1 | 1 | 0 |
| **APOC1** | 0 | 1.01592594254387 | 0.967 | 0.738 | 0 |
| **ZNF296** | 0 | 1.40776654877393 | 0.895 | 0.534 | 0 |
| **TRAPPC6A** | 0 | -1.15848671491593 | 0.641 | 0.949 | 0 |
| **ERCC1** | 0 | -1.01370001197502 | 0.729 | 0.953 | 0 |
| **SIX5** | 0 | -0.872315812924613 | 0.508 | 0.841 | 0 |
| **CCDC8** | 0 | -0.918480896398523 | 0.846 | 0.985 | 0 |
| **ARHGAP35** | 0 | -1.6996747896024 | 0.614 | 0.981 | 0 |
| **NPAS1** | 0 | 0.839836529577203 | 0.647 | 0.237 | 0 |
| **TMEM160** | 0 | 0.984372077249597 | 0.999 | 0.97 | 0 |
| **BBC3** | 0 | 1.14907495432137 | 0.98 | 0.732 | 0 |
| **C5AR1** | 0 | 0.900332812316943 | 0.671 | 0.198 | 0 |
| **MEIS3** | 0 | -2.08775322291767 | 0.498 | 0.88 | 0 |

| | | | | | |
|---|---|---|---|---|---|
| **ZSWIM9** | 0 | -0.869593092310949 | 0.12 | 0.586 | 0 |
| **CA11** | 0 | -1.93606808199439 | 0.043 | 0.762 | 0 |
| **RASIP1** | 0 | 1.01311075027004 | 0.659 | 0.153 | 0 |
| **RCN3** | 0 | 1.57769637426232 | 0.861 | 0.169 | 0 |
| **SCAF1** | 0 | -0.934652514624702 | 0.908 | 0.992 | 0 |
| **PTOV1** | 0 | -0.870497167110209 | 0.997 | 1 | 0 |
| **KCNC3** | 0 | 0.880251015139638 | 0.736 | 0.329 | 0 |
| **FAM71E1** | 0 | 1.03942814497078 | 0.723 | 0.194 | 0 |
| **KLK8** | 0 | 1.11170497241947 | 0.877 | 0.372 | 0 |
| **KLK13** | 0 | 1.40338076895295 | 0.856 | 0.29 | 0 |
| **IGLON5** | 0 | -1.20279490993391 | 0.29 | 0.798 | 0 |
| **LIM2** | 0 | 1.48931996467986 | 0.784 | 0.043 | 0 |
| **ZNF577** | 0 | 1.15043302313217 | 0.858 | 0.451 | 0 |
| **ZNF649** | 0 | 1.29171653802387 | 0.986 | 0.827 | 0 |
| **ZNF614** | 0 | 0.961693256019924 | 0.94 | 0.66 | 0 |
| **ZNF841** | 0 | 0.859869676880755 | 0.742 | 0.363 | 0 |
| **ZNF880** | 0 | 0.951516371604637 | 0.683 | 0.119 | 0 |
| **ZNF677** | 0 | 1.38664601463078 | 0.788 | 0.044 | 0 |
| **MYADM** | 0 | 0.929174512371645 | 1 | 0.995 | 0 |
| **MBOAT7** | 0 | -0.956272869505601 | 0.928 | 0.997 | 0 |
| **NLRP7** | 0 | 4.43422480435297 | 0.999 | 0.647 | 0 |
| **NLRP2** | 0 | 2.92569326755858 | 1 | 0.981 | 0 |
| **TNNI3** | 0 | 0.901450289587361 | 0.813 | 0.337 | 0 |
| **BRSK1** | 0 | -0.83814176943558 | 0.151 | 0.589 | 0 |
| **IL11** | 0 | 0.854918060175833 | 0.507 | 0.165 | 0 |
| **ISOC2** | 0 | 0.845469560786294 | 0.998 | 0.973 | 0 |
| **C19orf85** | 0 | 1.76096493636839 | 0.764 | 0.088 | 0 |
| **EPN1** | 0 | -0.827806189635847 | 0.985 | 0.999 | 0 |
| **ZNF667** | 0 | 1.08229182002947 | 0.632 | 0.021 | 0 |
| **PEG3** | 0 | 1.50564447917228 | 0.647 | 0.03 | 0 |
| **ZCCHC3** | 0 | -1.82998153777043 | 0.751 | 0.996 | 0 |
| **SOX12** | 0 | -1.16256801636416 | 0.678 | 0.951 | 0 |
| **NRSN2** | 0 | -0.983153202165654 | 0.558 | 0.876 | 0 |
| **TRIB3** | 0 | 2.89202087317261 | 0.983 | 0.626 | 0 |

| | | | | | |
|---|---|---|---|---|---|
| **RBCK1** | 0 | -0.831014079315038 | 0.425 | 0.77 | 0 |
| **SRXN1** | 0 | -1.1186387221681 | 0.743 | 0.938 | 0 |
| **CPXM1** | 0 | -1.06034294549773 | 0.89 | 0.983 | 0 |
| **PCED1A** | 0 | -1.54797090021009 | 0.223 | 0.823 | 0 |
| **PTPRA** | 0 | -1.17642818841241 | 0.737 | 0.973 | 0 |
| **UBOX5** | 0 | -0.864478083634771 | 0.226 | 0.667 | 0 |
| **CENPB** | 0 | -0.933224921187498 | 0.848 | 0.988 | 0 |
| **SLC23A2** | 0 | 1.24968350127035 | 0.972 | 0.849 | 0 |
| **CDS2** | 0 | -1.67048889345908 | 0.702 | 0.992 | 0 |
| **MCM8** | 0 | -0.901874046703333 | 0.975 | 0.998 | 0 |
| **BMP2** | 0 | -3.32059896744339 | 0.044 | 0.611 | 0 |
| **BTBD3** | 0 | -1.58381196997408 | 0.572 | 0.966 | 0 |
| **RRBP1** | 0 | -1.06723301146801 | 0.989 | 0.998 | 0 |
| **OVOL2** | 0 | 2.10378671062205 | 0.957 | 0.281 | 0 |
| **PET117** | 0 | -0.873112909337354 | 0.767 | 0.962 | 0 |
| **RBBP9** | 0 | 1.09553688074184 | 0.948 | 0.751 | 0 |
| **SMIM26** | 0 | -0.893773317153177 | 0.643 | 0.862 | 0 |
| **RIN2** | 0 | -1.57891732810637 | 0.408 | 0.876 | 0 |
| **RALGAPA2** | 0 | -0.956510706640355 | 0.286 | 0.706 | 0 |
| **XRN2** | 0 | -0.919055802352412 | 0.999 | 1 | 0 |
| **CST3** | 0 | -1.84900862898399 | 0.412 | 0.941 | 0 |
| **PYGB** | 0 | 1.37898512669095 | 0.999 | 0.998 | 0 |
| **ASXL1** | 0 | -1.92000632679231 | 0.518 | 0.977 | 0 |
| **NOL4L** | 0 | -1.12755336493008 | 0.417 | 0.856 | 0 |
| **COMMD7** | 0 | -1.09503116059996 | 0.755 | 0.946 | 0 |
| **CBFA2T2** | 0 | 1.04695371082241 | 0.991 | 0.92 | 0 |
| **E2F1** | 0 | -1.22009994344353 | 0.916 | 0.977 | 0 |
| **NCOA6** | 0 | -0.859819125517242 | 0.582 | 0.871 | 0 |
| **PROCR** | 0 | 2.69052136480171 | 0.957 | 0.476 | 0 |
| **MMP24** | 0 | 1.63029464839025 | 0.995 | 0.655 | 0 |
| **DLGAP4** | 0 | -1.00041282991188 | 0.938 | 0.995 | 0 |
| **MYL9** | 0 | -2.2597851103165 | 0.96 | 0.997 | 0 |
| **SAMHD1** | 0 | 3.32522292597173 | 0.997 | 0.591 | 0 |
| **SRC** | 0 | -1.56959338749268 | 0.714 | 0.987 | 0 |

| | | | | | |
|---|---|---|---|---|---|
| **BLCAP** | 0 | -0.910715017868703 | 0.881 | 0.988 | 0 |
| **NNAT** | 0 | 2.27215820225613 | 0.779 | 0.019 | 0 |
| **MYBL2** | 0 | 1.33383491458191 | 1 | 0.999 | 0 |
| **HNF4A** | 0 | 1.64344365820165 | 0.776 | 0.054 | 0 |
| **TTPAL** | 0 | -0.840738680039818 | 0.618 | 0.884 | 0 |
| **PKIG** | 0 | -1.11088519190842 | 0.766 | 0.955 | 0 |
| **STK4** | 0 | -1.0079020477519 | 0.835 | 0.976 | 0 |
| **SDC4** | 0 | 0.864041397148341 | 0.999 | 0.999 | 0 |
| **DBNDD2** | 0 | -1.68496536687284 | 0.31 | 0.879 | 0 |
| **PIGT** | 0 | -1.23161097268119 | 0.978 | 1 | 0 |
| **WFDC2** | 0 | -3.09798273404918 | 0.346 | 0.927 | 0 |
| **ZNF335** | 0 | 0.99760592129334 | 0.956 | 0.735 | 0 |
| **NCOA3** | 0 | 1.34154402203744 | 0.997 | 0.916 | 0 |
| **SULF2** | 0 | -1.23666170608554 | 0.911 | 0.99 | 0 |
| **PREX1** | 0 | -1.84509089997935 | 0.471 | 0.87 | 0 |
| **PTGIS** | 0 | -1.20229365748123 | 0.16 | 0.681 | 0 |
| **RNF114** | 0 | 1.14091775999467 | 1 | 1 | 0 |
| **UBE2V1** | 0 | -1.20802476114721 | 0.997 | 1 | 0 |
| **PTPN1** | 0 | -1.05585869659724 | 0.917 | 0.995 | 0 |
| **BCAS4** | 0 | -0.848711182328511 | 0.561 | 0.819 | 0 |
| **KCNG1** | 0 | -1.87734654799599 | 0.312 | 0.823 | 0 |
| **BMP7** | 0 | -2.37670513920801 | 0.446 | 0.952 | 0 |
| **NELFCD** | 0 | -1.21475056785766 | 0.953 | 0.998 | 0 |
| **CTSZ** | 0 | -1.24471680845015 | 0.58 | 0.868 | 0 |
| **ATP5F1E** | 0 | -1.62756476646429 | 0.98 | 1 | 0 |
| **FAM217B** | 0 | -0.869438791490469 | 0.663 | 0.917 | 0 |
| **LSM14B** | 0 | -0.893408114400564 | 0.958 | 0.997 | 0 |
| **LAMA5** | 0 | -0.844546116447892 | 0.987 | 0.992 | 0 |
| **CABLES2** | 0 | -0.895804971610441 | 0.204 | 0.651 | 0 |
| **KCNQ2** | 0 | 1.40831651966872 | 0.885 | 0.239 | 0 |
| **ARFRP1** | 0 | -0.838222099096197 | 0.727 | 0.939 | 0 |
| **SLC2A4RG** | 0 | -1.28839886614807 | 0.674 | 0.961 | 0 |
| **ZBTB46** | 0 | -0.818485429774797 | 0.49 | 0.772 | 0 |
| **DNAJC5** | 0 | -1.29474181898542 | 0.89 | 0.998 | 0 |

| | | | | | |
|---|---|---|---|---|---|
| **ZNF512B** | 0 | -1.09113075033189 | 0.595 | 0.918 | 0 |
| **TCEA2** | 0 | -1.13281405819402 | 0.542 | 0.903 | 0 |
| **NRIP1** | 0 | -1.07913214502574 | 0.498 | 0.86 | 0 |
| **APP** | 0 | -1.18503652548084 | 1 | 1 | 0 |
| **HUNK** | 0 | -1.49895157129614 | 0.232 | 0.834 | 0 |
| **CBR3** | 0 | 1.0315033155845 | 0.833 | 0.37 | 0 |
| **TTC3** | 0 | -1.05079778355904 | 0.827 | 0.978 | 0 |
| **BACE2** | 0 | -1.16519801092873 | 0.475 | 0.801 | 0 |
| **MX1** | 0 | 1.52694872229275 | 0.481 | 0.018 | 0 |
| **RIPK4** | 0 | 0.99085715510322 | 0.77 | 0.358 | 0 |
| **PDE9A** | 0 | -1.11965542166318 | 0.109 | 0.633 | 0 |
| **PDXK** | 0 | 0.956208616025776 | 1 | 0.996 | 0 |
| **DNMT3L** | 0 | 4.24456925750317 | 0.999 | 0.719 | 0 |
| **UBE2G2** | 0 | -1.06289258940014 | 0.669 | 0.947 | 0 |
| **SUMO3** | 0 | 0.869543474873748 | 1 | 0.99 | 0 |
| **COL18A1** | 0 | -3.59343501211139 | 0.564 | 0.995 | 0 |
| **COL6A1** | 0 | -1.97700021032112 | 0.946 | 0.947 | 0 |
| **COL6A2** | 0 | -2.23281658391091 | 0.886 | 0.979 | 0 |
| **SPATC1L** | 0 | -0.996980247708897 | 0.085 | 0.621 | 0 |
| **LSS** | 0 | -0.967682970403838 | 0.947 | 0.992 | 0 |
| **COMT** | 0 | -0.859263259445206 | 0.622 | 0.882 | 0 |
| **SCARF2** | 0 | -1.07413570945132 | 0.12 | 0.625 | 0 |
| **MED15** | 0 | 0.882755129797546 | 0.987 | 0.931 | 0 |
| **SDF2L1** | 0 | 1.20795281008838 | 0.984 | 0.93 | 0 |
| **DERL3** | 0 | 1.05107617827245 | 0.654 | 0.217 | 0 |
| **DDT** | 0 | 0.833506108079058 | 0.999 | 0.994 | 0 |
| **SUSD2** | 0 | 2.47638716864571 | 0.994 | 0.656 | 0 |
| **CHEK2** | 0 | 1.20299757351834 | 0.993 | 0.918 | 0 |
| **XBP1** | 0 | 1.34202602357196 | 1 | 0.982 | 0 |
| **NEFH** | 0 | 2.92857093350015 | 0.992 | 0.451 | 0 |
| **THOC5** | 0 | 0.995281640379178 | 0.998 | 0.965 | 0 |
| **NF2** | 0 | 1.13531380478963 | 1 | 0.992 | 0 |
| **TCN2** | 0 | -1.58263503597059 | 0.36 | 0.91 | 0 |
| **SELENOM** | 0 | 0.816942505471699 | 0.907 | 0.631 | 0 |

| | | | | | |
|---|---|---|---|---|---|
| **LIMK2** | 0 | 1.63365274378746 | 0.998 | 0.931 | 0 |
| **HMOX1** | 0 | 1.55876273665341 | 0.854 | 0.412 | 0 |
| **EIF3D** | 0 | 0.832648153118945 | 1 | 0.989 | 0 |
| **CDC42EP1** | 0 | -0.853805608985225 | 0.233 | 0.659 | 0 |
| **TRIOBP** | 0 | -1.15286263105945 | 0.726 | 0.966 | 0 |
| **GCAT** | 0 | 0.933226281039973 | 0.824 | 0.418 | 0 |
| **TOMM22** | 0 | 0.878399867783634 | 1 | 0.997 | 0 |
| **SYNGR1** | 0 | -1.36968991394798 | 0.312 | 0.822 | 0 |
| **MGAT3** | 0 | -1.0266564966681 | 0.102 | 0.568 | 0 |
| **ATF4** | 0 | 0.890142602882992 | 1 | 0.997 | 0 |
| **SNU13** | 0 | 1.39296917668255 | 1 | 1 | 0 |
| **NDUFA6** | 0 | 0.961970749198913 | 0.999 | 0.969 | 0 |
| **A4GALT** | 0 | 2.8386439862644 | 0.999 | 0.633 | 0 |
| **TTLL12** | 0 | 0.929119365473178 | 0.997 | 0.968 | 0 |
| **SULT4A1** | 0 | 0.871136386425274 | 0.705 | 0.285 | 0 |
| **PARVB** | 0 | 0.98007415409545 | 0.849 | 0.496 | 0 |
| **ARHGAP8** | 0 | 1.47089005175882 | 0.961 | 0.508 | 0 |
| **FBLN1** | 0 | -0.890839290791766 | 0.997 | 1 | 0 |
| **CRELD2** | 0 | 0.868604937426775 | 0.956 | 0.875 | 0 |
| **CSF2RA** | 0 | 0.964799633601415 | 0.718 | 0.336 | 0 |
| **ASMTL** | 0 | 1.14753861534137 | 0.983 | 0.895 | 0 |
| **DHRSX** | 0 | 2.03913767608669 | 0.995 | 0.906 | 0 |
| **CD99** | 0 | -2.8778065531743 | 0.121 | 0.952 | 0 |
| **GYG2** | 0 | -1.5252440573399 | 0.58 | 0.967 | 0 |
| **ARSE** | 0 | -0.925787538227578 | 0.17 | 0.6 | 0 |
| **NLGN4X** | 0 | -1.79024599490644 | 0.136 | 0.588 | 0 |
| **TBL1X** | 0 | -1.12834847399975 | 0.698 | 0.938 | 0 |
| **WWC3** | 0 | -1.6905845774712 | 0.496 | 0.933 | 0 |
| **TMSB4X** | 0 | -3.14521947031965 | 0.995 | 1 | 0 |
| **PIR** | 0 | -1.03537840297868 | 0.65 | 0.904 | 0 |
| **AP1S2** | 0 | -1.84548651514612 | 0.286 | 0.926 | 0 |
| **GRPR** | 0 | -1.36970816163594 | 0.013 | 0.475 | 0 |
| **CTPS2** | 0 | -1.29037646855427 | 0.749 | 0.988 | 0 |
| **RBBP7** | 0 | -1.01497755481152 | 0.861 | 0.986 | 0 |

| | | | | | |
|---|---|---|---|---|---|
| **SH3KBP1** | 0 | -1.59320233557037 | 0.529 | 0.956 | 0 |
| **SMS** | 0 | -2.11506717416583 | 0.943 | 1 | 0 |
| **EIF2S3** | 0 | 0.938572284750353 | 1 | 0.807 | 0 |
| **PCYT1B** | 0 | -1.28218162062626 | 0.319 | 0.799 | 0 |
| **MAGEB2** | 0 | 1.67564605705914 | 0.619 | 0.04 | 0 |
| **NR0B1** | 0 | -2.16571452495062 | 0.009 | 0.512 | 0 |
| **DMD** | 0 | -1.87077413512945 | 0.35 | 0.874 | 0 |
| **TSPAN7** | 0 | -2.00340882653617 | 0.054 | 0.623 | 0 |
| **CASK** | 0 | -0.821741127083591 | 0.961 | 0.996 | 0 |
| **JADE3** | 0 | -0.982827642676503 | 0.425 | 0.84 | 0 |
| **USP11** | 0 | -1.14779980869881 | 0.878 | 0.994 | 0 |
| **TIMP1** | 0 | -1.2703524398014 | 0.988 | 0.999 | 0 |
| **PIM2** | 0 | 1.66897065180074 | 0.998 | 0.964 | 0 |
| **GSPT2** | 0 | -1.40261770965578 | 0.406 | 0.918 | 0 |
| **MAGED1** | 0 | -2.29018581107025 | 0.955 | 1 | 0 |
| **WNK3** | 0 | -1.01438391161557 | 0.484 | 0.857 | 0 |
| **MAGED2** | 0 | -1.97103483910846 | 0.983 | 1 | 0 |
| **TRO** | 0 | -1.44533445904541 | 0.642 | 0.964 | 0 |
| **FAM104B** | 0 | -1.09952257867277 | 0.527 | 0.904 | 0 |
| **MAGEH1** | 0 | -1.70237999624276 | 0.254 | 0.905 | 0 |
| **USP51** | 0 | -1.05998562593017 | 0.163 | 0.696 | 0 |
| **UBQLN2** | 0 | -1.1763803415531 | 0.913 | 0.999 | 0 |
| **SPIN3** | 0 | -1.04538882090134 | 0.357 | 0.751 | 0 |
| **ARHGEF9** | 0 | -1.26364954523148 | 0.156 | 0.745 | 0 |
| **AMER1** | 0 | -1.35048930553943 | 0.291 | 0.85 | 0 |
| **ZC4H2** | 0 | -1.01636564452328 | 0.331 | 0.769 | 0 |
| **MSN** | 0 | -0.849680817486711 | 0.997 | 1 | 0 |
| **HEPH** | 0 | -2.11193394833536 | 0.023 | 0.889 | 0 |
| **STARD8** | 0 | -0.980764100909513 | 0.221 | 0.573 | 0 |
| **EFNB1** | 0 | -2.51382214769164 | 0.501 | 0.945 | 0 |
| **FOXO4** | 0 | -1.82840640985428 | 0.408 | 0.84 | 0 |
| **MED12** | 0 | -1.60165452008195 | 0.286 | 0.875 | 0 |
| **NHSL2** | 0 | -0.819018932018341 | 0.184 | 0.569 | 0 |
| **ERCC6L** | 0 | 0.853973797661586 | 0.974 | 0.914 | 0 |

| | | | | | |
|---|---|---|---|---|---|
| **PHKA1** | 0 | 3.04896820556405 | 1 | 0.741 | 0 |
| **SLC16A2** | 0 | -2.21791921675387 | 0.567 | 0.971 | 0 |
| **COX7B** | 0 | 1.02806709274289 | 0.999 | 0.9 | 0 |
| **TAF9B** | 0 | -1.14240274668408 | 0.476 | 0.869 | 0 |
| **TBX22** | 0 | -1.16677433550771 | 0.005 | 0.461 | 0 |
| **BRWD3** | 0 | 1.20707090265772 | 0.998 | 0.978 | 0 |
| **SH3BGRL** | 0 | -0.820409443001098 | 0.764 | 0.954 | 0 |
| **ZNF711** | 0 | -1.15637906276509 | 0.538 | 0.914 | 0 |
| **KLHL4** | 0 | -1.45085497163114 | 0.014 | 0.524 | 0 |
| **NAP1L3** | 0 | -1.52759237161153 | 0.228 | 0.825 | 0 |
| **DIAPH2** | 0 | -1.36703625894144 | 0.256 | 0.752 | 0 |
| **TRMT2B** | 0 | -0.970334658465415 | 0.544 | 0.886 | 0 |
| **CENPI** | 0 | -1.0540877899298 | 0.704 | 0.897 | 0 |
| **ARMCX1** | 0 | -0.991892745635042 | 0.378 | 0.775 | 0 |
| **ARMCX2** | 0 | -1.10952994525931 | 0.713 | 0.966 | 0 |
| **TMSB15A** | 0 | -2.15887752015032 | 0.197 | 0.803 | 0 |
| **BEX4** | 0 | -1.54567503525923 | 0.861 | 0.995 | 0 |
| **TCEAL8** | 0 | -0.88471273065267 | 0.98 | 1 | 0 |
| **TCEAL9** | 0 | -2.03702758907647 | 0.678 | 0.99 | 0 |
| **BEX3** | 0 | -2.54849527229884 | 0.968 | 1 | 0 |
| **TCEAL4** | 0 | -0.840011326381538 | 0.964 | 0.997 | 0 |
| **TMSB15B** | 0 | -0.863848490952243 | 0.083 | 0.496 | 0 |
| **FAM199X** | 0 | -0.957283565897074 | 0.777 | 0.971 | 0 |
| **NRK** | 0 | -2.77328247218165 | 0.25 | 0.728 | 0 |
| **MID2** | 0 | -0.995844516327787 | 0.171 | 0.627 | 0 |
| **COL4A6** | 0 | -1.12744247051441 | 0.027 | 0.583 | 0 |
| **COL4A5** | 0 | -1.73757881197414 | 0.065 | 0.804 | 0 |
| **LUZP4** | 0 | 0.956400969731971 | 0.497 | 0.015 | 0 |
| **WDR44** | 0 | 1.02548160491774 | 0.978 | 0.85 | 0 |
| **IL13RA1** | 0 | -1.6595713330293 | 0.181 | 0.685 | 0 |
| **PGRMC1** | 0 | -1.69339952577779 | 0.992 | 1 | 0 |
| **SLC25A5** | 0 | 1.06092528602982 | 0.988 | 0.698 | 0 |
| **SEPTIN6** | 0 | -1.01659195649497 | 0.556 | 0.893 | 0 |
| **ZBTB33** | 0 | -0.962159985353855 | 0.731 | 0.968 | 0 |

| | | | | | |
|---|---|---|---|---|---|
| **C1GALT1C1** | 0 | -0.980679743092405 | 0.502 | 0.851 | 0 |
| **SMARCA1** | 0 | -1.89960602712912 | 0.683 | 0.991 | 0 |
| **ELF4** | 0 | -0.870955465855266 | 0.12 | 0.506 | 0 |
| **IGSF1** | 0 | 1.31488204623192 | 0.876 | 0.551 | 0 |
| **MBNL3** | 0 | -1.05553889549788 | 0.567 | 0.876 | 0 |
| **HS6ST2** | 0 | -1.82891861593654 | 0.112 | 0.83 | 0 |
| **GPC4** | 0 | -1.49434519815412 | 0.892 | 0.996 | 0 |
| **FAM122B** | 0 | -1.36169699730667 | 0.666 | 0.951 | 0 |
| **RTL8C** | 0 | -0.940196695810745 | 0.823 | 0.941 | 0 |
| **SLC9A6** | 0 | -0.81924227813678 | 0.542 | 0.853 | 0 |
| **ZIC3** | 0 | -1.81879779638905 | 0.308 | 0.864 | 0 |
| **SOX3** | 0 | -1.18612040392014 | 0.097 | 0.452 | 0 |
| **AFF2** | 0 | 1.7641449972081 | 0.875 | 0.276 | 0 |
| **IDS** | 0 | -1.16008144209274 | 0.346 | 0.832 | 0 |
| **TMEM185A** | 0 | -1.46707084740623 | 0.804 | 0.988 | 0 |
| **CD99L2** | 0 | -1.24420444732765 | 0.146 | 0.633 | 0 |
| **VMA21** | 0 | -1.20368171639997 | 0.948 | 1 | 0 |
| **ZNF185** | 0 | -1.6696923502864 | 0.246 | 0.842 | 0 |
| **BGN** | 0 | -1.86660304696601 | 0.244 | 0.75 | 0 |
| **DUSP9** | 0 | 1.10566030370061 | 0.672 | 0.092 | 0 |
| **SLC6A8** | 0 | 1.51424621373349 | 0.995 | 0.877 | 0 |
| **PDZD4** | 0 | -1.04066115723297 | 0.351 | 0.775 | 0 |
| **L1CAM** | 0 | -1.13387006410033 | 0.676 | 0.888 | 0 |
| **GDI1** | 0 | -0.947392867695822 | 0.986 | 1 | 0 |
| **MPP1** | 0 | -0.954725895169991 | 0.57 | 0.858 | 0 |
| **TMLHE** | 0 | -1.21216699105862 | 0.188 | 0.767 | 0 |
| **DDX3Y** | 0 | 1.31639616725836 | 0.952 | 0.863 | 0 |
| **NOTUM** | 2.2780 | -0.83070599213262 | 0.1 | 0.432 | 3.8184440 |
| **PCBP4** | 2.7547 | -0.840199630984454 | 0.321 | 0.637 | 4.6175953 |
| **INSYN1** | 7.8499 | -0.861212849232039 | 0.148 | 0.482 | 1.3158141 |
| **RAB38** | 2.1192 | -0.886478460750462 | 0.186 | 0.515 | 3.5522470 |
| **PABPC4L** | 3.0967 | -0.947446616128329 | 0.348 | 0.664 | 5.1907672 |
| **MYC** | 7.2708 | -0.854698868099244 | 0.843 | 0.962 | 1.2187347 |
| **GRHL3** | 3.9098 | -0.944027312396627 | 0.103 | 0.431 | 6.5537191 |

| | | | | | |
|---|---|---|---|---|---|
| **SHISA3** | 1.7176 | -0.812329259059861 | 0.041 | 0.349 | 2.87907312 |
| **RGMA** | 3.8803 | -0.906937926489466 | 0.489 | 0.759 | 6.50419058 |
| **LHFPL2** | 9.3506 | -0.954134596065633 | 0.3 | 0.619 | 1.56735019 |
| **TGFB1I1** | 1.1636 | -0.846118571116937 | 0.324 | 0.638 | 1.95047404 |
| **PLAAT5** | 2.0723 | -0.855659990653382 | 0.401 | 0.698 | 3.47370891 |
| **CCDC85C** | 3.3322 | -0.904993068376143 | 0.873 | 0.925 | 5.58558730 |
| **APCDD1** | 4.4696 | -1.12323074852077 | 0.279 | 0.599 | 7.49205680 |
| **SCAND1** | 4.8933 | -0.852830910218768 | 0.828 | 0.923 | 8.20224660 |
| **MESP1** | 9.1596 | -0.970417788138216 | 0.008 | 0.292 | 1.53533591 |
| **ITM2A** | 2.0400 | -1.03463891889598 | 0.133 | 0.462 | 3.41955238 |
| **CCNG2** | 2.1539 | 0.882707179976743 | 0.727 | 0.441 | 3.61052720 |
| **ZNF165** | 7.0197 | 0.886393862752085 | 0.49 | 0.191 | 1.17664271 |
| **TBX6** | 3.3961 | -1.85427341147794 | 0.09 | 0.39 | 5.69262747 |
| **SLC52A3** | 5.5710 | -1.0711986903163 | 0.173 | 0.486 | 9.33816252 |
| **LRIG3** | 4.7105 | -1.66149205856723 | 0.592 | 0.782 | 7.89590281 |
| **FAM83D** | 2.0947 | 0.814497238377577 | 0.979 | 0.917 | 3.51126683 |
| **TRABD2B** | 2.5454 | -0.809992467042668 | 0.243 | 0.557 | 4.26660060 |
| **PDGFRA** | 4.0256 | -0.929029768513881 | 0.01 | 0.282 | 6.74771990 |
| **CLMP** | 6.9846 | -1.02245365756731 | 0.235 | 0.541 | 1.17076244 |
| **BMPER** | 3.0400 | -1.19631191630914 | 0.018 | 0.292 | 5.09568148 |
| **HAND1** | 3.3085 | -1.67889028801779 | 0.023 | 0.296 | 5.54574542 |
| **ATF3** | 6.4001 | 1.07937440561898 | 0.767 | 0.535 | 1.07279002 |
| **KCNK12** | 2.2417 | -0.929193775876399 | 0.096 | 0.392 | 3.75754398 |
| **MAGI3** | 2.6580 | -0.940207457495955 | 0.154 | 0.46 | 4.45539524 |
| **PALLD** | 4.5530 | -0.921543141175748 | 0.174 | 0.486 | 7.63185920 |
| **MVD** | 2.8047 | -0.807057095445649 | 0.944 | 0.972 | 4.70126642 |
| **FLRT3** | 2.6724 | -1.25099135615541 | 0.1 | 0.39 | 4.47951545 |
| **GLI3** | 7.3853 | -0.941571055885111 | 0.222 | 0.537 | 1.23793845 |
| **NRP1** | 3.6236 | -1.2257661052294 | 0.01 | 0.268 | 6.07389021 |
| **GAL** | 2.1332 | -1.53707874788394 | 0.997 | 0.985 | 3.57574500 |
| **B4GALT6** | 1.3207 | -0.832758959751816 | 0.72 | 0.862 | 2.21383730 |
| **SRSF8** | 1.4394 | 1.03954097929894 | 0.856 | 0.683 | 2.41279538 |
| **TMEM158** | 7.1072 | -0.832938083342585 | 0.214 | 0.51 | 1.19131639 |
| **SLC2A3** | 1.2222 | -0.938568083146456 | 0.93 | 0.97 | 2.04876475 |

| | | | | | |
|---|---|---|---|---|---|
| **MSGN1** | 1.8596 | -1.96594654835651 | 0.008 | 0.251 | 3.11707569 |
| **ATP1B2** | 3.8074 | -0.934887747579041 | 0.381 | 0.634 | 6.38201315 |
| **TFPI2** | 1.3010 | 0.856799024759919 | 0.67 | 0.401 | 2.18087955 |
| **ANOS1** | 1.4671 | -1.25904334865263 | 0.106 | 0.377 | 2.45928228 |
| **RXRG** | 5.9164 | -0.927867575123499 | 0.005 | 0.242 | 9.91712452 |
| **ACTC1** | 5.2957 | 0.80062727149701 | 0.286 | 0.063 | 8.87675357 |
| **LIX1L** | 3.9096 | -0.966861363634773 | 0.724 | 0.847 | 6.55343809 |
| **BTG2** | 2.4225 | -0.830229450704546 | 0.389 | 0.676 | 4.06072809 |
| **SEMA3A** | 1.1864 | -0.954646404782199 | 0.366 | 0.603 | 1.98875893 |
| **ARRB1** | 4.4448 | -0.814011744632423 | 0.942 | 0.951 | 7.45037451 |
| **EDN1** | 1.4929 | -0.927762287451583 | 0.025 | 0.266 | 2.50245220 |
| **GATA4** | 1.0025 | -1.16056347989125 | 0.137 | 0.399 | 1.68044422 |
| **TCF7L1** | 6.5085 | -0.814019147523413 | 0.48 | 0.693 | 1.09097087 |
| **B3GNT7** | 6.0036 | -1.1351558277479 | 0.434 | 0.625 | 1.00633299 |
| **FAM110B** | 5.7948 | -0.870530810578271 | 0.271 | 0.534 | 9.71329382 |
| **LPGAT1** | 1.1818 | -0.803409737276513 | 0.581 | 0.763 | 1.98097314 |
| **GSC** | 5.1634 | -1.22510546665277 | 0.046 | 0.272 | 8.65497103 |
| **LIX1** | 2.6786 | -0.894984111267579 | 0.002 | 0.198 | 4.48999022 |
| **LRRTM1** | 6.9474 | -0.930700320671211 | 0.027 | 0.241 | 1.16453748 |
| **FAM222A** | 2.8422 | -1.16483928613452 | 0.465 | 0.647 | 4.76421086 |
| **CCN2** | 1.8796 | -0.969220971558506 | 0.733 | 0.867 | 3.15064081 |
| **PCDH19** | 8.1338 | -1.70037808193382 | 0.146 | 0.385 | 1.36338952 |
| **NDUFA4L2** | 9.9334 | -0.812696720693956 | 0.048 | 0.263 | 1.66504108 |
| **TFPI** | 4.1777 | -1.00217888843891 | 0.385 | 0.607 | 7.00277567 |
| **EPHA4** | 4.0815 | -1.1355757133317 | 0.181 | 0.407 | 6.84151411 |
| **TRIM43** | 9.0002 | 1.07380900947978 | 0.114 | 0.004 | 1.50861403 |
| **FAM89A** | 1.4221 | -1.39909576478441 | 0.084 | 0.293 | 2.38387034 |
| **CDKN1A** | 1.8796 | 0.854621507420338 | 0.779 | 0.564 | 3.15063441 |
| **SEMA3F** | 1.8676 | -0.990554085026123 | 0.435 | 0.625 | 3.13053334 |
| **JCAD** | 8.0077 | -0.808309692521726 | 0.048 | 0.249 | 1.34225268 |
| **LRRN1** | 1.0460 | -1.00322550102164 | 0.428 | 0.638 | 1.75333518 |
| **LIFR** | 2.8239 | -1.33729482222349 | 0.339 | 0.528 | 4.73351180 |
| **LGR5** | 1.1114 | -0.850785599325533 | 0.009 | 0.169 | 1.86295199 |
| **CNN1** | 6.5082 | -0.898059400061619 | 0.735 | 0.857 | 1.09091885 |

| | | | | | |
|---|---|---|---|---|---|
| **EMILIN2** | 3.5156 | -0.837018363846569 | 0.412 | 0.599 | 5.8929421 |
| **SFN** | 1.1484 | 0.808668148312327 | 0.356 | 0.168 | 1.9250669 |
| **SLC39A8** | 1.3827 | -1.11574918299998 | 0.451 | 0.581 | 2.3176966 |
| **CDH11** | 2.7039 | -1.72240157927311 | 0.134 | 0.312 | 4.5324067 |
| **ELL2** | 3.8340 | -0.959834494453 | 0.414 | 0.586 | 6.4266408 |
| **CBX4** | 8.3497 | 0.936855258501698 | 0.74 | 0.665 | 1.3995911 |
| **SEMA6D** | 5.6620 | -0.820302499183839 | 0.152 | 0.297 | 9.4907765 |
| **MT-CO3** | 2.5355 | -1.08821031726446 | 0.879 | 0.927 | 4.2501029 |
| **ANKRD1** | 6.8081 | -0.966077355459164 | 0.137 | 0.282 | 1.1411793 |
| **MT-CYB** | 2.9038 | -0.913718205828065 | 0.767 | 0.798 | 4.8674639 |
| **HAS2** | 3.1930 | -1.15762211684325 | 0.907 | 0.74 | 5.3521248 |
| **HAPLN1** | 4.3342 | -0.8901776306205 | 0.08 | 0.171 | 7.2650074 |
| **DLL3** | 3.1730 | -2.1539042934335 | 0.748 | 0.702 | 5.3185950 |
| **APELA** | 8.1847 | -1.11877130155855 | 0.549 | 0.569 | 1.3719211 |
| **S1PR3** | 6.2330 | -0.810286774808908 | 0.34 | 0.411 | 1.0447881 |
| **TC2N** | 1.2092 | 1.13008818790503 | 0.683 | 0.586 | 2.0268671 |
| **DACT1** | 6.0679 | -1.13028742455618 | 0.738 | 0.719 | 1.0171044 |
| **SLC34A2** | 1.8359 | 1.09311962322457 | 0.113 | 0.074 | 3.0773605 |
| **HPGD** | 3.0446 | -0.896613653719854 | 0.755 | 0.603 | 5.1034699 |
| **ACTG2** | 0.0104 | -0.957036745853618 | 0.662 | 0.651 | 1 |

Supplemental Table S3_Sheet_1

| Genes | estimate | log2FC | statistic | p.value | adj_p | conf.low | conf.high |
|---|---|---|---|---|---|---|---|
| **RPL3** | -0.123445226 | NA | -13.09359563 | 1.66E-06 | 0.009545691 | -0.145364594 | -0.101525857 |

Supplemental Table S3_Sheet_2

| Genes | estimate | log2FC | statistic | p.value | adj_p | conf.low | conf.high |
|---|---|---|---|---|---|---|---|
| **TMA7** | 4.094049936 | 2.033528699 | 33.22639904 | 1.83E-09 | 1.06E-05 | 3.80695831 | 4.381141561 |
| **RBM25** | -0.409188785 | NA | -16.07689812 | 2.91E-07 | 5.59E-04 | -0.468135358 | -0.350242213 |
| **SELENOS** | 1.8600808 | 0.895365292 | 18.97140062 | 2.67E-07 | 5.59E-04 | 1.628468125 | 2.091693475 |
| **ATP5ME** | 0.420757682 | -1.248938483 | 12.63286976 | 1.62E-06 | 1.16E-03 | 0.343782401 | 0.497732963 |
| **CCT3** | 0.267855246 | -1.900474545 | 13.53176873 | 1.16E-06 | 1.16E-03 | 0.221941339 | 0.313769152 |
| **FAM162A** | 3.950478233 | 1.982027312 | 12.94925805 | 1.27E-06 | 1.16E-03 | 3.246122761 | 4.654833705 |
| **HAGH** | -0.528399272 | NA | -12.93325511 | 1.48E-06 | 1.16E-03 | -0.622994092 | -0.433804453 |
| **RAPH1** | 0.681560881 | -0.553085562 | 16.5588923 | 9.86E-07 | 1.16E-03 | 0.583577585 | 0.779544177 |
| **CTHRC1** | 3.674614112 | 1.877592754 | 14.39776811 | 2.83E-06 | 1.81E-03 | 3.065160971 | 4.284067254 |
| **FXN** | -0.707350467 | NA | -11.36132113 | 3.40E-06 | 1.96E-03 | -0.85106739 | -0.563633543 |
| **ARHGDIA** | -0.328429005 | NA | -12.3682625 | 4.10E-06 | 2.15E-03 | -0.390857818 | -0.266000192 |
| **AKAP12** | -0.272781046 | NA | -13.8527431 | 6.48E-06 | 2.33E-03 | -0.320532949 | -0.225029142 |
| **EIF3I** | 0.196627623 | -2.346462081 | 10.7105681 | 6.41E-06 | 2.33E-03 | 0.154061145 | 0.239194102 |
| **NCAPD3** | -0.266106877 | NA | -10.72624901 | 5.58E-06 | 2.33E-03 | -0.323457532 | -0.208756223 |
| **NT5C3B** | 1.827469845 | 0.8698476 | 11.97750042 | 4.87E-06 | 2.33E-03 | 1.469204291 | 2.185735399 |
| **RPL18A** | 0.420249719 | -1.25068124 | 10.81400261 | 6.35E-06 | 2.33E-03 | 0.330010648 | 0.51048879 |
| **CDC20** | 1.175737821 | 0.233566388 | 11.88661103 | 6.98E-06 | 2.36E-03 | 0.941668476 | 1.409807167 |
| **GOLM2** | 2.485602494 | 1.313595594 | 10.5386892 | 7.90E-06 | 2.53E-03 | 1.937515045 | 3.033689943 |
| **TPX2** | 0.441733545 | -1.178751701 | 10.60114341 | 9.11E-06 | 2.76E-03 | 0.344446703 | 0.539020387 |
| **EXOSC4** | 0.620823923 | -0.687743943 | 10.9024635 | 9.63E-06 | 2.77E-03 | 0.487000139 | 0.754647707 |
| **RTN3** | 3.967540567 | 1.988244974 | 16.02608803 | 1.27E-05 | 3.48E-03 | 3.338244734 | 4.596836399 |
| **TBL2** | 0.496634624 | -1.00974325 | 9.572417406 | 1.35E-05 | 3.54E-03 | 0.376560582 | 0.616708666 |
| **ARID1A** | -0.204395491 | NA | -10.43977875 | 1.49E-05 | 3.59E-03 | -0.250586547 | -0.158204436 |
| **CS** | 0.24489933 | -2.029739267 | 9.205748978 | 1.61E-05 | 3.59E-03 | 0.183513134 | 0.306285526 |
| **RPS20** | 0.7019114 | -0.51063916 | 10.49655886 | 1.52E-05 | 3.59E-03 | 0.54388623 | 0.859936569 |
| **ZNF578** | 3.980195081 | 1.992839143 | 11.31445833 | 1.62E-05 | 3.59E-03 | 3.135277957 | 4.825112204 |
| **DNAJA1** | 0.570885095 | -0.808727698 | 9.09705703 | 2.02E-05 | 4.30E-03 | 0.425514199 | 0.716255992 |
| **PBXIP1** | -0.364173301 | NA | -8.860399311 | 2.09E-05 | 4.30E-03 | -0.458965227 | -0.269381376 |
| **ADRM1** | 0.459883047 | -1.120661079 | 12.16611926 | 2.54E-05 | 4.90E-03 | 0.366416921 | 0.553349173 |
| **NECTIN2** | -0.583490657 | NA | -9.544894652 | 2.64E-05 | 4.90E-03 | -0.727609481 | -0.439371833 |
| **PHF6** | -0.530173852 | NA | -8.700518413 | 2.59E-05 | 4.90E-03 | -0.671034161 | -0.389313544 |
| **AKAP1** | -0.263945927 | NA | -8.770266769 | 3.65E-05 | 5.15E-03 | -0.334356771 | -0.193535083 |
| **BCL7C** | -1.740666666 | NA | -8.558082572 | 3.16E-05 | 5.15E-03 | -2.211940778 | -1.269392554 |
| **DDX52** | 0.43868819 | -1.188732228 | 8.516472987 | 3.63E-05 | 5.15E-03 | 0.318953006 | 0.558423374 |
| **KDM3B** | -0.297565028 | NA | -8.322072 | 3.46E-05 | 5.15E-03 | -0.380150637 | -0.21497942 |
| **MIF** | 1.688642068 | 0.75586356 | 10.12276811 | 3.02E-05 | 5.15E-03 | 1.28887279 | 2.088411346 |
| **NTN1** | 4.935918903 | 2.303318691 | 16.49432342 | 3.05E-05 | 5.15E-03 | 4.14398721 | 5.727850597 |

| | | | | | | | |
|---|---|---|---|---|---|---|---|
| **POLR1F** | 2.863136295 | 1.517596349 | 8.477515198 | 3.38E-05 | 5.15E-03 | 2.080590929 | 3.645681662 |
| **RPL15** | 0.596543568 | -0.745300587 | 13.51551789 | 3.62E-05 | 5.15E-03 | 0.483530821 | 0.709556315 |
| **TPM4** | -0.360232863 | NA | -8.87668861 | 2.96E-05 | 5.15E-03 | -0.454806555 | -0.265659172 |
| **U2AF2** | -0.34866547 | NA | -8.592592206 | 3.67E-05 | 5.15E-03 | -0.443197776 | -0.254133164 |
| **DUSP3** | 0.593884655 | -0.751745338 | 10.3104366 | 3.86E-05 | 5.29E-03 | 0.454140726 | 0.733628584 |
| **SELENOH** | -0.81374166 | NA | -8.095697227 | 4.01E-05 | 5.37E-03 | -1.045530475 | -0.581952845 |
| **BET1** | 1.785724599 | 0.8365096 | 8.536855424 | 4.67E-05 | 5.56E-03 | 1.295327808 | 2.276121391 |
| **CALB1** | 1.262184409 | 0.335922708 | 11.24221348 | 4.60E-05 | 5.56E-03 | 0.982890501 | 1.541478317 |
| **CTPS1** | 0.281978094 | -1.826345007 | 8.434585446 | 5.07E-05 | 5.56E-03 | 0.20359531 | 0.360360878 |
| **FUBP3** | -0.282161742 | NA | -7.868489423 | 5.12E-05 | 5.56E-03 | -0.364959554 | -0.19936393 |
| **MAP1B** | -0.283511045 | NA | -8.959386678 | 4.39E-05 | 5.56E-03 | -0.358337155 | -0.208684934 |
| **NFYC** | 0.574761039 | -0.798965826 | 8.067093448 | 4.89E-05 | 5.56E-03 | 0.409568253 | 0.739953824 |
| **NUBP1** | 2.473381829 | 1.306484973 | 8.817876747 | 4.82E-05 | 5.56E-03 | 1.810373611 | 3.136390047 |
| **YY1** | 0.516779308 | -0.952379789 | 8.188098721 | 5.00E-05 | 5.56E-03 | 0.369851987 | 0.66370663 |
| **ZNF384** | 0.520179651 | -0.942918132 | 8.136707333 | 5.12E-05 | 5.56E-03 | 0.371448738 | 0.668910564 |
| **ZNF593** | 0.27773091 | -1.848240342 | 7.927947349 | 4.86E-05 | 5.56E-03 | 0.196842487 | 0.358619334 |
| **SUB1** | -0.369410207 | NA | -7.761341834 | 5.43E-05 | 5.79E-03 | -0.479167402 | -0.259653011 |
| **ACAD9** | 0.320150943 | -1.643175835 | 8.100826653 | 6.00E-05 | 6.16E-03 | 0.227814767 | 0.412487119 |
| **GPX7** | 4.466971689 | 2.159297111 | 14.93136674 | 6.31E-05 | 6.16E-03 | 3.664296863 | 5.269646515 |
| **GRN** | -0.457767873 | NA | -8.168985701 | 6.29E-05 | 6.16E-03 | -0.589157776 | -0.32637797 |
| **NADK2** | 0.837345107 | -0.25610575 | 7.745111532 | 6.63E-05 | 6.16E-03 | 0.586512823 | 1.088177391 |
| **NASP** | -0.325726869 | NA | -7.560660509 | 6.56E-05 | 6.16E-03 | -0.425083567 | -0.22637017 |
| **NATD1** | 2.752368114 | 1.460673435 | 11.44952614 | 6.34E-05 | 6.16E-03 | 2.143933653 | 3.360802576 |
| **SPINT2** | -0.648640378 | NA | -7.646138584 | 6.20E-05 | 6.16E-03 | -0.84443248 | -0.452848275 |
| **UPF1** | 0.131221018 | -2.929929282 | 8.083013586 | 6.42E-05 | 6.16E-03 | 0.093222375 | 0.16921966 |
| **CTNNB1** | 0.294337287 | -1.764457779 | 9.048841241 | 6.89E-05 | 6.30E-03 | 0.215998172 | 0.372676402 |
| **PEX14** | 3.114704102 | 1.639095113 | 9.540242406 | 7.18E-05 | 6.46E-03 | 2.31752524 | 3.911882963 |
| **ANXA11** | -0.196677214 | NA | -7.440320317 | 7.35E-05 | 6.51E-03 | -0.257638173 | -0.135716255 |
| **EXOSC5** | 3.783577049 | 1.919750825 | 10.97651192 | 7.49E-05 | 6.54E-03 | 2.912986047 | 4.654168051 |
| **MPHOSPH** | 2.555683546 | 1.353709208 | 8.544615426 | 7.63E-05 | 6.55E-03 | 1.842102486 | 3.269264606 |
| **ATP5MK** | -2.971430802 | NA | -13.57264064 | 8.40E-05 | 6.93E-03 | -3.554950319 | -2.387911286 |
| **HNRNPUL1** | -0.34886616 | NA | -8.561403679 | 8.24E-05 | 6.93E-03 | -0.44641421 | -0.251318109 |
| **UBAC2** | -0.506083317 | NA | -8.114384445 | 8.43E-05 | 6.93E-03 | -0.653630707 | -0.358535927 |
| **PPP1R10** | 0.306076832 | -1.708034247 | 8.872537326 | 8.70E-05 | 7.06E-03 | 0.222615685 | 0.389537979 |
| **ATAD3A** | -0.499922375 | NA | -9.294609561 | 9.76E-05 | 7.54E-03 | -0.632122051 | -0.367722698 |
| **CANX** | -0.421117111 | NA | -7.625621697 | 9.55E-05 | 7.54E-03 | -0.550417427 | -0.291816795 |
| **NAA50** | 0.525474614 | -0.928307026 | 7.7487195 | 1.01E-04 | 7.54E-03 | 0.365750914 | 0.685198314 |
| **PEBP1** | -0.359626987 | NA | -7.481658969 | 9.95E-05 | 7.54E-03 | -0.471817908 | -0.247436066 |
| **SET** | -0.643387209 | NA | -11.7776549 | 1.01E-04 | 7.54E-03 | -0.785585879 | -0.50118854 |

| | | | | | | | |
|---|---|---|---|---|---|---|---|
| **SF3B6** | -0.425090051 | NA | -7.684037711 | 9.82E-05 | 7.54E-03 | -0.554994344 | -0.295185758 |
| **DCAKD** | 0.513056736 | -0.962809721 | 7.098952753 | 1.02E-04 | 7.54E-03 | 0.346395286 | 0.679718186 |
| **HAX1** | -0.256447259 | NA | -9.869870279 | 1.08E-04 | 7.78E-03 | -0.321528523 | -0.191365996 |
| **UBE2S** | 0.529244285 | -0.917994308 | 8.819871924 | 1.07E-04 | 7.78E-03 | 0.383002105 | 0.675486466 |
| **TTC4** | 0.710631185 | -0.492827094 | 7.707219167 | 1.13E-04 | 8.00E-03 | 0.492833045 | 0.928429325 |
| **LEFTY2** | 0.538659735 | -0.892553866 | 7.010191929 | 1.22E-04 | 8.48E-03 | 0.360869404 | 0.716450067 |
| **PODXL** | -0.367893918 | NA | -6.982750957 | 1.21E-04 | 8.48E-03 | -0.489629821 | -0.246158016 |
| **SDCBP** | 0.480822655 | -1.056423221 | 7.687330473 | 1.24E-04 | 8.51E-03 | 0.332594301 | 0.629051009 |
| **RPS15A** | 0.47477985 | -1.074669388 | 6.992298854 | 1.31E-04 | 8.80E-03 | 0.317348239 | 0.632211461 |
| **SOX2** | -0.718686312 | NA | -6.849498748 | 1.31E-04 | 8.80E-03 | -0.960645243 | -0.47672738 |
| **CKAP2** | 0.589212535 | -0.763139973 | 7.040151022 | 1.45E-04 | 9.61E-03 | 0.394079513 | 0.784345556 |
| **MISP** | -0.538430656 | NA | -9.31246239 | 1.50E-04 | 9.71E-03 | -0.683435572 | -0.393425739 |
| **POLE** | 0.385908259 | -1.373670177 | 6.721354172 | 1.50E-04 | 9.71E-03 | 0.253495096 | 0.518321421 |
| **CRYAB** | 0.713854317 | -0.486298415 | 6.696852161 | 1.54E-04 | 9.84E-03 | 0.468010993 | 0.959697641 |
| **PRDM14** | -0.844182844 | NA | -7.454848613 | 1.57E-04 | 9.93E-03 | -1.113014404 | -0.575351285 |
| **DNAJB1** | -0.267303097 | NA | -8.474497366 | 1.63E-04 | 9.99E-03 | -0.344826988 | -0.189779206 |
| **PTGFRN** | 0.374863478 | -1.41556282 | 6.682707994 | 1.62E-04 | 9.99E-03 | 0.245318466 | 0.504408491 |
| **RBP1** | 0.291926521 | -1.776322811 | 6.663724003 | 1.63E-04 | 9.99E-03 | 0.190795949 | 0.393057093 |
| **DYNC1I2** | -0.347824381 | NA | -6.617672656 | 1.77E-04 | 1.07E-02 | -0.469320441 | -0.226328321 |
| **TAGLN** | 0.491569585 | -1.024532441 | 6.573964812 | 1.79E-04 | 1.07E-02 | 0.318940723 | 0.664198446 |
| **MYDGF** | 0.327541502 | -1.610250374 | 6.502345164 | 1.89E-04 | 1.12E-02 | 0.211337967 | 0.443745038 |
| **H4C1** | 0.734306575 | -0.445545577 | 8.395924234 | 1.96E-04 | 1.14E-02 | 0.517942666 | 0.950670484 |
| **POLR1A** | 0.195815207 | -2.352435284 | 8.202976919 | 1.94E-04 | 1.14E-02 | 0.137146647 | 0.254483767 |
| **ALDOA** | -0.400994838 | NA | -7.276249929 | 2.09E-04 | 1.15E-02 | -0.532619903 | -0.269369773 |
| **ALG11** | -1.364253414 | NA | -6.570699633 | 2.03E-04 | 1.15E-02 | -1.845902691 | -0.882604137 |
| **FLNA** | -0.12048512 | NA | -6.417980883 | 2.05E-04 | 1.15E-02 | -0.163775961 | -0.07719428 |
| **RBMX** | 1.230671394 | 0.299445594 | 6.485332087 | 2.09E-04 | 1.15E-02 | 0.791492098 | 1.66985069 |
| **RPL7** | 0.183362252 | -2.447231429 | 6.496396434 | 2.07E-04 | 1.15E-02 | 0.118033592 | 0.248690911 |
| **TOP3B** | -0.451290815 | NA | -6.437842646 | 2.01E-04 | 1.15E-02 | -0.612951357 | -0.289630272 |
| **DPPA4** | -0.796493989 | NA | -8.367590124 | 2.13E-04 | 1.16E-02 | -1.032762166 | -0.560225811 |
| **PYM1** | -0.234401123 | NA | -7.156467241 | 2.24E-04 | 1.20E-02 | -0.312513163 | -0.156289082 |
| **BACH1** | 1.06725206 | 0.093900948 | 6.410580066 | 2.26E-04 | 1.20E-02 | 0.681954081 | 1.45255004 |
| **ANP32A** | -0.94052929 | NA | -7.357329254 | 2.32E-04 | 1.22E-02 | -1.248231177 | -0.632827403 |
| **HSPA8** | 0.187186189 | -2.417454101 | 6.899565777 | 2.40E-04 | 1.26E-02 | 0.12292373 | 0.251448648 |
| **CRNKL1** | 0.172804398 | -2.53278816 | 6.250905919 | 2.49E-04 | 1.29E-02 | 0.109012565 | 0.236596231 |
| **MCM6** | 0.234021936 | -2.095284328 | 6.710418227 | 2.56E-04 | 1.32E-02 | 0.151820794 | 0.316223078 |
| **ATP1B2** | 2.143353995 | 1.099870144 | 9.446912453 | 2.61E-04 | 1.33E-02 | 1.555140451 | 2.731567539 |
| **BTK** | 0.493631763 | -1.018492868 | 6.380863984 | 2.75E-04 | 1.35E-02 | 0.313301085 | 0.67396244 |
| **ERH** | -0.646914291 | NA | -6.862340305 | 2.74E-04 | 1.35E-02 | -0.871220055 | -0.422608528 |

| | | | | | | | |
|---|---|---|---|---|---|---|---|
| **FAH** | 0.831787151 | -0.265713696 | 6.560103639 | 2.67E-04 | 1.35E-02 | 0.534298889 | 1.129275412 |
| **FBL** | -0.201566924 | NA | -6.179271737 | 2.75E-04 | 1.35E-02 | -0.276900818 | -0.126233029 |
| **NUP93** | 0.307989987 | -1.699044647 | 6.716395438 | 2.76E-04 | 1.35E-02 | 0.199513194 | 0.416466779 |
| **MAP2K7** | 0.541423657 | -0.88517017 | 6.285699035 | 2.86E-04 | 1.39E-02 | 0.341151102 | 0.741696211 |
| **CC2D1A** | -0.278659654 | NA | -8.470704754 | 2.94E-04 | 1.39E-02 | -0.362002591 | -0.195316717 |
| **ISOC2** | -0.537289746 | NA | -6.167495395 | 2.92E-04 | 1.39E-02 | -0.738907084 | -0.335672408 |
| **RPL13** | 0.316172127 | -1.661217907 | 9.335588941 | 2.97E-04 | 1.39E-02 | 0.227978728 | 0.404365525 |
| **RPL6** | 0.22843142 | -2.130166991 | 6.101989821 | 2.99E-04 | 1.39E-02 | 0.141980872 | 0.314881969 |
| **S100A4** | 2.576604497 | 1.365471104 | 9.023763197 | 2.95E-04 | 1.39E-02 | 1.840210962 | 3.312998031 |
| **DLG3** | -0.474636924 | NA | -8.845855573 | 3.09E-04 | 1.41E-02 | -0.612624308 | -0.33664954 |
| **RPS14** | 0.187767403 | -2.412981467 | 8.329173958 | 3.08E-04 | 1.41E-02 | 0.130778559 | 0.244756247 |
| **BASP1** | -0.535504845 | NA | -6.825505076 | 3.16E-04 | 1.43E-02 | -0.723169387 | -0.347840303 |
| **FAM114A2** | 0.734391299 | -0.445379128 | 6.359018745 | 3.21E-04 | 1.45E-02 | 0.463561304 | 1.005221295 |
| **MTR** | 0.331112301 | -1.594607485 | 6.178031974 | 3.25E-04 | 1.45E-02 | 0.20642985 | 0.455794753 |
| **TMSB10** | -1.113656713 | NA | -6.556010872 | 3.34E-04 | 1.48E-02 | -1.516318281 | -0.710995144 |
| **EIPR1** | 0.57499349 | -0.798382474 | 8.932624355 | 3.42E-04 | 1.49E-02 | 0.40798306 | 0.742003919 |
| **LRRC47** | 0.22775442 | -2.134449041 | 6.026901226 | 3.45E-04 | 1.49E-02 | 0.140242051 | 0.315266789 |
| **RALGPS2** | 1.055404819 | 0.077796476 | 6.113003425 | 3.44E-04 | 1.49E-02 | 0.653951853 | 1.456857785 |
| **EWSR1** | -0.385532741 | NA | -7.031366922 | 3.72E-04 | 1.57E-02 | -0.518928928 | -0.252136555 |
| **RUVBL2** | 0.191693758 | -2.383124738 | 5.931410303 | 3.65E-04 | 1.57E-02 | 0.117019119 | 0.266368396 |
| **UNG** | 2.269143444 | 1.182147812 | 7.348050443 | 3.73E-04 | 1.57E-02 | 1.507952193 | 3.030334695 |
| **VPS45** | 0.593617925 | -0.752393438 | 5.905859946 | 3.70E-04 | 1.57E-02 | 0.361538102 | 0.825697748 |
| **LDHD** | -0.292783137 | NA | -5.885595531 | 3.77E-04 | 1.57E-02 | -0.407625185 | -0.177941088 |
| **CASC3** | -0.402437728 | NA | -5.906255826 | 3.80E-04 | 1.57E-02 | -0.55996373 | -0.244911727 |
| **HMGB2** | -0.371371935 | NA | -5.919108429 | 3.89E-04 | 1.60E-02 | -0.516677468 | -0.226066403 |
| **RPS3** | 0.332895365 | -1.586859311 | 6.193245938 | 3.96E-04 | 1.62E-02 | 0.206569248 | 0.459221482 |
| **CTSL** | 0.643522216 | -0.635938139 | 5.84623348 | 4.03E-04 | 1.62E-02 | 0.389134059 | 0.897910373 |
| **DCAF5** | -0.773587803 | NA | -6.856850553 | 4.01E-04 | 1.62E-02 | -1.047139318 | -0.500036289 |
| **PSMA7** | 0.194110054 | -2.36505325 | 5.809824358 | 4.08E-04 | 1.63E-02 | 0.116997831 | 0.271222277 |
| **CBX1** | -0.479773026 | NA | -5.923055849 | 4.16E-04 | 1.64E-02 | -0.668008445 | -0.291537608 |
| **PSMB3** | 0.36728018 | -1.445047052 | 6.343288934 | 4.13E-04 | 1.64E-02 | 0.229926749 | 0.50463361 |
| **BUD31** | 0.474033777 | -1.076938233 | 5.788470496 | 4.20E-04 | 1.64E-02 | 0.28498968 | 0.663077875 |
| **PURB** | 0.691692645 | -0.53179698 | 5.770192056 | 4.25E-04 | 1.64E-02 | 0.415082447 | 0.968302842 |
| **ZCCHC8** | -0.332292291 | NA | -6.093559231 | 4.26E-04 | 1.64E-02 | -0.460263785 | -0.204320797 |
| **GNG12** | -0.643660178 | NA | -6.270476613 | 4.32E-04 | 1.66E-02 | -0.886869027 | -0.400451329 |
| **ALDOC** | -0.355646011 | NA | -5.857498467 | 4.67E-04 | 1.67E-02 | -0.497041148 | -0.214250874 |
| **CKS1B** | 0.337570156 | -1.56674073 | 5.802990375 | 4.52E-04 | 1.67E-02 | 0.202710944 | 0.472429369 |
| **COMMD5** | 0.267682706 | -1.90140416 | 5.950680552 | 4.64E-04 | 1.67E-02 | 0.162433238 | 0.372932174 |
| **GTPBP4** | 0.484044023 | -1.04678983 | 7.212384002 | 4.66E-04 | 1.67E-02 | 0.317426961 | 0.650661086 |

| | | | | | | | |
|---|---|---|---|---|---|---|---|
| **GTPBP6** | 0.300194298 | -1.736031519 | 6.404114734 | 4.58E-04 | 1.67E-02 | 0.188076658 | 0.412311939 |
| **NFKB2** | -0.285351164 | NA | -6.273215253 | 4.58E-04 | 1.67E-02 | -0.393462839 | -0.17723949 |
| **NUP62** | -0.328128397 | NA | -5.81098101 | 4.63E-04 | 1.67E-02 | -0.459245849 | -0.197010945 |
| **PDAP1** | -0.475386222 | NA | -7.167532765 | 4.48E-04 | 1.67E-02 | -0.639356842 | -0.311415602 |
| **RPS11** | 0.153732434 | -2.701506518 | 6.164747819 | 4.64E-04 | 1.67E-02 | 0.094745988 | 0.212718881 |
| **SIKE1** | 0.853795555 | -0.228037443 | 5.860171707 | 4.48E-04 | 1.67E-02 | 0.515127339 | 1.192463771 |
| **ZC3H14** | -0.120560234 | NA | -5.891504174 | 4.39E-04 | 1.67E-02 | -0.168157361 | -0.072963107 |
| **SLC9A1** | 0.716542965 | -0.480874882 | 6.286290975 | 4.77E-04 | 1.70E-02 | 0.444893295 | 0.988192634 |
| **H1-6** | 0.670664128 | -0.576337657 | 5.64516995 | 4.95E-04 | 1.75E-02 | 0.396414054 | 0.944914202 |
| **EIF2B4** | 0.390061829 | -1.358225269 | 6.636573321 | 4.98E-04 | 1.75E-02 | 0.247280633 | 0.532843025 |
| **CALR** | 0.462874452 | -1.111307159 | 5.654846316 | 5.09E-04 | 1.75E-02 | 0.273551427 | 0.652197477 |
| **CCDC43** | 0.446735835 | -1.162506108 | 5.821189511 | 5.13E-04 | 1.75E-02 | 0.267512644 | 0.625959027 |
| **MOB4** | 0.399713573 | -1.32296153 | 8.531565052 | 5.19E-04 | 1.75E-02 | 0.27648081 | 0.522946337 |
| **OLFML3** | -0.508449292 | NA | -5.843796336 | 5.07E-04 | 1.75E-02 | -0.711763555 | -0.305135029 |
| **POLR2E** | 0.19495626 | -2.358777614 | 5.726974596 | 5.11E-04 | 1.75E-02 | 0.115887383 | 0.274025137 |
| **SRSF5** | -0.221780046 | NA | -5.607399108 | 5.19E-04 | 1.75E-02 | -0.313096144 | -0.130463948 |
| **VAV2** | -0.401125606 | NA | -6.478783528 | 5.18E-04 | 1.75E-02 | -0.550741599 | -0.251509613 |
| **RRP8** | -0.252761018 | NA | -6.717695367 | 5.25E-04 | 1.76E-02 | -0.344789378 | -0.160732657 |
| **HNRNPA1** | 0.386478745 | -1.371539021 | 6.066997289 | 5.28E-04 | 1.76E-02 | 0.235525734 | 0.537431756 |
| **ALAD** | 0.361970631 | -1.466055449 | 6.016619611 | 5.32E-04 | 1.76E-02 | 0.219724958 | 0.504216304 |
| **ARPC1B** | 0.356582426 | -1.487692493 | 6.582343969 | 5.49E-04 | 1.81E-02 | 0.22458593 | 0.488578922 |
| **CKAP2L** | 1.144621012 | 0.194869996 | 7.602969665 | 5.65E-04 | 1.85E-02 | 0.760239575 | 1.529002449 |
| **GTF3A** | 1.342643383 | 0.425076165 | 5.511183788 | 5.92E-04 | 1.93E-02 | 0.779586481 | 1.905700286 |
| **BUB1** | 0.53728548 | -0.896239246 | 6.199847843 | 6.06E-04 | 1.96E-02 | 0.32894083 | 0.745630129 |
| **CHAMP1** | -0.266551169 | NA | -5.69123029 | 6.12E-04 | 1.96E-02 | -0.376133073 | -0.156969266 |
| **ITSN1** | -0.358123128 | NA | -6.696351651 | 6.15E-04 | 1.96E-02 | -0.490029823 | -0.226216434 |
| **SMS** | -0.306713202 | NA | -5.562490436 | 6.10E-04 | 1.96E-02 | -0.434730614 | -0.178695791 |
| **ATG16L1** | 0.221407088 | -2.175226686 | 5.438923663 | 6.22E-04 | 1.97E-02 | 0.127493779 | 0.315320398 |
| **ARHGAP12** | 2.443787766 | 1.289118998 | 9.113042376 | 6.38E-04 | 1.99E-02 | 1.713035583 | 3.174539949 |
| **PITHD1** | 0.518972148 | -0.94627098 | 6.296977493 | 6.38E-04 | 1.99E-02 | 0.319235451 | 0.718708845 |
| **RPS16** | -0.291584431 | NA | -5.822703256 | 6.44E-04 | 1.99E-02 | -0.409952647 | -0.173216215 |
| **YJU2** | -0.696777147 | NA | -6.103083 | 6.44E-04 | 1.99E-02 | -0.970824764 | -0.42272953 |
| **CDC40** | 0.297819014 | -1.747492229 | 5.443733393 | 6.76E-04 | 2.06E-02 | 0.171022766 | 0.424615263 |
| **GLYR1** | -0.281128773 | NA | -7.212323486 | 6.73E-04 | 2.06E-02 | -0.380149531 | -0.182108015 |
| **ZYX** | -0.291013 | NA | -5.502242346 | 6.76E-04 | 2.06E-02 | -0.414038177 | -0.167987824 |
| **AIDA** | -0.453817343 | NA | -5.94398159 | 7.00E-04 | 2.12E-02 | -0.636377202 | -0.271257484 |
| **EIF3B** | 0.182345382 | -2.455254435 | 5.349251156 | 7.10E-04 | 2.12E-02 | 0.103600328 | 0.261090435 |
| **GPD2** | 0.210349994 | -2.249136321 | 6.379680527 | 7.05E-04 | 2.12E-02 | 0.129615936 | 0.291084051 |
| **PAWR** | -0.500314746 | NA | -5.463833188 | 7.07E-04 | 2.12E-02 | -0.713318467 | -0.287311024 |

| | | | | | | | |
|---|---|---|---|---|---|---|---|
| **ERVK-5** | -0.448701674 | NA | -5.699863641 | 7.18E-04 | 2.12E-02 | -0.634596304 | -0.262807043 |
| **RPL4** | 0.209323927 | -2.256190868 | 5.311694192 | 7.19E-04 | 2.12E-02 | 0.118444026 | 0.300203827 |
| **ANAPC2** | 0.545080443 | -0.875458936 | 6.188322257 | 7.49E-04 | 2.17E-02 | 0.330766112 | 0.759394774 |
| **H1-5** | 0.392586421 | -1.348917823 | 5.406249933 | 7.51E-04 | 2.17E-02 | 0.223722449 | 0.561450392 |
| **PRKAR2A** | 0.39954691 | -1.323563196 | 6.378297348 | 7.43E-04 | 2.17E-02 | 0.245673109 | 0.553420712 |
| **SMARCC1** | -0.286351245 | NA | -5.281161901 | 7.47E-04 | 2.17E-02 | -0.411400907 | -0.161301583 |
| **TPP1** | -0.87931497 | NA | -6.807766128 | 7.57E-04 | 2.18E-02 | -1.203883259 | -0.554746681 |
| **HNRNPA2E** | -0.329822871 | NA | -6.405830488 | 7.83E-04 | 2.24E-02 | -0.456902541 | -0.202743202 |
| **TMEM230** | 1.493298402 | 0.578502484 | 6.565815339 | 7.85E-04 | 2.24E-02 | 0.927207296 | 2.059389508 |
| **GSPT1** | 0.387731671 | -1.366869513 | 5.228880215 | 8.04E-04 | 2.28E-02 | 0.216624986 | 0.558838355 |
| **BROX** | 0.784063808 | -0.350957027 | 6.341302483 | 8.16E-04 | 2.29E-02 | 0.479103726 | 1.08902389 |
| **MFAP1** | -0.510977271 | NA | -5.225073721 | 8.13E-04 | 2.29E-02 | -0.736719742 | -0.285234801 |
| **CTTN** | -0.305541325 | NA | -5.97527015 | 8.31E-04 | 2.29E-02 | -0.429298916 | -0.181783735 |
| **RCN2** | -0.375140189 | NA | -5.236817629 | 8.37E-04 | 2.29E-02 | -0.540894282 | -0.209386096 |
| **RPL10** | -0.281538865 | NA | -5.194218832 | 8.29E-04 | 2.29E-02 | -0.406538868 | -0.156538862 |
| **RUVBL1** | 0.164055623 | -2.607743053 | 5.252346487 | 8.36E-04 | 2.29E-02 | 0.091713066 | 0.236398179 |
| **ZHX2** | 0.445971154 | -1.164977698 | 6.657804981 | 8.30E-04 | 2.29E-02 | 0.277847924 | 0.614094383 |
| **TPR** | -0.142294102 | NA | -5.204584853 | 8.45E-04 | 2.31E-02 | -0.205452221 | -0.079135983 |
| **EIF2S3** | -0.356208494 | NA | -6.845561638 | 8.71E-04 | 2.31E-02 | -0.488472327 | -0.223944661 |
| **ELP1** | 0.402769365 | -1.311974139 | 7.12397171 | 8.65E-04 | 2.31E-02 | 0.25719079 | 0.548347941 |
| **LARP7** | 0.303161817 | -1.721840036 | 5.371683492 | 8.67E-04 | 2.31E-02 | 0.171146929 | 0.435176705 |
| **RPS23** | 0.373983697 | -1.418952714 | 7.263861221 | 8.56E-04 | 2.31E-02 | 0.240658359 | 0.507309035 |
| **SACS** | 0.449339512 | -1.154122167 | 5.21002474 | 8.65E-04 | 2.31E-02 | 0.249770821 | 0.648908203 |
| **SDAD1** | 0.207253148 | -2.270534076 | 5.161963431 | 8.62E-04 | 2.31E-02 | 0.114663438 | 0.299842859 |
| **ANP32B** | -0.682393782 | NA | -5.414296034 | 8.89E-04 | 2.35E-02 | -0.978455459 | -0.386332105 |
| **GNG5** | 0.640007585 | -0.643839091 | 6.033128512 | 9.07E-04 | 2.39E-02 | 0.380984046 | 0.899031125 |
| **CCT4** | 0.244639697 | -2.031269568 | 5.451177888 | 9.33E-04 | 2.43E-02 | 0.13866557 | 0.350613825 |
| **CDCA7L** | 0.592729409 | -0.754554454 | 5.656355301 | 9.40E-04 | 2.43E-02 | 0.341949804 | 0.843509015 |
| **CNOT2** | -0.399475861 | NA | -6.816832708 | 9.40E-04 | 2.43E-02 | -0.549042312 | -0.249909409 |
| **SUMO2** | -0.861489335 | NA | -6.48516155 | 9.40E-04 | 2.43E-02 | -1.194802138 | -0.528176533 |
| **CLASP2** | 0.309214419 | -1.693320498 | 5.226915851 | 9.63E-04 | 2.46E-02 | 0.171322166 | 0.447106673 |
| **DPYSL3** | -0.191203261 | NA | -5.409790399 | 9.75E-04 | 2.46E-02 | -0.274661136 | -0.107745385 |
| **FLNB** | -0.152993933 | NA | -5.510011571 | 9.76E-04 | 2.46E-02 | -0.218989668 | -0.086998198 |
| **GTF2B** | 0.316127053 | -1.661423596 | 5.077132938 | 9.72E-04 | 2.46E-02 | 0.172411075 | 0.45984303 |
| **PPT1** | -0.384672029 | NA | -5.484051064 | 9.73E-04 | 2.46E-02 | -0.551076734 | -0.218267323 |
| **SRSF1** | -0.185077624 | NA | -5.225701198 | 9.86E-04 | 2.46E-02 | -0.267737436 | -0.102417813 |
| **UBE2D3** | 0.160711486 | -2.637455057 | 5.214474243 | 9.87E-04 | 2.46E-02 | 0.088829149 | 0.232593822 |
| **UBR2** | 0.328330696 | -1.606778459 | 5.441703781 | 9.87E-04 | 2.46E-02 | 0.185454137 | 0.471207256 |
| **TSPYL1** | 0.489402442 | -1.030906794 | 5.898088591 | 9.96E-04 | 2.47E-02 | 0.287151447 | 0.691653438 |

| | | | | | | | |
|---|---|---|---|---|---|---|---|
| **DENND4C** | 0.299579183 | -1.738990719 | 5.884451203 | 1.02E-03 | 2.52E-02 | 0.175386845 | 0.42377152 |
| **XRCC1** | 0.147862897 | -2.75766801 | 5.45547198 | 1.02E-03 | 2.52E-02 | 0.083483086 | 0.212242708 |
| **CNNM4** | 0.348580668 | -1.520435534 | 5.075155003 | 1.08E-03 | 2.63E-02 | 0.189097272 | 0.508064064 |
| **LAP3** | -0.124262402 | NA | -5.754679537 | 1.08E-03 | 2.63E-02 | -0.176707393 | -0.071817412 |
| **CBFA2T2** | -0.496850129 | NA | -5.028264635 | 1.12E-03 | 2.65E-02 | -0.725959891 | -0.267740366 |
| **COA3** | -0.254041507 | NA | -5.011893606 | 1.10E-03 | 2.65E-02 | -0.371337736 | -0.136745278 |
| **EDF1** | -0.684604424 | NA | -6.596340268 | 1.12E-03 | 2.65E-02 | -0.949900741 | -0.419308108 |
| **GYG1** | 0.387825738 | -1.366519544 | 5.068939472 | 1.09E-03 | 2.65E-02 | 0.210130175 | 0.565521302 |
| **HCCS** | -0.58690138 | NA | -6.548485883 | 1.11E-03 | 2.65E-02 | -0.81522642 | -0.358576341 |
| **POLR3B** | 0.232103233 | -2.107161475 | 5.173695899 | 1.13E-03 | 2.65E-02 | 0.126935821 | 0.337270646 |
| **SH3BP5L** | 0.396548245 | -1.334431696 | 4.955630076 | 1.12E-03 | 2.65E-02 | 0.21195322 | 0.581143271 |
| **TFIP11** | 0.259693453 | -1.945118452 | 5.066836685 | 1.12E-03 | 2.65E-02 | 0.140516286 | 0.37887062 |
| **TNPO3** | 0.48606776 | -1.04077065 | 5.216129163 | 1.13E-03 | 2.65E-02 | 0.266968994 | 0.705166525 |
| **ACP1** | -0.210974415 | NA | -5.983844705 | 1.14E-03 | 2.66E-02 | -0.298163765 | -0.123785065 |
| **HSPA4L** | 0.248377217 | -2.00939525 | 4.957892685 | 1.14E-03 | 2.66E-02 | 0.132686456 | 0.364067978 |
| **CLINT1** | -0.406438394 | NA | -5.464032385 | 1.17E-03 | 2.67E-02 | -0.584729677 | -0.22814711 |
| **DYNLL1** | -1.150582673 | NA | -5.102872186 | 1.15E-03 | 2.67E-02 | -1.677330179 | -0.623835167 |
| **RAB21** | 0.322014943 | -1.634800459 | 4.975501734 | 1.16E-03 | 2.67E-02 | 0.172153576 | 0.471876309 |
| **RTCB** | -0.261359482 | NA | -5.390222663 | 1.16E-03 | 2.67E-02 | -0.376977206 | -0.145741757 |
| **YWHAZ** | 0.214060124 | -2.223912029 | 7.392973367 | 1.16E-03 | 2.67E-02 | 0.136796154 | 0.291324093 |
| **GSTCD** | 0.487223304 | -1.037344956 | 5.312462466 | 1.17E-03 | 2.67E-02 | 0.269564318 | 0.70488229 |
| **RPL3** | -0.159796764 | NA | -6.794590822 | 1.18E-03 | 2.67E-02 | -0.220793185 | -0.098800343 |
| **ACO2** | -0.121545777 | NA | -7.714740932 | 1.19E-03 | 2.68E-02 | -0.164299047 | -0.078792507 |
| **MCM5** | 0.202509278 | -2.303940089 | 6.103130064 | 1.19E-03 | 2.69E-02 | 0.119604853 | 0.285413702 |
| **HDAC3** | 0.52910422 | -0.918376169 | 4.938724478 | 1.21E-03 | 2.70E-02 | 0.281183011 | 0.77702543 |
| **HCFC1** | -0.122856456 | NA | -6.339021298 | 1.22E-03 | 2.71E-02 | -0.172012781 | -0.073700131 |
| **EEF1A2** | -1.594235976 | NA | -5.486835399 | 1.22E-03 | 2.72E-02 | -2.293898828 | -0.894573123 |
| **RPL7L1** | 0.408764962 | -1.290656555 | 5.007030274 | 1.24E-03 | 2.75E-02 | 0.21851813 | 0.599011795 |
| **THAP11** | -0.77900477 | NA | -7.509017789 | 1.25E-03 | 2.75E-02 | -1.059122576 | -0.498886965 |
| **U2AF1** | -0.911324365 | NA | -5.640257487 | 1.25E-03 | 2.75E-02 | -1.304994519 | -0.517654211 |
| **ARMC8** | 0.483849834 | -1.047368726 | 5.960608418 | 1.26E-03 | 2.76E-02 | 0.28197712 | 0.685722549 |
| **AFG2B** | 0.407594921 | -1.294792021 | 4.903390095 | 1.29E-03 | 2.78E-02 | 0.214985873 | 0.600203969 |
| **DEPTOR** | -0.389194588 | NA | -4.938717486 | 1.29E-03 | 2.78E-02 | -0.572280432 | -0.206108743 |
| **HNRNPU** | 0.215888162 | -2.211643959 | 4.896095923 | 1.29E-03 | 2.78E-02 | 0.113768468 | 0.318007856 |
| **VAPA** | -0.434529169 | NA | -5.874019991 | 1.28E-03 | 2.78E-02 | -0.617757827 | -0.251300512 |
| **PSMB5** | 0.355704014 | -1.491250839 | 5.083068031 | 1.32E-03 | 2.84E-02 | 0.191044069 | 0.520363959 |
| **HCLS1** | -0.351536789 | NA | -5.214670399 | 1.37E-03 | 2.93E-02 | -0.512076806 | -0.190996772 |
| **TOMM7** | 2.520647883 | 1.333794598 | 7.556794454 | 1.38E-03 | 2.93E-02 | 1.610088365 | 3.4312074 |
| **DBN1** | -0.355907447 | NA | -4.987930189 | 1.40E-03 | 2.98E-02 | -0.523233194 | -0.1885817 |

| | | | | | | | |
|---|---|---|---|---|---|---|---|
| **3,00 ARL** | 0.380135463 | -1.395414475 | 5.199694629 | 1.46E-03 | 3.07E-02 | 0.205449419 | 0.554821506 |
| **CRABP1** | 0.520763301 | -0.941300313 | 4.74813802 | 1.46E-03 | 3.07E-02 | 0.267713907 | 0.773812694 |
| **HTRA2** | 0.661316864 | -0.596586404 | 4.927482267 | 1.47E-03 | 3.07E-02 | 0.347148772 | 0.975484956 |
| **PDIA5** | 0.318412967 | -1.651029004 | 5.274118925 | 1.47E-03 | 3.07E-02 | 0.173363907 | 0.463462028 |
| **HSD17B11** | 0.965261862 | -0.051007716 | 6.049130678 | 1.49E-03 | 3.07E-02 | 0.561123496 | 1.369400229 |
| **IGF2BP3** | -0.210544768 | NA | -5.195216826 | 1.48E-03 | 3.07E-02 | -0.307441647 | -0.113647889 |
| **PCGF6** | 1.794829081 | 0.843846465 | 5.894054674 | 1.48E-03 | 3.07E-02 | 1.031454538 | 2.558203624 |
| **POP1** | -0.186441475 | NA | -4.734101211 | 1.48E-03 | 3.07E-02 | -0.277292764 | -0.095590187 |
| **PACS1** | -0.341744485 | NA | -4.79626747 | 1.49E-03 | 3.07E-02 | -0.50702342 | -0.176465549 |
| **CNOT3** | -0.385890767 | NA | -5.12206577 | 1.51E-03 | 3.09E-02 | -0.565259814 | -0.206521721 |
| **12,00 LSM** | 0.280846266 | -1.832147473 | 4.75415586 | 1.52E-03 | 3.09E-02 | 0.144162281 | 0.417530251 |
| **NCDN** | 0.309275441 | -1.693035818 | 5.520672427 | 1.52E-03 | 3.10E-02 | 0.171947814 | 0.446603068 |
| **HMGB1** | -0.789070914 | NA | -6.486694772 | 1.53E-03 | 3.11E-02 | -1.106071298 | -0.472070531 |
| **SFPQ** | 0.152836694 | -2.70993714 | 4.953192661 | 1.55E-03 | 3.12E-02 | 0.080192947 | 0.225480441 |
| **SMAP1** | -2.509661321 | NA | -6.930700848 | 1.55E-03 | 3.12E-02 | -3.477672204 | -1.541650439 |
| **SSR3** | -0.484747476 | NA | -4.685091785 | 1.57E-03 | 3.15E-02 | -0.723343121 | -0.246151832 |
| **CCDC51** | -0.327414135 | NA | -4.710566968 | 1.58E-03 | 3.16E-02 | -0.488097711 | -0.16673056 |
| **KLC1** | -0.198370694 | NA | -4.692263415 | 1.61E-03 | 3.19E-02 | -0.296052516 | -0.100688872 |
| **NBN** | 0.200212988 | -2.32039253 | 4.868879364 | 1.61E-03 | 3.19E-02 | 0.103808219 | 0.296617757 |
| **SERPINH1** | 0.313831573 | -1.671937592 | 4.933823344 | 1.61E-03 | 3.19E-02 | 0.163910818 | 0.463752328 |
| **SNRNP70** | -0.1287416 | NA | -4.725376398 | 1.62E-03 | 3.19E-02 | -0.191902925 | -0.065580275 |
| **FAU** | 0.611136203 | -0.710434148 | 6.392842515 | 1.64E-03 | 3.20E-02 | 0.361914957 | 0.860357449 |
| **LYPLA2** | 0.237180119 | -2.075945011 | 4.710566005 | 1.64E-03 | 3.20E-02 | 0.120515961 | 0.353844277 |
| **THRAP3** | -0.228372484 | NA | -5.820236782 | 1.64E-03 | 3.20E-02 | -0.327093666 | -0.129651303 |
| **WDR1** | 0.101383005 | -3.302112262 | 4.660456385 | 1.64E-03 | 3.20E-02 | 0.05117584 | 0.15159017 |
| **NXF1** | 0.123257342 | -3.020254515 | 5.056281442 | 1.66E-03 | 3.22E-02 | 0.065122847 | 0.181391836 |
| **TCEA1** | -0.191614471 | NA | -5.346893058 | 1.68E-03 | 3.23E-02 | -0.279015382 | -0.10421356 |
| **ZW10** | 0.639875491 | -0.644136886 | 4.82431968 | 1.68E-03 | 3.23E-02 | 0.329103555 | 0.950647427 |
| **HEXA** | 0.230247536 | -2.118742379 | 4.663805274 | 1.71E-03 | 3.28E-02 | 0.115987897 | 0.344507175 |
| **GJA1** | -0.649966082 | NA | -4.768055761 | 1.72E-03 | 3.28E-02 | -0.968396185 | -0.331535979 |
| **VPS26B** | 1.049403387 | 0.069569352 | 6.756743362 | 1.73E-03 | 3.28E-02 | 0.633988059 | 1.464818715 |
| **ZNHIT6** | 0.557239049 | -0.843631736 | 4.66646672 | 1.73E-03 | 3.28E-02 | 0.28058269 | 0.833895408 |
| **CENPF** | 0.443740425 | -1.172212107 | 6.051205736 | 1.74E-03 | 3.30E-02 | 0.255558876 | 0.631921974 |
| **TK1** | 0.598653817 | -0.740206118 | 5.192247157 | 1.75E-03 | 3.31E-02 | 0.319696144 | 0.877611489 |
| **CDH1** | -0.642685043 | NA | -4.917375267 | 1.77E-03 | 3.32E-02 | -0.952380631 | -0.332989455 |
| **VCP** | -0.171679589 | NA | -4.599083119 | 1.77E-03 | 3.32E-02 | -0.257789422 | -0.085569756 |
| **ITPR3** | -0.716130256 | NA | -4.604770775 | 1.78E-03 | 3.33E-02 | -1.075223093 | -0.357037419 |
| **ATP5MG** | 0.361765624 | -1.466872771 | 5.451086924 | 1.82E-03 | 3.38E-02 | 0.197647866 | 0.525883381 |
| **CHERP** | 0.20755801 | -2.268413487 | 5.522291761 | 1.82E-03 | 3.38E-02 | 0.114147781 | 0.300968239 |

| | | | | | | | |
|---|---|---|---|---|---|---|---|
| **DUT** | 0.273825177 | -1.868672992 | 5.68377012 | 1.82E-03 | 3.38E-02 | 0.152689767 | 0.394960588 |
| **RPL22** | 0.229216088 | -2.125219788 | 4.73384544 | 1.83E-03 | 3.38E-02 | 0.115947463 | 0.342484713 |
| **ATP5PO** | 0.307045683 | -1.703474774 | 6.102127172 | 1.85E-03 | 3.40E-02 | 0.176822853 | 0.437268513 |
| **TRIM65** | 0.311693239 | -1.681801233 | 4.884173295 | 1.88E-03 | 3.45E-02 | 0.160211455 | 0.463175024 |
| **WDR3** | 0.417215452 | -1.261135505 | 4.790367759 | 1.91E-03 | 3.48E-02 | 0.211905145 | 0.622525759 |
| **RRP1B** | -0.144923127 | NA | -4.909586658 | 1.92E-03 | 3.49E-02 | -0.215241524 | -0.074604731 |
| **SHC1** | -0.263034645 | NA | -4.616898673 | 1.92E-03 | 3.49E-02 | -0.395411903 | -0.130657387 |
| **AHCYL2** | -0.234051804 | NA | -4.763327661 | 1.96E-03 | 3.49E-02 | -0.349836167 | -0.118267441 |
| **CNN2** | -0.167812976 | NA | -5.16306187 | 1.98E-03 | 3.49E-02 | -0.246999963 | -0.088625989 |
| **GPS1** | 0.32330709 | -1.629022949 | 4.745768165 | 1.95E-03 | 3.49E-02 | 0.163076586 | 0.483537595 |
| **KHSRP** | 0.123697106 | -3.015116352 | 4.536665804 | 1.96E-03 | 3.49E-02 | 0.06070122 | 0.186692991 |
| **MAT2A** | 0.204626908 | -2.288932223 | 5.488995751 | 1.95E-03 | 3.49E-02 | 0.111658661 | 0.297595156 |
| **NUP54** | -0.254006654 | NA | -5.097940847 | 1.97E-03 | 3.49E-02 | -0.374765386 | -0.133247922 |
| **SAP30** | 0.879105869 | -0.185891178 | 5.98717856 | 1.96E-03 | 3.49E-02 | 0.499988387 | 1.258223351 |
| **SLC35A4** | 2.696488263 | 1.431081754 | 6.468977644 | 1.98E-03 | 3.49E-02 | 1.586240911 | 3.806735614 |
| **TCP1** | 0.218485124 | -2.194393038 | 4.877332903 | 1.95E-03 | 3.49E-02 | 0.111955873 | 0.325014376 |
| **PJA2** | -0.978117891 | NA | -4.678602399 | 2.03E-03 | 3.55E-02 | -1.468528424 | -0.487707357 |
| **RBBP6** | -0.344759434 | NA | -4.88237565 | 2.02E-03 | 3.55E-02 | -0.513234653 | -0.176284214 |
| **SNAP23** | -0.446830222 | NA | -5.774512857 | 2.02E-03 | 3.55E-02 | -0.644339413 | -0.249321031 |
| **YWHAQ** | 0.231110263 | -2.113346767 | 6.083826748 | 2.03E-03 | 3.55E-02 | 0.132070914 | 0.330149613 |
| **PLXND1** | -1.36248012 | NA | -4.846443424 | 2.05E-03 | 3.57E-02 | -2.03195928 | -0.69300096 |
| **ANLN** | 0.330874302 | -1.595644847 | 4.933829317 | 2.08E-03 | 3.59E-02 | 0.169858372 | 0.491890232 |
| **HECTD1** | 0.22631116 | -2.143620366 | 5.701984012 | 2.07E-03 | 3.59E-02 | 0.125304857 | 0.327317463 |
| **DNPH1** | -0.28101076 | NA | -4.469253239 | 2.09E-03 | 3.60E-02 | -0.426007237 | -0.136014282 |
| **BAIAP2L1** | 0.255273484 | -1.969884408 | 4.463676376 | 2.13E-03 | 3.65E-02 | 0.123277215 | 0.387269752 |
| **NUDT9** | 0.242271085 | -2.045305868 | 4.511029733 | 2.13E-03 | 3.65E-02 | 0.117759702 | 0.366782467 |
| **LAMC1** | -0.260263708 | NA | -5.254671231 | 2.15E-03 | 3.68E-02 | -0.382626137 | -0.137901279 |
| **REPS1** | -0.322863943 | NA | -4.477858768 | 2.18E-03 | 3.71E-02 | -0.489797677 | -0.155930208 |
| **TRIM28** | 0.125997705 | -2.988530643 | 4.439508836 | 2.18E-03 | 3.71E-02 | 0.060534222 | 0.191461187 |
| **MED11** | -1.12276612 | NA | -4.933008815 | 2.20E-03 | 3.73E-02 | -1.671779016 | -0.573753224 |
| **TMSB4X** | -0.603854269 | NA | -4.455306735 | 2.21E-03 | 3.73E-02 | -0.91728846 | -0.290420077 |
| **ACTMAP** | -0.544565833 | NA | -4.660418761 | 2.24E-03 | 3.76E-02 | -0.820234187 | -0.268897479 |
| **ST6GAL1** | -0.385407991 | NA | -4.416886444 | 2.24E-03 | 3.76E-02 | -0.586679589 | -0.184136394 |
| **TAF10** | 2.150948738 | 1.104973143 | 6.140852575 | 2.24E-03 | 3.76E-02 | 1.226183965 | 3.075713511 |
| **TAX1BP1** | -1.931620841 | NA | -6.547613585 | 2.26E-03 | 3.78E-02 | -2.731843953 | -1.131397729 |
| **TTC1** | -0.13765943 | NA | -4.590704193 | 2.29E-03 | 3.81E-02 | -0.208055964 | -0.067262896 |
| **CENPH** | -0.469106505 | NA | -5.044438553 | 2.31E-03 | 3.81E-02 | -0.696365085 | -0.241847924 |
| **KIFC1** | 0.16984471 | -2.557711808 | 4.411044679 | 2.31E-03 | 3.81E-02 | 0.08088621 | 0.258803211 |
| **RPL27** | -0.241586262 | NA | -4.399858632 | 2.29E-03 | 3.81E-02 | -0.368234109 | -0.114938414 |

| | | | | | | | |
|---|---|---|---|---|---|---|---|
| **STK11IP** | -0.728907199 | NA | -4.795661485 | 2.30E-03 | 3.81E-02 | -1.09248284 | -0.365331559 |
| **LSM14A** | -0.85872372 | NA | -6.107249904 | 2.32E-03 | 3.81E-02 | -1.230503969 | -0.48694347 |
| **NOTCH2** | 0.344222638 | -1.538586113 | 4.641921806 | 2.34E-03 | 3.83E-02 | 0.169023235 | 0.519422041 |
| **FTH1** | -0.390849718 | NA | -4.728285668 | 2.37E-03 | 3.87E-02 | -0.587884642 | -0.193814793 |
| **BRPF1** | -1.172110627 | NA | -4.834086601 | 2.40E-03 | 3.90E-02 | -1.756078563 | -0.58814269 |
| **TEPSIN** | -0.725847411 | NA | -4.531986606 | 2.40E-03 | 3.90E-02 | -1.101178216 | -0.350516605 |
| **GINS3** | -0.245444683 | NA | -5.680709882 | 2.42E-03 | 3.91E-02 | -0.356780187 | -0.134109179 |
| **ACAT2** | 0.362533685 | -1.463813044 | 4.950683428 | 2.44E-03 | 3.94E-02 | 0.184151169 | 0.540916201 |
| **ROMO1** | -0.387210135 | NA | -5.014143219 | 2.46E-03 | 3.95E-02 | -0.576432515 | -0.197987755 |
| **GPX4** | -0.361947626 | NA | -4.911567152 | 2.47E-03 | 3.97E-02 | -0.541045313 | -0.182849939 |
| **ZWINT** | 0.303020998 | -1.722510324 | 4.563876955 | 2.48E-03 | 3.97E-02 | 0.146557518 | 0.459484479 |
| **SIN3A** | -0.15448231 | NA | -4.710435054 | 2.51E-03 | 4.00E-02 | -0.232879226 | -0.076085393 |
| **CD55** | 1.848576728 | 0.886414926 | 5.157973137 | 2.53E-03 | 4.02E-02 | 0.95771579 | 2.739437666 |
| **FILIP1L** | 2.457615248 | 1.297259072 | 4.886558667 | 2.55E-03 | 4.05E-02 | 1.234703283 | 3.680527213 |
| **EMC8** | -0.490359515 | NA | -4.397337896 | 2.60E-03 | 4.07E-02 | -0.749869921 | -0.230849109 |
| **MT1F** | 0.791520131 | -0.337302051 | 4.304848759 | 2.60E-03 | 4.07E-02 | 0.367497554 | 1.215542708 |
| **PLXNB2** | -0.27101892 | NA | -5.827433023 | 2.60E-03 | 4.07E-02 | -0.39298053 | -0.14905731 |
| **PSMC3** | -0.404380514 | NA | -6.037597401 | 2.58E-03 | 4.07E-02 | -0.58251374 | -0.226247288 |
| **UBQLN4** | -0.273883902 | NA | -4.360476953 | 2.58E-03 | 4.07E-02 | -0.419466965 | -0.128300838 |
| **MRPL12** | 0.34130738 | -1.550856487 | 4.310353136 | 2.61E-03 | 4.08E-02 | 0.158533155 | 0.524081605 |
| **EIF1AX** | -0.257272737 | NA | -4.506230542 | 2.63E-03 | 4.10E-02 | -0.39170431 | -0.122841164 |
| **NDUFA2** | -0.446564039 | NA | -4.302651839 | 2.64E-03 | 4.10E-02 | -0.686140723 | -0.206987354 |
| **RIOK1** | 0.456673295 | -1.130765669 | 4.309052681 | 2.66E-03 | 4.12E-02 | 0.211759096 | 0.701587494 |
| **ATP5F1D** | -0.379527589 | NA | -4.811979257 | 2.67E-03 | 4.12E-02 | -0.570769567 | -0.188285611 |
| **TACO1** | 0.468287391 | -1.094533903 | 4.281942132 | 2.68E-03 | 4.13E-02 | 0.216091515 | 0.720483267 |
| **UFD1** | 0.331971645 | -1.590868075 | 4.446045824 | 2.69E-03 | 4.14E-02 | 0.156869415 | 0.507073874 |
| **DDX24** | 0.300110621 | -1.736433716 | 4.292558302 | 2.71E-03 | 4.15E-02 | 0.138595033 | 0.461626209 |
| **SMYD5** | 0.278947822 | -1.84193281 | 4.286644349 | 2.77E-03 | 4.22E-02 | 0.12845665 | 0.429438993 |
| **PRXL2B** | 0.339837061 | -1.557084901 | 4.467718076 | 2.77E-03 | 4.23E-02 | 0.160662146 | 0.519011975 |
| **HEATR5A** | 0.366887119 | -1.44659184 | 4.670282075 | 2.79E-03 | 4.24E-02 | 0.178122083 | 0.555652156 |
| **BRAP** | 0.284966992 | -1.811133273 | 4.271671552 | 2.82E-03 | 4.26E-02 | 0.130700074 | 0.439233911 |
| **CAPZB** | 0.327746697 | -1.609346854 | 6.060918594 | 2.83E-03 | 4.26E-02 | 0.1822892 | 0.473204194 |
| **DAP3** | -0.220708536 | NA | -4.346707686 | 2.84E-03 | 4.26E-02 | -0.339093353 | -0.102323719 |
| **ENSA** | -0.467946917 | NA | -4.276207341 | 2.84E-03 | 4.26E-02 | -0.721295822 | -0.214598011 |
| **NME1** | 0.387236307 | -1.368713869 | 4.531785712 | 2.84E-03 | 4.26E-02 | 0.184312486 | 0.590160129 |
| **RPS2** | 0.214320974 | -2.222155055 | 4.40592624 | 2.85E-03 | 4.26E-02 | 0.100193044 | 0.328448903 |
| **NIBAN2** | 0.318759122 | -1.649461466 | 4.286220312 | 2.86E-03 | 4.27E-02 | 0.146322394 | 0.49119585 |
| **CARS1** | 0.153872449 | -2.700193157 | 4.238844849 | 2.87E-03 | 4.28E-02 | 0.070088591 | 0.237656306 |
| **FDPS** | -0.259158553 | NA | -4.602083025 | 2.89E-03 | 4.29E-02 | -0.39399249 | -0.124324616 |

| | | | | | | | |
|---|---|---|---|---|---|---|---|
| **FBLN1** | -1.173571894 | NA | -4.575508789 | 2.91E-03 | 4.30E-02 | -1.786423423 | -0.560720365 |
| **ZCCHC3** | 0.358788537 | -1.478794296 | 4.249305624 | 2.91E-03 | 4.30E-02 | 0.163499219 | 0.554077856 |
| **USO1** | -0.266814107 | NA | -4.2637988 | 2.95E-03 | 4.34E-02 | -0.411908575 | -0.121719639 |
| **SRP9** | -0.218873161 | NA | -4.438420786 | 2.97E-03 | 4.36E-02 | -0.335334834 | -0.102411489 |
| **NDUFB4** | 0.241864574 | -2.047728622 | 4.494528709 | 2.99E-03 | 4.37E-02 | 0.114006452 | 0.369722696 |
| **PDHA1** | -0.218268532 | NA | -4.339944026 | 2.99E-03 | 4.37E-02 | -0.335932293 | -0.10060477 |
| **KRT19** | -0.178040696 | NA | -4.216183526 | 3.01E-03 | 4.38E-02 | -0.275611731 | -0.080469661 |
| **NPTN** | -0.340089299 | NA | -4.197079872 | 3.02E-03 | 4.39E-02 | -0.526979778 | -0.15319882 |
| **DDOST** | 0.654780446 | -0.610916856 | 4.183482401 | 3.07E-03 | 4.42E-02 | 0.293834741 | 1.015726152 |
| **ERI1** | 0.226669427 | -2.141338281 | 4.306260719 | 3.06E-03 | 4.42E-02 | 0.10370907 | 0.349629783 |
| **SLC30A1** | 0.426882809 | -1.228088031 | 4.716741493 | 3.05E-03 | 4.42E-02 | 0.206789676 | 0.646975941 |
| **WDR82** | 0.402931562 | -1.311393277 | 4.853824036 | 3.07E-03 | 4.42E-02 | 0.198399061 | 0.607464064 |
| **MORC2** | -0.442498995 | NA | -4.349160137 | 3.09E-03 | 4.43E-02 | -0.681389911 | -0.203608079 |
| **LARP4B** | 0.279482879 | -1.839168186 | 4.191243766 | 3.10E-03 | 4.44E-02 | 0.125455273 | 0.433510485 |
| **FAM169A** | -0.400201549 | NA | -4.516644673 | 3.12E-03 | 4.46E-02 | -0.611994083 | -0.188409015 |
| **PRKAR1A** | 0.545497488 | -0.874355543 | 4.213938833 | 3.13E-03 | 4.46E-02 | 0.245516908 | 0.845478067 |
| **CKAP5** | 0.167167641 | -2.58063249 | 5.773177883 | 3.17E-03 | 4.47E-02 | 0.089956733 | 0.244378549 |
| **CNOT4** | 0.299278332 | -1.740440267 | 4.388616876 | 3.15E-03 | 4.47E-02 | 0.138232776 | 0.460323887 |
| **NUP153** | -0.108214381 | NA | -4.94665183 | 3.17E-03 | 4.47E-02 | -0.162723699 | -0.053705062 |
| **NUP88** | -0.208434434 | NA | -4.320896798 | 3.16E-03 | 4.47E-02 | -0.321596401 | -0.095272467 |
| **SNCA** | -0.261752412 | NA | -4.54441209 | 3.15E-03 | 4.47E-02 | -0.399921912 | -0.123582912 |
| **KTN1** | -0.160643465 | NA | -4.328027291 | 3.18E-03 | 4.47E-02 | -0.247825507 | -0.073461423 |
| **ARHGEF12** | -0.363134763 | NA | -4.178294242 | 3.24E-03 | 4.49E-02 | -0.564301992 | -0.161967535 |
| **MSRB2** | 0.217592771 | -2.200297469 | 4.167140215 | 3.21E-03 | 4.49E-02 | 0.096942373 | 0.338243169 |
| **PPP2R2D** | 0.699687007 | -0.515218393 | 4.187352126 | 3.23E-03 | 4.49E-02 | 0.312538903 | 1.086835112 |
| **SLC25A6** | 0.314486688 | -1.668929144 | 4.430900726 | 3.23E-03 | 4.49E-02 | 0.145802269 | 0.483171108 |
| **TFAM** | -0.132680548 | NA | -4.353933787 | 3.22E-03 | 4.49E-02 | -0.204515287 | -0.06084581 |
| **ENO1** | 0.332640869 | -1.587962662 | 5.665986548 | 3.26E-03 | 4.52E-02 | 0.17688678 | 0.488394958 |
| **FABP5** | -0.310003769 | NA | -4.373652905 | 3.29E-03 | 4.53E-02 | -0.477719456 | -0.142288081 |
| **TRIM24** | -0.176744249 | NA | -4.449594548 | 3.28E-03 | 4.53E-02 | -0.271448168 | -0.082040331 |
| **SNRPD2** | -0.253594405 | NA | -4.44630832 | 3.34E-03 | 4.58E-02 | -0.389761566 | -0.117427245 |
| **UBR3** | -0.568464922 | NA | -5.368634264 | 3.34E-03 | 4.58E-02 | -0.843480076 | -0.293449767 |
| **MGME1** | 0.396941711 | -1.333000923 | 4.404311674 | 3.36E-03 | 4.60E-02 | 0.182592458 | 0.611290965 |
| **FNDC3A** | 0.300544723 | -1.734348408 | 4.951938728 | 3.41E-03 | 4.64E-02 | 0.148166913 | 0.452922532 |
| **RRS1** | -0.134090571 | NA | -4.134353246 | 3.40E-03 | 4.64E-02 | -0.209105325 | -0.059075817 |
| **MAPK3** | 0.374183597 | -1.418181778 | 4.442880501 | 3.43E-03 | 4.65E-02 | 0.172733489 | 0.575633705 |
| **MTREX** | 0.199104698 | -2.328400832 | 5.442314628 | 3.45E-03 | 4.65E-02 | 0.103194222 | 0.295015174 |
| **NOP2** | -0.146269569 | NA | -4.267611368 | 3.45E-03 | 4.65E-02 | -0.226804518 | -0.065734621 |
| **RPS6KA3** | 0.464673631 | -1.105710316 | 4.781364139 | 3.44E-03 | 4.65E-02 | 0.224277176 | 0.705070086 |

| | | | | | | | |
|---|---|---|---|---|---|---|---|
| **PSMD4** | -0.093002088 | NA | -4.093138482 | 3.47E-03 | 4.66E-02 | -0.14539824 | -0.040605936 |
| **SKA3** | 0.226487187 | -2.142498658 | 4.420536059 | 3.46E-03 | 4.66E-02 | 0.104102413 | 0.348871962 |
| **DPY30** | -0.225103526 | NA | -5.150510941 | 3.51E-03 | 4.68E-02 | -0.337096995 | -0.113110056 |
| **PTGES3** | -0.35006932 | NA | -4.361810536 | 3.50E-03 | 4.68E-02 | -0.540781889 | -0.159356751 |
| **PIK3C3** | 0.314804276 | -1.66747296 | 4.087496271 | 3.52E-03 | 4.69E-02 | 0.137111283 | 0.492497268 |
| **COMMD7** | 1.117460604 | 0.16022397 | 4.103520136 | 3.55E-03 | 4.70E-02 | 0.487645834 | 1.747275374 |
| **PCM1** | -0.209667588 | NA | -4.596406764 | 3.54E-03 | 4.70E-02 | -0.320813382 | -0.098521794 |
| **CHEK1** | 0.428655764 | -1.222108552 | 4.189161634 | 3.57E-03 | 4.71E-02 | 0.189586279 | 0.667725249 |
| **CRIP1** | 0.213090524 | -2.230461657 | 4.089592852 | 3.57E-03 | 4.71E-02 | 0.092708484 | 0.333472564 |
| **CSTB** | -0.098469956 | NA | -4.071048742 | 3.61E-03 | 4.71E-02 | -0.154293706 | -0.042646207 |
| **NOL10** | 0.219953109 | -2.184732104 | 4.070647808 | 3.58E-03 | 4.71E-02 | 0.095347789 | 0.344558429 |
| **POLR3C** | 0.503687776 | -0.989398376 | 4.5830215 | 3.61E-03 | 4.71E-02 | 0.235782792 | 0.77159276 |
| **RENBP** | -0.789530193 | NA | -4.734992871 | 3.61E-03 | 4.71E-02 | -1.20206725 | -0.376993135 |
| **SERF2** | -1.962543486 | NA | -4.512422548 | 3.60E-03 | 4.71E-02 | -3.015025633 | -0.910061339 |
| **UBE2O** | 0.193482665 | -2.369723782 | 4.170373645 | 3.61E-03 | 4.71E-02 | 0.085207977 | 0.301757353 |
| **ADD2** | -0.188867041 | NA | -4.067574078 | 3.68E-03 | 4.74E-02 | -0.296147377 | -0.081586704 |
| **ATP5F1C** | 0.4040263 | -1.307478886 | 5.086644862 | 3.66E-03 | 4.74E-02 | 0.200707092 | 0.607345509 |
| **H1-10** | 0.488517077 | -1.033519099 | 4.12272864 | 3.69E-03 | 4.74E-02 | 0.212967478 | 0.764066676 |
| **HOMER1** | 0.256659008 | -1.962075199 | 4.119688392 | 3.69E-03 | 4.74E-02 | 0.111829147 | 0.401488869 |
| **N4BP2** | 0.215675227 | -2.21306762 | 4.064500073 | 3.69E-03 | 4.74E-02 | 0.093082312 | 0.338268142 |
| **RANBP3** | 0.148893669 | -2.747645688 | 4.326528886 | 3.66E-03 | 4.74E-02 | 0.067108265 | 0.230679072 |
| **SPIN1** | -0.264895439 | NA | -4.862649687 | 3.69E-03 | 4.74E-02 | -0.401629473 | -0.128161404 |
| **RPL22L1** | 0.377059756 | -1.407134917 | 4.151858521 | 3.71E-03 | 4.74E-02 | 0.165079097 | 0.589040415 |
| **MELK** | 0.3804429 | -1.394248158 | 4.237955225 | 3.74E-03 | 4.78E-02 | 0.168712042 | 0.592173757 |
| **N6AMT1** | 1.40190073 | 0.487384194 | 4.121414877 | 3.79E-03 | 4.83E-02 | 0.609096406 | 2.194705054 |
| **UBR5** | -0.156355298 | NA | -4.056065811 | 3.80E-03 | 4.83E-02 | -0.245540813 | -0.067169783 |
| **GTPBP1** | -0.213073843 | NA | -4.053390188 | 3.86E-03 | 4.87E-02 | -0.334814492 | -0.091333194 |
| **SMC5** | 0.308875677 | -1.694901828 | 4.163245228 | 3.86E-03 | 4.87E-02 | 0.134873935 | 0.482877419 |
| **VAPB** | -0.166117328 | NA | -4.028223192 | 3.85E-03 | 4.87E-02 | -0.261315884 | -0.070918772 |
| **IVD** | -0.422357635 | NA | -4.437566232 | 3.88E-03 | 4.87E-02 | -0.652505642 | -0.192209627 |
| **MAEA** | 0.506553466 | -0.981213544 | 4.396090409 | 3.88E-03 | 4.87E-02 | 0.229151781 | 0.783955151 |
| **WWOX** | -0.528643908 | NA | -4.592920024 | 3.88E-03 | 4.87E-02 | -0.811404653 | -0.245883163 |
| **CMAS** | 0.265577605 | -1.912794599 | 4.045974655 | 3.91E-03 | 4.88E-02 | 0.113514885 | 0.417640325 |
| **RAP2B** | -0.556901656 | NA | -4.322040485 | 3.91E-03 | 4.88E-02 | -0.864827089 | -0.248976223 |
| **SKP1** | -0.310073617 | NA | -4.019514879 | 3.91E-03 | 4.88E-02 | -0.488212495 | -0.131934739 |
| **CSK** | 0.356435589 | -1.488286701 | 4.531449838 | 3.97E-03 | 4.92E-02 | 0.163974249 | 0.548896929 |
| **RAD23B** | 0.289885937 | -1.786442748 | 4.691655845 | 3.96E-03 | 4.92E-02 | 0.136246571 | 0.443525304 |
| **COPG1** | 0.283201172 | -1.82010086 | 4.018620229 | 4.00E-03 | 4.94E-02 | 0.120151208 | 0.446251135 |
| **VWA1** | -0.73508736 | NA | -4.12361095 | 3.99E-03 | 4.94E-02 | -1.152562167 | -0.317612554 |

| | | | | | | | |
|---|---|---|---|---|---|---|---|
| **ARHGEF16** | -0.241102529 | NA | -4.01222784 | 4.01E-03 | 4.95E-02 | -0.380070977 | -0.102134081 |
| **DUSP12** | 0.270194044 | -1.887932223 | 4.080041641 | 4.02E-03 | 4.95E-02 | 0.115800515 | 0.424587572 |
| **RACK1** | 0.344332006 | -1.538127807 | 5.443788829 | 4.04E-03 | 4.96E-02 | 0.175513918 | 0.513150094 |
| **NSF** | 0.091812849 | -3.445160123 | 4.200998804 | 4.06E-03 | 4.98E-02 | 0.040098672 | 0.143527025 |
| **CHTF8** | 1.235733835 | 0.305368034 | 5.277875774 | 4.07E-03 | 4.98E-02 | 0.61917255 | 1.852295119 |
| **ZNF106** | -0.313794882 | NA | -4.767227572 | 4.09E-03 | 4.99E-02 | -0.479229114 | -0.14836065 |
| **FKBP7** | 0.321716784 | -1.636136894 | 3.972112319 | 4.11E-03 | 5.00E-02 | 0.134944536 | 0.508489031 |
| **MBIP** | 0.806227765 | -0.310740626 | 3.988386031 | 4.11E-03 | 5.00E-02 | 0.339114105 | 1.273341426 |
| **CERT1** | -0.165599394 | NA | -4.365632568 | 4.12E-03 | 5.00E-02 | -0.257138666 | -0.074060121 |

Supplemental Table S3_Sheet_3

| Genes | estimate | log2FC | statistic | p.value | adj_p | conf.low | conf.high |
|---|---|---|---|---|---|---|---|
| **DUSP3** | 0.784229738 | -0.350651744 | 21.51798596 | 3.18E-08 | 0.000183118 | 0.699803299 | 0.868656178 |
| **TMA7** | 4.06887971 | 2.02463163 | 21.95199824 | 1.87E-07 | 0.000538584 | 3.62586648 | 4.511892941 |
| **TBL2** | 0.621091546 | -0.687122164 | 13.93878358 | 8.64E-07 | 0.001658506 | 0.51787832 | 0.724304772 |
| **CPSF6** | -0.56328554 | NA | -14.39279104 | 1.27E-06 | 0.001829774 | -0.655068 | -0.47150308 |
| **PHKB** | 0.551882768 | -0.857566256 | 12.15440845 | 1.95E-06 | 0.0022413 | 0.447175097 | 0.656590439 |
| **FAM162A** | 3.479751956 | 1.798984471 | 11.86810615 | 2.98E-06 | 0.002597479 | 2.800088739 | 4.159415172 |
| **GARS1** | 0.302315323 | -1.72587399 | 11.47927456 | 3.16E-06 | 0.002597479 | 0.241519821 | 0.363110826 |
| **ASMTL** | 0.455883793 | -1.133261972 | 11.19634252 | 4.46E-06 | 0.002853526 | 0.361555376 | 0.55021221 |
| **NT5C3B** | 1.82605079 | 0.868726893 | 11.55566162 | 4.35E-06 | 0.002853526 | 1.458213282 | 2.193888298 |
| **KRAS** | -0.814074992 | NA | -11.00594742 | 5.68E-06 | 0.002971257 | -0.985894553 | -0.642255431 |
| **MMAB** | 0.428206524 | -1.22362132 | 11.80741334 | 5.41E-06 | 0.002971257 | 0.343037772 | 0.513375275 |
| **RTN3** | 3.65813827 | 1.871109607 | 16.18971382 | 7.97E-06 | 0.003826151 | 3.091755427 | 4.224521113 |
| **NTN1** | 5.995247068 | 2.583819209 | 19.79580845 | 9.24E-06 | 0.004094637 | 5.205119534 | 6.785374601 |
| **DCAKD** | 0.662243477 | -0.594566366 | 9.597401402 | 1.24E-05 | 0.005110423 | 0.502818047 | 0.821668908 |
| **FDXR** | 0.370818529 | -1.431214762 | 9.757781398 | 1.50E-05 | 0.005758303 | 0.282296161 | 0.459340896 |
| **CCT3** | 0.196498522 | -2.347409637 | 11.04619594 | 1.73E-05 | 0.005847671 | 0.153881941 | 0.239115102 |
| **NFYC** | 0.624451282 | -0.679339074 | 9.167550258 | 1.72E-05 | 0.005847671 | 0.467108201 | 0.781794363 |
| **TMSB10** | -1.145015733 | NA | -9.096841353 | 1.83E-05 | 0.005847671 | -1.435781216 | -0.85425025 |
| **FAH** | 0.894998196 | -0.16004332 | 9.826953355 | 1.96E-05 | 0.005938033 | 0.680944389 | 1.109052004 |
| **ANAPC2** | 0.512446638 | -0.964526314 | 8.562736674 | 2.68E-05 | 0.007011129 | 0.374426715 | 0.65046656 |
| **FILIP1L** | 2.767382356 | 1.468521988 | 9.358162 | 2.55E-05 | 0.007011129 | 2.073739662 | 3.461025051 |
| **ZNF106** | -0.316621652 | NA | -8.573951981 | 2.65E-05 | 0.007011129 | -0.401785874 | -0.23145743 |
| **GTF2A1** | -0.552683731 | NA | -8.384315839 | 3.24E-05 | 0.008108124 | -0.704872677 | -0.400494785 |
| **CTHRC1** | 3.562437771 | 1.832864813 | 15.82772528 | 3.41E-05 | 0.008151752 | 2.968748697 | 4.156126844 |
| **CTNNB1** | 0.326454095 | -1.615047958 | 9.991976281 | 3.54E-05 | 0.008151752 | 0.24794976 | 0.404958429 |
| **BET1** | 1.807246223 | 0.853793075 | 8.198786388 | 4.37E-05 | 0.008391597 | 1.296151228 | 2.318341217 |
| **FUBP3** | -0.341614203 | NA | -8.32940275 | 3.86E-05 | 0.008391597 | -0.436669484 | -0.246558922 |
| **HCCS** | -0.346253147 | NA | -8.385520943 | 4.02E-05 | 0.008391597 | -0.44221307 | -0.250293223 |
| **HTRA2** | 0.878673414 | -0.186601051 | 8.068387325 | 4.24E-05 | 0.008391597 | 0.627302046 | 1.130044782 |
| **PDIA5** | 0.327313014 | -1.611257128 | 8.319871396 | 4.34E-05 | 0.008391597 | 0.235818539 | 0.41880749 |
| **TPT1** | -0.380252144 | NA | -11.20995737 | 4.53E-05 | 0.008423945 | -0.464534354 | -0.295969935 |
| **ACAD9** | 0.394424197 | -1.342180034 | 8.954987285 | 5.14E-05 | 0.008552047 | 0.28972641 | 0.499121984 |
| **ADH7** | 0.780741402 | -0.357083318 | 7.790637291 | 5.50E-05 | 0.008552047 | 0.549347064 | 1.01213574 |
| **BTK** | 0.858118914 | -0.220750511 | 8.26548077 | 5.79E-05 | 0.008552047 | 0.614730945 | 1.101506883 |
| **DDX17** | -0.118759516 | NA | -8.276934875 | 5.78E-05 | 0.008552047 | -0.152405438 | -0.085113594 |
| **GPX7** | 4.332255699 | 2.115118397 | 14.27092065 | 5.28E-05 | 0.008552047 | 3.532767967 | 5.131743432 |
| **PPAN** | -0.245954525 | NA | -8.134657043 | 5.61E-05 | 0.008552047 | -0.316504133 | -0.175404917 |
| **PTGES3** | -0.496148474 | NA | -7.898459693 | 4.83E-05 | 0.008552047 | -0.641045603 | -0.351251345 |
| **UBA6** | 0.215821512 | -2.212089421 | 7.847586894 | 5.09E-05 | 0.008552047 | 0.152374262 | 0.279268762 |

| | | | | | | | |
|---|---|---|---|---|---|---|---|
| **ANP32A** | -0.790155455 | NA | -7.831155485 | 6.12E-05 | 0.008673443 | -1.024223136 | -0.556087775 |
| **PEX14** | 3.197804343 | 1.67708167 | 9.794446667 | 6.17E-05 | 0.008673443 | 2.400615978 | 3.994992707 |
| **GCA** | 0.464046802 | -1.107657778 | 9.342283385 | 6.99E-05 | 0.009552719 | 0.343475887 | 0.584617717 |
| **ZNF578** | 3.92812639 | 1.97384135 | 12.29389555 | 7.13E-05 | 0.009552719 | 3.102072362 | 4.754180419 |
| **NLRP7** | -0.288155082 | NA | -10.03721029 | 7.63E-05 | 0.009980793 | -0.35924984 | -0.217060324 |
| **ENY2** | -0.868922088 | NA | -7.347291324 | 8.15E-05 | 0.010023695 | -1.141794099 | -0.596050077 |
| **GOLM2** | 2.702034402 | 1.434046043 | 14.44055202 | 8.04E-05 | 0.010023695 | 2.197428227 | 3.206640577 |
| **HNRNPUL1** | -0.382406037 | NA | -8.91388239 | 8.18E-05 | 0.010023695 | -0.486055888 | -0.278756185 |
| **ALG11** | -2.09198442 | NA | -8.17431325 | 1.11E-04 | 0.010224272 | -2.704949325 | -1.479019515 |
| **ATP5ME** | 0.521185869 | -0.940130128 | 9.100921349 | 1.25E-04 | 0.010224272 | 0.379629809 | 0.662741928 |
| **ATP6V1F** | 0.317404715 | -1.655604536 | 7.029934792 | 1.10E-04 | 0.010224272 | 0.213256177 | 0.421553252 |
| **CALR** | 0.509823015 | -0.971931592 | 7.997114674 | 1.12E-04 | 0.010224272 | 0.357887556 | 0.661758474 |
| **DENND4C** | 0.368298696 | -1.441051804 | 8.066534882 | 1.16E-04 | 0.010224272 | 0.259074472 | 0.477522921 |
| **EXOSC4** | 0.429547326 | -1.219111004 | 7.605386739 | 1.15E-04 | 0.010224272 | 0.296464261 | 0.562630391 |
| **EXOSC5** | 3.480264388 | 1.799196909 | 9.963688088 | 9.60E-05 | 0.010224272 | 2.608069221 | 4.352459554 |
| **FUBP1** | -0.180285433 | NA | -8.337852728 | 1.24E-04 | 0.010224272 | -0.232580176 | -0.12799069 |
| **GRHPR** | -0.645568987 | NA | -9.094595012 | 1.17E-04 | 0.010224272 | -0.820509643 | -0.470628331 |
| **GTF2F1** | -0.30751557 | NA | -7.095776072 | 1.10E-04 | 0.010224272 | -0.407701253 | -0.207329888 |
| **GTPBP4** | 0.280808757 | -1.83234017 | 7.324562761 | 8.58E-05 | 0.010224272 | 0.192259578 | 0.369357936 |
| **LONP1** | 0.239560624 | -2.061537301 | 7.515875695 | 9.97E-05 | 0.010224272 | 0.165078808 | 0.314042439 |
| **N6AMT1** | 1.951570463 | 0.964635553 | 7.705911373 | 1.08E-04 | 0.010224272 | 1.354259719 | 2.548881206 |
| **NATD1** | 2.325220471 | 1.217367515 | 9.510666956 | 1.20E-04 | 0.010224272 | 1.715391737 | 2.935049206 |
| **NUBP1** | 2.164552837 | 1.114069018 | 7.694005321 | 1.13E-04 | 0.010224272 | 1.500285596 | 2.828820079 |
| **NUP153** | -0.155675577 | NA | -8.367823348 | 1.26E-04 | 0.010224272 | -0.200738575 | -0.110612579 |
| **ORC1** | -0.561817511 | NA | -7.564965221 | 1.18E-04 | 0.010224272 | -0.736763569 | -0.386871454 |
| **POLR1F** | 2.695833981 | 1.430731653 | 9.482666526 | 1.11E-04 | 0.010224272 | 1.989611509 | 3.402056452 |
| **PRPF3** | -0.332799655 | NA | -9.955335912 | 1.08E-04 | 0.010224272 | -0.416706627 | -0.248892682 |
| **RBM25** | -0.34449882 | NA | -9.144669306 | 1.16E-04 | 0.010224272 | -0.437445187 | -0.251552454 |
| **RPS20** | 0.535377074 | -0.901372734 | 8.480867692 | 1.24E-04 | 0.010224272 | 0.382045167 | 0.688708981 |
| **SFXN1** | 0.38778307 | -1.366678278 | 7.151325049 | 1.17E-04 | 0.010224272 | 0.261904754 | 0.513661385 |
| **SLC9A1** | 0.586695126 | -0.769317087 | 8.113980228 | 9.41E-05 | 0.010224272 | 0.414927115 | 0.758463137 |
| **TOX4** | 0.40107614 | -1.318051953 | 7.343854294 | 1.14E-04 | 0.010224272 | 0.273534635 | 0.528617644 |
| **EIF2S3** | -0.437447874 | NA | -7.09387416 | 1.33E-04 | 0.010516338 | -0.581030524 | -0.293865224 |
| **MED8** | -1.617955179 | NA | -6.837372618 | 1.33E-04 | 0.010516338 | -2.163637116 | -1.072273242 |
| **EFCAB14** | -0.585128895 | NA | -6.815172225 | 1.40E-04 | 0.010879734 | -0.783335857 | -0.386921933 |
| **RPS16** | -0.397925963 | NA | -6.746601218 | 1.46E-04 | 0.011181034 | -0.533939811 | -0.261912116 |
| **PPIG** | 0.477458659 | -1.066552272 | 6.759410859 | 1.52E-04 | 0.011483137 | 0.314239949 | 0.64067737 |
| **AKAP1** | -0.310556881 | NA | -7.267963409 | 1.59E-04 | 0.011532444 | -0.411376703 | -0.209737059 |
| **ARHGDIA** | -0.256098182 | NA | -6.943798977 | 1.83E-04 | 0.011532444 | -0.342586709 | -0.169609655 |
| **ATP1B2** | 2.115343699 | 1.08089209 | 9.019665049 | 1.80E-04 | 0.011532444 | 1.526605168 | 2.704082231 |
| **CAPZB** | 0.226327315 | -2.143517385 | 8.674568896 | 1.72E-04 | 0.011532444 | 0.161643738 | 0.291010892 |

| | | | | | | | |
|---|---|---|---|---|---|---|---|
| **CDC5L** | -0.296530698 | NA | -6.947984472 | 1.61E-04 | 0.011532444 | -0.396113717 | -0.196947678 |
| **DDX52** | 0.344346213 | -1.538068284 | 6.995204825 | 1.84E-04 | 0.011532444 | 0.228655242 | 0.460037184 |
| **FAM114A2** | 0.625875846 | -0.676051595 | 6.668941264 | 1.59E-04 | 0.011532444 | 0.409381693 | 0.842369999 |
| **FBL** | -0.219761633 | NA | -6.677321623 | 1.65E-04 | 0.011532444 | -0.295812117 | -0.143711149 |
| **FUCA1** | 1.329841329 | 0.41125412 | 6.760790209 | 1.66E-04 | 0.011532444 | 0.873667991 | 1.786014668 |
| **H1-5** | 0.399280487 | -1.324525525 | 6.580937583 | 1.84E-04 | 0.011532444 | 0.259025704 | 0.539535271 |
| **NFU1** | 0.343628655 | -1.541077749 | 6.625564454 | 1.65E-04 | 0.011532444 | 0.224027103 | 0.463230206 |
| **P3H1** | 0.246142763 | -2.02243277 | 6.936086432 | 1.74E-04 | 0.011532444 | 0.163135511 | 0.329150016 |
| **PSMG2** | 0.380944851 | -1.392345941 | 6.578915803 | 1.79E-04 | 0.011532444 | 0.247249113 | 0.514640588 |
| **RPS15A** | 0.379555907 | -1.397615692 | 6.705243072 | 1.77E-04 | 0.011532444 | 0.248241507 | 0.510870306 |
| **SARS1** | 0.27584193 | -1.858086321 | 6.835206657 | 1.77E-04 | 0.011532444 | 0.181737115 | 0.369946745 |
| **ZNF121** | 0.669112785 | -0.579678685 | 6.837364352 | 1.68E-04 | 0.011532444 | 0.441352496 | 0.896873074 |
| **NECTIN2** | -0.430237591 | NA | -6.83858562 | 1.88E-04 | 0.011586125 | -0.577307644 | -0.283167539 |
| **USP4** | -0.527172523 | NA | -6.607093763 | 1.89E-04 | 0.011586125 | -0.712024976 | -0.34232007 |
| **ALDH9A1** | 0.173562745 | -2.526470789 | 6.609532529 | 1.98E-04 | 0.011611675 | 0.112610884 | 0.234514605 |
| **MIF** | 1.370191589 | 0.454377634 | 8.784381592 | 1.97E-04 | 0.011611675 | 0.979612855 | 1.760770323 |
| **RPL10A** | -0.162897241 | NA | -6.970584077 | 1.92E-04 | 0.011611675 | -0.217865678 | -0.107928803 |
| **SLC25A6** | 0.356931317 | -1.486281605 | 6.473300376 | 1.94E-04 | 0.011611675 | 0.229761882 | 0.484100752 |
| **CALB1** | 1.335009392 | 0.416849892 | 7.123834708 | 2.08E-04 | 0.012001556 | 0.890050434 | 1.779968351 |
| **CLASP2** | 0.301953292 | -1.727602692 | 6.682123196 | 2.23E-04 | 0.012001556 | 0.196214947 | 0.407691637 |
| **GPD2** | 0.259241593 | -1.94763089 | 7.88351914 | 2.22E-04 | 0.012001556 | 0.178760789 | 0.339722397 |
| **HMGA1** | -0.47615534 | NA | -6.386360321 | 2.13E-04 | 0.012001556 | -0.648107553 | -0.304203127 |
| **JUN** | -0.922742426 | NA | -6.457460725 | 2.23E-04 | 0.012001556 | -1.253924127 | -0.591560726 |
| **MALT1** | 0.307630295 | -1.700730509 | 6.371524664 | 2.16E-04 | 0.012001556 | 0.196291053 | 0.418969538 |
| **NOL10** | 0.457275369 | -1.128864885 | 6.896618007 | 2.10E-04 | 0.012001556 | 0.301179758 | 0.613370979 |
| **PFDN6** | 0.194750753 | -2.360299185 | 6.545786933 | 2.21E-04 | 0.012001556 | 0.125556995 | 0.263944512 |
| **RPL27A** | -0.184197244 | NA | -7.08118617 | 2.11E-04 | 0.012001556 | -0.245888162 | -0.122506326 |
| **CDH1** | -0.592557555 | NA | -6.863545663 | 2.48E-04 | 0.012247973 | -0.797055676 | -0.388059434 |
| **CSK** | 0.324581679 | -1.623346526 | 6.239760329 | 2.50E-04 | 0.012247973 | 0.204604356 | 0.444559002 |
| **CTSL** | 0.685386485 | -0.545010351 | 7.042457322 | 2.35E-04 | 0.012247973 | 0.453791651 | 0.916981319 |
| **DDX5** | -0.251715779 | NA | -6.714560684 | 2.50E-04 | 0.012247973 | -0.340000804 | -0.163430754 |
| **GGCX** | 0.462319802 | -1.113036939 | 6.307218698 | 2.38E-04 | 0.012247973 | 0.293082853 | 0.63155675 |
| **L2HGDH** | 0.275459144 | -1.860089738 | 6.874741932 | 2.55E-04 | 0.012247973 | 0.180398494 | 0.370519794 |
| **MVB12A** | 0.46583397 | -1.102112244 | 8.358899781 | 2.45E-04 | 0.012247973 | 0.326547086 | 0.605120855 |
| **PITHD1** | 0.592210602 | -0.755817775 | 6.88838725 | 2.54E-04 | 0.012247973 | 0.388137027 | 0.796284178 |
| **SEC24D** | 0.250072303 | -1.999582815 | 6.612882235 | 2.41E-04 | 0.012247973 | 0.161548486 | 0.33859612 |
| **SNAP23** | -0.408141483 | NA | -7.350006195 | 2.53E-04 | 0.012247973 | -0.542334222 | -0.273948743 |
| **TAX1BP1** | -1.697900009 | NA | -9.72549992 | 2.55E-04 | 0.012247973 | -2.153535699 | -1.242264319 |
| **TPM4** | -0.433122845 | NA | -7.120645758 | 2.38E-04 | 0.012247973 | -0.578386659 | -0.287859032 |
| **ZW10** | 0.689477528 | -0.536424564 | 7.87700398 | 2.48E-04 | 0.012247973 | 0.474094112 | 0.904860943 |
| **FXN** | -0.634690175 | NA | -7.352183316 | 2.58E-04 | 0.012292023 | -0.84351605 | -0.425864301 |

| | | | | | | | |
|---|---|---|---|---|---|---|---|
| **MPHOSPH6** | 2.51218594 | 1.32894325 | 9.387176092 | 2.61E-04 | 0.012309664 | 1.819528736 | 3.204843144 |
| **DFFA** | -0.282848798 | NA | -6.254016175 | 2.69E-04 | 0.012378538 | -0.387555779 | -0.178141816 |
| **G3BP1** | -0.181601302 | NA | -7.165560011 | 2.71E-04 | 0.012378538 | -0.242609332 | -0.120593271 |
| **ZBTB40** | -0.954571812 | NA | -6.370100516 | 2.69E-04 | 0.012378538 | -1.303347252 | -0.605796372 |
| **ZPR1** | -0.432159964 | NA | -6.459867845 | 2.70E-04 | 0.012378538 | -0.588530618 | -0.27578931 |
| **ATAD3A** | -0.405453304 | NA | -6.1141278 | 2.88E-04 | 0.012724713 | -0.558444406 | -0.252462203 |
| **BRD9** | -0.885998149 | NA | -7.105821409 | 2.89E-04 | 0.012724713 | -1.186411364 | -0.585584934 |
| **EIPR1** | 0.585796634 | -0.771528191 | 9.042118812 | 2.88E-04 | 0.012724713 | 0.418867286 | 0.752725982 |
| **IFI16** | 0.281094402 | -1.830873371 | 6.132535742 | 2.88E-04 | 0.012724713 | 0.175249554 | 0.38693925 |
| **NASP** | -0.267960147 | NA | -6.113910922 | 2.85E-04 | 0.012724713 | -0.369029564 | -0.166890729 |
| **WDR12** | -0.4036886 | NA | -6.577666102 | 2.95E-04 | 0.012872773 | -0.548460589 | -0.258916612 |
| **PAK2** | -0.242702106 | NA | -6.150896335 | 3.01E-04 | 0.013016989 | -0.334064002 | -0.151340209 |
| **FXR2** | 0.285424707 | -1.808817874 | 6.207049416 | 3.07E-04 | 0.01308944 | 0.178572025 | 0.392277388 |
| **MYL3** | 0.447046937 | -1.161501783 | 6.159193967 | 3.06E-04 | 0.01308944 | 0.278790159 | 0.615303715 |
| **ATP5PO** | 0.386092407 | -1.372981911 | 6.084163691 | 3.15E-04 | 0.013339253 | 0.239324528 | 0.532860287 |
| **RABGEF1** | -0.31106566 | NA | -6.59735907 | 3.18E-04 | 0.013387441 | -0.422785777 | -0.199345542 |
| **CCT2** | -0.165562492 | NA | -7.014383509 | 3.29E-04 | 0.013621333 | -0.222582207 | -0.108542778 |
| **MYDGF** | 0.348529381 | -1.520647814 | 6.109630323 | 3.27E-04 | 0.013621333 | 0.216201397 | 0.480857365 |
| **BRD8** | 0.330640266 | -1.596665666 | 6.919293884 | 3.39E-04 | 0.013850736 | 0.215478347 | 0.445802185 |
| **DYNC1I2** | -0.318198856 | NA | -8.554437227 | 3.38E-04 | 0.013850736 | -0.413461852 | -0.222935861 |
| **STXBP3** | 0.148772895 | -2.748816392 | 6.047189844 | 3.46E-04 | 0.014013771 | 0.091738651 | 0.205807139 |
| **GEMIN7** | -0.826359025 | NA | -7.647332163 | 3.48E-04 | 0.014022245 | -1.094817109 | -0.55790094 |
| **PPT1** | -0.419134161 | NA | -6.30666147 | 3.58E-04 | 0.014314521 | -0.575403086 | -0.262865235 |
| **CARM1** | 0.378290098 | -1.402435081 | 5.934052025 | 3.62E-04 | 0.014394458 | 0.231016651 | 0.525563545 |
| **TKFC** | 0.373476188 | -1.420911832 | 6.686134051 | 3.72E-04 | 0.014673343 | 0.239578622 | 0.507373754 |
| **DDX1** | -0.219865952 | NA | -5.957640341 | 3.88E-04 | 0.015085128 | -0.305492595 | -0.134239309 |
| **PNPT1** | -0.245883045 | NA | -6.117309122 | 3.85E-04 | 0.015085128 | -0.339868013 | -0.151898078 |
| **HMGB2** | -0.399591846 | NA | -6.01293212 | 4.03E-04 | 0.015583834 | -0.554504019 | -0.244679672 |
| **TTC4** | 0.631523382 | -0.663091944 | 7.55682963 | 4.07E-04 | 0.015643355 | 0.42277571 | 0.840271054 |
| **ESRP1** | -0.210725548 | NA | -5.803758996 | 4.13E-04 | 0.015663904 | -0.294547819 | -0.126903278 |
| **PRKAR2A** | 0.431160347 | -1.213703592 | 6.657354907 | 4.13E-04 | 0.015663904 | 0.275262921 | 0.587057773 |
| **BTF3L4** | -0.616438233 | NA | -5.890679126 | 4.16E-04 | 0.015664642 | -0.859212804 | -0.373663663 |
| **HMGCR** | -0.503629104 | NA | -6.152426657 | 4.23E-04 | 0.015804833 | -0.696230953 | -0.311027255 |
| **EDF1** | -0.577421533 | NA | -7.094182966 | 4.33E-04 | 0.016082655 | -0.777630258 | -0.377212808 |
| **ACP1** | -0.174381583 | NA | -6.255664323 | 4.49E-04 | 0.016577781 | -0.240507716 | -0.10825545 |
| **ATP5MG** | 0.487549407 | -1.036379668 | 5.678410943 | 4.67E-04 | 0.016856266 | 0.289526843 | 0.685571972 |
| **ERI1** | 0.24155424 | -2.049580919 | 6.632167493 | 4.65E-04 | 0.016856266 | 0.153427976 | 0.329680504 |
| **KYAT3** | 0.435764868 | -1.198378205 | 5.698905789 | 4.68E-04 | 0.016856266 | 0.259192311 | 0.612337426 |
| **SIPA1L2** | 0.623130733 | -0.682393221 | 5.818925986 | 4.64E-04 | 0.016856266 | 0.374333869 | 0.871927597 |
| **DOHH** | -0.314050631 | NA | -5.682599382 | 4.77E-04 | 0.017052093 | -0.441663884 | -0.186437379 |
| **SEC16A** | -0.187410311 | NA | -5.759619084 | 4.82E-04 | 0.017144816 | -0.262907899 | -0.111912724 |

| | | | | | | | |
|---|---|---|---|---|---|---|---|
| **ALDOA** | -0.217420665 | NA | -5.639304326 | 5.18E-04 | 0.018316706 | -0.306596318 | -0.128245013 |
| **PTCD3** | -0.328798971 | NA | -5.578591854 | 5.25E-04 | 0.018440851 | -0.46473882 | -0.192859121 |
| **ADD2** | -0.262566087 | NA | -5.612671613 | 5.28E-04 | 0.018444435 | -0.370707909 | -0.154424265 |
| **MIS18A** | -0.360280731 | NA | -5.758775287 | 5.40E-04 | 0.018721331 | -0.506253788 | -0.214307674 |
| **OLFML3** | -0.384704713 | NA | -6.372293153 | 5.52E-04 | 0.019004258 | -0.530360408 | -0.239049017 |
| **SMARCC1** | -0.288608473 | NA | -5.537742105 | 5.54E-04 | 0.019004258 | -0.408850172 | -0.168366773 |
| **BUD31** | 0.441526562 | -1.179427862 | 5.600210995 | 5.60E-04 | 0.019075895 | 0.258876217 | 0.624176907 |
| **FKBP15** | -0.268773288 | NA | -5.50068142 | 5.76E-04 | 0.019405799 | -0.381477688 | -0.156068888 |
| **NAGK** | -0.407323862 | NA | -5.53869347 | 5.75E-04 | 0.019405799 | -0.577319577 | -0.237328146 |
| **FABP5** | -0.291955073 | NA | -6.571943353 | 5.82E-04 | 0.019481285 | -0.40051272 | -0.183397426 |
| **ARHGAP12** | 2.272826332 | 1.184487451 | 8.251075474 | 6.01E-04 | 0.020017883 | 1.548312238 | 2.997340426 |
| **IGSF1** | -0.459258824 | NA | -6.041833612 | 6.11E-04 | 0.02020925 | -0.640572444 | -0.277945203 |
| **ACADM** | 0.482990646 | -1.049932847 | 8.413749862 | 6.24E-04 | 0.020301597 | 0.330677139 | 0.635304152 |
| **ALDOC** | -0.319609083 | NA | -8.196669898 | 6.20E-04 | 0.020301597 | -0.422180673 | -0.217037493 |
| **RRAS** | -1.528416283 | NA | -6.241866131 | 6.22E-04 | 0.020301597 | -2.119291151 | -0.937541414 |
| **CACTIN** | -0.411373946 | NA | -5.421954489 | 6.38E-04 | 0.020536287 | -0.5864565 | -0.236291393 |
| **SCCPDH** | 0.292985551 | -1.771098577 | 5.975620319 | 6.38E-04 | 0.020536287 | 0.176161672 | 0.40980943 |
| **ATXN2L** | -0.249761229 | NA | -5.425436655 | 6.55E-04 | 0.020729397 | -0.356162004 | -0.143360454 |
| **PTRHD1** | -0.55322354 | NA | -5.398061431 | 6.48E-04 | 0.020729397 | -0.789564382 | -0.316882698 |
| **RBM17** | 0.240918387 | -2.053383591 | 6.086357685 | 6.52E-04 | 0.020729397 | 0.145925222 | 0.335911552 |
| **TPD52L2** | -0.324532286 | NA | -6.05707531 | 6.59E-04 | 0.020733749 | -0.45299224 | -0.196072331 |
| **AFG2A** | -0.538475659 | NA | -7.077183441 | 6.69E-04 | 0.02082333 | -0.730513704 | -0.346437613 |
| **ANO6** | 0.544446113 | -0.87713883 | 5.961874048 | 6.72E-04 | 0.02082333 | 0.326375208 | 0.762517019 |
| **CEBPZ** | -0.281759181 | NA | -5.361495128 | 6.79E-04 | 0.02082333 | -0.402971109 | -0.160547253 |
| **RPS3** | 0.290443449 | -1.783670803 | 5.586439841 | 6.80E-04 | 0.02082333 | 0.168862391 | 0.412024508 |
| **UBAC2** | -0.430464773 | NA | -6.153070839 | 6.78E-04 | 0.02082333 | -0.599345462 | -0.261584083 |
| **RPL9** | -0.291665723 | NA | -5.394800367 | 6.86E-04 | 0.020914931 | -0.416693083 | -0.166638362 |
| **POLR1A** | 0.246299914 | -2.02151197 | 5.979572305 | 6.96E-04 | 0.021100651 | 0.147646128 | 0.3449537 |
| **GNG12** | -1.003553262 | NA | -6.00729861 | 7.00E-04 | 0.021101461 | -1.404400536 | -0.602705989 |
| **ENO1** | 0.410047856 | -1.2861358 | 6.457545427 | 7.15E-04 | 0.021238416 | 0.253810733 | 0.56628498 |
| **PSMB5** | 0.314413258 | -1.669266042 | 5.328507616 | 7.15E-04 | 0.021238416 | 0.178227279 | 0.450599237 |
| **SELENOS** | 1.473716435 | 0.559458955 | 6.965958853 | 7.15E-04 | 0.021238416 | 0.940154935 | 2.007277935 |
| **AKAP12** | -0.232004924 | NA | -6.271686351 | 7.35E-04 | 0.0213283 | -0.322303797 | -0.141706051 |
| **COPS7B** | -0.253871379 | NA | -7.793517305 | 7.26E-04 | 0.0213283 | -0.33915775 | -0.168585008 |
| **HSD17B8** | 0.174665148 | -2.517336326 | 5.365093038 | 7.26E-04 | 0.0213283 | 0.099290448 | 0.250039848 |
| **KTI12** | -0.410025431 | NA | -5.537888824 | 7.38E-04 | 0.0213283 | -0.583437953 | -0.236612908 |
| **SYNCRIP** | -0.137553872 | NA | -6.264349277 | 7.41E-04 | 0.0213283 | -0.191160867 | -0.083946877 |
| **ZWILCH** | 0.292984449 | -1.771104003 | 5.311499983 | 7.30E-04 | 0.0213283 | 0.165674106 | 0.420294792 |
| **HSPH1** | -0.232684354 | NA | -6.691391743 | 7.47E-04 | 0.021412677 | -0.319496291 | -0.145872416 |
| **DNAJC7** | -0.169012481 | NA | -5.615469331 | 7.63E-04 | 0.021435375 | -0.239973764 | -0.098051197 |
| **KIF22** | -0.175504164 | NA | -5.287938282 | 7.60E-04 | 0.021435375 | -0.25215408 | -0.098854248 |

| | | | | | | | |
|---|---|---|---|---|---|---|---|
| **OAT** | 0.300895548 | -1.732665334 | 5.681339526 | 7.56E-04 | 0.021435375 | 0.175601997 | 0.426189099 |
| **THRAP3** | -0.247044427 | NA | -5.550406504 | 7.57E-04 | 0.021435375 | -0.351522201 | -0.142566652 |
| **ATP5F1C** | 0.504489481 | -0.987103906 | 5.620688201 | 7.99E-04 | 0.022074084 | 0.29224203 | 0.716736932 |
| **PALM2AKAP** | -0.295471673 | NA | -5.242651361 | 7.91E-04 | 0.022074084 | -0.425529113 | -0.165414234 |
| **PSMB3** | 0.497321114 | -1.007750411 | 6.542524859 | 7.98E-04 | 0.022074084 | 0.308149559 | 0.68649267 |
| **UNG** | 1.875313432 | 0.907131742 | 5.588675174 | 8.01E-04 | 0.022074084 | 1.08324538 | 2.667381483 |
| **CMAS** | 0.390503229 | -1.356593616 | 5.442231604 | 8.21E-04 | 0.022448388 | 0.222427605 | 0.558578854 |
| **CNPY2** | -0.36064306 | NA | -5.427276106 | 8.22E-04 | 0.022448388 | -0.516162569 | -0.20512355 |
| **TACO1** | 0.550634662 | -0.860832666 | 5.210626221 | 8.29E-04 | 0.022514714 | 0.306670128 | 0.794599196 |
| **DCXR** | 0.393131816 | -1.346914969 | 5.97396675 | 8.34E-04 | 0.022553814 | 0.233840946 | 0.552422687 |
| **ATOX1** | -0.85650083 | NA | -5.616488271 | 8.43E-04 | 0.022590311 | -1.21816687 | -0.49483479 |
| **RAPH1** | 0.728394828 | -0.457207417 | 7.920493569 | 8.41E-04 | 0.022590311 | 0.483631556 | 0.973158099 |
| **CBX1** | -0.394728836 | NA | -5.18428211 | 8.69E-04 | 0.023159228 | -0.570652033 | -0.21880564 |
| **ANXA2** | -0.310347238 | NA | -6.686635163 | 8.75E-04 | 0.023233254 | -0.4274356 | -0.193258875 |
| **PKN2** | 0.198881886 | -2.330016215 | 5.188289695 | 8.82E-04 | 0.023310423 | 0.110210468 | 0.287553304 |
| **AHNAK** | -0.36095405 | NA | -5.729589855 | 9.02E-04 | 0.023374599 | -0.512006563 | -0.209901538 |
| **ERP29** | 0.335079354 | -1.577425298 | 5.211501237 | 9.13E-04 | 0.023374599 | 0.185825699 | 0.484333009 |
| **FAM120A** | -0.23871864 | NA | -5.145812086 | 9.05E-04 | 0.023374599 | -0.345872211 | -0.131565068 |
| **H4C1** | 0.659851698 | -0.59978628 | 8.23533725 | 9.07E-04 | 0.023374599 | 0.442503209 | 0.877200188 |
| **LOX** | -1.5892447 | NA | -5.75624883 | 9.02E-04 | 0.023374599 | -2.25230138 | -0.926188021 |
| **MAP7D1** | 0.459778086 | -1.120990388 | 5.684990164 | 9.12E-04 | 0.023374599 | 0.266248426 | 0.653307747 |
| **NPTN** | -0.384412581 | NA | -5.20609312 | 8.89E-04 | 0.023374599 | -0.555493418 | -0.213331744 |
| **ISG15** | 0.441160879 | -1.180623232 | 8.401769125 | 9.19E-04 | 0.023411826 | 0.297623626 | 0.584698132 |
| **POLR2G** | 0.457244704 | -1.128961635 | 5.18941848 | 9.32E-04 | 0.023642451 | 0.252778354 | 0.661711054 |
| **ASAP1** | -0.310540963 | NA | -5.088981126 | 9.55E-04 | 0.023800415 | -0.451360031 | -0.169721895 |
| **FOLR1** | 0.704883587 | -0.504543081 | 6.369459873 | 9.54E-04 | 0.023800415 | 0.428633612 | 0.981133563 |
| **GNL3L** | -0.236623221 | NA | -5.079861185 | 9.53E-04 | 0.023800415 | -0.344039693 | -0.129206749 |
| **SAP30** | 0.973816779 | -0.038277736 | 6.407565484 | 9.44E-04 | 0.023800415 | 0.59393965 | 1.353693908 |
| **DUT** | 0.190687562 | -2.390717347 | 5.5068639 | 9.64E-04 | 0.023918041 | 0.108467756 | 0.272907369 |
| **MCM7** | -0.230607336 | NA | -5.449330956 | 9.85E-04 | 0.024168711 | -0.330853325 | -0.130361347 |
| **MRPL12** | 0.377786456 | -1.404357113 | 5.056241736 | 9.86E-04 | 0.024168711 | 0.205442942 | 0.550129971 |
| **PIK3R2** | 0.563780142 | -0.826795433 | 5.496757108 | 9.81E-04 | 0.024168711 | 0.320134024 | 0.807426259 |
| **MAN2B1** | 0.162160315 | -2.624507296 | 5.049416639 | 9.92E-04 | 0.024198166 | 0.088096682 | 0.236223949 |
| **TRAPPC6B** | 0.210446318 | -2.248475825 | 6.675933922 | 1.02E-03 | 0.024673246 | 0.130115013 | 0.290777624 |
| **S100A4** | 2.269933416 | 1.18264998 | 8.372393216 | 1.02E-03 | 0.024743966 | 1.522836783 | 3.01703005 |
| **PDE6D** | 0.207651539 | -2.267763527 | 5.050759637 | 1.05E-03 | 0.025379403 | 0.112494322 | 0.302808757 |
| **NOP2** | -0.156608118 | NA | -5.067197185 | 1.06E-03 | 0.025415549 | -0.228249673 | -0.084966562 |
| **PDLIM3** | 0.45984901 | -1.12076786 | 4.982996147 | 1.08E-03 | 0.025896831 | 0.246947032 | 0.672750988 |
| **TOMM7** | 2.347971358 | 1.23141481 | 6.686564156 | 1.09E-03 | 0.025943262 | 1.447856751 | 3.248085966 |
| **NUP35** | -0.136108931 | NA | -5.107501427 | 1.11E-03 | 0.026238782 | -0.198224455 | -0.073993407 |
| **PURB** | 0.611257653 | -0.710147473 | 6.533479121 | 1.12E-03 | 0.026359304 | 0.372944839 | 0.849570466 |

| | | | | | | | |
|---|---|---|---|---|---|---|---|
| **ITSN1** | -0.363961356 | NA | -6.21634477 | 1.13E-03 | 0.02645037 | -0.510578997 | -0.217343714 |
| **COPZ1** | -0.491698321 | NA | -4.923231332 | 1.16E-03 | 0.026762145 | -0.722021364 | -0.261375278 |
| **DLD** | -0.178965131 | NA | -5.115972006 | 1.17E-03 | 0.026762145 | -0.260812661 | -0.0971176 |
| **EEF1A2** | -1.595275075 | NA | -5.332172024 | 1.16E-03 | 0.026762145 | -2.305874152 | -0.884675998 |
| **SMARCAD1** | -0.293861914 | NA | -4.951285647 | 1.16E-03 | 0.026762145 | -0.431016289 | -0.156707539 |
| **STK39** | -0.377295357 | NA | -5.574612977 | 1.15E-03 | 0.026762145 | -0.540507539 | -0.214083175 |
| **UTF1** | -0.204442502 | NA | -4.918524413 | 1.17E-03 | 0.026762145 | -0.300294532 | -0.108590472 |
| **ZYX** | -0.319227845 | NA | -4.978131944 | 1.18E-03 | 0.026862395 | -0.467834545 | -0.170621146 |
| **TMED4** | -0.29178927 | NA | -4.964228563 | 1.20E-03 | 0.027391508 | -0.428056466 | -0.155522073 |
| **VPS26B** | 1.027902874 | 0.039703951 | 6.296812749 | 1.22E-03 | 0.027695074 | 0.614724158 | 1.44108159 |
| **15,00 ARL** | 0.865399087 | -0.208562496 | 5.409054979 | 1.23E-03 | 0.027779949 | 0.482003019 | 1.248795155 |
| **EMC4** | -0.433267748 | NA | -4.866935681 | 1.25E-03 | 0.028025871 | -0.638577605 | -0.227957891 |
| **F8A1** | 0.504356371 | -0.987484612 | 5.058272672 | 1.27E-03 | 0.028025871 | 0.270847775 | 0.737864968 |
| **RTCB** | -0.249566427 | NA | -5.263421444 | 1.25E-03 | 0.028025871 | -0.362187929 | -0.136944924 |
| **SOX2** | -0.471142095 | NA | -4.963486179 | 1.26E-03 | 0.028025871 | -0.691846576 | -0.250437615 |
| **UBTF** | -0.177230735 | NA | -5.524415668 | 1.26E-03 | 0.028025871 | -0.254848566 | -0.099612904 |
| **INTS4** | -0.493684921 | NA | -4.848546023 | 1.28E-03 | 0.028130912 | -0.728579811 | -0.258790031 |
| **MBIP** | 0.969284766 | -0.045007519 | 5.759428593 | 1.28E-03 | 0.028130912 | 0.555455764 | 1.383113767 |
| **NUP62** | -0.4101426 | NA | -5.54403481 | 1.28E-03 | 0.028130912 | -0.589569219 | -0.230715982 |
| **MTREX** | 0.2664873 | -1.907861317 | 4.900306526 | 1.29E-03 | 0.028173308 | 0.140466019 | 0.39250858 |
| **TMEM230** | 1.288728557 | 0.365948423 | 5.252405803 | 1.30E-03 | 0.028173308 | 0.705035755 | 1.872421359 |
| **VBP1** | 0.182534514 | -2.45375882 | 4.847938147 | 1.31E-03 | 0.028173308 | 0.095580636 | 0.269488391 |
| **WDR74** | -0.287317701 | NA | -5.08117945 | 1.30E-03 | 0.028173308 | -0.420225012 | -0.154410391 |
| **ADRM1** | 0.458911759 | -1.12371132 | 7.032645689 | 1.33E-03 | 0.028471661 | 0.285858413 | 0.631965105 |
| **GLYR1** | -0.130233173 | NA | -5.13173482 | 1.33E-03 | 0.028471661 | -0.190174712 | -0.070291635 |
| **SS18L2** | 0.483898791 | -1.04722276 | 5.565413586 | 1.34E-03 | 0.028508997 | 0.272130223 | 0.695667359 |
| **TOR1AIP1** | 0.270779246 | -1.884810929 | 4.800256381 | 1.36E-03 | 0.028796985 | 0.140699091 | 0.400859401 |
| **CRAT** | 0.504630373 | -0.986701052 | 6.215659615 | 1.38E-03 | 0.028941408 | 0.298201099 | 0.711059648 |
| **TAF10** | 2.306773434 | 1.205876313 | 6.442560742 | 1.37E-03 | 0.028941408 | 1.384724584 | 3.228822284 |
| **XIAP** | 0.194094806 | -2.365166583 | 4.894656514 | 1.37E-03 | 0.028941408 | 0.101892539 | 0.286297073 |
| **ACOT9** | 0.179431874 | -2.478491901 | 4.864653779 | 1.41E-03 | 0.029540903 | 0.093709908 | 0.26515384 |
| **RMDN1** | 0.355777666 | -1.490952148 | 6.50124135 | 1.42E-03 | 0.029540903 | 0.213998306 | 0.497557026 |
| **LARP7** | 0.276940224 | -1.852353481 | 5.081229976 | 1.43E-03 | 0.029633569 | 0.148053546 | 0.405826903 |
| **SCFD1** | -0.23419084 | NA | -4.902906208 | 1.43E-03 | 0.029633569 | -0.34561185 | -0.122769831 |
| **EEF2K** | 0.42526613 | -1.233562136 | 4.835785752 | 1.45E-03 | 0.029752467 | 0.221057022 | 0.629475238 |
| **TFIP11** | 0.265629273 | -1.912513948 | 4.936359254 | 1.44E-03 | 0.029752467 | 0.139692319 | 0.391566228 |
| **FTH1** | -0.367652026 | NA | -4.943459707 | 1.45E-03 | 0.02980439 | -0.541877579 | -0.193426474 |
| **DDX3X** | 0.428096849 | -1.223990877 | 5.411281669 | 1.46E-03 | 0.02984011 | 0.236211636 | 0.619982063 |
| **HNRNPA2B1** | -0.285204127 | NA | -4.82931893 | 1.51E-03 | 0.030700037 | -0.422643621 | -0.147764634 |
| **ENPP1** | -0.861779372 | NA | -5.684067341 | 1.53E-03 | 0.031017162 | -1.237694578 | -0.485864166 |
| **KDM3B** | -0.190224755 | NA | -4.833924651 | 1.55E-03 | 0.031278405 | -0.28199568 | -0.09845383 |

| | | | | | | | |
|---|---|---|---|---|---|---|---|
| **BROX** | 0.731061059 | -0.451936188 | 6.194989537 | 1.58E-03 | 0.031757034 | 0.427959326 | 1.034162792 |
| **DNAJB1** | -0.292018785 | NA | -6.374636178 | 1.59E-03 | 0.031757034 | -0.411008579 | -0.173028991 |
| **DNAJC11** | -0.326423892 | NA | -5.074831349 | 1.59E-03 | 0.031757034 | -0.479535479 | -0.173312305 |
| **POLR2E** | 0.156029056 | -2.68011338 | 4.723603599 | 1.59E-03 | 0.031757034 | 0.079542629 | 0.232515483 |
| **TRADD** | 0.54383998 | -0.878745882 | 5.282607413 | 1.62E-03 | 0.03213458 | 0.294582508 | 0.793097451 |
| **ARID1A** | -0.109273227 | NA | -4.6555489 | 1.65E-03 | 0.032448222 | -0.163425614 | -0.055120841 |
| **SDHA** | 0.268807771 | -1.89535325 | 4.99153488 | 1.64E-03 | 0.032448222 | 0.141125355 | 0.396490187 |
| **GAA** | 0.347620319 | -1.524415684 | 4.815733235 | 1.68E-03 | 0.032623118 | 0.178623942 | 0.516616695 |
| **INTS2** | -0.676916333 | NA | -5.404519519 | 1.67E-03 | 0.032623118 | -0.983524146 | -0.370308521 |
| **KLC1** | -0.299549857 | NA | -5.227723415 | 1.66E-03 | 0.032623118 | -0.43800569 | -0.161094025 |
| **SRP9** | -0.243451648 | NA | -4.862588496 | 1.67E-03 | 0.032623118 | -0.36109823 | -0.125805066 |
| **NACC1** | -0.166934519 | NA | -4.955981654 | 1.69E-03 | 0.032644916 | -0.246731291 | -0.087137746 |
| **SOAT1** | -0.326188097 | NA | -4.667388303 | 1.69E-03 | 0.032644916 | -0.487856671 | -0.164519523 |
| **ADA** | -0.179696946 | NA | -4.869905372 | 1.70E-03 | 0.032685899 | -0.266539337 | -0.092854555 |
| **JPT1** | -0.232683395 | NA | -5.346707453 | 1.71E-03 | 0.032844064 | -0.33898643 | -0.12638036 |
| **ADNP** | 0.130704629 | -2.935617861 | 4.622608657 | 1.72E-03 | 0.032868061 | 0.065465694 | 0.195943564 |
| **AP1G1** | 0.363405002 | -1.460349817 | 5.976086667 | 1.72E-03 | 0.032868061 | 0.208243177 | 0.518566827 |
| **PTGFRN** | 0.409713491 | -1.287312696 | 5.233811606 | 1.74E-03 | 0.03307637 | 0.219838337 | 0.599588646 |
| **LYPLA2** | 0.239540326 | -2.061659545 | 5.74036954 | 1.76E-03 | 0.033270479 | 0.13454186 | 0.344538792 |
| **ILK** | 0.242620952 | -2.043223949 | 4.636158935 | 1.77E-03 | 0.033277424 | 0.121512471 | 0.363729434 |
| **SSR4** | 0.235586964 | -2.085668385 | 4.667809607 | 1.77E-03 | 0.033277424 | 0.118459963 | 0.352713965 |
| **PYCR2** | 0.215691855 | -2.212956394 | 4.628821831 | 1.78E-03 | 0.033451139 | 0.107854284 | 0.323529427 |
| **IPMK** | -0.661900266 | NA | -4.659103458 | 1.85E-03 | 0.034467663 | -0.99236084 | -0.331439692 |
| **LAMC1** | -0.144446692 | NA | -4.570990309 | 1.85E-03 | 0.034467663 | -0.217379238 | -0.071514146 |
| **ADSS2** | -0.148089885 | NA | -4.542761431 | 1.90E-03 | 0.03507441 | -0.223284855 | -0.072894915 |
| **GNG5** | 0.631384243 | -0.663409838 | 4.555769821 | 1.89E-03 | 0.03507441 | 0.311422327 | 0.951346159 |
| **RBPJ** | 0.377050007 | -1.407172219 | 4.893793026 | 1.90E-03 | 0.03507441 | 0.193920125 | 0.560179889 |
| **C1orf122** | -0.363041642 | NA | -5.788840858 | 1.92E-03 | 0.035083312 | -0.522553996 | -0.203529287 |
| **CHTOP** | -0.143413042 | NA | -5.759297862 | 1.92E-03 | 0.035083312 | -0.206616044 | -0.08021004 |
| **CTPS1** | 0.163852826 | -2.609527535 | 4.563304734 | 1.93E-03 | 0.035083312 | 0.080793573 | 0.24691208 |
| **DDX18** | -0.156936091 | NA | -4.708880799 | 1.93E-03 | 0.035083312 | -0.235030994 | -0.078841189 |
| **GOLPH3** | -0.334290718 | NA | -4.723305474 | 1.93E-03 | 0.035083312 | -0.50033828 | -0.168243157 |
| **CNN2** | -0.357773301 | NA | -5.380768841 | 1.95E-03 | 0.035128614 | -0.522298871 | -0.193247731 |
| **LTN1** | -0.350385354 | NA | -5.099545106 | 1.95E-03 | 0.035128614 | -0.516772262 | -0.183998447 |
| **NPC1** | 0.654241332 | -0.612105191 | 4.528481584 | 1.96E-03 | 0.035128614 | 0.320735462 | 0.987747201 |
| **WDR3** | 0.437259644 | -1.193437891 | 4.547063879 | 1.94E-03 | 0.035128614 | 0.215032671 | 0.659486618 |
| **NUP54** | -0.288806418 | NA | -4.52212486 | 1.97E-03 | 0.035196971 | -0.436201734 | -0.141411101 |
| **PATJ** | 0.27833682 | -1.845096324 | 4.661106955 | 1.98E-03 | 0.035245798 | 0.138757926 | 0.417915715 |
| **AIDA** | -0.412274125 | NA | -5.065877288 | 1.99E-03 | 0.035315253 | -0.609108765 | -0.215439484 |
| **CDK12** | -0.30094441 | NA | -4.573144168 | 2.01E-03 | 0.035547855 | -0.453726722 | -0.148162097 |
| **CASC3** | -0.468726182 | NA | -4.876665922 | 2.02E-03 | 0.035652335 | -0.697947948 | -0.239504415 |

| | | | | | | | |
|---|---|---|---|---|---|---|---|
| **SERPINB6** | 0.247038715 | -2.017190943 | 4.64120584 | 2.02E-03 | 0.035652335 | 0.122630054 | 0.371447376 |
| **RALGPS2** | 0.624807276 | -0.678516842 | 4.643602745 | 2.05E-03 | 0.03591116 | 0.310017101 | 0.93959745 |
| **HIBCH** | 0.456470744 | -1.131405696 | 4.534559323 | 2.06E-03 | 0.03613463 | 0.223103467 | 0.689838022 |
| **PPP5C** | -0.170594826 | NA | -5.500503061 | 2.10E-03 | 0.036454785 | -0.248470465 | -0.092719187 |
| **TRAPPC8** | -0.31353648 | NA | -5.014541143 | 2.09E-03 | 0.036454785 | -0.464752125 | -0.162320835 |
| **PPP4R2** | 0.228698558 | -2.128480828 | 4.965539828 | 2.12E-03 | 0.036739745 | 0.117652942 | 0.339744174 |
| **NUMB** | 0.174345418 | -2.519979647 | 4.451016906 | 2.14E-03 | 0.036955957 | 0.084018286 | 0.26467255 |
| **BCLAF1** | -0.33069134 | NA | -4.451463816 | 2.15E-03 | 0.037029695 | -0.50207062 | -0.159312059 |
| **DDX51** | -0.204879894 | NA | -4.453367107 | 2.16E-03 | 0.037170914 | -0.311089207 | -0.098670581 |
| **VRK1** | -0.287117227 | NA | -5.794969552 | 2.17E-03 | 0.037170914 | -0.414548317 | -0.159686137 |
| **SEC24C** | 0.157657694 | -2.665132513 | 4.553728059 | 2.18E-03 | 0.037190347 | 0.076949359 | 0.238366029 |
| **ARPC5L** | -0.494069267 | NA | -5.90924721 | 2.24E-03 | 0.037381494 | -0.711451859 | -0.276686675 |
| **LUC7L2** | -0.162703456 | NA | -5.081248293 | 2.25E-03 | 0.037381494 | -0.241024798 | -0.084382114 |
| **MEA1** | 0.583134036 | -0.778100562 | 4.953565799 | 2.22E-03 | 0.037381494 | 0.298512019 | 0.867756054 |
| **NVL** | -0.243332712 | NA | -4.507037796 | 2.24E-03 | 0.037381494 | -0.368912307 | -0.117753118 |
| **PDS5A** | -0.2311466 | NA | -4.784654589 | 2.23E-03 | 0.037381494 | -0.346322315 | -0.115970886 |
| **SDF2** | -0.369578177 | NA | -4.638927256 | 2.21E-03 | 0.037381494 | -0.556941837 | -0.182214517 |
| **SPNS1** | 0.748787803 | -0.417371159 | 4.425508829 | 2.23E-03 | 0.037381494 | 0.358347092 | 1.139228515 |
| **STAT3** | -0.192143684 | NA | -4.753654642 | 2.25E-03 | 0.037381494 | -0.288313202 | -0.095974167 |
| **TAOK3** | -0.404704258 | NA | -5.719675689 | 2.23E-03 | 0.037381494 | -0.586196444 | -0.223212071 |
| **VPS35L** | -0.612402328 | NA | -6.403621961 | 2.20E-03 | 0.037381494 | -0.868777901 | -0.356026755 |
| **SIKE1** | 0.909188265 | -0.137349032 | 4.615691662 | 2.27E-03 | 0.03764776 | 0.445881168 | 1.372495361 |
| **CALU** | -0.330033324 | NA | -4.416662419 | 2.28E-03 | 0.037676949 | -0.502605379 | -0.157461268 |
| **PFDN2** | 0.721370947 | -0.471186774 | 4.583079343 | 2.31E-03 | 0.037935546 | 0.351886366 | 1.090855528 |
| **PLEKHA5** | 0.148864372 | -2.74792958 | 4.550129265 | 2.33E-03 | 0.038005954 | 0.072225601 | 0.225503144 |
| **SLC25A5** | 0.273339834 | -1.871232372 | 4.396755707 | 2.33E-03 | 0.038005954 | 0.129810392 | 0.416869277 |
| **TTC1** | -0.099344705 | NA | -4.521305003 | 2.32E-03 | 0.038005954 | -0.150652099 | -0.048037311 |
| **ZHX2** | 0.656451513 | -0.607239641 | 6.15522449 | 2.34E-03 | 0.038005954 | 0.373398727 | 0.939504299 |
| **MAPK9** | 0.596812222 | -0.744651014 | 6.325848514 | 2.34E-03 | 0.038011254 | 0.343550042 | 0.850074402 |
| **ANP32B** | -0.579738304 | NA | -4.573679577 | 2.38E-03 | 0.038312009 | -0.877755107 | -0.281721501 |
| **CDC37** | -0.222841118 | NA | -4.370260518 | 2.40E-03 | 0.038312009 | -0.340493002 | -0.105189234 |
| **DHX15** | 0.313785049 | -1.672151481 | 5.819738899 | 2.37E-03 | 0.038312009 | 0.17373136 | 0.453838738 |
| **PCGF6** | 1.676715172 | 0.745637635 | 5.658229288 | 2.41E-03 | 0.038312009 | 0.914580982 | 2.438849362 |
| **PPP1R8** | -0.445245774 | NA | -5.065397279 | 2.41E-03 | 0.038312009 | -0.661220078 | -0.229271469 |
| **REPS1** | -0.266985954 | NA | -4.457706933 | 2.40E-03 | 0.038312009 | -0.406380426 | -0.127591482 |
| **TAGLN** | 0.327964183 | -1.608389829 | 5.595997314 | 2.40E-03 | 0.038312009 | 0.177957344 | 0.477971022 |
| **WASHC2A** | -0.455901052 | NA | -4.537748144 | 2.39E-03 | 0.038312009 | -0.691378549 | -0.220423555 |
| **FAU** | 0.507254468 | -0.979218427 | 4.677488804 | 2.43E-03 | 0.038394412 | 0.249468725 | 0.765040211 |
| **SNTB2** | -0.329719824 | NA | -5.084086685 | 2.43E-03 | 0.038418733 | -0.489426002 | -0.170013647 |
| **ABHD10** | 0.452587588 | -1.143731075 | 4.356234339 | 2.49E-03 | 0.039106609 | 0.212571925 | 0.69260325 |
| **RHOT2** | 0.234636512 | -2.091500568 | 4.350590568 | 2.53E-03 | 0.039586212 | 0.109964227 | 0.359308796 |

| | | | | | | | |
|---|---|---|---|---|---|---|---|
| **SMC3** | -0.151889836 | NA | -4.32523946 | 2.53E-03 | 0.039586212 | -0.23287335 | -0.070906323 |
| **HNRNPH1** | -0.17144281 | NA | -5.605451944 | 2.55E-03 | 0.03960099 | -0.250216693 | -0.092668927 |
| **PGAM5** | 0.201120808 | -2.313865747 | 4.314988874 | 2.56E-03 | 0.03960099 | 0.093634372 | 0.308607243 |
| **SRM** | 0.193857244 | -2.366933445 | 4.337464327 | 2.56E-03 | 0.03960099 | 0.090576443 | 0.297138046 |
| **SRSF5** | -0.241683722 | NA | -4.701197808 | 2.56E-03 | 0.03960099 | -0.364693801 | -0.118673643 |
| **ZNF384** | 0.496419695 | -1.01036774 | 5.172416929 | 2.55E-03 | 0.03960099 | 0.257369729 | 0.735469661 |
| **CNOT2** | -0.291566879 | NA | -5.326393892 | 2.61E-03 | 0.039606148 | -0.429891272 | -0.153242487 |
| **H3-7** | 0.56699472 | -0.818592794 | 4.49749273 | 2.58E-03 | 0.039606148 | 0.270912857 | 0.863076583 |
| **MRPL33** | 0.707470035 | -0.49925905 | 4.618384793 | 2.60E-03 | 0.039606148 | 0.34331409 | 1.071625981 |
| **NHERF2** | -0.160385836 | NA | -4.425726561 | 2.61E-03 | 0.039606148 | -0.244983116 | -0.075788556 |
| **SSNA1** | -3.654010387 | NA | -6.080941084 | 2.59E-03 | 0.039606148 | -5.257247744 | -2.05077303 |
| **SYF2** | -0.321266233 | NA | -4.428306882 | 2.60E-03 | 0.039606148 | -0.490629237 | -0.151903229 |
| **TOP3B** | -0.446483166 | NA | -4.673516831 | 2.61E-03 | 0.039606148 | -0.67484555 | -0.218120781 |
| **SLC35A4** | 2.501300058 | 1.322678135 | 5.99492764 | 2.66E-03 | 0.040259285 | 1.391355897 | 3.611244219 |
| **AK3** | 0.366615776 | -1.447659227 | 5.134539541 | 2.70E-03 | 0.040647906 | 0.188420284 | 0.544811268 |
| **VCP** | -0.201970131 | NA | -4.48327824 | 2.70E-03 | 0.040647906 | -0.308027383 | -0.095912879 |
| **ERO1A** | 0.319683574 | -1.645283476 | 5.153415246 | 2.73E-03 | 0.040902955 | 0.164486722 | 0.474880427 |
| **SHC1** | -0.267770086 | NA | -4.267253979 | 2.73E-03 | 0.040902955 | -0.41247218 | -0.123067991 |
| **MED9** | -0.317019232 | NA | -6.408736928 | 2.75E-03 | 0.040936522 | -0.452797092 | -0.181241371 |
| **PEPD** | 0.263394566 | -1.924702514 | 4.577079768 | 2.75E-03 | 0.040936522 | 0.126498339 | 0.400290792 |
| **CLINT1** | -0.344942888 | NA | -4.507169419 | 2.81E-03 | 0.041780692 | -0.526125337 | -0.16376044 |
| **EML4** | -0.221651152 | NA | -5.138693248 | 2.90E-03 | 0.042873204 | -0.330020066 | -0.113282239 |
| **SIN3B** | 0.436763911 | -1.19507444 | 4.971986497 | 2.93E-03 | 0.043288929 | 0.218946208 | 0.654581615 |
| **BACH1** | 0.712427076 | -0.489185748 | 4.249747488 | 2.96E-03 | 0.043327398 | 0.324212543 | 1.100641608 |
| **LIMD1** | -0.586452553 | NA | -5.756547409 | 2.96E-03 | 0.043327398 | -0.855822006 | -0.3170831 |
| **RBM26** | 0.213278146 | -2.229191951 | 4.708532154 | 2.94E-03 | 0.043327398 | 0.103561047 | 0.322995245 |
| **SNX30** | 0.314729513 | -1.667815623 | 4.233870304 | 2.96E-03 | 0.043327398 | 0.142862575 | 0.486596451 |
| **ABCB7** | 0.560270931 | -0.835803453 | 4.478536421 | 2.99E-03 | 0.043576259 | 0.263472306 | 0.857069556 |
| **ANKS1A** | -0.31441748 | NA | -4.305420132 | 3.00E-03 | 0.043610267 | -0.484709491 | -0.144125469 |
| **KDM3A** | -0.180849893 | NA | -5.137866371 | 3.03E-03 | 0.043888931 | -0.269652756 | -0.09204703 |
| **TXNDC5** | 0.225378445 | -2.14957855 | 4.540672954 | 3.03E-03 | 0.043888931 | 0.106751751 | 0.34400514 |
| **TMSB4X** | -0.643552412 | NA | -4.209264301 | 3.05E-03 | 0.043954825 | -0.996913501 | -0.290191322 |
| **ZNF462** | 0.408901691 | -1.290174067 | 4.984162421 | 3.06E-03 | 0.044096153 | 0.204434612 | 0.61336877 |
| **CCDC137** | 0.345230048 | -1.534370055 | 4.192508283 | 3.10E-03 | 0.044213991 | 0.155005169 | 0.535454928 |
| **MED12** | 0.56380483 | -0.826732257 | 4.249349294 | 3.09E-03 | 0.044213991 | 0.255523325 | 0.872086336 |
| **MFAP1** | -0.388822794 | NA | -4.27080597 | 3.10E-03 | 0.044213991 | -0.600929585 | -0.176716003 |
| **TPM2** | -0.370316483 | NA | -4.288067558 | 3.09E-03 | 0.044213991 | -0.571819419 | -0.168813546 |
| **MRPL52** | -1.429862576 | NA | -4.216715589 | 3.14E-03 | 0.044551187 | -2.216138671 | -0.643586481 |
| **RPL4** | 0.150269883 | -2.734372202 | 4.946740604 | 3.14E-03 | 0.044551187 | 0.074639854 | 0.225899912 |
| **PNO1** | -0.563889556 | NA | -5.507486957 | 3.16E-03 | 0.044679609 | -0.831237961 | -0.296541151 |
| **HNRNPD** | -0.113555294 | NA | -4.156307855 | 3.19E-03 | 0.044688966 | -0.176570693 | -0.050539894 |

| | | | | | | | |
|---|---|---|---|---|---|---|---|
| **PPID** | -0.273212043 | NA | -4.717915834 | 3.17E-03 | 0.044688966 | -0.414527282 | -0.131896804 |
| **RPL15** | 0.351613245 | -1.507938678 | 4.485359442 | 3.18E-03 | 0.044688966 | 0.164513819 | 0.538712671 |
| **RPL18A** | 0.302369824 | -1.725613926 | 4.70828824 | 3.18E-03 | 0.044688966 | 0.145763619 | 0.458976029 |
| **UTP11** | 0.2843392 | -1.814315086 | 4.151715194 | 3.21E-03 | 0.044869109 | 0.126373983 | 0.442304418 |
| **BIN3** | 0.480451922 | -1.057536025 | 4.177105745 | 3.22E-03 | 0.044917664 | 0.214346297 | 0.746557547 |
| **DDI2** | -0.251384778 | NA | -4.686098452 | 3.24E-03 | 0.045063133 | -0.382158841 | -0.120610714 |
| **APLP2** | -0.306214044 | NA | -5.642432929 | 3.25E-03 | 0.045079735 | -0.449880073 | -0.162548014 |
| **SMC2** | -0.161881018 | NA | -5.29853927 | 3.26E-03 | 0.045079735 | -0.24056664 | -0.083195396 |
| **MYO1E** | -0.251759341 | NA | -4.750445703 | 3.27E-03 | 0.045120652 | -0.381844622 | -0.121674061 |
| **SLC25A4** | 0.403096413 | -1.31080315 | 5.506608136 | 3.29E-03 | 0.045357062 | 0.211128516 | 0.59506431 |
| **MEAF6** | -0.417391704 | NA | -4.842080529 | 3.31E-03 | 0.045441988 | -0.631028111 | -0.203755297 |
| **TRIM25** | -0.201545276 | NA | -4.144559938 | 3.32E-03 | 0.045518561 | -0.313922737 | -0.089167815 |
| **SETD3** | -0.366041996 | NA | -4.130167144 | 3.34E-03 | 0.045743011 | -0.570645158 | -0.161438833 |
| **DCAF7** | 0.192537402 | -2.376789368 | 4.168556235 | 3.37E-03 | 0.045841762 | 0.085393453 | 0.299681351 |
| **GIPC1** | -0.303525409 | NA | -4.246852951 | 3.36E-03 | 0.045841762 | -0.470723515 | -0.136327303 |
| **CNPY3** | -0.284114008 | NA | -4.111902478 | 3.41E-03 | 0.04619788 | -0.443568666 | -0.12465935 |
| **CORO2A** | 0.525032014 | -0.929522701 | 4.10883647 | 3.41E-03 | 0.04619788 | 0.230295092 | 0.819768935 |
| **FBXO2** | 0.372699641 | -1.423914666 | 4.152160227 | 3.42E-03 | 0.04619788 | 0.164602757 | 0.580796524 |
| **ANKRD13A** | 0.389752724 | -1.35936899 | 4.235471444 | 3.46E-03 | 0.046433489 | 0.174217502 | 0.605287946 |
| **GRK2** | 0.525890451 | -0.927165793 | 4.096989822 | 3.47E-03 | 0.046433489 | 0.229790585 | 0.821990318 |
| **SKIC2** | 0.201496014 | -2.311176798 | 4.475905706 | 3.45E-03 | 0.046433489 | 0.093385719 | 0.309606309 |
| **SLC25A24** | 0.185904721 | -2.42736469 | 4.576352665 | 3.47E-03 | 0.046433489 | 0.087318529 | 0.284490913 |
| **SDCBP** | 0.28337344 | -1.819223552 | 4.128030641 | 3.49E-03 | 0.046591797 | 0.124390027 | 0.442356853 |
| **MTM1** | -0.289782969 | NA | -4.168425552 | 3.50E-03 | 0.046634129 | -0.451557219 | -0.128008719 |
| **MTMR14** | 0.571678318 | -0.80672452 | 4.77445437 | 3.51E-03 | 0.046693773 | 0.275106453 | 0.868250184 |
| **MYH9** | -0.242695549 | NA | -4.576580092 | 3.53E-03 | 0.046844454 | -0.371609078 | -0.11378202 |
| **EMC2** | 0.124398301 | -3.006961318 | 4.078845745 | 3.56E-03 | 0.047135387 | 0.054030961 | 0.19476564 |
| **MOCS2** | 0.557341681 | -0.843366044 | 4.073361243 | 3.58E-03 | 0.047135387 | 0.241747968 | 0.872935395 |
| **STX8** | -0.339472338 | NA | -4.085437967 | 3.57E-03 | 0.047135387 | -0.531384474 | -0.147560201 |
| **MTHFSD** | -1.146248041 | NA | -4.4345619 | 3.59E-03 | 0.047152711 | -1.766613965 | -0.525882117 |
| **RPA1** | 0.120681552 | -3.050722937 | 4.415163914 | 3.59E-03 | 0.047152711 | 0.055211287 | 0.186151817 |
| **EHD1** | -0.559688351 | NA | -4.158298496 | 3.63E-03 | 0.047247106 | -0.873532348 | -0.245844354 |
| **GSDMD** | 0.56722069 | -0.818017938 | 4.053994229 | 3.66E-03 | 0.047247106 | 0.244565556 | 0.889875824 |
| **HCLS1** | -0.26026277 | NA | -4.215983522 | 3.64E-03 | 0.047247106 | -0.405150204 | -0.115375335 |
| **HMGCL** | 0.270499325 | -1.886303099 | 4.972251294 | 3.67E-03 | 0.047247106 | 0.132656641 | 0.40834201 |
| **IWS1** | -0.170913091 | NA | -4.311887014 | 3.62E-03 | 0.047247106 | -0.264893455 | -0.076932726 |
| **NDUFA2** | -0.479109976 | NA | -4.157638845 | 3.63E-03 | 0.047247106 | -0.74777496 | -0.210444993 |
| **PAXX** | 3.409195719 | 1.769431426 | 5.668960691 | 3.65E-03 | 0.047247106 | 1.793695697 | 5.02469574 |
| **RPL27** | -0.337462773 | NA | -4.452011543 | 3.65E-03 | 0.047247106 | -0.519993676 | -0.15493187 |
| **KHDC4** | -0.339147857 | NA | -4.477535049 | 3.68E-03 | 0.047333629 | -0.522153011 | -0.156142702 |
| **BAG6** | -0.190297717 | NA | -4.172257606 | 3.70E-03 | 0.047346221 | -0.296987862 | -0.083607571 |

| | | | | | | | |
|---|---|---|---|---|---|---|---|
| **TXNDC12** | -0.555646142 | NA | -4.048866491 | 3.70E-03 | 0.047346221 | -0.872176188 | -0.239116096 |
| **FAM120C** | 0.185318934 | -2.431917804 | 4.272161825 | 3.74E-03 | 0.047442556 | 0.082626655 | 0.288011214 |
| **MYEF2** | 0.228350824 | -2.130676099 | 4.066888007 | 3.76E-03 | 0.047442556 | 0.098408235 | 0.358293413 |
| **POLE** | 0.293467628 | -1.768726724 | 4.190703216 | 3.73E-03 | 0.047442556 | 0.129211696 | 0.45772356 |
| **PSMC4** | -0.362745967 | NA | -4.674634516 | 3.76E-03 | 0.047442556 | -0.554349312 | -0.171142621 |
| **PTPRA** | 0.594235294 | -0.7508938 | 4.973753843 | 3.76E-03 | 0.047442556 | 0.290722599 | 0.897747988 |
| **SALL4** | -0.245711946 | NA | -4.0994937 | 3.73E-03 | 0.047442556 | -0.38486933 | -0.106554562 |
| **FAM20B** | 0.356166708 | -1.489375425 | 4.998086388 | 3.78E-03 | 0.047468327 | 0.174598431 | 0.537734985 |
| **RANGAP1** | -0.116894796 | NA | -4.28573081 | 3.77E-03 | 0.047468327 | -0.181604982 | -0.05218461 |
| **SH3BP5L** | 0.346805783 | -1.52780014 | 4.044108808 | 3.78E-03 | 0.047468327 | 0.148780157 | 0.544831408 |
| **ATAD1** | -0.388838452 | NA | -4.86127424 | 3.82E-03 | 0.04775817 | -0.590261954 | -0.18741495 |
| **CCT4** | 0.201629311 | -2.310222713 | 4.040016368 | 3.82E-03 | 0.04775817 | 0.086316612 | 0.316942011 |
| **AP1S2** | 0.488096935 | -1.034760402 | 4.25013136 | 3.86E-03 | 0.048016007 | 0.216082844 | 0.760111026 |
| **DPPA4** | -0.738315683 | NA | -5.268065081 | 3.89E-03 | 0.048016007 | -1.105222642 | -0.371408725 |
| **IPO9** | -0.410503955 | NA | -4.282347713 | 3.90E-03 | 0.048016007 | -0.63856359 | -0.18244432 |
| **MAP4K3** | -0.370095727 | NA | -4.043322885 | 3.88E-03 | 0.048016007 | -0.58191417 | -0.158277284 |
| **MRPL15** | 0.219034345 | -2.19077099 | 5.016926884 | 3.90E-03 | 0.048016007 | 0.107239056 | 0.330829634 |
| **MRPL20** | 0.476502214 | -1.069445177 | 4.44035385 | 3.87E-03 | 0.048016007 | 0.217004728 | 0.7359997 |
| **PIN1** | -0.279696393 | NA | -4.366116875 | 3.89E-03 | 0.048016007 | -0.433436166 | -0.12595662 |
| **DPY30** | -0.209137287 | NA | -4.200299528 | 3.92E-03 | 0.048186177 | -0.326581254 | -0.09169332 |
| **SUCLG2** | 0.147944093 | -2.756876 | 4.045192634 | 3.95E-03 | 0.048340797 | 0.063163055 | 0.232725131 |
| **RIMOC1** | -0.76072622 | NA | -3.996861792 | 3.97E-03 | 0.048490686 | -1.19969736 | -0.321755079 |
| **SLTM** | -0.179905788 | NA | -4.037200555 | 3.97E-03 | 0.048490686 | -0.283171037 | -0.07664054 |
| **ABR** | -0.498670676 | NA | -4.737054531 | 4.01E-03 | 0.048669264 | -0.761934005 | -0.235407347 |
| **ASCC2** | -0.326415064 | NA | -3.98773708 | 4.03E-03 | 0.048669264 | -0.515225291 | -0.137604838 |
| **MORC2** | -0.311319382 | NA | -4.873007225 | 4.01E-03 | 0.048669264 | -0.473172631 | -0.149466132 |
| **POLR2A** | -0.134709191 | NA | -4.283835176 | 4.03E-03 | 0.048669264 | -0.209757282 | -0.059661099 |
| **RUVBL2** | 0.255534619 | -1.968409341 | 4.615188811 | 4.03E-03 | 0.048669264 | 0.118695381 | 0.392373857 |
| **HPRT1** | 0.245368188 | -2.026979877 | 4.154322474 | 4.06E-03 | 0.048720731 | 0.10635131 | 0.384385067 |
| **MCM3** | -0.106791018 | NA | -3.99657917 | 4.06E-03 | 0.048720731 | -0.168525061 | -0.045056976 |
| **SNIP1** | -0.268344492 | NA | -4.687293894 | 4.06E-03 | 0.048720731 | -0.410963364 | -0.125725621 |
| **ABRAXAS2** | 0.479009976 | -1.061872392 | 5.042298781 | 4.12E-03 | 0.04892909 | 0.23374768 | 0.724272273 |
| **PAWR** | -0.309335558 | NA | -3.980292227 | 4.11E-03 | 0.04892909 | -0.488742079 | -0.129929038 |
| **PSMD5** | -0.184772143 | NA | -4.287356789 | 4.11E-03 | 0.04892909 | -0.287862982 | -0.081681303 |
| **RACGAP1** | -0.358569542 | NA | -4.047821951 | 4.10E-03 | 0.04892909 | -0.564669333 | -0.15246975 |
| **TEX264** | 0.190942169 | -2.38879234 | 4.274318819 | 4.11E-03 | 0.04892909 | 0.084243526 | 0.297640813 |
| **WDR1** | 0.123510577 | -3.017293496 | 4.299703299 | 4.13E-03 | 0.048967306 | 0.054674071 | 0.192347084 |
| **ARL6IP6** | 0.700017118 | -0.514537893 | 5.127461373 | 4.15E-03 | 0.048989785 | 0.344494545 | 1.055539692 |
| **CCAR2** | -0.186806172 | NA | -5.265302261 | 4.16E-03 | 0.048989785 | -0.280363489 | -0.093248855 |
| **NEPRO** | -0.142961922 | NA | -4.165730895 | 4.17E-03 | 0.048989785 | -0.22404108 | -0.061882765 |
| **NOTCH2** | 0.297792243 | -1.747621921 | 4.130137625 | 4.15E-03 | 0.048989785 | 0.128214761 | 0.467369725 |

| | | | | | | | |
|---|---|---|---|---|---|---|---|
| **TLE3** | -0.165305344 | NA | -4.482805527 | 4.19E-03 | 0.048989785 | -0.2555478 | -0.075062888 |
| **VAPA** | -0.401574958 | NA | -3.980858308 | 4.18E-03 | 0.048989785 | -0.63480161 | -0.168348305 |
| **RBM42** | -0.372082751 | NA | -4.264697629 | 4.21E-03 | 0.049192834 | -0.580718203 | -0.163447299 |
| **RCN2** | -0.413487393 | NA | -4.35161206 | 4.22E-03 | 0.049250499 | -0.642975222 | -0.183999564 |
| **STON1** | 1.106017105 | 0.145373697 | 5.0359937 | 4.23E-03 | 0.049251523 | 0.537647963 | 1.674386247 |
| **ADISSP** | 0.586623397 | -0.769493482 | 4.961342376 | 4.28E-03 | 0.049302431 | 0.28239164 | 0.890855154 |
| **HNRNPA0** | -0.247168732 | NA | -3.989281812 | 4.25E-03 | 0.049302431 | -0.390762528 | -0.103574936 |
| **LRRC47** | 0.174321165 | -2.52018035 | 4.103815768 | 4.28E-03 | 0.049302431 | 0.0744577 | 0.27418463 |
| **SAT2** | 0.352507343 | -1.504274785 | 3.949287116 | 4.28E-03 | 0.049302431 | 0.146506527 | 0.558508158 |
| **TP53BP2** | -0.167133417 | NA | -4.192152458 | 4.27E-03 | 0.049302431 | -0.26181924 | -0.072447593 |
| **PHF6** | -0.534761003 | NA | -4.82293224 | 4.31E-03 | 0.049559173 | -0.81650594 | -0.253016067 |
| **SFSWAP** | 0.383026735 | -1.384483001 | 4.796302835 | 4.32E-03 | 0.049582604 | 0.180580085 | 0.585473384 |

Supplemental Table S4_Sheet_1

| | significant_4 | label | log2FC | p_value |
|---|---|---|---|---|
| 1 | down | CHAMP1_S297 | -0.807814 | 2.03307e-05 |

Supplemental Table S4_Sheet_2

| | significant_12 | label | log2FC | p_value |
|---|---|---|---|---|
| 1 | down | STX6_S2 | -0.756983 | 0.00525211 |
| 2 | down | AHCYL1_S2 | -1.34191 | 0.00645224 |
| 3 | down | EIF5B_S137 | -1.10206 | 0.00995484 |
| 4 | down | WBP4_S277 | -0.675967 | 0.00253812 |
| 5 | down | RPLP0;RPLP0P6_S304 | -0.956734 | 0.00170258 |
| 6 | down | KRT18_S399 | -0.703288 | 0.000751321 |
| 7 | down | TOP1_S10 | -0.925169 | 0.0178502 |
| 8 | down | XRCC6_S477 | -1.10611 | 0.00309438 |
| 9 | down | HNRNPL_S52 | -0.823498 | 0.00261182 |
| 10 | down | RAB3B_S190 | -1.85563 | 0.00952003 |
| 11 | down | FLNA_S1084 | -0.71388 | 0.00367353 |
| 12 | down | EEF1B2_S106 | -0.786134 | 0.00342134 |
| 13 | down | MAP1B_S1785 | -0.678971 | 0.0173808 |
| 14 | down | MARCKSL1_S151 | -0.983252 | 0.00286649 |
| 15 | down | MSH6_S227 | -0.589603 | 0.000767131 |
| 16 | down | RBM5_S59 | -0.802622 | 0.0227612 |
| 17 | down | MFAP1_S116 | -0.597189 | 0.00265701 |
| 18 | down | TPI1_S21 | -0.815141 | 0.0158729 |
| 19 | down | ACTB_S14 | -1.56389 | 0.00017164 |
| 20 | down | ACTG1_S14 | -0.722447 | 0.00126864 |
| 21 | down | PAFAH1B2_S2 | -0.880665 | 0.00239023 |
| 22 | down | PRKDC_S2612 | -0.838897 | 0.0221729 |
| 23 | down | HMGCS1_S4 | -0.718068 | 0.00229587 |
| 24 | down | AKAP12_S1395 | -0.767999 | 0.0108691 |
| 25 | down | TLE3_S203 | -0.635049 | 0.0111248 |
| 26 | down | SRSF11_S483 | -0.669233 | 0.0129728 |
| 27 | down | TP53BP1_S500 | -1.16279 | 0.00903567 |
| 28 | down | ORC1_S199 | -0.814608 | 0.000228522 |
| 29 | down | XRCC4_S328 | -1.11239 | 0.00992487 |
| 30 | down | SF3B2_S289 | -1.0978 | 0.00623281 |
| 31 | down | RBM39_S136 | -1.0499 | 0.00596769 |
| 32 | down | GSE1_S857 | -1.47252 | 0.0221065 |

| | | | | |
|---|---|---|---|---|
| 33 | down | PDIA6_S428 | -0.759262 | 0.0191504 |
| 34 | down | PTGES3_S151 | -0.725133 | 0.00299603 |
| 35 | down | TOMM34_S186 | -0.928213 | 0.00104272 |
| 36 | down | IFI16_S153 | -0.62464 | 0.0193645 |
| 37 | down | UGP2_S13 | -1.41103 | 0.0157862 |
| 38 | down | STRIP1_S335 | -1.12342 | 0.0165155 |
| 39 | down | RNF20_S138 | -0.628113 | 0.0137599 |
| 40 | down | DAB2IP_S747 | -1.148 | 0.0168249 |
| 41 | down | NIPBL_S2658 | -0.843631 | 0.0110339 |
| 42 | down | EIF3M_S2 | -1.05556 | 0.0221148 |
| 43 | down | HUWE1_S2362 | -0.616898 | 7.39006e-05 |
| 44 | down | SDCCAG8_S4 | -0.679996 | 0.0189704 |
| 45 | down | PATL1_S179 | -0.611607 | 0.0180718 |
| 46 | down | SMG6_S332 | -1.18808 | 0.00315477 |
| 47 | up | RALGPS2_S308 | 1.32785 | 2.9303e-06 |
| 48 | down | SERBP1_S25 | -0.59706 | 0.00534956 |
| 49 | down | NFATC2IP_S204 | -1.16637 | 0.00908979 |
| 50 | down | PRUNE2_S597 | -1.14633 | 0.0097658 |
| 51 | down | HDAC2_S424 | -0.812289 | 0.000634145 |
| 52 | down | PBXIP1_S3 | -1.41052 | 0.0157494 |
| 53 | down | DOCK7_S180 | -1.26972 | 0.00525337 |
| 54 | down | NEK1_S1052 | -0.612341 | 0.00992996 |
| 55 | up | SPEN_S1278 | 0.594888 | 0.00885052 |
| 56 | up | SPEN_S1918 | 1.21439 | 0.0118193 |
| 57 | down | TUBA1C_S439 | -0.641057 | 0.00191717 |
| 58 | down | UTP14A_S453 | -0.937603 | 0.0178164 |
| 59 | up | ADNP_S98 | 1.14901 | 0.0210398 |
| 60 | down | PNN_S381 | -0.897705 | 0.0128462 |
| 61 | down | PLEKHA5_S410 | -0.859874 | 0.00152502 |
| 62 | down | RTN4_S15 | -0.767933 | 0.0131867 |
| 63 | down | ACSS2_S267 | -1.09961 | 0.0038338 |
| 64 | up | TOMM22_S15 | 1.61858 | 0.00456758 |
| 65 | down | STAU2_S416 | -1.54838 | 0.00568864 |
| 66 | down | STIM2_S523 | -0.92016 | 0.00462929 |

| | | | | |
|---|---|---|---|---|
| 67 | down | NAGK_S76 | -0.748975 | 0.0129943 |
| 68 | down | SALL4_S789 | -0.972248 | 0.0148552 |
| 69 | down | GTF3C4_S611 | -0.611135 | 0.000807618 |
| 70 | down | TRIM33_S1105 | -0.598614 | 0.000204966 |
| 71 | down | PHF8_S857 | -0.634704 | 0.0173041 |
| 72 | down | THRAP3_S939 | -1.11944 | 0.0199375 |
| 73 | down | LUC7L2_S18 | -0.602098 | 0.006232 |
| 74 | up | CERT1_S373 | 0.678191 | 0.0168058 |
| 75 | down | IGF2BP2_S164 | -1.33535 | 0.000958266 |
| 76 | down | AKAP12_T1264 | -1.25558 | 0.00209152 |
| 77 | down | ZNF609_S358 | -1.07167 | 0.00209676 |
| 78 | down | KDM1A_S131 | -0.619193 | 0.00227466 |
| 79 | down | DKC1_S451 | -0.6785 | 0.00193613 |
| 80 | down | DKC1_S453 | -0.6785 | 0.00193613 |
| 81 | down | EIF3J_S11 | -0.672315 | 0.0209459 |
| 82 | down | EIF3J_S13 | -0.672315 | 0.0209459 |
| 83 | down | TOX4_S178 | -0.740878 | 0.0186095 |
| 84 | down | PDHA1;PDHA2_S293 | -0.716479 | 0.0132308 |
| 85 | down | RNH1_S2 | -0.946104 | 0.00262093 |
| 86 | down | RNH1_S7 | -0.923995 | 0.00466335 |
| 87 | down | PRKAR2A_S78 | -0.731647 | 0.000488688 |
| 88 | down | PRKAR2A_S80 | -0.731647 | 0.000488688 |
| 89 | down | GJA1_S368 | -1.0931 | 0.0149845 |
| 90 | down | LMNB1;LMNB2_S391 | -0.765707 | 0.00373172 |
| 91 | down | LMNB1;LMNB2_S393 | -0.765707 | 0.00373172 |
| 92 | down | FGFR3_S444 | -1.06202 | 0.000441401 |
| 93 | down | FGFR3_S445 | -1.06202 | 0.000441401 |
| 94 | down | MCM3_S711 | -0.814898 | 0.012828 |
| 95 | down | RFC1_S69 | -0.792497 | 0.0168803 |
| 96 | down | RFC1_S71 | -0.792497 | 0.0168803 |
| 97 | down | PPP1R2;PPP1R2B_S12[illegible] | -0.592701 | 0.00569107 |
| 98 | down | PPP1R2;PPP1R2B_S12[illegible] | -0.592701 | 0.00569107 |
| 99 | down | ATRX_S729 | -0.597314 | 0.00324504 |
| 10[illegible] | down | ATRX_S731 | -0.597314 | 0.00324504 |

| | | | | |
|---|---|---|---|---|
| 101 | down | MAP1B_S831 | -0.906628 | 0.019207 |
| 102 | down | MAP1B_S832 | -0.906628 | 0.019207 |
| 103 | down | CLK3_S224 | -0.827604 | 0.00190715 |
| 104 | down | CLK3_S226 | -0.827604 | 0.00190715 |
| 105 | down | RANBP2_S781 | -0.775375 | 0.000177201 |
| 106 | down | MFAP1_S52 | -0.748147 | 0.00153163 |
| 107 | down | MFAP1_S53 | -0.748147 | 0.00153163 |
| 108 | down | PRKDC_S2612 | -0.821096 | 0.0107642 |
| 109 | down | OTUD4_S1023 | -0.942772 | 0.0204214 |
| 110 | down | OTUD4_S1024 | -0.942772 | 0.0204214 |
| 111 | down | TOP2B_S1400 | -1.10935 | 0.00130221 |
| 112 | down | SRSF1_S199 | -0.806192 | 0.0155075 |
| 113 | down | SRSF1_S201 | -0.806192 | 0.0155075 |
| 114 | down | TP53BP1_S523 | -0.721944 | 0.0146063 |
| 115 | down | TP53BP1_S525 | -0.721944 | 0.0146063 |
| 116 | down | SRSF9_S211 | -1.12914 | 0.0139397 |
| 117 | down | SF3B2_S307 | -0.635516 | 0.000738127 |
| 118 | down | SF3B2_S309 | -0.635516 | 0.000738127 |
| 119 | down | BOP1_S126 | -0.607538 | 3.61177e-05 |
| 120 | down | BOP1_S127 | -0.607538 | 3.61177e-05 |
| 121 | down | PCM1_S65 | -0.667238 | 0.00128529 |
| 122 | down | FIP1L1_S492 | -0.734642 | 0.000384394 |
| 123 | down | FIP1L1_S500 | -0.734642 | 0.000384394 |
| 124 | down | RAB11FIP1_S345 | -0.691713 | 0.00167597 |
| 125 | down | PHLDB1_S520 | -1.14381 | 0.000744958 |
| 126 | down | SRRM1_S738 | -0.689459 | 0.00652095 |
| 127 | down | SRRM1_S740 | -0.689459 | 0.00652095 |
| 128 | down | PRUNE2_S595 | -0.917193 | 0.00367444 |
| 129 | down | PRUNE2_S597 | -0.917193 | 0.00367444 |
| 130 | down | RREB1_S1174 | -0.671358 | 0.000377055 |
| 131 | down | RREB1_S1175 | -0.671358 | 0.000377055 |
| 132 | down | IWS1_S438 | -0.605209 | 0.0112308 |
| 133 | down | IWS1_S440 | -0.605209 | 0.0112308 |
| 134 | down | RBM15_S670 | -0.705622 | 0.00765375 |

| | | | | |
|---|---|---|---|---|
| 135 | down | XRN2_S499 | -0.727564 | 0.000766022 |
| 136 | down | XRN2_S501 | -0.727564 | 0.000766022 |
| 137 | down | ADNP_S953 | -0.654807 | 0.00164843 |
| 138 | down | ADNP_S955 | -0.654807 | 0.00164843 |
| 139 | down | RTN4_S7 | -0.709298 | 0.00057353 |
| 140 | down | RCC2_S50 | -0.696971 | 0.0177881 |
| 141 | down | RCC2_S51 | -0.696971 | 0.0177881 |
| 142 | down | TJP2_S174 | -0.649711 | 0.0204136 |
| 143 | up | ACIN1_S240 | 0.637426 | 0.0164019 |
| 144 | down | PRR12_S1381 | -0.696104 | 0.0101642 |
| 145 | down | PRR12_S1382 | -0.696104 | 0.0101642 |
| 146 | down | LIMCH1_S233 | -0.851303 | 0.0093272 |
| 147 | down | ZC3H4_S1269 | -0.986696 | 0.0198259 |
| 148 | down | SRRM2_S2121 | -0.702905 | 0.00116923 |
| 149 | down | SRRM2_S2123 | -0.702905 | 0.00116923 |
| 150 | down | SRRM2_S295 | -0.641835 | 0.00498907 |
| 151 | down | SRRM2_S297 | -0.879532 | 0.00736193 |
| 152 | down | NOC2L_S672 | -0.593506 | 5.89665e-06 |
| 153 | down | NOC2L_S673 | -0.593506 | 5.89665e-06 |
| 154 | down | SAMHD1_S18 | -0.66348 | 0.0169808 |
| 155 | down | SAMHD1_S33 | -0.66348 | 0.0169808 |
| 156 | down | UTP18_S121 | -0.585215 | 0.000797722 |
| 157 | down | UTP18_S124 | -0.585215 | 0.000797722 |
| 158 | down | IGF2BP2_S162 | -0.831511 | 0.00333968 |
| 159 | down | IGF2BP2_S164 | -0.831511 | 0.00333968 |
| 160 | down | SF3B1_T223 | -0.798378 | 0.00796474 |
| 161 | down | SF3B1_T227 | -0.798378 | 0.00796474 |
| 162 | up | CDK1_T14 | 0.610237 | 0.0174816 |
| 163 | up | CDK1_Y15 | 0.610237 | 0.0174816 |
| 164 | down | EIF5B_S182 | -1.36876 | 0.0101438 |
| 165 | down | EIF5B_S183 | -1.36876 | 0.0101438 |
| 166 | down | GNL1_S51 | -1.0792 | 0.000301311 |
| 167 | down | CBX5_S11 | -3.37002 | 0.00878113 |
| 168 | down | CBX5_S12 | -3.39243 | 0.00977075 |

| | | | | |
|---|---|---|---|---|
| 169 | down | CBX5_S13 | -3.34307 | 0.00967852 |
| 170 | down | CBX5_S14 | -3.40442 | 0.00855299 |
| 171 | down | PRKDC_S2612 | -1.49867 | 0.00864667 |
| 172 | down | SPTBN1_S2165 | -0.617763 | 0.00399188 |
| 173 | down | TOP2B_S1336 | -1.13444 | 0.00274922 |
| 174 | down | TOP2B_S1340 | -1.13444 | 0.00274922 |
| 175 | down | TOP2B_S1342 | -1.13444 | 0.00274922 |
| 176 | down | TOP2B_S1344 | -1.13444 | 0.00274922 |
| 177 | down | PCM1_S1257 | -0.64673 | 0.00104783 |
| 178 | down | PCM1_S1260 | -0.64673 | 0.00104783 |
| 179 | down | PCM1_S1263 | -0.64673 | 0.00104783 |
| 180 | down | ANKRD11_S1852 | -2.00648 | 0.0156656 |
| 181 | down | NUCKS1_S54 | -0.711937 | 0.00157515 |
| 182 | down | NUCKS1_S58 | -0.711937 | 0.00157515 |
| 183 | down | NUCKS1_S61 | -0.711937 | 0.00157515 |
| 184 | down | ESF1_S312 | -1.33331 | 0.00150064 |
| 185 | down | ESF1_S313 | -1.33331 | 0.00150064 |
| 186 | down | GNL1_T48 | -1.0792 | 0.000301311 |
| 187 | down | GNL1_T50 | -1.0792 | 0.000301311 |
| 188 | down | CBX5_T8 | -3.27984 | 0.00233173 |
| 189 | down | PRKDC_T2615 | -1.18334 | 0.0013097 |
| 190 | down | ESF1_T311 | -1.33331 | 0.00150064 |

Supplemental Table S4_Sheet_3

| | significant_24 | label | log2FC | p_value |
|---|---|---|---|---|
| 1 | down | TRIM24_S1028 | -1.03039 | 0.00128777 |
| 2 | down | LMNA_S392 | -1.063 | 0.000257797 |
| 3 | down | TOP1_S10 | -1.5851 | 0.00158071 |
| 4 | down | HMGA1_S44 | -1.69371 | 0.000642383 |
| 5 | down | RPL14_S139 | -5.71664 | 5.18181e-05 |
| 6 | down | EIF3E_S399 | -2.35588 | 0.000415791 |
| 7 | down | ARHGEF5_S11 | -1.93672 | 0.0006968 |
| 8 | down | PPP1R1A_S6 | -1.40485 | 2.11362e-05 |
| 9 | up | KLC3_S497 | 0.926707 | 0.000237571 |
| 10 | down | BLOC1S3_S65 | -1.82597 | 0.000838835 |
| 11 | up | RALGPS2_S308 | 0.901591 | 0.000933691 |
| 12 | down | KMT2B_S821 | -1.706 | 0.00125498 |
| 13 | down | AKAP12_T1264 | -1.55683 | 0.000528208 |
| 14 | down | HMGA1_S99 | -1.03169 | 0.00164243 |
| 15 | down | ALS2_S483 | -2.20944 | 0.000336497 |
| 16 | down | ALS2_S492 | -2.07661 | 0.000502678 |
| 17 | down | REEP4_S202 | -0.925393 | 0.000680594 |
| 18 | down | ARHGAP35_S773 | -1.53399 | 0.000406781 |
| 19 | down | SRRM2_S297 | -0.899456 | 0.00165565 |
| 20 | down | THRAP3_S243 | -0.859608 | 0.000232238 |
| 21 | down | THRAP3_S248 | -0.859608 | 0.000232238 |
| 22 | down | GNL1_S51 | -2.10366 | 0.000310719 |
| 23 | down | GNL1_T48 | -2.28669 | 5.69227e-05 |
| 24 | down | GNL1_T50 | -2.28155 | 7.65703e-06 |
